# Loss of cell junctional components and matrix alterations drive cell desquamation and fibrotic changes in Idiopathic Pulmonary Fibrosis

**DOI:** 10.1101/2024.06.17.599411

**Authors:** Rachana R. Chandran, Preethi Vijayaraj, Rolando Garcia-Milian, Jade King, Kristen Castillo, Liang Chen, Yumi Kwon, Sarai William, Tammy M. Rickabaugh, Justin Langerman, Woosuk Choi, Chandani Sen, Jacelyn E.P Lever, Qian Li, Nikoleta Pavelkova, Erin J. Plosa, Steven M. Rowe, Kathrin Plath, Geremy Clair, Brigitte N. Gomperts

## Abstract

The distal bronchioles in Idiopathic Pulmonary Fibrosis (IPF) exhibit histopathological abnormalities such as bronchiolization, peribronchiolar fibrosis and honeycomb cysts that contribute to the overall architectural remodeling of lung tissue seen in the disease. Here we describe an additional histopathologic finding of epithelial desquamation in patients with IPF, wherein epithelial cells detach from the basement membrane of the distal bronchioles. To understand the mechanism driving this pathology, we performed spatial transcriptomics of the epithelial cells and spatial proteomics of the basement membrane of the distal bronchioles from IPF patients and patients with no prior history of lung disease. Our findings reveal a downregulation of cell junctional components, upregulation of epithelial-mesenchymal transition signatures and dysregulated basement membrane matrix in IPF distal bronchioles, facilitating epithelial desquamation. Further, functional assays identified regulation between Collagen IV in the matrix, and the junctional genes *JUP* and *PLEC*, that is crucial for maintaining distal bronchiolar homeostasis. In IPF, this balanced regulation between matrix and cell-junctions is disrupted, leading to loss of epithelial adhesion, peribronchiolar fibrosis and epithelial desquamation. Overall, our study suggests that in IPF the interplay between the loss of cell junctions and a dysregulated matrix results in desquamation of distal bronchiolar epithelium and lung remodeling, exacerbating the disease.

**One Sentence Summary:** Two-way regulation of cell junctional proteins and matrix proteins drives cellular desquamation and fibrosis in the distal bronchioles of patients with Idiopathic Pulmonary Fibrosis.

## INTRODUCTION

Idiopathic Pulmonary Fibrosis (IPF) is a disease characterized by irreversible scarring of the lungs, resulting in a progressive decline in lung function and ultimately death (*1*). Most studies on IPF focus on the remodeling and dysfunction of the alveolar compartment, however, there is a growing recognition of similar pathological changes occurring in the distal bronchioles (small airways with <2mm diameter). This includes epithelial changes, peribronchiolar fibrosis, aberrant bronchiolization and honeycomb cysts (*2-6*). However, there is no consensus regarding the mechanisms for the underlying changes in the distal bronchioles that drive these pathologies. One potential mechanism for these pathological changes is aberrant epithelial-matrix crosstalk in the distal bronchioles but this possibility has not been well explored. Understanding the mechanisms of distal bronchiolar pathologies is important for developing new therapeutic strategies to prevent or possibly reverse aberrant bronchiolar remodeling in IPF.

The distal bronchiolar wall consists of two major anatomical regions: the pseudostratified epithelia and the underlying basement membrane (BM). The pseudostratified epithelia consists of several cell types including basal, ciliated, goblet, club, and neuroendocrine cells (*7*). Exposure to toxins or diseases like IPF can alter the relative distribution of these cell types (*8-10*). Cell-cell and cell-BM connections in this epithelial layer primarily occur through tight junctions, desmosomes, adherens junctions and hemidesmosomes. Tight junctions, composed of Claudins and Occludins along with the scaffold proteins Zona Occludens (e.g. ZO-1), regulate epithelial barrier permeability (*11*). Adherens junctions, on the other hand, contain Cadherins (e.g. E-Cadherin) and the Nectin family of cell adhesion molecules that bind to cytoskeletal actin through proteins like Catenins (e.g. JUP [Gamma Catenin], p-120 Catenin) and Afadin. Desmosomes and hemidesmosomes maintain cell-cell adhesion and attach epithelial cells to the basement membrane, with proteins like Plectin playing a role in anchoring intermediate filaments and linking them to junctional adhesion complexes (*12-14*) Disruption of these junctional components or cytoskeletal proteins can lead to epithelial detachment and fragility disorders such as epidermolysis bullosa (EB) (*15-17*). The interaction between cells and their extracellular matrix (ECM) plays a crucial role in development, regeneration, and wound healing (*18*). In the distal bronchioles, epithelial cells are anchored to the biphasic matrix known as the BM, which consists primarily of collagens, particularly Collagen IV (COLIV), Laminin, Nidogen and Perlecan (*19*). The BM is a dynamic structure characterized by constant turnover of its constituent proteins, which is essential for maintaining developmental programs, cellular homeostasis, repair and regeneration (*20*). However, dysregulated BM can contribute to disease pathogenesis, primarily due to alterations in mechanical properties and subsequent changes in cell-matrix interactions (*21*).

Epithelial desquamation in distal bronchioles is the detachment of epithelial cells either individually or in sheets from the basement membrane. Desquamating epithelial cells have been identified in the IPF alveolar compartment, and in another interstitial lung disease called pleuroparenchymal fibroelastosis, as well as in upper airways of patients with asthma (*22, 23*). Nevertheless, whether epithelial desquamation constitutes a true pathological feature and whether desquamation can lead to airway remodeling in these diseases has been a subject of debate (*24*).

In this study, we demonstrate desquamation of regions of distal bronchiolar epithelium in IPF and highlight this phenotype as a true pathological feature through mechanistic findings. We show that this phenotype is associated with dysregulation of cell junction assembly and alterations in BM matrix in IPF distal bronchioles through the application of micro-scale global spatial transcriptomics and spatial proteomics. Our investigation further revealed a complex feedback loop that maintains the expression of specific matrix components and cell junctional molecules in the distal bronchioles during homeostasis. Specifically, the BM protein Collagen IV and junctional genes *JUP* and *PLEC* play an important role in this context. We identified dysregulation of the BM matrix and cell junctional genes, including *JUP* and *PLEC*, in IPF distal bronchioles, all of which can result in loss of epithelial adhesion and consequent desquamation. Remarkably, *JUP* and *PLEC* loss of function mutations are also known to be involved in skin detachment disorder epidermolysis bullosa. Our study further shows that loss of the cell junctional protein JUP leads to dysregulated matrix gene expression, particularly COLIV, and that dysregulated matrix downregulates cell junctional genes. This feedback loop likely contributes to the fibrotic burden in the lung during IPF pathogenesis, primarily causing peribronchiolar fibrosis and remodeling of the airway.

## RESULTS

### Extensive desquamation of the epithelium is detected in the IPF distal bronchiolar epithelium

In freshly excised IPF lung tissue in culture, we noted sheets of sloughed multi-ciliated epithelia spinning due to unidirectional beating of cilia (**Movie S1 and fig. S1A**). We suspected sloughing of the ciliated epithelium of the distal bronchioles and therefore examined histological sections of lungs from IPF patients obtained at the time of lung transplantation, and from patients who died with no prior history of lung disease (indicated as healthy lung). Histological analysis of the IPF distal bronchioles was conducted and compared to distal bronchioles at the same terminal bronchiolar level in healthy lung tissue. Shed epithelial cells, either individually or in sheets, in varying degrees, were observed in the distal bronchiolar lumen across all IPF samples that were analyzed (**Fig. 1A**). Of note, multiple areas within the distal bronchioles lacked an epithelial layer, suggesting that the shed epithelial cells in the lumen originated from the distal bronchioles themselves rather than being transported from other locations. There was heterogeneity across the bronchioles and not all bronchioles exhibited significant epithelial sloughing. Healthy lung bronchioles exhibited either no desquamation or significantly less desquamation compared to IPF bronchioles **(Fig. 1A, fig. S1B).** The quantification of non-intact epithelial coverage was performed on 41 bronchioles from six IPF patients and 31 bronchioles from five healthy individuals (Fig. 1B). Non-intact epithelia were defined by the absence of ciliated cells, or both basal cells and ciliated cells in the ciliated epithelium. On average, 60.5%±22.4 of the total perimeter of analyzed bronchioles in the IPF group lacked intact epithelia, whereas in healthy bronchioles, this percentage was only 2.6%±6.7 **(Fig. 1B).** We then sought to identify the specific cell-types undergoing sloughing in the IPF distal bronchioles. Immunofluorescence staining revealed that both KRT5+ basal-like cells and Acetyl-beta tubulin+ ciliated cells experienced sloughing **(Fig. 1C)**. Interestingly, KRT5+ cells in the distal bronchiolar wall also displayed regions of hyperplasia with stratified epithelium, likely indicating a reparative response (**fig. S1C**).

**Figure 1.**
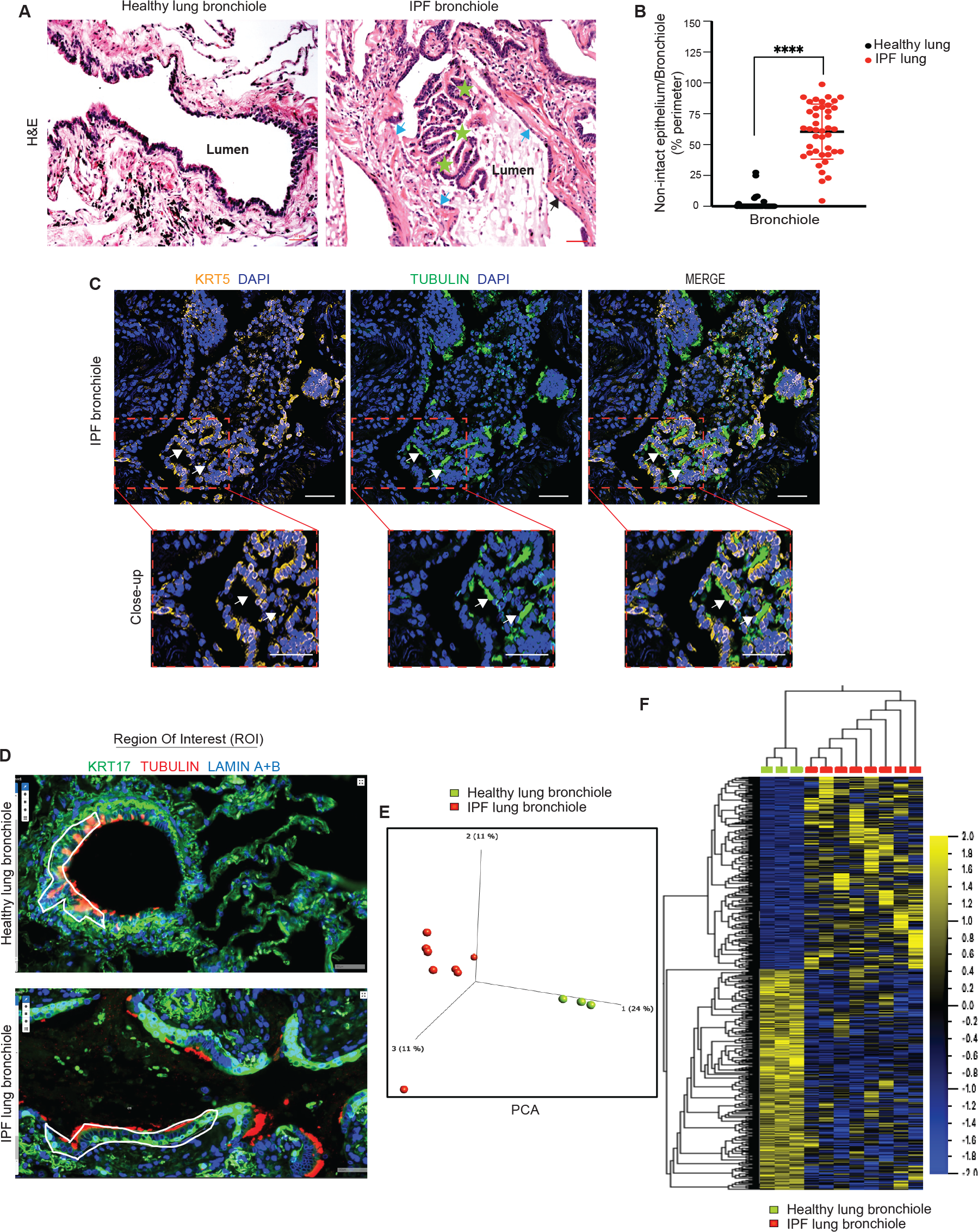
IPF distal bronchioles exhibit epithelial desquamation, and the spatial transcriptome demonstrates dysregulated gene expression in the distal bronchiolar epithelium. **(A)** Healthy and IPF lung sections were stained with hematoxylin and eosin. Representative images of the distal bronchioles are shown. Arrow indicates the bronchiolar wall with no epithelial lining and asterisk indicates sloughed (desquamated) epithelia in the lumen, n=5-6. Scale bar, 50μm. (**B**) The percentage of distal bronchioles with no intact epithelia across the perimeter divided by the total epithelial perimeter of the given bronchiole was quantified from IPF and healthy lung sections (n=5-6 individuals and 31-41 bronchioles total). Unpaired t-test was used with statistical significance **** indicates p<0.0001. (**C**) Lung cryosections were immunostained for the basal cell marker, KRT5 (orange) and cilia marker, Acetylated-Beta Tubulin (green), and nuclei with DAPI (blue) and imaged. Zoomed in images of boxed regions (outlined in red) are shown below. Arrows indicate detached basal and ciliated cells in the lumen. Scale bars, 50μm, (n=5-6). (**D**) Spatial transcriptomic profiling of distal bronchiolar epithelium was conducted on 3 healthy and 8 IPF bronchioles. Regions of interest in the distal bronchiolar epithelium were identified using airway specific antibodies against basal cells (KRT17-488) and ciliated cells (Acetyl-alpha Tubulin-555), and nuclear envelope marker (Lamin A+B-594). Representative images of the ‘Region of Interest’ (ROI) selection of distal bronchioles for transcriptomic profiling are shown. Scale bars, 50 μm. (**E**) Principal Component Analysis (PCA) indicated the distribution of healthy and IPF bronchioles based on the top variants. (**F**) Heat map representing 1351 differentially expressed genes between IPF and healthy distal bronchioles with fold change >2, FDR<0.06 and p<0.05. Scale describes the color code and the direction of differential expression.

### Spatial whole-genome transcriptomics shows distinct gene expression profiles in IPF distal bronchioles

To investigate the cellular pathways contributing to epithelial desquamation in the distal bronchioles of individuals with IPF, we utilized spatial whole-genome transcriptomic profiling with the GeoMx Digital Spatial Profiler. This approach enabled a comparative analysis of the transcriptomic profiles of regions of distal bronchiolar epithelium in IPF patients matched at the same level with bronchioles from healthy individuals. Our focus centered on the multi-ciliated terminal distal bronchiolar epithelium, given the pronounced desquamation within these bronchiolar regions. We ensured that the epithelium we profiled was not sloughed, thereby preventing any bias in the transcriptome linked to apoptotic signals following sloughing. For transcriptomic profiling, fixed lung sections were first subjected to oligo-tagged *in situ* RNA hybridization probes. The probes were designed uniquely to be attached to a UV photocleavable linker to facilitate the spatial technology. Subsequently, multi-ciliated distal bronchiolar epithelia were identified as Regions of Interest (ROI) based on immunostaining of lung sections with florescence-conjugated antibodies: Cytokeratin 17-488 (KRT17-488; a basal cell marker), Acetyl-alpha Tubulin-555 (a ciliated cell marker) and Lamin A+B-594 (a nuclear envelope marker) (**Fig. 1D**). ROIs, each comprising 50-100 cells, were selected from three bronchioles from one healthy lung and eight bronchioles from two different IPF patient samples. Notably, in IPF tissue, basal cells in distal bronchiolar epithelia exhibited higher expression of KRT17 and KRT5 compared to healthy lung tissue (**Fig. 1D, fig. S1D**). In addition, KRT5+ basal cells in IPF distal bronchioles displayed hyperplastic regions (**fig. S1C**). These hyperplastic areas were excluded from spatial transcriptomics ROI selection for two main reasons: a) no comparable hyperplastic KRT17+ enriched regions were present in the healthy lung bronchioles, potentially leading to an overrepresentation of basal cell signatures and interfering with downstream data analysis, and b) hyperplastic KRT17+ regions appeared mostly resistant to sloughing, with minimal observed epithelial shedding.

After selection, the ROIs were exposed to UV to photocleave the linker and release the oligos. The collected oligos underwent library preparation and subsequent sequencing. The normalized transcript counts between healthy and IPF bronchioles were then subjected to Principal Component Analysis (PCA) using Qlucore Omics Explorer, revealing a distinct separation between IPF and healthy controls (**Fig. 1E**). The heat map indicates 1351 significantly differentially expressed genes in IPF bronchioles compared to healthy bronchioles, encompassing both upregulated and downregulated genes (**Fig. 1F**). This analysis focused on genes with a fold change > ±2, FDR<0.06 and p<0.05, indicating significant differential expression.

### Distal bronchiolar epithelia exhibit downregulation of cell junctional molecules in IPF

To further understand the distinction between IPF and healthy distal bronchioles that may contribute to epithelial desquamation, we analyzed these differences in gene expression. Ingenuity pathway analysis (IPA) of the transcriptome revealed a significant downregulation of junctional gene transcripts and consequently predicted decreased cell junction assembly and formation in IPF distal bronchioles compared to healthy controls (Fold change ±2, p<0.05) (**Fig. 2A**). Similarly, utilizing differentially expressed genes with a false discovery rate (FDR)<0.25, Gene Set Enrichment Analysis (GSEA) also indicated a downregulated junctional profile for tight junctions and apical cell junctions. (**fig. S2A**). Among the significantly downregulated junctional genes in IPF bronchioles were Cingulin (*CGN*), Claudin-4 (*CLDN4*), p-120 catenin (*CTNND1*), Connexin 43 (*GJA1*), Junction Plakoglobin (*JUP*), Occludin (*OCLN*), Caveolin-1 (*CAV1*) and Plectin (*PLEC*) (**Fig. 2B**). The diminished expression of OCLN, PLEC and p-120 catenin was validated at the protein level in IPF bronchioles by immunofluorescent (IF) staining of lung tissue (**Fig. 2C-E**). Of note, while we observed a decrease in the levels of these junctional proteins in most bronchioles affected by IPF, there were also instances where some bronchioles showed no decrease in expression, re-emphasizing the heterogeneity observed within IPF lung tissue, and the importance of spatial transcriptomic approaches. Overall, these observations underscore the substantial downregulation of genes and proteins associated with cell junctions in distal bronchioles of individuals with IPF and suggest a mechanism for epithelial desquamation seen in the disease. This observation is consistent with a recent report that IPF distal airway epithelial cells are in an unjammed (fluid like-motile) state compared to the healthy distal airways (*25, 26*).

**Figure 2.**
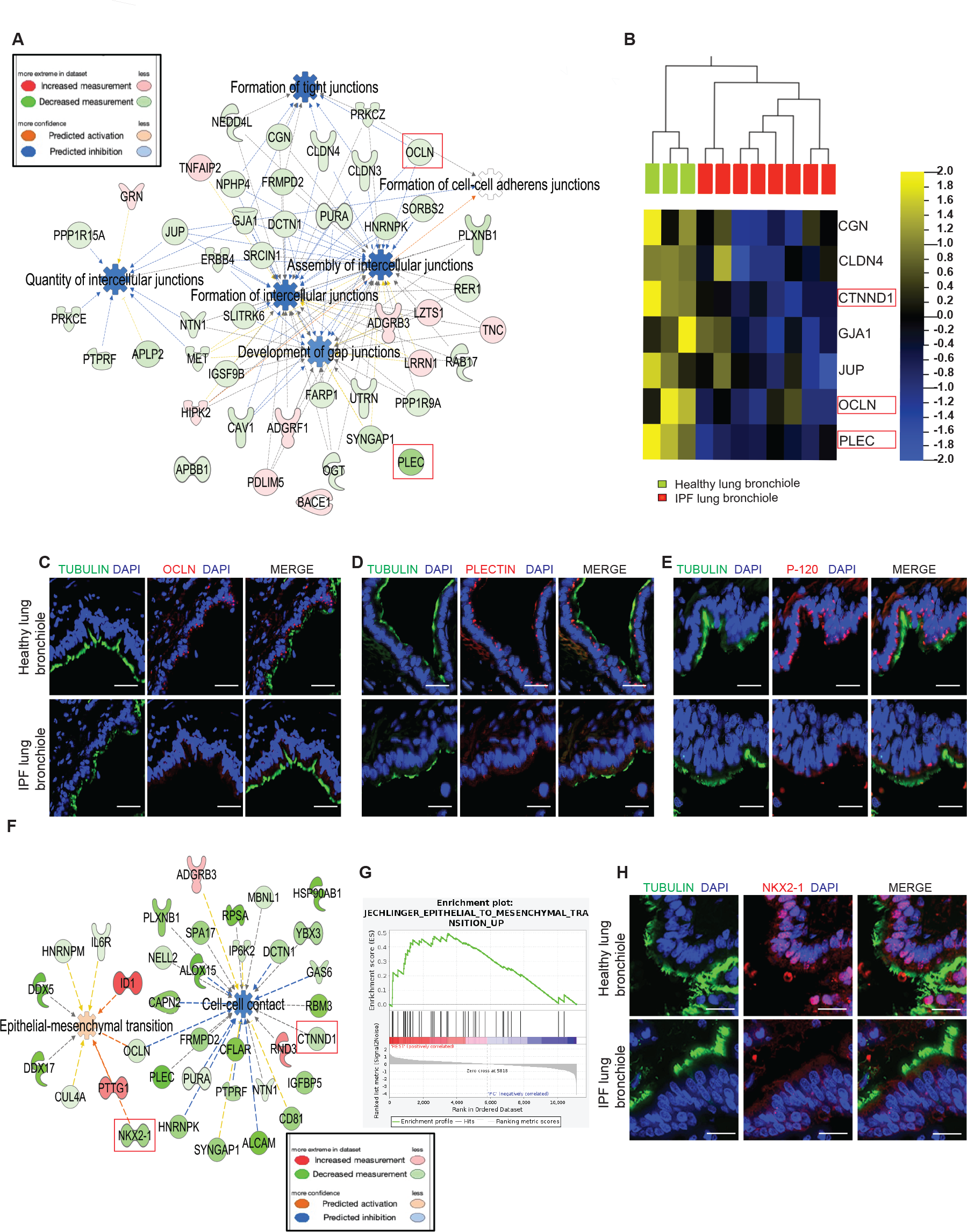
IPF bronchiolar epithelial cells demonstrate downregulation of cell junctional molecules and increased epithelial mesenchymal transition (EMT) signatures. (**A**) Global spatial transcriptomics of healthy and IPF distal bronchiolar epithelia were compared using Ingenuity Pathway Analysis (IPA). Differentially expressed genes with fold change >2 and p<0.05 were analyzed. The blue color of the nodes depicts the predicted inhibition of cell-junction associated pathways in IPF distal bronchioles as compared to healthy bronchioles, as indicated in the legend. (**B**) The heatmap indicated downregulation of select junctional genes in IPF distal bronchiolar epithelia (red samples) compared to healthy bronchioles (green samples). The scale depicts the color code and gene expression changes. (**C-E**) Immunofluorescence staining of healthy and IPF lung focusing on distal bronchioles, sections were stained for ciliated marker Acetyl Beta Tubulin and nuclei (DAPI). (**C**) Expression of junctional protein Occludin (OCLN) is shown, (n=4-6). Scale bar, 50 μm. (**D**) Expression of junctional protein Plectin (PLEC) is shown, (n=4-6). Scale bar, 20μm. (**E**)Expression of junctional protein P-120 is shown, (n=4-6). Scale bar, 20μm. (**F**) The spatial transcriptomes of healthy and IPF distal bronchiolar epithelia were analyzed using IPA for differentially expressed genes with fold change >2 and p<0.05. Red depicts predicted activation and blue depicts predicted inhibition in IPF compared to healthy transcriptome, as indicated in the legend. Inhibition of cell-cell contacts and activation of concomitant EMT associated signatures are shown. (**G**) Gene Set Enrichment Analysis (GSEA) enrichment plot for “Epithelial to Mesenchymal Transition_Up” associated signatures in IPF compared to healthy bronchioles with peak of the enrichment plot to the left indicates enrichment in IPF. (**H**) Immunofluorescence of healthy and IPF lung focusing on distal bronchioles, sections were stained for ciliated marker Acetylated Beta Tubulin, lung epithelial specific marker NKX2-1 and nuclei (DAPI). n=3-4. Scale bar, 20μm. Highlighted in red box in ‘B’ and ‘F’ are the corresponding transcript changes for Occludin, Plectin, P-120 Catenin and NKX2-1 in the spatial transcriptomic data.

### Distal bronchiolar epithelia exhibit increased EMT signatures in IPF

EMT often precedes disassembly and dysregulation of cell-cell junctions in epithelial cells and is characterized by reduced cell-cell adhesion and increased cell mobility along with matrix production (*27-30*). This prompted us to investigate whether IPF bronchiolar epithelia manifest elevated EMT-associated genetic signatures, further indicating enhanced epithelial cell mobility potentially leading to desquamation (*31*). In agreement with this hypothesis, IPA and GSEA indicated an enrichment of EMT-related gene signatures in IPF bronchiolar epithelia compared to healthy controls (**Fig. 2F, G**). Subsequently, the expression of specific EMT-related genes, including Snail family transcriptional repressor-1 (*SNAI1*), Complement component 1q subcomponent binding protein (*C1QBP*), Integrin subunit alpha 3 (*ITGA3*), Forkhead Box C2 (*FOXC2*), Tenascin C (*TNC*), Integrin subunit beta 4 (*ITGB4*), Decorin (*DCN*), *CD44*, and Inhibition DNA binding1 (*ID1*) were analyzed (*32-37*). These genes exhibited significant upregulation in the IPF cohort compared to healthy controls (**fig. S2B**). EMT-associated genes, Snail family transcriptional repressor-2 (*SLUG*) and Twist family basic helix loop helix transcription factor (*TWIST*) did not display significant changes. Validation through immunofluorescent staining confirmed the upregulation of C1QBP (HABP1), and ITGA3 subunit proteins in IPF tissue (**fig. S2C, D**). Notably, the upregulation of CD44, a receptor for Hyaluronic Acid (HA), and CIQBP, an HA-binding protein, correspond to an increase in the HA levels in IPF lung tissue (**fig. S2E**) (*38-41*). Consistent with these findings, the downregulation of epithelial-specific genes such as NKX2 homeobox 1(*Nkx2-1*) and E-cadherin (*CDH1*), as well as cell polarity-associated genes Pals1-associated tight junction protein (*PATJ1*) and Lethal (2) giant larvae (*LLGL2*), were observed **(fig. S2F**) (*27, 42, 43*). The downregulation of Nkx2-1 was validated at the protein level in lung histology sections from IPF patients with IF staining (**Fig. 2H**). Interestingly, a significant downregulation of the apoptotic marker Annexin A5 (*ANXA5*) and a drift towards increased expression in proliferation markers like MYB-proto-oncogene like 2 (*MYBL2*), Mitotic checkpoint serine/threonine-protein kinase (*BUB1*) and cyclin D1 (*CCND1*), were observed (**fig. S2F, G**) (*44*). Overall, these findings suggest that desquamation is not primarily linked to cellular apoptosis; instead, the cells exhibit active proliferation, heightened migratory behavior due to loss of cell-junctions and a phenotypic shift associated with diminished epithelial cell markers and increased EMT. EMT is often linked with increased matrix production, and that dysregulated matrix production by epithelial cells can lead to altered BM composition and epithelial detachment, thus we next examined the transcriptome of matrisome associated genes in distal bronchiolar epithelium within IPF lung tissue.

### Distal bronchiolar epithelia contribute to the altered composition of the basement membrane in IPF

The interaction between epithelial cells and BM relies on the appropriate composition of the extracellular matrix of the BM and the epithelial expression of their receptors, Integrins and DDRs. GSEA of the spatial transcriptome indicated a higher enrichment score for matrisome-associated genes in IPF (NABA-matrisome associated), which encompasses genes encoding ECM-associated proteins, ECM regulators and secreted factors (**Fig. 3A**). A significantly higher representation of collagen-associated pathways, including collagen biosynthesis, collagen fibril organization and metabolism was observed (**Fig. 3B**). Interestingly, a parallel increase was seen in the collagen degradation pathways with an increase in matrix metalloproteases, implying that overall collagen turnover is increased (**Fig. 3B**). Specific matrix proteins were identified as significantly differentially expressed in the transcriptomic data, with Collagen I (*COLI*), Collagen IV (*COLIV*), Collagen VI (*COLVI*) Collagen II (*COLII*) and Tenascin (*TNC*) among the upregulated matrix genes (**Fig. 3C**).

**Figure 3.**
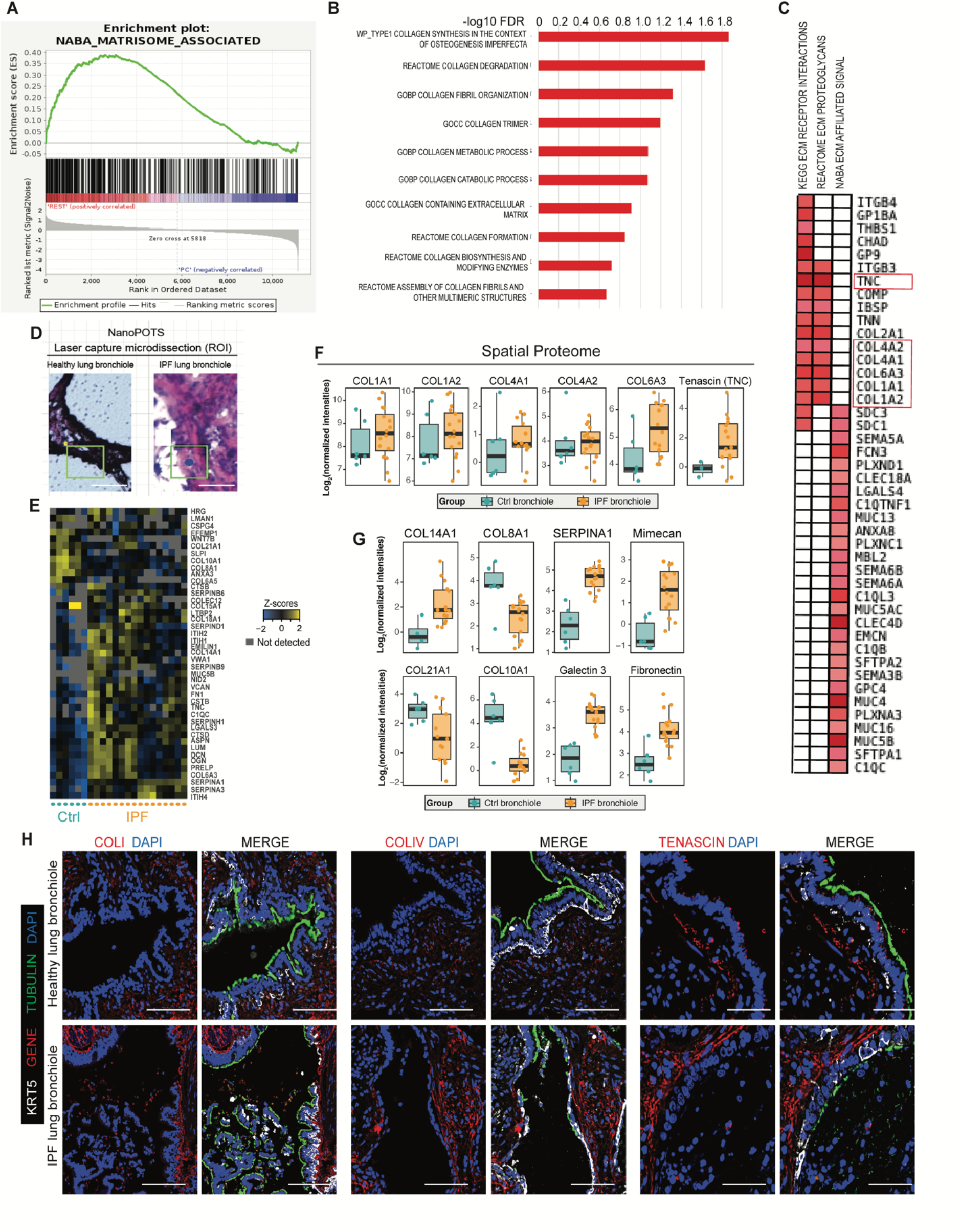
Spatial whole genome transcriptomics and spatial microscale proteomics show that bronchiolar epithelial cells play a key role in altering the composition of the basement membrane matrix in IPF. (**A**) Gene Set Enrichment Analysis (GSEA) enrichment plot for “NABA_matrisome_associated” signatures in IPF compared to healthy with a Normalized Enrichment Score (NES)= 2.05, and FDR (q-value) = 0.006. Peak of the enrichment plot to the left indicates enrichment in IPF. (**B**) GSEA identified pathways with collagen-enriched signatures. (**C**) Heatmap of three different curated matrix enriched gene lists (“Kegg ECM receptor”, “reactome ECM proteoglycans”, “NABA ECM affiliated signal”). Red indicates upregulated gene signatures in IPF distal bronchioles as compared to healthy bronchioles. (**D**) Spatial proteomic profiling of distal bronchioles was conducted for healthy and IPF bronchioles (these were different patient samples from those used for transcriptomic profiling). 50 μm^2^ of distal bronchiole (shown in outlined boxes) was dissected from histology sections on PEN slides using NanoPOTS Laser Capture Microdissection and subjected to Mass Spectrometry. Representative images of the ‘Region of Interest’ (ROI) selection for Laser Capture Microdissection is outlined in green. n=6 (healthy) and n=16 (IPF) regions of distal bronchioles, Scale bar, 50μm. (**E**) Heatmap representing the ECM proteins modulated or differentially expressed (*t*-test or binomial GLM *p*<0.05, respectively) in the IPF bronchiole compared to healthy bronchiolar sections. The MatrisomeDB (*64*) was used to identify ECM proteins for the generation of this Panel. (**F**) Boxplots and scatter plots demonstrate the proteomic abundances of Tenascin, and subunits of Collagens I, IV, and VI. These correspond with the gene expression data in Figure 3C. Note, similar changes in expression at both transcript and protein levels for these genes. Significant heterogeneity was observed. Mann Whitney test, p=0.37 for Col IA1 and ColIA2, p=0.65 for ColIVA1, p=0.63 for ColIVA2, p=0.01 for ColVIA3, and p=0.25 for TNC (**G**) Boxplot representing other ECM or ECM-associated proteins not identified from the transcriptomic data but significantly differentially expressed in IPF compared to control bronchiolar regions (Mann Whitney test, p<0.05 for all the genes shown). (**H**) Immunofluorescence staining of the histological sections from healthy and IPF lung samples focusing on distal bronchioles. Sections were stained for ciliated marker Acetylated Beta Tubulin, and basal cell marker KRT5 and nuclei (DAPI) in addition to COLIV, COLI or Tenascin as indicated, (n=5-6). Scale bars, 50 μm.

Integrins and DDRs, crucial cell surface proteins involved in cell adhesion to the BM, bind to specific matrix partners. We therefore analyzed their gene expression in IPF bronchioles. Interestingly, an inverse correlation in gene expression was observed between specific integrins and their binding partners among the matrix genes (**fig. S2H**). For instance, Integrin subunit alpha 10 (*ITGA10*) expression was significantly downregulated, but the corresponding matrix genes *COLIV* and *COLII*, which bind to ITGA10, exhibited increased expression (*45*). Similarly, *LAMA5*, the protein of which binds to ITGA3 and ITGB4 subunits, showed downregulated expression despite the upregulation of these integrin transcripts in IPF bronchioles. Additionally, the *DDR1* gene which encodes a non-integrin BM binding protein, was significantly downregulated, while COLI, COLIV and COLVI, to which DDR1 binds, were increased (**fig. S2H**). These observations reveal a dysregulation of cell-BM interactions, potentially facilitating epithelial desquamation in IPF.

Given that matrix secreted by epithelia contributes to BM composition, we investigated whether the matrix genes altered in our spatial transcriptomic data in IPF bronchiolar epithelia were consistent with the BM protein composition in IPF distal bronchioles. To this end, we performed spatial proteomic profiling of BM with NanoPOTS microscale laser capture microdissection combined with mass spectrometry. Specifically, 50μm^2^ of the basement membrane from the distal bronchioles of both healthy and IPF lungs was micro dissected and analyzed using mass spectrometry **(Fig. 3D).** We found forty-three significantly differentially expressed proteins in IPF distal bronchiolar BM in comparison to healthy (**Fig. 3E**). Consistent with the transcriptomic observations, the protein expression of COLIV (COLIVA1, COLIVA2), COL1(COL1A1, COL1A2), COLVI (COLVIA3), and TENASCIN exhibited an increasing trend in IPF distal bronchiolar BM (**Fig. 3F**). Of note, some of these proteins did not exhibit statistically significant alterations, emphasizing the inherent heterogeneity within IPF bronchiole. Other collagens with significantly altered expression in the proteomics data included the alpha1 subunits of COLVIII, COLXX1, COLX, COLXIV, and the alpha3 subunit of COLVI. While COLVIII, COLXX1, and COLX displayed downregulation, COLXIV and COLVI showed a significant upregulation in IPF samples (**Fig. 3E, G**). Other matrix proteins that exhibited increased expression include, Alpha-1-antichymotrypsin (SERPINA1), Galectin-3, Prolargin, Fibronectin, Lumican, Decorin, Mimecan, Versican and Asporin. (**Fig. 3E, G**). Immunofluorescent analysis of IPF and healthy lung bronchioles further confirmed increased expression of COLI, COLIV and TNC in the BM of IPF bronchioles. **(Fig. 3H).** Collectively, these findings showed a significant alteration in the BM composition of distal bronchioles in individuals with IPF. Given the dysregulation of cell junctional and BM proteins of the IPF distal bronchioles, we next sought to explore whether the junctional and matrix components interact with each other in IPF.

### Small airway epithelial cells cultured on higher Collagen IV concentrations have reduced cell junction assembly

Collagen IV (COLIV), a major component of the bronchiolar BM, demonstrated increased expression in both COLIVA1 and COLIVA2 at both the transcript and protein levels in the BM of IPF bronchioles compared to healthy controls (**Fig. 3**). Subsequently, we queried whether the downregulation of cell-junction associated genes in the epithelia of IPF bronchioles resulted from the increased expression of COLIV in the BM. To address this hypothesis, small airway epithelial cells (SAECs) isolated from human distal bronchioles were cultured on increasing concentrations of COLIV (0.1mg/ml, 0.3mg/ml, 0.6mg/ml and 1mg/ml). Forty-eight hours later, we found a significant reduction in the transcript levels of the cell junctional genes; *OCLN*, Tight junction protein 1 (ZO-1, *TJP1*), *CDH1, PLEC*, and *JUP* with increasing concentrations of COLIV matrix (**Fig. 4A-E**). This indicates that increased expression of COLIV in the matrix transcriptionally impacted cell junctional stability in SAECs. However, transcript levels of the Caveolae plasma membrane protein, Caveolin 1 (*CAV1*), the EMT related gene, *SNAI1*, and *DDR1* were not changed at higher concentrations of COLIV at this forty-eight-hour time point (**fig. S3A-C**). Downregulation of *OCLN, JUP*, and *PLEC* gene expression in SAECs cultured on higher COLIV-concentrations paralleled the transcriptomic profile we found by spatial transcriptomics of the IPF distal bronchioles, suggesting a mechanism involving COLIV-induced junctional downregulation in IPF distal bronchioles leading to desquamation **(Fig. 4F).** These *in vitro* observations were further validated and quantified at the protein level for PLEC and JUP by western blotting (**Fig. 4G, H**).

**Figure 4.**
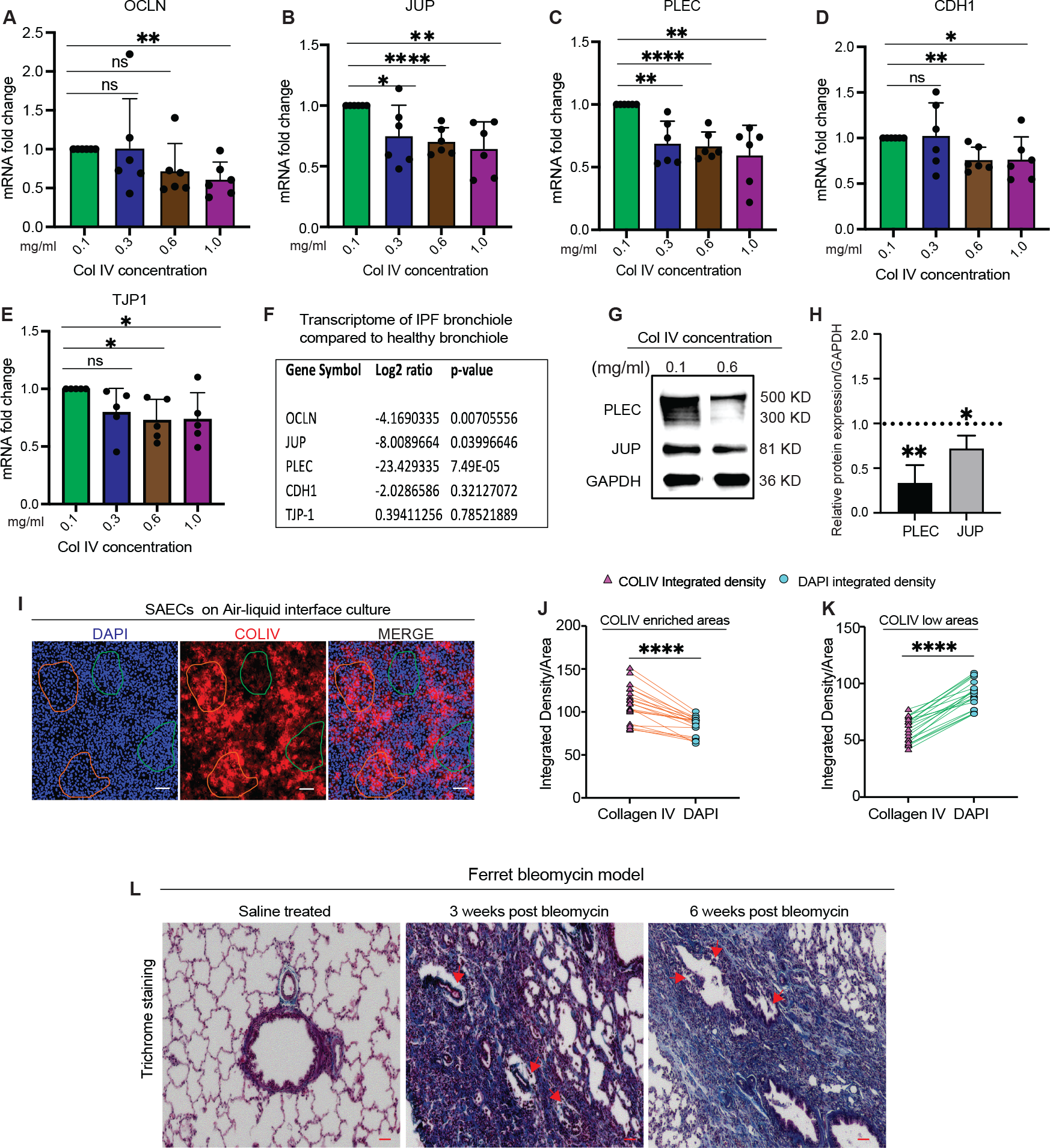
Small airway epithelial cells (SAECs) cultured on higher Collagen IV concentrations have reduced cell junction assembly indicating the role of matrix in maintaining cell-cell adhesion. (**A-E**) RT-qPCR analysis of the transcript levels in SAECs cultured on varying concentrations of Collagen IV matrix, measured relative to GAPDH and normalized to that of 0.1mg/ml COLIV. Transcript levels of (**A**) OCLN, (**B**) JUP, (**C**) PLEC, (**D**) CDH1 and (**E**) TJP1 are shown. n=6 for all transcripts analyzed, students’ *t*-test were used for all the analysis, error bars represent standard deviation. *p<0.05; **p<0.01; ***p<0.0001. (**F**) Analysis of spatial transcriptomic data of IPF bronchioles in comparison to healthy bronchioles of genes tested in **A-E** are shown. Note similar downregulation of some of the juctional genes *ex vivo* in IPF bronchioles, and *in vitro* upon culturing cells on higher COLIV concentrations. (**G**) Westen blot for SAECs cultured on 0.1 and 0.6mg/ml COLIV. JUP, PLEC and GAPDH protein levels are shown. (**H**) Densitometry of the ratio of protein JUP or PLEC to GAPDH. n=3, student’s *t*-test for statistical analysis, * p<0.05; ** p<0.01. (**I-K**) SAECs grown on Air Liquid Interface (ALI) for 28 days and subjected to immunofluorescence for COLIV, KRT5 and DAPI, imaged and quantified using imageJ. (**I**) IF for the endogenous COLIV secreted by cells are shown (in red). COLIV enriched areas are outlined in orange and COLIV low areas outlined in green. (**J**) Quantification of COLIV intensity and DAPI intensity in COLIV enriched areas. (**K**) Quantification of COLIV intensity and DAPI mean gray value (intensity) in COLIV low areas. The lines represent the corresponding COLIV and DAPI mean gray value in a given area and indicate the inverse correlation. Scale bar, 200 μm, n=4, ****p<0.0001. (**L**) Trichome stained lung sections of ferrets treated with saline or bleomycin for 3 or 6 weeks. Arrows indicate desquamated epithelia in the bronchioles. Scale bar, 100 μm, n=3

Furthermore, SAECs cultured on Collagen-I coated transwells and differentiated under air-liquid interface (ALI) conditions produced their own COLIV matrix in a patchy distribution (**Fig. 4I**). Remarkably, we quantified fewer SAECs in the epithelial regions above COLIV enriched patches than areas with less COLIV content where the epithelial cells are more tightly packed together and higher in number (**Fig. 4I-K**). This further suggests that higher COLIV induces migration or extrusion of cells. Similarly, in the context of IPF, it is conceivable that distal bronchiolar epithelial cells overlying COLIV enriched matrix may be extruded or migrated out, and this is likely facilitated by the downregulation of cell-junctional proteins (**Fig. 2, 4**).

### Desquamation of distal bronchiolar epithelia correlates with increased matrix deposition

Based on these observations, we hypothesized that desquamation in IPF patients may not necessarily be confined to end-stage disease, but rather could present in any distal bronchiole with aberrant matrix deposition in the BM, particularly with higher COLIV deposition. As the IPF lung sections obtained during transplantation typically represent end-stage disease, we utilized the ferret bleomycin model to investigate earlier stages of progressive fibrosis. Consistent with our hypothesis, we observed desquamation occurring in regions with higher matrix deposition, as shown by trichrome staining, with no correlation with disease-stage (**Fig. 4L**). To further understand the mechanism for desquamation we next functionally tested the junctional proteins that are significantly downregulated by the presence of higher COLIV matrix, and in the IPF distal bronchiolar epithelium, namely JUP and PLEC.

### JUP transcriptionally regulates matrix and other cell junctional components

*PLEC* and *JUP* loss-of-function mutations are tightly associated with the skin sloughing disorder Epidermolysis Bullosa (EB), which is occasionally associated with upper airway sloughing, highlighting their potential involvement in epithelial sloughing or desquamation in distal bronchioles (*17, 46, 47*). The significant downregulation of *PLEC* and *JUP* transcripts in IPF distal bronchioles and in SAECs cultured on increasing COLIV concentrations, further emphasizes their potential involvement in epithelial desquamation in IPF distal bronchioles. We therefore used a small interfering RNA (siRNA) approach to knockdown *JUP* in SAECs (**Fig. 5A, B, fig. S3D)** to study specific mechanism regulated by JUP. To this end, cultured SAECs were treated with either *JUP* siRNA or scrambled control and after 72 hours, the total RNA was collected and subjected to bulk-RNA sequencing. *JUP* knockdown led to a significant induction of transcriptional pathways related to collagen biosynthesis, collagen trimerization, and compensatory collagen degradation pathways **(Fig. 5C).** In line with these findings, there was an increase in the expression of many collagens, including *COLI* and *COLIV* (**Fig. 5D**). IPA further indicated an increase in Notch pathway components upon JUP knockdown, suggesting a role for Notch signaling in JUP regulation of collagen synthesis **(Fig. 5F**). Interestingly, Notch has previously been shown to regulate COLIV in vascular basement membrane (*48*). In addition, IPA showed downregulation of transcripts related to other cell junctional components including *CDH1, OCLN*, and *CDC42* with the decrease in JUP expression (**Fig. 5E, fig. S3E, Data file S1_JUP KD**). In summary, these findings indicate a significant role for JUP in determining the basement membrane matrix composition and the maintenance of cell-junction integrity of the epithelium, underscoring the intricate interplay between cell junctional and matrix components. We next explored whether PLEC has a similar regulatory role in the bronchiolar epithelium.

**Figure 5.**
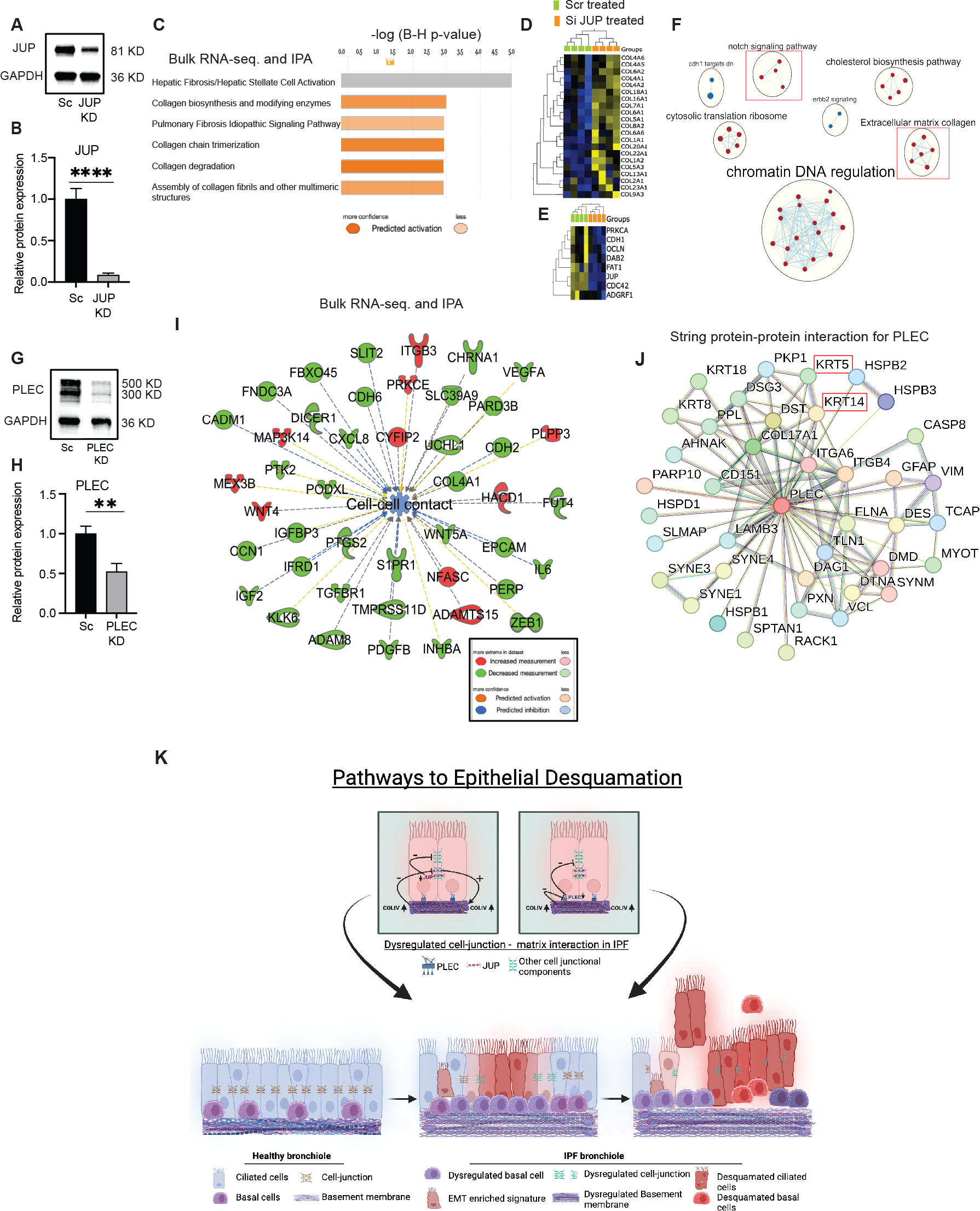
Junctional proteins, JUP and PLEC, transcriptionally regulate matrix and other junctional components, suggesting complex signaling between cell-junction and matrix proteins. (A)Knockdown (KD) efficiency of JUP siRNA in SAECs. Western blot analysis of cell lysates for JUP (81KD) and GAPDH (36KD) in Scrambled and JUP siRNA knockdown. (B) Densitometric analysis of JUP relative to total protein in corresponding samples. n=3; student’s *t-*test, ****p<0.0001. (C) siRNA knockdown of JUP or Scrambled (Scr) control in SAECs followed by bulk RNA sequencing, (n=4) for each Scr and JUP KD. IPA DESeq2 pairwise comparison of bulk RNA transcriptome for JUP KD utilizing differentially expressed genes with fold change >1.2, and FDR. 0.05. Top enriched pathways are shown, orange color code suggests predicted activation as indicated and grey depicts no predicted directionality. (D) Heatmap indicates differential expression of Collagen subunits in JUP KD compared to Scr. (E) Heatmap indicates differential expression of select cell junction-associated genes JUP KD compared to Scr. (F) Gene set enrichment map visualization through Cytoscape for the pathways enriched in JUP KD. Red dots indicate enrichment. Red boxed regions highlight the upregulation of Notch pathway and upregulation of Collagen synthesis. (G)Western blot analysis of cell lysates for PLEC (81KD) and GAPDH (36KD) in PLEC knockdown (KD) and Scrambled (Scr) treated SAECs. (H) Densitometric analysis of PLEC relative to total protein. n=3; student’s *t-*test, ** p<0.01. (I) RNA from PLEC KD and Scr treated samples were subjected to bulk RNA sequencing, n=4 for each Scr and PLEC KD. Transcriptome of Scr treated and PLEC KD SAECs were compared using IPA, differentially expressed genes with fold change >1.2 and p<0.05 were analyzed. Blue depicts predicted inhibition of cell-junction associated pathways by PLEC KD as indicated in the boxed legend. (J) String high confidence (0.700) protein-protein interaction network of PLEC. Boxed regions indicate physical interaction of PLEC with Keratins. (K) Cartoon generated by Biorender demonstrating (i) the role of JUP in regulating cell-cell junctions and collagens, particularly COLIV production, (ii) the role of PLEC in regulating cell-cell junctions and (iii) overall cellular pathways that leads to epithelial desquamation including dysregulation of basement membrane components, downregulation of cell-junctional components and increased EMT signatures suggesting that epithelial desquamation in IPF distal bronchioles is multifactorial.

### PLEC plays a key role in maintaining the cell junctional components and cellular structural framework

We performed siRNA knockdown of PLEC in SAECs and then subjected the total RNA to bulk-RNA sequencing (**Fig. 5G, H, fig. S3F)**. PLEC knockdown in SAECS significantly reduced gene expression of cell-junction-associated genes, including *EPCAM*, *CDH2*, *CCN1*, *PERP*, and *WNT5A* (**Fig. 5I**). IPA also predicted diminished cell-cell contact in the PLEC knockdown group compared to scrambled control (**fig. S3G, Data file S2_PLEC KD**). Of interest, unlike SAECs with JUP knockdown, SAECs with PLEC knockdown did not exhibit any significant upregulation of collagen synthesis pathways by 72 hours after knockdown. As noted earlier, Plectin loss from the epithelia is strongly associated with the skin sloughing disorder EB, likely due to its function in hemidesmosomes. Separately, keratin (KRT)5 and KRT14 loss-of-function mutations are also linked to EB (*49*). Intriguingly, the STRING database indicates a high-confidence physical association of Plectin with KRT5 and KRT14 (**Fig. 5J**), indicating how the interaction between Plectin and Keratin intermediate filaments maintains the cellular structural network and epithelial adhesion to the BM through hemidesmosomes. We propose that the loss of this physical interaction in the absence of Plectin, as in EB, may contribute to the sloughing of epithelia in distal bronchioles. Thus, these findings collectively suggest that Plectin plays a pivotal role in maintaining the cellular structural framework in the bronchiolar epithelium, and that the loss of Plectin could also be a key factor driving epithelial desquamation in IPF.

In summary, we have shown disruption of the tightly regulated interaction between airway epithelial cells and their BM in the distal bronchioles of IPF patients. Our findings suggest that epithelial desquamation in distal bronchioles of IPF patients is a complex and multifactorial process involving altered expression of a multitude of junctional and matrix proteins, within both cellular and BM compartments. Functional analysis suggests that the matrix protein, COLIV, and the junctional proteins JUP and PLEC have key roles in the pathobiology of epithelial desquamation in IPF (**Fig. 5K**).

## DISCUSSION

In this study, we highlight epithelial desquamation as a true pathological feature observed in the distal bronchioles of patients with IPF. We used spatial omics profiling of distal bronchioles, and functional assays, to identify dysregulated cellular pathways leading to this pathology in IPF. Our omics studies suggest that dysregulated cell junctional assembly, loss of epithelial characteristics represented by enriched EMT signatures, as well as an altered matrix profile led to peribronchiolar fibrosis, epithelial desquamation, and thus distal airway remodeling. In addition, our functional studies showed that increased matrix components of the bronchiolar basement membrane, particularly COLIV, exhibit a feed forward inhibition of junctional proteins, particularly JUP, and that this in turn upregulates matrix production. Additionally, our data suggest that downregulation of junctional genes, JUP and PLEC can further inhibit transcription of other cell-junctional genes, leading to loss of cell adhesion, cell extrusion and apoptosis. Interestingly, we observed downregulation of E-Cadherin and p-120 Catenin in distal bronchiolar epithelium in IPF, and these genes have been linked to cell extrusion (*50*). Thus, epithelial desquamation in IPF may in part be a cell-extrusion phenomenon that functions to remove aberrant or injured cells from the airway epithelium. But epithelial desquamation has significant clinical importance because it results in the loss of the epithelial barrier, allowing microbial invasion and exposure to toxins. And the loss of ciliary epithelium leads to impaired mucociliary function, causing the accumulation of cellular debris and mucus, which occludes the airway, and potentially hinders airflow. Ultimately, these factors exacerbate disease progression in patients with IPF.

It remains unclear whether mutations in junctional genes and genes encoded for BM proteins can drive IPF. Nevertheless, a recent Genome-Wide Association study revealed a notable correlation between Desmoplakin (DSP), a junctional gene, and IPF susceptibility (*51*). The biggest risk factors for IPF, namely cigarette smoking and aging, are both associated with loss of normal junctional proteins (*52, 53*). Likewise, the deletion of Lama3, a gene associated with BM protein Laminin α3, can cause junctional-EB in humans and has been shown to worsen the lung fibrosis induced by bleomycin and TGF-β in animal models (*54*). Plectin, a giant junctional protein, connects cytoskeletons, particularly intermediate filament Keratins, to cell-junctions, thereby maintaining the stability of hemidesmosomes. Loss of Keratins can lead to loss of Plectin from hemidesmosomes, causing the scattering of hemidesmosomes, increased EMT and migration of keratinocytes (*55*). Thus, both Plectin loss and loss of function mutations in Keratin genes (KRT5 and KRT14) result in EB. This disorder manifests as fragile skin that blisters easily following minor injury or friction, and this blistering can also extend to the epithelial layer of mouth, stomach, and upper airways (*12, 15, 47, 55, 56*). Similarly, mutations causing loss of function of Junctional Plakoglobin (JUP) lead to EB due to dysfunctional desmosomes (*17*). We demonstrate that JUP knockdown can induce collagen synthesis in SAECs. Conversely, SAECs grown on higher COLIV concentration exhibit significant downregulation of PLEC and JUP gene expression. This suggests a two-way regulation of junctional genes and matrix in the epithelial cells. Interestingly, in the context of our findings, in certain variants of EB, patients exhibit aberrant COLIV accumulation in subepidermal regions (*57, 58*). Overall, our data suggest that downregulation of both JUP and PLEC is likely involved in epithelial sloughing of distal bronchioles in IPF, mimicking their role in the skin sloughing disorder, EB.

Very little is known about the mechanisms involved in the alterations in distal bronchioles and their contribution to IPF pathogenesis. Similarly, whether desquamation is an injury response or an effort to regenerate damaged epithelium is not known. Interestingly, desquamation has also been documented in alveolar epithelial cells in IPF (*59*). In this context, the detachment of type I alveolar epithelial cells from alveoli, followed by replenishment by proliferation and differentiation of type II alveolar epithelial cells, characterizes a reparative process (*60*). Our findings suggest a similar phenomenon occurring in the epithelium of distal bronchioles in IPF, where epithelial shedding prompts basal cell proliferation likely to repair the epithelium resulting in hyperplastic basal cells within the distal bronchiolar epithelium (**fig. S1C**). Interestingly, junctional disruption has been linked to hyperproliferation of progenitor cells in the context of pulmonary fibrosis (*61*). It is also well accepted that alveolar epithelial cell injury stimulates fibroblasts to deposit matrix in the alveolar compartment (*5*). Similarly, our study demonstrated increased matrix deposition (peribronchiolar fibrosis) in the distal bronchiolar compartment of IPF patients. These observations suggest that both alveolar and distal bronchiolar compartments undergo similar pathological changes in IPF, encompassing epithelial sloughing, repair and regeneration, and excessive matrix deposition.

Overall, we have uncovered important mechanisms driving desquamation and matrix changes in the distal bronchioles in IPF, which could ultimately be targeted therapeutically. We anticipate that therapeutic approaches to stabilize cell junctions of the distal lung airways will reduce matrix deposition which, in turn, could further stabilize cell junctions to alleviate epithelial desquamation, distal bronchiolar remodeling, and IPF progression.

*Limitations to our study*: IPF is a progressive lung disease characterized by spatial and patient heterogeneity, and this heterogeneity is reflected in our data in both the transcriptomic and proteomic profiles of IPF samples. Another caveat is that we utilized the Nanostring GeoMx digital spatial profiler, which allowed us to detect unbiased transcriptomic alterations in specific regions comprising 50-100 cells of the ciliated distal bronchiolar epithelia. Newer technologies now offer more comprehensive approaches, such as unbiased single-cell spatial transcriptomics, which promise to further advance the field. Further, an intrinsic challenge of our approach lies in the profiling of small spatial regions within IPF. As our data captures a snapshot in time, different bronchioles are likely to be at varying stages of the disease.

## MATERIALS AND METHODS

### Human samples and ethical compliance

Lung tissue from deidentified healthy donors and patients with IPF were obtained at the time of lung transplantation from Ronald Reagan Medical Center and from Translational Pathology Core Laboratory at UCLA. Healthy lung samples were also obtained from the International Institute for the Advancement of Medicine (IIAM). These studies were exempt from full review by the UCLA institutional review board as the samples obtained were deidentified (IRB exemption number 21-000124).

### Ferret fibrosis model and ethical compliance

All animal protocols used in this work were reviewed and approved by the University of Alabama at Birmingham Institutional Animal Care and Use Committee. Age- and sex-matched wild-type ferrets (Marshall BioResources) were assigned to Bleomycin or saline control. A single dose of bleomycin sulfate (5U/kg) in sterile endotoxin-free saline (400 µl) or control saline (400 μl) was instilled with an intratracheal microsprayer (Biojane, China) through the trachea of anesthetized ferret, 2 cm from the carina as described previously(*62*). After the desired timeline, ferrets were euthanized with injectable anesthetics (dexmedetomidine and ketamine), lung lobes were inflated, fixed, and paraffin embedded. Masson’s trichrome staining was performed on paraffin-embedded lung sections. Images were obtained from Lionheart FX (Biotek) in the UAB High Resolution Imaging Facility and the UAB Cystic Fibrosis Research Center.

### Lung tissue preparation and Immunohistochemistry

#### Lung tissue preparation

Lung tissue was cut using sharp scissors into 0.5cm^3^ slices, washed in cold PBS with Ca^+^, Mg^+^ and 1% primocin, and then fixed in 4% PFA in 4^0^C. Lung slices were then washed with PBS, serially dehydrated in 50% ethanol (O/N) and then in 70% ethanol. Tissue blocks were paraffin embedded and the histological sections were prepared at 4um thickness.

#### Immunofluorescent staining

In preparation for staining, the sections were heated to 65^0^C for 1-2 hours to deparaffinize and treated with Histoclear II (Electron Microscopy Sciences) twice for 15 minutes each. They were then rehydrated into water through serial dilutions of ethanol. Rehydrated sections were washed with TNT (10 mM Tris-HCl, pH 8.0, 150 mM NaCl, 0.2% Tween-20) for 15 minutes. Sections were then incubated in prewarmed (65 °C) 1X Antigen retrieval buffer (DAKO) for 10 minutes in a microwave, then incubated in the slide container on ice for 30 minutes. After cooling down, sections were washed in TBST with 0.1% TritonX-100 for 10 minutes, blocked in Protein block (DAKO) for an hour, and incubated with primary antibodies overnight. The next day, sections were washed three times for 10 minutes each in TBST followed by a 10-minute wash in PBST and then PBS. Autofluorescence quenching was performed according to manufacturer’s instructions (Vectashield True VIEW quenching kit). After washing two times for 10 minutes in PBS, secondary fluorescent antibodies were added at 1:100 dilution along with DAPI. Slides were incubated in a humidified chamber in the dark for 1 hour, washed three times for 10 minutes in PBST and were mounted in Vectashield vibrance antifade mounting media. The primary antibodies used were KRT17-488 (ab185032, Abcam), Lamin A+B-594 (ab215324, Abcam), Acetyl Alpha Tubulin-555 (58241S, Cell Signaling), anti-KRT5 (905901, BioLegend), anti-HABP1 (24474-1-AP, Proteintech), anti-ITGA3 (sc-374242, Santa Cruz Biotechnology), anti-Occludin (91131S, Cell Signaling), anti-p120 (NBP2-45646, Novus), anti-Plectin (MAB5674, EMD Millipore Corp.), anti-Tenascin (AB108930, Abcam), anti-Collagen I (72026, Cell Signaling), anti-Collagen IV (AB6586, Abcam), anti-HA-FITC (MBS2014695, BioSource), and anti-Acetylated Tubulin (T7451-Sigma). Secondary antibodies (1:250-500) were raised in goat or donkey and conjugated to either Alexa-488, Alexa-594 or Alexa-647 (Invitrogen). Nuclei were visualized by DAPI (1:1000, Sigma). The images were taken using Zeiss Axio fluorescence microscope or Zeiss confocal LSM880 and image processing was done using ImageJ-version 2.9.0 or Adobe Photoshop-version 22.4.1.

### *Ex-vivo* lung slice cultures

Freshly obtained lungs from IPF patients at the time of transplantation were sliced into 8mm^3^ sections and cultured in DMEM with 10% FBS for 48 hours. The brightfield timelapse images were obtained using Zeiss Axio Vert.A1.

### SAEC culture and freezing

Small Airway Epithelial Cells were purchased from Lonza (CC-2547). To expand, the cells were cultured on culture dishes coated with 0.05mg/ml human placenta derived COLIV (Sigma) in Bronchial epithelial cell growth medium with added supplements according to the manufacturer’s recommendation (BEBM-Lonza) to 80% confluency. For subculture, cells were trypsinized in 0.25% trypsin-EDTA, and trypsin was neutralized in 10% FBS. Cells were pelleted and cultured in BEBM. To freeze the cells, cell pellet of ∼5x10^5^ cells are resuspend in 500ul BEBM media, transfered to a cryovial, and then 500ul of Bambanker Direct (Bambanker) is added. The cryovials were then kept on ice for 15 minutes, transferred to -80^0^C overnight and stored in liquid Nitrogen.

### Collagen IV gradient assay

To prepare the COLIV (derived from human placenta-Sigma) stock solution, 5mg COLIV powder was dissolved in 2.5ml of cell culture grade water. Five microliters (17.4M) of glacial acetic acid was added and then the solution was incubated for 2 hours at 37^0^C or until the COLIV was completely dissolved. The COLIV was filter sterilized using a 0.2um filter and stored at -20^0^C. Cell culture dishes were coated with COLIV at varying concentrations comprising 0.1mg/ml, 0.3mg/ml, 0.6mg/ml and 1mg/ml in a 12-well cell culture plate. Excess COLIV were removed immediately, and the plates were dried overnight at 4^0^C. We plated 2x10^5^ SAECs in each 12-well COLIV coated plate and incubated the cells in complete Bronchial Airway Epithelial Cell Media (BEBM-Lonza) at 37^0^C for 2 days. After 2 days, cells were washed in PBS and lysed for RNA or Protein isolation.

### RNA isolation, cDNA synthesis and quantitative Real Time-PCR (qRT-PCR)

Cultured SAECs were washed with PBS and total RNA from the samples were extracted using the RNeasy Mini Kit (Qiagen) according to the manufacturer’s instructions. The RNA concentrations and purity were measured on a NanoDrop ND-1000 spectrophotometer. We used 200ng of total RNA to synthesize single-stranded cDNA using iScript cDNA synthesis kit based on manufacturer’s directions (Bio-Rad). cDNA was then used for qRT-PCR analysis using SYBR Green qPCR master mix (Bio-Rad) to compare the gene expression between controls and experimental group using the StepOnePlus (Applied Biosystems). Relative fold change in gene expression was identified using gene specific primers and normalized to GAPDH. Primer sequences are listed in Table S1. Data generation and analysis was done using GraphPad Prism, version 9.5.1.

### Western blot analysis

SAECs were lysed in RIPA buffer containing phosphatases and proteases inhibitors (Gold Biotechnology), centrifuged at 13000xg for 10minutes, and the supernatant was collected. Protein concentrations were determined by the BCA assay (Thermo Scientific). To 10 μg protein, 6X Laemmli buffer (Bio-Rad) with β-mercaptoethanol was added and boiled at 95^0^C for 5 minutes. Proteins were separated using SDS-PAGE, transferred to nitrocellulose membrane using iBlot2 (Invitrogen) dry blotting for 7 minutes at 20V. The membrane was then blocked for an hour in 5% dry milk in TBST with 0.1% tween 20 and incubated with primary antibodies overnight at 4^0^C. primary antibodies used were anti-Plectin (MAB5674, EMD Millipore Corp.) anti-Jup (LS-B4364, LS Bio.) and anti-GAPDH (sc-47724, Santa Cruz Biotechnology). The membranes were then incubated with HRP-conjugated anti-mouse secondary antibodies (Cell signaling) for 1 hour. Detection was performed with Super Signal^TM^ West Femto (Thermo Fisher) and ChemiDoc^TM^ MP Imaging System (Bio-Rad), and image quantification was done using Image Lab software (Bio-Rad). Protein concentration was normalized to GAPDH or total protein.

### siRNA knockdown

SAECs from early passages (P3-P4) were collected after trypsinization, and the cells were treated with P3 primary cell 4D-nucleofector reagents (Lonza) according to the manufacturer’s directions. Cells in the nucleofection reagent were transferred to the Nucleocuvettes and treated with 50nM JUP siRNA, PLEC siRNA or Scrambled siRNA (Origene). Cells were then subjected to electroporation using the preset “Normal human bronchial epithelial cell” Nucleofector program. Cells were then incubated on ice for 10 min and plated on to COLIV coated cell culture plates. After 72 hours, cells were washed in 1X PBS, and then subjected to RNA or Protein isolation using respective lysis buffers.

### Air-liquid interface cultures

SAECs (Lonza-CC2547) were cultured on rat-tail collagen I coated (0.03mg/ml) 24-well transwells (Corning) at a seeding density of 50K cells/well. The cells were initially cultured for 3-days in ALI-growth medium (Lonza) based on manufacturer’s instructions. The culture was then airlifted and supplemented with ALI differentiation media (Lonza) in the bottom chamber of the transwell. After 28 or 30 days, the ALI cultures were PFA (4%) fixed and immunostained with antibodies of interest.

### Bulk-RNA sequencing

Total RNA was isolated from small airway epithelial cells (SAECs) and treated with Scrambled, siJUP or siPLEC, using QIAwave RNeasy Mini Kit (Qiagen). RNA quality was determined initially using nanodrop, followed by Agilent 2200 and TapeStation Analysis Software 3.2. All samples had excellent quality with RNA integrity number (RIN) 10. Libraries for RNA-Seq were prepared with KAPA mRNA-Seq Hyper Prep Kit. The workflow consists of (1) mRNA enrichment and fragmentation, (2) first strand cDNA synthesis using random priming followed by second strand synthesis converting cDNA:RNA hybrid to double-stranded cDNA (dscDNA), and (3) incorporating dUTP into the second cDNA strand. cDNA generation is followed by end repair to generate blunt ends, A-tailing, adaptor ligation and PCR amplification. Different adaptors were used for multiplexing samples in one lane. Sequencing was performed on Illumina NovaSeq6000 for a PE 2x100 run. Data quality check was done on Illumina SAV. Demultiplexing was performed with Illumina Bcl2fastq v2.19.1.403 software.

### Spatial transcriptomics

PFA (4%) fixed, paraffin embedded healthy lung and IPF lung blocks were freshly cut using microtomy. The GeoMx®slide was prepared and hybridized with Nanostring GeoMx® v.1.0 Human NGS Whole Transcriptome Atlas RNA Panel as per manufacturer instructions. The slide was stained with morphology markers as described in the results section. Regions of Interest (ROIs) were selected via the instrument interface and collected on microplates by the GeoMx® Digital Spatial Profiler (DSP) platform. One library was generated per ROI selected. Unique adaptors were used during library preparation to enable multiplexing samples. Sequencing was performed on the Illumina NovaSeq6000 platform using paired-end 2x150bp parameters. Data quality check was performed on Illumina SAV. Demultiplexing was performed with Illumina Bcl2fastq v2.19.1.403 software. Four genes with high background were removed from the initial data by Nanostring support staff.

### Laser Capture Microdissection

Fresh lung samples were embedded in Optical Cutting Temperature (OCT), and flash frozen in liquid nitrogen. Samples were sectioned and transferred to PEN membrane slides. The sections were used for microdissection. Sample isolation was performed using a PALM MicroBeam laser capture microdissection (LCM) microscope (Carl Zeiss MicroImaging, Munich, Germany) equipped with a RoboStage and a PALM RoboMover. A nanoPOTS chip was preloaded with 200-nL DMSO droplets to serve as a capture reagent. The region of interest (ROI) of the tissue section was selected by drawing a 50 × 50 µm^2^ tissue voxel under 63 × magnification using PalmRobo software. The tissue sections were cut at an energy level of 42 and a focus level of 21, followed by catapulting into the DMSO droplet with an energy of delta 25 and a focus level of delta 10. After sample collection, the nanoPOTS chip was heated to 70°C for 15 minutes to evaporate the DMSO droplet and stored at -20°C until use.

### NanoPOTS sample preparation

NanoPOTS sample preparation was slightly modified based on the previously described method (*63*). Briefly, 200 nL of cell lysis buffer containing 0.1% DDM and 1 mM TCEP in 50 mM ammonium bicarbonate (ABC) buffer was added to each nanowell. The nanoPOTS chip was then sealed with a glass cover to minimize liquid evaporation and incubated at 70°C in a humidity chamber for 60 minutes for cell lysis, protein extraction, and denaturation. Next, 50 nL of 10 mM CAA in 50 mM ABC buffer was added and incubated for 30 minutes at room temperature in the dark for the alkylation of protein sulfhydryl groups. Subsequently, 50 nL of a digestion solution containing 0.5 ng Lys-C and 2 ng trypsin in 50 mM ABC was added and incubated at 37 °C overnight. To stop the digestion, 50 nL of a formic acid solution (5%, v/v) was added. Finally, the droplets were completely dried in a vacuum desiccator. The nanoPOTS chip was covered, wrapped with aluminum foil, and stored at -20 °C before Liquid Chromatography-Mass Spectrometry (LC-MS) analysis.

### Liquid chromatography and Mass spectrometry

Dried peptide samples on chips were dissolved with mobile phase A (0.1 % FA in water), then injected to the SPE column for 5 minutes with a mobile phase A. After reverse washing, samples were eluted and separated at 100 nL/min using gradient of mobile phase B (0.1% FA in acetonitrile. A 30-minute and 100 nL/min linear gradient from 8% to 22% mobile phase B (0.1% formic acid in acetronitrile) followed 9-minutes linear gradient from 22% to 35% mobile phase B was used. A timsTOF SCP mass spectrometer equipped with Captive Spray source was used to collect the data. For fractionated library, DDA-PASEF were adopted with high-sensitivity mode enabled. For DDA-PASEF acquisition, one MS1 scan was followed by 7 PASEF MS/MS scans per acquisition cycle. The ion accumulation and ramp time was 166 ms and ion mobility (IM) range was set from 1/K0 = 0.6 Vs cm-2 to 1.6 Vs cm-2. Single charged ions were filtered with a polygon filter and precursors for MS/MS were picked at an intensity threshold of 500 a.u. re-sequenced until reaching a 10,000 a.u. with a dynamic exclusion of 0.4 min. Precursor ions for MS/MS analysis were isolated with a 2 Th window for m/z <700 and 3 Th for m/z>800 in a total m/z range 100-1,700.

The collision energy was starting from 59 eV at 1/K0 =1.6 Vs cm-2 to 20 eV at 1/K0 = 0.6 Vs cm-2. For DIA-PASEF analysis, we used a dia-PASEF method including four IM windows per dia-PASEF scan with variable isolation window widths adjusted to the precursor densities. Eight, 10, 8 and 4 dia-PASEF scans were deployed. The IM range was set from 1/K0 = 0.6 Vs cm-2 to 1.6 Vs cm-2. The accumulation and ramp times were specified as 166 ms. The collision energy was starting from 59 eV at 1/K0 =1.6 Vs cm-2 to 20 eV at 1/K0 = 0.6 Vs cm-2.

### Bioinformatics

#### Spatial transcriptomics data analysis

Q3 normalization was used to normalize read counts across samples. Differential analysis between the groups was done by two-Sample Independent *t-*Test. The cutoff value to select differentially expressed genes was Log2ratio |0.5|, FDR p<0.05. Qlucore Omics Explorer (Qlucore, Lund, Sweden), a dynamic, interactive visualization-guided bioinformatics program with a built-in statistical platform was used for differential analysis and data visualization including Principal Component Analysis and heatmaps, and volcano plots.

#### Bulk-RNA seq. data analysis

PartekTM FlowTM software (version 11) bulk RNA-seq pipeline was used for data analysis. In summary, trimmed reads were aligned to the hg38 genome reference using STAR (version 2.7.8), and subsequently Partek E/M algorithm was used to count reads mapping to the genes from Ensembl release 109. We applied DESeq2 normalization method and top 1000 genes by variance were analyzed for PCA. Heatmaps show row-normalized relative gene expression z-scores across columns. Differential expression was performed using the DESeq2 method. The cutoff value to select differentially expressed genes was Log2ratio |0.5|, FDR p<0.05.

#### Functional analysis

Overrepresented pathways, biological functions, upstream regulators, and biological networks were identified using Ingenuity Pathway Analysis (QIAGEN Redwood City, CA). For this, DEP identifiers are mapped and analyzed through one-tailed Fisher Exact Test, (FDR p< 0.05). In addition, Gene Set Enrichment Analysis (GSEA) was also used to identify significant signatures, pathways, and functions. Gene Set Enrichment Analysis (GSEA -Mac App Broad Institute Inc.) was performed using DESeq2-normalized counts with gene set permutation and otherwise default settings. Gene counts were tested against the Hallmak M2 subcollection CP: canonical pathways, and databases (version 7.5.1). The GSEA analysis was based on the C2 and C5 Canonical Pathways and Biological Process collections respectively (MSigDB, Broad Institute, and UC San Diego). Gene sets with an FDR (q value) ≤ 0.25 were considered significantly enriched. The overlap and connections between the resulting different gene sets were produced by the Enrichment Map Plugin (http://baderlab.org/Software/EnrichmentMap) for Cytoscape 3.8 (Shannon et al. 2003), considering a q value of FDR < 0.05. The nodes were joined if the overlap coefficient was ≥ 0.375. In addition, enriched pathways, upstream regulators, and functions were determined by Overrepresentation Analysis using Ingenuity Pathway Analysis (IPA, Qiagen, CA. 2024) and Fisher’s exact test Benjamini-Hochberg FDR p<0.05.

#### Mass spectrometry

The raw files from the Bruker TimsTOF scp instrument were analyzed using a spectral library for the identification of peptides. The library was constructed using the library fraction runs with MSFragger (v21.1) using default parameters against the UniProt non redundant database downloaded on February 20, 2024). The DIA-PASEF NanoPOTS runs were searched and quantified using DIA-NN (v1.8.1) in against the library built in MSFragger, all parameters were set to default except the number of miss cleavages set to “2” and the Neural network classifier that was set to “double-pass mode”. The resulting data was analyzed in R (v4.3.2) using the R package RomicsProcessorv1.2.0 (https://github.com/PNNL-Comp-Mass-Spec/RomicsProcessor). Only the proteins detected in fifty percent of the LCM-cuts from a slide (e.g., 3/6 or 2/4 cuts for the different bronchioles) were considered quantifiable. The data was log2 transformed and median centered, and principal component analysis was performed to verify that the sections from IPF bronchiole partitioned from the ones from control bronchioles. Student’s *t*-tests were performed to compare quantitative values for IPF and control bronchioles. To evaluate if the missingness in one condition compared to the other was random, we used a GLM binomial test. Enrichment analyses for the protein significantly modulated or detected more in one condition were performed using the package Protein MiniOn.

## List of Supplementary Materials

Figures S1 to S3

Table S1

Data file S1_JUP KD

Data file S2_PLEC KD

Movie S1

## Acknowledgments

**Funding:** This work was supported by National Heart, Lung, and Blood Institute (NHLBI) U01HL153000 (to BNG), (NHLBI) U01HL148860 (to GC) and by The Three Lakes Foundation and the Ablon Research Scholars Award (to BNG). Statistical analyses were supported by the NIH/National Center for Advancing Translational Science (NCATS) UCLA CTSI Grant Number UL1TR000124. We appreciate the UCLA Broad Stem Cell Research Center Microscopy Core, the JCCC Translational Pathology Core Laboratories and the Technology Center for Genomics and Bioinformatics. Parts of this work were performed in the Environmental Molecular Science Laboratory, a U.S. Department of Energy (DOE) national scientific user facility at Pacific Northwest National Laboratory (PNNL). Battelle operates PNNL for the DOE under contract DE-AC05-76RLO01830. This work was supported by National Heart, Lung, and Blood Institute (NHLBI) U01HL148860 (to G. Clair) and funding from the Three Lakes Foundation

## Author contributions

Conceptualization: RRC, PV, BNG

Methodology: RRC, JK, KC, LC, YK, SW, TMR, CS, WC, RGM, GC, SR, BNG

Investigation: RRC, PV, GC, RGM, JL, KP, SR, GC, BNG

Visualization: RRC, RGM, GC, BNG

Funding acquisition: GC, BNG

Project administration: RRC, BNG

Supervision: BNG

Writing – original draft: RRC, RGM, GC, BNG

## Data and materials availability

Data are available in the main figures and supplementary materials. The source data are available upon request to corresponding author.

