## Supplemental figs. and legends for "Loss of cell junctional components and matrix alterations drive cell desquamation and fibrotic changes in Idiopathic Pulmonary Fibrosis"

**List of Supplementary Materials**

Figures S1 to S3

Table S1

Data file S1_JUP KD

Data file S2_PLEC KD

Movie S1

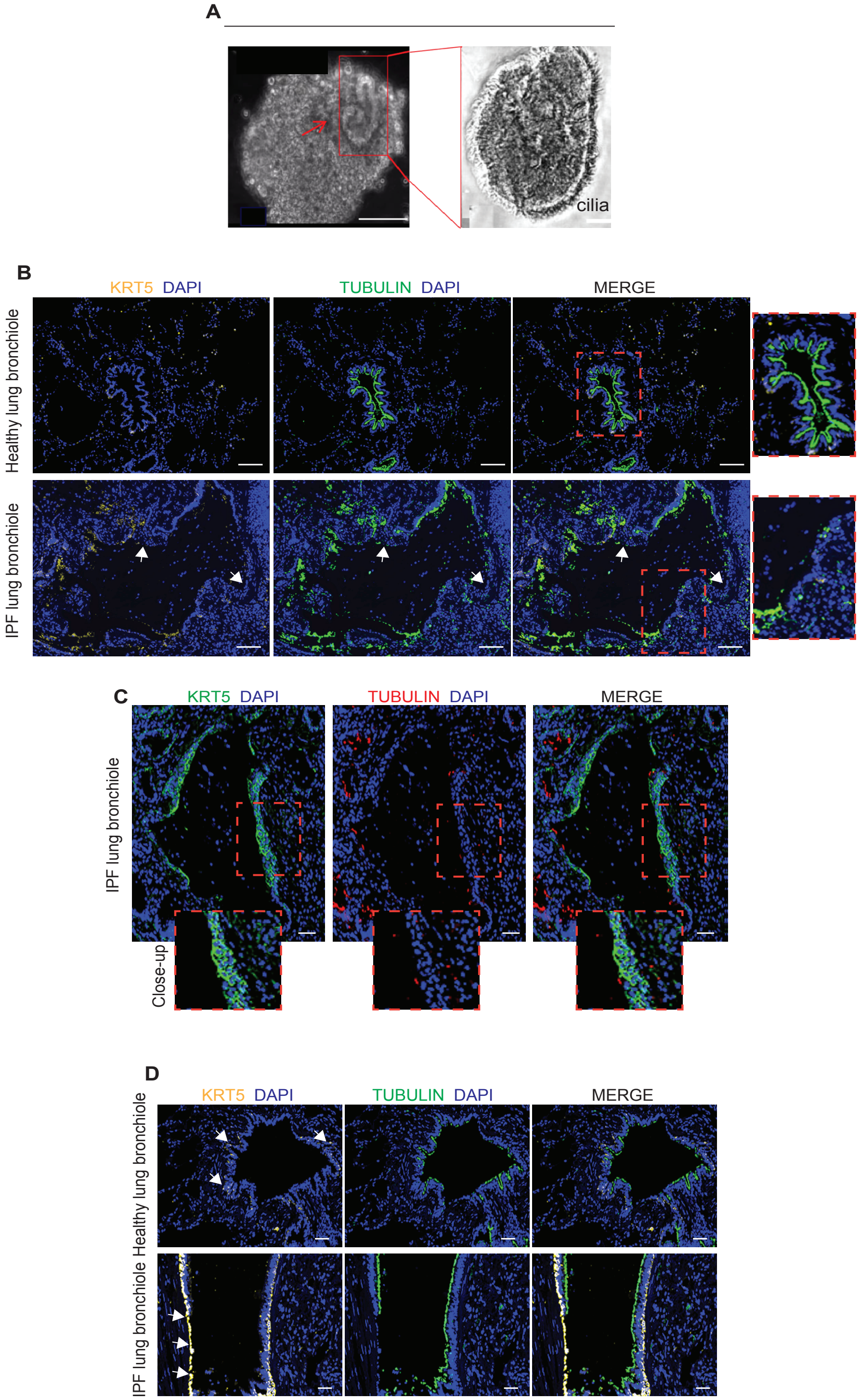
A
cilia
B
KRT5 DAPI
TUBULIN DAPI
MERGE
Healthy lung bronchiole
IPF lung bronchiole
C
KRT5 DAPI
TUBULIN DAPI
MERGE
IPF lung bronchiole
Close-up
D
KRT5 DAPI
TUBULIN DAPI
MERGE
Healthy lung bronchiole
IPF lung bronchiole

**Figure S1. IPF distal bronchioles exhibit epithelial desquamation and basal cell hyperplasia.** **(A-B),** *Ex vivo* IPF lung slice cultures and immunofluorescence analysis of distal bronchioles indicate epithelial desquamation in IPF. (**A)** IPF lung slices were removed using biopsy punch and cultured in DMEM for 48-hrs and imaged. Arrow indicates spinning epithelia as shown in supplementary videoS1. Higher magnification view of the boxed region in red is displayed to the right and indicates cilia around the spinning epithelial structures. (**B)** Lower magnification image of a bronchiole with desquamated/non-intact epithelial cells in IPF in comparison to healthy lung bronchioles. Representative images of the immunofluorescence analysis of distal bronchioles with KRT5 (yellow) Acetyl Beta Tubulin (green) and nuclei (DAPI) are shown. Arrows indicate non-intact epithelia. Boxed regions (in red) are enlarged and shown to the right. Scale bars, 100 µm, n=5-6. **(C-D),** Immunofluorescence of IPF distal bronchioles with KRT5 and Acetyl Beta Tubulin and DAPI **(C)** Representative images of distal bronchioles with KRT5 (green), Acetyl Beta Tubulin (red) and DAPI (blue) are shown. Hyperplastic KRT5 regions are indicated with boxed regions (in red). Close-ups are shown below. Scale bars, 50 µm, n=6. **(D)** Representative images of distal bronchioles with KRT5 (yellow), Acetyl Beta Tubulin (green) and DAPI (blue) are shown. Arrow indicates higher KRT5 expression in IPF compared to healthy. Scale bars, 50 µm, n=5.

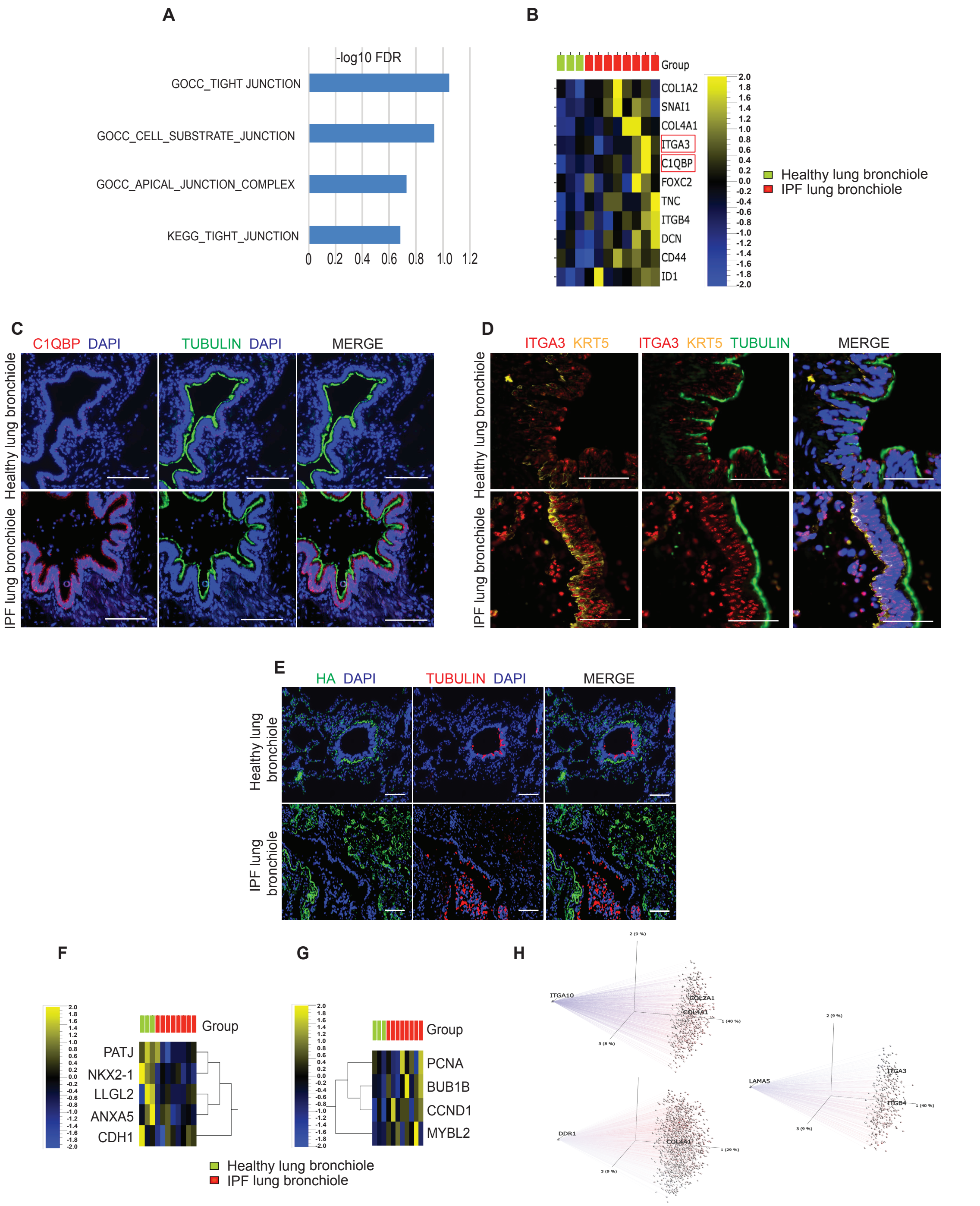
A
-log10 FDR
GOCC_TIGHT JUNCTION
GOCC_CELL_SUBSTRATE_JUNCTION
GOCC_APICAL_JUNCTION_COMPLEX
KEGG_TIGHT_JUNCTION
0 0.2 0.4 0.6 0.8 1.0 1.2
B
Group
COL1A2
SNAI1
COL4A1
ITGA3
C1QBP
FOXC2
TNC
ITGB4
DCN
CD44
ID1
2.0 1.8 1.6 1.4 1.2 1.0 0.8 0.6 0.4 0.2 0.0 -0.2 -0.4 -0.6 -0.8 -1.0 -1.2 -1.4 -1.6 -1.8 -2.0
Healthy lung bronchiole
IPF lung bronchiole
C
C1QBP DAPI
TUBULIN DAPI
MERGE
Healthy lung bronchiole
IPF lung bronchiole
D
ITGA3 KRT5
ITGA3 KRT5 TUBULIN
MERGE
Healthy lung bronchiole
IPF lung bronchiole
E
HA DAPI
TUBULIN DAPI
MERGE
Healthy lung bronchiole
IPF lung bronchiole
F
Group
PATJ
NKX2-1
LLGL2
ANXA5
CDH1
G
Group
PCNA
BUB1B
CCND1
MYBL2
Healthy lung bronchiole
IPF lung bronchiole
H
ITGA10
COL2A1
COL4A1
DDR1
LAMA5
ITGA3
ITGB4

**Figure S2. Spatial transcriptomics of IPF distal bronchioles suggest junctional downregulation, enhanced EMT and dysregulated cell-matrix interaction. (A)** GSEA of differentially expressed transcripts for cell-junction associated signatures. Bar graph indicate cell junction associated pathways, and blue depicts downregulation in IPF compared to healthy. **(B)** Heatmap indicates upregulation of select EMT associated genes in IPF distal bronchiolar epithelia, upregulation of these indicates enhanced EMT. Scale depicts the color code and the Z-score. **(C-D)** Immunofluorescence of the distal bronchioles in healthy and IPF lungs. Lung sections were stained for ciliated marker Acetyl Beta Tubulin (green), nuclei (DAPI), and select EMT related markers (red), and in addition KRT5 (yellow) in **'D'. (C)** Expression of C1QBP (HABP1) and, **(D)** ITGA3 are shown. Scale bars, 100 μm in '**C**' 50 μm in '**D**', n=4-6. Highlighted in red box in '**B**' are the corresponding transcript changes in spatial transcriptomics for C1QBP and ITGA3. **(E-F)** Heatmap depicting select proliferation, apoptosis, and epithelial specific genes. **(H)** Inverse correlation between expression of matrix genes and the corresponding receptors. Principal Component Analysis plot was generated with Qlucore Omics Explorer of connected pairs of variables by Pearson correlation.

Chandran et al., Fig. S3

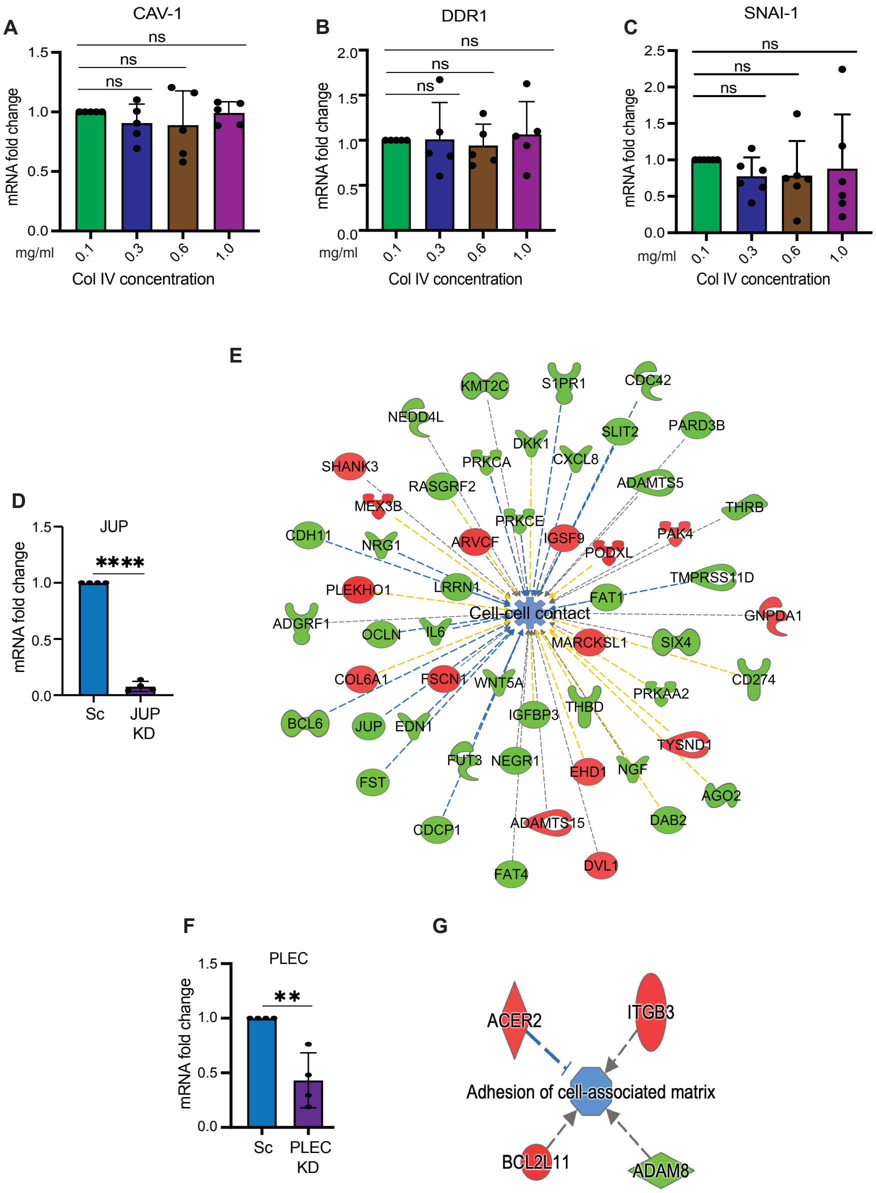

**Figure S3. Functional assays for cell-matrix interaction. (A-C),** SAECs were cultured on varying concentrations of COLIV (0.1,0.3,0.6 and 1mg/ml), two days later the cells were harvested and subjected to RNA or protein isolation and select genes including those differentially expressed in spatial transcriptomic data were analyzed. RT-qPCR analysis of the transcript levels measured relative to GAPDH and normalized to that of 0.1mg/ml COLIV. Transcript levels of **(A)** CAV-1, **(B)** DDR1, and **(C)** SNAI-1 are shown. n=6 for all transcripts analyzed, students' *t*-test were used for all the analysis, error bars represent standard deviation. 'ns' is non-significant. **(D-E)** SAECs were treated with Scrambled (Scr) RNA, siRNA for JUP (JUP KD) or siRNA for PLEC (PLEC KD) to knockdown JUP and PLEC respectively, followed by bulk RNA sequencing and transcriptome analysis, n=4 each for Scr, JUP KD and PLEC KD. **(D)** Knockdown efficiency of JUP vs. Scr at transcript level in SAECs, n=4, student's *t*-test, **** $p<0.0001$. **(E-F)** JUP KD samples were subjected to bulk RNA sequencing, followed by bioinformatic analysis. **(E)** IPA DESeq2 pairwise comparison of bulk RNA transcriptome for JUP KD utilizing differentially expressed genes with fold change $>1.2$ specifically for signatures for cell-cell contact. Blue color at the center indicates downregulation in JUP KD. **(F)** Knockdown efficiency of PLEC vs. Scr at transcript level in SAECs, n=4, student's *t*-test, ** $p<0.01$. **(G)** PLEC KD samples were subjected to bulk RNA sequencing, followed by IPA analysis for cell-matrix interaction. Blue indicates significant downregulation in PLEC KD.

**TableS1**: Primer sequences used for q-PCR

| Gene | Primers |
|---|---|
| GAPDH | Forward: GTCTCCTCTGACTTCAACAGCG<br>Reverse: ACCACCCTGTTGCTGTAGCCAA |
| OCLN | Forward: ATGGCAAAGTGAATGACAAGCGG<br>Reverse: CTGTAACGAGGCTGCCTGAAGT |
| TJP1 | Forward: GTCCAGAATCTCGGAAAAGTGCC<br>Reverse: CTTTCAGCGCACCATACCAACC |
| CDH1 | Forward: GCCTCCTGAAAAGAGAGTGGAAG<br>Reverse: TGGCAGTGTCTCTCCAAATCCG |
| PLEC | Forward: AGCGTGAGAAGGAGAAGCTCCA<br>Reverse: CAGAGAGGAAGCTTTGCTGCAG |
| SNAI1 | Forward: TGCCCTCAAGATGCACATCCGA<br>Reverse: GGGACAGGAGAAGGGCTTCTC |
| CAV1 | Forward: CCAAGGAGATCGACCTGGTCAA<br>Reverse: GCCGTCAAAACTGTGTGTCCCT |
| JUP | Forward: ACCAGCATCCTGCACAACCTCT<br>Reverse: GGTGATGGCATAGAACAGGACC |
| DDR1 | Forward: GATCTCGACTCCGCTTCAAGGA<br>Reverse: CAAAGGGTGTCCCTTACGCACA |
