## Supplementary file 2 for "Loss of cell junctional components and matrix alterations drive cell desquamation and fibrotic changes in Idiopathic Pulmonary Fibrosis"

| Gene ID | Gene name | P-value (PLE( | FDR step up | Ratio (PLEC-I | Log2(Ratio) ( | Fold change ( |
|---|---|---|---|---|---|---|
| ENSG000001 | PLEC | 2.64E-33 | 3.77E-29 | 2.65E-01 | -1.92E+00 | -3.78E+00 |
| ENSG000001 | ARL6IP1 | 1.87E-18 | 1.13E-14 | 1.23E+00 | 3.02E-01 | 1.23E+00 |
| ENSG000001 | GOLT1B | 2.39E-18 | 1.13E-14 | 7.56E-01 | -4.04E-01 | -1.32E+00 |
| ENSG000001 | HAS3 | 4.19E-15 | 1.49E-11 | 1.50E+00 | 5.84E-01 | 1.50E+00 |
| ENSG000001 | INSIG1 | 8.16E-14 | 2.33E-10 | 6.41E-01 | -6.41E-01 | -1.56E+00 |
| ENSG000001 | TMEM65 | 2.31E-13 | 4.93E-10 | 6.63E-01 | -5.94E-01 | -1.51E+00 |
| ENSG000001 | AKT3 | 2.42E-13 | 4.93E-10 | 6.21E-01 | -6.87E-01 | -1.61E+00 |
| ENSG000001 | ERLIN2 | 5.53E-13 | 9.50E-10 | 6.67E-01 | -5.84E-01 | -1.50E+00 |
| ENSG000001 | MMP1 | 6.00E-13 | 9.50E-10 | 5.38E-01 | -8.94E-01 | -1.86E+00 |
| ENSG000001 | GK5 | 1.40E-12 | 1.99E-09 | 6.97E-01 | -5.21E-01 | -1.43E+00 |
| ENSG000001 | ZNF721 | 1.64E-12 | 2.13E-09 | 6.34E-01 | -6.57E-01 | -1.58E+00 |
| ENSG000001 | ASAP2 | 2.07E-12 | 2.35E-09 | 6.82E-01 | -5.52E-01 | -1.47E+00 |
| ENSG000002 | --- | 2.15E-12 | 2.35E-09 | 1.30E+00 | 3.80E-01 | 1.30E+00 |
| ENSG000001 | CPEB2 | 9.40E-12 | 9.04E-09 | 7.02E-01 | -5.09E-01 | -1.42E+00 |
| ENSG000001 | IER3IP1 | 9.51E-12 | 9.04E-09 | 7.08E-01 | -4.98E-01 | -1.41E+00 |
| ENSG000001 | RBPJ | 1.03E-11 | 9.22E-09 | 7.27E-01 | -4.61E-01 | -1.38E+00 |
| ENSG000001 | FAM3C | 3.20E-11 | 2.68E-08 | 6.76E-01 | -5.66E-01 | -1.48E+00 |
| ENSG000001 | KCTD20 | 4.49E-11 | 3.56E-08 | 6.94E-01 | -5.26E-01 | -1.44E+00 |
| ENSG000001 | COA6 | 6.06E-11 | 4.55E-08 | 6.68E-01 | -5.82E-01 | -1.50E+00 |
| ENSG000001 | DDX6 | 7.94E-11 | 5.66E-08 | 8.53E-01 | -2.30E-01 | -1.17E+00 |
| ENSG000001 | JOSD1 | 1.03E-10 | 6.98E-08 | 8.07E-01 | -3.09E-01 | -1.24E+00 |
| ENSG000001 | CCND2 | 2.73E-10 | 1.77E-07 | 7.08E-01 | -4.98E-01 | -1.41E+00 |
| ENSG000001 | ABCD3 | 3.94E-10 | 2.44E-07 | 8.51E-01 | -2.34E-01 | -1.18E+00 |
| ENSG000002 | ATXN1L | 4.96E-10 | 2.95E-07 | 8.03E-01 | -3.16E-01 | -1.24E+00 |
| ENSG00000C | SLC39A9 | 6.28E-10 | 3.58E-07 | 6.83E-01 | -5.51E-01 | -1.46E+00 |
| ENSG000001 | PAMR1 | 1.19E-09 | 6.50E-07 | 2.02E+00 | 1.01E+00 | 2.02E+00 |
| ENSG000001 | LRRC58 | 2.59E-09 | 1.37E-06 | 7.39E-01 | -4.35E-01 | -1.35E+00 |
| ENSG000001 | PELI1 | 2.92E-09 | 1.48E-06 | 7.72E-01 | -3.74E-01 | -1.30E+00 |
| ENSG00000C | TAF2 | 4.08E-09 | 2.00E-06 | 8.04E-01 | -3.14E-01 | -1.24E+00 |
| ENSG000001 | LATS2 | 4.28E-09 | 2.02E-06 | 1.22E+00 | 2.90E-01 | 1.22E+00 |
| ENSG000001 | PERP | 4.39E-09 | 2.02E-06 | 7.35E-01 | -4.44E-01 | -1.36E+00 |
| ENSG000001 | FUT4 | 6.58E-09 | 2.93E-06 | 6.68E-01 | -5.83E-01 | -1.50E+00 |
| ENSG000001 | EPHA1 | 7.08E-09 | 3.06E-06 | 5.85E-01 | -7.75E-01 | -1.71E+00 |
| ENSG000001 | DDAH1 | 7.38E-09 | 3.10E-06 | 5.65E-01 | -8.23E-01 | -1.77E+00 |
| ENSG000001 | MICALL1 | 1.20E-08 | 4.87E-06 | 1.23E+00 | 2.95E-01 | 1.23E+00 |
| ENSG00000C | PTGS2 | 1.35E-08 | 5.35E-06 | 7.91E-01 | -3.37E-01 | -1.26E+00 |
| ENSG000001 | RAP2C | 1.47E-08 | 5.53E-06 | 8.18E-01 | -2.90E-01 | -1.22E+00 |
| ENSG000001 | PRR14L | 1.47E-08 | 5.53E-06 | 8.00E-01 | -3.22E-01 | -1.25E+00 |
| ENSG000001 | NMD3 | 1.77E-08 | 6.44E-06 | 7.63E-01 | -3.89E-01 | -1.31E+00 |
| ENSG000001 | MBOAT2 | 1.81E-08 | 6.44E-06 | 7.92E-01 | -3.37E-01 | -1.26E+00 |
| ENSG00000C | MAPK6 | 1.89E-08 | 6.59E-06 | 6.62E-01 | -5.95E-01 | -1.51E+00 |
| ENSG000001 | MMP10 | 2.01E-08 | 6.83E-06 | 6.72E-01 | -5.74E-01 | -1.49E+00 |
| ENSG000001 | OPA1 | 2.07E-08 | 6.88E-06 | 8.26E-01 | -2.75E-01 | -1.21E+00 |
| ENSG000001 | LPCAT1 | 2.59E-08 | 8.41E-06 | 8.43E-01 | -2.46E-01 | -1.19E+00 |

| | | | | | | |
|---|---|---|---|---|---|---|
| ENSG000001 | TP53 | 3.24E-08 | 1.01E-05 | 1.24E+00 | 3.12E-01 | 1.24E+00 |
| ENSG000001 | NFIA | 3.26E-08 | 1.01E-05 | 1.18E+00 | 2.35E-01 | 1.18E+00 |
| ENSG000001 | ZNF426 | 3.55E-08 | 1.07E-05 | 7.53E-01 | -4.10E-01 | -1.33E+00 |
| ENSG000001 | GRB10 | 3.59E-08 | 1.07E-05 | 8.08E-01 | -3.08E-01 | -1.24E+00 |
| ENSG000001 | SERPINE1 | 5.41E-08 | 1.57E-05 | 7.32E-01 | -4.51E-01 | -1.37E+00 |
| ENSG000002 | NRAS | 6.25E-08 | 1.78E-05 | 7.57E-01 | -4.02E-01 | -1.32E+00 |
| ENSG000001 | CMTM4 | 6.96E-08 | 1.91E-05 | 1.26E+00 | 3.31E-01 | 1.26E+00 |
| ENSG000001 | HNRNPF | 7.04E-08 | 1.91E-05 | 1.12E+00 | 1.62E-01 | 1.12E+00 |
| ENSG000001 | CHST3 | 7.16E-08 | 1.91E-05 | 8.55E-01 | -2.25E-01 | -1.17E+00 |
| ENSG000001 | VAMP7 | 7.22E-08 | 1.91E-05 | 7.64E-01 | -3.88E-01 | -1.31E+00 |
| ENSG000001 | TAF13 | 1.00E-07 | 2.60E-05 | 7.59E-01 | -3.97E-01 | -1.32E+00 |
| ENSG00000C | OSTM1 | 1.21E-07 | 3.07E-05 | 8.36E-01 | -2.58E-01 | -1.20E+00 |
| ENSG000001 | ARRB2 | 1.23E-07 | 3.08E-05 | 6.37E-01 | -6.52E-01 | -1.57E+00 |
| ENSG000001 | PTK2 | 1.45E-07 | 3.56E-05 | 8.24E-01 | -2.79E-01 | -1.21E+00 |
| ENSG00000C | IBTK | 1.60E-07 | 3.87E-05 | 8.41E-01 | -2.50E-01 | -1.19E+00 |
| ENSG000001 | MAPRE1 | 1.66E-07 | 3.90E-05 | 1.13E+00 | 1.79E-01 | 1.13E+00 |
| ENSG00000C | ST6GALNAC2 | 1.67E-07 | 3.90E-05 | 1.23E+00 | 2.99E-01 | 1.23E+00 |
| ENSG00000C | NDE1 | 1.69E-07 | 3.90E-05 | 1.20E+00 | 2.58E-01 | 1.20E+00 |
| ENSG00000C | EPB41L4B | 1.88E-07 | 4.22E-05 | 7.11E-01 | -4.91E-01 | -1.41E+00 |
| ENSG00000C | NSFL1C | 1.89E-07 | 4.22E-05 | 7.59E-01 | -3.99E-01 | -1.32E+00 |
| ENSG000001 | SPR | 2.19E-07 | 4.80E-05 | 1.21E+00 | 2.71E-01 | 1.21E+00 |
| ENSG000001 | MFSD14A | 2.32E-07 | 4.98E-05 | 8.12E-01 | -3.01E-01 | -1.23E+00 |
| ENSG000001 | TM4SF1 | 2.34E-07 | 4.98E-05 | 8.54E-01 | -2.28E-01 | -1.17E+00 |
| ENSG000001 | FLNB | 2.44E-07 | 5.11E-05 | 8.53E-01 | -2.29E-01 | -1.17E+00 |
| ENSG000001 | PDLIM1 | 2.54E-07 | 5.24E-05 | 1.13E+00 | 1.78E-01 | 1.13E+00 |
| ENSG000001 | PPIP5K2 | 2.64E-07 | 5.38E-05 | 8.27E-01 | -2.73E-01 | -1.21E+00 |
| ENSG000001 | TCEA1 | 2.84E-07 | 5.64E-05 | 7.95E-01 | -3.31E-01 | -1.26E+00 |
| ENSG000001 | SLC13A3 | 2.85E-07 | 5.64E-05 | 6.90E-01 | -5.36E-01 | -1.45E+00 |
| ENSG000002 | ANXA8 | 3.20E-07 | 6.24E-05 | 1.07E+00 | 1.01E-01 | 1.07E+00 |
| ENSG000001 | SOCS6 | 3.32E-07 | 6.40E-05 | 8.51E-01 | -2.33E-01 | -1.18E+00 |
| ENSG000001 | TTC33 | 3.51E-07 | 6.67E-05 | 6.77E-01 | -5.63E-01 | -1.48E+00 |
| ENSG000001 | CDK6 | 4.01E-07 | 7.52E-05 | 7.72E-01 | -3.73E-01 | -1.29E+00 |
| ENSG000001 | CHMP7 | 4.12E-07 | 7.62E-05 | 7.84E-01 | -3.52E-01 | -1.28E+00 |
| ENSG000001 | RSL24D1 | 4.54E-07 | 8.29E-05 | 1.24E+00 | 3.06E-01 | 1.24E+00 |
| ENSG000001 | DERL1 | 4.83E-07 | 8.64E-05 | 8.81E-01 | -1.83E-01 | -1.14E+00 |
| ENSG000001 | YTHDF2 | 4.85E-07 | 8.64E-05 | 8.79E-01 | -1.86E-01 | -1.14E+00 |
| ENSG000001 | KPNB1 | 4.97E-07 | 8.74E-05 | 8.48E-01 | -2.38E-01 | -1.18E+00 |
| ENSG000001 | G0S2 | 5.61E-07 | 9.65E-05 | 7.77E-01 | -3.64E-01 | -1.29E+00 |
| ENSG000001 | RAB5A | 5.62E-07 | 9.65E-05 | 8.11E-01 | -3.02E-01 | -1.23E+00 |
| ENSG000001 | CHRNA1 | 5.81E-07 | 9.85E-05 | 6.28E-01 | -6.70E-01 | -1.59E+00 |
| ENSG000001 | PDHX | 5.87E-07 | 9.85E-05 | 7.36E-01 | -4.42E-01 | -1.36E+00 |
| ENSG000001 | RNF7 | 6.92E-07 | 1.15E-04 | 1.22E+00 | 2.83E-01 | 1.22E+00 |
| ENSG000001 | TMEM263 | 7.08E-07 | 1.16E-04 | 8.10E-01 | -3.04E-01 | -1.23E+00 |
| ENSG000001 | HADH | 7.28E-07 | 1.18E-04 | 1.26E+00 | 3.30E-01 | 1.26E+00 |
| ENSG000001 | SEPTIN8 | 7.64E-07 | 1.22E-04 | 1.16E+00 | 2.11E-01 | 1.16E+00 |

| | | | | | | |
|---|---|---|---|---|---|---|
| ENSG00000C | NFYA | 8.40E-07 | 1.33E-04 | 8.33E-01 | -2.63E-01 | -1.20E+00 |
| ENSG000001 | RBM3 | 8.86E-07 | 1.39E-04 | 8.72E-01 | -1.98E-01 | -1.15E+00 |
| ENSG000001 | KATNAL1 | 9.23E-07 | 1.43E-04 | 7.57E-01 | -4.02E-01 | -1.32E+00 |
| ENSG00000C | DNAAF9 | 1.02E-06 | 1.56E-04 | 1.21E+00 | 2.71E-01 | 1.21E+00 |
| ENSG000001 | HBP1 | 1.03E-06 | 1.56E-04 | 7.79E-01 | -3.61E-01 | -1.28E+00 |
| ENSG000001 | TRAF6 | 1.12E-06 | 1.67E-04 | 8.03E-01 | -3.17E-01 | -1.25E+00 |
| ENSG000002 | TMED7-TICA | 1.22E-06 | 1.81E-04 | 8.19E-01 | -2.89E-01 | -1.22E+00 |
| ENSG000002 | SMIM7 | 1.28E-06 | 1.89E-04 | 1.19E+00 | 2.49E-01 | 1.19E+00 |
| ENSG000001 | GNAQ | 1.35E-06 | 1.97E-04 | 7.83E-01 | -3.53E-01 | -1.28E+00 |
| ENSG000001 | SPNS1 | 1.40E-06 | 2.00E-04 | 6.88E-01 | -5.40E-01 | -1.45E+00 |
| ENSG000001 | AKAP1 | 1.40E-06 | 2.00E-04 | 8.49E-01 | -2.35E-01 | -1.18E+00 |
| ENSG000001 | DICER1 | 1.47E-06 | 2.08E-04 | 7.88E-01 | -3.45E-01 | -1.27E+00 |
| ENSG000001 | IREB2 | 1.55E-06 | 2.16E-04 | 7.88E-01 | -3.44E-01 | -1.27E+00 |
| ENSG000001 | BACH2 | 1.61E-06 | 2.23E-04 | 1.41E+00 | 4.96E-01 | 1.41E+00 |
| ENSG000001 | KHNYN | 1.67E-06 | 2.29E-04 | 8.07E-01 | -3.09E-01 | -1.24E+00 |
| ENSG000001 | BTG2 | 1.69E-06 | 2.29E-04 | 1.25E+00 | 3.21E-01 | 1.25E+00 |
| ENSG000001 | AFF1 | 1.84E-06 | 2.46E-04 | 7.72E-01 | -3.74E-01 | -1.30E+00 |
| ENSG000001 | ZNF33B | 1.85E-06 | 2.46E-04 | 7.65E-01 | -3.87E-01 | -1.31E+00 |
| ENSG000001 | PLRG1 | 1.88E-06 | 2.48E-04 | 7.81E-01 | -3.57E-01 | -1.28E+00 |
| ENSG000001 | ANKRD46 | 1.91E-06 | 2.50E-04 | 7.56E-01 | -4.04E-01 | -1.32E+00 |
| ENSG000001 | CXCL14 | 1.95E-06 | 2.52E-04 | 1.39E+00 | 4.72E-01 | 1.39E+00 |
| ENSG000001 | AHCYL2 | 1.99E-06 | 2.55E-04 | 7.82E-01 | -3.55E-01 | -1.28E+00 |
| ENSG000001 | NHLRC3 | 2.00E-06 | 2.55E-04 | 7.44E-01 | -4.27E-01 | -1.34E+00 |
| ENSG00000C | PCBP4 | 2.15E-06 | 2.72E-04 | 1.29E+00 | 3.63E-01 | 1.29E+00 |
| ENSG000001 | NSG1 | 2.18E-06 | 2.73E-04 | 1.20E+00 | 2.62E-01 | 1.20E+00 |
| ENSG00000C | OSBPL3 | 2.31E-06 | 2.86E-04 | 8.48E-01 | -2.38E-01 | -1.18E+00 |
| ENSG000001 | HHLA3 | 2.34E-06 | 2.87E-04 | 1.41E+00 | 4.92E-01 | 1.41E+00 |
| ENSG000001 | MTA2 | 2.49E-06 | 3.03E-04 | 1.12E+00 | 1.64E-01 | 1.12E+00 |
| ENSG000001 | RPUSD4 | 2.50E-06 | 3.03E-04 | 8.32E-01 | -2.66E-01 | -1.20E+00 |
| ENSG000001 | FBXO21 | 2.53E-06 | 3.03E-04 | 1.13E+00 | 1.80E-01 | 1.13E+00 |
| ENSG000001 | TENT4A | 2.60E-06 | 3.08E-04 | 8.56E-01 | -2.24E-01 | -1.17E+00 |
| ENSG000001 | MYL9 | 2.73E-06 | 3.21E-04 | 1.34E+00 | 4.27E-01 | 1.34E+00 |
| ENSG000001 | RDH10 | 2.81E-06 | 3.28E-04 | 7.64E-01 | -3.89E-01 | -1.31E+00 |
| ENSG000001 | SEC24A | 2.88E-06 | 3.34E-04 | 8.22E-01 | -2.83E-01 | -1.22E+00 |
| ENSG000001 | NACC2 | 3.08E-06 | 3.54E-04 | 1.30E+00 | 3.80E-01 | 1.30E+00 |
| ENSG000001 | CCN1 | 3.24E-06 | 3.69E-04 | 7.74E-01 | -3.70E-01 | -1.29E+00 |
| ENSG000001 | BTF3L4 | 3.32E-06 | 3.75E-04 | 8.30E-01 | -2.69E-01 | -1.20E+00 |
| ENSG000001 | VDAC2 | 3.34E-06 | 3.75E-04 | 6.26E-01 | -6.75E-01 | -1.60E+00 |
| ENSG000001 | TACC1 | 3.61E-06 | 4.02E-04 | 8.51E-01 | -2.33E-01 | -1.18E+00 |
| ENSG000001 | THSD1 | 3.68E-06 | 4.06E-04 | 1.22E+00 | 2.91E-01 | 1.22E+00 |
| ENSG000001 | GXYLT1 | 3.70E-06 | 4.06E-04 | 7.89E-01 | -3.42E-01 | -1.27E+00 |
| ENSG000001 | MALL | 3.73E-06 | 4.06E-04 | 7.62E-01 | -3.92E-01 | -1.31E+00 |
| ENSG000001 | UGP2 | 3.86E-06 | 4.16E-04 | 8.34E-01 | -2.62E-01 | -1.20E+00 |
| ENSG000001 | EIF2S1 | 3.96E-06 | 4.23E-04 | 8.76E-01 | -1.90E-01 | -1.14E+00 |
| ENSG00000C | EPN3 | 3.99E-06 | 4.23E-04 | 1.26E+00 | 3.36E-01 | 1.26E+00 |

| | | | | | | |
|---|---|---|---|---|---|---|
| ENSG000001 | TMEM245 | 4.02E-06 | 4.23E-04 | 1.20E+00 | 2.57E-01 | 1.20E+00 |
| ENSG000001 | TRMT10A | 4.04E-06 | 4.23E-04 | 7.89E-01 | -3.42E-01 | -1.27E+00 |
| ENSG000001 | FNDC3A | 4.24E-06 | 4.41E-04 | 8.11E-01 | -3.03E-01 | -1.23E+00 |
| ENSG00000C | ZZEF1 | 4.40E-06 | 4.55E-04 | 8.68E-01 | -2.04E-01 | -1.15E+00 |
| ENSG000001 | PIK3R3 | 4.44E-06 | 4.55E-04 | 1.19E+00 | 2.50E-01 | 1.19E+00 |
| ENSG000001 | TVP23B | 4.49E-06 | 4.55E-04 | 8.19E-01 | -2.88E-01 | -1.22E+00 |
| ENSG000001 | SKP2 | 4.50E-06 | 4.55E-04 | 1.25E+00 | 3.27E-01 | 1.25E+00 |
| ENSG00000C | JARID2 | 4.95E-06 | 4.97E-04 | 8.38E-01 | -2.54E-01 | -1.19E+00 |
| ENSG000001 | CCDC68 | 5.11E-06 | 5.10E-04 | 7.90E-01 | -3.40E-01 | -1.27E+00 |
| ENSG000001 | POLR3A | 5.24E-06 | 5.18E-04 | 8.80E-01 | -1.84E-01 | -1.14E+00 |
| ENSG000001 | SERP1 | 5.45E-06 | 5.36E-04 | 8.84E-01 | -1.78E-01 | -1.13E+00 |
| ENSG00000C | SESN1 | 5.55E-06 | 5.42E-04 | 1.27E+00 | 3.40E-01 | 1.27E+00 |
| ENSG000001 | GSDME | 5.70E-06 | 5.53E-04 | 1.13E+00 | 1.71E-01 | 1.13E+00 |
| ENSG000001 | TNFRSF21 | 5.84E-06 | 5.63E-04 | 1.12E+00 | 1.64E-01 | 1.12E+00 |
| ENSG000001 | EDEM1 | 6.07E-06 | 5.80E-04 | 7.92E-01 | -3.36E-01 | -1.26E+00 |
| ENSG000001 | MAP3K2 | 6.25E-06 | 5.94E-04 | 7.90E-01 | -3.40E-01 | -1.27E+00 |
| ENSG00000C | CROT | 6.73E-06 | 6.35E-04 | 1.22E+00 | 2.87E-01 | 1.22E+00 |
| ENSG000001 | B3GLCT | 6.88E-06 | 6.45E-04 | 7.83E-01 | -3.53E-01 | -1.28E+00 |
| ENSG000001 | PANX1 | 7.03E-06 | 6.55E-04 | 8.81E-01 | -1.83E-01 | -1.14E+00 |
| ENSG000001 | S1PR1 | 7.11E-06 | 6.58E-04 | 6.35E-01 | -6.55E-01 | -1.58E+00 |
| ENSG000001 | LONRF1 | 7.26E-06 | 6.68E-04 | 8.04E-01 | -3.15E-01 | -1.24E+00 |
| ENSG000002 | ZCCHC3 | 7.33E-06 | 6.70E-04 | 1.21E+00 | 2.80E-01 | 1.21E+00 |
| ENSG00000C | CDK14 | 7.52E-06 | 6.83E-04 | 7.20E-01 | -4.74E-01 | -1.39E+00 |
| ENSG000001 | ARID1A | 7.76E-06 | 6.98E-04 | 8.53E-01 | -2.30E-01 | -1.17E+00 |
| ENSG000001 | ENOPH1 | 7.78E-06 | 6.98E-04 | 1.13E+00 | 1.79E-01 | 1.13E+00 |
| ENSG000002 | SEC22B | 8.28E-06 | 7.38E-04 | 8.88E-01 | -1.72E-01 | -1.13E+00 |
| ENSG000001 | CAMK1D | 8.40E-06 | 7.44E-04 | 1.30E+00 | 3.76E-01 | 1.30E+00 |
| ENSG000001 | ZPLD1 | 8.58E-06 | 7.55E-04 | 6.80E-01 | -5.57E-01 | -1.47E+00 |
| ENSG000001 | MTPN | 8.90E-06 | 7.78E-04 | 8.65E-01 | -2.08E-01 | -1.16E+00 |
| ENSG00000C | ZZZ3 | 9.30E-06 | 8.08E-04 | 8.20E-01 | -2.87E-01 | -1.22E+00 |
| ENSG00000C | AP5M1 | 9.87E-06 | 8.53E-04 | 8.33E-01 | -2.64E-01 | -1.20E+00 |
| ENSG000002 | --- | 9.93E-06 | 8.53E-04 | 6.82E-01 | -5.52E-01 | -1.47E+00 |
| ENSG000001 | TLR4 | 1.00E-05 | 8.55E-04 | 7.10E-01 | -4.93E-01 | -1.41E+00 |
| ENSG000001 | SULT1B1 | 1.02E-05 | 8.62E-04 | 5.53E-01 | -8.54E-01 | -1.81E+00 |
| ENSG00000C | IFRD1 | 1.12E-05 | 9.44E-04 | 8.26E-01 | -2.77E-01 | -1.21E+00 |
| ENSG000001 | COL4A1 | 1.19E-05 | 9.99E-04 | 7.33E-01 | -4.49E-01 | -1.36E+00 |
| ENSG000001 | PORCN | 1.26E-05 | 1.05E-03 | 8.33E-01 | -2.63E-01 | -1.20E+00 |
| ENSG000001 | IGSF3 | 1.28E-05 | 1.06E-03 | 1.15E+00 | 2.06E-01 | 1.15E+00 |
| ENSG000002 | CTAGE8 | 1.33E-05 | 1.10E-03 | 7.41E-01 | -4.33E-01 | -1.35E+00 |
| ENSG00000C | SERPINB1 | 1.42E-05 | 1.16E-03 | 8.10E-01 | -3.04E-01 | -1.23E+00 |
| ENSG000001 | SLC6A2 | 1.42E-05 | 1.16E-03 | 2.35E+00 | 1.24E+00 | 2.35E+00 |
| ENSG000001 | LYSMD3 | 1.43E-05 | 1.16E-03 | 7.92E-01 | -3.37E-01 | -1.26E+00 |
| ENSG000002 | TIMM23B | 1.46E-05 | 1.17E-03 | 7.59E-01 | -3.98E-01 | -1.32E+00 |
| ENSG000001 | PSMB2 | 1.46E-05 | 1.17E-03 | 8.96E-01 | -1.59E-01 | -1.12E+00 |
| ENSG000001 | COPS8 | 1.47E-05 | 1.17E-03 | 8.21E-01 | -2.84E-01 | -1.22E+00 |

| | | | | | | |
|---|---|---|---|---|---|---|
| ENSG00000C | UBE2D1 | 1.48E-05 | 1.17E-03 | 8.39E-01 | -2.54E-01 | -1.19E+00 |
| ENSG000001 | DRAM1 | 1.53E-05 | 1.21E-03 | 1.24E+00 | 3.09E-01 | 1.24E+00 |
| ENSG000001 | NQO1 | 1.55E-05 | 1.21E-03 | 7.27E-01 | -4.60E-01 | -1.38E+00 |
| ENSG000001 | NDUFS2 | 1.57E-05 | 1.22E-03 | 7.66E-01 | -3.84E-01 | -1.30E+00 |
| ENSG000001 | FZD6 | 1.61E-05 | 1.24E-03 | 1.11E+00 | 1.48E-01 | 1.11E+00 |
| ENSG000001 | SUGCT | 1.62E-05 | 1.25E-03 | 1.26E+00 | 3.34E-01 | 1.26E+00 |
| ENSG000002 | INSYN2B | 1.63E-05 | 1.25E-03 | 5.24E-01 | -9.31E-01 | -1.91E+00 |
| ENSG000001 | GLS | 1.64E-05 | 1.25E-03 | 8.90E-01 | -1.68E-01 | -1.12E+00 |
| ENSG000001 | TXNDC12 | 1.66E-05 | 1.26E-03 | 8.32E-01 | -2.65E-01 | -1.20E+00 |
| ENSG000001 | DNAJC3 | 1.70E-05 | 1.28E-03 | 8.56E-01 | -2.24E-01 | -1.17E+00 |
| ENSG000002 | LINC02660 | 1.71E-05 | 1.28E-03 | 1.45E+00 | 5.37E-01 | 1.45E+00 |
| ENSG000001 | ULBP2 | 1.72E-05 | 1.28E-03 | 8.08E-01 | -3.08E-01 | -1.24E+00 |
| ENSG00000C | MAST2 | 1.73E-05 | 1.28E-03 | 1.12E+00 | 1.66E-01 | 1.12E+00 |
| ENSG000001 | TBL1XR1 | 1.74E-05 | 1.28E-03 | 8.50E-01 | -2.34E-01 | -1.18E+00 |
| ENSG000001 | ADAMTS15 | 1.74E-05 | 1.28E-03 | 1.49E+00 | 5.74E-01 | 1.49E+00 |
| ENSG000001 | PSAT1 | 1.75E-05 | 1.28E-03 | 1.34E+00 | 4.27E-01 | 1.34E+00 |
| ENSG000002 | DENND1B | 1.79E-05 | 1.30E-03 | 7.64E-01 | -3.88E-01 | -1.31E+00 |
| ENSG000002 | ARFGAP3 | 1.81E-05 | 1.31E-03 | 8.68E-01 | -2.04E-01 | -1.15E+00 |
| ENSG000001 | RPL34 | 1.85E-05 | 1.33E-03 | 1.17E+00 | 2.27E-01 | 1.17E+00 |
| ENSG000001 | CYB5R1 | 1.86E-05 | 1.33E-03 | 1.18E+00 | 2.36E-01 | 1.18E+00 |
| ENSG000001 | TGFA | 1.89E-05 | 1.35E-03 | 7.39E-01 | -4.35E-01 | -1.35E+00 |
| ENSG000001 | GTF2F2 | 1.96E-05 | 1.39E-03 | 7.91E-01 | -3.39E-01 | -1.26E+00 |
| ENSG000001 | CPA4 | 1.96E-05 | 1.39E-03 | 1.37E+00 | 4.57E-01 | 1.37E+00 |
| ENSG000001 | TMEM41B | 2.00E-05 | 1.41E-03 | 8.66E-01 | -2.07E-01 | -1.15E+00 |
| ENSG00000C | NRDC | 2.02E-05 | 1.41E-03 | 9.24E-01 | -1.13E-01 | -1.08E+00 |
| ENSG000001 | ERO1A | 2.13E-05 | 1.48E-03 | 8.00E-01 | -3.23E-01 | -1.25E+00 |
| ENSG000001 | GPX8 | 2.20E-05 | 1.52E-03 | 7.71E-01 | -3.75E-01 | -1.30E+00 |
| ENSG000001 | TTC39B | 2.41E-05 | 1.66E-03 | 1.26E+00 | 3.39E-01 | 1.26E+00 |
| ENSG000001 | SP110 | 2.48E-05 | 1.70E-03 | 8.36E-01 | -2.59E-01 | -1.20E+00 |
| ENSG000001 | TRIM25 | 2.66E-05 | 1.82E-03 | 1.12E+00 | 1.68E-01 | 1.12E+00 |
| ENSG00000C | LAMA3 | 2.71E-05 | 1.84E-03 | 8.50E-01 | -2.35E-01 | -1.18E+00 |
| ENSG000001 | FBXO45 | 2.72E-05 | 1.84E-03 | 7.90E-01 | -3.40E-01 | -1.27E+00 |
| ENSG000001 | EIF3L | 2.95E-05 | 1.98E-03 | 1.12E+00 | 1.60E-01 | 1.12E+00 |
| ENSG000001 | GRAMD2A | 3.00E-05 | 2.01E-03 | 1.63E+00 | 7.06E-01 | 1.63E+00 |
| ENSG000001 | ARNT | 3.01E-05 | 2.01E-03 | 8.61E-01 | -2.15E-01 | -1.16E+00 |
| ENSG000001 | ZNF121 | 3.03E-05 | 2.01E-03 | 8.56E-01 | -2.25E-01 | -1.17E+00 |
| ENSG000001 | CCDC82 | 3.04E-05 | 2.01E-03 | 7.74E-01 | -3.69E-01 | -1.29E+00 |
| ENSG000001 | FRK | 3.09E-05 | 2.03E-03 | 7.44E-01 | -4.27E-01 | -1.34E+00 |
| ENSG00000C | OTUB2 | 3.16E-05 | 2.07E-03 | 7.22E-01 | -4.70E-01 | -1.38E+00 |
| ENSG000001 | PLCB4 | 3.19E-05 | 2.08E-03 | 7.94E-01 | -3.32E-01 | -1.26E+00 |
| ENSG000001 | RAB31 | 3.31E-05 | 2.14E-03 | 9.06E-01 | -1.43E-01 | -1.10E+00 |
| ENSG000001 | GPAM | 3.40E-05 | 2.19E-03 | 8.28E-01 | -2.73E-01 | -1.21E+00 |
| ENSG000001 | DYNC1LI2 | 3.40E-05 | 2.19E-03 | 9.10E-01 | -1.36E-01 | -1.10E+00 |
| ENSG000001 | NUDT15 | 3.61E-05 | 2.30E-03 | 8.44E-01 | -2.45E-01 | -1.18E+00 |
| ENSG000001 | BZW2 | 3.61E-05 | 2.30E-03 | 1.15E+00 | 2.02E-01 | 1.15E+00 |

| | | | | | | |
|---|---|---|---|---|---|---|
| ENSG000001 | NFKB1 | 3.66E-05 | 2.32E-03 | 1.13E+00 | 1.78E-01 | 1.13E+00 |
| ENSG000001 | PRRG4 | 3.71E-05 | 2.34E-03 | 1.32E+00 | 3.96E-01 | 1.32E+00 |
| ENSG00000C | MYH16 | 3.72E-05 | 2.34E-03 | 6.93E-01 | -5.30E-01 | -1.44E+00 |
| ENSG000001 | CSK | 3.79E-05 | 2.37E-03 | 8.43E-01 | -2.47E-01 | -1.19E+00 |
| ENSG00000C | STRADB | 3.86E-05 | 2.40E-03 | 1.19E+00 | 2.54E-01 | 1.19E+00 |
| ENSG000001 | NEDD9 | 3.89E-05 | 2.41E-03 | 1.39E+00 | 4.74E-01 | 1.39E+00 |
| ENSG000001 | TAF9B | 3.91E-05 | 2.41E-03 | 8.17E-01 | -2.91E-01 | -1.22E+00 |
| ENSG000001 | EPGN | 4.00E-05 | 2.46E-03 | 6.45E-01 | -6.32E-01 | -1.55E+00 |
| ENSG000001 | FNBP1L | 4.21E-05 | 2.57E-03 | 7.94E-01 | -3.33E-01 | -1.26E+00 |
| ENSG000002 | RN7SL1 | 4.22E-05 | 2.57E-03 | 4.29E+00 | 2.10E+00 | 4.29E+00 |
| ENSG000001 | ENTPD7 | 4.24E-05 | 2.57E-03 | 1.21E+00 | 2.76E-01 | 1.21E+00 |
| ENSG000001 | CADM1 | 4.33E-05 | 2.62E-03 | 8.04E-01 | -3.14E-01 | -1.24E+00 |
| ENSG00000C | CNN2 | 4.46E-05 | 2.68E-03 | 1.13E+00 | 1.76E-01 | 1.13E+00 |
| ENSG000001 | KCNJ5 | 4.84E-05 | 2.89E-03 | 1.63E+00 | 7.03E-01 | 1.63E+00 |
| ENSG000001 | BLOC1S2 | 4.85E-05 | 2.89E-03 | 1.12E+00 | 1.66E-01 | 1.12E+00 |
| ENSG000001 | TMEM254 | 4.87E-05 | 2.89E-03 | 1.19E+00 | 2.53E-01 | 1.19E+00 |
| ENSG000001 | POPDC3 | 5.30E-05 | 3.14E-03 | 7.49E-01 | -4.16E-01 | -1.33E+00 |
| ENSG000001 | TNFRSF10D | 5.42E-05 | 3.19E-03 | 1.13E+00 | 1.80E-01 | 1.13E+00 |
| ENSG000001 | PIWIL4 | 5.57E-05 | 3.27E-03 | 5.44E-01 | -8.78E-01 | -1.84E+00 |
| ENSG00000C | TM7SF3 | 5.62E-05 | 3.28E-03 | 1.14E+00 | 1.84E-01 | 1.14E+00 |
| ENSG00000C | DGKG | 5.72E-05 | 3.33E-03 | 5.98E-01 | -7.42E-01 | -1.67E+00 |
| ENSG000001 | COQ8A | 5.81E-05 | 3.36E-03 | 1.34E+00 | 4.25E-01 | 1.34E+00 |
| ENSG000001 | BCL7A | 5.84E-05 | 3.36E-03 | 1.26E+00 | 3.32E-01 | 1.26E+00 |
| ENSG000001 | ACER2 | 5.84E-05 | 3.36E-03 | 1.24E+00 | 3.11E-01 | 1.24E+00 |
| ENSG000001 | PRICKLE2 | 5.90E-05 | 3.38E-03 | 1.17E+00 | 2.27E-01 | 1.17E+00 |
| ENSG000001 | ALDH1B1 | 6.05E-05 | 3.45E-03 | 1.18E+00 | 2.39E-01 | 1.18E+00 |
| ENSG000001 | ABHD4 | 6.16E-05 | 3.50E-03 | 1.20E+00 | 2.58E-01 | 1.20E+00 |
| ENSG00000C | YBX3 | 6.19E-05 | 3.50E-03 | 1.09E+00 | 1.19E-01 | 1.09E+00 |
| ENSG000001 | DCP2 | 6.27E-05 | 3.53E-03 | 8.30E-01 | -2.69E-01 | -1.20E+00 |
| ENSG00000C | NTN4 | 6.39E-05 | 3.59E-03 | 8.09E-01 | -3.06E-01 | -1.24E+00 |
| ENSG00000C | ME1 | 6.45E-05 | 3.61E-03 | 8.62E-01 | -2.14E-01 | -1.16E+00 |
| ENSG000002 | LINC00294 | 6.49E-05 | 3.62E-03 | 1.17E+00 | 2.32E-01 | 1.17E+00 |
| ENSG000002 | RPL13AP20 | 6.52E-05 | 3.62E-03 | 6.36E-01 | -6.52E-01 | -1.57E+00 |
| ENSG000001 | CCDC93 | 7.03E-05 | 3.89E-03 | 8.80E-01 | -1.85E-01 | -1.14E+00 |
| ENSG000001 | ARHGEF10 | 7.20E-05 | 3.96E-03 | 8.96E-01 | -1.58E-01 | -1.12E+00 |
| ENSG000001 | PTP4A2 | 7.31E-05 | 3.99E-03 | 8.14E-01 | -2.97E-01 | -1.23E+00 |
| ENSG000001 | SDC1 | 7.33E-05 | 3.99E-03 | 7.50E-01 | -4.16E-01 | -1.33E+00 |
| ENSG000001 | TPRG1 | 7.33E-05 | 3.99E-03 | 7.57E-01 | -4.02E-01 | -1.32E+00 |
| ENSG000001 | CD46 | 7.38E-05 | 3.99E-03 | 8.74E-01 | -1.94E-01 | -1.14E+00 |
| ENSG000001 | C21orf91 | 7.40E-05 | 3.99E-03 | 1.23E+00 | 2.93E-01 | 1.23E+00 |
| ENSG000002 | SLC7A11-AS1 | 7.50E-05 | ? | 4.24E-01 | -1.24E+00 | -2.36E+00 |
| ENSG00000C | CYFIP2 | 7.84E-05 | 4.22E-03 | 1.24E+00 | 3.12E-01 | 1.24E+00 |
| ENSG000001 | TMEM156 | 8.05E-05 | 4.32E-03 | 8.06E-01 | -3.12E-01 | -1.24E+00 |
| ENSG000001 | SOWAHB | 8.15E-05 | 4.35E-03 | 1.43E+00 | 5.15E-01 | 1.43E+00 |
| ENSG000001 | RAB32 | 8.40E-05 | 4.47E-03 | 1.26E+00 | 3.34E-01 | 1.26E+00 |

| | | | | | | |
|---|---|---|---|---|---|---|
| ENSG000001 | COPZ1 | 8.44E-05 | 4.47E-03 | 8.50E-01 | -2.34E-01 | -1.18E+00 |
| ENSG000001 | RPL7A | 8.54E-05 | 4.51E-03 | 1.14E+00 | 1.87E-01 | 1.14E+00 |
| ENSG000001 | ABCA12 | 8.85E-05 | 4.65E-03 | 1.19E+00 | 2.56E-01 | 1.19E+00 |
| ENSG000001 | ABAT | 8.87E-05 | 4.65E-03 | 1.25E+00 | 3.21E-01 | 1.25E+00 |
| ENSG000001 | PAFAH1B2 | 8.91E-05 | 4.65E-03 | 8.45E-01 | -2.43E-01 | -1.18E+00 |
| ENSG000002 | FTX | 9.04E-05 | 4.70E-03 | 8.19E-01 | -2.88E-01 | -1.22E+00 |
| ENSG00000C | NUP160 | 9.17E-05 | 4.76E-03 | 1.12E+00 | 1.64E-01 | 1.12E+00 |
| ENSG000001 | NAV2 | 9.39E-05 | 4.85E-03 | 1.16E+00 | 2.17E-01 | 1.16E+00 |
| ENSG000001 | SERPINB6 | 9.43E-05 | 4.85E-03 | 8.70E-01 | -2.01E-01 | -1.15E+00 |
| ENSG000001 | PRSS12 | 9.67E-05 | 4.96E-03 | 1.16E+00 | 2.15E-01 | 1.16E+00 |
| ENSG000002 | LINC02783 | 9.82E-05 | 5.02E-03 | 6.71E-01 | -5.76E-01 | -1.49E+00 |
| ENSG000001 | GGA3 | 9.90E-05 | 5.04E-03 | 8.33E-01 | -2.64E-01 | -1.20E+00 |
| ENSG000001 | STC2 | 9.99E-05 | 5.06E-03 | 8.13E-01 | -2.99E-01 | -1.23E+00 |
| ENSG000001 | SLC38A1 | 1.00E-04 | 5.06E-03 | 8.96E-01 | -1.59E-01 | -1.12E+00 |
| ENSG00000C | AIFM2 | 1.01E-04 | 5.06E-03 | 7.83E-01 | -3.52E-01 | -1.28E+00 |
| ENSG000001 | VEGFA | 1.01E-04 | 5.06E-03 | 7.47E-01 | -4.22E-01 | -1.34E+00 |
| ENSG000001 | NHEJ1 | 1.01E-04 | 5.06E-03 | 8.31E-01 | -2.67E-01 | -1.20E+00 |
| ENSG00000C | IKZF5 | 1.03E-04 | 5.11E-03 | 8.31E-01 | -2.68E-01 | -1.20E+00 |
| ENSG00000C | PPP1R12B | 1.03E-04 | 5.11E-03 | 1.41E+00 | 4.97E-01 | 1.41E+00 |
| ENSG000001 | TPCN1 | 1.05E-04 | 5.21E-03 | 1.14E+00 | 1.92E-01 | 1.14E+00 |
| ENSG000001 | ARHGEF6 | 1.10E-04 | 5.43E-03 | 1.36E+00 | 4.47E-01 | 1.36E+00 |
| ENSG000001 | C6orf62 | 1.13E-04 | 5.54E-03 | 8.70E-01 | -2.01E-01 | -1.15E+00 |
| ENSG000001 | JPH1 | 1.20E-04 | 5.88E-03 | 1.25E+00 | 3.21E-01 | 1.25E+00 |
| ENSG000001 | PANK1 | 1.21E-04 | 5.93E-03 | 7.75E-01 | -3.68E-01 | -1.29E+00 |
| ENSG000001 | VAMP8 | 1.25E-04 | 6.11E-03 | 1.14E+00 | 1.85E-01 | 1.14E+00 |
| ENSG000001 | SESN3 | 1.28E-04 | 6.19E-03 | 1.30E+00 | 3.81E-01 | 1.30E+00 |
| ENSG00000C | HAUS4 | 1.28E-04 | 6.19E-03 | 1.39E+00 | 4.78E-01 | 1.39E+00 |
| ENSG000002 | --- | 1.31E-04 | 6.29E-03 | 2.07E+00 | 1.05E+00 | 2.07E+00 |
| ENSG000001 | SAMD9L | 1.32E-04 | 6.32E-03 | 6.78E-01 | -5.61E-01 | -1.48E+00 |
| ENSG000001 | ZDHHC9 | 1.36E-04 | 6.51E-03 | 1.14E+00 | 1.85E-01 | 1.14E+00 |
| ENSG000001 | FYTTD1 | 1.37E-04 | 6.53E-03 | 8.86E-01 | -1.75E-01 | -1.13E+00 |
| ENSG00000C | PRR11 | 1.39E-04 | 6.60E-03 | 7.99E-01 | -3.24E-01 | -1.25E+00 |
| ENSG000001 | POMK | 1.43E-04 | 6.79E-03 | 8.03E-01 | -3.17E-01 | -1.25E+00 |
| ENSG000001 | HNRNPA3 | 1.44E-04 | 6.81E-03 | 9.00E-01 | -1.53E-01 | -1.11E+00 |
| ENSG000001 | VDAC2P5 | 1.45E-04 | 6.81E-03 | 6.03E-01 | -7.29E-01 | -1.66E+00 |
| ENSG000001 | SESN2 | 1.46E-04 | 6.85E-03 | 1.22E+00 | 2.83E-01 | 1.22E+00 |
| ENSG000001 | HACD1 | 1.49E-04 | 6.93E-03 | 1.21E+00 | 2.81E-01 | 1.21E+00 |
| ENSG000002 | NMRAL2P | 1.49E-04 | 6.93E-03 | 6.68E-01 | -5.82E-01 | -1.50E+00 |
| ENSG000001 | RPS8 | 1.50E-04 | 6.93E-03 | 1.10E+00 | 1.33E-01 | 1.10E+00 |
| ENSG000001 | SLC30A5 | 1.50E-04 | 6.93E-03 | 8.79E-01 | -1.86E-01 | -1.14E+00 |
| ENSG000001 | DUSP6 | 1.52E-04 | 7.00E-03 | 8.46E-01 | -2.41E-01 | -1.18E+00 |
| ENSG00000C | GCLM | 1.54E-04 | 7.08E-03 | 8.31E-01 | -2.67E-01 | -1.20E+00 |
| ENSG000001 | AGFG1 | 1.58E-04 | 7.27E-03 | 8.92E-01 | -1.65E-01 | -1.12E+00 |
| ENSG000001 | SLX4IP | 1.61E-04 | 7.35E-03 | 8.35E-01 | -2.60E-01 | -1.20E+00 |
| ENSG000001 | NUDT11 | 1.63E-04 | 7.43E-03 | 1.19E+00 | 2.51E-01 | 1.19E+00 |

| | | | | | | |
|---|---|---|---|---|---|---|
| ENSG00000C | CBFB | 1.66E-04 | 7.53E-03 | 8.58E-01 | -2.21E-01 | -1.17E+00 |
| ENSG000001 | XIAP | 1.66E-04 | 7.53E-03 | 9.02E-01 | -1.49E-01 | -1.11E+00 |
| ENSG000001 | MAP2K5 | 1.79E-04 | 8.06E-03 | 8.20E-01 | -2.87E-01 | -1.22E+00 |
| ENSG000001 | CORO2B | 1.81E-04 | 8.16E-03 | 8.33E-01 | -2.64E-01 | -1.20E+00 |
| ENSG000001 | AFDN | 1.84E-04 | 8.25E-03 | 8.60E-01 | -2.18E-01 | -1.16E+00 |
| ENSG000001 | SELENON | 1.88E-04 | 8.41E-03 | 1.12E+00 | 1.68E-01 | 1.12E+00 |
| ENSG000001 | INTS4 | 1.89E-04 | 8.42E-03 | 8.45E-01 | -2.43E-01 | -1.18E+00 |
| ENSG000001 | POLH | 1.91E-04 | 8.48E-03 | 1.13E+00 | 1.71E-01 | 1.13E+00 |
| ENSG000001 | WNT4 | 1.94E-04 | 8.60E-03 | 1.60E+00 | 6.81E-01 | 1.60E+00 |
| ENSG000001 | HSF2BP | 2.02E-04 | 8.93E-03 | 1.19E+00 | 2.54E-01 | 1.19E+00 |
| ENSG000001 | RAB38 | 2.04E-04 | 8.96E-03 | 1.21E+00 | 2.80E-01 | 1.21E+00 |
| ENSG000001 | ZHX3 | 2.05E-04 | 9.01E-03 | 8.50E-01 | -2.34E-01 | -1.18E+00 |
| ENSG000001 | NRBP1 | 2.08E-04 | 9.08E-03 | 1.11E+00 | 1.50E-01 | 1.11E+00 |
| ENSG00000C | BZW1 | 2.09E-04 | 9.11E-03 | 8.81E-01 | -1.83E-01 | -1.14E+00 |
| ENSG000001 | LATS1 | 2.10E-04 | 9.11E-03 | 8.76E-01 | -1.91E-01 | -1.14E+00 |
| ENSG000001 | GBP1 | 2.18E-04 | 9.44E-03 | 1.36E+00 | 4.48E-01 | 1.36E+00 |
| ENSG000001 | CPT2 | 2.20E-04 | 9.48E-03 | 1.19E+00 | 2.56E-01 | 1.19E+00 |
| ENSG000001 | MTURN | 2.20E-04 | 9.48E-03 | 1.12E+00 | 1.57E-01 | 1.12E+00 |
| ENSG000001 | IL1A | 2.21E-04 | 9.48E-03 | 8.35E-01 | -2.61E-01 | -1.20E+00 |
| ENSG000001 | STK32A | 2.22E-04 | 9.50E-03 | 1.24E+00 | 3.16E-01 | 1.24E+00 |
| ENSG000001 | SGO2 | 2.23E-04 | 9.50E-03 | 7.89E-01 | -3.41E-01 | -1.27E+00 |
| ENSG000001 | RFX5 | 2.23E-04 | 9.50E-03 | 1.10E+00 | 1.44E-01 | 1.10E+00 |
| ENSG000001 | POMGNT2 | 2.26E-04 | 9.61E-03 | 1.30E+00 | 3.76E-01 | 1.30E+00 |
| ENSG000001 | SLC25A44 | 2.27E-04 | 9.61E-03 | 8.63E-01 | -2.13E-01 | -1.16E+00 |
| ENSG000001 | CFAP300 | 2.28E-04 | 9.61E-03 | 1.52E+00 | 6.08E-01 | 1.52E+00 |
| ENSG000001 | CEP20 | 2.32E-04 | 9.76E-03 | 8.49E-01 | -2.36E-01 | -1.18E+00 |
| ENSG000001 | C6orf120 | 2.34E-04 | 9.82E-03 | 1.16E+00 | 2.17E-01 | 1.16E+00 |
| ENSG000001 | WDR82 | 2.35E-04 | 9.84E-03 | 1.10E+00 | 1.39E-01 | 1.10E+00 |
| ENSG000001 | TBC1D14 | 2.37E-04 | 9.87E-03 | 1.12E+00 | 1.69E-01 | 1.12E+00 |
| ENSG000001 | OLR1 | 2.42E-04 | 1.00E-02 | 1.66E+00 | 7.28E-01 | 1.66E+00 |
| ENSG000001 | PGK1 | 2.44E-04 | 1.01E-02 | 8.50E-01 | -2.35E-01 | -1.18E+00 |
| ENSG000001 | RPS4X | 2.44E-04 | 1.01E-02 | 1.10E+00 | 1.31E-01 | 1.10E+00 |
| ENSG000001 | SH2D4A | 2.48E-04 | 1.02E-02 | 8.21E-01 | -2.84E-01 | -1.22E+00 |
| ENSG000001 | RWDD2B | 2.50E-04 | 1.03E-02 | 1.17E+00 | 2.22E-01 | 1.17E+00 |
| ENSG000001 | ZNF41 | 2.53E-04 | 1.03E-02 | 8.36E-01 | -2.58E-01 | -1.20E+00 |
| ENSG000001 | MFSD9 | 2.54E-04 | 1.04E-02 | 8.59E-01 | -2.19E-01 | -1.16E+00 |
| ENSG000001 | CDK4 | 2.58E-04 | 1.05E-02 | 1.21E+00 | 2.81E-01 | 1.21E+00 |
| ENSG000001 | EREG | 2.60E-04 | 1.06E-02 | 7.12E-01 | -4.90E-01 | -1.40E+00 |
| ENSG000001 | LCOR | 2.66E-04 | 1.08E-02 | 8.23E-01 | -2.81E-01 | -1.22E+00 |
| ENSG000001 | HEG1 | 2.71E-04 | 1.09E-02 | 1.20E+00 | 2.66E-01 | 1.20E+00 |
| ENSG000001 | SCD5 | 2.71E-04 | 1.09E-02 | 1.25E+00 | 3.27E-01 | 1.25E+00 |
| ENSG000001 | NIPA1 | 2.72E-04 | 1.09E-02 | 8.88E-01 | -1.71E-01 | -1.13E+00 |
| ENSG000001 | ASPH | 2.75E-04 | 1.10E-02 | 8.76E-01 | -1.90E-01 | -1.14E+00 |
| ENSG000001 | RPL38 | 2.78E-04 | 1.11E-02 | 1.08E+00 | 1.18E-01 | 1.08E+00 |
| ENSG000001 | SCEL | 2.80E-04 | 1.11E-02 | 8.51E-01 | -2.32E-01 | -1.17E+00 |

| | | | | | | |
|---|---|---|---|---|---|---|
| ENSG000001 | SLC35G2 | 2.81E-04 | 1.11E-02 | 8.13E-01 | -2.99E-01 | -1.23E+00 |
| ENSG000001 | TEX2 | 2.81E-04 | 1.11E-02 | 1.12E+00 | 1.62E-01 | 1.12E+00 |
| ENSG000001 | FZD1 | 2.84E-04 | 1.12E-02 | 1.22E+00 | 2.86E-01 | 1.22E+00 |
| ENSG000002 | IL10RB | 2.87E-04 | 1.13E-02 | 8.33E-01 | -2.63E-01 | -1.20E+00 |
| ENSG000002 | RN7SL2 | 2.90E-04 | 1.14E-02 | 3.73E+00 | 1.90E+00 | 3.73E+00 |
| ENSG000001 | PDGFB | 2.90E-04 | 1.14E-02 | 7.32E-01 | -4.50E-01 | -1.37E+00 |
| ENSG000001 | HIPK1 | 2.94E-04 | 1.15E-02 | 8.94E-01 | -1.62E-01 | -1.12E+00 |
| ENSG00000C | NUDC | 2.98E-04 | 1.16E-02 | 9.23E-01 | -1.15E-01 | -1.08E+00 |
| ENSG000001 | GAR1 | 2.99E-04 | 1.16E-02 | 8.32E-01 | -2.65E-01 | -1.20E+00 |
| ENSG000001 | BCL2L11 | 3.06E-04 | 1.19E-02 | 1.33E+00 | 4.14E-01 | 1.33E+00 |
| ENSG000001 | IMPA2 | 3.10E-04 | 1.20E-02 | 1.50E+00 | 5.82E-01 | 1.50E+00 |
| ENSG000001 | CDH2 | 3.11E-04 | 1.20E-02 | 7.99E-01 | -3.23E-01 | -1.25E+00 |
| ENSG000001 | B4GALT2 | 3.15E-04 | 1.21E-02 | 1.21E+00 | 2.78E-01 | 1.21E+00 |
| ENSG000001 | LARGE2 | 3.15E-04 | 1.21E-02 | 1.44E+00 | 5.24E-01 | 1.44E+00 |
| ENSG000001 | CEP128 | 3.16E-04 | 1.21E-02 | 7.32E-01 | -4.49E-01 | -1.37E+00 |
| ENSG000001 | MPDZ | 3.18E-04 | 1.21E-02 | 8.76E-01 | -1.90E-01 | -1.14E+00 |
| ENSG000001 | UBE3A | 3.26E-04 | 1.24E-02 | 8.84E-01 | -1.78E-01 | -1.13E+00 |
| ENSG000001 | GZF1 | 3.30E-04 | 1.25E-02 | 8.81E-01 | -1.83E-01 | -1.14E+00 |
| ENSG00000C | EXOC5 | 3.30E-04 | 1.25E-02 | 8.76E-01 | -1.91E-01 | -1.14E+00 |
| ENSG000001 | PTK7 | 3.34E-04 | 1.26E-02 | 1.18E+00 | 2.37E-01 | 1.18E+00 |
| ENSG00000C | MAP3K14 | 3.37E-04 | 1.27E-02 | 1.33E+00 | 4.12E-01 | 1.33E+00 |
| ENSG000002 | ERCC6 | 3.43E-04 | 1.29E-02 | 8.91E-01 | -1.66E-01 | -1.12E+00 |
| ENSG000001 | CEP135 | 3.45E-04 | 1.29E-02 | 8.14E-01 | -2.97E-01 | -1.23E+00 |
| ENSG000001 | ZEB1 | 3.46E-04 | 1.29E-02 | 7.63E-01 | -3.89E-01 | -1.31E+00 |
| ENSG000001 | SGMS1 | 3.47E-04 | 1.29E-02 | 8.85E-01 | -1.76E-01 | -1.13E+00 |
| ENSG000001 | ADGRF4 | 3.51E-04 | 1.30E-02 | 1.12E+00 | 1.69E-01 | 1.12E+00 |
| ENSG000001 | WEE1 | 3.51E-04 | 1.30E-02 | 1.31E+00 | 3.86E-01 | 1.31E+00 |
| ENSG000001 | SLC39A11 | 3.52E-04 | 1.30E-02 | 8.09E-01 | -3.06E-01 | -1.24E+00 |
| ENSG000001 | ZFAND1 | 3.54E-04 | 1.30E-02 | 8.25E-01 | -2.78E-01 | -1.21E+00 |
| ENSG000002 | --- | 3.54E-04 | ? | 1.06E-01 | -3.24E+00 | -9.46E+00 |
| ENSG00000C | TTC38 | 3.54E-04 | 1.30E-02 | 1.18E+00 | 2.36E-01 | 1.18E+00 |
| ENSG000001 | MFAP5 | 3.56E-04 | 1.31E-02 | 1.16E+00 | 2.09E-01 | 1.16E+00 |
| ENSG000001 | SLC11A2 | 3.61E-04 | 1.32E-02 | 1.12E+00 | 1.59E-01 | 1.12E+00 |
| ENSG000001 | EI24 | 3.62E-04 | 1.32E-02 | 1.08E+00 | 1.07E-01 | 1.08E+00 |
| ENSG000002 | MOCS2-DT | 3.66E-04 | 1.33E-02 | 1.28E+00 | 3.61E-01 | 1.28E+00 |
| ENSG000001 | TNKS1BP1 | 3.66E-04 | 1.33E-02 | 1.14E+00 | 1.90E-01 | 1.14E+00 |
| ENSG000002 | --- | 3.67E-04 | ? | 3.71E-02 | -4.75E+00 | -2.70E+01 |
| ENSG000001 | TNS3 | 3.70E-04 | 1.33E-02 | 8.67E-01 | -2.06E-01 | -1.15E+00 |
| ENSG000001 | USP8 | 3.71E-04 | 1.33E-02 | 8.68E-01 | -2.04E-01 | -1.15E+00 |
| ENSG000001 | RBBP5 | 3.71E-04 | 1.33E-02 | 8.84E-01 | -1.78E-01 | -1.13E+00 |
| ENSG000001 | ASNSD1 | 3.72E-04 | 1.33E-02 | 8.40E-01 | -2.52E-01 | -1.19E+00 |
| ENSG00000C | SEMA3A | 3.73E-04 | 1.33E-02 | 8.07E-01 | -3.10E-01 | -1.24E+00 |
| ENSG000001 | ODF2L | 3.73E-04 | 1.33E-02 | 8.18E-01 | -2.90E-01 | -1.22E+00 |
| ENSG000001 | COMMD6 | 3.74E-04 | 1.33E-02 | 1.14E+00 | 1.83E-01 | 1.14E+00 |
| ENSG000002 | --- | 3.75E-04 | 1.33E-02 | 7.92E-01 | -3.36E-01 | -1.26E+00 |

| | | | | | | |
|---|---|---|---|---|---|---|
| ENSG000001 | SP2 | 3.75E-04 | 1.33E-02 | 8.41E-01 | -2.50E-01 | -1.19E+00 |
| ENSG000001 | ECPAS | 3.79E-04 | 1.34E-02 | 9.30E-01 | -1.05E-01 | -1.08E+00 |
| ENSG000002 | PIP4K2B | 3.84E-04 | 1.35E-02 | 8.32E-01 | -2.66E-01 | -1.20E+00 |
| ENSG00000C | LDAF1 | 3.85E-04 | 1.35E-02 | 1.21E+00 | 2.76E-01 | 1.21E+00 |
| ENSG000001 | WNT9A | 3.85E-04 | 1.35E-02 | 7.68E-01 | -3.81E-01 | -1.30E+00 |
| ENSG00000C | ERO1B | 3.92E-04 | 1.37E-02 | 7.84E-01 | -3.52E-01 | -1.28E+00 |
| ENSG000002 | ZNF425 | 3.95E-04 | 1.38E-02 | 1.32E+00 | 3.99E-01 | 1.32E+00 |
| ENSG000001 | RAB4A | 4.00E-04 | 1.40E-02 | 1.11E+00 | 1.57E-01 | 1.11E+00 |
| ENSG000002 | PDE7A | 4.07E-04 | 1.42E-02 | 8.59E-01 | -2.19E-01 | -1.16E+00 |
| ENSG00000C | GANAB | 4.08E-04 | 1.42E-02 | 8.01E-01 | -3.21E-01 | -1.25E+00 |
| ENSG000001 | EXT2 | 4.10E-04 | 1.42E-02 | 1.15E+00 | 2.02E-01 | 1.15E+00 |
| ENSG000001 | PEX10 | 4.13E-04 | 1.43E-02 | 1.22E+00 | 2.81E-01 | 1.22E+00 |
| ENSG000002 | ZNF512 | 4.15E-04 | 1.43E-02 | 8.10E-01 | -3.05E-01 | -1.24E+00 |
| ENSG000001 | ORAI3 | 4.17E-04 | 1.43E-02 | 1.27E+00 | 3.48E-01 | 1.27E+00 |
| ENSG000001 | MIA2 | 4.20E-04 | 1.44E-02 | 8.60E-01 | -2.17E-01 | -1.16E+00 |
| ENSG000001 | ARPC5 | 4.24E-04 | 1.45E-02 | 9.18E-01 | -1.24E-01 | -1.09E+00 |
| ENSG00000C | TMEM38B | 4.27E-04 | 1.46E-02 | 7.88E-01 | -3.44E-01 | -1.27E+00 |
| ENSG000002 | LINC02693 | 4.28E-04 | 1.46E-02 | 8.87E-01 | -1.72E-01 | -1.13E+00 |
| ENSG000002 | RPSAP52 | 4.29E-04 | 1.46E-02 | 7.59E-01 | -3.98E-01 | -1.32E+00 |
| ENSG000002 | SYNRG | 4.33E-04 | 1.47E-02 | 1.15E+00 | 2.07E-01 | 1.15E+00 |
| ENSG000001 | GFPT1 | 4.35E-04 | 1.47E-02 | 8.75E-01 | -1.92E-01 | -1.14E+00 |
| ENSG000002 | FTCDNL1 | 4.37E-04 | ? | 1.92E+00 | 9.37E-01 | 1.92E+00 |
| ENSG000001 | SIPA1L2 | 4.38E-04 | 1.48E-02 | 1.19E+00 | 2.51E-01 | 1.19E+00 |
| ENSG000002 | LINC02210 | 4.40E-04 | 1.48E-02 | 8.30E-01 | -2.68E-01 | -1.20E+00 |
| ENSG000002 | FAM66B | 4.43E-04 | 1.49E-02 | 1.61E+00 | 6.83E-01 | 1.61E+00 |
| ENSG000001 | TENT5A | 4.44E-04 | 1.49E-02 | 7.70E-01 | -3.78E-01 | -1.30E+00 |
| ENSG000001 | PLA2R1 | 4.47E-04 | 1.49E-02 | 8.24E-01 | -2.80E-01 | -1.21E+00 |
| ENSG000001 | ATP5MC2 | 4.49E-04 | 1.50E-02 | 1.15E+00 | 2.06E-01 | 1.15E+00 |
| ENSG000001 | IL6ST | 4.53E-04 | 1.51E-02 | 8.67E-01 | -2.07E-01 | -1.15E+00 |
| ENSG000001 | PNPO | 4.59E-04 | 1.52E-02 | 1.12E+00 | 1.69E-01 | 1.12E+00 |
| ENSG000001 | FOSL1 | 4.65E-04 | 1.54E-02 | 8.04E-01 | -3.15E-01 | -1.24E+00 |
| ENSG000001 | CHODL | 4.68E-04 | ? | 2.19E+00 | 1.13E+00 | 2.19E+00 |
| ENSG000001 | STX12 | 4.69E-04 | 1.55E-02 | 1.11E+00 | 1.54E-01 | 1.11E+00 |
| ENSG000001 | MORF4L1 | 4.72E-04 | 1.55E-02 | 1.09E+00 | 1.23E-01 | 1.09E+00 |
| ENSG000001 | DNAJB14 | 4.73E-04 | 1.55E-02 | 8.49E-01 | -2.36E-01 | -1.18E+00 |
| ENSG000001 | SREK1IP1 | 4.76E-04 | 1.56E-02 | 8.73E-01 | -1.97E-01 | -1.15E+00 |
| ENSG000001 | EFNA5 | 4.77E-04 | 1.56E-02 | 8.58E-01 | -2.22E-01 | -1.17E+00 |
| ENSG00000C | LAMC2 | 4.79E-04 | 1.56E-02 | 9.08E-01 | -1.39E-01 | -1.10E+00 |
| ENSG000001 | CD1D | 4.85E-04 | 1.58E-02 | 1.34E+00 | 4.24E-01 | 1.34E+00 |
| ENSG000001 | NUS1 | 4.85E-04 | 1.58E-02 | 8.91E-01 | -1.66E-01 | -1.12E+00 |
| ENSG000001 | STRIP2 | 4.93E-04 | 1.60E-02 | 8.75E-01 | -1.93E-01 | -1.14E+00 |
| ENSG000001 | NEK1 | 4.96E-04 | 1.60E-02 | 8.95E-01 | -1.60E-01 | -1.12E+00 |
| ENSG00000C | THOC3 | 4.99E-04 | 1.61E-02 | 1.12E+00 | 1.57E-01 | 1.12E+00 |
| ENSG000001 | ALDOC | 5.03E-04 | 1.62E-02 | 1.23E+00 | 3.00E-01 | 1.23E+00 |
| ENSG000001 | TMX3 | 5.11E-04 | 1.64E-02 | 8.63E-01 | -2.13E-01 | -1.16E+00 |

| | | | | | | |
|---|---|---|---|---|---|---|
| ENSG000002 | SH3PXD2A-A | 5.12E-04 | 1.64E-02 | 6.89E-01 | -5.38E-01 | -1.45E+00 |
| ENSG00000C | CCNT2 | 5.14E-04 | 1.64E-02 | 8.51E-01 | -2.33E-01 | -1.18E+00 |
| ENSG000001 | AGGF1 | 5.15E-04 | 1.64E-02 | 8.54E-01 | -2.28E-01 | -1.17E+00 |
| ENSG000001 | RAP2A | 5.19E-04 | 1.64E-02 | 8.67E-01 | -2.06E-01 | -1.15E+00 |
| ENSG000002 | ZDHHC18 | 5.19E-04 | 1.64E-02 | 8.05E-01 | -3.13E-01 | -1.24E+00 |
| ENSG000001 | RPL13A | 5.19E-04 | 1.64E-02 | 1.15E+00 | 1.96E-01 | 1.15E+00 |
| ENSG000002 | --- | 5.21E-04 | ? | 6.18E+00 | 2.63E+00 | 6.18E+00 |
| ENSG00000C | SZRD1 | 5.34E-04 | 1.68E-02 | 1.11E+00 | 1.53E-01 | 1.11E+00 |
| ENSG000001 | PCGF3 | 5.35E-04 | 1.68E-02 | 8.93E-01 | -1.63E-01 | -1.12E+00 |
| ENSG000002 | TRIM59 | 5.35E-04 | 1.68E-02 | 8.44E-01 | -2.45E-01 | -1.18E+00 |
| ENSG00000C | FOXN3 | 5.37E-04 | 1.69E-02 | 8.69E-01 | -2.03E-01 | -1.15E+00 |
| ENSG000001 | CBX1 | 5.39E-04 | 1.69E-02 | 1.17E+00 | 2.26E-01 | 1.17E+00 |
| ENSG000001 | PTPRD | 5.41E-04 | 1.69E-02 | 1.22E+00 | 2.85E-01 | 1.22E+00 |
| ENSG000001 | GOSR2 | 5.54E-04 | 1.72E-02 | 9.06E-01 | -1.42E-01 | -1.10E+00 |
| ENSG000001 | CLOCK | 5.54E-04 | 1.72E-02 | 8.42E-01 | -2.48E-01 | -1.19E+00 |
| ENSG000001 | HSD17B7 | 5.56E-04 | 1.72E-02 | 1.13E+00 | 1.73E-01 | 1.13E+00 |
| ENSG000001 | CLN8 | 5.58E-04 | 1.72E-02 | 1.15E+00 | 2.00E-01 | 1.15E+00 |
| ENSG000001 | DYNLT3 | 5.58E-04 | 1.72E-02 | 8.58E-01 | -2.22E-01 | -1.17E+00 |
| ENSG00000C | RUFY3 | 5.59E-04 | 1.72E-02 | 8.59E-01 | -2.20E-01 | -1.16E+00 |
| ENSG00000C | RPS20 | 5.61E-04 | 1.73E-02 | 1.09E+00 | 1.31E-01 | 1.09E+00 |
| ENSG000001 | PGAP6 | 5.62E-04 | 1.73E-02 | 7.71E-01 | -3.76E-01 | -1.30E+00 |
| ENSG000001 | PTGR3 | 5.66E-04 | 1.74E-02 | 8.72E-01 | -1.97E-01 | -1.15E+00 |
| ENSG000001 | VPS53 | 5.68E-04 | 1.74E-02 | 8.87E-01 | -1.73E-01 | -1.13E+00 |
| ENSG000001 | BEX3 | 5.75E-04 | 1.75E-02 | 1.12E+00 | 1.59E-01 | 1.12E+00 |
| ENSG000001 | PSTPIP2 | 5.83E-04 | 1.78E-02 | 1.15E+00 | 1.98E-01 | 1.15E+00 |
| ENSG00000C | WBP11 | 5.86E-04 | 1.78E-02 | 8.41E-01 | -2.50E-01 | -1.19E+00 |
| ENSG000002 | BACE1-AS | 5.89E-04 | 1.79E-02 | 8.34E-01 | -2.61E-01 | -1.20E+00 |
| ENSG000001 | NCR3LG1 | 5.96E-04 | 1.80E-02 | 1.29E+00 | 3.62E-01 | 1.29E+00 |
| ENSG000001 | SLC7A11 | 6.00E-04 | 1.81E-02 | 6.85E-01 | -5.46E-01 | -1.46E+00 |
| ENSG000002 | MRPS17 | 6.04E-04 | 1.82E-02 | 1.13E+00 | 1.81E-01 | 1.13E+00 |
| ENSG000001 | RPL14 | 6.13E-04 | 1.84E-02 | 1.07E+00 | 9.93E-02 | 1.07E+00 |
| ENSG000001 | RUNX1 | 6.21E-04 | 1.86E-02 | 8.76E-01 | -1.91E-01 | -1.14E+00 |
| ENSG000001 | GID8 | 6.21E-04 | 1.86E-02 | 1.09E+00 | 1.19E-01 | 1.09E+00 |
| ENSG000001 | WBP4 | 6.22E-04 | 1.86E-02 | 8.47E-01 | -2.39E-01 | -1.18E+00 |
| ENSG000001 | PDSS2 | 6.23E-04 | 1.86E-02 | 8.37E-01 | -2.57E-01 | -1.19E+00 |
| ENSG000002 | NDUFAF8 | 6.29E-04 | 1.87E-02 | 1.21E+00 | 2.72E-01 | 1.21E+00 |
| ENSG000001 | ZNF761 | 6.32E-04 | 1.88E-02 | 8.77E-01 | -1.89E-01 | -1.14E+00 |
| ENSG000001 | TIGD2 | 6.39E-04 | 1.90E-02 | 1.16E+00 | 2.18E-01 | 1.16E+00 |
| ENSG000001 | ZMAT3 | 6.43E-04 | 1.90E-02 | 1.16E+00 | 2.10E-01 | 1.16E+00 |
| ENSG000001 | CCM2 | 6.44E-04 | 1.90E-02 | 1.17E+00 | 2.22E-01 | 1.17E+00 |
| ENSG000001 | GLB1L2 | 6.50E-04 | 1.92E-02 | 1.26E+00 | 3.37E-01 | 1.26E+00 |
| ENSG000001 | CCDC113 | 6.56E-04 | 1.93E-02 | 1.15E+00 | 2.04E-01 | 1.15E+00 |
| ENSG000001 | ADIRF | 6.66E-04 | 1.95E-02 | 7.62E-01 | -3.92E-01 | -1.31E+00 |
| ENSG000001 | KBTBD7 | 6.81E-04 | 1.99E-02 | 1.23E+00 | 2.98E-01 | 1.23E+00 |
| ENSG00000C | FAM76A | 7.02E-04 | 2.05E-02 | 8.36E-01 | -2.58E-01 | -1.20E+00 |

| | | | | | | |
|---|---|---|---|---|---|---|
| ENSG000002 | ZBED1 | 7.11E-04 | 2.07E-02 | 1.17E+00 | 2.25E-01 | 1.17E+00 |
| ENSG000002 | FKBP9P1 | 7.16E-04 | 2.08E-02 | 8.05E-01 | -3.14E-01 | -1.24E+00 |
| ENSG000001 | RPL37 | 7.21E-04 | 2.09E-02 | 1.09E+00 | 1.20E-01 | 1.09E+00 |
| ENSG000001 | ATP1B1 | 7.24E-04 | 2.10E-02 | 8.95E-01 | -1.60E-01 | -1.12E+00 |
| ENSG000001 | ACSL1 | 7.28E-04 | 2.11E-02 | 1.12E+00 | 1.67E-01 | 1.12E+00 |
| ENSG000001 | GSS | 7.30E-04 | 2.11E-02 | 1.15E+00 | 1.96E-01 | 1.15E+00 |
| ENSG000001 | CSGALNACT2 | 7.33E-04 | 2.11E-02 | 8.46E-01 | -2.42E-01 | -1.18E+00 |
| ENSG000001 | MBP | 7.34E-04 | 2.11E-02 | 8.67E-01 | -2.06E-01 | -1.15E+00 |
| ENSG000001 | SNRPD2 | 7.41E-04 | 2.13E-02 | 1.15E+00 | 1.97E-01 | 1.15E+00 |
| ENSG000001 | CSNK1A1 | 7.48E-04 | 2.14E-02 | 9.43E-01 | -8.47E-02 | -1.06E+00 |
| ENSG000001 | EIF2B2 | 7.52E-04 | 2.15E-02 | 8.70E-01 | -2.00E-01 | -1.15E+00 |
| ENSG000000 | NLRP2 | 7.54E-04 | 2.15E-02 | 1.15E+00 | 2.08E-01 | 1.15E+00 |
| ENSG000001 | ADORA2B | 7.57E-04 | 2.15E-02 | 1.17E+00 | 2.26E-01 | 1.17E+00 |
| ENSG000001 | SLC20A2 | 7.58E-04 | 2.15E-02 | 1.16E+00 | 2.15E-01 | 1.16E+00 |
| ENSG000002 | STIMATE-MU | 7.60E-04 | 2.15E-02 | 7.03E-01 | -5.09E-01 | -1.42E+00 |
| ENSG000001 | HPRT1 | 7.81E-04 | 2.21E-02 | 1.16E+00 | 2.09E-01 | 1.16E+00 |
| ENSG000001 | STXBP3 | 7.88E-04 | 2.22E-02 | 8.58E-01 | -2.22E-01 | -1.17E+00 |
| ENSG000001 | TDRP | 7.88E-04 | 2.22E-02 | 1.10E+00 | 1.33E-01 | 1.10E+00 |
| ENSG000001 | TOMM5 | 7.97E-04 | 2.24E-02 | 1.12E+00 | 1.59E-01 | 1.12E+00 |
| ENSG000002 | RN7SL3 | 8.12E-04 | ? | 3.93E+00 | 1.98E+00 | 3.93E+00 |
| ENSG000000 | MAPRE3 | 8.30E-04 | 2.33E-02 | 1.14E+00 | 1.84E-01 | 1.14E+00 |
| ENSG000001 | RIMS2 | 8.34E-04 | 2.34E-02 | 8.11E-01 | -3.03E-01 | -1.23E+00 |
| ENSG000002 | CCNY-AS1 | 8.36E-04 | ? | 3.21E+00 | 1.68E+00 | 3.21E+00 |
| ENSG000001 | RPS15A | 8.42E-04 | 2.35E-02 | 1.12E+00 | 1.58E-01 | 1.12E+00 |
| ENSG000001 | FBXL17 | 8.48E-04 | 2.37E-02 | 8.85E-01 | -1.77E-01 | -1.13E+00 |
| ENSG000001 | UQCR10 | 8.50E-04 | 2.37E-02 | 1.13E+00 | 1.71E-01 | 1.13E+00 |
| ENSG000002 | --- | 8.53E-04 | ? | 5.50E-01 | -8.62E-01 | -1.82E+00 |
| ENSG000002 | MKKS | 8.53E-04 | 2.37E-02 | 1.39E+00 | 4.78E-01 | 1.39E+00 |
| ENSG000001 | ANKRD29 | 8.55E-04 | 2.37E-02 | 1.22E+00 | 2.92E-01 | 1.22E+00 |
| ENSG000001 | VEGFC | 8.57E-04 | 2.37E-02 | 8.64E-01 | -2.11E-01 | -1.16E+00 |
| ENSG000001 | SNN | 8.59E-04 | 2.37E-02 | 1.18E+00 | 2.36E-01 | 1.18E+00 |
| ENSG000001 | NTPCR | 8.60E-04 | 2.37E-02 | 1.12E+00 | 1.62E-01 | 1.12E+00 |
| ENSG000001 | SLC49A4 | 8.62E-04 | 2.37E-02 | 1.18E+00 | 2.36E-01 | 1.18E+00 |
| ENSG000001 | UGT1A6 | 8.64E-04 | 2.37E-02 | 1.12E+00 | 1.69E-01 | 1.12E+00 |
| ENSG000001 | TP53AIP1 | 8.68E-04 | 2.38E-02 | 1.29E+00 | 3.63E-01 | 1.29E+00 |
| ENSG000000 | HSP90AA1 | 8.73E-04 | 2.39E-02 | 9.07E-01 | -1.41E-01 | -1.10E+00 |
| ENSG000002 | ASAH2B | 8.74E-04 | 2.39E-02 | 8.55E-01 | -2.27E-01 | -1.17E+00 |
| ENSG000001 | DNAJB6 | 8.77E-04 | 2.39E-02 | 1.10E+00 | 1.40E-01 | 1.10E+00 |
| ENSG000001 | GCNT4 | 8.79E-04 | 2.39E-02 | 1.17E+00 | 2.32E-01 | 1.17E+00 |
| ENSG000000 | PDCD7 | 8.82E-04 | 2.39E-02 | 8.46E-01 | -2.41E-01 | -1.18E+00 |
| ENSG000001 | NDFIP1 | 8.83E-04 | 2.39E-02 | 9.20E-01 | -1.20E-01 | -1.09E+00 |
| ENSG000001 | CLCN3 | 8.93E-04 | 2.42E-02 | 1.10E+00 | 1.33E-01 | 1.10E+00 |
| ENSG000001 | XPNPEP3 | 8.97E-04 | 2.42E-02 | 1.26E+00 | 3.30E-01 | 1.26E+00 |
| ENSG000001 | DYNLT1 | 9.01E-04 | 2.42E-02 | 1.09E+00 | 1.28E-01 | 1.09E+00 |
| ENSG000001 | HPS5 | 9.01E-04 | 2.42E-02 | 1.13E+00 | 1.81E-01 | 1.13E+00 |

| | | | | | | |
|---|---|---|---|---|---|---|
| ENSG000002 | ATP5MF | 9.03E-04 | 2.43E-02 | 1.12E+00 | 1.59E-01 | 1.12E+00 |
| ENSG000001 | TSPAN7 | 9.08E-04 | 2.43E-02 | 6.69E-01 | -5.80E-01 | -1.50E+00 |
| ENSG000001 | LIMK1 | 9.11E-04 | 2.43E-02 | 1.14E+00 | 1.87E-01 | 1.14E+00 |
| ENSG000001 | NEK6 | 9.11E-04 | 2.43E-02 | 1.15E+00 | 1.96E-01 | 1.15E+00 |
| ENSG000001 | TRIM32 | 9.22E-04 | 2.46E-02 | 1.11E+00 | 1.47E-01 | 1.11E+00 |
| ENSG000001 | OSBPL11 | 9.36E-04 | 2.49E-02 | 8.98E-01 | -1.55E-01 | -1.11E+00 |
| ENSG00000C | TBXAS1 | 9.36E-04 | 2.49E-02 | 6.70E-01 | -5.77E-01 | -1.49E+00 |
| ENSG000001 | TMEM170A | 9.41E-04 | 2.49E-02 | 8.83E-01 | -1.79E-01 | -1.13E+00 |
| ENSG000001 | BNIP2 | 9.51E-04 | 2.51E-02 | 8.97E-01 | -1.58E-01 | -1.12E+00 |
| ENSG000001 | GM2A | 9.57E-04 | 2.53E-02 | 1.15E+00 | 1.96E-01 | 1.15E+00 |
| ENSG000001 | ZNF33A | 9.61E-04 | 2.53E-02 | 8.50E-01 | -2.35E-01 | -1.18E+00 |
| ENSG000001 | DNPH1 | 9.72E-04 | 2.56E-02 | 1.18E+00 | 2.44E-01 | 1.18E+00 |
| ENSG000001 | FANCM | 9.74E-04 | 2.56E-02 | 8.55E-01 | -2.26E-01 | -1.17E+00 |
| ENSG000001 | MED28 | 9.94E-04 | 2.61E-02 | 1.16E+00 | 2.17E-01 | 1.16E+00 |
| ENSG000001 | RHBDL2 | 9.97E-04 | 2.61E-02 | 1.17E+00 | 2.30E-01 | 1.17E+00 |
| ENSG000001 | KRT17 | 1.01E-03 | 2.63E-02 | 1.09E+00 | 1.27E-01 | 1.09E+00 |
| ENSG000001 | INA | 1.02E-03 | 2.65E-02 | 8.29E-01 | -2.70E-01 | -1.21E+00 |
| ENSG000002 | EEF1G | 1.02E-03 | 2.66E-02 | 1.12E+00 | 1.66E-01 | 1.12E+00 |
| ENSG000001 | B4GALNT3 | 1.04E-03 | 2.70E-02 | 1.22E+00 | 2.90E-01 | 1.22E+00 |
| ENSG000001 | PARP1 | 1.05E-03 | 2.72E-02 | 1.11E+00 | 1.50E-01 | 1.11E+00 |
| ENSG000001 | NKAPD1 | 1.05E-03 | 2.72E-02 | 8.73E-01 | -1.97E-01 | -1.15E+00 |
| ENSG000001 | CCNT1 | 1.05E-03 | 2.72E-02 | 8.92E-01 | -1.65E-01 | -1.12E+00 |
| ENSG00000C | TIMM21 | 1.05E-03 | 2.72E-02 | 1.11E+00 | 1.53E-01 | 1.11E+00 |
| ENSG000001 | TMEM54 | 1.07E-03 | 2.74E-02 | 1.22E+00 | 2.86E-01 | 1.22E+00 |
| ENSG000002 | --- | 1.07E-03 | 2.74E-02 | 6.40E-01 | -6.45E-01 | -1.56E+00 |
| ENSG000001 | NADK2 | 1.07E-03 | 2.74E-02 | 8.82E-01 | -1.82E-01 | -1.13E+00 |
| ENSG000001 | RARG | 1.07E-03 | 2.74E-02 | 1.21E+00 | 2.71E-01 | 1.21E+00 |
| ENSG000001 | ANXA3 | 1.08E-03 | 2.75E-02 | 1.14E+00 | 1.85E-01 | 1.14E+00 |
| ENSG00000C | DGCR2 | 1.09E-03 | 2.77E-02 | 1.18E+00 | 2.39E-01 | 1.18E+00 |
| ENSG000001 | SMARCA1 | 1.09E-03 | 2.78E-02 | 8.69E-01 | -2.03E-01 | -1.15E+00 |
| ENSG000001 | RAG1 | 1.09E-03 | 2.78E-02 | 1.19E+00 | 2.50E-01 | 1.19E+00 |
| ENSG000001 | CAPZA1 | 1.10E-03 | 2.80E-02 | 9.06E-01 | -1.42E-01 | -1.10E+00 |
| ENSG000001 | RPS27 | 1.11E-03 | 2.80E-02 | 1.10E+00 | 1.43E-01 | 1.10E+00 |
| ENSG000001 | NFKBIA | 1.11E-03 | 2.82E-02 | 1.14E+00 | 1.91E-01 | 1.14E+00 |
| ENSG000001 | TENT4B | 1.12E-03 | 2.83E-02 | 8.90E-01 | -1.68E-01 | -1.12E+00 |
| ENSG000001 | ZNF678 | 1.13E-03 | 2.85E-02 | 7.96E-01 | -3.30E-01 | -1.26E+00 |
| ENSG000001 | CXADR | 1.13E-03 | 2.85E-02 | 8.96E-01 | -1.58E-01 | -1.12E+00 |
| ENSG00000C | YIPF1 | 1.14E-03 | 2.85E-02 | 8.90E-01 | -1.68E-01 | -1.12E+00 |
| ENSG000002 | MZT1 | 1.14E-03 | 2.85E-02 | 8.27E-01 | -2.74E-01 | -1.21E+00 |
| ENSG000001 | GPX4 | 1.15E-03 | 2.86E-02 | 1.13E+00 | 1.74E-01 | 1.13E+00 |
| ENSG000001 | CEP290 | 1.15E-03 | 2.86E-02 | 8.30E-01 | -2.69E-01 | -1.21E+00 |
| ENSG000001 | IGFBP3 | 1.15E-03 | 2.87E-02 | 7.46E-01 | -4.23E-01 | -1.34E+00 |
| ENSG000002 | --- | 1.15E-03 | 2.87E-02 | 1.22E+00 | 2.82E-01 | 1.22E+00 |
| ENSG000001 | MRPL1 | 1.16E-03 | 2.88E-02 | 8.54E-01 | -2.27E-01 | -1.17E+00 |
| ENSG000002 | --- | 1.16E-03 | 2.88E-02 | 6.49E-01 | -6.24E-01 | -1.54E+00 |

| | | | | | | |
|---|---|---|---|---|---|---|
| ENSG000001 | FBH1 | 1.17E-03 | 2.89E-02 | 1.09E+00 | 1.22E-01 | 1.09E+00 |
| ENSG00000C | FERMT2 | 1.17E-03 | 2.89E-02 | 1.19E+00 | 2.56E-01 | 1.19E+00 |
| ENSG000001 | SHLD2 | 1.17E-03 | 2.89E-02 | 8.63E-01 | -2.13E-01 | -1.16E+00 |
| ENSG000001 | LRIG3 | 1.18E-03 | 2.89E-02 | 1.20E+00 | 2.61E-01 | 1.20E+00 |
| ENSG000001 | DIAPH1 | 1.18E-03 | 2.90E-02 | 9.36E-01 | -9.54E-02 | -1.07E+00 |
| ENSG000001 | TMC7 | 1.18E-03 | 2.91E-02 | 1.19E+00 | 2.54E-01 | 1.19E+00 |
| ENSG000001 | AXL | 1.19E-03 | 2.91E-02 | 8.84E-01 | -1.77E-01 | -1.13E+00 |
| ENSG000001 | DPY19L1 | 1.20E-03 | 2.93E-02 | 8.38E-01 | -2.56E-01 | -1.19E+00 |
| ENSG000001 | RPL12 | 1.20E-03 | 2.94E-02 | 1.09E+00 | 1.27E-01 | 1.09E+00 |
| ENSG00000C | CD44 | 1.21E-03 | 2.94E-02 | 9.12E-01 | -1.33E-01 | -1.10E+00 |
| ENSG000001 | SCHIP1 | 1.21E-03 | 2.94E-02 | 1.35E+00 | 4.34E-01 | 1.35E+00 |
| ENSG000001 | RGS10 | 1.22E-03 | 2.95E-02 | 1.15E+00 | 1.99E-01 | 1.15E+00 |
| ENSG000001 | CBX6 | 1.22E-03 | 2.95E-02 | 1.28E+00 | 3.52E-01 | 1.28E+00 |
| ENSG000001 | VSNL1 | 1.23E-03 | 2.97E-02 | 1.21E+00 | 2.75E-01 | 1.21E+00 |
| ENSG000002 | --- | 1.23E-03 | ? | 1.74E+00 | 8.02E-01 | 1.74E+00 |
| ENSG000001 | RPS17 | 1.23E-03 | 2.98E-02 | 1.07E+00 | 9.79E-02 | 1.07E+00 |
| ENSG000001 | ENDOD1 | 1.24E-03 | 2.99E-02 | 1.13E+00 | 1.74E-01 | 1.13E+00 |
| ENSG000001 | FAM3C2P | 1.24E-03 | 2.99E-02 | 6.98E-01 | -5.19E-01 | -1.43E+00 |
| ENSG000001 | RPS16 | 1.25E-03 | 3.00E-02 | 1.11E+00 | 1.49E-01 | 1.11E+00 |
| ENSG000001 | MTCL1 | 1.25E-03 | 3.01E-02 | 1.14E+00 | 1.90E-01 | 1.14E+00 |
| ENSG000001 | SLC38A2 | 1.26E-03 | 3.02E-02 | 9.14E-01 | -1.30E-01 | -1.09E+00 |
| ENSG000001 | ZNF827 | 1.26E-03 | 3.02E-02 | 8.66E-01 | -2.08E-01 | -1.16E+00 |
| ENSG000001 | ALG10B | 1.28E-03 | 3.06E-02 | 8.09E-01 | -3.06E-01 | -1.24E+00 |
| ENSG000001 | ATXN1 | 1.28E-03 | 3.06E-02 | 8.07E-01 | -3.09E-01 | -1.24E+00 |
| ENSG000001 | CMSS1 | 1.29E-03 | 3.06E-02 | 8.77E-01 | -1.90E-01 | -1.14E+00 |
| ENSG000001 | ANO1 | 1.30E-03 | 3.09E-02 | 7.05E-01 | -5.04E-01 | -1.42E+00 |
| ENSG000001 | PDP1 | 1.30E-03 | 3.09E-02 | 8.44E-01 | -2.44E-01 | -1.18E+00 |
| ENSG000001 | SLIT2 | 1.31E-03 | 3.11E-02 | 7.38E-01 | -4.38E-01 | -1.35E+00 |
| ENSG000001 | ACOX1 | 1.32E-03 | 3.11E-02 | 1.10E+00 | 1.33E-01 | 1.10E+00 |
| ENSG000001 | CHST11 | 1.32E-03 | 3.12E-02 | 8.39E-01 | -2.52E-01 | -1.19E+00 |
| ENSG000001 | DUSP16 | 1.32E-03 | 3.12E-02 | 1.13E+00 | 1.76E-01 | 1.13E+00 |
| ENSG000001 | BDKRB2 | 1.33E-03 | 3.12E-02 | 1.18E+00 | 2.40E-01 | 1.18E+00 |
| ENSG000001 | TRIM8 | 1.33E-03 | 3.12E-02 | 8.93E-01 | -1.63E-01 | -1.12E+00 |
| ENSG000001 | NABP1 | 1.33E-03 | 3.12E-02 | 7.97E-01 | -3.27E-01 | -1.25E+00 |
| ENSG00000C | DNAJB11 | 1.34E-03 | 3.12E-02 | 9.04E-01 | -1.46E-01 | -1.11E+00 |
| ENSG000001 | CPEB4 | 1.34E-03 | 3.12E-02 | 8.53E-01 | -2.29E-01 | -1.17E+00 |
| ENSG000002 | MMP28 | 1.34E-03 | 3.14E-02 | 1.24E+00 | 3.07E-01 | 1.24E+00 |
| ENSG000001 | G3BP1 | 1.35E-03 | 3.14E-02 | 9.06E-01 | -1.42E-01 | -1.10E+00 |
| ENSG000001 | TRIM23 | 1.35E-03 | 3.14E-02 | 1.13E+00 | 1.79E-01 | 1.13E+00 |
| ENSG000001 | RGS18 | 1.35E-03 | 3.14E-02 | 7.96E-01 | -3.29E-01 | -1.26E+00 |
| ENSG000001 | FITM2 | 1.37E-03 | 3.17E-02 | 1.14E+00 | 1.87E-01 | 1.14E+00 |
| ENSG000001 | ETV3 | 1.37E-03 | 3.17E-02 | 9.10E-01 | -1.36E-01 | -1.10E+00 |
| ENSG000001 | ZNF490 | 1.37E-03 | 3.17E-02 | 8.24E-01 | -2.80E-01 | -1.21E+00 |
| ENSG000001 | CDH8 | 1.38E-03 | 3.17E-02 | 1.68E+00 | 7.44E-01 | 1.68E+00 |
| ENSG000001 | SLC41A2 | 1.38E-03 | 3.17E-02 | 8.37E-01 | -2.57E-01 | -1.19E+00 |

| | | | | | | |
|---|---|---|---|---|---|---|
| ENSG000001 | PLSCR3 | 1.38E-03 | 3.17E-02 | 1.15E+00 | 2.02E-01 | 1.15E+00 |
| ENSG000001 | SCN9A | 1.39E-03 | 3.18E-02 | 7.75E-01 | -3.67E-01 | -1.29E+00 |
| ENSG000001 | RPL35A | 1.40E-03 | 3.20E-02 | 1.11E+00 | 1.53E-01 | 1.11E+00 |
| ENSG000001 | NIBAN1 | 1.42E-03 | 3.26E-02 | 1.30E+00 | 3.74E-01 | 1.30E+00 |
| ENSG000001 | RNF141 | 1.43E-03 | 3.26E-02 | 1.10E+00 | 1.33E-01 | 1.10E+00 |
| ENSG000001 | MINDY2 | 1.43E-03 | 3.26E-02 | 8.64E-01 | -2.10E-01 | -1.16E+00 |
| ENSG000001 | SERTAD3 | 1.44E-03 | 3.28E-02 | 1.17E+00 | 2.29E-01 | 1.17E+00 |
| ENSG000001 | BTN2A1 | 1.44E-03 | 3.28E-02 | 8.63E-01 | -2.13E-01 | -1.16E+00 |
| ENSG00000C | AASS | 1.46E-03 | 3.31E-02 | 8.33E-01 | -2.64E-01 | -1.20E+00 |
| ENSG000001 | MCTP2 | 1.46E-03 | 3.31E-02 | 8.92E-01 | -1.64E-01 | -1.12E+00 |
| ENSG000002 | CYS1 | 1.49E-03 | 3.37E-02 | 1.62E+00 | 6.94E-01 | 1.62E+00 |
| ENSG000001 | PIGL | 1.49E-03 | 3.37E-02 | 8.21E-01 | -2.84E-01 | -1.22E+00 |
| ENSG000001 | TAX1BP1 | 1.49E-03 | 3.37E-02 | 8.76E-01 | -1.92E-01 | -1.14E+00 |
| ENSG000001 | SLC35D2 | 1.51E-03 | 3.39E-02 | 1.15E+00 | 2.05E-01 | 1.15E+00 |
| ENSG000001 | NAPEPLD | 1.52E-03 | 3.40E-02 | 8.38E-01 | -2.55E-01 | -1.19E+00 |
| ENSG00000C | PDIA5 | 1.52E-03 | 3.40E-02 | 1.12E+00 | 1.69E-01 | 1.12E+00 |
| ENSG000001 | IGFN1 | 1.52E-03 | 3.40E-02 | 5.31E-01 | -9.13E-01 | -1.88E+00 |
| ENSG000001 | CALU | 1.52E-03 | 3.41E-02 | 8.76E-01 | -1.91E-01 | -1.14E+00 |
| ENSG000001 | TMED8 | 1.53E-03 | 3.42E-02 | 8.18E-01 | -2.90E-01 | -1.22E+00 |
| ENSG000001 | IFNAR1 | 1.54E-03 | 3.42E-02 | 1.09E+00 | 1.18E-01 | 1.09E+00 |
| ENSG000001 | TRAFD1 | 1.54E-03 | 3.42E-02 | 1.14E+00 | 1.90E-01 | 1.14E+00 |
| ENSG000001 | IPPK | 1.54E-03 | 3.42E-02 | 1.15E+00 | 2.02E-01 | 1.15E+00 |
| ENSG00000C | PVR | 1.54E-03 | 3.42E-02 | 9.14E-01 | -1.30E-01 | -1.09E+00 |
| ENSG000001 | NEK7 | 1.54E-03 | 3.42E-02 | 8.88E-01 | -1.71E-01 | -1.13E+00 |
| ENSG000001 | ACOT2 | 1.56E-03 | 3.46E-02 | 1.32E+00 | 4.01E-01 | 1.32E+00 |
| ENSG000002 | ARHGAP19 | 1.57E-03 | 3.46E-02 | 8.39E-01 | -2.52E-01 | -1.19E+00 |
| ENSG000001 | MAPK8IP1 | 1.57E-03 | 3.46E-02 | 1.29E+00 | 3.69E-01 | 1.29E+00 |
| ENSG000001 | DPY19L3 | 1.57E-03 | 3.46E-02 | 8.16E-01 | -2.93E-01 | -1.22E+00 |
| ENSG00000C | PHF20 | 1.57E-03 | 3.46E-02 | 1.08E+00 | 1.15E-01 | 1.08E+00 |
| ENSG000001 | TCF4 | 1.57E-03 | 3.46E-02 | 1.25E+00 | 3.22E-01 | 1.25E+00 |
| ENSG000001 | STAU1 | 1.58E-03 | 3.47E-02 | 8.82E-01 | -1.81E-01 | -1.13E+00 |
| ENSG000001 | JAG1 | 1.60E-03 | 3.49E-02 | 8.84E-01 | -1.79E-01 | -1.13E+00 |
| ENSG000001 | RPL36 | 1.60E-03 | 3.49E-02 | 1.16E+00 | 2.16E-01 | 1.16E+00 |
| ENSG000001 | MRAP2 | 1.60E-03 | 3.49E-02 | 1.26E+00 | 3.35E-01 | 1.26E+00 |
| ENSG000002 | GASK1B-AS1 | 1.61E-03 | 3.51E-02 | 1.18E+00 | 2.33E-01 | 1.18E+00 |
| ENSG000001 | BTBD3 | 1.61E-03 | 3.51E-02 | 1.14E+00 | 1.90E-01 | 1.14E+00 |
| ENSG000001 | RRAS2 | 1.62E-03 | 3.53E-02 | 8.89E-01 | -1.70E-01 | -1.13E+00 |
| ENSG000002 | --- | 1.63E-03 | 3.54E-02 | 1.21E+00 | 2.78E-01 | 1.21E+00 |
| ENSG000001 | H2BC12 | 1.63E-03 | 3.54E-02 | 1.21E+00 | 2.81E-01 | 1.21E+00 |
| ENSG000001 | CDKN3 | 1.64E-03 | 3.54E-02 | 6.45E-01 | -6.33E-01 | -1.55E+00 |
| ENSG000001 | SMO | 1.64E-03 | 3.55E-02 | 1.40E+00 | 4.86E-01 | 1.40E+00 |
| ENSG000002 | PPP2R2A | 1.65E-03 | 3.56E-02 | 9.04E-01 | -1.46E-01 | -1.11E+00 |
| ENSG000001 | FAM217B | 1.67E-03 | 3.59E-02 | 8.68E-01 | -2.05E-01 | -1.15E+00 |
| ENSG000001 | H2BC5 | 1.69E-03 | 3.64E-02 | 8.53E-01 | -2.30E-01 | -1.17E+00 |
| ENSG00000C | SLC1A3 | 1.70E-03 | 3.64E-02 | 1.40E+00 | 4.83E-01 | 1.40E+00 |

| | | | | | | |
|---|---|---|---|---|---|---|
| ENSG000001 | RELB | 1.70E-03 | 3.64E-02 | 1.22E+00 | 2.87E-01 | 1.22E+00 |
| ENSG00000C | PFN2 | 1.70E-03 | 3.64E-02 | 1.08E+00 | 1.17E-01 | 1.08E+00 |
| ENSG000001 | SANBR | 1.70E-03 | 3.64E-02 | 8.63E-01 | -2.12E-01 | -1.16E+00 |
| ENSG000001 | E2F3 | 1.71E-03 | 3.65E-02 | 9.02E-01 | -1.49E-01 | -1.11E+00 |
| ENSG000001 | SQLE | 1.72E-03 | 3.67E-02 | 9.04E-01 | -1.45E-01 | -1.11E+00 |
| ENSG00000C | ABCC9 | 1.73E-03 | 3.68E-02 | 7.35E-01 | -4.44E-01 | -1.36E+00 |
| ENSG000001 | PDCD1LG2 | 1.75E-03 | 3.72E-02 | 1.16E+00 | 2.08E-01 | 1.16E+00 |
| ENSG000001 | FLRT3 | 1.76E-03 | 3.74E-02 | 1.16E+00 | 2.13E-01 | 1.16E+00 |
| ENSG000001 | CCNG1 | 1.78E-03 | 3.76E-02 | 1.17E+00 | 2.28E-01 | 1.17E+00 |
| ENSG000001 | RPL32 | 1.78E-03 | 3.77E-02 | 1.08E+00 | 1.09E-01 | 1.08E+00 |
| ENSG000001 | ANXA4 | 1.80E-03 | 3.80E-02 | 1.08E+00 | 1.16E-01 | 1.08E+00 |
| ENSG00000C | TRAM2 | 1.80E-03 | 3.80E-02 | 1.11E+00 | 1.54E-01 | 1.11E+00 |
| ENSG000001 | CPE | 1.82E-03 | 3.82E-02 | 1.25E+00 | 3.26E-01 | 1.25E+00 |
| ENSG000001 | RPS6 | 1.82E-03 | 3.82E-02 | 1.09E+00 | 1.21E-01 | 1.09E+00 |
| ENSG000002 | ATP6V1E2 | 1.82E-03 | 3.82E-02 | 1.19E+00 | 2.47E-01 | 1.19E+00 |
| ENSG000002 | PSMG3-AS1 | 1.82E-03 | 3.82E-02 | 1.30E+00 | 3.74E-01 | 1.30E+00 |
| ENSG000001 | RPS11 | 1.83E-03 | 3.82E-02 | 1.08E+00 | 1.17E-01 | 1.08E+00 |
| ENSG000001 | MYBL1 | 1.83E-03 | 3.83E-02 | 7.52E-01 | -4.12E-01 | -1.33E+00 |
| ENSG000002 | TSPY26P | 1.84E-03 | 3.84E-02 | 1.56E+00 | 6.41E-01 | 1.56E+00 |
| ENSG00000C | KLF6 | 1.85E-03 | 3.85E-02 | 9.40E-01 | -8.99E-02 | -1.06E+00 |
| ENSG000001 | ZNF138 | 1.85E-03 | 3.85E-02 | 7.96E-01 | -3.29E-01 | -1.26E+00 |
| ENSG000001 | HKDC1 | 1.86E-03 | 3.87E-02 | 7.43E-01 | -4.28E-01 | -1.35E+00 |
| ENSG000002 | APOBEC3C | 1.86E-03 | 3.87E-02 | 1.10E+00 | 1.40E-01 | 1.10E+00 |
| ENSG000001 | RCAN1 | 1.89E-03 | 3.90E-02 | 1.13E+00 | 1.70E-01 | 1.13E+00 |
| ENSG000001 | PTPRG | 1.89E-03 | 3.90E-02 | 8.34E-01 | -2.62E-01 | -1.20E+00 |
| ENSG000001 | RASEF | 1.89E-03 | 3.90E-02 | 1.25E+00 | 3.27E-01 | 1.25E+00 |
| ENSG00000C | ATP8B1 | 1.89E-03 | 3.90E-02 | 1.15E+00 | 1.96E-01 | 1.15E+00 |
| ENSG000002 | RPL7P1 | 1.90E-03 | 3.91E-02 | 1.33E+00 | 4.08E-01 | 1.33E+00 |
| ENSG000001 | NDN | 1.91E-03 | 3.93E-02 | 1.20E+00 | 2.64E-01 | 1.20E+00 |
| ENSG000001 | MLKL | 1.91E-03 | 3.93E-02 | 8.27E-01 | -2.73E-01 | -1.21E+00 |
| ENSG000002 | --- | 1.91E-03 | ? | 3.38E+00 | 1.76E+00 | 3.38E+00 |
| ENSG000001 | RBM47 | 1.92E-03 | 3.93E-02 | 1.11E+00 | 1.55E-01 | 1.11E+00 |
| ENSG000001 | UBE2D3 | 1.93E-03 | 3.94E-02 | 9.16E-01 | -1.26E-01 | -1.09E+00 |
| ENSG000001 | RPL5 | 1.93E-03 | 3.95E-02 | 1.08E+00 | 1.10E-01 | 1.08E+00 |
| ENSG000001 | DDX60L | 1.94E-03 | 3.96E-02 | 8.42E-01 | -2.49E-01 | -1.19E+00 |
| ENSG000001 | RPL27A | 1.94E-03 | 3.96E-02 | 1.09E+00 | 1.30E-01 | 1.09E+00 |
| ENSG000001 | SYK | 1.95E-03 | 3.97E-02 | 1.09E+00 | 1.20E-01 | 1.09E+00 |
| ENSG000002 | RTL5 | 1.95E-03 | 3.97E-02 | 1.32E+00 | 4.03E-01 | 1.32E+00 |
| ENSG000002 | LINC03033 | 1.96E-03 | ? | 4.91E-01 | -1.03E+00 | -2.04E+00 |
| ENSG000001 | NXPE3 | 1.98E-03 | 4.01E-02 | 8.90E-01 | -1.68E-01 | -1.12E+00 |
| ENSG00000C | ICAM1 | 1.98E-03 | 4.01E-02 | 8.34E-01 | -2.62E-01 | -1.20E+00 |
| ENSG000001 | FBN2 | 2.00E-03 | 4.04E-02 | 8.56E-01 | -2.24E-01 | -1.17E+00 |
| ENSG000002 | JPX | 2.01E-03 | 4.06E-02 | 8.40E-01 | -2.52E-01 | -1.19E+00 |
| ENSG000001 | SPRED3 | 2.01E-03 | 4.06E-02 | 8.61E-01 | -2.17E-01 | -1.16E+00 |
| ENSG000001 | ZNF70 | 2.01E-03 | 4.06E-02 | 1.20E+00 | 2.68E-01 | 1.20E+00 |

| | | | | | | |
|---|---|---|---|---|---|---|
| ENSG000001 | TGFBI | 2.02E-03 | 4.06E-02 | 7.87E-01 | -3.45E-01 | -1.27E+00 |
| ENSG00000C | ADAT1 | 2.02E-03 | 4.06E-02 | 1.10E+00 | 1.40E-01 | 1.10E+00 |
| ENSG00000C | WDR1 | 2.02E-03 | 4.06E-02 | 1.08E+00 | 1.10E-01 | 1.08E+00 |
| ENSG000002 | --- | 2.02E-03 | 4.06E-02 | 1.60E+00 | 6.79E-01 | 1.60E+00 |
| ENSG00000C | ATG5 | 2.03E-03 | 4.07E-02 | 1.10E+00 | 1.35E-01 | 1.10E+00 |
| ENSG000001 | C17orf58 | 2.03E-03 | 4.07E-02 | 1.18E+00 | 2.45E-01 | 1.18E+00 |
| ENSG000001 | IL6R | 2.04E-03 | 4.08E-02 | 8.48E-01 | -2.38E-01 | -1.18E+00 |
| ENSG00000C | INPP5A | 2.05E-03 | 4.09E-02 | 8.93E-01 | -1.64E-01 | -1.12E+00 |
| ENSG000001 | PWWP2A | 2.07E-03 | 4.11E-02 | 8.50E-01 | -2.34E-01 | -1.18E+00 |
| ENSG00000C | PIGV | 2.07E-03 | 4.11E-02 | 1.16E+00 | 2.17E-01 | 1.16E+00 |
| ENSG000001 | CTSZ | 2.07E-03 | 4.11E-02 | 8.93E-01 | -1.63E-01 | -1.12E+00 |
| ENSG00000C | LYRM2 | 2.08E-03 | 4.13E-02 | 1.10E+00 | 1.44E-01 | 1.10E+00 |
| ENSG000001 | PHC2 | 2.09E-03 | 4.14E-02 | 1.15E+00 | 2.01E-01 | 1.15E+00 |
| ENSG000001 | PGM2L1 | 2.10E-03 | 4.15E-02 | 8.20E-01 | -2.87E-01 | -1.22E+00 |
| ENSG000001 | TMPRSS11D | 2.11E-03 | 4.16E-02 | 6.06E-01 | -7.21E-01 | -1.65E+00 |
| ENSG000001 | MTR | 2.12E-03 | 4.17E-02 | 8.87E-01 | -1.74E-01 | -1.13E+00 |
| ENSG000001 | NTNG1 | 2.12E-03 | 4.17E-02 | 7.96E-01 | -3.29E-01 | -1.26E+00 |
| ENSG000002 | --- | 2.12E-03 | 4.17E-02 | 1.37E+00 | 4.59E-01 | 1.37E+00 |
| ENSG000001 | USP12 | 2.12E-03 | 4.17E-02 | 9.09E-01 | -1.38E-01 | -1.10E+00 |
| ENSG000002 | FMN1 | 2.14E-03 | 4.20E-02 | 7.96E-01 | -3.28E-01 | -1.26E+00 |
| ENSG000001 | WNT7B | 2.15E-03 | 4.21E-02 | 8.08E-01 | -3.07E-01 | -1.24E+00 |
| ENSG000001 | ERAP2 | 2.16E-03 | 4.23E-02 | 9.10E-01 | -1.36E-01 | -1.10E+00 |
| ENSG000001 | HMGN4 | 2.17E-03 | 4.24E-02 | 1.07E+00 | 1.02E-01 | 1.07E+00 |
| ENSG000002 | --- | 2.18E-03 | 4.24E-02 | 8.52E-01 | -2.31E-01 | -1.17E+00 |
| ENSG000001 | LPAR5 | 2.18E-03 | 4.24E-02 | 8.38E-01 | -2.56E-01 | -1.19E+00 |
| ENSG00000C | KLHL20 | 2.18E-03 | 4.25E-02 | 8.86E-01 | -1.75E-01 | -1.13E+00 |
| ENSG000001 | TMEM167A | 2.20E-03 | 4.27E-02 | 9.22E-01 | -1.17E-01 | -1.08E+00 |
| ENSG000002 | EXOSC6 | 2.21E-03 | 4.28E-02 | 8.85E-01 | -1.77E-01 | -1.13E+00 |
| ENSG000002 | --- | 2.23E-03 | 4.32E-02 | 1.58E+00 | 6.62E-01 | 1.58E+00 |
| ENSG000002 | TOB1-AS1 | 2.25E-03 | ? | 2.19E+00 | 1.13E+00 | 2.19E+00 |
| ENSG000001 | CTPS1 | 2.25E-03 | 4.36E-02 | 1.10E+00 | 1.43E-01 | 1.10E+00 |
| ENSG000001 | TRIM56 | 2.26E-03 | 4.36E-02 | 9.06E-01 | -1.42E-01 | -1.10E+00 |
| ENSG000001 | TUBA1B | 2.27E-03 | 4.37E-02 | 9.05E-01 | -1.45E-01 | -1.11E+00 |
| ENSG000001 | RPL4 | 2.28E-03 | 4.39E-02 | 1.08E+00 | 1.11E-01 | 1.08E+00 |
| ENSG000001 | PLAG1 | 2.28E-03 | 4.39E-02 | 1.19E+00 | 2.49E-01 | 1.19E+00 |
| ENSG000001 | HMOX1 | 2.28E-03 | 4.39E-02 | 7.22E-01 | -4.71E-01 | -1.39E+00 |
| ENSG000002 | --- | 2.29E-03 | 4.40E-02 | 7.71E-01 | -3.76E-01 | -1.30E+00 |
| ENSG000001 | CASP7 | 2.30E-03 | 4.41E-02 | 8.92E-01 | -1.64E-01 | -1.12E+00 |
| ENSG000001 | SAMD12 | 2.30E-03 | 4.41E-02 | 8.57E-01 | -2.23E-01 | -1.17E+00 |
| ENSG000001 | NPR2 | 2.31E-03 | 4.41E-02 | 8.70E-01 | -2.00E-01 | -1.15E+00 |
| ENSG000001 | PHLDB3 | 2.32E-03 | 4.43E-02 | 1.20E+00 | 2.67E-01 | 1.20E+00 |
| ENSG000002 | --- | 2.32E-03 | 4.43E-02 | 8.61E-01 | -2.16E-01 | -1.16E+00 |
| ENSG000001 | PNMA8A | 2.33E-03 | 4.43E-02 | 1.16E+00 | 2.13E-01 | 1.16E+00 |
| ENSG00000C | SLC25A24 | 2.33E-03 | 4.43E-02 | 8.83E-01 | -1.79E-01 | -1.13E+00 |
| ENSG000001 | LINC02915 | 2.33E-03 | ? | 5.71E-01 | -8.07E-01 | -1.75E+00 |

| | | | | | | |
|---|---|---|---|---|---|---|
| ENSG000002 | --- | 2.34E-03 | ? | 2.71E-01 | -1.88E+00 | -3.69E+00 |
| ENSG00000C | MYO9B | 2.36E-03 | 4.47E-02 | 1.09E+00 | 1.19E-01 | 1.09E+00 |
| ENSG00000C | VPS41 | 2.39E-03 | 4.51E-02 | 8.91E-01 | -1.67E-01 | -1.12E+00 |
| ENSG000002 | --- | 2.39E-03 | 4.51E-02 | 1.57E+00 | 6.53E-01 | 1.57E+00 |
| ENSG000001 | TSPYL1 | 2.39E-03 | 4.51E-02 | 9.30E-01 | -1.04E-01 | -1.07E+00 |
| ENSG000002 | --- | 2.42E-03 | ? | 3.96E+00 | 1.98E+00 | 3.96E+00 |
| ENSG000002 | GTF2H5 | 2.43E-03 | 4.59E-02 | 1.11E+00 | 1.52E-01 | 1.11E+00 |
| ENSG000001 | GLMP | 2.44E-03 | 4.59E-02 | 1.17E+00 | 2.25E-01 | 1.17E+00 |
| ENSG000001 | TXNDC15 | 2.44E-03 | 4.59E-02 | 1.11E+00 | 1.46E-01 | 1.11E+00 |
| ENSG000001 | ANKLE2 | 2.44E-03 | 4.59E-02 | 9.17E-01 | -1.25E-01 | -1.09E+00 |
| ENSG000001 | ADGRA3 | 2.45E-03 | 4.59E-02 | 1.08E+00 | 1.11E-01 | 1.08E+00 |
| ENSG000001 | PEX5 | 2.47E-03 | 4.62E-02 | 1.14E+00 | 1.93E-01 | 1.14E+00 |
| ENSG00000C | DCUN1D1 | 2.47E-03 | 4.63E-02 | 8.72E-01 | -1.98E-01 | -1.15E+00 |
| ENSG000001 | TACSTD2 | 2.48E-03 | 4.65E-02 | 1.14E+00 | 1.95E-01 | 1.14E+00 |
| ENSG000001 | BTBD9 | 2.51E-03 | 4.69E-02 | 9.15E-01 | -1.29E-01 | -1.09E+00 |
| ENSG000001 | NIPAL4 | 2.51E-03 | 4.69E-02 | 1.24E+00 | 3.13E-01 | 1.24E+00 |
| ENSG000001 | PEX11B | 2.52E-03 | 4.70E-02 | 1.13E+00 | 1.81E-01 | 1.13E+00 |
| ENSG000002 | UTP14C | 2.54E-03 | 4.73E-02 | 1.82E+00 | 8.65E-01 | 1.82E+00 |
| ENSG000002 | --- | 2.54E-03 | ? | 4.83E-01 | -1.05E+00 | -2.07E+00 |
| ENSG000001 | RPS3 | 2.56E-03 | 4.76E-02 | 1.14E+00 | 1.95E-01 | 1.14E+00 |
| ENSG00000C | PEBP1 | 2.59E-03 | 4.81E-02 | 1.12E+00 | 1.61E-01 | 1.12E+00 |
| ENSG000001 | UCHL1 | 2.60E-03 | 4.81E-02 | 8.01E-01 | -3.20E-01 | -1.25E+00 |
| ENSG000001 | HSD17B3 | 2.60E-03 | ? | 2.05E+00 | 1.03E+00 | 2.05E+00 |
| ENSG000002 | NBPF1 | 2.60E-03 | 4.81E-02 | 8.93E-01 | -1.63E-01 | -1.12E+00 |
| ENSG000001 | GASK1B | 2.60E-03 | 4.81E-02 | 1.15E+00 | 2.02E-01 | 1.15E+00 |
| ENSG000001 | CTSB | 2.61E-03 | 4.82E-02 | 8.82E-01 | -1.80E-01 | -1.13E+00 |
| ENSG00000C | NSF | 2.65E-03 | 4.88E-02 | 1.09E+00 | 1.29E-01 | 1.09E+00 |
| ENSG000001 | TMX1 | 2.66E-03 | 4.90E-02 | 9.04E-01 | -1.45E-01 | -1.11E+00 |
| ENSG000001 | SLC7A1 | 2.68E-03 | 4.92E-02 | 1.11E+00 | 1.54E-01 | 1.11E+00 |
| ENSG000001 | ALG14 | 2.69E-03 | 4.93E-02 | 8.53E-01 | -2.29E-01 | -1.17E+00 |
| ENSG000001 | USF3 | 2.70E-03 | 4.96E-02 | 8.15E-01 | -2.95E-01 | -1.23E+00 |
| ENSG000001 | SSR3 | 2.73E-03 | 5.01E-02 | 8.97E-01 | -1.56E-01 | -1.11E+00 |
| ENSG000001 | SHTN1 | 2.75E-03 | 5.04E-02 | 8.60E-01 | -2.18E-01 | -1.16E+00 |
| ENSG000002 | TINCR | 2.78E-03 | 5.07E-02 | 1.47E+00 | 5.52E-01 | 1.47E+00 |
| ENSG000001 | ZNF322 | 2.79E-03 | 5.10E-02 | 8.95E-01 | -1.60E-01 | -1.12E+00 |
| ENSG000001 | NMNAT2 | 2.80E-03 | 5.10E-02 | 7.58E-01 | -3.99E-01 | -1.32E+00 |
| ENSG000001 | CRADD | 2.80E-03 | 5.10E-02 | 7.69E-01 | -3.79E-01 | -1.30E+00 |
| ENSG000001 | ALDH4A1 | 2.82E-03 | 5.13E-02 | 1.24E+00 | 3.12E-01 | 1.24E+00 |
| ENSG000002 | --- | 2.83E-03 | ? | 5.03E+00 | 2.33E+00 | 5.03E+00 |
| ENSG000001 | KIFBP | 2.87E-03 | 5.21E-02 | 9.05E-01 | -1.43E-01 | -1.10E+00 |
| ENSG000001 | CHST10 | 2.87E-03 | 5.21E-02 | 1.20E+00 | 2.57E-01 | 1.20E+00 |
| ENSG000001 | S100A2 | 2.89E-03 | 5.24E-02 | 1.21E+00 | 2.81E-01 | 1.21E+00 |
| ENSG000001 | PTGFRN | 2.90E-03 | 5.24E-02 | 1.09E+00 | 1.31E-01 | 1.09E+00 |
| ENSG000001 | TIPRL | 2.90E-03 | 5.24E-02 | 8.67E-01 | -2.06E-01 | -1.15E+00 |
| ENSG000001 | SLC40A1 | 2.91E-03 | 5.24E-02 | 1.39E+00 | 4.76E-01 | 1.39E+00 |

| | | | | | | |
|---|---|---|---|---|---|---|
| ENSG000001 | UCHL5 | 2.91E-03 | 5.24E-02 | 8.50E-01 | -2.35E-01 | -1.18E+00 |
| ENSG000001 | HNRNPA1L2 | 2.91E-03 | ? | 4.78E+00 | 2.26E+00 | 4.78E+00 |
| ENSG000001 | RBBP4 | 2.91E-03 | 5.24E-02 | 9.20E-01 | -1.20E-01 | -1.09E+00 |
| ENSG000001 | SLC66A3 | 2.92E-03 | 5.25E-02 | 1.15E+00 | 1.99E-01 | 1.15E+00 |
| ENSG00000C | REXO2 | 2.93E-03 | 5.26E-02 | 1.08E+00 | 1.05E-01 | 1.08E+00 |
| ENSG000001 | ZNF518B | 2.94E-03 | 5.28E-02 | 8.91E-01 | -1.67E-01 | -1.12E+00 |
| ENSG000001 | EIF4E2 | 2.95E-03 | 5.29E-02 | 9.17E-01 | -1.25E-01 | -1.09E+00 |
| ENSG000001 | C4orf3 | 2.96E-03 | 5.29E-02 | 1.10E+00 | 1.43E-01 | 1.10E+00 |
| ENSG000002 | ITGB2-AS1 | 3.00E-03 | 5.36E-02 | 5.81E-01 | -7.83E-01 | -1.72E+00 |
| ENSG000001 | KCNA2 | 3.03E-03 | ? | 2.19E+00 | 1.13E+00 | 2.19E+00 |
| ENSG000001 | TUSC2 | 3.04E-03 | 5.42E-02 | 8.69E-01 | -2.03E-01 | -1.15E+00 |
| ENSG000001 | BCOR | 3.04E-03 | 5.42E-02 | 9.16E-01 | -1.27E-01 | -1.09E+00 |
| ENSG000001 | STAM2 | 3.04E-03 | 5.42E-02 | 8.83E-01 | -1.79E-01 | -1.13E+00 |
| ENSG000002 | CRYZP1 | 3.05E-03 | ? | 3.98E-01 | -1.33E+00 | -2.51E+00 |
| ENSG000001 | STAG1 | 3.06E-03 | 5.45E-02 | 9.07E-01 | -1.41E-01 | -1.10E+00 |
| ENSG000001 | GUSBP1 | 3.08E-03 | ? | 2.55E+00 | 1.35E+00 | 2.55E+00 |
| ENSG000002 | LINC02541 | 3.09E-03 | 5.49E-02 | 1.29E+00 | 3.71E-01 | 1.29E+00 |
| ENSG000001 | GPX2 | 3.10E-03 | 5.50E-02 | 8.09E-01 | -3.06E-01 | -1.24E+00 |
| ENSG00000C | HADHA | 3.15E-03 | 5.58E-02 | 1.06E+00 | 8.42E-02 | 1.06E+00 |
| ENSG000001 | GALNT7 | 3.26E-03 | 5.76E-02 | 1.10E+00 | 1.40E-01 | 1.10E+00 |
| ENSG000002 | --- | 3.30E-03 | 5.84E-02 | 7.27E-01 | -4.61E-01 | -1.38E+00 |
| ENSG000001 | ACAA2 | 3.32E-03 | 5.86E-02 | 1.13E+00 | 1.77E-01 | 1.13E+00 |
| ENSG000001 | F11R | 3.33E-03 | 5.86E-02 | 1.08E+00 | 1.10E-01 | 1.08E+00 |
| ENSG000001 | LDHB | 3.34E-03 | 5.88E-02 | 1.12E+00 | 1.60E-01 | 1.12E+00 |
| ENSG000001 | SUPT20H | 3.35E-03 | 5.89E-02 | 9.14E-01 | -1.29E-01 | -1.09E+00 |
| ENSG000002 | --- | 3.36E-03 | 5.90E-02 | 4.61E-01 | -1.12E+00 | -2.17E+00 |
| ENSG00000C | GDE1 | 3.38E-03 | 5.92E-02 | 1.07E+00 | 1.04E-01 | 1.07E+00 |
| ENSG000001 | HSPD1 | 3.40E-03 | 5.95E-02 | 8.74E-01 | -1.94E-01 | -1.14E+00 |
| ENSG000001 | SCYL2 | 3.41E-03 | 5.96E-02 | 9.02E-01 | -1.49E-01 | -1.11E+00 |
| ENSG000001 | BIRC2 | 3.41E-03 | 5.96E-02 | 9.23E-01 | -1.16E-01 | -1.08E+00 |
| ENSG000001 | IDH1 | 3.42E-03 | 5.96E-02 | 8.88E-01 | -1.72E-01 | -1.13E+00 |
| ENSG000002 | NOP14-AS1 | 3.42E-03 | 5.96E-02 | 9.11E-01 | -1.35E-01 | -1.10E+00 |
| ENSG00000C | CLNS1A | 3.45E-03 | 6.00E-02 | 1.12E+00 | 1.58E-01 | 1.12E+00 |
| ENSG000001 | XYLT1 | 3.45E-03 | 6.00E-02 | 1.42E+00 | 5.04E-01 | 1.42E+00 |
| ENSG000002 | LINC00342 | 3.47E-03 | 6.03E-02 | 7.47E-01 | -4.20E-01 | -1.34E+00 |
| ENSG000001 | FBN1 | 3.49E-03 | 6.05E-02 | 1.28E+00 | 3.61E-01 | 1.28E+00 |
| ENSG000001 | RPS3A | 3.49E-03 | 6.05E-02 | 1.10E+00 | 1.33E-01 | 1.10E+00 |
| ENSG000001 | EHBP1 | 3.50E-03 | 6.05E-02 | 9.19E-01 | -1.22E-01 | -1.09E+00 |
| ENSG000001 | TTC14 | 3.51E-03 | 6.07E-02 | 8.86E-01 | -1.75E-01 | -1.13E+00 |
| ENSG000001 | GNS | 3.53E-03 | 6.09E-02 | 1.08E+00 | 1.08E-01 | 1.08E+00 |
| ENSG000002 | PRCD | 3.54E-03 | 6.11E-02 | 1.34E+00 | 4.26E-01 | 1.34E+00 |
| ENSG000001 | MPV17 | 3.57E-03 | 6.15E-02 | 1.11E+00 | 1.46E-01 | 1.11E+00 |
| ENSG000001 | FAM171A1 | 3.59E-03 | 6.18E-02 | 8.97E-01 | -1.57E-01 | -1.12E+00 |
| ENSG000001 | KLHL9 | 3.61E-03 | 6.21E-02 | 1.10E+00 | 1.34E-01 | 1.10E+00 |
| ENSG000001 | MET | 3.62E-03 | 6.21E-02 | 1.10E+00 | 1.38E-01 | 1.10E+00 |

| | | | | | | |
|---|---|---|---|---|---|---|
| ENSG000001 | IRF6 | 3.63E-03 | 6.21E-02 | 1.08E+00 | 1.10E-01 | 1.08E+00 |
| ENSG000001 | MARVELD3 | 3.64E-03 | 6.22E-02 | 1.16E+00 | 2.20E-01 | 1.16E+00 |
| ENSG000001 | TMED10 | 3.64E-03 | 6.22E-02 | 9.24E-01 | -1.14E-01 | -1.08E+00 |
| ENSG00000C | SLC4A11 | 3.66E-03 | 6.24E-02 | 8.43E-01 | -2.47E-01 | -1.19E+00 |
| ENSG000001 | NFKBIE | 3.67E-03 | 6.25E-02 | 1.30E+00 | 3.74E-01 | 1.30E+00 |
| ENSG00000C | SEL1L | 3.68E-03 | 6.27E-02 | 8.95E-01 | -1.60E-01 | -1.12E+00 |
| ENSG000001 | GLRX | 3.72E-03 | 6.32E-02 | 8.68E-01 | -2.04E-01 | -1.15E+00 |
| ENSG000001 | FAH | 3.72E-03 | 6.33E-02 | 1.15E+00 | 2.00E-01 | 1.15E+00 |
| ENSG000001 | S100A10 | 3.73E-03 | 6.33E-02 | 1.08E+00 | 1.12E-01 | 1.08E+00 |
| ENSG000001 | NDUFV3 | 3.74E-03 | 6.34E-02 | 1.10E+00 | 1.36E-01 | 1.10E+00 |
| ENSG000001 | SMPD1 | 3.74E-03 | 6.34E-02 | 1.19E+00 | 2.47E-01 | 1.19E+00 |
| ENSG000002 | --- | 3.75E-03 | 6.34E-02 | 1.26E+00 | 3.34E-01 | 1.26E+00 |
| ENSG000001 | CDK8 | 3.75E-03 | 6.34E-02 | 1.08E+00 | 1.17E-01 | 1.08E+00 |
| ENSG000002 | --- | 3.76E-03 | 6.34E-02 | 7.61E-01 | -3.94E-01 | -1.31E+00 |
| ENSG000001 | HEXIM1 | 3.77E-03 | 6.36E-02 | 8.83E-01 | -1.79E-01 | -1.13E+00 |
| ENSG00000C | RBM22 | 3.78E-03 | 6.36E-02 | 9.12E-01 | -1.33E-01 | -1.10E+00 |
| ENSG000001 | AMFR | 3.79E-03 | 6.37E-02 | 1.11E+00 | 1.52E-01 | 1.11E+00 |
| ENSG000002 | --- | 3.79E-03 | ? | 2.45E+00 | 1.29E+00 | 2.45E+00 |
| ENSG00000C | MNAT1 | 3.80E-03 | 6.38E-02 | 8.48E-01 | -2.39E-01 | -1.18E+00 |
| ENSG000001 | PTMA | 3.80E-03 | 6.38E-02 | 8.57E-01 | -2.22E-01 | -1.17E+00 |
| ENSG000001 | CCAR2 | 3.81E-03 | 6.38E-02 | 1.09E+00 | 1.31E-01 | 1.09E+00 |
| ENSG000001 | SGPL1 | 3.83E-03 | 6.41E-02 | 1.08E+00 | 1.14E-01 | 1.08E+00 |
| ENSG000002 | --- | 3.84E-03 | 6.41E-02 | 6.59E-01 | -6.01E-01 | -1.52E+00 |
| ENSG000001 | CDHR1 | 3.85E-03 | 6.42E-02 | 1.17E+00 | 2.29E-01 | 1.17E+00 |
| ENSG000001 | AHR | 3.85E-03 | 6.42E-02 | 1.16E+00 | 2.09E-01 | 1.16E+00 |
| ENSG000001 | TACO1 | 3.86E-03 | 6.43E-02 | 1.12E+00 | 1.70E-01 | 1.12E+00 |
| ENSG000001 | CTSF | 3.86E-03 | 6.43E-02 | 1.23E+00 | 2.98E-01 | 1.23E+00 |
| ENSG000001 | FAM222B | 3.87E-03 | 6.44E-02 | 1.08E+00 | 1.17E-01 | 1.08E+00 |
| ENSG000001 | UXS1 | 3.89E-03 | 6.45E-02 | 1.11E+00 | 1.57E-01 | 1.11E+00 |
| ENSG000001 | PFKM | 3.89E-03 | 6.45E-02 | 1.13E+00 | 1.74E-01 | 1.13E+00 |
| ENSG000001 | TCFL5 | 3.93E-03 | 6.51E-02 | 1.17E+00 | 2.24E-01 | 1.17E+00 |
| ENSG000001 | PARD6G | 3.93E-03 | 6.51E-02 | 1.15E+00 | 2.01E-01 | 1.15E+00 |
| ENSG000001 | GAREM1 | 3.94E-03 | 6.51E-02 | 1.17E+00 | 2.27E-01 | 1.17E+00 |
| ENSG000001 | CYP2U1 | 3.97E-03 | 6.54E-02 | 1.24E+00 | 3.05E-01 | 1.24E+00 |
| ENSG000001 | TAGLN | 3.98E-03 | 6.56E-02 | 1.27E+00 | 3.50E-01 | 1.27E+00 |
| ENSG000001 | DHRS7 | 3.99E-03 | 6.56E-02 | 9.27E-01 | -1.10E-01 | -1.08E+00 |
| ENSG00000C | SNAPC1 | 3.99E-03 | 6.56E-02 | 8.78E-01 | -1.87E-01 | -1.14E+00 |
| ENSG000002 | NHSL2 | 3.99E-03 | ? | 1.79E+00 | 8.36E-01 | 1.79E+00 |
| ENSG000001 | ZNFX1 | 4.00E-03 | 6.57E-02 | 1.09E+00 | 1.23E-01 | 1.09E+00 |
| ENSG000001 | CCDC91 | 4.00E-03 | 6.57E-02 | 8.66E-01 | -2.07E-01 | -1.15E+00 |
| ENSG000001 | ASMTL | 4.02E-03 | 6.57E-02 | 1.17E+00 | 2.28E-01 | 1.17E+00 |
| ENSG000001 | ZDHHC5 | 4.02E-03 | 6.57E-02 | 9.16E-01 | -1.26E-01 | -1.09E+00 |
| ENSG000001 | OXSR1 | 4.02E-03 | 6.57E-02 | 9.40E-01 | -8.94E-02 | -1.06E+00 |
| ENSG000001 | APOL1 | 4.03E-03 | 6.57E-02 | 8.05E-01 | -3.13E-01 | -1.24E+00 |
| ENSG000001 | ERBB2 | 4.03E-03 | 6.57E-02 | 1.13E+00 | 1.70E-01 | 1.13E+00 |

| | | | | | | |
|---|---|---|---|---|---|---|
| ENSG000001 | MOB4 | 4.05E-03 | 6.58E-02 | 9.01E-01 | -1.51E-01 | -1.11E+00 |
| ENSG000001 | NYAP2 | 4.05E-03 | 6.58E-02 | 7.50E-01 | -4.15E-01 | -1.33E+00 |
| ENSG000001 | SENP6 | 4.05E-03 | 6.58E-02 | 9.02E-01 | -1.49E-01 | -1.11E+00 |
| ENSG000001 | IGIP | 4.05E-03 | 6.58E-02 | 1.23E+00 | 3.01E-01 | 1.23E+00 |
| ENSG000001 | TTYH2 | 4.06E-03 | 6.58E-02 | 1.36E+00 | 4.39E-01 | 1.36E+00 |
| ENSG000001 | ZNF84 | 4.07E-03 | 6.59E-02 | 8.62E-01 | -2.15E-01 | -1.16E+00 |
| ENSG000001 | FLACC1 | 4.07E-03 | ? | 4.11E-01 | -1.28E+00 | -2.43E+00 |
| ENSG000001 | GLRX3 | 4.08E-03 | 6.60E-02 | 9.23E-01 | -1.16E-01 | -1.08E+00 |
| ENSG000001 | PLPP3 | 4.12E-03 | 6.67E-02 | 1.24E+00 | 3.14E-01 | 1.24E+00 |
| ENSG000002 | MIR100HG | 4.14E-03 | 6.69E-02 | 8.15E-01 | -2.95E-01 | -1.23E+00 |
| ENSG000001 | SMARCC1 | 4.17E-03 | 6.72E-02 | 9.24E-01 | -1.14E-01 | -1.08E+00 |
| ENSG000001 | TMEM106C | 4.17E-03 | 6.72E-02 | 1.15E+00 | 2.03E-01 | 1.15E+00 |
| ENSG000001 | GPR108 | 4.20E-03 | 6.75E-02 | 1.12E+00 | 1.64E-01 | 1.12E+00 |
| ENSG000001 | GALNT5 | 4.20E-03 | 6.75E-02 | 1.13E+00 | 1.82E-01 | 1.13E+00 |
| ENSG000001 | BCAS4 | 4.22E-03 | 6.77E-02 | 1.25E+00 | 3.18E-01 | 1.25E+00 |
| ENSG000002 | LINC02547 | 4.24E-03 | ? | 4.22E-01 | -1.24E+00 | -2.37E+00 |
| ENSG00000C | DNAJA1 | 4.24E-03 | 6.80E-02 | 9.17E-01 | -1.24E-01 | -1.09E+00 |
| ENSG00000C | KIF16B | 4.25E-03 | 6.81E-02 | 1.09E+00 | 1.28E-01 | 1.09E+00 |
| ENSG00000C | HNRNPH3 | 4.26E-03 | 6.81E-02 | 8.88E-01 | -1.72E-01 | -1.13E+00 |
| ENSG000001 | LIMK2 | 4.27E-03 | 6.83E-02 | 1.11E+00 | 1.53E-01 | 1.11E+00 |
| ENSG000001 | TUBA4A | 4.29E-03 | 6.85E-02 | 8.83E-01 | -1.80E-01 | -1.13E+00 |
| ENSG000001 | CSNK1G1 | 4.29E-03 | 6.85E-02 | 9.21E-01 | -1.19E-01 | -1.09E+00 |
| ENSG000001 | MPZL3 | 4.31E-03 | 6.85E-02 | 9.14E-01 | -1.30E-01 | -1.09E+00 |
| ENSG000001 | DNAJA4 | 4.32E-03 | 6.85E-02 | 8.52E-01 | -2.31E-01 | -1.17E+00 |
| ENSG00000C | STK17B | 4.32E-03 | 6.85E-02 | 8.67E-01 | -2.06E-01 | -1.15E+00 |
| ENSG000001 | GNL3L | 4.32E-03 | 6.85E-02 | 1.20E+00 | 2.63E-01 | 1.20E+00 |
| ENSG000001 | H2BC21 | 4.33E-03 | 6.85E-02 | 1.15E+00 | 2.01E-01 | 1.15E+00 |
| ENSG000001 | RTN4 | 4.33E-03 | 6.85E-02 | 9.19E-01 | -1.23E-01 | -1.09E+00 |
| ENSG00000C | DGKD | 4.33E-03 | 6.85E-02 | 8.88E-01 | -1.72E-01 | -1.13E+00 |
| ENSG000001 | PHLDA1 | 4.34E-03 | 6.86E-02 | 8.87E-01 | -1.73E-01 | -1.13E+00 |
| ENSG00000C | MARCO | 4.35E-03 | ? | 4.85E-01 | -1.04E+00 | -2.06E+00 |
| ENSG00000C | HYAL2 | 4.38E-03 | 6.92E-02 | 1.17E+00 | 2.22E-01 | 1.17E+00 |
| ENSG000001 | KCTD12 | 4.39E-03 | 6.92E-02 | 8.60E-01 | -2.18E-01 | -1.16E+00 |
| ENSG000001 | RSKR | 4.39E-03 | 6.92E-02 | 7.81E-01 | -3.57E-01 | -1.28E+00 |
| ENSG00000C | VTA1 | 4.44E-03 | 6.99E-02 | 8.95E-01 | -1.60E-01 | -1.12E+00 |
| ENSG000001 | EPCAM | 4.46E-03 | 7.00E-02 | 8.26E-01 | -2.75E-01 | -1.21E+00 |
| ENSG000001 | MAPRE2 | 4.46E-03 | 7.00E-02 | 8.96E-01 | -1.59E-01 | -1.12E+00 |
| ENSG000001 | WASHC4 | 4.47E-03 | 7.00E-02 | 9.04E-01 | -1.45E-01 | -1.11E+00 |
| ENSG000002 | RBM15B | 4.47E-03 | 7.00E-02 | 1.08E+00 | 1.14E-01 | 1.08E+00 |
| ENSG000001 | CHD1L | 4.48E-03 | 7.00E-02 | 1.09E+00 | 1.24E-01 | 1.09E+00 |
| ENSG000001 | LNX2 | 4.48E-03 | 7.00E-02 | 1.11E+00 | 1.47E-01 | 1.11E+00 |
| ENSG000001 | MYH15 | 4.50E-03 | 7.02E-02 | 7.51E-01 | -4.14E-01 | -1.33E+00 |
| ENSG000002 | --- | 4.50E-03 | 7.02E-02 | 8.79E-01 | -1.86E-01 | -1.14E+00 |
| ENSG000002 | PLPP6 | 4.54E-03 | 7.08E-02 | 1.13E+00 | 1.78E-01 | 1.13E+00 |
| ENSG000001 | LPIN2 | 4.55E-03 | 7.08E-02 | 1.16E+00 | 2.09E-01 | 1.16E+00 |

| | | | | | | |
|---|---|---|---|---|---|---|
| ENSG00000C | TXNL1 | 4.55E-03 | 7.08E-02 | 9.11E-01 | -1.34E-01 | -1.10E+00 |
| ENSG000001 | ADAM8 | 4.57E-03 | 7.10E-02 | 8.03E-01 | -3.17E-01 | -1.25E+00 |
| ENSG000001 | GOT2 | 4.58E-03 | 7.11E-02 | 1.09E+00 | 1.27E-01 | 1.09E+00 |
| ENSG00000C | SCFD1 | 4.61E-03 | 7.14E-02 | 9.08E-01 | -1.39E-01 | -1.10E+00 |
| ENSG000001 | SPP1 | 4.62E-03 | 7.15E-02 | 5.89E-01 | -7.64E-01 | -1.70E+00 |
| ENSG000002 | NPTN-IT1 | 4.63E-03 | 7.15E-02 | 7.58E-01 | -3.99E-01 | -1.32E+00 |
| ENSG000002 | --- | 4.63E-03 | 7.15E-02 | 7.86E-01 | -3.48E-01 | -1.27E+00 |
| ENSG000001 | SLC38A9 | 4.63E-03 | 7.15E-02 | 1.10E+00 | 1.31E-01 | 1.10E+00 |
| ENSG000001 | CASK | 4.64E-03 | 7.16E-02 | 8.92E-01 | -1.64E-01 | -1.12E+00 |
| ENSG000002 | --- | 4.65E-03 | 7.17E-02 | 7.43E-01 | -4.29E-01 | -1.35E+00 |
| ENSG00000C | AP3D1 | 4.70E-03 | 7.22E-02 | 9.12E-01 | -1.33E-01 | -1.10E+00 |
| ENSG000001 | ECT2 | 4.70E-03 | 7.23E-02 | 9.16E-01 | -1.26E-01 | -1.09E+00 |
| ENSG000001 | CD163L1 | 4.72E-03 | 7.24E-02 | 7.82E-01 | -3.55E-01 | -1.28E+00 |
| ENSG000001 | EDARADD | 4.73E-03 | 7.25E-02 | 1.15E+00 | 2.03E-01 | 1.15E+00 |
| ENSG000001 | NAP1L5 | 4.75E-03 | 7.28E-02 | 1.23E+00 | 3.03E-01 | 1.23E+00 |
| ENSG000001 | PLS3 | 4.78E-03 | 7.30E-02 | 1.09E+00 | 1.19E-01 | 1.09E+00 |
| ENSG000001 | EDEM3 | 4.81E-03 | 7.35E-02 | 9.05E-01 | -1.44E-01 | -1.10E+00 |
| ENSG000001 | PIP4K2A | 4.84E-03 | 7.39E-02 | 8.99E-01 | -1.53E-01 | -1.11E+00 |
| ENSG000002 | NEU1 | 4.85E-03 | 7.39E-02 | 1.17E+00 | 2.28E-01 | 1.17E+00 |
| ENSG000001 | TPP1 | 4.86E-03 | 7.41E-02 | 8.75E-01 | -1.92E-01 | -1.14E+00 |
| ENSG000002 | TMEM35B | 4.91E-03 | 7.47E-02 | 1.30E+00 | 3.81E-01 | 1.30E+00 |
| ENSG000001 | CDCA4 | 4.92E-03 | 7.47E-02 | 1.17E+00 | 2.23E-01 | 1.17E+00 |
| ENSG000002 | RPS18 | 4.92E-03 | 7.47E-02 | 1.16E+00 | 2.13E-01 | 1.16E+00 |
| ENSG000001 | CPT1A | 4.94E-03 | 7.47E-02 | 1.14E+00 | 1.94E-01 | 1.14E+00 |
| ENSG00000C | NME1-NME2 | 4.94E-03 | 7.47E-02 | 1.21E+00 | 2.73E-01 | 1.21E+00 |
| ENSG000001 | USP42 | 4.94E-03 | 7.47E-02 | 8.76E-01 | -1.92E-01 | -1.14E+00 |
| ENSG000001 | SPHK1 | 4.95E-03 | 7.47E-02 | 1.17E+00 | 2.25E-01 | 1.17E+00 |
| ENSG00000C | ZNF76 | 4.95E-03 | 7.47E-02 | 1.16E+00 | 2.13E-01 | 1.16E+00 |
| ENSG000001 | HOXA1 | 4.95E-03 | 7.47E-02 | 1.25E+00 | 3.23E-01 | 1.25E+00 |
| ENSG000001 | TLE1 | 4.97E-03 | 7.50E-02 | 1.11E+00 | 1.54E-01 | 1.11E+00 |
| ENSG000001 | C18orf25 | 4.99E-03 | 7.50E-02 | 9.22E-01 | -1.16E-01 | -1.08E+00 |
| ENSG000002 | --- | 4.99E-03 | 7.50E-02 | 1.32E+00 | 4.04E-01 | 1.32E+00 |
| ENSG000001 | NRG1 | 5.00E-03 | 7.50E-02 | 8.77E-01 | -1.89E-01 | -1.14E+00 |
| ENSG000001 | CDC42EP5 | 5.01E-03 | 7.52E-02 | 1.32E+00 | 4.02E-01 | 1.32E+00 |
| ENSG000001 | MED18 | 5.06E-03 | 7.58E-02 | 1.13E+00 | 1.75E-01 | 1.13E+00 |
| ENSG000001 | GPR87 | 5.06E-03 | 7.58E-02 | 1.09E+00 | 1.23E-01 | 1.09E+00 |
| ENSG00000C | NAT14 | 5.08E-03 | 7.58E-02 | 1.30E+00 | 3.74E-01 | 1.30E+00 |
| ENSG000001 | HMGCR | 5.09E-03 | 7.58E-02 | 1.09E+00 | 1.22E-01 | 1.09E+00 |
| ENSG000002 | COMETT | 5.09E-03 | 7.58E-02 | 7.38E-01 | -4.38E-01 | -1.35E+00 |
| ENSG000001 | PPTC7 | 5.09E-03 | 7.58E-02 | 9.10E-01 | -1.37E-01 | -1.10E+00 |
| ENSG000001 | IFT20 | 5.10E-03 | 7.58E-02 | 1.13E+00 | 1.77E-01 | 1.13E+00 |
| ENSG000002 | UCA1 | 5.10E-03 | 7.58E-02 | 6.89E-01 | -5.37E-01 | -1.45E+00 |
| ENSG000001 | RPA1 | 5.10E-03 | 7.58E-02 | 1.08E+00 | 1.08E-01 | 1.08E+00 |
| ENSG00000C | PREP | 5.12E-03 | 7.59E-02 | 1.08E+00 | 1.07E-01 | 1.08E+00 |
| ENSG000001 | DNAI3 | 5.12E-03 | 7.59E-02 | 1.22E+00 | 2.82E-01 | 1.22E+00 |

| | | | | | | |
|---|---|---|---|---|---|---|
| ENSG000001 | RCBTB2 | 5.12E-03 | 7.59E-02 | 8.63E-01 | -2.13E-01 | -1.16E+00 |
| ENSG000001 | LITAF | 5.14E-03 | 7.60E-02 | 1.11E+00 | 1.56E-01 | 1.11E+00 |
| ENSG000001 | LAMTOR2 | 5.15E-03 | 7.60E-02 | 1.14E+00 | 1.89E-01 | 1.14E+00 |
| ENSG000001 | TXNDC11 | 5.15E-03 | 7.60E-02 | 8.75E-01 | -1.93E-01 | -1.14E+00 |
| ENSG000002 | KRT8P3 | 5.15E-03 | 7.60E-02 | 7.78E-01 | -3.63E-01 | -1.29E+00 |
| ENSG000001 | RPP25L | 5.16E-03 | 7.60E-02 | 1.29E+00 | 3.67E-01 | 1.29E+00 |
| ENSG000002 | AGAP12P | 5.18E-03 | ? | 1.88E-01 | -2.41E+00 | -5.32E+00 |
| ENSG000001 | ATPSCKMT | 5.18E-03 | 7.63E-02 | 8.48E-01 | -2.37E-01 | -1.18E+00 |
| ENSG000001 | TSPAN31 | 5.21E-03 | 7.66E-02 | 8.99E-01 | -1.53E-01 | -1.11E+00 |
| ENSG00000C | RPS5 | 5.26E-03 | 7.73E-02 | 1.11E+00 | 1.55E-01 | 1.11E+00 |
| ENSG000001 | FMOD | 5.29E-03 | 7.76E-02 | 1.37E+00 | 4.51E-01 | 1.37E+00 |
| ENSG00000C | ARF5 | 5.29E-03 | 7.76E-02 | 1.08E+00 | 1.18E-01 | 1.08E+00 |
| ENSG000001 | MYCT1 | 5.30E-03 | 7.76E-02 | 5.96E-01 | -7.46E-01 | -1.68E+00 |
| ENSG000001 | MMS19 | 5.31E-03 | 7.77E-02 | 1.11E+00 | 1.53E-01 | 1.11E+00 |
| ENSG000001 | GNPDA1 | 5.32E-03 | 7.77E-02 | 1.18E+00 | 2.43E-01 | 1.18E+00 |
| ENSG000001 | ZNF597 | 5.35E-03 | 7.81E-02 | 8.69E-01 | -2.03E-01 | -1.15E+00 |
| ENSG000001 | HBEGF | 5.36E-03 | 7.82E-02 | 8.47E-01 | -2.40E-01 | -1.18E+00 |
| ENSG000001 | TRAM1L1 | 5.37E-03 | 7.82E-02 | 1.55E+00 | 6.28E-01 | 1.55E+00 |
| ENSG000002 | --- | 5.38E-03 | 7.84E-02 | 5.66E-01 | -8.22E-01 | -1.77E+00 |
| ENSG000002 | SDHD | 5.40E-03 | 7.86E-02 | 1.08E+00 | 1.13E-01 | 1.08E+00 |
| ENSG00000C | L2HGDH | 5.41E-03 | 7.86E-02 | 1.18E+00 | 2.35E-01 | 1.18E+00 |
| ENSG000001 | SERPINI1 | 5.43E-03 | 7.89E-02 | 8.34E-01 | -2.61E-01 | -1.20E+00 |
| ENSG00000C | COBLL1 | 5.45E-03 | 7.91E-02 | 8.69E-01 | -2.02E-01 | -1.15E+00 |
| ENSG000001 | DNAJC30 | 5.47E-03 | 7.93E-02 | 1.18E+00 | 2.38E-01 | 1.18E+00 |
| ENSG000001 | FERMT1 | 5.48E-03 | 7.94E-02 | 9.06E-01 | -1.42E-01 | -1.10E+00 |
| ENSG000001 | PITRM1 | 5.51E-03 | 7.97E-02 | 9.39E-01 | -9.04E-02 | -1.06E+00 |
| ENSG00000C | CDH10 | 5.54E-03 | 8.00E-02 | 1.31E+00 | 3.91E-01 | 1.31E+00 |
| ENSG000001 | SIAH2 | 5.54E-03 | 8.00E-02 | 1.09E+00 | 1.29E-01 | 1.09E+00 |
| ENSG000001 | OSBP | 5.55E-03 | 8.00E-02 | 9.37E-01 | -9.44E-02 | -1.07E+00 |
| ENSG000001 | CENPF | 5.55E-03 | 8.00E-02 | 8.09E-01 | -3.06E-01 | -1.24E+00 |
| ENSG000001 | MAP1LC3B | 5.56E-03 | 8.00E-02 | 9.31E-01 | -1.04E-01 | -1.07E+00 |
| ENSG000001 | UBE2Z | 5.59E-03 | 8.04E-02 | 1.08E+00 | 1.06E-01 | 1.08E+00 |
| ENSG000001 | INTU | 5.61E-03 | 8.06E-02 | 1.13E+00 | 1.81E-01 | 1.13E+00 |
| ENSG000002 | --- | 5.62E-03 | ? | 1.90E+00 | 9.24E-01 | 1.90E+00 |
| ENSG000001 | BTRC | 5.63E-03 | 8.08E-02 | 9.02E-01 | -1.49E-01 | -1.11E+00 |
| ENSG000001 | DENND2D | 5.67E-03 | 8.12E-02 | 1.15E+00 | 1.98E-01 | 1.15E+00 |
| ENSG00000C | SEC62 | 5.68E-03 | 8.13E-02 | 9.04E-01 | -1.46E-01 | -1.11E+00 |
| ENSG000002 | LINC00641 | 5.69E-03 | 8.13E-02 | 8.56E-01 | -2.24E-01 | -1.17E+00 |
| ENSG000001 | TBC1D5 | 5.69E-03 | 8.13E-02 | 9.12E-01 | -1.33E-01 | -1.10E+00 |
| ENSG000001 | GABARAP | 5.70E-03 | 8.13E-02 | 1.14E+00 | 1.89E-01 | 1.14E+00 |
| ENSG000001 | LTN1 | 5.71E-03 | 8.13E-02 | 9.07E-01 | -1.40E-01 | -1.10E+00 |
| ENSG000001 | ATOX1 | 5.71E-03 | 8.13E-02 | 1.14E+00 | 1.91E-01 | 1.14E+00 |
| ENSG000001 | RREB1 | 5.75E-03 | 8.18E-02 | 9.04E-01 | -1.45E-01 | -1.11E+00 |
| ENSG000001 | FHDC1 | 5.79E-03 | 8.22E-02 | 8.53E-01 | -2.29E-01 | -1.17E+00 |
| ENSG000001 | TP53TG1 | 5.81E-03 | 8.24E-02 | 1.26E+00 | 3.36E-01 | 1.26E+00 |

| | | | | | | |
|---|---|---|---|---|---|---|
| ENSG000001 | UBLCP1 | 5.82E-03 | 8.26E-02 | 8.88E-01 | -1.72E-01 | -1.13E+00 |
| ENSG000001 | LDB1 | 5.84E-03 | 8.28E-02 | 1.09E+00 | 1.24E-01 | 1.09E+00 |
| ENSG000001 | PHACTR1 | 5.87E-03 | 8.31E-02 | 8.43E-01 | -2.46E-01 | -1.19E+00 |
| ENSG00000C | TFRC | 5.91E-03 | 8.34E-02 | 9.24E-01 | -1.14E-01 | -1.08E+00 |
| ENSG000001 | SIRPA | 5.92E-03 | 8.34E-02 | 1.11E+00 | 1.50E-01 | 1.11E+00 |
| ENSG000002 | ABCF1 | 5.92E-03 | 8.34E-02 | 1.07E+00 | 9.85E-02 | 1.07E+00 |
| ENSG000001 | CYB5D2 | 5.93E-03 | 8.34E-02 | 1.25E+00 | 3.20E-01 | 1.25E+00 |
| ENSG000001 | TSPYL5 | 5.93E-03 | 8.34E-02 | 8.83E-01 | -1.80E-01 | -1.13E+00 |
| ENSG000001 | BBC3 | 5.93E-03 | 8.34E-02 | 1.40E+00 | 4.87E-01 | 1.40E+00 |
| ENSG000001 | OGFOD2 | 5.94E-03 | 8.35E-02 | 1.32E+00 | 3.96E-01 | 1.32E+00 |
| ENSG00000C | CDIP1 | 5.97E-03 | 8.37E-02 | 1.14E+00 | 1.88E-01 | 1.14E+00 |
| ENSG000001 | THG1L | 5.97E-03 | 8.37E-02 | 1.13E+00 | 1.71E-01 | 1.13E+00 |
| ENSG000001 | CAMKK2 | 5.97E-03 | 8.37E-02 | 1.10E+00 | 1.32E-01 | 1.10E+00 |
| ENSG000001 | ZNF567 | 5.99E-03 | 8.39E-02 | 8.52E-01 | -2.32E-01 | -1.17E+00 |
| ENSG000001 | ETFB | 6.00E-03 | 8.40E-02 | 1.14E+00 | 1.95E-01 | 1.14E+00 |
| ENSG000001 | RPS27A | 6.01E-03 | 8.40E-02 | 1.13E+00 | 1.78E-01 | 1.13E+00 |
| ENSG000001 | LSM7 | 6.03E-03 | 8.42E-02 | 1.21E+00 | 2.81E-01 | 1.21E+00 |
| ENSG000001 | ABO | 6.03E-03 | 8.42E-02 | 7.86E-01 | -3.48E-01 | -1.27E+00 |
| ENSG000002 | --- | 6.05E-03 | ? | 1.94E+00 | 9.58E-01 | 1.94E+00 |
| ENSG000001 | AVL9 | 6.05E-03 | 8.44E-02 | 1.11E+00 | 1.48E-01 | 1.11E+00 |
| ENSG000001 | MRGPRX3 | 6.06E-03 | 8.44E-02 | 1.61E+00 | 6.87E-01 | 1.61E+00 |
| ENSG000002 | HMGA1P8 | 6.09E-03 | ? | 2.74E+00 | 1.46E+00 | 2.74E+00 |
| ENSG000001 | GSTCD | 6.12E-03 | 8.51E-02 | 8.37E-01 | -2.58E-01 | -1.20E+00 |
| ENSG000001 | SAXO2 | 6.13E-03 | 8.52E-02 | 1.45E+00 | 5.33E-01 | 1.45E+00 |
| ENSG000001 | HIF3A | 6.15E-03 | ? | 3.66E-01 | -1.45E+00 | -2.73E+00 |
| ENSG00000C | MAST4 | 6.16E-03 | 8.55E-02 | 1.10E+00 | 1.40E-01 | 1.10E+00 |
| ENSG000001 | LPAR6 | 6.20E-03 | 8.59E-02 | 1.20E+00 | 2.66E-01 | 1.20E+00 |
| ENSG00000C | BCL2L13 | 6.20E-03 | 8.59E-02 | 1.08E+00 | 1.12E-01 | 1.08E+00 |
| ENSG000002 | --- | 6.20E-03 | ? | 2.04E+00 | 1.03E+00 | 2.04E+00 |
| ENSG00000C | DLG1 | 6.21E-03 | 8.60E-02 | 8.70E-01 | -2.01E-01 | -1.15E+00 |
| ENSG000001 | HNRNPA0 | 6.22E-03 | 8.60E-02 | 1.12E+00 | 1.68E-01 | 1.12E+00 |
| ENSG000001 | ADAMTS13 | 6.24E-03 | 8.62E-02 | 1.63E+00 | 7.08E-01 | 1.63E+00 |
| ENSG000002 | --- | 6.27E-03 | ? | 6.57E-01 | -6.06E-01 | -1.52E+00 |
| ENSG000001 | SLC2A12 | 6.28E-03 | 8.66E-02 | 1.34E+00 | 4.27E-01 | 1.34E+00 |
| ENSG000001 | PTGR1 | 6.28E-03 | 8.66E-02 | 8.96E-01 | -1.58E-01 | -1.12E+00 |
| ENSG000001 | CHD3 | 6.29E-03 | 8.66E-02 | 9.39E-01 | -9.01E-02 | -1.06E+00 |
| ENSG000002 | --- | 6.31E-03 | 8.69E-02 | 1.32E+00 | 4.02E-01 | 1.32E+00 |
| ENSG000002 | LIX1L | 6.32E-03 | 8.69E-02 | 1.10E+00 | 1.41E-01 | 1.10E+00 |
| ENSG000001 | NPC2 | 6.34E-03 | 8.71E-02 | 1.11E+00 | 1.51E-01 | 1.11E+00 |
| ENSG000002 | ZNF542P | 6.35E-03 | 8.71E-02 | 8.00E-01 | -3.23E-01 | -1.25E+00 |
| ENSG000001 | MCU | 6.36E-03 | 8.72E-02 | 8.92E-01 | -1.64E-01 | -1.12E+00 |
| ENSG000001 | INTS14 | 6.37E-03 | 8.72E-02 | 1.09E+00 | 1.21E-01 | 1.09E+00 |
| ENSG000001 | RARA | 6.39E-03 | 8.74E-02 | 8.92E-01 | -1.65E-01 | -1.12E+00 |
| ENSG00000C | CYP24A1 | 6.39E-03 | 8.74E-02 | 8.51E-01 | -2.33E-01 | -1.18E+00 |
| ENSG000001 | NOD1 | 6.41E-03 | 8.75E-02 | 1.17E+00 | 2.33E-01 | 1.17E+00 |

| | | | | | | |
|---|---|---|---|---|---|---|
| ENSG000001 | DAB2IP | 6.42E-03 | 8.75E-02 | 1.11E+00 | 1.52E-01 | 1.11E+00 |
| ENSG000002 | TBCE | 6.43E-03 | 8.76E-02 | 8.18E-01 | -2.90E-01 | -1.22E+00 |
| ENSG00000C | RALA | 6.46E-03 | 8.79E-02 | 9.23E-01 | -1.16E-01 | -1.08E+00 |
| ENSG000002 | ZNF542P | 6.46E-03 | 8.79E-02 | 8.91E-01 | -1.67E-01 | -1.12E+00 |
| ENSG000001 | RPS12 | 6.49E-03 | 8.81E-02 | 1.18E+00 | 2.37E-01 | 1.18E+00 |
| ENSG000001 | ZYX | 6.49E-03 | 8.81E-02 | 1.08E+00 | 1.14E-01 | 1.08E+00 |
| ENSG000001 | DNAJB9 | 6.50E-03 | 8.81E-02 | 9.18E-01 | -1.24E-01 | -1.09E+00 |
| ENSG00000C | ST6GAL1 | 6.50E-03 | 8.81E-02 | 1.11E+00 | 1.55E-01 | 1.11E+00 |
| ENSG000002 | LUCAT1 | 6.52E-03 | 8.82E-02 | 6.38E-01 | -6.49E-01 | -1.57E+00 |
| ENSG000001 | RPLP1 | 6.52E-03 | 8.82E-02 | 1.19E+00 | 2.56E-01 | 1.19E+00 |
| ENSG000001 | TECPR2 | 6.53E-03 | 8.82E-02 | 1.11E+00 | 1.51E-01 | 1.11E+00 |
| ENSG000001 | IER3 | 6.54E-03 | 8.83E-02 | 8.82E-01 | -1.81E-01 | -1.13E+00 |
| ENSG000001 | STN1 | 6.56E-03 | 8.85E-02 | 8.91E-01 | -1.67E-01 | -1.12E+00 |
| ENSG000001 | ZNF618 | 6.57E-03 | 8.85E-02 | 8.88E-01 | -1.71E-01 | -1.13E+00 |
| ENSG000001 | ZNF791 | 6.60E-03 | 8.89E-02 | 8.92E-01 | -1.65E-01 | -1.12E+00 |
| ENSG000001 | WNT7A | 6.62E-03 | 8.91E-02 | 8.90E-01 | -1.67E-01 | -1.12E+00 |
| ENSG000001 | PAK6 | 6.63E-03 | 8.91E-02 | 1.17E+00 | 2.24E-01 | 1.17E+00 |
| ENSG00000C | EDEM2 | 6.64E-03 | 8.92E-02 | 1.13E+00 | 1.77E-01 | 1.13E+00 |
| ENSG000001 | GFM1 | 6.67E-03 | 8.94E-02 | 9.13E-01 | -1.31E-01 | -1.10E+00 |
| ENSG000001 | GTPBP4 | 6.67E-03 | 8.94E-02 | 9.03E-01 | -1.47E-01 | -1.11E+00 |
| ENSG000001 | NXN | 6.70E-03 | 8.97E-02 | 1.13E+00 | 1.74E-01 | 1.13E+00 |
| ENSG000002 | MLLT11 | 6.72E-03 | 8.98E-02 | 8.91E-01 | -1.67E-01 | -1.12E+00 |
| ENSG000002 | KRT8P45 | 6.72E-03 | 8.98E-02 | 6.76E-01 | -5.65E-01 | -1.48E+00 |
| ENSG000001 | ITGB8 | 6.73E-03 | 8.98E-02 | 1.16E+00 | 2.12E-01 | 1.16E+00 |
| ENSG000001 | SLC20A1 | 6.74E-03 | 8.99E-02 | 9.15E-01 | -1.28E-01 | -1.09E+00 |
| ENSG00000C | TRAF1 | 6.78E-03 | 9.03E-02 | 1.36E+00 | 4.41E-01 | 1.36E+00 |
| ENSG000002 | CYP4F26P | 6.81E-03 | 9.05E-02 | 1.39E+00 | 4.80E-01 | 1.39E+00 |
| ENSG00000C | SNX29 | 6.81E-03 | 9.05E-02 | 1.09E+00 | 1.24E-01 | 1.09E+00 |
| ENSG000001 | PGAP3 | 6.81E-03 | 9.05E-02 | 1.30E+00 | 3.77E-01 | 1.30E+00 |
| ENSG000002 | SYT15-AS1 | 6.83E-03 | 9.06E-02 | 1.33E+00 | 4.09E-01 | 1.33E+00 |
| ENSG00000C | SKAP2 | 6.85E-03 | 9.08E-02 | 1.11E+00 | 1.46E-01 | 1.11E+00 |
| ENSG000001 | SCAMP5 | 6.85E-03 | 9.08E-02 | 1.32E+00 | 3.99E-01 | 1.32E+00 |
| ENSG000001 | VEPH1 | 6.88E-03 | 9.11E-02 | 7.79E-01 | -3.61E-01 | -1.28E+00 |
| ENSG000001 | COLGALT1 | 6.88E-03 | 9.11E-02 | 1.11E+00 | 1.50E-01 | 1.11E+00 |
| ENSG000001 | ZNF140 | 6.95E-03 | 9.19E-02 | 8.83E-01 | -1.80E-01 | -1.13E+00 |
| ENSG000001 | MYEF2 | 6.98E-03 | 9.21E-02 | 8.40E-01 | -2.51E-01 | -1.19E+00 |
| ENSG000001 | PCTP | 7.00E-03 | 9.23E-02 | 1.08E+00 | 1.15E-01 | 1.08E+00 |
| ENSG000001 | MYH14 | 7.00E-03 | 9.23E-02 | 1.35E+00 | 4.37E-01 | 1.35E+00 |
| ENSG000001 | EVI2B | 7.01E-03 | 9.23E-02 | 6.33E-01 | -6.59E-01 | -1.58E+00 |
| ENSG000002 | SHISA9 | 7.01E-03 | ? | 2.45E+00 | 1.29E+00 | 2.45E+00 |
| ENSG000001 | SLC35G1 | 7.05E-03 | 9.28E-02 | 1.20E+00 | 2.61E-01 | 1.20E+00 |
| ENSG000001 | DHFR2 | 7.10E-03 | 9.33E-02 | 1.19E+00 | 2.51E-01 | 1.19E+00 |
| ENSG000001 | CDH6 | 7.11E-03 | 9.33E-02 | 7.22E-01 | -4.70E-01 | -1.39E+00 |
| ENSG000001 | EIF3K | 7.12E-03 | 9.33E-02 | 1.10E+00 | 1.44E-01 | 1.10E+00 |
| ENSG000001 | UBR3 | 7.12E-03 | 9.33E-02 | 8.98E-01 | -1.55E-01 | -1.11E+00 |

| | | | | | | |
|---|---|---|---|---|---|---|
| ENSG000002 | MIR3936HG | 7.12E-03 | 9.33E-02 | 7.86E-01 | -3.47E-01 | -1.27E+00 |
| ENSG000001 | ZNF267 | 7.14E-03 | 9.33E-02 | 8.60E-01 | -2.18E-01 | -1.16E+00 |
| ENSG000001 | LRRC56 | 7.14E-03 | 9.33E-02 | 1.48E+00 | 5.62E-01 | 1.48E+00 |
| ENSG000001 | TPST2 | 7.15E-03 | 9.33E-02 | 1.09E+00 | 1.24E-01 | 1.09E+00 |
| ENSG000001 | MRPL11 | 7.19E-03 | 9.38E-02 | 1.15E+00 | 2.03E-01 | 1.15E+00 |
| ENSG000001 | ZNF512B | 7.21E-03 | 9.40E-02 | 1.15E+00 | 1.97E-01 | 1.15E+00 |
| ENSG000001 | PARVB | 7.25E-03 | 9.44E-02 | 1.11E+00 | 1.54E-01 | 1.11E+00 |
| ENSG000001 | DOCK4 | 7.27E-03 | 9.44E-02 | 8.58E-01 | -2.21E-01 | -1.17E+00 |
| ENSG000002 | MACO1 | 7.27E-03 | 9.44E-02 | 8.90E-01 | -1.68E-01 | -1.12E+00 |
| ENSG000001 | TOR1AIP1 | 7.29E-03 | 9.46E-02 | 9.02E-01 | -1.49E-01 | -1.11E+00 |
| ENSG000002 | LUARIS | 7.30E-03 | 9.46E-02 | 1.29E+00 | 3.68E-01 | 1.29E+00 |
| ENSG000001 | BRD4 | 7.30E-03 | 9.46E-02 | 8.76E-01 | -1.90E-01 | -1.14E+00 |
| ENSG000001 | LARP1B | 7.31E-03 | 9.47E-02 | 8.63E-01 | -2.13E-01 | -1.16E+00 |
| ENSG000001 | ARAP1 | 7.32E-03 | 9.48E-02 | 1.13E+00 | 1.79E-01 | 1.13E+00 |
| ENSG000001 | SOWAHC | 7.35E-03 | 9.50E-02 | 8.76E-01 | -1.91E-01 | -1.14E+00 |
| ENSG000001 | RPL37A | 7.37E-03 | 9.51E-02 | 1.12E+00 | 1.64E-01 | 1.12E+00 |
| ENSG000001 | ZNF680 | 7.37E-03 | 9.51E-02 | 8.23E-01 | -2.81E-01 | -1.22E+00 |
| ENSG000001 | ELF4 | 7.39E-03 | 9.52E-02 | 1.07E+00 | 9.92E-02 | 1.07E+00 |
| ENSG00000C | PHKA1 | 7.43E-03 | 9.57E-02 | 1.11E+00 | 1.51E-01 | 1.11E+00 |
| ENSG000001 | ZNG1C | 7.47E-03 | 9.62E-02 | 8.72E-01 | -1.97E-01 | -1.15E+00 |
| ENSG000001 | AP3M1 | 7.50E-03 | 9.64E-02 | 9.29E-01 | -1.06E-01 | -1.08E+00 |
| ENSG000001 | ZKSCAN8 | 7.53E-03 | 9.67E-02 | 9.01E-01 | -1.50E-01 | -1.11E+00 |
| ENSG000002 | CHCHD10 | 7.54E-03 | 9.67E-02 | 1.37E+00 | 4.57E-01 | 1.37E+00 |
| ENSG000001 | CCNH | 7.55E-03 | 9.68E-02 | 9.06E-01 | -1.42E-01 | -1.10E+00 |
| ENSG00000C | NDST1 | 7.56E-03 | 9.69E-02 | 1.12E+00 | 1.68E-01 | 1.12E+00 |
| ENSG000002 | SERPINB4 | 7.58E-03 | 9.70E-02 | 6.74E-01 | -5.69E-01 | -1.48E+00 |
| ENSG000001 | GJB6 | 7.59E-03 | 9.70E-02 | 1.13E+00 | 1.73E-01 | 1.13E+00 |
| ENSG000001 | FMNL1 | 7.60E-03 | 9.70E-02 | 7.85E-01 | -3.50E-01 | -1.27E+00 |
| ENSG00000C | MRTO4 | 7.61E-03 | 9.70E-02 | 1.08E+00 | 1.11E-01 | 1.08E+00 |
| ENSG000001 | PPIF | 7.61E-03 | 9.70E-02 | 1.12E+00 | 1.61E-01 | 1.12E+00 |
| ENSG000001 | TBC1D4 | 7.61E-03 | 9.70E-02 | 1.09E+00 | 1.29E-01 | 1.09E+00 |
| ENSG000001 | LGR6 | 7.66E-03 | 9.74E-02 | 6.77E-01 | -5.62E-01 | -1.48E+00 |
| ENSG000001 | LNPK | 7.66E-03 | 9.74E-02 | 9.08E-01 | -1.39E-01 | -1.10E+00 |
| ENSG000001 | DUSP7 | 7.70E-03 | 9.79E-02 | 1.09E+00 | 1.23E-01 | 1.09E+00 |
| ENSG000002 | --- | 7.72E-03 | 9.80E-02 | 7.37E-01 | -4.40E-01 | -1.36E+00 |
| ENSG000001 | ENTPD6 | 7.76E-03 | 9.84E-02 | 1.14E+00 | 1.86E-01 | 1.14E+00 |
| ENSG00000C | MACROH2A2 | 7.76E-03 | 9.84E-02 | 1.13E+00 | 1.82E-01 | 1.13E+00 |
| ENSG000001 | SPRY4 | 7.77E-03 | 9.84E-02 | 8.99E-01 | -1.53E-01 | -1.11E+00 |
| ENSG000001 | PANK3 | 7.78E-03 | 9.84E-02 | 9.07E-01 | -1.41E-01 | -1.10E+00 |
| ENSG000001 | INPP4B | 7.80E-03 | 9.86E-02 | 8.76E-01 | -1.91E-01 | -1.14E+00 |
| ENSG000001 | KRT8 | 7.82E-03 | 9.87E-02 | 8.25E-01 | -2.77E-01 | -1.21E+00 |
| ENSG000001 | PI4K2A | 7.82E-03 | 9.87E-02 | 1.11E+00 | 1.47E-01 | 1.11E+00 |
| ENSG000001 | ZNF329 | 7.83E-03 | 9.87E-02 | 1.10E+00 | 1.38E-01 | 1.10E+00 |
| ENSG000001 | ING2 | 7.84E-03 | 9.87E-02 | 8.73E-01 | -1.96E-01 | -1.15E+00 |
| ENSG000001 | PKP4 | 7.85E-03 | 9.87E-02 | 9.26E-01 | -1.11E-01 | -1.08E+00 |

| | | | | | | |
|---|---|---|---|---|---|---|
| ENSG000001 | CASP1 | 7.85E-03 | 9.87E-02 | 1.16E+00 | 2.08E-01 | 1.16E+00 |
| ENSG000001 | ATP5MJ | 7.93E-03 | 9.96E-02 | 1.11E+00 | 1.45E-01 | 1.11E+00 |
| ENSG000002 | NPTXR | 7.96E-03 | 9.98E-02 | 1.36E+00 | 4.42E-01 | 1.36E+00 |
| ENSG000001 | TBCEL | 7.96E-03 | 9.98E-02 | 1.11E+00 | 1.49E-01 | 1.11E+00 |
| ENSG000001 | UCN2 | 7.96E-03 | 9.98E-02 | 8.14E-01 | -2.96E-01 | -1.23E+00 |
| ENSG000001 | PRORP | 7.98E-03 | 9.98E-02 | 8.79E-01 | -1.86E-01 | -1.14E+00 |
| ENSG00000C | HSD17B2 | 7.98E-03 | 9.98E-02 | 7.65E-01 | -3.86E-01 | -1.31E+00 |
| ENSG000001 | RPL19 | 7.99E-03 | 9.98E-02 | 1.08E+00 | 1.06E-01 | 1.08E+00 |
| ENSG000001 | IL18 | 8.02E-03 | 1.00E-01 | 1.09E+00 | 1.28E-01 | 1.09E+00 |
| ENSG00000C | COL23A1 | 8.04E-03 | ? | 1.98E+00 | 9.85E-01 | 1.98E+00 |
| ENSG000001 | RPL29 | 8.08E-03 | 1.01E-01 | 1.16E+00 | 2.15E-01 | 1.16E+00 |
| ENSG000001 | OLFM1 | 8.10E-03 | 1.01E-01 | 1.52E+00 | 6.02E-01 | 1.52E+00 |
| ENSG000001 | ZDHHC16 | 8.11E-03 | 1.01E-01 | 9.07E-01 | -1.41E-01 | -1.10E+00 |
| ENSG000001 | RPL10A | 8.11E-03 | 1.01E-01 | 1.09E+00 | 1.29E-01 | 1.09E+00 |
| ENSG000001 | GPATCH3 | 8.16E-03 | 1.01E-01 | 1.12E+00 | 1.66E-01 | 1.12E+00 |
| ENSG000001 | ERF | 8.19E-03 | 1.02E-01 | 1.12E+00 | 1.67E-01 | 1.12E+00 |
| ENSG000001 | ZNF697 | 8.21E-03 | 1.02E-01 | 9.28E-01 | -1.08E-01 | -1.08E+00 |
| ENSG000001 | HSD11B1 | 8.25E-03 | 1.02E-01 | 6.88E-01 | -5.39E-01 | -1.45E+00 |
| ENSG000001 | FNBP1 | 8.26E-03 | 1.02E-01 | 8.91E-01 | -1.66E-01 | -1.12E+00 |
| ENSG000002 | --- | 8.26E-03 | ? | 6.57E-01 | -6.06E-01 | -1.52E+00 |
| ENSG000001 | PHYH | 8.32E-03 | 1.03E-01 | 1.14E+00 | 1.88E-01 | 1.14E+00 |
| ENSG000001 | KMT5C | 8.38E-03 | 1.04E-01 | 1.29E+00 | 3.64E-01 | 1.29E+00 |
| ENSG000001 | LRATD1 | 8.49E-03 | 1.05E-01 | 1.16E+00 | 2.10E-01 | 1.16E+00 |
| ENSG00000C | ATRN | 8.49E-03 | 1.05E-01 | 1.09E+00 | 1.22E-01 | 1.09E+00 |
| ENSG000001 | OTUD1 | 8.51E-03 | 1.05E-01 | 1.18E+00 | 2.36E-01 | 1.18E+00 |
| ENSG000001 | RPS10 | 8.51E-03 | 1.05E-01 | 1.15E+00 | 2.03E-01 | 1.15E+00 |
| ENSG000001 | KPNA5 | 8.52E-03 | 1.05E-01 | 8.49E-01 | -2.35E-01 | -1.18E+00 |
| ENSG000001 | GOLGA4 | 8.53E-03 | 1.05E-01 | 8.67E-01 | -2.06E-01 | -1.15E+00 |
| ENSG000001 | MARCHF6 | 8.53E-03 | 1.05E-01 | 9.48E-01 | -7.64E-02 | -1.05E+00 |
| ENSG000001 | DCAF7 | 8.56E-03 | 1.05E-01 | 1.07E+00 | 1.04E-01 | 1.07E+00 |
| ENSG000001 | STARD13 | 8.60E-03 | 1.05E-01 | 9.23E-01 | -1.16E-01 | -1.08E+00 |
| ENSG000001 | CEMIP2 | 8.61E-03 | 1.05E-01 | 8.00E-01 | -3.22E-01 | -1.25E+00 |
| ENSG000001 | NUMB | 8.67E-03 | 1.06E-01 | 9.22E-01 | -1.17E-01 | -1.08E+00 |
| ENSG000001 | MAP3K12 | 8.68E-03 | 1.06E-01 | 1.20E+00 | 2.64E-01 | 1.20E+00 |
| ENSG00000C | MGLL | 8.69E-03 | 1.06E-01 | 8.80E-01 | -1.85E-01 | -1.14E+00 |
| ENSG000001 | PSMD12 | 8.70E-03 | 1.06E-01 | 9.16E-01 | -1.26E-01 | -1.09E+00 |
| ENSG000001 | HNRNPH1 | 8.71E-03 | 1.06E-01 | 8.64E-01 | -2.11E-01 | -1.16E+00 |
| ENSG000001 | KIF20A | 8.73E-03 | 1.06E-01 | 7.60E-01 | -3.95E-01 | -1.32E+00 |
| ENSG000001 | RPL23 | 8.73E-03 | 1.06E-01 | 1.09E+00 | 1.18E-01 | 1.09E+00 |
| ENSG00000C | CLCN4 | 8.77E-03 | 1.07E-01 | 1.23E+00 | 2.98E-01 | 1.23E+00 |
| ENSG000001 | HSD17B12 | 8.78E-03 | 1.07E-01 | 9.17E-01 | -1.25E-01 | -1.09E+00 |
| ENSG000001 | ZDHHC4 | 8.78E-03 | 1.07E-01 | 1.14E+00 | 1.87E-01 | 1.14E+00 |
| ENSG00000C | RHOA | 8.79E-03 | 1.07E-01 | 1.06E+00 | 8.28E-02 | 1.06E+00 |
| ENSG000001 | BANK1 | 8.87E-03 | 1.08E-01 | 1.33E+00 | 4.17E-01 | 1.33E+00 |
| ENSG00000C | H6PD | 8.89E-03 | 1.08E-01 | 1.15E+00 | 2.07E-01 | 1.15E+00 |

| | | | | | | |
|---|---|---|---|---|---|---|
| ENSG000001 | AADAC | 8.89E-03 | 1.08E-01 | 6.62E-01 | -5.96E-01 | -1.51E+00 |
| ENSG000000 | ETV7 | 8.90E-03 | ? | 3.21E+00 | 1.68E+00 | 3.21E+00 |
| ENSG000001 | YARS2 | 8.91E-03 | 1.08E-01 | 1.10E+00 | 1.37E-01 | 1.10E+00 |
| ENSG000000 | TIGAR | 8.91E-03 | 1.08E-01 | 1.17E+00 | 2.29E-01 | 1.17E+00 |
| ENSG000001 | CPVL | 8.93E-03 | 1.08E-01 | 1.55E+00 | 6.32E-01 | 1.55E+00 |
| ENSG000002 | LINC00926 | 8.95E-03 | 1.08E-01 | 6.55E-01 | -6.11E-01 | -1.53E+00 |
| ENSG000002 | SCO2 | 8.96E-03 | 1.08E-01 | 1.31E+00 | 3.87E-01 | 1.31E+00 |
| ENSG000001 | PODXL | 9.01E-03 | 1.09E-01 | 6.62E-01 | -5.96E-01 | -1.51E+00 |
| ENSG000001 | UNC5B | 9.04E-03 | 1.09E-01 | 1.19E+00 | 2.54E-01 | 1.19E+00 |
| ENSG000001 | CASP4 | 9.05E-03 | 1.09E-01 | 9.08E-01 | -1.38E-01 | -1.10E+00 |
| ENSG000001 | FAM83A | 9.10E-03 | 1.09E-01 | 8.86E-01 | -1.75E-01 | -1.13E+00 |
| ENSG000001 | PRPF38B | 9.10E-03 | 1.09E-01 | 8.65E-01 | -2.09E-01 | -1.16E+00 |
| ENSG000001 | VHL | 9.14E-03 | 1.10E-01 | 1.12E+00 | 1.65E-01 | 1.12E+00 |
| ENSG000001 | LAMP1 | 9.14E-03 | 1.10E-01 | 1.07E+00 | 9.81E-02 | 1.07E+00 |
| ENSG000001 | HMOX2 | 9.18E-03 | 1.10E-01 | 1.08E+00 | 1.08E-01 | 1.08E+00 |
| ENSG000001 | FIP1L1 | 9.21E-03 | 1.10E-01 | 8.97E-01 | -1.57E-01 | -1.12E+00 |
| ENSG000001 | IL1RAP | 9.24E-03 | 1.10E-01 | 1.16E+00 | 2.09E-01 | 1.16E+00 |
| ENSG000001 | SIRT3 | 9.24E-03 | 1.10E-01 | 1.19E+00 | 2.47E-01 | 1.19E+00 |
| ENSG000000 | BAK1 | 9.26E-03 | 1.11E-01 | 1.19E+00 | 2.50E-01 | 1.19E+00 |
| ENSG000001 | MEIOC | 9.27E-03 | ? | 5.34E-01 | -9.05E-01 | -1.87E+00 |
| ENSG000001 | CSRNP2 | 9.29E-03 | 1.11E-01 | 9.24E-01 | -1.14E-01 | -1.08E+00 |
| ENSG000000 | FAP | 9.32E-03 | 1.11E-01 | 8.13E-01 | -2.99E-01 | -1.23E+00 |
| ENSG000001 | TACC2 | 9.33E-03 | 1.11E-01 | 1.09E+00 | 1.18E-01 | 1.09E+00 |
| ENSG000001 | RAB5B | 9.34E-03 | 1.11E-01 | 1.06E+00 | 8.20E-02 | 1.06E+00 |
| ENSG000001 | BACE2 | 9.38E-03 | 1.12E-01 | 1.08E+00 | 1.11E-01 | 1.08E+00 |
| ENSG000001 | MT1G | 9.38E-03 | ? | 5.59E-01 | -8.39E-01 | -1.79E+00 |
| ENSG000001 | KLF10 | 9.39E-03 | 1.12E-01 | 9.19E-01 | -1.21E-01 | -1.09E+00 |
| ENSG000000 | MTMR7 | 9.40E-03 | 1.12E-01 | 7.62E-01 | -3.92E-01 | -1.31E+00 |
| ENSG000001 | ANKFY1 | 9.42E-03 | 1.12E-01 | 1.07E+00 | 1.00E-01 | 1.07E+00 |
| ENSG000001 | APBB1 | 9.43E-03 | 1.12E-01 | 1.26E+00 | 3.38E-01 | 1.26E+00 |
| ENSG000002 | C2orf81 | 9.44E-03 | 1.12E-01 | 1.34E+00 | 4.23E-01 | 1.34E+00 |
| ENSG000001 | SPRING1 | 9.45E-03 | 1.12E-01 | 1.06E+00 | 8.41E-02 | 1.06E+00 |
| ENSG000002 | RAB7B | 9.45E-03 | 1.12E-01 | 1.18E+00 | 2.43E-01 | 1.18E+00 |
| ENSG000001 | NLRX1 | 9.46E-03 | 1.12E-01 | 1.19E+00 | 2.54E-01 | 1.19E+00 |
| ENSG000001 | SPINK6 | 9.47E-03 | ? | 2.77E+00 | 1.47E+00 | 2.77E+00 |
| ENSG000001 | RPSA | 9.49E-03 | 1.12E-01 | 1.09E+00 | 1.20E-01 | 1.09E+00 |
| ENSG000001 | LOXL2 | 9.50E-03 | 1.12E-01 | 8.51E-01 | -2.33E-01 | -1.18E+00 |
| ENSG000001 | LINC02910 | 9.51E-03 | 1.12E-01 | 8.22E-01 | -2.83E-01 | -1.22E+00 |
| ENSG000000 | CFH | 9.55E-03 | 1.12E-01 | 9.04E-01 | -1.45E-01 | -1.11E+00 |
| ENSG000002 | --- | 9.57E-03 | 1.13E-01 | 1.17E+00 | 2.23E-01 | 1.17E+00 |
| ENSG000000 | PAG1 | 9.60E-03 | 1.13E-01 | 8.70E-01 | -2.01E-01 | -1.15E+00 |
| ENSG000001 | PARD3B | 9.61E-03 | 1.13E-01 | 7.82E-01 | -3.54E-01 | -1.28E+00 |
| ENSG000001 | SELPLG | 9.62E-03 | 1.13E-01 | 8.25E-01 | -2.78E-01 | -1.21E+00 |
| ENSG000001 | TNNI1 | 9.66E-03 | 1.13E-01 | 1.24E+00 | 3.10E-01 | 1.24E+00 |
| ENSG000001 | ATP5PB | 9.67E-03 | 1.13E-01 | 1.06E+00 | 8.79E-02 | 1.06E+00 |

| | | | | | | | |
|---|---|---|---|---|---|---|---|
| ENSG000002 | TMEM141 | 9.71E-03 | | 1.14E-01 | 1.20E+00 | 2.66E-01 | 1.20E+00 |
| ENSG000001 | MPI | 9.72E-03 | | 1.14E-01 | 1.11E+00 | 1.50E-01 | 1.11E+00 |
| ENSG000001 | MEX3B | 9.73E-03 | | 1.14E-01 | 1.29E+00 | 3.70E-01 | 1.29E+00 |
| ENSG000001 | ARHGAP18 | 9.73E-03 | | 1.14E-01 | 8.46E-01 | -2.41E-01 | -1.18E+00 |
| ENSG000002 | NUDT19 | 9.74E-03 | | 1.14E-01 | 8.94E-01 | -1.61E-01 | -1.12E+00 |
| ENSG000002 | --- | 9.75E-03 | ? | | 2.45E+00 | 1.29E+00 | 2.45E+00 |
| ENSG00000C | SSH1 | 9.79E-03 | | 1.14E-01 | 1.08E+00 | 1.09E-01 | 1.08E+00 |
| ENSG000001 | EXOC6B | 9.79E-03 | | 1.14E-01 | 1.07E+00 | 9.55E-02 | 1.07E+00 |
| ENSG000001 | THBS1 | 9.79E-03 | | 1.14E-01 | 8.90E-01 | -1.68E-01 | -1.12E+00 |
| ENSG000001 | PRICKLE1 | 9.82E-03 | | 1.14E-01 | 1.28E+00 | 3.55E-01 | 1.28E+00 |
| ENSG000001 | DDX10 | 9.82E-03 | | 1.14E-01 | 8.73E-01 | -1.96E-01 | -1.15E+00 |
| ENSG000001 | RPS13 | 9.84E-03 | | 1.14E-01 | 1.10E+00 | 1.39E-01 | 1.10E+00 |
| ENSG000001 | UTP15 | 9.84E-03 | | 1.14E-01 | 9.10E-01 | -1.36E-01 | -1.10E+00 |
| ENSG000002 | --- | 9.86E-03 | ? | | 3.94E-01 | -1.34E+00 | -2.54E+00 |
| ENSG000001 | AREL1 | 9.88E-03 | | 1.14E-01 | 1.08E+00 | 1.15E-01 | 1.08E+00 |
| ENSG000001 | HSPA13 | 9.88E-03 | | 1.14E-01 | 9.16E-01 | -1.27E-01 | -1.09E+00 |
| ENSG000002 | TRIM6-TRIM | 9.90E-03 | ? | | 3.07E+00 | 1.62E+00 | 3.07E+00 |
| ENSG000001 | KIF6 | 9.93E-03 | ? | | 1.85E+00 | 8.90E-01 | 1.85E+00 |
| ENSG000002 | --- | 9.93E-03 | ? | | 6.34E-01 | -6.57E-01 | -1.58E+00 |
| ENSG000001 | COMMD1 | 9.94E-03 | | 1.15E-01 | 1.12E+00 | 1.65E-01 | 1.12E+00 |
| ENSG000001 | SRSF4 | 9.98E-03 | | 1.15E-01 | 9.01E-01 | -1.51E-01 | -1.11E+00 |
| ENSG000002 | DUSP5-DT | 9.98E-03 | ? | | 1.73E+00 | 7.94E-01 | 1.73E+00 |
| ENSG000002 | ARHGAP5-AS | 9.99E-03 | | 1.15E-01 | 1.22E+00 | 2.87E-01 | 1.22E+00 |
| ENSG000001 | IWS1 | 1.00E-02 | | 1.15E-01 | 9.27E-01 | -1.09E-01 | -1.08E+00 |
| ENSG000002 | RPL41 | 1.00E-02 | | 1.15E-01 | 1.12E+00 | 1.58E-01 | 1.12E+00 |
| ENSG000001 | SRD5A1 | 1.00E-02 | | 1.15E-01 | 9.09E-01 | -1.38E-01 | -1.10E+00 |
| ENSG000001 | RABGEF1 | 1.00E-02 | | 1.15E-01 | 9.34E-01 | -9.89E-02 | -1.07E+00 |
| ENSG000001 | AMN1 | 1.00E-02 | | 1.15E-01 | 8.05E-01 | -3.13E-01 | -1.24E+00 |
| ENSG00000C | HEATR6 | 1.00E-02 | | 1.15E-01 | 1.09E+00 | 1.29E-01 | 1.09E+00 |
| ENSG000001 | RPS19 | 1.01E-02 | | 1.16E-01 | 1.18E+00 | 2.35E-01 | 1.18E+00 |
| ENSG000001 | RNF152 | 1.01E-02 | | 1.16E-01 | 1.14E+00 | 1.93E-01 | 1.14E+00 |
| ENSG000001 | IFNGR2 | 1.01E-02 | | 1.16E-01 | 1.11E+00 | 1.47E-01 | 1.11E+00 |
| ENSG00000C | CC2D2A | 1.01E-02 | | 1.16E-01 | 1.18E+00 | 2.37E-01 | 1.18E+00 |
| ENSG000002 | RACK1 | 1.01E-02 | | 1.16E-01 | 1.11E+00 | 1.55E-01 | 1.11E+00 |
| ENSG000002 | FZD10-AS1 | 1.01E-02 | ? | | 1.78E+00 | 8.34E-01 | 1.78E+00 |
| ENSG00000C | ZCCHC8 | 1.02E-02 | | 1.16E-01 | 8.92E-01 | -1.64E-01 | -1.12E+00 |
| ENSG000001 | DIPK1A | 1.02E-02 | | 1.16E-01 | 1.11E+00 | 1.49E-01 | 1.11E+00 |
| ENSG000001 | SLC16A1 | 1.02E-02 | | 1.16E-01 | 1.07E+00 | 9.84E-02 | 1.07E+00 |
| ENSG000001 | OPCML | 1.02E-02 | ? | | 1.61E+00 | 6.83E-01 | 1.61E+00 |
| ENSG000001 | TRIML2 | 1.02E-02 | | 1.16E-01 | 8.01E-01 | -3.19E-01 | -1.25E+00 |
| ENSG000001 | SMIM10L2A | 1.02E-02 | | 1.16E-01 | 1.21E+00 | 2.75E-01 | 1.21E+00 |
| ENSG000001 | ZNF362 | 1.02E-02 | | 1.16E-01 | 1.20E+00 | 2.65E-01 | 1.20E+00 |
| ENSG000001 | CTDSPL | 1.02E-02 | | 1.17E-01 | 1.10E+00 | 1.34E-01 | 1.10E+00 |
| ENSG000001 | APBB2 | 1.03E-02 | | 1.17E-01 | 1.09E+00 | 1.26E-01 | 1.09E+00 |
| ENSG000001 | ACO1 | 1.03E-02 | | 1.17E-01 | 1.07E+00 | 1.02E-01 | 1.07E+00 |

| | | | | | | |
|---|---|---|---|---|---|---|
| ENSG000002 | ALDH1A3-AS | 1.03E-02 | 1.17E-01 | 8.12E-01 | -3.01E-01 | -1.23E+00 |
| ENSG000001 | RPL3 | 1.03E-02 | 1.17E-01 | 1.12E+00 | 1.63E-01 | 1.12E+00 |
| ENSG000001 | UBR1 | 1.03E-02 | 1.17E-01 | 9.13E-01 | -1.31E-01 | -1.10E+00 |
| ENSG00000C | CEP68 | 1.03E-02 | 1.17E-01 | 1.12E+00 | 1.59E-01 | 1.12E+00 |
| ENSG000002 | CNOT6LP1 | 1.03E-02 | 1.17E-01 | 1.39E+00 | 4.75E-01 | 1.39E+00 |
| ENSG000002 | --- | 1.04E-02 | ? | 2.83E+00 | 1.50E+00 | 2.83E+00 |
| ENSG000001 | INHBA | 1.04E-02 | 1.18E-01 | 8.12E-01 | -3.01E-01 | -1.23E+00 |
| ENSG000001 | MTDH | 1.04E-02 | 1.18E-01 | 9.26E-01 | -1.12E-01 | -1.08E+00 |
| ENSG000002 | RPL23AP49 | 1.04E-02 | 1.18E-01 | 1.37E+00 | 4.56E-01 | 1.37E+00 |
| ENSG000001 | AHCY | 1.04E-02 | 1.18E-01 | 1.12E+00 | 1.58E-01 | 1.12E+00 |
| ENSG000001 | THADA | 1.05E-02 | 1.18E-01 | 1.08E+00 | 1.10E-01 | 1.08E+00 |
| ENSG00000C | PPP1R3F | 1.05E-02 | 1.18E-01 | 1.48E+00 | 5.69E-01 | 1.48E+00 |
| ENSG000002 | --- | 1.05E-02 | 1.18E-01 | 6.71E-01 | -5.75E-01 | -1.49E+00 |
| ENSG00000C | HMGB3 | 1.05E-02 | 1.18E-01 | 8.34E-01 | -2.61E-01 | -1.20E+00 |
| ENSG00000C | RBM41 | 1.05E-02 | 1.18E-01 | 8.89E-01 | -1.70E-01 | -1.13E+00 |
| ENSG00000C | GAB2 | 1.05E-02 | 1.18E-01 | 8.55E-01 | -2.27E-01 | -1.17E+00 |
| ENSG000001 | BCAT2 | 1.05E-02 | 1.18E-01 | 1.13E+00 | 1.78E-01 | 1.13E+00 |
| ENSG000001 | MICU2 | 1.06E-02 | 1.19E-01 | 9.08E-01 | -1.40E-01 | -1.10E+00 |
| ENSG000001 | RAB13 | 1.06E-02 | 1.19E-01 | 1.06E+00 | 8.85E-02 | 1.06E+00 |
| ENSG000001 | CCNC | 1.06E-02 | 1.19E-01 | 9.24E-01 | -1.15E-01 | -1.08E+00 |
| ENSG00000C | NUAK1 | 1.06E-02 | 1.19E-01 | 1.13E+00 | 1.73E-01 | 1.13E+00 |
| ENSG000001 | KRBOX5 | 1.07E-02 | 1.19E-01 | 8.90E-01 | -1.68E-01 | -1.12E+00 |
| ENSG000002 | --- | 1.07E-02 | 1.19E-01 | 7.78E-01 | -3.62E-01 | -1.29E+00 |
| ENSG000001 | EIF4E | 1.07E-02 | 1.19E-01 | 9.26E-01 | -1.11E-01 | -1.08E+00 |
| ENSG000001 | BMS1 | 1.07E-02 | 1.19E-01 | 9.00E-01 | -1.52E-01 | -1.11E+00 |
| ENSG000001 | LANCL1 | 1.07E-02 | 1.19E-01 | 9.27E-01 | -1.09E-01 | -1.08E+00 |
| ENSG000001 | RPS6KA1 | 1.07E-02 | 1.20E-01 | 1.12E+00 | 1.66E-01 | 1.12E+00 |
| ENSG000001 | CAMK2N1 | 1.08E-02 | 1.20E-01 | 8.66E-01 | -2.08E-01 | -1.15E+00 |
| ENSG000001 | ODR4 | 1.08E-02 | 1.20E-01 | 9.00E-01 | -1.52E-01 | -1.11E+00 |
| ENSG000001 | TBC1D9 | 1.08E-02 | 1.21E-01 | 9.29E-01 | -1.06E-01 | -1.08E+00 |
| ENSG000001 | ZBED3 | 1.09E-02 | 1.21E-01 | 1.29E+00 | 3.70E-01 | 1.29E+00 |
| ENSG00000C | SERPINB3 | 1.09E-02 | 1.21E-01 | 6.74E-01 | -5.70E-01 | -1.48E+00 |
| ENSG000001 | PMAIP1 | 1.09E-02 | 1.21E-01 | 8.35E-01 | -2.61E-01 | -1.20E+00 |
| ENSG000001 | MAP7 | 1.10E-02 | 1.21E-01 | 9.03E-01 | -1.47E-01 | -1.11E+00 |
| ENSG000001 | ZNF385A | 1.10E-02 | 1.21E-01 | 1.08E+00 | 1.17E-01 | 1.08E+00 |
| ENSG000001 | CORO1C | 1.10E-02 | 1.21E-01 | 1.07E+00 | 9.68E-02 | 1.07E+00 |
| ENSG000001 | IL7R | 1.10E-02 | 1.21E-01 | 8.84E-01 | -1.78E-01 | -1.13E+00 |
| ENSG000002 | --- | 1.10E-02 | 1.21E-01 | 7.84E-01 | -3.51E-01 | -1.28E+00 |
| ENSG00000C | PHF23 | 1.10E-02 | 1.22E-01 | 1.08E+00 | 1.13E-01 | 1.08E+00 |
| ENSG000001 | BOC | 1.10E-02 | 1.22E-01 | 1.28E+00 | 3.52E-01 | 1.28E+00 |
| ENSG000001 | RPS25 | 1.11E-02 | 1.22E-01 | 1.09E+00 | 1.30E-01 | 1.09E+00 |
| ENSG000001 | NACA | 1.11E-02 | 1.22E-01 | 1.07E+00 | 1.01E-01 | 1.07E+00 |
| ENSG000001 | NR1D1 | 1.11E-02 | 1.22E-01 | 7.54E-01 | -4.08E-01 | -1.33E+00 |
| ENSG000001 | COG7 | 1.12E-02 | 1.23E-01 | 8.77E-01 | -1.89E-01 | -1.14E+00 |
| ENSG000002 | HAUS3 | 1.12E-02 | 1.23E-01 | 8.73E-01 | -1.97E-01 | -1.15E+00 |

| | | | | | | |
|---|---|---|---|---|---|---|
| ENSG00000C | DAZAP1 | 1.12E-02 | 1.23E-01 | 9.33E-01 | -1.01E-01 | -1.07E+00 |
| ENSG000001 | FUT3 | 1.12E-02 | 1.23E-01 | 8.40E-01 | -2.52E-01 | -1.19E+00 |
| ENSG000001 | MRPL44 | 1.13E-02 | 1.23E-01 | 1.09E+00 | 1.28E-01 | 1.09E+00 |
| ENSG000001 | CCDC112 | 1.13E-02 | 1.23E-01 | 8.21E-01 | -2.84E-01 | -1.22E+00 |
| ENSG00000C | IFT80 | 1.13E-02 | 1.23E-01 | 8.71E-01 | -1.98E-01 | -1.15E+00 |
| ENSG000001 | SLC16A13 | 1.13E-02 | 1.23E-01 | 1.21E+00 | 2.78E-01 | 1.21E+00 |
| ENSG00000C | BAX | 1.13E-02 | 1.24E-01 | 1.12E+00 | 1.65E-01 | 1.12E+00 |
| ENSG000002 | FGD5-AS1 | 1.13E-02 | 1.24E-01 | 1.08E+00 | 1.07E-01 | 1.08E+00 |
| ENSG000001 | TNFRSF10B | 1.13E-02 | 1.24E-01 | 1.07E+00 | 1.03E-01 | 1.07E+00 |
| ENSG000001 | MOB3A | 1.14E-02 | 1.24E-01 | 9.10E-01 | -1.37E-01 | -1.10E+00 |
| ENSG000001 | SNCA | 1.15E-02 | 1.25E-01 | 1.13E+00 | 1.72E-01 | 1.13E+00 |
| ENSG000001 | RAB36 | 1.15E-02 | 1.25E-01 | 1.14E+00 | 1.93E-01 | 1.14E+00 |
| ENSG000002 | CAPNS2 | 1.15E-02 | 1.25E-01 | 1.27E+00 | 3.40E-01 | 1.27E+00 |
| ENSG000001 | USP18 | 1.15E-02 | 1.25E-01 | 7.51E-01 | -4.12E-01 | -1.33E+00 |
| ENSG00000C | MGST2 | 1.15E-02 | 1.25E-01 | 1.12E+00 | 1.57E-01 | 1.12E+00 |
| ENSG000002 | ZBED6 | 1.16E-02 | 1.25E-01 | 8.85E-01 | -1.76E-01 | -1.13E+00 |
| ENSG000001 | RMND5A | 1.16E-02 | 1.25E-01 | 9.18E-01 | -1.24E-01 | -1.09E+00 |
| ENSG000001 | PRKX | 1.16E-02 | 1.26E-01 | 1.09E+00 | 1.18E-01 | 1.09E+00 |
| ENSG000001 | FGFBP1 | 1.16E-02 | 1.26E-01 | 1.13E+00 | 1.79E-01 | 1.13E+00 |
| ENSG000001 | PSD4 | 1.16E-02 | 1.26E-01 | 1.09E+00 | 1.30E-01 | 1.09E+00 |
| ENSG000002 | LINC00707 | 1.17E-02 | 1.26E-01 | 6.96E-01 | -5.23E-01 | -1.44E+00 |
| ENSG000001 | PLA2G4C | 1.17E-02 | 1.26E-01 | 1.24E+00 | 3.09E-01 | 1.24E+00 |
| ENSG000001 | TJP2 | 1.17E-02 | 1.26E-01 | 9.09E-01 | -1.38E-01 | -1.10E+00 |
| ENSG000002 | --- | 1.17E-02 | 1.26E-01 | 1.47E+00 | 5.55E-01 | 1.47E+00 |
| ENSG000001 | BCL9 | 1.17E-02 | 1.26E-01 | 8.99E-01 | -1.54E-01 | -1.11E+00 |
| ENSG000001 | GABPB2 | 1.17E-02 | 1.26E-01 | 1.12E+00 | 1.67E-01 | 1.12E+00 |
| ENSG00000C | ALAS1 | 1.17E-02 | 1.26E-01 | 9.22E-01 | -1.18E-01 | -1.09E+00 |
| ENSG000001 | PFKFB4 | 1.17E-02 | 1.26E-01 | 8.60E-01 | -2.17E-01 | -1.16E+00 |
| ENSG000001 | EDRF1 | 1.17E-02 | 1.26E-01 | 9.06E-01 | -1.42E-01 | -1.10E+00 |
| ENSG000002 | --- | 1.18E-02 | ? | 5.98E-01 | -7.41E-01 | -1.67E+00 |
| ENSG000001 | ILF3 | 1.18E-02 | 1.27E-01 | 1.05E+00 | 7.50E-02 | 1.05E+00 |
| ENSG000001 | LRP4 | 1.18E-02 | 1.27E-01 | 1.16E+00 | 2.16E-01 | 1.16E+00 |
| ENSG000002 | LINC00973 | 1.18E-02 | 1.27E-01 | 7.74E-01 | -3.70E-01 | -1.29E+00 |
| ENSG000001 | NIPSNAP1 | 1.18E-02 | 1.27E-01 | 1.08E+00 | 1.14E-01 | 1.08E+00 |
| ENSG000001 | TRMT12 | 1.18E-02 | 1.27E-01 | 1.14E+00 | 1.87E-01 | 1.14E+00 |
| ENSG000001 | CHTF8 | 1.19E-02 | 1.27E-01 | 1.18E+00 | 2.33E-01 | 1.18E+00 |
| ENSG000001 | MTF1 | 1.19E-02 | 1.27E-01 | 9.23E-01 | -1.15E-01 | -1.08E+00 |
| ENSG000001 | NDUFA3 | 1.19E-02 | 1.27E-01 | 1.14E+00 | 1.93E-01 | 1.14E+00 |
| ENSG000002 | RPL32P29 | 1.19E-02 | 1.27E-01 | 1.53E+00 | 6.12E-01 | 1.53E+00 |
| ENSG000001 | POLR2A | 1.19E-02 | 1.27E-01 | 8.81E-01 | -1.83E-01 | -1.14E+00 |
| ENSG000001 | HNRNPH2 | 1.20E-02 | 1.27E-01 | 1.08E+00 | 1.05E-01 | 1.08E+00 |
| ENSG000001 | EPHX1 | 1.20E-02 | 1.27E-01 | 8.56E-01 | -2.24E-01 | -1.17E+00 |
| ENSG00000C | PRDM11 | 1.20E-02 | 1.28E-01 | 1.13E+00 | 1.81E-01 | 1.13E+00 |
| ENSG000001 | NINJ1 | 1.20E-02 | 1.28E-01 | 1.20E+00 | 2.60E-01 | 1.20E+00 |
| ENSG000001 | RAB8B | 1.21E-02 | 1.28E-01 | 1.14E+00 | 1.87E-01 | 1.14E+00 |

| | | | | | | |
|---|---|---|---|---|---|---|
| ENSG000002 | SUGT1-DT | 1.21E-02 | ? | 2.04E+00 | 1.03E+00 | 2.04E+00 |
| ENSG000001 | C2orf69 | 1.21E-02 | 1.28E-01 | 8.90E-01 | -1.68E-01 | -1.12E+00 |
| ENSG000001 | GRHL3 | 1.21E-02 | 1.28E-01 | 1.25E+00 | 3.18E-01 | 1.25E+00 |
| ENSG000001 | MMP9 | 1.21E-02 | 1.28E-01 | 7.40E-01 | -4.34E-01 | -1.35E+00 |
| ENSG000002 | --- | 1.21E-02 | 1.28E-01 | 1.27E+00 | 3.44E-01 | 1.27E+00 |
| ENSG000002 | RPTN | 1.22E-02 | ? | 2.04E+00 | 1.03E+00 | 2.04E+00 |
| ENSG000002 | PRR13 | 1.23E-02 | 1.30E-01 | 1.06E+00 | 8.16E-02 | 1.06E+00 |
| ENSG000002 | FMNL1-DT | 1.23E-02 | 1.30E-01 | 6.81E-01 | -5.55E-01 | -1.47E+00 |
| ENSG000001 | RNF8 | 1.23E-02 | 1.30E-01 | 1.09E+00 | 1.22E-01 | 1.09E+00 |
| ENSG000001 | NAA25 | 1.23E-02 | 1.30E-01 | 9.30E-01 | -1.04E-01 | -1.07E+00 |
| ENSG000001 | CDS2 | 1.24E-02 | 1.30E-01 | 1.05E+00 | 7.58E-02 | 1.05E+00 |
| ENSG000002 | TRABD2B | 1.24E-02 | 1.30E-01 | 1.24E+00 | 3.07E-01 | 1.24E+00 |
| ENSG000001 | MAGOH | 1.24E-02 | 1.30E-01 | 1.11E+00 | 1.52E-01 | 1.11E+00 |
| ENSG00000C | PGM3 | 1.24E-02 | 1.30E-01 | 9.32E-01 | -1.02E-01 | -1.07E+00 |
| ENSG000002 | --- | 1.24E-02 | 1.31E-01 | 1.17E+00 | 2.30E-01 | 1.17E+00 |
| ENSG000002 | ABALON | 1.24E-02 | ? | 1.54E+00 | 6.24E-01 | 1.54E+00 |
| ENSG000002 | FLNB-AS1 | 1.24E-02 | 1.31E-01 | 7.89E-01 | -3.41E-01 | -1.27E+00 |
| ENSG000001 | MAPKBP1 | 1.24E-02 | 1.31E-01 | 1.12E+00 | 1.57E-01 | 1.12E+00 |
| ENSG000002 | HNRNPUL2 | 1.24E-02 | 1.31E-01 | 9.42E-01 | -8.69E-02 | -1.06E+00 |
| ENSG000001 | SLC16A9 | 1.25E-02 | 1.31E-01 | 1.29E+00 | 3.72E-01 | 1.29E+00 |
| ENSG00000C | UBE2K | 1.25E-02 | 1.31E-01 | 9.14E-01 | -1.30E-01 | -1.09E+00 |
| ENSG000001 | WNT2B | 1.25E-02 | 1.31E-01 | 7.94E-01 | -3.34E-01 | -1.26E+00 |
| ENSG000001 | HS3ST3A1 | 1.25E-02 | 1.31E-01 | 6.95E-01 | -5.24E-01 | -1.44E+00 |
| ENSG000001 | KIF3B | 1.25E-02 | 1.31E-01 | 1.08E+00 | 1.05E-01 | 1.08E+00 |
| ENSG000001 | HERPUD2 | 1.25E-02 | 1.31E-01 | 9.14E-01 | -1.30E-01 | -1.09E+00 |
| ENSG000001 | SNHG29 | 1.26E-02 | 1.31E-01 | 1.09E+00 | 1.29E-01 | 1.09E+00 |
| ENSG000001 | WIPF2 | 1.26E-02 | 1.32E-01 | 9.51E-01 | -7.29E-02 | -1.05E+00 |
| ENSG000001 | CALHM6 | 1.26E-02 | ? | 4.91E-01 | -1.03E+00 | -2.04E+00 |
| ENSG000001 | B4GAT1 | 1.26E-02 | 1.32E-01 | 1.21E+00 | 2.80E-01 | 1.21E+00 |
| ENSG000002 | --- | 1.26E-02 | ? | 1.92E+00 | 9.41E-01 | 1.92E+00 |
| ENSG000001 | CEBPG | 1.26E-02 | 1.32E-01 | 1.09E+00 | 1.19E-01 | 1.09E+00 |
| ENSG000001 | TWF1P1 | 1.27E-02 | 1.32E-01 | 7.76E-01 | -3.66E-01 | -1.29E+00 |
| ENSG000001 | LDLRAD3 | 1.27E-02 | 1.32E-01 | 1.12E+00 | 1.59E-01 | 1.12E+00 |
| ENSG00000C | PGM1 | 1.27E-02 | 1.32E-01 | 1.12E+00 | 1.65E-01 | 1.12E+00 |
| ENSG000001 | ZNF562 | 1.27E-02 | 1.32E-01 | 9.15E-01 | -1.28E-01 | -1.09E+00 |
| ENSG000001 | CACNA1B | 1.28E-02 | ? | 2.04E+00 | 1.03E+00 | 2.04E+00 |
| ENSG000001 | PAM | 1.28E-02 | 1.33E-01 | 1.08E+00 | 1.17E-01 | 1.08E+00 |
| ENSG000001 | MSL1 | 1.28E-02 | 1.33E-01 | 9.20E-01 | -1.20E-01 | -1.09E+00 |
| ENSG000001 | KCNJ5-AS1 | 1.28E-02 | 1.33E-01 | 1.12E+00 | 1.67E-01 | 1.12E+00 |
| ENSG000001 | IFFO2 | 1.28E-02 | 1.33E-01 | 1.12E+00 | 1.59E-01 | 1.12E+00 |
| ENSG000002 | RIMOC1 | 1.28E-02 | 1.33E-01 | 8.84E-01 | -1.79E-01 | -1.13E+00 |
| ENSG000002 | --- | 1.29E-02 | 1.33E-01 | 1.32E+00 | 3.98E-01 | 1.32E+00 |
| ENSG000001 | BZW1P2 | 1.29E-02 | 1.33E-01 | 8.90E-01 | -1.69E-01 | -1.12E+00 |
| ENSG000001 | NOP10 | 1.29E-02 | 1.33E-01 | 1.10E+00 | 1.36E-01 | 1.10E+00 |
| ENSG000001 | HSDL2 | 1.29E-02 | 1.33E-01 | 1.10E+00 | 1.37E-01 | 1.10E+00 |

| | | | | | | |
|---|---|---|---|---|---|---|
| ENSG000001 | CC2D1B | 1.29E-02 | 1.33E-01 | 1.11E+00 | 1.54E-01 | 1.11E+00 |
| ENSG000000 | DAPP1 | 1.29E-02 | 1.33E-01 | 1.11E+00 | 1.56E-01 | 1.11E+00 |
| ENSG000001 | FJX1 | 1.29E-02 | 1.33E-01 | 1.17E+00 | 2.22E-01 | 1.17E+00 |
| ENSG000000 | NT5C2 | 1.29E-02 | 1.33E-01 | 9.23E-01 | -1.15E-01 | -1.08E+00 |
| ENSG000001 | PDE2A | 1.29E-02 | 1.33E-01 | 8.00E-01 | -3.22E-01 | -1.25E+00 |
| ENSG000001 | CTSK | 1.30E-02 | 1.33E-01 | 7.70E-01 | -3.78E-01 | -1.30E+00 |
| ENSG000001 | TGFBR1 | 1.30E-02 | 1.33E-01 | 6.62E-01 | -5.95E-01 | -1.51E+00 |
| ENSG000001 | ALDH9A1 | 1.30E-02 | 1.33E-01 | 1.09E+00 | 1.25E-01 | 1.09E+00 |
| ENSG000000 | DNAJC10 | 1.30E-02 | 1.33E-01 | 9.10E-01 | -1.37E-01 | -1.10E+00 |
| ENSG000002 | --- | 1.30E-02 | 1.33E-01 | 1.26E+00 | 3.36E-01 | 1.26E+00 |
| ENSG000000 | ROCK1 | 1.30E-02 | 1.34E-01 | 8.98E-01 | -1.55E-01 | -1.11E+00 |
| ENSG000001 | CELF1 | 1.31E-02 | 1.34E-01 | 9.25E-01 | -1.12E-01 | -1.08E+00 |
| ENSG000002 | TUBA3FP | 1.31E-02 | ? | 1.85E+00 | 8.88E-01 | 1.85E+00 |
| ENSG000002 | --- | 1.31E-02 | 1.34E-01 | 1.27E+00 | 3.49E-01 | 1.27E+00 |
| ENSG000001 | TNFAIP1 | 1.32E-02 | 1.34E-01 | 1.06E+00 | 7.84E-02 | 1.06E+00 |
| ENSG000001 | RAB29 | 1.32E-02 | 1.35E-01 | 1.09E+00 | 1.18E-01 | 1.09E+00 |
| ENSG000001 | CYB561A3 | 1.32E-02 | 1.35E-01 | 1.16E+00 | 2.10E-01 | 1.16E+00 |
| ENSG000002 | --- | 1.32E-02 | ? | 6.09E-01 | -7.16E-01 | -1.64E+00 |
| ENSG000001 | SPOCD1 | 1.32E-02 | 1.35E-01 | 1.25E+00 | 3.17E-01 | 1.25E+00 |
| ENSG000001 | MYO5C | 1.32E-02 | 1.35E-01 | 8.91E-01 | -1.66E-01 | -1.12E+00 |
| ENSG000001 | RNF170 | 1.32E-02 | 1.35E-01 | 1.10E+00 | 1.34E-01 | 1.10E+00 |
| ENSG000002 | EEF1AKMT4 | 1.33E-02 | 1.35E-01 | 1.26E+00 | 3.31E-01 | 1.26E+00 |
| ENSG000001 | PLAGL2 | 1.33E-02 | 1.35E-01 | 1.09E+00 | 1.19E-01 | 1.09E+00 |
| ENSG000001 | LINC00173 | 1.33E-02 | 1.35E-01 | 7.17E-01 | -4.81E-01 | -1.40E+00 |
| ENSG000001 | B3GALNT2 | 1.34E-02 | 1.36E-01 | 1.09E+00 | 1.29E-01 | 1.09E+00 |
| ENSG000002 | SBDSP1 | 1.34E-02 | 1.36E-01 | 9.06E-01 | -1.43E-01 | -1.10E+00 |
| ENSG000002 | RPS12P26 | 1.34E-02 | ? | 3.90E-01 | -1.36E+00 | -2.56E+00 |
| ENSG000000 | FAM76B | 1.34E-02 | 1.36E-01 | 8.49E-01 | -2.37E-01 | -1.18E+00 |
| ENSG000001 | RASA2 | 1.34E-02 | 1.36E-01 | 9.02E-01 | -1.49E-01 | -1.11E+00 |
| ENSG000001 | FAM110B | 1.34E-02 | ? | 1.91E+00 | 9.37E-01 | 1.91E+00 |
| ENSG000001 | ARL10 | 1.34E-02 | 1.36E-01 | 1.16E+00 | 2.10E-01 | 1.16E+00 |
| ENSG000001 | DHX35 | 1.35E-02 | 1.36E-01 | 1.14E+00 | 1.86E-01 | 1.14E+00 |
| ENSG000001 | GALE | 1.35E-02 | 1.36E-01 | 1.14E+00 | 1.86E-01 | 1.14E+00 |
| ENSG000001 | GJB4 | 1.35E-02 | 1.36E-01 | 8.87E-01 | -1.73E-01 | -1.13E+00 |
| ENSG000002 | FAM86EP | 1.35E-02 | 1.37E-01 | 1.30E+00 | 3.80E-01 | 1.30E+00 |
| ENSG000001 | SORCS2 | 1.35E-02 | 1.37E-01 | 7.03E-01 | -5.09E-01 | -1.42E+00 |
| ENSG000002 | --- | 1.36E-02 | 1.37E-01 | 8.17E-01 | -2.91E-01 | -1.22E+00 |
| ENSG000001 | FAM162A | 1.36E-02 | 1.37E-01 | 1.08E+00 | 1.09E-01 | 1.08E+00 |
| ENSG000000 | RPLP0 | 1.36E-02 | 1.37E-01 | 1.13E+00 | 1.72E-01 | 1.13E+00 |
| ENSG000001 | PLAT | 1.36E-02 | 1.37E-01 | 7.56E-01 | -4.03E-01 | -1.32E+00 |
| ENSG000001 | BBS1 | 1.37E-02 | 1.38E-01 | 1.15E+00 | 2.08E-01 | 1.15E+00 |
| ENSG000001 | INTS9 | 1.37E-02 | 1.38E-01 | 8.99E-01 | -1.54E-01 | -1.11E+00 |
| ENSG000001 | PSMC3 | 1.37E-02 | 1.38E-01 | 9.24E-01 | -1.15E-01 | -1.08E+00 |
| ENSG000001 | TYW3 | 1.37E-02 | 1.38E-01 | 9.07E-01 | -1.42E-01 | -1.10E+00 |
| ENSG000002 | --- | 1.37E-02 | 1.38E-01 | 8.48E-01 | -2.38E-01 | -1.18E+00 |

| | | | | | | |
|---|---|---|---|---|---|---|
| ENSG000002 | BRD2 | 1.38E-02 | 1.38E-01 | 9.23E-01 | -1.16E-01 | -1.08E+00 |
| ENSG000001 | FSIP2 | 1.38E-02 | 1.38E-01 | 7.87E-01 | -3.46E-01 | -1.27E+00 |
| ENSG000002 | --- | 1.38E-02 | 1.38E-01 | 7.59E-01 | -3.98E-01 | -1.32E+00 |
| ENSG000001 | VDR | 1.38E-02 | 1.38E-01 | 8.81E-01 | -1.82E-01 | -1.13E+00 |
| ENSG000001 | NPIPB13 | 1.38E-02 | 1.38E-01 | 7.81E-01 | -3.57E-01 | -1.28E+00 |
| ENSG000001 | ECH1 | 1.39E-02 | 1.39E-01 | 1.14E+00 | 1.87E-01 | 1.14E+00 |
| ENSG000002 | ITGB3 | 1.39E-02 | 1.39E-01 | 1.29E+00 | 3.64E-01 | 1.29E+00 |
| ENSG000001 | MED20 | 1.39E-02 | 1.39E-01 | 1.11E+00 | 1.55E-01 | 1.11E+00 |
| ENSG000001 | ENSA | 1.40E-02 | 1.40E-01 | 9.30E-01 | -1.04E-01 | -1.07E+00 |
| ENSG000001 | PI4KB | 1.41E-02 | 1.40E-01 | 1.10E+00 | 1.31E-01 | 1.10E+00 |
| ENSG000001 | YES1 | 1.41E-02 | 1.40E-01 | 9.15E-01 | -1.28E-01 | -1.09E+00 |
| ENSG000001 | SNX4 | 1.41E-02 | 1.40E-01 | 1.10E+00 | 1.38E-01 | 1.10E+00 |
| ENSG000001 | GTF3C4 | 1.41E-02 | 1.40E-01 | 1.07E+00 | 9.40E-02 | 1.07E+00 |
| ENSG000001 | MYOM1 | 1.41E-02 | 1.40E-01 | 1.43E+00 | 5.18E-01 | 1.43E+00 |
| ENSG000001 | PPCDC | 1.41E-02 | 1.40E-01 | 1.17E+00 | 2.21E-01 | 1.17E+00 |
| ENSG000001 | USP16 | 1.42E-02 | 1.40E-01 | 9.18E-01 | -1.23E-01 | -1.09E+00 |
| ENSG000002 | HLA-L | 1.42E-02 | 1.40E-01 | 1.45E+00 | 5.38E-01 | 1.45E+00 |
| ENSG000002 | FANCG | 1.42E-02 | 1.40E-01 | 1.22E+00 | 2.83E-01 | 1.22E+00 |
| ENSG000001 | SPTY2D1 | 1.42E-02 | 1.41E-01 | 9.17E-01 | -1.25E-01 | -1.09E+00 |
| ENSG00000C | COL16A1 | 1.42E-02 | 1.41E-01 | 1.16E+00 | 2.12E-01 | 1.16E+00 |
| ENSG00000C | STPG1 | 1.42E-02 | 1.41E-01 | 9.21E-01 | -1.19E-01 | -1.09E+00 |
| ENSG000001 | MYL12A | 1.42E-02 | 1.41E-01 | 1.06E+00 | 8.41E-02 | 1.06E+00 |
| ENSG000001 | STARD5 | 1.43E-02 | 1.41E-01 | 1.15E+00 | 2.08E-01 | 1.15E+00 |
| ENSG000001 | RBM48 | 1.43E-02 | 1.41E-01 | 8.61E-01 | -2.16E-01 | -1.16E+00 |
| ENSG000002 | RBBP4P1 | 1.44E-02 | ? | 4.41E-01 | -1.18E+00 | -2.27E+00 |
| ENSG00000C | IGSF9 | 1.44E-02 | 1.42E-01 | 1.18E+00 | 2.33E-01 | 1.18E+00 |
| ENSG000002 | ZFAND2A-DT | 1.44E-02 | 1.42E-01 | 1.22E+00 | 2.86E-01 | 1.22E+00 |
| ENSG000001 | FBXW4 | 1.44E-02 | 1.42E-01 | 1.17E+00 | 2.27E-01 | 1.17E+00 |
| ENSG000001 | MR1 | 1.44E-02 | 1.42E-01 | 1.11E+00 | 1.52E-01 | 1.11E+00 |
| ENSG000001 | GUCD1 | 1.44E-02 | 1.42E-01 | 1.09E+00 | 1.30E-01 | 1.09E+00 |
| ENSG000001 | RPS27L | 1.44E-02 | 1.42E-01 | 1.07E+00 | 9.94E-02 | 1.07E+00 |
| ENSG000001 | FAM234A | 1.44E-02 | 1.42E-01 | 1.14E+00 | 1.91E-01 | 1.14E+00 |
| ENSG000002 | --- | 1.45E-02 | 1.42E-01 | 7.60E-01 | -3.96E-01 | -1.32E+00 |
| ENSG000001 | ZSCAN21 | 1.45E-02 | 1.42E-01 | 1.14E+00 | 1.89E-01 | 1.14E+00 |
| ENSG000001 | MCM9 | 1.45E-02 | 1.42E-01 | 1.14E+00 | 1.90E-01 | 1.14E+00 |
| ENSG000001 | SLC25A37 | 1.45E-02 | 1.42E-01 | 8.87E-01 | -1.73E-01 | -1.13E+00 |
| ENSG000002 | --- | 1.46E-02 | 1.43E-01 | 8.18E-01 | -2.90E-01 | -1.22E+00 |
| ENSG000002 | HSP90AA2P | 1.46E-02 | 1.43E-01 | 7.95E-01 | -3.32E-01 | -1.26E+00 |
| ENSG000002 | --- | 1.47E-02 | ? | 2.08E+00 | 1.06E+00 | 2.08E+00 |
| ENSG000001 | PLPP5 | 1.47E-02 | 1.43E-01 | 8.99E-01 | -1.54E-01 | -1.11E+00 |
| ENSG000001 | TRAPPC2 | 1.47E-02 | 1.44E-01 | 9.08E-01 | -1.39E-01 | -1.10E+00 |
| ENSG000001 | CALD1 | 1.47E-02 | 1.44E-01 | 1.10E+00 | 1.39E-01 | 1.10E+00 |
| ENSG000001 | CSMD1 | 1.47E-02 | 1.44E-01 | 1.43E+00 | 5.13E-01 | 1.43E+00 |
| ENSG000001 | FEZ1 | 1.47E-02 | 1.44E-01 | 1.11E+00 | 1.50E-01 | 1.11E+00 |
| ENSG00000C | ABL1 | 1.48E-02 | 1.44E-01 | 1.09E+00 | 1.24E-01 | 1.09E+00 |

| | | | | | | |
|---|---|---|---|---|---|---|
| ENSG000001 | ORMDL3 | 1.49E-02 | 1.44E-01 | 1.15E+00 | 1.98E-01 | 1.15E+00 |
| ENSG000001 | DNAL4 | 1.49E-02 | 1.45E-01 | 1.15E+00 | 2.02E-01 | 1.15E+00 |
| ENSG000002 | --- | 1.49E-02 | ? | 2.29E+00 | 1.19E+00 | 2.29E+00 |
| ENSG000001 | PIK3IP1 | 1.49E-02 | 1.45E-01 | 1.20E+00 | 2.61E-01 | 1.20E+00 |
| ENSG000002 | TMEM256 | 1.49E-02 | 1.45E-01 | 1.28E+00 | 3.58E-01 | 1.28E+00 |
| ENSG000001 | VPS8 | 1.49E-02 | 1.45E-01 | 9.29E-01 | -1.06E-01 | -1.08E+00 |
| ENSG000001 | VPS37A | 1.50E-02 | 1.45E-01 | 9.13E-01 | -1.32E-01 | -1.10E+00 |
| ENSG000001 | PPP2R2D | 1.50E-02 | 1.45E-01 | 9.30E-01 | -1.04E-01 | -1.07E+00 |
| ENSG000001 | GLYR1 | 1.50E-02 | 1.45E-01 | 9.39E-01 | -9.04E-02 | -1.06E+00 |
| ENSG000001 | SLC31A2 | 1.50E-02 | 1.45E-01 | 1.13E+00 | 1.73E-01 | 1.13E+00 |
| ENSG000001 | FOXE1 | 1.51E-02 | 1.46E-01 | 1.21E+00 | 2.71E-01 | 1.21E+00 |
| ENSG000001 | PHLDA2 | 1.51E-02 | 1.46E-01 | 8.61E-01 | -2.16E-01 | -1.16E+00 |
| ENSG000001 | GNB4 | 1.51E-02 | 1.46E-01 | 1.07E+00 | 1.04E-01 | 1.07E+00 |
| ENSG000001 | RRAGA | 1.52E-02 | 1.46E-01 | 1.08E+00 | 1.11E-01 | 1.08E+00 |
| ENSG000001 | ESCO1 | 1.52E-02 | 1.46E-01 | 8.92E-01 | -1.64E-01 | -1.12E+00 |
| ENSG000001 | EVI5L | 1.52E-02 | 1.46E-01 | 1.16E+00 | 2.18E-01 | 1.16E+00 |
| ENSG000001 | COA1 | 1.52E-02 | 1.46E-01 | 9.09E-01 | -1.38E-01 | -1.10E+00 |
| ENSG000002 | --- | 1.52E-02 | 1.46E-01 | 8.27E-01 | -2.74E-01 | -1.21E+00 |
| ENSG000002 | --- | 1.52E-02 | ? | 6.01E-01 | -7.34E-01 | -1.66E+00 |
| ENSG000001 | TNFRSF19 | 1.53E-02 | 1.47E-01 | 7.40E-01 | -4.34E-01 | -1.35E+00 |
| ENSG000001 | TSSK4 | 1.53E-02 | 1.47E-01 | 6.34E-01 | -6.58E-01 | -1.58E+00 |
| ENSG000001 | GOLGA1 | 1.53E-02 | 1.47E-01 | 9.07E-01 | -1.41E-01 | -1.10E+00 |
| ENSG000001 | ZFP92 | 1.53E-02 | 1.47E-01 | 1.20E+00 | 2.59E-01 | 1.20E+00 |
| ENSG000002 | PSG1 | 1.53E-02 | ? | 6.22E-01 | -6.86E-01 | -1.61E+00 |
| ENSG000002 | --- | 1.54E-02 | 1.48E-01 | 1.81E+00 | 8.53E-01 | 1.81E+00 |
| ENSG000001 | ENTPD1 | 1.55E-02 | 1.48E-01 | 8.27E-01 | -2.75E-01 | -1.21E+00 |
| ENSG000001 | GGH | 1.55E-02 | 1.48E-01 | 8.68E-01 | -2.05E-01 | -1.15E+00 |
| ENSG000001 | FAM89B | 1.55E-02 | 1.48E-01 | 1.16E+00 | 2.16E-01 | 1.16E+00 |
| ENSG00000C | CALCRL | 1.55E-02 | 1.48E-01 | 6.79E-01 | -5.59E-01 | -1.47E+00 |
| ENSG000001 | ZNF703 | 1.55E-02 | 1.48E-01 | 8.97E-01 | -1.56E-01 | -1.11E+00 |
| ENSG000001 | OAS3 | 1.55E-02 | 1.48E-01 | 1.09E+00 | 1.18E-01 | 1.09E+00 |
| ENSG000001 | SUN2 | 1.56E-02 | 1.49E-01 | 1.13E+00 | 1.77E-01 | 1.13E+00 |
| ENSG000001 | TTLL7 | 1.56E-02 | 1.49E-01 | 8.83E-01 | -1.80E-01 | -1.13E+00 |
| ENSG000001 | ANAPC15 | 1.56E-02 | 1.49E-01 | 1.09E+00 | 1.25E-01 | 1.09E+00 |
| ENSG000001 | KBTBD4 | 1.57E-02 | 1.49E-01 | 1.13E+00 | 1.75E-01 | 1.13E+00 |
| ENSG000001 | TMEM39A | 1.57E-02 | 1.49E-01 | 8.97E-01 | -1.57E-01 | -1.11E+00 |
| ENSG00000C | RAP1GAP | 1.57E-02 | 1.49E-01 | 1.33E+00 | 4.10E-01 | 1.33E+00 |
| ENSG000001 | COL6A3 | 1.58E-02 | ? | 6.32E-01 | -6.62E-01 | -1.58E+00 |
| ENSG000001 | ZC3H14 | 1.58E-02 | 1.50E-01 | 9.00E-01 | -1.53E-01 | -1.11E+00 |
| ENSG000001 | ETHE1 | 1.59E-02 | 1.51E-01 | 1.16E+00 | 2.14E-01 | 1.16E+00 |
| ENSG000001 | MAGED4 | 1.59E-02 | 1.51E-01 | 1.34E+00 | 4.24E-01 | 1.34E+00 |
| ENSG000002 | FAM66D | 1.59E-02 | 1.51E-01 | 1.45E+00 | 5.32E-01 | 1.45E+00 |
| ENSG000001 | ACVR1B | 1.59E-02 | 1.51E-01 | 1.08E+00 | 1.09E-01 | 1.08E+00 |
| ENSG000001 | RPLP2 | 1.60E-02 | 1.51E-01 | 1.14E+00 | 1.85E-01 | 1.14E+00 |
| ENSG000001 | CPSF4 | 1.60E-02 | 1.51E-01 | 1.13E+00 | 1.77E-01 | 1.13E+00 |

| | | | | | | |
|---|---|---|---|---|---|---|
| ENSG000001 | RGP1 | 1.60E-02 | 1.51E-01 | 9.39E-01 | -9.13E-02 | -1.07E+00 |
| ENSG000001 | TMEM187 | 1.60E-02 | 1.51E-01 | 1.28E+00 | 3.58E-01 | 1.28E+00 |
| ENSG000002 | PIK3CD-AS2 | 1.60E-02 | ? | 2.15E+00 | 1.11E+00 | 2.15E+00 |
| ENSG000002 | --- | 1.60E-02 | 1.51E-01 | 1.46E+00 | 5.44E-01 | 1.46E+00 |
| ENSG000001 | MED26 | 1.61E-02 | 1.52E-01 | 1.12E+00 | 1.67E-01 | 1.12E+00 |
| ENSG000001 | MIX23 | 1.61E-02 | 1.52E-01 | 8.61E-01 | -2.16E-01 | -1.16E+00 |
| ENSG000001 | ALCAM | 1.61E-02 | 1.52E-01 | 9.00E-01 | -1.52E-01 | -1.11E+00 |
| ENSG000001 | SLC39A8 | 1.61E-02 | 1.52E-01 | 8.97E-01 | -1.57E-01 | -1.11E+00 |
| ENSG000001 | PSMD1 | 1.61E-02 | 1.52E-01 | 9.36E-01 | -9.60E-02 | -1.07E+00 |
| ENSG000002 | ARHGEF35 | 1.62E-02 | 1.52E-01 | 9.08E-01 | -1.39E-01 | -1.10E+00 |
| ENSG000001 | RNF24 | 1.62E-02 | 1.52E-01 | 9.41E-01 | -8.81E-02 | -1.06E+00 |
| ENSG000001 | PIK3R2 | 1.62E-02 | 1.52E-01 | 1.08E+00 | 1.12E-01 | 1.08E+00 |
| ENSG000001 | RITA1 | 1.62E-02 | 1.52E-01 | 1.13E+00 | 1.79E-01 | 1.13E+00 |
| ENSG000002 | SMIM10L1 | 1.62E-02 | 1.52E-01 | 9.17E-01 | -1.24E-01 | -1.09E+00 |
| ENSG000001 | ADCK2 | 1.62E-02 | 1.52E-01 | 1.18E+00 | 2.43E-01 | 1.18E+00 |
| ENSG000001 | TMEM43 | 1.62E-02 | 1.52E-01 | 1.07E+00 | 1.01E-01 | 1.07E+00 |
| ENSG000001 | GEMIN4 | 1.62E-02 | 1.52E-01 | 1.14E+00 | 1.84E-01 | 1.14E+00 |
| ENSG000001 | TAF1D | 1.62E-02 | 1.52E-01 | 8.78E-01 | -1.87E-01 | -1.14E+00 |
| ENSG000001 | FAM83F | 1.62E-02 | 1.52E-01 | 1.13E+00 | 1.78E-01 | 1.13E+00 |
| ENSG000001 | PLEKHG1 | 1.63E-02 | 1.52E-01 | 1.11E+00 | 1.47E-01 | 1.11E+00 |
| ENSG000001 | CEP97 | 1.63E-02 | 1.52E-01 | 9.03E-01 | -1.47E-01 | -1.11E+00 |
| ENSG000001 | MAP1B | 1.63E-02 | 1.52E-01 | 8.38E-01 | -2.55E-01 | -1.19E+00 |
| ENSG000001 | EPHX2 | 1.63E-02 | 1.52E-01 | 1.26E+00 | 3.35E-01 | 1.26E+00 |
| ENSG000001 | TKFC | 1.63E-02 | 1.52E-01 | 1.19E+00 | 2.47E-01 | 1.19E+00 |
| ENSG000001 | DPP7 | 1.64E-02 | 1.53E-01 | 1.17E+00 | 2.30E-01 | 1.17E+00 |
| ENSG000001 | ZNF571 | 1.64E-02 | 1.53E-01 | 8.30E-01 | -2.69E-01 | -1.20E+00 |
| ENSG000001 | MRS2 | 1.64E-02 | 1.53E-01 | 9.22E-01 | -1.17E-01 | -1.08E+00 |
| ENSG000001 | SRGAP2B | 1.64E-02 | 1.53E-01 | 8.74E-01 | -1.95E-01 | -1.14E+00 |
| ENSG000001 | KRT80 | 1.65E-02 | 1.53E-01 | 8.16E-01 | -2.93E-01 | -1.23E+00 |
| ENSG000002 | --- | 1.66E-02 | ? | 2.81E+00 | 1.49E+00 | 2.81E+00 |
| ENSG000001 | GJB5 | 1.66E-02 | 1.54E-01 | 1.08E+00 | 1.04E-01 | 1.08E+00 |
| ENSG000002 | SLC20A1-DT | 1.66E-02 | 1.54E-01 | 8.43E-01 | -2.46E-01 | -1.19E+00 |
| ENSG000001 | UBE2N | 1.66E-02 | 1.54E-01 | 9.18E-01 | -1.24E-01 | -1.09E+00 |
| ENSG000002 | --- | 1.67E-02 | ? | 1.44E+00 | 5.21E-01 | 1.44E+00 |
| ENSG000001 | CLIP1 | 1.67E-02 | 1.55E-01 | 8.98E-01 | -1.56E-01 | -1.11E+00 |
| ENSG000002 | PEG10 | 1.68E-02 | 1.56E-01 | 8.67E-01 | -2.06E-01 | -1.15E+00 |
| ENSG000001 | FRG1BP | 1.68E-02 | 1.56E-01 | 7.94E-01 | -3.34E-01 | -1.26E+00 |
| ENSG000001 | MAP1LC3A | 1.68E-02 | 1.56E-01 | 1.39E+00 | 4.72E-01 | 1.39E+00 |
| ENSG000001 | IVL | 1.68E-02 | 1.56E-01 | 1.47E+00 | 5.57E-01 | 1.47E+00 |
| ENSG000001 | PYM1 | 1.69E-02 | 1.57E-01 | 1.10E+00 | 1.31E-01 | 1.10E+00 |
| ENSG000002 | TMA7 | 1.70E-02 | 1.57E-01 | 1.09E+00 | 1.30E-01 | 1.09E+00 |
| ENSG000001 | ZNF16 | 1.71E-02 | 1.57E-01 | 1.19E+00 | 2.48E-01 | 1.19E+00 |
| ENSG000001 | SMC4 | 1.71E-02 | 1.57E-01 | 8.72E-01 | -1.98E-01 | -1.15E+00 |
| ENSG000001 | MEGF9 | 1.71E-02 | 1.58E-01 | 1.09E+00 | 1.21E-01 | 1.09E+00 |
| ENSG000001 | PLCD1 | 1.71E-02 | 1.58E-01 | 1.19E+00 | 2.57E-01 | 1.19E+00 |

| | | | | | | |
|---|---|---|---|---|---|---|
| ENSG000001 | TMEM177 | 1.72E-02 | 1.58E-01 | 1.18E+00 | 2.38E-01 | 1.18E+00 |
| ENSG00000C | CARMIL1 | 1.72E-02 | 1.58E-01 | 1.09E+00 | 1.18E-01 | 1.09E+00 |
| ENSG000002 | --- | 1.72E-02 | 1.58E-01 | 5.82E-01 | -7.80E-01 | -1.72E+00 |
| ENSG000001 | ITPK1 | 1.72E-02 | 1.58E-01 | 1.11E+00 | 1.56E-01 | 1.11E+00 |
| ENSG00000C | SLC11A1 | 1.73E-02 | ? | 1.49E+00 | 5.78E-01 | 1.49E+00 |
| ENSG000001 | ACOT9 | 1.73E-02 | 1.59E-01 | 9.38E-01 | -9.31E-02 | -1.07E+00 |
| ENSG000001 | STEAP4 | 1.73E-02 | 1.59E-01 | 7.24E-01 | -4.66E-01 | -1.38E+00 |
| ENSG000001 | MYBL2 | 1.73E-02 | 1.59E-01 | 7.15E-01 | -4.84E-01 | -1.40E+00 |
| ENSG000001 | TGFBRAP1 | 1.74E-02 | 1.59E-01 | 1.08E+00 | 1.07E-01 | 1.08E+00 |
| ENSG000001 | LSM6 | 1.74E-02 | 1.60E-01 | 1.13E+00 | 1.77E-01 | 1.13E+00 |
| ENSG000001 | SLC35A4 | 1.75E-02 | 1.60E-01 | 1.10E+00 | 1.38E-01 | 1.10E+00 |
| ENSG000001 | MFAP2 | 1.75E-02 | 1.60E-01 | 1.09E+00 | 1.30E-01 | 1.09E+00 |
| ENSG000001 | EAF1 | 1.75E-02 | 1.60E-01 | 1.06E+00 | 8.05E-02 | 1.06E+00 |
| ENSG000001 | SCARA3 | 1.76E-02 | 1.61E-01 | 1.27E+00 | 3.43E-01 | 1.27E+00 |
| ENSG000002 | ATP6V1G2-D | 1.76E-02 | 1.61E-01 | 6.36E-01 | -6.53E-01 | -1.57E+00 |
| ENSG000001 | NAGS | 1.76E-02 | 1.61E-01 | 8.50E-01 | -2.34E-01 | -1.18E+00 |
| ENSG00000C | OFD1 | 1.76E-02 | 1.61E-01 | 8.95E-01 | -1.60E-01 | -1.12E+00 |
| ENSG000001 | RAB11FIP4 | 1.76E-02 | 1.61E-01 | 1.16E+00 | 2.19E-01 | 1.16E+00 |
| ENSG00000C | PRSS8 | 1.77E-02 | 1.61E-01 | 1.13E+00 | 1.82E-01 | 1.13E+00 |
| ENSG000001 | RPL22 | 1.77E-02 | 1.61E-01 | 1.06E+00 | 8.78E-02 | 1.06E+00 |
| ENSG000001 | TMEM30B | 1.77E-02 | 1.61E-01 | 1.07E+00 | 9.93E-02 | 1.07E+00 |
| ENSG000001 | CRYBG2 | 1.78E-02 | 1.62E-01 | 1.13E+00 | 1.79E-01 | 1.13E+00 |
| ENSG000001 | CYTH4 | 1.78E-02 | ? | 5.60E-01 | -8.38E-01 | -1.79E+00 |
| ENSG00000C | ACTN1 | 1.79E-02 | 1.62E-01 | 1.07E+00 | 9.80E-02 | 1.07E+00 |
| ENSG000001 | FRRS1 | 1.79E-02 | 1.62E-01 | 1.12E+00 | 1.67E-01 | 1.12E+00 |
| ENSG000001 | MMP24 | 1.79E-02 | 1.62E-01 | 1.25E+00 | 3.18E-01 | 1.25E+00 |
| ENSG000002 | ST3GAL6-AS | 1.79E-02 | 1.62E-01 | 7.06E-01 | -5.02E-01 | -1.42E+00 |
| ENSG000001 | PTPRK | 1.79E-02 | 1.62E-01 | 9.03E-01 | -1.48E-01 | -1.11E+00 |
| ENSG000001 | TRAK1 | 1.79E-02 | 1.62E-01 | 9.49E-01 | -7.56E-02 | -1.05E+00 |
| ENSG000001 | --- | 1.79E-02 | ? | 1.76E+00 | 8.14E-01 | 1.76E+00 |
| ENSG000002 | S1PR2 | 1.80E-02 | 1.62E-01 | 1.18E+00 | 2.44E-01 | 1.18E+00 |
| ENSG000001 | DYRK3 | 1.80E-02 | 1.62E-01 | 1.09E+00 | 1.25E-01 | 1.09E+00 |
| ENSG000001 | PDPN | 1.80E-02 | 1.62E-01 | 8.88E-01 | -1.71E-01 | -1.13E+00 |
| ENSG000001 | CCDC32 | 1.80E-02 | 1.63E-01 | 8.89E-01 | -1.70E-01 | -1.12E+00 |
| ENSG000001 | KCTD10 | 1.81E-02 | 1.63E-01 | 1.05E+00 | 6.51E-02 | 1.05E+00 |
| ENSG000001 | KCNK1 | 1.81E-02 | 1.63E-01 | 8.64E-01 | -2.11E-01 | -1.16E+00 |
| ENSG00000C | PRICKLE3 | 1.81E-02 | 1.63E-01 | 1.11E+00 | 1.46E-01 | 1.11E+00 |
| ENSG000001 | CA10 | 1.81E-02 | ? | 6.34E-01 | -6.57E-01 | -1.58E+00 |
| ENSG000002 | --- | 1.81E-02 | ? | 4.75E-01 | -1.07E+00 | -2.10E+00 |
| ENSG000001 | DLGAP5 | 1.82E-02 | 1.63E-01 | 7.30E-01 | -4.55E-01 | -1.37E+00 |
| ENSG000001 | PTTG1IP | 1.82E-02 | 1.63E-01 | 9.24E-01 | -1.14E-01 | -1.08E+00 |
| ENSG000001 | IRF1 | 1.82E-02 | 1.63E-01 | 8.77E-01 | -1.89E-01 | -1.14E+00 |
| ENSG000002 | --- | 1.82E-02 | 1.63E-01 | 9.01E-01 | -1.51E-01 | -1.11E+00 |
| ENSG000002 | TUBB3 | 1.82E-02 | 1.63E-01 | 8.57E-01 | -2.22E-01 | -1.17E+00 |
| ENSG000002 | TMEM265 | 1.82E-02 | 1.63E-01 | 6.12E+00 | 2.61E+00 | 6.12E+00 |

| | | | | | | |
|---|---|---|---|---|---|---|
| ENSG00000C | PLOD1 | 1.82E-02 | 1.63E-01 | 1.11E+00 | 1.56E-01 | 1.11E+00 |
| ENSG000001 | P2RY6 | 1.82E-02 | 1.63E-01 | 7.03E-01 | -5.08E-01 | -1.42E+00 |
| ENSG000001 | ATL3 | 1.82E-02 | 1.63E-01 | 9.56E-01 | -6.51E-02 | -1.05E+00 |
| ENSG00000C | PRDM1 | 1.83E-02 | 1.64E-01 | 8.17E-01 | -2.91E-01 | -1.22E+00 |
| ENSG000001 | ASCC2 | 1.83E-02 | 1.64E-01 | 1.06E+00 | 8.59E-02 | 1.06E+00 |
| ENSG000002 | ZBTB9 | 1.83E-02 | 1.64E-01 | 1.11E+00 | 1.50E-01 | 1.11E+00 |
| ENSG000001 | TFEB | 1.83E-02 | 1.64E-01 | 1.20E+00 | 2.58E-01 | 1.20E+00 |
| ENSG000001 | DYNC1LI1 | 1.83E-02 | 1.64E-01 | 9.24E-01 | -1.15E-01 | -1.08E+00 |
| ENSG000002 | LAMTOR5-AS | 1.83E-02 | ? | 1.47E+00 | 5.55E-01 | 1.47E+00 |
| ENSG000001 | WNT5A | 1.84E-02 | 1.64E-01 | 8.00E-01 | -3.22E-01 | -1.25E+00 |
| ENSG000001 | DLC1 | 1.84E-02 | 1.64E-01 | 8.52E-01 | -2.31E-01 | -1.17E+00 |
| ENSG000001 | FAM210A | 1.84E-02 | 1.64E-01 | 9.18E-01 | -1.23E-01 | -1.09E+00 |
| ENSG00000C | METTL13 | 1.85E-02 | 1.65E-01 | 1.10E+00 | 1.31E-01 | 1.10E+00 |
| ENSG000001 | BACH1 | 1.85E-02 | 1.65E-01 | 1.09E+00 | 1.24E-01 | 1.09E+00 |
| ENSG000001 | IRF2BP2 | 1.85E-02 | 1.65E-01 | 9.38E-01 | -9.30E-02 | -1.07E+00 |
| ENSG000001 | UBE2V2 | 1.85E-02 | 1.65E-01 | 9.21E-01 | -1.19E-01 | -1.09E+00 |
| ENSG000001 | KIAA1549 | 1.85E-02 | 1.65E-01 | 1.15E+00 | 2.00E-01 | 1.15E+00 |
| ENSG000001 | NRIP1 | 1.86E-02 | 1.65E-01 | 9.21E-01 | -1.19E-01 | -1.09E+00 |
| ENSG000001 | SNAPIN | 1.86E-02 | 1.65E-01 | 1.10E+00 | 1.35E-01 | 1.10E+00 |
| ENSG000002 | --- | 1.86E-02 | 1.65E-01 | 1.15E+00 | 1.98E-01 | 1.15E+00 |
| ENSG00000C | LAMB1 | 1.86E-02 | 1.65E-01 | 9.10E-01 | -1.36E-01 | -1.10E+00 |
| ENSG000001 | ATP5PD | 1.87E-02 | 1.65E-01 | 1.06E+00 | 8.41E-02 | 1.06E+00 |
| ENSG000001 | UNC13B | 1.87E-02 | 1.65E-01 | 9.21E-01 | -1.18E-01 | -1.09E+00 |
| ENSG000001 | MAP2K1 | 1.87E-02 | 1.65E-01 | 9.02E-01 | -1.49E-01 | -1.11E+00 |
| ENSG00000C | CD82 | 1.88E-02 | 1.66E-01 | 1.14E+00 | 1.87E-01 | 1.14E+00 |
| ENSG000001 | CCDC115 | 1.88E-02 | 1.66E-01 | 1.14E+00 | 1.92E-01 | 1.14E+00 |
| ENSG000002 | B3GALT9 | 1.88E-02 | 1.66E-01 | 1.14E+00 | 1.90E-01 | 1.14E+00 |
| ENSG000001 | ICAM4 | 1.88E-02 | 1.66E-01 | 1.33E+00 | 4.09E-01 | 1.33E+00 |
| ENSG000001 | GNPNAT1 | 1.89E-02 | 1.66E-01 | 9.14E-01 | -1.31E-01 | -1.09E+00 |
| ENSG000002 | --- | 1.89E-02 | 1.66E-01 | 8.67E-01 | -2.05E-01 | -1.15E+00 |
| ENSG000002 | --- | 1.89E-02 | ? | 4.26E-01 | -1.23E+00 | -2.35E+00 |
| ENSG000002 | --- | 1.89E-02 | 1.67E-01 | 7.70E-01 | -3.76E-01 | -1.30E+00 |
| ENSG000001 | COL4A5 | 1.90E-02 | 1.67E-01 | 1.15E+00 | 1.99E-01 | 1.15E+00 |
| ENSG00000C | ARSD | 1.90E-02 | 1.67E-01 | 1.10E+00 | 1.32E-01 | 1.10E+00 |
| ENSG000002 | DISP3 | 1.90E-02 | ? | 1.86E+00 | 8.92E-01 | 1.86E+00 |
| ENSG000001 | CLCC1 | 1.90E-02 | 1.67E-01 | 9.12E-01 | -1.34E-01 | -1.10E+00 |
| ENSG000001 | MRPS28 | 1.90E-02 | 1.67E-01 | 1.10E+00 | 1.43E-01 | 1.10E+00 |
| ENSG000001 | TMBIM6 | 1.90E-02 | 1.67E-01 | 1.07E+00 | 9.59E-02 | 1.07E+00 |
| ENSG000001 | DCTN5 | 1.90E-02 | 1.67E-01 | 1.05E+00 | 7.45E-02 | 1.05E+00 |
| ENSG000001 | CANT1 | 1.90E-02 | 1.67E-01 | 1.10E+00 | 1.43E-01 | 1.10E+00 |
| ENSG000001 | RHBDF2 | 1.91E-02 | 1.68E-01 | 1.13E+00 | 1.71E-01 | 1.13E+00 |
| ENSG000002 | --- | 1.91E-02 | 1.68E-01 | 1.23E+00 | 2.97E-01 | 1.23E+00 |
| ENSG000002 | FAM83A-AS1 | 1.92E-02 | 1.68E-01 | 8.09E-01 | -3.05E-01 | -1.24E+00 |
| ENSG000002 | --- | 1.92E-02 | ? | 2.48E+00 | 1.31E+00 | 2.48E+00 |
| ENSG000002 | EPOP | 1.92E-02 | 1.68E-01 | 1.16E+00 | 2.12E-01 | 1.16E+00 |

| | | | | | | |
|---|---|---|---|---|---|---|
| ENSG000000 | CDK13 | 1.92E-02 | 1.68E-01 | 9.31E-01 | -1.04E-01 | -1.07E+00 |
| ENSG000002 | --- | 1.92E-02 | 1.68E-01 | 1.18E+00 | 2.37E-01 | 1.18E+00 |
| ENSG000002 | ZNF252P | 1.92E-02 | 1.68E-01 | 9.08E-01 | -1.39E-01 | -1.10E+00 |
| ENSG000002 | RHOXF1-AS1 | 1.93E-02 | ? | 2.07E+00 | 1.05E+00 | 2.07E+00 |
| ENSG000001 | GGPS1 | 1.93E-02 | 1.69E-01 | 9.30E-01 | -1.04E-01 | -1.07E+00 |
| ENSG000000 | ITGA6 | 1.93E-02 | 1.69E-01 | 9.24E-01 | -1.14E-01 | -1.08E+00 |
| ENSG000000 | BCAT1 | 1.94E-02 | 1.69E-01 | 8.64E-01 | -2.11E-01 | -1.16E+00 |
| ENSG000001 | ERMAP | 1.94E-02 | 1.69E-01 | 1.11E+00 | 1.56E-01 | 1.11E+00 |
| ENSG000001 | BRD3 | 1.94E-02 | 1.69E-01 | 1.11E+00 | 1.47E-01 | 1.11E+00 |
| ENSG000001 | SATL1 | 1.95E-02 | ? | 3.09E-01 | -1.69E+00 | -3.23E+00 |
| ENSG000001 | SCNN1B | 1.95E-02 | ? | 2.02E+00 | 1.01E+00 | 2.02E+00 |
| ENSG000000 | UBA6 | 1.95E-02 | 1.70E-01 | 9.28E-01 | -1.08E-01 | -1.08E+00 |
| ENSG000000 | OSBPL8 | 1.96E-02 | 1.70E-01 | 9.15E-01 | -1.28E-01 | -1.09E+00 |
| ENSG000002 | BTBD19 | 1.96E-02 | 1.70E-01 | 8.33E-01 | -2.63E-01 | -1.20E+00 |
| ENSG000002 | SRCIN1 | 1.97E-02 | 1.71E-01 | 1.32E+00 | 4.02E-01 | 1.32E+00 |
| ENSG000001 | KLHDC3 | 1.97E-02 | 1.71E-01 | 1.13E+00 | 1.74E-01 | 1.13E+00 |
| ENSG000001 | COX6C | 1.97E-02 | 1.71E-01 | 1.08E+00 | 1.12E-01 | 1.08E+00 |
| ENSG000001 | MAPKAPK2 | 1.97E-02 | 1.71E-01 | 9.23E-01 | -1.16E-01 | -1.08E+00 |
| ENSG000002 | --- | 1.98E-02 | 1.71E-01 | 1.33E+00 | 4.17E-01 | 1.33E+00 |
| ENSG000001 | SMARCD2 | 1.98E-02 | 1.72E-01 | 1.10E+00 | 1.34E-01 | 1.10E+00 |
| ENSG000000 | ANKRD10 | 1.99E-02 | 1.72E-01 | 8.92E-01 | -1.64E-01 | -1.12E+00 |
| ENSG000001 | TREX2 | 2.00E-02 | ? | 2.03E+00 | 1.02E+00 | 2.03E+00 |
| ENSG000001 | TRUB1 | 2.00E-02 | 1.73E-01 | 1.08E+00 | 1.06E-01 | 1.08E+00 |
| ENSG000001 | ISCA2 | 2.01E-02 | 1.73E-01 | 1.14E+00 | 1.95E-01 | 1.14E+00 |
| ENSG000001 | SNX12 | 2.01E-02 | 1.73E-01 | 1.09E+00 | 1.20E-01 | 1.09E+00 |
| ENSG000000 | RFXANK | 2.01E-02 | 1.74E-01 | 1.16E+00 | 2.12E-01 | 1.16E+00 |
| ENSG000001 | INPP5K | 2.01E-02 | 1.74E-01 | 1.11E+00 | 1.50E-01 | 1.11E+00 |
| ENSG000001 | TRIP10 | 2.02E-02 | 1.74E-01 | 1.08E+00 | 1.12E-01 | 1.08E+00 |
| ENSG000001 | TMEM168 | 2.03E-02 | 1.75E-01 | 1.09E+00 | 1.24E-01 | 1.09E+00 |
| ENSG000001 | HIP1R | 2.04E-02 | 1.75E-01 | 1.12E+00 | 1.69E-01 | 1.12E+00 |
| ENSG000001 | DCUN1D3 | 2.04E-02 | 1.75E-01 | 9.15E-01 | -1.28E-01 | -1.09E+00 |
| ENSG000000 | SMOX | 2.04E-02 | 1.75E-01 | 8.39E-01 | -2.53E-01 | -1.19E+00 |
| ENSG000001 | CDH11 | 2.04E-02 | 1.75E-01 | 8.93E-01 | -1.63E-01 | -1.12E+00 |
| ENSG000001 | TRMT13 | 2.04E-02 | 1.75E-01 | 8.75E-01 | -1.93E-01 | -1.14E+00 |
| ENSG000001 | RNASE1 | 2.04E-02 | ? | 2.00E+00 | 9.98E-01 | 2.00E+00 |
| ENSG000001 | TIMM8B | 2.04E-02 | 1.75E-01 | 1.10E+00 | 1.33E-01 | 1.10E+00 |
| ENSG000001 | ZFAND2A | 2.05E-02 | 1.76E-01 | 9.05E-01 | -1.44E-01 | -1.10E+00 |
| ENSG000001 | GGA2 | 2.06E-02 | 1.77E-01 | 1.09E+00 | 1.24E-01 | 1.09E+00 |
| ENSG000000 | DPP8 | 2.07E-02 | 1.77E-01 | 9.39E-01 | -9.00E-02 | -1.06E+00 |
| ENSG000001 | PEX6 | 2.07E-02 | 1.77E-01 | 1.19E+00 | 2.50E-01 | 1.19E+00 |
| ENSG000001 | LOXL1 | 2.08E-02 | 1.78E-01 | 1.18E+00 | 2.39E-01 | 1.18E+00 |
| ENSG000001 | ITSN2 | 2.08E-02 | 1.78E-01 | 8.99E-01 | -1.54E-01 | -1.11E+00 |
| ENSG000001 | PATJ | 2.08E-02 | 1.78E-01 | 9.19E-01 | -1.23E-01 | -1.09E+00 |
| ENSG000001 | IL1RAPL1 | 2.09E-02 | ? | 4.27E-01 | -1.23E+00 | -2.34E+00 |
| ENSG000002 | SUNO1 | 2.09E-02 | ? | 1.78E+00 | 8.35E-01 | 1.78E+00 |

| | | | | | | |
|---|---|---|---|---|---|---|
| ENSG000001 | GOLGA7B | 2.09E-02 | 1.79E-01 | 1.18E+00 | 2.42E-01 | 1.18E+00 |
| ENSG00000C | POLA2 | 2.09E-02 | 1.79E-01 | 1.21E+00 | 2.76E-01 | 1.21E+00 |
| ENSG000001 | ANK3 | 2.09E-02 | 1.79E-01 | 1.13E+00 | 1.74E-01 | 1.13E+00 |
| ENSG00000C | ZNF37A | 2.10E-02 | 1.79E-01 | 8.96E-01 | -1.58E-01 | -1.12E+00 |
| ENSG000001 | ZNF497 | 2.10E-02 | ? | 1.76E+00 | 8.20E-01 | 1.76E+00 |
| ENSG000001 | ATRAID | 2.11E-02 | 1.79E-01 | 1.09E+00 | 1.28E-01 | 1.09E+00 |
| ENSG000001 | EEF2KMT | 2.11E-02 | 1.79E-01 | 1.15E+00 | 2.08E-01 | 1.15E+00 |
| ENSG000001 | CHD7 | 2.11E-02 | 1.79E-01 | 9.12E-01 | -1.32E-01 | -1.10E+00 |
| ENSG00000C | EPB41L3 | 2.11E-02 | 1.79E-01 | 7.72E-01 | -3.73E-01 | -1.29E+00 |
| ENSG000001 | LCE1E | 2.11E-02 | ? | 1.85E+00 | 8.86E-01 | 1.85E+00 |
| ENSG000001 | GRIA3 | 2.12E-02 | 1.80E-01 | 1.51E+00 | 5.90E-01 | 1.51E+00 |
| ENSG000002 | --- | 2.12E-02 | ? | 6.82E-01 | -5.52E-01 | -1.47E+00 |
| ENSG000001 | GSDMC | 2.12E-02 | 1.80E-01 | 8.79E-01 | -1.86E-01 | -1.14E+00 |
| ENSG00000C | SMARCE1 | 2.12E-02 | 1.80E-01 | 9.04E-01 | -1.46E-01 | -1.11E+00 |
| ENSG000001 | PLIN4 | 2.12E-02 | 1.80E-01 | 1.26E+00 | 3.31E-01 | 1.26E+00 |
| ENSG000001 | LRRC39 | 2.12E-02 | ? | 6.50E-01 | -6.21E-01 | -1.54E+00 |
| ENSG000001 | MED10 | 2.13E-02 | 1.81E-01 | 9.22E-01 | -1.18E-01 | -1.09E+00 |
| ENSG000001 | HEY2 | 2.13E-02 | ? | 1.74E+00 | 8.01E-01 | 1.74E+00 |
| ENSG000001 | SOX2 | 2.13E-02 | 1.81E-01 | 7.72E-01 | -3.73E-01 | -1.29E+00 |
| ENSG000001 | TMEM171 | 2.13E-02 | 1.81E-01 | 1.28E+00 | 3.59E-01 | 1.28E+00 |
| ENSG000001 | EPB41L4A | 2.13E-02 | 1.81E-01 | 8.90E-01 | -1.67E-01 | -1.12E+00 |
| ENSG000001 | CARD11 | 2.13E-02 | 1.81E-01 | 9.13E-01 | -1.32E-01 | -1.10E+00 |
| ENSG00000C | RPL31 | 2.14E-02 | 1.81E-01 | 1.09E+00 | 1.27E-01 | 1.09E+00 |
| ENSG000001 | SINHCAF | 2.14E-02 | 1.81E-01 | 1.09E+00 | 1.20E-01 | 1.09E+00 |
| ENSG00000C | ZNF207 | 2.14E-02 | 1.81E-01 | 9.50E-01 | -7.40E-02 | -1.05E+00 |
| ENSG000001 | --- | 2.15E-02 | ? | 4.22E-01 | -1.24E+00 | -2.37E+00 |
| ENSG000001 | MARCHF7 | 2.15E-02 | 1.81E-01 | 9.15E-01 | -1.28E-01 | -1.09E+00 |
| ENSG000001 | NKTR | 2.16E-02 | 1.82E-01 | 8.64E-01 | -2.11E-01 | -1.16E+00 |
| ENSG000001 | TIMM22 | 2.16E-02 | 1.82E-01 | 1.08E+00 | 1.09E-01 | 1.08E+00 |
| ENSG000001 | LRRC57 | 2.17E-02 | 1.83E-01 | 1.08E+00 | 1.10E-01 | 1.08E+00 |
| ENSG00000C | ST3GAL1 | 2.17E-02 | 1.83E-01 | 9.30E-01 | -1.05E-01 | -1.08E+00 |
| ENSG00000C | NOA1 | 2.18E-02 | 1.83E-01 | 1.09E+00 | 1.29E-01 | 1.09E+00 |
| ENSG00000C | DPYSL2 | 2.18E-02 | 1.83E-01 | 9.05E-01 | -1.44E-01 | -1.10E+00 |
| ENSG000002 | CNTF | 2.18E-02 | ? | 5.62E-01 | -8.31E-01 | -1.78E+00 |
| ENSG000001 | DBT | 2.18E-02 | 1.84E-01 | 1.10E+00 | 1.41E-01 | 1.10E+00 |
| ENSG000002 | N4BP2L2 | 2.19E-02 | 1.84E-01 | 8.92E-01 | -1.64E-01 | -1.12E+00 |
| ENSG000001 | ADM2 | 2.20E-02 | 1.84E-01 | 7.44E-01 | -4.28E-01 | -1.34E+00 |
| ENSG000001 | PYCR1 | 2.20E-02 | 1.84E-01 | 1.16E+00 | 2.13E-01 | 1.16E+00 |
| ENSG00000C | TRAF3IP2 | 2.20E-02 | 1.84E-01 | 1.06E+00 | 8.41E-02 | 1.06E+00 |
| ENSG000001 | COX7C | 2.20E-02 | 1.84E-01 | 1.10E+00 | 1.38E-01 | 1.10E+00 |
| ENSG000001 | LRRC42 | 2.21E-02 | 1.85E-01 | 1.14E+00 | 1.91E-01 | 1.14E+00 |
| ENSG000001 | HSPH1 | 2.21E-02 | 1.85E-01 | 9.19E-01 | -1.22E-01 | -1.09E+00 |
| ENSG000001 | IL36G | 2.21E-02 | 1.85E-01 | 1.31E+00 | 3.94E-01 | 1.31E+00 |
| ENSG000002 | --- | 2.21E-02 | 1.85E-01 | 1.26E+00 | 3.36E-01 | 1.26E+00 |
| ENSG000001 | ZBTB4 | 2.22E-02 | 1.85E-01 | 1.06E+00 | 8.35E-02 | 1.06E+00 |

| | | | | | | |
|---|---|---|---|---|---|---|
| ENSG000002 | LINC03038 | 2.22E-02 | ? | 4.67E-01 | -1.10E+00 | -2.14E+00 |
| ENSG000001 | GINS1 | 2.22E-02 | 1.86E-01 | 8.59E-01 | -2.19E-01 | -1.16E+00 |
| ENSG00000C | NTN1 | 2.22E-02 | 1.86E-01 | 1.64E+00 | 7.11E-01 | 1.64E+00 |
| ENSG000001 | KLF5 | 2.22E-02 | 1.86E-01 | 8.88E-01 | -1.72E-01 | -1.13E+00 |
| ENSG000002 | NUDT18 | 2.23E-02 | 1.86E-01 | 1.19E+00 | 2.51E-01 | 1.19E+00 |
| ENSG000001 | CDKN1B | 2.23E-02 | 1.86E-01 | 1.10E+00 | 1.43E-01 | 1.10E+00 |
| ENSG000001 | LAPTM4B | 2.23E-02 | 1.86E-01 | 1.07E+00 | 1.00E-01 | 1.07E+00 |
| ENSG00000C | IL32 | 2.23E-02 | 1.86E-01 | 1.12E+00 | 1.59E-01 | 1.12E+00 |
| ENSG000001 | NIPA2 | 2.23E-02 | 1.86E-01 | 9.22E-01 | -1.16E-01 | -1.08E+00 |
| ENSG00000C | SLK | 2.24E-02 | 1.86E-01 | 8.94E-01 | -1.62E-01 | -1.12E+00 |
| ENSG000001 | RGL1 | 2.25E-02 | 1.87E-01 | 1.24E+00 | 3.09E-01 | 1.24E+00 |
| ENSG000001 | PLEKHM2 | 2.25E-02 | 1.87E-01 | 1.12E+00 | 1.61E-01 | 1.12E+00 |
| ENSG000001 | HMGCL | 2.25E-02 | 1.87E-01 | 1.11E+00 | 1.48E-01 | 1.11E+00 |
| ENSG000001 | SGIP1 | 2.25E-02 | 1.87E-01 | 1.30E+00 | 3.74E-01 | 1.30E+00 |
| ENSG000002 | PDE2A-AS2 | 2.25E-02 | 1.87E-01 | 7.67E-01 | -3.83E-01 | -1.30E+00 |
| ENSG000001 | TMEM81 | 2.25E-02 | 1.87E-01 | 7.65E-01 | -3.87E-01 | -1.31E+00 |
| ENSG000001 | TMEM134 | 2.25E-02 | 1.87E-01 | 1.18E+00 | 2.36E-01 | 1.18E+00 |
| ENSG000001 | INKA1 | 2.25E-02 | 1.87E-01 | 1.25E+00 | 3.20E-01 | 1.25E+00 |
| ENSG000001 | IRAK4 | 2.26E-02 | 1.87E-01 | 1.11E+00 | 1.44E-01 | 1.11E+00 |
| ENSG000001 | KDELR2 | 2.26E-02 | 1.87E-01 | 9.43E-01 | -8.45E-02 | -1.06E+00 |
| ENSG000001 | ADCY1 | 2.26E-02 | 1.87E-01 | 1.23E+00 | 2.97E-01 | 1.23E+00 |
| ENSG000001 | NFASC | 2.26E-02 | 1.87E-01 | 1.26E+00 | 3.36E-01 | 1.26E+00 |
| ENSG000001 | FAM171B | 2.26E-02 | 1.87E-01 | 1.18E+00 | 2.38E-01 | 1.18E+00 |
| ENSG000001 | ILK | 2.26E-02 | 1.87E-01 | 1.08E+00 | 1.11E-01 | 1.08E+00 |
| ENSG000001 | SLC25A33 | 2.26E-02 | 1.87E-01 | 1.16E+00 | 2.16E-01 | 1.16E+00 |
| ENSG000001 | STK32C | 2.26E-02 | 1.87E-01 | 1.21E+00 | 2.76E-01 | 1.21E+00 |
| ENSG000001 | LRATD2 | 2.27E-02 | 1.87E-01 | 9.35E-01 | -9.64E-02 | -1.07E+00 |
| ENSG000001 | SNU13 | 2.27E-02 | 1.87E-01 | 1.09E+00 | 1.23E-01 | 1.09E+00 |
| ENSG000001 | SLC27A2 | 2.27E-02 | ? | 2.32E+00 | 1.22E+00 | 2.32E+00 |
| ENSG000001 | RYR1 | 2.27E-02 | ? | 1.46E+00 | 5.49E-01 | 1.46E+00 |
| ENSG000001 | UMPS | 2.28E-02 | 1.87E-01 | 1.09E+00 | 1.19E-01 | 1.09E+00 |
| ENSG000001 | DCBLD1 | 2.28E-02 | 1.87E-01 | 1.10E+00 | 1.38E-01 | 1.10E+00 |
| ENSG00000C | FGFR2 | 2.28E-02 | 1.87E-01 | 1.12E+00 | 1.65E-01 | 1.12E+00 |
| ENSG000001 | BANF1 | 2.28E-02 | 1.87E-01 | 1.07E+00 | 1.01E-01 | 1.07E+00 |
| ENSG000002 | DDR1 | 2.29E-02 | 1.88E-01 | 1.12E+00 | 1.68E-01 | 1.12E+00 |
| ENSG000001 | ZDHHC2 | 2.30E-02 | 1.89E-01 | 8.86E-01 | -1.74E-01 | -1.13E+00 |
| ENSG000001 | DUS2 | 2.30E-02 | 1.89E-01 | 1.13E+00 | 1.78E-01 | 1.13E+00 |
| ENSG000001 | PRKCE | 2.31E-02 | 1.89E-01 | 1.21E+00 | 2.70E-01 | 1.21E+00 |
| ENSG000002 | --- | 2.31E-02 | ? | 4.44E-01 | -1.17E+00 | -2.25E+00 |
| ENSG000001 | GALNT18 | 2.32E-02 | 1.90E-01 | 1.11E+00 | 1.52E-01 | 1.11E+00 |
| ENSG000002 | --- | 2.32E-02 | ? | 1.59E+00 | 6.74E-01 | 1.59E+00 |
| ENSG000001 | ARHGEF2 | 2.33E-02 | 1.90E-01 | 9.39E-01 | -9.01E-02 | -1.06E+00 |
| ENSG000001 | NT5C3A | 2.33E-02 | 1.90E-01 | 8.92E-01 | -1.65E-01 | -1.12E+00 |
| ENSG000002 | LINC00847 | 2.33E-02 | 1.91E-01 | 1.20E+00 | 2.61E-01 | 1.20E+00 |
| ENSG000001 | PROSER1 | 2.33E-02 | 1.91E-01 | 8.90E-01 | -1.68E-01 | -1.12E+00 |

| | | | | | | |
|---|---|---|---|---|---|---|
| ENSG00000C | PNPLA4 | 2.33E-02 | 1.91E-01 | 1.08E+00 | 1.16E-01 | 1.08E+00 |
| ENSG000001 | TMEM200A | 2.34E-02 | ? | 5.99E-01 | -7.40E-01 | -1.67E+00 |
| ENSG000001 | TOPORS | 2.34E-02 | 1.91E-01 | 9.07E-01 | -1.41E-01 | -1.10E+00 |
| ENSG000001 | HCFC1 | 2.34E-02 | 1.91E-01 | 9.30E-01 | -1.05E-01 | -1.08E+00 |
| ENSG000001 | EDIL3 | 2.35E-02 | 1.91E-01 | 8.17E-01 | -2.92E-01 | -1.22E+00 |
| ENSG000002 | MTND5P11 | 2.35E-02 | ? | 4.79E-01 | -1.06E+00 | -2.09E+00 |
| ENSG000001 | TFEC | 2.35E-02 | 1.92E-01 | 7.83E-01 | -3.53E-01 | -1.28E+00 |
| ENSG000001 | TLCD5 | 2.36E-02 | 1.92E-01 | 1.19E+00 | 2.52E-01 | 1.19E+00 |
| ENSG000002 | --- | 2.37E-02 | 1.93E-01 | 8.50E-01 | -2.34E-01 | -1.18E+00 |
| ENSG000002 | --- | 2.37E-02 | 1.93E-01 | 7.82E-01 | -3.55E-01 | -1.28E+00 |
| ENSG000002 | --- | 2.38E-02 | 1.93E-01 | 5.87E-01 | -7.69E-01 | -1.70E+00 |
| ENSG000001 | MCTP1 | 2.38E-02 | 1.93E-01 | 9.00E-01 | -1.53E-01 | -1.11E+00 |
| ENSG000001 | C10orf82 | 2.38E-02 | ? | 5.83E-01 | -7.80E-01 | -1.72E+00 |
| ENSG00000C | CPNE3 | 2.38E-02 | 1.93E-01 | 9.33E-01 | -1.00E-01 | -1.07E+00 |
| ENSG000001 | RPL35 | 2.39E-02 | 1.94E-01 | 1.07E+00 | 9.52E-02 | 1.07E+00 |
| ENSG000001 | CUL4A | 2.39E-02 | 1.94E-01 | 9.39E-01 | -9.08E-02 | -1.06E+00 |
| ENSG000001 | DKK1 | 2.40E-02 | 1.95E-01 | 8.54E-01 | -2.28E-01 | -1.17E+00 |
| ENSG000002 | --- | 2.40E-02 | ? | 5.02E-01 | -9.96E-01 | -1.99E+00 |
| ENSG000001 | MTHFR | 2.41E-02 | 1.95E-01 | 1.12E+00 | 1.66E-01 | 1.12E+00 |
| ENSG000001 | LYPD6B | 2.41E-02 | ? | 1.51E+00 | 5.94E-01 | 1.51E+00 |
| ENSG000002 | GTF2IP4 | 2.41E-02 | 1.95E-01 | 1.11E+00 | 1.53E-01 | 1.11E+00 |
| ENSG000002 | ZNF525 | 2.41E-02 | 1.95E-01 | 8.97E-01 | -1.57E-01 | -1.12E+00 |
| ENSG000001 | GJC1 | 2.41E-02 | 1.95E-01 | 1.12E+00 | 1.62E-01 | 1.12E+00 |
| ENSG000001 | RAB40C | 2.42E-02 | 1.96E-01 | 1.16E+00 | 2.11E-01 | 1.16E+00 |
| ENSG000001 | UBIAD1 | 2.43E-02 | 1.96E-01 | 1.09E+00 | 1.25E-01 | 1.09E+00 |
| ENSG000001 | CBR3 | 2.43E-02 | 1.96E-01 | 7.34E-01 | -4.46E-01 | -1.36E+00 |
| ENSG000002 | --- | 2.43E-02 | ? | 2.99E+00 | 1.58E+00 | 2.99E+00 |
| ENSG000001 | MPZ | 2.43E-02 | 1.96E-01 | 7.69E-01 | -3.79E-01 | -1.30E+00 |
| ENSG000001 | DCAKD | 2.44E-02 | 1.97E-01 | 1.13E+00 | 1.75E-01 | 1.13E+00 |
| ENSG000001 | ZNF691 | 2.45E-02 | 1.97E-01 | 8.40E-01 | -2.52E-01 | -1.19E+00 |
| ENSG000001 | RAB14 | 2.45E-02 | 1.97E-01 | 1.06E+00 | 8.52E-02 | 1.06E+00 |
| ENSG000002 | --- | 2.45E-02 | 1.98E-01 | 1.19E+00 | 2.48E-01 | 1.19E+00 |
| ENSG000001 | AIP | 2.46E-02 | 1.98E-01 | 1.15E+00 | 1.97E-01 | 1.15E+00 |
| ENSG000002 | --- | 2.46E-02 | ? | 2.71E+00 | 1.44E+00 | 2.71E+00 |
| ENSG000002 | ANKRD34A | 2.46E-02 | 1.98E-01 | 1.33E+00 | 4.14E-01 | 1.33E+00 |
| ENSG000001 | ERCC5 | 2.46E-02 | 1.98E-01 | 9.26E-01 | -1.11E-01 | -1.08E+00 |
| ENSG000001 | ZNF714 | 2.46E-02 | 1.98E-01 | 8.63E-01 | -2.12E-01 | -1.16E+00 |
| ENSG00000C | POLR1H | 2.47E-02 | 1.98E-01 | 1.16E+00 | 2.18E-01 | 1.16E+00 |
| ENSG000001 | MPRIP | 2.47E-02 | 1.98E-01 | 9.40E-01 | -8.85E-02 | -1.06E+00 |
| ENSG000001 | EIF2A | 2.47E-02 | 1.98E-01 | 9.08E-01 | -1.39E-01 | -1.10E+00 |
| ENSG000001 | NKAIN2 | 2.48E-02 | ? | 3.74E-01 | -1.42E+00 | -2.68E+00 |
| ENSG000001 | PLIN1 | 2.48E-02 | ? | 2.74E-01 | -1.87E+00 | -3.65E+00 |
| ENSG000002 | MRPL20 | 2.48E-02 | 1.99E-01 | 1.08E+00 | 1.12E-01 | 1.08E+00 |
| ENSG000001 | CDC37 | 2.49E-02 | 1.99E-01 | 9.49E-01 | -7.57E-02 | -1.05E+00 |
| ENSG00000C | DYNLL1 | 2.49E-02 | 1.99E-01 | 1.06E+00 | 8.34E-02 | 1.06E+00 |

| | | | | | | |
|---|---|---|---|---|---|---|
| ENSG000001 | RPS2 | 2.49E-02 | 1.99E-01 | 1.08E+00 | 1.06E-01 | 1.08E+00 |
| ENSG00000C | RNF13 | 2.49E-02 | 1.99E-01 | 1.07E+00 | 9.27E-02 | 1.07E+00 |
| ENSG000001 | SLC2A1 | 2.50E-02 | 1.99E-01 | 1.10E+00 | 1.39E-01 | 1.10E+00 |
| ENSG00000C | FAM168A | 2.50E-02 | 1.99E-01 | 1.09E+00 | 1.20E-01 | 1.09E+00 |
| ENSG000001 | SPATA2 | 2.50E-02 | 1.99E-01 | 1.13E+00 | 1.78E-01 | 1.13E+00 |
| ENSG00000C | NDUFB4 | 2.50E-02 | 1.99E-01 | 1.06E+00 | 8.75E-02 | 1.06E+00 |
| ENSG000001 | TRANK1 | 2.50E-02 | 1.99E-01 | 1.14E+00 | 1.85E-01 | 1.14E+00 |
| ENSG000001 | DPYD | 2.50E-02 | 1.99E-01 | 8.29E-01 | -2.71E-01 | -1.21E+00 |
| ENSG000001 | RPL23A | 2.50E-02 | 1.99E-01 | 1.09E+00 | 1.25E-01 | 1.09E+00 |
| ENSG000001 | PSMA2 | 2.51E-02 | 2.00E-01 | 1.09E+00 | 1.28E-01 | 1.09E+00 |
| ENSG000001 | OVOL1 | 2.51E-02 | 2.00E-01 | 1.37E+00 | 4.56E-01 | 1.37E+00 |
| ENSG000002 | --- | 2.51E-02 | 2.00E-01 | 8.18E-01 | -2.90E-01 | -1.22E+00 |
| ENSG00000C | TIMP2 | 2.51E-02 | 2.00E-01 | 1.13E+00 | 1.71E-01 | 1.13E+00 |
| ENSG00000C | ATP6V0A1 | 2.52E-02 | 2.00E-01 | 9.11E-01 | -1.34E-01 | -1.10E+00 |
| ENSG000001 | C16orf91 | 2.52E-02 | 2.00E-01 | 1.17E+00 | 2.32E-01 | 1.17E+00 |
| ENSG000001 | XKR6 | 2.52E-02 | 2.00E-01 | 7.45E-01 | -4.25E-01 | -1.34E+00 |
| ENSG00000C | RPL18 | 2.52E-02 | 2.00E-01 | 1.13E+00 | 1.83E-01 | 1.13E+00 |
| ENSG000001 | ZNF217 | 2.52E-02 | 2.00E-01 | 1.10E+00 | 1.37E-01 | 1.10E+00 |
| ENSG000001 | ZDHHC1 | 2.52E-02 | 2.00E-01 | 1.25E+00 | 3.17E-01 | 1.25E+00 |
| ENSG000002 | YWHABP2 | 2.52E-02 ? | | 3.22E-01 | -1.64E+00 | -3.11E+00 |
| ENSG000002 | --- | 2.53E-02 | 2.00E-01 | 1.29E+00 | 3.64E-01 | 1.29E+00 |
| ENSG000001 | ZNF589 | 2.53E-02 | 2.00E-01 | 8.71E-01 | -2.00E-01 | -1.15E+00 |
| ENSG000001 | AP1B1 | 2.53E-02 | 2.00E-01 | 1.08E+00 | 1.07E-01 | 1.08E+00 |
| ENSG000001 | HS1BP3 | 2.53E-02 | 2.00E-01 | 1.13E+00 | 1.72E-01 | 1.13E+00 |
| ENSG00000C | SLC26A4 | 2.53E-02 ? | | 6.21E-01 | -6.88E-01 | -1.61E+00 |
| ENSG000001 | PDE8B | 2.53E-02 | 2.00E-01 | 7.22E-01 | -4.70E-01 | -1.39E+00 |
| ENSG000001 | ALKBH7 | 2.54E-02 | 2.00E-01 | 1.33E+00 | 4.12E-01 | 1.33E+00 |
| ENSG00000C | PHPT1 | 2.54E-02 | 2.00E-01 | 1.12E+00 | 1.60E-01 | 1.12E+00 |
| ENSG000002 | NKILA | 2.54E-02 | 2.00E-01 | 8.14E-01 | -2.97E-01 | -1.23E+00 |
| ENSG000001 | DDR2 | 2.54E-02 ? | | 6.54E-01 | -6.13E-01 | -1.53E+00 |
| ENSG000001 | ZNF816 | 2.54E-02 | 2.00E-01 | 1.14E+00 | 1.85E-01 | 1.14E+00 |
| ENSG000001 | PHF20L1 | 2.54E-02 | 2.00E-01 | 9.12E-01 | -1.32E-01 | -1.10E+00 |
| ENSG000002 | CCNQ | 2.55E-02 | 2.01E-01 | 1.12E+00 | 1.70E-01 | 1.12E+00 |
| ENSG000001 | ASB13 | 2.55E-02 | 2.01E-01 | 1.18E+00 | 2.37E-01 | 1.18E+00 |
| ENSG000001 | FAM98A | 2.55E-02 | 2.01E-01 | 8.98E-01 | -1.55E-01 | -1.11E+00 |
| ENSG000001 | TBCK | 2.55E-02 | 2.01E-01 | 1.09E+00 | 1.24E-01 | 1.09E+00 |
| ENSG000001 | PLXNC1 | 2.56E-02 | 2.01E-01 | 1.20E+00 | 2.66E-01 | 1.20E+00 |
| ENSG000001 | L3MBTL2 | 2.56E-02 | 2.01E-01 | 1.10E+00 | 1.32E-01 | 1.10E+00 |
| ENSG000002 | RDH14 | 2.56E-02 | 2.01E-01 | 9.14E-01 | -1.30E-01 | -1.09E+00 |
| ENSG000002 | MT-TS1 | 2.56E-02 ? | | 3.07E-01 | -1.70E+00 | -3.25E+00 |
| ENSG000001 | SWT1 | 2.57E-02 | 2.01E-01 | 8.58E-01 | -2.21E-01 | -1.17E+00 |
| ENSG00000C | CDHR2 | 2.57E-02 ? | | 1.69E+00 | 7.57E-01 | 1.69E+00 |
| ENSG000001 | TXN2 | 2.57E-02 | 2.01E-01 | 1.07E+00 | 9.29E-02 | 1.07E+00 |
| ENSG000001 | ID4 | 2.57E-02 ? | | 1.82E+00 | 8.64E-01 | 1.82E+00 |
| ENSG000001 | SLC10A3 | 2.57E-02 | 2.02E-01 | 1.14E+00 | 1.84E-01 | 1.14E+00 |

| | | | | | | |
|---|---|---|---|---|---|---|
| ENSG000002 | --- | 2.58E-02 | ? | 3.23E+00 | 1.69E+00 | 3.23E+00 |
| ENSG000001 | KIF5A | 2.58E-02 | 2.02E-01 | 7.53E-01 | -4.09E-01 | -1.33E+00 |
| ENSG000001 | TNFRSF10C | 2.58E-02 | 2.02E-01 | 1.22E+00 | 2.84E-01 | 1.22E+00 |
| ENSG00000C | ZFAT | 2.59E-02 | 2.02E-01 | 1.12E+00 | 1.66E-01 | 1.12E+00 |
| ENSG000001 | PLGLB2 | 2.59E-02 | ? | 5.18E-01 | -9.50E-01 | -1.93E+00 |
| ENSG000001 | LCORL | 2.59E-02 | 2.02E-01 | 8.61E-01 | -2.16E-01 | -1.16E+00 |
| ENSG000001 | ST18 | 2.59E-02 | ? | 6.51E-01 | -6.19E-01 | -1.54E+00 |
| ENSG000001 | FABP5 | 2.59E-02 | 2.02E-01 | 8.70E-01 | -2.00E-01 | -1.15E+00 |
| ENSG000002 | ZNF345 | 2.59E-02 | 2.02E-01 | 7.63E-01 | -3.90E-01 | -1.31E+00 |
| ENSG000001 | MAFF | 2.59E-02 | 2.02E-01 | 9.14E-01 | -1.29E-01 | -1.09E+00 |
| ENSG000002 | ATP5PO | 2.59E-02 | 2.02E-01 | 1.09E+00 | 1.19E-01 | 1.09E+00 |
| ENSG00000C | PPP1R12A | 2.60E-02 | 2.02E-01 | 9.11E-01 | -1.34E-01 | -1.10E+00 |
| ENSG000001 | SCN2A | 2.60E-02 | 2.02E-01 | 1.30E+00 | 3.84E-01 | 1.30E+00 |
| ENSG000001 | KXD1 | 2.60E-02 | 2.02E-01 | 9.28E-01 | -1.08E-01 | -1.08E+00 |
| ENSG000001 | HSPB1 | 2.60E-02 | 2.02E-01 | 1.16E+00 | 2.14E-01 | 1.16E+00 |
| ENSG000001 | TSC22D2 | 2.60E-02 | 2.02E-01 | 9.27E-01 | -1.10E-01 | -1.08E+00 |
| ENSG000002 | ITPRIPL2 | 2.60E-02 | 2.02E-01 | 1.12E+00 | 1.65E-01 | 1.12E+00 |
| ENSG000001 | GAS2L3 | 2.61E-02 | 2.02E-01 | 7.87E-01 | -3.46E-01 | -1.27E+00 |
| ENSG000001 | TRIB1 | 2.61E-02 | 2.03E-01 | 8.88E-01 | -1.72E-01 | -1.13E+00 |
| ENSG000002 | POU5F1B | 2.61E-02 | 2.03E-01 | 6.98E-01 | -5.19E-01 | -1.43E+00 |
| ENSG000001 | NHLRC2 | 2.61E-02 | 2.03E-01 | 8.89E-01 | -1.70E-01 | -1.13E+00 |
| ENSG000002 | NKX1-2 | 2.61E-02 | 2.03E-01 | 1.25E+00 | 3.23E-01 | 1.25E+00 |
| ENSG00000C | MRPS24 | 2.62E-02 | 2.03E-01 | 1.12E+00 | 1.64E-01 | 1.12E+00 |
| ENSG000001 | TUBGCP3 | 2.62E-02 | 2.03E-01 | 1.10E+00 | 1.33E-01 | 1.10E+00 |
| ENSG000001 | SERPINB8 | 2.62E-02 | 2.03E-01 | 9.33E-01 | -1.01E-01 | -1.07E+00 |
| ENSG000002 | --- | 2.62E-02 | ? | 3.38E+00 | 1.76E+00 | 3.38E+00 |
| ENSG000001 | TLCD3A | 2.62E-02 | 2.03E-01 | 1.07E+00 | 9.99E-02 | 1.07E+00 |
| ENSG000001 | TPM3 | 2.62E-02 | 2.03E-01 | 9.08E-01 | -1.39E-01 | -1.10E+00 |
| ENSG000002 | TMSB4XP6 | 2.63E-02 | 2.03E-01 | 7.28E-01 | -4.57E-01 | -1.37E+00 |
| ENSG000001 | PADI1 | 2.63E-02 | 2.03E-01 | 7.76E-01 | -3.66E-01 | -1.29E+00 |
| ENSG000001 | OTX1 | 2.64E-02 | 2.04E-01 | 8.50E-01 | -2.34E-01 | -1.18E+00 |
| ENSG000001 | FAM199X | 2.64E-02 | 2.04E-01 | 9.25E-01 | -1.12E-01 | -1.08E+00 |
| ENSG000001 | RNF227 | 2.64E-02 | 2.04E-01 | 1.18E+00 | 2.38E-01 | 1.18E+00 |
| ENSG000001 | DHRS4L2 | 2.65E-02 | 2.04E-01 | 1.12E+00 | 1.69E-01 | 1.12E+00 |
| ENSG000001 | UNC119B | 2.66E-02 | 2.05E-01 | 1.12E+00 | 1.58E-01 | 1.12E+00 |
| ENSG000001 | PIGZ | 2.66E-02 | 2.05E-01 | 1.16E+00 | 2.11E-01 | 1.16E+00 |
| ENSG000001 | KTN1 | 2.67E-02 | 2.05E-01 | 8.99E-01 | -1.54E-01 | -1.11E+00 |
| ENSG000002 | RPS29 | 2.67E-02 | 2.05E-01 | 1.21E+00 | 2.77E-01 | 1.21E+00 |
| ENSG00000C | LIMA1 | 2.67E-02 | 2.05E-01 | 1.10E+00 | 1.35E-01 | 1.10E+00 |
| ENSG000001 | PROCR | 2.68E-02 | 2.06E-01 | 1.12E+00 | 1.64E-01 | 1.12E+00 |
| ENSG000001 | SLC18B1 | 2.68E-02 | 2.06E-01 | 8.96E-01 | -1.58E-01 | -1.12E+00 |
| ENSG000002 | --- | 2.68E-02 | ? | 4.14E+00 | 2.05E+00 | 4.14E+00 |
| ENSG000001 | SLTM | 2.70E-02 | 2.07E-01 | 9.08E-01 | -1.39E-01 | -1.10E+00 |
| ENSG000001 | CHEK1 | 2.70E-02 | 2.07E-01 | 8.71E-01 | -1.99E-01 | -1.15E+00 |
| ENSG000001 | XXYLT1 | 2.70E-02 | 2.07E-01 | 1.11E+00 | 1.53E-01 | 1.11E+00 |

| | | | | | | |
|---|---|---|---|---|---|---|
| ENSG000001 | MRPL24 | 2.71E-02 | 2.07E-01 | 1.08E+00 | 1.06E-01 | 1.08E+00 |
| ENSG000002 | ARHGEF28 | 2.71E-02 | 2.07E-01 | 9.09E-01 | -1.38E-01 | -1.10E+00 |
| ENSG000001 | SLC25A20 | 2.71E-02 | 2.07E-01 | 1.13E+00 | 1.75E-01 | 1.13E+00 |
| ENSG000001 | PTPN1 | 2.71E-02 | 2.07E-01 | 9.33E-01 | -9.96E-02 | -1.07E+00 |
| ENSG00000C | HLTF | 2.71E-02 | 2.07E-01 | 9.01E-01 | -1.51E-01 | -1.11E+00 |
| ENSG000001 | RPL14P1 | 2.72E-02 | 2.08E-01 | 1.14E+00 | 1.94E-01 | 1.14E+00 |
| ENSG00000C | WDR76 | 2.72E-02 | 2.08E-01 | 6.99E-01 | -5.17E-01 | -1.43E+00 |
| ENSG000001 | ZMIZ2 | 2.73E-02 | 2.08E-01 | 9.21E-01 | -1.19E-01 | -1.09E+00 |
| ENSG000002 | --- | 2.73E-02 | 2.08E-01 | 8.37E-01 | -2.57E-01 | -1.19E+00 |
| ENSG000001 | S100PBP | 2.73E-02 | 2.08E-01 | 1.10E+00 | 1.33E-01 | 1.10E+00 |
| ENSG000001 | STAT6 | 2.73E-02 | 2.08E-01 | 1.08E+00 | 1.07E-01 | 1.08E+00 |
| ENSG000001 | ZC3H4 | 2.73E-02 | 2.08E-01 | 1.08E+00 | 1.06E-01 | 1.08E+00 |
| ENSG000001 | SLC1A4 | 2.74E-02 | 2.09E-01 | 1.10E+00 | 1.34E-01 | 1.10E+00 |
| ENSG000001 | IQGAP3 | 2.74E-02 | 2.09E-01 | 8.14E-01 | -2.97E-01 | -1.23E+00 |
| ENSG000001 | MRPL34 | 2.74E-02 | 2.09E-01 | 1.15E+00 | 2.05E-01 | 1.15E+00 |
| ENSG000002 | PISD | 2.75E-02 | 2.09E-01 | 1.09E+00 | 1.29E-01 | 1.09E+00 |
| ENSG00000C | DDT | 2.75E-02 | 2.09E-01 | 1.15E+00 | 1.99E-01 | 1.15E+00 |
| ENSG000001 | FUOM | 2.75E-02 | 2.09E-01 | 1.39E+00 | 4.78E-01 | 1.39E+00 |
| ENSG000001 | HS6ST1 | 2.76E-02 | 2.09E-01 | 1.14E+00 | 1.95E-01 | 1.14E+00 |
| ENSG000002 | ERFL | 2.76E-02 | ? | 2.64E+00 | 1.40E+00 | 2.64E+00 |
| ENSG000002 | FAM111A-DT | 2.76E-02 | 2.09E-01 | 1.25E+00 | 3.25E-01 | 1.25E+00 |
| ENSG000001 | NDUFA4L2 | 2.77E-02 | 2.10E-01 | 1.13E+00 | 1.76E-01 | 1.13E+00 |
| ENSG000002 | CDKN2B-AS1 | 2.78E-02 | ? | 1.48E+00 | 5.70E-01 | 1.48E+00 |
| ENSG000001 | PLEKHA2 | 2.78E-02 | 2.11E-01 | 1.10E+00 | 1.37E-01 | 1.10E+00 |
| ENSG000001 | BGN | 2.78E-02 | ? | 5.29E-01 | -9.20E-01 | -1.89E+00 |
| ENSG000001 | DYSF | 2.78E-02 | 2.11E-01 | 9.04E-01 | -1.45E-01 | -1.11E+00 |
| ENSG000002 | C12orf75 | 2.79E-02 | 2.11E-01 | 8.81E-01 | -1.83E-01 | -1.13E+00 |
| ENSG000001 | CANX | 2.80E-02 | 2.11E-01 | 9.42E-01 | -8.61E-02 | -1.06E+00 |
| ENSG000001 | ADAMTS1 | 2.80E-02 | 2.11E-01 | 8.92E-01 | -1.65E-01 | -1.12E+00 |
| ENSG000001 | PRELID3A | 2.80E-02 | 2.12E-01 | 8.31E-01 | -2.68E-01 | -1.20E+00 |
| ENSG000001 | COX7A2L | 2.80E-02 | 2.12E-01 | 1.07E+00 | 9.82E-02 | 1.07E+00 |
| ENSG00000C | TRMT6 | 2.80E-02 | 2.12E-01 | 8.88E-01 | -1.72E-01 | -1.13E+00 |
| ENSG000001 | KBTBD6 | 2.81E-02 | 2.12E-01 | 1.09E+00 | 1.22E-01 | 1.09E+00 |
| ENSG000001 | SLC19A2 | 2.81E-02 | 2.12E-01 | 1.11E+00 | 1.56E-01 | 1.11E+00 |
| ENSG000002 | --- | 2.81E-02 | 2.12E-01 | 8.48E-01 | -2.37E-01 | -1.18E+00 |
| ENSG000001 | BCLAF3 | 2.81E-02 | 2.12E-01 | 9.02E-01 | -1.50E-01 | -1.11E+00 |
| ENSG000001 | ZNF740 | 2.82E-02 | 2.12E-01 | 1.10E+00 | 1.39E-01 | 1.10E+00 |
| ENSG000001 | IGF2 | 2.82E-02 | 2.12E-01 | 8.23E-01 | -2.82E-01 | -1.22E+00 |
| ENSG000001 | TCP1 | 2.82E-02 | 2.12E-01 | 9.38E-01 | -9.27E-02 | -1.07E+00 |
| ENSG000001 | RAPH1 | 2.83E-02 | 2.12E-01 | 9.09E-01 | -1.38E-01 | -1.10E+00 |
| ENSG00000C | CAD | 2.83E-02 | 2.13E-01 | 1.09E+00 | 1.30E-01 | 1.09E+00 |
| ENSG00000C | U2AF2 | 2.84E-02 | 2.13E-01 | 1.09E+00 | 1.27E-01 | 1.09E+00 |
| ENSG00000C | SEMA3C | 2.84E-02 | 2.13E-01 | 9.14E-01 | -1.30E-01 | -1.09E+00 |
| ENSG000001 | BAP1 | 2.84E-02 | 2.13E-01 | 1.11E+00 | 1.49E-01 | 1.11E+00 |
| ENSG000002 | --- | 2.84E-02 | 2.13E-01 | 6.49E-01 | -6.23E-01 | -1.54E+00 |

| | | | | | | | |
|---|---|---|---|---|---|---|---|
| ENSG000001 | AIMP1 | 2.84E-02 | | 2.13E-01 | 9.31E-01 | -1.04E-01 | -1.07E+00 |
| ENSG000002 | EGFR-AS1 | 2.86E-02 | ? | | 5.12E-01 | -9.65E-01 | -1.95E+00 |
| ENSG00000C | SH2D2A | 2.86E-02 | | 2.14E-01 | 7.81E-01 | -3.56E-01 | -1.28E+00 |
| ENSG000001 | DNAJB12 | 2.86E-02 | | 2.14E-01 | 9.37E-01 | -9.41E-02 | -1.07E+00 |
| ENSG000001 | EIF4ENIF1 | 2.86E-02 | | 2.14E-01 | 9.30E-01 | -1.04E-01 | -1.07E+00 |
| ENSG000001 | EIF5 | 2.86E-02 | | 2.14E-01 | 9.26E-01 | -1.11E-01 | -1.08E+00 |
| ENSG000001 | TSC22D1 | 2.87E-02 | | 2.14E-01 | 9.48E-01 | -7.69E-02 | -1.05E+00 |
| ENSG000001 | LRP11 | 2.87E-02 | | 2.14E-01 | 1.11E+00 | 1.53E-01 | 1.11E+00 |
| ENSG000001 | ARHGEF9 | 2.87E-02 | | 2.14E-01 | 1.09E+00 | 1.26E-01 | 1.09E+00 |
| ENSG00000C | THOC1 | 2.87E-02 | | 2.14E-01 | 9.00E-01 | -1.53E-01 | -1.11E+00 |
| ENSG000001 | PRKCA | 2.87E-02 | | 2.14E-01 | 8.72E-01 | -1.97E-01 | -1.15E+00 |
| ENSG000002 | --- | 2.87E-02 | | 2.14E-01 | 7.46E-01 | -4.23E-01 | -1.34E+00 |
| ENSG000001 | RRBP1 | 2.89E-02 | | 2.16E-01 | 9.44E-01 | -8.29E-02 | -1.06E+00 |
| ENSG000001 | SAP130 | 2.90E-02 | | 2.16E-01 | 9.04E-01 | -1.45E-01 | -1.11E+00 |
| ENSG000002 | AGAP6 | 2.90E-02 | | 2.16E-01 | 7.95E-01 | -3.32E-01 | -1.26E+00 |
| ENSG000002 | CD302 | 2.90E-02 | ? | | 1.79E+00 | 8.38E-01 | 1.79E+00 |
| ENSG00000C | SEC63 | 2.90E-02 | | 2.16E-01 | 9.12E-01 | -1.33E-01 | -1.10E+00 |
| ENSG000001 | BRI3 | 2.91E-02 | | 2.16E-01 | 1.15E+00 | 2.07E-01 | 1.15E+00 |
| ENSG000002 | TNFRSF6B | 2.91E-02 | | 2.16E-01 | 6.16E-01 | -6.98E-01 | -1.62E+00 |
| ENSG000001 | NECTIN2 | 2.91E-02 | | 2.16E-01 | 1.08E+00 | 1.15E-01 | 1.08E+00 |
| ENSG00000C | TNC | 2.91E-02 | | 2.16E-01 | 1.15E+00 | 1.99E-01 | 1.15E+00 |
| ENSG000001 | QTRT2 | 2.91E-02 | | 2.16E-01 | 9.10E-01 | -1.36E-01 | -1.10E+00 |
| ENSG000001 | MRPL27 | 2.92E-02 | | 2.16E-01 | 1.09E+00 | 1.26E-01 | 1.09E+00 |
| ENSG000001 | PCK2 | 2.92E-02 | | 2.16E-01 | 1.19E+00 | 2.54E-01 | 1.19E+00 |
| ENSG000001 | NPIPB4 | 2.92E-02 | | 2.16E-01 | 8.22E-01 | -2.83E-01 | -1.22E+00 |
| ENSG000001 | ICE2 | 2.92E-02 | | 2.16E-01 | 9.21E-01 | -1.19E-01 | -1.09E+00 |
| ENSG000001 | RPL30 | 2.92E-02 | | 2.16E-01 | 1.07E+00 | 9.41E-02 | 1.07E+00 |
| ENSG00000C | NUP188 | 2.93E-02 | | 2.16E-01 | 1.07E+00 | 9.45E-02 | 1.07E+00 |
| ENSG000001 | HDGFL3 | 2.93E-02 | | 2.16E-01 | 9.14E-01 | -1.30E-01 | -1.09E+00 |
| ENSG000002 | HERC2P7 | 2.93E-02 | | 2.16E-01 | 8.01E-01 | -3.20E-01 | -1.25E+00 |
| ENSG000002 | --- | 2.94E-02 | | 2.17E-01 | 1.11E+00 | 1.46E-01 | 1.11E+00 |
| ENSG000002 | LINC00963 | 2.94E-02 | | 2.17E-01 | 1.12E+00 | 1.67E-01 | 1.12E+00 |
| ENSG000002 | MYOSLID | 2.95E-02 | | 2.17E-01 | 9.12E-01 | -1.33E-01 | -1.10E+00 |
| ENSG000001 | ZNF34 | 2.95E-02 | | 2.17E-01 | 1.27E+00 | 3.40E-01 | 1.27E+00 |
| ENSG000001 | GCDH | 2.95E-02 | | 2.17E-01 | 1.16E+00 | 2.19E-01 | 1.16E+00 |
| ENSG000001 | KRT18 | 2.95E-02 | | 2.17E-01 | 8.86E-01 | -1.74E-01 | -1.13E+00 |
| ENSG000001 | GPR107 | 2.96E-02 | | 2.18E-01 | 9.45E-01 | -8.16E-02 | -1.06E+00 |
| ENSG00000C | NSUN2 | 2.96E-02 | | 2.18E-01 | 9.37E-01 | -9.43E-02 | -1.07E+00 |
| ENSG000002 | --- | 2.96E-02 | ? | | 2.70E+00 | 1.43E+00 | 2.70E+00 |
| ENSG000001 | CNDP2 | 2.96E-02 | | 2.18E-01 | 1.07E+00 | 9.87E-02 | 1.07E+00 |
| ENSG000002 | MAFB | 2.96E-02 | | 2.18E-01 | 8.30E-01 | -2.69E-01 | -1.20E+00 |
| ENSG00000C | R3HDM1 | 2.97E-02 | | 2.18E-01 | 9.09E-01 | -1.38E-01 | -1.10E+00 |
| ENSG000002 | --- | 2.97E-02 | ? | | 5.83E-01 | -7.79E-01 | -1.72E+00 |
| ENSG000002 | LINC02635 | 2.97E-02 | ? | | 1.72E+00 | 7.80E-01 | 1.72E+00 |
| ENSG000001 | FBXO2 | 2.97E-02 | | 2.18E-01 | 1.12E+00 | 1.66E-01 | 1.12E+00 |

| | | | | | | |
|---|---|---|---|---|---|---|
| ENSG000002 | --- | 2.98E-02 | ? | 3.18E-01 | -1.65E+00 | -3.14E+00 |
| ENSG00000C | RANBP9 | 2.98E-02 | 2.18E-01 | 9.32E-01 | -1.01E-01 | -1.07E+00 |
| ENSG000001 | TSPYL2 | 2.98E-02 | 2.19E-01 | 8.79E-01 | -1.86E-01 | -1.14E+00 |
| ENSG00000C | FDFT1 | 2.99E-02 | 2.19E-01 | 1.06E+00 | 8.49E-02 | 1.06E+00 |
| ENSG000001 | CEP152 | 3.00E-02 | 2.19E-01 | 8.53E-01 | -2.30E-01 | -1.17E+00 |
| ENSG000001 | KIAA1191 | 3.00E-02 | 2.19E-01 | 9.42E-01 | -8.68E-02 | -1.06E+00 |
| ENSG000001 | SFR1 | 3.00E-02 | 2.19E-01 | 8.80E-01 | -1.85E-01 | -1.14E+00 |
| ENSG000001 | SH3GL3 | 3.00E-02 | ? | 2.57E+00 | 1.36E+00 | 2.57E+00 |
| ENSG000001 | PTBP3 | 3.01E-02 | 2.20E-01 | 9.19E-01 | -1.23E-01 | -1.09E+00 |
| ENSG000002 | EBLN3P | 3.01E-02 | 2.20E-01 | 9.20E-01 | -1.20E-01 | -1.09E+00 |
| ENSG000001 | RDH13 | 3.02E-02 | 2.20E-01 | 1.15E+00 | 2.02E-01 | 1.15E+00 |
| ENSG000001 | USP37 | 3.02E-02 | 2.20E-01 | 8.82E-01 | -1.81E-01 | -1.13E+00 |
| ENSG000002 | --- | 3.02E-02 | ? | 4.02E-01 | -1.32E+00 | -2.49E+00 |
| ENSG000002 | TAPBP | 3.02E-02 | 2.21E-01 | 9.38E-01 | -9.25E-02 | -1.07E+00 |
| ENSG000001 | AP2M1 | 3.03E-02 | 2.21E-01 | 1.06E+00 | 8.60E-02 | 1.06E+00 |
| ENSG000001 | SRRM3 | 3.04E-02 | ? | 2.50E+00 | 1.32E+00 | 2.50E+00 |
| ENSG000002 | WHAMMP1 | 3.04E-02 | 2.22E-01 | 8.25E-01 | -2.78E-01 | -1.21E+00 |
| ENSG000001 | SOS2 | 3.04E-02 | 2.22E-01 | 9.08E-01 | -1.40E-01 | -1.10E+00 |
| ENSG000001 | INIP | 3.04E-02 | 2.22E-01 | 9.21E-01 | -1.19E-01 | -1.09E+00 |
| ENSG000002 | ARMCX5-GPI | 3.04E-02 | ? | 2.03E+00 | 1.02E+00 | 2.03E+00 |
| ENSG000001 | RPS9 | 3.05E-02 | 2.22E-01 | 1.14E+00 | 1.87E-01 | 1.14E+00 |
| ENSG000002 | MALAT1 | 3.05E-02 | 2.22E-01 | 7.78E-01 | -3.63E-01 | -1.29E+00 |
| ENSG000001 | FKBP2 | 3.06E-02 | 2.22E-01 | 1.10E+00 | 1.34E-01 | 1.10E+00 |
| ENSG000001 | PAFAH2 | 3.06E-02 | 2.22E-01 | 1.08E+00 | 1.11E-01 | 1.08E+00 |
| ENSG000001 | TYW5 | 3.06E-02 | 2.22E-01 | 9.16E-01 | -1.26E-01 | -1.09E+00 |
| ENSG000001 | FAR1 | 3.06E-02 | 2.23E-01 | 9.25E-01 | -1.12E-01 | -1.08E+00 |
| ENSG000001 | DNAAF3 | 3.07E-02 | ? | 6.25E-01 | -6.78E-01 | -1.60E+00 |
| ENSG000002 | CTAGE15 | 3.07E-02 | ? | 6.54E-01 | -6.12E-01 | -1.53E+00 |
| ENSG000001 | PSKH1 | 3.07E-02 | 2.23E-01 | 1.14E+00 | 1.89E-01 | 1.14E+00 |
| ENSG00000C | MAP4 | 3.08E-02 | 2.23E-01 | 9.48E-01 | -7.77E-02 | -1.06E+00 |
| ENSG000001 | ABHD2 | 3.08E-02 | 2.23E-01 | 1.07E+00 | 9.36E-02 | 1.07E+00 |
| ENSG00000C | CLEC16A | 3.08E-02 | 2.23E-01 | 9.14E-01 | -1.29E-01 | -1.09E+00 |
| ENSG000001 | CREB3L1 | 3.08E-02 | 2.23E-01 | 7.64E-01 | -3.88E-01 | -1.31E+00 |
| ENSG000001 | MOV10 | 3.08E-02 | 2.23E-01 | 1.12E+00 | 1.63E-01 | 1.12E+00 |
| ENSG000001 | RPS14 | 3.09E-02 | 2.23E-01 | 1.14E+00 | 1.86E-01 | 1.14E+00 |
| ENSG000001 | CSPG4 | 3.09E-02 | 2.23E-01 | 8.14E-01 | -2.97E-01 | -1.23E+00 |
| ENSG000001 | GPNMB | 3.09E-02 | 2.23E-01 | 1.11E+00 | 1.54E-01 | 1.11E+00 |
| ENSG000002 | --- | 3.09E-02 | 2.23E-01 | 7.75E-01 | -3.69E-01 | -1.29E+00 |
| ENSG00000C | SLC7A8 | 3.10E-02 | 2.24E-01 | 1.08E+00 | 1.12E-01 | 1.08E+00 |
| ENSG000001 | DNAJC16 | 3.11E-02 | 2.24E-01 | 1.08E+00 | 1.14E-01 | 1.08E+00 |
| ENSG000001 | B3GNT4 | 3.11E-02 | 2.24E-01 | 1.19E+00 | 2.51E-01 | 1.19E+00 |
| ENSG000001 | PTPN12 | 3.11E-02 | 2.24E-01 | 9.14E-01 | -1.30E-01 | -1.09E+00 |
| ENSG000001 | CASP3 | 3.11E-02 | 2.24E-01 | 9.38E-01 | -9.16E-02 | -1.07E+00 |
| ENSG000002 | --- | 3.12E-02 | 2.24E-01 | 8.02E-01 | -3.18E-01 | -1.25E+00 |
| ENSG00000C | M6PR | 3.12E-02 | 2.25E-01 | 1.05E+00 | 6.66E-02 | 1.05E+00 |

| | | | | | | |
|---|---|---|---|---|---|---|
| ENSG000002 | FOXD1 | 3.13E-02 | 2.25E-01 | 1.19E+00 | 2.47E-01 | 1.19E+00 |
| ENSG000002 | FBXL19-AS1 | 3.13E-02 | 2.25E-01 | 8.79E-01 | -1.86E-01 | -1.14E+00 |
| ENSG000001 | PGAM1 | 3.13E-02 | 2.25E-01 | 1.09E+00 | 1.24E-01 | 1.09E+00 |
| ENSG000002 | TATDN2P2 | 3.13E-02 | 2.25E-01 | 1.24E+00 | 3.15E-01 | 1.24E+00 |
| ENSG00000C | GALC | 3.13E-02 | 2.25E-01 | 1.10E+00 | 1.38E-01 | 1.10E+00 |
| ENSG000001 | MYO18A | 3.13E-02 | 2.25E-01 | 1.07E+00 | 9.67E-02 | 1.07E+00 |
| ENSG000001 | RNF214 | 3.13E-02 | 2.25E-01 | 9.01E-01 | -1.51E-01 | -1.11E+00 |
| ENSG000001 | RPL39 | 3.13E-02 | 2.25E-01 | 1.11E+00 | 1.48E-01 | 1.11E+00 |
| ENSG000001 | CCDC50 | 3.14E-02 | 2.25E-01 | 9.20E-01 | -1.20E-01 | -1.09E+00 |
| ENSG000001 | MYL7 | 3.14E-02 | ? | 2.07E+00 | 1.05E+00 | 2.07E+00 |
| ENSG000002 | --- | 3.14E-02 | 2.25E-01 | 1.41E+00 | 4.97E-01 | 1.41E+00 |
| ENSG000001 | PIK3CA | 3.14E-02 | 2.25E-01 | 9.14E-01 | -1.29E-01 | -1.09E+00 |
| ENSG00000C | GRHL2 | 3.15E-02 | 2.25E-01 | 1.12E+00 | 1.59E-01 | 1.12E+00 |
| ENSG000002 | --- | 3.15E-02 | 2.25E-01 | 8.82E-01 | -1.82E-01 | -1.13E+00 |
| ENSG000001 | DEXI | 3.15E-02 | 2.25E-01 | 1.13E+00 | 1.77E-01 | 1.13E+00 |
| ENSG00000C | NAV3 | 3.15E-02 | 2.25E-01 | 8.27E-01 | -2.75E-01 | -1.21E+00 |
| ENSG00000C | ADGRL1 | 3.16E-02 | 2.26E-01 | 1.19E+00 | 2.55E-01 | 1.19E+00 |
| ENSG000001 | POP7 | 3.16E-02 | 2.26E-01 | 1.12E+00 | 1.64E-01 | 1.12E+00 |
| ENSG000002 | --- | 3.17E-02 | ? | 1.93E+00 | 9.50E-01 | 1.93E+00 |
| ENSG000002 | LGR4 | 3.18E-02 | 2.27E-01 | 1.08E+00 | 1.05E-01 | 1.08E+00 |
| ENSG000001 | CAVIN2 | 3.18E-02 | 2.27E-01 | 8.84E-01 | -1.78E-01 | -1.13E+00 |
| ENSG00000C | RAD18 | 3.19E-02 | 2.28E-01 | 9.08E-01 | -1.39E-01 | -1.10E+00 |
| ENSG000002 | --- | 3.20E-02 | ? | 2.72E+00 | 1.44E+00 | 2.72E+00 |
| ENSG000001 | PSMB5 | 3.20E-02 | 2.28E-01 | 1.11E+00 | 1.45E-01 | 1.11E+00 |
| ENSG00000C | ATP12A | 3.20E-02 | ? | 4.98E-01 | -1.00E+00 | -2.01E+00 |
| ENSG000001 | PTPN13 | 3.20E-02 | 2.28E-01 | 1.09E+00 | 1.22E-01 | 1.09E+00 |
| ENSG000001 | GIT1 | 3.21E-02 | 2.28E-01 | 1.08E+00 | 1.09E-01 | 1.08E+00 |
| ENSG000002 | --- | 3.21E-02 | ? | 1.74E+00 | 7.99E-01 | 1.74E+00 |
| ENSG000001 | SERPINB9 | 3.21E-02 | 2.28E-01 | 7.81E-01 | -3.56E-01 | -1.28E+00 |
| ENSG000001 | SCN3B | 3.21E-02 | 2.28E-01 | 8.14E-01 | -2.97E-01 | -1.23E+00 |
| ENSG000001 | STAT3 | 3.21E-02 | 2.28E-01 | 9.32E-01 | -1.02E-01 | -1.07E+00 |
| ENSG000001 | ASB2 | 3.21E-02 | 2.28E-01 | 1.24E+00 | 3.07E-01 | 1.24E+00 |
| ENSG000001 | CEP83 | 3.21E-02 | 2.28E-01 | 8.70E-01 | -2.01E-01 | -1.15E+00 |
| ENSG000001 | DEGS1 | 3.22E-02 | 2.28E-01 | 1.06E+00 | 8.43E-02 | 1.06E+00 |
| ENSG000002 | --- | 3.22E-02 | 2.28E-01 | 7.82E-01 | -3.55E-01 | -1.28E+00 |
| ENSG00000C | RRP15 | 3.22E-02 | 2.28E-01 | 9.01E-01 | -1.50E-01 | -1.11E+00 |
| ENSG000001 | SLC15A2 | 3.23E-02 | 2.29E-01 | 1.20E+00 | 2.61E-01 | 1.20E+00 |
| ENSG000001 | RPL9 | 3.23E-02 | 2.29E-01 | 1.09E+00 | 1.18E-01 | 1.09E+00 |
| ENSG000001 | EVA1A | 3.23E-02 | 2.29E-01 | 9.26E-01 | -1.11E-01 | -1.08E+00 |
| ENSG000001 | MAGEH1 | 3.24E-02 | 2.29E-01 | 1.16E+00 | 2.12E-01 | 1.16E+00 |
| ENSG000001 | MCPH1 | 3.24E-02 | 2.29E-01 | 9.24E-01 | -1.14E-01 | -1.08E+00 |
| ENSG000001 | FKBP9 | 3.24E-02 | 2.29E-01 | 8.95E-01 | -1.60E-01 | -1.12E+00 |
| ENSG000001 | NF1 | 3.24E-02 | 2.29E-01 | 8.98E-01 | -1.55E-01 | -1.11E+00 |
| ENSG000001 | RWDD4 | 3.24E-02 | 2.29E-01 | 8.97E-01 | -1.57E-01 | -1.11E+00 |
| ENSG000001 | RPL21 | 3.24E-02 | 2.29E-01 | 1.07E+00 | 9.94E-02 | 1.07E+00 |

| | | | | | | |
|---|---|---|---|---|---|---|
| ENSG000000 | ERBB3 | 3.25E-02 | 2.29E-01 | 1.15E+00 | 2.04E-01 | 1.15E+00 |
| ENSG000001 | DGCR6L | 3.25E-02 | 2.29E-01 | 1.18E+00 | 2.33E-01 | 1.18E+00 |
| ENSG000000 | PTHLH | 3.25E-02 | 2.29E-01 | 8.85E-01 | -1.76E-01 | -1.13E+00 |
| ENSG000002 | ZC3H11C | 3.25E-02 | 2.29E-01 | 7.58E-01 | -4.00E-01 | -1.32E+00 |
| ENSG000001 | PHB1 | 3.25E-02 | 2.29E-01 | 1.08E+00 | 1.08E-01 | 1.08E+00 |
| ENSG000001 | CLIC4 | 3.26E-02 | 2.29E-01 | 1.08E+00 | 1.14E-01 | 1.08E+00 |
| ENSG000001 | APMAP | 3.26E-02 | 2.29E-01 | 1.06E+00 | 8.96E-02 | 1.06E+00 |
| ENSG000001 | NSUN4 | 3.26E-02 | 2.29E-01 | 9.31E-01 | -1.03E-01 | -1.07E+00 |
| ENSG000000 | NOP16 | 3.27E-02 | 2.30E-01 | 1.07E+00 | 9.11E-02 | 1.07E+00 |
| ENSG000001 | MDC1 | 3.27E-02 | 2.30E-01 | 9.23E-01 | -1.16E-01 | -1.08E+00 |
| ENSG000001 | COX6A1 | 3.28E-02 | 2.30E-01 | 1.09E+00 | 1.31E-01 | 1.09E+00 |
| ENSG000002 | STRADA | 3.28E-02 | 2.30E-01 | 1.06E+00 | 8.65E-02 | 1.06E+00 |
| ENSG000002 | ST13P4 | 3.29E-02 | ? | 4.52E-01 | -1.14E+00 | -2.21E+00 |
| ENSG000001 | FAM131A | 3.29E-02 | 2.30E-01 | 1.15E+00 | 1.98E-01 | 1.15E+00 |
| ENSG000001 | BLCAP | 3.29E-02 | 2.30E-01 | 1.07E+00 | 1.02E-01 | 1.07E+00 |
| ENSG000001 | EBP | 3.29E-02 | 2.30E-01 | 1.10E+00 | 1.36E-01 | 1.10E+00 |
| ENSG000001 | TM7SF2 | 3.29E-02 | 2.30E-01 | 1.15E+00 | 2.05E-01 | 1.15E+00 |
| ENSG000001 | ENKUR | 3.29E-02 | 2.30E-01 | 6.90E-01 | -5.36E-01 | -1.45E+00 |
| ENSG000000 | PSMA4 | 3.30E-02 | 2.31E-01 | 9.44E-01 | -8.31E-02 | -1.06E+00 |
| ENSG000001 | PCNT | 3.30E-02 | 2.31E-01 | 9.02E-01 | -1.49E-01 | -1.11E+00 |
| ENSG000001 | TMOD2 | 3.30E-02 | 2.31E-01 | 9.18E-01 | -1.23E-01 | -1.09E+00 |
| ENSG000002 | --- | 3.31E-02 | ? | 2.10E+00 | 1.07E+00 | 2.10E+00 |
| ENSG000001 | IL17RA | 3.31E-02 | 2.31E-01 | 1.06E+00 | 8.62E-02 | 1.06E+00 |
| ENSG000000 | BLVRB | 3.32E-02 | 2.32E-01 | 1.17E+00 | 2.26E-01 | 1.17E+00 |
| ENSG000001 | KNTC1 | 3.32E-02 | 2.32E-01 | 8.74E-01 | -1.94E-01 | -1.14E+00 |
| ENSG000001 | FAU | 3.33E-02 | 2.32E-01 | 1.15E+00 | 2.02E-01 | 1.15E+00 |
| ENSG000002 | --- | 3.33E-02 | ? | 5.27E-01 | -9.24E-01 | -1.90E+00 |
| ENSG000002 | C4B-AS1 | 3.33E-02 | ? | 5.15E-01 | -9.56E-01 | -1.94E+00 |
| ENSG000002 | C4A-AS1 | 3.33E-02 | ? | 5.15E-01 | -9.56E-01 | -1.94E+00 |
| ENSG000001 | RBFOX2 | 3.33E-02 | 2.32E-01 | 1.06E+00 | 8.07E-02 | 1.06E+00 |
| ENSG000001 | C18orf54 | 3.33E-02 | 2.32E-01 | 8.92E-01 | -1.65E-01 | -1.12E+00 |
| ENSG000001 | TTC6 | 3.34E-02 | ? | 5.38E-01 | -8.94E-01 | -1.86E+00 |
| ENSG000001 | PPP4R2 | 3.34E-02 | 2.32E-01 | 9.37E-01 | -9.36E-02 | -1.07E+00 |
| ENSG000002 | RPLP0P9 | 3.34E-02 | ? | 1.71E+00 | 7.74E-01 | 1.71E+00 |
| ENSG000001 | TMEM63B | 3.34E-02 | 2.33E-01 | 1.05E+00 | 6.63E-02 | 1.05E+00 |
| ENSG000002 | --- | 3.35E-02 | ? | 1.70E+00 | 7.69E-01 | 1.70E+00 |
| ENSG000002 | --- | 3.35E-02 | 2.33E-01 | 1.18E+00 | 2.42E-01 | 1.18E+00 |
| ENSG000001 | GPHA2 | 3.36E-02 | ? | 1.95E+00 | 9.64E-01 | 1.95E+00 |
| ENSG000001 | GJA1 | 3.36E-02 | 2.33E-01 | 1.08E+00 | 1.11E-01 | 1.08E+00 |
| ENSG000001 | KCTD5 | 3.36E-02 | 2.34E-01 | 1.07E+00 | 9.74E-02 | 1.07E+00 |
| ENSG000001 | AREG | 3.37E-02 | 2.34E-01 | 9.05E-01 | -1.44E-01 | -1.11E+00 |
| ENSG000001 | GOLGA2 | 3.37E-02 | 2.34E-01 | 9.18E-01 | -1.23E-01 | -1.09E+00 |
| ENSG000001 | KCTD9 | 3.38E-02 | 2.35E-01 | 9.27E-01 | -1.10E-01 | -1.08E+00 |
| ENSG000001 | CAVIN3 | 3.38E-02 | 2.35E-01 | 1.22E+00 | 2.89E-01 | 1.22E+00 |
| ENSG000001 | KATNBL1 | 3.39E-02 | 2.35E-01 | 9.36E-01 | -9.55E-02 | -1.07E+00 |

| | | | | | | |
|---|---|---|---|---|---|---|
| ENSG000001 | ZPBP2 | 3.39E-02 | ? | 1.89E+00 | 9.15E-01 | 1.89E+00 |
| ENSG00000C | KIF4A | 3.39E-02 | 2.35E-01 | 7.54E-01 | -4.07E-01 | -1.33E+00 |
| ENSG00000C | CREM | 3.40E-02 | 2.35E-01 | 8.56E-01 | -2.24E-01 | -1.17E+00 |
| ENSG000001 | TRAF7 | 3.41E-02 | 2.36E-01 | 1.10E+00 | 1.42E-01 | 1.10E+00 |
| ENSG000002 | --- | 3.41E-02 | ? | 1.81E+00 | 8.54E-01 | 1.81E+00 |
| ENSG000001 | MEX3C | 3.41E-02 | 2.36E-01 | 1.07E+00 | 9.85E-02 | 1.07E+00 |
| ENSG000001 | FBXO6 | 3.42E-02 | 2.36E-01 | 1.17E+00 | 2.21E-01 | 1.17E+00 |
| ENSG000001 | ZNF100 | 3.42E-02 | 2.36E-01 | 8.69E-01 | -2.03E-01 | -1.15E+00 |
| ENSG000002 | --- | 3.42E-02 | ? | 6.56E-01 | -6.08E-01 | -1.52E+00 |
| ENSG000001 | LRRC15 | 3.43E-02 | ? | 3.30E-01 | -1.60E+00 | -3.03E+00 |
| ENSG000001 | STAMBPL1 | 3.43E-02 | 2.37E-01 | 8.47E-01 | -2.40E-01 | -1.18E+00 |
| ENSG000001 | SLC22A23 | 3.44E-02 | 2.38E-01 | 1.08E+00 | 1.18E-01 | 1.08E+00 |
| ENSG000001 | MYORG | 3.44E-02 | 2.38E-01 | 1.08E+00 | 1.10E-01 | 1.08E+00 |
| ENSG000001 | FAM104A | 3.44E-02 | 2.38E-01 | 1.08E+00 | 1.13E-01 | 1.08E+00 |
| ENSG000002 | --- | 3.45E-02 | ? | 6.73E-01 | -5.72E-01 | -1.49E+00 |
| ENSG000001 | PON3 | 3.45E-02 | ? | 4.75E-01 | -1.07E+00 | -2.11E+00 |
| ENSG000001 | CYB561D1 | 3.45E-02 | 2.38E-01 | 8.90E-01 | -1.69E-01 | -1.12E+00 |
| ENSG000001 | URM1 | 3.46E-02 | 2.38E-01 | 1.11E+00 | 1.55E-01 | 1.11E+00 |
| ENSG000001 | GUCY1A1 | 3.46E-02 | ? | 1.69E+00 | 7.55E-01 | 1.69E+00 |
| ENSG000001 | SHPK | 3.46E-02 | 2.38E-01 | 1.16E+00 | 2.12E-01 | 1.16E+00 |
| ENSG000001 | AP5S1 | 3.46E-02 | 2.38E-01 | 1.12E+00 | 1.67E-01 | 1.12E+00 |
| ENSG000002 | LSM2 | 3.46E-02 | 2.38E-01 | 1.12E+00 | 1.67E-01 | 1.12E+00 |
| ENSG000001 | RBSN | 3.46E-02 | 2.38E-01 | 1.08E+00 | 1.06E-01 | 1.08E+00 |
| ENSG000001 | SUMF1 | 3.47E-02 | 2.38E-01 | 1.10E+00 | 1.37E-01 | 1.10E+00 |
| ENSG000001 | FKBP11 | 3.47E-02 | 2.39E-01 | 8.63E-01 | -2.13E-01 | -1.16E+00 |
| ENSG000001 | SMTN | 3.48E-02 | 2.39E-01 | 9.10E-01 | -1.36E-01 | -1.10E+00 |
| ENSG000001 | GTPBP1 | 3.48E-02 | 2.39E-01 | 1.12E+00 | 1.62E-01 | 1.12E+00 |
| ENSG000001 | QDPR | 3.49E-02 | 2.39E-01 | 1.09E+00 | 1.24E-01 | 1.09E+00 |
| ENSG000001 | TTC12 | 3.49E-02 | 2.39E-01 | 1.09E+00 | 1.28E-01 | 1.09E+00 |
| ENSG000001 | OTULIN | 3.49E-02 | 2.39E-01 | 1.07E+00 | 1.02E-01 | 1.07E+00 |
| ENSG000001 | SLC36A4 | 3.50E-02 | 2.40E-01 | 9.07E-01 | -1.41E-01 | -1.10E+00 |
| ENSG000001 | MED9 | 3.50E-02 | 2.40E-01 | 1.13E+00 | 1.79E-01 | 1.13E+00 |
| ENSG000001 | CNTNAP1 | 3.52E-02 | 2.41E-01 | 1.20E+00 | 2.60E-01 | 1.20E+00 |
| ENSG000001 | GARRE1 | 3.52E-02 | 2.41E-01 | 9.08E-01 | -1.39E-01 | -1.10E+00 |
| ENSG000001 | PRDM10 | 3.52E-02 | 2.41E-01 | 9.17E-01 | -1.26E-01 | -1.09E+00 |
| ENSG00000C | PRKCQ | 3.52E-02 | 2.41E-01 | 1.16E+00 | 2.20E-01 | 1.16E+00 |
| ENSG000002 | --- | 3.53E-02 | 2.41E-01 | 5.59E-01 | -8.40E-01 | -1.79E+00 |
| ENSG000002 | SLC48A1 | 3.53E-02 | 2.41E-01 | 8.68E-01 | -2.04E-01 | -1.15E+00 |
| ENSG000001 | FLII | 3.53E-02 | 2.41E-01 | 1.08E+00 | 1.08E-01 | 1.08E+00 |
| ENSG000002 | --- | 3.53E-02 | ? | 2.97E+00 | 1.57E+00 | 2.97E+00 |
| ENSG000002 | RPS28 | 3.54E-02 | 2.41E-01 | 1.11E+00 | 1.53E-01 | 1.11E+00 |
| ENSG000001 | PPP1R2 | 3.54E-02 | 2.42E-01 | 9.13E-01 | -1.31E-01 | -1.10E+00 |
| ENSG000001 | NPAS2 | 3.55E-02 | 2.42E-01 | 1.12E+00 | 1.58E-01 | 1.12E+00 |
| ENSG000001 | DHCR7 | 3.56E-02 | 2.43E-01 | 1.10E+00 | 1.32E-01 | 1.10E+00 |
| ENSG000001 | LRRC75B | 3.56E-02 | 2.43E-01 | 1.20E+00 | 2.68E-01 | 1.20E+00 |

| | | | | | | |
|---|---|---|---|---|---|---|
| ENSG000001 | SH3RF2 | 3.56E-02 | 2.43E-01 | 1.10E+00 | 1.31E-01 | 1.10E+00 |
| ENSG000001 | C9orf152 | 3.56E-02 | ? | 6.06E-01 | -7.23E-01 | -1.65E+00 |
| ENSG000002 | --- | 3.57E-02 | ? | 5.79E-01 | -7.88E-01 | -1.73E+00 |
| ENSG000001 | DDB1 | 3.57E-02 | 2.43E-01 | 1.06E+00 | 8.27E-02 | 1.06E+00 |
| ENSG000001 | CA5B | 3.57E-02 | 2.43E-01 | 8.59E-01 | -2.20E-01 | -1.16E+00 |
| ENSG000002 | CLDN23 | 3.57E-02 | 2.43E-01 | 1.36E+00 | 4.48E-01 | 1.36E+00 |
| ENSG000001 | ATP6V0A4 | 3.57E-02 | 2.43E-01 | 8.42E-01 | -2.48E-01 | -1.19E+00 |
| ENSG000001 | ACTN4 | 3.58E-02 | 2.43E-01 | 1.05E+00 | 6.94E-02 | 1.05E+00 |
| ENSG000001 | PCNA | 3.58E-02 | 2.43E-01 | 1.09E+00 | 1.30E-01 | 1.09E+00 |
| ENSG000001 | USO1 | 3.58E-02 | 2.43E-01 | 9.34E-01 | -9.90E-02 | -1.07E+00 |
| ENSG000001 | NECTIN4 | 3.58E-02 | 2.43E-01 | 1.20E+00 | 2.59E-01 | 1.20E+00 |
| ENSG000001 | PRKAA2 | 3.59E-02 | 2.43E-01 | 8.78E-01 | -1.88E-01 | -1.14E+00 |
| ENSG000001 | SRR | 3.59E-02 | 2.43E-01 | 1.13E+00 | 1.77E-01 | 1.13E+00 |
| ENSG000001 | CCT5 | 3.59E-02 | 2.43E-01 | 9.33E-01 | -1.01E-01 | -1.07E+00 |
| ENSG000001 | HOMER1 | 3.59E-02 | 2.43E-01 | 9.14E-01 | -1.30E-01 | -1.09E+00 |
| ENSG000001 | TPO | 3.60E-02 | ? | 5.21E-01 | -9.39E-01 | -1.92E+00 |
| ENSG000001 | EIF2B5 | 3.60E-02 | 2.44E-01 | 1.06E+00 | 9.03E-02 | 1.06E+00 |
| ENSG000001 | GEMIN6 | 3.60E-02 | 2.44E-01 | 1.09E+00 | 1.31E-01 | 1.09E+00 |
| ENSG000001 | POLR2F | 3.61E-02 | 2.44E-01 | 1.09E+00 | 1.30E-01 | 1.09E+00 |
| ENSG000001 | RBM18 | 3.61E-02 | 2.44E-01 | 9.20E-01 | -1.20E-01 | -1.09E+00 |
| ENSG000001 | DAB2 | 3.61E-02 | 2.44E-01 | 8.80E-01 | -1.85E-01 | -1.14E+00 |
| ENSG000002 | SPDYE10 | 3.62E-02 | ? | 2.54E+00 | 1.34E+00 | 2.54E+00 |
| ENSG00000C | ELAVL1 | 3.62E-02 | 2.44E-01 | 9.37E-01 | -9.45E-02 | -1.07E+00 |
| ENSG000002 | --- | 3.62E-02 | ? | 2.48E+00 | 1.31E+00 | 2.48E+00 |
| ENSG000001 | TATDN2 | 3.62E-02 | 2.44E-01 | 1.09E+00 | 1.29E-01 | 1.09E+00 |
| ENSG000001 | CXCL8 | 3.62E-02 | 2.44E-01 | 5.57E-01 | -8.45E-01 | -1.80E+00 |
| ENSG000001 | TRIM37 | 3.62E-02 | 2.44E-01 | 1.09E+00 | 1.19E-01 | 1.09E+00 |
| ENSG000001 | ZFAND4 | 3.62E-02 | 2.44E-01 | 8.11E-01 | -3.03E-01 | -1.23E+00 |
| ENSG000002 | SERBP1P1 | 3.62E-02 | ? | 6.06E-01 | -7.23E-01 | -1.65E+00 |
| ENSG000001 | THEM6 | 3.63E-02 | 2.45E-01 | 1.25E+00 | 3.19E-01 | 1.25E+00 |
| ENSG000001 | KLF13 | 3.63E-02 | 2.45E-01 | 1.09E+00 | 1.20E-01 | 1.09E+00 |
| ENSG000001 | PCDH9 | 3.63E-02 | 2.45E-01 | 8.04E-01 | -3.15E-01 | -1.24E+00 |
| ENSG000001 | RIC8A | 3.64E-02 | 2.45E-01 | 1.07E+00 | 1.02E-01 | 1.07E+00 |
| ENSG000001 | RNF32-DT | 3.64E-02 | 2.45E-01 | 1.22E+00 | 2.89E-01 | 1.22E+00 |
| ENSG000001 | SAMM50 | 3.64E-02 | 2.45E-01 | 1.09E+00 | 1.26E-01 | 1.09E+00 |
| ENSG000001 | ZNF700 | 3.65E-02 | 2.45E-01 | 8.93E-01 | -1.64E-01 | -1.12E+00 |
| ENSG000002 | --- | 3.65E-02 | ? | 4.62E-01 | -1.11E+00 | -2.16E+00 |
| ENSG000001 | NAA35 | 3.65E-02 | 2.45E-01 | 9.40E-01 | -8.99E-02 | -1.06E+00 |
| ENSG000001 | STXBP1 | 3.66E-02 | 2.45E-01 | 9.31E-01 | -1.03E-01 | -1.07E+00 |
| ENSG000001 | CLDN16 | 3.66E-02 | 2.46E-01 | 1.20E+00 | 2.65E-01 | 1.20E+00 |
| ENSG000002 | --- | 3.67E-02 | ? | 6.59E-01 | -6.01E-01 | -1.52E+00 |
| ENSG000002 | --- | 3.67E-02 | ? | 1.94E+00 | 9.57E-01 | 1.94E+00 |
| ENSG000002 | --- | 3.68E-02 | 2.46E-01 | 1.26E+00 | 3.29E-01 | 1.26E+00 |
| ENSG000001 | HSP90B1 | 3.68E-02 | 2.46E-01 | 9.13E-01 | -1.32E-01 | -1.10E+00 |
| ENSG000002 | INTS4P1 | 3.68E-02 | ? | 4.34E-01 | -1.20E+00 | -2.30E+00 |

| | | | | | | |
|---|---|---|---|---|---|---|
| ENSG000001 | NAA20 | 3.70E-02 | 2.48E-01 | 9.35E-01 | -9.70E-02 | -1.07E+00 |
| ENSG00000( | HECTD1 | 3.70E-02 | 2.48E-01 | 9.33E-01 | -9.97E-02 | -1.07E+00 |
| ENSG000001 | CIAO1 | 3.71E-02 | 2.48E-01 | 1.06E+00 | 9.01E-02 | 1.06E+00 |
| ENSG000001 | ABCD4 | 3.71E-02 | 2.48E-01 | 9.01E-01 | -1.50E-01 | -1.11E+00 |
| ENSG000001 | GPR135 | 3.71E-02 | 2.48E-01 | 8.28E-01 | -2.72E-01 | -1.21E+00 |
| ENSG00000( | ACER3 | 3.71E-02 | 2.48E-01 | 9.07E-01 | -1.40E-01 | -1.10E+00 |
| ENSG000001 | SETBP1 | 3.71E-02 | 2.48E-01 | 8.81E-01 | -1.83E-01 | -1.14E+00 |
| ENSG000002 | --- | 3.72E-02 | ? | 3.30E-01 | -1.60E+00 | -3.03E+00 |
| ENSG000001 | DDB2 | 3.72E-02 | 2.48E-01 | 1.10E+00 | 1.37E-01 | 1.10E+00 |
| ENSG000001 | RPAP1 | 3.72E-02 | 2.48E-01 | 1.10E+00 | 1.43E-01 | 1.10E+00 |
| ENSG000002 | EEF1A1P19 | 3.73E-02 | ? | 1.80E+00 | 8.50E-01 | 1.80E+00 |
| ENSG000001 | MAT1A | 3.73E-02 | ? | 2.25E+00 | 1.17E+00 | 2.25E+00 |
| ENSG000002 | UBE2CP5 | 3.73E-02 | 2.49E-01 | 8.26E-01 | -2.75E-01 | -1.21E+00 |
| ENSG000001 | ERCC6L2-AS1 | 3.74E-02 | 2.49E-01 | 1.21E+00 | 2.79E-01 | 1.21E+00 |
| ENSG000001 | HNRNPDL | 3.74E-02 | 2.49E-01 | 9.15E-01 | -1.28E-01 | -1.09E+00 |
| ENSG000001 | CALM1 | 3.74E-02 | 2.49E-01 | 9.36E-01 | -9.59E-02 | -1.07E+00 |
| ENSG000001 | NDUFC1 | 3.75E-02 | 2.49E-01 | 1.11E+00 | 1.55E-01 | 1.11E+00 |
| ENSG000002 | GGNBP2 | 3.75E-02 | 2.49E-01 | 9.37E-01 | -9.42E-02 | -1.07E+00 |
| ENSG00000( | DEPDC1 | 3.76E-02 | 2.50E-01 | 7.60E-01 | -3.96E-01 | -1.32E+00 |
| ENSG000002 | GCSHP5 | 3.76E-02 | ? | 1.75E+00 | 8.04E-01 | 1.75E+00 |
| ENSG000001 | E2F8 | 3.76E-02 | ? | 6.63E-01 | -5.92E-01 | -1.51E+00 |
| ENSG000001 | DCTN2 | 3.77E-02 | 2.50E-01 | 1.05E+00 | 7.56E-02 | 1.05E+00 |
| ENSG000001 | QKI | 3.77E-02 | 2.50E-01 | 9.18E-01 | -1.23E-01 | -1.09E+00 |
| ENSG000001 | PCIF1 | 3.77E-02 | 2.50E-01 | 1.12E+00 | 1.67E-01 | 1.12E+00 |
| ENSG000002 | CARD16 | 3.77E-02 | 2.50E-01 | 1.24E+00 | 3.10E-01 | 1.24E+00 |
| ENSG000001 | ZNF765 | 3.77E-02 | 2.50E-01 | 9.11E-01 | -1.34E-01 | -1.10E+00 |
| ENSG000001 | KEL | 3.77E-02 | ? | 2.08E+00 | 1.05E+00 | 2.08E+00 |
| ENSG000001 | RNF217 | 3.78E-02 | 2.51E-01 | 8.98E-01 | -1.54E-01 | -1.11E+00 |
| ENSG000001 | HRG | 3.78E-02 | ? | 6.77E-01 | -5.62E-01 | -1.48E+00 |
| ENSG000001 | PRMT6 | 3.78E-02 | 2.51E-01 | 1.11E+00 | 1.49E-01 | 1.11E+00 |
| ENSG000002 | BCL2L2-PABI | 3.79E-02 | 2.51E-01 | 1.18E+00 | 2.37E-01 | 1.18E+00 |
| ENSG000001 | AQP10 | 3.79E-02 | ? | 1.50E+00 | 5.85E-01 | 1.50E+00 |
| ENSG000002 | HERC2P3 | 3.80E-02 | 2.52E-01 | 8.25E-01 | -2.78E-01 | -1.21E+00 |
| ENSG000001 | ECI1 | 3.80E-02 | 2.52E-01 | 1.15E+00 | 1.98E-01 | 1.15E+00 |
| ENSG00000( | TNPO1 | 3.81E-02 | 2.52E-01 | 9.33E-01 | -9.95E-02 | -1.07E+00 |
| ENSG000001 | SLC38A4 | 3.81E-02 | 2.52E-01 | 8.80E-01 | -1.84E-01 | -1.14E+00 |
| ENSG000001 | GPAT3 | 3.81E-02 | 2.52E-01 | 7.01E-01 | -5.13E-01 | -1.43E+00 |
| ENSG000001 | VWA8 | 3.81E-02 | 2.52E-01 | 9.29E-01 | -1.06E-01 | -1.08E+00 |
| ENSG000001 | SNTB2 | 3.81E-02 | 2.52E-01 | 1.11E+00 | 1.56E-01 | 1.11E+00 |
| ENSG000001 | RGMA | 3.81E-02 | 2.52E-01 | 1.44E+00 | 5.27E-01 | 1.44E+00 |
| ENSG000001 | FNBP4 | 3.82E-02 | 2.52E-01 | 8.85E-01 | -1.77E-01 | -1.13E+00 |
| ENSG000001 | EBAG9 | 3.82E-02 | 2.52E-01 | 8.96E-01 | -1.58E-01 | -1.12E+00 |
| ENSG000002 | ZNG1F | 3.82E-02 | 2.52E-01 | 8.25E-01 | -2.78E-01 | -1.21E+00 |
| ENSG000001 | TMEM184C | 3.83E-02 | 2.52E-01 | 9.17E-01 | -1.25E-01 | -1.09E+00 |
| ENSG000001 | ANP32E | 3.83E-02 | 2.52E-01 | 9.16E-01 | -1.26E-01 | -1.09E+00 |

| | | | | | | |
|---|---|---|---|---|---|---|
| ENSG000001 | POLR2G | 3.84E-02 | 2.52E-01 | 1.14E+00 | 1.92E-01 | 1.14E+00 |
| ENSG000001 | MARCHF9 | 3.84E-02 | 2.53E-01 | 1.12E+00 | 1.62E-01 | 1.12E+00 |
| ENSG000001 | TNFRSF10A | 3.84E-02 | 2.53E-01 | 9.39E-01 | -9.11E-02 | -1.07E+00 |
| ENSG000001 | AKAP8 | 3.84E-02 | 2.53E-01 | 9.22E-01 | -1.18E-01 | -1.09E+00 |
| ENSG000002 | --- | 3.85E-02 | ? | 2.79E+00 | 1.48E+00 | 2.79E+00 |
| ENSG000002 | --- | 3.85E-02 | 2.53E-01 | 9.15E-01 | -1.28E-01 | -1.09E+00 |
| ENSG000001 | --- | 3.85E-02 | 2.53E-01 | 1.19E+00 | 2.48E-01 | 1.19E+00 |
| ENSG000001 | FRAS1 | 3.85E-02 | 2.53E-01 | 1.16E+00 | 2.11E-01 | 1.16E+00 |
| ENSG000001 | MRPL50 | 3.85E-02 | 2.53E-01 | 1.08E+00 | 1.05E-01 | 1.08E+00 |
| ENSG000001 | SNRPF | 3.87E-02 | 2.54E-01 | 1.10E+00 | 1.34E-01 | 1.10E+00 |
| ENSG000002 | RBM14-RBM | 3.87E-02 | 2.54E-01 | 6.97E-01 | -5.21E-01 | -1.43E+00 |
| ENSG000002 | --- | 3.87E-02 | ? | 1.81E+00 | 8.53E-01 | 1.81E+00 |
| ENSG000002 | --- | 3.87E-02 | ? | 4.84E-01 | -1.05E+00 | -2.06E+00 |
| ENSG000002 | --- | 3.88E-02 | ? | 1.44E+00 | 5.29E-01 | 1.44E+00 |
| ENSG000001 | SLC30A6 | 3.88E-02 | 2.54E-01 | 9.39E-01 | -9.13E-02 | -1.07E+00 |
| ENSG000002 | TPM1-AS | 3.88E-02 | 2.54E-01 | 1.40E+00 | 4.90E-01 | 1.40E+00 |
| ENSG000001 | UBE2O | 3.88E-02 | 2.54E-01 | 1.07E+00 | 1.04E-01 | 1.07E+00 |
| ENSG000001 | KCNF1 | 3.88E-02 | ? | 1.69E+00 | 7.54E-01 | 1.69E+00 |
| ENSG000001 | DHRS12 | 3.89E-02 | 2.54E-01 | 1.21E+00 | 2.71E-01 | 1.21E+00 |
| ENSG000002 | MIR222HG | 3.89E-02 | 2.54E-01 | 8.35E-01 | -2.60E-01 | -1.20E+00 |
| ENSG000001 | GALNT2 | 3.89E-02 | 2.54E-01 | 1.10E+00 | 1.32E-01 | 1.10E+00 |
| ENSG000001 | PNPLA8 | 3.89E-02 | 2.54E-01 | 9.42E-01 | -8.63E-02 | -1.06E+00 |
| ENSG000001 | COA7 | 3.89E-02 | 2.54E-01 | 1.07E+00 | 1.04E-01 | 1.07E+00 |
| ENSG000001 | ATP1A1 | 3.90E-02 | 2.54E-01 | 1.06E+00 | 7.84E-02 | 1.06E+00 |
| ENSG000001 | RHOC | 3.90E-02 | 2.54E-01 | 1.08E+00 | 1.09E-01 | 1.08E+00 |
| ENSG000001 | CA13 | 3.90E-02 | 2.55E-01 | 8.56E-01 | -2.25E-01 | -1.17E+00 |
| ENSG000001 | TSPAN13 | 3.91E-02 | 2.55E-01 | 9.43E-01 | -8.51E-02 | -1.06E+00 |
| ENSG000001 | COL1A1 | 3.91E-02 | 2.55E-01 | 8.79E-01 | -1.85E-01 | -1.14E+00 |
| ENSG000001 | VPS25 | 3.92E-02 | 2.55E-01 | 1.07E+00 | 9.30E-02 | 1.07E+00 |
| ENSG00000C | PLD1 | 3.92E-02 | 2.55E-01 | 1.14E+00 | 1.86E-01 | 1.14E+00 |
| ENSG000001 | ANPEP | 3.94E-02 | ? | 6.59E-01 | -6.02E-01 | -1.52E+00 |
| ENSG000001 | WIPF1 | 3.95E-02 | 2.57E-01 | 8.62E-01 | -2.15E-01 | -1.16E+00 |
| ENSG00000C | KIFAP3 | 3.95E-02 | 2.57E-01 | 9.25E-01 | -1.13E-01 | -1.08E+00 |
| ENSG000001 | MTSS1 | 3.95E-02 | 2.57E-01 | 8.96E-01 | -1.59E-01 | -1.12E+00 |
| ENSG000001 | DHCR24 | 3.95E-02 | 2.57E-01 | 1.11E+00 | 1.53E-01 | 1.11E+00 |
| ENSG00000C | IL11 | 3.96E-02 | 2.57E-01 | 8.31E-01 | -2.67E-01 | -1.20E+00 |
| ENSG000001 | AFF3 | 3.96E-02 | ? | 2.41E+00 | 1.27E+00 | 2.41E+00 |
| ENSG000001 | PINK1-AS | 3.97E-02 | 2.57E-01 | 1.08E+00 | 1.08E-01 | 1.08E+00 |
| ENSG000001 | IL1R1 | 3.97E-02 | 2.57E-01 | 9.02E-01 | -1.48E-01 | -1.11E+00 |
| ENSG000002 | --- | 3.97E-02 | 2.58E-01 | 1.15E+00 | 2.04E-01 | 1.15E+00 |
| ENSG000002 | --- | 3.98E-02 | ? | 4.37E-01 | -1.19E+00 | -2.29E+00 |
| ENSG000001 | VGLL1 | 3.98E-02 | 2.58E-01 | 1.38E+00 | 4.68E-01 | 1.38E+00 |
| ENSG000001 | AP1M2 | 3.98E-02 | 2.58E-01 | 1.11E+00 | 1.44E-01 | 1.11E+00 |
| ENSG000001 | SERPINH1 | 3.98E-02 | 2.58E-01 | 1.08E+00 | 1.06E-01 | 1.08E+00 |
| ENSG000001 | C9orf72 | 3.99E-02 | 2.58E-01 | 8.97E-01 | -1.57E-01 | -1.11E+00 |

| | | | | | | |
|---|---|---|---|---|---|---|
| ENSG00000C | CCDC80 | 3.99E-02 | 2.58E-01 | 8.97E-01 | -1.56E-01 | -1.11E+00 |
| ENSG000001 | NRROS | 3.99E-02 | ? | 5.85E-01 | -7.75E-01 | -1.71E+00 |
| ENSG000001 | IMPA1 | 3.99E-02 | 2.58E-01 | 9.01E-01 | -1.51E-01 | -1.11E+00 |
| ENSG000001 | CABLES1 | 4.00E-02 | 2.58E-01 | 1.36E+00 | 4.44E-01 | 1.36E+00 |
| ENSG000002 | LINC00592 | 4.00E-02 | ? | 4.33E-01 | -1.21E+00 | -2.31E+00 |
| ENSG000002 | TAP2 | 4.00E-02 | 2.58E-01 | 1.06E+00 | 8.61E-02 | 1.06E+00 |
| ENSG000002 | --- | 4.00E-02 | 2.59E-01 | 7.10E-01 | -4.94E-01 | -1.41E+00 |
| ENSG00000C | CDKL3 | 4.01E-02 | 2.59E-01 | 7.04E-01 | -5.05E-01 | -1.42E+00 |
| ENSG000001 | CTNNB1 | 4.01E-02 | 2.59E-01 | 1.06E+00 | 8.28E-02 | 1.06E+00 |
| ENSG000001 | H4C8 | 4.01E-02 | 2.59E-01 | 8.34E-01 | -2.62E-01 | -1.20E+00 |
| ENSG000001 | TRPS1 | 4.02E-02 | 2.59E-01 | 9.19E-01 | -1.22E-01 | -1.09E+00 |
| ENSG000001 | SSR2 | 4.02E-02 | 2.59E-01 | 1.06E+00 | 8.06E-02 | 1.06E+00 |
| ENSG000001 | PYROXD1 | 4.02E-02 | 2.59E-01 | 9.11E-01 | -1.34E-01 | -1.10E+00 |
| ENSG000002 | CD99P1 | 4.02E-02 | 2.59E-01 | 1.14E+00 | 1.85E-01 | 1.14E+00 |
| ENSG000002 | --- | 4.02E-02 | 2.59E-01 | 7.69E-01 | -3.79E-01 | -1.30E+00 |
| ENSG000001 | SLC37A1 | 4.03E-02 | 2.59E-01 | 9.03E-01 | -1.47E-01 | -1.11E+00 |
| ENSG000001 | ZWINT | 4.04E-02 | 2.60E-01 | 8.47E-01 | -2.39E-01 | -1.18E+00 |
| ENSG000001 | B9D1 | 4.04E-02 | 2.60E-01 | 1.17E+00 | 2.23E-01 | 1.17E+00 |
| ENSG000001 | TEX264 | 4.04E-02 | 2.60E-01 | 1.12E+00 | 1.69E-01 | 1.12E+00 |
| ENSG00000C | PPP2R5C | 4.05E-02 | 2.60E-01 | 9.44E-01 | -8.25E-02 | -1.06E+00 |
| ENSG000002 | --- | 4.05E-02 | 2.60E-01 | 8.87E-01 | -1.73E-01 | -1.13E+00 |
| ENSG000002 | FAM21EP | 4.05E-02 | ? | 1.95E+00 | 9.60E-01 | 1.95E+00 |
| ENSG000001 | RAP1A | 4.05E-02 | 2.60E-01 | 9.21E-01 | -1.18E-01 | -1.09E+00 |
| ENSG000001 | MCFD2 | 4.06E-02 | 2.60E-01 | 1.07E+00 | 9.99E-02 | 1.07E+00 |
| ENSG000001 | THY1 | 4.06E-02 | 2.60E-01 | 1.37E+00 | 4.58E-01 | 1.37E+00 |
| ENSG000001 | PDGFRA | 4.06E-02 | 2.60E-01 | 7.50E-01 | -4.15E-01 | -1.33E+00 |
| ENSG000002 | --- | 4.06E-02 | ? | 5.26E-01 | -9.28E-01 | -1.90E+00 |
| ENSG000001 | CCDC180 | 4.07E-02 | 2.61E-01 | 7.62E-01 | -3.91E-01 | -1.31E+00 |
| ENSG000002 | --- | 4.08E-02 | 2.61E-01 | 8.77E-01 | -1.89E-01 | -1.14E+00 |
| ENSG000002 | LYRM9 | 4.08E-02 | 2.61E-01 | 1.20E+00 | 2.64E-01 | 1.20E+00 |
| ENSG000001 | LMNB1 | 4.08E-02 | 2.61E-01 | 7.58E-01 | -4.00E-01 | -1.32E+00 |
| ENSG000001 | ZNF215 | 4.09E-02 | 2.61E-01 | 8.73E-01 | -1.95E-01 | -1.15E+00 |
| ENSG000001 | WDR33 | 4.09E-02 | 2.61E-01 | 1.05E+00 | 6.92E-02 | 1.05E+00 |
| ENSG000001 | MPC2 | 4.09E-02 | 2.61E-01 | 1.09E+00 | 1.30E-01 | 1.09E+00 |
| ENSG000001 | TRAPPC2L | 4.09E-02 | 2.61E-01 | 1.09E+00 | 1.18E-01 | 1.09E+00 |
| ENSG000002 | --- | 4.09E-02 | 2.61E-01 | 7.99E-01 | -3.24E-01 | -1.25E+00 |
| ENSG000001 | ARID2 | 4.10E-02 | 2.61E-01 | 9.17E-01 | -1.25E-01 | -1.09E+00 |
| ENSG000001 | NKD1 | 4.11E-02 | 2.62E-01 | 1.26E+00 | 3.39E-01 | 1.26E+00 |
| ENSG000002 | --- | 4.11E-02 | ? | 1.41E+00 | 4.91E-01 | 1.41E+00 |
| ENSG000001 | PTGER4 | 4.11E-02 | 2.62E-01 | 1.18E+00 | 2.35E-01 | 1.18E+00 |
| ENSG00000C | CRKL | 4.11E-02 | 2.62E-01 | 1.05E+00 | 6.89E-02 | 1.05E+00 |
| ENSG000001 | COX11 | 4.11E-02 | 2.62E-01 | 9.26E-01 | -1.10E-01 | -1.08E+00 |
| ENSG000001 | SYVN1 | 4.11E-02 | 2.62E-01 | 9.11E-01 | -1.35E-01 | -1.10E+00 |
| ENSG000002 | --- | 4.11E-02 | 2.62E-01 | 7.91E-01 | -3.39E-01 | -1.26E+00 |
| ENSG000001 | HPS6 | 4.12E-02 | 2.62E-01 | 1.15E+00 | 2.03E-01 | 1.15E+00 |

| | | | | | | |
|---|---|---|---|---|---|---|
| ENSG000001 | SIGMAR1 | 4.12E-02 | 2.62E-01 | 1.06E+00 | 8.73E-02 | 1.06E+00 |
| ENSG00000C | DLD | 4.12E-02 | 2.62E-01 | 9.44E-01 | -8.26E-02 | -1.06E+00 |
| ENSG000001 | DHRS2 | 4.13E-02 | 2.62E-01 | 6.63E-01 | -5.93E-01 | -1.51E+00 |
| ENSG000001 | CHP1 | 4.13E-02 | 2.62E-01 | 1.05E+00 | 7.54E-02 | 1.05E+00 |
| ENSG000001 | CCDC134 | 4.14E-02 | 2.62E-01 | 9.07E-01 | -1.41E-01 | -1.10E+00 |
| ENSG000001 | EFCAB6 | 4.14E-02 | 2.63E-01 | 1.30E+00 | 3.84E-01 | 1.30E+00 |
| ENSG000001 | ADNP2 | 4.14E-02 | 2.63E-01 | 1.07E+00 | 9.88E-02 | 1.07E+00 |
| ENSG000001 | CACYBP | 4.14E-02 | 2.63E-01 | 9.14E-01 | -1.30E-01 | -1.09E+00 |
| ENSG000001 | TTC3 | 4.15E-02 | 2.63E-01 | 9.32E-01 | -1.01E-01 | -1.07E+00 |
| ENSG000001 | CCDC117 | 4.16E-02 | 2.63E-01 | 1.09E+00 | 1.21E-01 | 1.09E+00 |
| ENSG000001 | CEP57L1 | 4.16E-02 | 2.63E-01 | 9.02E-01 | -1.49E-01 | -1.11E+00 |
| ENSG00000C | CRAT | 4.16E-02 | 2.63E-01 | 1.10E+00 | 1.34E-01 | 1.10E+00 |
| ENSG000002 | HBA1 | 4.17E-02 ? | | 6.24E-01 | -6.80E-01 | -1.60E+00 |
| ENSG00000C | CELSR3 | 4.17E-02 | 2.64E-01 | 1.26E+00 | 3.34E-01 | 1.26E+00 |
| ENSG00000C | CD38 | 4.18E-02 ? | | 1.86E+00 | 8.93E-01 | 1.86E+00 |
| ENSG000001 | HVCN1 | 4.18E-02 | 2.64E-01 | 8.11E-01 | -3.03E-01 | -1.23E+00 |
| ENSG000001 | RAB23 | 4.18E-02 | 2.64E-01 | 1.10E+00 | 1.43E-01 | 1.10E+00 |
| ENSG000001 | ANKDD1A | 4.18E-02 | 2.64E-01 | 7.69E-01 | -3.79E-01 | -1.30E+00 |
| ENSG000001 | COX6B1 | 4.18E-02 | 2.64E-01 | 1.08E+00 | 1.17E-01 | 1.08E+00 |
| ENSG000001 | COPB1 | 4.19E-02 | 2.64E-01 | 9.54E-01 | -6.78E-02 | -1.05E+00 |
| ENSG000001 | CTDSPL2 | 4.19E-02 | 2.64E-01 | 8.87E-01 | -1.74E-01 | -1.13E+00 |
| ENSG000001 | NFAT5 | 4.19E-02 | 2.64E-01 | 8.74E-01 | -1.95E-01 | -1.14E+00 |
| ENSG000001 | COQ2 | 4.19E-02 | 2.64E-01 | 1.10E+00 | 1.42E-01 | 1.10E+00 |
| ENSG000001 | CAPRIN2 | 4.20E-02 | 2.65E-01 | 8.53E-01 | -2.30E-01 | -1.17E+00 |
| ENSG00000C | PALMD | 4.21E-02 | 2.65E-01 | 1.22E+00 | 2.83E-01 | 1.22E+00 |
| ENSG000001 | STXBP5 | 4.21E-02 | 2.65E-01 | 9.35E-01 | -9.75E-02 | -1.07E+00 |
| ENSG000001 | NFE2L2 | 4.21E-02 | 2.65E-01 | 1.09E+00 | 1.20E-01 | 1.09E+00 |
| ENSG00000C | STRAP | 4.23E-02 | 2.66E-01 | 9.51E-01 | -7.24E-02 | -1.05E+00 |
| ENSG000001 | ERCC3 | 4.23E-02 | 2.66E-01 | 9.30E-01 | -1.04E-01 | -1.07E+00 |
| ENSG000002 | RIPPLY2 | 4.24E-02 ? | | 1.67E+00 | 7.43E-01 | 1.67E+00 |
| ENSG000002 | MROH6 | 4.24E-02 | 2.66E-01 | 8.65E-01 | -2.10E-01 | -1.16E+00 |
| ENSG000001 | CRLF3 | 4.24E-02 | 2.66E-01 | 9.20E-01 | -1.21E-01 | -1.09E+00 |
| ENSG000001 | TMEM86A | 4.25E-02 | 2.66E-01 | 1.26E+00 | 3.34E-01 | 1.26E+00 |
| ENSG00000C | PICALM | 4.26E-02 | 2.67E-01 | 9.34E-01 | -9.79E-02 | -1.07E+00 |
| ENSG000002 | AADACP1 | 4.27E-02 | 2.68E-01 | 7.82E-01 | -3.54E-01 | -1.28E+00 |
| ENSG00000C | RAB27B | 4.27E-02 | 2.68E-01 | 1.08E+00 | 1.17E-01 | 1.08E+00 |
| ENSG000001 | MAN2A1 | 4.27E-02 | 2.68E-01 | 1.07E+00 | 9.39E-02 | 1.07E+00 |
| ENSG000002 | --- | 4.28E-02 ? | | 2.03E+00 | 1.02E+00 | 2.03E+00 |
| ENSG000001 | S100A16 | 4.29E-02 | 2.68E-01 | 1.08E+00 | 1.12E-01 | 1.08E+00 |
| ENSG000001 | LSS | 4.29E-02 | 2.68E-01 | 1.10E+00 | 1.32E-01 | 1.10E+00 |
| ENSG000001 | ANTXR2 | 4.29E-02 | 2.68E-01 | 9.24E-01 | -1.14E-01 | -1.08E+00 |
| ENSG00000C | ATXN7L3 | 4.29E-02 | 2.68E-01 | 1.11E+00 | 1.52E-01 | 1.11E+00 |
| ENSG000001 | MAN2B1 | 4.29E-02 | 2.68E-01 | 1.08E+00 | 1.11E-01 | 1.08E+00 |
| ENSG000002 | --- | 4.29E-02 | 2.68E-01 | 1.31E+00 | 3.88E-01 | 1.31E+00 |
| ENSG000001 | POGK | 4.31E-02 | 2.69E-01 | 9.52E-01 | -7.15E-02 | -1.05E+00 |

| | | | | | | |
|---|---|---|---|---|---|---|
| ENSG000001 | FOXN1 | 4.31E-02 | ? | 1.83E+00 | 8.72E-01 | 1.83E+00 |
| ENSG000002 | TBILA | 4.31E-02 | 2.69E-01 | 8.02E-01 | -3.18E-01 | -1.25E+00 |
| ENSG000001 | E2F6 | 4.31E-02 | 2.69E-01 | 1.10E+00 | 1.36E-01 | 1.10E+00 |
| ENSG000001 | ZFP14 | 4.32E-02 | 2.69E-01 | 1.16E+00 | 2.16E-01 | 1.16E+00 |
| ENSG000001 | DSCC1 | 4.32E-02 | 2.70E-01 | 7.51E-01 | -4.13E-01 | -1.33E+00 |
| ENSG000002 | --- | 4.33E-02 | ? | 4.91E-01 | -1.02E+00 | -2.03E+00 |
| ENSG000002 | AKT1S1 | 4.33E-02 | 2.70E-01 | 9.31E-01 | -1.03E-01 | -1.07E+00 |
| ENSG000001 | TSSK6 | 4.34E-02 | ? | 1.89E+00 | 9.17E-01 | 1.89E+00 |
| ENSG000001 | PSMD11 | 4.35E-02 | 2.71E-01 | 9.38E-01 | -9.30E-02 | -1.07E+00 |
| ENSG00000C | SLC25A13 | 4.35E-02 | 2.71E-01 | 9.30E-01 | -1.05E-01 | -1.08E+00 |
| ENSG000001 | ZHX2 | 4.36E-02 | 2.71E-01 | 9.30E-01 | -1.05E-01 | -1.08E+00 |
| ENSG000001 | GRHL1 | 4.36E-02 | 2.71E-01 | 1.11E+00 | 1.55E-01 | 1.11E+00 |
| ENSG00000C | SLC25A39 | 4.36E-02 | 2.71E-01 | 1.10E+00 | 1.35E-01 | 1.10E+00 |
| ENSG000001 | LSM3 | 4.37E-02 | 2.71E-01 | 9.20E-01 | -1.21E-01 | -1.09E+00 |
| ENSG000002 | --- | 4.37E-02 | ? | 2.45E+00 | 1.29E+00 | 2.45E+00 |
| ENSG000001 | EFNA1 | 4.37E-02 | 2.72E-01 | 1.11E+00 | 1.49E-01 | 1.11E+00 |
| ENSG000002 | --- | 4.37E-02 | ? | 6.62E-01 | -5.95E-01 | -1.51E+00 |
| ENSG000001 | ZNHIT6 | 4.38E-02 | 2.72E-01 | 9.16E-01 | -1.27E-01 | -1.09E+00 |
| ENSG000001 | FBLN5 | 4.38E-02 | 2.72E-01 | 1.23E+00 | 3.01E-01 | 1.23E+00 |
| ENSG000001 | TMPO | 4.39E-02 | 2.72E-01 | 9.10E-01 | -1.36E-01 | -1.10E+00 |
| ENSG000002 | --- | 4.39E-02 | ? | 4.60E+00 | 2.20E+00 | 4.60E+00 |
| ENSG000001 | PSMC6 | 4.39E-02 | 2.72E-01 | 9.40E-01 | -8.90E-02 | -1.06E+00 |
| ENSG00000C | ANAPC5 | 4.39E-02 | 2.72E-01 | 1.06E+00 | 8.04E-02 | 1.06E+00 |
| ENSG000001 | JAGN1 | 4.40E-02 | 2.72E-01 | 1.15E+00 | 1.96E-01 | 1.15E+00 |
| ENSG000002 | --- | 4.40E-02 | 2.72E-01 | 7.94E-01 | -3.33E-01 | -1.26E+00 |
| ENSG000001 | ZFP69B | 4.40E-02 | 2.72E-01 | 8.51E-01 | -2.33E-01 | -1.18E+00 |
| ENSG000001 | RBM25 | 4.41E-02 | 2.72E-01 | 9.02E-01 | -1.49E-01 | -1.11E+00 |
| ENSG000001 | GSTK1 | 4.41E-02 | 2.72E-01 | 1.06E+00 | 8.16E-02 | 1.06E+00 |
| ENSG000001 | IPMK | 4.41E-02 | 2.72E-01 | 1.10E+00 | 1.35E-01 | 1.10E+00 |
| ENSG000001 | CYGB | 4.42E-02 | 2.73E-01 | 1.15E+00 | 1.99E-01 | 1.15E+00 |
| ENSG000001 | GRB14 | 4.42E-02 | 2.73E-01 | 1.23E+00 | 2.93E-01 | 1.23E+00 |
| ENSG000001 | PCDHB9 | 4.42E-02 | ? | 2.83E+00 | 1.50E+00 | 2.83E+00 |
| ENSG000001 | KCNAB3 | 4.43E-02 | 2.73E-01 | 7.49E-01 | -4.18E-01 | -1.34E+00 |
| ENSG000001 | FAM169A | 4.43E-02 | 2.73E-01 | 1.14E+00 | 1.86E-01 | 1.14E+00 |
| ENSG000002 | USP46-DT | 4.43E-02 | 2.73E-01 | 1.26E+00 | 3.37E-01 | 1.26E+00 |
| ENSG00000C | MED29 | 4.43E-02 | 2.73E-01 | 9.39E-01 | -9.12E-02 | -1.07E+00 |
| ENSG000002 | LINC01322 | 4.43E-02 | 2.73E-01 | 8.60E-01 | -2.17E-01 | -1.16E+00 |
| ENSG000001 | PXMP4 | 4.44E-02 | 2.73E-01 | 1.13E+00 | 1.81E-01 | 1.13E+00 |
| ENSG000001 | CCDC138 | 4.44E-02 | 2.73E-01 | 8.16E-01 | -2.93E-01 | -1.23E+00 |
| ENSG000002 | --- | 4.44E-02 | ? | 3.06E-01 | -1.71E+00 | -3.27E+00 |
| ENSG000001 | FGF11 | 4.44E-02 | 2.73E-01 | 1.10E+00 | 1.32E-01 | 1.10E+00 |
| ENSG000001 | EPHB2 | 4.44E-02 | 2.73E-01 | 9.22E-01 | -1.17E-01 | -1.08E+00 |
| ENSG000002 | HMGN1 | 4.45E-02 | 2.73E-01 | 9.33E-01 | -1.01E-01 | -1.07E+00 |
| ENSG000001 | CD274 | 4.45E-02 | 2.74E-01 | 9.01E-01 | -1.51E-01 | -1.11E+00 |
| ENSG000001 | HACD4 | 4.46E-02 | 2.74E-01 | 1.10E+00 | 1.38E-01 | 1.10E+00 |

| | | | | | | |
|---|---|---|---|---|---|---|
| ENSG000001 | IGSF11 | 4.47E-02 | ? | 4.28E-01 | -1.23E+00 | -2.34E+00 |
| ENSG000002 | DLEU2 | 4.49E-02 | 2.76E-01 | 6.91E-01 | -5.34E-01 | -1.45E+00 |
| ENSG000001 | NDUFA4 | 4.49E-02 | 2.76E-01 | 1.10E+00 | 1.39E-01 | 1.10E+00 |
| ENSG000002 | --- | 4.49E-02 | 2.76E-01 | 7.47E-01 | -4.20E-01 | -1.34E+00 |
| ENSG00000C | FBLN1 | 4.49E-02 | 2.76E-01 | 1.11E+00 | 1.49E-01 | 1.11E+00 |
| ENSG000001 | RCAN3 | 4.50E-02 | 2.76E-01 | 1.08E+00 | 1.12E-01 | 1.08E+00 |
| ENSG000001 | HJURP | 4.50E-02 | 2.76E-01 | 8.04E-01 | -3.15E-01 | -1.24E+00 |
| ENSG000001 | ANXA6 | 4.50E-02 | 2.76E-01 | 9.09E-01 | -1.38E-01 | -1.10E+00 |
| ENSG000002 | YJEFN3 | 4.51E-02 | 2.76E-01 | 8.35E-01 | -2.60E-01 | -1.20E+00 |
| ENSG000001 | COG4 | 4.51E-02 | 2.76E-01 | 1.08E+00 | 1.16E-01 | 1.08E+00 |
| ENSG000001 | C2CD2L | 4.51E-02 | 2.76E-01 | 9.06E-01 | -1.42E-01 | -1.10E+00 |
| ENSG000001 | COL12A1 | 4.52E-02 | 2.76E-01 | 8.84E-01 | -1.78E-01 | -1.13E+00 |
| ENSG000001 | DLAT | 4.52E-02 | 2.76E-01 | 9.15E-01 | -1.28E-01 | -1.09E+00 |
| ENSG00000C | PTPN21 | 4.52E-02 | 2.76E-01 | 9.17E-01 | -1.24E-01 | -1.09E+00 |
| ENSG000001 | RPL11 | 4.52E-02 | 2.76E-01 | 1.07E+00 | 9.66E-02 | 1.07E+00 |
| ENSG000001 | SLC29A2 | 4.52E-02 | 2.76E-01 | 1.16E+00 | 2.18E-01 | 1.16E+00 |
| ENSG000001 | ELL2 | 4.52E-02 | 2.76E-01 | 9.27E-01 | -1.10E-01 | -1.08E+00 |
| ENSG000001 | CXCL17 | 4.53E-02 | 2.76E-01 | 8.06E-01 | -3.11E-01 | -1.24E+00 |
| ENSG000001 | CFAP36 | 4.53E-02 | 2.76E-01 | 8.64E-01 | -2.10E-01 | -1.16E+00 |
| ENSG000001 | PIERCE1 | 4.53E-02 | ? | 1.42E+00 | 5.10E-01 | 1.42E+00 |
| ENSG000001 | IQCD | 4.53E-02 | ? | 1.70E+00 | 7.63E-01 | 1.70E+00 |
| ENSG000002 | --- | 4.53E-02 | ? | 1.92E+00 | 9.45E-01 | 1.92E+00 |
| ENSG000001 | PPFIA1 | 4.54E-02 | 2.76E-01 | 9.42E-01 | -8.60E-02 | -1.06E+00 |
| ENSG000001 | RDX | 4.54E-02 | 2.76E-01 | 1.09E+00 | 1.27E-01 | 1.09E+00 |
| ENSG000002 | H2BC6 | 4.54E-02 | 2.76E-01 | 6.95E-01 | -5.25E-01 | -1.44E+00 |
| ENSG000001 | PSMD3 | 4.54E-02 | 2.76E-01 | 1.06E+00 | 8.67E-02 | 1.06E+00 |
| ENSG000002 | RPS2P5 | 4.54E-02 | 2.76E-01 | 1.09E+00 | 1.22E-01 | 1.09E+00 |
| ENSG000002 | --- | 4.55E-02 | 2.76E-01 | 1.14E+00 | 1.84E-01 | 1.14E+00 |
| ENSG00000C | MYL6 | 4.55E-02 | 2.76E-01 | 1.03E+00 | 4.94E-02 | 1.03E+00 |
| ENSG000001 | TMEM47 | 4.55E-02 | 2.77E-01 | 8.34E-01 | -2.62E-01 | -1.20E+00 |
| ENSG000001 | SET | 4.55E-02 | 2.77E-01 | 9.52E-01 | -7.04E-02 | -1.05E+00 |
| ENSG000002 | PEX26 | 4.56E-02 | 2.77E-01 | 1.06E+00 | 8.43E-02 | 1.06E+00 |
| ENSG000001 | GAB1 | 4.56E-02 | 2.77E-01 | 9.09E-01 | -1.37E-01 | -1.10E+00 |
| ENSG000001 | APLF | 4.56E-02 | 2.77E-01 | 1.17E+00 | 2.26E-01 | 1.17E+00 |
| ENSG00000C | AGBL5 | 4.57E-02 | 2.77E-01 | 1.08E+00 | 1.06E-01 | 1.08E+00 |
| ENSG000001 | PPARGC1B | 4.57E-02 | 2.77E-01 | 1.15E+00 | 2.01E-01 | 1.15E+00 |
| ENSG000002 | --- | 4.57E-02 | ? | 3.68E-01 | -1.44E+00 | -2.71E+00 |
| ENSG000001 | CBL | 4.58E-02 | 2.78E-01 | 1.09E+00 | 1.27E-01 | 1.09E+00 |
| ENSG000001 | RAB24 | 4.59E-02 | 2.78E-01 | 1.16E+00 | 2.18E-01 | 1.16E+00 |
| ENSG000001 | ZNF318 | 4.59E-02 | 2.78E-01 | 9.29E-01 | -1.06E-01 | -1.08E+00 |
| ENSG000002 | CFAP44 | 4.59E-02 | 2.78E-01 | 8.20E-01 | -2.85E-01 | -1.22E+00 |
| ENSG000002 | COL28A1 | 4.60E-02 | ? | 6.10E-01 | -7.12E-01 | -1.64E+00 |
| ENSG000001 | SEC61B | 4.61E-02 | 2.79E-01 | 1.08E+00 | 1.14E-01 | 1.08E+00 |
| ENSG000001 | HEXIM2 | 4.62E-02 | 2.79E-01 | 1.32E+00 | 3.97E-01 | 1.32E+00 |
| ENSG000001 | KIF23 | 4.62E-02 | 2.79E-01 | 8.67E-01 | -2.06E-01 | -1.15E+00 |

| | | | | | | |
|---|---|---|---|---|---|---|
| ENSG00000C | ERCC1 | 4.62E-02 | 2.79E-01 | 9.33E-01 | -1.01E-01 | -1.07E+00 |
| ENSG000001 | UAP1 | 4.63E-02 | 2.79E-01 | 9.36E-01 | -9.61E-02 | -1.07E+00 |
| ENSG000001 | CCNA2 | 4.63E-02 | 2.79E-01 | 7.98E-01 | -3.25E-01 | -1.25E+00 |
| ENSG000001 | PSMA1 | 4.63E-02 | 2.79E-01 | 9.37E-01 | -9.36E-02 | -1.07E+00 |
| ENSG000001 | OTUD6B | 4.63E-02 | 2.79E-01 | 8.87E-01 | -1.73E-01 | -1.13E+00 |
| ENSG000002 | TMEM167B | 4.63E-02 | 2.79E-01 | 1.08E+00 | 1.15E-01 | 1.08E+00 |
| ENSG000001 | ANAPC16 | 4.63E-02 | 2.79E-01 | 1.06E+00 | 7.91E-02 | 1.06E+00 |
| ENSG000001 | RPL18A | 4.63E-02 | 2.79E-01 | 1.13E+00 | 1.82E-01 | 1.13E+00 |
| ENSG000001 | DNAH3 | 4.64E-02 | ? | 7.27E-01 | -4.59E-01 | -1.37E+00 |
| ENSG000001 | C19orf33 | 4.64E-02 | 2.79E-01 | 1.08E+00 | 1.12E-01 | 1.08E+00 |
| ENSG000002 | FAM153CP | 4.64E-02 | ? | 6.43E-01 | -6.37E-01 | -1.56E+00 |
| ENSG00000C | RRAGD | 4.64E-02 | 2.79E-01 | 1.24E+00 | 3.13E-01 | 1.24E+00 |
| ENSG000001 | FOLR3 | 4.65E-02 | 2.79E-01 | 1.15E+00 | 2.02E-01 | 1.15E+00 |
| ENSG000002 | SUDS3P1 | 4.65E-02 | ? | 1.50E+00 | 5.89E-01 | 1.50E+00 |
| ENSG000001 | BMP4 | 4.65E-02 | 2.80E-01 | 1.32E+00 | 3.95E-01 | 1.32E+00 |
| ENSG000001 | ERMARD | 4.65E-02 | 2.80E-01 | 1.10E+00 | 1.35E-01 | 1.10E+00 |
| ENSG000002 | SRXN1 | 4.66E-02 | 2.80E-01 | 9.14E-01 | -1.30E-01 | -1.09E+00 |
| ENSG000001 | GALNT11 | 4.66E-02 | 2.80E-01 | 1.07E+00 | 9.89E-02 | 1.07E+00 |
| ENSG000001 | FKBP1C | 4.66E-02 | 2.80E-01 | 1.19E+00 | 2.50E-01 | 1.19E+00 |
| ENSG000001 | SF3B5 | 4.67E-02 | 2.80E-01 | 1.12E+00 | 1.67E-01 | 1.12E+00 |
| ENSG000001 | MRPL10 | 4.67E-02 | 2.80E-01 | 1.07E+00 | 1.01E-01 | 1.07E+00 |
| ENSG000002 | GTF2I | 4.68E-02 | 2.80E-01 | 1.04E+00 | 6.27E-02 | 1.04E+00 |
| ENSG000002 | --- | 4.68E-02 | ? | 1.44E+00 | 5.31E-01 | 1.44E+00 |
| ENSG000001 | TAF1B | 4.69E-02 | 2.81E-01 | 9.24E-01 | -1.14E-01 | -1.08E+00 |
| ENSG00000C | METTL2A | 4.70E-02 | 2.81E-01 | 9.26E-01 | -1.11E-01 | -1.08E+00 |
| ENSG000001 | GRTP1 | 4.70E-02 | 2.81E-01 | 1.30E+00 | 3.81E-01 | 1.30E+00 |
| ENSG000002 | PKP4-AS1 | 4.71E-02 | ? | 5.43E-01 | -8.80E-01 | -1.84E+00 |
| ENSG000001 | RBM26 | 4.71E-02 | 2.82E-01 | 9.08E-01 | -1.39E-01 | -1.10E+00 |
| ENSG000002 | LINC01278 | 4.71E-02 | 2.82E-01 | 8.82E-01 | -1.80E-01 | -1.13E+00 |
| ENSG000001 | ZNF32 | 4.71E-02 | 2.82E-01 | 1.13E+00 | 1.79E-01 | 1.13E+00 |
| ENSG000001 | CLIP3 | 4.71E-02 | 2.82E-01 | 1.21E+00 | 2.71E-01 | 1.21E+00 |
| ENSG00000C | BCAP29 | 4.72E-02 | 2.82E-01 | 9.27E-01 | -1.09E-01 | -1.08E+00 |
| ENSG000001 | AMOT | 4.72E-02 | 2.82E-01 | 1.33E+00 | 4.13E-01 | 1.33E+00 |
| ENSG000001 | ZC3HAV1L | 4.72E-02 | 2.82E-01 | 8.85E-01 | -1.77E-01 | -1.13E+00 |
| ENSG000001 | MTMR3 | 4.73E-02 | 2.82E-01 | 1.07E+00 | 1.04E-01 | 1.07E+00 |
| ENSG000002 | --- | 4.74E-02 | 2.82E-01 | 1.30E+00 | 3.77E-01 | 1.30E+00 |
| ENSG000001 | CUEDC2 | 4.74E-02 | 2.82E-01 | 1.10E+00 | 1.44E-01 | 1.10E+00 |
| ENSG000001 | PCYT1B | 4.74E-02 | 2.83E-01 | 1.18E+00 | 2.39E-01 | 1.18E+00 |
| ENSG000001 | ZNF658B | 4.74E-02 | 2.83E-01 | 7.46E-01 | -4.22E-01 | -1.34E+00 |
| ENSG000001 | EIF3H | 4.76E-02 | 2.83E-01 | 1.04E+00 | 5.83E-02 | 1.04E+00 |
| ENSG000001 | TMPRSS13 | 4.76E-02 | 2.83E-01 | 1.29E+00 | 3.73E-01 | 1.29E+00 |
| ENSG000001 | WLS | 4.76E-02 | 2.83E-01 | 1.16E+00 | 2.11E-01 | 1.16E+00 |
| ENSG00000C | SLC23A2 | 4.77E-02 | 2.84E-01 | 1.07E+00 | 9.55E-02 | 1.07E+00 |
| ENSG000001 | FDXR | 4.77E-02 | 2.84E-01 | 1.18E+00 | 2.42E-01 | 1.18E+00 |
| ENSG000001 | WDR44 | 4.78E-02 | 2.84E-01 | 9.26E-01 | -1.11E-01 | -1.08E+00 |

| | | | | | | |
|---|---|---|---|---|---|---|
| ENSG000001 | SLC2A9 | 4.78E-02 | 2.84E-01 | 1.12E+00 | 1.68E-01 | 1.12E+00 |
| ENSG000001 | LGALS1 | 4.78E-02 | 2.84E-01 | 1.13E+00 | 1.79E-01 | 1.13E+00 |
| ENSG000002 | ANKUB1 | 4.80E-02 | 2.85E-01 | 1.16E+00 | 2.18E-01 | 1.16E+00 |
| ENSG000001 | PKDCC | 4.80E-02 | ? | 2.16E+00 | 1.11E+00 | 2.16E+00 |
| ENSG00000C | CUL3 | 4.80E-02 | 2.85E-01 | 9.32E-01 | -1.02E-01 | -1.07E+00 |
| ENSG000001 | KLK6 | 4.80E-02 | 2.85E-01 | 6.44E-01 | -6.35E-01 | -1.55E+00 |
| ENSG000002 | RPA4 | 4.81E-02 | ? | 4.11E-01 | -1.28E+00 | -2.43E+00 |
| ENSG000002 | NME1 | 4.81E-02 | 2.85E-01 | 1.08E+00 | 1.18E-01 | 1.08E+00 |
| ENSG000002 | TIRAP-AS1 | 4.81E-02 | ? | 1.98E+00 | 9.88E-01 | 1.98E+00 |
| ENSG000001 | RASSF4 | 4.82E-02 | 2.86E-01 | 8.14E-01 | -2.97E-01 | -1.23E+00 |
| ENSG00000C | RIOK2 | 4.82E-02 | 2.86E-01 | 9.33E-01 | -1.00E-01 | -1.07E+00 |
| ENSG000001 | SREBF2 | 4.83E-02 | 2.86E-01 | 1.08E+00 | 1.05E-01 | 1.08E+00 |
| ENSG000002 | --- | 4.83E-02 | ? | 4.03E-01 | -1.31E+00 | -2.48E+00 |
| ENSG000001 | TTI1 | 4.83E-02 | 2.86E-01 | 1.06E+00 | 8.94E-02 | 1.06E+00 |
| ENSG000001 | PCDH18 | 4.83E-02 | ? | 2.39E+00 | 1.25E+00 | 2.39E+00 |
| ENSG000001 | SLC44A2 | 4.83E-02 | 2.86E-01 | 1.09E+00 | 1.22E-01 | 1.09E+00 |
| ENSG000002 | --- | 4.83E-02 | 2.86E-01 | 6.72E-01 | -5.73E-01 | -1.49E+00 |
| ENSG000001 | PGAP1 | 4.83E-02 | 2.86E-01 | 1.10E+00 | 1.35E-01 | 1.10E+00 |
| ENSG000001 | ORMDL2 | 4.83E-02 | 2.86E-01 | 1.07E+00 | 1.01E-01 | 1.07E+00 |
| ENSG000001 | RND1 | 4.84E-02 | ? | 6.42E-01 | -6.40E-01 | -1.56E+00 |
| ENSG00000C | STAU2 | 4.85E-02 | 2.86E-01 | 9.24E-01 | -1.14E-01 | -1.08E+00 |
| ENSG000001 | CYP2D6 | 4.86E-02 | ? | 6.52E-01 | -6.16E-01 | -1.53E+00 |
| ENSG000001 | RPL13 | 4.86E-02 | 2.87E-01 | 1.13E+00 | 1.70E-01 | 1.13E+00 |
| ENSG000001 | GLTP | 4.86E-02 | 2.87E-01 | 9.25E-01 | -1.13E-01 | -1.08E+00 |
| ENSG000001 | GOLGA8R | 4.86E-02 | 2.87E-01 | 8.09E-01 | -3.06E-01 | -1.24E+00 |
| ENSG000001 | PRKACB | 4.86E-02 | 2.87E-01 | 9.14E-01 | -1.30E-01 | -1.09E+00 |
| ENSG00000C | CDC14A | 4.86E-02 | 2.87E-01 | 7.81E-01 | -3.56E-01 | -1.28E+00 |
| ENSG000002 | ZNF561-AS1 | 4.87E-02 | 2.87E-01 | 9.20E-01 | -1.20E-01 | -1.09E+00 |
| ENSG000001 | TMEM203 | 4.87E-02 | 2.87E-01 | 1.12E+00 | 1.65E-01 | 1.12E+00 |
| ENSG000001 | KRT74 | 4.88E-02 | ? | 1.77E+00 | 8.21E-01 | 1.77E+00 |
| ENSG000001 | TPT1 | 4.88E-02 | 2.87E-01 | 1.07E+00 | 9.11E-02 | 1.07E+00 |
| ENSG000001 | SLC16A2 | 4.89E-02 | 2.88E-01 | 1.08E+00 | 1.10E-01 | 1.08E+00 |
| ENSG000001 | SUSD1 | 4.90E-02 | 2.88E-01 | 9.19E-01 | -1.22E-01 | -1.09E+00 |
| ENSG000001 | BMPR1A | 4.90E-02 | 2.88E-01 | 9.19E-01 | -1.23E-01 | -1.09E+00 |
| ENSG000001 | HM13 | 4.90E-02 | 2.88E-01 | 9.60E-01 | -5.83E-02 | -1.04E+00 |
| ENSG00000C | DNAH9 | 4.90E-02 | ? | 1.49E+00 | 5.79E-01 | 1.49E+00 |
| ENSG00000C | RB1CC1 | 4.90E-02 | 2.88E-01 | 9.17E-01 | -1.25E-01 | -1.09E+00 |
| ENSG000002 | C11orf98 | 4.90E-02 | 2.88E-01 | 1.08E+00 | 1.11E-01 | 1.08E+00 |
| ENSG00000C | LRRC23 | 4.91E-02 | 2.88E-01 | 1.18E+00 | 2.44E-01 | 1.18E+00 |
| ENSG000001 | POLDIP3 | 4.91E-02 | 2.88E-01 | 1.08E+00 | 1.17E-01 | 1.08E+00 |
| ENSG000001 | TTC9 | 4.91E-02 | 2.88E-01 | 8.62E-01 | -2.14E-01 | -1.16E+00 |
| ENSG000001 | DUT | 4.91E-02 | 2.88E-01 | 1.09E+00 | 1.20E-01 | 1.09E+00 |
| ENSG000001 | ABCA13 | 4.91E-02 | 2.88E-01 | 8.58E-01 | -2.21E-01 | -1.17E+00 |
| ENSG000001 | ARL2BP | 4.92E-02 | 2.88E-01 | 1.06E+00 | 8.70E-02 | 1.06E+00 |
| ENSG000001 | ASIC1 | 4.92E-02 | 2.88E-01 | 1.18E+00 | 2.36E-01 | 1.18E+00 |

| | | | | | | |
|---|---|---|---|---|---|---|
| ENSG00000C | TMED2 | 4.92E-02 | 2.88E-01 | 9.42E-01 | -8.59E-02 | -1.06E+00 |
| ENSG000001 | PRCP | 4.92E-02 | 2.88E-01 | 1.09E+00 | 1.21E-01 | 1.09E+00 |
| ENSG000002 | ST3GAL1P1 | 4.92E-02 | ? | 3.08E+00 | 1.62E+00 | 3.08E+00 |
| ENSG000001 | CLASP2 | 4.93E-02 | 2.89E-01 | 9.38E-01 | -9.21E-02 | -1.07E+00 |
| ENSG000001 | FAM174A | 4.94E-02 | 2.89E-01 | 8.95E-01 | -1.60E-01 | -1.12E+00 |
| ENSG000002 | --- | 4.95E-02 | ? | 3.23E-01 | -1.63E+00 | -3.09E+00 |
| ENSG000002 | H2AJ | 4.95E-02 | 2.89E-01 | 1.17E+00 | 2.25E-01 | 1.17E+00 |
| ENSG000002 | --- | 4.96E-02 | ? | 4.61E-01 | -1.12E+00 | -2.17E+00 |
| ENSG000001 | ANKRD33B | 4.96E-02 | 2.90E-01 | 1.39E+00 | 4.78E-01 | 1.39E+00 |
| ENSG00000C | EXTL3 | 4.97E-02 | 2.90E-01 | 1.08E+00 | 1.13E-01 | 1.08E+00 |
| ENSG000001 | SLBP | 4.97E-02 | 2.90E-01 | 1.06E+00 | 7.83E-02 | 1.06E+00 |
| ENSG000001 | PPAT | 4.98E-02 | 2.90E-01 | 1.08E+00 | 1.16E-01 | 1.08E+00 |
| ENSG000001 | UACA | 4.98E-02 | 2.91E-01 | 8.93E-01 | -1.63E-01 | -1.12E+00 |
| ENSG00000C | TOLLIP | 4.98E-02 | 2.91E-01 | 1.11E+00 | 1.54E-01 | 1.11E+00 |
| ENSG000001 | CDK9 | 4.99E-02 | 2.91E-01 | 9.33E-01 | -1.01E-01 | -1.07E+00 |
| ENSG000001 | TOP3A | 4.99E-02 | 2.91E-01 | 1.08E+00 | 1.10E-01 | 1.08E+00 |
| ENSG000002 | ECI1-AS1 | 4.99E-02 | ? | 6.31E-01 | -6.65E-01 | -1.59E+00 |
| ENSG000001 | ZNF160 | 5.00E-02 | 2.91E-01 | 9.16E-01 | -1.26E-01 | -1.09E+00 |
| ENSG000001 | SH3BP5 | 5.00E-02 | 2.91E-01 | 9.10E-01 | -1.36E-01 | -1.10E+00 |
| ENSG000001 | UQCRH | 5.00E-02 | 2.91E-01 | 1.08E+00 | 1.11E-01 | 1.08E+00 |
| ENSG000001 | DCAF8 | 5.01E-02 | 2.91E-01 | 1.05E+00 | 7.69E-02 | 1.05E+00 |
| ENSG000001 | SIRT5 | 5.01E-02 | 2.91E-01 | 9.10E-01 | -1.36E-01 | -1.10E+00 |
| ENSG000002 | --- | 5.03E-02 | ? | 6.82E-01 | -5.51E-01 | -1.47E+00 |
| ENSG000002 | --- | 5.04E-02 | ? | 4.14E-01 | -1.27E+00 | -2.41E+00 |
| ENSG000001 | SDR39U1 | 5.04E-02 | 2.93E-01 | 1.13E+00 | 1.77E-01 | 1.13E+00 |
| ENSG000001 | RNF138 | 5.04E-02 | 2.93E-01 | 9.18E-01 | -1.24E-01 | -1.09E+00 |
| ENSG000001 | IL1RL1 | 5.05E-02 | 2.93E-01 | 6.87E-01 | -5.41E-01 | -1.46E+00 |
| ENSG000002 | --- | 5.05E-02 | 2.93E-01 | 8.44E-01 | -2.45E-01 | -1.18E+00 |
| ENSG000002 | LINC01269 | 5.05E-02 | ? | 6.21E-01 | -6.87E-01 | -1.61E+00 |
| ENSG000001 | RNF41 | 5.05E-02 | 2.93E-01 | 1.06E+00 | 8.86E-02 | 1.06E+00 |
| ENSG000001 | SMPD2 | 5.06E-02 | 2.93E-01 | 1.15E+00 | 2.02E-01 | 1.15E+00 |
| ENSG000001 | NIPSNAP3A | 5.06E-02 | 2.93E-01 | 1.08E+00 | 1.07E-01 | 1.08E+00 |
| ENSG000001 | DOLK | 5.06E-02 | 2.93E-01 | 1.09E+00 | 1.26E-01 | 1.09E+00 |
| ENSG000002 | UBQLN1-AS1 | 5.06E-02 | 2.93E-01 | 8.15E-01 | -2.96E-01 | -1.23E+00 |
| ENSG000002 | --- | 5.07E-02 | 2.94E-01 | 6.71E-01 | -5.75E-01 | -1.49E+00 |
| ENSG000001 | PIK3R4 | 5.08E-02 | 2.94E-01 | 1.05E+00 | 7.47E-02 | 1.05E+00 |
| ENSG000001 | NKAP | 5.09E-02 | 2.94E-01 | 9.41E-01 | -8.70E-02 | -1.06E+00 |
| ENSG00000C | ZNF532 | 5.09E-02 | 2.94E-01 | 9.30E-01 | -1.04E-01 | -1.07E+00 |
| ENSG000002 | NPHP3-ACAD | 5.09E-02 | ? | 2.30E+00 | 1.20E+00 | 2.30E+00 |
| ENSG000002 | TMEM250 | 5.09E-02 | 2.94E-01 | 1.12E+00 | 1.66E-01 | 1.12E+00 |
| ENSG000001 | RAET1E | 5.10E-02 | 2.95E-01 | 1.19E+00 | 2.53E-01 | 1.19E+00 |
| ENSG000001 | KCNE4 | 5.10E-02 | ? | 1.48E+00 | 5.65E-01 | 1.48E+00 |
| ENSG000002 | UBA52 | 5.10E-02 | 2.95E-01 | 1.09E+00 | 1.21E-01 | 1.09E+00 |
| ENSG000001 | SMC5 | 5.11E-02 | 2.95E-01 | 9.11E-01 | -1.35E-01 | -1.10E+00 |
| ENSG000001 | MSRB2 | 5.12E-02 | 2.95E-01 | 1.15E+00 | 2.05E-01 | 1.15E+00 |

| | | | | | | |
|---|---|---|---|---|---|---|
| ENSG000001 | PGAP2 | 5.12E-02 | 2.95E-01 | 1.11E+00 | 1.51E-01 | 1.11E+00 |
| ENSG000001 | SLC7A6 | 5.12E-02 | 2.95E-01 | 1.11E+00 | 1.47E-01 | 1.11E+00 |
| ENSG000001 | IL24 | 5.12E-02 | 2.95E-01 | 8.17E-01 | -2.91E-01 | -1.22E+00 |
| ENSG000001 | ETF1 | 5.13E-02 | 2.95E-01 | 9.53E-01 | -6.91E-02 | -1.05E+00 |
| ENSG000001 | TPT1-AS1 | 5.13E-02 | 2.95E-01 | 8.97E-01 | -1.56E-01 | -1.11E+00 |
| ENSG00000C | SYNJ2 | 5.13E-02 | 2.95E-01 | 9.08E-01 | -1.40E-01 | -1.10E+00 |
| ENSG000002 | ZNF254 | 5.14E-02 | 2.95E-01 | 8.86E-01 | -1.74E-01 | -1.13E+00 |
| ENSG000001 | LBR | 5.14E-02 | 2.95E-01 | 1.10E+00 | 1.37E-01 | 1.10E+00 |
| ENSG000001 | EMD | 5.14E-02 | 2.95E-01 | 1.09E+00 | 1.18E-01 | 1.09E+00 |
| ENSG000002 | CERS6-AS1 | 5.14E-02 | ? | 3.26E-01 | -1.62E+00 | -3.07E+00 |
| ENSG000001 | COG5 | 5.15E-02 | 2.96E-01 | 9.35E-01 | -9.77E-02 | -1.07E+00 |
| ENSG00000C | ARHGEF5 | 5.16E-02 | 2.96E-01 | 9.47E-01 | -7.85E-02 | -1.06E+00 |
| ENSG000001 | BROX | 5.16E-02 | 2.96E-01 | 9.47E-01 | -7.89E-02 | -1.06E+00 |
| ENSG000002 | --- | 5.16E-02 | ? | 2.03E+00 | 1.02E+00 | 2.03E+00 |
| ENSG000001 | GALNT1 | 5.17E-02 | 2.97E-01 | 1.07E+00 | 1.02E-01 | 1.07E+00 |
| ENSG000001 | ZNF524 | 5.17E-02 | 2.97E-01 | 1.23E+00 | 2.95E-01 | 1.23E+00 |
| ENSG000001 | NDUFB8 | 5.17E-02 | 2.97E-01 | 1.09E+00 | 1.28E-01 | 1.09E+00 |
| ENSG000001 | TMEM18 | 5.17E-02 | 2.97E-01 | 9.14E-01 | -1.31E-01 | -1.09E+00 |
| ENSG000001 | STAT5A | 5.18E-02 | 2.97E-01 | 1.27E+00 | 3.45E-01 | 1.27E+00 |
| ENSG000001 | ENPP5 | 5.18E-02 | 2.97E-01 | 8.55E-01 | -2.27E-01 | -1.17E+00 |
| ENSG000002 | SETBP1-DT | 5.19E-02 | ? | 1.54E+00 | 6.19E-01 | 1.54E+00 |
| ENSG000002 | --- | 5.19E-02 | ? | 5.49E-01 | -8.65E-01 | -1.82E+00 |
| ENSG000002 | INSIG1-DT | 5.19E-02 | 2.97E-01 | 1.32E+00 | 3.97E-01 | 1.32E+00 |
| ENSG000001 | ACTR1A | 5.20E-02 | 2.97E-01 | 1.07E+00 | 1.04E-01 | 1.07E+00 |
| ENSG000001 | NUDT12 | 5.20E-02 | 2.98E-01 | 1.11E+00 | 1.44E-01 | 1.11E+00 |
| ENSG000001 | PAPSS1 | 5.21E-02 | 2.98E-01 | 1.07E+00 | 9.09E-02 | 1.07E+00 |
| ENSG00000C | JADE1 | 5.21E-02 | 2.98E-01 | 9.09E-01 | -1.37E-01 | -1.10E+00 |
| ENSG000002 | ERVMER34-1 | 5.21E-02 | 2.98E-01 | 1.12E+00 | 1.58E-01 | 1.12E+00 |
| ENSG000001 | HDDC2 | 5.22E-02 | 2.98E-01 | 1.08E+00 | 1.15E-01 | 1.08E+00 |
| ENSG000001 | CITED4 | 5.22E-02 | 2.98E-01 | 1.15E+00 | 2.01E-01 | 1.15E+00 |
| ENSG00000C | FUS | 5.22E-02 | 2.98E-01 | 8.99E-01 | -1.54E-01 | -1.11E+00 |
| ENSG000001 | CTSH | 5.22E-02 | 2.98E-01 | 1.10E+00 | 1.39E-01 | 1.10E+00 |
| ENSG000001 | CRACD | 5.22E-02 | 2.98E-01 | 1.29E+00 | 3.67E-01 | 1.29E+00 |
| ENSG000001 | HIF1A | 5.22E-02 | 2.98E-01 | 9.19E-01 | -1.22E-01 | -1.09E+00 |
| ENSG000002 | CAPN10-DT | 5.23E-02 | 2.98E-01 | 1.27E+00 | 3.46E-01 | 1.27E+00 |
| ENSG000001 | PARK7 | 5.23E-02 | 2.98E-01 | 1.08E+00 | 1.07E-01 | 1.08E+00 |
| ENSG000001 | YIPF5 | 5.24E-02 | 2.98E-01 | 9.53E-01 | -6.96E-02 | -1.05E+00 |
| ENSG00000C | SLC27A5 | 5.25E-02 | 2.98E-01 | 1.34E+00 | 4.26E-01 | 1.34E+00 |
| ENSG000001 | WDR5 | 5.25E-02 | 2.98E-01 | 1.07E+00 | 1.01E-01 | 1.07E+00 |
| ENSG000001 | EMSY | 5.25E-02 | 2.98E-01 | 9.15E-01 | -1.28E-01 | -1.09E+00 |
| ENSG000001 | MAOA | 5.25E-02 | 2.98E-01 | 9.26E-01 | -1.10E-01 | -1.08E+00 |
| ENSG000001 | PAAF1 | 5.25E-02 | 2.99E-01 | 1.07E+00 | 9.46E-02 | 1.07E+00 |
| ENSG000001 | SLC6A11 | 5.26E-02 | 2.99E-01 | 1.11E+00 | 1.44E-01 | 1.11E+00 |
| ENSG000001 | SNUPN | 5.26E-02 | 2.99E-01 | 1.07E+00 | 9.71E-02 | 1.07E+00 |
| ENSG000001 | ACOX2 | 5.26E-02 | ? | 4.01E-01 | -1.32E+00 | -2.49E+00 |

| | | | | | | | |
|---|---|---|---|---|---|---|---|
| ENSG000001 | AGO2 | 5.27E-02 | | 2.99E-01 | 9.10E-01 | -1.36E-01 | -1.10E+00 |
| ENSG000001 | OSBPL1A | 5.27E-02 | | 2.99E-01 | 1.11E+00 | 1.49E-01 | 1.11E+00 |
| ENSG000001 | SEMA7A | 5.27E-02 | | 2.99E-01 | 8.98E-01 | -1.54E-01 | -1.11E+00 |
| ENSG00000C | SCML1 | 5.27E-02 | | 2.99E-01 | 1.21E+00 | 2.72E-01 | 1.21E+00 |
| ENSG000001 | HAS2 | 5.29E-02 | | 3.00E-01 | 7.48E-01 | -4.18E-01 | -1.34E+00 |
| ENSG000001 | ID3 | 5.29E-02 | | 3.00E-01 | 1.16E+00 | 2.18E-01 | 1.16E+00 |
| ENSG000001 | APEX1 | 5.29E-02 | | 3.00E-01 | 1.06E+00 | 7.80E-02 | 1.06E+00 |
| ENSG000001 | NAP1L4P1 | 5.30E-02 | ? | | 5.68E-01 | -8.15E-01 | -1.76E+00 |
| ENSG000001 | HIKESHI | 5.30E-02 | | 3.00E-01 | 1.10E+00 | 1.38E-01 | 1.10E+00 |
| ENSG000001 | TMEM251 | 5.31E-02 | | 3.00E-01 | 1.11E+00 | 1.52E-01 | 1.11E+00 |
| ENSG000001 | RRH | 5.31E-02 | ? | | 5.31E-01 | -9.12E-01 | -1.88E+00 |
| ENSG000001 | TXNRD3 | 5.31E-02 | | 3.00E-01 | 1.12E+00 | 1.66E-01 | 1.12E+00 |
| ENSG000002 | LINC01637 | 5.32E-02 | ? | | 1.63E+00 | 7.03E-01 | 1.63E+00 |
| ENSG000001 | OLAH | 5.32E-02 | ? | | 7.16E-01 | -4.83E-01 | -1.40E+00 |
| ENSG000002 | --- | 5.32E-02 | ? | | 2.19E+00 | 1.13E+00 | 2.19E+00 |
| ENSG000001 | SYTL2 | 5.34E-02 | | 3.01E-01 | 1.13E+00 | 1.80E-01 | 1.13E+00 |
| ENSG000001 | DKC1 | 5.34E-02 | | 3.01E-01 | 9.35E-01 | -9.64E-02 | -1.07E+00 |
| ENSG000001 | TCP11L1 | 5.34E-02 | | 3.01E-01 | 1.10E+00 | 1.35E-01 | 1.10E+00 |
| ENSG000001 | GOSR1 | 5.34E-02 | | 3.01E-01 | 9.50E-01 | -7.33E-02 | -1.05E+00 |
| ENSG000001 | NUMBL | 5.34E-02 | | 3.02E-01 | 9.07E-01 | -1.41E-01 | -1.10E+00 |
| ENSG000001 | C9orf85 | 5.35E-02 | | 3.02E-01 | 1.09E+00 | 1.20E-01 | 1.09E+00 |
| ENSG000002 | --- | 5.35E-02 | ? | | 4.01E-01 | -1.32E+00 | -2.49E+00 |
| ENSG000001 | AAMP | 5.35E-02 | | 3.02E-01 | 1.09E+00 | 1.30E-01 | 1.09E+00 |
| ENSG000001 | SCG5 | 5.35E-02 | | 3.02E-01 | 8.28E-01 | -2.72E-01 | -1.21E+00 |
| ENSG000001 | MAPK8 | 5.36E-02 | | 3.02E-01 | 9.18E-01 | -1.24E-01 | -1.09E+00 |
| ENSG000002 | DUSP14 | 5.37E-02 | | 3.02E-01 | 1.13E+00 | 1.81E-01 | 1.13E+00 |
| ENSG000001 | PSMC2 | 5.37E-02 | | 3.02E-01 | 9.57E-01 | -6.38E-02 | -1.05E+00 |
| ENSG000001 | RNF185 | 5.37E-02 | | 3.02E-01 | 1.10E+00 | 1.40E-01 | 1.10E+00 |
| ENSG000001 | ANKRD20A5I | 5.37E-02 | ? | | 5.91E-01 | -7.59E-01 | -1.69E+00 |
| ENSG000001 | TCTN3 | 5.37E-02 | | 3.02E-01 | 1.08E+00 | 1.11E-01 | 1.08E+00 |
| ENSG000001 | CCDC8 | 5.37E-02 | | 3.02E-01 | 1.12E+00 | 1.62E-01 | 1.12E+00 |
| ENSG000001 | CEP57 | 5.38E-02 | | 3.03E-01 | 9.18E-01 | -1.24E-01 | -1.09E+00 |
| ENSG00000C | SARS1 | 5.39E-02 | | 3.03E-01 | 1.04E+00 | 6.20E-02 | 1.04E+00 |
| ENSG000001 | ZNF277 | 5.39E-02 | | 3.03E-01 | 1.09E+00 | 1.27E-01 | 1.09E+00 |
| ENSG000002 | ANXA8L1 | 5.39E-02 | | 3.03E-01 | 1.04E+00 | 6.13E-02 | 1.04E+00 |
| ENSG000002 | NOTCH2NLB | 5.39E-02 | | 3.03E-01 | 1.19E+00 | 2.48E-01 | 1.19E+00 |
| ENSG000001 | PCSK9 | 5.39E-02 | | 3.03E-01 | 1.17E+00 | 2.27E-01 | 1.17E+00 |
| ENSG000001 | LINC02363 | 5.40E-02 | ? | | 2.03E+00 | 1.02E+00 | 2.03E+00 |
| ENSG000001 | CD96 | 5.40E-02 | ? | | 5.94E-01 | -7.52E-01 | -1.68E+00 |
| ENSG000001 | GPR89B | 5.40E-02 | | 3.03E-01 | 9.00E-01 | -1.52E-01 | -1.11E+00 |
| ENSG000001 | YLPM1 | 5.40E-02 | | 3.03E-01 | 9.25E-01 | -1.13E-01 | -1.08E+00 |
| ENSG000001 | ARAP3 | 5.41E-02 | | 3.03E-01 | 1.09E+00 | 1.26E-01 | 1.09E+00 |
| ENSG000001 | DTWD2 | 5.41E-02 | | 3.03E-01 | 9.09E-01 | -1.38E-01 | -1.10E+00 |
| ENSG000001 | DBI | 5.42E-02 | | 3.03E-01 | 1.07E+00 | 9.71E-02 | 1.07E+00 |
| ENSG000002 | C1QTNF3-AN | 5.43E-02 | ? | | 3.13E-01 | -1.68E+00 | -3.20E+00 |

| | | | | | | |
|---|---|---|---|---|---|---|
| ENSG000001 | ARF3 | 5.43E-02 | 3.04E-01 | 1.04E+00 | 6.26E-02 | 1.04E+00 |
| ENSG000001 | ZBTB5 | 5.44E-02 | 3.04E-01 | 1.08E+00 | 1.09E-01 | 1.08E+00 |
| ENSG000001 | CYRIA | 5.44E-02 | 3.04E-01 | 1.15E+00 | 2.07E-01 | 1.15E+00 |
| ENSG000001 | ZNF395 | 5.45E-02 | 3.04E-01 | 1.11E+00 | 1.52E-01 | 1.11E+00 |
| ENSG000001 | TMEM70 | 5.45E-02 | 3.04E-01 | 1.09E+00 | 1.29E-01 | 1.09E+00 |
| ENSG00000C | KEAP1 | 5.45E-02 | 3.04E-01 | 1.11E+00 | 1.50E-01 | 1.11E+00 |
| ENSG000001 | NAXE | 5.45E-02 | 3.04E-01 | 1.08E+00 | 1.12E-01 | 1.08E+00 |
| ENSG00000C | ELOA | 5.45E-02 | 3.04E-01 | 9.45E-01 | -8.14E-02 | -1.06E+00 |
| ENSG00000C | HSD17B10 | 5.46E-02 | 3.04E-01 | 1.09E+00 | 1.30E-01 | 1.09E+00 |
| ENSG000002 | ALG1L9P | 5.46E-02 | ? | 1.60E+00 | 6.79E-01 | 1.60E+00 |
| ENSG00000C | WDFY1 | 5.46E-02 | 3.05E-01 | 1.06E+00 | 8.36E-02 | 1.06E+00 |
| ENSG000001 | PDE12 | 5.46E-02 | 3.05E-01 | 9.48E-01 | -7.76E-02 | -1.06E+00 |
| ENSG000001 | COPB2 | 5.46E-02 | 3.05E-01 | 9.52E-01 | -7.03E-02 | -1.05E+00 |
| ENSG000001 | C1orf116 | 5.47E-02 | 3.05E-01 | 1.06E+00 | 8.32E-02 | 1.06E+00 |
| ENSG000002 | LYPLAL1-DT | 5.47E-02 | ? | 6.30E-01 | -6.66E-01 | -1.59E+00 |
| ENSG000002 | LINC01524 | 5.47E-02 | ? | 6.95E-01 | -5.24E-01 | -1.44E+00 |
| ENSG000001 | CCDC146 | 5.48E-02 | 3.05E-01 | 7.86E-01 | -3.47E-01 | -1.27E+00 |
| ENSG000001 | GOLGA6L9 | 5.49E-02 | 3.06E-01 | 8.26E-01 | -2.75E-01 | -1.21E+00 |
| ENSG000001 | MRPL54 | 5.50E-02 | 3.06E-01 | 1.13E+00 | 1.82E-01 | 1.13E+00 |
| ENSG000001 | SLC38A7 | 5.52E-02 | 3.07E-01 | 1.07E+00 | 1.02E-01 | 1.07E+00 |
| ENSG000002 | --- | 5.53E-02 | ? | 2.51E+00 | 1.33E+00 | 2.51E+00 |
| ENSG000002 | --- | 5.54E-02 | ? | 7.03E-01 | -5.08E-01 | -1.42E+00 |
| ENSG000001 | TNS4 | 5.54E-02 | 3.08E-01 | 8.80E-01 | -1.84E-01 | -1.14E+00 |
| ENSG000001 | PPCS | 5.54E-02 | 3.08E-01 | 1.08E+00 | 1.07E-01 | 1.08E+00 |
| ENSG000001 | PDS5A | 5.56E-02 | 3.09E-01 | 9.39E-01 | -9.12E-02 | -1.07E+00 |
| ENSG000001 | FOXQ1 | 5.56E-02 | 3.09E-01 | 7.70E-01 | -3.77E-01 | -1.30E+00 |
| ENSG000001 | RP2 | 5.56E-02 | 3.09E-01 | 1.06E+00 | 9.01E-02 | 1.06E+00 |
| ENSG000001 | TBC1D7 | 5.57E-02 | 3.09E-01 | 1.09E+00 | 1.30E-01 | 1.09E+00 |
| ENSG000002 | ATP6V0D1-D | 5.59E-02 | ? | 6.88E-01 | -5.39E-01 | -1.45E+00 |
| ENSG000001 | MKI67 | 5.59E-02 | 3.10E-01 | 7.79E-01 | -3.61E-01 | -1.28E+00 |
| ENSG000002 | PLCXD2 | 5.59E-02 | 3.10E-01 | 1.10E+00 | 1.32E-01 | 1.10E+00 |
| ENSG000001 | PHIP | 5.59E-02 | 3.10E-01 | 9.12E-01 | -1.33E-01 | -1.10E+00 |
| ENSG000002 | APPAT | 5.60E-02 | 3.10E-01 | 7.88E-01 | -3.44E-01 | -1.27E+00 |
| ENSG000001 | H1-2 | 5.61E-02 | 3.11E-01 | 1.09E+00 | 1.25E-01 | 1.09E+00 |
| ENSG000001 | WDTC1 | 5.62E-02 | 3.11E-01 | 1.07E+00 | 1.04E-01 | 1.07E+00 |
| ENSG000001 | TMEM138 | 5.62E-02 | 3.11E-01 | 1.07E+00 | 9.98E-02 | 1.07E+00 |
| ENSG000001 | ZNF584 | 5.63E-02 | 3.11E-01 | 1.12E+00 | 1.70E-01 | 1.12E+00 |
| ENSG000001 | MED21 | 5.63E-02 | 3.11E-01 | 9.11E-01 | -1.34E-01 | -1.10E+00 |
| ENSG000001 | DMTF1 | 5.64E-02 | 3.12E-01 | 8.94E-01 | -1.62E-01 | -1.12E+00 |
| ENSG000001 | DENND6A | 5.65E-02 | 3.12E-01 | 1.09E+00 | 1.27E-01 | 1.09E+00 |
| ENSG000001 | KIF14 | 5.65E-02 | 3.12E-01 | 8.13E-01 | -2.99E-01 | -1.23E+00 |
| ENSG000001 | KSR1 | 5.66E-02 | 3.12E-01 | 1.13E+00 | 1.82E-01 | 1.13E+00 |
| ENSG000001 | ZNF776 | 5.67E-02 | 3.12E-01 | 9.19E-01 | -1.22E-01 | -1.09E+00 |
| ENSG000002 | LRRC37A4P | 5.67E-02 | 3.12E-01 | 7.48E-01 | -4.18E-01 | -1.34E+00 |
| ENSG000002 | LINC00900 | 5.67E-02 | 3.12E-01 | 1.27E+00 | 3.47E-01 | 1.27E+00 |

| | | | | | | |
|---|---|---|---|---|---|---|
| ENSG000001 | EGR1 | 5.67E-02 | 3.12E-01 | 8.56E-01 | -2.25E-01 | -1.17E+00 |
| ENSG000001 | USP54 | 5.67E-02 | 3.12E-01 | 9.16E-01 | -1.26E-01 | -1.09E+00 |
| ENSG000002 | EFCAB2 | 5.68E-02 | 3.13E-01 | 8.32E-01 | -2.66E-01 | -1.20E+00 |
| ENSG000001 | TTC9C | 5.68E-02 | 3.13E-01 | 9.09E-01 | -1.37E-01 | -1.10E+00 |
| ENSG000001 | TRAPPC9 | 5.68E-02 | 3.13E-01 | 1.09E+00 | 1.25E-01 | 1.09E+00 |
| ENSG000001 | SGMS2 | 5.68E-02 | 3.13E-01 | 9.30E-01 | -1.05E-01 | -1.08E+00 |
| ENSG000001 | ZP3 | 5.69E-02 | 3.13E-01 | 1.22E+00 | 2.86E-01 | 1.22E+00 |
| ENSG000001 | SLCO2A1 | 5.70E-02 | 3.13E-01 | 8.56E-01 | -2.24E-01 | -1.17E+00 |
| ENSG000001 | BMERB1 | 5.70E-02 | 3.13E-01 | 1.09E+00 | 1.21E-01 | 1.09E+00 |
| ENSG000002 | --- | 5.71E-02 | 3.14E-01 | 7.53E-01 | -4.10E-01 | -1.33E+00 |
| ENSG000001 | BCL11B | 5.71E-02 | 3.14E-01 | 8.85E-01 | -1.76E-01 | -1.13E+00 |
| ENSG000001 | GABRR2 | 5.71E-02 | ? | 6.07E-01 | -7.20E-01 | -1.65E+00 |
| ENSG000002 | KIF9-AS1 | 5.72E-02 | 3.14E-01 | 1.13E+00 | 1.77E-01 | 1.13E+00 |
| ENSG000002 | COA6-AS1 | 5.73E-02 | ? | 1.49E+00 | 5.75E-01 | 1.49E+00 |
| ENSG000002 | --- | 5.73E-02 | 3.15E-01 | 8.36E-01 | -2.58E-01 | -1.20E+00 |
| ENSG000001 | HERC6 | 5.73E-02 | 3.15E-01 | 1.10E+00 | 1.41E-01 | 1.10E+00 |
| ENSG00000C | AK6 | 5.74E-02 | 3.15E-01 | 9.17E-01 | -1.24E-01 | -1.09E+00 |
| ENSG000001 | ASAH2 | 5.74E-02 | 3.15E-01 | 8.33E-01 | -2.64E-01 | -1.20E+00 |
| ENSG000002 | MAP7-AS1 | 5.74E-02 | ? | 1.46E+00 | 5.42E-01 | 1.46E+00 |
| ENSG000002 | LYRM4-AS1 | 5.75E-02 | 3.15E-01 | 1.11E+00 | 1.52E-01 | 1.11E+00 |
| ENSG000001 | UQCRQ | 5.75E-02 | 3.15E-01 | 1.07E+00 | 1.01E-01 | 1.07E+00 |
| ENSG000002 | --- | 5.76E-02 | 3.15E-01 | 8.33E-01 | -2.63E-01 | -1.20E+00 |
| ENSG000002 | RPL10AP6 | 5.76E-02 | ? | 1.44E+00 | 5.27E-01 | 1.44E+00 |
| ENSG000001 | WDFY3-AS2 | 5.76E-02 | 3.15E-01 | 7.58E-01 | -4.00E-01 | -1.32E+00 |
| ENSG000001 | SBF2 | 5.77E-02 | 3.16E-01 | 1.08E+00 | 1.12E-01 | 1.08E+00 |
| ENSG000001 | ITGAM | 5.77E-02 | ? | 1.46E+00 | 5.44E-01 | 1.46E+00 |
| ENSG000001 | BMPER | 5.77E-02 | 3.16E-01 | 1.22E+00 | 2.82E-01 | 1.22E+00 |
| ENSG00000C | CYB5R4 | 5.78E-02 | 3.16E-01 | 9.25E-01 | -1.13E-01 | -1.08E+00 |
| ENSG000001 | CCL2 | 5.78E-02 | ? | 5.50E-01 | -8.62E-01 | -1.82E+00 |
| ENSG000001 | FAM78B | 5.78E-02 | ? | 4.81E-01 | -1.05E+00 | -2.08E+00 |
| ENSG00000C | GTSE1 | 5.78E-02 | 3.16E-01 | 7.97E-01 | -3.28E-01 | -1.26E+00 |
| ENSG000001 | CLINT1 | 5.78E-02 | 3.16E-01 | 9.46E-01 | -8.07E-02 | -1.06E+00 |
| ENSG000001 | NT5E | 5.78E-02 | 3.16E-01 | 9.29E-01 | -1.06E-01 | -1.08E+00 |
| ENSG00000C | NIPAL3 | 5.79E-02 | 3.16E-01 | 9.28E-01 | -1.07E-01 | -1.08E+00 |
| ENSG000001 | ZNF470 | 5.80E-02 | 3.16E-01 | 8.96E-01 | -1.58E-01 | -1.12E+00 |
| ENSG000001 | VSIG10 | 5.82E-02 | 3.17E-01 | 1.11E+00 | 1.51E-01 | 1.11E+00 |
| ENSG000002 | --- | 5.82E-02 | ? | 6.07E-01 | -7.20E-01 | -1.65E+00 |
| ENSG000001 | TNFSF9 | 5.82E-02 | 3.17E-01 | 1.12E+00 | 1.69E-01 | 1.12E+00 |
| ENSG000002 | --- | 5.82E-02 | ? | 2.38E+00 | 1.25E+00 | 2.38E+00 |
| ENSG000001 | CENPU | 5.82E-02 | 3.17E-01 | 8.81E-01 | -1.83E-01 | -1.14E+00 |
| ENSG000001 | CAPS2 | 5.82E-02 | 3.17E-01 | 7.63E-01 | -3.91E-01 | -1.31E+00 |
| ENSG000002 | --- | 5.82E-02 | ? | 6.81E-01 | -5.55E-01 | -1.47E+00 |
| ENSG00000C | GLP2R | 5.83E-02 | 3.17E-01 | 6.99E-01 | -5.18E-01 | -1.43E+00 |
| ENSG000002 | --- | 5.83E-02 | ? | 5.10E-01 | -9.72E-01 | -1.96E+00 |
| ENSG000001 | TRAPPC6B | 5.84E-02 | 3.18E-01 | 9.16E-01 | -1.27E-01 | -1.09E+00 |

| | | | | | | |
|---|---|---|---|---|---|---|
| ENSG000001 | NCOA2 | 5.84E-02 | 3.18E-01 | 9.03E-01 | -1.48E-01 | -1.11E+00 |
| ENSG000001 | ATM | 5.85E-02 | 3.18E-01 | 9.11E-01 | -1.34E-01 | -1.10E+00 |
| ENSG000001 | SLC45A3 | 5.85E-02 | 3.18E-01 | 8.76E-01 | -1.91E-01 | -1.14E+00 |
| ENSG000001 | CDC26 | 5.86E-02 | 3.18E-01 | 9.10E-01 | -1.35E-01 | -1.10E+00 |
| ENSG000001 | ITGB1BP1 | 5.86E-02 | 3.18E-01 | 9.35E-01 | -9.66E-02 | -1.07E+00 |
| ENSG000001 | PLB1 | 5.87E-02 | 3.18E-01 | 1.12E+00 | 1.61E-01 | 1.12E+00 |
| ENSG000001 | AUNIP | 5.87E-02 | 3.18E-01 | 1.18E+00 | 2.34E-01 | 1.18E+00 |
| ENSG00000C | GPATCH2 | 5.87E-02 | 3.18E-01 | 9.21E-01 | -1.19E-01 | -1.09E+00 |
| ENSG000001 | PRPSAP1 | 5.87E-02 | 3.18E-01 | 1.09E+00 | 1.20E-01 | 1.09E+00 |
| ENSG000001 | ADGRL4 | 5.88E-02 | ? | 5.67E-01 | -8.19E-01 | -1.76E+00 |
| ENSG00000C | EXOC1 | 5.90E-02 | 3.20E-01 | 9.25E-01 | -1.13E-01 | -1.08E+00 |
| ENSG000001 | PRPF39 | 5.90E-02 | 3.20E-01 | 9.07E-01 | -1.41E-01 | -1.10E+00 |
| ENSG000001 | CLSTN1 | 5.90E-02 | 3.20E-01 | 1.06E+00 | 8.96E-02 | 1.06E+00 |
| ENSG00000C | COL17A1 | 5.91E-02 | 3.20E-01 | 1.06E+00 | 8.37E-02 | 1.06E+00 |
| ENSG000001 | PIP4P1 | 5.91E-02 | 3.20E-01 | 1.11E+00 | 1.50E-01 | 1.11E+00 |
| ENSG000001 | LCN12 | 5.91E-02 | ? | 4.47E-01 | -1.16E+00 | -2.24E+00 |
| ENSG000001 | KDM6B | 5.91E-02 | 3.20E-01 | 9.00E-01 | -1.51E-01 | -1.11E+00 |
| ENSG00000C | NFE2L3 | 5.91E-02 | 3.20E-01 | 1.11E+00 | 1.48E-01 | 1.11E+00 |
| ENSG000001 | PSPN | 5.92E-02 | 3.20E-01 | 1.17E+00 | 2.31E-01 | 1.17E+00 |
| ENSG000002 | STARD10 | 5.92E-02 | 3.20E-01 | 1.13E+00 | 1.76E-01 | 1.13E+00 |
| ENSG000002 | GTF2H4 | 5.92E-02 | 3.20E-01 | 1.10E+00 | 1.40E-01 | 1.10E+00 |
| ENSG000001 | ZNF843 | 5.93E-02 | ? | 5.25E-01 | -9.29E-01 | -1.90E+00 |
| ENSG000001 | CCNL1 | 5.93E-02 | 3.20E-01 | 8.82E-01 | -1.80E-01 | -1.13E+00 |
| ENSG000001 | JADE3 | 5.93E-02 | 3.20E-01 | 1.07E+00 | 9.89E-02 | 1.07E+00 |
| ENSG000001 | XPC | 5.94E-02 | 3.20E-01 | 1.06E+00 | 8.07E-02 | 1.06E+00 |
| ENSG000001 | ASXL3 | 5.94E-02 | ? | 4.50E-01 | -1.15E+00 | -2.22E+00 |
| ENSG000002 | LINC01704 | 5.94E-02 | 3.20E-01 | 1.14E+00 | 1.92E-01 | 1.14E+00 |
| ENSG000001 | RHOD | 5.94E-02 | 3.20E-01 | 1.15E+00 | 1.96E-01 | 1.15E+00 |
| ENSG000002 | CAPN1-AS1 | 5.95E-02 | 3.21E-01 | 8.60E-01 | -2.17E-01 | -1.16E+00 |
| ENSG000002 | --- | 5.95E-02 | 3.21E-01 | 1.37E+00 | 4.52E-01 | 1.37E+00 |
| ENSG000001 | SLIT3 | 5.95E-02 | 3.21E-01 | 1.13E+00 | 1.71E-01 | 1.13E+00 |
| ENSG000001 | RBAK | 5.95E-02 | 3.21E-01 | 9.17E-01 | -1.24E-01 | -1.09E+00 |
| ENSG000001 | CCDC3 | 5.95E-02 | ? | 1.65E+00 | 7.18E-01 | 1.65E+00 |
| ENSG000001 | CCDC43 | 5.96E-02 | 3.21E-01 | 9.39E-01 | -9.05E-02 | -1.06E+00 |
| ENSG000001 | CCDC126 | 5.96E-02 | 3.21E-01 | 8.86E-01 | -1.74E-01 | -1.13E+00 |
| ENSG000001 | SOX9 | 5.96E-02 | 3.21E-01 | 9.07E-01 | -1.40E-01 | -1.10E+00 |
| ENSG000002 | DGCR5 | 5.96E-02 | 3.21E-01 | 1.24E+00 | 3.14E-01 | 1.24E+00 |
| ENSG000001 | SPTAN1 | 5.97E-02 | 3.21E-01 | 9.59E-01 | -6.10E-02 | -1.04E+00 |
| ENSG000001 | TMCC2 | 5.97E-02 | 3.21E-01 | 1.20E+00 | 2.65E-01 | 1.20E+00 |
| ENSG000001 | CD36 | 5.98E-02 | ? | 1.51E+00 | 5.92E-01 | 1.51E+00 |
| ENSG000001 | MMAB | 5.98E-02 | 3.21E-01 | 1.09E+00 | 1.30E-01 | 1.09E+00 |
| ENSG000002 | UBE2V1P2 | 5.98E-02 | ? | 2.10E+00 | 1.07E+00 | 2.10E+00 |
| ENSG000001 | TMEM255A | 5.99E-02 | ? | 1.96E+00 | 9.67E-01 | 1.96E+00 |
| ENSG000001 | ZNF566 | 5.99E-02 | 3.22E-01 | 8.86E-01 | -1.75E-01 | -1.13E+00 |
| ENSG000002 | MVP-DT | 6.00E-02 | 3.22E-01 | 7.81E-01 | -3.56E-01 | -1.28E+00 |

| | | | | | | |
|---|---|---|---|---|---|---|
| ENSG000001 | WNK3 | 6.00E-02 | 3.22E-01 | 8.43E-01 | -2.46E-01 | -1.19E+00 |
| ENSG000001 | RPS2P46 | 6.00E-02 | 3.22E-01 | 1.13E+00 | 1.74E-01 | 1.13E+00 |
| ENSG000001 | KLHDC8B | 6.01E-02 | 3.22E-01 | 1.17E+00 | 2.30E-01 | 1.17E+00 |
| ENSG000001 | PPT1 | 6.01E-02 | 3.22E-01 | 1.05E+00 | 7.29E-02 | 1.05E+00 |
| ENSG000001 | CHD1 | 6.01E-02 | 3.22E-01 | 9.17E-01 | -1.25E-01 | -1.09E+00 |
| ENSG000001 | C11orf52 | 6.01E-02 | 3.22E-01 | 1.28E+00 | 3.59E-01 | 1.28E+00 |
| ENSG000002 | MRPS21 | 6.01E-02 | 3.22E-01 | 1.06E+00 | 8.47E-02 | 1.06E+00 |
| ENSG000001 | HBA2 | 6.02E-02 | ? | 5.35E-01 | -9.04E-01 | -1.87E+00 |
| ENSG000002 | --- | 6.02E-02 | 3.22E-01 | 8.20E-01 | -2.87E-01 | -1.22E+00 |
| ENSG000001 | TUT1 | 6.03E-02 | 3.22E-01 | 1.14E+00 | 1.85E-01 | 1.14E+00 |
| ENSG000001 | ZNF529 | 6.03E-02 | 3.22E-01 | 9.11E-01 | -1.34E-01 | -1.10E+00 |
| ENSG000001 | NEK4 | 6.03E-02 | 3.22E-01 | 9.23E-01 | -1.16E-01 | -1.08E+00 |
| ENSG000002 | RPL6P27 | 6.03E-02 | 3.22E-01 | 1.12E+00 | 1.60E-01 | 1.12E+00 |
| ENSG000001 | KRT15 | 6.04E-02 | 3.23E-01 | 1.25E+00 | 3.24E-01 | 1.25E+00 |
| ENSG000002 | --- | 6.04E-02 | ? | 2.33E-01 | -2.10E+00 | -4.30E+00 |
| ENSG000002 | TMEM178B | 6.05E-02 | ? | 2.11E+00 | 1.08E+00 | 2.11E+00 |
| ENSG00000C | AGA | 6.05E-02 | 3.23E-01 | 9.08E-01 | -1.38E-01 | -1.10E+00 |
| ENSG000001 | SNRK | 6.05E-02 | 3.23E-01 | 9.46E-01 | -8.07E-02 | -1.06E+00 |
| ENSG000001 | SNAPC2 | 6.05E-02 | 3.23E-01 | 9.18E-01 | -1.24E-01 | -1.09E+00 |
| ENSG000002 | SOD2 | 6.05E-02 | 3.23E-01 | 9.22E-01 | -1.17E-01 | -1.08E+00 |
| ENSG000001 | MTFR1L | 6.05E-02 | 3.23E-01 | 1.09E+00 | 1.24E-01 | 1.09E+00 |
| ENSG000001 | BOLA3 | 6.06E-02 | 3.23E-01 | 1.08E+00 | 1.14E-01 | 1.08E+00 |
| ENSG000002 | ZNF625-ZNF | 6.06E-02 | 3.23E-01 | 8.25E-01 | -2.77E-01 | -1.21E+00 |
| ENSG000001 | OXSM | 6.06E-02 | 3.23E-01 | 1.11E+00 | 1.56E-01 | 1.11E+00 |
| ENSG000001 | PLEKHF1 | 6.07E-02 | 3.23E-01 | 1.17E+00 | 2.23E-01 | 1.17E+00 |
| ENSG000001 | AMIGO2 | 6.08E-02 | 3.23E-01 | 1.06E+00 | 8.57E-02 | 1.06E+00 |
| ENSG00000C | ZNF264 | 6.08E-02 | 3.23E-01 | 9.18E-01 | -1.24E-01 | -1.09E+00 |
| ENSG000001 | ANK2 | 6.08E-02 | 3.23E-01 | 1.20E+00 | 2.59E-01 | 1.20E+00 |
| ENSG000001 | CCNG2 | 6.08E-02 | 3.23E-01 | 9.36E-01 | -9.56E-02 | -1.07E+00 |
| ENSG000002 | --- | 6.09E-02 | ? | 4.08E-01 | -1.29E+00 | -2.45E+00 |
| ENSG000001 | FKBP1B | 6.09E-02 | 3.24E-01 | 1.12E+00 | 1.63E-01 | 1.12E+00 |
| ENSG000002 | LINC02550 | 6.09E-02 | ? | 3.61E-01 | -1.47E+00 | -2.77E+00 |
| ENSG000001 | KDM2A | 6.10E-02 | 3.24E-01 | 9.57E-01 | -6.32E-02 | -1.04E+00 |
| ENSG00000C | FAT2 | 6.10E-02 | 3.24E-01 | 1.12E+00 | 1.68E-01 | 1.12E+00 |
| ENSG000001 | LINC01559 | 6.11E-02 | 3.24E-01 | 8.31E-01 | -2.67E-01 | -1.20E+00 |
| ENSG000001 | CCL25 | 6.11E-02 | ? | 2.68E+00 | 1.42E+00 | 2.68E+00 |
| ENSG00000C | SLC25A3 | 6.11E-02 | 3.24E-01 | 1.07E+00 | 1.04E-01 | 1.07E+00 |
| ENSG000001 | RAD54B | 6.11E-02 | 3.24E-01 | 8.67E-01 | -2.06E-01 | -1.15E+00 |
| ENSG000001 | GJB3 | 6.12E-02 | 3.24E-01 | 1.06E+00 | 8.96E-02 | 1.06E+00 |
| ENSG000002 | SEMA6A-AS2 | 6.12E-02 | ? | 2.04E+00 | 1.03E+00 | 2.04E+00 |
| ENSG000002 | DNAJC27-AS | 6.12E-02 | ? | 1.74E+00 | 8.01E-01 | 1.74E+00 |
| ENSG000001 | MYOF | 6.12E-02 | 3.24E-01 | 1.06E+00 | 8.53E-02 | 1.06E+00 |
| ENSG000001 | PHAF1 | 6.12E-02 | 3.24E-01 | 1.06E+00 | 8.56E-02 | 1.06E+00 |
| ENSG000001 | NDUFAF3 | 6.12E-02 | 3.24E-01 | 1.15E+00 | 1.99E-01 | 1.15E+00 |
| ENSG00000C | SNAP91 | 6.12E-02 | ? | 1.47E+00 | 5.52E-01 | 1.47E+00 |

| | | | | | | |
|---|---|---|---|---|---|---|
| ENSG000001 | PPP2R5D | 6.13E-02 | 3.24E-01 | 9.38E-01 | -9.18E-02 | -1.07E+00 |
| ENSG000001 | BRIP1 | 6.13E-02 | 3.24E-01 | 1.19E+00 | 2.55E-01 | 1.19E+00 |
| ENSG000002 | --- | 6.13E-02 | 3.24E-01 | 8.70E-01 | -2.01E-01 | -1.15E+00 |
| ENSG000001 | RPS6KL1 | 6.13E-02 | 3.24E-01 | 1.21E+00 | 2.81E-01 | 1.21E+00 |
| ENSG000001 | TEDC2 | 6.14E-02 | 3.24E-01 | 1.34E+00 | 4.20E-01 | 1.34E+00 |
| ENSG00000C | GNA15 | 6.15E-02 | 3.25E-01 | 1.07E+00 | 9.85E-02 | 1.07E+00 |
| ENSG000002 | --- | 6.15E-02 | ? | 1.55E+00 | 6.37E-01 | 1.55E+00 |
| ENSG000001 | INPP5D | 6.15E-02 | 3.25E-01 | 1.09E+00 | 1.22E-01 | 1.09E+00 |
| ENSG000002 | CATSPER2P1 | 6.15E-02 | 3.25E-01 | 1.28E+00 | 3.60E-01 | 1.28E+00 |
| ENSG000001 | HPCAL1 | 6.16E-02 | 3.25E-01 | 8.91E-01 | -1.67E-01 | -1.12E+00 |
| ENSG000001 | ZNF583 | 6.17E-02 | 3.25E-01 | 8.91E-01 | -1.66E-01 | -1.12E+00 |
| ENSG000001 | GOLGA7 | 6.17E-02 | 3.26E-01 | 9.30E-01 | -1.05E-01 | -1.08E+00 |
| ENSG000001 | ZNF462 | 6.18E-02 | 3.26E-01 | 9.13E-01 | -1.32E-01 | -1.10E+00 |
| ENSG000001 | ABHD12 | 6.18E-02 | 3.26E-01 | 1.13E+00 | 1.74E-01 | 1.13E+00 |
| ENSG000001 | CWH43 | 6.18E-02 | 3.26E-01 | 1.33E+00 | 4.11E-01 | 1.33E+00 |
| ENSG000001 | AKNA | 6.18E-02 | 3.26E-01 | 1.12E+00 | 1.63E-01 | 1.12E+00 |
| ENSG000001 | EPAS1 | 6.19E-02 | 3.26E-01 | 8.91E-01 | -1.66E-01 | -1.12E+00 |
| ENSG000001 | KPNA4 | 6.19E-02 | 3.26E-01 | 9.31E-01 | -1.04E-01 | -1.07E+00 |
| ENSG000001 | CIPC | 6.19E-02 | 3.26E-01 | 1.07E+00 | 1.01E-01 | 1.07E+00 |
| ENSG00000C | ITPR3 | 6.20E-02 | 3.26E-01 | 9.50E-01 | -7.41E-02 | -1.05E+00 |
| ENSG000001 | ZNF219 | 6.20E-02 | 3.26E-01 | 1.16E+00 | 2.09E-01 | 1.16E+00 |
| ENSG000001 | BDNF | 6.20E-02 | 3.26E-01 | 1.40E+00 | 4.82E-01 | 1.40E+00 |
| ENSG000001 | C2CD5 | 6.21E-02 | 3.26E-01 | 1.10E+00 | 1.35E-01 | 1.10E+00 |
| ENSG000001 | DCTPP1 | 6.21E-02 | 3.26E-01 | 1.13E+00 | 1.77E-01 | 1.13E+00 |
| ENSG000002 | MIR194-2HG | 6.21E-02 | ? | 2.47E+00 | 1.30E+00 | 2.47E+00 |
| ENSG000002 | --- | 6.21E-02 | 3.26E-01 | 8.55E-01 | -2.26E-01 | -1.17E+00 |
| ENSG000002 | LINC02614 | 6.22E-02 | 3.26E-01 | 1.33E+00 | 4.11E-01 | 1.33E+00 |
| ENSG000001 | FAM91A1 | 6.22E-02 | 3.26E-01 | 9.40E-01 | -8.89E-02 | -1.06E+00 |
| ENSG000001 | IGF2BP3 | 6.23E-02 | 3.27E-01 | 1.08E+00 | 1.08E-01 | 1.08E+00 |
| ENSG000001 | EEF1A1 | 6.24E-02 | 3.27E-01 | 1.09E+00 | 1.25E-01 | 1.09E+00 |
| ENSG000001 | CMKLR2 | 6.24E-02 | 3.27E-01 | 1.29E+00 | 3.69E-01 | 1.29E+00 |
| ENSG000001 | KPTN | 6.24E-02 | 3.27E-01 | 1.13E+00 | 1.79E-01 | 1.13E+00 |
| ENSG000001 | SLX1B | 6.24E-02 | 3.27E-01 | 1.38E+00 | 4.63E-01 | 1.38E+00 |
| ENSG000001 | PRSS23 | 6.27E-02 | 3.28E-01 | 1.08E+00 | 1.09E-01 | 1.08E+00 |
| ENSG00000C | RRM2B | 6.27E-02 | 3.28E-01 | 1.07E+00 | 9.53E-02 | 1.07E+00 |
| ENSG00000C | CHMP5 | 6.28E-02 | 3.28E-01 | 9.35E-01 | -9.69E-02 | -1.07E+00 |
| ENSG000002 | --- | 6.28E-02 | ? | 2.22E+00 | 1.15E+00 | 2.22E+00 |
| ENSG000001 | DMAC1 | 6.28E-02 | 3.29E-01 | 1.10E+00 | 1.41E-01 | 1.10E+00 |
| ENSG00000C | WDR37 | 6.29E-02 | 3.29E-01 | 1.09E+00 | 1.19E-01 | 1.09E+00 |
| ENSG000002 | --- | 6.29E-02 | ? | 7.14E-01 | -4.85E-01 | -1.40E+00 |
| ENSG000002 | --- | 6.29E-02 | ? | 1.49E+00 | 5.71E-01 | 1.49E+00 |
| ENSG000002 | LINC01436 | 6.29E-02 | ? | 1.69E+00 | 7.58E-01 | 1.69E+00 |
| ENSG000001 | ADAR | 6.30E-02 | 3.29E-01 | 1.04E+00 | 5.36E-02 | 1.04E+00 |
| ENSG000001 | BCHE | 6.32E-02 | ? | 6.02E-01 | -7.33E-01 | -1.66E+00 |
| ENSG000001 | TMEM234 | 6.32E-02 | 3.30E-01 | 1.10E+00 | 1.43E-01 | 1.10E+00 |

| | | | | | | |
|---|---|---|---|---|---|---|
| ENSG000001 | SLC43A2 | 6.32E-02 | 3.30E-01 | 1.15E+00 | 1.96E-01 | 1.15E+00 |
| ENSG000001 | BEST3 | 6.32E-02 | ? | 6.31E-01 | -6.64E-01 | -1.58E+00 |
| ENSG000001 | KRR1 | 6.34E-02 | 3.31E-01 | 9.18E-01 | -1.23E-01 | -1.09E+00 |
| ENSG000001 | PACS2 | 6.35E-02 | 3.31E-01 | 1.08E+00 | 1.13E-01 | 1.08E+00 |
| ENSG000001 | MAD2L1BP | 6.35E-02 | 3.31E-01 | 1.09E+00 | 1.22E-01 | 1.09E+00 |
| ENSG000001 | PMEL | 6.36E-02 | 3.32E-01 | 1.31E+00 | 3.91E-01 | 1.31E+00 |
| ENSG000001 | GTF2A1 | 6.36E-02 | 3.32E-01 | 9.14E-01 | -1.30E-01 | -1.09E+00 |
| ENSG000001 | ZXDB | 6.37E-02 | 3.32E-01 | 1.07E+00 | 9.18E-02 | 1.07E+00 |
| ENSG000002 | --- | 6.37E-02 | 3.32E-01 | 1.45E+00 | 5.40E-01 | 1.45E+00 |
| ENSG000001 | TUBA1A | 6.38E-02 | 3.32E-01 | 9.39E-01 | -9.04E-02 | -1.06E+00 |
| ENSG00000C | RHOBTB1 | 6.38E-02 | 3.32E-01 | 1.09E+00 | 1.29E-01 | 1.09E+00 |
| ENSG000001 | CGNL1 | 6.38E-02 | 3.32E-01 | 1.42E+00 | 5.05E-01 | 1.42E+00 |
| ENSG000001 | KIAA0232 | 6.39E-02 | 3.32E-01 | 9.31E-01 | -1.03E-01 | -1.07E+00 |
| ENSG00000C | KIF1B | 6.41E-02 | 3.33E-01 | 9.47E-01 | -7.85E-02 | -1.06E+00 |
| ENSG000001 | PRPF6 | 6.42E-02 | 3.33E-01 | 9.52E-01 | -7.08E-02 | -1.05E+00 |
| ENSG00000C | RRP12 | 6.42E-02 | 3.33E-01 | 9.25E-01 | -1.13E-01 | -1.08E+00 |
| ENSG00000C | CS | 6.42E-02 | 3.33E-01 | 1.04E+00 | 5.41E-02 | 1.04E+00 |
| ENSG00000C | RIPOR1 | 6.42E-02 | 3.33E-01 | 9.27E-01 | -1.09E-01 | -1.08E+00 |
| ENSG000002 | MUC19 | 6.42E-02 | 3.33E-01 | 1.32E+00 | 4.04E-01 | 1.32E+00 |
| ENSG000001 | TENT5B | 6.43E-02 | 3.34E-01 | 1.30E+00 | 3.76E-01 | 1.30E+00 |
| ENSG000001 | TRNAU1AP | 6.44E-02 | 3.34E-01 | 1.08E+00 | 1.06E-01 | 1.08E+00 |
| ENSG000001 | ANKRD36B | 6.44E-02 | 3.34E-01 | 8.99E-01 | -1.54E-01 | -1.11E+00 |
| ENSG000001 | EXT1 | 6.44E-02 | 3.34E-01 | 1.06E+00 | 7.88E-02 | 1.06E+00 |
| ENSG000001 | APH1A | 6.45E-02 | 3.34E-01 | 1.12E+00 | 1.58E-01 | 1.12E+00 |
| ENSG000001 | TMEM236 | 6.45E-02 | 3.34E-01 | 8.16E-01 | -2.93E-01 | -1.23E+00 |
| ENSG000001 | NPM3 | 6.45E-02 | 3.34E-01 | 1.11E+00 | 1.56E-01 | 1.11E+00 |
| ENSG00000C | ALDH18A1 | 6.45E-02 | 3.34E-01 | 1.09E+00 | 1.19E-01 | 1.09E+00 |
| ENSG000002 | ZDHHC20P4 | 6.45E-02 | ? | 3.13E-01 | -1.68E+00 | -3.20E+00 |
| ENSG000001 | MRPL39 | 6.45E-02 | 3.34E-01 | 1.07E+00 | 1.02E-01 | 1.07E+00 |
| ENSG000001 | ZUP1 | 6.46E-02 | 3.34E-01 | 9.15E-01 | -1.28E-01 | -1.09E+00 |
| ENSG000001 | HPS4 | 6.46E-02 | 3.34E-01 | 9.28E-01 | -1.07E-01 | -1.08E+00 |
| ENSG000002 | ODCP | 6.47E-02 | ? | 5.91E-01 | -7.59E-01 | -1.69E+00 |
| ENSG000001 | PFKL | 6.47E-02 | 3.34E-01 | 1.12E+00 | 1.67E-01 | 1.12E+00 |
| ENSG000001 | RAD50 | 6.47E-02 | 3.34E-01 | 9.18E-01 | -1.23E-01 | -1.09E+00 |
| ENSG000002 | TRIM39-RPP2 | 6.47E-02 | ? | 6.40E+00 | 2.68E+00 | 6.40E+00 |
| ENSG000001 | AMER1 | 6.47E-02 | 3.34E-01 | 1.15E+00 | 2.01E-01 | 1.15E+00 |
| ENSG000002 | --- | 6.48E-02 | ? | 1.34E+00 | 4.28E-01 | 1.34E+00 |
| ENSG000002 | --- | 6.48E-02 | ? | 4.36E-01 | -1.20E+00 | -2.29E+00 |
| ENSG000001 | PXYLP1 | 6.48E-02 | 3.34E-01 | 1.13E+00 | 1.71E-01 | 1.13E+00 |
| ENSG000001 | C12orf76 | 6.48E-02 | 3.34E-01 | 1.16E+00 | 2.16E-01 | 1.16E+00 |
| ENSG000001 | ST6GALNAC4 | 6.49E-02 | 3.34E-01 | 1.16E+00 | 2.20E-01 | 1.16E+00 |
| ENSG000001 | BTBD16 | 6.49E-02 | ? | 1.67E+00 | 7.43E-01 | 1.67E+00 |
| ENSG000001 | SDHAF2 | 6.49E-02 | 3.35E-01 | 1.07E+00 | 1.02E-01 | 1.07E+00 |
| ENSG000001 | PPP4R1L | 6.49E-02 | ? | 2.28E+00 | 1.19E+00 | 2.28E+00 |
| ENSG000001 | ARFGAP2 | 6.50E-02 | 3.35E-01 | 1.09E+00 | 1.26E-01 | 1.09E+00 |

| | | | | | | |
|---|---|---|---|---|---|---|
| ENSG000002 | --- | 6.50E-02 | ? | 1.63E+00 | 7.08E-01 | 1.63E+00 |
| ENSG000001 | CFAP299 | 6.50E-02 | ? | 4.69E-01 | -1.09E+00 | -2.13E+00 |
| ENSG000001 | WDSUB1 | 6.51E-02 | 3.35E-01 | 1.15E+00 | 2.05E-01 | 1.15E+00 |
| ENSG000002 | --- | 6.51E-02 | 3.35E-01 | 7.86E-01 | -3.47E-01 | -1.27E+00 |
| ENSG000001 | ZNF454 | 6.51E-02 | 3.35E-01 | 1.31E+00 | 3.87E-01 | 1.31E+00 |
| ENSG000001 | SCN2B | 6.52E-02 | 3.35E-01 | 1.20E+00 | 2.57E-01 | 1.20E+00 |
| ENSG000001 | ATP6V1D | 6.52E-02 | 3.35E-01 | 9.46E-01 | -7.96E-02 | -1.06E+00 |
| ENSG000001 | SAE1 | 6.52E-02 | 3.35E-01 | 1.05E+00 | 7.06E-02 | 1.05E+00 |
| ENSG000001 | PHYHIPL | 6.53E-02 | 3.35E-01 | 8.40E-01 | -2.52E-01 | -1.19E+00 |
| ENSG000001 | VWA5A | 6.53E-02 | 3.35E-01 | 1.07E+00 | 1.02E-01 | 1.07E+00 |
| ENSG000002 | PDGFA-DT | 6.53E-02 | ? | 1.61E+00 | 6.90E-01 | 1.61E+00 |
| ENSG000001 | P4HA1 | 6.53E-02 | 3.35E-01 | 9.33E-01 | -9.99E-02 | -1.07E+00 |
| ENSG000001 | NELFCD | 6.53E-02 | 3.35E-01 | 1.06E+00 | 7.87E-02 | 1.06E+00 |
| ENSG000001 | GOLGA6L3 | 6.54E-02 | ? | 4.84E-01 | -1.05E+00 | -2.06E+00 |
| ENSG000001 | PPFIA3 | 6.54E-02 | 3.36E-01 | 1.34E+00 | 4.24E-01 | 1.34E+00 |
| ENSG00000C | LRP6 | 6.55E-02 | 3.36E-01 | 9.24E-01 | -1.14E-01 | -1.08E+00 |
| ENSG000001 | YWHAZ | 6.55E-02 | 3.36E-01 | 9.51E-01 | -7.18E-02 | -1.05E+00 |
| ENSG000001 | ULK1 | 6.55E-02 | 3.36E-01 | 8.78E-01 | -1.88E-01 | -1.14E+00 |
| ENSG000001 | NCEH1 | 6.56E-02 | 3.36E-01 | 8.93E-01 | -1.63E-01 | -1.12E+00 |
| ENSG000001 | VTCN1 | 6.56E-02 | ? | 1.55E+00 | 6.31E-01 | 1.55E+00 |
| ENSG00000C | COQ9 | 6.56E-02 | 3.36E-01 | 1.07E+00 | 9.55E-02 | 1.07E+00 |
| ENSG000001 | MMS22L | 6.56E-02 | 3.36E-01 | 9.19E-01 | -1.22E-01 | -1.09E+00 |
| ENSG000001 | MSX1 | 6.57E-02 | ? | 6.76E-01 | -5.64E-01 | -1.48E+00 |
| ENSG000001 | FARP1 | 6.57E-02 | 3.36E-01 | 9.47E-01 | -7.80E-02 | -1.06E+00 |
| ENSG000001 | ZNF587 | 6.57E-02 | 3.36E-01 | 9.23E-01 | -1.15E-01 | -1.08E+00 |
| ENSG000001 | FST | 6.58E-02 | 3.36E-01 | 1.14E+00 | 1.88E-01 | 1.14E+00 |
| ENSG000001 | AK8 | 6.58E-02 | ? | 1.97E+00 | 9.79E-01 | 1.97E+00 |
| ENSG000001 | GOLGB1 | 6.58E-02 | 3.37E-01 | 9.08E-01 | -1.40E-01 | -1.10E+00 |
| ENSG00000C | PIGS | 6.58E-02 | 3.37E-01 | 1.08E+00 | 1.16E-01 | 1.08E+00 |
| ENSG00000C | CTPS2 | 6.59E-02 | 3.37E-01 | 9.19E-01 | -1.22E-01 | -1.09E+00 |
| ENSG000002 | ZBED5 | 6.59E-02 | 3.37E-01 | 9.41E-01 | -8.72E-02 | -1.06E+00 |
| ENSG000002 | ADAM21P1 | 6.59E-02 | ? | 1.76E+00 | 8.16E-01 | 1.76E+00 |
| ENSG000001 | GHR | 6.59E-02 | 3.37E-01 | 1.15E+00 | 1.96E-01 | 1.15E+00 |
| ENSG000001 | KRT7 | 6.60E-02 | 3.37E-01 | 9.18E-01 | -1.23E-01 | -1.09E+00 |
| ENSG000001 | AURKAIP1 | 6.60E-02 | 3.37E-01 | 1.09E+00 | 1.25E-01 | 1.09E+00 |
| ENSG000001 | FIBCD1 | 6.60E-02 | ? | 6.49E-01 | -6.25E-01 | -1.54E+00 |
| ENSG000001 | PRR16 | 6.61E-02 | ? | 1.44E+00 | 5.26E-01 | 1.44E+00 |
| ENSG000001 | IL36RN | 6.61E-02 | ? | 1.55E+00 | 6.28E-01 | 1.55E+00 |
| ENSG000002 | FAM27C | 6.61E-02 | 3.37E-01 | 1.25E+00 | 3.17E-01 | 1.25E+00 |
| ENSG000002 | FAM192BP | 6.62E-02 | ? | 5.24E-01 | -9.32E-01 | -1.91E+00 |
| ENSG000001 | TCN1 | 6.62E-02 | 3.37E-01 | 7.43E-01 | -4.29E-01 | -1.35E+00 |
| ENSG000002 | ZNF224 | 6.62E-02 | 3.37E-01 | 8.52E-01 | -2.31E-01 | -1.17E+00 |
| ENSG000001 | RFXAP | 6.62E-02 | 3.37E-01 | 1.23E+00 | 3.01E-01 | 1.23E+00 |
| ENSG000002 | CALM2P2 | 6.62E-02 | ? | 6.85E-01 | -5.46E-01 | -1.46E+00 |
| ENSG000002 | ASDURF | 6.62E-02 | ? | 5.31E-01 | -9.12E-01 | -1.88E+00 |

| | | | | | | |
|---|---|---|---|---|---|---|
| ENSG000001 | EDAR | 6.62E-02 | 3.37E-01 | 1.29E+00 | 3.72E-01 | 1.29E+00 |
| ENSG000002 | CHROMR | 6.63E-02 | 3.37E-01 | 9.01E-01 | -1.50E-01 | -1.11E+00 |
| ENSG000002 | --- | 6.63E-02 | 3.37E-01 | 1.61E+00 | 6.87E-01 | 1.61E+00 |
| ENSG000001 | PPP1CC | 6.63E-02 | 3.37E-01 | 9.50E-01 | -7.47E-02 | -1.05E+00 |
| ENSG000002 | --- | 6.63E-02 | ? | 7.10E-01 | -4.95E-01 | -1.41E+00 |
| ENSG000002 | ZNF496-DT | 6.65E-02 | 3.38E-01 | 1.13E+00 | 1.73E-01 | 1.13E+00 |
| ENSG000002 | SH3GL1P2 | 6.65E-02 | ? | 4.02E-01 | -1.32E+00 | -2.49E+00 |
| ENSG000001 | FN3K | 6.65E-02 | 3.38E-01 | 8.80E-01 | -1.84E-01 | -1.14E+00 |
| ENSG000001 | SLC26A11 | 6.65E-02 | 3.38E-01 | 1.19E+00 | 2.51E-01 | 1.19E+00 |
| ENSG000001 | PUDP | 6.66E-02 | 3.38E-01 | 9.41E-01 | -8.81E-02 | -1.06E+00 |
| ENSG000001 | PLD5 | 6.66E-02 | 3.38E-01 | 8.66E-01 | -2.08E-01 | -1.16E+00 |
| ENSG000001 | ERAP1 | 6.66E-02 | 3.38E-01 | 9.43E-01 | -8.46E-02 | -1.06E+00 |
| ENSG000001 | NREP | 6.66E-02 | 3.38E-01 | 1.13E+00 | 1.78E-01 | 1.13E+00 |
| ENSG000001 | FAM32A | 6.66E-02 | 3.38E-01 | 9.31E-01 | -1.03E-01 | -1.07E+00 |
| ENSG000002 | HSD17B8 | 6.67E-02 | 3.38E-01 | 1.20E+00 | 2.67E-01 | 1.20E+00 |
| ENSG000001 | GNAI1 | 6.68E-02 | 3.38E-01 | 1.06E+00 | 8.70E-02 | 1.06E+00 |
| ENSG000001 | HPS1 | 6.68E-02 | 3.38E-01 | 1.10E+00 | 1.43E-01 | 1.10E+00 |
| ENSG000001 | SMU1 | 6.68E-02 | 3.38E-01 | 1.04E+00 | 6.35E-02 | 1.04E+00 |
| ENSG000002 | --- | 6.68E-02 | 3.38E-01 | 8.00E-01 | -3.23E-01 | -1.25E+00 |
| ENSG000001 | KIT | 6.68E-02 | ? | 1.86E+00 | 8.97E-01 | 1.86E+00 |
| ENSG000001 | NDUFA2 | 6.69E-02 | 3.38E-01 | 1.13E+00 | 1.78E-01 | 1.13E+00 |
| ENSG000002 | QTRT1 | 6.69E-02 | 3.38E-01 | 1.16E+00 | 2.12E-01 | 1.16E+00 |
| ENSG000002 | FBXW10B | 6.69E-02 | 3.38E-01 | 8.33E-01 | -2.63E-01 | -1.20E+00 |
| ENSG000002 | RPL13AP5 | 6.69E-02 | 3.38E-01 | 1.15E+00 | 1.96E-01 | 1.15E+00 |
| ENSG000001 | BSDC1 | 6.69E-02 | 3.38E-01 | 1.08E+00 | 1.07E-01 | 1.08E+00 |
| ENSG000001 | RPL27 | 6.70E-02 | 3.38E-01 | 1.07E+00 | 9.20E-02 | 1.07E+00 |
| ENSG000001 | ACLY | 6.70E-02 | 3.38E-01 | 1.07E+00 | 1.03E-01 | 1.07E+00 |
| ENSG000001 | SLC25A32 | 6.70E-02 | 3.38E-01 | 9.16E-01 | -1.27E-01 | -1.09E+00 |
| ENSG000001 | C1QTNF6 | 6.70E-02 | 3.38E-01 | 1.06E+00 | 8.75E-02 | 1.06E+00 |
| ENSG000002 | --- | 6.70E-02 | 3.38E-01 | 9.08E-01 | -1.39E-01 | -1.10E+00 |
| ENSG000001 | CNKSR3 | 6.70E-02 | 3.38E-01 | 1.15E+00 | 2.06E-01 | 1.15E+00 |
| ENSG000001 | VPS37B | 6.71E-02 | 3.38E-01 | 1.07E+00 | 1.00E-01 | 1.07E+00 |
| ENSG000002 | SAP30L-AS1 | 6.71E-02 | 3.38E-01 | 1.10E+00 | 1.41E-01 | 1.10E+00 |
| ENSG000001 | STIL | 6.72E-02 | 3.38E-01 | 8.93E-01 | -1.64E-01 | -1.12E+00 |
| ENSG000001 | DFFB | 6.72E-02 | 3.39E-01 | 1.15E+00 | 2.02E-01 | 1.15E+00 |
| ENSG000001 | TAF3 | 6.73E-02 | 3.39E-01 | 9.12E-01 | -1.32E-01 | -1.10E+00 |
| ENSG000002 | LINC00958 | 6.73E-02 | 3.39E-01 | 1.07E+00 | 9.89E-02 | 1.07E+00 |
| ENSG000001 | SURF2 | 6.74E-02 | 3.39E-01 | 1.09E+00 | 1.25E-01 | 1.09E+00 |
| ENSG000001 | LIPA | 6.74E-02 | 3.39E-01 | 1.07E+00 | 9.53E-02 | 1.07E+00 |
| ENSG00000C | RGS17 | 6.74E-02 | 3.39E-01 | 8.27E-01 | -2.75E-01 | -1.21E+00 |
| ENSG000001 | COIL | 6.74E-02 | 3.39E-01 | 1.07E+00 | 9.63E-02 | 1.07E+00 |
| ENSG00000C | MBTD1 | 6.75E-02 | 3.39E-01 | 9.03E-01 | -1.47E-01 | -1.11E+00 |
| ENSG000001 | HILPDA | 6.75E-02 | 3.39E-01 | 1.13E+00 | 1.73E-01 | 1.13E+00 |
| ENSG000001 | SHROOM3 | 6.75E-02 | 3.39E-01 | 8.97E-01 | -1.57E-01 | -1.12E+00 |
| ENSG000001 | RBBP7 | 6.75E-02 | 3.39E-01 | 1.06E+00 | 7.96E-02 | 1.06E+00 |

| | | | | | | |
|---|---|---|---|---|---|---|
| ENSG000001 | NAMPT | 6.76E-02 | 3.39E-01 | 9.28E-01 | -1.08E-01 | -1.08E+00 |
| ENSG00000C | SOX30 | 6.76E-02 | ? | 5.83E-01 | -7.79E-01 | -1.72E+00 |
| ENSG000001 | STXBP4 | 6.76E-02 | 3.39E-01 | 9.22E-01 | -1.16E-01 | -1.08E+00 |
| ENSG000001 | ZNF354B | 6.76E-02 | 3.39E-01 | 8.51E-01 | -2.33E-01 | -1.18E+00 |
| ENSG000001 | TERF2IP | 6.77E-02 | 3.39E-01 | 9.22E-01 | -1.17E-01 | -1.08E+00 |
| ENSG000001 | TGM2 | 6.77E-02 | 3.39E-01 | 7.76E-01 | -3.66E-01 | -1.29E+00 |
| ENSG000001 | CAMLG | 6.77E-02 | 3.39E-01 | 9.42E-01 | -8.60E-02 | -1.06E+00 |
| ENSG000001 | PAPOLG | 6.77E-02 | 3.39E-01 | 9.22E-01 | -1.17E-01 | -1.08E+00 |
| ENSG000001 | WASHC3 | 6.78E-02 | 3.39E-01 | 1.07E+00 | 9.22E-02 | 1.07E+00 |
| ENSG000001 | PPP4C | 6.78E-02 | 3.39E-01 | 1.11E+00 | 1.56E-01 | 1.11E+00 |
| ENSG000001 | RASGRF2 | 6.78E-02 | 3.39E-01 | 8.76E-01 | -1.91E-01 | -1.14E+00 |
| ENSG00000C | EVC | 6.79E-02 | 3.40E-01 | 1.07E+00 | 1.01E-01 | 1.07E+00 |
| ENSG000001 | RNF38 | 6.79E-02 | 3.40E-01 | 1.10E+00 | 1.34E-01 | 1.10E+00 |
| ENSG000001 | FA2H | 6.80E-02 | ? | 5.85E-01 | -7.74E-01 | -1.71E+00 |
| ENSG000001 | NODAL | 6.80E-02 | ? | 1.38E+00 | 4.68E-01 | 1.38E+00 |
| ENSG000001 | PRR5L | 6.81E-02 | 3.40E-01 | 1.07E+00 | 1.02E-01 | 1.07E+00 |
| ENSG000001 | ZNF385D | 6.81E-02 | ? | 1.61E+00 | 6.87E-01 | 1.61E+00 |
| ENSG000001 | GFOD1 | 6.81E-02 | 3.40E-01 | 9.20E-01 | -1.21E-01 | -1.09E+00 |
| ENSG000001 | SATB1 | 6.81E-02 | 3.40E-01 | 1.11E+00 | 1.50E-01 | 1.11E+00 |
| ENSG00000C | FXYD3 | 6.82E-02 | 3.40E-01 | 1.16E+00 | 2.14E-01 | 1.16E+00 |
| ENSG000001 | GBP2 | 6.82E-02 | ? | 1.40E+00 | 4.88E-01 | 1.40E+00 |
| ENSG000002 | SLC23A3 | 6.82E-02 | 3.40E-01 | 8.04E-01 | -3.15E-01 | -1.24E+00 |
| ENSG000002 | XIST | 6.82E-02 | 3.40E-01 | 8.81E-01 | -1.83E-01 | -1.14E+00 |
| ENSG000001 | CLSTN2 | 6.82E-02 | ? | 1.45E+00 | 5.37E-01 | 1.45E+00 |
| ENSG000002 | --- | 6.83E-02 | ? | 1.77E+00 | 8.24E-01 | 1.77E+00 |
| ENSG000001 | TTYH1 | 6.84E-02 | ? | 1.61E+00 | 6.84E-01 | 1.61E+00 |
| ENSG000001 | PSMD7 | 6.85E-02 | 3.41E-01 | 9.54E-01 | -6.82E-02 | -1.05E+00 |
| ENSG000001 | NCS1 | 6.85E-02 | 3.41E-01 | 9.10E-01 | -1.36E-01 | -1.10E+00 |
| ENSG000001 | PP7080 | 6.85E-02 | 3.41E-01 | 1.17E+00 | 2.22E-01 | 1.17E+00 |
| ENSG000001 | CPM | 6.85E-02 | 3.41E-01 | 9.21E-01 | -1.19E-01 | -1.09E+00 |
| ENSG000001 | NKIRAS2 | 6.86E-02 | 3.42E-01 | 9.34E-01 | -9.89E-02 | -1.07E+00 |
| ENSG000002 | PTMAP5 | 6.87E-02 | 3.42E-01 | 8.10E-01 | -3.04E-01 | -1.23E+00 |
| ENSG000001 | SHANK2 | 6.88E-02 | 3.42E-01 | 9.23E-01 | -1.15E-01 | -1.08E+00 |
| ENSG000001 | ATN1 | 6.88E-02 | 3.42E-01 | 9.47E-01 | -7.86E-02 | -1.06E+00 |
| ENSG000002 | RPL17-C18or | 6.88E-02 | 3.42E-01 | 9.40E-01 | -9.00E-02 | -1.06E+00 |
| ENSG000002 | ANKRD39 | 6.88E-02 | 3.42E-01 | 8.53E-01 | -2.29E-01 | -1.17E+00 |
| ENSG00000C | LPAR2 | 6.88E-02 | 3.42E-01 | 1.11E+00 | 1.52E-01 | 1.11E+00 |
| ENSG000002 | HSPA5P1 | 6.88E-02 | ? | 2.05E+00 | 1.04E+00 | 2.05E+00 |
| ENSG000001 | CTRL | 6.89E-02 | ? | 2.16E+00 | 1.11E+00 | 2.16E+00 |
| ENSG000001 | MGAT1 | 6.89E-02 | 3.42E-01 | 1.11E+00 | 1.48E-01 | 1.11E+00 |
| ENSG000001 | PRDM8 | 6.90E-02 | 3.43E-01 | 8.67E-01 | -2.06E-01 | -1.15E+00 |
| ENSG000002 | --- | 6.90E-02 | ? | 5.00E-01 | -1.00E+00 | -2.00E+00 |
| ENSG000001 | EOGT | 6.90E-02 | 3.43E-01 | 9.25E-01 | -1.12E-01 | -1.08E+00 |
| ENSG000001 | KLHL11 | 6.91E-02 | 3.43E-01 | 1.08E+00 | 1.16E-01 | 1.08E+00 |
| ENSG000001 | ZHX1 | 6.91E-02 | 3.43E-01 | 1.07E+00 | 9.32E-02 | 1.07E+00 |

| | | | | | | |
|---|---|---|---|---|---|---|
| ENSG000002 | TXNIP | 6.92E-02 | 3.43E-01 | 1.07E+00 | 9.96E-02 | 1.07E+00 |
| ENSG000001 | ZNF770 | 6.92E-02 | 3.43E-01 | 9.41E-01 | -8.81E-02 | -1.06E+00 |
| ENSG000001 | STAM | 6.92E-02 | 3.43E-01 | 9.36E-01 | -9.56E-02 | -1.07E+00 |
| ENSG000002 | --- | 6.93E-02 | ? | 1.61E+00 | 6.86E-01 | 1.61E+00 |
| ENSG000001 | MEAK7 | 6.94E-02 | 3.44E-01 | 1.07E+00 | 1.02E-01 | 1.07E+00 |
| ENSG000001 | SCAP | 6.94E-02 | 3.44E-01 | 1.10E+00 | 1.44E-01 | 1.10E+00 |
| ENSG000002 | ZNF529-AS1 | 6.95E-02 | 3.44E-01 | 7.83E-01 | -3.52E-01 | -1.28E+00 |
| ENSG000001 | ATP5F1E | 6.95E-02 | 3.44E-01 | 1.07E+00 | 9.38E-02 | 1.07E+00 |
| ENSG000002 | LINC02899 | 6.95E-02 | ? | 5.39E-01 | -8.92E-01 | -1.86E+00 |
| ENSG000001 | PRPF40A | 6.96E-02 | 3.44E-01 | 9.18E-01 | -1.24E-01 | -1.09E+00 |
| ENSG000001 | MTIF3 | 6.96E-02 | 3.44E-01 | 9.17E-01 | -1.26E-01 | -1.09E+00 |
| ENSG000001 | UBE3B | 6.96E-02 | 3.44E-01 | 1.04E+00 | 6.18E-02 | 1.04E+00 |
| ENSG000001 | MPZL2 | 6.96E-02 | 3.44E-01 | 1.16E+00 | 2.17E-01 | 1.16E+00 |
| ENSG000001 | NALCN | 6.96E-02 | 3.44E-01 | 1.30E+00 | 3.83E-01 | 1.30E+00 |
| ENSG000002 | CENPW | 6.97E-02 | 3.44E-01 | 8.45E-01 | -2.43E-01 | -1.18E+00 |
| ENSG000002 | --- | 6.97E-02 | ? | 1.98E+00 | 9.87E-01 | 1.98E+00 |
| ENSG000001 | EML4 | 6.98E-02 | 3.45E-01 | 9.35E-01 | -9.73E-02 | -1.07E+00 |
| ENSG00000C | PLEKHO1 | 6.99E-02 | 3.45E-01 | 1.21E+00 | 2.80E-01 | 1.21E+00 |
| ENSG000001 | PRSS16 | 6.99E-02 | 3.45E-01 | 9.06E-01 | -1.43E-01 | -1.10E+00 |
| ENSG000001 | TEPP | 6.99E-02 | 3.45E-01 | 1.27E+00 | 3.40E-01 | 1.27E+00 |
| ENSG000001 | EPB41L5 | 6.99E-02 | 3.45E-01 | 9.17E-01 | -1.25E-01 | -1.09E+00 |
| ENSG000001 | TNIK | 7.00E-02 | 3.45E-01 | 8.80E-01 | -1.85E-01 | -1.14E+00 |
| ENSG000001 | SLC27A3 | 7.01E-02 | 3.45E-01 | 1.14E+00 | 1.90E-01 | 1.14E+00 |
| ENSG000001 | TBRG1 | 7.01E-02 | 3.45E-01 | 9.39E-01 | -9.03E-02 | -1.06E+00 |
| ENSG000001 | GNRH1 | 7.01E-02 | 3.45E-01 | 7.94E-01 | -3.33E-01 | -1.26E+00 |
| ENSG000001 | NDUFA1 | 7.01E-02 | 3.45E-01 | 1.10E+00 | 1.31E-01 | 1.10E+00 |
| ENSG000001 | CCDC71 | 7.01E-02 | 3.45E-01 | 1.11E+00 | 1.50E-01 | 1.11E+00 |
| ENSG000002 | KRT16P3 | 7.02E-02 | ? | 5.18E-01 | -9.50E-01 | -1.93E+00 |
| ENSG000002 | --- | 7.02E-02 | ? | 5.90E-01 | -7.60E-01 | -1.69E+00 |
| ENSG00000C | ZNF446 | 7.02E-02 | 3.45E-01 | 1.18E+00 | 2.36E-01 | 1.18E+00 |
| ENSG000002 | FIS1 | 7.03E-02 | 3.45E-01 | 1.07E+00 | 9.51E-02 | 1.07E+00 |
| ENSG00000C | FAM114A2 | 7.03E-02 | 3.45E-01 | 9.38E-01 | -9.22E-02 | -1.07E+00 |
| ENSG000002 | SLC22A31 | 7.03E-02 | ? | 2.44E+00 | 1.29E+00 | 2.44E+00 |
| ENSG000002 | AGAP13P | 7.03E-02 | 3.45E-01 | 7.74E-01 | -3.70E-01 | -1.29E+00 |
| ENSG000001 | AP1S1 | 7.03E-02 | 3.45E-01 | 1.06E+00 | 8.41E-02 | 1.06E+00 |
| ENSG000001 | LMBR1L | 7.04E-02 | 3.45E-01 | 1.12E+00 | 1.61E-01 | 1.12E+00 |
| ENSG000002 | TMEM231 | 7.04E-02 | 3.45E-01 | 1.09E+00 | 1.25E-01 | 1.09E+00 |
| ENSG000001 | ACAP2 | 7.05E-02 | 3.46E-01 | 9.44E-01 | -8.32E-02 | -1.06E+00 |
| ENSG000002 | C16orf95 | 7.05E-02 | ? | 1.43E+00 | 5.20E-01 | 1.43E+00 |
| ENSG000001 | DTWD1 | 7.05E-02 | 3.46E-01 | 9.10E-01 | -1.37E-01 | -1.10E+00 |
| ENSG000001 | ABHD15 | 7.06E-02 | 3.46E-01 | 1.09E+00 | 1.28E-01 | 1.09E+00 |
| ENSG000002 | TMCO1-AS1 | 7.06E-02 | ? | 2.01E+00 | 1.01E+00 | 2.01E+00 |
| ENSG000001 | MGARP | 7.06E-02 | ? | 1.70E+00 | 7.65E-01 | 1.70E+00 |
| ENSG000002 | --- | 7.07E-02 | ? | 1.98E+00 | 9.88E-01 | 1.98E+00 |
| ENSG000001 | IFI44 | 7.07E-02 | ? | 6.70E-01 | -5.77E-01 | -1.49E+00 |

| | | | | | | |
|---|---|---|---|---|---|---|
| ENSG000001 | STX1A | 7.07E-02 | 3.46E-01 | 9.24E-01 | -1.14E-01 | -1.08E+00 |
| ENSG000001 | PEDS1-UBE2 | 7.07E-02 | 3.46E-01 | 1.44E+00 | 5.28E-01 | 1.44E+00 |
| ENSG000002 | --- | 7.07E-02 | ? | 1.63E+00 | 7.04E-01 | 1.63E+00 |
| ENSG000001 | LUZP1 | 7.07E-02 | 3.46E-01 | 1.06E+00 | 8.31E-02 | 1.06E+00 |
| ENSG00000C | SP140 | 7.08E-02 | 3.47E-01 | 8.22E-01 | -2.83E-01 | -1.22E+00 |
| ENSG000001 | ANKHD1 | 7.08E-02 | 3.47E-01 | 9.08E-01 | -1.39E-01 | -1.10E+00 |
| ENSG00000C | REEP1 | 7.08E-02 | ? | 6.22E-01 | -6.84E-01 | -1.61E+00 |
| ENSG000001 | CNNM2 | 7.09E-02 | 3.47E-01 | 1.13E+00 | 1.78E-01 | 1.13E+00 |
| ENSG000002 | --- | 7.09E-02 | ? | 3.79E-01 | -1.40E+00 | -2.64E+00 |
| ENSG000001 | RAB40B | 7.09E-02 | 3.47E-01 | 1.10E+00 | 1.36E-01 | 1.10E+00 |
| ENSG000001 | P4HTM | 7.10E-02 | 3.47E-01 | 1.14E+00 | 1.95E-01 | 1.14E+00 |
| ENSG00000C | FAR2 | 7.11E-02 | 3.47E-01 | 1.13E+00 | 1.71E-01 | 1.13E+00 |
| ENSG000001 | CELF4 | 7.11E-02 | ? | 5.64E-01 | -8.27E-01 | -1.77E+00 |
| ENSG000001 | OBI1 | 7.11E-02 | 3.48E-01 | 9.10E-01 | -1.37E-01 | -1.10E+00 |
| ENSG000001 | TRIM36 | 7.12E-02 | 3.48E-01 | 8.35E-01 | -2.60E-01 | -1.20E+00 |
| ENSG000001 | MICU3 | 7.12E-02 | 3.48E-01 | 1.16E+00 | 2.10E-01 | 1.16E+00 |
| ENSG000001 | ZSCAN12 | 7.13E-02 | 3.48E-01 | 8.90E-01 | -1.67E-01 | -1.12E+00 |
| ENSG000001 | LCK | 7.13E-02 | ? | 5.82E-01 | -7.81E-01 | -1.72E+00 |
| ENSG000001 | HMGA1 | 7.14E-02 | 3.48E-01 | 9.24E-01 | -1.14E-01 | -1.08E+00 |
| ENSG000001 | GORASP2 | 7.14E-02 | 3.48E-01 | 9.39E-01 | -9.05E-02 | -1.06E+00 |
| ENSG000001 | MRPL15 | 7.14E-02 | 3.48E-01 | 1.05E+00 | 7.63E-02 | 1.05E+00 |
| ENSG000001 | WHAMM | 7.15E-02 | 3.48E-01 | 9.30E-01 | -1.05E-01 | -1.08E+00 |
| ENSG000001 | RBMS1 | 7.15E-02 | 3.48E-01 | 9.40E-01 | -8.98E-02 | -1.06E+00 |
| ENSG000002 | MRPL45 | 7.15E-02 | 3.48E-01 | 9.43E-01 | -8.51E-02 | -1.06E+00 |
| ENSG00000C | EIF4G3 | 7.15E-02 | 3.48E-01 | 9.29E-01 | -1.06E-01 | -1.08E+00 |
| ENSG000001 | PGAP4 | 7.17E-02 | 3.49E-01 | 1.06E+00 | 8.70E-02 | 1.06E+00 |
| ENSG000001 | RBBP6 | 7.17E-02 | 3.49E-01 | 9.15E-01 | -1.29E-01 | -1.09E+00 |
| ENSG00000C | UBE2A | 7.17E-02 | 3.49E-01 | 9.52E-01 | -7.10E-02 | -1.05E+00 |
| ENSG000001 | NFKBIZ | 7.17E-02 | 3.49E-01 | 9.07E-01 | -1.40E-01 | -1.10E+00 |
| ENSG000002 | --- | 7.17E-02 | ? | 2.60E+00 | 1.38E+00 | 2.60E+00 |
| ENSG000001 | QRFP | 7.18E-02 | ? | 1.71E+00 | 7.74E-01 | 1.71E+00 |
| ENSG000002 | GASAL1 | 7.18E-02 | 3.49E-01 | 8.31E-01 | -2.67E-01 | -1.20E+00 |
| ENSG000001 | CHORDC1 | 7.18E-02 | 3.49E-01 | 9.02E-01 | -1.48E-01 | -1.11E+00 |
| ENSG000001 | GAS8 | 7.18E-02 | 3.49E-01 | 9.24E-01 | -1.14E-01 | -1.08E+00 |
| ENSG000002 | --- | 7.19E-02 | ? | 1.45E+00 | 5.33E-01 | 1.45E+00 |
| ENSG000001 | FAM182B | 7.20E-02 | ? | 2.53E+00 | 1.34E+00 | 2.53E+00 |
| ENSG000001 | RNASEK-C17 | 7.20E-02 | 3.49E-01 | 1.48E+00 | 5.70E-01 | 1.48E+00 |
| ENSG00000C | MAP3K20 | 7.21E-02 | 3.50E-01 | 1.06E+00 | 8.96E-02 | 1.06E+00 |
| ENSG000001 | ATP6V0D1 | 7.21E-02 | 3.50E-01 | 1.07E+00 | 9.71E-02 | 1.07E+00 |
| ENSG000001 | MANEAL | 7.21E-02 | 3.50E-01 | 1.29E+00 | 3.65E-01 | 1.29E+00 |
| ENSG000001 | PPP2R1B | 7.22E-02 | 3.50E-01 | 1.06E+00 | 8.49E-02 | 1.06E+00 |
| ENSG000001 | KCTD18 | 7.22E-02 | 3.50E-01 | 1.07E+00 | 1.04E-01 | 1.07E+00 |
| ENSG000002 | MIR1302-9H | 7.22E-02 | 3.50E-01 | 8.14E-01 | -2.97E-01 | -1.23E+00 |
| ENSG00000C | TUT7 | 7.23E-02 | 3.50E-01 | 9.25E-01 | -1.12E-01 | -1.08E+00 |
| ENSG000001 | UBL5 | 7.23E-02 | 3.50E-01 | 1.11E+00 | 1.56E-01 | 1.11E+00 |

| | | | | | | |
|---|---|---|---|---|---|---|
| ENSG00000C | DZANK1 | 7.23E-02 | 3.50E-01 | 8.62E-01 | -2.14E-01 | -1.16E+00 |
| ENSG000001 | TBX3 | 7.23E-02 | 3.50E-01 | 8.91E-01 | -1.67E-01 | -1.12E+00 |
| ENSG000001 | GCNT2 | 7.24E-02 | 3.50E-01 | 9.37E-01 | -9.38E-02 | -1.07E+00 |
| ENSG000001 | MRFAP1 | 7.24E-02 | 3.50E-01 | 9.58E-01 | -6.19E-02 | -1.04E+00 |
| ENSG000001 | PRRC1 | 7.24E-02 | 3.50E-01 | 9.34E-01 | -9.79E-02 | -1.07E+00 |
| ENSG000001 | SUMO3 | 7.24E-02 | 3.50E-01 | 1.04E+00 | 6.30E-02 | 1.04E+00 |
| ENSG000001 | CACNB2 | 7.25E-02 | ? | 7.33E-01 | -4.48E-01 | -1.36E+00 |
| ENSG000001 | CDK5R1 | 7.25E-02 | 3.50E-01 | 8.85E-01 | -1.76E-01 | -1.13E+00 |
| ENSG00000C | DERA | 7.25E-02 | 3.50E-01 | 1.06E+00 | 9.06E-02 | 1.06E+00 |
| ENSG00000C | ASPM | 7.26E-02 | 3.50E-01 | 8.13E-01 | -2.99E-01 | -1.23E+00 |
| ENSG000001 | SLC9A6 | 7.27E-02 | 3.51E-01 | 1.06E+00 | 8.76E-02 | 1.06E+00 |
| ENSG000001 | ANKRD36 | 7.28E-02 | 3.51E-01 | 8.91E-01 | -1.67E-01 | -1.12E+00 |
| ENSG000001 | LINC01106 | 7.28E-02 | 3.51E-01 | 1.13E+00 | 1.73E-01 | 1.13E+00 |
| ENSG000001 | MOSPD2 | 7.29E-02 | 3.51E-01 | 9.17E-01 | -1.25E-01 | -1.09E+00 |
| ENSG000002 | ACAP2-IT1 | 7.29E-02 | ? | 4.38E-01 | -1.19E+00 | -2.28E+00 |
| ENSG000001 | EZH2 | 7.30E-02 | 3.52E-01 | 9.01E-01 | -1.50E-01 | -1.11E+00 |
| ENSG000001 | TCF20 | 7.31E-02 | 3.52E-01 | 9.60E-01 | -5.84E-02 | -1.04E+00 |
| ENSG000002 | TATDN3 | 7.32E-02 | 3.53E-01 | 1.09E+00 | 1.29E-01 | 1.09E+00 |
| ENSG000002 | RPL23AP1 | 7.32E-02 | ? | 3.33E+00 | 1.74E+00 | 3.33E+00 |
| ENSG000001 | SON | 7.32E-02 | 3.53E-01 | 9.56E-01 | -6.53E-02 | -1.05E+00 |
| ENSG000002 | NNT-AS1 | 7.35E-02 | 3.54E-01 | 9.10E-01 | -1.37E-01 | -1.10E+00 |
| ENSG000001 | SPTBN4 | 7.35E-02 | ? | 1.56E+00 | 6.44E-01 | 1.56E+00 |
| ENSG000001 | ATP8B2 | 7.36E-02 | 3.54E-01 | 1.12E+00 | 1.62E-01 | 1.12E+00 |
| ENSG00000C | CDH3 | 7.36E-02 | 3.54E-01 | 1.06E+00 | 8.60E-02 | 1.06E+00 |
| ENSG000001 | G6PC3 | 7.37E-02 | 3.54E-01 | 1.08E+00 | 1.15E-01 | 1.08E+00 |
| ENSG000001 | ARMC8 | 7.37E-02 | 3.54E-01 | 1.07E+00 | 9.23E-02 | 1.07E+00 |
| ENSG000001 | HBG2 | 7.37E-02 | ? | 4.25E-01 | -1.24E+00 | -2.35E+00 |
| ENSG000002 | LINC00942 | 7.37E-02 | ? | 6.58E-01 | -6.04E-01 | -1.52E+00 |
| ENSG00000C | TTC7A | 7.38E-02 | 3.54E-01 | 1.08E+00 | 1.16E-01 | 1.08E+00 |
| ENSG000002 | CENPBD2P | 7.38E-02 | 3.54E-01 | 9.21E-01 | -1.19E-01 | -1.09E+00 |
| ENSG000002 | CHURC1 | 7.38E-02 | 3.54E-01 | 1.05E+00 | 7.35E-02 | 1.05E+00 |
| ENSG000002 | --- | 7.38E-02 | ? | 3.82E-01 | -1.39E+00 | -2.62E+00 |
| ENSG00000C | MRPS18A | 7.38E-02 | 3.54E-01 | 1.08E+00 | 1.16E-01 | 1.08E+00 |
| ENSG000001 | WDR81 | 7.39E-02 | 3.55E-01 | 1.14E+00 | 1.87E-01 | 1.14E+00 |
| ENSG000001 | SPOP | 7.39E-02 | 3.55E-01 | 1.07E+00 | 9.23E-02 | 1.07E+00 |
| ENSG000001 | IFNLR1 | 7.40E-02 | 3.55E-01 | 1.11E+00 | 1.55E-01 | 1.11E+00 |
| ENSG000001 | HARS2 | 7.41E-02 | 3.55E-01 | 1.06E+00 | 8.32E-02 | 1.06E+00 |
| ENSG000002 | --- | 7.42E-02 | 3.55E-01 | 1.45E+00 | 5.34E-01 | 1.45E+00 |
| ENSG000001 | GXYLT2 | 7.42E-02 | 3.55E-01 | 1.17E+00 | 2.29E-01 | 1.17E+00 |
| ENSG000001 | DCAF12 | 7.42E-02 | 3.56E-01 | 1.05E+00 | 6.53E-02 | 1.05E+00 |
| ENSG000001 | EIF2S3 | 7.43E-02 | 3.56E-01 | 1.06E+00 | 8.26E-02 | 1.06E+00 |
| ENSG000001 | KLHL5 | 7.43E-02 | 3.56E-01 | 1.10E+00 | 1.32E-01 | 1.10E+00 |
| ENSG000001 | COMMD3 | 7.43E-02 | 3.56E-01 | 1.08E+00 | 1.06E-01 | 1.08E+00 |
| ENSG000001 | GABARAPL1 | 7.43E-02 | 3.56E-01 | 1.09E+00 | 1.25E-01 | 1.09E+00 |
| ENSG000001 | MTHFD2L | 7.44E-02 | 3.56E-01 | 8.06E-01 | -3.11E-01 | -1.24E+00 |

| | | | | | | |
|---|---|---|---|---|---|---|
| ENSG000001 | NIN | 7.44E-02 | 3.56E-01 | 9.18E-01 | -1.24E-01 | -1.09E+00 |
| ENSG000002 | MIR193BHG | 7.45E-02 | 3.56E-01 | 8.72E-01 | -1.98E-01 | -1.15E+00 |
| ENSG000002 | GREP1 | 7.46E-02 | ? | 4.81E-01 | -1.06E+00 | -2.08E+00 |
| ENSG000001 | ZBTB47 | 7.46E-02 | 3.56E-01 | 1.11E+00 | 1.57E-01 | 1.11E+00 |
| ENSG000001 | WDR83OS | 7.47E-02 | 3.57E-01 | 1.16E+00 | 2.16E-01 | 1.16E+00 |
| ENSG000001 | MYEOV | 7.47E-02 | ? | 5.11E-01 | -9.70E-01 | -1.96E+00 |
| ENSG000002 | --- | 7.47E-02 | 3.57E-01 | 1.31E+00 | 3.87E-01 | 1.31E+00 |
| ENSG000001 | NAAA | 7.47E-02 | 3.57E-01 | 1.12E+00 | 1.62E-01 | 1.12E+00 |
| ENSG000001 | FBXO44 | 7.48E-02 | 3.57E-01 | 1.08E+00 | 1.11E-01 | 1.08E+00 |
| ENSG000001 | YWHAE | 7.48E-02 | 3.57E-01 | 9.47E-01 | -7.80E-02 | -1.06E+00 |
| ENSG000000 | FBXW11 | 7.49E-02 | 3.57E-01 | 1.05E+00 | 7.11E-02 | 1.05E+00 |
| ENSG000001 | MTRF1L | 7.49E-02 | 3.57E-01 | 9.37E-01 | -9.46E-02 | -1.07E+00 |
| ENSG000002 | --- | 7.49E-02 | 3.57E-01 | 1.16E+00 | 2.10E-01 | 1.16E+00 |
| ENSG000002 | POLG-DT | 7.49E-02 | ? | 1.98E+00 | 9.86E-01 | 1.98E+00 |
| ENSG000002 | --- | 7.49E-02 | ? | 4.93E-01 | -1.02E+00 | -2.03E+00 |
| ENSG000001 | SHC4 | 7.50E-02 | 3.57E-01 | 8.34E-01 | -2.62E-01 | -1.20E+00 |
| ENSG000001 | DCLRE1C | 7.50E-02 | 3.57E-01 | 9.15E-01 | -1.28E-01 | -1.09E+00 |
| ENSG000001 | MATN2 | 7.50E-02 | 3.57E-01 | 1.15E+00 | 2.05E-01 | 1.15E+00 |
| ENSG000002 | ST7-OT4 | 7.51E-02 | ? | 2.24E+00 | 1.17E+00 | 2.24E+00 |
| ENSG000001 | MARCKSL1 | 7.51E-02 | 3.57E-01 | 9.20E-01 | -1.21E-01 | -1.09E+00 |
| ENSG000001 | PRR14 | 7.51E-02 | 3.57E-01 | 9.22E-01 | -1.17E-01 | -1.08E+00 |
| ENSG000000 | CDK11A | 7.52E-02 | 3.58E-01 | 9.22E-01 | -1.18E-01 | -1.09E+00 |
| ENSG000002 | --- | 7.52E-02 | 3.58E-01 | 1.23E+00 | 2.93E-01 | 1.23E+00 |
| ENSG000002 | --- | 7.53E-02 | 3.58E-01 | 7.54E-01 | -4.08E-01 | -1.33E+00 |
| ENSG000001 | ZNF256 | 7.54E-02 | 3.58E-01 | 1.12E+00 | 1.65E-01 | 1.12E+00 |
| ENSG000001 | FASTK | 7.54E-02 | 3.58E-01 | 1.11E+00 | 1.53E-01 | 1.11E+00 |
| ENSG000001 | PRDX3 | 7.55E-02 | 3.58E-01 | 1.09E+00 | 1.23E-01 | 1.09E+00 |
| ENSG000001 | PSEN2 | 7.56E-02 | 3.59E-01 | 9.13E-01 | -1.31E-01 | -1.09E+00 |
| ENSG000001 | ELOB | 7.57E-02 | 3.59E-01 | 1.12E+00 | 1.63E-01 | 1.12E+00 |
| ENSG000002 | --- | 7.57E-02 | ? | 6.33E-01 | -6.60E-01 | -1.58E+00 |
| ENSG000001 | FICD | 7.58E-02 | 3.59E-01 | 8.62E-01 | -2.14E-01 | -1.16E+00 |
| ENSG000001 | TCAF1 | 7.58E-02 | 3.59E-01 | 1.07E+00 | 9.40E-02 | 1.07E+00 |
| ENSG000001 | TRMT10B | 7.58E-02 | 3.59E-01 | 8.84E-01 | -1.79E-01 | -1.13E+00 |
| ENSG000001 | CSMD3 | 7.58E-02 | 3.59E-01 | 9.18E-01 | -1.24E-01 | -1.09E+00 |
| ENSG000000 | TSG101 | 7.58E-02 | 3.59E-01 | 9.54E-01 | -6.80E-02 | -1.05E+00 |
| ENSG000002 | --- | 7.59E-02 | ? | 5.18E-01 | -9.50E-01 | -1.93E+00 |
| ENSG000001 | TMEM39B | 7.60E-02 | 3.60E-01 | 1.14E+00 | 1.84E-01 | 1.14E+00 |
| ENSG000001 | PRIMPOL | 7.60E-02 | 3.60E-01 | 1.11E+00 | 1.46E-01 | 1.11E+00 |
| ENSG000001 | PAPPA2 | 7.61E-02 | 3.60E-01 | 7.47E-01 | -4.22E-01 | -1.34E+00 |
| ENSG000002 | --- | 7.62E-02 | ? | 1.34E+00 | 4.19E-01 | 1.34E+00 |
| ENSG000002 | --- | 7.62E-02 | ? | 6.35E-01 | -6.56E-01 | -1.58E+00 |
| ENSG000001 | MEN1 | 7.62E-02 | 3.60E-01 | 1.06E+00 | 8.72E-02 | 1.06E+00 |
| ENSG000001 | C1orf56 | 7.62E-02 | 3.60E-01 | 8.46E-01 | -2.41E-01 | -1.18E+00 |
| ENSG000001 | POM121 | 7.63E-02 | 3.60E-01 | 9.53E-01 | -6.93E-02 | -1.05E+00 |
| ENSG000001 | HINT3 | 7.63E-02 | 3.60E-01 | 9.35E-01 | -9.67E-02 | -1.07E+00 |

| | | | | | | |
|---|---|---|---|---|---|---|
| ENSG000002 | --- | 7.63E-02 | ? | 1.51E+00 | 5.95E-01 | 1.51E+00 |
| ENSG000001 | EEF1A1P5 | 7.63E-02 | 3.61E-01 | 1.11E+00 | 1.49E-01 | 1.11E+00 |
| ENSG000001 | LGALS9C | 7.64E-02 | 3.61E-01 | 1.21E+00 | 2.77E-01 | 1.21E+00 |
| ENSG000001 | DROSHA | 7.64E-02 | 3.61E-01 | 9.21E-01 | -1.18E-01 | -1.09E+00 |
| ENSG00000C | MYO3B | 7.65E-02 | 3.61E-01 | 1.13E+00 | 1.81E-01 | 1.13E+00 |
| ENSG00000C | MAP4K4 | 7.65E-02 | 3.61E-01 | 9.37E-01 | -9.31E-02 | -1.07E+00 |
| ENSG000001 | GOLM1 | 7.65E-02 | 3.61E-01 | 9.41E-01 | -8.81E-02 | -1.06E+00 |
| ENSG00000C | EFNB1 | 7.66E-02 | 3.61E-01 | 1.10E+00 | 1.31E-01 | 1.10E+00 |
| ENSG00000C | GEMIN8 | 7.66E-02 | 3.61E-01 | 1.08E+00 | 1.15E-01 | 1.08E+00 |
| ENSG000001 | OSGIN1 | 7.66E-02 | 3.61E-01 | 8.85E-01 | -1.77E-01 | -1.13E+00 |
| ENSG000001 | PSD3 | 7.66E-02 | 3.61E-01 | 1.08E+00 | 1.08E-01 | 1.08E+00 |
| ENSG000002 | MGAM | 7.66E-02 | 3.61E-01 | 1.17E+00 | 2.23E-01 | 1.17E+00 |
| ENSG000001 | NOL9 | 7.66E-02 | 3.61E-01 | 1.11E+00 | 1.57E-01 | 1.11E+00 |
| ENSG000001 | GRK3 | 7.66E-02 | 3.61E-01 | 1.08E+00 | 1.16E-01 | 1.08E+00 |
| ENSG000001 | GAS1 | 7.67E-02 | ? | 2.16E+00 | 1.11E+00 | 2.16E+00 |
| ENSG000001 | PSMG3 | 7.67E-02 | 3.61E-01 | 1.12E+00 | 1.60E-01 | 1.12E+00 |
| ENSG000001 | STAMBP | 7.68E-02 | 3.61E-01 | 9.46E-01 | -7.93E-02 | -1.06E+00 |
| ENSG000001 | EFNB2 | 7.68E-02 | 3.61E-01 | 9.43E-01 | -8.40E-02 | -1.06E+00 |
| ENSG000002 | LINC00106 | 7.68E-02 | ? | 1.66E+00 | 7.28E-01 | 1.66E+00 |
| ENSG000001 | GLI3 | 7.68E-02 | 3.61E-01 | 1.11E+00 | 1.55E-01 | 1.11E+00 |
| ENSG000001 | PIGM | 7.69E-02 | 3.61E-01 | 1.07E+00 | 9.84E-02 | 1.07E+00 |
| ENSG000002 | FAM66C | 7.70E-02 | ? | 1.65E+00 | 7.21E-01 | 1.65E+00 |
| ENSG000002 | COL5A2 | 7.71E-02 | 3.62E-01 | 1.08E+00 | 1.13E-01 | 1.08E+00 |
| ENSG000001 | RASSF6 | 7.71E-02 | 3.62E-01 | 1.09E+00 | 1.26E-01 | 1.09E+00 |
| ENSG000002 | SLC35F6 | 7.72E-02 | 3.62E-01 | 1.06E+00 | 8.62E-02 | 1.06E+00 |
| ENSG000002 | VLDLR-AS1 | 7.72E-02 | ? | 6.09E-01 | -7.15E-01 | -1.64E+00 |
| ENSG000001 | KCTD1 | 7.73E-02 | 3.62E-01 | 1.05E+00 | 6.39E-02 | 1.05E+00 |
| ENSG000001 | HADHB | 7.73E-02 | 3.63E-01 | 1.05E+00 | 6.61E-02 | 1.05E+00 |
| ENSG000002 | MICE | 7.74E-02 | ? | 1.57E+00 | 6.47E-01 | 1.57E+00 |
| ENSG00000C | IGBP1 | 7.74E-02 | 3.63E-01 | 1.08E+00 | 1.07E-01 | 1.08E+00 |
| ENSG000002 | --- | 7.74E-02 | ? | 6.56E-01 | -6.09E-01 | -1.53E+00 |
| ENSG000001 | TMEM258 | 7.75E-02 | 3.63E-01 | 1.15E+00 | 1.98E-01 | 1.15E+00 |
| ENSG000002 | LINC00381 | 7.75E-02 | 3.63E-01 | 1.21E+00 | 2.74E-01 | 1.21E+00 |
| ENSG000001 | TRAF2 | 7.75E-02 | 3.63E-01 | 1.10E+00 | 1.35E-01 | 1.10E+00 |
| ENSG000001 | EBPL | 7.76E-02 | 3.63E-01 | 1.07E+00 | 9.90E-02 | 1.07E+00 |
| ENSG000001 | CMTM7 | 7.76E-02 | 3.63E-01 | 1.06E+00 | 9.02E-02 | 1.06E+00 |
| ENSG000001 | FANCL | 7.76E-02 | 3.63E-01 | 9.29E-01 | -1.07E-01 | -1.08E+00 |
| ENSG000002 | --- | 7.76E-02 | 3.63E-01 | 1.14E+00 | 1.94E-01 | 1.14E+00 |
| ENSG000002 | ATF6B | 7.76E-02 | 3.63E-01 | 1.05E+00 | 7.29E-02 | 1.05E+00 |
| ENSG000001 | ZNF431 | 7.77E-02 | 3.63E-01 | 9.11E-01 | -1.34E-01 | -1.10E+00 |
| ENSG00000C | MDH1 | 7.77E-02 | 3.63E-01 | 1.07E+00 | 9.60E-02 | 1.07E+00 |
| ENSG000001 | UNC5CL | 7.78E-02 | 3.63E-01 | 7.63E-01 | -3.90E-01 | -1.31E+00 |
| ENSG000001 | CCDC149 | 7.78E-02 | 3.63E-01 | 1.10E+00 | 1.32E-01 | 1.10E+00 |
| ENSG000001 | WSB1 | 7.78E-02 | 3.63E-01 | 8.91E-01 | -1.67E-01 | -1.12E+00 |
| ENSG000002 | LEPROT | 7.78E-02 | 3.63E-01 | 1.06E+00 | 7.79E-02 | 1.06E+00 |

| | | | | | | | |
|---|---|---|---|---|---|---|---|
| ENSG00000C | TP63 | 7.79E-02 | | 3.63E-01 | 1.06E+00 | 8.08E-02 | 1.06E+00 |
| ENSG000001 | UNC5D | 7.81E-02 | ? | | 5.99E-01 | -7.39E-01 | -1.67E+00 |
| ENSG00000C | TEAD2 | 7.81E-02 | | 3.64E-01 | 1.08E+00 | 1.09E-01 | 1.08E+00 |
| ENSG000001 | CMC4 | 7.81E-02 | | 3.64E-01 | 1.13E+00 | 1.81E-01 | 1.13E+00 |
| ENSG000001 | CUL5 | 7.82E-02 | | 3.64E-01 | 9.15E-01 | -1.28E-01 | -1.09E+00 |
| ENSG000001 | IMP3 | 7.82E-02 | | 3.64E-01 | 1.11E+00 | 1.49E-01 | 1.11E+00 |
| ENSG00000C | ACAA1 | 7.82E-02 | | 3.64E-01 | 1.09E+00 | 1.20E-01 | 1.09E+00 |
| ENSG000001 | NPL | 7.83E-02 | | 3.64E-01 | 1.19E+00 | 2.55E-01 | 1.19E+00 |
| ENSG000001 | SHROOM2 | 7.83E-02 | | 3.64E-01 | 1.09E+00 | 1.30E-01 | 1.09E+00 |
| ENSG000001 | S100A7A | 7.83E-02 | ? | | 6.00E-01 | -7.37E-01 | -1.67E+00 |
| ENSG000001 | MSX2 | 7.84E-02 | | 3.64E-01 | 1.29E+00 | 3.69E-01 | 1.29E+00 |
| ENSG000001 | SRSF2 | 7.85E-02 | | 3.65E-01 | 9.56E-01 | -6.50E-02 | -1.05E+00 |
| ENSG000002 | --- | 7.86E-02 | | 3.65E-01 | 1.27E+00 | 3.43E-01 | 1.27E+00 |
| ENSG000002 | --- | 7.86E-02 | | 3.65E-01 | 1.09E+00 | 1.20E-01 | 1.09E+00 |
| ENSG000001 | ACADSB | 7.86E-02 | | 3.65E-01 | 1.11E+00 | 1.49E-01 | 1.11E+00 |
| ENSG000001 | MSRB1 | 7.86E-02 | | 3.65E-01 | 1.11E+00 | 1.54E-01 | 1.11E+00 |
| ENSG000001 | BLMH | 7.87E-02 | | 3.65E-01 | 1.09E+00 | 1.27E-01 | 1.09E+00 |
| ENSG000001 | EEF1AKMT3 | 7.87E-02 | | 3.65E-01 | 8.94E-01 | -1.62E-01 | -1.12E+00 |
| ENSG000002 | --- | 7.88E-02 | ? | | 1.86E+00 | 8.97E-01 | 1.86E+00 |
| ENSG000001 | SLC31A1 | 7.88E-02 | | 3.65E-01 | 1.05E+00 | 7.20E-02 | 1.05E+00 |
| ENSG000001 | USP10 | 7.88E-02 | | 3.65E-01 | 1.04E+00 | 6.07E-02 | 1.04E+00 |
| ENSG000001 | SDC3 | 7.88E-02 | | 3.65E-01 | 1.08E+00 | 1.08E-01 | 1.08E+00 |
| ENSG000001 | GTF2E1 | 7.89E-02 | | 3.65E-01 | 1.10E+00 | 1.40E-01 | 1.10E+00 |
| ENSG000001 | FAM86B2 | 7.89E-02 | ? | | 1.51E+00 | 5.99E-01 | 1.51E+00 |
| ENSG000001 | SERF2 | 7.89E-02 | | 3.65E-01 | 1.04E+00 | 6.20E-02 | 1.04E+00 |
| ENSG000001 | NAGLU | 7.89E-02 | | 3.65E-01 | 1.19E+00 | 2.52E-01 | 1.19E+00 |
| ENSG000001 | ETS1 | 7.90E-02 | | 3.66E-01 | 9.46E-01 | -8.02E-02 | -1.06E+00 |
| ENSG000002 | --- | 7.90E-02 | ? | | 2.68E+00 | 1.42E+00 | 2.68E+00 |
| ENSG000001 | CCBE1 | 7.91E-02 | | 3.66E-01 | 8.98E-01 | -1.55E-01 | -1.11E+00 |
| ENSG000001 | ZNF675 | 7.92E-02 | | 3.66E-01 | 8.78E-01 | -1.88E-01 | -1.14E+00 |
| ENSG000001 | SLC35A1 | 7.92E-02 | | 3.66E-01 | 1.08E+00 | 1.10E-01 | 1.08E+00 |
| ENSG000001 | COMMD5 | 7.92E-02 | | 3.66E-01 | 1.16E+00 | 2.16E-01 | 1.16E+00 |
| ENSG000002 | H2AC18 | 7.92E-02 | | 3.66E-01 | 1.17E+00 | 2.25E-01 | 1.17E+00 |
| ENSG000001 | BNIP3P1 | 7.93E-02 | ? | | 1.72E+00 | 7.81E-01 | 1.72E+00 |
| ENSG000001 | SCN4B | 7.94E-02 | | 3.67E-01 | 1.16E+00 | 2.10E-01 | 1.16E+00 |
| ENSG00000C | CP | 7.95E-02 | ? | | 6.34E-01 | -6.57E-01 | -1.58E+00 |
| ENSG000001 | B3GNT3 | 7.96E-02 | | 3.68E-01 | 9.13E-01 | -1.31E-01 | -1.09E+00 |
| ENSG00000C | CHMP2B | 7.96E-02 | | 3.68E-01 | 9.20E-01 | -1.20E-01 | -1.09E+00 |
| ENSG00000C | MPND | 7.97E-02 | | 3.68E-01 | 1.20E+00 | 2.65E-01 | 1.20E+00 |
| ENSG000001 | RHPN2 | 7.97E-02 | | 3.68E-01 | 8.87E-01 | -1.73E-01 | -1.13E+00 |
| ENSG000002 | MBNL1-AS1 | 7.97E-02 | | 3.68E-01 | 1.21E+00 | 2.74E-01 | 1.21E+00 |
| ENSG000001 | MAP7D2 | 7.98E-02 | | 3.68E-01 | 1.28E+00 | 3.61E-01 | 1.28E+00 |
| ENSG000001 | NANP | 7.98E-02 | | 3.68E-01 | 9.13E-01 | -1.31E-01 | -1.10E+00 |
| ENSG000001 | DCAF11 | 7.98E-02 | | 3.68E-01 | 1.06E+00 | 8.15E-02 | 1.06E+00 |
| ENSG000002 | FAM201A | 7.98E-02 | | 3.68E-01 | 7.55E-01 | -4.05E-01 | -1.32E+00 |

| | | | | | | |
|---|---|---|---|---|---|---|
| ENSG000001 | CREB3L2 | 7.99E-02 | 3.68E-01 | 9.30E-01 | -1.05E-01 | -1.08E+00 |
| ENSG000001 | DENR | 7.99E-02 | 3.68E-01 | 9.47E-01 | -7.83E-02 | -1.06E+00 |
| ENSG000001 | TCF19 | 7.99E-02 | 3.68E-01 | 8.53E-01 | -2.29E-01 | -1.17E+00 |
| ENSG000001 | FAM117B | 7.99E-02 | 3.68E-01 | 1.09E+00 | 1.22E-01 | 1.09E+00 |
| ENSG000002 | --- | 8.01E-02 | ? | 5.44E-01 | -8.78E-01 | -1.84E+00 |
| ENSG000002 | --- | 8.01E-02 | ? | 1.60E+00 | 6.76E-01 | 1.60E+00 |
| ENSG000001 | HAPLN3 | 8.01E-02 | 3.69E-01 | 1.27E+00 | 3.40E-01 | 1.27E+00 |
| ENSG000002 | ERVV-1 | 8.01E-02 | ? | 1.75E+00 | 8.04E-01 | 1.75E+00 |
| ENSG000001 | NRDE2 | 8.01E-02 | 3.69E-01 | 1.08E+00 | 1.07E-01 | 1.08E+00 |
| ENSG000001 | SORT1 | 8.03E-02 | 3.69E-01 | 9.19E-01 | -1.21E-01 | -1.09E+00 |
| ENSG000001 | SPINT2 | 8.03E-02 | 3.69E-01 | 1.10E+00 | 1.38E-01 | 1.10E+00 |
| ENSG000001 | TAF4 | 8.04E-02 | 3.69E-01 | 9.21E-01 | -1.18E-01 | -1.09E+00 |
| ENSG000001 | PLOD2 | 8.04E-02 | 3.69E-01 | 9.12E-01 | -1.33E-01 | -1.10E+00 |
| ENSG000001 | CLDN14 | 8.04E-02 | ? | 6.43E-01 | -6.37E-01 | -1.56E+00 |
| ENSG000002 | --- | 8.05E-02 | 3.69E-01 | 1.11E+00 | 1.49E-01 | 1.11E+00 |
| ENSG000001 | SLC22A14 | 8.05E-02 | ? | 6.05E-01 | -7.24E-01 | -1.65E+00 |
| ENSG00000C | DKK3 | 8.05E-02 | 3.70E-01 | 1.07E+00 | 1.04E-01 | 1.07E+00 |
| ENSG000001 | SLC25A36 | 8.06E-02 | 3.70E-01 | 9.40E-01 | -8.87E-02 | -1.06E+00 |
| ENSG000001 | SMYD4 | 8.07E-02 | 3.70E-01 | 9.21E-01 | -1.18E-01 | -1.09E+00 |
| ENSG000001 | MTMR10 | 8.07E-02 | 3.70E-01 | 9.42E-01 | -8.67E-02 | -1.06E+00 |
| ENSG000001 | ATP6V0E1 | 8.07E-02 | 3.70E-01 | 1.05E+00 | 7.12E-02 | 1.05E+00 |
| ENSG000001 | ZNF483 | 8.08E-02 | 3.70E-01 | 8.46E-01 | -2.41E-01 | -1.18E+00 |
| ENSG000002 | LINC01123 | 8.08E-02 | 3.70E-01 | 1.13E+00 | 1.75E-01 | 1.13E+00 |
| ENSG00000C | HDHD5 | 8.08E-02 | 3.70E-01 | 1.15E+00 | 2.04E-01 | 1.15E+00 |
| ENSG000001 | XRN1 | 8.09E-02 | 3.70E-01 | 9.28E-01 | -1.08E-01 | -1.08E+00 |
| ENSG000002 | TALAM1 | 8.09E-02 | 3.70E-01 | 8.31E-01 | -2.67E-01 | -1.20E+00 |
| ENSG000001 | DYNC2I2 | 8.09E-02 | 3.70E-01 | 1.14E+00 | 1.89E-01 | 1.14E+00 |
| ENSG000001 | NANOS1 | 8.09E-02 | 3.70E-01 | 1.12E+00 | 1.61E-01 | 1.12E+00 |
| ENSG00000C | RIMS1 | 8.10E-02 | ? | 4.75E-01 | -1.07E+00 | -2.10E+00 |
| ENSG000002 | PITX1-AS1 | 8.11E-02 | 3.70E-01 | 1.25E+00 | 3.18E-01 | 1.25E+00 |
| ENSG000001 | RPF1 | 8.11E-02 | 3.70E-01 | 9.47E-01 | -7.88E-02 | -1.06E+00 |
| ENSG00000C | MKNK2 | 8.11E-02 | 3.70E-01 | 1.11E+00 | 1.48E-01 | 1.11E+00 |
| ENSG000001 | TBRG4 | 8.11E-02 | 3.71E-01 | 1.08E+00 | 1.11E-01 | 1.08E+00 |
| ENSG000002 | --- | 8.12E-02 | ? | 5.41E-01 | -8.87E-01 | -1.85E+00 |
| ENSG000002 | NTF4 | 8.13E-02 | 3.71E-01 | 1.14E+00 | 1.94E-01 | 1.14E+00 |
| ENSG000002 | --- | 8.13E-02 | ? | 2.11E+00 | 1.08E+00 | 2.11E+00 |
| ENSG000002 | --- | 8.13E-02 | ? | 5.34E-01 | -9.04E-01 | -1.87E+00 |
| ENSG000001 | DDI2 | 8.13E-02 | 3.71E-01 | 1.07E+00 | 9.16E-02 | 1.07E+00 |
| ENSG000001 | SNX17 | 8.13E-02 | 3.71E-01 | 1.12E+00 | 1.67E-01 | 1.12E+00 |
| ENSG000001 | LEPROTL1 | 8.13E-02 | 3.71E-01 | 1.06E+00 | 7.98E-02 | 1.06E+00 |
| ENSG000001 | SRSF11 | 8.14E-02 | 3.71E-01 | 9.33E-01 | -1.00E-01 | -1.07E+00 |
| ENSG000001 | GLE1 | 8.14E-02 | 3.71E-01 | 9.58E-01 | -6.20E-02 | -1.04E+00 |
| ENSG000001 | FANCD2 | 8.15E-02 | 3.71E-01 | 8.70E-01 | -2.01E-01 | -1.15E+00 |
| ENSG000002 | ZNF731P | 8.15E-02 | 3.71E-01 | 8.82E-01 | -1.82E-01 | -1.13E+00 |
| ENSG000001 | PANK4 | 8.16E-02 | 3.71E-01 | 1.09E+00 | 1.18E-01 | 1.09E+00 |

| | | | | | | |
|---|---|---|---|---|---|---|
| ENSG00000C | MSL3 | 8.16E-02 | 3.71E-01 | 9.34E-01 | -9.89E-02 | -1.07E+00 |
| ENSG00000C | CX3CL1 | 8.16E-02 | 3.71E-01 | 1.35E+00 | 4.29E-01 | 1.35E+00 |
| ENSG000001 | PRRC2C | 8.16E-02 | 3.71E-01 | 9.08E-01 | -1.39E-01 | -1.10E+00 |
| ENSG000001 | NKX2-1 | 8.16E-02 | 3.71E-01 | 1.13E+00 | 1.78E-01 | 1.13E+00 |
| ENSG000002 | --- | 8.16E-02 | ? | 6.21E-01 | -6.88E-01 | -1.61E+00 |
| ENSG000001 | PEPD | 8.17E-02 | 3.71E-01 | 1.11E+00 | 1.47E-01 | 1.11E+00 |
| ENSG000002 | RIPK2-DT | 8.17E-02 | 3.71E-01 | 8.62E-01 | -2.15E-01 | -1.16E+00 |
| ENSG000001 | CMTM3 | 8.17E-02 | 3.71E-01 | 1.11E+00 | 1.51E-01 | 1.11E+00 |
| ENSG000001 | RUNDC3A | 8.17E-02 | 3.71E-01 | 1.19E+00 | 2.50E-01 | 1.19E+00 |
| ENSG000002 | ITPRID2-DT | 8.18E-02 | ? | 1.80E+00 | 8.46E-01 | 1.80E+00 |
| ENSG000001 | ZSCAN20 | 8.18E-02 | 3.71E-01 | 9.30E-01 | -1.04E-01 | -1.07E+00 |
| ENSG000001 | RPS21 | 8.18E-02 | 3.71E-01 | 1.12E+00 | 1.66E-01 | 1.12E+00 |
| ENSG000001 | CTNNBIP1 | 8.19E-02 | 3.72E-01 | 1.09E+00 | 1.26E-01 | 1.09E+00 |
| ENSG000001 | DNHD1 | 8.20E-02 | 3.72E-01 | 8.83E-01 | -1.79E-01 | -1.13E+00 |
| ENSG000001 | IFT22 | 8.20E-02 | 3.72E-01 | 1.10E+00 | 1.41E-01 | 1.10E+00 |
| ENSG000002 | FADS3 | 8.20E-02 | 3.72E-01 | 1.09E+00 | 1.29E-01 | 1.09E+00 |
| ENSG000002 | --- | 8.21E-02 | 3.72E-01 | 8.47E-01 | -2.40E-01 | -1.18E+00 |
| ENSG000001 | OBSL1 | 8.22E-02 | 3.72E-01 | 1.13E+00 | 1.82E-01 | 1.13E+00 |
| ENSG000001 | SBSN | 8.22E-02 | 3.72E-01 | 1.16E+00 | 2.16E-01 | 1.16E+00 |
| ENSG000001 | TIMMDC1 | 8.22E-02 | 3.72E-01 | 1.05E+00 | 7.17E-02 | 1.05E+00 |
| ENSG000002 | HCP5 | 8.23E-02 | 3.72E-01 | 1.25E+00 | 3.18E-01 | 1.25E+00 |
| ENSG000001 | SIX1 | 8.23E-02 | 3.73E-01 | 8.96E-01 | -1.58E-01 | -1.12E+00 |
| ENSG000002 | --- | 8.23E-02 | ? | 1.49E+00 | 5.73E-01 | 1.49E+00 |
| ENSG000001 | LAMC1 | 8.24E-02 | 3.73E-01 | 1.06E+00 | 7.99E-02 | 1.06E+00 |
| ENSG000001 | ANKRD11 | 8.24E-02 | 3.73E-01 | 9.44E-01 | -8.27E-02 | -1.06E+00 |
| ENSG000002 | --- | 8.24E-02 | 3.73E-01 | 1.14E+00 | 1.85E-01 | 1.14E+00 |
| ENSG000001 | IRX2-DT | 8.24E-02 | 3.73E-01 | 1.16E+00 | 2.10E-01 | 1.16E+00 |
| ENSG000001 | TMEM105 | 8.24E-02 | ? | 7.08E-01 | -4.97E-01 | -1.41E+00 |
| ENSG000001 | PDE4DIP | 8.24E-02 | 3.73E-01 | 9.36E-01 | -9.47E-02 | -1.07E+00 |
| ENSG000002 | --- | 8.25E-02 | ? | 5.15E+00 | 2.37E+00 | 5.15E+00 |
| ENSG000001 | SYP | 8.26E-02 | ? | 7.06E-01 | -5.03E-01 | -1.42E+00 |
| ENSG000002 | --- | 8.27E-02 | ? | 5.04E-01 | -9.89E-01 | -1.99E+00 |
| ENSG000001 | ZFP91 | 8.27E-02 | 3.73E-01 | 9.34E-01 | -9.79E-02 | -1.07E+00 |
| ENSG000001 | MRPS30 | 8.27E-02 | 3.73E-01 | 9.37E-01 | -9.35E-02 | -1.07E+00 |
| ENSG00000C | CDC27 | 8.27E-02 | 3.73E-01 | 9.36E-01 | -9.54E-02 | -1.07E+00 |
| ENSG000001 | ARHGEF40 | 8.27E-02 | 3.73E-01 | 1.09E+00 | 1.24E-01 | 1.09E+00 |
| ENSG000001 | DOCK1 | 8.27E-02 | 3.73E-01 | 9.55E-01 | -6.62E-02 | -1.05E+00 |
| ENSG00000C | CAPN1 | 8.27E-02 | 3.73E-01 | 1.09E+00 | 1.18E-01 | 1.09E+00 |
| ENSG000002 | --- | 8.28E-02 | ? | 2.09E+00 | 1.06E+00 | 2.09E+00 |
| ENSG000002 | --- | 8.28E-02 | 3.74E-01 | 1.23E+00 | 2.99E-01 | 1.23E+00 |
| ENSG000002 | --- | 8.29E-02 | ? | 4.89E-01 | -1.03E+00 | -2.05E+00 |
| ENSG000002 | --- | 8.30E-02 | 3.74E-01 | 1.22E+00 | 2.91E-01 | 1.22E+00 |
| ENSG000001 | TANK | 8.30E-02 | 3.74E-01 | 9.32E-01 | -1.01E-01 | -1.07E+00 |
| ENSG000001 | CAMKMT | 8.30E-02 | 3.74E-01 | 8.64E-01 | -2.11E-01 | -1.16E+00 |
| ENSG000001 | FANCE | 8.31E-02 | 3.74E-01 | 1.22E+00 | 2.87E-01 | 1.22E+00 |

| | | | | | | |
|---|---|---|---|---|---|---|
| ENSG00000C | GRIPAP1 | 8.31E-02 | 3.74E-01 | 9.20E-01 | -1.20E-01 | -1.09E+00 |
| ENSG000001 | RNF220 | 8.31E-02 | 3.74E-01 | 1.07E+00 | 9.32E-02 | 1.07E+00 |
| ENSG000002 | PGAM1P8 | 8.32E-02 | 3.74E-01 | 8.01E-01 | -3.20E-01 | -1.25E+00 |
| ENSG00000C | SEC22C | 8.34E-02 | 3.75E-01 | 1.05E+00 | 7.14E-02 | 1.05E+00 |
| ENSG000001 | FAM81A | 8.35E-02 | 3.75E-01 | 1.16E+00 | 2.09E-01 | 1.16E+00 |
| ENSG000001 | DSC2 | 8.35E-02 | 3.75E-01 | 1.10E+00 | 1.42E-01 | 1.10E+00 |
| ENSG000001 | NEIL3 | 8.35E-02 | ? | 6.92E-01 | -5.32E-01 | -1.45E+00 |
| ENSG000001 | DNASE2 | 8.35E-02 | 3.75E-01 | 1.12E+00 | 1.59E-01 | 1.12E+00 |
| ENSG000001 | BHLHE40 | 8.35E-02 | 3.75E-01 | 9.44E-01 | -8.37E-02 | -1.06E+00 |
| ENSG000001 | NDUFA6 | 8.35E-02 | 3.75E-01 | 1.07E+00 | 9.44E-02 | 1.07E+00 |
| ENSG000001 | PIK3C2B | 8.36E-02 | 3.75E-01 | 1.13E+00 | 1.80E-01 | 1.13E+00 |
| ENSG000002 | LTB | 8.36E-02 | ? | 1.47E+00 | 5.61E-01 | 1.47E+00 |
| ENSG000001 | TMEM120B | 8.38E-02 | 3.76E-01 | 1.08E+00 | 1.16E-01 | 1.08E+00 |
| ENSG000001 | FAM156B | 8.38E-02 | 3.76E-01 | 1.65E+00 | 7.22E-01 | 1.65E+00 |
| ENSG000001 | P2RY1 | 8.39E-02 | 3.76E-01 | 1.10E+00 | 1.42E-01 | 1.10E+00 |
| ENSG000002 | KLLN | 8.39E-02 | 3.76E-01 | 1.24E+00 | 3.07E-01 | 1.24E+00 |
| ENSG00000C | P4HA2 | 8.40E-02 | 3.76E-01 | 9.39E-01 | -9.04E-02 | -1.06E+00 |
| ENSG000001 | RAB9A | 8.40E-02 | 3.76E-01 | 9.25E-01 | -1.13E-01 | -1.08E+00 |
| ENSG00000C | ADIPOR2 | 8.41E-02 | 3.77E-01 | 9.58E-01 | -6.17E-02 | -1.04E+00 |
| ENSG000001 | PRDM2 | 8.41E-02 | 3.77E-01 | 9.27E-01 | -1.10E-01 | -1.08E+00 |
| ENSG000001 | CDC42SE1 | 8.42E-02 | 3.77E-01 | 1.06E+00 | 8.45E-02 | 1.06E+00 |
| ENSG000001 | SEPHS2 | 8.43E-02 | 3.77E-01 | 1.06E+00 | 8.07E-02 | 1.06E+00 |
| ENSG000002 | --- | 8.44E-02 | ? | 1.59E+00 | 6.70E-01 | 1.59E+00 |
| ENSG000002 | HLA-F | 8.44E-02 | 3.77E-01 | 1.13E+00 | 1.76E-01 | 1.13E+00 |
| ENSG000001 | CIR1 | 8.44E-02 | 3.77E-01 | 9.16E-01 | -1.27E-01 | -1.09E+00 |
| ENSG000002 | RNF103 | 8.44E-02 | 3.77E-01 | 1.06E+00 | 8.57E-02 | 1.06E+00 |
| ENSG000001 | PFKFB2 | 8.44E-02 | 3.77E-01 | 1.11E+00 | 1.47E-01 | 1.11E+00 |
| ENSG000001 | GPD1L | 8.44E-02 | 3.77E-01 | 1.09E+00 | 1.28E-01 | 1.09E+00 |
| ENSG000001 | PRKRA | 8.44E-02 | 3.77E-01 | 9.39E-01 | -9.06E-02 | -1.06E+00 |
| ENSG000001 | PHLDB2 | 8.44E-02 | 3.77E-01 | 1.07E+00 | 9.33E-02 | 1.07E+00 |
| ENSG000001 | POP5 | 8.44E-02 | 3.77E-01 | 1.10E+00 | 1.34E-01 | 1.10E+00 |
| ENSG000001 | H2AC25 | 8.45E-02 | 3.77E-01 | 1.18E+00 | 2.35E-01 | 1.18E+00 |
| ENSG000001 | FOXJ3 | 8.46E-02 | 3.77E-01 | 9.54E-01 | -6.72E-02 | -1.05E+00 |
| ENSG000002 | --- | 8.46E-02 | 3.77E-01 | 7.79E-01 | -3.61E-01 | -1.28E+00 |
| ENSG000001 | SKA3 | 8.46E-02 | 3.77E-01 | 7.79E-01 | -3.60E-01 | -1.28E+00 |
| ENSG00000C | TREM2 | 8.48E-02 | 3.78E-01 | 1.17E+00 | 2.29E-01 | 1.17E+00 |
| ENSG000001 | CKAP4 | 8.48E-02 | 3.78E-01 | 1.07E+00 | 1.00E-01 | 1.07E+00 |
| ENSG000001 | SLC39A13 | 8.49E-02 | 3.78E-01 | 1.11E+00 | 1.46E-01 | 1.11E+00 |
| ENSG000001 | ESRP2 | 8.49E-02 | 3.78E-01 | 1.07E+00 | 9.30E-02 | 1.07E+00 |
| ENSG000002 | --- | 8.50E-02 | ? | 1.51E+00 | 5.99E-01 | 1.51E+00 |
| ENSG000002 | LINC02246 | 8.50E-02 | ? | 2.06E+00 | 1.04E+00 | 2.06E+00 |
| ENSG000001 | POLR1D | 8.51E-02 | 3.79E-01 | 1.04E+00 | 5.96E-02 | 1.04E+00 |
| ENSG000001 | PTPRF | 8.51E-02 | 3.79E-01 | 1.09E+00 | 1.19E-01 | 1.09E+00 |
| ENSG00000C | SYT1 | 8.51E-02 | 3.79E-01 | 8.78E-01 | -1.87E-01 | -1.14E+00 |
| ENSG000001 | ACSF2 | 8.52E-02 | 3.79E-01 | 1.08E+00 | 1.13E-01 | 1.08E+00 |

| | | | | | | |
|---|---|---|---|---|---|---|
| ENSG000000 | TRAM1 | 8.52E-02 | 3.79E-01 | 9.29E-01 | -1.07E-01 | -1.08E+00 |
| ENSG000000 | CCNK | 8.53E-02 | 3.79E-01 | 9.41E-01 | -8.77E-02 | -1.06E+00 |
| ENSG000001 | NRARP | 8.53E-02 | 3.79E-01 | 1.19E+00 | 2.54E-01 | 1.19E+00 |
| ENSG000001 | DLGAP1-AS1 | 8.53E-02 | 3.79E-01 | 1.12E+00 | 1.65E-01 | 1.12E+00 |
| ENSG000000 | BAZ2A | 8.54E-02 | 3.79E-01 | 1.04E+00 | 4.98E-02 | 1.04E+00 |
| ENSG000001 | AKR1C1 | 8.55E-02 | 3.79E-01 | 8.31E-01 | -2.67E-01 | -1.20E+00 |
| ENSG000001 | SPOUT1 | 8.55E-02 | 3.79E-01 | 1.08E+00 | 1.11E-01 | 1.08E+00 |
| ENSG000001 | KCNE5 | 8.55E-02 | ? | 6.65E-01 | -5.89E-01 | -1.50E+00 |
| ENSG000001 | PSME2 | 8.55E-02 | 3.79E-01 | 1.07E+00 | 9.32E-02 | 1.07E+00 |
| ENSG000001 | FBXO38 | 8.55E-02 | 3.79E-01 | 9.43E-01 | -8.40E-02 | -1.06E+00 |
| ENSG000001 | PPA2 | 8.56E-02 | 3.79E-01 | 1.07E+00 | 1.03E-01 | 1.07E+00 |
| ENSG000001 | RGS3 | 8.56E-02 | 3.79E-01 | 1.09E+00 | 1.28E-01 | 1.09E+00 |
| ENSG000001 | NFATC4 | 8.57E-02 | 3.80E-01 | 1.14E+00 | 1.84E-01 | 1.14E+00 |
| ENSG000001 | RAB3B | 8.57E-02 | 3.80E-01 | 8.78E-01 | -1.88E-01 | -1.14E+00 |
| ENSG000001 | MAPK13 | 8.59E-02 | 3.80E-01 | 9.39E-01 | -9.08E-02 | -1.06E+00 |
| ENSG000001 | ZNF35 | 8.59E-02 | 3.80E-01 | 9.19E-01 | -1.23E-01 | -1.09E+00 |
| ENSG000001 | ARHGEF26 | 8.59E-02 | 3.80E-01 | 8.29E-01 | -2.71E-01 | -1.21E+00 |
| ENSG000001 | RPL10 | 8.59E-02 | 3.80E-01 | 1.14E+00 | 1.85E-01 | 1.14E+00 |
| ENSG000002 | --- | 8.60E-02 | ? | 6.61E-01 | -5.98E-01 | -1.51E+00 |
| ENSG000002 | --- | 8.60E-02 | 3.80E-01 | 7.43E-01 | -4.29E-01 | -1.35E+00 |
| ENSG000001 | PATZ1 | 8.61E-02 | 3.80E-01 | 1.13E+00 | 1.75E-01 | 1.13E+00 |
| ENSG000002 | NRAV | 8.61E-02 | 3.80E-01 | 1.11E+00 | 1.44E-01 | 1.11E+00 |
| ENSG000002 | AP5B1 | 8.61E-02 | 3.80E-01 | 1.08E+00 | 1.12E-01 | 1.08E+00 |
| ENSG000001 | SPATS2L | 8.61E-02 | 3.80E-01 | 9.47E-01 | -7.85E-02 | -1.06E+00 |
| ENSG000002 | --- | 8.61E-02 | ? | 4.47E-01 | -1.16E+00 | -2.24E+00 |
| ENSG000001 | ERRFI1 | 8.62E-02 | 3.80E-01 | 9.22E-01 | -1.17E-01 | -1.08E+00 |
| ENSG000000 | NOS1 | 8.62E-02 | 3.80E-01 | 1.17E+00 | 2.27E-01 | 1.17E+00 |
| ENSG000001 | LMBRD1 | 8.62E-02 | 3.80E-01 | 1.07E+00 | 9.51E-02 | 1.07E+00 |
| ENSG000002 | RPL26P19 | 8.63E-02 | 3.80E-01 | 1.19E+00 | 2.47E-01 | 1.19E+00 |
| ENSG000002 | ZNF10 | 8.63E-02 | 3.80E-01 | 8.56E-01 | -2.24E-01 | -1.17E+00 |
| ENSG000001 | CALHM2 | 8.63E-02 | 3.80E-01 | 8.23E-01 | -2.81E-01 | -1.22E+00 |
| ENSG000001 | ZNF644 | 8.63E-02 | 3.80E-01 | 1.08E+00 | 1.09E-01 | 1.08E+00 |
| ENSG000001 | ARPP19 | 8.64E-02 | 3.80E-01 | 9.61E-01 | -5.67E-02 | -1.04E+00 |
| ENSG000001 | S100A14 | 8.64E-02 | 3.80E-01 | 1.04E+00 | 6.00E-02 | 1.04E+00 |
| ENSG000000 | PARP3 | 8.64E-02 | 3.81E-01 | 1.10E+00 | 1.39E-01 | 1.10E+00 |
| ENSG000002 | --- | 8.65E-02 | 3.81E-01 | 8.22E-01 | -2.82E-01 | -1.22E+00 |
| ENSG000002 | RBMS2P1 | 8.66E-02 | ? | 7.35E-01 | -4.45E-01 | -1.36E+00 |
| ENSG000001 | CSRP1 | 8.66E-02 | 3.81E-01 | 1.06E+00 | 8.68E-02 | 1.06E+00 |
| ENSG000002 | LIFR-AS1 | 8.66E-02 | ? | 4.74E-01 | -1.08E+00 | -2.11E+00 |
| ENSG000001 | CARM1 | 8.66E-02 | 3.81E-01 | 1.08E+00 | 1.05E-01 | 1.08E+00 |
| ENSG000001 | SH3GL1 | 8.67E-02 | 3.81E-01 | 1.08E+00 | 1.12E-01 | 1.08E+00 |
| ENSG000001 | ROBO3 | 8.67E-02 | 3.81E-01 | 8.94E-01 | -1.62E-01 | -1.12E+00 |
| ENSG000001 | UHRF2 | 8.67E-02 | 3.81E-01 | 9.36E-01 | -9.54E-02 | -1.07E+00 |
| ENSG000001 | SKIC8 | 8.68E-02 | 3.81E-01 | 1.07E+00 | 9.52E-02 | 1.07E+00 |
| ENSG000000 | TMEM14A | 8.68E-02 | 3.81E-01 | 1.08E+00 | 1.09E-01 | 1.08E+00 |

| | | | | | | |
|---|---|---|---|---|---|---|
| ENSG000001 | TFCP2L1 | 8.68E-02 | 3.81E-01 | 9.30E-01 | -1.05E-01 | -1.08E+00 |
| ENSG000001 | RIMS3 | 8.68E-02 | 3.81E-01 | 8.51E-01 | -2.33E-01 | -1.17E+00 |
| ENSG000001 | URB1 | 8.68E-02 | 3.81E-01 | 1.06E+00 | 8.09E-02 | 1.06E+00 |
| ENSG000001 | SMC2 | 8.69E-02 | 3.81E-01 | 9.23E-01 | -1.15E-01 | -1.08E+00 |
| ENSG000002 | CNEP1R1 | 8.69E-02 | 3.81E-01 | 9.33E-01 | -1.01E-01 | -1.07E+00 |
| ENSG000001 | ZNF414 | 8.69E-02 | 3.81E-01 | 1.21E+00 | 2.70E-01 | 1.21E+00 |
| ENSG000001 | LBX2 | 8.69E-02 | ? | 1.72E+00 | 7.85E-01 | 1.72E+00 |
| ENSG000001 | SF3B6 | 8.70E-02 | 3.81E-01 | 9.58E-01 | -6.26E-02 | -1.04E+00 |
| ENSG000001 | ZC3H12A | 8.70E-02 | 3.81E-01 | 1.10E+00 | 1.42E-01 | 1.10E+00 |
| ENSG000001 | PITPNC1 | 8.70E-02 | 3.81E-01 | 9.26E-01 | -1.10E-01 | -1.08E+00 |
| ENSG000001 | RFT1 | 8.70E-02 | 3.81E-01 | 1.08E+00 | 1.06E-01 | 1.08E+00 |
| ENSG00000C | REV3L | 8.71E-02 | 3.81E-01 | 1.09E+00 | 1.19E-01 | 1.09E+00 |
| ENSG000002 | BTBD18 | 8.71E-02 | ? | 3.82E-01 | -1.39E+00 | -2.62E+00 |
| ENSG00000C | MED24 | 8.71E-02 | 3.81E-01 | 1.09E+00 | 1.18E-01 | 1.09E+00 |
| ENSG000001 | ATG9A | 8.72E-02 | 3.81E-01 | 1.07E+00 | 1.00E-01 | 1.07E+00 |
| ENSG000002 | LINC03014 | 8.72E-02 | ? | 6.92E-01 | -5.31E-01 | -1.44E+00 |
| ENSG000001 | TOMM7 | 8.73E-02 | 3.82E-01 | 1.15E+00 | 2.04E-01 | 1.15E+00 |
| ENSG000001 | PUS3 | 8.73E-02 | 3.82E-01 | 1.08E+00 | 1.11E-01 | 1.08E+00 |
| ENSG000001 | GIT2 | 8.74E-02 | 3.82E-01 | 9.47E-01 | -7.88E-02 | -1.06E+00 |
| ENSG000001 | AURKC | 8.74E-02 | ? | 7.10E-01 | -4.95E-01 | -1.41E+00 |
| ENSG000001 | KHK | 8.74E-02 | 3.82E-01 | 7.84E-01 | -3.50E-01 | -1.27E+00 |
| ENSG00000C | DNAJA2 | 8.74E-02 | 3.82E-01 | 9.56E-01 | -6.49E-02 | -1.05E+00 |
| ENSG000001 | SNAI1 | 8.75E-02 | ? | 6.02E-01 | -7.33E-01 | -1.66E+00 |
| ENSG000001 | FZD2 | 8.75E-02 | 3.82E-01 | 1.24E+00 | 3.10E-01 | 1.24E+00 |
| ENSG000001 | FKBP3 | 8.76E-02 | 3.82E-01 | 9.47E-01 | -7.83E-02 | -1.06E+00 |
| ENSG000001 | ERAL1 | 8.76E-02 | 3.82E-01 | 1.09E+00 | 1.21E-01 | 1.09E+00 |
| ENSG000001 | RNF216P1 | 8.76E-02 | 3.82E-01 | 1.07E+00 | 1.03E-01 | 1.07E+00 |
| ENSG000001 | RPL7 | 8.76E-02 | 3.82E-01 | 1.05E+00 | 6.87E-02 | 1.05E+00 |
| ENSG000001 | GRHPR | 8.77E-02 | 3.83E-01 | 1.08E+00 | 1.15E-01 | 1.08E+00 |
| ENSG000002 | --- | 8.78E-02 | 3.83E-01 | 8.50E-01 | -2.35E-01 | -1.18E+00 |
| ENSG000001 | ARGLU1 | 8.78E-02 | 3.83E-01 | 9.06E-01 | -1.42E-01 | -1.10E+00 |
| ENSG00000C | MAOB | 8.79E-02 | ? | 7.59E-01 | -3.98E-01 | -1.32E+00 |
| ENSG000001 | MTCH1 | 8.79E-02 | 3.83E-01 | 1.06E+00 | 8.45E-02 | 1.06E+00 |
| ENSG00000C | DOP1A | 8.79E-02 | 3.83E-01 | 1.06E+00 | 8.78E-02 | 1.06E+00 |
| ENSG000001 | ATMIN | 8.79E-02 | 3.83E-01 | 9.50E-01 | -7.39E-02 | -1.05E+00 |
| ENSG000001 | GARS1 | 8.79E-02 | 3.83E-01 | 9.51E-01 | -7.21E-02 | -1.05E+00 |
| ENSG000001 | CHCHD3 | 8.81E-02 | 3.83E-01 | 9.34E-01 | -9.85E-02 | -1.07E+00 |
| ENSG000001 | CNNM1 | 8.81E-02 | ? | 1.54E+00 | 6.21E-01 | 1.54E+00 |
| ENSG00000C | CPS1 | 8.81E-02 | 3.83E-01 | 9.02E-01 | -1.48E-01 | -1.11E+00 |
| ENSG000001 | LAMTOR1 | 8.81E-02 | 3.83E-01 | 1.10E+00 | 1.33E-01 | 1.10E+00 |
| ENSG000001 | TBX1 | 8.82E-02 | 3.83E-01 | 8.78E-01 | -1.88E-01 | -1.14E+00 |
| ENSG000001 | MRPL57 | 8.83E-02 | 3.83E-01 | 1.08E+00 | 1.10E-01 | 1.08E+00 |
| ENSG000001 | AP3S2 | 8.83E-02 | 3.83E-01 | 9.35E-01 | -9.76E-02 | -1.07E+00 |
| ENSG000001 | UROD | 8.83E-02 | 3.83E-01 | 1.06E+00 | 8.81E-02 | 1.06E+00 |
| ENSG00000C | ZRANB1 | 8.83E-02 | 3.83E-01 | 9.49E-01 | -7.50E-02 | -1.05E+00 |

| | | | | | | |
|---|---|---|---|---|---|---|
| ENSG000001 | NAT10 | 8.84E-02 | 3.83E-01 | 1.05E+00 | 7.16E-02 | 1.05E+00 |
| ENSG000002 | TAS2R6P | 8.84E-02 | ? | 5.98E-01 | -7.42E-01 | -1.67E+00 |
| ENSG000001 | KCTD17 | 8.84E-02 | 3.84E-01 | 1.11E+00 | 1.52E-01 | 1.11E+00 |
| ENSG000002 | --- | 8.84E-02 | 3.84E-01 | 1.32E+00 | 4.00E-01 | 1.32E+00 |
| ENSG000002 | --- | 8.86E-02 | 3.84E-01 | 1.20E+00 | 2.64E-01 | 1.20E+00 |
| ENSG000002 | SERPINB5 | 8.86E-02 | 3.84E-01 | 1.04E+00 | 5.09E-02 | 1.04E+00 |
| ENSG000002 | MSANTD3-TI | 8.86E-02 | 3.84E-01 | 8.58E-01 | -2.21E-01 | -1.17E+00 |
| ENSG000001 | ABI1 | 8.87E-02 | 3.84E-01 | 1.05E+00 | 7.45E-02 | 1.05E+00 |
| ENSG000002 | --- | 8.87E-02 | ? | 2.54E+00 | 1.34E+00 | 2.54E+00 |
| ENSG000002 | --- | 8.87E-02 | 3.84E-01 | 8.51E-01 | -2.33E-01 | -1.18E+00 |
| ENSG000001 | NAA50 | 8.87E-02 | 3.84E-01 | 1.05E+00 | 6.81E-02 | 1.05E+00 |
| ENSG000001 | COL6A1 | 8.87E-02 | 3.84E-01 | 1.09E+00 | 1.26E-01 | 1.09E+00 |
| ENSG000001 | CUL9 | 8.88E-02 | 3.84E-01 | 1.10E+00 | 1.42E-01 | 1.10E+00 |
| ENSG000001 | CFAP20DC | 8.88E-02 | 3.84E-01 | 8.86E-01 | -1.75E-01 | -1.13E+00 |
| ENSG000001 | VOPP1 | 8.88E-02 | 3.84E-01 | 1.07E+00 | 1.01E-01 | 1.07E+00 |
| ENSG000001 | GNA12 | 8.88E-02 | 3.84E-01 | 1.06E+00 | 8.70E-02 | 1.06E+00 |
| ENSG000002 | PLA2G4B | 8.89E-02 | 3.84E-01 | 1.38E+00 | 4.69E-01 | 1.38E+00 |
| ENSG000001 | NAA38 | 8.90E-02 | 3.84E-01 | 1.11E+00 | 1.54E-01 | 1.11E+00 |
| ENSG000001 | AIG1 | 8.90E-02 | 3.84E-01 | 1.06E+00 | 7.87E-02 | 1.06E+00 |
| ENSG000001 | PCOLCE | 8.90E-02 | 3.84E-01 | 7.56E-01 | -4.03E-01 | -1.32E+00 |
| ENSG000001 | C12orf56 | 8.90E-02 | ? | 4.57E-01 | -1.13E+00 | -2.19E+00 |
| ENSG000002 | --- | 8.91E-02 | ? | 3.85E-01 | -1.38E+00 | -2.60E+00 |
| ENSG000002 | --- | 8.91E-02 | ? | 2.38E+00 | 1.25E+00 | 2.38E+00 |
| ENSG000001 | CENPBD1P | 8.91E-02 | 3.85E-01 | 1.08E+00 | 1.13E-01 | 1.08E+00 |
| ENSG000001 | METTL15P1 | 8.91E-02 | ? | 4.73E-01 | -1.08E+00 | -2.12E+00 |
| ENSG000001 | TAF1A | 8.92E-02 | 3.85E-01 | 8.88E-01 | -1.72E-01 | -1.13E+00 |
| ENSG000001 | VAMP4 | 8.92E-02 | 3.85E-01 | 9.20E-01 | -1.21E-01 | -1.09E+00 |
| ENSG000001 | ACSS1 | 8.93E-02 | 3.85E-01 | 1.32E+00 | 4.03E-01 | 1.32E+00 |
| ENSG000001 | EHF | 8.94E-02 | 3.85E-01 | 9.53E-01 | -6.99E-02 | -1.05E+00 |
| ENSG000001 | CDK20 | 8.94E-02 | 3.85E-01 | 1.16E+00 | 2.10E-01 | 1.16E+00 |
| ENSG000001 | MIF4GD | 8.94E-02 | 3.85E-01 | 1.12E+00 | 1.57E-01 | 1.12E+00 |
| ENSG000001 | LRRIQ1 | 8.94E-02 | 3.85E-01 | 8.53E-01 | -2.30E-01 | -1.17E+00 |
| ENSG000002 | --- | 8.95E-02 | ? | 1.40E+00 | 4.85E-01 | 1.40E+00 |
| ENSG000002 | NME2 | 8.97E-02 | 3.86E-01 | 1.09E+00 | 1.27E-01 | 1.09E+00 |
| ENSG000001 | CORO1B | 8.97E-02 | 3.86E-01 | 1.12E+00 | 1.61E-01 | 1.12E+00 |
| ENSG000002 | PITRM1-AS1 | 8.97E-02 | 3.86E-01 | 7.75E-01 | -3.68E-01 | -1.29E+00 |
| ENSG000001 | DARS2 | 8.97E-02 | 3.86E-01 | 1.05E+00 | 7.44E-02 | 1.05E+00 |
| ENSG000001 | SOX7 | 8.97E-02 | 3.86E-01 | 8.86E-01 | -1.74E-01 | -1.13E+00 |
| ENSG00000C | ING3 | 8.97E-02 | 3.86E-01 | 9.00E-01 | -1.52E-01 | -1.11E+00 |
| ENSG00000C | NLK | 8.98E-02 | 3.86E-01 | 9.27E-01 | -1.09E-01 | -1.08E+00 |
| ENSG00000C | RAB18 | 8.98E-02 | 3.86E-01 | 9.42E-01 | -8.66E-02 | -1.06E+00 |
| ENSG000002 | RBM14 | 8.99E-02 | 3.86E-01 | 1.08E+00 | 1.16E-01 | 1.08E+00 |
| ENSG000001 | PUF60 | 8.99E-02 | 3.86E-01 | 1.09E+00 | 1.25E-01 | 1.09E+00 |
| ENSG000002 | --- | 8.99E-02 | ? | 7.65E-01 | -3.86E-01 | -1.31E+00 |
| ENSG000001 | CNGA1 | 8.99E-02 | ? | 7.40E-01 | -4.34E-01 | -1.35E+00 |

| | | | | | | | |
|---|---|---|---|---|---|---|---|
| ENSG000001 | AEN | 9.00E-02 | | 3.86E-01 | 1.07E+00 | 9.61E-02 | 1.07E+00 |
| ENSG000001 | ATP10D | 9.01E-02 | | 3.86E-01 | 1.05E+00 | 7.42E-02 | 1.05E+00 |
| ENSG000001 | C12orf54 | 9.01E-02 | | 3.87E-01 | 1.17E+00 | 2.27E-01 | 1.17E+00 |
| ENSG000001 | MTPAP | 9.02E-02 | | 3.87E-01 | 1.06E+00 | 8.35E-02 | 1.06E+00 |
| ENSG000002 | SDHAF1 | 9.02E-02 | | 3.87E-01 | 1.14E+00 | 1.90E-01 | 1.14E+00 |
| ENSG000001 | DYDC2 | 9.03E-02 | ? | | 5.69E-01 | -8.13E-01 | -1.76E+00 |
| ENSG000002 | LINC02535 | 9.04E-02 | | 3.87E-01 | 9.00E-01 | -1.52E-01 | -1.11E+00 |
| ENSG000001 | HSD11B1L | 9.04E-02 | | 3.87E-01 | 1.30E+00 | 3.79E-01 | 1.30E+00 |
| ENSG000001 | NDUFAF4 | 9.04E-02 | | 3.87E-01 | 9.21E-01 | -1.19E-01 | -1.09E+00 |
| ENSG000001 | PIGP | 9.04E-02 | | 3.87E-01 | 1.10E+00 | 1.40E-01 | 1.10E+00 |
| ENSG000001 | ZNF304 | 9.05E-02 | | 3.87E-01 | 9.33E-01 | -1.01E-01 | -1.07E+00 |
| ENSG000002 | ENPP7P12 | 9.05E-02 | ? | | 1.62E+00 | 6.96E-01 | 1.62E+00 |
| ENSG00000C | EIF3I | 9.05E-02 | | 3.87E-01 | 1.05E+00 | 7.62E-02 | 1.05E+00 |
| ENSG000002 | RPL28P2 | 9.05E-02 | ? | | 5.17E-01 | -9.51E-01 | -1.93E+00 |
| ENSG00000C | CIC | 9.06E-02 | | 3.88E-01 | 1.11E+00 | 1.55E-01 | 1.11E+00 |
| ENSG000002 | --- | 9.06E-02 | | 3.88E-01 | 7.92E-01 | -3.36E-01 | -1.26E+00 |
| ENSG000001 | TYMSOS | 9.06E-02 | ? | | 1.52E+00 | 6.04E-01 | 1.52E+00 |
| ENSG000002 | TRIM26 | 9.06E-02 | | 3.88E-01 | 1.05E+00 | 6.64E-02 | 1.05E+00 |
| ENSG000001 | ADPGK | 9.07E-02 | | 3.88E-01 | 9.14E-01 | -1.30E-01 | -1.09E+00 |
| ENSG000001 | ANKRD26 | 9.07E-02 | | 3.88E-01 | 9.03E-01 | -1.47E-01 | -1.11E+00 |
| ENSG000002 | --- | 9.07E-02 | ? | | 5.83E-01 | -7.79E-01 | -1.72E+00 |
| ENSG000002 | --- | 9.08E-02 | | 3.88E-01 | 1.09E+00 | 1.30E-01 | 1.09E+00 |
| ENSG000001 | ZNF74 | 9.08E-02 | | 3.88E-01 | 9.01E-01 | -1.50E-01 | -1.11E+00 |
| ENSG000002 | H3C6 | 9.09E-02 | | 3.88E-01 | 1.18E+00 | 2.36E-01 | 1.18E+00 |
| ENSG000002 | --- | 9.09E-02 | ? | | 3.62E-01 | -1.47E+00 | -2.76E+00 |
| ENSG000002 | --- | 9.10E-02 | | 3.88E-01 | 7.95E-01 | -3.32E-01 | -1.26E+00 |
| ENSG000001 | ZNF860 | 9.10E-02 | | 3.88E-01 | 8.61E-01 | -2.16E-01 | -1.16E+00 |
| ENSG00000C | FNDC3B | 9.10E-02 | | 3.88E-01 | 9.45E-01 | -8.11E-02 | -1.06E+00 |
| ENSG000001 | CCDC13-AS1 | 9.11E-02 | ? | | 2.29E+00 | 1.20E+00 | 2.29E+00 |
| ENSG000001 | CALML3 | 9.12E-02 | | 3.89E-01 | 1.16E+00 | 2.10E-01 | 1.16E+00 |
| ENSG00000C | PIAS1 | 9.12E-02 | | 3.89E-01 | 9.26E-01 | -1.10E-01 | -1.08E+00 |
| ENSG000002 | NAIP | 9.13E-02 | | 3.89E-01 | 8.10E-01 | -3.03E-01 | -1.23E+00 |
| ENSG000001 | ANTKMT | 9.13E-02 | | 3.89E-01 | 1.29E+00 | 3.72E-01 | 1.29E+00 |
| ENSG000001 | C2orf66 | 9.14E-02 | | 3.89E-01 | 7.54E-01 | -4.07E-01 | -1.33E+00 |
| ENSG00000C | WDR18 | 9.14E-02 | | 3.89E-01 | 1.17E+00 | 2.23E-01 | 1.17E+00 |
| ENSG000002 | RN7SKP70 | 9.14E-02 | ? | | 2.27E+00 | 1.18E+00 | 2.27E+00 |
| ENSG000001 | MOB1B | 9.14E-02 | | 3.89E-01 | 9.32E-01 | -1.01E-01 | -1.07E+00 |
| ENSG000002 | --- | 9.14E-02 | ? | | 5.58E-01 | -8.42E-01 | -1.79E+00 |
| ENSG000001 | TENM4 | 9.15E-02 | | 3.89E-01 | 1.11E+00 | 1.55E-01 | 1.11E+00 |
| ENSG000001 | SLC39A3 | 9.15E-02 | | 3.89E-01 | 1.13E+00 | 1.82E-01 | 1.13E+00 |
| ENSG000001 | SLC35A2 | 9.15E-02 | | 3.89E-01 | 1.10E+00 | 1.42E-01 | 1.10E+00 |
| ENSG000001 | LGI3 | 9.16E-02 | ? | | 1.63E+00 | 7.05E-01 | 1.63E+00 |
| ENSG000001 | KHDC1 | 9.16E-02 | | 3.90E-01 | 1.21E+00 | 2.74E-01 | 1.21E+00 |
| ENSG000001 | AK5 | 9.16E-02 | ? | | 5.25E-01 | -9.31E-01 | -1.91E+00 |
| ENSG000002 | --- | 9.18E-02 | | 3.90E-01 | 1.95E+00 | 9.65E-01 | 1.95E+00 |

| | | | | | | |
|---|---|---|---|---|---|---|
| ENSG000001 | RAB42 | 9.18E-02 | ? | 1.63E+00 | 7.08E-01 | 1.63E+00 |
| ENSG000001 | KCMF1 | 9.18E-02 | 3.90E-01 | 9.45E-01 | -8.22E-02 | -1.06E+00 |
| ENSG000001 | SPTSSB | 9.19E-02 | ? | 1.71E+00 | 7.74E-01 | 1.71E+00 |
| ENSG000001 | HSPG2 | 9.19E-02 | 3.90E-01 | 1.13E+00 | 1.79E-01 | 1.13E+00 |
| ENSG000001 | PRKAR2A | 9.20E-02 | 3.91E-01 | 9.56E-01 | -6.45E-02 | -1.05E+00 |
| ENSG000001 | GAMT | 9.21E-02 | 3.91E-01 | 1.21E+00 | 2.76E-01 | 1.21E+00 |
| ENSG000001 | LIG4 | 9.21E-02 | 3.91E-01 | 9.35E-01 | -9.67E-02 | -1.07E+00 |
| ENSG00000C | SCMH1 | 9.21E-02 | 3.91E-01 | 1.09E+00 | 1.26E-01 | 1.09E+00 |
| ENSG000001 | TASOR | 9.21E-02 | 3.91E-01 | 9.47E-01 | -7.83E-02 | -1.06E+00 |
| ENSG000001 | C11orf80 | 9.22E-02 | 3.91E-01 | 9.27E-01 | -1.09E-01 | -1.08E+00 |
| ENSG000002 | --- | 9.22E-02 | ? | 1.83E+00 | 8.72E-01 | 1.83E+00 |
| ENSG000002 | ST7-AS1 | 9.23E-02 | ? | 4.63E-01 | -1.11E+00 | -2.16E+00 |
| ENSG000002 | ZNF488 | 9.23E-02 | 3.91E-01 | 1.26E+00 | 3.32E-01 | 1.26E+00 |
| ENSG000001 | RALGDS | 9.23E-02 | 3.91E-01 | 1.07E+00 | 1.02E-01 | 1.07E+00 |
| ENSG000001 | TREML2 | 9.23E-02 | ? | 5.02E-01 | -9.93E-01 | -1.99E+00 |
| ENSG000001 | ZNF451 | 9.24E-02 | 3.91E-01 | 9.43E-01 | -8.43E-02 | -1.06E+00 |
| ENSG000001 | CHAMP1 | 9.24E-02 | 3.91E-01 | 9.29E-01 | -1.06E-01 | -1.08E+00 |
| ENSG000002 | LSP1P5 | 9.24E-02 | 3.91E-01 | 1.23E+00 | 2.93E-01 | 1.23E+00 |
| ENSG000001 | INSR | 9.25E-02 | 3.92E-01 | 1.12E+00 | 1.60E-01 | 1.12E+00 |
| ENSG000001 | SEC23B | 9.25E-02 | 3.92E-01 | 1.05E+00 | 7.02E-02 | 1.05E+00 |
| ENSG000001 | GALNT6 | 9.26E-02 | 3.92E-01 | 1.06E+00 | 8.65E-02 | 1.06E+00 |
| ENSG000001 | COPS6 | 9.26E-02 | 3.92E-01 | 1.06E+00 | 8.37E-02 | 1.06E+00 |
| ENSG000001 | EPS15L1 | 9.27E-02 | 3.92E-01 | 1.05E+00 | 7.37E-02 | 1.05E+00 |
| ENSG000002 | --- | 9.27E-02 | ? | 2.10E+00 | 1.07E+00 | 2.10E+00 |
| ENSG000001 | VBP1 | 9.28E-02 | 3.92E-01 | 9.53E-01 | -6.92E-02 | -1.05E+00 |
| ENSG00000C | STK10 | 9.28E-02 | 3.92E-01 | 1.04E+00 | 6.03E-02 | 1.04E+00 |
| ENSG000001 | BCL2A1 | 9.28E-02 | 3.92E-01 | 6.45E-01 | -6.33E-01 | -1.55E+00 |
| ENSG000001 | STRA6 | 9.29E-02 | ? | 6.53E-01 | -6.15E-01 | -1.53E+00 |
| ENSG000002 | --- | 9.29E-02 | ? | 5.86E-01 | -7.71E-01 | -1.71E+00 |
| ENSG000001 | DDX27 | 9.30E-02 | 3.93E-01 | 9.48E-01 | -7.73E-02 | -1.06E+00 |
| ENSG000002 | LINC02193 | 9.30E-02 | 3.93E-01 | 7.62E-01 | -3.92E-01 | -1.31E+00 |
| ENSG000001 | AHSG | 9.30E-02 | ? | 2.25E+00 | 1.17E+00 | 2.25E+00 |
| ENSG000001 | PRKG2 | 9.31E-02 | 3.93E-01 | 8.54E-01 | -2.28E-01 | -1.17E+00 |
| ENSG000002 | LINC02334 | 9.31E-02 | ? | 1.50E+00 | 5.83E-01 | 1.50E+00 |
| ENSG000002 | --- | 9.32E-02 | ? | 4.22E-01 | -1.25E+00 | -2.37E+00 |
| ENSG000001 | RNF43 | 9.32E-02 | 3.93E-01 | 1.07E+00 | 1.03E-01 | 1.07E+00 |
| ENSG000001 | MIPOL1 | 9.32E-02 | 3.93E-01 | 9.22E-01 | -1.17E-01 | -1.08E+00 |
| ENSG000002 | TCTN1 | 9.32E-02 | 3.93E-01 | 1.08E+00 | 1.17E-01 | 1.08E+00 |
| ENSG000002 | LINC02984 | 9.32E-02 | 3.93E-01 | 1.24E+00 | 3.15E-01 | 1.24E+00 |
| ENSG000001 | MRPL36 | 9.33E-02 | 3.93E-01 | 1.08E+00 | 1.09E-01 | 1.08E+00 |
| ENSG000001 | INTS10 | 9.33E-02 | 3.93E-01 | 1.05E+00 | 7.42E-02 | 1.05E+00 |
| ENSG00000C | IFFO1 | 9.33E-02 | 3.93E-01 | 8.17E-01 | -2.91E-01 | -1.22E+00 |
| ENSG000002 | --- | 9.34E-02 | ? | 6.99E-01 | -5.16E-01 | -1.43E+00 |
| ENSG00000C | RPAP3 | 9.34E-02 | 3.93E-01 | 9.31E-01 | -1.04E-01 | -1.07E+00 |
| ENSG000001 | LRR1 | 9.35E-02 | 3.94E-01 | 1.11E+00 | 1.49E-01 | 1.11E+00 |

| | | | | | | |
|---|---|---|---|---|---|---|
| ENSG000001 | RIOX2 | 9.36E-02 | 3.94E-01 | 1.05E+00 | 6.83E-02 | 1.05E+00 |
| ENSG000001 | INHBB | 9.36E-02 | ? | 3.58E-01 | -1.48E+00 | -2.79E+00 |
| ENSG000002 | GUSBP2 | 9.36E-02 | ? | 5.14E-01 | -9.60E-01 | -1.95E+00 |
| ENSG00000C | RFX2 | 9.36E-02 | 3.94E-01 | 1.15E+00 | 1.99E-01 | 1.15E+00 |
| ENSG000001 | PLCB1 | 9.37E-02 | 3.94E-01 | 1.19E+00 | 2.46E-01 | 1.19E+00 |
| ENSG000001 | CAB39L | 9.38E-02 | 3.94E-01 | 9.13E-01 | -1.31E-01 | -1.09E+00 |
| ENSG000001 | KLHL12 | 9.38E-02 | 3.94E-01 | 1.05E+00 | 7.28E-02 | 1.05E+00 |
| ENSG000001 | STEAP2 | 9.39E-02 | 3.94E-01 | 1.15E+00 | 2.03E-01 | 1.15E+00 |
| ENSG000002 | --- | 9.39E-02 | ? | 6.00E-01 | -7.37E-01 | -1.67E+00 |
| ENSG00000C | MAPK8IP2 | 9.39E-02 | 3.95E-01 | 1.23E+00 | 2.97E-01 | 1.23E+00 |
| ENSG000001 | DPP4 | 9.40E-02 | 3.95E-01 | 8.07E-01 | -3.09E-01 | -1.24E+00 |
| ENSG000001 | NDEL1 | 9.42E-02 | 3.95E-01 | 9.48E-01 | -7.71E-02 | -1.05E+00 |
| ENSG000001 | ZSWIM6 | 9.42E-02 | 3.95E-01 | 9.16E-01 | -1.27E-01 | -1.09E+00 |
| ENSG000002 | LINC01010 | 9.43E-02 | ? | 1.75E+00 | 8.04E-01 | 1.75E+00 |
| ENSG00000C | DIS3 | 9.43E-02 | 3.96E-01 | 9.41E-01 | -8.84E-02 | -1.06E+00 |
| ENSG000002 | PPP2R5CP | 9.43E-02 | ? | 6.12E-01 | -7.08E-01 | -1.63E+00 |
| ENSG000001 | FRMPD2 | 9.44E-02 | ? | 1.39E+00 | 4.79E-01 | 1.39E+00 |
| ENSG000001 | KIAA1522 | 9.45E-02 | 3.96E-01 | 1.08E+00 | 1.06E-01 | 1.08E+00 |
| ENSG000001 | TIMM10 | 9.45E-02 | 3.96E-01 | 1.08E+00 | 1.15E-01 | 1.08E+00 |
| ENSG000001 | ROMO1 | 9.45E-02 | 3.96E-01 | 1.17E+00 | 2.20E-01 | 1.17E+00 |
| ENSG000001 | ZNF330 | 9.46E-02 | 3.96E-01 | 9.41E-01 | -8.72E-02 | -1.06E+00 |
| ENSG000002 | PDXP-DT | 9.46E-02 | ? | 6.78E-01 | -5.60E-01 | -1.47E+00 |
| ENSG000002 | TAPT1-AS1 | 9.47E-02 | ? | 1.40E+00 | 4.90E-01 | 1.40E+00 |
| ENSG00000C | SRI | 9.47E-02 | 3.96E-01 | 1.05E+00 | 7.72E-02 | 1.05E+00 |
| ENSG000001 | ATPAF2 | 9.47E-02 | 3.96E-01 | 1.08E+00 | 1.11E-01 | 1.08E+00 |
| ENSG000001 | GSTM3 | 9.47E-02 | 3.96E-01 | 1.08E+00 | 1.06E-01 | 1.08E+00 |
| ENSG000001 | HMGN5 | 9.47E-02 | 3.97E-01 | 8.39E-01 | -2.54E-01 | -1.19E+00 |
| ENSG000001 | BCR | 9.48E-02 | 3.97E-01 | 1.07E+00 | 1.02E-01 | 1.07E+00 |
| ENSG000001 | CMTR1 | 9.49E-02 | 3.97E-01 | 1.08E+00 | 1.05E-01 | 1.08E+00 |
| ENSG000001 | TPPP | 9.49E-02 | 3.97E-01 | 1.09E+00 | 1.30E-01 | 1.09E+00 |
| ENSG000001 | LCE3E | 9.50E-02 | ? | 6.44E-01 | -6.35E-01 | -1.55E+00 |
| ENSG000002 | --- | 9.51E-02 | ? | 1.33E+00 | 4.11E-01 | 1.33E+00 |
| ENSG00000C | VPS50 | 9.51E-02 | 3.98E-01 | 9.36E-01 | -9.50E-02 | -1.07E+00 |
| ENSG00000C | FAM120A | 9.52E-02 | 3.98E-01 | 9.54E-01 | -6.78E-02 | -1.05E+00 |
| ENSG000001 | RPA3 | 9.52E-02 | 3.98E-01 | 1.12E+00 | 1.66E-01 | 1.12E+00 |
| ENSG000002 | ATXN7L3B | 9.53E-02 | 3.98E-01 | 1.07E+00 | 9.63E-02 | 1.07E+00 |
| ENSG000001 | EIF3E | 9.53E-02 | 3.98E-01 | 1.06E+00 | 7.76E-02 | 1.06E+00 |
| ENSG000001 | LGALS3 | 9.53E-02 | 3.98E-01 | 1.06E+00 | 8.08E-02 | 1.06E+00 |
| ENSG000001 | PALLD | 9.54E-02 | 3.98E-01 | 1.08E+00 | 1.06E-01 | 1.08E+00 |
| ENSG000001 | DUSP18 | 9.54E-02 | 3.98E-01 | 1.08E+00 | 1.16E-01 | 1.08E+00 |
| ENSG00000C | LMCD1 | 9.54E-02 | 3.98E-01 | 1.23E+00 | 3.04E-01 | 1.23E+00 |
| ENSG000001 | RBM23 | 9.54E-02 | 3.98E-01 | 1.06E+00 | 7.93E-02 | 1.06E+00 |
| ENSG000002 | SNRPGP15 | 9.54E-02 | ? | 7.49E-01 | -4.17E-01 | -1.33E+00 |
| ENSG000002 | LINC00638 | 9.54E-02 | ? | 6.79E-01 | -5.58E-01 | -1.47E+00 |
| ENSG000001 | SP4 | 9.55E-02 | 3.98E-01 | 8.72E-01 | -1.97E-01 | -1.15E+00 |

| | | | | | | |
|---|---|---|---|---|---|---|
| ENSG000001 | TMEM267 | 9.55E-02 | 3.98E-01 | 1.08E+00 | 1.09E-01 | 1.08E+00 |
| ENSG00000C | PHLDB1 | 9.55E-02 | 3.98E-01 | 1.06E+00 | 8.17E-02 | 1.06E+00 |
| ENSG000001 | RARS2 | 9.55E-02 | 3.98E-01 | 9.38E-01 | -9.20E-02 | -1.07E+00 |
| ENSG000001 | RPS6KB2 | 9.56E-02 | 3.98E-01 | 1.09E+00 | 1.23E-01 | 1.09E+00 |
| ENSG000001 | MT-ND6 | 9.56E-02 | 3.98E-01 | 8.25E-01 | -2.77E-01 | -1.21E+00 |
| ENSG000001 | TARS2 | 9.56E-02 | 3.98E-01 | 9.32E-01 | -1.01E-01 | -1.07E+00 |
| ENSG000001 | CHAC2 | 9.56E-02 | 3.98E-01 | 8.98E-01 | -1.56E-01 | -1.11E+00 |
| ENSG000001 | CRY2 | 9.57E-02 | 3.98E-01 | 1.11E+00 | 1.45E-01 | 1.11E+00 |
| ENSG000001 | OARD1 | 9.57E-02 | 3.98E-01 | 9.29E-01 | -1.06E-01 | -1.08E+00 |
| ENSG000002 | --- | 9.57E-02 | ? | 6.48E-01 | -6.27E-01 | -1.54E+00 |
| ENSG000002 | --- | 9.57E-02 | ? | 1.54E+00 | 6.19E-01 | 1.54E+00 |
| ENSG000001 | SMG7 | 9.57E-02 | 3.98E-01 | 9.36E-01 | -9.60E-02 | -1.07E+00 |
| ENSG000001 | SEPTIN10 | 9.57E-02 | 3.98E-01 | 9.40E-01 | -8.91E-02 | -1.06E+00 |
| ENSG00000C | TACR2 | 9.58E-02 | 3.98E-01 | 8.36E-01 | -2.58E-01 | -1.20E+00 |
| ENSG000001 | PNPT1 | 9.58E-02 | 3.98E-01 | 9.32E-01 | -1.01E-01 | -1.07E+00 |
| ENSG00000C | KCNQ1 | 9.59E-02 | ? | 5.87E-01 | -7.67E-01 | -1.70E+00 |
| ENSG000001 | CAVIN1 | 9.59E-02 | 3.98E-01 | 1.08E+00 | 1.06E-01 | 1.08E+00 |
| ENSG00000C | RBL1 | 9.60E-02 | 3.98E-01 | 8.53E-01 | -2.30E-01 | -1.17E+00 |
| ENSG000002 | TEN1 | 9.60E-02 | 3.98E-01 | 1.21E+00 | 2.80E-01 | 1.21E+00 |
| ENSG00000C | EML1 | 9.60E-02 | 3.98E-01 | 1.10E+00 | 1.36E-01 | 1.10E+00 |
| ENSG000001 | BORCS7 | 9.61E-02 | 3.98E-01 | 9.30E-01 | -1.04E-01 | -1.07E+00 |
| ENSG000001 | SELENOS | 9.61E-02 | 3.98E-01 | 9.50E-01 | -7.41E-02 | -1.05E+00 |
| ENSG000002 | LINC00865 | 9.61E-02 | 3.98E-01 | 1.17E+00 | 2.23E-01 | 1.17E+00 |
| ENSG00000C | SLC25A40 | 9.62E-02 | 3.98E-01 | 8.93E-01 | -1.63E-01 | -1.12E+00 |
| ENSG000002 | MANCR | 9.62E-02 | 3.98E-01 | 8.78E-01 | -1.88E-01 | -1.14E+00 |
| ENSG000001 | ABCG4 | 9.62E-02 | ? | 1.78E+00 | 8.32E-01 | 1.78E+00 |
| ENSG000001 | BABAM1 | 9.62E-02 | 3.98E-01 | 1.05E+00 | 7.48E-02 | 1.05E+00 |
| ENSG000002 | LINC00390 | 9.62E-02 | ? | 1.95E+00 | 9.64E-01 | 1.95E+00 |
| ENSG00000C | SLC4A7 | 9.62E-02 | 3.98E-01 | 9.34E-01 | -9.78E-02 | -1.07E+00 |
| ENSG00000C | CYBRD1 | 9.63E-02 | 3.98E-01 | 1.09E+00 | 1.18E-01 | 1.09E+00 |
| ENSG000001 | TESK1 | 9.63E-02 | 3.98E-01 | 9.20E-01 | -1.20E-01 | -1.09E+00 |
| ENSG000001 | GPCPD1 | 9.64E-02 | 3.98E-01 | 1.12E+00 | 1.69E-01 | 1.12E+00 |
| ENSG000002 | SNHG21 | 9.64E-02 | ? | 7.03E-01 | -5.08E-01 | -1.42E+00 |
| ENSG000001 | MIEF2 | 9.64E-02 | 3.98E-01 | 1.14E+00 | 1.94E-01 | 1.14E+00 |
| ENSG000002 | --- | 9.64E-02 | 3.98E-01 | 1.23E+00 | 2.99E-01 | 1.23E+00 |
| ENSG000001 | CGREF1 | 9.64E-02 | 3.98E-01 | 1.30E+00 | 3.75E-01 | 1.30E+00 |
| ENSG00000C | SLC30A9 | 9.65E-02 | 3.99E-01 | 9.56E-01 | -6.55E-02 | -1.05E+00 |
| ENSG000002 | RPL17 | 9.67E-02 | 3.99E-01 | 1.05E+00 | 6.55E-02 | 1.05E+00 |
| ENSG000001 | MED14 | 9.67E-02 | 3.99E-01 | 9.47E-01 | -7.85E-02 | -1.06E+00 |
| ENSG000001 | KMT5B | 9.68E-02 | 4.00E-01 | 9.51E-01 | -7.20E-02 | -1.05E+00 |
| ENSG000001 | UBTD2 | 9.69E-02 | 4.00E-01 | 1.07E+00 | 1.01E-01 | 1.07E+00 |
| ENSG000001 | QRICH2 | 9.70E-02 | 4.00E-01 | 1.12E+00 | 1.68E-01 | 1.12E+00 |
| ENSG000001 | CAPN2 | 9.70E-02 | 4.00E-01 | 9.71E-01 | -4.22E-02 | -1.03E+00 |
| ENSG000002 | APOL6 | 9.71E-02 | 4.00E-01 | 8.80E-01 | -1.85E-01 | -1.14E+00 |
| ENSG000001 | SARAF | 9.71E-02 | 4.00E-01 | 1.05E+00 | 6.47E-02 | 1.05E+00 |

| | | | | | | |
|---|---|---|---|---|---|---|
| ENSG000001 | ATP6V1A | 9.71E-02 | 4.00E-01 | 1.04E+00 | 5.72E-02 | 1.04E+00 |
| ENSG000001 | TFG | 9.72E-02 | 4.00E-01 | 9.62E-01 | -5.55E-02 | -1.04E+00 |
| ENSG000001 | KIF2C | 9.72E-02 | 4.00E-01 | 7.97E-01 | -3.27E-01 | -1.25E+00 |
| ENSG000001 | PBX1 | 9.72E-02 | 4.00E-01 | 1.12E+00 | 1.62E-01 | 1.12E+00 |
| ENSG000001 | GABRE | 9.72E-02 | 4.00E-01 | 1.19E+00 | 2.45E-01 | 1.19E+00 |
| ENSG000001 | ING4 | 9.72E-02 | 4.00E-01 | 1.08E+00 | 1.14E-01 | 1.08E+00 |
| ENSG000001 | TBCA | 9.76E-02 | 4.01E-01 | 9.50E-01 | -7.35E-02 | -1.05E+00 |
| ENSG000001 | CCDC125 | 9.76E-02 | 4.01E-01 | 9.19E-01 | -1.22E-01 | -1.09E+00 |
| ENSG000001 | ARID3A | 9.76E-02 | 4.01E-01 | 1.11E+00 | 1.50E-01 | 1.11E+00 |
| ENSG000002 | SRP14-DT | 9.76E-02 | 4.01E-01 | 8.34E-01 | -2.61E-01 | -1.20E+00 |
| ENSG000001 | SEC11C | 9.78E-02 | 4.01E-01 | 9.27E-01 | -1.09E-01 | -1.08E+00 |
| ENSG000001 | FAM153A | 9.78E-02 | 4.01E-01 | 7.87E-01 | -3.46E-01 | -1.27E+00 |
| ENSG000001 | TM2D3 | 9.78E-02 | 4.01E-01 | 1.06E+00 | 8.89E-02 | 1.06E+00 |
| ENSG000001 | IL13RA2 | 9.78E-02 | 4.01E-01 | 7.66E-01 | -3.85E-01 | -1.31E+00 |
| ENSG000002 | KLRC2 | 9.78E-02 | ? | 1.60E+00 | 6.78E-01 | 1.60E+00 |
| ENSG000002 | MYZAP | 9.78E-02 | 4.01E-01 | 1.27E+00 | 3.43E-01 | 1.27E+00 |
| ENSG000002 | FLJ16779 | 9.78E-02 | ? | 5.55E-01 | -8.50E-01 | -1.80E+00 |
| ENSG000001 | USP31 | 9.78E-02 | 4.01E-01 | 1.06E+00 | 8.39E-02 | 1.06E+00 |
| ENSG000002 | NPIPA3 | 9.79E-02 | 4.02E-01 | 6.69E-01 | -5.80E-01 | -1.49E+00 |
| ENSG000001 | SP140L | 9.79E-02 | 4.02E-01 | 9.37E-01 | -9.36E-02 | -1.07E+00 |
| ENSG000001 | PHF13 | 9.80E-02 | 4.02E-01 | 1.10E+00 | 1.40E-01 | 1.10E+00 |
| ENSG000002 | SHANK3 | 9.80E-02 | 4.02E-01 | 1.11E+00 | 1.54E-01 | 1.11E+00 |
| ENSG000001 | B3GNT2 | 9.81E-02 | 4.02E-01 | 9.36E-01 | -9.51E-02 | -1.07E+00 |
| ENSG000001 | PRPF19 | 9.82E-02 | 4.02E-01 | 1.06E+00 | 8.12E-02 | 1.06E+00 |
| ENSG000002 | LINC02561 | 9.82E-02 | ? | 6.42E-01 | -6.40E-01 | -1.56E+00 |
| ENSG000001 | EPG5 | 9.82E-02 | 4.02E-01 | 1.10E+00 | 1.38E-01 | 1.10E+00 |
| ENSG000001 | ZNF23 | 9.82E-02 | 4.02E-01 | 9.11E-01 | -1.35E-01 | -1.10E+00 |
| ENSG000001 | MFN2 | 9.83E-02 | 4.02E-01 | 1.06E+00 | 9.07E-02 | 1.06E+00 |
| ENSG000002 | SHROOM3-A | 9.84E-02 | ? | 5.45E-01 | -8.74E-01 | -1.83E+00 |
| ENSG00000C | DLGAP4 | 9.84E-02 | 4.02E-01 | 9.41E-01 | -8.73E-02 | -1.06E+00 |
| ENSG000001 | PER2 | 9.85E-02 | 4.02E-01 | 1.12E+00 | 1.63E-01 | 1.12E+00 |
| ENSG000001 | CERCAM | 9.85E-02 | 4.02E-01 | 1.10E+00 | 1.34E-01 | 1.10E+00 |
| ENSG000001 | INTS5 | 9.85E-02 | 4.02E-01 | 1.13E+00 | 1.73E-01 | 1.13E+00 |
| ENSG00000C | SUPT16H | 9.85E-02 | 4.02E-01 | 9.38E-01 | -9.18E-02 | -1.07E+00 |
| ENSG000001 | IVD | 9.85E-02 | 4.02E-01 | 1.07E+00 | 9.97E-02 | 1.07E+00 |
| ENSG000001 | ADI1 | 9.86E-02 | 4.02E-01 | 1.06E+00 | 7.97E-02 | 1.06E+00 |
| ENSG000002 | --- | 9.86E-02 | ? | 5.98E-01 | -7.42E-01 | -1.67E+00 |
| ENSG000001 | SNPH | 9.87E-02 | 4.03E-01 | 1.12E+00 | 1.60E-01 | 1.12E+00 |
| ENSG000002 | DIO2 | 9.87E-02 | ? | 5.27E-01 | -9.24E-01 | -1.90E+00 |
| ENSG000001 | HHIP | 9.87E-02 | ? | 1.55E+00 | 6.32E-01 | 1.55E+00 |
| ENSG000002 | --- | 9.87E-02 | 4.03E-01 | 8.69E-01 | -2.03E-01 | -1.15E+00 |
| ENSG000002 | --- | 9.88E-02 | ? | 1.83E+00 | 8.71E-01 | 1.83E+00 |
| ENSG000002 | JPT2 | 9.88E-02 | 4.03E-01 | 9.65E-01 | -5.11E-02 | -1.04E+00 |
| ENSG000001 | VLDLR | 9.88E-02 | 4.03E-01 | 1.07E+00 | 1.01E-01 | 1.07E+00 |
| ENSG000001 | KIAA1614 | 9.88E-02 | 4.03E-01 | 8.53E-01 | -2.29E-01 | -1.17E+00 |

| | | | | | | |
|---|---|---|---|---|---|---|
| ENSG000001 | ACBD3 | 9.88E-02 | 4.03E-01 | 9.42E-01 | -8.68E-02 | -1.06E+00 |
| ENSG00000C | UNG | 9.88E-02 | 4.03E-01 | 9.20E-01 | -1.20E-01 | -1.09E+00 |
| ENSG000001 | SHROOM4 | 9.88E-02 | 4.03E-01 | 9.21E-01 | -1.19E-01 | -1.09E+00 |
| ENSG000001 | SETD6 | 9.89E-02 | 4.03E-01 | 9.18E-01 | -1.23E-01 | -1.09E+00 |
| ENSG000001 | ATG2A | 9.89E-02 | 4.03E-01 | 1.12E+00 | 1.62E-01 | 1.12E+00 |
| ENSG000002 | CICP27 | 9.90E-02 | ? | 5.77E-01 | -7.93E-01 | -1.73E+00 |
| ENSG000001 | CNTN5 | 9.90E-02 | 4.03E-01 | 7.46E-01 | -4.22E-01 | -1.34E+00 |
| ENSG000001 | RNPEP | 9.91E-02 | 4.03E-01 | 1.06E+00 | 8.52E-02 | 1.06E+00 |
| ENSG000001 | TIMM23B-A( | 9.91E-02 | 4.03E-01 | 1.28E+00 | 3.53E-01 | 1.28E+00 |
| ENSG000001 | TBCCD1 | 9.92E-02 | 4.03E-01 | 1.07E+00 | 1.03E-01 | 1.07E+00 |
| ENSG000001 | HIGD2A | 9.92E-02 | 4.03E-01 | 1.12E+00 | 1.59E-01 | 1.12E+00 |
| ENSG000001 | KDM6A | 9.93E-02 | 4.04E-01 | 1.07E+00 | 1.04E-01 | 1.07E+00 |
| ENSG000001 | SMAD4 | 9.94E-02 | 4.04E-01 | 9.56E-01 | -6.48E-02 | -1.05E+00 |
| ENSG000001 | ZNF708 | 9.95E-02 | 4.04E-01 | 8.97E-01 | -1.56E-01 | -1.11E+00 |
| ENSG000001 | GCM1 | 9.95E-02 | ? | 1.73E+00 | 7.94E-01 | 1.73E+00 |
| ENSG000001 | HINT1 | 9.95E-02 | 4.04E-01 | 1.06E+00 | 8.58E-02 | 1.06E+00 |
| ENSG000001 | SH3D19 | 9.96E-02 | 4.05E-01 | 9.42E-01 | -8.64E-02 | -1.06E+00 |
| ENSG000002 | --- | 9.97E-02 | ? | 3.96E-01 | -1.34E+00 | -2.52E+00 |
| ENSG000002 | --- | 9.97E-02 | ? | 2.89E+00 | 1.53E+00 | 2.89E+00 |
| ENSG00000C | TRAF5 | 9.97E-02 | 4.05E-01 | 8.82E-01 | -1.80E-01 | -1.13E+00 |
| ENSG000002 | CCDC144BP | 9.98E-02 | 4.05E-01 | 8.47E-01 | -2.40E-01 | -1.18E+00 |
| ENSG000001 | KCNB2 | 9.98E-02 | ? | 6.34E-01 | -6.58E-01 | -1.58E+00 |
| ENSG000002 | STUM | 9.98E-02 | ? | 1.51E+00 | 5.90E-01 | 1.51E+00 |
| ENSG000001 | ESRRA | 9.99E-02 | 4.05E-01 | 1.13E+00 | 1.82E-01 | 1.13E+00 |
| ENSG000002 | PCDHGC5 | 9.99E-02 | 4.05E-01 | 7.97E-01 | -3.27E-01 | -1.25E+00 |
| ENSG000002 | --- | 1.00E-01 | 4.05E-01 | 6.80E-01 | -5.56E-01 | -1.47E+00 |
| ENSG000001 | AKAP5 | 1.00E-01 | ? | 6.44E-01 | -6.34E-01 | -1.55E+00 |
| ENSG00000C | RPS6KA2 | 1.00E-01 | 4.05E-01 | 8.85E-01 | -1.76E-01 | -1.13E+00 |
| ENSG000001 | ZNF165 | 1.00E-01 | 4.05E-01 | 8.75E-01 | -1.93E-01 | -1.14E+00 |
| ENSG000001 | GSN | 1.00E-01 | 4.05E-01 | 1.09E+00 | 1.27E-01 | 1.09E+00 |
| ENSG000001 | ZNF367 | 1.00E-01 | 4.05E-01 | 8.01E-01 | -3.20E-01 | -1.25E+00 |
| ENSG000002 | --- | 1.00E-01 | ? | 2.05E+00 | 1.04E+00 | 2.05E+00 |
| ENSG000002 | --- | 1.00E-01 | 4.06E-01 | 1.22E+00 | 2.89E-01 | 1.22E+00 |
| ENSG00000C | ARAF | 1.00E-01 | 4.06E-01 | 1.06E+00 | 9.02E-02 | 1.06E+00 |
| ENSG000001 | SH3BP4 | 1.00E-01 | 4.06E-01 | 1.04E+00 | 6.18E-02 | 1.04E+00 |
| ENSG00000C | HCCS | 1.00E-01 | 4.06E-01 | 9.49E-01 | -7.50E-02 | -1.05E+00 |
| ENSG000002 | --- | 1.00E-01 | ? | 1.98E+00 | 9.83E-01 | 1.98E+00 |
| ENSG000001 | NUDT13 | 1.00E-01 | ? | 1.34E+00 | 4.28E-01 | 1.34E+00 |
| ENSG000001 | PPP2R1A | 1.00E-01 | 4.06E-01 | 1.09E+00 | 1.27E-01 | 1.09E+00 |
| ENSG000001 | BLOC1S6 | 1.00E-01 | 4.06E-01 | 1.05E+00 | 6.59E-02 | 1.05E+00 |
| ENSG00000C | PLXND1 | 1.00E-01 | ? | 1.46E+00 | 5.45E-01 | 1.46E+00 |
| ENSG000001 | KLHL29 | 1.01E-01 | 4.06E-01 | 1.06E+00 | 8.85E-02 | 1.06E+00 |
| ENSG000002 | HEXA | 1.01E-01 | 4.07E-01 | 1.08E+00 | 1.13E-01 | 1.08E+00 |
| ENSG000001 | ELAC1 | 1.01E-01 | 4.07E-01 | 1.12E+00 | 1.69E-01 | 1.12E+00 |
| ENSG000001 | PRDX2 | 1.01E-01 | 4.07E-01 | 1.08E+00 | 1.11E-01 | 1.08E+00 |

| | | | | | | |
|---|---|---|---|---|---|---|
| ENSG00000C | SIKE1 | 1.01E-01 | 4.07E-01 | 9.57E-01 | -6.35E-02 | -1.05E+00 |
| ENSG000002 | ADGRL1-AS1 | 1.01E-01 | 4.07E-01 | 8.32E-01 | -2.66E-01 | -1.20E+00 |
| ENSG000001 | MARVELD1 | 1.01E-01 | 4.07E-01 | 1.09E+00 | 1.27E-01 | 1.09E+00 |
| ENSG000001 | --- | 1.01E-01 | ? | 4.37E-01 | -1.19E+00 | -2.29E+00 |
| ENSG00000C | SIRT1 | 1.01E-01 | 4.07E-01 | 9.16E-01 | -1.27E-01 | -1.09E+00 |
| ENSG000001 | LOX | 1.01E-01 | 4.07E-01 | 8.39E-01 | -2.54E-01 | -1.19E+00 |
| ENSG000001 | VPS4A | 1.01E-01 | 4.07E-01 | 1.05E+00 | 6.70E-02 | 1.05E+00 |
| ENSG00000C | CHPF2 | 1.01E-01 | 4.07E-01 | 1.11E+00 | 1.57E-01 | 1.11E+00 |
| ENSG000002 | DM1-AS | 1.01E-01 | ? | 2.44E+00 | 1.28E+00 | 2.44E+00 |
| ENSG000001 | AMOTL2 | 1.01E-01 | 4.07E-01 | 1.08E+00 | 1.05E-01 | 1.08E+00 |
| ENSG000001 | SKA1 | 1.01E-01 | 4.07E-01 | 7.95E-01 | -3.31E-01 | -1.26E+00 |
| ENSG000001 | NMRK1 | 1.01E-01 | 4.07E-01 | 1.09E+00 | 1.20E-01 | 1.09E+00 |
| ENSG000001 | --- | 1.01E-01 | 4.07E-01 | 7.54E-01 | -4.08E-01 | -1.33E+00 |
| ENSG000001 | EIF4G2 | 1.01E-01 | 4.07E-01 | 9.59E-01 | -5.99E-02 | -1.04E+00 |
| ENSG000001 | GAS6 | 1.01E-01 | 4.07E-01 | 1.11E+00 | 1.52E-01 | 1.11E+00 |
| ENSG000001 | NSDHL | 1.01E-01 | 4.07E-01 | 1.06E+00 | 8.26E-02 | 1.06E+00 |
| ENSG000001 | LYRM7 | 1.01E-01 | 4.07E-01 | 9.38E-01 | -9.22E-02 | -1.07E+00 |
| ENSG000002 | --- | 1.01E-01 | 4.07E-01 | 7.89E-01 | -3.41E-01 | -1.27E+00 |
| ENSG000002 | LNCATV | 1.01E-01 | ? | 6.18E-01 | -6.93E-01 | -1.62E+00 |
| ENSG000001 | MGAT5 | 1.01E-01 | 4.08E-01 | 9.45E-01 | -8.10E-02 | -1.06E+00 |
| ENSG000002 | --- | 1.01E-01 | ? | 7.19E-01 | -4.75E-01 | -1.39E+00 |
| ENSG000001 | TMEM109 | 1.02E-01 | 4.08E-01 | 1.08E+00 | 1.04E-01 | 1.08E+00 |
| ENSG000001 | SSC4D | 1.02E-01 | 4.08E-01 | 1.18E+00 | 2.38E-01 | 1.18E+00 |
| ENSG000001 | SCP2 | 1.02E-01 | 4.09E-01 | 1.05E+00 | 7.26E-02 | 1.05E+00 |
| ENSG000002 | --- | 1.02E-01 | ? | 1.38E+00 | 4.64E-01 | 1.38E+00 |
| ENSG000001 | LMF1 | 1.02E-01 | 4.10E-01 | 1.28E+00 | 3.52E-01 | 1.28E+00 |
| ENSG000001 | TXNRD1 | 1.02E-01 | 4.10E-01 | 9.05E-01 | -1.43E-01 | -1.10E+00 |
| ENSG000001 | GRK6 | 1.02E-01 | 4.10E-01 | 1.08E+00 | 1.17E-01 | 1.08E+00 |
| ENSG000001 | PIGC | 1.02E-01 | 4.10E-01 | 1.08E+00 | 1.06E-01 | 1.08E+00 |
| ENSG000001 | IMPDH2 | 1.02E-01 | 4.10E-01 | 1.08E+00 | 1.10E-01 | 1.08E+00 |
| ENSG00000C | CFAP20 | 1.02E-01 | 4.10E-01 | 9.29E-01 | -1.06E-01 | -1.08E+00 |
| ENSG000002 | --- | 1.02E-01 | ? | 1.34E+00 | 4.21E-01 | 1.34E+00 |
| ENSG000001 | SLC9B1 | 1.02E-01 | ? | 2.09E+00 | 1.07E+00 | 2.09E+00 |
| ENSG000002 | --- | 1.02E-01 | 4.11E-01 | 7.92E-01 | -3.36E-01 | -1.26E+00 |
| ENSG000002 | --- | 1.03E-01 | 4.11E-01 | 7.75E-01 | -3.68E-01 | -1.29E+00 |
| ENSG000001 | SLC30A1 | 1.03E-01 | 4.12E-01 | 9.43E-01 | -8.51E-02 | -1.06E+00 |
| ENSG000001 | SFPQ | 1.03E-01 | 4.12E-01 | 9.37E-01 | -9.39E-02 | -1.07E+00 |
| ENSG000002 | --- | 1.03E-01 | ? | 7.19E-01 | -4.75E-01 | -1.39E+00 |
| ENSG000002 | PSG4 | 1.03E-01 | 4.12E-01 | 1.17E+00 | 2.30E-01 | 1.17E+00 |
| ENSG000001 | COX17 | 1.03E-01 | 4.12E-01 | 9.17E-01 | -1.24E-01 | -1.09E+00 |
| ENSG00000C | LIMCH1 | 1.03E-01 | 4.12E-01 | 1.14E+00 | 1.90E-01 | 1.14E+00 |
| ENSG000002 | LINC02958 | 1.03E-01 | 4.12E-01 | 7.73E-01 | -3.72E-01 | -1.29E+00 |
| ENSG000001 | PRODH | 1.03E-01 | 4.12E-01 | 8.56E-01 | -2.24E-01 | -1.17E+00 |
| ENSG000001 | D2HGDH | 1.03E-01 | 4.12E-01 | 1.13E+00 | 1.83E-01 | 1.13E+00 |
| ENSG000001 | MRPL40 | 1.03E-01 | 4.12E-01 | 1.07E+00 | 1.01E-01 | 1.07E+00 |

| | | | | | | |
|---|---|---|---|---|---|---|
| ENSG000001 | PGM2 | 1.03E-01 | 4.12E-01 | 1.06E+00 | 8.99E-02 | 1.06E+00 |
| ENSG000002 | --- | 1.03E-01 | ? | 6.05E-01 | -7.24E-01 | -1.65E+00 |
| ENSG000001 | MTCH2 | 1.03E-01 | 4.12E-01 | 1.05E+00 | 6.68E-02 | 1.05E+00 |
| ENSG000001 | CWC15 | 1.03E-01 | 4.12E-01 | 9.34E-01 | -9.80E-02 | -1.07E+00 |
| ENSG00000C | SLC25A5 | 1.03E-01 | 4.12E-01 | 1.07E+00 | 1.03E-01 | 1.07E+00 |
| ENSG000002 | NYNRIN | 1.03E-01 | 4.12E-01 | 1.09E+00 | 1.18E-01 | 1.09E+00 |
| ENSG000001 | DENND4A | 1.03E-01 | 4.12E-01 | 9.31E-01 | -1.04E-01 | -1.07E+00 |
| ENSG000001 | SUMF2 | 1.03E-01 | 4.12E-01 | 1.09E+00 | 1.28E-01 | 1.09E+00 |
| ENSG000001 | NEDD1 | 1.03E-01 | 4.12E-01 | 9.36E-01 | -9.62E-02 | -1.07E+00 |
| ENSG000001 | CYP27B1 | 1.03E-01 | 4.12E-01 | 8.44E-01 | -2.45E-01 | -1.19E+00 |
| ENSG000002 | PPP1CB | 1.04E-01 | 4.13E-01 | 9.41E-01 | -8.85E-02 | -1.06E+00 |
| ENSG000001 | WDCP | 1.04E-01 | 4.13E-01 | 1.06E+00 | 9.01E-02 | 1.06E+00 |
| ENSG000002 | POM121B | 1.04E-01 | ? | 1.82E+00 | 8.62E-01 | 1.82E+00 |
| ENSG000001 | NPIPA5 | 1.04E-01 | 4.13E-01 | 7.91E-01 | -3.37E-01 | -1.26E+00 |
| ENSG000002 | FOXG1-AS1 | 1.04E-01 | ? | 6.40E-01 | -6.43E-01 | -1.56E+00 |
| ENSG000001 | DRG1 | 1.04E-01 | 4.13E-01 | 1.07E+00 | 9.59E-02 | 1.07E+00 |
| ENSG000001 | PDC | 1.04E-01 | ? | 6.53E-01 | -6.16E-01 | -1.53E+00 |
| ENSG000002 | C6orf15 | 1.04E-01 | ? | 1.89E+00 | 9.16E-01 | 1.89E+00 |
| ENSG000002 | LINC01229 | 1.04E-01 | ? | 5.19E-01 | -9.46E-01 | -1.93E+00 |
| ENSG000001 | PRDX6 | 1.04E-01 | 4.13E-01 | 1.07E+00 | 9.35E-02 | 1.07E+00 |
| ENSG000001 | F3 | 1.04E-01 | 4.13E-01 | 9.32E-01 | -1.02E-01 | -1.07E+00 |
| ENSG000002 | MBTPS1-DT | 1.04E-01 | ? | 2.45E+00 | 1.29E+00 | 2.45E+00 |
| ENSG000002 | MSC-AS1 | 1.04E-01 | 4.13E-01 | 7.87E-01 | -3.46E-01 | -1.27E+00 |
| ENSG000001 | FAM83B | 1.04E-01 | 4.13E-01 | 1.05E+00 | 7.52E-02 | 1.05E+00 |
| ENSG000001 | MIA3 | 1.04E-01 | 4.13E-01 | 9.44E-01 | -8.29E-02 | -1.06E+00 |
| ENSG000002 | --- | 1.04E-01 | ? | 2.21E+00 | 1.14E+00 | 2.21E+00 |
| ENSG000002 | CCPG1 | 1.04E-01 | 4.14E-01 | 9.35E-01 | -9.67E-02 | -1.07E+00 |
| ENSG000001 | ZNF333 | 1.04E-01 | 4.14E-01 | 8.96E-01 | -1.58E-01 | -1.12E+00 |
| ENSG000002 | ECI2-DT | 1.04E-01 | ? | 1.57E+00 | 6.50E-01 | 1.57E+00 |
| ENSG000001 | OR1F1 | 1.04E-01 | ? | 1.46E+00 | 5.44E-01 | 1.46E+00 |
| ENSG000001 | BRMS1L | 1.04E-01 | 4.14E-01 | 1.08E+00 | 1.07E-01 | 1.08E+00 |
| ENSG000001 | HTR7 | 1.04E-01 | 4.14E-01 | 1.07E+00 | 9.69E-02 | 1.07E+00 |
| ENSG000001 | COX7A2 | 1.04E-01 | 4.14E-01 | 1.07E+00 | 9.33E-02 | 1.07E+00 |
| ENSG00000C | VASH1 | 1.04E-01 | 4.14E-01 | 1.24E+00 | 3.13E-01 | 1.24E+00 |
| ENSG000002 | RPS28P7 | 1.04E-01 | ? | 1.47E+00 | 5.60E-01 | 1.47E+00 |
| ENSG000001 | FBXO11 | 1.04E-01 | 4.15E-01 | 9.41E-01 | -8.74E-02 | -1.06E+00 |
| ENSG000002 | OXLD1 | 1.05E-01 | 4.15E-01 | 1.14E+00 | 1.85E-01 | 1.14E+00 |
| ENSG00000C | PDK3 | 1.05E-01 | 4.15E-01 | 9.29E-01 | -1.06E-01 | -1.08E+00 |
| ENSG000001 | TSEN34 | 1.05E-01 | 4.15E-01 | 1.11E+00 | 1.54E-01 | 1.11E+00 |
| ENSG000001 | TOR1B | 1.05E-01 | 4.15E-01 | 9.40E-01 | -8.85E-02 | -1.06E+00 |
| ENSG000001 | SLC25A23 | 1.05E-01 | 4.15E-01 | 9.25E-01 | -1.13E-01 | -1.08E+00 |
| ENSG00000C | PDK4 | 1.05E-01 | ? | 1.83E+00 | 8.71E-01 | 1.83E+00 |
| ENSG000001 | TADA3 | 1.05E-01 | 4.16E-01 | 1.05E+00 | 7.54E-02 | 1.05E+00 |
| ENSG000001 | COG6 | 1.05E-01 | 4.16E-01 | 9.42E-01 | -8.66E-02 | -1.06E+00 |
| ENSG000001 | MANBAL | 1.05E-01 | 4.16E-01 | 1.04E+00 | 6.27E-02 | 1.04E+00 |

| | | | | | | |
|---|---|---|---|---|---|---|
| ENSG000001 | DDX19A | 1.05E-01 | 4.16E-01 | 1.05E+00 | 6.36E-02 | 1.05E+00 |
| ENSG00000C | PPP2R5B | 1.05E-01 | 4.16E-01 | 1.10E+00 | 1.38E-01 | 1.10E+00 |
| ENSG000001 | COA3 | 1.05E-01 | 4.16E-01 | 1.09E+00 | 1.20E-01 | 1.09E+00 |
| ENSG000001 | SSRP1 | 1.05E-01 | 4.16E-01 | 1.05E+00 | 6.47E-02 | 1.05E+00 |
| ENSG000002 | KBTBD11-OT | 1.05E-01 | 4.16E-01 | 1.25E+00 | 3.22E-01 | 1.25E+00 |
| ENSG000001 | RYBP | 1.05E-01 | 4.16E-01 | 9.52E-01 | -7.09E-02 | -1.05E+00 |
| ENSG000001 | LSM14B | 1.06E-01 | 4.17E-01 | 1.06E+00 | 8.18E-02 | 1.06E+00 |
| ENSG00000C | NRIP2 | 1.06E-01 | 4.17E-01 | 7.58E-01 | -3.99E-01 | -1.32E+00 |
| ENSG000002 | --- | 1.06E-01 | ? | 2.98E-01 | -1.75E+00 | -3.36E+00 |
| ENSG000001 | ZDHHC12 | 1.06E-01 | 4.18E-01 | 1.13E+00 | 1.77E-01 | 1.13E+00 |
| ENSG000001 | TRMO | 1.06E-01 | 4.18E-01 | 1.09E+00 | 1.28E-01 | 1.09E+00 |
| ENSG000002 | --- | 1.06E-01 | 4.18E-01 | 7.56E-01 | -4.03E-01 | -1.32E+00 |
| ENSG000001 | ORAI2 | 1.06E-01 | 4.18E-01 | 1.13E+00 | 1.81E-01 | 1.13E+00 |
| ENSG000002 | LINC01503 | 1.06E-01 | 4.18E-01 | 9.16E-01 | -1.27E-01 | -1.09E+00 |
| ENSG000001 | FAAH | 1.06E-01 | 4.18E-01 | 1.12E+00 | 1.58E-01 | 1.12E+00 |
| ENSG000001 | KSR2 | 1.06E-01 | ? | 2.54E+00 | 1.34E+00 | 2.54E+00 |
| ENSG000002 | LNCOC1 | 1.06E-01 | ? | 2.09E+00 | 1.06E+00 | 2.09E+00 |
| ENSG000002 | --- | 1.06E-01 | 4.19E-01 | 7.59E-01 | -3.98E-01 | -1.32E+00 |
| ENSG000001 | POU2F1 | 1.06E-01 | 4.19E-01 | 9.04E-01 | -1.45E-01 | -1.11E+00 |
| ENSG000001 | PRRG1 | 1.06E-01 | 4.19E-01 | 1.07E+00 | 1.01E-01 | 1.07E+00 |
| ENSG000001 | AGPAT5 | 1.06E-01 | 4.19E-01 | 1.06E+00 | 7.75E-02 | 1.06E+00 |
| ENSG00000C | CSNK2A2 | 1.06E-01 | 4.19E-01 | 9.54E-01 | -6.73E-02 | -1.05E+00 |
| ENSG000001 | MAGED2 | 1.06E-01 | 4.19E-01 | 1.09E+00 | 1.24E-01 | 1.09E+00 |
| ENSG000001 | PCSK4 | 1.06E-01 | ? | 6.34E-01 | -6.58E-01 | -1.58E+00 |
| ENSG00000C | BIRC5 | 1.06E-01 | 4.19E-01 | 8.14E-01 | -2.97E-01 | -1.23E+00 |
| ENSG00000C | PKM | 1.06E-01 | 4.19E-01 | 9.67E-01 | -4.77E-02 | -1.03E+00 |
| ENSG000001 | PDCD6IP | 1.06E-01 | 4.19E-01 | 9.55E-01 | -6.69E-02 | -1.05E+00 |
| ENSG000001 | SPRYD7 | 1.07E-01 | 4.19E-01 | 9.49E-01 | -7.50E-02 | -1.05E+00 |
| ENSG000002 | ZBTB47-AS1 | 1.07E-01 | 4.20E-01 | 8.26E-01 | -2.75E-01 | -1.21E+00 |
| ENSG000001 | PTN | 1.07E-01 | ? | 3.87E-01 | -1.37E+00 | -2.59E+00 |
| ENSG000002 | NBPF11 | 1.07E-01 | 4.20E-01 | 8.97E-01 | -1.56E-01 | -1.11E+00 |
| ENSG000001 | ATIC | 1.07E-01 | 4.20E-01 | 1.05E+00 | 6.81E-02 | 1.05E+00 |
| ENSG000001 | BCL7B | 1.07E-01 | 4.20E-01 | 1.08E+00 | 1.13E-01 | 1.08E+00 |
| ENSG000002 | LINC01277 | 1.07E-01 | ? | 6.24E-01 | -6.80E-01 | -1.60E+00 |
| ENSG000002 | LINC01348 | 1.07E-01 | ? | 7.65E-01 | -3.86E-01 | -1.31E+00 |
| ENSG000001 | FAM114A1 | 1.07E-01 | 4.21E-01 | 9.25E-01 | -1.12E-01 | -1.08E+00 |
| ENSG000001 | CAMK2D | 1.07E-01 | 4.21E-01 | 1.10E+00 | 1.39E-01 | 1.10E+00 |
| ENSG000001 | TRUB2 | 1.07E-01 | 4.21E-01 | 1.06E+00 | 8.72E-02 | 1.06E+00 |
| ENSG000001 | ZPR1 | 1.07E-01 | 4.21E-01 | 9.45E-01 | -8.12E-02 | -1.06E+00 |
| ENSG000002 | C1RL-AS1 | 1.07E-01 | 4.21E-01 | 8.84E-01 | -1.79E-01 | -1.13E+00 |
| ENSG000001 | PSMC3IP | 1.07E-01 | 4.21E-01 | 8.50E-01 | -2.35E-01 | -1.18E+00 |
| ENSG00000C | RPL6 | 1.07E-01 | 4.21E-01 | 1.04E+00 | 5.93E-02 | 1.04E+00 |
| ENSG000001 | KDF1 | 1.07E-01 | 4.21E-01 | 1.10E+00 | 1.44E-01 | 1.10E+00 |
| ENSG000001 | SPATS2 | 1.07E-01 | 4.21E-01 | 1.05E+00 | 7.71E-02 | 1.05E+00 |
| ENSG000001 | SIDT2 | 1.07E-01 | 4.21E-01 | 1.08E+00 | 1.09E-01 | 1.08E+00 |

| | | | | | | |
|---|---|---|---|---|---|---|
| ENSG000002 | --- | 1.07E-01 | ? | 1.42E+00 | 5.04E-01 | 1.42E+00 |
| ENSG000001 | PSMD9 | 1.07E-01 | 4.21E-01 | 1.06E+00 | 8.94E-02 | 1.06E+00 |
| ENSG000002 | UBE2FP3 | 1.08E-01 | ? | 6.90E-01 | -5.36E-01 | -1.45E+00 |
| ENSG000002 | --- | 1.08E-01 | ? | 6.79E-01 | -5.58E-01 | -1.47E+00 |
| ENSG000002 | --- | 1.08E-01 | 4.21E-01 | 1.24E+00 | 3.08E-01 | 1.24E+00 |
| ENSG000001 | EP300 | 1.08E-01 | 4.21E-01 | 9.25E-01 | -1.12E-01 | -1.08E+00 |
| ENSG000002 | IFRD2 | 1.08E-01 | 4.21E-01 | 1.09E+00 | 1.26E-01 | 1.09E+00 |
| ENSG000001 | GLRX5 | 1.08E-01 | 4.22E-01 | 1.07E+00 | 1.03E-01 | 1.07E+00 |
| ENSG000001 | NHLH1 | 1.08E-01 | ? | 5.96E-01 | -7.46E-01 | -1.68E+00 |
| ENSG00000C | TESK2 | 1.08E-01 | 4.22E-01 | 1.28E+00 | 3.54E-01 | 1.28E+00 |
| ENSG000001 | UBE2H | 1.08E-01 | 4.22E-01 | 9.66E-01 | -5.00E-02 | -1.04E+00 |
| ENSG000001 | SDHAF3 | 1.08E-01 | 4.22E-01 | 8.98E-01 | -1.55E-01 | -1.11E+00 |
| ENSG000001 | TTN | 1.08E-01 | 4.22E-01 | 8.78E-01 | -1.87E-01 | -1.14E+00 |
| ENSG000002 | --- | 1.08E-01 | 4.22E-01 | 7.44E-01 | -4.27E-01 | -1.34E+00 |
| ENSG000001 | LEPR | 1.08E-01 | 4.22E-01 | 9.19E-01 | -1.23E-01 | -1.09E+00 |
| ENSG000002 | PDCD4-AS1 | 1.08E-01 | 4.22E-01 | 7.81E-01 | -3.56E-01 | -1.28E+00 |
| ENSG000001 | BECN1 | 1.08E-01 | 4.22E-01 | 1.04E+00 | 5.41E-02 | 1.04E+00 |
| ENSG000001 | ATG13 | 1.08E-01 | 4.22E-01 | 1.04E+00 | 6.29E-02 | 1.04E+00 |
| ENSG00000C | TPX2 | 1.08E-01 | 4.23E-01 | 8.99E-01 | -1.54E-01 | -1.11E+00 |
| ENSG000002 | UPK3B | 1.08E-01 | 4.23E-01 | 1.25E+00 | 3.17E-01 | 1.25E+00 |
| ENSG000001 | CCDC51 | 1.08E-01 | 4.23E-01 | 1.09E+00 | 1.18E-01 | 1.09E+00 |
| ENSG000001 | DZIP1L | 1.08E-01 | ? | 5.75E-01 | -7.97E-01 | -1.74E+00 |
| ENSG000002 | --- | 1.08E-01 | ? | 2.60E+00 | 1.38E+00 | 2.60E+00 |
| ENSG000001 | ANKRD52 | 1.08E-01 | 4.23E-01 | 1.07E+00 | 9.25E-02 | 1.07E+00 |
| ENSG000001 | KLF4 | 1.08E-01 | 4.23E-01 | 1.10E+00 | 1.39E-01 | 1.10E+00 |
| ENSG000001 | EMP3 | 1.09E-01 | 4.23E-01 | 1.09E+00 | 1.23E-01 | 1.09E+00 |
| ENSG000001 | C12orf43 | 1.09E-01 | 4.23E-01 | 1.07E+00 | 9.16E-02 | 1.07E+00 |
| ENSG000002 | SPDYE18 | 1.09E-01 | ? | 6.86E-01 | -5.43E-01 | -1.46E+00 |
| ENSG000001 | PRDX5 | 1.09E-01 | 4.24E-01 | 1.07E+00 | 9.43E-02 | 1.07E+00 |
| ENSG000001 | VAPA | 1.09E-01 | 4.24E-01 | 9.59E-01 | -6.06E-02 | -1.04E+00 |
| ENSG000001 | NR2C2AP | 1.09E-01 | 4.24E-01 | 1.13E+00 | 1.76E-01 | 1.13E+00 |
| ENSG000002 | --- | 1.09E-01 | ? | 4.40E-01 | -1.18E+00 | -2.27E+00 |
| ENSG000001 | LINC00319 | 1.09E-01 | 4.24E-01 | 1.19E+00 | 2.46E-01 | 1.19E+00 |
| ENSG000001 | ERBB4 | 1.09E-01 | ? | 1.50E+00 | 5.88E-01 | 1.50E+00 |
| ENSG000001 | LINC00324 | 1.09E-01 | ? | 1.63E+00 | 7.01E-01 | 1.63E+00 |
| ENSG00000C | RNH1 | 1.09E-01 | 4.24E-01 | 1.10E+00 | 1.42E-01 | 1.10E+00 |
| ENSG000001 | GVQW3 | 1.09E-01 | 4.24E-01 | 7.96E-01 | -3.29E-01 | -1.26E+00 |
| ENSG000001 | TMEM161B | 1.09E-01 | 4.24E-01 | 9.22E-01 | -1.17E-01 | -1.08E+00 |
| ENSG000001 | PELI2 | 1.09E-01 | ? | 1.83E+00 | 8.70E-01 | 1.83E+00 |
| ENSG000001 | CNTNAP3 | 1.09E-01 | 4.24E-01 | 9.17E-01 | -1.25E-01 | -1.09E+00 |
| ENSG000001 | SMPDL3A | 1.09E-01 | 4.24E-01 | 1.15E+00 | 1.98E-01 | 1.15E+00 |
| ENSG000001 | KRT19 | 1.09E-01 | 4.24E-01 | 9.42E-01 | -8.65E-02 | -1.06E+00 |
| ENSG000001 | GNL3 | 1.09E-01 | 4.24E-01 | 9.36E-01 | -9.56E-02 | -1.07E+00 |
| ENSG000002 | --- | 1.09E-01 | ? | 5.10E-01 | -9.72E-01 | -1.96E+00 |
| ENSG000001 | TEF | 1.09E-01 | 4.24E-01 | 1.15E+00 | 2.00E-01 | 1.15E+00 |

| | | | | | | |
|---|---|---|---|---|---|---|
| ENSG000001 | GPSM2 | 1.09E-01 | 4.24E-01 | 1.06E+00 | 8.15E-02 | 1.06E+00 |
| ENSG000001 | SYBU | 1.09E-01 | 4.25E-01 | 1.11E+00 | 1.53E-01 | 1.11E+00 |
| ENSG000002 | KRTAP5-AS1 | 1.09E-01 | ? | 6.35E-01 | -6.56E-01 | -1.58E+00 |
| ENSG000000 | HUWE1 | 1.09E-01 | 4.25E-01 | 9.15E-01 | -1.28E-01 | -1.09E+00 |
| ENSG000002 | ZNHIT3 | 1.09E-01 | 4.25E-01 | 1.06E+00 | 8.65E-02 | 1.06E+00 |
| ENSG000002 | --- | 1.10E-01 | ? | 1.52E+00 | 6.02E-01 | 1.52E+00 |
| ENSG000001 | TMEM42 | 1.10E-01 | 4.25E-01 | 1.13E+00 | 1.74E-01 | 1.13E+00 |
| ENSG000001 | TMEM45B | 1.10E-01 | ? | 7.48E-01 | -4.20E-01 | -1.34E+00 |
| ENSG000002 | --- | 1.10E-01 | ? | 2.25E+00 | 1.17E+00 | 2.25E+00 |
| ENSG000001 | EXOC7 | 1.10E-01 | 4.25E-01 | 1.05E+00 | 6.72E-02 | 1.05E+00 |
| ENSG000002 | --- | 1.10E-01 | ? | 6.39E-01 | -6.46E-01 | -1.56E+00 |
| ENSG000002 | MATR3 | 1.10E-01 | 4.25E-01 | 1.07E+00 | 9.30E-02 | 1.07E+00 |
| ENSG000002 | --- | 1.10E-01 | ? | 1.38E+00 | 4.65E-01 | 1.38E+00 |
| ENSG000002 | --- | 1.10E-01 | 4.26E-01 | 1.16E+00 | 2.12E-01 | 1.16E+00 |
| ENSG000002 | PBX2 | 1.10E-01 | 4.26E-01 | 1.05E+00 | 7.42E-02 | 1.05E+00 |
| ENSG000001 | ECHS1 | 1.10E-01 | 4.26E-01 | 1.05E+00 | 6.90E-02 | 1.05E+00 |
| ENSG000001 | FBXL18 | 1.10E-01 | 4.26E-01 | 1.08E+00 | 1.17E-01 | 1.08E+00 |
| ENSG000001 | ITGB2 | 1.10E-01 | 4.26E-01 | 8.25E-01 | -2.77E-01 | -1.21E+00 |
| ENSG000001 | DOK3 | 1.10E-01 | 4.26E-01 | 8.97E-01 | -1.57E-01 | -1.12E+00 |
| ENSG000001 | TMEM123 | 1.10E-01 | 4.26E-01 | 9.47E-01 | -7.90E-02 | -1.06E+00 |
| ENSG000001 | NUCB1 | 1.10E-01 | 4.26E-01 | 1.07E+00 | 9.22E-02 | 1.07E+00 |
| ENSG000001 | IL13RA1 | 1.10E-01 | 4.27E-01 | 1.05E+00 | 7.31E-02 | 1.05E+00 |
| ENSG000001 | TPI1 | 1.10E-01 | 4.27E-01 | 1.04E+00 | 5.76E-02 | 1.04E+00 |
| ENSG000001 | MYO1C | 1.10E-01 | 4.27E-01 | 1.06E+00 | 8.21E-02 | 1.06E+00 |
| ENSG000001 | RMND1 | 1.10E-01 | 4.27E-01 | 1.06E+00 | 7.75E-02 | 1.06E+00 |
| ENSG000001 | TBC1D16 | 1.10E-01 | 4.27E-01 | 1.07E+00 | 9.72E-02 | 1.07E+00 |
| ENSG000001 | SRRM2 | 1.10E-01 | 4.27E-01 | 9.40E-01 | -8.98E-02 | -1.06E+00 |
| ENSG000001 | TOE1 | 1.10E-01 | 4.27E-01 | 1.13E+00 | 1.76E-01 | 1.13E+00 |
| ENSG000002 | GAS2L1P2 | 1.11E-01 | ? | 2.24E+00 | 1.16E+00 | 2.24E+00 |
| ENSG000002 | --- | 1.11E-01 | 4.27E-01 | 1.16E+00 | 2.18E-01 | 1.16E+00 |
| ENSG000001 | ZNF66 | 1.11E-01 | 4.27E-01 | 8.45E-01 | -2.44E-01 | -1.18E+00 |
| ENSG000001 | MAP3K3 | 1.11E-01 | 4.27E-01 | 1.05E+00 | 7.32E-02 | 1.05E+00 |
| ENSG000001 | ERG28 | 1.11E-01 | 4.27E-01 | 1.06E+00 | 8.23E-02 | 1.06E+00 |
| ENSG000002 | NCK1-DT | 1.11E-01 | 4.27E-01 | 1.20E+00 | 2.60E-01 | 1.20E+00 |
| ENSG000000 | AGO1 | 1.11E-01 | 4.27E-01 | 1.05E+00 | 7.21E-02 | 1.05E+00 |
| ENSG000001 | MINPP1 | 1.11E-01 | 4.27E-01 | 1.07E+00 | 9.39E-02 | 1.07E+00 |
| ENSG000002 | STK19B | 1.11E-01 | ? | 1.83E+00 | 8.73E-01 | 1.83E+00 |
| ENSG000001 | AQP3 | 1.11E-01 | 4.27E-01 | 7.90E-01 | -3.41E-01 | -1.27E+00 |
| ENSG000001 | ANKRD27 | 1.11E-01 | 4.27E-01 | 1.06E+00 | 8.15E-02 | 1.06E+00 |
| ENSG000000 | KLHL22 | 1.11E-01 | 4.27E-01 | 1.13E+00 | 1.73E-01 | 1.13E+00 |
| ENSG000001 | TICAM1 | 1.11E-01 | 4.27E-01 | 1.08E+00 | 1.11E-01 | 1.08E+00 |
| ENSG000001 | RIN2 | 1.11E-01 | 4.27E-01 | 1.04E+00 | 5.78E-02 | 1.04E+00 |
| ENSG000001 | COX5B | 1.11E-01 | 4.27E-01 | 1.08E+00 | 1.15E-01 | 1.08E+00 |
| ENSG000001 | SDAD1 | 1.11E-01 | 4.27E-01 | 9.47E-01 | -7.78E-02 | -1.06E+00 |
| ENSG000001 | PARP16 | 1.11E-01 | 4.27E-01 | 1.12E+00 | 1.61E-01 | 1.12E+00 |

| | | | | | | |
|---|---|---|---|---|---|---|
| ENSG000001 | AP1S3 | 1.11E-01 | 4.27E-01 | 1.07E+00 | 1.03E-01 | 1.07E+00 |
| ENSG000002 | LINP1 | 1.11E-01 | 4.28E-01 | 8.73E-01 | -1.96E-01 | -1.15E+00 |
| ENSG00000C | AFF4 | 1.11E-01 | 4.28E-01 | 9.38E-01 | -9.30E-02 | -1.07E+00 |
| ENSG000001 | ABHD14B | 1.11E-01 | 4.28E-01 | 1.09E+00 | 1.22E-01 | 1.09E+00 |
| ENSG000001 | TUSC3 | 1.11E-01 | 4.28E-01 | 1.06E+00 | 8.02E-02 | 1.06E+00 |
| ENSG000001 | HES6 | 1.11E-01 | ? | 1.48E+00 | 5.68E-01 | 1.48E+00 |
| ENSG000001 | NPIPB3 | 1.11E-01 | 4.28E-01 | 8.75E-01 | -1.93E-01 | -1.14E+00 |
| ENSG000001 | HINT2 | 1.11E-01 | 4.28E-01 | 1.19E+00 | 2.45E-01 | 1.19E+00 |
| ENSG000002 | --- | 1.12E-01 | ? | 1.62E+00 | 6.96E-01 | 1.62E+00 |
| ENSG000001 | ZNF233 | 1.12E-01 | ? | 1.53E+00 | 6.13E-01 | 1.53E+00 |
| ENSG000001 | GLT8D2 | 1.12E-01 | 4.28E-01 | 1.13E+00 | 1.77E-01 | 1.13E+00 |
| ENSG000001 | CNN3 | 1.12E-01 | 4.28E-01 | 9.28E-01 | -1.08E-01 | -1.08E+00 |
| ENSG000001 | GSTZ1 | 1.12E-01 | 4.28E-01 | 1.10E+00 | 1.42E-01 | 1.10E+00 |
| ENSG000001 | SENP3 | 1.12E-01 | 4.28E-01 | 1.06E+00 | 8.21E-02 | 1.06E+00 |
| ENSG000001 | LRP8 | 1.12E-01 | 4.28E-01 | 9.26E-01 | -1.11E-01 | -1.08E+00 |
| ENSG000001 | RUVBL1 | 1.12E-01 | 4.28E-01 | 1.06E+00 | 8.57E-02 | 1.06E+00 |
| ENSG000001 | UBE2Q2 | 1.12E-01 | 4.28E-01 | 1.04E+00 | 5.39E-02 | 1.04E+00 |
| ENSG000001 | SLC28A2 | 1.12E-01 | ? | 2.52E+00 | 1.34E+00 | 2.52E+00 |
| ENSG000002 | STMP1 | 1.12E-01 | 4.28E-01 | 1.05E+00 | 6.97E-02 | 1.05E+00 |
| ENSG000001 | CASZ1 | 1.12E-01 | 4.28E-01 | 1.09E+00 | 1.18E-01 | 1.09E+00 |
| ENSG000001 | POLM | 1.12E-01 | 4.29E-01 | 9.04E-01 | -1.46E-01 | -1.11E+00 |
| ENSG000001 | LINC00265 | 1.12E-01 | 4.29E-01 | 8.96E-01 | -1.59E-01 | -1.12E+00 |
| ENSG000001 | FBXL15 | 1.12E-01 | 4.29E-01 | 1.23E+00 | 3.01E-01 | 1.23E+00 |
| ENSG00000C | ATG2B | 1.12E-01 | 4.29E-01 | 9.42E-01 | -8.60E-02 | -1.06E+00 |
| ENSG000001 | CFAP251 | 1.12E-01 | 4.29E-01 | 1.11E+00 | 1.46E-01 | 1.11E+00 |
| ENSG000002 | --- | 1.12E-01 | ? | 7.51E-01 | -4.14E-01 | -1.33E+00 |
| ENSG000001 | NLRP10 | 1.12E-01 | 4.29E-01 | 8.23E-01 | -2.81E-01 | -1.21E+00 |
| ENSG000001 | CHM | 1.12E-01 | 4.29E-01 | 9.40E-01 | -8.93E-02 | -1.06E+00 |
| ENSG000001 | RASL11A | 1.12E-01 | ? | 1.33E+00 | 4.15E-01 | 1.33E+00 |
| ENSG000001 | ACOT1 | 1.12E-01 | 4.29E-01 | 1.32E+00 | 4.01E-01 | 1.32E+00 |
| ENSG000002 | --- | 1.12E-01 | 4.29E-01 | 8.65E-01 | -2.09E-01 | -1.16E+00 |
| ENSG000001 | GMPR2 | 1.12E-01 | 4.29E-01 | 1.08E+00 | 1.09E-01 | 1.08E+00 |
| ENSG000001 | THUMPD2 | 1.12E-01 | 4.29E-01 | 9.08E-01 | -1.39E-01 | -1.10E+00 |
| ENSG000001 | TRAPPC10 | 1.12E-01 | 4.29E-01 | 9.61E-01 | -5.79E-02 | -1.04E+00 |
| ENSG00000C | EXD2 | 1.12E-01 | 4.29E-01 | 9.59E-01 | -6.01E-02 | -1.04E+00 |
| ENSG000001 | PIK3CD | 1.12E-01 | 4.29E-01 | 9.37E-01 | -9.33E-02 | -1.07E+00 |
| ENSG000001 | CARF | 1.13E-01 | 4.30E-01 | 8.98E-01 | -1.55E-01 | -1.11E+00 |
| ENSG000002 | PPIAP91 | 1.13E-01 | ? | 4.41E-01 | -1.18E+00 | -2.27E+00 |
| ENSG000001 | PRKN | 1.13E-01 | ? | 1.60E+00 | 6.82E-01 | 1.60E+00 |
| ENSG000001 | ACP6 | 1.13E-01 | 4.30E-01 | 9.02E-01 | -1.49E-01 | -1.11E+00 |
| ENSG000002 | --- | 1.13E-01 | ? | 4.85E-01 | -1.04E+00 | -2.06E+00 |
| ENSG000001 | CA2 | 1.13E-01 | 4.30E-01 | 1.08E+00 | 1.06E-01 | 1.08E+00 |
| ENSG000001 | ARHGAP11A | 1.13E-01 | 4.30E-01 | 8.74E-01 | -1.95E-01 | -1.14E+00 |
| ENSG000002 | --- | 1.13E-01 | ? | 5.06E-01 | -9.82E-01 | -1.98E+00 |
| ENSG000002 | TAX1BP3 | 1.13E-01 | 4.30E-01 | 1.13E+00 | 1.83E-01 | 1.13E+00 |

| | | | | | | |
|---|---|---|---|---|---|---|
| ENSG000002 | RNFT1-DT | 1.13E-01 | ? | 6.95E-01 | -5.24E-01 | -1.44E+00 |
| ENSG000001 | ELAPOR1 | 1.13E-01 | 4.31E-01 | 1.15E+00 | 2.04E-01 | 1.15E+00 |
| ENSG000001 | RBIS | 1.13E-01 | 4.31E-01 | 9.34E-01 | -9.87E-02 | -1.07E+00 |
| ENSG000002 | --- | 1.13E-01 | ? | 6.38E-01 | -6.49E-01 | -1.57E+00 |
| ENSG000001 | TOR2A | 1.13E-01 | 4.31E-01 | 1.12E+00 | 1.69E-01 | 1.12E+00 |
| ENSG000002 | --- | 1.13E-01 | ? | 5.85E-01 | -7.73E-01 | -1.71E+00 |
| ENSG000002 | LINC01068 | 1.13E-01 | ? | 4.77E-01 | -1.07E+00 | -2.09E+00 |
| ENSG000001 | TLK2 | 1.13E-01 | 4.31E-01 | 9.36E-01 | -9.62E-02 | -1.07E+00 |
| ENSG00000C | SPTLC1 | 1.13E-01 | 4.31E-01 | 9.63E-01 | -5.37E-02 | -1.04E+00 |
| ENSG00000C | AACS | 1.13E-01 | 4.31E-01 | 1.07E+00 | 9.11E-02 | 1.07E+00 |
| ENSG000001 | NME5 | 1.13E-01 | 4.31E-01 | 1.20E+00 | 2.60E-01 | 1.20E+00 |
| ENSG000001 | RBM45 | 1.13E-01 | 4.31E-01 | 9.33E-01 | -9.96E-02 | -1.07E+00 |
| ENSG000002 | --- | 1.13E-01 | 4.31E-01 | 7.98E-01 | -3.26E-01 | -1.25E+00 |
| ENSG000001 | MOCS3 | 1.13E-01 | 4.31E-01 | 1.08E+00 | 1.16E-01 | 1.08E+00 |
| ENSG000001 | ANKRD9 | 1.13E-01 | 4.31E-01 | 1.12E+00 | 1.62E-01 | 1.12E+00 |
| ENSG000001 | NOB1 | 1.14E-01 | 4.31E-01 | 1.06E+00 | 8.59E-02 | 1.06E+00 |
| ENSG000002 | --- | 1.14E-01 | 4.31E-01 | 7.23E-01 | -4.68E-01 | -1.38E+00 |
| ENSG000001 | SRRD | 1.14E-01 | 4.31E-01 | 1.07E+00 | 9.16E-02 | 1.07E+00 |
| ENSG000001 | PDZD8 | 1.14E-01 | 4.31E-01 | 9.39E-01 | -9.03E-02 | -1.06E+00 |
| ENSG000001 | EVL | 1.14E-01 | 4.31E-01 | 1.10E+00 | 1.36E-01 | 1.10E+00 |
| ENSG000002 | RGS5 | 1.14E-01 | ? | 7.39E-01 | -4.36E-01 | -1.35E+00 |
| ENSG000001 | HSPA8 | 1.14E-01 | 4.31E-01 | 9.45E-01 | -8.22E-02 | -1.06E+00 |
| ENSG000001 | SVEP1 | 1.14E-01 | 4.31E-01 | 1.13E+00 | 1.81E-01 | 1.13E+00 |
| ENSG000001 | NECAP2 | 1.14E-01 | 4.31E-01 | 1.06E+00 | 8.71E-02 | 1.06E+00 |
| ENSG000001 | NRP2 | 1.14E-01 | 4.31E-01 | 9.21E-01 | -1.18E-01 | -1.09E+00 |
| ENSG000001 | TRPM4 | 1.14E-01 | 4.31E-01 | 1.12E+00 | 1.66E-01 | 1.12E+00 |
| ENSG000001 | RAP1GAP2 | 1.14E-01 | 4.31E-01 | 1.06E+00 | 8.53E-02 | 1.06E+00 |
| ENSG000002 | --- | 1.14E-01 | ? | 1.65E+00 | 7.22E-01 | 1.65E+00 |
| ENSG000001 | GRSF1 | 1.14E-01 | 4.32E-01 | 9.61E-01 | -5.75E-02 | -1.04E+00 |
| ENSG000002 | BLOC1S5-TXI | 1.14E-01 | 4.32E-01 | 1.32E+00 | 3.96E-01 | 1.32E+00 |
| ENSG000001 | PLEKHH2 | 1.14E-01 | ? | 6.87E-01 | -5.42E-01 | -1.46E+00 |
| ENSG000002 | --- | 1.14E-01 | ? | 6.03E-01 | -7.30E-01 | -1.66E+00 |
| ENSG000002 | --- | 1.14E-01 | 4.32E-01 | 1.25E+00 | 3.22E-01 | 1.25E+00 |
| ENSG000001 | RNF145 | 1.14E-01 | 4.32E-01 | 9.35E-01 | -9.70E-02 | -1.07E+00 |
| ENSG000001 | TMEM97 | 1.14E-01 | 4.32E-01 | 1.10E+00 | 1.31E-01 | 1.10E+00 |
| ENSG000001 | SREK1 | 1.14E-01 | 4.32E-01 | 9.29E-01 | -1.07E-01 | -1.08E+00 |
| ENSG000001 | SPRY2 | 1.14E-01 | 4.32E-01 | 9.16E-01 | -1.27E-01 | -1.09E+00 |
| ENSG000002 | --- | 1.14E-01 | ? | 2.86E-01 | -1.80E+00 | -3.49E+00 |
| ENSG000001 | SCAND1 | 1.14E-01 | 4.32E-01 | 1.18E+00 | 2.35E-01 | 1.18E+00 |
| ENSG000001 | RPS23 | 1.14E-01 | 4.32E-01 | 1.04E+00 | 5.94E-02 | 1.04E+00 |
| ENSG000001 | P2RX4 | 1.14E-01 | 4.32E-01 | 1.15E+00 | 2.03E-01 | 1.15E+00 |
| ENSG00000C | CRLS1 | 1.14E-01 | 4.32E-01 | 9.31E-01 | -1.03E-01 | -1.07E+00 |
| ENSG000001 | RPL8 | 1.14E-01 | 4.32E-01 | 1.07E+00 | 9.29E-02 | 1.07E+00 |
| ENSG000001 | LRRN4 | 1.14E-01 | 4.32E-01 | 8.50E-01 | -2.35E-01 | -1.18E+00 |
| ENSG000001 | ANKH | 1.14E-01 | 4.32E-01 | 1.05E+00 | 6.70E-02 | 1.05E+00 |

| | | | | | | |
|---|---|---|---|---|---|---|
| ENSG000001 | KIAA0319L | 1.14E-01 | 4.32E-01 | 1.05E+00 | 7.20E-02 | 1.05E+00 |
| ENSG000002 | --- | 1.15E-01 | ? | 5.11E-01 | -9.68E-01 | -1.96E+00 |
| ENSG00000C | SMARCB1 | 1.15E-01 | 4.32E-01 | 1.05E+00 | 6.61E-02 | 1.05E+00 |
| ENSG000001 | OLFML3 | 1.15E-01 | ? | 1.35E+00 | 4.31E-01 | 1.35E+00 |
| ENSG000001 | SNRNP35 | 1.15E-01 | 4.32E-01 | 9.11E-01 | -1.34E-01 | -1.10E+00 |
| ENSG000002 | --- | 1.15E-01 | ? | 5.91E-01 | -7.60E-01 | -1.69E+00 |
| ENSG000001 | IL20RB | 1.15E-01 | 4.32E-01 | 1.08E+00 | 1.08E-01 | 1.08E+00 |
| ENSG000002 | RBMS3-AS3 | 1.15E-01 | ? | 2.36E+00 | 1.24E+00 | 2.36E+00 |
| ENSG000001 | DTD2 | 1.15E-01 | 4.33E-01 | 1.09E+00 | 1.25E-01 | 1.09E+00 |
| ENSG000002 | --- | 1.15E-01 | ? | 5.00E-01 | -9.99E-01 | -2.00E+00 |
| ENSG000001 | TSHZ1 | 1.15E-01 | 4.33E-01 | 1.09E+00 | 1.26E-01 | 1.09E+00 |
| ENSG000002 | CHKB-DT | 1.15E-01 | 4.33E-01 | 1.25E+00 | 3.19E-01 | 1.25E+00 |
| ENSG000001 | SRSF5 | 1.15E-01 | 4.33E-01 | 9.15E-01 | -1.28E-01 | -1.09E+00 |
| ENSG000001 | PGLYRP3 | 1.15E-01 | ? | 1.35E+00 | 4.29E-01 | 1.35E+00 |
| ENSG000001 | PPP1R9B | 1.15E-01 | 4.33E-01 | 9.49E-01 | -7.53E-02 | -1.05E+00 |
| ENSG00000C | KIF9 | 1.15E-01 | 4.33E-01 | 8.66E-01 | -2.08E-01 | -1.15E+00 |
| ENSG000002 | --- | 1.15E-01 | 4.33E-01 | 8.72E-01 | -1.98E-01 | -1.15E+00 |
| ENSG000001 | PFKFB3 | 1.15E-01 | 4.33E-01 | 8.99E-01 | -1.54E-01 | -1.11E+00 |
| ENSG000002 | --- | 1.15E-01 | ? | 1.51E+00 | 5.93E-01 | 1.51E+00 |
| ENSG000001 | SLC46A3 | 1.15E-01 | 4.33E-01 | 9.34E-01 | -9.89E-02 | -1.07E+00 |
| ENSG000001 | SUSD6 | 1.15E-01 | 4.33E-01 | 1.07E+00 | 9.84E-02 | 1.07E+00 |
| ENSG000002 | BAIAP2-DT | 1.15E-01 | 4.33E-01 | 1.10E+00 | 1.33E-01 | 1.10E+00 |
| ENSG000001 | MYL12B | 1.15E-01 | 4.33E-01 | 1.04E+00 | 5.87E-02 | 1.04E+00 |
| ENSG000001 | ACE2 | 1.15E-01 | 4.34E-01 | 1.25E+00 | 3.24E-01 | 1.25E+00 |
| ENSG000001 | SEPTIN11 | 1.16E-01 | 4.34E-01 | 9.56E-01 | -6.54E-02 | -1.05E+00 |
| ENSG000001 | KRBA1 | 1.16E-01 | 4.34E-01 | 1.13E+00 | 1.82E-01 | 1.13E+00 |
| ENSG000001 | CIB2 | 1.16E-01 | 4.34E-01 | 1.19E+00 | 2.51E-01 | 1.19E+00 |
| ENSG000001 | RTKN | 1.16E-01 | 4.34E-01 | 1.07E+00 | 9.61E-02 | 1.07E+00 |
| ENSG000001 | RETREG2 | 1.16E-01 | 4.34E-01 | 1.07E+00 | 9.51E-02 | 1.07E+00 |
| ENSG000001 | MTMR9 | 1.16E-01 | 4.34E-01 | 1.05E+00 | 7.23E-02 | 1.05E+00 |
| ENSG000002 | CKAP2-DT | 1.16E-01 | ? | 1.71E+00 | 7.74E-01 | 1.71E+00 |
| ENSG000001 | PLEKHJ1 | 1.16E-01 | 4.35E-01 | 1.13E+00 | 1.82E-01 | 1.13E+00 |
| ENSG000001 | RAB11FIP5 | 1.16E-01 | 4.35E-01 | 1.07E+00 | 9.63E-02 | 1.07E+00 |
| ENSG000001 | KCNH1 | 1.16E-01 | ? | 1.77E+00 | 8.20E-01 | 1.77E+00 |
| ENSG000002 | NCOA4 | 1.16E-01 | 4.35E-01 | 1.04E+00 | 6.10E-02 | 1.04E+00 |
| ENSG00000C | COMT | 1.16E-01 | 4.35E-01 | 1.09E+00 | 1.29E-01 | 1.09E+00 |
| ENSG000002 | --- | 1.16E-01 | ? | 6.51E-01 | -6.19E-01 | -1.54E+00 |
| ENSG000001 | ZBTB41 | 1.16E-01 | 4.35E-01 | 9.33E-01 | -1.00E-01 | -1.07E+00 |
| ENSG000001 | TAOK3 | 1.16E-01 | 4.35E-01 | 9.24E-01 | -1.14E-01 | -1.08E+00 |
| ENSG000002 | --- | 1.16E-01 | ? | 4.13E-01 | -1.28E+00 | -2.42E+00 |
| ENSG000002 | HNRNPA3P6 | 1.16E-01 | 4.35E-01 | 8.30E-01 | -2.69E-01 | -1.20E+00 |
| ENSG000002 | INSYN1 | 1.16E-01 | ? | 1.43E+00 | 5.14E-01 | 1.43E+00 |
| ENSG000001 | CYP2C18 | 1.16E-01 | ? | 1.98E+00 | 9.83E-01 | 1.98E+00 |
| ENSG00000C | FAM234B | 1.16E-01 | 4.36E-01 | 1.11E+00 | 1.47E-01 | 1.11E+00 |
| ENSG000001 | CAP1 | 1.16E-01 | 4.36E-01 | 9.66E-01 | -4.94E-02 | -1.03E+00 |

| | | | | | | |
|---|---|---|---|---|---|---|
| ENSG000002 | POC1B-AS1 | 1.16E-01 | 4.36E-01 | 8.41E-01 | -2.50E-01 | -1.19E+00 |
| ENSG00000C | MPC1 | 1.17E-01 | 4.36E-01 | 9.17E-01 | -1.25E-01 | -1.09E+00 |
| ENSG000002 | INE1 | 1.17E-01 | 4.36E-01 | 7.85E-01 | -3.49E-01 | -1.27E+00 |
| ENSG000002 | MSL3P1 | 1.17E-01 | 4.36E-01 | 8.35E-01 | -2.60E-01 | -1.20E+00 |
| ENSG00000C | ACSM3 | 1.17E-01 | ? | 1.44E+00 | 5.22E-01 | 1.44E+00 |
| ENSG000001 | PLEKHA1 | 1.17E-01 | 4.36E-01 | 1.05E+00 | 7.20E-02 | 1.05E+00 |
| ENSG000001 | ABCE1 | 1.17E-01 | 4.36E-01 | 9.65E-01 | -5.21E-02 | -1.04E+00 |
| ENSG000002 | SMIM3 | 1.17E-01 | 4.36E-01 | 1.12E+00 | 1.68E-01 | 1.12E+00 |
| ENSG000001 | PSMD4 | 1.17E-01 | 4.36E-01 | 1.05E+00 | 6.89E-02 | 1.05E+00 |
| ENSG000001 | USP11 | 1.17E-01 | 4.37E-01 | 1.06E+00 | 8.89E-02 | 1.06E+00 |
| ENSG00000C | TTC17 | 1.17E-01 | 4.37E-01 | 9.50E-01 | -7.44E-02 | -1.05E+00 |
| ENSG000001 | MSL2 | 1.17E-01 | 4.37E-01 | 9.40E-01 | -8.92E-02 | -1.06E+00 |
| ENSG000002 | --- | 1.17E-01 | ? | 2.05E+00 | 1.04E+00 | 2.05E+00 |
| ENSG000002 | RN7SKP16 | 1.17E-01 | ? | 4.83E-01 | -1.05E+00 | -2.07E+00 |
| ENSG000002 | --- | 1.17E-01 | ? | 2.79E+00 | 1.48E+00 | 2.79E+00 |
| ENSG000002 | --- | 1.17E-01 | 4.37E-01 | 7.92E-01 | -3.36E-01 | -1.26E+00 |
| ENSG000001 | CNOT9 | 1.17E-01 | 4.37E-01 | 1.05E+00 | 7.65E-02 | 1.05E+00 |
| ENSG000001 | SASS6 | 1.17E-01 | 4.37E-01 | 9.02E-01 | -1.49E-01 | -1.11E+00 |
| ENSG000001 | DNAJC4 | 1.17E-01 | 4.37E-01 | 1.10E+00 | 1.34E-01 | 1.10E+00 |
| ENSG000001 | FBXO5 | 1.18E-01 | 4.38E-01 | 8.40E-01 | -2.51E-01 | -1.19E+00 |
| ENSG000002 | --- | 1.18E-01 | ? | 1.37E+00 | 4.53E-01 | 1.37E+00 |
| ENSG000001 | LAMA1 | 1.18E-01 | 4.38E-01 | 1.16E+00 | 2.10E-01 | 1.16E+00 |
| ENSG000001 | ATG4D | 1.18E-01 | 4.38E-01 | 1.14E+00 | 1.90E-01 | 1.14E+00 |
| ENSG000001 | NCAM1 | 1.18E-01 | ? | 5.62E-01 | -8.31E-01 | -1.78E+00 |
| ENSG000001 | FGF14 | 1.18E-01 | ? | 6.18E-01 | -6.94E-01 | -1.62E+00 |
| ENSG00000C | FAS | 1.18E-01 | 4.38E-01 | 1.05E+00 | 6.65E-02 | 1.05E+00 |
| ENSG000002 | --- | 1.18E-01 | ? | 5.68E-01 | -8.16E-01 | -1.76E+00 |
| ENSG000001 | TMEM60 | 1.18E-01 | 4.39E-01 | 1.10E+00 | 1.31E-01 | 1.10E+00 |
| ENSG000001 | MT-ND5 | 1.18E-01 | 4.39E-01 | 8.41E-01 | -2.50E-01 | -1.19E+00 |
| ENSG00000C | PGS1 | 1.18E-01 | 4.39E-01 | 1.09E+00 | 1.30E-01 | 1.09E+00 |
| ENSG00000C | RRAGB | 1.18E-01 | 4.39E-01 | 1.10E+00 | 1.41E-01 | 1.10E+00 |
| ENSG000002 | PCDHGA6 | 1.18E-01 | 4.39E-01 | 1.28E+00 | 3.56E-01 | 1.28E+00 |
| ENSG000001 | CNIH4 | 1.18E-01 | 4.39E-01 | 1.05E+00 | 7.53E-02 | 1.05E+00 |
| ENSG000002 | --- | 1.18E-01 | ? | 5.32E-01 | -9.11E-01 | -1.88E+00 |
| ENSG000001 | PLCD3 | 1.18E-01 | 4.39E-01 | 1.08E+00 | 1.08E-01 | 1.08E+00 |
| ENSG000001 | SLC39A10 | 1.18E-01 | 4.39E-01 | 9.31E-01 | -1.03E-01 | -1.07E+00 |
| ENSG000002 | --- | 1.18E-01 | ? | 7.25E-01 | -4.65E-01 | -1.38E+00 |
| ENSG000001 | DNAJB1 | 1.18E-01 | 4.40E-01 | 1.06E+00 | 8.01E-02 | 1.06E+00 |
| ENSG000002 | DUBR | 1.18E-01 | 4.40E-01 | 1.09E+00 | 1.19E-01 | 1.09E+00 |
| ENSG000001 | TAGLN2 | 1.18E-01 | 4.40E-01 | 1.07E+00 | 9.50E-02 | 1.07E+00 |
| ENSG000001 | SF3B4 | 1.19E-01 | 4.40E-01 | 9.55E-01 | -6.58E-02 | -1.05E+00 |
| ENSG000001 | CELF5 | 1.19E-01 | ? | 1.33E+00 | 4.13E-01 | 1.33E+00 |
| ENSG000001 | MAN1A1 | 1.19E-01 | 4.40E-01 | 9.01E-01 | -1.50E-01 | -1.11E+00 |
| ENSG000001 | NUP58 | 1.19E-01 | 4.40E-01 | 1.05E+00 | 6.95E-02 | 1.05E+00 |
| ENSG000001 | ALDH1A1 | 1.19E-01 | ? | 6.85E-01 | -5.45E-01 | -1.46E+00 |

| | | | | | | |
|---|---|---|---|---|---|---|
| ENSG000001 | SARDH | 1.19E-01 | ? | 1.43E+00 | 5.14E-01 | 1.43E+00 |
| ENSG000002 | SMU1P1 | 1.19E-01 | ? | 2.11E+00 | 1.08E+00 | 2.11E+00 |
| ENSG000002 | LINC01001 | 1.19E-01 | 4.41E-01 | 8.52E-01 | -2.32E-01 | -1.17E+00 |
| ENSG000001 | LTV1 | 1.19E-01 | 4.41E-01 | 9.38E-01 | -9.25E-02 | -1.07E+00 |
| ENSG00000C | CD200 | 1.19E-01 | ? | 4.85E-01 | -1.04E+00 | -2.06E+00 |
| ENSG000001 | ZC4H2 | 1.19E-01 | 4.42E-01 | 1.10E+00 | 1.33E-01 | 1.10E+00 |
| ENSG00000C | FNIP2 | 1.19E-01 | 4.42E-01 | 9.01E-01 | -1.51E-01 | -1.11E+00 |
| ENSG000001 | HSP90AB3P | 1.19E-01 | 4.42E-01 | 7.91E-01 | -3.38E-01 | -1.26E+00 |
| ENSG000002 | ATP5MC1P4 | 1.19E-01 | ? | 1.62E+00 | 6.98E-01 | 1.62E+00 |
| ENSG000001 | KIAA0930 | 1.19E-01 | 4.42E-01 | 1.07E+00 | 9.70E-02 | 1.07E+00 |
| ENSG000001 | CCDC110 | 1.19E-01 | ? | 7.17E-01 | -4.80E-01 | -1.39E+00 |
| ENSG000001 | MAF | 1.20E-01 | 4.43E-01 | 1.11E+00 | 1.46E-01 | 1.11E+00 |
| ENSG00000C | MATK | 1.20E-01 | ? | 2.13E+00 | 1.09E+00 | 2.13E+00 |
| ENSG000002 | --- | 1.20E-01 | 4.43E-01 | 8.51E-01 | -2.33E-01 | -1.18E+00 |
| ENSG000002 | KRBOX1 | 1.20E-01 | ? | 1.57E+00 | 6.52E-01 | 1.57E+00 |
| ENSG000001 | SHQ1 | 1.20E-01 | 4.43E-01 | 1.06E+00 | 8.59E-02 | 1.06E+00 |
| ENSG000001 | REPS1 | 1.20E-01 | 4.43E-01 | 9.48E-01 | -7.74E-02 | -1.06E+00 |
| ENSG000002 | --- | 1.20E-01 | ? | 4.89E-01 | -1.03E+00 | -2.04E+00 |
| ENSG000001 | NEXN | 1.20E-01 | 4.43E-01 | 8.34E-01 | -2.62E-01 | -1.20E+00 |
| ENSG000001 | SCCPDH | 1.20E-01 | 4.43E-01 | 1.08E+00 | 1.10E-01 | 1.08E+00 |
| ENSG000001 | GYS1 | 1.20E-01 | 4.43E-01 | 1.06E+00 | 8.02E-02 | 1.06E+00 |
| ENSG000002 | CSPG4P11 | 1.20E-01 | 4.43E-01 | 1.24E+00 | 3.10E-01 | 1.24E+00 |
| ENSG000001 | KLHL25 | 1.20E-01 | 4.43E-01 | 1.12E+00 | 1.66E-01 | 1.12E+00 |
| ENSG000001 | PITPNM1 | 1.20E-01 | 4.43E-01 | 1.11E+00 | 1.55E-01 | 1.11E+00 |
| ENSG000002 | DLGAP4-AS1 | 1.20E-01 | ? | 1.84E+00 | 8.80E-01 | 1.84E+00 |
| ENSG000001 | ATP6V0E2 | 1.20E-01 | 4.43E-01 | 1.21E+00 | 2.80E-01 | 1.21E+00 |
| ENSG000001 | GRB7 | 1.20E-01 | 4.43E-01 | 1.11E+00 | 1.53E-01 | 1.11E+00 |
| ENSG000001 | PLPBP | 1.20E-01 | 4.43E-01 | 1.05E+00 | 6.45E-02 | 1.05E+00 |
| ENSG000001 | SCYL1 | 1.20E-01 | 4.43E-01 | 1.07E+00 | 9.75E-02 | 1.07E+00 |
| ENSG000001 | LYSMD2 | 1.20E-01 | 4.43E-01 | 1.16E+00 | 2.20E-01 | 1.16E+00 |
| ENSG000001 | EXOSC10 | 1.20E-01 | 4.43E-01 | 9.45E-01 | -8.19E-02 | -1.06E+00 |
| ENSG000001 | HDAC3 | 1.20E-01 | 4.43E-01 | 1.06E+00 | 8.99E-02 | 1.06E+00 |
| ENSG000001 | UBR4 | 1.20E-01 | 4.43E-01 | 9.09E-01 | -1.38E-01 | -1.10E+00 |
| ENSG00000C | ATP2B4 | 1.20E-01 | 4.43E-01 | 1.10E+00 | 1.33E-01 | 1.10E+00 |
| ENSG000002 | C8orf58 | 1.20E-01 | 4.43E-01 | 1.09E+00 | 1.29E-01 | 1.09E+00 |
| ENSG000001 | COX18 | 1.20E-01 | 4.43E-01 | 1.08E+00 | 1.08E-01 | 1.08E+00 |
| ENSG000001 | HCLS1 | 1.20E-01 | 4.43E-01 | 9.10E-01 | -1.36E-01 | -1.10E+00 |
| ENSG000001 | WASF3 | 1.20E-01 | 4.43E-01 | 1.07E+00 | 1.02E-01 | 1.07E+00 |
| ENSG000002 | DDX50P1 | 1.20E-01 | ? | 6.48E-01 | -6.26E-01 | -1.54E+00 |
| ENSG000001 | TRMT112 | 1.20E-01 | 4.43E-01 | 1.05E+00 | 6.52E-02 | 1.05E+00 |
| ENSG00000C | UBA5 | 1.20E-01 | 4.43E-01 | 9.39E-01 | -9.05E-02 | -1.06E+00 |
| ENSG000001 | MMGT1 | 1.21E-01 | 4.43E-01 | 9.40E-01 | -8.88E-02 | -1.06E+00 |
| ENSG000001 | CHD4 | 1.21E-01 | 4.43E-01 | 9.61E-01 | -5.68E-02 | -1.04E+00 |
| ENSG000001 | BMP6 | 1.21E-01 | ? | 1.79E+00 | 8.37E-01 | 1.79E+00 |
| ENSG000002 | MRNIP-DT | 1.21E-01 | ? | 7.13E-01 | -4.88E-01 | -1.40E+00 |

| | | | | | | |
|---|---|---|---|---|---|---|
| ENSG000001 | --- | 1.21E-01 | ? | 1.30E+00 | 3.82E-01 | 1.30E+00 |
| ENSG00000C | PLAUR | 1.21E-01 | 4.44E-01 | 9.12E-01 | -1.33E-01 | -1.10E+00 |
| ENSG000002 | CFI | 1.21E-01 | ? | 6.56E-01 | -6.09E-01 | -1.52E+00 |
| ENSG000001 | TMEM255B | 1.21E-01 | 4.44E-01 | 1.09E+00 | 1.23E-01 | 1.09E+00 |
| ENSG000001 | CEP112 | 1.21E-01 | 4.44E-01 | 9.00E-01 | -1.51E-01 | -1.11E+00 |
| ENSG000001 | GTPBP10 | 1.21E-01 | 4.44E-01 | 9.28E-01 | -1.07E-01 | -1.08E+00 |
| ENSG00000C | BLTP3A | 1.21E-01 | 4.44E-01 | 1.07E+00 | 1.00E-01 | 1.07E+00 |
| ENSG000002 | --- | 1.21E-01 | ? | 1.35E+00 | 4.37E-01 | 1.35E+00 |
| ENSG000002 | UBA6-DT | 1.21E-01 | 4.44E-01 | 9.36E-01 | -9.62E-02 | -1.07E+00 |
| ENSG00000C | MTMR1 | 1.21E-01 | 4.44E-01 | 1.05E+00 | 7.06E-02 | 1.05E+00 |
| ENSG000002 | --- | 1.21E-01 | ? | 2.30E+00 | 1.20E+00 | 2.30E+00 |
| ENSG000002 | --- | 1.21E-01 | 4.45E-01 | 7.90E-01 | -3.40E-01 | -1.27E+00 |
| ENSG000001 | TADA2B | 1.21E-01 | 4.45E-01 | 1.06E+00 | 8.59E-02 | 1.06E+00 |
| ENSG000001 | DAGLA | 1.21E-01 | 4.45E-01 | 8.08E-01 | -3.08E-01 | -1.24E+00 |
| ENSG000002 | ZNF564 | 1.21E-01 | 4.45E-01 | 8.89E-01 | -1.70E-01 | -1.12E+00 |
| ENSG000002 | DDOST | 1.21E-01 | 4.45E-01 | 1.07E+00 | 1.02E-01 | 1.07E+00 |
| ENSG000002 | PI4KAP1 | 1.21E-01 | 4.45E-01 | 8.54E-01 | -2.28E-01 | -1.17E+00 |
| ENSG000001 | EMILIN3 | 1.21E-01 | ? | 5.02E-01 | -9.94E-01 | -1.99E+00 |
| ENSG00000C | XYLT2 | 1.21E-01 | 4.45E-01 | 1.06E+00 | 8.60E-02 | 1.06E+00 |
| ENSG000001 | G6PD | 1.21E-01 | 4.45E-01 | 9.28E-01 | -1.07E-01 | -1.08E+00 |
| ENSG000002 | --- | 1.22E-01 | 4.45E-01 | 7.74E-01 | -3.69E-01 | -1.29E+00 |
| ENSG000002 | EID3 | 1.22E-01 | ? | 5.87E-01 | -7.68E-01 | -1.70E+00 |
| ENSG000002 | --- | 1.22E-01 | ? | 1.82E+00 | 8.65E-01 | 1.82E+00 |
| ENSG000002 | --- | 1.22E-01 | 4.46E-01 | 8.64E-01 | -2.11E-01 | -1.16E+00 |
| ENSG000001 | WDR13 | 1.22E-01 | 4.46E-01 | 1.09E+00 | 1.18E-01 | 1.09E+00 |
| ENSG00000C | RBM6 | 1.22E-01 | 4.46E-01 | 9.23E-01 | -1.15E-01 | -1.08E+00 |
| ENSG000001 | UST | 1.22E-01 | 4.46E-01 | 1.14E+00 | 1.86E-01 | 1.14E+00 |
| ENSG000002 | --- | 1.22E-01 | 4.46E-01 | 7.48E-01 | -4.18E-01 | -1.34E+00 |
| ENSG000001 | TSPAN11 | 1.22E-01 | ? | 1.45E+00 | 5.40E-01 | 1.45E+00 |
| ENSG000001 | PKNOX2 | 1.22E-01 | ? | 2.03E+00 | 1.02E+00 | 2.03E+00 |
| ENSG000001 | SSR4 | 1.22E-01 | 4.46E-01 | 1.11E+00 | 1.44E-01 | 1.11E+00 |
| ENSG000001 | ZNF621 | 1.22E-01 | 4.47E-01 | 9.34E-01 | -9.80E-02 | -1.07E+00 |
| ENSG00000C | AGPAT4 | 1.22E-01 | 4.47E-01 | 9.30E-01 | -1.05E-01 | -1.08E+00 |
| ENSG000002 | ATXN2 | 1.23E-01 | 4.48E-01 | 9.16E-01 | -1.27E-01 | -1.09E+00 |
| ENSG000001 | SLF1 | 1.23E-01 | 4.48E-01 | 8.87E-01 | -1.72E-01 | -1.13E+00 |
| ENSG000002 | LINC00824 | 1.23E-01 | ? | 7.00E-01 | -5.15E-01 | -1.43E+00 |
| ENSG000001 | ARRB1 | 1.23E-01 | 4.48E-01 | 1.28E+00 | 3.60E-01 | 1.28E+00 |
| ENSG000001 | TANGO6 | 1.23E-01 | 4.48E-01 | 9.39E-01 | -9.15E-02 | -1.07E+00 |
| ENSG00000C | NEO1 | 1.23E-01 | 4.48E-01 | 1.06E+00 | 8.77E-02 | 1.06E+00 |
| ENSG000002 | OTUD6B-AS1 | 1.23E-01 | 4.48E-01 | 9.30E-01 | -1.05E-01 | -1.08E+00 |
| ENSG00000C | KDM7A | 1.23E-01 | 4.49E-01 | 9.37E-01 | -9.44E-02 | -1.07E+00 |
| ENSG000001 | SMAD5 | 1.23E-01 | 4.49E-01 | 1.04E+00 | 6.14E-02 | 1.04E+00 |
| ENSG000001 | SEC24D | 1.23E-01 | 4.49E-01 | 9.40E-01 | -8.97E-02 | -1.06E+00 |
| ENSG000002 | PCDHB1-AS1 | 1.23E-01 | 4.49E-01 | 1.31E+00 | 3.85E-01 | 1.31E+00 |
| ENSG000001 | CCDC184 | 1.23E-01 | ? | 2.32E+00 | 1.22E+00 | 2.32E+00 |

| | | | | | | |
|---|---|---|---|---|---|---|
| ENSG000002 | --- | 1.23E-01 | ? | 7.41E-01 | -4.32E-01 | -1.35E+00 |
| ENSG000001 | RPF2 | 1.23E-01 | 4.49E-01 | 9.33E-01 | -1.00E-01 | -1.07E+00 |
| ENSG00000C | RAI14 | 1.23E-01 | 4.49E-01 | 1.06E+00 | 8.80E-02 | 1.06E+00 |
| ENSG000002 | --- | 1.23E-01 | 4.49E-01 | 7.28E-01 | -4.59E-01 | -1.37E+00 |
| ENSG000001 | APOL2 | 1.23E-01 | 4.49E-01 | 9.31E-01 | -1.03E-01 | -1.07E+00 |
| ENSG000001 | TBC1D32 | 1.23E-01 | 4.49E-01 | 1.12E+00 | 1.66E-01 | 1.12E+00 |
| ENSG000002 | --- | 1.23E-01 | 4.49E-01 | 7.70E-01 | -3.77E-01 | -1.30E+00 |
| ENSG000002 | LAYN | 1.23E-01 | 4.49E-01 | 1.33E+00 | 4.12E-01 | 1.33E+00 |
| ENSG000001 | CRTC1 | 1.24E-01 | 4.49E-01 | 1.10E+00 | 1.36E-01 | 1.10E+00 |
| ENSG00000C | LPCAT2 | 1.24E-01 | 4.49E-01 | 9.03E-01 | -1.48E-01 | -1.11E+00 |
| ENSG000001 | ATP5MC1 | 1.24E-01 | 4.49E-01 | 1.09E+00 | 1.22E-01 | 1.09E+00 |
| ENSG000001 | BTBD6 | 1.24E-01 | 4.49E-01 | 1.09E+00 | 1.26E-01 | 1.09E+00 |
| ENSG000002 | --- | 1.24E-01 | ? | 2.29E+00 | 1.20E+00 | 2.29E+00 |
| ENSG000001 | BBOX1 | 1.24E-01 | ? | 1.46E+00 | 5.49E-01 | 1.46E+00 |
| ENSG000001 | PROM2 | 1.24E-01 | 4.49E-01 | 1.07E+00 | 9.48E-02 | 1.07E+00 |
| ENSG000001 | KDELR3 | 1.24E-01 | 4.49E-01 | 8.91E-01 | -1.67E-01 | -1.12E+00 |
| ENSG000002 | RGMB-AS1 | 1.24E-01 | 4.49E-01 | 8.32E-01 | -2.65E-01 | -1.20E+00 |
| ENSG000002 | SRSF9P1 | 1.24E-01 | ? | 2.03E+00 | 1.02E+00 | 2.03E+00 |
| ENSG000001 | ADD3 | 1.24E-01 | 4.50E-01 | 9.49E-01 | -7.49E-02 | -1.05E+00 |
| ENSG000001 | MAN1C1 | 1.24E-01 | ? | 1.37E+00 | 4.56E-01 | 1.37E+00 |
| ENSG00000C | ARVCF | 1.24E-01 | 4.51E-01 | 1.14E+00 | 1.87E-01 | 1.14E+00 |
| ENSG000002 | --- | 1.24E-01 | ? | 5.34E-01 | -9.06E-01 | -1.87E+00 |
| ENSG000002 | --- | 1.24E-01 | 4.51E-01 | 1.19E+00 | 2.56E-01 | 1.19E+00 |
| ENSG000001 | GALNT12 | 1.24E-01 | 4.51E-01 | 1.21E+00 | 2.74E-01 | 1.21E+00 |
| ENSG000002 | --- | 1.24E-01 | ? | 1.77E+00 | 8.27E-01 | 1.77E+00 |
| ENSG00000C | COX15 | 1.24E-01 | 4.51E-01 | 1.06E+00 | 8.27E-02 | 1.06E+00 |
| ENSG000001 | SOX4 | 1.24E-01 | 4.51E-01 | 9.47E-01 | -7.79E-02 | -1.06E+00 |
| ENSG000002 | --- | 1.24E-01 | ? | 6.44E-01 | -6.35E-01 | -1.55E+00 |
| ENSG000002 | --- | 1.24E-01 | ? | 5.61E-01 | -8.33E-01 | -1.78E+00 |
| ENSG000002 | PSMA3-AS1 | 1.24E-01 | 4.51E-01 | 9.04E-01 | -1.46E-01 | -1.11E+00 |
| ENSG00000C | ATP6AP1 | 1.24E-01 | 4.51E-01 | 1.07E+00 | 9.29E-02 | 1.07E+00 |
| ENSG000001 | VPS26B | 1.25E-01 | 4.51E-01 | 1.05E+00 | 7.51E-02 | 1.05E+00 |
| ENSG000001 | TP53TG5 | 1.25E-01 | ? | 6.52E-01 | -6.17E-01 | -1.53E+00 |
| ENSG000002 | --- | 1.25E-01 | ? | 1.90E+00 | 9.22E-01 | 1.90E+00 |
| ENSG000001 | ORC5 | 1.25E-01 | 4.51E-01 | 1.07E+00 | 9.63E-02 | 1.07E+00 |
| ENSG00000C | TMEM98 | 1.25E-01 | 4.51E-01 | 1.08E+00 | 1.05E-01 | 1.08E+00 |
| ENSG000001 | BMT2 | 1.25E-01 | 4.51E-01 | 1.08E+00 | 1.12E-01 | 1.08E+00 |
| ENSG000001 | ZNF83 | 1.25E-01 | 4.51E-01 | 9.10E-01 | -1.36E-01 | -1.10E+00 |
| ENSG000001 | TRAP1 | 1.25E-01 | 4.52E-01 | 1.06E+00 | 8.41E-02 | 1.06E+00 |
| ENSG000002 | MBD5 | 1.25E-01 | 4.52E-01 | 9.41E-01 | -8.70E-02 | -1.06E+00 |
| ENSG00000C | TRAF3IP3 | 1.25E-01 | 4.52E-01 | 8.77E-01 | -1.90E-01 | -1.14E+00 |
| ENSG000002 | LINC00571 | 1.25E-01 | ? | 1.30E+00 | 3.79E-01 | 1.30E+00 |
| ENSG000002 | LINC02035 | 1.25E-01 | 4.52E-01 | 1.15E+00 | 2.03E-01 | 1.15E+00 |
| ENSG000002 | --- | 1.25E-01 | 4.52E-01 | 7.89E-01 | -3.41E-01 | -1.27E+00 |
| ENSG00000C | EFR3B | 1.25E-01 | 4.52E-01 | 8.55E-01 | -2.26E-01 | -1.17E+00 |

| | | | | | | |
|---|---|---|---|---|---|---|
| ENSG000001 | COX5A | 1.25E-01 | 4.52E-01 | 1.06E+00 | 8.23E-02 | 1.06E+00 |
| ENSG000001 | DMAC2L | 1.25E-01 | 4.52E-01 | 1.11E+00 | 1.51E-01 | 1.11E+00 |
| ENSG000001 | SLC5A3 | 1.25E-01 | 4.52E-01 | 1.06E+00 | 9.08E-02 | 1.06E+00 |
| ENSG000001 | PGLYRP4 | 1.25E-01 | 4.52E-01 | 1.23E+00 | 2.94E-01 | 1.23E+00 |
| ENSG000002 | --- | 1.25E-01 | 4.52E-01 | 9.09E-01 | -1.37E-01 | -1.10E+00 |
| ENSG000001 | ZDHHC7 | 1.25E-01 | 4.52E-01 | 1.04E+00 | 5.85E-02 | 1.04E+00 |
| ENSG000000 | PCDHB4 | 1.25E-01 | ? | 1.71E+00 | 7.72E-01 | 1.71E+00 |
| ENSG000002 | RPL36A | 1.25E-01 | 4.52E-01 | 1.05E+00 | 7.08E-02 | 1.05E+00 |
| ENSG000002 | NFS1 | 1.25E-01 | 4.52E-01 | 9.37E-01 | -9.43E-02 | -1.07E+00 |
| ENSG000002 | --- | 1.25E-01 | ? | 6.41E-01 | -6.41E-01 | -1.56E+00 |
| ENSG000001 | PAXBP1 | 1.25E-01 | 4.52E-01 | 9.27E-01 | -1.09E-01 | -1.08E+00 |
| ENSG000000 | ALKBH5 | 1.25E-01 | 4.52E-01 | 9.34E-01 | -9.87E-02 | -1.07E+00 |
| ENSG000002 | --- | 1.25E-01 | ? | 7.47E-01 | -4.20E-01 | -1.34E+00 |
| ENSG000002 | --- | 1.26E-01 | ? | 7.19E-01 | -4.76E-01 | -1.39E+00 |
| ENSG000001 | LRRTM2 | 1.26E-01 | ? | 2.40E+00 | 1.27E+00 | 2.40E+00 |
| ENSG000001 | GIPC1 | 1.26E-01 | 4.53E-01 | 1.10E+00 | 1.42E-01 | 1.10E+00 |
| ENSG000001 | HPSE2 | 1.26E-01 | ? | 5.14E-01 | -9.59E-01 | -1.94E+00 |
| ENSG000001 | STEAP3 | 1.26E-01 | 4.53E-01 | 9.23E-01 | -1.16E-01 | -1.08E+00 |
| ENSG000001 | GLB1 | 1.26E-01 | 4.53E-01 | 1.06E+00 | 8.30E-02 | 1.06E+00 |
| ENSG000001 | CIART | 1.26E-01 | 4.54E-01 | 8.34E-01 | -2.62E-01 | -1.20E+00 |
| ENSG000002 | ZNF737 | 1.26E-01 | 4.54E-01 | 8.78E-01 | -1.88E-01 | -1.14E+00 |
| ENSG000001 | SRM | 1.26E-01 | 4.55E-01 | 1.08E+00 | 1.15E-01 | 1.08E+00 |
| ENSG000002 | --- | 1.26E-01 | ? | 2.35E+00 | 1.23E+00 | 2.35E+00 |
| ENSG000001 | WDR20 | 1.26E-01 | 4.55E-01 | 1.04E+00 | 6.17E-02 | 1.04E+00 |
| ENSG000002 | SNAI3-AS1 | 1.26E-01 | 4.55E-01 | 1.15E+00 | 2.00E-01 | 1.15E+00 |
| ENSG000002 | --- | 1.26E-01 | ? | 4.05E-01 | -1.30E+00 | -2.47E+00 |
| ENSG000001 | TOMM20 | 1.26E-01 | 4.55E-01 | 9.30E-01 | -1.04E-01 | -1.07E+00 |
| ENSG000001 | ZNF20 | 1.27E-01 | 4.55E-01 | 1.35E+00 | 4.32E-01 | 1.35E+00 |
| ENSG000001 | ALG5 | 1.27E-01 | 4.55E-01 | 9.51E-01 | -7.29E-02 | -1.05E+00 |
| ENSG000002 | LINC00261 | 1.27E-01 | 4.55E-01 | 8.05E-01 | -3.13E-01 | -1.24E+00 |
| ENSG000001 | ZNF706 | 1.27E-01 | 4.56E-01 | 1.06E+00 | 7.91E-02 | 1.06E+00 |
| ENSG000002 | HSPD1P1 | 1.27E-01 | ? | 7.13E-01 | -4.88E-01 | -1.40E+00 |
| ENSG000002 | SEPTIN9-DT | 1.27E-01 | ? | 1.29E+00 | 3.62E-01 | 1.29E+00 |
| ENSG000001 | --- | 1.27E-01 | 4.56E-01 | 9.26E-01 | -1.11E-01 | -1.08E+00 |
| ENSG000001 | OPLAH | 1.27E-01 | 4.56E-01 | 1.23E+00 | 3.03E-01 | 1.23E+00 |
| ENSG000001 | TRIM62 | 1.27E-01 | 4.56E-01 | 1.08E+00 | 1.15E-01 | 1.08E+00 |
| ENSG000002 | --- | 1.27E-01 | ? | 2.45E+00 | 1.29E+00 | 2.45E+00 |
| ENSG000001 | CSF3 | 1.27E-01 | ? | 5.27E-01 | -9.24E-01 | -1.90E+00 |
| ENSG000002 | BNIP3P11 | 1.27E-01 | ? | 2.15E+00 | 1.10E+00 | 2.15E+00 |
| ENSG000001 | CXXC1 | 1.27E-01 | 4.56E-01 | 1.05E+00 | 7.72E-02 | 1.05E+00 |
| ENSG000001 | MTARC1 | 1.27E-01 | 4.56E-01 | 9.18E-01 | -1.23E-01 | -1.09E+00 |
| ENSG000001 | ZCCHC2 | 1.27E-01 | 4.56E-01 | 9.38E-01 | -9.26E-02 | -1.07E+00 |
| ENSG000001 | VEZF1 | 1.27E-01 | 4.56E-01 | 9.51E-01 | -7.26E-02 | -1.05E+00 |
| ENSG000001 | SGSM3 | 1.27E-01 | 4.57E-01 | 1.11E+00 | 1.52E-01 | 1.11E+00 |
| ENSG000001 | DPYSL4 | 1.27E-01 | 4.57E-01 | 1.16E+00 | 2.20E-01 | 1.16E+00 |

| | | | | | | |
|---|---|---|---|---|---|---|
| ENSG000001 | GJD3 | 1.27E-01 | ? | 1.56E+00 | 6.44E-01 | 1.56E+00 |
| ENSG000001 | KCNK3 | 1.27E-01 | ? | 1.57E+00 | 6.47E-01 | 1.57E+00 |
| ENSG000001 | SLC30A4 | 1.27E-01 | 4.57E-01 | 1.06E+00 | 7.96E-02 | 1.06E+00 |
| ENSG000001 | TRPV3 | 1.28E-01 | 4.57E-01 | 8.42E-01 | -2.48E-01 | -1.19E+00 |
| ENSG00000C | WWC3 | 1.28E-01 | 4.57E-01 | 1.05E+00 | 7.17E-02 | 1.05E+00 |
| ENSG00000C | PAPOLA | 1.28E-01 | 4.57E-01 | 9.55E-01 | -6.67E-02 | -1.05E+00 |
| ENSG000002 | --- | 1.28E-01 | ? | 1.34E+00 | 4.22E-01 | 1.34E+00 |
| ENSG000001 | NFATC1 | 1.28E-01 | 4.57E-01 | 1.23E+00 | 2.96E-01 | 1.23E+00 |
| ENSG000002 | --- | 1.28E-01 | 4.57E-01 | 9.05E-01 | -1.44E-01 | -1.10E+00 |
| ENSG000001 | CDC25B | 1.28E-01 | 4.58E-01 | 9.38E-01 | -9.30E-02 | -1.07E+00 |
| ENSG000002 | GATC | 1.28E-01 | 4.58E-01 | 1.04E+00 | 6.14E-02 | 1.04E+00 |
| ENSG000002 | --- | 1.28E-01 | 4.58E-01 | 1.12E+00 | 1.63E-01 | 1.12E+00 |
| ENSG000001 | LAPTM5 | 1.28E-01 | 4.58E-01 | 8.86E-01 | -1.75E-01 | -1.13E+00 |
| ENSG000002 | BMS1P1 | 1.28E-01 | 4.58E-01 | 8.53E-01 | -2.29E-01 | -1.17E+00 |
| ENSG000001 | TFCP2 | 1.28E-01 | 4.58E-01 | 9.52E-01 | -7.07E-02 | -1.05E+00 |
| ENSG000001 | PRKAG2 | 1.28E-01 | 4.58E-01 | 9.27E-01 | -1.10E-01 | -1.08E+00 |
| ENSG000002 | LINC01139 | 1.28E-01 | ? | 6.89E-01 | -5.38E-01 | -1.45E+00 |
| ENSG000002 | PARG | 1.28E-01 | 4.58E-01 | 9.52E-01 | -7.10E-02 | -1.05E+00 |
| ENSG000001 | GCAT | 1.28E-01 | 4.58E-01 | 1.15E+00 | 2.01E-01 | 1.15E+00 |
| ENSG000001 | CDKN1A | 1.28E-01 | 4.58E-01 | 1.09E+00 | 1.22E-01 | 1.09E+00 |
| ENSG000001 | TBCC | 1.28E-01 | 4.58E-01 | 1.07E+00 | 1.04E-01 | 1.07E+00 |
| ENSG000002 | CDSN | 1.28E-01 | ? | 2.63E+00 | 1.40E+00 | 2.63E+00 |
| ENSG000001 | TTC32 | 1.28E-01 | 4.58E-01 | 8.20E-01 | -2.87E-01 | -1.22E+00 |
| ENSG000001 | PPP1R7 | 1.28E-01 | 4.58E-01 | 1.04E+00 | 5.24E-02 | 1.04E+00 |
| ENSG000002 | LINC00957 | 1.28E-01 | 4.58E-01 | 1.22E+00 | 2.91E-01 | 1.22E+00 |
| ENSG000001 | FZD5 | 1.28E-01 | 4.58E-01 | 1.08E+00 | 1.16E-01 | 1.08E+00 |
| ENSG00000C | SLC7A2 | 1.29E-01 | 4.58E-01 | 8.14E-01 | -2.97E-01 | -1.23E+00 |
| ENSG000002 | LINC02598 | 1.29E-01 | ? | 1.60E+00 | 6.76E-01 | 1.60E+00 |
| ENSG000002 | FAM47E-STB | 1.29E-01 | ? | 3.43E-01 | -1.54E+00 | -2.92E+00 |
| ENSG000001 | UVSSA | 1.29E-01 | 4.59E-01 | 9.36E-01 | -9.50E-02 | -1.07E+00 |
| ENSG000002 | DUS4L-BCAP | 1.29E-01 | 4.59E-01 | 9.30E-01 | -1.05E-01 | -1.08E+00 |
| ENSG000001 | MTM1 | 1.29E-01 | 4.59E-01 | 9.37E-01 | -9.45E-02 | -1.07E+00 |
| ENSG000001 | CHCHD4 | 1.29E-01 | 4.59E-01 | 9.27E-01 | -1.10E-01 | -1.08E+00 |
| ENSG000001 | ERI3 | 1.29E-01 | 4.59E-01 | 1.08E+00 | 1.17E-01 | 1.08E+00 |
| ENSG000001 | RPN2 | 1.29E-01 | 4.59E-01 | 9.64E-01 | -5.23E-02 | -1.04E+00 |
| ENSG000001 | MLXIP | 1.29E-01 | 4.59E-01 | 1.05E+00 | 7.10E-02 | 1.05E+00 |
| ENSG000002 | ATP2A1-AS1 | 1.29E-01 | ? | 7.06E-01 | -5.03E-01 | -1.42E+00 |
| ENSG000001 | KIF12 | 1.29E-01 | 4.59E-01 | 7.87E-01 | -3.46E-01 | -1.27E+00 |
| ENSG000002 | LINC02762 | 1.29E-01 | 4.60E-01 | 9.06E-01 | -1.42E-01 | -1.10E+00 |
| ENSG000002 | RPL26L1-AS1 | 1.29E-01 | 4.60E-01 | 1.29E+00 | 3.68E-01 | 1.29E+00 |
| ENSG000001 | NARS2 | 1.29E-01 | 4.60E-01 | 1.07E+00 | 9.18E-02 | 1.07E+00 |
| ENSG000002 | --- | 1.29E-01 | ? | 1.57E+00 | 6.55E-01 | 1.57E+00 |
| ENSG000002 | --- | 1.29E-01 | ? | 2.08E+00 | 1.05E+00 | 2.08E+00 |
| ENSG00000C | GADD45B | 1.29E-01 | 4.60E-01 | 8.88E-01 | -1.71E-01 | -1.13E+00 |
| ENSG000001 | KDM4B | 1.29E-01 | 4.60E-01 | 1.10E+00 | 1.35E-01 | 1.10E+00 |

| | | | | | | |
|---|---|---|---|---|---|---|
| ENSG000001 | METAP1 | 1.29E-01 | 4.60E-01 | 1.03E+00 | 4.79E-02 | 1.03E+00 |
| ENSG00000C | G2E3 | 1.29E-01 | 4.60E-01 | 9.44E-01 | -8.31E-02 | -1.06E+00 |
| ENSG00000C | CAPG | 1.30E-01 | 4.60E-01 | 1.08E+00 | 1.08E-01 | 1.08E+00 |
| ENSG000002 | --- | 1.30E-01 | 4.60E-01 | 8.81E-01 | -1.83E-01 | -1.14E+00 |
| ENSG000001 | TRIM73 | 1.30E-01 | 4.60E-01 | 1.24E+00 | 3.12E-01 | 1.24E+00 |
| ENSG000001 | FOXM1 | 1.30E-01 | 4.60E-01 | 8.40E-01 | -2.52E-01 | -1.19E+00 |
| ENSG000002 | --- | 1.30E-01 | 4.60E-01 | 1.19E+00 | 2.47E-01 | 1.19E+00 |
| ENSG000001 | DNAH10 | 1.30E-01 | ? | 1.66E+00 | 7.29E-01 | 1.66E+00 |
| ENSG000002 | RPSAP9 | 1.30E-01 | ? | 1.57E+00 | 6.53E-01 | 1.57E+00 |
| ENSG000001 | MUC1 | 1.30E-01 | 4.61E-01 | 8.58E-01 | -2.21E-01 | -1.17E+00 |
| ENSG000001 | SLC25A34 | 1.30E-01 | 4.61E-01 | 1.28E+00 | 3.58E-01 | 1.28E+00 |
| ENSG000002 | FNDC10 | 1.30E-01 | 4.61E-01 | 1.16E+00 | 2.14E-01 | 1.16E+00 |
| ENSG000001 | DNAAF5 | 1.30E-01 | 4.61E-01 | 1.08E+00 | 1.12E-01 | 1.08E+00 |
| ENSG000002 | --- | 1.30E-01 | ? | 1.70E+00 | 7.66E-01 | 1.70E+00 |
| ENSG000002 | --- | 1.30E-01 | ? | 5.93E-01 | -7.54E-01 | -1.69E+00 |
| ENSG000001 | ZNG1A | 1.30E-01 | 4.61E-01 | 9.59E-01 | -5.99E-02 | -1.04E+00 |
| ENSG000001 | ALDH3A1 | 1.30E-01 | 4.61E-01 | 7.98E-01 | -3.26E-01 | -1.25E+00 |
| ENSG000002 | ATP8B1-AS1 | 1.30E-01 | 4.61E-01 | 1.16E+00 | 2.12E-01 | 1.16E+00 |
| ENSG00000C | MKNK1 | 1.30E-01 | 4.61E-01 | 9.40E-01 | -8.87E-02 | -1.06E+00 |
| ENSG000002 | --- | 1.30E-01 | 4.61E-01 | 8.12E-01 | -3.00E-01 | -1.23E+00 |
| ENSG000002 | --- | 1.30E-01 | 4.61E-01 | 1.32E+00 | 4.03E-01 | 1.32E+00 |
| ENSG000002 | --- | 1.31E-01 | 4.62E-01 | 8.35E-01 | -2.60E-01 | -1.20E+00 |
| ENSG000001 | KREMEN2 | 1.31E-01 | ? | 1.48E+00 | 5.63E-01 | 1.48E+00 |
| ENSG000001 | PDCD2L | 1.31E-01 | 4.62E-01 | 1.15E+00 | 2.03E-01 | 1.15E+00 |
| ENSG000001 | NSL1 | 1.31E-01 | 4.62E-01 | 9.40E-01 | -8.94E-02 | -1.06E+00 |
| ENSG000001 | POLR2D | 1.31E-01 | 4.63E-01 | 9.47E-01 | -7.80E-02 | -1.06E+00 |
| ENSG000001 | WDR17 | 1.31E-01 | 4.63E-01 | 1.21E+00 | 2.74E-01 | 1.21E+00 |
| ENSG000002 | RHEBP2 | 1.31E-01 | ? | 1.77E+00 | 8.23E-01 | 1.77E+00 |
| ENSG000001 | ZNF713 | 1.31E-01 | 4.64E-01 | 8.92E-01 | -1.65E-01 | -1.12E+00 |
| ENSG000002 | PPP1R2B | 1.31E-01 | ? | 6.61E-01 | -5.96E-01 | -1.51E+00 |
| ENSG000002 | SENP3-EIF4A | 1.31E-01 | 4.64E-01 | 1.31E+00 | 3.92E-01 | 1.31E+00 |
| ENSG00000C | CCAR1 | 1.31E-01 | 4.64E-01 | 9.39E-01 | -9.13E-02 | -1.07E+00 |
| ENSG000002 | AP2B1P1 | 1.31E-01 | ? | 1.96E+00 | 9.72E-01 | 1.96E+00 |
| ENSG000002 | --- | 1.31E-01 | ? | 5.17E-01 | -9.53E-01 | -1.94E+00 |
| ENSG000001 | CDADC1 | 1.31E-01 | 4.64E-01 | 9.09E-01 | -1.38E-01 | -1.10E+00 |
| ENSG00000C | MVP | 1.31E-01 | 4.64E-01 | 1.07E+00 | 1.02E-01 | 1.07E+00 |
| ENSG000001 | XPO5 | 1.31E-01 | 4.64E-01 | 1.05E+00 | 6.76E-02 | 1.05E+00 |
| ENSG00000C | ITPKC | 1.31E-01 | 4.64E-01 | 1.09E+00 | 1.22E-01 | 1.09E+00 |
| ENSG000001 | PLS1 | 1.31E-01 | 4.64E-01 | 9.42E-01 | -8.69E-02 | -1.06E+00 |
| ENSG000001 | SMYD5 | 1.32E-01 | 4.64E-01 | 1.08E+00 | 1.13E-01 | 1.08E+00 |
| ENSG00000C | HERPUD1 | 1.32E-01 | 4.64E-01 | 9.56E-01 | -6.56E-02 | -1.05E+00 |
| ENSG00000C | SNX13 | 1.32E-01 | 4.64E-01 | 9.42E-01 | -8.61E-02 | -1.06E+00 |
| ENSG000001 | ZBTB34 | 1.32E-01 | 4.64E-01 | 9.26E-01 | -1.11E-01 | -1.08E+00 |
| ENSG000001 | MTLN | 1.32E-01 | 4.64E-01 | 1.21E+00 | 2.71E-01 | 1.21E+00 |
| ENSG000001 | A1BG | 1.32E-01 | ? | 7.54E-01 | -4.08E-01 | -1.33E+00 |

| | | | | | | |
|---|---|---|---|---|---|---|
| ENSG000001 | GATAD2B | 1.32E-01 | 4.65E-01 | 9.47E-01 | -7.83E-02 | -1.06E+00 |
| ENSG000001 | RSRC2 | 1.32E-01 | 4.65E-01 | 9.26E-01 | -1.11E-01 | -1.08E+00 |
| ENSG00000C | WDR62 | 1.32E-01 | 4.65E-01 | 8.45E-01 | -2.43E-01 | -1.18E+00 |
| ENSG000001 | KDR | 1.32E-01 | ? | 1.34E+00 | 4.25E-01 | 1.34E+00 |
| ENSG00000C | CASP10 | 1.32E-01 | 4.65E-01 | 1.07E+00 | 1.04E-01 | 1.07E+00 |
| ENSG000002 | --- | 1.32E-01 | ? | 5.12E-01 | -9.65E-01 | -1.95E+00 |
| ENSG000001 | TRAPPC13 | 1.32E-01 | 4.65E-01 | 9.37E-01 | -9.35E-02 | -1.07E+00 |
| ENSG000002 | NRBF2P6 | 1.32E-01 | ? | 7.84E-01 | -3.51E-01 | -1.28E+00 |
| ENSG00000C | CNTLN | 1.32E-01 | 4.65E-01 | 8.70E-01 | -2.00E-01 | -1.15E+00 |
| ENSG000001 | ZNF273 | 1.32E-01 | 4.65E-01 | 8.59E-01 | -2.19E-01 | -1.16E+00 |
| ENSG000001 | MICOS10 | 1.32E-01 | 4.65E-01 | 1.08E+00 | 1.15E-01 | 1.08E+00 |
| ENSG000002 | HK2-DT | 1.32E-01 | 4.65E-01 | 1.17E+00 | 2.25E-01 | 1.17E+00 |
| ENSG000002 | --- | 1.32E-01 | ? | 1.56E+00 | 6.42E-01 | 1.56E+00 |
| ENSG000001 | THAP8 | 1.32E-01 | 4.65E-01 | 1.17E+00 | 2.31E-01 | 1.17E+00 |
| ENSG000002 | --- | 1.32E-01 | 4.65E-01 | 8.64E-01 | -2.10E-01 | -1.16E+00 |
| ENSG000001 | MAML1 | 1.32E-01 | 4.65E-01 | 9.56E-01 | -6.47E-02 | -1.05E+00 |
| ENSG000002 | LINC02178 | 1.32E-01 | ? | 1.76E+00 | 8.17E-01 | 1.76E+00 |
| ENSG00000C | NADK | 1.33E-01 | 4.66E-01 | 1.08E+00 | 1.11E-01 | 1.08E+00 |
| ENSG000002 | CNN2P4 | 1.33E-01 | ? | 1.69E+00 | 7.54E-01 | 1.69E+00 |
| ENSG000002 | --- | 1.33E-01 | ? | 1.34E+00 | 4.22E-01 | 1.34E+00 |
| ENSG000001 | CENPM | 1.33E-01 | ? | 6.42E-01 | -6.40E-01 | -1.56E+00 |
| ENSG000002 | MILR1 | 1.33E-01 | ? | 7.17E-01 | -4.80E-01 | -1.39E+00 |
| ENSG000002 | --- | 1.33E-01 | ? | 1.79E+00 | 8.43E-01 | 1.79E+00 |
| ENSG000002 | C8orf34-AS1 | 1.33E-01 | ? | 1.63E+00 | 7.08E-01 | 1.63E+00 |
| ENSG000001 | PPA1 | 1.33E-01 | 4.67E-01 | 1.04E+00 | 5.56E-02 | 1.04E+00 |
| ENSG000001 | HNRNPUL1 | 1.33E-01 | 4.67E-01 | 9.65E-01 | -5.13E-02 | -1.04E+00 |
| ENSG000002 | BBIP1 | 1.33E-01 | 4.67E-01 | 9.20E-01 | -1.20E-01 | -1.09E+00 |
| ENSG000002 | GOLGA6L5P | 1.33E-01 | 4.67E-01 | 8.37E-01 | -2.56E-01 | -1.19E+00 |
| ENSG000001 | HECW2 | 1.33E-01 | 4.67E-01 | 8.34E-01 | -2.62E-01 | -1.20E+00 |
| ENSG000002 | LINC00601 | 1.33E-01 | ? | 6.61E-01 | -5.98E-01 | -1.51E+00 |
| ENSG000001 | ZDHHC21 | 1.33E-01 | 4.67E-01 | 9.12E-01 | -1.32E-01 | -1.10E+00 |
| ENSG000002 | PPAN-P2RY1 | 1.33E-01 | ? | 4.99E-01 | -1.00E+00 | -2.01E+00 |
| ENSG00000C | IGSF9B | 1.33E-01 | 4.67E-01 | 1.23E+00 | 3.00E-01 | 1.23E+00 |
| ENSG00000C | CPOX | 1.33E-01 | 4.67E-01 | 9.42E-01 | -8.57E-02 | -1.06E+00 |
| ENSG000002 | --- | 1.33E-01 | 4.67E-01 | 7.74E-01 | -3.69E-01 | -1.29E+00 |
| ENSG000001 | BOD1 | 1.33E-01 | 4.67E-01 | 1.06E+00 | 7.79E-02 | 1.06E+00 |
| ENSG000001 | MTMR4 | 1.33E-01 | 4.67E-01 | 1.08E+00 | 1.12E-01 | 1.08E+00 |
| ENSG000001 | ARHGAP17 | 1.33E-01 | 4.67E-01 | 9.51E-01 | -7.24E-02 | -1.05E+00 |
| ENSG000001 | ZNF790 | 1.34E-01 | 4.68E-01 | 9.22E-01 | -1.18E-01 | -1.09E+00 |
| ENSG000001 | ZNF746 | 1.34E-01 | 4.68E-01 | 9.45E-01 | -8.16E-02 | -1.06E+00 |
| ENSG000001 | USP32 | 1.34E-01 | 4.68E-01 | 9.55E-01 | -6.57E-02 | -1.05E+00 |
| ENSG000001 | IFIT3 | 1.34E-01 | 4.68E-01 | 8.69E-01 | -2.02E-01 | -1.15E+00 |
| ENSG000002 | --- | 1.34E-01 | ? | 5.73E-01 | -8.04E-01 | -1.75E+00 |
| ENSG000001 | TTLL1 | 1.34E-01 | 4.68E-01 | 1.14E+00 | 1.83E-01 | 1.14E+00 |
| ENSG000002 | LINC00339 | 1.34E-01 | 4.68E-01 | 1.16E+00 | 2.13E-01 | 1.16E+00 |

| | | | | | | |
|---|---|---|---|---|---|---|
| ENSG000002 | KLHDC7B-DT | 1.34E-01 | ? | 1.73E+00 | 7.93E-01 | 1.73E+00 |
| ENSG000001 | KAT14 | 1.34E-01 | 4.68E-01 | 1.09E+00 | 1.20E-01 | 1.09E+00 |
| ENSG000001 | MRPS22 | 1.34E-01 | 4.68E-01 | 9.49E-01 | -7.55E-02 | -1.05E+00 |
| ENSG000000 | NFKB2 | 1.34E-01 | 4.68E-01 | 1.05E+00 | 7.29E-02 | 1.05E+00 |
| ENSG000001 | LRPAP1 | 1.34E-01 | 4.68E-01 | 1.07E+00 | 9.16E-02 | 1.07E+00 |
| ENSG000001 | ZBTB3 | 1.34E-01 | 4.68E-01 | 1.11E+00 | 1.55E-01 | 1.11E+00 |
| ENSG000001 | MED30 | 1.34E-01 | 4.68E-01 | 1.13E+00 | 1.78E-01 | 1.13E+00 |
| ENSG000001 | COX14 | 1.34E-01 | 4.68E-01 | 1.13E+00 | 1.79E-01 | 1.13E+00 |
| ENSG000002 | --- | 1.34E-01 | ? | 5.76E-01 | -7.95E-01 | -1.74E+00 |
| ENSG000001 | PEMT | 1.34E-01 | 4.68E-01 | 1.15E+00 | 2.06E-01 | 1.15E+00 |
| ENSG000001 | DSE | 1.34E-01 | 4.68E-01 | 1.05E+00 | 6.89E-02 | 1.05E+00 |
| ENSG000001 | ZNF516 | 1.34E-01 | 4.68E-01 | 1.07E+00 | 9.27E-02 | 1.07E+00 |
| ENSG000001 | RAB37 | 1.34E-01 | ? | 1.81E+00 | 8.52E-01 | 1.81E+00 |
| ENSG000001 | ADAM12 | 1.34E-01 | 4.68E-01 | 9.09E-01 | -1.37E-01 | -1.10E+00 |
| ENSG000000 | EPB41L1 | 1.34E-01 | 4.68E-01 | 9.28E-01 | -1.08E-01 | -1.08E+00 |
| ENSG000001 | RECK | 1.34E-01 | 4.68E-01 | 1.14E+00 | 1.85E-01 | 1.14E+00 |
| ENSG000001 | CKS2 | 1.34E-01 | 4.68E-01 | 9.37E-01 | -9.46E-02 | -1.07E+00 |
| ENSG000001 | MSH6 | 1.34E-01 | 4.68E-01 | 9.54E-01 | -6.86E-02 | -1.05E+00 |
| ENSG000002 | NCBP2AS2 | 1.34E-01 | 4.68E-01 | 1.11E+00 | 1.48E-01 | 1.11E+00 |
| ENSG000002 | OIP5-AS1 | 1.34E-01 | 4.68E-01 | 1.07E+00 | 9.35E-02 | 1.07E+00 |
| ENSG000000 | DHRS9 | 1.34E-01 | ? | 1.48E+00 | 5.69E-01 | 1.48E+00 |
| ENSG000001 | NUP93 | 1.35E-01 | 4.68E-01 | 1.06E+00 | 7.93E-02 | 1.06E+00 |
| ENSG000002 | --- | 1.35E-01 | ? | 2.28E+00 | 1.19E+00 | 2.28E+00 |
| ENSG000000 | NFE2L1 | 1.35E-01 | 4.68E-01 | 9.60E-01 | -5.82E-02 | -1.04E+00 |
| ENSG000001 | PCBP1 | 1.35E-01 | 4.68E-01 | 1.05E+00 | 6.76E-02 | 1.05E+00 |
| ENSG000002 | --- | 1.35E-01 | ? | 6.80E-01 | -5.57E-01 | -1.47E+00 |
| ENSG000001 | SYNE4 | 1.35E-01 | 4.69E-01 | 1.20E+00 | 2.64E-01 | 1.20E+00 |
| ENSG000000 | TRPC5 | 1.35E-01 | ? | 6.87E-01 | -5.43E-01 | -1.46E+00 |
| ENSG000002 | --- | 1.35E-01 | ? | 7.38E-01 | -4.39E-01 | -1.36E+00 |
| ENSG000001 | TOMM40L | 1.35E-01 | 4.69E-01 | 1.08E+00 | 1.17E-01 | 1.08E+00 |
| ENSG000001 | GPBP1L1 | 1.35E-01 | 4.69E-01 | 9.67E-01 | -4.90E-02 | -1.03E+00 |
| ENSG000002 | IL10RB-DT | 1.35E-01 | ? | 6.94E-01 | -5.27E-01 | -1.44E+00 |
| ENSG000002 | RPL10P13 | 1.35E-01 | ? | 1.46E+00 | 5.45E-01 | 1.46E+00 |
| ENSG000001 | CDK16 | 1.35E-01 | 4.69E-01 | 1.03E+00 | 4.78E-02 | 1.03E+00 |
| ENSG000001 | NOSTRIN | 1.35E-01 | 4.69E-01 | 8.43E-01 | -2.47E-01 | -1.19E+00 |
| ENSG000000 | RFC1 | 1.35E-01 | 4.69E-01 | 9.48E-01 | -7.74E-02 | -1.06E+00 |
| ENSG000001 | CD2AP | 1.35E-01 | 4.69E-01 | 9.27E-01 | -1.10E-01 | -1.08E+00 |
| ENSG000002 | ZNF845 | 1.35E-01 | 4.69E-01 | 9.36E-01 | -9.58E-02 | -1.07E+00 |
| ENSG000002 | VPS16 | 1.35E-01 | 4.69E-01 | 1.07E+00 | 9.23E-02 | 1.07E+00 |
| ENSG000001 | ABRAXAS2 | 1.35E-01 | 4.69E-01 | 9.40E-01 | -8.88E-02 | -1.06E+00 |
| ENSG000001 | PIDD1 | 1.35E-01 | 4.70E-01 | 1.16E+00 | 2.13E-01 | 1.16E+00 |
| ENSG000001 | CIZ1 | 1.35E-01 | 4.70E-01 | 9.48E-01 | -7.76E-02 | -1.06E+00 |
| ENSG000001 | ZNF480 | 1.35E-01 | 4.70E-01 | 9.16E-01 | -1.26E-01 | -1.09E+00 |
| ENSG000001 | MMUT | 1.36E-01 | 4.70E-01 | 1.06E+00 | 8.38E-02 | 1.06E+00 |
| ENSG000002 | --- | 1.36E-01 | 4.70E-01 | 8.50E-01 | -2.34E-01 | -1.18E+00 |

| | | | | | | |
|---|---|---|---|---|---|---|
| ENSG00000C | DNASE1L1 | 1.36E-01 | 4.70E-01 | 1.06E+00 | 8.66E-02 | 1.06E+00 |
| ENSG000001 | CKAP5 | 1.36E-01 | 4.70E-01 | 9.61E-01 | -5.79E-02 | -1.04E+00 |
| ENSG000002 | --- | 1.36E-01 | ? | 1.62E+00 | 6.96E-01 | 1.62E+00 |
| ENSG000002 | LINC02019 | 1.36E-01 | ? | 5.45E-01 | -8.75E-01 | -1.83E+00 |
| ENSG000001 | SHOC1 | 1.36E-01 | ? | 6.93E-01 | -5.28E-01 | -1.44E+00 |
| ENSG000002 | CEBPA-DT | 1.36E-01 | 4.70E-01 | 1.30E+00 | 3.80E-01 | 1.30E+00 |
| ENSG000001 | GOLPH3L | 1.36E-01 | 4.70E-01 | 1.06E+00 | 8.36E-02 | 1.06E+00 |
| ENSG000001 | SHISAL1 | 1.36E-01 | 4.70E-01 | 8.58E-01 | -2.21E-01 | -1.17E+00 |
| ENSG000001 | VPS35L | 1.36E-01 | 4.70E-01 | 1.04E+00 | 5.90E-02 | 1.04E+00 |
| ENSG000001 | TTI2 | 1.36E-01 | 4.70E-01 | 1.08E+00 | 1.17E-01 | 1.08E+00 |
| ENSG000001 | NUDT21 | 1.36E-01 | 4.70E-01 | 1.06E+00 | 8.83E-02 | 1.06E+00 |
| ENSG000001 | RASGRP1 | 1.36E-01 | ? | 7.53E-01 | -4.09E-01 | -1.33E+00 |
| ENSG000002 | CARD8-AS1 | 1.36E-01 | 4.71E-01 | 8.88E-01 | -1.71E-01 | -1.13E+00 |
| ENSG000001 | ZNF180 | 1.36E-01 | 4.71E-01 | 9.26E-01 | -1.11E-01 | -1.08E+00 |
| ENSG00000C | ZC3H11A | 1.36E-01 | 4.71E-01 | 9.38E-01 | -9.17E-02 | -1.07E+00 |
| ENSG00000C | LETMD1 | 1.36E-01 | 4.71E-01 | 1.09E+00 | 1.18E-01 | 1.09E+00 |
| ENSG000001 | KATNB1 | 1.36E-01 | 4.71E-01 | 1.10E+00 | 1.37E-01 | 1.10E+00 |
| ENSG000001 | OSR2 | 1.36E-01 | 4.71E-01 | 1.28E+00 | 3.57E-01 | 1.28E+00 |
| ENSG000001 | APEX2 | 1.36E-01 | 4.71E-01 | 1.08E+00 | 1.09E-01 | 1.08E+00 |
| ENSG000002 | --- | 1.36E-01 | ? | 3.95E-01 | -1.34E+00 | -2.53E+00 |
| ENSG000001 | ZNF730 | 1.36E-01 | ? | 7.41E-01 | -4.33E-01 | -1.35E+00 |
| ENSG000001 | ZMIZ1 | 1.36E-01 | 4.71E-01 | 9.45E-01 | -8.21E-02 | -1.06E+00 |
| ENSG000001 | MCC | 1.37E-01 | 4.71E-01 | 1.05E+00 | 7.10E-02 | 1.05E+00 |
| ENSG000001 | RRP8 | 1.37E-01 | 4.72E-01 | 1.06E+00 | 7.85E-02 | 1.06E+00 |
| ENSG000001 | CDAN1 | 1.37E-01 | 4.72E-01 | 1.09E+00 | 1.20E-01 | 1.09E+00 |
| ENSG000002 | GOLGA6L17I | 1.37E-01 | ? | 6.17E-01 | -6.96E-01 | -1.62E+00 |
| ENSG000001 | FKBP14 | 1.37E-01 | 4.72E-01 | 9.48E-01 | -7.67E-02 | -1.05E+00 |
| ENSG000001 | PRDM5 | 1.37E-01 | 4.72E-01 | 8.88E-01 | -1.72E-01 | -1.13E+00 |
| ENSG000001 | KLHDC8A | 1.37E-01 | ? | 6.92E-01 | -5.32E-01 | -1.45E+00 |
| ENSG000001 | ZNF786 | 1.37E-01 | 4.72E-01 | 1.09E+00 | 1.28E-01 | 1.09E+00 |
| ENSG000001 | GTF3C2 | 1.37E-01 | 4.73E-01 | 1.04E+00 | 6.01E-02 | 1.04E+00 |
| ENSG000002 | --- | 1.37E-01 | ? | 2.05E+00 | 1.04E+00 | 2.05E+00 |
| ENSG000001 | PNISR | 1.37E-01 | 4.73E-01 | 9.11E-01 | -1.35E-01 | -1.10E+00 |
| ENSG00000C | GRN | 1.37E-01 | 4.73E-01 | 1.09E+00 | 1.18E-01 | 1.09E+00 |
| ENSG000002 | ARHGAP11A- | 1.37E-01 | 4.73E-01 | 1.28E+00 | 3.56E-01 | 1.28E+00 |
| ENSG000001 | SENP5 | 1.37E-01 | 4.73E-01 | 1.04E+00 | 4.98E-02 | 1.04E+00 |
| ENSG00000C | CLCA4 | 1.37E-01 | ? | 7.21E-01 | -4.71E-01 | -1.39E+00 |
| ENSG000001 | ATF5 | 1.37E-01 | 4.73E-01 | 9.38E-01 | -9.17E-02 | -1.07E+00 |
| ENSG000002 | --- | 1.37E-01 | ? | 6.26E-01 | -6.75E-01 | -1.60E+00 |
| ENSG000001 | ZNF608 | 1.37E-01 | 4.73E-01 | 1.16E+00 | 2.10E-01 | 1.16E+00 |
| ENSG000001 | FZD10 | 1.37E-01 | 4.73E-01 | 8.71E-01 | -1.99E-01 | -1.15E+00 |
| ENSG000001 | ANKRD12 | 1.37E-01 | 4.73E-01 | 9.26E-01 | -1.11E-01 | -1.08E+00 |
| ENSG000001 | PPP1R14C | 1.37E-01 | 4.73E-01 | 1.06E+00 | 9.07E-02 | 1.06E+00 |
| ENSG000002 | FRG1DP | 1.37E-01 | ? | 6.13E-01 | -7.05E-01 | -1.63E+00 |
| ENSG000001 | DNAJC7 | 1.37E-01 | 4.73E-01 | 9.48E-01 | -7.63E-02 | -1.05E+00 |

| | | | | | | | |
|---|---|---|---|---|---|---|---|
| ENSG000002 | NCKIPSD | 1.37E-01 | | 4.73E-01 | 1.09E+00 | 1.20E-01 | 1.09E+00 |
| ENSG000002 | --- | 1.38E-01 | ? | | 6.73E-01 | -5.72E-01 | -1.49E+00 |
| ENSG000001 | KIF15 | 1.38E-01 | | 4.73E-01 | 7.78E-01 | -3.62E-01 | -1.29E+00 |
| ENSG000001 | PIAS3 | 1.38E-01 | | 4.73E-01 | 1.05E+00 | 7.52E-02 | 1.05E+00 |
| ENSG000001 | GALK1 | 1.38E-01 | | 4.74E-01 | 1.22E+00 | 2.83E-01 | 1.22E+00 |
| ENSG000002 | POU5F1 | 1.38E-01 | ? | | 6.98E-01 | -5.18E-01 | -1.43E+00 |
| ENSG000002 | --- | 1.38E-01 | ? | | 6.96E-01 | -5.22E-01 | -1.44E+00 |
| ENSG000001 | FBF1 | 1.38E-01 | | 4.74E-01 | 1.14E+00 | 1.95E-01 | 1.14E+00 |
| ENSG000001 | VRK3 | 1.38E-01 | | 4.74E-01 | 1.07E+00 | 9.85E-02 | 1.07E+00 |
| ENSG000002 | LINC02986 | 1.38E-01 | ? | | 6.29E-01 | -6.68E-01 | -1.59E+00 |
| ENSG000001 | YJU2B | 1.38E-01 | | 4.74E-01 | 9.23E-01 | -1.16E-01 | -1.08E+00 |
| ENSG000001 | TUSC1 | 1.38E-01 | | 4.74E-01 | 1.15E+00 | 1.96E-01 | 1.15E+00 |
| ENSG000001 | UPF2 | 1.38E-01 | | 4.74E-01 | 9.39E-01 | -9.01E-02 | -1.06E+00 |
| ENSG000001 | DYNC2I1 | 1.38E-01 | | 4.74E-01 | 9.31E-01 | -1.03E-01 | -1.07E+00 |
| ENSG000001 | MORN4 | 1.38E-01 | | 4.74E-01 | 1.08E+00 | 1.13E-01 | 1.08E+00 |
| ENSG000001 | NDUFV1 | 1.38E-01 | | 4.74E-01 | 1.09E+00 | 1.24E-01 | 1.09E+00 |
| ENSG00000C | TBX15 | 1.38E-01 | ? | | 6.19E-01 | -6.91E-01 | -1.61E+00 |
| ENSG000001 | FMNL3 | 1.38E-01 | | 4.74E-01 | 9.17E-01 | -1.24E-01 | -1.09E+00 |
| ENSG000001 | DIAPH2 | 1.38E-01 | | 4.74E-01 | 9.30E-01 | -1.05E-01 | -1.08E+00 |
| ENSG000001 | ATP7B | 1.38E-01 | | 4.74E-01 | 9.42E-01 | -8.70E-02 | -1.06E+00 |
| ENSG000001 | LSM8 | 1.38E-01 | | 4.74E-01 | 9.16E-01 | -1.27E-01 | -1.09E+00 |
| ENSG000001 | COPS9 | 1.38E-01 | | 4.74E-01 | 1.09E+00 | 1.28E-01 | 1.09E+00 |
| ENSG000001 | RHOBTB3 | 1.39E-01 | | 4.74E-01 | 1.06E+00 | 7.87E-02 | 1.06E+00 |
| ENSG000001 | USP22 | 1.39E-01 | | 4.74E-01 | 1.04E+00 | 5.57E-02 | 1.04E+00 |
| ENSG000002 | --- | 1.39E-01 | ? | | 1.62E+00 | 6.95E-01 | 1.62E+00 |
| ENSG00000C | HSPB11 | 1.39E-01 | | 4.75E-01 | 1.06E+00 | 9.08E-02 | 1.06E+00 |
| ENSG000001 | VCP | 1.39E-01 | | 4.75E-01 | 1.05E+00 | 7.34E-02 | 1.05E+00 |
| ENSG000001 | LRRC8B | 1.39E-01 | | 4.75E-01 | 1.07E+00 | 9.78E-02 | 1.07E+00 |
| ENSG000002 | RPS24P8 | 1.39E-01 | ? | | 1.44E+00 | 5.28E-01 | 1.44E+00 |
| ENSG000001 | KLHDC9 | 1.39E-01 | ? | | 1.72E+00 | 7.84E-01 | 1.72E+00 |
| ENSG000002 | RPSAP58 | 1.39E-01 | | 4.75E-01 | 1.12E+00 | 1.62E-01 | 1.12E+00 |
| ENSG000001 | CMAS | 1.39E-01 | | 4.75E-01 | 9.63E-01 | -5.48E-02 | -1.04E+00 |
| ENSG000001 | PPIH | 1.39E-01 | | 4.75E-01 | 1.06E+00 | 8.26E-02 | 1.06E+00 |
| ENSG000001 | DZIP3 | 1.39E-01 | | 4.75E-01 | 1.08E+00 | 1.08E-01 | 1.08E+00 |
| ENSG000002 | TMSB10P1 | 1.39E-01 | ? | | 7.12E-01 | -4.90E-01 | -1.40E+00 |
| ENSG000001 | FANCF | 1.39E-01 | | 4.75E-01 | 1.09E+00 | 1.19E-01 | 1.09E+00 |
| ENSG000001 | MMP19 | 1.39E-01 | | 4.75E-01 | 1.18E+00 | 2.34E-01 | 1.18E+00 |
| ENSG00000C | SLC6A15 | 1.39E-01 | | 4.75E-01 | 1.07E+00 | 9.83E-02 | 1.07E+00 |
| ENSG000002 | BRD3OS | 1.39E-01 | | 4.75E-01 | 1.07E+00 | 9.57E-02 | 1.07E+00 |
| ENSG000001 | LRRC14 | 1.39E-01 | | 4.75E-01 | 1.10E+00 | 1.33E-01 | 1.10E+00 |
| ENSG000001 | RPL24 | 1.39E-01 | | 4.75E-01 | 1.04E+00 | 5.99E-02 | 1.04E+00 |
| ENSG000001 | FPGS | 1.39E-01 | | 4.75E-01 | 1.08E+00 | 1.10E-01 | 1.08E+00 |
| ENSG000001 | CCDC18 | 1.39E-01 | | 4.75E-01 | 8.76E-01 | -1.90E-01 | -1.14E+00 |
| ENSG000001 | RIGI | 1.39E-01 | | 4.75E-01 | 8.82E-01 | -1.81E-01 | -1.13E+00 |
| ENSG00000C | MAP2K3 | 1.39E-01 | | 4.75E-01 | 1.10E+00 | 1.31E-01 | 1.10E+00 |

| | | | | | | |
|---|---|---|---|---|---|---|
| ENSG000001 | CCNE1 | 1.39E-01 | 4.75E-01 | 1.09E+00 | 1.20E-01 | 1.09E+00 |
| ENSG000001 | ACBD4 | 1.39E-01 | 4.75E-01 | 1.14E+00 | 1.95E-01 | 1.14E+00 |
| ENSG000001 | RAB2A | 1.39E-01 | 4.75E-01 | 9.54E-01 | -6.84E-02 | -1.05E+00 |
| ENSG000001 | ZNF568 | 1.39E-01 | 4.75E-01 | 9.25E-01 | -1.13E-01 | -1.08E+00 |
| ENSG000001 | LRP2BP | 1.39E-01 | 4.75E-01 | 8.18E-01 | -2.90E-01 | -1.22E+00 |
| ENSG00000C | TRO | 1.39E-01 | ? | 1.32E+00 | 4.01E-01 | 1.32E+00 |
| ENSG000001 | FBXL5 | 1.39E-01 | 4.75E-01 | 1.04E+00 | 5.47E-02 | 1.04E+00 |
| ENSG000002 | WBP1 | 1.39E-01 | 4.75E-01 | 1.09E+00 | 1.24E-01 | 1.09E+00 |
| ENSG000001 | SPATA33 | 1.40E-01 | 4.75E-01 | 1.12E+00 | 1.62E-01 | 1.12E+00 |
| ENSG000001 | DDIT3 | 1.40E-01 | 4.75E-01 | 8.94E-01 | -1.61E-01 | -1.12E+00 |
| ENSG000002 | --- | 1.40E-01 | ? | 4.88E-01 | -1.04E+00 | -2.05E+00 |
| ENSG000001 | EFCAB13 | 1.40E-01 | 4.75E-01 | 7.72E-01 | -3.74E-01 | -1.30E+00 |
| ENSG000002 | SEC14L6 | 1.40E-01 | ? | 6.65E-01 | -5.89E-01 | -1.50E+00 |
| ENSG000002 | CCT8P1 | 1.40E-01 | ? | 1.31E+00 | 3.88E-01 | 1.31E+00 |
| ENSG000002 | --- | 1.40E-01 | 4.75E-01 | 1.16E+00 | 2.16E-01 | 1.16E+00 |
| ENSG000002 | --- | 1.40E-01 | 4.75E-01 | 8.64E-01 | -2.11E-01 | -1.16E+00 |
| ENSG000001 | DTD1 | 1.40E-01 | 4.75E-01 | 1.05E+00 | 6.76E-02 | 1.05E+00 |
| ENSG00000C | CIAPIN1 | 1.40E-01 | 4.75E-01 | 1.07E+00 | 1.02E-01 | 1.07E+00 |
| ENSG000002 | --- | 1.40E-01 | ? | 5.02E-01 | -9.94E-01 | -1.99E+00 |
| ENSG000001 | EPS8L2 | 1.40E-01 | 4.75E-01 | 1.08E+00 | 1.07E-01 | 1.08E+00 |
| ENSG000001 | ABCC11 | 1.40E-01 | 4.75E-01 | 1.20E+00 | 2.65E-01 | 1.20E+00 |
| ENSG000002 | NAMPTP1 | 1.40E-01 | 4.75E-01 | 8.50E-01 | -2.34E-01 | -1.18E+00 |
| ENSG00000C | PCM1 | 1.40E-01 | 4.75E-01 | 9.39E-01 | -9.09E-02 | -1.07E+00 |
| ENSG000002 | MIR23AHG | 1.40E-01 | 4.75E-01 | 1.18E+00 | 2.34E-01 | 1.18E+00 |
| ENSG000002 | --- | 1.40E-01 | ? | 7.71E-01 | -3.76E-01 | -1.30E+00 |
| ENSG000002 | FAM151B-DT | 1.40E-01 | 4.75E-01 | 1.17E+00 | 2.24E-01 | 1.17E+00 |
| ENSG000001 | PTPN11 | 1.40E-01 | 4.75E-01 | 9.57E-01 | -6.35E-02 | -1.05E+00 |
| ENSG000001 | HOOK1 | 1.40E-01 | 4.75E-01 | 1.10E+00 | 1.33E-01 | 1.10E+00 |
| ENSG00000C | MFSD11 | 1.40E-01 | 4.75E-01 | 1.07E+00 | 1.04E-01 | 1.07E+00 |
| ENSG000001 | VPS72 | 1.40E-01 | 4.75E-01 | 1.05E+00 | 6.71E-02 | 1.05E+00 |
| ENSG00000C | FBXO42 | 1.40E-01 | 4.75E-01 | 1.05E+00 | 7.18E-02 | 1.05E+00 |
| ENSG000001 | AFG3L2 | 1.40E-01 | 4.76E-01 | 1.03E+00 | 4.44E-02 | 1.03E+00 |
| ENSG000001 | TINAGL1 | 1.41E-01 | 4.76E-01 | 1.09E+00 | 1.20E-01 | 1.09E+00 |
| ENSG000001 | BRCA2 | 1.41E-01 | 4.76E-01 | 8.39E-01 | -2.53E-01 | -1.19E+00 |
| ENSG000002 | LINC00472 | 1.41E-01 | 4.76E-01 | 8.81E-01 | -1.83E-01 | -1.14E+00 |
| ENSG000002 | SNHG18 | 1.41E-01 | 4.76E-01 | 1.14E+00 | 1.93E-01 | 1.14E+00 |
| ENSG000001 | CALML6 | 1.41E-01 | ? | 1.51E+00 | 5.94E-01 | 1.51E+00 |
| ENSG00000C | ITM2A | 1.41E-01 | ? | 1.40E+00 | 4.91E-01 | 1.40E+00 |
| ENSG00000C | ARID4B | 1.41E-01 | 4.76E-01 | 9.48E-01 | -7.68E-02 | -1.05E+00 |
| ENSG000001 | BRWD1 | 1.41E-01 | 4.77E-01 | 9.37E-01 | -9.42E-02 | -1.07E+00 |
| ENSG000002 | BMS1P10 | 1.41E-01 | ? | 6.99E-01 | -5.16E-01 | -1.43E+00 |
| ENSG000002 | ALDH7A1P1 | 1.41E-01 | ? | 1.74E+00 | 7.98E-01 | 1.74E+00 |
| ENSG000002 | PA2G4P4 | 1.41E-01 | ? | 6.84E-01 | -5.48E-01 | -1.46E+00 |
| ENSG000001 | SLC41A3 | 1.41E-01 | 4.77E-01 | 1.07E+00 | 1.01E-01 | 1.07E+00 |
| ENSG000002 | UBE2FP1 | 1.41E-01 | ? | 1.60E+00 | 6.77E-01 | 1.60E+00 |

| | | | | | | | |
|---|---|---|---|---|---|---|---|
| ENSG000001 | POGLUT3 | 1.41E-01 | | 4.77E-01 | 1.10E+00 | 1.41E-01 | 1.10E+00 |
| ENSG000001 | SRGAP3 | 1.41E-01 | | 4.77E-01 | 1.08E+00 | 1.17E-01 | 1.08E+00 |
| ENSG000001 | NFATC2IP | 1.41E-01 | | 4.77E-01 | 9.40E-01 | -8.89E-02 | -1.06E+00 |
| ENSG000001 | ARL4A | 1.41E-01 | | 4.77E-01 | 9.33E-01 | -1.00E-01 | -1.07E+00 |
| ENSG000002 | LINC00852 | 1.41E-01 | ? | | 7.79E-01 | -3.60E-01 | -1.28E+00 |
| ENSG000001 | LMNB2 | 1.41E-01 | | 4.77E-01 | 1.08E+00 | 1.08E-01 | 1.08E+00 |
| ENSG00000C | FKBP4 | 1.41E-01 | | 4.77E-01 | 9.54E-01 | -6.82E-02 | -1.05E+00 |
| ENSG000001 | RMND5B | 1.41E-01 | | 4.77E-01 | 1.09E+00 | 1.21E-01 | 1.09E+00 |
| ENSG000001 | FHL3 | 1.41E-01 | | 4.77E-01 | 1.08E+00 | 1.15E-01 | 1.08E+00 |
| ENSG000001 | LRIF1 | 1.41E-01 | | 4.77E-01 | 9.34E-01 | -9.82E-02 | -1.07E+00 |
| ENSG000001 | HMGA2-AS1 | 1.41E-01 | | 4.77E-01 | 7.94E-01 | -3.34E-01 | -1.26E+00 |
| ENSG000001 | CPA6 | 1.41E-01 | ? | | 6.61E-01 | -5.98E-01 | -1.51E+00 |
| ENSG00000C | FAM135A | 1.41E-01 | | 4.77E-01 | 1.06E+00 | 8.85E-02 | 1.06E+00 |
| ENSG000002 | --- | 1.41E-01 | ? | | 2.20E+00 | 1.14E+00 | 2.20E+00 |
| ENSG000001 | MCL1 | 1.41E-01 | | 4.77E-01 | 1.04E+00 | 5.67E-02 | 1.04E+00 |
| ENSG000002 | --- | 1.42E-01 | ? | | 1.93E+00 | 9.48E-01 | 1.93E+00 |
| ENSG000001 | URI1 | 1.42E-01 | | 4.77E-01 | 9.47E-01 | -7.82E-02 | -1.06E+00 |
| ENSG000001 | ARIH1 | 1.42E-01 | | 4.77E-01 | 9.50E-01 | -7.41E-02 | -1.05E+00 |
| ENSG00000C | HEBP2 | 1.42E-01 | | 4.77E-01 | 1.04E+00 | 6.12E-02 | 1.04E+00 |
| ENSG000001 | HDAC1 | 1.42E-01 | | 4.77E-01 | 9.57E-01 | -6.31E-02 | -1.04E+00 |
| ENSG000002 | ETFRF1 | 1.42E-01 | | 4.77E-01 | 9.09E-01 | -1.37E-01 | -1.10E+00 |
| ENSG000001 | SOX12 | 1.42E-01 | | 4.77E-01 | 1.10E+00 | 1.39E-01 | 1.10E+00 |
| ENSG000001 | GYG1 | 1.42E-01 | | 4.77E-01 | 1.06E+00 | 8.04E-02 | 1.06E+00 |
| ENSG000002 | --- | 1.42E-01 | ? | | 1.54E+00 | 6.20E-01 | 1.54E+00 |
| ENSG000001 | PIP4K2C | 1.42E-01 | | 4.77E-01 | 9.55E-01 | -6.65E-02 | -1.05E+00 |
| ENSG000002 | --- | 1.42E-01 | ? | | 6.07E-01 | -7.21E-01 | -1.65E+00 |
| ENSG000001 | PMVK | 1.42E-01 | | 4.78E-01 | 1.12E+00 | 1.65E-01 | 1.12E+00 |
| ENSG000001 | PELI3 | 1.42E-01 | | 4.78E-01 | 1.17E+00 | 2.31E-01 | 1.17E+00 |
| ENSG000001 | ZDHHC17 | 1.42E-01 | | 4.78E-01 | 9.47E-01 | -7.89E-02 | -1.06E+00 |
| ENSG000001 | C11orf68 | 1.42E-01 | | 4.78E-01 | 1.10E+00 | 1.39E-01 | 1.10E+00 |
| ENSG000002 | RPL5P12 | 1.42E-01 | ? | | 6.27E-01 | -6.74E-01 | -1.60E+00 |
| ENSG000001 | IFT57 | 1.42E-01 | | 4.78E-01 | 1.06E+00 | 8.56E-02 | 1.06E+00 |
| ENSG00000C | PDZD4 | 1.42E-01 | ? | | 1.67E+00 | 7.36E-01 | 1.67E+00 |
| ENSG00000C | RAPGEF4 | 1.42E-01 | | 4.78E-01 | 8.53E-01 | -2.29E-01 | -1.17E+00 |
| ENSG000002 | HNRNPA1P7 | 1.42E-01 | | 4.78E-01 | 1.18E+00 | 2.44E-01 | 1.18E+00 |
| ENSG000002 | --- | 1.42E-01 | ? | | 3.92E-01 | -1.35E+00 | -2.55E+00 |
| ENSG000001 | TNFAIP8 | 1.42E-01 | | 4.79E-01 | 9.33E-01 | -9.94E-02 | -1.07E+00 |
| ENSG000001 | AKAP6 | 1.42E-01 | | 4.79E-01 | 1.10E+00 | 1.34E-01 | 1.10E+00 |
| ENSG000001 | CENPO | 1.43E-01 | | 4.79E-01 | 8.89E-01 | -1.70E-01 | -1.13E+00 |
| ENSG000002 | --- | 1.43E-01 | ? | | 4.62E-01 | -1.11E+00 | -2.16E+00 |
| ENSG000002 | --- | 1.43E-01 | | 4.79E-01 | 1.08E+00 | 1.13E-01 | 1.08E+00 |
| ENSG000001 | FBXW7 | 1.43E-01 | | 4.79E-01 | 1.06E+00 | 8.85E-02 | 1.06E+00 |
| ENSG000002 | RPL26P6 | 1.43E-01 | ? | | 4.80E-01 | -1.06E+00 | -2.08E+00 |
| ENSG000001 | ZBTB2 | 1.43E-01 | | 4.79E-01 | 9.40E-01 | -8.90E-02 | -1.06E+00 |
| ENSG000001 | SLC35B2 | 1.43E-01 | | 4.79E-01 | 1.07E+00 | 1.02E-01 | 1.07E+00 |

| | | | | | | |
|---|---|---|---|---|---|---|
| ENSG000001 | AJUBA | 1.43E-01 | 4.79E-01 | 1.06E+00 | 8.09E-02 | 1.06E+00 |
| ENSG000002 | PINX1 | 1.43E-01 | 4.79E-01 | 1.05E+00 | 7.09E-02 | 1.05E+00 |
| ENSG000001 | POGLUT2 | 1.43E-01 | 4.79E-01 | 1.11E+00 | 1.53E-01 | 1.11E+00 |
| ENSG000001 | DCAF5 | 1.43E-01 | 4.79E-01 | 1.04E+00 | 4.98E-02 | 1.04E+00 |
| ENSG000001 | NIPSNAP2 | 1.43E-01 | 4.79E-01 | 1.06E+00 | 8.77E-02 | 1.06E+00 |
| ENSG000001 | ACP7 | 1.43E-01 | 4.79E-01 | 1.16E+00 | 2.16E-01 | 1.16E+00 |
| ENSG000002 | CASTOR1 | 1.43E-01 | 4.79E-01 | 1.13E+00 | 1.74E-01 | 1.13E+00 |
| ENSG00000C | NOTCH3 | 1.43E-01 | 4.79E-01 | 1.11E+00 | 1.54E-01 | 1.11E+00 |
| ENSG000001 | CFAP70 | 1.43E-01 | 4.79E-01 | 8.63E-01 | -2.13E-01 | -1.16E+00 |
| ENSG000001 | PDHA1 | 1.43E-01 | 4.79E-01 | 9.57E-01 | -6.39E-02 | -1.05E+00 |
| ENSG000001 | UCN | 1.43E-01 | ? | 2.20E+00 | 1.14E+00 | 2.20E+00 |
| ENSG000001 | EFCAB10 | 1.43E-01 | ? | 1.31E+00 | 3.88E-01 | 1.31E+00 |
| ENSG000002 | LINC00539 | 1.43E-01 | ? | 1.70E+00 | 7.67E-01 | 1.70E+00 |
| ENSG000001 | MST1 | 1.43E-01 | 4.80E-01 | 1.17E+00 | 2.24E-01 | 1.17E+00 |
| ENSG000001 | TPRA1 | 1.43E-01 | 4.80E-01 | 1.12E+00 | 1.63E-01 | 1.12E+00 |
| ENSG000001 | TNNI2 | 1.43E-01 | ? | 5.70E-01 | -8.12E-01 | -1.76E+00 |
| ENSG000001 | CDC25A | 1.44E-01 | 4.80E-01 | 8.81E-01 | -1.83E-01 | -1.14E+00 |
| ENSG000001 | TWF1 | 1.44E-01 | 4.80E-01 | 9.48E-01 | -7.69E-02 | -1.05E+00 |
| ENSG00000C | RHOBTB2 | 1.44E-01 | 4.80E-01 | 1.21E+00 | 2.69E-01 | 1.21E+00 |
| ENSG000001 | SHH | 1.44E-01 | ? | 5.74E-01 | -8.00E-01 | -1.74E+00 |
| ENSG000002 | ZBED5-AS1 | 1.44E-01 | 4.81E-01 | 1.09E+00 | 1.25E-01 | 1.09E+00 |
| ENSG000002 | GSTM2 | 1.44E-01 | 4.81E-01 | 1.13E+00 | 1.81E-01 | 1.13E+00 |
| ENSG000002 | AGAP2-AS1 | 1.44E-01 | 4.81E-01 | 1.12E+00 | 1.68E-01 | 1.12E+00 |
| ENSG000001 | AP1AR | 1.44E-01 | 4.81E-01 | 9.39E-01 | -9.15E-02 | -1.07E+00 |
| ENSG000001 | POSTN | 1.44E-01 | 4.81E-01 | 1.12E+00 | 1.60E-01 | 1.12E+00 |
| ENSG000001 | FOS | 1.44E-01 | 4.81E-01 | 6.36E-01 | -6.53E-01 | -1.57E+00 |
| ENSG000001 | USP43 | 1.44E-01 | 4.81E-01 | 1.08E+00 | 1.08E-01 | 1.08E+00 |
| ENSG00000C | ROS1 | 1.44E-01 | 4.82E-01 | 8.76E-01 | -1.91E-01 | -1.14E+00 |
| ENSG000001 | ERC2 | 1.44E-01 | ? | 7.73E-01 | -3.71E-01 | -1.29E+00 |
| ENSG000001 | COMMD4 | 1.44E-01 | 4.82E-01 | 1.11E+00 | 1.49E-01 | 1.11E+00 |
| ENSG00000C | LAMP3 | 1.44E-01 | 4.82E-01 | 1.20E+00 | 2.63E-01 | 1.20E+00 |
| ENSG000001 | ZNF672 | 1.44E-01 | 4.82E-01 | 1.12E+00 | 1.57E-01 | 1.12E+00 |
| ENSG000002 | --- | 1.45E-01 | ? | 6.42E-01 | -6.40E-01 | -1.56E+00 |
| ENSG000001 | KIRREL1 | 1.45E-01 | 4.82E-01 | 1.05E+00 | 6.88E-02 | 1.05E+00 |
| ENSG000001 | ABRAXAS1 | 1.45E-01 | 4.82E-01 | 9.07E-01 | -1.40E-01 | -1.10E+00 |
| ENSG000001 | CACTIN | 1.45E-01 | 4.82E-01 | 1.08E+00 | 1.05E-01 | 1.08E+00 |
| ENSG000001 | UROS | 1.45E-01 | 4.82E-01 | 1.05E+00 | 7.46E-02 | 1.05E+00 |
| ENSG00000C | ADAM11 | 1.45E-01 | ? | 4.42E-01 | -1.18E+00 | -2.26E+00 |
| ENSG000001 | NOP56 | 1.45E-01 | 4.82E-01 | 9.54E-01 | -6.72E-02 | -1.05E+00 |
| ENSG000001 | SSH2 | 1.45E-01 | 4.82E-01 | 1.05E+00 | 7.22E-02 | 1.05E+00 |
| ENSG000002 | --- | 1.45E-01 | 4.82E-01 | 8.74E-01 | -1.94E-01 | -1.14E+00 |
| ENSG000002 | --- | 1.45E-01 | ? | 7.30E-01 | -4.54E-01 | -1.37E+00 |
| ENSG000001 | DDX17 | 1.45E-01 | 4.82E-01 | 9.52E-01 | -7.12E-02 | -1.05E+00 |
| ENSG000001 | PIN4 | 1.45E-01 | 4.82E-01 | 9.39E-01 | -9.04E-02 | -1.06E+00 |
| ENSG00000C | HIVEP1 | 1.45E-01 | 4.82E-01 | 9.33E-01 | -1.00E-01 | -1.07E+00 |

| | | | | | | |
|---|---|---|---|---|---|---|
| ENSG000001 | SEZ6L2 | 1.45E-01 | 4.82E-01 | 1.11E+00 | 1.53E-01 | 1.11E+00 |
| ENSG000001 | UBXN7 | 1.45E-01 | 4.82E-01 | 9.46E-01 | -7.99E-02 | -1.06E+00 |
| ENSG000002 | MRPS31P4 | 1.45E-01 | ? | 6.71E-01 | -5.75E-01 | -1.49E+00 |
| ENSG000002 | MED28-DT | 1.45E-01 | ? | 6.55E-01 | -6.12E-01 | -1.53E+00 |
| ENSG000002 | PSMD6-AS2 | 1.45E-01 | 4.82E-01 | 8.26E-01 | -2.76E-01 | -1.21E+00 |
| ENSG000001 | ST3GAL3 | 1.45E-01 | 4.82E-01 | 9.16E-01 | -1.26E-01 | -1.09E+00 |
| ENSG000001 | EHMT1 | 1.45E-01 | 4.82E-01 | 9.51E-01 | -7.26E-02 | -1.05E+00 |
| ENSG000001 | ZKSCAN5 | 1.45E-01 | 4.82E-01 | 1.05E+00 | 7.01E-02 | 1.05E+00 |
| ENSG000002 | ZNF280B | 1.45E-01 | 4.83E-01 | 1.13E+00 | 1.82E-01 | 1.13E+00 |
| ENSG000002 | FAM86C2P | 1.45E-01 | 4.83E-01 | 1.15E+00 | 1.98E-01 | 1.15E+00 |
| ENSG000002 | --- | 1.45E-01 | 4.83E-01 | 1.23E+00 | 3.03E-01 | 1.23E+00 |
| ENSG00000C | SAR1A | 1.45E-01 | 4.83E-01 | 9.66E-01 | -4.98E-02 | -1.04E+00 |
| ENSG000001 | DPCD | 1.45E-01 | 4.83E-01 | 1.07E+00 | 1.03E-01 | 1.07E+00 |
| ENSG000002 | --- | 1.45E-01 | ? | 6.45E-01 | -6.33E-01 | -1.55E+00 |
| ENSG000002 | VGLL3 | 1.46E-01 | 4.83E-01 | 1.12E+00 | 1.60E-01 | 1.12E+00 |
| ENSG000001 | B4GALNT1 | 1.46E-01 | 4.83E-01 | 8.25E-01 | -2.77E-01 | -1.21E+00 |
| ENSG000002 | MLLT6 | 1.46E-01 | 4.83E-01 | 1.08E+00 | 1.07E-01 | 1.08E+00 |
| ENSG00000C | EBF4 | 1.46E-01 | ? | 1.67E+00 | 7.37E-01 | 1.67E+00 |
| ENSG000001 | BBS9 | 1.46E-01 | 4.83E-01 | 9.33E-01 | -1.00E-01 | -1.07E+00 |
| ENSG000002 | --- | 1.46E-01 | 4.83E-01 | 8.62E-01 | -2.14E-01 | -1.16E+00 |
| ENSG000001 | SLC19A1 | 1.46E-01 | 4.83E-01 | 1.12E+00 | 1.62E-01 | 1.12E+00 |
| ENSG000002 | --- | 1.46E-01 | ? | 5.46E-01 | -8.74E-01 | -1.83E+00 |
| ENSG000001 | VWF | 1.46E-01 | 4.83E-01 | 1.29E+00 | 3.71E-01 | 1.29E+00 |
| ENSG000002 | RTL10 | 1.46E-01 | 4.83E-01 | 1.06E+00 | 7.85E-02 | 1.06E+00 |
| ENSG000001 | PHYHIP | 1.46E-01 | ? | 1.30E+00 | 3.76E-01 | 1.30E+00 |
| ENSG000001 | PLPPR2 | 1.46E-01 | 4.83E-01 | 1.09E+00 | 1.28E-01 | 1.09E+00 |
| ENSG000001 | DCAF4L1 | 1.46E-01 | 4.83E-01 | 7.72E-01 | -3.74E-01 | -1.30E+00 |
| ENSG000001 | ZNF358 | 1.46E-01 | 4.83E-01 | 1.10E+00 | 1.43E-01 | 1.10E+00 |
| ENSG000001 | SLC27A1 | 1.46E-01 | 4.84E-01 | 1.11E+00 | 1.53E-01 | 1.11E+00 |
| ENSG000002 | HLA-DPB1 | 1.46E-01 | ? | 2.17E+00 | 1.12E+00 | 2.17E+00 |
| ENSG000001 | PATL1 | 1.46E-01 | 4.84E-01 | 9.49E-01 | -7.56E-02 | -1.05E+00 |
| ENSG000001 | PSMB4 | 1.46E-01 | 4.84E-01 | 1.06E+00 | 8.08E-02 | 1.06E+00 |
| ENSG000001 | FAM222A | 1.46E-01 | ? | 7.32E-01 | -4.49E-01 | -1.37E+00 |
| ENSG000002 | FAM86DP | 1.46E-01 | 4.84E-01 | 1.09E+00 | 1.22E-01 | 1.09E+00 |
| ENSG000001 | RABL6 | 1.46E-01 | 4.84E-01 | 1.07E+00 | 9.33E-02 | 1.07E+00 |
| ENSG000001 | PART1 | 1.46E-01 | ? | 6.68E-01 | -5.82E-01 | -1.50E+00 |
| ENSG000001 | ANGEL2 | 1.46E-01 | 4.84E-01 | 9.35E-01 | -9.66E-02 | -1.07E+00 |
| ENSG000002 | PCED1B-AS1 | 1.46E-01 | ? | 6.19E-01 | -6.91E-01 | -1.61E+00 |
| ENSG000001 | CDK12 | 1.46E-01 | 4.84E-01 | 9.55E-01 | -6.59E-02 | -1.05E+00 |
| ENSG000002 | POM121C | 1.46E-01 | 4.84E-01 | 9.58E-01 | -6.21E-02 | -1.04E+00 |
| ENSG000001 | HSPA9 | 1.46E-01 | 4.84E-01 | 9.52E-01 | -7.16E-02 | -1.05E+00 |
| ENSG00000C | BICRA | 1.47E-01 | 4.84E-01 | 9.08E-01 | -1.39E-01 | -1.10E+00 |
| ENSG000001 | ZSCAN26 | 1.47E-01 | 4.84E-01 | 9.26E-01 | -1.11E-01 | -1.08E+00 |
| ENSG000001 | ARL6 | 1.47E-01 | 4.84E-01 | 9.06E-01 | -1.42E-01 | -1.10E+00 |
| ENSG000001 | KDM3B | 1.47E-01 | 4.84E-01 | 1.04E+00 | 5.64E-02 | 1.04E+00 |

| | | | | | | |
|---|---|---|---|---|---|---|
| ENSG000002 | IRF9 | 1.47E-01 | 4.84E-01 | 9.25E-01 | -1.12E-01 | -1.08E+00 |
| ENSG000001 | INTS8 | 1.47E-01 | 4.84E-01 | 9.52E-01 | -7.05E-02 | -1.05E+00 |
| ENSG000001 | WASL | 1.47E-01 | 4.84E-01 | 9.49E-01 | -7.54E-02 | -1.05E+00 |
| ENSG000001 | SGTB | 1.47E-01 | 4.84E-01 | 9.27E-01 | -1.10E-01 | -1.08E+00 |
| ENSG000001 | COPS4 | 1.47E-01 | 4.84E-01 | 9.53E-01 | -7.01E-02 | -1.05E+00 |
| ENSG000001 | KREMEN1 | 1.47E-01 | 4.84E-01 | 1.09E+00 | 1.22E-01 | 1.09E+00 |
| ENSG000002 | PTPN20 | 1.47E-01 | ? | 7.59E-01 | -3.98E-01 | -1.32E+00 |
| ENSG000002 | --- | 1.47E-01 | 4.84E-01 | 1.06E+00 | 8.76E-02 | 1.06E+00 |
| ENSG000001 | PTPMT1 | 1.47E-01 | 4.85E-01 | 1.06E+00 | 8.43E-02 | 1.06E+00 |
| ENSG000001 | DYRK2 | 1.47E-01 | 4.85E-01 | 1.07E+00 | 9.80E-02 | 1.07E+00 |
| ENSG000001 | PRPS2 | 1.47E-01 | 4.85E-01 | 1.06E+00 | 8.31E-02 | 1.06E+00 |
| ENSG000001 | OBSCN | 1.47E-01 | 4.85E-01 | 9.06E-01 | -1.42E-01 | -1.10E+00 |
| ENSG000001 | AKAP3 | 1.47E-01 | 4.85E-01 | 1.18E+00 | 2.39E-01 | 1.18E+00 |
| ENSG000002 | --- | 1.47E-01 | ? | 1.73E+00 | 7.87E-01 | 1.73E+00 |
| ENSG000001 | CNOT10 | 1.48E-01 | 4.86E-01 | 9.31E-01 | -1.04E-01 | -1.07E+00 |
| ENSG000001 | CXCL16 | 1.48E-01 | 4.86E-01 | 1.10E+00 | 1.39E-01 | 1.10E+00 |
| ENSG000001 | PRDM4 | 1.48E-01 | 4.86E-01 | 1.04E+00 | 5.60E-02 | 1.04E+00 |
| ENSG000001 | ASTN2 | 1.48E-01 | ? | 1.61E+00 | 6.87E-01 | 1.61E+00 |
| ENSG000002 | FAM200B | 1.48E-01 | 4.86E-01 | 1.08E+00 | 1.09E-01 | 1.08E+00 |
| ENSG000001 | TOP3B | 1.48E-01 | 4.86E-01 | 9.28E-01 | -1.07E-01 | -1.08E+00 |
| ENSG000001 | TLN1 | 1.48E-01 | 4.86E-01 | 9.59E-01 | -6.00E-02 | -1.04E+00 |
| ENSG000001 | RPL15 | 1.48E-01 | 4.86E-01 | 1.04E+00 | 5.93E-02 | 1.04E+00 |
| ENSG000001 | TOM1 | 1.48E-01 | 4.86E-01 | 1.07E+00 | 9.80E-02 | 1.07E+00 |
| ENSG000001 | GNL2 | 1.48E-01 | 4.86E-01 | 9.48E-01 | -7.69E-02 | -1.05E+00 |
| ENSG000001 | EYS | 1.48E-01 | 4.86E-01 | 8.00E-01 | -3.23E-01 | -1.25E+00 |
| ENSG000001 | APOOL | 1.48E-01 | 4.86E-01 | 9.35E-01 | -9.72E-02 | -1.07E+00 |
| ENSG00000C | CERS4 | 1.48E-01 | 4.86E-01 | 1.21E+00 | 2.75E-01 | 1.21E+00 |
| ENSG000001 | ZDHHC20 | 1.48E-01 | 4.86E-01 | 9.51E-01 | -7.22E-02 | -1.05E+00 |
| ENSG000002 | DHFR | 1.48E-01 | 4.86E-01 | 9.02E-01 | -1.49E-01 | -1.11E+00 |
| ENSG000001 | PADI2 | 1.48E-01 | ? | 7.33E-01 | -4.48E-01 | -1.36E+00 |
| ENSG000002 | GLYCTK-AS1 | 1.48E-01 | ? | 1.68E+00 | 7.51E-01 | 1.68E+00 |
| ENSG000002 | --- | 1.48E-01 | 4.86E-01 | 8.48E-01 | -2.37E-01 | -1.18E+00 |
| ENSG000002 | RPSAP15 | 1.48E-01 | 4.86E-01 | 1.15E+00 | 2.04E-01 | 1.15E+00 |
| ENSG000001 | C2orf49 | 1.48E-01 | 4.86E-01 | 9.36E-01 | -9.61E-02 | -1.07E+00 |
| ENSG00000C | NANS | 1.48E-01 | 4.86E-01 | 9.52E-01 | -7.08E-02 | -1.05E+00 |
| ENSG000001 | PMS2P5 | 1.48E-01 | 4.86E-01 | 8.13E-01 | -2.99E-01 | -1.23E+00 |
| ENSG000001 | ISCU | 1.49E-01 | 4.86E-01 | 1.04E+00 | 6.27E-02 | 1.04E+00 |
| ENSG000001 | TMEM129 | 1.49E-01 | 4.86E-01 | 1.15E+00 | 2.06E-01 | 1.15E+00 |
| ENSG00000C | AP2S1 | 1.49E-01 | 4.86E-01 | 1.09E+00 | 1.24E-01 | 1.09E+00 |
| ENSG000002 | --- | 1.49E-01 | 4.86E-01 | 1.15E+00 | 1.97E-01 | 1.15E+00 |
| ENSG000002 | COA8 | 1.49E-01 | 4.86E-01 | 9.54E-01 | -6.83E-02 | -1.05E+00 |
| ENSG000001 | MARK1 | 1.49E-01 | 4.86E-01 | 1.08E+00 | 1.06E-01 | 1.08E+00 |
| ENSG000001 | REEP3 | 1.49E-01 | 4.86E-01 | 9.56E-01 | -6.44E-02 | -1.05E+00 |
| ENSG000001 | CDR2L | 1.49E-01 | 4.86E-01 | 1.06E+00 | 9.04E-02 | 1.06E+00 |
| ENSG000001 | SETD2 | 1.49E-01 | 4.86E-01 | 9.39E-01 | -9.11E-02 | -1.07E+00 |

| | | | | | | |
|---|---|---|---|---|---|---|
| ENSG000001 | DESI1 | 1.49E-01 | 4.86E-01 | 1.06E+00 | 8.29E-02 | 1.06E+00 |
| ENSG000001 | COL22A1 | 1.49E-01 | 4.86E-01 | 8.24E-01 | -2.79E-01 | -1.21E+00 |
| ENSG000001 | ZNF12 | 1.49E-01 | 4.86E-01 | 1.05E+00 | 6.68E-02 | 1.05E+00 |
| ENSG000002 | HCG11 | 1.49E-01 | 4.86E-01 | 1.09E+00 | 1.19E-01 | 1.09E+00 |
| ENSG000002 | --- | 1.49E-01 | ? | 1.97E+00 | 9.75E-01 | 1.97E+00 |
| ENSG000001 | SGK1 | 1.49E-01 | 4.87E-01 | 1.08E+00 | 1.17E-01 | 1.08E+00 |
| ENSG000002 | --- | 1.49E-01 | ? | 6.63E-01 | -5.94E-01 | -1.51E+00 |
| ENSG000001 | ZNF704 | 1.49E-01 | 4.87E-01 | 1.10E+00 | 1.43E-01 | 1.10E+00 |
| ENSG000001 | ANXA7 | 1.49E-01 | 4.87E-01 | 1.04E+00 | 5.26E-02 | 1.04E+00 |
| ENSG000001 | UQCRFS1 | 1.49E-01 | 4.87E-01 | 1.05E+00 | 6.93E-02 | 1.05E+00 |
| ENSG000001 | OXTR | 1.49E-01 | ? | 7.34E-01 | -4.46E-01 | -1.36E+00 |
| ENSG000002 | --- | 1.49E-01 | ? | 2.20E+00 | 1.13E+00 | 2.20E+00 |
| ENSG000001 | CARNMT1 | 1.49E-01 | 4.87E-01 | 9.25E-01 | -1.13E-01 | -1.08E+00 |
| ENSG000001 | ZNHIT2 | 1.49E-01 | 4.87E-01 | 1.23E+00 | 3.00E-01 | 1.23E+00 |
| ENSG000001 | SYNDIG1 | 1.49E-01 | ? | 5.71E-01 | -8.09E-01 | -1.75E+00 |
| ENSG000001 | MIS18BP1 | 1.49E-01 | 4.87E-01 | 9.28E-01 | -1.08E-01 | -1.08E+00 |
| ENSG00000C | MBD3 | 1.49E-01 | 4.87E-01 | 1.14E+00 | 1.88E-01 | 1.14E+00 |
| ENSG000002 | --- | 1.49E-01 | ? | 1.46E+00 | 5.50E-01 | 1.46E+00 |
| ENSG000002 | PIERCE2 | 1.49E-01 | 4.87E-01 | 8.45E-01 | -2.43E-01 | -1.18E+00 |
| ENSG000001 | ZBTB40 | 1.49E-01 | 4.87E-01 | 9.54E-01 | -6.75E-02 | -1.05E+00 |
| ENSG00000C | LRRC40 | 1.49E-01 | 4.87E-01 | 1.07E+00 | 9.27E-02 | 1.07E+00 |
| ENSG000001 | ERCC4 | 1.49E-01 | 4.87E-01 | 9.48E-01 | -7.71E-02 | -1.05E+00 |
| ENSG000001 | BTBD2 | 1.49E-01 | 4.87E-01 | 1.08E+00 | 1.13E-01 | 1.08E+00 |
| ENSG000002 | --- | 1.49E-01 | ? | 6.44E-01 | -6.34E-01 | -1.55E+00 |
| ENSG000001 | MROH8 | 1.50E-01 | ? | 1.50E+00 | 5.86E-01 | 1.50E+00 |
| ENSG000002 | RRAGC-DT | 1.50E-01 | ? | 1.41E+00 | 4.98E-01 | 1.41E+00 |
| ENSG000002 | LDHAP7 | 1.50E-01 | ? | 7.62E-01 | -3.92E-01 | -1.31E+00 |
| ENSG00000C | JMJD4 | 1.50E-01 | 4.88E-01 | 1.13E+00 | 1.78E-01 | 1.13E+00 |
| ENSG000002 | --- | 1.50E-01 | ? | 7.30E-01 | -4.53E-01 | -1.37E+00 |
| ENSG000002 | --- | 1.50E-01 | 4.88E-01 | 8.36E-01 | -2.58E-01 | -1.20E+00 |
| ENSG000001 | SPACA6 | 1.50E-01 | 4.89E-01 | 8.29E-01 | -2.71E-01 | -1.21E+00 |
| ENSG00000C | RRN3 | 1.50E-01 | 4.89E-01 | 1.04E+00 | 5.67E-02 | 1.04E+00 |
| ENSG000001 | CEP170 | 1.50E-01 | 4.89E-01 | 9.28E-01 | -1.08E-01 | -1.08E+00 |
| ENSG000002 | ZNF880 | 1.50E-01 | 4.89E-01 | 1.07E+00 | 1.03E-01 | 1.07E+00 |
| ENSG00000C | WDR54 | 1.50E-01 | 4.89E-01 | 1.09E+00 | 1.31E-01 | 1.09E+00 |
| ENSG000001 | DQX1 | 1.50E-01 | 4.89E-01 | 1.10E+00 | 1.41E-01 | 1.10E+00 |
| ENSG000001 | NCAPG | 1.50E-01 | 4.89E-01 | 8.56E-01 | -2.23E-01 | -1.17E+00 |
| ENSG000002 | --- | 1.50E-01 | 4.89E-01 | 8.69E-01 | -2.02E-01 | -1.15E+00 |
| ENSG000001 | CCDC66 | 1.51E-01 | 4.89E-01 | 9.22E-01 | -1.18E-01 | -1.09E+00 |
| ENSG00000C | NEDD4L | 1.51E-01 | 4.89E-01 | 9.38E-01 | -9.28E-02 | -1.07E+00 |
| ENSG000002 | --- | 1.51E-01 | ? | 1.31E+00 | 3.87E-01 | 1.31E+00 |
| ENSG000002 | MAPKAPK5-A | 1.51E-01 | 4.89E-01 | 9.11E-01 | -1.35E-01 | -1.10E+00 |
| ENSG000002 | --- | 1.51E-01 | ? | 8.07E-01 | -3.09E-01 | -1.24E+00 |
| ENSG000001 | SORCS3 | 1.51E-01 | ? | 1.87E+00 | 9.04E-01 | 1.87E+00 |
| ENSG000001 | MARCHF4 | 1.51E-01 | 4.89E-01 | 9.30E-01 | -1.04E-01 | -1.07E+00 |

| | | | | | | |
|---|---|---|---|---|---|---|
| ENSG000001 | WASH2P | 1.51E-01 | ? | 2.23E+00 | 1.16E+00 | 2.23E+00 |
| ENSG000001 | CREBZF | 1.51E-01 | 4.89E-01 | 9.34E-01 | -9.86E-02 | -1.07E+00 |
| ENSG000002 | CWC25 | 1.51E-01 | 4.89E-01 | 9.36E-01 | -9.51E-02 | -1.07E+00 |
| ENSG000002 | HMGB1P31 | 1.51E-01 | ? | 1.94E+00 | 9.54E-01 | 1.94E+00 |
| ENSG000001 | CUL4B | 1.51E-01 | 4.89E-01 | 1.04E+00 | 5.82E-02 | 1.04E+00 |
| ENSG000002 | --- | 1.51E-01 | ? | 6.82E-01 | -5.52E-01 | -1.47E+00 |
| ENSG000001 | MYO1D | 1.51E-01 | 4.90E-01 | 1.05E+00 | 7.44E-02 | 1.05E+00 |
| ENSG000001 | DHX57 | 1.51E-01 | 4.90E-01 | 1.06E+00 | 8.64E-02 | 1.06E+00 |
| ENSG000002 | LINC02154 | 1.51E-01 | 4.90E-01 | 7.57E-01 | -4.01E-01 | -1.32E+00 |
| ENSG000001 | STAP2 | 1.51E-01 | 4.90E-01 | 1.09E+00 | 1.18E-01 | 1.09E+00 |
| ENSG000001 | DUSP3 | 1.51E-01 | 4.90E-01 | 1.05E+00 | 7.16E-02 | 1.05E+00 |
| ENSG000001 | ELF2 | 1.51E-01 | 4.90E-01 | 1.06E+00 | 8.87E-02 | 1.06E+00 |
| ENSG000002 | --- | 1.51E-01 | ? | 5.88E-01 | -7.66E-01 | -1.70E+00 |
| ENSG000001 | SH3GL2 | 1.51E-01 | ? | 5.83E-01 | -7.79E-01 | -1.72E+00 |
| ENSG000001 | USP19 | 1.51E-01 | 4.90E-01 | 1.08E+00 | 1.05E-01 | 1.08E+00 |
| ENSG00000C | MECOM | 1.51E-01 | 4.90E-01 | 9.45E-01 | -8.19E-02 | -1.06E+00 |
| ENSG00000C | FUT8 | 1.51E-01 | 4.91E-01 | 1.05E+00 | 7.61E-02 | 1.05E+00 |
| ENSG000001 | GET3 | 1.52E-01 | 4.91E-01 | 1.07E+00 | 1.01E-01 | 1.07E+00 |
| ENSG000001 | SH3PXD2A | 1.52E-01 | 4.91E-01 | 1.05E+00 | 7.59E-02 | 1.05E+00 |
| ENSG000001 | ZBTB8OS | 1.52E-01 | 4.91E-01 | 9.34E-01 | -9.90E-02 | -1.07E+00 |
| ENSG000001 | MYH11 | 1.52E-01 | 4.91E-01 | 1.25E+00 | 3.19E-01 | 1.25E+00 |
| ENSG00000C | RECQL | 1.52E-01 | 4.91E-01 | 9.65E-01 | -5.20E-02 | -1.04E+00 |
| ENSG000002 | SORD2P | 1.52E-01 | 4.91E-01 | 8.53E-01 | -2.30E-01 | -1.17E+00 |
| ENSG000001 | CIAO3 | 1.52E-01 | 4.91E-01 | 1.11E+00 | 1.53E-01 | 1.11E+00 |
| ENSG000001 | SLFN11 | 1.52E-01 | ? | 7.84E-01 | -3.51E-01 | -1.28E+00 |
| ENSG000001 | GFM2 | 1.52E-01 | 4.91E-01 | 9.54E-01 | -6.73E-02 | -1.05E+00 |
| ENSG000001 | ZNF157 | 1.52E-01 | ? | 5.05E-01 | -9.86E-01 | -1.98E+00 |
| ENSG000001 | ESRRG | 1.52E-01 | ? | 5.86E-01 | -7.71E-01 | -1.71E+00 |
| ENSG000001 | ZMYM5 | 1.52E-01 | 4.91E-01 | 9.38E-01 | -9.23E-02 | -1.07E+00 |
| ENSG000001 | PLEK | 1.52E-01 | ? | 1.95E+00 | 9.60E-01 | 1.95E+00 |
| ENSG000001 | ABCB10 | 1.52E-01 | 4.91E-01 | 9.41E-01 | -8.75E-02 | -1.06E+00 |
| ENSG000002 | LPP-AS2 | 1.52E-01 | 4.91E-01 | 1.20E+00 | 2.68E-01 | 1.20E+00 |
| ENSG000001 | LHX4 | 1.52E-01 | 4.92E-01 | 8.47E-01 | -2.39E-01 | -1.18E+00 |
| ENSG000002 | IQANK1 | 1.52E-01 | 4.92E-01 | 1.08E+00 | 1.11E-01 | 1.08E+00 |
| ENSG000001 | KIAA2026 | 1.52E-01 | 4.92E-01 | 9.32E-01 | -1.02E-01 | -1.07E+00 |
| ENSG000001 | FBLIM1 | 1.52E-01 | 4.92E-01 | 1.04E+00 | 5.50E-02 | 1.04E+00 |
| ENSG000001 | DUOX1 | 1.52E-01 | 4.92E-01 | 1.08E+00 | 1.05E-01 | 1.08E+00 |
| ENSG000001 | FTSJ3 | 1.53E-01 | 4.92E-01 | 9.42E-01 | -8.62E-02 | -1.06E+00 |
| ENSG000002 | --- | 1.53E-01 | ? | 6.33E-01 | -6.60E-01 | -1.58E+00 |
| ENSG000001 | HMGA2 | 1.53E-01 | 4.92E-01 | 1.06E+00 | 7.99E-02 | 1.06E+00 |
| ENSG000001 | ZNF415 | 1.53E-01 | 4.92E-01 | 9.07E-01 | -1.40E-01 | -1.10E+00 |
| ENSG000001 | SRSF10 | 1.53E-01 | 4.93E-01 | 9.61E-01 | -5.80E-02 | -1.04E+00 |
| ENSG00000C | TLE2 | 1.53E-01 | ? | 1.39E+00 | 4.73E-01 | 1.39E+00 |
| ENSG000001 | PDSS1 | 1.53E-01 | 4.93E-01 | 9.17E-01 | -1.25E-01 | -1.09E+00 |
| ENSG000001 | TIAM1 | 1.53E-01 | 4.93E-01 | 1.06E+00 | 8.34E-02 | 1.06E+00 |

| | | | | | | | |
|---|---|---|---|---|---|---|---|
| ENSG000001 | PNO1 | 1.53E-01 | | 4.93E-01 | 1.04E+00 | 5.97E-02 | 1.04E+00 |
| ENSG000001 | PKP3 | 1.53E-01 | | 4.93E-01 | 1.10E+00 | 1.34E-01 | 1.10E+00 |
| ENSG000002 | YEATS2-AS1 | 1.53E-01 | | 4.93E-01 | 7.64E-01 | -3.88E-01 | -1.31E+00 |
| ENSG00000C | WDR47 | 1.53E-01 | | 4.93E-01 | 1.04E+00 | 5.43E-02 | 1.04E+00 |
| ENSG000001 | CHCHD2 | 1.53E-01 | | 4.93E-01 | 1.07E+00 | 9.87E-02 | 1.07E+00 |
| ENSG000002 | --- | 1.53E-01 | ? | | 6.95E-01 | -5.26E-01 | -1.44E+00 |
| ENSG000001 | SGK2 | 1.53E-01 | ? | | 1.69E+00 | 7.59E-01 | 1.69E+00 |
| ENSG000001 | ZNF503 | 1.53E-01 | | 4.93E-01 | 9.24E-01 | -1.15E-01 | -1.08E+00 |
| ENSG000001 | PARP15 | 1.53E-01 | ? | | 5.98E-01 | -7.42E-01 | -1.67E+00 |
| ENSG000001 | MMP16 | 1.53E-01 | ? | | 6.89E-01 | -5.38E-01 | -1.45E+00 |
| ENSG000001 | RIMKLA | 1.53E-01 | ? | | 6.51E-01 | -6.19E-01 | -1.54E+00 |
| ENSG000001 | SURF1 | 1.53E-01 | | 4.94E-01 | 1.08E+00 | 1.16E-01 | 1.08E+00 |
| ENSG000001 | LIPH | 1.54E-01 | | 4.94E-01 | 9.14E-01 | -1.29E-01 | -1.09E+00 |
| ENSG000002 | NPIPA9 | 1.54E-01 | ? | | 7.39E-01 | -4.37E-01 | -1.35E+00 |
| ENSG000001 | AAR2 | 1.54E-01 | | 4.94E-01 | 1.08E+00 | 1.13E-01 | 1.08E+00 |
| ENSG000001 | BRIX1 | 1.54E-01 | | 4.94E-01 | 9.37E-01 | -9.45E-02 | -1.07E+00 |
| ENSG000001 | GID4 | 1.54E-01 | | 4.94E-01 | 1.06E+00 | 7.98E-02 | 1.06E+00 |
| ENSG000001 | AMMECR1 | 1.54E-01 | | 4.94E-01 | 1.07E+00 | 9.12E-02 | 1.07E+00 |
| ENSG00000C | CLASP1 | 1.54E-01 | | 4.94E-01 | 9.64E-01 | -5.32E-02 | -1.04E+00 |
| ENSG000001 | CLASRP | 1.54E-01 | | 4.94E-01 | 9.41E-01 | -8.79E-02 | -1.06E+00 |
| ENSG000001 | SKIDA1 | 1.54E-01 | ? | | 1.26E+00 | 3.30E-01 | 1.26E+00 |
| ENSG000002 | ARHGEF33 | 1.54E-01 | ? | | 6.85E-01 | -5.45E-01 | -1.46E+00 |
| ENSG000002 | TRAF3IP2-AS | 1.54E-01 | ? | | 7.50E-01 | -4.15E-01 | -1.33E+00 |
| ENSG000002 | --- | 1.54E-01 | ? | | 1.29E+00 | 3.68E-01 | 1.29E+00 |
| ENSG000001 | KLHDC2 | 1.54E-01 | | 4.95E-01 | 9.37E-01 | -9.42E-02 | -1.07E+00 |
| ENSG000001 | PPID | 1.54E-01 | | 4.95E-01 | 9.46E-01 | -7.98E-02 | -1.06E+00 |
| ENSG000001 | NUPR1 | 1.54E-01 | | 4.95E-01 | 1.10E+00 | 1.41E-01 | 1.10E+00 |
| ENSG000001 | ZNF639 | 1.54E-01 | | 4.95E-01 | 1.06E+00 | 8.59E-02 | 1.06E+00 |
| ENSG000002 | TADA2A | 1.54E-01 | | 4.95E-01 | 1.08E+00 | 1.07E-01 | 1.08E+00 |
| ENSG000002 | NOL7 | 1.54E-01 | | 4.95E-01 | 9.55E-01 | -6.58E-02 | -1.05E+00 |
| ENSG00000C | DLL3 | 1.54E-01 | ? | | 1.31E+00 | 3.87E-01 | 1.31E+00 |
| ENSG000002 | LINC01963 | 1.54E-01 | | 4.95E-01 | 9.09E-01 | -1.38E-01 | -1.10E+00 |
| ENSG000002 | --- | 1.54E-01 | ? | | 5.94E-01 | -7.51E-01 | -1.68E+00 |
| ENSG000001 | MT-ND4 | 1.54E-01 | | 4.95E-01 | 8.60E-01 | -2.17E-01 | -1.16E+00 |
| ENSG000002 | NSRP1P1 | 1.54E-01 | ? | | 2.67E+00 | 1.42E+00 | 2.67E+00 |
| ENSG000001 | BFAR | 1.54E-01 | | 4.95E-01 | 1.04E+00 | 6.28E-02 | 1.04E+00 |
| ENSG000002 | --- | 1.54E-01 | ? | | 5.58E-01 | -8.42E-01 | -1.79E+00 |
| ENSG000001 | SLC16A4 | 1.54E-01 | | 4.95E-01 | 9.04E-01 | -1.45E-01 | -1.11E+00 |
| ENSG000001 | SQOR | 1.55E-01 | | 4.95E-01 | 1.05E+00 | 6.56E-02 | 1.05E+00 |
| ENSG000001 | PPARG | 1.55E-01 | | 4.95E-01 | 1.12E+00 | 1.58E-01 | 1.12E+00 |
| ENSG000001 | RLIM | 1.55E-01 | | 4.95E-01 | 1.05E+00 | 6.48E-02 | 1.05E+00 |
| ENSG000001 | R3HCC1 | 1.55E-01 | | 4.95E-01 | 1.06E+00 | 8.91E-02 | 1.06E+00 |
| ENSG000001 | ASAH1 | 1.55E-01 | | 4.95E-01 | 1.05E+00 | 6.66E-02 | 1.05E+00 |
| ENSG000002 | CYP2D7 | 1.55E-01 | | 4.95E-01 | 1.17E+00 | 2.22E-01 | 1.17E+00 |
| ENSG000001 | IL15 | 1.55E-01 | | 4.95E-01 | 7.77E-01 | -3.64E-01 | -1.29E+00 |

| | | | | | | |
|---|---|---|---|---|---|---|
| ENSG000001 | SNX33 | 1.55E-01 | 4.95E-01 | 1.05E+00 | 7.52E-02 | 1.05E+00 |
| ENSG000001 | SUGT1 | 1.55E-01 | 4.95E-01 | 9.54E-01 | -6.76E-02 | -1.05E+00 |
| ENSG000001 | SNTB1 | 1.55E-01 | ? | 1.27E+00 | 3.48E-01 | 1.27E+00 |
| ENSG000002 | PPP1R3E | 1.55E-01 | 4.95E-01 | 1.16E+00 | 2.16E-01 | 1.16E+00 |
| ENSG000001 | ZNF44 | 1.55E-01 | 4.95E-01 | 9.17E-01 | -1.25E-01 | -1.09E+00 |
| ENSG00000C | KHSRP | 1.55E-01 | 4.95E-01 | 9.53E-01 | -6.97E-02 | -1.05E+00 |
| ENSG000001 | STT3A | 1.55E-01 | 4.95E-01 | 9.61E-01 | -5.72E-02 | -1.04E+00 |
| ENSG000001 | MEAF6 | 1.55E-01 | 4.95E-01 | 1.05E+00 | 6.69E-02 | 1.05E+00 |
| ENSG000001 | ZNF396 | 1.55E-01 | ? | 1.28E+00 | 3.57E-01 | 1.28E+00 |
| ENSG000001 | SLC25A26 | 1.55E-01 | 4.96E-01 | 1.08E+00 | 1.14E-01 | 1.08E+00 |
| ENSG000001 | ITPRIPL1 | 1.55E-01 | ? | 1.38E+00 | 4.68E-01 | 1.38E+00 |
| ENSG000001 | ESD | 1.55E-01 | 4.96E-01 | 1.04E+00 | 5.87E-02 | 1.04E+00 |
| ENSG000001 | STK36 | 1.55E-01 | 4.96E-01 | 1.07E+00 | 9.93E-02 | 1.07E+00 |
| ENSG000001 | WDFY4 | 1.55E-01 | ? | 1.48E+00 | 5.63E-01 | 1.48E+00 |
| ENSG000001 | LINC00174 | 1.55E-01 | 4.96E-01 | 8.98E-01 | -1.55E-01 | -1.11E+00 |
| ENSG000001 | MIDN | 1.55E-01 | 4.96E-01 | 9.39E-01 | -9.04E-02 | -1.06E+00 |
| ENSG000001 | ZBTB1 | 1.55E-01 | 4.96E-01 | 9.24E-01 | -1.13E-01 | -1.08E+00 |
| ENSG000001 | RAB28 | 1.55E-01 | 4.96E-01 | 1.07E+00 | 9.90E-02 | 1.07E+00 |
| ENSG000002 | --- | 1.56E-01 | ? | 1.67E+00 | 7.36E-01 | 1.67E+00 |
| ENSG000001 | PPP4R4 | 1.56E-01 | 4.96E-01 | 1.07E+00 | 9.56E-02 | 1.07E+00 |
| ENSG00000C | WASHC2A | 1.56E-01 | 4.96E-01 | 9.62E-01 | -5.55E-02 | -1.04E+00 |
| ENSG000001 | GTF2H2C | 1.56E-01 | 4.96E-01 | 9.52E-01 | -7.17E-02 | -1.05E+00 |
| ENSG000001 | TMC1 | 1.56E-01 | ? | 1.82E+00 | 8.63E-01 | 1.82E+00 |
| ENSG000001 | CYP1A2 | 1.56E-01 | ? | 1.64E+00 | 7.11E-01 | 1.64E+00 |
| ENSG000001 | PC | 1.56E-01 | 4.97E-01 | 1.12E+00 | 1.58E-01 | 1.12E+00 |
| ENSG000001 | KCNIP2 | 1.56E-01 | ? | 6.79E-01 | -5.58E-01 | -1.47E+00 |
| ENSG000002 | --- | 1.56E-01 | ? | 1.68E+00 | 7.50E-01 | 1.68E+00 |
| ENSG000001 | ADCY3 | 1.56E-01 | 4.97E-01 | 1.07E+00 | 9.47E-02 | 1.07E+00 |
| ENSG000001 | ZNF317 | 1.56E-01 | 4.97E-01 | 9.53E-01 | -6.89E-02 | -1.05E+00 |
| ENSG00000C | RBM5 | 1.56E-01 | 4.97E-01 | 9.40E-01 | -8.95E-02 | -1.06E+00 |
| ENSG000001 | TMEM64 | 1.56E-01 | 4.97E-01 | 1.06E+00 | 8.75E-02 | 1.06E+00 |
| ENSG000001 | PLK1 | 1.56E-01 | 4.97E-01 | 8.68E-01 | -2.05E-01 | -1.15E+00 |
| ENSG000001 | DIAPH3 | 1.56E-01 | 4.97E-01 | 9.19E-01 | -1.21E-01 | -1.09E+00 |
| ENSG000001 | COA5 | 1.56E-01 | 4.97E-01 | 1.08E+00 | 1.07E-01 | 1.08E+00 |
| ENSG00000C | ADAMTS6 | 1.56E-01 | 4.97E-01 | 9.18E-01 | -1.24E-01 | -1.09E+00 |
| ENSG000001 | TRIM41 | 1.56E-01 | 4.97E-01 | 9.42E-01 | -8.61E-02 | -1.06E+00 |
| ENSG00000C | FRY | 1.56E-01 | 4.97E-01 | 8.11E-01 | -3.03E-01 | -1.23E+00 |
| ENSG000001 | AZI2 | 1.56E-01 | 4.97E-01 | 9.51E-01 | -7.25E-02 | -1.05E+00 |
| ENSG000001 | ATXN10 | 1.56E-01 | 4.97E-01 | 1.04E+00 | 5.30E-02 | 1.04E+00 |
| ENSG000002 | RAD51AP2 | 1.57E-01 | ? | 7.24E-01 | -4.66E-01 | -1.38E+00 |
| ENSG000001 | TARDBP | 1.57E-01 | 4.97E-01 | 9.54E-01 | -6.81E-02 | -1.05E+00 |
| ENSG00000C | EXOSC7 | 1.57E-01 | 4.97E-01 | 1.06E+00 | 8.50E-02 | 1.06E+00 |
| ENSG000001 | GNPDA2 | 1.57E-01 | 4.97E-01 | 9.35E-01 | -9.66E-02 | -1.07E+00 |
| ENSG000001 | BICRAL | 1.57E-01 | 4.98E-01 | 1.06E+00 | 8.93E-02 | 1.06E+00 |
| ENSG000002 | LURAP1L-AS | 1.57E-01 | 4.98E-01 | 8.12E-01 | -3.00E-01 | -1.23E+00 |

| | | | | | | |
|---|---|---|---|---|---|---|
| ENSG000001 | RNF114 | 1.57E-01 | 4.98E-01 | 9.54E-01 | -6.74E-02 | -1.05E+00 |
| ENSG000001 | SMCR8 | 1.57E-01 | 4.98E-01 | 1.04E+00 | 6.21E-02 | 1.04E+00 |
| ENSG000002 | --- | 1.57E-01 | ? | 6.83E-01 | -5.50E-01 | -1.46E+00 |
| ENSG000001 | ATP23 | 1.57E-01 | 4.98E-01 | 1.16E+00 | 2.11E-01 | 1.16E+00 |
| ENSG000001 | PHLDA3 | 1.57E-01 | 4.98E-01 | 1.10E+00 | 1.43E-01 | 1.10E+00 |
| ENSG000001 | ABCG2 | 1.57E-01 | 4.98E-01 | 1.16E+00 | 2.19E-01 | 1.16E+00 |
| ENSG000002 | --- | 1.57E-01 | ? | 1.83E+00 | 8.71E-01 | 1.83E+00 |
| ENSG000001 | SAFB2 | 1.57E-01 | 4.99E-01 | 9.33E-01 | -9.94E-02 | -1.07E+00 |
| ENSG000001 | UBE4A | 1.57E-01 | 4.99E-01 | 9.59E-01 | -6.05E-02 | -1.04E+00 |
| ENSG000001 | LIG1 | 1.57E-01 | 4.99E-01 | 1.09E+00 | 1.29E-01 | 1.09E+00 |
| ENSG000001 | TMEM209 | 1.57E-01 | 4.99E-01 | 9.37E-01 | -9.44E-02 | -1.07E+00 |
| ENSG000001 | CELF6 | 1.57E-01 | ? | 1.62E+00 | 6.94E-01 | 1.62E+00 |
| ENSG000001 | STOML2 | 1.57E-01 | 4.99E-01 | 1.07E+00 | 9.99E-02 | 1.07E+00 |
| ENSG000001 | BICC1 | 1.57E-01 | 4.99E-01 | 9.41E-01 | -8.79E-02 | -1.06E+00 |
| ENSG000001 | ZGRF1 | 1.58E-01 | 4.99E-01 | 8.52E-01 | -2.31E-01 | -1.17E+00 |
| ENSG000001 | CPNE4 | 1.58E-01 | ? | 5.62E-01 | -8.30E-01 | -1.78E+00 |
| ENSG00000C | PITPNM3 | 1.58E-01 | 4.99E-01 | 1.09E+00 | 1.21E-01 | 1.09E+00 |
| ENSG000002 | GOLGA8B | 1.58E-01 | 4.99E-01 | 8.87E-01 | -1.73E-01 | -1.13E+00 |
| ENSG000001 | RNF144B | 1.58E-01 | 4.99E-01 | 1.07E+00 | 9.84E-02 | 1.07E+00 |
| ENSG000002 | DTX2P1-UPK | 1.58E-01 | 4.99E-01 | 9.28E-01 | -1.09E-01 | -1.08E+00 |
| ENSG000001 | CBX3 | 1.58E-01 | 4.99E-01 | 1.04E+00 | 5.57E-02 | 1.04E+00 |
| ENSG000001 | UCK2 | 1.58E-01 | 4.99E-01 | 9.56E-01 | -6.50E-02 | -1.05E+00 |
| ENSG000002 | FAM88E | 1.58E-01 | ? | 1.80E+00 | 8.46E-01 | 1.80E+00 |
| ENSG000001 | STARD8 | 1.58E-01 | ? | 6.50E-01 | -6.22E-01 | -1.54E+00 |
| ENSG000001 | AEBP1 | 1.58E-01 | 5.00E-01 | 8.35E-01 | -2.59E-01 | -1.20E+00 |
| ENSG00000C | MAN2B2 | 1.58E-01 | 5.00E-01 | 1.04E+00 | 6.26E-02 | 1.04E+00 |
| ENSG000001 | SIGLEC15 | 1.58E-01 | ? | 1.61E+00 | 6.85E-01 | 1.61E+00 |
| ENSG000001 | LDAH | 1.58E-01 | 5.00E-01 | 9.42E-01 | -8.64E-02 | -1.06E+00 |
| ENSG000001 | FGD6 | 1.58E-01 | 5.00E-01 | 1.06E+00 | 8.85E-02 | 1.06E+00 |
| ENSG000002 | --- | 1.58E-01 | 5.00E-01 | 9.03E-01 | -1.48E-01 | -1.11E+00 |
| ENSG000002 | ATXN1-AS1 | 1.58E-01 | 5.00E-01 | 1.23E+00 | 2.93E-01 | 1.23E+00 |
| ENSG000001 | TRPC4AP | 1.58E-01 | 5.00E-01 | 1.07E+00 | 9.54E-02 | 1.07E+00 |
| ENSG000002 | SKP1P2 | 1.58E-01 | ? | 2.47E+00 | 1.30E+00 | 2.47E+00 |
| ENSG000002 | --- | 1.58E-01 | ? | 6.09E-01 | -7.15E-01 | -1.64E+00 |
| ENSG000001 | GALNT14 | 1.58E-01 | 5.00E-01 | 1.06E+00 | 8.07E-02 | 1.06E+00 |
| ENSG000002 | --- | 1.58E-01 | ? | 7.09E-01 | -4.97E-01 | -1.41E+00 |
| ENSG000001 | TCHP | 1.58E-01 | 5.00E-01 | 9.47E-01 | -7.79E-02 | -1.06E+00 |
| ENSG000001 | FILIP1L | 1.58E-01 | 5.00E-01 | 1.19E+00 | 2.49E-01 | 1.19E+00 |
| ENSG000002 | DDAH2 | 1.58E-01 | 5.00E-01 | 1.12E+00 | 1.61E-01 | 1.12E+00 |
| ENSG000001 | MTX1 | 1.59E-01 | 5.00E-01 | 1.08E+00 | 1.12E-01 | 1.08E+00 |
| ENSG000001 | HIGD1A | 1.59E-01 | 5.00E-01 | 9.68E-01 | -4.71E-02 | -1.03E+00 |
| ENSG000001 | NHSL1 | 1.59E-01 | 5.00E-01 | 9.43E-01 | -8.43E-02 | -1.06E+00 |
| ENSG000002 | LINC00896 | 1.59E-01 | ? | 1.57E+00 | 6.47E-01 | 1.57E+00 |
| ENSG000001 | REEP2 | 1.59E-01 | 5.00E-01 | 1.26E+00 | 3.33E-01 | 1.26E+00 |
| ENSG00000C | PKD2L2 | 1.59E-01 | ? | 2.13E+00 | 1.09E+00 | 2.13E+00 |

| | | | | | | |
|---|---|---|---|---|---|---|
| ENSG000000 | XRCC1 | 1.59E-01 | 5.00E-01 | 1.07E+00 | 9.42E-02 | 1.07E+00 |
| ENSG000001 | ALDH1L1 | 1.59E-01 | ? | 1.57E+00 | 6.48E-01 | 1.57E+00 |
| ENSG000002 | --- | 1.59E-01 | ? | 4.68E-01 | -1.10E+00 | -2.14E+00 |
| ENSG000000 | CABIN1 | 1.59E-01 | 5.00E-01 | 1.07E+00 | 9.22E-02 | 1.07E+00 |
| ENSG000000 | ASAP3 | 1.59E-01 | 5.00E-01 | 1.18E+00 | 2.34E-01 | 1.18E+00 |
| ENSG000001 | LUC7L3 | 1.59E-01 | 5.00E-01 | 9.26E-01 | -1.11E-01 | -1.08E+00 |
| ENSG000001 | DDIT4L | 1.59E-01 | 5.00E-01 | 8.20E-01 | -2.87E-01 | -1.22E+00 |
| ENSG000001 | TRIM14 | 1.59E-01 | 5.01E-01 | 1.05E+00 | 7.62E-02 | 1.05E+00 |
| ENSG000000 | TDP1 | 1.59E-01 | 5.01E-01 | 1.06E+00 | 8.36E-02 | 1.06E+00 |
| ENSG000000 | SLC22A17 | 1.59E-01 | ? | 1.31E+00 | 3.84E-01 | 1.31E+00 |
| ENSG000001 | CETN2 | 1.59E-01 | 5.01E-01 | 1.05E+00 | 7.10E-02 | 1.05E+00 |
| ENSG000001 | NEFH | 1.59E-01 | 5.01E-01 | 8.88E-01 | -1.71E-01 | -1.13E+00 |
| ENSG000001 | SFT2D1 | 1.59E-01 | 5.01E-01 | 1.05E+00 | 7.03E-02 | 1.05E+00 |
| ENSG000001 | IGF2BP1 | 1.59E-01 | ? | 5.82E-01 | -7.81E-01 | -1.72E+00 |
| ENSG000000 | PRSS22 | 1.59E-01 | 5.01E-01 | 8.49E-01 | -2.37E-01 | -1.18E+00 |
| ENSG000002 | --- | 1.60E-01 | 5.01E-01 | 1.22E+00 | 2.81E-01 | 1.22E+00 |
| ENSG000002 | MT-ATP8 | 1.60E-01 | 5.01E-01 | 8.09E-01 | -3.05E-01 | -1.24E+00 |
| ENSG000001 | ZNF117 | 1.60E-01 | 5.01E-01 | 8.78E-01 | -1.88E-01 | -1.14E+00 |
| ENSG000002 | --- | 1.60E-01 | 5.01E-01 | 1.18E+00 | 2.38E-01 | 1.18E+00 |
| ENSG000002 | H2BP1 | 1.60E-01 | ? | 1.95E+00 | 9.65E-01 | 1.95E+00 |
| ENSG000002 | SLC25A25-AS | 1.60E-01 | 5.02E-01 | 8.61E-01 | -2.16E-01 | -1.16E+00 |
| ENSG000002 | --- | 1.60E-01 | ? | 6.47E-01 | -6.29E-01 | -1.55E+00 |
| ENSG000001 | FTO | 1.60E-01 | 5.02E-01 | 9.55E-01 | -6.67E-02 | -1.05E+00 |
| ENSG000000 | SERTAD4 | 1.60E-01 | 5.02E-01 | 1.09E+00 | 1.26E-01 | 1.09E+00 |
| ENSG000001 | NT5DC2 | 1.60E-01 | 5.02E-01 | 9.45E-01 | -8.21E-02 | -1.06E+00 |
| ENSG000001 | FAXDC2 | 1.60E-01 | 5.02E-01 | 1.11E+00 | 1.53E-01 | 1.11E+00 |
| ENSG000001 | KCNIP3 | 1.60E-01 | 5.02E-01 | 1.19E+00 | 2.49E-01 | 1.19E+00 |
| ENSG000001 | EFR3A | 1.60E-01 | 5.02E-01 | 1.06E+00 | 7.89E-02 | 1.06E+00 |
| ENSG000001 | PCYOX1L | 1.60E-01 | 5.02E-01 | 9.31E-01 | -1.03E-01 | -1.07E+00 |
| ENSG000001 | TXNRD2 | 1.60E-01 | 5.02E-01 | 1.10E+00 | 1.31E-01 | 1.10E+00 |
| ENSG000001 | TUBB1 | 1.60E-01 | ? | 6.45E-01 | -6.33E-01 | -1.55E+00 |
| ENSG000001 | TDRD6 | 1.60E-01 | ? | 7.12E-01 | -4.89E-01 | -1.40E+00 |
| ENSG000002 | --- | 1.60E-01 | ? | 1.50E+00 | 5.82E-01 | 1.50E+00 |
| ENSG000002 | DOC2B | 1.60E-01 | 5.02E-01 | 1.20E+00 | 2.63E-01 | 1.20E+00 |
| ENSG000000 | B4GALT7 | 1.60E-01 | 5.02E-01 | 1.10E+00 | 1.43E-01 | 1.10E+00 |
| ENSG000001 | AKIP1 | 1.60E-01 | 5.02E-01 | 1.07E+00 | 1.00E-01 | 1.07E+00 |
| ENSG000001 | RNF187 | 1.60E-01 | 5.02E-01 | 1.06E+00 | 8.39E-02 | 1.06E+00 |
| ENSG000002 | --- | 1.60E-01 | ? | 6.36E-01 | -6.53E-01 | -1.57E+00 |
| ENSG000001 | SPC24 | 1.60E-01 | 5.02E-01 | 7.53E-01 | -4.09E-01 | -1.33E+00 |
| ENSG000001 | PTGES2 | 1.61E-01 | 5.02E-01 | 1.08E+00 | 1.08E-01 | 1.08E+00 |
| ENSG000002 | HEIH | 1.61E-01 | 5.02E-01 | 1.09E+00 | 1.29E-01 | 1.09E+00 |
| ENSG000001 | STRN3 | 1.61E-01 | 5.02E-01 | 9.53E-01 | -6.88E-02 | -1.05E+00 |
| ENSG000001 | RILPL2 | 1.61E-01 | 5.02E-01 | 1.09E+00 | 1.22E-01 | 1.09E+00 |
| ENSG000001 | PIP5KL1 | 1.61E-01 | 5.02E-01 | 1.16E+00 | 2.11E-01 | 1.16E+00 |
| ENSG000002 | CFAP58-DT | 1.61E-01 | ? | 4.86E-01 | -1.04E+00 | -2.06E+00 |

| | | | | | | |
|---|---|---|---|---|---|---|
| ENSG000002 | DDX47 | 1.61E-01 | 5.02E-01 | 1.04E+00 | 6.14E-02 | 1.04E+00 |
| ENSG000001 | SPICE1 | 1.61E-01 | 5.02E-01 | 9.29E-01 | -1.06E-01 | -1.08E+00 |
| ENSG00000C | RALBP1 | 1.61E-01 | 5.02E-01 | 1.03E+00 | 4.70E-02 | 1.03E+00 |
| ENSG000001 | TMPPE | 1.61E-01 | 5.02E-01 | 8.83E-01 | -1.79E-01 | -1.13E+00 |
| ENSG000001 | CHGA | 1.61E-01 | ? | 2.02E+00 | 1.01E+00 | 2.02E+00 |
| ENSG000001 | ACADS | 1.61E-01 | 5.02E-01 | 1.17E+00 | 2.31E-01 | 1.17E+00 |
| ENSG000001 | TMEM208 | 1.61E-01 | 5.02E-01 | 1.11E+00 | 1.50E-01 | 1.11E+00 |
| ENSG000001 | GSTM1 | 1.61E-01 | 5.02E-01 | 1.08E+00 | 1.08E-01 | 1.08E+00 |
| ENSG000001 | ABLIM3 | 1.61E-01 | 5.02E-01 | 9.46E-01 | -8.06E-02 | -1.06E+00 |
| ENSG00000C | PPP2R2C | 1.61E-01 | 5.02E-01 | 1.08E+00 | 1.13E-01 | 1.08E+00 |
| ENSG000001 | ACCS | 1.61E-01 | 5.02E-01 | 1.13E+00 | 1.71E-01 | 1.13E+00 |
| ENSG000001 | CNBP | 1.61E-01 | 5.02E-01 | 1.02E+00 | 3.45E-02 | 1.02E+00 |
| ENSG000001 | PDP2 | 1.61E-01 | 5.02E-01 | 1.06E+00 | 8.35E-02 | 1.06E+00 |
| ENSG000001 | PCGF6 | 1.61E-01 | 5.03E-01 | 9.12E-01 | -1.32E-01 | -1.10E+00 |
| ENSG000001 | HNRNPL | 1.61E-01 | 5.03E-01 | 1.05E+00 | 7.06E-02 | 1.05E+00 |
| ENSG000002 | TRBC2 | 1.61E-01 | 5.03E-01 | 8.30E-01 | -2.69E-01 | -1.20E+00 |
| ENSG000001 | H3-3B | 1.61E-01 | 5.03E-01 | 1.03E+00 | 4.88E-02 | 1.03E+00 |
| ENSG00000C | LY75 | 1.61E-01 | 5.03E-01 | 9.13E-01 | -1.31E-01 | -1.09E+00 |
| ENSG000001 | XKRX | 1.61E-01 | ? | 1.73E+00 | 7.89E-01 | 1.73E+00 |
| ENSG000002 | FDX2 | 1.62E-01 | 5.03E-01 | 1.07E+00 | 9.52E-02 | 1.07E+00 |
| ENSG000001 | ATF4 | 1.62E-01 | 5.03E-01 | 9.65E-01 | -5.17E-02 | -1.04E+00 |
| ENSG000002 | COPB2-DT | 1.62E-01 | 5.03E-01 | 1.15E+00 | 2.01E-01 | 1.15E+00 |
| ENSG000001 | C8orf33 | 1.62E-01 | 5.03E-01 | 1.06E+00 | 8.15E-02 | 1.06E+00 |
| ENSG00000C | GSDMB | 1.62E-01 | 5.03E-01 | 8.91E-01 | -1.66E-01 | -1.12E+00 |
| ENSG000001 | H1-10 | 1.62E-01 | 5.03E-01 | 1.12E+00 | 1.63E-01 | 1.12E+00 |
| ENSG000002 | KDSR-DT | 1.62E-01 | ? | 1.57E+00 | 6.53E-01 | 1.57E+00 |
| ENSG000001 | FHIP2B | 1.62E-01 | 5.03E-01 | 1.07E+00 | 9.69E-02 | 1.07E+00 |
| ENSG00000C | PSME1 | 1.62E-01 | 5.03E-01 | 1.06E+00 | 8.43E-02 | 1.06E+00 |
| ENSG000001 | MSI2 | 1.62E-01 | 5.03E-01 | 1.08E+00 | 1.15E-01 | 1.08E+00 |
| ENSG000001 | YME1L1 | 1.62E-01 | 5.03E-01 | 1.04E+00 | 5.04E-02 | 1.04E+00 |
| ENSG00000C | CREBBP | 1.62E-01 | 5.03E-01 | 9.43E-01 | -8.53E-02 | -1.06E+00 |
| ENSG000001 | DENND5B | 1.62E-01 | 5.03E-01 | 1.12E+00 | 1.63E-01 | 1.12E+00 |
| ENSG000001 | NUDT10 | 1.62E-01 | 5.03E-01 | 1.20E+00 | 2.69E-01 | 1.20E+00 |
| ENSG000001 | TNFAIP8L1 | 1.62E-01 | 5.03E-01 | 1.12E+00 | 1.67E-01 | 1.12E+00 |
| ENSG000001 | DLEU1 | 1.62E-01 | 5.03E-01 | 1.14E+00 | 1.89E-01 | 1.14E+00 |
| ENSG000002 | --- | 1.62E-01 | ? | 5.90E-01 | -7.60E-01 | -1.69E+00 |
| ENSG000002 | --- | 1.62E-01 | ? | 5.89E-01 | -7.65E-01 | -1.70E+00 |
| ENSG000002 | AGAP10P | 1.62E-01 | ? | 1.42E+00 | 5.08E-01 | 1.42E+00 |
| ENSG000001 | ABCB6 | 1.62E-01 | 5.04E-01 | 1.12E+00 | 1.60E-01 | 1.12E+00 |
| ENSG000002 | TBC1D22A-D | 1.62E-01 | ? | 7.84E-01 | -3.51E-01 | -1.28E+00 |
| ENSG000001 | PACS1 | 1.62E-01 | 5.04E-01 | 1.06E+00 | 7.93E-02 | 1.06E+00 |
| ENSG00000C | TMEM161A | 1.62E-01 | 5.04E-01 | 1.13E+00 | 1.80E-01 | 1.13E+00 |
| ENSG000001 | LARP7 | 1.62E-01 | 5.04E-01 | 9.45E-01 | -8.14E-02 | -1.06E+00 |
| ENSG000001 | PASK | 1.62E-01 | 5.04E-01 | 1.09E+00 | 1.28E-01 | 1.09E+00 |
| ENSG000001 | CUL2 | 1.62E-01 | 5.04E-01 | 9.62E-01 | -5.52E-02 | -1.04E+00 |

| | | | | | | |
|---|---|---|---|---|---|---|
| ENSG000001 | SDR16C5 | 1.62E-01 | 5.04E-01 | 1.08E+00 | 1.10E-01 | 1.08E+00 |
| ENSG000001 | PMPCB | 1.63E-01 | 5.04E-01 | 9.47E-01 | -7.92E-02 | -1.06E+00 |
| ENSG000001 | FOXO3 | 1.63E-01 | 5.04E-01 | 1.05E+00 | 7.70E-02 | 1.05E+00 |
| ENSG000001 | C19orf54 | 1.63E-01 | 5.04E-01 | 9.23E-01 | -1.16E-01 | -1.08E+00 |
| ENSG000001 | CRIP2 | 1.63E-01 | 5.04E-01 | 1.11E+00 | 1.56E-01 | 1.11E+00 |
| ENSG000001 | RMDN3 | 1.63E-01 | 5.04E-01 | 1.07E+00 | 1.02E-01 | 1.07E+00 |
| ENSG000001 | CPSF7 | 1.63E-01 | 5.04E-01 | 9.61E-01 | -5.74E-02 | -1.04E+00 |
| ENSG000001 | BCL10 | 1.63E-01 | 5.04E-01 | 1.07E+00 | 9.24E-02 | 1.07E+00 |
| ENSG000002 | --- | 1.63E-01 | ? | 1.37E+00 | 4.52E-01 | 1.37E+00 |
| ENSG000001 | CEP192 | 1.63E-01 | 5.04E-01 | 1.06E+00 | 7.86E-02 | 1.06E+00 |
| ENSG000002 | COX19 | 1.63E-01 | 5.04E-01 | 1.05E+00 | 6.90E-02 | 1.05E+00 |
| ENSG000002 | --- | 1.63E-01 | ? | 1.35E+00 | 4.36E-01 | 1.35E+00 |
| ENSG000001 | FLAD1 | 1.63E-01 | 5.04E-01 | 1.07E+00 | 9.10E-02 | 1.07E+00 |
| ENSG00000C | ARHGAP10 | 1.63E-01 | 5.04E-01 | 1.08E+00 | 1.07E-01 | 1.08E+00 |
| ENSG000001 | MID1 | 1.63E-01 | 5.04E-01 | 1.06E+00 | 8.03E-02 | 1.06E+00 |
| ENSG000001 | RAB25 | 1.63E-01 | 5.04E-01 | 1.07E+00 | 1.03E-01 | 1.07E+00 |
| ENSG000002 | RPL5P30 | 1.63E-01 | ? | 6.89E-01 | -5.38E-01 | -1.45E+00 |
| ENSG000001 | BTN2A3P | 1.63E-01 | ? | 7.71E-01 | -3.75E-01 | -1.30E+00 |
| ENSG000001 | DEPP1 | 1.63E-01 | 5.04E-01 | 8.12E-01 | -3.00E-01 | -1.23E+00 |
| ENSG000001 | BRMS1 | 1.63E-01 | 5.04E-01 | 1.05E+00 | 7.04E-02 | 1.05E+00 |
| ENSG00000C | GINM1 | 1.63E-01 | 5.04E-01 | 1.04E+00 | 5.77E-02 | 1.04E+00 |
| ENSG000001 | BRWD3 | 1.63E-01 | 5.04E-01 | 9.28E-01 | -1.07E-01 | -1.08E+00 |
| ENSG000001 | EEF1B2 | 1.63E-01 | 5.04E-01 | 1.05E+00 | 7.25E-02 | 1.05E+00 |
| ENSG000002 | EIF1AXP1 | 1.63E-01 | 5.04E-01 | 1.21E+00 | 2.70E-01 | 1.21E+00 |
| ENSG000001 | TOR3A | 1.63E-01 | 5.04E-01 | 1.10E+00 | 1.35E-01 | 1.10E+00 |
| ENSG000001 | SSU72 | 1.63E-01 | 5.04E-01 | 9.60E-01 | -5.95E-02 | -1.04E+00 |
| ENSG000001 | RAB4B | 1.63E-01 | 5.04E-01 | 1.14E+00 | 1.90E-01 | 1.14E+00 |
| ENSG000001 | TGFBR2 | 1.63E-01 | 5.04E-01 | 9.48E-01 | -7.76E-02 | -1.06E+00 |
| ENSG000002 | STAG3L5P-P\ | 1.63E-01 | 5.04E-01 | 1.13E+00 | 1.79E-01 | 1.13E+00 |
| ENSG000002 | GP1BB | 1.63E-01 | ? | 1.68E+00 | 7.52E-01 | 1.68E+00 |
| ENSG000001 | NUDT2 | 1.63E-01 | 5.04E-01 | 1.08E+00 | 1.12E-01 | 1.08E+00 |
| ENSG000001 | COL4A2 | 1.63E-01 | 5.04E-01 | 1.09E+00 | 1.29E-01 | 1.09E+00 |
| ENSG000002 | GAPDHP62 | 1.63E-01 | ? | 2.09E+00 | 1.06E+00 | 2.09E+00 |
| ENSG000001 | ZFYVE1 | 1.63E-01 | 5.04E-01 | 1.08E+00 | 1.14E-01 | 1.08E+00 |
| ENSG000002 | LINC01703 | 1.63E-01 | ? | 2.23E+00 | 1.16E+00 | 2.23E+00 |
| ENSG000002 | ZBTB45P2 | 1.63E-01 | 5.04E-01 | 1.15E+00 | 2.07E-01 | 1.15E+00 |
| ENSG000001 | TNFSF11 | 1.63E-01 | ? | 1.80E+00 | 8.46E-01 | 1.80E+00 |
| ENSG000002 | --- | 1.64E-01 | ? | 1.82E+00 | 8.61E-01 | 1.82E+00 |
| ENSG000001 | PTBP2 | 1.64E-01 | 5.04E-01 | 8.88E-01 | -1.71E-01 | -1.13E+00 |
| ENSG000001 | PPM1K | 1.64E-01 | 5.04E-01 | 9.30E-01 | -1.04E-01 | -1.07E+00 |
| ENSG000002 | LINC01410 | 1.64E-01 | 5.04E-01 | 8.58E-01 | -2.21E-01 | -1.17E+00 |
| ENSG000001 | ARID3B | 1.64E-01 | 5.04E-01 | 1.07E+00 | 1.01E-01 | 1.07E+00 |
| ENSG000001 | MAD2L2 | 1.64E-01 | 5.04E-01 | 1.10E+00 | 1.38E-01 | 1.10E+00 |
| ENSG000001 | CRTC2 | 1.64E-01 | 5.04E-01 | 9.47E-01 | -7.82E-02 | -1.06E+00 |
| ENSG000001 | DYRK1B | 1.64E-01 | 5.04E-01 | 1.08E+00 | 1.12E-01 | 1.08E+00 |

| | | | | | | |
|---|---|---|---|---|---|---|
| ENSG000001 | CTBP2 | 1.64E-01 | 5.04E-01 | 9.63E-01 | -5.38E-02 | -1.04E+00 |
| ENSG000001 | WDR31 | 1.64E-01 | 5.04E-01 | 1.13E+00 | 1.75E-01 | 1.13E+00 |
| ENSG00000C | ABCF2 | 1.64E-01 | 5.04E-01 | 1.06E+00 | 8.57E-02 | 1.06E+00 |
| ENSG000001 | TAB3 | 1.64E-01 | 5.04E-01 | 9.40E-01 | -8.98E-02 | -1.06E+00 |
| ENSG000001 | CRYAB | 1.64E-01 | 5.05E-01 | 1.22E+00 | 2.93E-01 | 1.22E+00 |
| ENSG000001 | WBP1L | 1.64E-01 | 5.05E-01 | 1.04E+00 | 5.73E-02 | 1.04E+00 |
| ENSG000001 | DCUN1D2 | 1.64E-01 | 5.05E-01 | 9.06E-01 | -1.43E-01 | -1.10E+00 |
| ENSG000001 | MST1P2 | 1.64E-01 | ? | 1.39E+00 | 4.79E-01 | 1.39E+00 |
| ENSG000001 | PPFIBP2 | 1.64E-01 | 5.05E-01 | 1.11E+00 | 1.45E-01 | 1.11E+00 |
| ENSG000001 | CEP104 | 1.64E-01 | 5.05E-01 | 9.51E-01 | -7.23E-02 | -1.05E+00 |
| ENSG000002 | --- | 1.64E-01 | ? | 1.55E+00 | 6.28E-01 | 1.55E+00 |
| ENSG000001 | TDRD7 | 1.64E-01 | 5.05E-01 | 1.06E+00 | 8.67E-02 | 1.06E+00 |
| ENSG00000C | RAB21 | 1.64E-01 | 5.05E-01 | 9.46E-01 | -7.98E-02 | -1.06E+00 |
| ENSG000002 | --- | 1.64E-01 | ? | 5.83E-01 | -7.79E-01 | -1.72E+00 |
| ENSG00000C | GLT8D1 | 1.64E-01 | 5.05E-01 | 1.04E+00 | 6.18E-02 | 1.04E+00 |
| ENSG000001 | RRAS | 1.64E-01 | 5.05E-01 | 1.06E+00 | 9.07E-02 | 1.06E+00 |
| ENSG000002 | PI4KA | 1.65E-01 | 5.05E-01 | 1.05E+00 | 6.59E-02 | 1.05E+00 |
| ENSG000001 | COX8A | 1.65E-01 | 5.05E-01 | 1.13E+00 | 1.78E-01 | 1.13E+00 |
| ENSG000002 | PEF1-AS1 | 1.65E-01 | ? | 7.09E-01 | -4.95E-01 | -1.41E+00 |
| ENSG000001 | H2AZ2 | 1.65E-01 | 5.05E-01 | 9.63E-01 | -5.49E-02 | -1.04E+00 |
| ENSG00000C | TBXA2R | 1.65E-01 | ? | 6.66E-01 | -5.87E-01 | -1.50E+00 |
| ENSG000001 | SERINC3 | 1.65E-01 | 5.06E-01 | 9.60E-01 | -5.95E-02 | -1.04E+00 |
| ENSG000001 | FASTKD2 | 1.65E-01 | 5.06E-01 | 1.04E+00 | 6.15E-02 | 1.04E+00 |
| ENSG000002 | EIF6 | 1.65E-01 | 5.06E-01 | 1.07E+00 | 9.91E-02 | 1.07E+00 |
| ENSG000001 | AAGAB | 1.65E-01 | 5.06E-01 | 1.05E+00 | 6.42E-02 | 1.05E+00 |
| ENSG00000C | TXLNA | 1.65E-01 | 5.06E-01 | 9.69E-01 | -4.52E-02 | -1.03E+00 |
| ENSG000001 | KRTCAP3 | 1.65E-01 | 5.06E-01 | 1.12E+00 | 1.65E-01 | 1.12E+00 |
| ENSG000002 | --- | 1.65E-01 | 5.06E-01 | 8.86E-01 | -1.75E-01 | -1.13E+00 |
| ENSG000002 | MIR497HG | 1.65E-01 | ? | 6.34E-01 | -6.57E-01 | -1.58E+00 |
| ENSG000001 | ERG | 1.65E-01 | 5.06E-01 | 8.31E-01 | -2.66E-01 | -1.20E+00 |
| ENSG000001 | SREBF2-AS1 | 1.65E-01 | ? | 5.65E-01 | -8.24E-01 | -1.77E+00 |
| ENSG000001 | WDR6 | 1.65E-01 | 5.07E-01 | 1.07E+00 | 1.02E-01 | 1.07E+00 |
| ENSG000001 | IL2RG | 1.65E-01 | ? | 5.39E-01 | -8.91E-01 | -1.85E+00 |
| ENSG000001 | WASH6P | 1.66E-01 | 5.07E-01 | 9.08E-01 | -1.39E-01 | -1.10E+00 |
| ENSG000002 | --- | 1.66E-01 | ? | 7.32E-01 | -4.50E-01 | -1.37E+00 |
| ENSG000001 | TTC26 | 1.66E-01 | 5.07E-01 | 1.08E+00 | 1.16E-01 | 1.08E+00 |
| ENSG000002 | --- | 1.66E-01 | ? | 7.80E-01 | -3.59E-01 | -1.28E+00 |
| ENSG000001 | TSPYL4 | 1.66E-01 | 5.07E-01 | 1.07E+00 | 9.50E-02 | 1.07E+00 |
| ENSG000001 | EGLN1 | 1.66E-01 | 5.07E-01 | 1.06E+00 | 7.76E-02 | 1.06E+00 |
| ENSG000001 | DYNLRB1 | 1.66E-01 | 5.07E-01 | 9.18E-01 | -1.24E-01 | -1.09E+00 |
| ENSG000001 | ISYNA1 | 1.66E-01 | 5.07E-01 | 1.11E+00 | 1.55E-01 | 1.11E+00 |
| ENSG000001 | CDCA2 | 1.66E-01 | 5.07E-01 | 8.71E-01 | -1.99E-01 | -1.15E+00 |
| ENSG000002 | --- | 1.66E-01 | ? | 1.94E+00 | 9.59E-01 | 1.94E+00 |
| ENSG000002 | PRR7-AS1 | 1.66E-01 | ? | 7.74E-01 | -3.70E-01 | -1.29E+00 |
| ENSG000001 | ZRANB2 | 1.66E-01 | 5.07E-01 | 9.42E-01 | -8.67E-02 | -1.06E+00 |

| | | | | | | |
|---|---|---|---|---|---|---|
| ENSG000001 | UBQLN4 | 1.66E-01 | 5.08E-01 | 1.04E+00 | 5.52E-02 | 1.04E+00 |
| ENSG000001 | BDP1 | 1.66E-01 | 5.08E-01 | 9.19E-01 | -1.22E-01 | -1.09E+00 |
| ENSG000001 | ATP6V0C | 1.66E-01 | 5.08E-01 | 9.17E-01 | -1.25E-01 | -1.09E+00 |
| ENSG000001 | RORB | 1.66E-01 | ? | 5.81E-01 | -7.84E-01 | -1.72E+00 |
| ENSG000001 | NHS | 1.66E-01 | 5.08E-01 | 9.22E-01 | -1.18E-01 | -1.08E+00 |
| ENSG000001 | NUTM2B | 1.67E-01 | ? | 1.39E+00 | 4.70E-01 | 1.39E+00 |
| ENSG000002 | SNHG32 | 1.67E-01 | 5.09E-01 | 1.12E+00 | 1.65E-01 | 1.12E+00 |
| ENSG000002 | --- | 1.67E-01 | 5.09E-01 | 7.74E-01 | -3.70E-01 | -1.29E+00 |
| ENSG000001 | EIF3A | 1.67E-01 | 5.09E-01 | 9.49E-01 | -7.56E-02 | -1.05E+00 |
| ENSG000001 | --- | 1.67E-01 | 5.09E-01 | 7.41E-01 | -4.33E-01 | -1.35E+00 |
| ENSG000001 | CAPN7 | 1.67E-01 | 5.09E-01 | 9.57E-01 | -6.28E-02 | -1.04E+00 |
| ENSG000001 | CBX7 | 1.67E-01 | 5.10E-01 | 1.26E+00 | 3.31E-01 | 1.26E+00 |
| ENSG000002 | KIFC1 | 1.67E-01 | 5.10E-01 | 8.43E-01 | -2.46E-01 | -1.19E+00 |
| ENSG000001 | ARF4 | 1.67E-01 | 5.10E-01 | 9.74E-01 | -3.87E-02 | -1.03E+00 |
| ENSG000002 | NPIPA7 | 1.67E-01 | 5.10E-01 | 5.61E-01 | -8.33E-01 | -1.78E+00 |
| ENSG000001 | RAPSN | 1.67E-01 | ? | 6.65E-01 | -5.88E-01 | -1.50E+00 |
| ENSG000002 | STK19 | 1.68E-01 | 5.11E-01 | 1.09E+00 | 1.27E-01 | 1.09E+00 |
| ENSG000001 | GPR160 | 1.68E-01 | 5.11E-01 | 8.11E-01 | -3.02E-01 | -1.23E+00 |
| ENSG000002 | --- | 1.68E-01 | ? | 1.42E+00 | 5.01E-01 | 1.42E+00 |
| ENSG000001 | FAM161A | 1.68E-01 | 5.11E-01 | 8.76E-01 | -1.91E-01 | -1.14E+00 |
| ENSG000002 | CTAGE4 | 1.68E-01 | 5.11E-01 | 8.67E-01 | -2.06E-01 | -1.15E+00 |
| ENSG000001 | DSTN | 1.68E-01 | 5.11E-01 | 1.03E+00 | 4.03E-02 | 1.03E+00 |
| ENSG000001 | ZNF785 | 1.68E-01 | 5.11E-01 | 1.08E+00 | 1.15E-01 | 1.08E+00 |
| ENSG000001 | RCN2 | 1.68E-01 | 5.11E-01 | 9.47E-01 | -7.80E-02 | -1.06E+00 |
| ENSG000002 | --- | 1.68E-01 | ? | 1.95E+00 | 9.61E-01 | 1.95E+00 |
| ENSG000001 | TRIM52 | 1.68E-01 | 5.11E-01 | 9.09E-01 | -1.38E-01 | -1.10E+00 |
| ENSG000002 | --- | 1.68E-01 | 5.11E-01 | 1.35E+00 | 4.29E-01 | 1.35E+00 |
| ENSG000001 | FAM83C | 1.68E-01 | ? | 1.56E+00 | 6.40E-01 | 1.56E+00 |
| ENSG000001 | SCNN1A | 1.68E-01 | 5.11E-01 | 8.19E-01 | -2.88E-01 | -1.22E+00 |
| ENSG000001 | ATE1 | 1.68E-01 | 5.11E-01 | 9.43E-01 | -8.53E-02 | -1.06E+00 |
| ENSG000002 | --- | 1.68E-01 | 5.11E-01 | 7.79E-01 | -3.60E-01 | -1.28E+00 |
| ENSG000001 | KBTBD3 | 1.68E-01 | 5.11E-01 | 1.12E+00 | 1.70E-01 | 1.12E+00 |
| ENSG000002 | --- | 1.68E-01 | 5.11E-01 | 8.86E-01 | -1.75E-01 | -1.13E+00 |
| ENSG000001 | SPAG1 | 1.68E-01 | 5.11E-01 | 9.13E-01 | -1.31E-01 | -1.09E+00 |
| ENSG000002 | KPRP | 1.68E-01 | ? | 6.00E-01 | -7.36E-01 | -1.67E+00 |
| ENSG000001 | TMED9 | 1.68E-01 | 5.11E-01 | 1.08E+00 | 1.16E-01 | 1.08E+00 |
| ENSG000001 | PRXL2A | 1.68E-01 | 5.12E-01 | 1.05E+00 | 7.31E-02 | 1.05E+00 |
| ENSG000001 | DNAJC28 | 1.69E-01 | 5.12E-01 | 7.77E-01 | -3.63E-01 | -1.29E+00 |
| ENSG000001 | TMEM183A | 1.69E-01 | 5.12E-01 | 1.03E+00 | 4.72E-02 | 1.03E+00 |
| ENSG000001 | CDIN1 | 1.69E-01 | 5.12E-01 | 9.34E-01 | -9.90E-02 | -1.07E+00 |
| ENSG000002 | --- | 1.69E-01 | ? | 1.56E+00 | 6.42E-01 | 1.56E+00 |
| ENSG000002 | HSD17B1-AS | 1.69E-01 | 5.12E-01 | 1.08E+00 | 1.06E-01 | 1.08E+00 |
| ENSG000001 | EML3 | 1.69E-01 | 5.12E-01 | 1.11E+00 | 1.55E-01 | 1.11E+00 |
| ENSG000002 | ZBTB11-AS1 | 1.69E-01 | 5.12E-01 | 1.14E+00 | 1.92E-01 | 1.14E+00 |
| ENSG000001 | CBLC | 1.69E-01 | 5.13E-01 | 1.08E+00 | 1.14E-01 | 1.08E+00 |

| | | | | | | |
|---|---|---|---|---|---|---|
| ENSG000002 | LINC02273 | 1.69E-01 | 5.13E-01 | 8.28E-01 | -2.73E-01 | -1.21E+00 |
| ENSG000001 | RUBCN | 1.69E-01 | 5.13E-01 | 9.49E-01 | -7.57E-02 | -1.05E+00 |
| ENSG000001 | --- | 1.69E-01 | ? | 1.44E+00 | 5.22E-01 | 1.44E+00 |
| ENSG000002 | --- | 1.69E-01 | ? | 7.34E-01 | -4.47E-01 | -1.36E+00 |
| ENSG000001 | ACADL | 1.69E-01 | ? | 1.26E+00 | 3.32E-01 | 1.26E+00 |
| ENSG000002 | --- | 1.69E-01 | ? | 7.54E-01 | -4.07E-01 | -1.33E+00 |
| ENSG000002 | --- | 1.69E-01 | ? | 5.63E-01 | -8.29E-01 | -1.78E+00 |
| ENSG000001 | ERGIC3 | 1.70E-01 | 5.14E-01 | 1.05E+00 | 7.02E-02 | 1.05E+00 |
| ENSG000002 | --- | 1.70E-01 | ? | 4.68E-01 | -1.10E+00 | -2.14E+00 |
| ENSG000002 | YWHAZP4 | 1.70E-01 | ? | 6.31E-01 | -6.65E-01 | -1.59E+00 |
| ENSG000001 | ZMYND8 | 1.70E-01 | 5.14E-01 | 9.29E-01 | -1.06E-01 | -1.08E+00 |
| ENSG000001 | SWSAP1 | 1.70E-01 | 5.14E-01 | 1.13E+00 | 1.76E-01 | 1.13E+00 |
| ENSG000001 | SUSD5 | 1.70E-01 | 5.14E-01 | 8.62E-01 | -2.14E-01 | -1.16E+00 |
| ENSG000001 | HELZ | 1.70E-01 | 5.14E-01 | 1.08E+00 | 1.13E-01 | 1.08E+00 |
| ENSG000001 | ATP6V1C1 | 1.70E-01 | 5.14E-01 | 9.54E-01 | -6.86E-02 | -1.05E+00 |
| ENSG000001 | ARHGEF37 | 1.70E-01 | 5.14E-01 | 1.10E+00 | 1.33E-01 | 1.10E+00 |
| ENSG000001 | MRPL51 | 1.70E-01 | 5.15E-01 | 1.05E+00 | 6.63E-02 | 1.05E+00 |
| ENSG000002 | ASB16-AS1 | 1.70E-01 | 5.15E-01 | 1.11E+00 | 1.51E-01 | 1.11E+00 |
| ENSG000002 | TLE1P1 | 1.70E-01 | ? | 1.45E+00 | 5.37E-01 | 1.45E+00 |
| ENSG000002 | --- | 1.70E-01 | 5.15E-01 | 1.23E+00 | 2.94E-01 | 1.23E+00 |
| ENSG000002 | --- | 1.70E-01 | ? | 1.25E+00 | 3.22E-01 | 1.25E+00 |
| ENSG000000 | ANGPT2 | 1.70E-01 | ? | 1.63E+00 | 7.00E-01 | 1.63E+00 |
| ENSG000002 | HMGN2P5 | 1.70E-01 | 5.15E-01 | 8.63E-01 | -2.12E-01 | -1.16E+00 |
| ENSG000001 | CLTC | 1.70E-01 | 5.15E-01 | 9.71E-01 | -4.29E-02 | -1.03E+00 |
| ENSG000002 | NOC2LP1 | 1.70E-01 | ? | 1.30E+00 | 3.75E-01 | 1.30E+00 |
| ENSG000000 | BAIAP3 | 1.70E-01 | ? | 6.34E-01 | -6.58E-01 | -1.58E+00 |
| ENSG000001 | OSER1 | 1.70E-01 | 5.15E-01 | 1.06E+00 | 7.93E-02 | 1.06E+00 |
| ENSG000002 | WDR5B-DT | 1.70E-01 | 5.15E-01 | 1.16E+00 | 2.12E-01 | 1.16E+00 |
| ENSG000001 | SLC25A16 | 1.70E-01 | 5.15E-01 | 9.48E-01 | -7.70E-02 | -1.05E+00 |
| ENSG000002 | SCARF2 | 1.70E-01 | ? | 1.36E+00 | 4.48E-01 | 1.36E+00 |
| ENSG000001 | FBXL13 | 1.70E-01 | ? | 7.25E-01 | -4.63E-01 | -1.38E+00 |
| ENSG000002 | MYCBP | 1.71E-01 | 5.15E-01 | 1.05E+00 | 7.68E-02 | 1.05E+00 |
| ENSG000002 | --- | 1.71E-01 | 5.15E-01 | 9.49E-01 | -7.61E-02 | -1.05E+00 |
| ENSG000001 | GRPEL2 | 1.71E-01 | 5.15E-01 | 9.46E-01 | -8.02E-02 | -1.06E+00 |
| ENSG000001 | ZNF320 | 1.71E-01 | 5.16E-01 | 9.31E-01 | -1.03E-01 | -1.07E+00 |
| ENSG000001 | MT-ND2 | 1.71E-01 | 5.16E-01 | 8.73E-01 | -1.96E-01 | -1.15E+00 |
| ENSG000001 | BTBD8 | 1.71E-01 | 5.16E-01 | 9.28E-01 | -1.08E-01 | -1.08E+00 |
| ENSG000001 | CD276 | 1.71E-01 | 5.16E-01 | 1.05E+00 | 6.41E-02 | 1.05E+00 |
| ENSG000001 | VPS18 | 1.71E-01 | 5.16E-01 | 1.06E+00 | 8.62E-02 | 1.06E+00 |
| ENSG000002 | SAMD13 | 1.71E-01 | ? | 1.37E+00 | 4.55E-01 | 1.37E+00 |
| ENSG000001 | DNAJC2 | 1.71E-01 | 5.16E-01 | 9.27E-01 | -1.09E-01 | -1.08E+00 |
| ENSG000001 | HCST | 1.71E-01 | ? | 4.61E-01 | -1.12E+00 | -2.17E+00 |
| ENSG000002 | --- | 1.71E-01 | 5.16E-01 | 8.14E-01 | -2.97E-01 | -1.23E+00 |
| ENSG000002 | LINC00519 | 1.71E-01 | 5.16E-01 | 8.70E-01 | -2.02E-01 | -1.15E+00 |
| ENSG000002 | --- | 1.71E-01 | 5.16E-01 | 8.78E-01 | -1.88E-01 | -1.14E+00 |

| | | | | | | |
|---|---|---|---|---|---|---|
| ENSG000001 | MYO1F | 1.71E-01 | ? | 1.43E+00 | 5.18E-01 | 1.43E+00 |
| ENSG000001 | RASSF10 | 1.71E-01 | 5.16E-01 | 1.13E+00 | 1.70E-01 | 1.13E+00 |
| ENSG000001 | SDSL | 1.71E-01 | 5.16E-01 | 1.17E+00 | 2.21E-01 | 1.17E+00 |
| ENSG000001 | KIF17 | 1.71E-01 | 5.16E-01 | 8.44E-01 | -2.45E-01 | -1.18E+00 |
| ENSG000002 | --- | 1.71E-01 | 5.16E-01 | 1.26E+00 | 3.32E-01 | 1.26E+00 |
| ENSG00000C | ATP2C1 | 1.71E-01 | 5.16E-01 | 1.03E+00 | 4.17E-02 | 1.03E+00 |
| ENSG000002 | --- | 1.72E-01 | ? | 2.09E+00 | 1.07E+00 | 2.09E+00 |
| ENSG000001 | PEG3 | 1.72E-01 | 5.16E-01 | 1.13E+00 | 1.82E-01 | 1.13E+00 |
| ENSG000001 | ASF1A | 1.72E-01 | 5.16E-01 | 9.50E-01 | -7.42E-02 | -1.05E+00 |
| ENSG000002 | --- | 1.72E-01 | 5.16E-01 | 8.77E-01 | -1.89E-01 | -1.14E+00 |
| ENSG000001 | SETD9 | 1.72E-01 | 5.16E-01 | 1.16E+00 | 2.09E-01 | 1.16E+00 |
| ENSG000001 | DNAJC18 | 1.72E-01 | 5.16E-01 | 1.07E+00 | 9.85E-02 | 1.07E+00 |
| ENSG000001 | SNX25 | 1.72E-01 | 5.16E-01 | 9.46E-01 | -7.97E-02 | -1.06E+00 |
| ENSG000001 | PEAR1 | 1.72E-01 | 5.16E-01 | 9.43E-01 | -8.54E-02 | -1.06E+00 |
| ENSG000001 | RBM38 | 1.72E-01 | 5.16E-01 | 1.10E+00 | 1.43E-01 | 1.10E+00 |
| ENSG00000C | DHPS | 1.72E-01 | 5.16E-01 | 1.08E+00 | 1.06E-01 | 1.08E+00 |
| ENSG000002 | SACM1L | 1.72E-01 | 5.16E-01 | 9.59E-01 | -5.98E-02 | -1.04E+00 |
| ENSG000001 | H3-3A | 1.72E-01 | 5.16E-01 | 1.05E+00 | 7.15E-02 | 1.05E+00 |
| ENSG000001 | ZNF189 | 1.72E-01 | 5.16E-01 | 9.45E-01 | -8.21E-02 | -1.06E+00 |
| ENSG000001 | STUB1 | 1.72E-01 | 5.17E-01 | 1.07E+00 | 9.96E-02 | 1.07E+00 |
| ENSG000001 | DENND1A | 1.72E-01 | 5.17E-01 | 1.06E+00 | 9.02E-02 | 1.06E+00 |
| ENSG000001 | SV2A | 1.72E-01 | ? | 1.39E+00 | 4.80E-01 | 1.39E+00 |
| ENSG000001 | ZNF473 | 1.72E-01 | 5.17E-01 | 1.06E+00 | 8.73E-02 | 1.06E+00 |
| ENSG000001 | HHEX | 1.72E-01 | ? | 1.62E+00 | 7.00E-01 | 1.62E+00 |
| ENSG000001 | PLP2 | 1.72E-01 | 5.17E-01 | 9.60E-01 | -5.91E-02 | -1.04E+00 |
| ENSG000001 | MDH2 | 1.72E-01 | 5.17E-01 | 1.05E+00 | 6.80E-02 | 1.05E+00 |
| ENSG000001 | UPF3A | 1.72E-01 | 5.17E-01 | 9.62E-01 | -5.65E-02 | -1.04E+00 |
| ENSG000001 | TOB2 | 1.72E-01 | 5.17E-01 | 1.04E+00 | 5.38E-02 | 1.04E+00 |
| ENSG000001 | LAMTOR3 | 1.72E-01 | 5.17E-01 | 1.04E+00 | 6.11E-02 | 1.04E+00 |
| ENSG000001 | ACTR3C | 1.72E-01 | 5.17E-01 | 7.91E-01 | -3.38E-01 | -1.26E+00 |
| ENSG000001 | PAWR | 1.73E-01 | 5.17E-01 | 9.57E-01 | -6.41E-02 | -1.05E+00 |
| ENSG000001 | MFAP4 | 1.73E-01 | ? | 6.25E-01 | -6.77E-01 | -1.60E+00 |
| ENSG000001 | MBOAT1 | 1.73E-01 | 5.17E-01 | 8.69E-01 | -2.03E-01 | -1.15E+00 |
| ENSG000001 | TCERG1 | 1.73E-01 | 5.17E-01 | 9.42E-01 | -8.56E-02 | -1.06E+00 |
| ENSG000001 | SOCS2 | 1.73E-01 | 5.17E-01 | 1.13E+00 | 1.71E-01 | 1.13E+00 |
| ENSG000001 | TCEA2 | 1.73E-01 | 5.17E-01 | 1.10E+00 | 1.42E-01 | 1.10E+00 |
| ENSG000002 | --- | 1.73E-01 | ? | 7.62E-01 | -3.92E-01 | -1.31E+00 |
| ENSG00000C | GSTP1 | 1.73E-01 | 5.17E-01 | 1.06E+00 | 8.42E-02 | 1.06E+00 |
| ENSG00000C | PABPC4 | 1.73E-01 | 5.17E-01 | 1.04E+00 | 5.07E-02 | 1.04E+00 |
| ENSG000001 | RTN4RL1 | 1.73E-01 | 5.17E-01 | 1.14E+00 | 1.87E-01 | 1.14E+00 |
| ENSG00000C | OGFR | 1.73E-01 | 5.17E-01 | 1.08E+00 | 1.17E-01 | 1.08E+00 |
| ENSG000001 | MRPL4 | 1.73E-01 | 5.18E-01 | 1.06E+00 | 8.58E-02 | 1.06E+00 |
| ENSG000001 | IL27RA | 1.73E-01 | 5.18E-01 | 1.10E+00 | 1.39E-01 | 1.10E+00 |
| ENSG000002 | --- | 1.73E-01 | ? | 1.39E+00 | 4.76E-01 | 1.39E+00 |
| ENSG00000C | ATOSB | 1.73E-01 | 5.18E-01 | 9.58E-01 | -6.21E-02 | -1.04E+00 |

| | | | | | | | |
|---|---|---|---|---|---|---|---|
| ENSG000001 | PAIP2B | 1.73E-01 | | 5.18E-01 | 1.09E+00 | 1.23E-01 | 1.09E+00 |
| ENSG000000 | RAPGEF3 | 1.73E-01 | | 5.18E-01 | 8.81E-01 | -1.83E-01 | -1.14E+00 |
| ENSG000001 | RNF6 | 1.73E-01 | | 5.18E-01 | 9.52E-01 | -7.09E-02 | -1.05E+00 |
| ENSG000002 | --- | 1.73E-01 | | 5.18E-01 | 7.41E-01 | -4.33E-01 | -1.35E+00 |
| ENSG000002 | LINC02827 | 1.74E-01 | ? | | 4.76E-01 | -1.07E+00 | -2.10E+00 |
| ENSG000001 | TMEM62 | 1.74E-01 | | 5.18E-01 | 1.07E+00 | 9.82E-02 | 1.07E+00 |
| ENSG000002 | NR2F1-AS1 | 1.74E-01 | | 5.19E-01 | 8.47E-01 | -2.40E-01 | -1.18E+00 |
| ENSG000002 | --- | 1.74E-01 | ? | | 1.56E+00 | 6.39E-01 | 1.56E+00 |
| ENSG000000 | ANLN | 1.74E-01 | | 5.19E-01 | 8.80E-01 | -1.84E-01 | -1.14E+00 |
| ENSG000002 | RBMS1P1 | 1.74E-01 | ? | | 7.08E-01 | -4.97E-01 | -1.41E+00 |
| ENSG000001 | MPHOSPH10 | 1.74E-01 | | 5.19E-01 | 9.35E-01 | -9.76E-02 | -1.07E+00 |
| ENSG000001 | CNIH3 | 1.74E-01 | ? | | 7.72E-01 | -3.73E-01 | -1.29E+00 |
| ENSG000001 | CRTAP | 1.74E-01 | | 5.20E-01 | 1.05E+00 | 6.67E-02 | 1.05E+00 |
| ENSG000002 | --- | 1.74E-01 | | 5.20E-01 | 8.47E-01 | -2.40E-01 | -1.18E+00 |
| ENSG000001 | ZSCAN23 | 1.74E-01 | | 5.20E-01 | 8.42E-01 | -2.48E-01 | -1.19E+00 |
| ENSG000001 | RELL2 | 1.74E-01 | | 5.20E-01 | 1.09E+00 | 1.27E-01 | 1.09E+00 |
| ENSG000001 | CEP55 | 1.74E-01 | | 5.20E-01 | 8.60E-01 | -2.18E-01 | -1.16E+00 |
| ENSG000002 | PRKCH-AS1 | 1.74E-01 | | 5.20E-01 | 8.59E-01 | -2.19E-01 | -1.16E+00 |
| ENSG000001 | HSPA4 | 1.74E-01 | | 5.20E-01 | 9.65E-01 | -5.09E-02 | -1.04E+00 |
| ENSG000002 | OLMALINC | 1.74E-01 | | 5.20E-01 | 9.02E-01 | -1.48E-01 | -1.11E+00 |
| ENSG000002 | ITGB8-AS1 | 1.74E-01 | ? | | 1.73E+00 | 7.94E-01 | 1.73E+00 |
| ENSG000001 | DGUOK | 1.74E-01 | | 5.20E-01 | 1.06E+00 | 8.78E-02 | 1.06E+00 |
| ENSG000001 | FOXP1 | 1.75E-01 | | 5.20E-01 | 9.45E-01 | -8.14E-02 | -1.06E+00 |
| ENSG000001 | MAIP1 | 1.75E-01 | | 5.20E-01 | 1.09E+00 | 1.24E-01 | 1.09E+00 |
| ENSG000000 | UFD1 | 1.75E-01 | | 5.20E-01 | 9.60E-01 | -5.84E-02 | -1.04E+00 |
| ENSG000001 | ZNF391 | 1.75E-01 | | 5.20E-01 | 9.19E-01 | -1.22E-01 | -1.09E+00 |
| ENSG000001 | SERPINB2 | 1.75E-01 | | 5.20E-01 | 8.06E-01 | -3.12E-01 | -1.24E+00 |
| ENSG000001 | PLXNB1 | 1.75E-01 | | 5.20E-01 | 1.08E+00 | 1.16E-01 | 1.08E+00 |
| ENSG000002 | LINC00216 | 1.75E-01 | ? | | 6.43E-01 | -6.38E-01 | -1.56E+00 |
| ENSG000000 | SEMA5B | 1.75E-01 | ? | | 1.37E+00 | 4.59E-01 | 1.37E+00 |
| ENSG000000 | TMEM101 | 1.75E-01 | | 5.21E-01 | 1.08E+00 | 1.17E-01 | 1.08E+00 |
| ENSG000001 | HPCAL4 | 1.75E-01 | ? | | 1.28E+00 | 3.61E-01 | 1.28E+00 |
| ENSG000000 | KPNA6 | 1.75E-01 | | 5.21E-01 | 1.04E+00 | 5.79E-02 | 1.04E+00 |
| ENSG000001 | ANKRD30B | 1.75E-01 | ? | | 4.90E-01 | -1.03E+00 | -2.04E+00 |
| ENSG000002 | INO80B-WBI | 1.75E-01 | | 5.21E-01 | 1.08E+00 | 1.16E-01 | 1.08E+00 |
| ENSG000000 | NSMAF | 1.75E-01 | | 5.21E-01 | 1.04E+00 | 6.15E-02 | 1.04E+00 |
| ENSG000001 | RTP4 | 1.75E-01 | ? | | 1.28E+00 | 3.61E-01 | 1.28E+00 |
| ENSG000001 | FGFR1OP2 | 1.75E-01 | | 5.21E-01 | 9.34E-01 | -9.81E-02 | -1.07E+00 |
| ENSG000001 | FAM177A1 | 1.76E-01 | | 5.22E-01 | 9.37E-01 | -9.45E-02 | -1.07E+00 |
| ENSG000001 | ZNF561 | 1.76E-01 | | 5.22E-01 | 9.54E-01 | -6.83E-02 | -1.05E+00 |
| ENSG000001 | UGDH | 1.76E-01 | | 5.22E-01 | 9.49E-01 | -7.56E-02 | -1.05E+00 |
| ENSG000001 | CCNI | 1.76E-01 | | 5.22E-01 | 9.63E-01 | -5.38E-02 | -1.04E+00 |
| ENSG000001 | C17orf99 | 1.76E-01 | ? | | 7.16E-01 | -4.82E-01 | -1.40E+00 |
| ENSG000001 | SLC3A1 | 1.76E-01 | ? | | 5.82E-01 | -7.82E-01 | -1.72E+00 |
| ENSG000001 | LGI4 | 1.76E-01 | ? | | 6.33E-01 | -6.60E-01 | -1.58E+00 |

| | | | | | | |
|---|---|---|---|---|---|---|
| ENSG000001 | RAD17 | 1.76E-01 | 5.22E-01 | 9.51E-01 | -7.20E-02 | -1.05E+00 |
| ENSG000001 | MACROD1 | 1.76E-01 | ? | 6.96E-01 | -5.23E-01 | -1.44E+00 |
| ENSG000001 | CD99L2 | 1.76E-01 | 5.23E-01 | 1.04E+00 | 6.30E-02 | 1.04E+00 |
| ENSG000001 | SMDT1 | 1.76E-01 | 5.23E-01 | 1.10E+00 | 1.32E-01 | 1.10E+00 |
| ENSG000001 | AKR1C2 | 1.76E-01 | 5.23E-01 | 8.46E-01 | -2.41E-01 | -1.18E+00 |
| ENSG000001 | ACP5 | 1.76E-01 | 5.23E-01 | 1.10E+00 | 1.42E-01 | 1.10E+00 |
| ENSG000001 | HEATR1 | 1.76E-01 | 5.23E-01 | 9.59E-01 | -6.07E-02 | -1.04E+00 |
| ENSG000001 | HNRNPK | 1.76E-01 | 5.23E-01 | 1.02E+00 | 3.55E-02 | 1.02E+00 |
| ENSG000002 | LINC00630 | 1.76E-01 | 5.23E-01 | 1.11E+00 | 1.55E-01 | 1.11E+00 |
| ENSG000002 | WAC-AS1 | 1.77E-01 | 5.23E-01 | 1.06E+00 | 8.14E-02 | 1.06E+00 |
| ENSG000002 | HLA-E | 1.77E-01 | 5.23E-01 | 1.05E+00 | 7.21E-02 | 1.05E+00 |
| ENSG000002 | KCNU1 | 1.77E-01 | ? | 7.87E-01 | -3.46E-01 | -1.27E+00 |
| ENSG000001 | SELENOW | 1.77E-01 | 5.23E-01 | 1.08E+00 | 1.08E-01 | 1.08E+00 |
| ENSG000002 | --- | 1.77E-01 | ? | 5.83E-01 | -7.79E-01 | -1.72E+00 |
| ENSG000001 | TIMM8A | 1.77E-01 | 5.23E-01 | 1.08E+00 | 1.15E-01 | 1.08E+00 |
| ENSG000002 | --- | 1.77E-01 | 5.23E-01 | 1.14E+00 | 1.83E-01 | 1.14E+00 |
| ENSG000002 | RHEBP1 | 1.77E-01 | ? | 4.26E-01 | -1.23E+00 | -2.35E+00 |
| ENSG000002 | --- | 1.77E-01 | ? | 2.05E+00 | 1.04E+00 | 2.05E+00 |
| ENSG000001 | ABTB1 | 1.77E-01 | 5.24E-01 | 1.12E+00 | 1.59E-01 | 1.12E+00 |
| ENSG000001 | MT2A | 1.77E-01 | 5.24E-01 | 9.45E-01 | -8.21E-02 | -1.06E+00 |
| ENSG000001 | SOX13 | 1.77E-01 | 5.24E-01 | 9.42E-01 | -8.64E-02 | -1.06E+00 |
| ENSG000002 | --- | 1.77E-01 | ? | 1.64E+00 | 7.11E-01 | 1.64E+00 |
| ENSG000001 | AKR1A1 | 1.77E-01 | 5.24E-01 | 1.08E+00 | 1.05E-01 | 1.08E+00 |
| ENSG000002 | GPX3 | 1.77E-01 | 5.24E-01 | 9.32E-01 | -1.02E-01 | -1.07E+00 |
| ENSG00000C | ZNF416 | 1.77E-01 | 5.24E-01 | 1.08E+00 | 1.08E-01 | 1.08E+00 |
| ENSG000002 | --- | 1.77E-01 | ? | 5.84E-01 | -7.76E-01 | -1.71E+00 |
| ENSG000001 | ARMC6 | 1.77E-01 | 5.25E-01 | 1.09E+00 | 1.25E-01 | 1.09E+00 |
| ENSG000002 | RPS12P3 | 1.78E-01 | ? | 1.92E+00 | 9.40E-01 | 1.92E+00 |
| ENSG00000C | SETD1A | 1.78E-01 | 5.25E-01 | 9.46E-01 | -8.01E-02 | -1.06E+00 |
| ENSG000001 | RLF | 1.78E-01 | 5.26E-01 | 1.06E+00 | 8.24E-02 | 1.06E+00 |
| ENSG000001 | C12orf60 | 1.78E-01 | 5.26E-01 | 8.62E-01 | -2.14E-01 | -1.16E+00 |
| ENSG000001 | RAB1A | 1.78E-01 | 5.26E-01 | 1.03E+00 | 4.19E-02 | 1.03E+00 |
| ENSG000001 | EMP1 | 1.78E-01 | 5.26E-01 | 9.54E-01 | -6.78E-02 | -1.05E+00 |
| ENSG000001 | GTPBP6 | 1.78E-01 | 5.26E-01 | 1.07E+00 | 1.00E-01 | 1.07E+00 |
| ENSG000002 | E2F4 | 1.78E-01 | 5.26E-01 | 1.06E+00 | 7.90E-02 | 1.06E+00 |
| ENSG000001 | CCDC170 | 1.78E-01 | ? | 1.50E+00 | 5.88E-01 | 1.50E+00 |
| ENSG000001 | PRR36 | 1.78E-01 | ? | 7.47E-01 | -4.21E-01 | -1.34E+00 |
| ENSG000001 | SNX9 | 1.78E-01 | 5.27E-01 | 9.63E-01 | -5.40E-02 | -1.04E+00 |
| ENSG000002 | --- | 1.79E-01 | ? | 1.37E+00 | 4.58E-01 | 1.37E+00 |
| ENSG000001 | BTN3A2 | 1.79E-01 | 5.27E-01 | 1.08E+00 | 1.12E-01 | 1.08E+00 |
| ENSG000001 | PLIN2 | 1.79E-01 | 5.27E-01 | 1.17E+00 | 2.28E-01 | 1.17E+00 |
| ENSG00000C | TBC1D25 | 1.79E-01 | 5.28E-01 | 1.07E+00 | 9.84E-02 | 1.07E+00 |
| ENSG000002 | RAET1K | 1.79E-01 | ? | 1.54E+00 | 6.23E-01 | 1.54E+00 |
| ENSG000001 | ERVK3-1 | 1.79E-01 | 5.28E-01 | 9.43E-01 | -8.52E-02 | -1.06E+00 |
| ENSG000002 | --- | 1.79E-01 | ? | 1.61E+00 | 6.91E-01 | 1.61E+00 |

| | | | | | | | |
|---|---|---|---|---|---|---|---|
| ENSG000002 | KIAA1614-AS | 1.79E-01 | ? | | 1.31E+00 | 3.87E-01 | 1.31E+00 |
| ENSG000001 | PPP1R14B | 1.79E-01 | | 5.28E-01 | 1.06E+00 | 7.89E-02 | 1.06E+00 |
| ENSG000002 | --- | 1.79E-01 | ? | | 1.67E+00 | 7.36E-01 | 1.67E+00 |
| ENSG000002 | KANSL1-AS1 | 1.79E-01 | ? | | 6.00E-01 | -7.37E-01 | -1.67E+00 |
| ENSG000001 | EIF2S2P4 | 1.79E-01 | | 5.28E-01 | 8.98E-01 | -1.55E-01 | -1.11E+00 |
| ENSG000001 | PDZD2 | 1.79E-01 | | 5.29E-01 | 9.50E-01 | -7.47E-02 | -1.05E+00 |
| ENSG000001 | VPS51 | 1.80E-01 | | 5.29E-01 | 1.11E+00 | 1.48E-01 | 1.11E+00 |
| ENSG000001 | SLC47A1 | 1.80E-01 | ? | | 1.26E+00 | 3.33E-01 | 1.26E+00 |
| ENSG000001 | RANGRF | 1.80E-01 | | 5.29E-01 | 1.12E+00 | 1.62E-01 | 1.12E+00 |
| ENSG000001 | EXO1 | 1.80E-01 | | 5.29E-01 | 8.11E-01 | -3.02E-01 | -1.23E+00 |
| ENSG000001 | PEX5L | 1.80E-01 | ? | | 7.82E-01 | -3.54E-01 | -1.28E+00 |
| ENSG000002 | RPS18P12 | 1.80E-01 | ? | | 1.67E+00 | 7.43E-01 | 1.67E+00 |
| ENSG000002 | GAN | 1.80E-01 | | 5.29E-01 | 1.09E+00 | 1.24E-01 | 1.09E+00 |
| ENSG000002 | SUGT1P4-ST | 1.80E-01 | | 5.29E-01 | 8.36E-01 | -2.59E-01 | -1.20E+00 |
| ENSG000002 | --- | 1.80E-01 | ? | | 1.85E+00 | 8.88E-01 | 1.85E+00 |
| ENSG000002 | NOTCH2NLA | 1.80E-01 | | 5.30E-01 | 1.07E+00 | 1.04E-01 | 1.07E+00 |
| ENSG000001 | LONRF3 | 1.80E-01 | ? | | 7.65E-01 | -3.86E-01 | -1.31E+00 |
| ENSG00000C | LYPLA2 | 1.80E-01 | | 5.30E-01 | 1.07E+00 | 1.02E-01 | 1.07E+00 |
| ENSG000001 | MIEF1 | 1.80E-01 | | 5.30E-01 | 9.58E-01 | -6.15E-02 | -1.04E+00 |
| ENSG000001 | SLC23A1 | 1.80E-01 | ? | | 6.51E-01 | -6.20E-01 | -1.54E+00 |
| ENSG000001 | PPP1R21 | 1.80E-01 | | 5.30E-01 | 9.52E-01 | -7.15E-02 | -1.05E+00 |
| ENSG000001 | ZNF133 | 1.80E-01 | | 5.30E-01 | 1.06E+00 | 8.88E-02 | 1.06E+00 |
| ENSG000002 | --- | 1.80E-01 | ? | | 1.56E+00 | 6.38E-01 | 1.56E+00 |
| ENSG000001 | CCNYL1 | 1.80E-01 | | 5.30E-01 | 9.50E-01 | -7.39E-02 | -1.05E+00 |
| ENSG000002 | PRR3 | 1.80E-01 | | 5.30E-01 | 8.91E-01 | -1.66E-01 | -1.12E+00 |
| ENSG000001 | GPR146 | 1.80E-01 | ? | | 1.81E+00 | 8.57E-01 | 1.81E+00 |
| ENSG000001 | NKRF | 1.80E-01 | | 5.30E-01 | 1.06E+00 | 8.09E-02 | 1.06E+00 |
| ENSG000001 | RAD21 | 1.80E-01 | | 5.30E-01 | 9.66E-01 | -5.03E-02 | -1.04E+00 |
| ENSG000001 | DDX41 | 1.80E-01 | | 5.30E-01 | 1.04E+00 | 5.98E-02 | 1.04E+00 |
| ENSG000002 | APCDD1L-DT | 1.80E-01 | ? | | 1.42E+00 | 5.04E-01 | 1.42E+00 |
| ENSG000002 | --- | 1.80E-01 | ? | | 1.37E+00 | 4.52E-01 | 1.37E+00 |
| ENSG000001 | DHDH | 1.80E-01 | ? | | 1.52E+00 | 6.04E-01 | 1.52E+00 |
| ENSG000002 | --- | 1.80E-01 | ? | | 5.84E-01 | -7.75E-01 | -1.71E+00 |
| ENSG000002 | RIMBP3 | 1.80E-01 | ? | | 5.40E-01 | -8.89E-01 | -1.85E+00 |
| ENSG000001 | GLUL | 1.80E-01 | | 5.30E-01 | 1.08E+00 | 1.06E-01 | 1.08E+00 |
| ENSG000001 | NIF3L1 | 1.81E-01 | | 5.31E-01 | 9.45E-01 | -8.12E-02 | -1.06E+00 |
| ENSG000002 | CSPG4P10 | 1.81E-01 | | 5.31E-01 | 7.88E-01 | -3.44E-01 | -1.27E+00 |
| ENSG000001 | TOMM40 | 1.81E-01 | | 5.31E-01 | 1.08E+00 | 1.05E-01 | 1.08E+00 |
| ENSG000002 | --- | 1.81E-01 | ? | | 1.58E+00 | 6.60E-01 | 1.58E+00 |
| ENSG000002 | SEC14L1P1 | 1.81E-01 | ? | | 7.80E-01 | -3.58E-01 | -1.28E+00 |
| ENSG000001 | HHIPL1 | 1.81E-01 | | 5.31E-01 | 1.18E+00 | 2.42E-01 | 1.18E+00 |
| ENSG000001 | SDCBP2 | 1.81E-01 | | 5.31E-01 | 1.10E+00 | 1.38E-01 | 1.10E+00 |
| ENSG000001 | PELO | 1.81E-01 | | 5.31E-01 | 1.05E+00 | 6.79E-02 | 1.05E+00 |
| ENSG000001 | MAP2K6 | 1.81E-01 | ? | | 6.27E-01 | -6.73E-01 | -1.59E+00 |
| ENSG000001 | TLCD2 | 1.81E-01 | | 5.31E-01 | 1.08E+00 | 1.11E-01 | 1.08E+00 |

| | | | | | | |
|---|---|---|---|---|---|---|
| ENSG000002 | ZNF718 | 1.81E-01 | 5.31E-01 | 1.13E+00 | 1.71E-01 | 1.13E+00 |
| ENSG000001 | ENTREP1 | 1.81E-01 | 5.31E-01 | 9.12E-01 | -1.34E-01 | -1.10E+00 |
| ENSG000002 | --- | 1.81E-01 | ? | 7.23E-01 | -4.68E-01 | -1.38E+00 |
| ENSG000001 | ITGA2 | 1.81E-01 | 5.31E-01 | 9.48E-01 | -7.68E-02 | -1.05E+00 |
| ENSG000002 | --- | 1.81E-01 | 5.31E-01 | 1.19E+00 | 2.50E-01 | 1.19E+00 |
| ENSG000001 | CLN6 | 1.81E-01 | 5.31E-01 | 1.08E+00 | 1.08E-01 | 1.08E+00 |
| ENSG000001 | TNFRSF11A | 1.81E-01 | 5.32E-01 | 1.17E+00 | 2.33E-01 | 1.17E+00 |
| ENSG000002 | MFSD13B | 1.81E-01 | ? | 2.26E+00 | 1.18E+00 | 2.26E+00 |
| ENSG000002 | CNN2P9 | 1.82E-01 | ? | 2.04E+00 | 1.03E+00 | 2.04E+00 |
| ENSG000002 | MAGI2-AS3 | 1.82E-01 | 5.32E-01 | 9.02E-01 | -1.48E-01 | -1.11E+00 |
| ENSG000002 | --- | 1.82E-01 | 5.32E-01 | 8.85E-01 | -1.75E-01 | -1.13E+00 |
| ENSG000002 | ANKRD36BP1 | 1.82E-01 | ? | 3.63E-01 | -1.46E+00 | -2.76E+00 |
| ENSG000001 | ING1 | 1.82E-01 | 5.32E-01 | 9.27E-01 | -1.09E-01 | -1.08E+00 |
| ENSG000001 | HECA | 1.82E-01 | 5.32E-01 | 9.56E-01 | -6.52E-02 | -1.05E+00 |
| ENSG000001 | SLITRK6 | 1.82E-01 | 5.32E-01 | 8.83E-01 | -1.79E-01 | -1.13E+00 |
| ENSG000001 | PEA15 | 1.82E-01 | 5.32E-01 | 9.39E-01 | -9.03E-02 | -1.06E+00 |
| ENSG000001 | MRPS12 | 1.82E-01 | 5.32E-01 | 1.11E+00 | 1.54E-01 | 1.11E+00 |
| ENSG000001 | NOG | 1.82E-01 | ? | 1.29E+00 | 3.66E-01 | 1.29E+00 |
| ENSG000001 | ATAD1 | 1.82E-01 | 5.32E-01 | 9.58E-01 | -6.12E-02 | -1.04E+00 |
| ENSG000001 | TSEN54 | 1.82E-01 | 5.32E-01 | 1.11E+00 | 1.45E-01 | 1.11E+00 |
| ENSG000002 | MAPK6P3 | 1.82E-01 | ? | 1.82E+00 | 8.61E-01 | 1.82E+00 |
| ENSG000002 | TVP23C-CDRT | 1.82E-01 | 5.32E-01 | 9.01E-01 | -1.51E-01 | -1.11E+00 |
| ENSG000001 | SLF2 | 1.82E-01 | 5.32E-01 | 9.39E-01 | -9.14E-02 | -1.07E+00 |
| ENSG000000 | AGPS | 1.82E-01 | 5.32E-01 | 9.54E-01 | -6.83E-02 | -1.05E+00 |
| ENSG000001 | PTPRN2 | 1.82E-01 | ? | 1.48E+00 | 5.64E-01 | 1.48E+00 |
| ENSG000001 | MINAR1 | 1.82E-01 | ? | 7.60E-01 | -3.97E-01 | -1.32E+00 |
| ENSG000000 | TMEM132A | 1.82E-01 | 5.32E-01 | 1.10E+00 | 1.31E-01 | 1.10E+00 |
| ENSG000001 | RHOG | 1.82E-01 | 5.32E-01 | 1.08E+00 | 1.11E-01 | 1.08E+00 |
| ENSG000002 | PAXBP1-AS1 | 1.82E-01 | 5.32E-01 | 8.56E-01 | -2.24E-01 | -1.17E+00 |
| ENSG000001 | GPT | 1.82E-01 | ? | 1.74E+00 | 7.98E-01 | 1.74E+00 |
| ENSG000002 | TUBB8P1 | 1.82E-01 | ? | 1.84E+00 | 8.83E-01 | 1.84E+00 |
| ENSG000002 | RGL2 | 1.82E-01 | 5.32E-01 | 1.07E+00 | 9.42E-02 | 1.07E+00 |
| ENSG000001 | MORF4L2 | 1.82E-01 | 5.32E-01 | 9.71E-01 | -4.18E-02 | -1.03E+00 |
| ENSG000002 | LINC00540 | 1.82E-01 | ? | 7.43E-01 | -4.29E-01 | -1.35E+00 |
| ENSG000000 | DTX2 | 1.82E-01 | 5.32E-01 | 1.05E+00 | 7.35E-02 | 1.05E+00 |
| ENSG000001 | IGSF10 | 1.82E-01 | ? | 6.04E-01 | -7.28E-01 | -1.66E+00 |
| ENSG000002 | AGBL1 | 1.82E-01 | ? | 7.81E-01 | -3.56E-01 | -1.28E+00 |
| ENSG000002 | AFG3L1P | 1.82E-01 | 5.33E-01 | 9.15E-01 | -1.27E-01 | -1.09E+00 |
| ENSG000001 | TRIM35 | 1.82E-01 | 5.33E-01 | 9.32E-01 | -1.02E-01 | -1.07E+00 |
| ENSG000001 | CASP2 | 1.82E-01 | 5.33E-01 | 9.60E-01 | -5.87E-02 | -1.04E+00 |
| ENSG000001 | CRABP2 | 1.83E-01 | 5.33E-01 | 1.16E+00 | 2.10E-01 | 1.16E+00 |
| ENSG000002 | TAT-AS1 | 1.83E-01 | ? | 6.58E-01 | -6.05E-01 | -1.52E+00 |
| ENSG000001 | CYB5D1 | 1.83E-01 | 5.33E-01 | 9.22E-01 | -1.17E-01 | -1.08E+00 |
| ENSG000002 | --- | 1.83E-01 | ? | 1.62E+00 | 6.98E-01 | 1.62E+00 |
| ENSG000001 | TK1 | 1.83E-01 | 5.33E-01 | 8.79E-01 | -1.86E-01 | -1.14E+00 |

| | | | | | | |
|---|---|---|---|---|---|---|
| ENSG000002 | --- | 1.83E-01 | ? | 6.95E-01 | -5.25E-01 | -1.44E+00 |
| ENSG000001 | NEK9 | 1.83E-01 | 5.33E-01 | 1.05E+00 | 6.53E-02 | 1.05E+00 |
| ENSG000000 | CLTCL1 | 1.83E-01 | 5.33E-01 | 1.06E+00 | 8.07E-02 | 1.06E+00 |
| ENSG000001 | KMT5A | 1.83E-01 | 5.33E-01 | 1.03E+00 | 4.85E-02 | 1.03E+00 |
| ENSG000000 | WAC | 1.83E-01 | 5.33E-01 | 9.48E-01 | -7.64E-02 | -1.05E+00 |
| ENSG000002 | --- | 1.83E-01 | ? | 6.59E-01 | -6.01E-01 | -1.52E+00 |
| ENSG000002 | PMS2P3 | 1.83E-01 | 5.33E-01 | 8.82E-01 | -1.81E-01 | -1.13E+00 |
| ENSG000002 | DPP9-AS1 | 1.83E-01 | ? | 7.71E-01 | -3.75E-01 | -1.30E+00 |
| ENSG000001 | SNW1 | 1.83E-01 | 5.33E-01 | 9.49E-01 | -7.57E-02 | -1.05E+00 |
| ENSG000001 | RAVER2 | 1.83E-01 | 5.33E-01 | 1.06E+00 | 8.20E-02 | 1.06E+00 |
| ENSG000001 | FLG | 1.83E-01 | 5.33E-01 | 1.12E+00 | 1.67E-01 | 1.12E+00 |
| ENSG000001 | MAGED4B | 1.83E-01 | 5.33E-01 | 7.75E-01 | -3.67E-01 | -1.29E+00 |
| ENSG000002 | --- | 1.83E-01 | 5.33E-01 | 8.14E-01 | -2.97E-01 | -1.23E+00 |
| ENSG000001 | UBE2C | 1.83E-01 | 5.33E-01 | 8.25E-01 | -2.78E-01 | -1.21E+00 |
| ENSG000002 | --- | 1.83E-01 | ? | 7.29E-01 | -4.56E-01 | -1.37E+00 |
| ENSG000002 | --- | 1.83E-01 | ? | 4.13E-01 | -1.28E+00 | -2.42E+00 |
| ENSG000001 | PBDC1 | 1.83E-01 | 5.34E-01 | 9.51E-01 | -7.25E-02 | -1.05E+00 |
| ENSG000001 | SPRED1 | 1.83E-01 | 5.34E-01 | 9.36E-01 | -9.56E-02 | -1.07E+00 |
| ENSG000001 | BMI1 | 1.84E-01 | 5.34E-01 | 9.51E-01 | -7.21E-02 | -1.05E+00 |
| ENSG000001 | RPS6KB1 | 1.84E-01 | 5.34E-01 | 9.46E-01 | -8.03E-02 | -1.06E+00 |
| ENSG000001 | RAB4B-EGLN | 1.84E-01 | 5.34E-01 | 1.55E+00 | 6.36E-01 | 1.55E+00 |
| ENSG000001 | VPS37D | 1.84E-01 | ? | 6.77E-01 | -5.63E-01 | -1.48E+00 |
| ENSG000001 | AK7 | 1.84E-01 | ? | 7.06E-01 | -5.03E-01 | -1.42E+00 |
| ENSG000001 | S100A3 | 1.84E-01 | 5.34E-01 | 1.17E+00 | 2.27E-01 | 1.17E+00 |
| ENSG000001 | SLC37A4 | 1.84E-01 | 5.34E-01 | 1.10E+00 | 1.35E-01 | 1.10E+00 |
| ENSG000002 | --- | 1.84E-01 | ? | 7.54E-01 | -4.08E-01 | -1.33E+00 |
| ENSG000002 | LINC01847 | 1.84E-01 | ? | 5.06E-01 | -9.84E-01 | -1.98E+00 |
| ENSG000002 | --- | 1.84E-01 | ? | 6.30E-01 | -6.67E-01 | -1.59E+00 |
| ENSG000001 | U2AF1L4 | 1.84E-01 | 5.34E-01 | 8.93E-01 | -1.63E-01 | -1.12E+00 |
| ENSG000001 | MT-ND1 | 1.84E-01 | 5.34E-01 | 8.73E-01 | -1.97E-01 | -1.15E+00 |
| ENSG000001 | ALDH6A1 | 1.84E-01 | 5.34E-01 | 1.08E+00 | 1.11E-01 | 1.08E+00 |
| ENSG000001 | CACYBPP2 | 1.84E-01 | ? | 7.63E-01 | -3.90E-01 | -1.31E+00 |
| ENSG000002 | DARS1-AS1 | 1.84E-01 | 5.34E-01 | 1.21E+00 | 2.75E-01 | 1.21E+00 |
| ENSG000001 | KIF24 | 1.84E-01 | 5.34E-01 | 1.13E+00 | 1.79E-01 | 1.13E+00 |
| ENSG000000 | ARHGAP44 | 1.84E-01 | 5.34E-01 | 1.11E+00 | 1.55E-01 | 1.11E+00 |
| ENSG000001 | SLC17A5 | 1.84E-01 | 5.34E-01 | 1.05E+00 | 6.37E-02 | 1.05E+00 |
| ENSG000002 | --- | 1.84E-01 | ? | 1.47E+00 | 5.60E-01 | 1.47E+00 |
| ENSG000001 | MRPL58 | 1.84E-01 | 5.34E-01 | 9.44E-01 | -8.24E-02 | -1.06E+00 |
| ENSG000001 | ADGRE5 | 1.84E-01 | 5.34E-01 | 1.07E+00 | 9.17E-02 | 1.07E+00 |
| ENSG000001 | SDC2 | 1.84E-01 | 5.34E-01 | 1.06E+00 | 9.09E-02 | 1.06E+00 |
| ENSG000001 | EPHA2 | 1.84E-01 | 5.34E-01 | 9.39E-01 | -9.06E-02 | -1.06E+00 |
| ENSG000001 | TFAP2A | 1.84E-01 | 5.34E-01 | 9.44E-01 | -8.34E-02 | -1.06E+00 |
| ENSG000001 | ELK3 | 1.84E-01 | 5.34E-01 | 1.06E+00 | 7.93E-02 | 1.06E+00 |
| ENSG000002 | PLEKHM1 | 1.84E-01 | 5.34E-01 | 1.08E+00 | 1.06E-01 | 1.08E+00 |
| ENSG000001 | DUOXA1 | 1.84E-01 | 5.34E-01 | 1.11E+00 | 1.56E-01 | 1.11E+00 |

| | | | | | | |
|---|---|---|---|---|---|---|
| ENSG000001 | TXNDC9 | 1.85E-01 | 5.34E-01 | 9.58E-01 | -6.21E-02 | -1.04E+00 |
| ENSG000002 | LINC00705 | 1.85E-01 | 5.34E-01 | 8.49E-01 | -2.36E-01 | -1.18E+00 |
| ENSG000001 | ACADM | 1.85E-01 | 5.34E-01 | 1.05E+00 | 7.55E-02 | 1.05E+00 |
| ENSG000001 | MFSD4B | 1.85E-01 | 5.34E-01 | 1.14E+00 | 1.83E-01 | 1.14E+00 |
| ENSG000002 | JMJD7 | 1.85E-01 | 5.34E-01 | 1.19E+00 | 2.46E-01 | 1.19E+00 |
| ENSG000002 | RBM12 | 1.85E-01 | 5.34E-01 | 9.51E-01 | -7.18E-02 | -1.05E+00 |
| ENSG00000C | TLL2 | 1.85E-01 | 5.34E-01 | 8.22E-01 | -2.83E-01 | -1.22E+00 |
| ENSG000001 | PRR15L | 1.85E-01 | ? | 1.47E+00 | 5.58E-01 | 1.47E+00 |
| ENSG000001 | SPATA4 | 1.85E-01 | ? | 5.73E-01 | -8.03E-01 | -1.74E+00 |
| ENSG000001 | SPOCK1 | 1.85E-01 | 5.35E-01 | 9.55E-01 | -6.65E-02 | -1.05E+00 |
| ENSG000001 | PBLD | 1.85E-01 | 5.35E-01 | 1.10E+00 | 1.39E-01 | 1.10E+00 |
| ENSG000001 | ZIK1 | 1.85E-01 | 5.35E-01 | 1.09E+00 | 1.21E-01 | 1.09E+00 |
| ENSG000002 | TBC1D8 | 1.85E-01 | 5.35E-01 | 1.06E+00 | 8.24E-02 | 1.06E+00 |
| ENSG000001 | GTDC1 | 1.85E-01 | 5.35E-01 | 1.06E+00 | 8.07E-02 | 1.06E+00 |
| ENSG000002 | C2orf72 | 1.85E-01 | ? | 1.34E+00 | 4.23E-01 | 1.34E+00 |
| ENSG000001 | ZNF624 | 1.85E-01 | 5.35E-01 | 9.09E-01 | -1.37E-01 | -1.10E+00 |
| ENSG000001 | CELSR2 | 1.85E-01 | 5.35E-01 | 1.06E+00 | 8.96E-02 | 1.06E+00 |
| ENSG000001 | SYNM | 1.85E-01 | 5.35E-01 | 1.22E+00 | 2.87E-01 | 1.22E+00 |
| ENSG000001 | CHD8 | 1.85E-01 | 5.35E-01 | 9.58E-01 | -6.14E-02 | -1.04E+00 |
| ENSG000001 | TSEN2 | 1.85E-01 | 5.35E-01 | 1.09E+00 | 1.27E-01 | 1.09E+00 |
| ENSG000002 | --- | 1.85E-01 | ? | 7.74E-01 | -3.69E-01 | -1.29E+00 |
| ENSG000001 | SUV39H1 | 1.85E-01 | 5.35E-01 | 1.11E+00 | 1.51E-01 | 1.11E+00 |
| ENSG00000C | TASP1 | 1.85E-01 | 5.35E-01 | 1.08E+00 | 1.06E-01 | 1.08E+00 |
| ENSG000001 | LETM1 | 1.85E-01 | 5.35E-01 | 9.56E-01 | -6.43E-02 | -1.05E+00 |
| ENSG000002 | CFAP206 | 1.85E-01 | ? | 1.84E+00 | 8.77E-01 | 1.84E+00 |
| ENSG000002 | --- | 1.85E-01 | 5.35E-01 | 9.00E-01 | -1.52E-01 | -1.11E+00 |
| ENSG000002 | --- | 1.86E-01 | 5.35E-01 | 1.15E+00 | 2.00E-01 | 1.15E+00 |
| ENSG000002 | --- | 1.86E-01 | ? | 7.42E-01 | -4.31E-01 | -1.35E+00 |
| ENSG000002 | ZSCAN16-AS | 1.86E-01 | 5.35E-01 | 1.16E+00 | 2.16E-01 | 1.16E+00 |
| ENSG000001 | PTP4A1 | 1.86E-01 | 5.35E-01 | 9.28E-01 | -1.08E-01 | -1.08E+00 |
| ENSG000001 | PARP9 | 1.86E-01 | 5.36E-01 | 9.37E-01 | -9.32E-02 | -1.07E+00 |
| ENSG000002 | --- | 1.86E-01 | ? | 1.37E+00 | 4.59E-01 | 1.37E+00 |
| ENSG00000C | EIF4B | 1.86E-01 | 5.36E-01 | 1.04E+00 | 5.82E-02 | 1.04E+00 |
| ENSG000001 | ORC3 | 1.86E-01 | 5.36E-01 | 9.58E-01 | -6.16E-02 | -1.04E+00 |
| ENSG000002 | --- | 1.86E-01 | ? | 6.00E-01 | -7.36E-01 | -1.67E+00 |
| ENSG000001 | RNF135 | 1.86E-01 | 5.36E-01 | 1.09E+00 | 1.25E-01 | 1.09E+00 |
| ENSG000001 | UBAC1 | 1.86E-01 | 5.36E-01 | 1.04E+00 | 5.94E-02 | 1.04E+00 |
| ENSG000001 | MINK1 | 1.86E-01 | 5.36E-01 | 1.04E+00 | 5.82E-02 | 1.04E+00 |
| ENSG000002 | --- | 1.86E-01 | 5.36E-01 | 1.11E+00 | 1.52E-01 | 1.11E+00 |
| ENSG000001 | CAND1 | 1.86E-01 | 5.37E-01 | 1.04E+00 | 5.42E-02 | 1.04E+00 |
| ENSG000002 | PPP1R10 | 1.86E-01 | 5.37E-01 | 9.59E-01 | -6.03E-02 | -1.04E+00 |
| ENSG000002 | ZSWIM8-AS1 | 1.87E-01 | ? | 6.66E-01 | -5.87E-01 | -1.50E+00 |
| ENSG000001 | CENPE | 1.87E-01 | 5.37E-01 | 9.11E-01 | -1.34E-01 | -1.10E+00 |
| ENSG000002 | MIR3677HG | 1.87E-01 | ? | 1.60E+00 | 6.76E-01 | 1.60E+00 |
| ENSG000001 | GNAT2 | 1.87E-01 | ? | 7.15E-01 | -4.83E-01 | -1.40E+00 |

| | | | | | | |
|---|---|---|---|---|---|---|
| ENSG000001 | PIGG | 1.87E-01 | 5.38E-01 | 9.61E-01 | -5.71E-02 | -1.04E+00 |
| ENSG000001 | CTNNBL1 | 1.87E-01 | 5.38E-01 | 1.04E+00 | 5.49E-02 | 1.04E+00 |
| ENSG000002 | OTUD7B | 1.87E-01 | 5.38E-01 | 9.61E-01 | -5.72E-02 | -1.04E+00 |
| ENSG000002 | --- | 1.87E-01 | 5.38E-01 | 1.12E+00 | 1.60E-01 | 1.12E+00 |
| ENSG000001 | SLC45A1 | 1.87E-01 | ? | 1.49E+00 | 5.78E-01 | 1.49E+00 |
| ENSG000002 | --- | 1.87E-01 | ? | 1.63E+00 | 7.09E-01 | 1.63E+00 |
| ENSG000002 | ELN-AS1 | 1.87E-01 | ? | 6.58E-01 | -6.04E-01 | -1.52E+00 |
| ENSG000001 | SELP | 1.87E-01 | ? | 7.42E-01 | -4.30E-01 | -1.35E+00 |
| ENSG000001 | INO80D | 1.88E-01 | 5.39E-01 | 9.27E-01 | -1.09E-01 | -1.08E+00 |
| ENSG000002 | --- | 1.88E-01 | 5.39E-01 | 8.66E-01 | -2.08E-01 | -1.16E+00 |
| ENSG000001 | TMEM45A | 1.88E-01 | 5.39E-01 | 9.25E-01 | -1.13E-01 | -1.08E+00 |
| ENSG000001 | C20orf27 | 1.88E-01 | 5.39E-01 | 1.10E+00 | 1.44E-01 | 1.10E+00 |
| ENSG000001 | TSPAN3 | 1.88E-01 | 5.39E-01 | 9.64E-01 | -5.31E-02 | -1.04E+00 |
| ENSG000002 | NDUFA7 | 1.88E-01 | 5.39E-01 | 1.08E+00 | 1.10E-01 | 1.08E+00 |
| ENSG000002 | --- | 1.88E-01 | 5.39E-01 | 1.07E+00 | 1.03E-01 | 1.07E+00 |
| ENSG000002 | TAS2R20 | 1.88E-01 | ? | 7.20E-01 | -4.74E-01 | -1.39E+00 |
| ENSG000001 | CXCL11 | 1.88E-01 | ? | 1.93E+00 | 9.45E-01 | 1.93E+00 |
| ENSG00000C | USP33 | 1.88E-01 | 5.39E-01 | 9.60E-01 | -5.85E-02 | -1.04E+00 |
| ENSG000002 | DPH5-DT | 1.88E-01 | 5.39E-01 | 1.10E+00 | 1.32E-01 | 1.10E+00 |
| ENSG000001 | ANO6 | 1.88E-01 | 5.39E-01 | 1.06E+00 | 8.32E-02 | 1.06E+00 |
| ENSG000001 | SLC25A30 | 1.88E-01 | 5.39E-01 | 9.58E-01 | -6.26E-02 | -1.04E+00 |
| ENSG000002 | RBBP4P2 | 1.88E-01 | ? | 4.79E-01 | -1.06E+00 | -2.09E+00 |
| ENSG000001 | RFC5 | 1.88E-01 | 5.39E-01 | 9.24E-01 | -1.15E-01 | -1.08E+00 |
| ENSG000001 | ZNRD2 | 1.88E-01 | 5.39E-01 | 1.09E+00 | 1.25E-01 | 1.09E+00 |
| ENSG00000C | LSG1 | 1.88E-01 | 5.39E-01 | 9.53E-01 | -6.88E-02 | -1.05E+00 |
| ENSG000002 | ZNF674 | 1.88E-01 | 5.39E-01 | 1.11E+00 | 1.53E-01 | 1.11E+00 |
| ENSG000001 | TCEAL1 | 1.88E-01 | 5.39E-01 | 1.12E+00 | 1.60E-01 | 1.12E+00 |
| ENSG000001 | PRMT1 | 1.88E-01 | 5.39E-01 | 1.05E+00 | 6.45E-02 | 1.05E+00 |
| ENSG000002 | --- | 1.88E-01 | ? | 1.93E+00 | 9.48E-01 | 1.93E+00 |
| ENSG000001 | TMC4 | 1.88E-01 | 5.39E-01 | 1.11E+00 | 1.48E-01 | 1.11E+00 |
| ENSG000001 | RTCA | 1.88E-01 | 5.39E-01 | 9.61E-01 | -5.80E-02 | -1.04E+00 |
| ENSG000001 | LY6E | 1.88E-01 | 5.39E-01 | 1.13E+00 | 1.75E-01 | 1.13E+00 |
| ENSG000002 | --- | 1.88E-01 | ? | 1.33E+00 | 4.10E-01 | 1.33E+00 |
| ENSG000002 | --- | 1.88E-01 | ? | 1.39E+00 | 4.70E-01 | 1.39E+00 |
| ENSG000001 | ALDH7A1 | 1.88E-01 | 5.39E-01 | 1.08E+00 | 1.16E-01 | 1.08E+00 |
| ENSG000001 | BCDIN3D | 1.88E-01 | 5.39E-01 | 1.09E+00 | 1.18E-01 | 1.09E+00 |
| ENSG000001 | PRPF3 | 1.88E-01 | 5.39E-01 | 9.42E-01 | -8.67E-02 | -1.06E+00 |
| ENSG000001 | TMC8 | 1.89E-01 | 5.39E-01 | 8.23E-01 | -2.81E-01 | -1.22E+00 |
| ENSG000001 | ADH5 | 1.89E-01 | 5.39E-01 | 1.04E+00 | 5.75E-02 | 1.04E+00 |
| ENSG000001 | NR1D2 | 1.89E-01 | 5.39E-01 | 8.83E-01 | -1.79E-01 | -1.13E+00 |
| ENSG000001 | APOBEC3F | 1.89E-01 | 5.40E-01 | 1.12E+00 | 1.62E-01 | 1.12E+00 |
| ENSG00000C | SNAP29 | 1.89E-01 | 5.40E-01 | 1.04E+00 | 5.97E-02 | 1.04E+00 |
| ENSG000001 | E2F1 | 1.89E-01 | 5.40E-01 | 8.08E-01 | -3.08E-01 | -1.24E+00 |
| ENSG000002 | CBR3-AS1 | 1.89E-01 | 5.40E-01 | 1.14E+00 | 1.94E-01 | 1.14E+00 |
| ENSG000001 | THBS2 | 1.89E-01 | 5.40E-01 | 1.11E+00 | 1.52E-01 | 1.11E+00 |

| | | | | | | |
|---|---|---|---|---|---|---|
| ENSG000001 | IGSF22 | 1.89E-01 | ? | 6.98E-01 | -5.18E-01 | -1.43E+00 |
| ENSG000001 | IKBIP | 1.89E-01 | 5.41E-01 | 1.05E+00 | 6.77E-02 | 1.05E+00 |
| ENSG000001 | APOLD1 | 1.89E-01 | 5.41E-01 | 8.69E-01 | -2.03E-01 | -1.15E+00 |
| ENSG000002 | LINC00885 | 1.89E-01 | ? | 1.54E+00 | 6.26E-01 | 1.54E+00 |
| ENSG000002 | --- | 1.89E-01 | ? | 4.98E-01 | -1.01E+00 | -2.01E+00 |
| ENSG00000C | ARAP2 | 1.89E-01 | 5.41E-01 | 9.45E-01 | -8.18E-02 | -1.06E+00 |
| ENSG00000C | RNF10 | 1.89E-01 | 5.41E-01 | 9.59E-01 | -6.10E-02 | -1.04E+00 |
| ENSG000002 | --- | 1.89E-01 | ? | 7.67E-01 | -3.82E-01 | -1.30E+00 |
| ENSG000002 | --- | 1.89E-01 | ? | 2.44E-01 | -2.03E+00 | -4.10E+00 |
| ENSG00000C | ARG2 | 1.89E-01 | 5.41E-01 | 1.11E+00 | 1.53E-01 | 1.11E+00 |
| ENSG000002 | HSP90B3P | 1.89E-01 | ? | 1.89E+00 | 9.17E-01 | 1.89E+00 |
| ENSG000001 | ATG14 | 1.90E-01 | 5.41E-01 | 9.36E-01 | -9.55E-02 | -1.07E+00 |
| ENSG000002 | --- | 1.90E-01 | ? | 1.30E+00 | 3.83E-01 | 1.30E+00 |
| ENSG000001 | CD63 | 1.90E-01 | 5.41E-01 | 1.03E+00 | 4.49E-02 | 1.03E+00 |
| ENSG000001 | KNL1 | 1.90E-01 | 5.41E-01 | 8.75E-01 | -1.93E-01 | -1.14E+00 |
| ENSG000002 | SGMS1-AS1 | 1.90E-01 | 5.41E-01 | 1.14E+00 | 1.84E-01 | 1.14E+00 |
| ENSG000002 | --- | 1.90E-01 | ? | 1.52E+00 | 6.00E-01 | 1.52E+00 |
| ENSG000001 | BLOC1S4 | 1.90E-01 | 5.42E-01 | 1.09E+00 | 1.25E-01 | 1.09E+00 |
| ENSG000002 | CEBPZOS | 1.90E-01 | 5.42E-01 | 1.06E+00 | 8.87E-02 | 1.06E+00 |
| ENSG000001 | LACTB | 1.90E-01 | 5.42E-01 | 1.05E+00 | 7.27E-02 | 1.05E+00 |
| ENSG000001 | ZACN | 1.90E-01 | ? | 6.15E-01 | -7.01E-01 | -1.63E+00 |
| ENSG000002 | ERVK9-11 | 1.90E-01 | ? | 7.34E-01 | -4.46E-01 | -1.36E+00 |
| ENSG000002 | --- | 1.90E-01 | ? | 6.38E-01 | -6.47E-01 | -1.57E+00 |
| ENSG000001 | LRP1B | 1.90E-01 | ? | 2.08E+00 | 1.05E+00 | 2.08E+00 |
| ENSG000002 | --- | 1.90E-01 | ? | 1.76E+00 | 8.18E-01 | 1.76E+00 |
| ENSG000001 | DLG5 | 1.90E-01 | 5.42E-01 | 1.04E+00 | 5.13E-02 | 1.04E+00 |
| ENSG000001 | ELOC | 1.90E-01 | 5.42E-01 | 9.56E-01 | -6.47E-02 | -1.05E+00 |
| ENSG000001 | GNB2 | 1.90E-01 | 5.42E-01 | 1.08E+00 | 1.12E-01 | 1.08E+00 |
| ENSG000002 | --- | 1.90E-01 | ? | 2.90E+00 | 1.54E+00 | 2.90E+00 |
| ENSG000001 | MMP24OS | 1.91E-01 | 5.42E-01 | 1.05E+00 | 7.64E-02 | 1.05E+00 |
| ENSG000001 | RALY | 1.91E-01 | 5.42E-01 | 1.05E+00 | 7.09E-02 | 1.05E+00 |
| ENSG000001 | CLUAP1 | 1.91E-01 | 5.42E-01 | 9.41E-01 | -8.72E-02 | -1.06E+00 |
| ENSG000001 | ZNF821 | 1.91E-01 | 5.42E-01 | 1.14E+00 | 1.84E-01 | 1.14E+00 |
| ENSG000002 | PSMC1P1 | 1.91E-01 | 5.42E-01 | 9.27E-01 | -1.09E-01 | -1.08E+00 |
| ENSG000001 | DEDD | 1.91E-01 | 5.42E-01 | 1.05E+00 | 6.82E-02 | 1.05E+00 |
| ENSG00000C | POLD1 | 1.91E-01 | 5.42E-01 | 1.12E+00 | 1.59E-01 | 1.12E+00 |
| ENSG000001 | C1RL | 1.91E-01 | 5.42E-01 | 1.06E+00 | 8.76E-02 | 1.06E+00 |
| ENSG000001 | ZC3H13 | 1.91E-01 | 5.42E-01 | 9.35E-01 | -9.64E-02 | -1.07E+00 |
| ENSG000001 | HYCC2 | 1.91E-01 | 5.42E-01 | 1.07E+00 | 1.04E-01 | 1.07E+00 |
| ENSG000001 | CHMP6 | 1.91E-01 | 5.42E-01 | 1.08E+00 | 1.08E-01 | 1.08E+00 |
| ENSG000001 | LINC00467 | 1.91E-01 | 5.42E-01 | 9.14E-01 | -1.31E-01 | -1.09E+00 |
| ENSG000002 | --- | 1.91E-01 | 5.42E-01 | 7.60E-01 | -3.96E-01 | -1.32E+00 |
| ENSG000002 | RPL7AP6 | 1.91E-01 | 5.42E-01 | 1.10E+00 | 1.36E-01 | 1.10E+00 |
| ENSG000002 | GTF2IRD1P1 | 1.91E-01 | ? | 1.57E+00 | 6.55E-01 | 1.57E+00 |
| ENSG000001 | HIVEP3 | 1.91E-01 | 5.42E-01 | 8.94E-01 | -1.61E-01 | -1.12E+00 |

| | | | | | | |
|---|---|---|---|---|---|---|
| ENSG000001 | RPS15 | 1.91E-01 | 5.42E-01 | 1.08E+00 | 1.13E-01 | 1.08E+00 |
| ENSG000002 | BTF3-DT | 1.91E-01 | ? | 1.82E+00 | 8.61E-01 | 1.82E+00 |
| ENSG000001 | SUDS3 | 1.91E-01 | 5.42E-01 | 9.57E-01 | -6.32E-02 | -1.04E+00 |
| ENSG000001 | AVPI1 | 1.91E-01 | 5.42E-01 | 1.19E+00 | 2.49E-01 | 1.19E+00 |
| ENSG000001 | SCOC | 1.91E-01 | 5.42E-01 | 9.56E-01 | -6.42E-02 | -1.05E+00 |
| ENSG000001 | PKD2 | 1.91E-01 | 5.42E-01 | 1.05E+00 | 7.23E-02 | 1.05E+00 |
| ENSG000001 | NOP53 | 1.91E-01 | 5.43E-01 | 1.08E+00 | 1.14E-01 | 1.08E+00 |
| ENSG000001 | TLCD4 | 1.91E-01 | ? | 7.04E-01 | -5.06E-01 | -1.42E+00 |
| ENSG000001 | IMPACT | 1.91E-01 | 5.43E-01 | 1.06E+00 | 7.88E-02 | 1.06E+00 |
| ENSG000002 | --- | 1.92E-01 | ? | 6.91E-01 | -5.34E-01 | -1.45E+00 |
| ENSG000001 | SNAI3 | 1.92E-01 | ? | 1.30E+00 | 3.82E-01 | 1.30E+00 |
| ENSG000001 | FOXL2 | 1.92E-01 | 5.43E-01 | 1.11E+00 | 1.48E-01 | 1.11E+00 |
| ENSG000001 | CBLL1 | 1.92E-01 | 5.43E-01 | 9.46E-01 | -7.96E-02 | -1.06E+00 |
| ENSG000001 | UNC45A | 1.92E-01 | 5.43E-01 | 1.06E+00 | 8.14E-02 | 1.06E+00 |
| ENSG000002 | FDPSP7 | 1.92E-01 | ? | 1.45E+00 | 5.35E-01 | 1.45E+00 |
| ENSG000001 | ATP6V0B | 1.92E-01 | 5.44E-01 | 1.09E+00 | 1.22E-01 | 1.09E+00 |
| ENSG000001 | PGD | 1.92E-01 | 5.44E-01 | 9.45E-01 | -8.13E-02 | -1.06E+00 |
| ENSG000001 | MOCS1 | 1.92E-01 | 5.44E-01 | 1.08E+00 | 1.18E-01 | 1.08E+00 |
| ENSG000001 | ATG12 | 1.92E-01 | 5.44E-01 | 9.65E-01 | -5.11E-02 | -1.04E+00 |
| ENSG000001 | ERLIN1 | 1.92E-01 | 5.44E-01 | 9.54E-01 | -6.74E-02 | -1.05E+00 |
| ENSG000001 | ATXN7 | 1.92E-01 | 5.44E-01 | 9.50E-01 | -7.36E-02 | -1.05E+00 |
| ENSG000001 | CNPY3 | 1.92E-01 | 5.44E-01 | 1.07E+00 | 9.55E-02 | 1.07E+00 |
| ENSG000002 | --- | 1.92E-01 | ? | 1.65E+00 | 7.27E-01 | 1.65E+00 |
| ENSG000002 | PRECSIT | 1.92E-01 | ? | 1.39E+00 | 4.75E-01 | 1.39E+00 |
| ENSG000002 | --- | 1.92E-01 | 5.44E-01 | 1.24E+00 | 3.15E-01 | 1.24E+00 |
| ENSG000002 | KRT18P4 | 1.92E-01 | ? | 1.83E+00 | 8.71E-01 | 1.83E+00 |
| ENSG000002 | SHQ1P1 | 1.92E-01 | ? | 6.38E-01 | -6.47E-01 | -1.57E+00 |
| ENSG000001 | RBX1 | 1.92E-01 | 5.44E-01 | 1.06E+00 | 8.26E-02 | 1.06E+00 |
| ENSG00000C | ERGIC2 | 1.92E-01 | 5.44E-01 | 9.44E-01 | -8.39E-02 | -1.06E+00 |
| ENSG000001 | HR | 1.92E-01 | 5.44E-01 | 8.68E-01 | -2.05E-01 | -1.15E+00 |
| ENSG000002 | MT-ND4L | 1.92E-01 | 5.44E-01 | 8.70E-01 | -2.00E-01 | -1.15E+00 |
| ENSG000002 | PCDHGA10 | 1.93E-01 | 5.44E-01 | 8.94E-01 | -1.61E-01 | -1.12E+00 |
| ENSG000001 | REV1 | 1.93E-01 | 5.44E-01 | 1.04E+00 | 6.20E-02 | 1.04E+00 |
| ENSG000002 | --- | 1.93E-01 | 5.44E-01 | 7.11E-01 | -4.93E-01 | -1.41E+00 |
| ENSG000002 | --- | 1.93E-01 | 5.44E-01 | 7.11E-01 | -4.93E-01 | -1.41E+00 |
| ENSG000001 | CTBS | 1.93E-01 | 5.44E-01 | 9.42E-01 | -8.67E-02 | -1.06E+00 |
| ENSG000001 | NBR1 | 1.93E-01 | 5.44E-01 | 9.64E-01 | -5.32E-02 | -1.04E+00 |
| ENSG000001 | NCAPH | 1.93E-01 | 5.44E-01 | 8.40E-01 | -2.51E-01 | -1.19E+00 |
| ENSG000002 | POLR2J2 | 1.93E-01 | ? | 6.89E-01 | -5.37E-01 | -1.45E+00 |
| ENSG000001 | C1orf52 | 1.93E-01 | 5.44E-01 | 9.50E-01 | -7.41E-02 | -1.05E+00 |
| ENSG000001 | ALMS1 | 1.93E-01 | 5.45E-01 | 9.26E-01 | -1.10E-01 | -1.08E+00 |
| ENSG000001 | ZFP36 | 1.93E-01 | 5.45E-01 | 9.16E-01 | -1.26E-01 | -1.09E+00 |
| ENSG00000C | DNMT3B | 1.93E-01 | 5.45E-01 | 8.66E-01 | -2.07E-01 | -1.15E+00 |
| ENSG000002 | --- | 1.93E-01 | 5.45E-01 | 7.81E-01 | -3.56E-01 | -1.28E+00 |
| ENSG000001 | CSNK1G3 | 1.93E-01 | 5.45E-01 | 9.48E-01 | -7.77E-02 | -1.06E+00 |

| | | | | | | | |
|---|---|---|---|---|---|---|---|
| ENSG000001 | TP53INP1 | 1.93E-01 | | 5.45E-01 | 1.07E+00 | 1.02E-01 | 1.07E+00 |
| ENSG000002 | --- | 1.93E-01 | | 5.45E-01 | 1.22E+00 | 2.83E-01 | 1.22E+00 |
| ENSG000001 | OAF | 1.93E-01 | | 5.45E-01 | 1.05E+00 | 7.51E-02 | 1.05E+00 |
| ENSG000001 | PPIL4 | 1.93E-01 | | 5.45E-01 | 9.34E-01 | -9.83E-02 | -1.07E+00 |
| ENSG000001 | DEAF1 | 1.93E-01 | | 5.45E-01 | 1.07E+00 | 9.58E-02 | 1.07E+00 |
| ENSG000001 | CCN3 | 1.93E-01 | ? | | 7.46E-01 | -4.23E-01 | -1.34E+00 |
| ENSG000001 | PTGDS | 1.93E-01 | ? | | 2.40E+00 | 1.27E+00 | 2.40E+00 |
| ENSG000001 | PLXNA3 | 1.93E-01 | | 5.45E-01 | 9.38E-01 | -9.29E-02 | -1.07E+00 |
| ENSG000001 | AP3B1 | 1.93E-01 | | 5.45E-01 | 9.63E-01 | -5.38E-02 | -1.04E+00 |
| ENSG000001 | SMIM29 | 1.94E-01 | | 5.45E-01 | 1.08E+00 | 1.11E-01 | 1.08E+00 |
| ENSG000002 | NUTM2A-AS | 1.94E-01 | | 5.45E-01 | 9.48E-01 | -7.72E-02 | -1.06E+00 |
| ENSG000001 | WDFY2 | 1.94E-01 | | 5.45E-01 | 9.67E-01 | -4.81E-02 | -1.03E+00 |
| ENSG000001 | PTGES | 1.94E-01 | | 5.45E-01 | 1.19E+00 | 2.57E-01 | 1.19E+00 |
| ENSG000002 | SAMD9 | 1.94E-01 | | 5.45E-01 | 9.17E-01 | -1.24E-01 | -1.09E+00 |
| ENSG000001 | MALT1 | 1.94E-01 | | 5.45E-01 | 9.53E-01 | -6.89E-02 | -1.05E+00 |
| ENSG000001 | ZNF622 | 1.94E-01 | | 5.45E-01 | 9.59E-01 | -5.98E-02 | -1.04E+00 |
| ENSG000002 | SRGAP2D | 1.94E-01 | | 5.45E-01 | 1.18E+00 | 2.42E-01 | 1.18E+00 |
| ENSG000002 | MT-RNR1 | 1.94E-01 | | 5.45E-01 | 1.09E+00 | 1.20E-01 | 1.09E+00 |
| ENSG000002 | --- | 1.94E-01 | ? | | 6.39E-01 | -6.47E-01 | -1.57E+00 |
| ENSG000002 | VDAC1 | 1.94E-01 | | 5.45E-01 | 1.03E+00 | 4.31E-02 | 1.03E+00 |
| ENSG000001 | BCL6 | 1.94E-01 | | 5.45E-01 | 9.35E-01 | -9.75E-02 | -1.07E+00 |
| ENSG000001 | FRAT1 | 1.94E-01 | ? | | 1.23E+00 | 2.99E-01 | 1.23E+00 |
| ENSG000001 | RABL3 | 1.94E-01 | | 5.45E-01 | 9.60E-01 | -5.88E-02 | -1.04E+00 |
| ENSG000001 | OSBPL9 | 1.94E-01 | | 5.45E-01 | 9.58E-01 | -6.13E-02 | -1.04E+00 |
| ENSG000002 | --- | 1.94E-01 | ? | | 1.57E+00 | 6.54E-01 | 1.57E+00 |
| ENSG000002 | FPGT-TNNI3 | 1.94E-01 | ? | | 6.83E-01 | -5.51E-01 | -1.46E+00 |
| ENSG000002 | TMEM30BP1 | 1.94E-01 | ? | | 1.49E+00 | 5.72E-01 | 1.49E+00 |
| ENSG000002 | PARTICL | 1.94E-01 | ? | | 7.65E-01 | -3.86E-01 | -1.31E+00 |
| ENSG000002 | GALNT4 | 1.94E-01 | | 5.46E-01 | 1.07E+00 | 9.34E-02 | 1.07E+00 |
| ENSG000002 | SMKR1 | 1.94E-01 | ? | | 1.44E+00 | 5.29E-01 | 1.44E+00 |
| ENSG00000C | PIK3CB | 1.94E-01 | | 5.46E-01 | 9.61E-01 | -5.69E-02 | -1.04E+00 |
| ENSG00000C | BRINP1 | 1.94E-01 | | 5.46E-01 | 1.14E+00 | 1.88E-01 | 1.14E+00 |
| ENSG000001 | RASD1 | 1.94E-01 | | 5.46E-01 | 8.60E-01 | -2.17E-01 | -1.16E+00 |
| ENSG000002 | --- | 1.95E-01 | | 5.46E-01 | 8.51E-01 | -2.32E-01 | -1.17E+00 |
| ENSG000002 | RBM12B-AS: | 1.95E-01 | ? | | 1.40E+00 | 4.86E-01 | 1.40E+00 |
| ENSG000001 | NIP7 | 1.95E-01 | | 5.46E-01 | 9.52E-01 | -7.16E-02 | -1.05E+00 |
| ENSG000001 | SPINDOC | 1.95E-01 | | 5.46E-01 | 1.06E+00 | 8.51E-02 | 1.06E+00 |
| ENSG000001 | LAMTOR4 | 1.95E-01 | | 5.46E-01 | 1.09E+00 | 1.19E-01 | 1.09E+00 |
| ENSG000002 | --- | 1.95E-01 | ? | | 4.87E-01 | -1.04E+00 | -2.05E+00 |
| ENSG000001 | IFI27L2 | 1.95E-01 | | 5.46E-01 | 1.10E+00 | 1.34E-01 | 1.10E+00 |
| ENSG000001 | TWIST1 | 1.95E-01 | ? | | 1.28E+00 | 3.56E-01 | 1.28E+00 |
| ENSG000002 | ZNF132-DT | 1.95E-01 | | 5.46E-01 | 1.24E+00 | 3.08E-01 | 1.24E+00 |
| ENSG000001 | BEGAIN | 1.95E-01 | ? | | 1.53E+00 | 6.12E-01 | 1.53E+00 |
| ENSG000001 | RFC3 | 1.95E-01 | | 5.46E-01 | 9.20E-01 | -1.21E-01 | -1.09E+00 |
| ENSG00000C | COL5A3 | 1.95E-01 | | 5.46E-01 | 1.16E+00 | 2.18E-01 | 1.16E+00 |

| | | | | | | |
|---|---|---|---|---|---|---|
| ENSG000001 | GRAMD2B | 1.95E-01 | 5.46E-01 | 9.50E-01 | -7.47E-02 | -1.05E+00 |
| ENSG000001 | ZMAT5 | 1.95E-01 | 5.46E-01 | 1.09E+00 | 1.26E-01 | 1.09E+00 |
| ENSG000001 | COLGALT2 | 1.95E-01 | ? | 1.87E+00 | 8.99E-01 | 1.87E+00 |
| ENSG000001 | QARS1 | 1.95E-01 | 5.46E-01 | 1.05E+00 | 6.85E-02 | 1.05E+00 |
| ENSG000002 | --- | 1.95E-01 | ? | 1.74E+00 | 8.02E-01 | 1.74E+00 |
| ENSG000002 | AGAP9 | 1.95E-01 | 5.46E-01 | 8.84E-01 | -1.78E-01 | -1.13E+00 |
| ENSG000002 | --- | 1.95E-01 | ? | 6.19E-01 | -6.92E-01 | -1.62E+00 |
| ENSG00000C | TACC3 | 1.95E-01 | 5.46E-01 | 8.90E-01 | -1.68E-01 | -1.12E+00 |
| ENSG000001 | GPER1 | 1.95E-01 | ? | 6.53E-01 | -6.15E-01 | -1.53E+00 |
| ENSG000001 | WRAP73 | 1.95E-01 | 5.46E-01 | 1.09E+00 | 1.25E-01 | 1.09E+00 |
| ENSG000001 | CPSF6 | 1.95E-01 | 5.46E-01 | 9.54E-01 | -6.80E-02 | -1.05E+00 |
| ENSG000001 | ELP6 | 1.95E-01 | 5.46E-01 | 1.06E+00 | 8.84E-02 | 1.06E+00 |
| ENSG000001 | PTPRO | 1.95E-01 | ? | 7.83E-01 | -3.53E-01 | -1.28E+00 |
| ENSG000001 | THOP1 | 1.95E-01 | 5.46E-01 | 1.07E+00 | 9.79E-02 | 1.07E+00 |
| ENSG000002 | PCOTH | 1.95E-01 | 5.46E-01 | 8.30E-01 | -2.69E-01 | -1.20E+00 |
| ENSG000002 | --- | 1.95E-01 | ? | 6.33E-01 | -6.59E-01 | -1.58E+00 |
| ENSG00000C | ANKS1A | 1.95E-01 | 5.46E-01 | 1.03E+00 | 4.63E-02 | 1.03E+00 |
| ENSG000001 | FXR1 | 1.95E-01 | 5.46E-01 | 9.59E-01 | -6.09E-02 | -1.04E+00 |
| ENSG000001 | NBEA | 1.95E-01 | 5.46E-01 | 1.16E+00 | 2.08E-01 | 1.16E+00 |
| ENSG000002 | --- | 1.96E-01 | ? | 1.51E+00 | 5.92E-01 | 1.51E+00 |
| ENSG000001 | FSTL1 | 1.96E-01 | 5.47E-01 | 9.60E-01 | -5.92E-02 | -1.04E+00 |
| ENSG00000C | MAP3K9 | 1.96E-01 | 5.47E-01 | 1.05E+00 | 6.97E-02 | 1.05E+00 |
| ENSG000002 | --- | 1.96E-01 | ? | 1.73E+00 | 7.91E-01 | 1.73E+00 |
| ENSG000001 | FZD3 | 1.96E-01 | 5.48E-01 | 9.37E-01 | -9.43E-02 | -1.07E+00 |
| ENSG000001 | PAOX | 1.96E-01 | 5.48E-01 | 1.30E+00 | 3.80E-01 | 1.30E+00 |
| ENSG000001 | SPPL2A | 1.96E-01 | 5.48E-01 | 1.04E+00 | 4.97E-02 | 1.04E+00 |
| ENSG000002 | --- | 1.96E-01 | ? | 6.14E-01 | -7.04E-01 | -1.63E+00 |
| ENSG000001 | CGN | 1.96E-01 | 5.48E-01 | 1.12E+00 | 1.59E-01 | 1.12E+00 |
| ENSG000002 | --- | 1.96E-01 | 5.48E-01 | 8.24E-01 | -2.79E-01 | -1.21E+00 |
| ENSG000002 | LINC02925 | 1.96E-01 | 5.48E-01 | 1.17E+00 | 2.27E-01 | 1.17E+00 |
| ENSG000001 | SFN | 1.96E-01 | 5.48E-01 | 1.06E+00 | 7.79E-02 | 1.06E+00 |
| ENSG000001 | TAF6 | 1.97E-01 | 5.48E-01 | 1.05E+00 | 7.26E-02 | 1.05E+00 |
| ENSG000002 | --- | 1.97E-01 | ? | 1.33E+00 | 4.16E-01 | 1.33E+00 |
| ENSG000002 | --- | 1.97E-01 | ? | 1.56E+00 | 6.38E-01 | 1.56E+00 |
| ENSG000002 | --- | 1.97E-01 | ? | 1.53E+00 | 6.12E-01 | 1.53E+00 |
| ENSG000001 | SLC9A3R1 | 1.97E-01 | 5.48E-01 | 9.43E-01 | -8.43E-02 | -1.06E+00 |
| ENSG000002 | RBM8B | 1.97E-01 | ? | 6.59E-01 | -6.01E-01 | -1.52E+00 |
| ENSG000001 | RAB3IL1 | 1.97E-01 | ? | 1.30E+00 | 3.76E-01 | 1.30E+00 |
| ENSG000001 | RPS6KA4 | 1.97E-01 | 5.48E-01 | 1.09E+00 | 1.23E-01 | 1.09E+00 |
| ENSG00000C | C5orf22 | 1.97E-01 | 5.48E-01 | 1.05E+00 | 7.63E-02 | 1.05E+00 |
| ENSG000001 | IP6K1 | 1.97E-01 | 5.48E-01 | 1.08E+00 | 1.05E-01 | 1.08E+00 |
| ENSG000001 | XBP1 | 1.97E-01 | 5.48E-01 | 9.64E-01 | -5.23E-02 | -1.04E+00 |
| ENSG000002 | --- | 1.97E-01 | 5.49E-01 | 8.14E-01 | -2.96E-01 | -1.23E+00 |
| ENSG000001 | C11orf71 | 1.97E-01 | 5.49E-01 | 1.20E+00 | 2.60E-01 | 1.20E+00 |
| ENSG000001 | ATG7 | 1.97E-01 | 5.49E-01 | 1.07E+00 | 9.41E-02 | 1.07E+00 |

| | | | | | | |
|---|---|---|---|---|---|---|
| ENSG000001 | REEP6 | 1.97E-01 | 5.49E-01 | 1.12E+00 | 1.60E-01 | 1.12E+00 |
| ENSG000002 | LINC02029 | 1.97E-01 | 5.49E-01 | 8.81E-01 | -1.83E-01 | -1.14E+00 |
| ENSG000002 | --- | 1.97E-01 | 5.49E-01 | 1.13E+00 | 1.78E-01 | 1.13E+00 |
| ENSG000001 | CMIP | 1.97E-01 | 5.49E-01 | 9.51E-01 | -7.22E-02 | -1.05E+00 |
| ENSG000002 | --- | 1.97E-01 | ? | 1.79E+00 | 8.42E-01 | 1.79E+00 |
| ENSG000001 | SORBS2 | 1.97E-01 | 5.49E-01 | 8.29E-01 | -2.70E-01 | -1.21E+00 |
| ENSG000001 | SEC14L1 | 1.97E-01 | 5.49E-01 | 1.07E+00 | 9.88E-02 | 1.07E+00 |
| ENSG000001 | LARGE1 | 1.97E-01 | 5.49E-01 | 1.07E+00 | 9.88E-02 | 1.07E+00 |
| ENSG00000C | KMT2C | 1.97E-01 | 5.49E-01 | 9.12E-01 | -1.33E-01 | -1.10E+00 |
| ENSG000002 | LDHAP3 | 1.97E-01 | ? | 1.53E+00 | 6.15E-01 | 1.53E+00 |
| ENSG000001 | TDRD9 | 1.98E-01 | ? | 1.36E+00 | 4.44E-01 | 1.36E+00 |
| ENSG000001 | TMEM117 | 1.98E-01 | 5.49E-01 | 1.10E+00 | 1.43E-01 | 1.10E+00 |
| ENSG000001 | NCALD | 1.98E-01 | ? | 7.73E-01 | -3.72E-01 | -1.29E+00 |
| ENSG000001 | STAT4 | 1.98E-01 | 5.49E-01 | 1.08E+00 | 1.13E-01 | 1.08E+00 |
| ENSG000002 | --- | 1.98E-01 | ? | 6.43E-01 | -6.37E-01 | -1.56E+00 |
| ENSG000002 | GAS6-DT | 1.98E-01 | 5.50E-01 | 1.10E+00 | 1.39E-01 | 1.10E+00 |
| ENSG000001 | KCNB1 | 1.98E-01 | ? | 6.31E-01 | -6.64E-01 | -1.58E+00 |
| ENSG000002 | CCDC144NL-/ | 1.98E-01 | 5.50E-01 | 1.27E+00 | 3.44E-01 | 1.27E+00 |
| ENSG000001 | BMX | 1.98E-01 | ? | 6.39E-01 | -6.46E-01 | -1.56E+00 |
| ENSG000001 | THAP1 | 1.98E-01 | 5.50E-01 | 9.42E-01 | -8.63E-02 | -1.06E+00 |
| ENSG00000C | JAK2 | 1.98E-01 | 5.50E-01 | 1.06E+00 | 8.63E-02 | 1.06E+00 |
| ENSG000001 | ATXN7L1 | 1.98E-01 | 5.50E-01 | 9.20E-01 | -1.20E-01 | -1.09E+00 |
| ENSG000002 | RPL7AP30 | 1.98E-01 | ? | 1.35E+00 | 4.37E-01 | 1.35E+00 |
| ENSG000001 | BRCC3 | 1.98E-01 | 5.50E-01 | 9.46E-01 | -8.03E-02 | -1.06E+00 |
| ENSG00000C | TTC39A | 1.98E-01 | 5.50E-01 | 1.25E+00 | 3.21E-01 | 1.25E+00 |
| ENSG000001 | NOL10 | 1.98E-01 | 5.50E-01 | 9.57E-01 | -6.28E-02 | -1.04E+00 |
| ENSG00000C | ACP3 | 1.98E-01 | ? | 1.36E+00 | 4.46E-01 | 1.36E+00 |
| ENSG000001 | HDAC10 | 1.98E-01 | 5.50E-01 | 1.13E+00 | 1.82E-01 | 1.13E+00 |
| ENSG000001 | ZNF793 | 1.98E-01 | 5.50E-01 | 9.33E-01 | -1.01E-01 | -1.07E+00 |
| ENSG000002 | ZNF691-DT | 1.98E-01 | 5.50E-01 | 8.33E-01 | -2.64E-01 | -1.20E+00 |
| ENSG000002 | NUTM2D | 1.98E-01 | 5.50E-01 | 1.10E+00 | 1.33E-01 | 1.10E+00 |
| ENSG000002 | FAM229B | 1.98E-01 | 5.50E-01 | 1.19E+00 | 2.47E-01 | 1.19E+00 |
| ENSG000002 | --- | 1.98E-01 | ? | 7.32E-01 | -4.50E-01 | -1.37E+00 |
| ENSG000001 | THYN1 | 1.98E-01 | 5.50E-01 | 1.07E+00 | 9.68E-02 | 1.07E+00 |
| ENSG000001 | PCED1B | 1.98E-01 | 5.50E-01 | 9.13E-01 | -1.32E-01 | -1.10E+00 |
| ENSG000001 | SLC66A2 | 1.99E-01 | 5.50E-01 | 1.10E+00 | 1.36E-01 | 1.10E+00 |
| ENSG000001 | GJA9 | 1.99E-01 | ? | 1.79E+00 | 8.41E-01 | 1.79E+00 |
| ENSG000001 | TMEM92 | 1.99E-01 | 5.50E-01 | 9.40E-01 | -8.97E-02 | -1.06E+00 |
| ENSG000001 | FAM53B | 1.99E-01 | 5.50E-01 | 1.08E+00 | 1.08E-01 | 1.08E+00 |
| ENSG000001 | TNIP1 | 1.99E-01 | 5.51E-01 | 1.03E+00 | 4.92E-02 | 1.03E+00 |
| ENSG000001 | TAF8 | 1.99E-01 | 5.51E-01 | 1.04E+00 | 5.66E-02 | 1.04E+00 |
| ENSG000002 | FOXD1-AS1 | 1.99E-01 | 5.51E-01 | 1.21E+00 | 2.69E-01 | 1.21E+00 |
| ENSG000002 | PSG5 | 1.99E-01 | 5.51E-01 | 1.24E+00 | 3.07E-01 | 1.24E+00 |
| ENSG000002 | --- | 1.99E-01 | 5.51E-01 | 1.19E+00 | 2.48E-01 | 1.19E+00 |
| ENSG000002 | --- | 1.99E-01 | ? | 6.45E-01 | -6.32E-01 | -1.55E+00 |

| | | | | | | |
|---|---|---|---|---|---|---|
| ENSG000001 | IRGQ | 1.99E-01 | 5.51E-01 | 9.62E-01 | -5.65E-02 | -1.04E+00 |
| ENSG000001 | FBXO31 | 1.99E-01 | 5.51E-01 | 1.05E+00 | 7.56E-02 | 1.05E+00 |
| ENSG000001 | AMZ2 | 1.99E-01 | 5.51E-01 | 9.55E-01 | -6.60E-02 | -1.05E+00 |
| ENSG000001 | PCNX3 | 1.99E-01 | 5.52E-01 | 1.08E+00 | 1.07E-01 | 1.08E+00 |
| ENSG000001 | C7orf25 | 1.99E-01 | 5.52E-01 | 8.56E-01 | -2.25E-01 | -1.17E+00 |
| ENSG000002 | ADPGK-AS1 | 2.00E-01 | ? | 6.11E-01 | -7.10E-01 | -1.64E+00 |
| ENSG000001 | CXCR1 | 2.00E-01 | ? | 7.36E-01 | -4.42E-01 | -1.36E+00 |
| ENSG00000C | NEBL | 2.00E-01 | 5.52E-01 | 1.12E+00 | 1.60E-01 | 1.12E+00 |
| ENSG000002 | --- | 2.00E-01 | ? | 5.94E-01 | -7.51E-01 | -1.68E+00 |
| ENSG000001 | C1orf54 | 2.00E-01 | 5.52E-01 | 1.20E+00 | 2.67E-01 | 1.20E+00 |
| ENSG000001 | RMDN1 | 2.00E-01 | 5.52E-01 | 9.49E-01 | -7.55E-02 | -1.05E+00 |
| ENSG000002 | XBP1P1 | 2.00E-01 | ? | 6.51E-01 | -6.20E-01 | -1.54E+00 |
| ENSG000001 | WDR25 | 2.00E-01 | 5.52E-01 | 1.09E+00 | 1.29E-01 | 1.09E+00 |
| ENSG000001 | ATP6V1G1 | 2.00E-01 | 5.53E-01 | 9.50E-01 | -7.43E-02 | -1.05E+00 |
| ENSG000001 | MAPK10 | 2.00E-01 | 5.53E-01 | 1.11E+00 | 1.57E-01 | 1.11E+00 |
| ENSG000001 | SEC61G | 2.00E-01 | 5.53E-01 | 9.50E-01 | -7.36E-02 | -1.05E+00 |
| ENSG000001 | TMEM14C | 2.00E-01 | 5.53E-01 | 1.06E+00 | 8.09E-02 | 1.06E+00 |
| ENSG000001 | TRMT44 | 2.00E-01 | 5.53E-01 | 1.08E+00 | 1.05E-01 | 1.08E+00 |
| ENSG000002 | NBPF14 | 2.00E-01 | 5.53E-01 | 9.25E-01 | -1.12E-01 | -1.08E+00 |
| ENSG000002 | --- | 2.00E-01 | 5.53E-01 | 8.36E-01 | -2.59E-01 | -1.20E+00 |
| ENSG000001 | BTG3 | 2.00E-01 | 5.53E-01 | 1.06E+00 | 8.45E-02 | 1.06E+00 |
| ENSG000001 | ASXL2 | 2.00E-01 | 5.53E-01 | 9.51E-01 | -7.27E-02 | -1.05E+00 |
| ENSG00000C | TP73 | 2.00E-01 | ? | 1.80E+00 | 8.45E-01 | 1.80E+00 |
| ENSG000002 | --- | 2.01E-01 | ? | 1.58E+00 | 6.63E-01 | 1.58E+00 |
| ENSG00000C | DYNC1I2 | 2.01E-01 | 5.53E-01 | 1.03E+00 | 3.75E-02 | 1.03E+00 |
| ENSG000001 | GSE1 | 2.01E-01 | 5.53E-01 | 9.51E-01 | -7.28E-02 | -1.05E+00 |
| ENSG000002 | KIAA0040 | 2.01E-01 | 5.53E-01 | 1.07E+00 | 1.01E-01 | 1.07E+00 |
| ENSG000002 | --- | 2.01E-01 | 5.53E-01 | 9.29E-01 | -1.07E-01 | -1.08E+00 |
| ENSG000002 | --- | 2.01E-01 | ? | 7.62E-01 | -3.93E-01 | -1.31E+00 |
| ENSG000001 | AFG1L | 2.01E-01 | 5.53E-01 | 9.23E-01 | -1.15E-01 | -1.08E+00 |
| ENSG00000C | CDK17 | 2.01E-01 | 5.53E-01 | 9.44E-01 | -8.32E-02 | -1.06E+00 |
| ENSG000002 | LINC01012 | 2.01E-01 | 5.54E-01 | 1.18E+00 | 2.36E-01 | 1.18E+00 |
| ENSG000002 | LSM12P1 | 2.01E-01 | 5.54E-01 | 1.13E+00 | 1.76E-01 | 1.13E+00 |
| ENSG000002 | RNF32-AS1 | 2.01E-01 | ? | 7.24E-01 | -4.67E-01 | -1.38E+00 |
| ENSG000001 | GUCA1B | 2.01E-01 | 5.54E-01 | 8.26E-01 | -2.76E-01 | -1.21E+00 |
| ENSG000001 | FOSB | 2.01E-01 | ? | 7.92E-01 | -3.36E-01 | -1.26E+00 |
| ENSG000001 | KLRG2 | 2.01E-01 | ? | 1.24E+00 | 3.12E-01 | 1.24E+00 |
| ENSG000001 | PEAK1 | 2.01E-01 | 5.54E-01 | 9.11E-01 | -1.34E-01 | -1.10E+00 |
| ENSG000001 | SRP68 | 2.01E-01 | 5.54E-01 | 1.03E+00 | 3.84E-02 | 1.03E+00 |
| ENSG000001 | LCE3D | 2.01E-01 | ? | 1.41E+00 | 4.94E-01 | 1.41E+00 |
| ENSG000002 | DYNLT2B | 2.01E-01 | 5.54E-01 | 1.08E+00 | 1.06E-01 | 1.08E+00 |
| ENSG000002 | --- | 2.01E-01 | 5.54E-01 | 1.11E+00 | 1.46E-01 | 1.11E+00 |
| ENSG000001 | KCTD16 | 2.01E-01 | ? | 1.39E+00 | 4.76E-01 | 1.39E+00 |
| ENSG000001 | TNRC18 | 2.01E-01 | 5.54E-01 | 1.07E+00 | 9.22E-02 | 1.07E+00 |
| ENSG000001 | ADAMTSL4 | 2.01E-01 | 5.54E-01 | 1.10E+00 | 1.42E-01 | 1.10E+00 |

| | | | | | | |
|---|---|---|---|---|---|---|
| ENSG00000C | BAD | 2.01E-01 | 5.54E-01 | 1.14E+00 | 1.94E-01 | 1.14E+00 |
| ENSG000002 | MIR4453HG | 2.01E-01 | 5.54E-01 | 8.95E-01 | -1.60E-01 | -1.12E+00 |
| ENSG000001 | ZFPL1 | 2.02E-01 | 5.54E-01 | 9.40E-01 | -8.90E-02 | -1.06E+00 |
| ENSG000001 | TICRR | 2.02E-01 | 5.54E-01 | 9.07E-01 | -1.41E-01 | -1.10E+00 |
| ENSG000001 | YEATS2 | 2.02E-01 | 5.54E-01 | 9.68E-01 | -4.68E-02 | -1.03E+00 |
| ENSG000002 | --- | 2.02E-01 | ? | 1.65E+00 | 7.25E-01 | 1.65E+00 |
| ENSG000002 | MIOS-DT | 2.02E-01 | 5.54E-01 | 8.10E-01 | -3.04E-01 | -1.23E+00 |
| ENSG00000C | CTNNA1 | 2.02E-01 | 5.54E-01 | 1.03E+00 | 3.70E-02 | 1.03E+00 |
| ENSG000001 | LRGUK | 2.02E-01 | ? | 1.48E+00 | 5.70E-01 | 1.48E+00 |
| ENSG000001 | DYNLT4 | 2.02E-01 | 5.55E-01 | 8.75E-01 | -1.93E-01 | -1.14E+00 |
| ENSG000001 | WWC2 | 2.02E-01 | 5.55E-01 | 9.54E-01 | -6.82E-02 | -1.05E+00 |
| ENSG000002 | CSKMT | 2.02E-01 | 5.55E-01 | 1.13E+00 | 1.76E-01 | 1.13E+00 |
| ENSG00000C | IKZF2 | 2.02E-01 | 5.55E-01 | 1.08E+00 | 1.14E-01 | 1.08E+00 |
| ENSG000002 | TTC23L | 2.02E-01 | ? | 1.75E+00 | 8.09E-01 | 1.75E+00 |
| ENSG000001 | NDUFS6 | 2.02E-01 | 5.55E-01 | 1.04E+00 | 6.12E-02 | 1.04E+00 |
| ENSG000002 | SLFN12L | 2.03E-01 | 5.55E-01 | 8.68E-01 | -2.04E-01 | -1.15E+00 |
| ENSG000001 | TRPV6 | 2.03E-01 | ? | 1.39E+00 | 4.71E-01 | 1.39E+00 |
| ENSG000001 | HCAR2 | 2.03E-01 | 5.55E-01 | 9.30E-01 | -1.05E-01 | -1.08E+00 |
| ENSG000001 | PPP3CC | 2.03E-01 | 5.55E-01 | 9.52E-01 | -7.12E-02 | -1.05E+00 |
| ENSG000001 | VNN1 | 2.03E-01 | ? | 7.63E-01 | -3.91E-01 | -1.31E+00 |
| ENSG000001 | MIGA2 | 2.03E-01 | 5.56E-01 | 1.10E+00 | 1.40E-01 | 1.10E+00 |
| ENSG000001 | LRRIQ3 | 2.03E-01 | 5.56E-01 | 1.17E+00 | 2.24E-01 | 1.17E+00 |
| ENSG000001 | SIL1 | 2.03E-01 | 5.56E-01 | 1.05E+00 | 6.82E-02 | 1.05E+00 |
| ENSG00000C | NCOA1 | 2.03E-01 | 5.56E-01 | 9.41E-01 | -8.70E-02 | -1.06E+00 |
| ENSG000001 | FHIP1A | 2.03E-01 | 5.56E-01 | 1.07E+00 | 1.03E-01 | 1.07E+00 |
| ENSG000001 | FIBP | 2.03E-01 | 5.56E-01 | 1.05E+00 | 7.20E-02 | 1.05E+00 |
| ENSG00000C | ENTPD2 | 2.03E-01 | 5.56E-01 | 1.23E+00 | 2.96E-01 | 1.23E+00 |
| ENSG000001 | MED13 | 2.03E-01 | 5.56E-01 | 9.49E-01 | -7.52E-02 | -1.05E+00 |
| ENSG000001 | DCDC2 | 2.03E-01 | ? | 7.93E-01 | -3.34E-01 | -1.26E+00 |
| ENSG000001 | FAM20C | 2.03E-01 | 5.56E-01 | 9.06E-01 | -1.42E-01 | -1.10E+00 |
| ENSG000002 | --- | 2.03E-01 | ? | 7.65E-01 | -3.86E-01 | -1.31E+00 |
| ENSG000001 | THAP4 | 2.03E-01 | 5.56E-01 | 1.05E+00 | 7.46E-02 | 1.05E+00 |
| ENSG000001 | CNTNAP3B | 2.03E-01 | 5.56E-01 | 8.88E-01 | -1.72E-01 | -1.13E+00 |
| ENSG000001 | GSPT1 | 2.03E-01 | 5.56E-01 | 9.57E-01 | -6.34E-02 | -1.04E+00 |
| ENSG000002 | --- | 2.03E-01 | 5.56E-01 | 9.10E-01 | -1.36E-01 | -1.10E+00 |
| ENSG000001 | MDM2 | 2.03E-01 | 5.56E-01 | 9.55E-01 | -6.65E-02 | -1.05E+00 |
| ENSG000001 | WDR77 | 2.03E-01 | 5.56E-01 | 1.05E+00 | 7.68E-02 | 1.05E+00 |
| ENSG000002 | --- | 2.04E-01 | 5.56E-01 | 1.31E+00 | 3.89E-01 | 1.31E+00 |
| ENSG000002 | SLC35G6 | 2.04E-01 | ? | 1.57E+00 | 6.53E-01 | 1.57E+00 |
| ENSG000001 | SLC22A4 | 2.04E-01 | ? | 1.27E+00 | 3.45E-01 | 1.27E+00 |
| ENSG000001 | RAD51D | 2.04E-01 | 5.56E-01 | 9.19E-01 | -1.22E-01 | -1.09E+00 |
| ENSG000001 | TTC30A | 2.04E-01 | 5.56E-01 | 1.07E+00 | 9.21E-02 | 1.07E+00 |
| ENSG000002 | RASL10B | 2.04E-01 | ? | 1.83E+00 | 8.70E-01 | 1.83E+00 |
| ENSG000002 | --- | 2.04E-01 | 5.56E-01 | 1.13E+00 | 1.72E-01 | 1.13E+00 |
| ENSG000001 | IRF1-AS1 | 2.04E-01 | ? | 1.26E+00 | 3.28E-01 | 1.26E+00 |

| | | | | | | | |
|---|---|---|---|---|---|---|---|
| ENSG000001 | GEMIN7 | 2.04E-01 | | 5.56E-01 | 1.09E+00 | 1.20E-01 | 1.09E+00 |
| ENSG000001 | AKAP9 | 2.04E-01 | | 5.56E-01 | 9.21E-01 | -1.18E-01 | -1.09E+00 |
| ENSG000001 | APBA1 | 2.04E-01 | | 5.56E-01 | 1.12E+00 | 1.67E-01 | 1.12E+00 |
| ENSG000001 | IRF2BP1 | 2.04E-01 | | 5.56E-01 | 1.13E+00 | 1.72E-01 | 1.13E+00 |
| ENSG000002 | --- | 2.04E-01 | | 5.56E-01 | 1.24E+00 | 3.11E-01 | 1.24E+00 |
| ENSG000001 | THOC6 | 2.04E-01 | | 5.56E-01 | 1.10E+00 | 1.36E-01 | 1.10E+00 |
| ENSG000001 | CASD1 | 2.04E-01 | | 5.56E-01 | 1.06E+00 | 8.29E-02 | 1.06E+00 |
| ENSG000002 | MTND2P28 | 2.04E-01 | | 5.57E-01 | 8.48E-01 | -2.39E-01 | -1.18E+00 |
| ENSG000001 | DHX37 | 2.04E-01 | | 5.57E-01 | 9.28E-01 | -1.08E-01 | -1.08E+00 |
| ENSG00000C | GAS7 | 2.04E-01 | ? | | 1.46E+00 | 5.47E-01 | 1.46E+00 |
| ENSG00000C | GMCL1 | 2.04E-01 | | 5.58E-01 | 9.39E-01 | -9.08E-02 | -1.06E+00 |
| ENSG000001 | PKD1L2 | 2.05E-01 | ? | | 6.12E-01 | -7.08E-01 | -1.63E+00 |
| ENSG000001 | TGM4 | 2.05E-01 | | 5.58E-01 | 8.96E-01 | -1.59E-01 | -1.12E+00 |
| ENSG000001 | TIFA | 2.05E-01 | | 5.58E-01 | 1.08E+00 | 1.10E-01 | 1.08E+00 |
| ENSG000001 | HGSNAT | 2.05E-01 | | 5.58E-01 | 1.04E+00 | 5.93E-02 | 1.04E+00 |
| ENSG000001 | MICOS13 | 2.05E-01 | | 5.58E-01 | 1.10E+00 | 1.38E-01 | 1.10E+00 |
| ENSG000002 | LINC01301 | 2.05E-01 | ? | | 7.47E-01 | -4.20E-01 | -1.34E+00 |
| ENSG000001 | PRNP | 2.05E-01 | | 5.58E-01 | 1.03E+00 | 4.85E-02 | 1.03E+00 |
| ENSG000002 | LINC02604 | 2.05E-01 | | 5.58E-01 | 8.79E-01 | -1.87E-01 | -1.14E+00 |
| ENSG000001 | GCNT1 | 2.05E-01 | | 5.58E-01 | 1.06E+00 | 9.06E-02 | 1.06E+00 |
| ENSG000001 | STYX | 2.05E-01 | | 5.58E-01 | 9.42E-01 | -8.61E-02 | -1.06E+00 |
| ENSG000001 | MRPL14 | 2.05E-01 | | 5.59E-01 | 1.08E+00 | 1.07E-01 | 1.08E+00 |
| ENSG000001 | LNPEP | 2.05E-01 | | 5.59E-01 | 1.07E+00 | 9.59E-02 | 1.07E+00 |
| ENSG000002 | --- | 2.05E-01 | ? | | 6.36E-01 | -6.54E-01 | -1.57E+00 |
| ENSG00000C | MAEA | 2.05E-01 | | 5.59E-01 | 1.04E+00 | 5.58E-02 | 1.04E+00 |
| ENSG000001 | IPO13 | 2.05E-01 | | 5.59E-01 | 1.06E+00 | 7.87E-02 | 1.06E+00 |
| ENSG000001 | PDCD4 | 2.05E-01 | | 5.59E-01 | 9.37E-01 | -9.46E-02 | -1.07E+00 |
| ENSG000001 | NAA10 | 2.05E-01 | | 5.59E-01 | 1.05E+00 | 6.41E-02 | 1.05E+00 |
| ENSG000001 | ACBD5 | 2.06E-01 | | 5.59E-01 | 9.52E-01 | -7.14E-02 | -1.05E+00 |
| ENSG000001 | GBP6 | 2.06E-01 | | 5.59E-01 | 9.43E-01 | -8.39E-02 | -1.06E+00 |
| ENSG000002 | LBX2-AS1 | 2.06E-01 | | 5.59E-01 | 8.13E-01 | -2.99E-01 | -1.23E+00 |
| ENSG00000C | RWDD2A | 2.06E-01 | | 5.59E-01 | 1.07E+00 | 1.04E-01 | 1.07E+00 |
| ENSG000001 | SPIN4 | 2.06E-01 | | 5.59E-01 | 9.03E-01 | -1.47E-01 | -1.11E+00 |
| ENSG00000C | ANKIB1 | 2.06E-01 | | 5.59E-01 | 9.67E-01 | -4.88E-02 | -1.03E+00 |
| ENSG00000C | ESF1 | 2.06E-01 | | 5.59E-01 | 9.37E-01 | -9.34E-02 | -1.07E+00 |
| ENSG000002 | CLIC1 | 2.06E-01 | | 5.59E-01 | 1.04E+00 | 5.32E-02 | 1.04E+00 |
| ENSG000002 | --- | 2.06E-01 | ? | | 1.67E+00 | 7.40E-01 | 1.67E+00 |
| ENSG000001 | NDUFS4 | 2.06E-01 | | 5.59E-01 | 9.53E-01 | -6.96E-02 | -1.05E+00 |
| ENSG000001 | ADAMTS19 | 2.06E-01 | ? | | 1.29E+00 | 3.68E-01 | 1.29E+00 |
| ENSG000002 | --- | 2.06E-01 | | 5.59E-01 | 7.84E-01 | -3.51E-01 | -1.28E+00 |
| ENSG000001 | PPP1R1C | 2.06E-01 | | 5.59E-01 | 8.70E-01 | -2.01E-01 | -1.15E+00 |
| ENSG00000C | TFAP2C | 2.06E-01 | | 5.59E-01 | 9.06E-01 | -1.43E-01 | -1.10E+00 |
| ENSG000001 | HIC1 | 2.06E-01 | | 5.59E-01 | 1.15E+00 | 2.04E-01 | 1.15E+00 |
| ENSG000002 | TMEM240 | 2.06E-01 | ? | | 7.73E-01 | -3.72E-01 | -1.29E+00 |
| ENSG000001 | FOXA1 | 2.06E-01 | | 5.59E-01 | 8.89E-01 | -1.70E-01 | -1.12E+00 |

| | | | | | | |
|---|---|---|---|---|---|---|
| ENSG000002 | OST4 | 2.06E-01 | 5.59E-01 | 1.04E+00 | 4.99E-02 | 1.04E+00 |
| ENSG000001 | AIM2 | 2.06E-01 | ? | 6.06E-01 | -7.23E-01 | -1.65E+00 |
| ENSG000001 | ATF6 | 2.06E-01 | 5.59E-01 | 9.56E-01 | -6.45E-02 | -1.05E+00 |
| ENSG000002 | --- | 2.06E-01 | 5.59E-01 | 8.96E-01 | -1.58E-01 | -1.12E+00 |
| ENSG000002 | UGT1A4 | 2.06E-01 | ? | 1.34E+00 | 4.22E-01 | 1.34E+00 |
| ENSG000001 | UTP11 | 2.06E-01 | 5.59E-01 | 9.43E-01 | -8.49E-02 | -1.06E+00 |
| ENSG000001 | RAB43 | 2.06E-01 | 5.59E-01 | 1.17E+00 | 2.28E-01 | 1.17E+00 |
| ENSG000001 | MYRIP | 2.06E-01 | ? | 5.41E-01 | -8.88E-01 | -1.85E+00 |
| ENSG00000C | NUB1 | 2.07E-01 | 5.59E-01 | 9.57E-01 | -6.27E-02 | -1.04E+00 |
| ENSG000001 | ZNF185 | 2.07E-01 | 5.59E-01 | 9.65E-01 | -5.20E-02 | -1.04E+00 |
| ENSG00000C | ACADVL | 2.07E-01 | 5.59E-01 | 1.07E+00 | 9.92E-02 | 1.07E+00 |
| ENSG000002 | NQO2-AS1 | 2.07E-01 | ? | 2.05E+00 | 1.03E+00 | 2.05E+00 |
| ENSG000002 | BDNF-AS | 2.07E-01 | 5.59E-01 | 1.20E+00 | 2.58E-01 | 1.20E+00 |
| ENSG000002 | --- | 2.07E-01 | ? | 7.11E-01 | -4.91E-01 | -1.41E+00 |
| ENSG000001 | HERC4 | 2.07E-01 | 5.59E-01 | 9.69E-01 | -4.48E-02 | -1.03E+00 |
| ENSG000001 | CTBP1-DT | 2.07E-01 | 5.59E-01 | 1.10E+00 | 1.32E-01 | 1.10E+00 |
| ENSG000001 | ARMC7 | 2.07E-01 | 5.59E-01 | 9.18E-01 | -1.23E-01 | -1.09E+00 |
| ENSG000001 | ADNP | 2.07E-01 | 5.59E-01 | 9.57E-01 | -6.33E-02 | -1.04E+00 |
| ENSG000001 | PRPF4B | 2.07E-01 | 5.59E-01 | 9.51E-01 | -7.20E-02 | -1.05E+00 |
| ENSG000001 | SERTAD1 | 2.07E-01 | 5.59E-01 | 1.12E+00 | 1.66E-01 | 1.12E+00 |
| ENSG000002 | RPL13P5 | 2.07E-01 | 5.59E-01 | 8.07E-01 | -3.10E-01 | -1.24E+00 |
| ENSG000001 | CCDC186 | 2.07E-01 | 5.59E-01 | 9.33E-01 | -1.01E-01 | -1.07E+00 |
| ENSG000001 | CTDSP2 | 2.07E-01 | 5.59E-01 | 1.07E+00 | 1.04E-01 | 1.07E+00 |
| ENSG00000C | AKR7A2 | 2.07E-01 | 5.59E-01 | 1.06E+00 | 8.01E-02 | 1.06E+00 |
| ENSG000002 | TEX52 | 2.07E-01 | ? | 1.55E+00 | 6.29E-01 | 1.55E+00 |
| ENSG000001 | SOS1 | 2.07E-01 | 5.60E-01 | 9.45E-01 | -8.14E-02 | -1.06E+00 |
| ENSG000001 | LRRC37B | 2.07E-01 | 5.60E-01 | 8.98E-01 | -1.55E-01 | -1.11E+00 |
| ENSG000001 | CUEDC1 | 2.07E-01 | 5.60E-01 | 1.05E+00 | 7.51E-02 | 1.05E+00 |
| ENSG000001 | AASDHPPT | 2.07E-01 | 5.60E-01 | 1.05E+00 | 6.43E-02 | 1.05E+00 |
| ENSG000002 | --- | 2.07E-01 | 5.60E-01 | 1.21E+00 | 2.75E-01 | 1.21E+00 |
| ENSG000001 | LCMT2 | 2.07E-01 | 5.60E-01 | 1.07E+00 | 9.90E-02 | 1.07E+00 |
| ENSG000002 | USP32P2 | 2.07E-01 | ? | 1.94E+00 | 9.54E-01 | 1.94E+00 |
| ENSG000001 | PTPRS | 2.07E-01 | 5.60E-01 | 1.09E+00 | 1.20E-01 | 1.09E+00 |
| ENSG000002 | BISPR | 2.07E-01 | ? | 2.09E+00 | 1.06E+00 | 2.09E+00 |
| ENSG000001 | TRMT1L | 2.07E-01 | 5.60E-01 | 1.04E+00 | 5.92E-02 | 1.04E+00 |
| ENSG00000C | VDAC1P1 | 2.07E-01 | ? | 7.49E-01 | -4.18E-01 | -1.34E+00 |
| ENSG000002 | GPR199P | 2.08E-01 | ? | 1.72E+00 | 7.79E-01 | 1.72E+00 |
| ENSG000001 | KIFC2 | 2.08E-01 | 5.60E-01 | 1.08E+00 | 1.16E-01 | 1.08E+00 |
| ENSG000002 | LINC00888 | 2.08E-01 | 5.60E-01 | 9.00E-01 | -1.52E-01 | -1.11E+00 |
| ENSG000001 | DALRD3 | 2.08E-01 | 5.60E-01 | 1.09E+00 | 1.21E-01 | 1.09E+00 |
| ENSG000002 | --- | 2.08E-01 | ? | 1.46E+00 | 5.43E-01 | 1.46E+00 |
| ENSG000001 | GPR37L1 | 2.08E-01 | ? | 1.23E+00 | 3.00E-01 | 1.23E+00 |
| ENSG000001 | TTLL5 | 2.08E-01 | 5.61E-01 | 9.45E-01 | -8.11E-02 | -1.06E+00 |
| ENSG000001 | RFTN1 | 2.08E-01 | 5.61E-01 | 1.03E+00 | 4.73E-02 | 1.03E+00 |
| ENSG000001 | DCLK1 | 2.08E-01 | ? | 1.34E+00 | 4.17E-01 | 1.34E+00 |

| | | | | | | |
|---|---|---|---|---|---|---|
| ENSG000001 | C8orf76 | 2.08E-01 | 5.61E-01 | 9.46E-01 | -7.96E-02 | -1.06E+00 |
| ENSG000002 | --- | 2.08E-01 | ? | 1.70E+00 | 7.67E-01 | 1.70E+00 |
| ENSG000001 | ATAT1 | 2.08E-01 | 5.61E-01 | 9.28E-01 | -1.08E-01 | -1.08E+00 |
| ENSG000001 | TAS1R3 | 2.08E-01 | ? | 7.36E-01 | -4.42E-01 | -1.36E+00 |
| ENSG000002 | CRIM1-DT | 2.08E-01 | ? | 1.37E+00 | 4.51E-01 | 1.37E+00 |
| ENSG00000C | ALDH3A2 | 2.08E-01 | 5.61E-01 | 9.67E-01 | -4.77E-02 | -1.03E+00 |
| ENSG000001 | NEDD8 | 2.08E-01 | 5.61E-01 | 1.04E+00 | 5.45E-02 | 1.04E+00 |
| ENSG000002 | LCE1F | 2.08E-01 | ? | 1.50E+00 | 5.89E-01 | 1.50E+00 |
| ENSG000002 | TLCD4-RWDI | 2.08E-01 | ? | 1.31E+00 | 3.86E-01 | 1.31E+00 |
| ENSG000001 | FAM193A | 2.09E-01 | 5.61E-01 | 9.51E-01 | -7.20E-02 | -1.05E+00 |
| ENSG000002 | --- | 2.09E-01 | 5.61E-01 | 1.19E+00 | 2.54E-01 | 1.19E+00 |
| ENSG000001 | SPAG16 | 2.09E-01 | 5.61E-01 | 9.20E-01 | -1.21E-01 | -1.09E+00 |
| ENSG00000C | CYTH3 | 2.09E-01 | 5.61E-01 | 1.07E+00 | 9.74E-02 | 1.07E+00 |
| ENSG000001 | GAREM2 | 2.09E-01 | 5.61E-01 | 1.18E+00 | 2.43E-01 | 1.18E+00 |
| ENSG000002 | SELENOP | 2.09E-01 | 5.62E-01 | 8.22E-01 | -2.83E-01 | -1.22E+00 |
| ENSG000002 | --- | 2.09E-01 | 5.62E-01 | 9.01E-01 | -1.50E-01 | -1.11E+00 |
| ENSG000001 | VSIR | 2.09E-01 | 5.62E-01 | 1.13E+00 | 1.81E-01 | 1.13E+00 |
| ENSG00000C | LZTR1 | 2.09E-01 | 5.62E-01 | 1.07E+00 | 9.68E-02 | 1.07E+00 |
| ENSG000001 | ANKRD17 | 2.09E-01 | 5.62E-01 | 9.49E-01 | -7.50E-02 | -1.05E+00 |
| ENSG000001 | RNF168 | 2.09E-01 | 5.62E-01 | 1.05E+00 | 7.54E-02 | 1.05E+00 |
| ENSG000001 | C2orf50 | 2.09E-01 | ? | 7.86E-01 | -3.48E-01 | -1.27E+00 |
| ENSG000001 | GPS2 | 2.09E-01 | 5.62E-01 | 9.23E-01 | -1.16E-01 | -1.08E+00 |
| ENSG000001 | CYP4F2 | 2.09E-01 | ? | 4.98E-01 | -1.01E+00 | -2.01E+00 |
| ENSG000002 | --- | 2.09E-01 | 5.63E-01 | 7.93E-01 | -3.34E-01 | -1.26E+00 |
| ENSG000001 | POLR1E | 2.10E-01 | 5.63E-01 | 1.06E+00 | 8.19E-02 | 1.06E+00 |
| ENSG000001 | AKTIP | 2.10E-01 | 5.63E-01 | 1.06E+00 | 8.74E-02 | 1.06E+00 |
| ENSG000001 | JUND | 2.10E-01 | 5.63E-01 | 1.08E+00 | 1.13E-01 | 1.08E+00 |
| ENSG000001 | TMEM8B | 2.10E-01 | 5.63E-01 | 1.09E+00 | 1.20E-01 | 1.09E+00 |
| ENSG000001 | TMTC4 | 2.10E-01 | 5.63E-01 | 1.07E+00 | 1.02E-01 | 1.07E+00 |
| ENSG00000C | SEMA3B | 2.10E-01 | 5.63E-01 | 8.79E-01 | -1.87E-01 | -1.14E+00 |
| ENSG00000C | SNX11 | 2.10E-01 | 5.63E-01 | 1.07E+00 | 9.96E-02 | 1.07E+00 |
| ENSG000002 | --- | 2.10E-01 | ? | 6.65E-01 | -5.89E-01 | -1.50E+00 |
| ENSG000002 | ZFHX3-AS1 | 2.10E-01 | 5.63E-01 | 8.84E-01 | -1.78E-01 | -1.13E+00 |
| ENSG000001 | PRR19 | 2.10E-01 | 5.63E-01 | 1.18E+00 | 2.40E-01 | 1.18E+00 |
| ENSG000002 | TOLLIP-DT | 2.10E-01 | ? | 5.47E-01 | -8.71E-01 | -1.83E+00 |
| ENSG000001 | ESPL1 | 2.10E-01 | 5.63E-01 | 8.13E-01 | -2.99E-01 | -1.23E+00 |
| ENSG00000C | NOP14 | 2.10E-01 | 5.63E-01 | 1.05E+00 | 6.38E-02 | 1.05E+00 |
| ENSG000001 | DCLRE1A | 2.10E-01 | 5.63E-01 | 9.48E-01 | -7.76E-02 | -1.06E+00 |
| ENSG00000C | WDR70 | 2.10E-01 | 5.63E-01 | 1.05E+00 | 6.86E-02 | 1.05E+00 |
| ENSG000001 | MFSD8 | 2.10E-01 | 5.63E-01 | 9.47E-01 | -7.80E-02 | -1.06E+00 |
| ENSG00000C | AGK | 2.10E-01 | 5.63E-01 | 9.47E-01 | -7.89E-02 | -1.06E+00 |
| ENSG000002 | UBAP1L | 2.10E-01 | 5.63E-01 | 8.62E-01 | -2.15E-01 | -1.16E+00 |
| ENSG000002 | TMEM161B- | 2.10E-01 | 5.63E-01 | 8.93E-01 | -1.63E-01 | -1.12E+00 |
| ENSG000002 | --- | 2.10E-01 | ? | 7.52E-01 | -4.11E-01 | -1.33E+00 |
| ENSG000001 | GRIN2D | 2.10E-01 | ? | 7.78E-01 | -3.61E-01 | -1.28E+00 |

| | | | | | | |
|---|---|---|---|---|---|---|
| ENSG000001 | RNF139 | 2.10E-01 | 5.63E-01 | 1.04E+00 | 5.91E-02 | 1.04E+00 |
| ENSG00000( | ZNF586 | 2.11E-01 | 5.64E-01 | 1.08E+00 | 1.11E-01 | 1.08E+00 |
| ENSG000001 | ST14 | 2.11E-01 | 5.64E-01 | 1.06E+00 | 8.52E-02 | 1.06E+00 |
| ENSG000001 | C14orf39 | 2.11E-01 | 5.64E-01 | 1.15E+00 | 1.99E-01 | 1.15E+00 |
| ENSG000002 | --- | 2.11E-01 | ? | 5.38E-01 | -8.95E-01 | -1.86E+00 |
| ENSG000002 | RAB11B-AS1 | 2.11E-01 | 5.64E-01 | 1.15E+00 | 1.96E-01 | 1.15E+00 |
| ENSG000001 | RAPGEFL1 | 2.11E-01 | 5.64E-01 | 1.08E+00 | 1.13E-01 | 1.08E+00 |
| ENSG000001 | TMCO3 | 2.11E-01 | 5.64E-01 | 1.04E+00 | 5.18E-02 | 1.04E+00 |
| ENSG000001 | TNS2 | 2.11E-01 | 5.64E-01 | 9.20E-01 | -1.20E-01 | -1.09E+00 |
| ENSG000002 | NEURL4 | 2.11E-01 | 5.64E-01 | 1.07E+00 | 1.01E-01 | 1.07E+00 |
| ENSG000001 | C2CD4C | 2.11E-01 | ? | 1.30E+00 | 3.79E-01 | 1.30E+00 |
| ENSG000001 | CCDC14 | 2.11E-01 | 5.64E-01 | 9.29E-01 | -1.06E-01 | -1.08E+00 |
| ENSG000002 | CTNS-AS1 | 2.11E-01 | 5.64E-01 | 8.09E-01 | -3.06E-01 | -1.24E+00 |
| ENSG000001 | SIMC1 | 2.11E-01 | 5.64E-01 | 1.07E+00 | 9.10E-02 | 1.07E+00 |
| ENSG000002 | --- | 2.11E-01 | 5.64E-01 | 1.12E+00 | 1.59E-01 | 1.12E+00 |
| ENSG000001 | MOGS | 2.11E-01 | 5.64E-01 | 1.06E+00 | 8.67E-02 | 1.06E+00 |
| ENSG00000( | MPP4 | 2.11E-01 | ? | 6.40E-01 | -6.43E-01 | -1.56E+00 |
| ENSG000001 | ZXDA | 2.11E-01 | 5.65E-01 | 1.08E+00 | 1.16E-01 | 1.08E+00 |
| ENSG000001 | SECISBP2 | 2.11E-01 | 5.65E-01 | 1.04E+00 | 6.33E-02 | 1.04E+00 |
| ENSG000001 | CXCL10 | 2.11E-01 | ? | 2.05E+00 | 1.03E+00 | 2.05E+00 |
| ENSG000001 | CHRNA7 | 2.12E-01 | ? | 1.41E+00 | 4.92E-01 | 1.41E+00 |
| ENSG00000( | SPATA7 | 2.12E-01 | 5.65E-01 | 1.08E+00 | 1.13E-01 | 1.08E+00 |
| ENSG000002 | --- | 2.12E-01 | ? | 6.95E-01 | -5.26E-01 | -1.44E+00 |
| ENSG000001 | LGI2 | 2.12E-01 | ? | 5.03E-01 | -9.92E-01 | -1.99E+00 |
| ENSG000002 | ADAT3 | 2.12E-01 | ? | 6.79E-01 | -5.58E-01 | -1.47E+00 |
| ENSG000001 | ITGB3BP | 2.12E-01 | 5.65E-01 | 8.96E-01 | -1.58E-01 | -1.12E+00 |
| ENSG000001 | CYP51A1-AS1 | 2.12E-01 | ? | 5.27E-01 | -9.24E-01 | -1.90E+00 |
| ENSG000001 | NPAS3 | 2.12E-01 | ? | 7.93E-01 | -3.35E-01 | -1.26E+00 |
| ENSG000001 | DTNA | 2.12E-01 | 5.65E-01 | 1.05E+00 | 7.07E-02 | 1.05E+00 |
| ENSG000001 | COG2 | 2.12E-01 | 5.65E-01 | 1.04E+00 | 5.36E-02 | 1.04E+00 |
| ENSG000001 | PDXDC2P-NP | 2.12E-01 | 5.65E-01 | 9.24E-01 | -1.14E-01 | -1.08E+00 |
| ENSG00000( | UBE3C | 2.12E-01 | 5.65E-01 | 1.04E+00 | 5.38E-02 | 1.04E+00 |
| ENSG000001 | CMTM8 | 2.12E-01 | 5.65E-01 | 1.17E+00 | 2.23E-01 | 1.17E+00 |
| ENSG000001 | FLI1 | 2.12E-01 | 5.65E-01 | 1.07E+00 | 9.75E-02 | 1.07E+00 |
| ENSG000002 | --- | 2.12E-01 | ? | 1.39E+00 | 4.78E-01 | 1.39E+00 |
| ENSG000002 | --- | 2.12E-01 | ? | 7.25E-01 | -4.64E-01 | -1.38E+00 |
| ENSG000001 | FAM120B | 2.12E-01 | 5.65E-01 | 1.03E+00 | 4.28E-02 | 1.03E+00 |
| ENSG000001 | ZG16B | 2.12E-01 | ? | 1.49E+00 | 5.75E-01 | 1.49E+00 |
| ENSG000001 | SMC3 | 2.12E-01 | 5.65E-01 | 9.37E-01 | -9.33E-02 | -1.07E+00 |
| ENSG000002 | --- | 2.12E-01 | ? | 1.30E+00 | 3.73E-01 | 1.30E+00 |
| ENSG00000( | MUL1 | 2.12E-01 | 5.66E-01 | 1.07E+00 | 1.01E-01 | 1.07E+00 |
| ENSG000001 | ZNF436 | 2.12E-01 | 5.66E-01 | 1.05E+00 | 6.66E-02 | 1.05E+00 |
| ENSG000002 | --- | 2.12E-01 | 5.66E-01 | 8.22E-01 | -2.83E-01 | -1.22E+00 |
| ENSG000001 | RHOV | 2.12E-01 | 5.66E-01 | 1.22E+00 | 2.89E-01 | 1.22E+00 |
| ENSG000001 | PPP1R14A | 2.12E-01 | 5.66E-01 | 1.23E+00 | 2.98E-01 | 1.23E+00 |

| | | | | | | |
|---|---|---|---|---|---|---|
| ENSG000001 | C1orf122 | 2.12E-01 | 5.66E-01 | 1.12E+00 | 1.60E-01 | 1.12E+00 |
| ENSG000001 | NDUFB3 | 2.12E-01 | 5.66E-01 | 1.04E+00 | 6.00E-02 | 1.04E+00 |
| ENSG000001 | GBGT1 | 2.13E-01 | 5.66E-01 | 1.17E+00 | 2.30E-01 | 1.17E+00 |
| ENSG000002 | SMG1P2 | 2.13E-01 | 5.66E-01 | 8.05E-01 | -3.13E-01 | -1.24E+00 |
| ENSG000001 | NUDT7 | 2.13E-01 | 5.66E-01 | 1.17E+00 | 2.25E-01 | 1.17E+00 |
| ENSG000001 | CRIM1 | 2.13E-01 | 5.66E-01 | 1.05E+00 | 7.30E-02 | 1.05E+00 |
| ENSG000001 | BBLN | 2.13E-01 | 5.66E-01 | 1.16E+00 | 2.09E-01 | 1.16E+00 |
| ENSG000001 | FEM1B | 2.13E-01 | 5.66E-01 | 9.48E-01 | -7.65E-02 | -1.05E+00 |
| ENSG00000C | UBE2D4 | 2.13E-01 | 5.66E-01 | 1.07E+00 | 9.10E-02 | 1.07E+00 |
| ENSG000002 | --- | 2.13E-01 | ? | 7.85E-01 | -3.48E-01 | -1.27E+00 |
| ENSG000002 | MPV17L2 | 2.13E-01 | 5.66E-01 | 9.34E-01 | -9.90E-02 | -1.07E+00 |
| ENSG000001 | PHF6 | 2.13E-01 | 5.66E-01 | 1.04E+00 | 5.53E-02 | 1.04E+00 |
| ENSG000001 | EXOSC9 | 2.13E-01 | 5.66E-01 | 9.46E-01 | -7.95E-02 | -1.06E+00 |
| ENSG000001 | YJU2 | 2.13E-01 | 5.66E-01 | 9.34E-01 | -9.81E-02 | -1.07E+00 |
| ENSG000001 | FKBP8 | 2.13E-01 | 5.66E-01 | 1.09E+00 | 1.25E-01 | 1.09E+00 |
| ENSG000001 | SEMA4F | 2.13E-01 | 5.66E-01 | 1.08E+00 | 1.08E-01 | 1.08E+00 |
| ENSG00000C | ATP5F1D | 2.13E-01 | 5.66E-01 | 1.11E+00 | 1.56E-01 | 1.11E+00 |
| ENSG000002 | HLA-K | 2.13E-01 | ? | 5.84E-01 | -7.76E-01 | -1.71E+00 |
| ENSG000002 | LINC01546 | 2.14E-01 | 5.66E-01 | 1.15E+00 | 2.06E-01 | 1.15E+00 |
| ENSG00000C | SNX24 | 2.14E-01 | 5.66E-01 | 1.08E+00 | 1.11E-01 | 1.08E+00 |
| ENSG000002 | --- | 2.14E-01 | 5.66E-01 | 1.15E+00 | 2.01E-01 | 1.15E+00 |
| ENSG000002 | STIMATE | 2.14E-01 | 5.66E-01 | 1.06E+00 | 7.84E-02 | 1.06E+00 |
| ENSG000001 | UBALD1 | 2.14E-01 | 5.66E-01 | 1.10E+00 | 1.33E-01 | 1.10E+00 |
| ENSG000001 | CCDC59 | 2.14E-01 | 5.66E-01 | 9.37E-01 | -9.32E-02 | -1.07E+00 |
| ENSG000001 | PKDREJ | 2.14E-01 | ? | 6.89E-01 | -5.38E-01 | -1.45E+00 |
| ENSG000001 | GDPGP1 | 2.14E-01 | 5.66E-01 | 8.48E-01 | -2.38E-01 | -1.18E+00 |
| ENSG000001 | SEC16B | 2.14E-01 | 5.66E-01 | 7.49E-01 | -4.17E-01 | -1.33E+00 |
| ENSG000001 | MFF | 2.14E-01 | 5.67E-01 | 9.58E-01 | -6.18E-02 | -1.04E+00 |
| ENSG000001 | CAMSAP2 | 2.14E-01 | 5.67E-01 | 9.50E-01 | -7.47E-02 | -1.05E+00 |
| ENSG000001 | MEST | 2.14E-01 | 5.67E-01 | 1.08E+00 | 1.12E-01 | 1.08E+00 |
| ENSG000002 | --- | 2.14E-01 | ? | 1.34E+00 | 4.20E-01 | 1.34E+00 |
| ENSG000001 | NEK10 | 2.14E-01 | 5.67E-01 | 8.60E-01 | -2.17E-01 | -1.16E+00 |
| ENSG000001 | RAB3GAP2 | 2.14E-01 | 5.67E-01 | 9.70E-01 | -4.36E-02 | -1.03E+00 |
| ENSG000002 | --- | 2.14E-01 | 5.68E-01 | 7.24E-01 | -4.65E-01 | -1.38E+00 |
| ENSG000001 | SAFB | 2.14E-01 | 5.68E-01 | 9.59E-01 | -6.07E-02 | -1.04E+00 |
| ENSG000001 | CDC16 | 2.14E-01 | 5.68E-01 | 9.70E-01 | -4.32E-02 | -1.03E+00 |
| ENSG000002 | --- | 2.15E-01 | 5.68E-01 | 8.62E-01 | -2.13E-01 | -1.16E+00 |
| ENSG000001 | ZNF701 | 2.15E-01 | 5.68E-01 | 9.22E-01 | -1.17E-01 | -1.08E+00 |
| ENSG000001 | TUB | 2.15E-01 | 5.68E-01 | 1.18E+00 | 2.44E-01 | 1.18E+00 |
| ENSG000002 | GTF3C2-AS1 | 2.15E-01 | ? | 1.59E+00 | 6.70E-01 | 1.59E+00 |
| ENSG000001 | NARF | 2.15E-01 | 5.68E-01 | 9.49E-01 | -7.61E-02 | -1.05E+00 |
| ENSG00000C | ALG6 | 2.15E-01 | 5.68E-01 | 9.34E-01 | -9.91E-02 | -1.07E+00 |
| ENSG000001 | CLDN6 | 2.15E-01 | ? | 7.49E-01 | -4.16E-01 | -1.33E+00 |
| ENSG000002 | --- | 2.15E-01 | 5.68E-01 | 8.24E-01 | -2.79E-01 | -1.21E+00 |
| ENSG000002 | --- | 2.15E-01 | ? | 8.21E-01 | -2.84E-01 | -1.22E+00 |

| | | | | | | | |
|---|---|---|---|---|---|---|---|
| ENSG000001 | EIF4E3 | 2.15E-01 | | 5.69E-01 | 9.41E-01 | -8.83E-02 | -1.06E+00 |
| ENSG000002 | PMF1-BGLAI | 2.15E-01 | | 5.69E-01 | 7.78E-01 | -3.63E-01 | -1.29E+00 |
| ENSG000001 | COPS7A | 2.15E-01 | | 5.69E-01 | 1.05E+00 | 6.78E-02 | 1.05E+00 |
| ENSG000002 | --- | 2.15E-01 | | 5.69E-01 | 1.12E+00 | 1.61E-01 | 1.12E+00 |
| ENSG000002 | --- | 2.15E-01 | | 5.69E-01 | 1.14E+00 | 1.90E-01 | 1.14E+00 |
| ENSG000002 | CD99P1 | 2.15E-01 | ? | | 2.75E-01 | -1.86E+00 | -3.64E+00 |
| ENSG000002 | --- | 2.15E-01 | ? | | 7.18E-01 | -4.79E-01 | -1.39E+00 |
| ENSG000001 | TRA2A | 2.15E-01 | | 5.69E-01 | 9.60E-01 | -5.87E-02 | -1.04E+00 |
| ENSG000002 | FER1L6 | 2.15E-01 | | 5.69E-01 | 8.63E-01 | -2.12E-01 | -1.16E+00 |
| ENSG000001 | PCCB | 2.16E-01 | | 5.69E-01 | 1.06E+00 | 8.41E-02 | 1.06E+00 |
| ENSG000001 | SLC4A5 | 2.16E-01 | | 5.69E-01 | 8.49E-01 | -2.37E-01 | -1.18E+00 |
| ENSG000001 | ZCCHC9 | 2.16E-01 | | 5.69E-01 | 9.42E-01 | -8.68E-02 | -1.06E+00 |
| ENSG000001 | EXOC3 | 2.16E-01 | | 5.69E-01 | 1.05E+00 | 6.65E-02 | 1.05E+00 |
| ENSG00000C | MBNL3 | 2.16E-01 | | 5.69E-01 | 9.11E-01 | -1.34E-01 | -1.10E+00 |
| ENSG000002 | ZNF727 | 2.16E-01 | | 5.69E-01 | 1.17E+00 | 2.28E-01 | 1.17E+00 |
| ENSG000002 | --- | 2.16E-01 | ? | | 1.36E+00 | 4.42E-01 | 1.36E+00 |
| ENSG000002 | LINC00680 | 2.16E-01 | | 5.69E-01 | 9.10E-01 | -1.36E-01 | -1.10E+00 |
| ENSG000001 | PLD2 | 2.16E-01 | | 5.69E-01 | 1.06E+00 | 9.03E-02 | 1.06E+00 |
| ENSG000001 | DNAH2 | 2.16E-01 | | 5.69E-01 | 1.15E+00 | 2.06E-01 | 1.15E+00 |
| ENSG000001 | ARMCX1 | 2.16E-01 | | 5.69E-01 | 9.57E-01 | -6.39E-02 | -1.05E+00 |
| ENSG000001 | PNLDC1 | 2.16E-01 | ? | | 1.86E+00 | 8.97E-01 | 1.86E+00 |
| ENSG000001 | TMEM237 | 2.16E-01 | | 5.69E-01 | 9.47E-01 | -7.91E-02 | -1.06E+00 |
| ENSG000001 | HABP4 | 2.16E-01 | | 5.69E-01 | 9.50E-01 | -7.33E-02 | -1.05E+00 |
| ENSG000001 | SLC25A10 | 2.16E-01 | | 5.69E-01 | 1.22E+00 | 2.90E-01 | 1.22E+00 |
| ENSG000002 | CDPF1 | 2.16E-01 | | 5.69E-01 | 1.12E+00 | 1.61E-01 | 1.12E+00 |
| ENSG00000C | NDC1 | 2.16E-01 | | 5.69E-01 | 9.40E-01 | -8.88E-02 | -1.06E+00 |
| ENSG000002 | CCDC39 | 2.16E-01 | | 5.69E-01 | 8.30E-01 | -2.69E-01 | -1.20E+00 |
| ENSG00000C | POLR2E | 2.16E-01 | | 5.70E-01 | 1.07E+00 | 1.04E-01 | 1.07E+00 |
| ENSG000001 | OGFRL1 | 2.16E-01 | | 5.70E-01 | 9.42E-01 | -8.69E-02 | -1.06E+00 |
| ENSG00000C | EHD2 | 2.16E-01 | | 5.70E-01 | 1.04E+00 | 6.07E-02 | 1.04E+00 |
| ENSG000001 | LY6D | 2.16E-01 | | 5.70E-01 | 1.21E+00 | 2.77E-01 | 1.21E+00 |
| ENSG000002 | --- | 2.17E-01 | ? | | 1.31E+00 | 3.86E-01 | 1.31E+00 |
| ENSG00000C | DNTTIP2 | 2.17E-01 | | 5.70E-01 | 9.55E-01 | -6.63E-02 | -1.05E+00 |
| ENSG000001 | CSTA | 2.17E-01 | | 5.70E-01 | 1.09E+00 | 1.25E-01 | 1.09E+00 |
| ENSG000002 | --- | 2.17E-01 | ? | | 1.31E+00 | 3.86E-01 | 1.31E+00 |
| ENSG000001 | AHCTF1 | 2.17E-01 | | 5.70E-01 | 9.48E-01 | -7.67E-02 | -1.05E+00 |
| ENSG000001 | APEH | 2.17E-01 | | 5.70E-01 | 1.07E+00 | 9.54E-02 | 1.07E+00 |
| ENSG000001 | ZFHX2-AS1 | 2.17E-01 | | 5.70E-01 | 8.63E-01 | -2.13E-01 | -1.16E+00 |
| ENSG000001 | SH3RF1 | 2.17E-01 | | 5.70E-01 | 9.50E-01 | -7.43E-02 | -1.05E+00 |
| ENSG00000C | ETV1 | 2.17E-01 | | 5.70E-01 | 8.10E-01 | -3.03E-01 | -1.23E+00 |
| ENSG00000C | EIF2B3 | 2.17E-01 | | 5.70E-01 | 1.05E+00 | 6.61E-02 | 1.05E+00 |
| ENSG000001 | TRMU | 2.17E-01 | | 5.70E-01 | 9.33E-01 | -1.01E-01 | -1.07E+00 |
| ENSG000002 | PHACTR4 | 2.17E-01 | | 5.70E-01 | 1.05E+00 | 6.78E-02 | 1.05E+00 |
| ENSG000002 | CALML3-AS1 | 2.17E-01 | | 5.70E-01 | 8.76E-01 | -1.91E-01 | -1.14E+00 |
| ENSG00000C | COL9A3 | 2.17E-01 | ? | | 1.74E+00 | 7.99E-01 | 1.74E+00 |

| | | | | | | | |
|---|---|---|---|---|---|---|---|
| ENSG00000C | GPM6B | 2.17E-01 | | 5.70E-01 | 8.15E-01 | -2.95E-01 | -1.23E+00 |
| ENSG000001 | UQCR11 | 2.17E-01 | | 5.70E-01 | 1.08E+00 | 1.15E-01 | 1.08E+00 |
| ENSG000001 | ZNF410 | 2.17E-01 | | 5.71E-01 | 1.06E+00 | 7.84E-02 | 1.06E+00 |
| ENSG000001 | ZNF26 | 2.18E-01 | | 5.71E-01 | 9.22E-01 | -1.18E-01 | -1.09E+00 |
| ENSG00000C | DHX8 | 2.18E-01 | | 5.71E-01 | 1.03E+00 | 4.71E-02 | 1.03E+00 |
| ENSG000001 | RRM2 | 2.18E-01 | | 5.71E-01 | 8.08E-01 | -3.08E-01 | -1.24E+00 |
| ENSG000002 | NOC2LP2 | 2.18E-01 | ? | | 5.81E-01 | -7.83E-01 | -1.72E+00 |
| ENSG000002 | --- | 2.18E-01 | ? | | 6.89E-01 | -5.37E-01 | -1.45E+00 |
| ENSG00000C | SLC66A1 | 2.18E-01 | | 5.71E-01 | 1.10E+00 | 1.33E-01 | 1.10E+00 |
| ENSG000002 | H3-7 | 2.18E-01 | | 5.71E-01 | 9.34E-01 | -9.83E-02 | -1.07E+00 |
| ENSG000001 | TSFM | 2.18E-01 | | 5.71E-01 | 9.27E-01 | -1.09E-01 | -1.08E+00 |
| ENSG000001 | ZC2HC1A | 2.18E-01 | | 5.71E-01 | 9.24E-01 | -1.14E-01 | -1.08E+00 |
| ENSG000001 | CCT3 | 2.18E-01 | | 5.71E-01 | 1.04E+00 | 6.02E-02 | 1.04E+00 |
| ENSG00000C | SFSWAP | 2.18E-01 | | 5.71E-01 | 9.56E-01 | -6.49E-02 | -1.05E+00 |
| ENSG00000C | XRN2 | 2.18E-01 | | 5.71E-01 | 9.71E-01 | -4.27E-02 | -1.03E+00 |
| ENSG000001 | ARTN | 2.18E-01 | | 5.71E-01 | 1.09E+00 | 1.18E-01 | 1.09E+00 |
| ENSG000002 | --- | 2.18E-01 | ? | | 6.19E-01 | -6.91E-01 | -1.61E+00 |
| ENSG000001 | TCHHL1 | 2.18E-01 | ? | | 4.43E-01 | -1.18E+00 | -2.26E+00 |
| ENSG000002 | --- | 2.18E-01 | ? | | 7.06E-01 | -5.03E-01 | -1.42E+00 |
| ENSG000001 | KIF20B | 2.18E-01 | | 5.72E-01 | 9.15E-01 | -1.27E-01 | -1.09E+00 |
| ENSG000001 | ZNF14 | 2.18E-01 | | 5.72E-01 | 9.22E-01 | -1.18E-01 | -1.08E+00 |
| ENSG000001 | MYCL | 2.19E-01 | | 5.72E-01 | 1.08E+00 | 1.17E-01 | 1.08E+00 |
| ENSG000001 | LINGO2 | 2.19E-01 | ? | | 6.91E-01 | -5.34E-01 | -1.45E+00 |
| ENSG000001 | LPXN | 2.19E-01 | | 5.73E-01 | 9.46E-01 | -7.99E-02 | -1.06E+00 |
| ENSG000002 | CDC20-DT | 2.19E-01 | ? | | 1.52E+00 | 6.02E-01 | 1.52E+00 |
| ENSG000001 | TGFB1I1 | 2.19E-01 | | 5.73E-01 | 1.07E+00 | 1.02E-01 | 1.07E+00 |
| ENSG000002 | --- | 2.19E-01 | ? | | 1.33E+00 | 4.15E-01 | 1.33E+00 |
| ENSG000002 | --- | 2.19E-01 | ? | | 1.88E+00 | 9.12E-01 | 1.88E+00 |
| ENSG000001 | ZNF439 | 2.19E-01 | | 5.73E-01 | 8.82E-01 | -1.81E-01 | -1.13E+00 |
| ENSG000001 | RAB4A-AS1 | 2.19E-01 | | 5.74E-01 | 1.07E+00 | 9.66E-02 | 1.07E+00 |
| ENSG000001 | SMG8 | 2.19E-01 | | 5.74E-01 | 1.04E+00 | 5.66E-02 | 1.04E+00 |
| ENSG00000C | CATSPERG | 2.19E-01 | | 5.74E-01 | 8.55E-01 | -2.26E-01 | -1.17E+00 |
| ENSG000001 | CHRAC1 | 2.19E-01 | | 5.74E-01 | 9.39E-01 | -9.10E-02 | -1.07E+00 |
| ENSG000001 | TIA1 | 2.20E-01 | | 5.74E-01 | 9.61E-01 | -5.68E-02 | -1.04E+00 |
| ENSG00000C | TRIP13 | 2.20E-01 | | 5.74E-01 | 1.09E+00 | 1.23E-01 | 1.09E+00 |
| ENSG000002 | PLEKHO2 | 2.20E-01 | | 5.74E-01 | 9.49E-01 | -7.53E-02 | -1.05E+00 |
| ENSG000002 | CFAP57 | 2.20E-01 | ? | | 1.46E+00 | 5.46E-01 | 1.46E+00 |
| ENSG000002 | --- | 2.20E-01 | ? | | 1.48E+00 | 5.62E-01 | 1.48E+00 |
| ENSG00000C | CYP46A1 | 2.20E-01 | ? | | 1.45E+00 | 5.38E-01 | 1.45E+00 |
| ENSG000001 | NPEPPS | 2.20E-01 | | 5.75E-01 | 1.04E+00 | 6.23E-02 | 1.04E+00 |
| ENSG000001 | SF1 | 2.20E-01 | | 5.75E-01 | 9.74E-01 | -3.84E-02 | -1.03E+00 |
| ENSG000002 | ZBED9 | 2.20E-01 | | 5.75E-01 | 1.05E+00 | 7.25E-02 | 1.05E+00 |
| ENSG00000C | BORCS8-MEI | 2.20E-01 | | 5.75E-01 | 1.22E+00 | 2.82E-01 | 1.22E+00 |
| ENSG000002 | OR7E126P | 2.20E-01 | ? | | 7.32E-01 | -4.49E-01 | -1.37E+00 |
| ENSG000001 | BEND3 | 2.20E-01 | | 5.75E-01 | 1.06E+00 | 8.44E-02 | 1.06E+00 |

| | | | | | | |
|---|---|---|---|---|---|---|
| ENSG000002 | CCDC13 | 2.20E-01 | ? | 5.98E-01 | -7.41E-01 | -1.67E+00 |
| ENSG000001 | NPIPB9 | 2.20E-01 | ? | 1.51E+00 | 5.93E-01 | 1.51E+00 |
| ENSG000001 | PECR | 2.21E-01 | 5.76E-01 | 8.95E-01 | -1.60E-01 | -1.12E+00 |
| ENSG000002 | --- | 2.21E-01 | ? | 1.87E+00 | 8.99E-01 | 1.87E+00 |
| ENSG000002 | CD109-AS1 | 2.21E-01 | ? | 1.52E+00 | 6.01E-01 | 1.52E+00 |
| ENSG000001 | PCDH12 | 2.21E-01 | 5.76E-01 | 8.73E-01 | -1.96E-01 | -1.15E+00 |
| ENSG00000C | BTN3A1 | 2.21E-01 | 5.76E-01 | 1.09E+00 | 1.27E-01 | 1.09E+00 |
| ENSG000002 | --- | 2.21E-01 | ? | 1.36E+00 | 4.41E-01 | 1.36E+00 |
| ENSG000001 | TM9SF2 | 2.21E-01 | 5.76E-01 | 9.76E-01 | -3.58E-02 | -1.03E+00 |
| ENSG000001 | CYYR1-AS1 | 2.21E-01 | ? | 7.85E-01 | -3.49E-01 | -1.27E+00 |
| ENSG000001 | DAW1 | 2.21E-01 | ? | 1.23E+00 | 3.02E-01 | 1.23E+00 |
| ENSG000001 | ARRDC1 | 2.21E-01 | 5.77E-01 | 1.09E+00 | 1.31E-01 | 1.09E+00 |
| ENSG000002 | --- | 2.21E-01 | 5.77E-01 | 8.86E-01 | -1.75E-01 | -1.13E+00 |
| ENSG000002 | --- | 2.21E-01 | ? | 1.43E+00 | 5.19E-01 | 1.43E+00 |
| ENSG000001 | INKA2 | 2.21E-01 | 5.77E-01 | 1.07E+00 | 1.00E-01 | 1.07E+00 |
| ENSG00000C | SENP1 | 2.21E-01 | 5.77E-01 | 9.42E-01 | -8.67E-02 | -1.06E+00 |
| ENSG000001 | SPPL3 | 2.21E-01 | 5.77E-01 | 1.04E+00 | 6.15E-02 | 1.04E+00 |
| ENSG000001 | SPC25 | 2.21E-01 | ? | 7.73E-01 | -3.71E-01 | -1.29E+00 |
| ENSG000001 | HSPBAP1 | 2.21E-01 | 5.77E-01 | 1.12E+00 | 1.63E-01 | 1.12E+00 |
| ENSG000001 | TRIM29 | 2.21E-01 | 5.77E-01 | 1.05E+00 | 6.81E-02 | 1.05E+00 |
| ENSG000002 | --- | 2.22E-01 | ? | 1.73E+00 | 7.87E-01 | 1.73E+00 |
| ENSG000001 | EMC9 | 2.22E-01 | 5.77E-01 | 1.12E+00 | 1.68E-01 | 1.12E+00 |
| ENSG000001 | CXCL1 | 2.22E-01 | 5.77E-01 | 8.36E-01 | -2.58E-01 | -1.20E+00 |
| ENSG000001 | MEIS3P1 | 2.22E-01 | 5.77E-01 | 1.17E+00 | 2.32E-01 | 1.17E+00 |
| ENSG00000C | DDX24 | 2.22E-01 | 5.77E-01 | 1.03E+00 | 4.46E-02 | 1.03E+00 |
| ENSG000001 | TMEM178A | 2.22E-01 | ? | 7.66E-01 | -3.85E-01 | -1.31E+00 |
| ENSG000001 | PSCA | 2.22E-01 | ? | 1.65E+00 | 7.26E-01 | 1.65E+00 |
| ENSG000001 | CD47 | 2.22E-01 | 5.78E-01 | 9.55E-01 | -6.63E-02 | -1.05E+00 |
| ENSG000001 | ELP3 | 2.22E-01 | 5.78E-01 | 9.59E-01 | -5.98E-02 | -1.04E+00 |
| ENSG000001 | FBXO4 | 2.22E-01 | 5.78E-01 | 1.14E+00 | 1.88E-01 | 1.14E+00 |
| ENSG000002 | SCAMP1-AS1 | 2.22E-01 | 5.78E-01 | 1.13E+00 | 1.73E-01 | 1.13E+00 |
| ENSG000002 | RPL4P5 | 2.22E-01 | 5.78E-01 | 1.21E+00 | 2.81E-01 | 1.21E+00 |
| ENSG000001 | NEK3 | 2.22E-01 | 5.78E-01 | 9.22E-01 | -1.18E-01 | -1.08E+00 |
| ENSG000002 | FGGY-DT | 2.22E-01 | ? | 5.75E-01 | -7.97E-01 | -1.74E+00 |
| ENSG000002 | --- | 2.22E-01 | ? | 6.39E-01 | -6.47E-01 | -1.57E+00 |
| ENSG000002 | EIF5AL1 | 2.22E-01 | 5.78E-01 | 8.75E-01 | -1.92E-01 | -1.14E+00 |
| ENSG000002 | TMEM238 | 2.22E-01 | ? | 1.76E+00 | 8.12E-01 | 1.76E+00 |
| ENSG000002 | --- | 2.23E-01 | 5.78E-01 | 1.15E+00 | 2.08E-01 | 1.15E+00 |
| ENSG000002 | KRTAP2-3 | 2.23E-01 | 5.78E-01 | 8.26E-01 | -2.76E-01 | -1.21E+00 |
| ENSG000002 | SNAP25-AS1 | 2.23E-01 | ? | 1.43E+00 | 5.15E-01 | 1.43E+00 |
| ENSG00000C | IPO5 | 2.23E-01 | 5.78E-01 | 1.04E+00 | 5.02E-02 | 1.04E+00 |
| ENSG000001 | NAA16 | 2.23E-01 | 5.78E-01 | 9.39E-01 | -9.06E-02 | -1.06E+00 |
| ENSG000001 | TOGARAM1 | 2.23E-01 | 5.78E-01 | 1.05E+00 | 7.64E-02 | 1.05E+00 |
| ENSG000001 | JPT1 | 2.23E-01 | 5.78E-01 | 1.05E+00 | 7.04E-02 | 1.05E+00 |
| ENSG000001 | APBB3 | 2.23E-01 | 5.78E-01 | 1.16E+00 | 2.17E-01 | 1.16E+00 |

| | | | | | | |
|---|---|---|---|---|---|---|
| ENSG000001 | SMYD2 | 2.23E-01 | 5.78E-01 | 1.04E+00 | 5.52E-02 | 1.04E+00 |
| ENSG000001 | ASB9 | 2.23E-01 | 5.78E-01 | 1.16E+00 | 2.17E-01 | 1.16E+00 |
| ENSG000002 | --- | 2.23E-01 | 5.78E-01 | 9.02E-01 | -1.49E-01 | -1.11E+00 |
| ENSG000001 | LDHA | 2.23E-01 | 5.78E-01 | 9.77E-01 | -3.33E-02 | -1.02E+00 |
| ENSG000002 | --- | 2.23E-01 | ? | 1.49E+00 | 5.74E-01 | 1.49E+00 |
| ENSG000001 | ZRANB3 | 2.23E-01 | 5.78E-01 | 1.08E+00 | 1.07E-01 | 1.08E+00 |
| ENSG00000C | EFHC1 | 2.23E-01 | 5.78E-01 | 1.09E+00 | 1.26E-01 | 1.09E+00 |
| ENSG000002 | MTCP1 | 2.23E-01 | 5.78E-01 | 8.30E-01 | -2.69E-01 | -1.20E+00 |
| ENSG000001 | ZMYM6 | 2.23E-01 | 5.79E-01 | 9.63E-01 | -5.46E-02 | -1.04E+00 |
| ENSG00000C | FXYD5 | 2.23E-01 | 5.79E-01 | 9.34E-01 | -9.92E-02 | -1.07E+00 |
| ENSG000001 | TTF1 | 2.23E-01 | 5.79E-01 | 9.54E-01 | -6.83E-02 | -1.05E+00 |
| ENSG000001 | PAQR5 | 2.23E-01 | 5.79E-01 | 8.28E-01 | -2.72E-01 | -1.21E+00 |
| ENSG000001 | NDRG1 | 2.23E-01 | 5.79E-01 | 9.37E-01 | -9.38E-02 | -1.07E+00 |
| ENSG000002 | NONOP2 | 2.23E-01 | ? | 7.98E-01 | -3.26E-01 | -1.25E+00 |
| ENSG000001 | GPAT2 | 2.24E-01 | 5.79E-01 | 1.16E+00 | 2.19E-01 | 1.16E+00 |
| ENSG000002 | --- | 2.24E-01 | 5.80E-01 | 7.27E-01 | -4.59E-01 | -1.37E+00 |
| ENSG000001 | TYSND1 | 2.24E-01 | 5.80E-01 | 1.10E+00 | 1.43E-01 | 1.10E+00 |
| ENSG000001 | NGLY1 | 2.24E-01 | 5.80E-01 | 9.57E-01 | -6.41E-02 | -1.05E+00 |
| ENSG000001 | PTCH1 | 2.24E-01 | 5.80E-01 | 1.10E+00 | 1.35E-01 | 1.10E+00 |
| ENSG000001 | INHBE | 2.24E-01 | ? | 5.30E-01 | -9.15E-01 | -1.89E+00 |
| ENSG000001 | ZNF662 | 2.24E-01 | 5.80E-01 | 1.17E+00 | 2.28E-01 | 1.17E+00 |
| ENSG000001 | SMAGP | 2.24E-01 | 5.80E-01 | 1.07E+00 | 1.02E-01 | 1.07E+00 |
| ENSG000001 | CREBL2 | 2.24E-01 | 5.80E-01 | 9.57E-01 | -6.31E-02 | -1.04E+00 |
| ENSG000002 | NT5M | 2.24E-01 | ? | 1.51E+00 | 5.92E-01 | 1.51E+00 |
| ENSG000002 | --- | 2.24E-01 | ? | 7.03E-01 | -5.08E-01 | -1.42E+00 |
| ENSG000001 | FXN | 2.24E-01 | 5.80E-01 | 9.50E-01 | -7.36E-02 | -1.05E+00 |
| ENSG000001 | RIC3 | 2.24E-01 | 5.80E-01 | 1.15E+00 | 1.98E-01 | 1.15E+00 |
| ENSG000001 | CRIPT | 2.24E-01 | 5.80E-01 | 9.35E-01 | -9.76E-02 | -1.07E+00 |
| ENSG000001 | UBE2F | 2.24E-01 | 5.80E-01 | 9.59E-01 | -6.06E-02 | -1.04E+00 |
| ENSG000002 | --- | 2.24E-01 | ? | 7.96E-01 | -3.29E-01 | -1.26E+00 |
| ENSG000001 | PCF11 | 2.24E-01 | 5.80E-01 | 9.50E-01 | -7.43E-02 | -1.05E+00 |
| ENSG000002 | --- | 2.25E-01 | ? | 1.70E+00 | 7.65E-01 | 1.70E+00 |
| ENSG000001 | KDM5C | 2.25E-01 | 5.80E-01 | 1.04E+00 | 5.74E-02 | 1.04E+00 |
| ENSG000002 | LINC03015 | 2.25E-01 | 5.80E-01 | 8.76E-01 | -1.90E-01 | -1.14E+00 |
| ENSG000001 | LARS1 | 2.25E-01 | 5.80E-01 | 9.70E-01 | -4.38E-02 | -1.03E+00 |
| ENSG000001 | SEC14L2 | 2.25E-01 | 5.80E-01 | 1.07E+00 | 9.88E-02 | 1.07E+00 |
| ENSG000001 | B3GAT3 | 2.25E-01 | 5.80E-01 | 1.10E+00 | 1.38E-01 | 1.10E+00 |
| ENSG000002 | --- | 2.25E-01 | 5.80E-01 | 8.64E-01 | -2.12E-01 | -1.16E+00 |
| ENSG000001 | SAYSD1 | 2.25E-01 | 5.80E-01 | 1.06E+00 | 8.86E-02 | 1.06E+00 |
| ENSG000001 | PREB | 2.25E-01 | 5.80E-01 | 1.05E+00 | 7.27E-02 | 1.05E+00 |
| ENSG000001 | DDX60 | 2.25E-01 | 5.80E-01 | 1.08E+00 | 1.05E-01 | 1.08E+00 |
| ENSG000001 | PHF10 | 2.25E-01 | 5.80E-01 | 9.45E-01 | -8.22E-02 | -1.06E+00 |
| ENSG000001 | NLGN4X | 2.25E-01 | ? | 1.29E+00 | 3.62E-01 | 1.29E+00 |
| ENSG000002 | PPM1F-AS1 | 2.25E-01 | 5.80E-01 | 9.24E-01 | -1.14E-01 | -1.08E+00 |
| ENSG000001 | TBC1D10B | 2.25E-01 | 5.81E-01 | 9.52E-01 | -7.03E-02 | -1.05E+00 |

| | | | | | | |
|---|---|---|---|---|---|---|
| ENSG000001 | LSM5 | 2.25E-01 | 5.81E-01 | 1.06E+00 | 7.87E-02 | 1.06E+00 |
| ENSG000001 | CORO7-PAM | 2.25E-01 | 5.81E-01 | 7.28E-01 | -4.58E-01 | -1.37E+00 |
| ENSG000002 | VDAC1P8 | 2.25E-01 | 5.81E-01 | 8.57E-01 | -2.23E-01 | -1.17E+00 |
| ENSG000002 | --- | 2.25E-01 | 5.81E-01 | 8.70E-01 | -2.01E-01 | -1.15E+00 |
| ENSG000001 | SCN1B | 2.25E-01 | ? | 1.26E+00 | 3.29E-01 | 1.26E+00 |
| ENSG00000C | SH2D3C | 2.25E-01 | ? | 1.73E+00 | 7.95E-01 | 1.73E+00 |
| ENSG000002 | CYP4A22-AS1 | 2.25E-01 | ? | 6.78E-01 | -5.60E-01 | -1.47E+00 |
| ENSG000001 | NUSAP1 | 2.25E-01 | 5.81E-01 | 8.82E-01 | -1.81E-01 | -1.13E+00 |
| ENSG000002 | --- | 2.25E-01 | ? | 7.55E-01 | -4.05E-01 | -1.32E+00 |
| ENSG000002 | THAP9-AS1 | 2.26E-01 | 5.81E-01 | 9.25E-01 | -1.13E-01 | -1.08E+00 |
| ENSG000001 | TRAPPC14 | 2.26E-01 | 5.81E-01 | 1.07E+00 | 9.55E-02 | 1.07E+00 |
| ENSG000002 | PRRT4 | 2.26E-01 | ? | 1.67E+00 | 7.38E-01 | 1.67E+00 |
| ENSG00000C | SEMA6A | 2.26E-01 | 5.81E-01 | 9.34E-01 | -9.78E-02 | -1.07E+00 |
| ENSG000001 | WDR41 | 2.26E-01 | 5.81E-01 | 9.57E-01 | -6.28E-02 | -1.04E+00 |
| ENSG000001 | TTC39C | 2.26E-01 | 5.81E-01 | 1.06E+00 | 8.45E-02 | 1.06E+00 |
| ENSG000001 | OSCP1 | 2.26E-01 | 5.81E-01 | 1.13E+00 | 1.72E-01 | 1.13E+00 |
| ENSG000002 | TRIQK | 2.26E-01 | 5.81E-01 | 9.56E-01 | -6.54E-02 | -1.05E+00 |
| ENSG000001 | --- | 2.26E-01 | ? | 7.32E-01 | -4.50E-01 | -1.37E+00 |
| ENSG000001 | KIF21B | 2.26E-01 | 5.81E-01 | 8.53E-01 | -2.30E-01 | -1.17E+00 |
| ENSG000001 | PSMA6 | 2.26E-01 | 5.81E-01 | 9.66E-01 | -4.93E-02 | -1.03E+00 |
| ENSG000002 | --- | 2.26E-01 | ? | 7.09E-01 | -4.96E-01 | -1.41E+00 |
| ENSG000001 | DBF4B | 2.26E-01 | 5.81E-01 | 9.09E-01 | -1.38E-01 | -1.10E+00 |
| ENSG000002 | TLK2P1 | 2.26E-01 | ? | 7.91E-01 | -3.38E-01 | -1.26E+00 |
| ENSG00000C | AKAP11 | 2.26E-01 | 5.81E-01 | 9.50E-01 | -7.43E-02 | -1.05E+00 |
| ENSG000001 | CCDC137 | 2.26E-01 | 5.81E-01 | 1.05E+00 | 6.74E-02 | 1.05E+00 |
| ENSG000001 | POLR2L | 2.26E-01 | 5.81E-01 | 1.11E+00 | 1.51E-01 | 1.11E+00 |
| ENSG000002 | --- | 2.26E-01 | ? | 7.85E-01 | -3.50E-01 | -1.27E+00 |
| ENSG000001 | MAP9 | 2.26E-01 | 5.81E-01 | 9.42E-01 | -8.70E-02 | -1.06E+00 |
| ENSG000002 | --- | 2.26E-01 | ? | 1.59E+00 | 6.72E-01 | 1.59E+00 |
| ENSG000001 | SLC2A6 | 2.26E-01 | 5.81E-01 | 1.10E+00 | 1.42E-01 | 1.10E+00 |
| ENSG000002 | RAB6D | 2.26E-01 | ? | 1.50E+00 | 5.86E-01 | 1.50E+00 |
| ENSG000002 | --- | 2.26E-01 | ? | 1.24E+00 | 3.10E-01 | 1.24E+00 |
| ENSG000001 | GRK2 | 2.26E-01 | 5.81E-01 | 1.09E+00 | 1.18E-01 | 1.09E+00 |
| ENSG000001 | MOAP1 | 2.26E-01 | 5.81E-01 | 1.05E+00 | 6.47E-02 | 1.05E+00 |
| ENSG000001 | LINC02871 | 2.26E-01 | ? | 1.47E+00 | 5.55E-01 | 1.47E+00 |
| ENSG00000C | LUC7L | 2.26E-01 | 5.81E-01 | 9.42E-01 | -8.63E-02 | -1.06E+00 |
| ENSG000001 | NRGN | 2.26E-01 | ? | 7.33E-01 | -4.48E-01 | -1.36E+00 |
| ENSG000002 | --- | 2.26E-01 | 5.81E-01 | 1.17E+00 | 2.31E-01 | 1.17E+00 |
| ENSG000002 | SUCLG2P2 | 2.26E-01 | ? | 1.72E+00 | 7.84E-01 | 1.72E+00 |
| ENSG000001 | SF3B3 | 2.26E-01 | 5.81E-01 | 1.03E+00 | 4.24E-02 | 1.03E+00 |
| ENSG000001 | FNDC11 | 2.26E-01 | ? | 6.64E-01 | -5.92E-01 | -1.51E+00 |
| ENSG000001 | TRIM16L | 2.26E-01 | ? | 4.20E-01 | -1.25E+00 | -2.38E+00 |
| ENSG000002 | GOLGA2P5 | 2.27E-01 | 5.82E-01 | 8.96E-01 | -1.59E-01 | -1.12E+00 |
| ENSG000002 | ZBTB12 | 2.27E-01 | ? | 8.16E-01 | -2.93E-01 | -1.23E+00 |
| ENSG00000C | EIPR1 | 2.27E-01 | 5.82E-01 | 1.07E+00 | 9.71E-02 | 1.07E+00 |

| | | | | | | |
|---|---|---|---|---|---|---|
| ENSG000002 | --- | 2.27E-01 | 5.82E-01 | 1.25E+00 | 3.16E-01 | 1.25E+00 |
| ENSG000002 | HLA-DQB1-A | 2.27E-01 | ? | 5.13E-01 | -9.62E-01 | -1.95E+00 |
| ENSG000001 | ACVR2B | 2.27E-01 | 5.82E-01 | 1.07E+00 | 9.32E-02 | 1.07E+00 |
| ENSG00000C | DDX1 | 2.27E-01 | 5.82E-01 | 9.55E-01 | -6.57E-02 | -1.05E+00 |
| ENSG000002 | --- | 2.27E-01 | ? | 2.06E+00 | 1.04E+00 | 2.06E+00 |
| ENSG000001 | TUBA1C | 2.27E-01 | 5.82E-01 | 9.62E-01 | -5.59E-02 | -1.04E+00 |
| ENSG000002 | --- | 2.27E-01 | ? | 1.68E+00 | 7.53E-01 | 1.68E+00 |
| ENSG000001 | PSAP | 2.27E-01 | 5.83E-01 | 1.06E+00 | 8.02E-02 | 1.06E+00 |
| ENSG000001 | ZBTB37 | 2.27E-01 | 5.83E-01 | 9.23E-01 | -1.16E-01 | -1.08E+00 |
| ENSG000002 | --- | 2.27E-01 | ? | 8.05E-01 | -3.12E-01 | -1.24E+00 |
| ENSG00000C | GPC4 | 2.27E-01 | 5.83E-01 | 9.43E-01 | -8.53E-02 | -1.06E+00 |
| ENSG000001 | MFN1 | 2.27E-01 | 5.83E-01 | 9.62E-01 | -5.61E-02 | -1.04E+00 |
| ENSG000001 | FCRLA | 2.27E-01 | 5.83E-01 | 1.13E+00 | 1.76E-01 | 1.13E+00 |
| ENSG000002 | ZNNT1 | 2.27E-01 | 5.83E-01 | 8.88E-01 | -1.71E-01 | -1.13E+00 |
| ENSG000002 | LINC02457 | 2.27E-01 | ? | 7.73E-01 | -3.71E-01 | -1.29E+00 |
| ENSG000001 | ANXA11 | 2.27E-01 | 5.83E-01 | 1.04E+00 | 5.72E-02 | 1.04E+00 |
| ENSG000001 | SMAP1 | 2.28E-01 | 5.83E-01 | 9.66E-01 | -5.02E-02 | -1.04E+00 |
| ENSG000001 | NGF | 2.28E-01 | 5.83E-01 | 8.88E-01 | -1.71E-01 | -1.13E+00 |
| ENSG000001 | CDCA3 | 2.28E-01 | 5.83E-01 | 8.52E-01 | -2.31E-01 | -1.17E+00 |
| ENSG00000C | DTNBP1 | 2.28E-01 | 5.83E-01 | 1.09E+00 | 1.27E-01 | 1.09E+00 |
| ENSG000001 | NIBAN2 | 2.28E-01 | 5.83E-01 | 1.07E+00 | 9.75E-02 | 1.07E+00 |
| ENSG000002 | --- | 2.28E-01 | ? | 6.84E-01 | -5.49E-01 | -1.46E+00 |
| ENSG000001 | RBM10 | 2.28E-01 | 5.83E-01 | 1.04E+00 | 5.31E-02 | 1.04E+00 |
| ENSG000002 | RN7SKP116 | 2.28E-01 | ? | 6.34E-01 | -6.57E-01 | -1.58E+00 |
| ENSG000001 | NSRP1 | 2.28E-01 | 5.83E-01 | 9.31E-01 | -1.03E-01 | -1.07E+00 |
| ENSG000001 | IDH3A | 2.28E-01 | 5.83E-01 | 9.56E-01 | -6.56E-02 | -1.05E+00 |
| ENSG000001 | LAMB3 | 2.28E-01 | 5.83E-01 | 9.45E-01 | -8.17E-02 | -1.06E+00 |
| ENSG000002 | PPATP1 | 2.28E-01 | ? | 1.96E+00 | 9.73E-01 | 1.96E+00 |
| ENSG000001 | ELOVL7 | 2.28E-01 | 5.83E-01 | 1.10E+00 | 1.43E-01 | 1.10E+00 |
| ENSG00000C | METTL1 | 2.28E-01 | 5.83E-01 | 1.08E+00 | 1.11E-01 | 1.08E+00 |
| ENSG000001 | GFOD2 | 2.28E-01 | 5.83E-01 | 1.05E+00 | 6.81E-02 | 1.05E+00 |
| ENSG000002 | RPL26P30 | 2.28E-01 | ? | 1.45E+00 | 5.38E-01 | 1.45E+00 |
| ENSG000002 | RPL13P12 | 2.28E-01 | 5.83E-01 | 1.21E+00 | 2.75E-01 | 1.21E+00 |
| ENSG000001 | ASCC1 | 2.28E-01 | 5.83E-01 | 1.05E+00 | 7.02E-02 | 1.05E+00 |
| ENSG000002 | --- | 2.28E-01 | ? | 1.45E+00 | 5.35E-01 | 1.45E+00 |
| ENSG000001 | CXXC5 | 2.28E-01 | 5.83E-01 | 1.10E+00 | 1.36E-01 | 1.10E+00 |
| ENSG000001 | ZNF449 | 2.28E-01 | 5.83E-01 | 1.07E+00 | 9.30E-02 | 1.07E+00 |
| ENSG00000C | ATP11B | 2.28E-01 | 5.83E-01 | 9.58E-01 | -6.13E-02 | -1.04E+00 |
| ENSG000001 | NCF4 | 2.28E-01 | 5.83E-01 | 8.18E-01 | -2.90E-01 | -1.22E+00 |
| ENSG000002 | TMEM202-A: | 2.28E-01 | 5.83E-01 | 1.31E+00 | 3.89E-01 | 1.31E+00 |
| ENSG000002 | XPOTP1 | 2.28E-01 | ? | 1.83E+00 | 8.72E-01 | 1.83E+00 |
| ENSG00000C | CUL7 | 2.29E-01 | 5.83E-01 | 1.06E+00 | 8.52E-02 | 1.06E+00 |
| ENSG000002 | PRADX | 2.29E-01 | ? | 1.46E+00 | 5.47E-01 | 1.46E+00 |
| ENSG000001 | PKD1L1 | 2.29E-01 | ? | 7.92E-01 | -3.37E-01 | -1.26E+00 |
| ENSG000001 | TP53RK | 2.29E-01 | 5.84E-01 | 1.05E+00 | 6.85E-02 | 1.05E+00 |

| | | | | | | |
|---|---|---|---|---|---|---|
| ENSG000002 | LINC02977 | 2.29E-01 | 5.84E-01 | 9.08E-01 | -1.39E-01 | -1.10E+00 |
| ENSG000001 | PYCARD | 2.29E-01 | 5.84E-01 | 1.10E+00 | 1.38E-01 | 1.10E+00 |
| ENSG000001 | ZNF174 | 2.29E-01 | 5.84E-01 | 1.08E+00 | 1.13E-01 | 1.08E+00 |
| ENSG000002 | ZNF232-AS1 | 2.29E-01 | 5.84E-01 | 1.21E+00 | 2.71E-01 | 1.21E+00 |
| ENSG000002 | C2orf74 | 2.29E-01 | 5.84E-01 | 9.11E-01 | -1.34E-01 | -1.10E+00 |
| ENSG00000C | USP40 | 2.29E-01 | 5.84E-01 | 1.04E+00 | 5.05E-02 | 1.04E+00 |
| ENSG000002 | --- | 2.29E-01 | ? | 1.27E+00 | 3.45E-01 | 1.27E+00 |
| ENSG000001 | VIP | 2.29E-01 | ? | 6.54E-01 | -6.13E-01 | -1.53E+00 |
| ENSG000002 | TXNDC5 | 2.29E-01 | 5.84E-01 | 1.05E+00 | 6.79E-02 | 1.05E+00 |
| ENSG000001 | KLHDC10 | 2.29E-01 | 5.84E-01 | 9.64E-01 | -5.25E-02 | -1.04E+00 |
| ENSG000002 | --- | 2.29E-01 | 5.84E-01 | 9.22E-01 | -1.18E-01 | -1.08E+00 |
| ENSG000001 | FCF1 | 2.29E-01 | 5.84E-01 | 1.03E+00 | 4.50E-02 | 1.03E+00 |
| ENSG00000C | CLDND1 | 2.29E-01 | 5.84E-01 | 1.06E+00 | 8.85E-02 | 1.06E+00 |
| ENSG000001 | GNPAT | 2.29E-01 | 5.84E-01 | 1.04E+00 | 6.32E-02 | 1.04E+00 |
| ENSG000001 | SLC25A1 | 2.29E-01 | 5.84E-01 | 1.08E+00 | 1.13E-01 | 1.08E+00 |
| ENSG000001 | FAM220A | 2.29E-01 | 5.84E-01 | 1.06E+00 | 8.88E-02 | 1.06E+00 |
| ENSG000001 | NAB2 | 2.30E-01 | 5.84E-01 | 1.08E+00 | 1.10E-01 | 1.08E+00 |
| ENSG000001 | RANBP10 | 2.30E-01 | 5.84E-01 | 1.04E+00 | 5.38E-02 | 1.04E+00 |
| ENSG000001 | RGS5 | 2.30E-01 | 5.84E-01 | 9.00E-01 | -1.52E-01 | -1.11E+00 |
| ENSG000001 | TMED7 | 2.30E-01 | 5.84E-01 | 9.62E-01 | -5.63E-02 | -1.04E+00 |
| ENSG000002 | --- | 2.30E-01 | ? | 1.72E+00 | 7.79E-01 | 1.72E+00 |
| ENSG000002 | PIGW | 2.30E-01 | 5.84E-01 | 9.42E-01 | -8.58E-02 | -1.06E+00 |
| ENSG000002 | TYW1B | 2.30E-01 | 5.84E-01 | 8.58E-01 | -2.21E-01 | -1.17E+00 |
| ENSG000001 | C14orf119 | 2.30E-01 | 5.84E-01 | 1.04E+00 | 5.71E-02 | 1.04E+00 |
| ENSG000001 | PNKD | 2.30E-01 | 5.84E-01 | 1.06E+00 | 7.89E-02 | 1.06E+00 |
| ENSG000001 | SPATA32 | 2.30E-01 | ? | 1.41E+00 | 5.00E-01 | 1.41E+00 |
| ENSG000001 | POLK | 2.30E-01 | 5.84E-01 | 9.47E-01 | -7.83E-02 | -1.06E+00 |
| ENSG000001 | NOL3 | 2.30E-01 | 5.84E-01 | 1.08E+00 | 1.07E-01 | 1.08E+00 |
| ENSG000002 | VARS1 | 2.30E-01 | 5.84E-01 | 1.06E+00 | 8.55E-02 | 1.06E+00 |
| ENSG000002 | --- | 2.30E-01 | 5.84E-01 | 8.23E-01 | -2.81E-01 | -1.21E+00 |
| ENSG000001 | RIOX1 | 2.30E-01 | 5.84E-01 | 1.05E+00 | 7.71E-02 | 1.05E+00 |
| ENSG000002 | CPNE8-AS1 | 2.30E-01 | ? | 5.21E-01 | -9.41E-01 | -1.92E+00 |
| ENSG000001 | PURA | 2.30E-01 | 5.84E-01 | 1.06E+00 | 7.73E-02 | 1.06E+00 |
| ENSG000002 | --- | 2.30E-01 | ? | 1.40E+00 | 4.80E-01 | 1.40E+00 |
| ENSG000001 | --- | 2.30E-01 | 5.84E-01 | 8.07E-01 | -3.10E-01 | -1.24E+00 |
| ENSG000001 | COA4 | 2.30E-01 | 5.84E-01 | 1.05E+00 | 7.19E-02 | 1.05E+00 |
| ENSG000001 | SCARB2 | 2.30E-01 | 5.84E-01 | 1.04E+00 | 5.03E-02 | 1.04E+00 |
| ENSG000001 | DST | 2.30E-01 | 5.85E-01 | 1.05E+00 | 6.94E-02 | 1.05E+00 |
| ENSG000002 | --- | 2.30E-01 | 5.85E-01 | 1.32E+00 | 3.98E-01 | 1.32E+00 |
| ENSG000002 | --- | 2.30E-01 | 5.85E-01 | 8.48E-01 | -2.37E-01 | -1.18E+00 |
| ENSG000001 | TSPAN12 | 2.31E-01 | 5.85E-01 | 1.12E+00 | 1.66E-01 | 1.12E+00 |
| ENSG00000C | JKAMP | 2.31E-01 | 5.85E-01 | 9.63E-01 | -5.51E-02 | -1.04E+00 |
| ENSG000001 | TNFAIP3 | 2.31E-01 | 5.85E-01 | 1.08E+00 | 1.09E-01 | 1.08E+00 |
| ENSG000002 | NUDT19-DT | 2.31E-01 | ? | 1.22E+00 | 2.89E-01 | 1.22E+00 |
| ENSG000001 | CAPN5 | 2.31E-01 | 5.85E-01 | 9.18E-01 | -1.24E-01 | -1.09E+00 |

| | | | | | | |
|---|---|---|---|---|---|---|
| ENSG000002 | RPLP1P6 | 2.31E-01 | ? | 1.92E+00 | 9.41E-01 | 1.92E+00 |
| ENSG00000C | SIRT2 | 2.31E-01 | 5.85E-01 | 1.06E+00 | 7.90E-02 | 1.06E+00 |
| ENSG00000C | BCS1L | 2.31E-01 | 5.85E-01 | 1.07E+00 | 9.92E-02 | 1.07E+00 |
| ENSG000002 | DNAAF10 | 2.31E-01 | 5.85E-01 | 1.05E+00 | 7.16E-02 | 1.05E+00 |
| ENSG000001 | TMEM243 | 2.31E-01 | 5.85E-01 | 1.09E+00 | 1.24E-01 | 1.09E+00 |
| ENSG000001 | PARP14 | 2.31E-01 | 5.85E-01 | 9.49E-01 | -7.59E-02 | -1.05E+00 |
| ENSG000002 | MRM1 | 2.31E-01 | 5.85E-01 | 1.12E+00 | 1.68E-01 | 1.12E+00 |
| ENSG000002 | --- | 2.31E-01 | ? | 1.49E+00 | 5.73E-01 | 1.49E+00 |
| ENSG000002 | --- | 2.31E-01 | 5.85E-01 | 1.15E+00 | 2.04E-01 | 1.15E+00 |
| ENSG000002 | MIR4280HG | 2.31E-01 | 5.85E-01 | 8.12E-01 | -3.00E-01 | -1.23E+00 |
| ENSG000001 | SLC6A17 | 2.31E-01 | 5.86E-01 | 1.11E+00 | 1.45E-01 | 1.11E+00 |
| ENSG000001 | MT-ND3 | 2.31E-01 | 5.86E-01 | 9.24E-01 | -1.14E-01 | -1.08E+00 |
| ENSG000002 | CEBPD | 2.31E-01 | 5.86E-01 | 1.16E+00 | 2.11E-01 | 1.16E+00 |
| ENSG000002 | GMNC | 2.31E-01 | ? | 6.10E-01 | -7.13E-01 | -1.64E+00 |
| ENSG000002 | --- | 2.31E-01 | 5.86E-01 | 9.33E-01 | -1.01E-01 | -1.07E+00 |
| ENSG000002 | SCYL2P1 | 2.32E-01 | ? | 4.55E-01 | -1.14E+00 | -2.20E+00 |
| ENSG000002 | --- | 2.32E-01 | 5.86E-01 | 1.18E+00 | 2.35E-01 | 1.18E+00 |
| ENSG000001 | GALK2 | 2.32E-01 | 5.86E-01 | 9.47E-01 | -7.90E-02 | -1.06E+00 |
| ENSG000002 | KRT7-AS | 2.32E-01 | 5.86E-01 | 8.92E-01 | -1.64E-01 | -1.12E+00 |
| ENSG000001 | SLC25A38 | 2.32E-01 | 5.86E-01 | 1.05E+00 | 6.86E-02 | 1.05E+00 |
| ENSG00000C | SH3YL1 | 2.32E-01 | 5.86E-01 | 1.05E+00 | 6.99E-02 | 1.05E+00 |
| ENSG000001 | DDX49 | 2.32E-01 | 5.86E-01 | 1.07E+00 | 9.48E-02 | 1.07E+00 |
| ENSG00000C | SLC35C2 | 2.32E-01 | 5.86E-01 | 1.07E+00 | 9.56E-02 | 1.07E+00 |
| ENSG000001 | VGLL4 | 2.32E-01 | 5.86E-01 | 1.04E+00 | 5.45E-02 | 1.04E+00 |
| ENSG000001 | IRX5 | 2.32E-01 | 5.86E-01 | 9.19E-01 | -1.22E-01 | -1.09E+00 |
| ENSG000002 | --- | 2.32E-01 | 5.86E-01 | 8.39E-01 | -2.53E-01 | -1.19E+00 |
| ENSG000001 | SRSF1 | 2.32E-01 | 5.86E-01 | 9.67E-01 | -4.85E-02 | -1.03E+00 |
| ENSG000001 | KIN | 2.32E-01 | 5.86E-01 | 9.44E-01 | -8.36E-02 | -1.06E+00 |
| ENSG000002 | PKN2-AS1 | 2.32E-01 | ? | 1.52E+00 | 6.04E-01 | 1.52E+00 |
| ENSG000002 | --- | 2.32E-01 | ? | 6.36E-01 | -6.52E-01 | -1.57E+00 |
| ENSG000002 | UMAD1 | 2.32E-01 | 5.86E-01 | 1.05E+00 | 7.67E-02 | 1.05E+00 |
| ENSG000002 | RPLP0P6 | 2.32E-01 | 5.86E-01 | 1.11E+00 | 1.51E-01 | 1.11E+00 |
| ENSG000002 | LINC00663 | 2.32E-01 | 5.86E-01 | 1.23E+00 | 3.03E-01 | 1.23E+00 |
| ENSG000002 | LINC01801 | 2.32E-01 | ? | 1.57E+00 | 6.55E-01 | 1.57E+00 |
| ENSG000001 | ACSS2 | 2.32E-01 | 5.86E-01 | 1.06E+00 | 8.97E-02 | 1.06E+00 |
| ENSG000002 | --- | 2.32E-01 | ? | 1.39E+00 | 4.76E-01 | 1.39E+00 |
| ENSG000001 | MXD4 | 2.32E-01 | 5.86E-01 | 1.07E+00 | 1.02E-01 | 1.07E+00 |
| ENSG000002 | CFLAR-AS1 | 2.32E-01 | ? | 7.70E-01 | -3.78E-01 | -1.30E+00 |
| ENSG000001 | BMP8A | 2.33E-01 | 5.86E-01 | 8.52E-01 | -2.32E-01 | -1.17E+00 |
| ENSG000002 | CEP95 | 2.33E-01 | 5.86E-01 | 9.41E-01 | -8.76E-02 | -1.06E+00 |
| ENSG000002 | SLURP2 | 2.33E-01 | ? | 5.82E-01 | -7.81E-01 | -1.72E+00 |
| ENSG000002 | --- | 2.33E-01 | 5.86E-01 | 7.40E-01 | -4.35E-01 | -1.35E+00 |
| ENSG000001 | CISD1 | 2.33E-01 | 5.86E-01 | 1.05E+00 | 7.08E-02 | 1.05E+00 |
| ENSG00000C | IL20RA | 2.33E-01 | 5.86E-01 | 1.20E+00 | 2.58E-01 | 1.20E+00 |
| ENSG000002 | TMEM14B-D | 2.33E-01 | ? | 1.46E+00 | 5.46E-01 | 1.46E+00 |

| | | | | | | |
|---|---|---|---|---|---|---|
| ENSG000002 | ARL16 | 2.33E-01 | 5.86E-01 | 9.42E-01 | -8.57E-02 | -1.06E+00 |
| ENSG000001 | NBAS | 2.33E-01 | 5.86E-01 | 9.62E-01 | -5.53E-02 | -1.04E+00 |
| ENSG000002 | KRT8P36 | 2.33E-01 | 5.86E-01 | 8.72E-01 | -1.97E-01 | -1.15E+00 |
| ENSG000001 | LRP3 | 2.33E-01 | 5.86E-01 | 1.15E+00 | 2.03E-01 | 1.15E+00 |
| ENSG000001 | TLE5 | 2.33E-01 | 5.87E-01 | 1.05E+00 | 6.47E-02 | 1.05E+00 |
| ENSG000002 | DDX3X | 2.33E-01 | 5.87E-01 | 9.72E-01 | -4.14E-02 | -1.03E+00 |
| ENSG000002 | PRR22 | 2.33E-01 | ? | 7.66E-01 | -3.84E-01 | -1.31E+00 |
| ENSG000001 | RNF123 | 2.33E-01 | 5.87E-01 | 1.07E+00 | 1.03E-01 | 1.07E+00 |
| ENSG000001 | PKIG | 2.33E-01 | 5.87E-01 | 1.11E+00 | 1.55E-01 | 1.11E+00 |
| ENSG000001 | TAGAP | 2.33E-01 | ? | 1.66E+00 | 7.33E-01 | 1.66E+00 |
| ENSG000001 | NDUFAF5 | 2.33E-01 | 5.87E-01 | 9.37E-01 | -9.36E-02 | -1.07E+00 |
| ENSG000001 | FAXC | 2.33E-01 | 5.87E-01 | 9.33E-01 | -9.98E-02 | -1.07E+00 |
| ENSG000002 | --- | 2.33E-01 | 5.87E-01 | 9.02E-01 | -1.49E-01 | -1.11E+00 |
| ENSG000001 | CHN1 | 2.33E-01 | 5.87E-01 | 8.68E-01 | -2.04E-01 | -1.15E+00 |
| ENSG000002 | LINC01293 | 2.33E-01 | ? | 6.83E-01 | -5.51E-01 | -1.47E+00 |
| ENSG000001 | PCBD1 | 2.33E-01 | 5.87E-01 | 1.05E+00 | 6.82E-02 | 1.05E+00 |
| ENSG000001 | LLGL1 | 2.33E-01 | 5.87E-01 | 1.06E+00 | 8.82E-02 | 1.06E+00 |
| ENSG000002 | ALKBH3-AS1 | 2.33E-01 | ? | 1.57E+00 | 6.49E-01 | 1.57E+00 |
| ENSG000002 | FIRRE | 2.33E-01 | ? | 8.19E-01 | -2.88E-01 | -1.22E+00 |
| ENSG000001 | DEFB103A | 2.33E-01 | ? | 6.95E-01 | -5.26E-01 | -1.44E+00 |
| ENSG000001 | DSC3 | 2.33E-01 | 5.87E-01 | 9.57E-01 | -6.34E-02 | -1.04E+00 |
| ENSG000001 | SEPTIN1 | 2.34E-01 | ? | 1.59E+00 | 6.65E-01 | 1.59E+00 |
| ENSG000001 | SGCE | 2.34E-01 | 5.87E-01 | 1.10E+00 | 1.35E-01 | 1.10E+00 |
| ENSG000001 | NAPG | 2.34E-01 | 5.87E-01 | 9.43E-01 | -8.48E-02 | -1.06E+00 |
| ENSG000002 | --- | 2.34E-01 | 5.87E-01 | 1.08E+00 | 1.12E-01 | 1.08E+00 |
| ENSG000002 | LINC00115 | 2.34E-01 | ? | 7.89E-01 | -3.42E-01 | -1.27E+00 |
| ENSG00000C | ELMO2 | 2.34E-01 | 5.87E-01 | 1.03E+00 | 4.60E-02 | 1.03E+00 |
| ENSG000001 | PRKCZ-AS1 | 2.34E-01 | 5.87E-01 | 8.31E-01 | -2.67E-01 | -1.20E+00 |
| ENSG000001 | ALDH3B2 | 2.34E-01 | ? | 1.27E+00 | 3.44E-01 | 1.27E+00 |
| ENSG000001 | ALDH1A2 | 2.34E-01 | ? | 1.40E+00 | 4.83E-01 | 1.40E+00 |
| ENSG000001 | UBE2G2 | 2.34E-01 | 5.87E-01 | 9.58E-01 | -6.23E-02 | -1.04E+00 |
| ENSG000002 | --- | 2.34E-01 | ? | 7.66E-01 | -3.84E-01 | -1.30E+00 |
| ENSG000002 | --- | 2.34E-01 | ? | 7.10E-01 | -4.95E-01 | -1.41E+00 |
| ENSG000001 | MRPL37 | 2.34E-01 | 5.87E-01 | 9.63E-01 | -5.37E-02 | -1.04E+00 |
| ENSG000002 | TRIM16 | 2.34E-01 | 5.87E-01 | 1.06E+00 | 8.14E-02 | 1.06E+00 |
| ENSG00000C | TBC1D2 | 2.34E-01 | 5.87E-01 | 1.06E+00 | 8.42E-02 | 1.06E+00 |
| ENSG00000C | CHI3L2 | 2.34E-01 | 5.87E-01 | 1.21E+00 | 2.79E-01 | 1.21E+00 |
| ENSG000002 | SCX | 2.34E-01 | ? | 7.51E-01 | -4.14E-01 | -1.33E+00 |
| ENSG000001 | CNGA3 | 2.34E-01 | ? | 1.61E+00 | 6.84E-01 | 1.61E+00 |
| ENSG000002 | --- | 2.34E-01 | ? | 1.53E+00 | 6.14E-01 | 1.53E+00 |
| ENSG000001 | CTSL | 2.34E-01 | 5.87E-01 | 9.70E-01 | -4.35E-02 | -1.03E+00 |
| ENSG000001 | CDCA5 | 2.34E-01 | 5.87E-01 | 8.55E-01 | -2.25E-01 | -1.17E+00 |
| ENSG000002 | NPIPB5 | 2.34E-01 | 5.87E-01 | 8.71E-01 | -2.00E-01 | -1.15E+00 |
| ENSG000001 | SF3A3 | 2.34E-01 | 5.87E-01 | 1.04E+00 | 6.16E-02 | 1.04E+00 |
| ENSG000001 | SATB2 | 2.34E-01 | 5.87E-01 | 9.27E-01 | -1.09E-01 | -1.08E+00 |

| | | | | | | | |
|---|---|---|---|---|---|---|---|
| ENSG000001 | ZNF365 | 2.34E-01 | | 5.87E-01 | 8.76E-01 | -1.91E-01 | -1.14E+00 |
| ENSG000001 | MOCS2 | 2.34E-01 | | 5.87E-01 | 9.46E-01 | -8.02E-02 | -1.06E+00 |
| ENSG000001 | GIGYF1 | 2.34E-01 | | 5.87E-01 | 1.08E+00 | 1.17E-01 | 1.08E+00 |
| ENSG000002 | LYRM4 | 2.34E-01 | | 5.87E-01 | 1.06E+00 | 8.22E-02 | 1.06E+00 |
| ENSG000002 | IQCJ-SCHIP1 | 2.34E-01 | | 5.87E-01 | 1.06E+00 | 9.06E-02 | 1.06E+00 |
| ENSG000002 | --- | 2.34E-01 | ? | | 6.07E-01 | -7.20E-01 | -1.65E+00 |
| ENSG000001 | TMEM262 | 2.34E-01 | | 5.87E-01 | 8.00E-01 | -3.21E-01 | -1.25E+00 |
| ENSG000001 | LY96 | 2.34E-01 | | 5.87E-01 | 7.92E-01 | -3.36E-01 | -1.26E+00 |
| ENSG000001 | ANGPTL4 | 2.34E-01 | | 5.87E-01 | 8.32E-01 | -2.65E-01 | -1.20E+00 |
| ENSG000001 | SUPT7L | 2.34E-01 | | 5.87E-01 | 9.55E-01 | -6.58E-02 | -1.05E+00 |
| ENSG000002 | --- | 2.35E-01 | ? | | 6.81E-01 | -5.54E-01 | -1.47E+00 |
| ENSG000001 | HTT | 2.35E-01 | | 5.87E-01 | 1.05E+00 | 6.40E-02 | 1.05E+00 |
| ENSG000001 | BCAP31 | 2.35E-01 | | 5.87E-01 | 1.04E+00 | 5.63E-02 | 1.04E+00 |
| ENSG000001 | RASSF3 | 2.35E-01 | | 5.87E-01 | 1.05E+00 | 7.33E-02 | 1.05E+00 |
| ENSG000001 | C16orf46 | 2.35E-01 | ? | | 1.46E+00 | 5.42E-01 | 1.46E+00 |
| ENSG000002 | --- | 2.35E-01 | ? | | 1.25E+00 | 3.21E-01 | 1.25E+00 |
| ENSG000002 | ZNF426-DT | 2.35E-01 | | 5.87E-01 | 8.19E-01 | -2.89E-01 | -1.22E+00 |
| ENSG000002 | OR2H2 | 2.35E-01 | ? | | 5.38E-01 | -8.96E-01 | -1.86E+00 |
| ENSG000002 | S1PR3 | 2.35E-01 | | 5.87E-01 | 1.08E+00 | 1.06E-01 | 1.08E+00 |
| ENSG000001 | SEMA4A | 2.35E-01 | | 5.87E-01 | 1.08E+00 | 1.07E-01 | 1.08E+00 |
| ENSG000001 | AKAP10 | 2.35E-01 | | 5.87E-01 | 9.67E-01 | -4.89E-02 | -1.03E+00 |
| ENSG000002 | --- | 2.35E-01 | | 5.87E-01 | 8.29E-01 | -2.70E-01 | -1.21E+00 |
| ENSG000001 | RUVBL2 | 2.35E-01 | | 5.87E-01 | 1.05E+00 | 6.73E-02 | 1.05E+00 |
| ENSG000001 | FAM43A | 2.35E-01 | | 5.87E-01 | 1.21E+00 | 2.78E-01 | 1.21E+00 |
| ENSG000001 | UBFD1 | 2.35E-01 | | 5.87E-01 | 1.03E+00 | 4.77E-02 | 1.03E+00 |
| ENSG000002 | LINC02680 | 2.35E-01 | ? | | 6.76E-01 | -5.65E-01 | -1.48E+00 |
| ENSG000001 | PXDN | 2.35E-01 | | 5.87E-01 | 1.04E+00 | 5.13E-02 | 1.04E+00 |
| ENSG000002 | --- | 2.35E-01 | ? | | 7.94E-01 | -3.34E-01 | -1.26E+00 |
| ENSG000001 | DUSP8 | 2.35E-01 | | 5.87E-01 | 1.10E+00 | 1.41E-01 | 1.10E+00 |
| ENSG000001 | PRR32 | 2.35E-01 | ? | | 1.60E+00 | 6.76E-01 | 1.60E+00 |
| ENSG000001 | NT5C | 2.35E-01 | | 5.87E-01 | 1.07E+00 | 9.92E-02 | 1.07E+00 |
| ENSG000001 | ARHGAP20 | 2.35E-01 | ? | | 6.32E-01 | -6.63E-01 | -1.58E+00 |
| ENSG000002 | ECSCR | 2.35E-01 | ? | | 5.34E-01 | -9.06E-01 | -1.87E+00 |
| ENSG000001 | COPS5 | 2.35E-01 | | 5.87E-01 | 1.04E+00 | 6.24E-02 | 1.04E+00 |
| ENSG000001 | BCAS2 | 2.35E-01 | | 5.87E-01 | 9.53E-01 | -6.96E-02 | -1.05E+00 |
| ENSG000001 | MEIS2 | 2.35E-01 | | 5.87E-01 | 1.08E+00 | 1.10E-01 | 1.08E+00 |
| ENSG000001 | TRIP11 | 2.35E-01 | | 5.87E-01 | 9.45E-01 | -8.14E-02 | -1.06E+00 |
| ENSG000001 | RRS1 | 2.35E-01 | | 5.87E-01 | 1.05E+00 | 6.62E-02 | 1.05E+00 |
| ENSG000001 | GCSAM | 2.36E-01 | | 5.87E-01 | 8.87E-01 | -1.73E-01 | -1.13E+00 |
| ENSG000002 | SNHG8 | 2.36E-01 | | 5.88E-01 | 1.09E+00 | 1.18E-01 | 1.09E+00 |
| ENSG00000C | CENPQ | 2.36E-01 | | 5.88E-01 | 9.12E-01 | -1.32E-01 | -1.10E+00 |
| ENSG000001 | TPRG1L | 2.36E-01 | | 5.88E-01 | 1.04E+00 | 5.45E-02 | 1.04E+00 |
| ENSG000001 | PPM1J | 2.36E-01 | | 5.88E-01 | 1.10E+00 | 1.41E-01 | 1.10E+00 |
| ENSG000001 | PTRH1 | 2.36E-01 | | 5.88E-01 | 1.16E+00 | 2.12E-01 | 1.16E+00 |
| ENSG000002 | CD44-AS1 | 2.36E-01 | ? | | 7.61E-01 | -3.94E-01 | -1.31E+00 |

| | | | | | | |
|---|---|---|---|---|---|---|
| ENSG000001 | GBE1 | 2.36E-01 | 5.88E-01 | 9.59E-01 | -6.10E-02 | -1.04E+00 |
| ENSG000001 | NR2F2 | 2.36E-01 | 5.88E-01 | 9.57E-01 | -6.30E-02 | -1.04E+00 |
| ENSG000002 | VSTM5 | 2.36E-01 | 5.88E-01 | 8.55E-01 | -2.26E-01 | -1.17E+00 |
| ENSG000001 | FAM111B | 2.36E-01 | 5.88E-01 | 8.30E-01 | -2.69E-01 | -1.21E+00 |
| ENSG000002 | --- | 2.37E-01 | ? | 1.38E+00 | 4.68E-01 | 1.38E+00 |
| ENSG000001 | SLC43A3 | 2.37E-01 | 5.89E-01 | 9.32E-01 | -1.02E-01 | -1.07E+00 |
| ENSG000002 | --- | 2.37E-01 | 5.89E-01 | 8.13E-01 | -2.98E-01 | -1.23E+00 |
| ENSG000002 | PRRT1 | 2.37E-01 | ? | 6.35E-01 | -6.55E-01 | -1.57E+00 |
| ENSG000001 | SMURF2 | 2.37E-01 | 5.89E-01 | 9.37E-01 | -9.32E-02 | -1.07E+00 |
| ENSG000002 | H2BC18 | 2.37E-01 | ? | 8.23E-01 | -2.81E-01 | -1.22E+00 |
| ENSG00000C | MCOLN1 | 2.37E-01 | 5.89E-01 | 9.39E-01 | -9.13E-02 | -1.07E+00 |
| ENSG000002 | --- | 2.37E-01 | ? | 5.35E-01 | -9.03E-01 | -1.87E+00 |
| ENSG000002 | --- | 2.37E-01 | ? | 6.80E-01 | -5.56E-01 | -1.47E+00 |
| ENSG000001 | MAT2A | 2.37E-01 | 5.89E-01 | 9.50E-01 | -7.33E-02 | -1.05E+00 |
| ENSG000001 | EMC8 | 2.37E-01 | 5.89E-01 | 1.05E+00 | 7.22E-02 | 1.05E+00 |
| ENSG000002 | --- | 2.37E-01 | ? | 5.34E-01 | -9.06E-01 | -1.87E+00 |
| ENSG000001 | BNIP3L | 2.37E-01 | 5.89E-01 | 9.59E-01 | -5.99E-02 | -1.04E+00 |
| ENSG000001 | CDYL2 | 2.37E-01 | 5.89E-01 | 9.52E-01 | -7.02E-02 | -1.05E+00 |
| ENSG000001 | TRA2B | 2.37E-01 | 5.89E-01 | 9.65E-01 | -5.16E-02 | -1.04E+00 |
| ENSG000001 | C6orf52 | 2.37E-01 | ? | 1.41E+00 | 4.98E-01 | 1.41E+00 |
| ENSG000002 | PGM5P2 | 2.37E-01 | 5.89E-01 | 1.19E+00 | 2.46E-01 | 1.19E+00 |
| ENSG000001 | NEFM | 2.37E-01 | 5.90E-01 | 8.70E-01 | -2.01E-01 | -1.15E+00 |
| ENSG00000C | GNAO1 | 2.37E-01 | ? | 1.33E+00 | 4.09E-01 | 1.33E+00 |
| ENSG000001 | PEX11A | 2.37E-01 | 5.90E-01 | 1.10E+00 | 1.38E-01 | 1.10E+00 |
| ENSG000001 | AMDHD2 | 2.38E-01 | 5.90E-01 | 1.11E+00 | 1.49E-01 | 1.11E+00 |
| ENSG000001 | PDLIM3 | 2.38E-01 | ? | 1.59E+00 | 6.65E-01 | 1.59E+00 |
| ENSG000002 | TMEM18-DT | 2.38E-01 | ? | 1.85E+00 | 8.89E-01 | 1.85E+00 |
| ENSG00000C | MRC2 | 2.38E-01 | 5.90E-01 | 1.08E+00 | 1.10E-01 | 1.08E+00 |
| ENSG000001 | CDT1 | 2.38E-01 | 5.90E-01 | 8.39E-01 | -2.54E-01 | -1.19E+00 |
| ENSG000001 | RAB35 | 2.38E-01 | 5.90E-01 | 1.04E+00 | 5.34E-02 | 1.04E+00 |
| ENSG00000C | ST7 | 2.38E-01 | 5.90E-01 | 9.46E-01 | -8.04E-02 | -1.06E+00 |
| ENSG00000C | DHX32 | 2.38E-01 | 5.90E-01 | 1.04E+00 | 5.53E-02 | 1.04E+00 |
| ENSG000002 | --- | 2.38E-01 | ? | 6.09E-01 | -7.15E-01 | -1.64E+00 |
| ENSG000001 | FAM110C | 2.38E-01 | 5.90E-01 | 9.40E-01 | -8.87E-02 | -1.06E+00 |
| ENSG000002 | --- | 2.38E-01 | ? | 5.92E-01 | -7.56E-01 | -1.69E+00 |
| ENSG000002 | --- | 2.38E-01 | 5.90E-01 | 1.39E+00 | 4.79E-01 | 1.39E+00 |
| ENSG000001 | SCN8A | 2.38E-01 | ? | 1.24E+00 | 3.06E-01 | 1.24E+00 |
| ENSG000001 | DCAF16 | 2.38E-01 | 5.91E-01 | 1.04E+00 | 6.17E-02 | 1.04E+00 |
| ENSG000002 | ITGB1-DT | 2.38E-01 | 5.91E-01 | 8.82E-01 | -1.80E-01 | -1.13E+00 |
| ENSG000001 | DAP3 | 2.38E-01 | 5.91E-01 | 1.03E+00 | 3.68E-02 | 1.03E+00 |
| ENSG000002 | --- | 2.38E-01 | ? | 6.53E-01 | -6.15E-01 | -1.53E+00 |
| ENSG000002 | --- | 2.38E-01 | ? | 1.41E+00 | 4.99E-01 | 1.41E+00 |
| ENSG000001 | FBXO46 | 2.38E-01 | 5.91E-01 | 1.07E+00 | 1.02E-01 | 1.07E+00 |
| ENSG000001 | THEG | 2.38E-01 | 5.91E-01 | 1.23E+00 | 2.96E-01 | 1.23E+00 |
| ENSG00000C | RUNX3 | 2.38E-01 | 5.91E-01 | 8.71E-01 | -2.00E-01 | -1.15E+00 |

| | | | | | | |
|---|---|---|---|---|---|---|
| ENSG000001 | DHRS13 | 2.38E-01 | ? | 7.32E-01 | -4.50E-01 | -1.37E+00 |
| ENSG000001 | BFSP1 | 2.39E-01 | 5.91E-01 | 1.06E+00 | 9.06E-02 | 1.06E+00 |
| ENSG000001 | TMPRSS4 | 2.39E-01 | 5.91E-01 | 1.13E+00 | 1.75E-01 | 1.13E+00 |
| ENSG000001 | TRIM28 | 2.39E-01 | 5.91E-01 | 1.06E+00 | 9.01E-02 | 1.06E+00 |
| ENSG000002 | NUDT16L2P | 2.39E-01 | 5.91E-01 | 1.09E+00 | 1.25E-01 | 1.09E+00 |
| ENSG000001 | YTHDF3 | 2.39E-01 | 5.91E-01 | 9.62E-01 | -5.63E-02 | -1.04E+00 |
| ENSG000002 | --- | 2.39E-01 | ? | 7.68E-01 | -3.82E-01 | -1.30E+00 |
| ENSG000002 | --- | 2.39E-01 | ? | 1.42E+00 | 5.04E-01 | 1.42E+00 |
| ENSG000002 | --- | 2.39E-01 | ? | 1.28E+00 | 3.54E-01 | 1.28E+00 |
| ENSG000002 | --- | 2.39E-01 | ? | 6.97E-01 | -5.21E-01 | -1.44E+00 |
| ENSG000001 | SNTA1 | 2.39E-01 | 5.91E-01 | 1.11E+00 | 1.44E-01 | 1.11E+00 |
| ENSG000001 | YARS1 | 2.39E-01 | 5.91E-01 | 1.03E+00 | 4.56E-02 | 1.03E+00 |
| ENSG000002 | ZNF321P | 2.39E-01 | 5.91E-01 | 1.13E+00 | 1.82E-01 | 1.13E+00 |
| ENSG000002 | DUSP8P5 | 2.39E-01 | ? | 7.95E-01 | -3.30E-01 | -1.26E+00 |
| ENSG000001 | AAK1 | 2.39E-01 | 5.91E-01 | 1.06E+00 | 8.37E-02 | 1.06E+00 |
| ENSG000002 | BZW1-AS1 | 2.39E-01 | ? | 7.44E-01 | -4.27E-01 | -1.34E+00 |
| ENSG000001 | ASS1 | 2.39E-01 | 5.92E-01 | 9.16E-01 | -1.27E-01 | -1.09E+00 |
| ENSG000002 | --- | 2.39E-01 | ? | 7.55E-01 | -4.06E-01 | -1.32E+00 |
| ENSG000001 | NUDT8 | 2.39E-01 | 5.92E-01 | 1.18E+00 | 2.37E-01 | 1.18E+00 |
| ENSG000001 | TBC1D8B | 2.39E-01 | 5.92E-01 | 9.55E-01 | -6.69E-02 | -1.05E+00 |
| ENSG000002 | TTC4 | 2.39E-01 | 5.92E-01 | 1.04E+00 | 5.09E-02 | 1.04E+00 |
| ENSG000001 | MT1X | 2.40E-01 | 5.92E-01 | 8.82E-01 | -1.81E-01 | -1.13E+00 |
| ENSG000002 | STON1 | 2.40E-01 | 5.92E-01 | 1.23E+00 | 3.01E-01 | 1.23E+00 |
| ENSG000001 | GJB2 | 2.40E-01 | 5.92E-01 | 9.12E-01 | -1.33E-01 | -1.10E+00 |
| ENSG000001 | NRG2 | 2.40E-01 | ? | 7.01E-01 | -5.13E-01 | -1.43E+00 |
| ENSG000001 | SPSB2 | 2.40E-01 | 5.92E-01 | 1.11E+00 | 1.53E-01 | 1.11E+00 |
| ENSG000001 | C17orf80 | 2.40E-01 | 5.92E-01 | 9.53E-01 | -6.94E-02 | -1.05E+00 |
| ENSG000001 | EXOC6 | 2.40E-01 | 5.92E-01 | 9.43E-01 | -8.49E-02 | -1.06E+00 |
| ENSG000001 | NID1 | 2.40E-01 | ? | 1.23E+00 | 2.97E-01 | 1.23E+00 |
| ENSG000001 | ACTR1B | 2.40E-01 | 5.92E-01 | 1.08E+00 | 1.12E-01 | 1.08E+00 |
| ENSG000001 | ZBTB49 | 2.40E-01 | 5.92E-01 | 8.88E-01 | -1.72E-01 | -1.13E+00 |
| ENSG000001 | FAM53A | 2.40E-01 | ? | 1.26E+00 | 3.38E-01 | 1.26E+00 |
| ENSG000002 | RAP2C-AS1 | 2.40E-01 | 5.93E-01 | 9.03E-01 | -1.48E-01 | -1.11E+00 |
| ENSG000001 | NRL | 2.40E-01 | ? | 7.70E-01 | -3.78E-01 | -1.30E+00 |
| ENSG000002 | WDR46 | 2.40E-01 | 5.93E-01 | 1.05E+00 | 6.48E-02 | 1.05E+00 |
| ENSG000001 | KLF2 | 2.40E-01 | ? | 1.72E+00 | 7.79E-01 | 1.72E+00 |
| ENSG000001 | FZD8 | 2.40E-01 | 5.93E-01 | 9.04E-01 | -1.45E-01 | -1.11E+00 |
| ENSG00000C | OPHN1 | 2.40E-01 | 5.93E-01 | 9.18E-01 | -1.23E-01 | -1.09E+00 |
| ENSG000001 | C1orf43 | 2.40E-01 | 5.93E-01 | 1.05E+00 | 6.52E-02 | 1.05E+00 |
| ENSG000001 | LINC02870 | 2.40E-01 | ? | 7.83E-01 | -3.53E-01 | -1.28E+00 |
| ENSG000001 | CYP27C1 | 2.40E-01 | 5.93E-01 | 8.95E-01 | -1.60E-01 | -1.12E+00 |
| ENSG000001 | CTIF | 2.40E-01 | 5.93E-01 | 1.06E+00 | 7.96E-02 | 1.06E+00 |
| ENSG000002 | --- | 2.40E-01 | 5.93E-01 | 8.24E-01 | -2.80E-01 | -1.21E+00 |
| ENSG00000C | REST | 2.40E-01 | 5.93E-01 | 9.69E-01 | -4.61E-02 | -1.03E+00 |
| ENSG000002 | PPP1R26-AS1 | 2.41E-01 | ? | 1.24E+00 | 3.09E-01 | 1.24E+00 |

| | | | | | | |
|---|---|---|---|---|---|---|
| ENSG000001 | HSD3BP5 | 2.41E-01 | ? | 7.23E-01 | -4.68E-01 | -1.38E+00 |
| ENSG000001 | LRRC70 | 2.41E-01 | ? | 1.89E+00 | 9.21E-01 | 1.89E+00 |
| ENSG00000C | YTHDC1 | 2.41E-01 | 5.93E-01 | 9.51E-01 | -7.25E-02 | -1.05E+00 |
| ENSG000002 | PPP1R11 | 2.41E-01 | 5.93E-01 | 9.67E-01 | -4.80E-02 | -1.03E+00 |
| ENSG000001 | PKIB | 2.41E-01 | 5.93E-01 | 1.13E+00 | 1.72E-01 | 1.13E+00 |
| ENSG000001 | ST3GAL2 | 2.41E-01 | 5.93E-01 | 1.06E+00 | 7.79E-02 | 1.06E+00 |
| ENSG000001 | SPDYA | 2.41E-01 | ? | 7.44E-01 | -4.26E-01 | -1.34E+00 |
| ENSG000001 | CAPZA2 | 2.41E-01 | 5.93E-01 | 1.03E+00 | 4.37E-02 | 1.03E+00 |
| ENSG000001 | PKNOX1 | 2.41E-01 | 5.93E-01 | 1.04E+00 | 5.11E-02 | 1.04E+00 |
| ENSG000002 | --- | 2.41E-01 | ? | 1.32E+00 | 4.04E-01 | 1.32E+00 |
| ENSG000001 | IFI6 | 2.41E-01 | 5.93E-01 | 8.33E-01 | -2.64E-01 | -1.20E+00 |
| ENSG00000C | MAP4K5 | 2.41E-01 | 5.93E-01 | 9.59E-01 | -6.02E-02 | -1.04E+00 |
| ENSG000001 | PEF1 | 2.41E-01 | 5.93E-01 | 1.04E+00 | 5.88E-02 | 1.04E+00 |
| ENSG000002 | SAGSIN1 | 2.41E-01 | 5.93E-01 | 1.06E+00 | 8.72E-02 | 1.06E+00 |
| ENSG000001 | TMTC2 | 2.41E-01 | 5.93E-01 | 9.29E-01 | -1.06E-01 | -1.08E+00 |
| ENSG00000C | STX1B | 2.41E-01 | 5.93E-01 | 1.19E+00 | 2.45E-01 | 1.19E+00 |
| ENSG000002 | CCDC188 | 2.41E-01 | ? | 6.13E-01 | -7.07E-01 | -1.63E+00 |
| ENSG000002 | PCDHGC3 | 2.41E-01 | 5.93E-01 | 1.06E+00 | 8.32E-02 | 1.06E+00 |
| ENSG000002 | PCDHGA1 | 2.41E-01 | ? | 1.43E+00 | 5.18E-01 | 1.43E+00 |
| ENSG000002 | --- | 2.41E-01 | 5.93E-01 | 9.12E-01 | -1.33E-01 | -1.10E+00 |
| ENSG00000C | CNTN1 | 2.41E-01 | 5.93E-01 | 8.96E-01 | -1.59E-01 | -1.12E+00 |
| ENSG000001 | ARHGAP27 | 2.41E-01 | 5.93E-01 | 1.06E+00 | 7.79E-02 | 1.06E+00 |
| ENSG000001 | MYD88 | 2.41E-01 | 5.93E-01 | 9.75E-01 | -3.70E-02 | -1.03E+00 |
| ENSG000001 | KIF18B | 2.41E-01 | 5.93E-01 | 8.24E-01 | -2.80E-01 | -1.21E+00 |
| ENSG000001 | OGFRP1 | 2.41E-01 | 5.93E-01 | 1.06E+00 | 9.06E-02 | 1.06E+00 |
| ENSG000001 | SDHB | 2.42E-01 | 5.94E-01 | 9.60E-01 | -5.87E-02 | -1.04E+00 |
| ENSG000001 | POGZ | 2.42E-01 | 5.94E-01 | 9.60E-01 | -5.88E-02 | -1.04E+00 |
| ENSG000002 | EML2-AS1 | 2.42E-01 | ? | 2.32E+00 | 1.22E+00 | 2.32E+00 |
| ENSG000001 | ARMCX6 | 2.42E-01 | 5.94E-01 | 1.05E+00 | 7.70E-02 | 1.05E+00 |
| ENSG000002 | --- | 2.42E-01 | ? | 5.35E-01 | -9.02E-01 | -1.87E+00 |
| ENSG000001 | GSK3A | 2.42E-01 | 5.94E-01 | 1.04E+00 | 4.97E-02 | 1.04E+00 |
| ENSG000001 | SIM2 | 2.42E-01 | ? | 1.34E+00 | 4.25E-01 | 1.34E+00 |
| ENSG000002 | --- | 2.42E-01 | ? | 1.62E+00 | 6.95E-01 | 1.62E+00 |
| ENSG000001 | AK9 | 2.42E-01 | 5.94E-01 | 8.93E-01 | -1.64E-01 | -1.12E+00 |
| ENSG000001 | ID1 | 2.42E-01 | 5.94E-01 | 7.70E-01 | -3.76E-01 | -1.30E+00 |
| ENSG000001 | AP2A1 | 2.42E-01 | 5.94E-01 | 1.07E+00 | 9.13E-02 | 1.07E+00 |
| ENSG000001 | SYT2 | 2.42E-01 | ? | 7.54E-01 | -4.07E-01 | -1.33E+00 |
| ENSG000002 | --- | 2.42E-01 | 5.95E-01 | 8.49E-01 | -2.36E-01 | -1.18E+00 |
| ENSG000001 | SLC50A1 | 2.42E-01 | 5.95E-01 | 1.06E+00 | 8.05E-02 | 1.06E+00 |
| ENSG000002 | --- | 2.42E-01 | ? | 1.61E+00 | 6.85E-01 | 1.61E+00 |
| ENSG000002 | IKBKE | 2.42E-01 | 5.95E-01 | 1.07E+00 | 9.95E-02 | 1.07E+00 |
| ENSG000001 | SNAP47 | 2.43E-01 | 5.95E-01 | 1.08E+00 | 1.06E-01 | 1.08E+00 |
| ENSG000001 | C1S | 2.43E-01 | 5.95E-01 | 9.11E-01 | -1.35E-01 | -1.10E+00 |
| ENSG000001 | NETO1 | 2.43E-01 | ? | 1.34E+00 | 4.26E-01 | 1.34E+00 |
| ENSG000001 | TMEM87A | 2.43E-01 | 5.95E-01 | 9.71E-01 | -4.20E-02 | -1.03E+00 |

| | | | | | | |
|---|---|---|---|---|---|---|
| ENSG000001 | SMAD1 | 2.43E-01 | 5.95E-01 | 1.05E+00 | 6.43E-02 | 1.05E+00 |
| ENSG00000C | TSPAN17 | 2.43E-01 | 5.95E-01 | 1.05E+00 | 7.03E-02 | 1.05E+00 |
| ENSG000002 | THBS2-AS1 | 2.43E-01 | 5.96E-01 | 8.51E-01 | -2.32E-01 | -1.17E+00 |
| ENSG000002 | GARIN1A | 2.43E-01 | 5.96E-01 | 8.26E-01 | -2.76E-01 | -1.21E+00 |
| ENSG000002 | WASH7P | 2.43E-01 | 5.96E-01 | 9.04E-01 | -1.45E-01 | -1.11E+00 |
| ENSG000001 | TMEM11 | 2.43E-01 | 5.96E-01 | 1.04E+00 | 5.26E-02 | 1.04E+00 |
| ENSG000002 | --- | 2.43E-01 | 5.96E-01 | 9.42E-01 | -8.63E-02 | -1.06E+00 |
| ENSG000002 | --- | 2.43E-01 | ? | 6.44E-01 | -6.34E-01 | -1.55E+00 |
| ENSG000002 | RNF213-AS1 | 2.43E-01 | ? | 1.24E+00 | 3.07E-01 | 1.24E+00 |
| ENSG000002 | ARHGEF34P | 2.43E-01 | 5.96E-01 | 9.62E-01 | -5.64E-02 | -1.04E+00 |
| ENSG000001 | SERP2 | 2.43E-01 | ? | 8.12E-01 | -3.01E-01 | -1.23E+00 |
| ENSG000001 | POLR3H | 2.43E-01 | 5.96E-01 | 1.06E+00 | 8.97E-02 | 1.06E+00 |
| ENSG000001 | SUCLA2 | 2.44E-01 | 5.96E-01 | 9.62E-01 | -5.55E-02 | -1.04E+00 |
| ENSG000001 | ZMYM2 | 2.44E-01 | 5.96E-01 | 9.61E-01 | -5.81E-02 | -1.04E+00 |
| ENSG000001 | EPB41 | 2.44E-01 | 5.96E-01 | 1.05E+00 | 6.41E-02 | 1.05E+00 |
| ENSG000002 | UBE2E4P | 2.44E-01 | ? | 1.49E+00 | 5.77E-01 | 1.49E+00 |
| ENSG000002 | --- | 2.44E-01 | ? | 7.44E-01 | -4.27E-01 | -1.34E+00 |
| ENSG00000C | SH3BP2 | 2.44E-01 | 5.97E-01 | 9.43E-01 | -8.42E-02 | -1.06E+00 |
| ENSG000001 | ARSL | 2.44E-01 | ? | 8.19E-01 | -2.88E-01 | -1.22E+00 |
| ENSG000001 | PAPLN | 2.44E-01 | 5.97E-01 | 1.11E+00 | 1.55E-01 | 1.11E+00 |
| ENSG000001 | RASSF9 | 2.44E-01 | 5.97E-01 | 1.05E+00 | 6.87E-02 | 1.05E+00 |
| ENSG000001 | FGF13 | 2.44E-01 | ? | 1.88E+00 | 9.09E-01 | 1.88E+00 |
| ENSG000001 | ZNF143 | 2.44E-01 | 5.97E-01 | 9.58E-01 | -6.14E-02 | -1.04E+00 |
| ENSG000001 | PPM1H | 2.44E-01 | 5.97E-01 | 1.17E+00 | 2.22E-01 | 1.17E+00 |
| ENSG000001 | CLN3 | 2.44E-01 | 5.97E-01 | 1.07E+00 | 9.40E-02 | 1.07E+00 |
| ENSG000001 | RASGEF1A | 2.44E-01 | 5.97E-01 | 9.44E-01 | -8.29E-02 | -1.06E+00 |
| ENSG000002 | --- | 2.44E-01 | ? | 6.98E-01 | -5.18E-01 | -1.43E+00 |
| ENSG000001 | HOATZ | 2.44E-01 | ? | 1.40E+00 | 4.81E-01 | 1.40E+00 |
| ENSG000001 | TPK1 | 2.44E-01 | 5.97E-01 | 9.13E-01 | -1.31E-01 | -1.09E+00 |
| ENSG000001 | ZNF214 | 2.44E-01 | 5.97E-01 | 8.86E-01 | -1.74E-01 | -1.13E+00 |
| ENSG000002 | --- | 2.44E-01 | ? | 5.81E-01 | -7.84E-01 | -1.72E+00 |
| ENSG000001 | PIGX | 2.44E-01 | 5.97E-01 | 1.06E+00 | 7.89E-02 | 1.06E+00 |
| ENSG000002 | --- | 2.45E-01 | ? | 1.34E+00 | 4.21E-01 | 1.34E+00 |
| ENSG000001 | SYF2 | 2.45E-01 | 5.97E-01 | 9.43E-01 | -8.52E-02 | -1.06E+00 |
| ENSG000001 | NEMP2 | 2.45E-01 | 5.97E-01 | 8.80E-01 | -1.84E-01 | -1.14E+00 |
| ENSG000001 | PSMA3 | 2.45E-01 | 5.97E-01 | 9.66E-01 | -4.93E-02 | -1.03E+00 |
| ENSG000002 | --- | 2.45E-01 | 5.97E-01 | 1.14E+00 | 1.87E-01 | 1.14E+00 |
| ENSG000001 | ABCB9 | 2.45E-01 | 5.98E-01 | 1.11E+00 | 1.55E-01 | 1.11E+00 |
| ENSG000001 | FAM87A | 2.45E-01 | 5.98E-01 | 1.15E+00 | 2.06E-01 | 1.15E+00 |
| ENSG000001 | CCDC121 | 2.45E-01 | 5.98E-01 | 1.16E+00 | 2.11E-01 | 1.16E+00 |
| ENSG000002 | HCG27 | 2.45E-01 | ? | 1.32E+00 | 4.01E-01 | 1.32E+00 |
| ENSG000002 | --- | 2.45E-01 | 5.98E-01 | 8.42E-01 | -2.48E-01 | -1.19E+00 |
| ENSG000002 | SRP54-AS1 | 2.45E-01 | ? | 6.83E-01 | -5.49E-01 | -1.46E+00 |
| ENSG000002 | PCMTD1-DT | 2.45E-01 | 5.98E-01 | 1.09E+00 | 1.21E-01 | 1.09E+00 |
| ENSG000001 | VAT1 | 2.45E-01 | 5.98E-01 | 9.59E-01 | -6.05E-02 | -1.04E+00 |

| | | | | | | |
|---|---|---|---|---|---|---|
| ENSG000002 | CPB2-AS1 | 2.45E-01 | ? | 1.35E+00 | 4.32E-01 | 1.35E+00 |
| ENSG000001 | CAMK2N2 | 2.45E-01 | ? | 7.89E-01 | -3.42E-01 | -1.27E+00 |
| ENSG000002 | PAGR1 | 2.45E-01 | 5.98E-01 | 9.30E-01 | -1.04E-01 | -1.07E+00 |
| ENSG000001 | ZMYM4 | 2.45E-01 | 5.98E-01 | 9.63E-01 | -5.46E-02 | -1.04E+00 |
| ENSG000001 | SYNE3 | 2.45E-01 | 5.98E-01 | 1.13E+00 | 1.80E-01 | 1.13E+00 |
| ENSG000001 | TGS1 | 2.46E-01 | 5.98E-01 | 1.05E+00 | 6.43E-02 | 1.05E+00 |
| ENSG000001 | BAHD1 | 2.46E-01 | 5.98E-01 | 1.05E+00 | 6.47E-02 | 1.05E+00 |
| ENSG000001 | ABCA2 | 2.46E-01 | 5.98E-01 | 1.08E+00 | 1.09E-01 | 1.08E+00 |
| ENSG000002 | MEG8 | 2.46E-01 | 5.98E-01 | 8.83E-01 | -1.80E-01 | -1.13E+00 |
| ENSG000001 | HACL1 | 2.46E-01 | 5.98E-01 | 1.05E+00 | 6.61E-02 | 1.05E+00 |
| ENSG000001 | MYSM1 | 2.46E-01 | 5.98E-01 | 9.45E-01 | -8.09E-02 | -1.06E+00 |
| ENSG00000C | ENO1 | 2.46E-01 | 5.99E-01 | 1.04E+00 | 5.10E-02 | 1.04E+00 |
| ENSG000001 | CXorf58 | 2.46E-01 | 5.99E-01 | 9.04E-01 | -1.45E-01 | -1.11E+00 |
| ENSG000001 | BTN2A2 | 2.46E-01 | 5.99E-01 | 1.09E+00 | 1.23E-01 | 1.09E+00 |
| ENSG000002 | --- | 2.46E-01 | ? | 1.84E+00 | 8.82E-01 | 1.84E+00 |
| ENSG00000C | TOX4 | 2.46E-01 | 5.99E-01 | 9.74E-01 | -3.84E-02 | -1.03E+00 |
| ENSG000001 | TIMM44 | 2.46E-01 | 5.99E-01 | 9.54E-01 | -6.84E-02 | -1.05E+00 |
| ENSG000001 | NBN | 2.46E-01 | 5.99E-01 | 9.59E-01 | -5.99E-02 | -1.04E+00 |
| ENSG00000C | ZC3H3 | 2.46E-01 | 5.99E-01 | 1.06E+00 | 8.43E-02 | 1.06E+00 |
| ENSG000002 | KMT5AP1 | 2.46E-01 | ? | 1.53E+00 | 6.09E-01 | 1.53E+00 |
| ENSG000002 | TAGAP-AS1 | 2.46E-01 | ? | 1.22E+00 | 2.83E-01 | 1.22E+00 |
| ENSG000001 | UTS2B | 2.46E-01 | ? | 7.59E-01 | -3.98E-01 | -1.32E+00 |
| ENSG000001 | RNF112 | 2.46E-01 | ? | 6.64E-01 | -5.91E-01 | -1.51E+00 |
| ENSG000002 | SH3GL1P1 | 2.46E-01 | ? | 7.68E-01 | -3.82E-01 | -1.30E+00 |
| ENSG000002 | FAM131B-AS | 2.47E-01 | 5.99E-01 | 1.07E+00 | 1.02E-01 | 1.07E+00 |
| ENSG000001 | CCL28 | 2.47E-01 | 5.99E-01 | 1.08E+00 | 1.17E-01 | 1.08E+00 |
| ENSG000001 | LRRK1 | 2.47E-01 | 5.99E-01 | 1.03E+00 | 4.85E-02 | 1.03E+00 |
| ENSG000001 | MASTL | 2.47E-01 | 5.99E-01 | 9.25E-01 | -1.12E-01 | -1.08E+00 |
| ENSG000001 | ZNF43 | 2.47E-01 | 5.99E-01 | 9.34E-01 | -9.79E-02 | -1.07E+00 |
| ENSG000002 | TSN | 2.47E-01 | 5.99E-01 | 1.03E+00 | 4.29E-02 | 1.03E+00 |
| ENSG000002 | --- | 2.47E-01 | 5.99E-01 | 8.88E-01 | -1.71E-01 | -1.13E+00 |
| ENSG000001 | CCDC92 | 2.47E-01 | 6.00E-01 | 9.40E-01 | -8.87E-02 | -1.06E+00 |
| ENSG000002 | --- | 2.47E-01 | ? | 5.78E-01 | -7.92E-01 | -1.73E+00 |
| ENSG000001 | MRTFB | 2.47E-01 | 6.00E-01 | 9.56E-01 | -6.48E-02 | -1.05E+00 |
| ENSG000002 | --- | 2.47E-01 | 6.00E-01 | 1.10E+00 | 1.43E-01 | 1.10E+00 |
| ENSG000001 | TMEM218 | 2.47E-01 | 6.00E-01 | 1.07E+00 | 9.20E-02 | 1.07E+00 |
| ENSG000001 | UPF3B | 2.47E-01 | 6.00E-01 | 1.07E+00 | 9.79E-02 | 1.07E+00 |
| ENSG000001 | MEGF8 | 2.47E-01 | 6.00E-01 | 1.06E+00 | 8.83E-02 | 1.06E+00 |
| ENSG000001 | BAG5 | 2.47E-01 | 6.00E-01 | 1.03E+00 | 4.44E-02 | 1.03E+00 |
| ENSG000001 | VTI1A | 2.47E-01 | 6.00E-01 | 9.63E-01 | -5.40E-02 | -1.04E+00 |
| ENSG00000C | ZNF200 | 2.47E-01 | 6.00E-01 | 9.46E-01 | -8.04E-02 | -1.06E+00 |
| ENSG000001 | ATP13A2 | 2.47E-01 | 6.00E-01 | 1.07E+00 | 9.51E-02 | 1.07E+00 |
| ENSG000001 | EEFSEC | 2.47E-01 | 6.00E-01 | 1.06E+00 | 8.68E-02 | 1.06E+00 |
| ENSG000002 | TSKU-AS1 | 2.47E-01 | ? | 6.07E-01 | -7.19E-01 | -1.65E+00 |
| ENSG000001 | ZSCAN25 | 2.48E-01 | 6.00E-01 | 1.06E+00 | 8.15E-02 | 1.06E+00 |

| | | | | | | |
|---|---|---|---|---|---|---|
| ENSG000002 | --- | 2.48E-01 | ? | 1.32E+00 | 3.99E-01 | 1.32E+00 |
| ENSG000002 | LINC02341 | 2.48E-01 | ? | 2.00E+00 | 1.00E+00 | 2.00E+00 |
| ENSG000001 | MICAL1 | 2.48E-01 | 6.00E-01 | 9.47E-01 | -7.83E-02 | -1.06E+00 |
| ENSG000002 | MTFP1 | 2.48E-01 | 6.00E-01 | 1.11E+00 | 1.45E-01 | 1.11E+00 |
| ENSG000001 | ZNF141 | 2.48E-01 | 6.00E-01 | 9.12E-01 | -1.32E-01 | -1.10E+00 |
| ENSG000001 | KCNQ3 | 2.48E-01 | 6.00E-01 | 9.36E-01 | -9.62E-02 | -1.07E+00 |
| ENSG000001 | IGFBP7 | 2.48E-01 | 6.00E-01 | 1.04E+00 | 6.08E-02 | 1.04E+00 |
| ENSG000001 | STMN1 | 2.48E-01 | 6.00E-01 | 9.54E-01 | -6.84E-02 | -1.05E+00 |
| ENSG000001 | SP3 | 2.48E-01 | 6.00E-01 | 1.04E+00 | 5.55E-02 | 1.04E+00 |
| ENSG000002 | --- | 2.48E-01 | 6.00E-01 | 8.78E-01 | -1.88E-01 | -1.14E+00 |
| ENSG000002 | --- | 2.48E-01 | 6.00E-01 | 1.07E+00 | 1.00E-01 | 1.07E+00 |
| ENSG000001 | CYYR1 | 2.48E-01 | 6.01E-01 | 1.10E+00 | 1.37E-01 | 1.10E+00 |
| ENSG000002 | --- | 2.48E-01 | ? | 1.36E+00 | 4.40E-01 | 1.36E+00 |
| ENSG000001 | FAM83D | 2.48E-01 | 6.01E-01 | 8.64E-01 | -2.11E-01 | -1.16E+00 |
| ENSG00000C | GSTO2 | 2.48E-01 | 6.01E-01 | 1.06E+00 | 8.36E-02 | 1.06E+00 |
| ENSG000001 | NCK1 | 2.48E-01 | 6.01E-01 | 9.59E-01 | -6.06E-02 | -1.04E+00 |
| ENSG000001 | SS18L1 | 2.48E-01 | 6.01E-01 | 9.45E-01 | -8.14E-02 | -1.06E+00 |
| ENSG000001 | DUOX2 | 2.49E-01 | 6.01E-01 | 1.15E+00 | 1.95E-01 | 1.15E+00 |
| ENSG000002 | POLR2M | 2.49E-01 | 6.01E-01 | 9.64E-01 | -5.22E-02 | -1.04E+00 |
| ENSG000001 | TMEM87B | 2.49E-01 | 6.01E-01 | 9.60E-01 | -5.94E-02 | -1.04E+00 |
| ENSG00000C | IARS2 | 2.49E-01 | 6.01E-01 | 9.72E-01 | -4.13E-02 | -1.03E+00 |
| ENSG000001 | ENC1 | 2.49E-01 | 6.01E-01 | 1.07E+00 | 9.22E-02 | 1.07E+00 |
| ENSG000001 | ZNF600 | 2.49E-01 | 6.01E-01 | 9.37E-01 | -9.34E-02 | -1.07E+00 |
| ENSG000001 | UBE2D2 | 2.49E-01 | 6.01E-01 | 9.69E-01 | -4.60E-02 | -1.03E+00 |
| ENSG000001 | BIN3 | 2.49E-01 | 6.01E-01 | 9.48E-01 | -7.64E-02 | -1.05E+00 |
| ENSG000002 | --- | 2.49E-01 | ? | 7.54E-01 | -4.07E-01 | -1.33E+00 |
| ENSG000001 | PRPF18 | 2.49E-01 | 6.01E-01 | 9.56E-01 | -6.49E-02 | -1.05E+00 |
| ENSG000002 | --- | 2.49E-01 | 6.01E-01 | 9.03E-01 | -1.47E-01 | -1.11E+00 |
| ENSG00000C | PPP1R15A | 2.49E-01 | 6.01E-01 | 9.26E-01 | -1.12E-01 | -1.08E+00 |
| ENSG000001 | ZNF829 | 2.49E-01 | 6.01E-01 | 9.23E-01 | -1.16E-01 | -1.08E+00 |
| ENSG000001 | RGL4 | 2.49E-01 | ? | 1.87E+00 | 9.02E-01 | 1.87E+00 |
| ENSG000001 | ZNF766 | 2.49E-01 | 6.01E-01 | 9.54E-01 | -6.85E-02 | -1.05E+00 |
| ENSG000002 | HLA-DQB2 | 2.49E-01 | ? | 1.35E+00 | 4.36E-01 | 1.35E+00 |
| ENSG000001 | TSHZ3 | 2.49E-01 | 6.01E-01 | 1.06E+00 | 8.14E-02 | 1.06E+00 |
| ENSG000001 | WRNIP1 | 2.49E-01 | 6.01E-01 | 1.04E+00 | 5.78E-02 | 1.04E+00 |
| ENSG000002 | --- | 2.49E-01 | ? | 7.61E-01 | -3.94E-01 | -1.31E+00 |
| ENSG000001 | SLC2A10 | 2.49E-01 | 6.01E-01 | 1.10E+00 | 1.33E-01 | 1.10E+00 |
| ENSG000002 | --- | 2.49E-01 | ? | 6.07E-01 | -7.21E-01 | -1.65E+00 |
| ENSG000002 | ILF3-DT | 2.49E-01 | 6.01E-01 | 1.11E+00 | 1.53E-01 | 1.11E+00 |
| ENSG00000C | IL4R | 2.50E-01 | 6.02E-01 | 1.05E+00 | 6.84E-02 | 1.05E+00 |
| ENSG000001 | SESTD1 | 2.50E-01 | 6.02E-01 | 9.52E-01 | -7.07E-02 | -1.05E+00 |
| ENSG00000C | SNAI2 | 2.50E-01 | 6.02E-01 | 9.55E-01 | -6.67E-02 | -1.05E+00 |
| ENSG000001 | NHLRC1 | 2.50E-01 | 6.02E-01 | 1.13E+00 | 1.73E-01 | 1.13E+00 |
| ENSG000001 | LPCAT3 | 2.50E-01 | 6.02E-01 | 1.05E+00 | 6.67E-02 | 1.05E+00 |
| ENSG000001 | SEPTIN4 | 2.50E-01 | ? | 7.23E-01 | -4.69E-01 | -1.38E+00 |

| | | | | | | |
|---|---|---|---|---|---|---|
| ENSG00000C | NGEF | 2.50E-01 | ? | 5.82E-01 | -7.82E-01 | -1.72E+00 |
| ENSG000001 | COL18A1 | 2.50E-01 | 6.02E-01 | 1.14E+00 | 1.90E-01 | 1.14E+00 |
| ENSG000001 | ANKS6 | 2.50E-01 | 6.02E-01 | 1.05E+00 | 6.85E-02 | 1.05E+00 |
| ENSG000002 | --- | 2.50E-01 | ? | 1.30E+00 | 3.78E-01 | 1.30E+00 |
| ENSG000002 | PAN3-AS1 | 2.50E-01 | ? | 7.88E-01 | -3.45E-01 | -1.27E+00 |
| ENSG000002 | --- | 2.50E-01 | ? | 1.86E+00 | 8.93E-01 | 1.86E+00 |
| ENSG000001 | PML | 2.50E-01 | 6.02E-01 | 1.04E+00 | 6.01E-02 | 1.04E+00 |
| ENSG000001 | STX8 | 2.50E-01 | 6.02E-01 | 9.56E-01 | -6.46E-02 | -1.05E+00 |
| ENSG000001 | ZNRF1 | 2.50E-01 | 6.02E-01 | 1.05E+00 | 7.57E-02 | 1.05E+00 |
| ENSG000002 | RN7SL535P | 2.50E-01 | ? | 1.97E+00 | 9.77E-01 | 1.97E+00 |
| ENSG000001 | CBX8 | 2.50E-01 | 6.03E-01 | 8.95E-01 | -1.60E-01 | -1.12E+00 |
| ENSG000001 | NR2F1 | 2.50E-01 | ? | 8.30E-01 | -2.69E-01 | -1.20E+00 |
| ENSG000002 | MAGOH-DT | 2.50E-01 | ? | 1.37E+00 | 4.57E-01 | 1.37E+00 |
| ENSG000001 | RMI2 | 2.50E-01 | 6.03E-01 | 1.11E+00 | 1.56E-01 | 1.11E+00 |
| ENSG000001 | ATP5ME | 2.50E-01 | 6.03E-01 | 1.04E+00 | 5.81E-02 | 1.04E+00 |
| ENSG000002 | --- | 2.50E-01 | ? | 7.22E-01 | -4.69E-01 | -1.38E+00 |
| ENSG000001 | ETV2 | 2.51E-01 | ? | 7.30E-01 | -4.54E-01 | -1.37E+00 |
| ENSG000001 | ELAVL2 | 2.51E-01 | 6.03E-01 | 1.22E+00 | 2.81E-01 | 1.22E+00 |
| ENSG000001 | GTF2H2 | 2.51E-01 | 6.03E-01 | 9.35E-01 | -9.77E-02 | -1.07E+00 |
| ENSG000001 | UBXN8 | 2.51E-01 | 6.03E-01 | 9.46E-01 | -8.00E-02 | -1.06E+00 |
| ENSG000002 | --- | 2.51E-01 | ? | 7.63E-01 | -3.90E-01 | -1.31E+00 |
| ENSG000001 | LINC00910 | 2.51E-01 | 6.03E-01 | 8.82E-01 | -1.81E-01 | -1.13E+00 |
| ENSG000001 | MIER3 | 2.51E-01 | 6.03E-01 | 9.54E-01 | -6.82E-02 | -1.05E+00 |
| ENSG000002 | --- | 2.51E-01 | ? | 1.40E+00 | 4.82E-01 | 1.40E+00 |
| ENSG000002 | --- | 2.51E-01 | ? | 1.52E+00 | 6.07E-01 | 1.52E+00 |
| ENSG00000C | PARP12 | 2.51E-01 | 6.04E-01 | 9.34E-01 | -9.81E-02 | -1.07E+00 |
| ENSG000001 | GPALPP1 | 2.51E-01 | 6.04E-01 | 9.48E-01 | -7.68E-02 | -1.05E+00 |
| ENSG000001 | RPL7P9 | 2.51E-01 | 6.04E-01 | 1.07E+00 | 1.03E-01 | 1.07E+00 |
| ENSG000002 | OTULIN-DT | 2.51E-01 | ? | 7.38E-01 | -4.38E-01 | -1.35E+00 |
| ENSG000002 | AK4P3 | 2.51E-01 | ? | 7.91E-01 | -3.38E-01 | -1.26E+00 |
| ENSG000001 | CCNDBP1 | 2.51E-01 | 6.04E-01 | 1.03E+00 | 4.69E-02 | 1.03E+00 |
| ENSG000001 | CCDC30 | 2.51E-01 | 6.04E-01 | 8.40E-01 | -2.51E-01 | -1.19E+00 |
| ENSG000001 | MAD2L1 | 2.51E-01 | 6.04E-01 | 9.09E-01 | -1.38E-01 | -1.10E+00 |
| ENSG000001 | MRPL35 | 2.51E-01 | 6.04E-01 | 9.54E-01 | -6.72E-02 | -1.05E+00 |
| ENSG000001 | FIZ1 | 2.52E-01 | 6.04E-01 | 1.07E+00 | 1.03E-01 | 1.07E+00 |
| ENSG000002 | --- | 2.52E-01 | ? | 1.28E+00 | 3.55E-01 | 1.28E+00 |
| ENSG000002 | --- | 2.52E-01 | ? | 7.55E-01 | -4.06E-01 | -1.33E+00 |
| ENSG000001 | UBAP1 | 2.52E-01 | 6.04E-01 | 1.03E+00 | 4.29E-02 | 1.03E+00 |
| ENSG000001 | SH3KBP1 | 2.52E-01 | 6.04E-01 | 9.47E-01 | -7.88E-02 | -1.06E+00 |
| ENSG000002 | NDUFV2P1 | 2.52E-01 | 6.04E-01 | 1.18E+00 | 2.34E-01 | 1.18E+00 |
| ENSG000002 | --- | 2.52E-01 | ? | 1.52E+00 | 6.06E-01 | 1.52E+00 |
| ENSG000001 | COMMD8 | 2.52E-01 | 6.04E-01 | 9.55E-01 | -6.63E-02 | -1.05E+00 |
| ENSG000001 | SCRN3 | 2.52E-01 | 6.04E-01 | 9.43E-01 | -8.46E-02 | -1.06E+00 |
| ENSG000001 | ZNF554 | 2.52E-01 | 6.04E-01 | 1.10E+00 | 1.39E-01 | 1.10E+00 |
| ENSG000002 | TMEM276 | 2.52E-01 | 6.04E-01 | 1.15E+00 | 2.05E-01 | 1.15E+00 |

| | | | | | | |
|---|---|---|---|---|---|---|
| ENSG000002 | TUBAP2 | 2.52E-01 | ? | 7.34E-01 | -4.46E-01 | -1.36E+00 |
| ENSG000000 | TBC1D10A | 2.52E-01 | 6.05E-01 | 1.07E+00 | 1.01E-01 | 1.07E+00 |
| ENSG000000 | MTA3 | 2.52E-01 | 6.05E-01 | 1.05E+00 | 6.76E-02 | 1.05E+00 |
| ENSG000002 | ZNF710-AS1 | 2.52E-01 | 6.05E-01 | 1.31E+00 | 3.91E-01 | 1.31E+00 |
| ENSG000002 | RBMXP4 | 2.52E-01 | ? | 1.64E+00 | 7.10E-01 | 1.64E+00 |
| ENSG000002 | --- | 2.52E-01 | ? | 7.94E-01 | -3.33E-01 | -1.26E+00 |
| ENSG000000 | CAMSAP3 | 2.52E-01 | 6.05E-01 | 9.20E-01 | -1.21E-01 | -1.09E+00 |
| ENSG000001 | AKNAD1 | 2.53E-01 | 6.05E-01 | 1.17E+00 | 2.25E-01 | 1.17E+00 |
| ENSG000001 | GOLGA6L4 | 2.53E-01 | 6.06E-01 | 9.01E-01 | -1.51E-01 | -1.11E+00 |
| ENSG000000 | ZMPSTE24 | 2.53E-01 | 6.06E-01 | 9.56E-01 | -6.49E-02 | -1.05E+00 |
| ENSG000002 | --- | 2.53E-01 | ? | 7.25E-01 | -4.64E-01 | -1.38E+00 |
| ENSG000002 | LINC00365 | 2.53E-01 | ? | 7.00E-01 | -5.14E-01 | -1.43E+00 |
| ENSG000002 | PTCHD4 | 2.53E-01 | 6.06E-01 | 9.17E-01 | -1.25E-01 | -1.09E+00 |
| ENSG000002 | COX20 | 2.53E-01 | 6.06E-01 | 9.66E-01 | -4.92E-02 | -1.03E+00 |
| ENSG000001 | PCNX1 | 2.53E-01 | 6.06E-01 | 1.05E+00 | 7.41E-02 | 1.05E+00 |
| ENSG000001 | SDF2 | 2.53E-01 | 6.06E-01 | 1.05E+00 | 7.05E-02 | 1.05E+00 |
| ENSG000002 | MTND1P23 | 2.53E-01 | ? | 7.96E-01 | -3.29E-01 | -1.26E+00 |
| ENSG000002 | --- | 2.53E-01 | ? | 1.60E+00 | 6.79E-01 | 1.60E+00 |
| ENSG000001 | QRICH1 | 2.53E-01 | 6.06E-01 | 9.69E-01 | -4.60E-02 | -1.03E+00 |
| ENSG000002 | --- | 2.53E-01 | ? | 1.29E+00 | 3.62E-01 | 1.29E+00 |
| ENSG000001 | MRPL19 | 2.53E-01 | 6.07E-01 | 9.63E-01 | -5.42E-02 | -1.04E+00 |
| ENSG000001 | USP50 | 2.54E-01 | ? | 7.17E-01 | -4.81E-01 | -1.40E+00 |
| ENSG000002 | --- | 2.54E-01 | ? | 8.12E-01 | -3.00E-01 | -1.23E+00 |
| ENSG000002 | --- | 2.54E-01 | 6.07E-01 | 1.17E+00 | 2.32E-01 | 1.17E+00 |
| ENSG000002 | HMSD | 2.54E-01 | ? | 1.98E+00 | 9.87E-01 | 1.98E+00 |
| ENSG000000 | PXN | 2.54E-01 | 6.07E-01 | 1.04E+00 | 5.15E-02 | 1.04E+00 |
| ENSG000001 | UBC | 2.54E-01 | 6.07E-01 | 1.03E+00 | 4.71E-02 | 1.03E+00 |
| ENSG000002 | VAMP2 | 2.54E-01 | 6.07E-01 | 9.57E-01 | -6.38E-02 | -1.05E+00 |
| ENSG000001 | ESYT1 | 2.54E-01 | 6.07E-01 | 9.64E-01 | -5.36E-02 | -1.04E+00 |
| ENSG000001 | YWHAB | 2.54E-01 | 6.07E-01 | 9.80E-01 | -2.89E-02 | -1.02E+00 |
| ENSG000001 | UBE2E3 | 2.54E-01 | 6.07E-01 | 1.05E+00 | 6.40E-02 | 1.05E+00 |
| ENSG000001 | ITPKB | 2.54E-01 | 6.07E-01 | 9.43E-01 | -8.46E-02 | -1.06E+00 |
| ENSG000001 | SLC10A7 | 2.54E-01 | 6.07E-01 | 9.37E-01 | -9.38E-02 | -1.07E+00 |
| ENSG000002 | ARPC1A | 2.54E-01 | 6.07E-01 | 1.03E+00 | 4.66E-02 | 1.03E+00 |
| ENSG000002 | SEPTIN4-AS1 | 2.54E-01 | ? | 1.53E+00 | 6.17E-01 | 1.53E+00 |
| ENSG000002 | COG8 | 2.54E-01 | 6.07E-01 | 9.59E-01 | -6.10E-02 | -1.04E+00 |
| ENSG000001 | TNKS2 | 2.54E-01 | 6.07E-01 | 9.77E-01 | -3.41E-02 | -1.02E+00 |
| ENSG000002 | --- | 2.54E-01 | ? | 1.59E+00 | 6.68E-01 | 1.59E+00 |
| ENSG000001 | EYA4 | 2.54E-01 | 6.07E-01 | 9.52E-01 | -7.15E-02 | -1.05E+00 |
| ENSG000001 | PDE4C | 2.54E-01 | ? | 1.62E+00 | 6.92E-01 | 1.62E+00 |
| ENSG000001 | NTM | 2.54E-01 | 6.07E-01 | 9.52E-01 | -7.09E-02 | -1.05E+00 |
| ENSG000002 | --- | 2.55E-01 | ? | 6.74E-01 | -5.70E-01 | -1.48E+00 |
| ENSG000001 | LINC01116 | 2.55E-01 | ? | 8.11E-01 | -3.03E-01 | -1.23E+00 |
| ENSG000001 | EIF2AK3 | 2.55E-01 | 6.08E-01 | 9.64E-01 | -5.34E-02 | -1.04E+00 |
| ENSG000001 | DACT3 | 2.55E-01 | ? | 1.51E+00 | 5.93E-01 | 1.51E+00 |

| | | | | | | |
|---|---|---|---|---|---|---|
| ENSG000001 | C6orf89 | 2.55E-01 | 6.08E-01 | 1.03E+00 | 3.94E-02 | 1.03E+00 |
| ENSG000001 | PEX16 | 2.55E-01 | 6.08E-01 | 1.09E+00 | 1.20E-01 | 1.09E+00 |
| ENSG000001 | EEF2 | 2.55E-01 | 6.09E-01 | 1.05E+00 | 6.57E-02 | 1.05E+00 |
| ENSG000001 | POC1B | 2.55E-01 | 6.09E-01 | 9.44E-01 | -8.36E-02 | -1.06E+00 |
| ENSG000002 | --- | 2.55E-01 | ? | 1.82E+00 | 8.61E-01 | 1.82E+00 |
| ENSG000001 | GPRASP1 | 2.55E-01 | 6.09E-01 | 8.70E-01 | -2.01E-01 | -1.15E+00 |
| ENSG00000C | CDC5L | 2.55E-01 | 6.09E-01 | 9.52E-01 | -7.04E-02 | -1.05E+00 |
| ENSG000001 | UFC1 | 2.55E-01 | 6.09E-01 | 1.03E+00 | 4.16E-02 | 1.03E+00 |
| ENSG000001 | ZNF92 | 2.56E-01 | 6.09E-01 | 9.22E-01 | -1.17E-01 | -1.08E+00 |
| ENSG000001 | ALKBH3 | 2.56E-01 | 6.09E-01 | 9.46E-01 | -8.05E-02 | -1.06E+00 |
| ENSG000002 | --- | 2.56E-01 | 6.09E-01 | 1.07E+00 | 1.03E-01 | 1.07E+00 |
| ENSG000002 | SEMA3F-AS1 | 2.56E-01 | 6.09E-01 | 8.11E-01 | -3.02E-01 | -1.23E+00 |
| ENSG000001 | ARL17A | 2.56E-01 | 6.09E-01 | 9.49E-01 | -7.58E-02 | -1.05E+00 |
| ENSG000001 | MPP2 | 2.56E-01 | 6.09E-01 | 1.14E+00 | 1.94E-01 | 1.14E+00 |
| ENSG00000C | SRRT | 2.56E-01 | 6.09E-01 | 9.61E-01 | -5.68E-02 | -1.04E+00 |
| ENSG000001 | DRAM2 | 2.56E-01 | 6.09E-01 | 1.05E+00 | 6.35E-02 | 1.05E+00 |
| ENSG000001 | RELN | 2.56E-01 | ? | 1.29E+00 | 3.68E-01 | 1.29E+00 |
| ENSG000002 | --- | 2.56E-01 | ? | 1.30E+00 | 3.81E-01 | 1.30E+00 |
| ENSG000001 | LRRFIP1 | 2.56E-01 | 6.09E-01 | 9.58E-01 | -6.23E-02 | -1.04E+00 |
| ENSG000002 | LINC02832 | 2.56E-01 | ? | 6.14E-01 | -7.03E-01 | -1.63E+00 |
| ENSG000001 | PTAR1 | 2.56E-01 | 6.09E-01 | 9.54E-01 | -6.84E-02 | -1.05E+00 |
| ENSG000002 | --- | 2.56E-01 | 6.09E-01 | 8.79E-01 | -1.85E-01 | -1.14E+00 |
| ENSG000002 | RPL7P23 | 2.56E-01 | ? | 1.40E+00 | 4.87E-01 | 1.40E+00 |
| ENSG000001 | KRT5 | 2.56E-01 | 6.10E-01 | 1.04E+00 | 5.14E-02 | 1.04E+00 |
| ENSG000001 | RPP40 | 2.56E-01 | 6.10E-01 | 9.36E-01 | -9.51E-02 | -1.07E+00 |
| ENSG000002 | NPM1P26 | 2.56E-01 | ? | 6.02E-01 | -7.31E-01 | -1.66E+00 |
| ENSG000001 | MME | 2.57E-01 | 6.10E-01 | 1.11E+00 | 1.47E-01 | 1.11E+00 |
| ENSG000001 | SLC25A46 | 2.57E-01 | 6.10E-01 | 9.65E-01 | -5.15E-02 | -1.04E+00 |
| ENSG000001 | TNFSF4 | 2.57E-01 | 6.10E-01 | 8.18E-01 | -2.90E-01 | -1.22E+00 |
| ENSG000002 | CORO7 | 2.57E-01 | 6.10E-01 | 1.12E+00 | 1.63E-01 | 1.12E+00 |
| ENSG000001 | CCDC62 | 2.57E-01 | ? | 8.28E-01 | -2.73E-01 | -1.21E+00 |
| ENSG000002 | LINC01135 | 2.57E-01 | ? | 7.08E-01 | -4.98E-01 | -1.41E+00 |
| ENSG000001 | ERI1 | 2.57E-01 | 6.10E-01 | 9.49E-01 | -7.48E-02 | -1.05E+00 |
| ENSG000001 | SUPT3H | 2.57E-01 | 6.10E-01 | 9.33E-01 | -1.01E-01 | -1.07E+00 |
| ENSG000002 | RASSF5 | 2.57E-01 | 6.10E-01 | 1.09E+00 | 1.24E-01 | 1.09E+00 |
| ENSG000002 | ZNF585B | 2.57E-01 | 6.10E-01 | 9.40E-01 | -8.90E-02 | -1.06E+00 |
| ENSG000001 | KHDC4 | 2.57E-01 | 6.10E-01 | 9.34E-01 | -9.92E-02 | -1.07E+00 |
| ENSG000001 | LFNG | 2.57E-01 | 6.11E-01 | 1.12E+00 | 1.61E-01 | 1.12E+00 |
| ENSG00000C | NECAP1 | 2.57E-01 | 6.11E-01 | 9.59E-01 | -6.07E-02 | -1.04E+00 |
| ENSG00000C | TRIO | 2.57E-01 | 6.11E-01 | 1.04E+00 | 6.06E-02 | 1.04E+00 |
| ENSG000001 | SERGEF | 2.57E-01 | 6.11E-01 | 1.07E+00 | 1.01E-01 | 1.07E+00 |
| ENSG000001 | ZNF177 | 2.57E-01 | 6.11E-01 | 1.15E+00 | 1.98E-01 | 1.15E+00 |
| ENSG000001 | LEF1 | 2.57E-01 | ? | 1.24E+00 | 3.16E-01 | 1.24E+00 |
| ENSG000001 | ZNF282 | 2.57E-01 | 6.11E-01 | 1.06E+00 | 8.98E-02 | 1.06E+00 |
| ENSG00000C | ANKRD24 | 2.58E-01 | ? | 1.30E+00 | 3.78E-01 | 1.30E+00 |

| | | | | | | |
|---|---|---|---|---|---|---|
| ENSG000001 | SLC37A2 | 2.58E-01 | 6.11E-01 | 9.43E-01 | -8.44E-02 | -1.06E+00 |
| ENSG00000C | SPG21 | 2.58E-01 | 6.11E-01 | 1.04E+00 | 5.67E-02 | 1.04E+00 |
| ENSG000002 | --- | 2.58E-01 | ? | 1.35E+00 | 4.28E-01 | 1.35E+00 |
| ENSG000001 | ANKRD6 | 2.58E-01 | 6.11E-01 | 9.58E-01 | -6.24E-02 | -1.04E+00 |
| ENSG000001 | NUDT4 | 2.58E-01 | 6.11E-01 | 1.04E+00 | 5.48E-02 | 1.04E+00 |
| ENSG000002 | --- | 2.58E-01 | ? | 1.27E+00 | 3.48E-01 | 1.27E+00 |
| ENSG000001 | HSCB | 2.58E-01 | 6.11E-01 | 1.10E+00 | 1.43E-01 | 1.10E+00 |
| ENSG00000C | AP1M1 | 2.58E-01 | 6.11E-01 | 1.06E+00 | 7.97E-02 | 1.06E+00 |
| ENSG00000C | OSBPL6 | 2.58E-01 | 6.11E-01 | 1.06E+00 | 7.75E-02 | 1.06E+00 |
| ENSG000001 | HIC2 | 2.58E-01 | 6.11E-01 | 1.07E+00 | 9.47E-02 | 1.07E+00 |
| ENSG000001 | TYK2 | 2.58E-01 | 6.11E-01 | 1.06E+00 | 9.02E-02 | 1.06E+00 |
| ENSG000002 | RPL5P23 | 2.58E-01 | ? | 6.72E-01 | -5.74E-01 | -1.49E+00 |
| ENSG00000C | MAGI3 | 2.58E-01 | 6.12E-01 | 9.49E-01 | -7.61E-02 | -1.05E+00 |
| ENSG000002 | --- | 2.58E-01 | 6.12E-01 | 8.69E-01 | -2.03E-01 | -1.15E+00 |
| ENSG000001 | NDUFA12 | 2.58E-01 | 6.12E-01 | 1.05E+00 | 7.35E-02 | 1.05E+00 |
| ENSG000001 | S1PR5 | 2.58E-01 | 6.12E-01 | 1.09E+00 | 1.22E-01 | 1.09E+00 |
| ENSG000001 | BRI3BP | 2.58E-01 | 6.12E-01 | 1.06E+00 | 8.97E-02 | 1.06E+00 |
| ENSG000001 | FUT11 | 2.58E-01 | 6.12E-01 | 1.04E+00 | 5.91E-02 | 1.04E+00 |
| ENSG000001 | CLTB | 2.59E-01 | 6.12E-01 | 1.04E+00 | 5.81E-02 | 1.04E+00 |
| ENSG000002 | HLA-H | 2.59E-01 | 6.12E-01 | 8.42E-01 | -2.48E-01 | -1.19E+00 |
| ENSG000001 | TRPC1 | 2.59E-01 | 6.12E-01 | 1.07E+00 | 9.28E-02 | 1.07E+00 |
| ENSG000002 | --- | 2.59E-01 | ? | 6.92E-01 | -5.31E-01 | -1.45E+00 |
| ENSG000002 | DDX19A-DT | 2.59E-01 | ? | 7.84E-01 | -3.50E-01 | -1.27E+00 |
| ENSG000001 | MED19 | 2.59E-01 | 6.12E-01 | 1.07E+00 | 1.01E-01 | 1.07E+00 |
| ENSG000001 | EIF4H | 2.59E-01 | 6.12E-01 | 1.04E+00 | 5.56E-02 | 1.04E+00 |
| ENSG000001 | LETM2 | 2.59E-01 | 6.12E-01 | 1.07E+00 | 1.02E-01 | 1.07E+00 |
| ENSG000001 | PLEKHG4 | 2.59E-01 | 6.12E-01 | 1.15E+00 | 2.01E-01 | 1.15E+00 |
| ENSG000001 | HSDL1 | 2.59E-01 | 6.12E-01 | 9.55E-01 | -6.65E-02 | -1.05E+00 |
| ENSG000001 | RNF180 | 2.59E-01 | 6.12E-01 | 9.09E-01 | -1.38E-01 | -1.10E+00 |
| ENSG000001 | SUPT5H | 2.59E-01 | 6.12E-01 | 1.03E+00 | 3.86E-02 | 1.03E+00 |
| ENSG000002 | UBAC2-AS1 | 2.59E-01 | ? | 1.25E+00 | 3.26E-01 | 1.25E+00 |
| ENSG000002 | --- | 2.59E-01 | ? | 6.57E-01 | -6.07E-01 | -1.52E+00 |
| ENSG000001 | SAT2 | 2.59E-01 | 6.13E-01 | 1.07E+00 | 9.99E-02 | 1.07E+00 |
| ENSG000001 | LMOD1 | 2.59E-01 | ? | 7.26E-01 | -4.61E-01 | -1.38E+00 |
| ENSG000001 | MRPS23 | 2.59E-01 | 6.13E-01 | 1.04E+00 | 6.14E-02 | 1.04E+00 |
| ENSG000002 | ZSCAN31 | 2.59E-01 | 6.13E-01 | 9.38E-01 | -9.21E-02 | -1.07E+00 |
| ENSG000002 | RPL36A-HNR | 2.59E-01 | 6.13E-01 | 1.61E+00 | 6.84E-01 | 1.61E+00 |
| ENSG000002 | THBS1-AS1 | 2.59E-01 | 6.13E-01 | 8.19E-01 | -2.89E-01 | -1.22E+00 |
| ENSG000002 | --- | 2.59E-01 | ? | 6.96E-01 | -5.23E-01 | -1.44E+00 |
| ENSG000002 | --- | 2.59E-01 | 6.13E-01 | 8.32E-01 | -2.66E-01 | -1.20E+00 |
| ENSG000001 | GCHFR | 2.60E-01 | ? | 7.31E-01 | -4.53E-01 | -1.37E+00 |
| ENSG000002 | CCDC200 | 2.60E-01 | ? | 7.51E-01 | -4.13E-01 | -1.33E+00 |
| ENSG000001 | TPRN | 2.60E-01 | 6.13E-01 | 8.95E-01 | -1.59E-01 | -1.12E+00 |
| ENSG000002 | ARHGAP23 | 2.60E-01 | 6.13E-01 | 1.05E+00 | 6.61E-02 | 1.05E+00 |
| ENSG00000C | CCL26 | 2.60E-01 | ? | 1.46E+00 | 5.44E-01 | 1.46E+00 |

| | | | | | | |
|---|---|---|---|---|---|---|
| ENSG000001 | DYNC1I1 | 2.60E-01 | 6.13E-01 | 8.74E-01 | -1.95E-01 | -1.14E+00 |
| ENSG000002 | ZNF487 | 2.60E-01 | 6.13E-01 | 1.13E+00 | 1.71E-01 | 1.13E+00 |
| ENSG000002 | MT-TY | 2.60E-01 | ? | 1.40E+00 | 4.90E-01 | 1.40E+00 |
| ENSG000001 | NIM1K | 2.60E-01 | ? | 1.32E+00 | 4.01E-01 | 1.32E+00 |
| ENSG000001 | ABI3BP | 2.60E-01 | 6.14E-01 | 8.85E-01 | -1.76E-01 | -1.13E+00 |
| ENSG000002 | --- | 2.60E-01 | ? | 5.67E-01 | -8.19E-01 | -1.76E+00 |
| ENSG000002 | ADIRF-AS1 | 2.60E-01 | 6.14E-01 | 8.69E-01 | -2.03E-01 | -1.15E+00 |
| ENSG000001 | N4BP1 | 2.60E-01 | 6.14E-01 | 1.04E+00 | 5.46E-02 | 1.04E+00 |
| ENSG000001 | BMP2 | 2.61E-01 | 6.15E-01 | 9.41E-01 | -8.79E-02 | -1.06E+00 |
| ENSG000001 | ELP2 | 2.61E-01 | 6.15E-01 | 1.03E+00 | 4.57E-02 | 1.03E+00 |
| ENSG000002 | DENND11 | 2.61E-01 | 6.15E-01 | 9.61E-01 | -5.72E-02 | -1.04E+00 |
| ENSG000002 | SMG1P1 | 2.61E-01 | 6.15E-01 | 7.85E-01 | -3.48E-01 | -1.27E+00 |
| ENSG000001 | ABCA10 | 2.61E-01 | 6.15E-01 | 8.79E-01 | -1.86E-01 | -1.14E+00 |
| ENSG000001 | ZNF225-AS1 | 2.61E-01 | 6.15E-01 | 9.09E-01 | -1.37E-01 | -1.10E+00 |
| ENSG000002 | JTB-DT | 2.61E-01 | ? | 8.33E-01 | -2.63E-01 | -1.20E+00 |
| ENSG000002 | STXBP5-AS1 | 2.61E-01 | 6.15E-01 | 9.37E-01 | -9.33E-02 | -1.07E+00 |
| ENSG000002 | --- | 2.61E-01 | ? | 7.27E-01 | -4.60E-01 | -1.38E+00 |
| ENSG000001 | ADCY10P1 | 2.61E-01 | ? | 1.51E+00 | 5.98E-01 | 1.51E+00 |
| ENSG00000C | MCM6 | 2.61E-01 | 6.15E-01 | 1.07E+00 | 1.00E-01 | 1.07E+00 |
| ENSG000001 | FBXL8 | 2.61E-01 | 6.15E-01 | 1.15E+00 | 2.07E-01 | 1.15E+00 |
| ENSG000001 | ATP6V1F | 2.61E-01 | 6.15E-01 | 1.05E+00 | 6.63E-02 | 1.05E+00 |
| ENSG000001 | PPIG | 2.61E-01 | 6.15E-01 | 9.44E-01 | -8.25E-02 | -1.06E+00 |
| ENSG000002 | LINC02245 | 2.61E-01 | ? | 1.54E+00 | 6.26E-01 | 1.54E+00 |
| ENSG000002 | LNCRNA-IUR | 2.61E-01 | 6.15E-01 | 1.19E+00 | 2.47E-01 | 1.19E+00 |
| ENSG00000C | EIF2AK2 | 2.61E-01 | 6.15E-01 | 9.50E-01 | -7.45E-02 | -1.05E+00 |
| ENSG00000C | ZNF671 | 2.62E-01 | 6.15E-01 | 9.18E-01 | -1.23E-01 | -1.09E+00 |
| ENSG000002 | --- | 2.62E-01 | 6.15E-01 | 1.16E+00 | 2.19E-01 | 1.16E+00 |
| ENSG000001 | TMEM147 | 2.62E-01 | 6.15E-01 | 1.08E+00 | 1.15E-01 | 1.08E+00 |
| ENSG000002 | --- | 2.62E-01 | ? | 8.07E-01 | -3.09E-01 | -1.24E+00 |
| ENSG000001 | MEP1A | 2.62E-01 | ? | 8.46E-01 | -2.41E-01 | -1.18E+00 |
| ENSG000002 | --- | 2.62E-01 | ? | 7.21E-01 | -4.73E-01 | -1.39E+00 |
| ENSG00000C | PLEKHB1 | 2.62E-01 | ? | 1.23E+00 | 2.98E-01 | 1.23E+00 |
| ENSG000001 | CRBN | 2.62E-01 | 6.16E-01 | 9.51E-01 | -7.30E-02 | -1.05E+00 |
| ENSG00000C | SRCAP | 2.62E-01 | 6.16E-01 | 9.41E-01 | -8.73E-02 | -1.06E+00 |
| ENSG000001 | PNCK | 2.62E-01 | 6.16E-01 | 1.21E+00 | 2.77E-01 | 1.21E+00 |
| ENSG000001 | H2BC11 | 2.62E-01 | 6.17E-01 | 1.12E+00 | 1.58E-01 | 1.12E+00 |
| ENSG000002 | --- | 2.62E-01 | ? | 4.68E-01 | -1.09E+00 | -2.14E+00 |
| ENSG000002 | --- | 2.62E-01 | ? | 7.17E-01 | -4.79E-01 | -1.39E+00 |
| ENSG000002 | --- | 2.62E-01 | ? | 1.67E+00 | 7.41E-01 | 1.67E+00 |
| ENSG000002 | SPESP1 | 2.62E-01 | ? | 7.74E-01 | -3.70E-01 | -1.29E+00 |
| ENSG000001 | SMAD7 | 2.63E-01 | 6.17E-01 | 9.48E-01 | -7.68E-02 | -1.05E+00 |
| ENSG00000C | PDRG1 | 2.63E-01 | 6.17E-01 | 1.07E+00 | 1.01E-01 | 1.07E+00 |
| ENSG000001 | RAPGEF2 | 2.63E-01 | 6.17E-01 | 1.04E+00 | 5.89E-02 | 1.04E+00 |
| ENSG000001 | KATNAL2 | 2.63E-01 | 6.17E-01 | 8.50E-01 | -2.34E-01 | -1.18E+00 |
| ENSG000002 | SP2-AS1 | 2.63E-01 | ? | 1.22E+00 | 2.90E-01 | 1.22E+00 |

| | | | | | | |
|---|---|---|---|---|---|---|
| ENSG00000C | TAB2 | 2.63E-01 | 6.17E-01 | 9.53E-01 | -6.94E-02 | -1.05E+00 |
| ENSG000002 | --- | 2.63E-01 | 6.17E-01 | 8.46E-01 | -2.41E-01 | -1.18E+00 |
| ENSG000001 | CORO2A | 2.63E-01 | 6.17E-01 | 1.06E+00 | 8.07E-02 | 1.06E+00 |
| ENSG000001 | RCOR2 | 2.63E-01 | ? | 1.24E+00 | 3.14E-01 | 1.24E+00 |
| ENSG000001 | HSPBP1 | 2.63E-01 | 6.17E-01 | 1.09E+00 | 1.28E-01 | 1.09E+00 |
| ENSG000001 | SORBS3 | 2.63E-01 | 6.17E-01 | 1.05E+00 | 7.26E-02 | 1.05E+00 |
| ENSG000002 | SMIM27 | 2.63E-01 | 6.17E-01 | 1.15E+00 | 2.02E-01 | 1.15E+00 |
| ENSG000002 | NORAD | 2.63E-01 | 6.17E-01 | 9.68E-01 | -4.63E-02 | -1.03E+00 |
| ENSG000001 | SUOX | 2.63E-01 | 6.17E-01 | 9.36E-01 | -9.55E-02 | -1.07E+00 |
| ENSG000001 | GREB1L | 2.63E-01 | 6.17E-01 | 1.12E+00 | 1.58E-01 | 1.12E+00 |
| ENSG000001 | SLC2A5 | 2.63E-01 | 6.17E-01 | 1.17E+00 | 2.25E-01 | 1.17E+00 |
| ENSG000002 | --- | 2.63E-01 | ? | 6.17E-01 | -6.96E-01 | -1.62E+00 |
| ENSG00000C | PHKA2 | 2.63E-01 | 6.17E-01 | 1.06E+00 | 8.91E-02 | 1.06E+00 |
| ENSG000002 | H3C15 | 2.63E-01 | ? | 1.91E+00 | 9.32E-01 | 1.91E+00 |
| ENSG000001 | CTF1 | 2.63E-01 | 6.17E-01 | 1.13E+00 | 1.73E-01 | 1.13E+00 |
| ENSG000001 | STARD4 | 2.63E-01 | 6.17E-01 | 9.53E-01 | -6.97E-02 | -1.05E+00 |
| ENSG000001 | CDC37L1 | 2.63E-01 | 6.17E-01 | 9.61E-01 | -5.78E-02 | -1.04E+00 |
| ENSG000002 | TMEM167B- | 2.63E-01 | 6.17E-01 | 1.12E+00 | 1.58E-01 | 1.12E+00 |
| ENSG000001 | ARIH2 | 2.63E-01 | 6.17E-01 | 9.67E-01 | -4.89E-02 | -1.03E+00 |
| ENSG000001 | SMARCC2 | 2.63E-01 | 6.17E-01 | 1.04E+00 | 5.25E-02 | 1.04E+00 |
| ENSG000002 | --- | 2.63E-01 | 6.17E-01 | 8.96E-01 | -1.58E-01 | -1.12E+00 |
| ENSG000002 | SPINT1-AS1 | 2.63E-01 | 6.17E-01 | 1.13E+00 | 1.74E-01 | 1.13E+00 |
| ENSG000002 | LINC01270 | 2.64E-01 | 6.17E-01 | 8.81E-01 | -1.82E-01 | -1.13E+00 |
| ENSG000002 | --- | 2.64E-01 | ? | 7.26E-01 | -4.61E-01 | -1.38E+00 |
| ENSG000001 | NARS1 | 2.64E-01 | 6.17E-01 | 9.74E-01 | -3.81E-02 | -1.03E+00 |
| ENSG000002 | --- | 2.64E-01 | 6.17E-01 | 1.14E+00 | 1.93E-01 | 1.14E+00 |
| ENSG000001 | VARS2 | 2.64E-01 | 6.17E-01 | 1.08E+00 | 1.10E-01 | 1.08E+00 |
| ENSG000001 | TPH1 | 2.64E-01 | ? | 7.87E-01 | -3.46E-01 | -1.27E+00 |
| ENSG000002 | SLC16A4-AS1 | 2.64E-01 | ? | 8.10E-01 | -3.05E-01 | -1.24E+00 |
| ENSG000001 | CYP4V2 | 2.64E-01 | 6.18E-01 | 9.29E-01 | -1.06E-01 | -1.08E+00 |
| ENSG00000C | SOAT1 | 2.64E-01 | 6.18E-01 | 1.06E+00 | 7.86E-02 | 1.06E+00 |
| ENSG000001 | TMEM116 | 2.64E-01 | 6.18E-01 | 1.11E+00 | 1.55E-01 | 1.11E+00 |
| ENSG000001 | CCDC88B | 2.64E-01 | 6.18E-01 | 1.11E+00 | 1.57E-01 | 1.11E+00 |
| ENSG000002 | ASPDH | 2.64E-01 | ? | 6.09E-01 | -7.15E-01 | -1.64E+00 |
| ENSG000001 | DUS4L | 2.64E-01 | 6.18E-01 | 1.08E+00 | 1.10E-01 | 1.08E+00 |
| ENSG000001 | TMEM175 | 2.65E-01 | 6.18E-01 | 1.07E+00 | 9.60E-02 | 1.07E+00 |
| ENSG000001 | FBXO10 | 2.65E-01 | 6.18E-01 | 1.06E+00 | 9.02E-02 | 1.06E+00 |
| ENSG00000C | RTEL1-TNFRS | 2.65E-01 | 6.18E-01 | 8.65E-01 | -2.10E-01 | -1.16E+00 |
| ENSG000001 | ZNF75A | 2.65E-01 | 6.18E-01 | 9.29E-01 | -1.06E-01 | -1.08E+00 |
| ENSG000002 | SUMO2P21 | 2.65E-01 | ? | 6.35E-01 | -6.55E-01 | -1.57E+00 |
| ENSG000001 | FBL | 2.65E-01 | 6.19E-01 | 1.06E+00 | 8.45E-02 | 1.06E+00 |
| ENSG000001 | GABPB1 | 2.65E-01 | 6.19E-01 | 9.51E-01 | -7.30E-02 | -1.05E+00 |
| ENSG000002 | --- | 2.65E-01 | 6.19E-01 | 1.14E+00 | 1.92E-01 | 1.14E+00 |
| ENSG000001 | LURAP1L | 2.65E-01 | 6.19E-01 | 1.08E+00 | 1.14E-01 | 1.08E+00 |
| ENSG000002 | CYP2U1-AS1 | 2.65E-01 | ? | 8.30E-01 | -2.68E-01 | -1.20E+00 |

| | | | | | | |
|---|---|---|---|---|---|---|
| ENSG000001 | RRP36 | 2.65E-01 | 6.19E-01 | 1.03E+00 | 4.42E-02 | 1.03E+00 |
| ENSG000001 | TIMM17B | 2.65E-01 | 6.19E-01 | 1.07E+00 | 9.39E-02 | 1.07E+00 |
| ENSG000001 | KIAA1755 | 2.65E-01 | ? | 6.89E-01 | -5.38E-01 | -1.45E+00 |
| ENSG000002 | CRYBB2P1 | 2.66E-01 | 6.20E-01 | 9.34E-01 | -9.93E-02 | -1.07E+00 |
| ENSG000002 | --- | 2.66E-01 | 6.20E-01 | 8.78E-01 | -1.87E-01 | -1.14E+00 |
| ENSG000001 | LINC00336 | 2.66E-01 | ? | 6.50E-01 | -6.22E-01 | -1.54E+00 |
| ENSG000001 | CD320 | 2.66E-01 | 6.20E-01 | 1.10E+00 | 1.41E-01 | 1.10E+00 |
| ENSG000001 | XG | 2.66E-01 | 6.20E-01 | 1.07E+00 | 9.10E-02 | 1.07E+00 |
| ENSG000001 | DNAJB5 | 2.66E-01 | 6.20E-01 | 1.05E+00 | 7.21E-02 | 1.05E+00 |
| ENSG000001 | COL24A1 | 2.66E-01 | ? | 1.27E+00 | 3.43E-01 | 1.27E+00 |
| ENSG000002 | TAS2R14 | 2.66E-01 | ? | 7.65E-01 | -3.87E-01 | -1.31E+00 |
| ENSG000001 | AMOTL1 | 2.66E-01 | 6.20E-01 | 9.64E-01 | -5.22E-02 | -1.04E+00 |
| ENSG000001 | AKIRIN2 | 2.66E-01 | 6.20E-01 | 9.52E-01 | -7.03E-02 | -1.05E+00 |
| ENSG00000C | MLF2 | 2.66E-01 | 6.20E-01 | 1.04E+00 | 5.82E-02 | 1.04E+00 |
| ENSG000002 | LINC02449 | 2.66E-01 | ? | 1.24E+00 | 3.05E-01 | 1.24E+00 |
| ENSG000001 | SAMD4B | 2.66E-01 | 6.20E-01 | 9.73E-01 | -3.88E-02 | -1.03E+00 |
| ENSG000001 | CLDN19 | 2.66E-01 | ? | 6.59E-01 | -6.02E-01 | -1.52E+00 |
| ENSG000002 | --- | 2.66E-01 | ? | 5.15E-01 | -9.57E-01 | -1.94E+00 |
| ENSG000002 | ZNF37BP | 2.66E-01 | 6.20E-01 | 9.30E-01 | -1.05E-01 | -1.08E+00 |
| ENSG000001 | EIF3F | 2.66E-01 | 6.20E-01 | 1.04E+00 | 5.11E-02 | 1.04E+00 |
| ENSG000002 | PDCD6-AHRR | 2.66E-01 | ? | 1.54E+00 | 6.26E-01 | 1.54E+00 |
| ENSG000001 | FLRT1 | 2.66E-01 | ? | 6.91E-01 | -5.34E-01 | -1.45E+00 |
| ENSG000001 | STC1 | 2.66E-01 | 6.20E-01 | 7.35E-01 | -4.44E-01 | -1.36E+00 |
| ENSG000001 | COPE | 2.66E-01 | 6.20E-01 | 1.07E+00 | 1.01E-01 | 1.07E+00 |
| ENSG000001 | ITGA11 | 2.66E-01 | ? | 1.48E+00 | 5.70E-01 | 1.48E+00 |
| ENSG000001 | TST | 2.66E-01 | 6.20E-01 | 1.11E+00 | 1.45E-01 | 1.11E+00 |
| ENSG000001 | DLX1 | 2.66E-01 | ? | 7.81E-01 | -3.57E-01 | -1.28E+00 |
| ENSG000001 | CLPX | 2.67E-01 | 6.20E-01 | 9.73E-01 | -3.96E-02 | -1.03E+00 |
| ENSG000002 | --- | 2.67E-01 | 6.20E-01 | 8.59E-01 | -2.19E-01 | -1.16E+00 |
| ENSG000001 | H2BC26 | 2.67E-01 | 6.20E-01 | 1.16E+00 | 2.20E-01 | 1.16E+00 |
| ENSG000002 | --- | 2.67E-01 | ? | 1.73E+00 | 7.90E-01 | 1.73E+00 |
| ENSG000002 | --- | 2.67E-01 | 6.20E-01 | 8.45E-01 | -2.42E-01 | -1.18E+00 |
| ENSG000001 | IGHMBP2 | 2.67E-01 | 6.20E-01 | 9.49E-01 | -7.57E-02 | -1.05E+00 |
| ENSG000002 | RPSAP53 | 2.67E-01 | ? | 6.70E-01 | -5.78E-01 | -1.49E+00 |
| ENSG000001 | TIMP3 | 2.67E-01 | 6.20E-01 | 9.26E-01 | -1.11E-01 | -1.08E+00 |
| ENSG000001 | ZNF876P | 2.67E-01 | ? | 6.23E-01 | -6.83E-01 | -1.61E+00 |
| ENSG000001 | ZNF593 | 2.67E-01 | 6.20E-01 | 1.10E+00 | 1.32E-01 | 1.10E+00 |
| ENSG000002 | --- | 2.67E-01 | 6.20E-01 | 1.17E+00 | 2.27E-01 | 1.17E+00 |
| ENSG000001 | UNC80 | 2.67E-01 | 6.20E-01 | 8.52E-01 | -2.31E-01 | -1.17E+00 |
| ENSG000001 | ST6GALNAC6 | 2.67E-01 | 6.21E-01 | 1.06E+00 | 7.78E-02 | 1.06E+00 |
| ENSG000001 | WASF1 | 2.67E-01 | 6.21E-01 | 1.05E+00 | 6.43E-02 | 1.05E+00 |
| ENSG000002 | --- | 2.67E-01 | ? | 5.04E-01 | -9.89E-01 | -1.99E+00 |
| ENSG000001 | DEPDC4 | 2.67E-01 | 6.21E-01 | 1.15E+00 | 2.03E-01 | 1.15E+00 |
| ENSG00000C | LAG3 | 2.67E-01 | ? | 7.77E-01 | -3.64E-01 | -1.29E+00 |
| ENSG000001 | DONSON | 2.67E-01 | 6.21E-01 | 9.28E-01 | -1.08E-01 | -1.08E+00 |

| | | | | | | |
|---|---|---|---|---|---|---|
| ENSG000001 | RCC2 | 2.67E-01 | 6.21E-01 | 9.75E-01 | -3.59E-02 | -1.03E+00 |
| ENSG000002 | CD2BP2-DT | 2.67E-01 | 6.21E-01 | 1.18E+00 | 2.33E-01 | 1.18E+00 |
| ENSG000001 | MYO10 | 2.67E-01 | 6.21E-01 | 9.48E-01 | -7.65E-02 | -1.05E+00 |
| ENSG000001 | C1QL3 | 2.67E-01 ? | | 8.17E-01 | -2.91E-01 | -1.22E+00 |
| ENSG000001 | TARS3 | 2.68E-01 | 6.21E-01 | 9.40E-01 | -8.89E-02 | -1.06E+00 |
| ENSG000001 | CSF2 | 2.68E-01 | 6.21E-01 | 8.41E-01 | -2.50E-01 | -1.19E+00 |
| ENSG000001 | ACSL6 | 2.68E-01 ? | | 1.43E+00 | 5.16E-01 | 1.43E+00 |
| ENSG000001 | EEF1D | 2.68E-01 | 6.21E-01 | 1.04E+00 | 6.24E-02 | 1.04E+00 |
| ENSG000001 | CRYM | 2.68E-01 ? | | 1.63E+00 | 7.03E-01 | 1.63E+00 |
| ENSG000001 | GPRASP3 | 2.68E-01 | 6.21E-01 | 1.09E+00 | 1.24E-01 | 1.09E+00 |
| ENSG000002 | --- | 2.68E-01 | 6.21E-01 | 8.22E-01 | -2.83E-01 | -1.22E+00 |
| ENSG000001 | VGF | 2.68E-01 ? | | 6.98E-01 | -5.19E-01 | -1.43E+00 |
| ENSG000001 | HSD17B4 | 2.68E-01 | 6.21E-01 | 1.03E+00 | 4.93E-02 | 1.03E+00 |
| ENSG000001 | MPHOSPH8 | 2.68E-01 | 6.21E-01 | 9.57E-01 | -6.28E-02 | -1.04E+00 |
| ENSG000001 | ZFP28 | 2.68E-01 | 6.22E-01 | 9.51E-01 | -7.18E-02 | -1.05E+00 |
| ENSG000001 | GPR39 | 2.68E-01 | 6.22E-01 | 1.07E+00 | 9.80E-02 | 1.07E+00 |
| ENSG000002 | LINC01004 | 2.68E-01 | 6.22E-01 | 8.27E-01 | -2.74E-01 | -1.21E+00 |
| ENSG000001 | KLC3 | 2.68E-01 | 6.22E-01 | 1.10E+00 | 1.38E-01 | 1.10E+00 |
| ENSG000002 | --- | 2.68E-01 ? | | 1.79E+00 | 8.37E-01 | 1.79E+00 |
| ENSG000001 | XCR1 | 2.68E-01 ? | | 1.67E+00 | 7.38E-01 | 1.67E+00 |
| ENSG00000C | TRIP6 | 2.69E-01 | 6.23E-01 | 1.08E+00 | 1.07E-01 | 1.08E+00 |
| ENSG000001 | HAUS6 | 2.69E-01 | 6.23E-01 | 9.65E-01 | -5.20E-02 | -1.04E+00 |
| ENSG000002 | LINC01985 | 2.69E-01 | 6.23E-01 | 8.46E-01 | -2.41E-01 | -1.18E+00 |
| ENSG000002 | FAM13A-AS1 | 2.69E-01 | 6.23E-01 | 8.81E-01 | -1.82E-01 | -1.13E+00 |
| ENSG000002 | --- | 2.69E-01 ? | | 1.35E+00 | 4.30E-01 | 1.35E+00 |
| ENSG000001 | RAC2 | 2.69E-01 | 6.23E-01 | 1.04E+00 | 5.06E-02 | 1.04E+00 |
| ENSG000002 | --- | 2.69E-01 ? | | 5.74E-01 | -8.00E-01 | -1.74E+00 |
| ENSG00000C | AURKA | 2.69E-01 | 6.23E-01 | 9.29E-01 | -1.06E-01 | -1.08E+00 |
| ENSG000002 | EDIL3-DT | 2.69E-01 ? | | 7.99E-01 | -3.23E-01 | -1.25E+00 |
| ENSG000002 | ZNF702P | 2.69E-01 | 6.24E-01 | 9.21E-01 | -1.18E-01 | -1.09E+00 |
| ENSG000001 | TOP1 | 2.69E-01 | 6.24E-01 | 9.53E-01 | -6.97E-02 | -1.05E+00 |
| ENSG000001 | LYPLAL1 | 2.69E-01 | 6.24E-01 | 9.49E-01 | -7.59E-02 | -1.05E+00 |
| ENSG000001 | GMPS | 2.69E-01 | 6.24E-01 | 1.04E+00 | 4.99E-02 | 1.04E+00 |
| ENSG000001 | METTL25B | 2.70E-01 | 6.24E-01 | 1.09E+00 | 1.23E-01 | 1.09E+00 |
| ENSG000002 | --- | 2.70E-01 ? | | 1.44E+00 | 5.26E-01 | 1.44E+00 |
| ENSG000001 | ZNF836 | 2.70E-01 | 6.24E-01 | 9.38E-01 | -9.25E-02 | -1.07E+00 |
| ENSG000001 | ZFAND2B | 2.70E-01 | 6.24E-01 | 1.05E+00 | 7.59E-02 | 1.05E+00 |
| ENSG000002 | LY6G5B | 2.70E-01 | 6.24E-01 | 8.83E-01 | -1.79E-01 | -1.13E+00 |
| ENSG000002 | RAET1G | 2.70E-01 | 6.24E-01 | 1.09E+00 | 1.27E-01 | 1.09E+00 |
| ENSG000002 | UPF3AP3 | 2.70E-01 ? | | 7.43E-01 | -4.28E-01 | -1.35E+00 |
| ENSG000001 | FGD4 | 2.70E-01 | 6.24E-01 | 9.53E-01 | -6.96E-02 | -1.05E+00 |
| ENSG000001 | PPIC | 2.70E-01 | 6.24E-01 | 1.04E+00 | 5.38E-02 | 1.04E+00 |
| ENSG000002 | CACTIN-AS1 | 2.70E-01 ? | | 7.11E-01 | -4.91E-01 | -1.41E+00 |
| ENSG000002 | BAZ2B-AS1 | 2.70E-01 | 6.24E-01 | 1.14E+00 | 1.85E-01 | 1.14E+00 |
| ENSG000001 | MTBP | 2.70E-01 | 6.24E-01 | 9.01E-01 | -1.50E-01 | -1.11E+00 |

| | | | | | | |
|---|---|---|---|---|---|---|
| ENSG00000C | GNPTG | 2.70E-01 | 6.24E-01 | 1.04E+00 | 6.15E-02 | 1.04E+00 |
| ENSG000001 | RFESD | 2.70E-01 | 6.24E-01 | 8.61E-01 | -2.16E-01 | -1.16E+00 |
| ENSG000002 | --- | 2.70E-01 | ? | 5.81E-01 | -7.83E-01 | -1.72E+00 |
| ENSG000001 | FAF2 | 2.70E-01 | 6.24E-01 | 1.03E+00 | 3.82E-02 | 1.03E+00 |
| ENSG000001 | SEC11A | 2.70E-01 | 6.24E-01 | 9.65E-01 | -5.08E-02 | -1.04E+00 |
| ENSG000002 | MRPL20-DT | 2.70E-01 | 6.24E-01 | 1.14E+00 | 1.94E-01 | 1.14E+00 |
| ENSG000001 | KLK13 | 2.70E-01 | ? | 7.83E-01 | -3.53E-01 | -1.28E+00 |
| ENSG000001 | CENPL | 2.70E-01 | 6.24E-01 | 9.33E-01 | -9.96E-02 | -1.07E+00 |
| ENSG000001 | SYT16 | 2.70E-01 | 6.24E-01 | 8.76E-01 | -1.91E-01 | -1.14E+00 |
| ENSG000002 | B4GAT1-DT | 2.70E-01 | ? | 1.22E+00 | 2.90E-01 | 1.22E+00 |
| ENSG000001 | SNRNP200 | 2.70E-01 | 6.24E-01 | 1.02E+00 | 2.84E-02 | 1.02E+00 |
| ENSG000002 | HOTAIRM1 | 2.70E-01 | 6.24E-01 | 1.09E+00 | 1.19E-01 | 1.09E+00 |
| ENSG000002 | --- | 2.70E-01 | 6.24E-01 | 8.55E-01 | -2.26E-01 | -1.17E+00 |
| ENSG000001 | ANAPC7 | 2.71E-01 | 6.24E-01 | 9.61E-01 | -5.74E-02 | -1.04E+00 |
| ENSG000002 | --- | 2.71E-01 | ? | 7.54E-01 | -4.08E-01 | -1.33E+00 |
| ENSG000001 | CNTD1 | 2.71E-01 | 6.24E-01 | 8.58E-01 | -2.22E-01 | -1.17E+00 |
| ENSG000001 | DNAJC27 | 2.71E-01 | 6.24E-01 | 9.28E-01 | -1.08E-01 | -1.08E+00 |
| ENSG000001 | RASGEF1B | 2.71E-01 | ? | 1.27E+00 | 3.40E-01 | 1.27E+00 |
| ENSG00000C | SLC12A2 | 2.71E-01 | 6.24E-01 | 9.56E-01 | -6.44E-02 | -1.05E+00 |
| ENSG000001 | FBXO41 | 2.71E-01 | 6.24E-01 | 1.06E+00 | 8.51E-02 | 1.06E+00 |
| ENSG00000C | STARD3NL | 2.71E-01 | 6.24E-01 | 9.57E-01 | -6.35E-02 | -1.04E+00 |
| ENSG000002 | DLG5-AS1 | 2.71E-01 | ? | 1.35E+00 | 4.38E-01 | 1.35E+00 |
| ENSG000001 | PRELID2 | 2.71E-01 | 6.25E-01 | 1.09E+00 | 1.25E-01 | 1.09E+00 |
| ENSG000002 | EOLA1-DT | 2.71E-01 | ? | 8.13E-01 | -2.99E-01 | -1.23E+00 |
| ENSG000001 | RELCH | 2.71E-01 | 6.25E-01 | 9.66E-01 | -4.98E-02 | -1.04E+00 |
| ENSG000001 | KLHL24 | 2.71E-01 | 6.25E-01 | 9.32E-01 | -1.01E-01 | -1.07E+00 |
| ENSG00000C | SPATA20 | 2.71E-01 | 6.25E-01 | 1.06E+00 | 8.89E-02 | 1.06E+00 |
| ENSG000002 | TAF12-DT | 2.71E-01 | ? | 1.68E+00 | 7.49E-01 | 1.68E+00 |
| ENSG00000C | SMARCD3 | 2.71E-01 | 6.25E-01 | 9.35E-01 | -9.63E-02 | -1.07E+00 |
| ENSG000002 | --- | 2.71E-01 | ? | 7.72E-01 | -3.74E-01 | -1.30E+00 |
| ENSG000001 | KIF1A | 2.71E-01 | ? | 1.64E+00 | 7.13E-01 | 1.64E+00 |
| ENSG00000C | VAMP3 | 2.71E-01 | 6.25E-01 | 9.73E-01 | -3.98E-02 | -1.03E+00 |
| ENSG000001 | ANAPC11 | 2.71E-01 | 6.25E-01 | 1.10E+00 | 1.35E-01 | 1.10E+00 |
| ENSG000001 | ISCA1 | 2.71E-01 | 6.25E-01 | 9.66E-01 | -5.05E-02 | -1.04E+00 |
| ENSG000001 | AFAP1L2 | 2.71E-01 | 6.25E-01 | 1.02E+00 | 3.54E-02 | 1.02E+00 |
| ENSG000001 | TMEM181 | 2.71E-01 | 6.25E-01 | 1.05E+00 | 6.39E-02 | 1.05E+00 |
| ENSG000001 | IL10RA | 2.71E-01 | ? | 1.67E+00 | 7.39E-01 | 1.67E+00 |
| ENSG000002 | EPB41L4A-D | 2.71E-01 | ? | 1.33E+00 | 4.11E-01 | 1.33E+00 |
| ENSG000001 | CSPP1 | 2.71E-01 | 6.25E-01 | 9.41E-01 | -8.70E-02 | -1.06E+00 |
| ENSG000001 | TMEM74B | 2.72E-01 | 6.25E-01 | 1.20E+00 | 2.58E-01 | 1.20E+00 |
| ENSG000002 | --- | 2.72E-01 | ? | 6.60E-01 | -5.99E-01 | -1.51E+00 |
| ENSG00000C | MTIF2 | 2.72E-01 | 6.25E-01 | 9.70E-01 | -4.44E-02 | -1.03E+00 |
| ENSG00000C | PLEKHA5 | 2.72E-01 | 6.25E-01 | 9.56E-01 | -6.56E-02 | -1.05E+00 |
| ENSG000002 | LINC01235 | 2.72E-01 | ? | 1.57E+00 | 6.54E-01 | 1.57E+00 |
| ENSG000001 | APIP | 2.72E-01 | 6.25E-01 | 9.31E-01 | -1.02E-01 | -1.07E+00 |

| | | | | | | |
|---|---|---|---|---|---|---|
| ENSG000001 | NEK2 | 2.72E-01 | 6.25E-01 | 8.19E-01 | -2.89E-01 | -1.22E+00 |
| ENSG000001 | CIDEB | 2.72E-01 | ? | 7.34E-01 | -4.47E-01 | -1.36E+00 |
| ENSG000001 | PGPEP1 | 2.72E-01 | 6.25E-01 | 1.06E+00 | 8.26E-02 | 1.06E+00 |
| ENSG000002 | --- | 2.72E-01 | ? | 7.02E-01 | -5.10E-01 | -1.42E+00 |
| ENSG000002 | --- | 2.72E-01 | ? | 1.51E+00 | 5.96E-01 | 1.51E+00 |
| ENSG00000C | PRTFDC1 | 2.72E-01 | 6.26E-01 | 1.16E+00 | 2.14E-01 | 1.16E+00 |
| ENSG000001 | SMUG1 | 2.72E-01 | 6.26E-01 | 1.09E+00 | 1.19E-01 | 1.09E+00 |
| ENSG000002 | PAICSP1 | 2.72E-01 | ? | 1.45E+00 | 5.37E-01 | 1.45E+00 |
| ENSG000001 | TAS2R4 | 2.72E-01 | 6.26E-01 | 8.52E-01 | -2.31E-01 | -1.17E+00 |
| ENSG00000C | JADE2 | 2.73E-01 | 6.26E-01 | 9.27E-01 | -1.10E-01 | -1.08E+00 |
| ENSG000002 | KCNMB2-AS1 | 2.73E-01 | ? | 1.75E+00 | 8.07E-01 | 1.75E+00 |
| ENSG000001 | BANP | 2.73E-01 | 6.26E-01 | 9.42E-01 | -8.62E-02 | -1.06E+00 |
| ENSG000001 | RDH5 | 2.73E-01 | ? | 7.34E-01 | -4.47E-01 | -1.36E+00 |
| ENSG000001 | SAPCD2 | 2.73E-01 | 6.27E-01 | 8.04E-01 | -3.14E-01 | -1.24E+00 |
| ENSG000001 | NME9 | 2.73E-01 | ? | 7.29E-01 | -4.57E-01 | -1.37E+00 |
| ENSG00000C | LMAN1 | 2.73E-01 | 6.27E-01 | 9.38E-01 | -9.27E-02 | -1.07E+00 |
| ENSG000002 | SND1-IT1 | 2.73E-01 | ? | 1.65E+00 | 7.26E-01 | 1.65E+00 |
| ENSG000001 | HECTD4 | 2.73E-01 | 6.27E-01 | 9.39E-01 | -9.04E-02 | -1.06E+00 |
| ENSG000002 | LENG9 | 2.73E-01 | 6.27E-01 | 8.70E-01 | -2.01E-01 | -1.15E+00 |
| ENSG000001 | LRRC41 | 2.73E-01 | 6.27E-01 | 1.05E+00 | 6.81E-02 | 1.05E+00 |
| ENSG000001 | PCMT1 | 2.73E-01 | 6.27E-01 | 9.75E-01 | -3.67E-02 | -1.03E+00 |
| ENSG000001 | C3orf33 | 2.73E-01 | 6.27E-01 | 8.99E-01 | -1.54E-01 | -1.11E+00 |
| ENSG000001 | SLC25A6 | 2.74E-01 | 6.27E-01 | 1.07E+00 | 9.83E-02 | 1.07E+00 |
| ENSG000001 | YAP1 | 2.74E-01 | 6.27E-01 | 1.04E+00 | 5.69E-02 | 1.04E+00 |
| ENSG000001 | INTS6 | 2.74E-01 | 6.27E-01 | 9.54E-01 | -6.82E-02 | -1.05E+00 |
| ENSG000002 | MTATP6P1 | 2.74E-01 | 6.27E-01 | 8.47E-01 | -2.40E-01 | -1.18E+00 |
| ENSG000001 | TRIM6 | 2.74E-01 | 6.28E-01 | 9.53E-01 | -6.96E-02 | -1.05E+00 |
| ENSG00000C | IDS | 2.74E-01 | 6.28E-01 | 9.54E-01 | -6.79E-02 | -1.05E+00 |
| ENSG000001 | INTS3 | 2.74E-01 | 6.28E-01 | 1.04E+00 | 5.17E-02 | 1.04E+00 |
| ENSG000001 | POLR3K | 2.74E-01 | 6.28E-01 | 1.11E+00 | 1.47E-01 | 1.11E+00 |
| ENSG000002 | ACY1 | 2.74E-01 | 6.28E-01 | 1.09E+00 | 1.18E-01 | 1.09E+00 |
| ENSG00000C | POLDIP2 | 2.74E-01 | 6.28E-01 | 1.04E+00 | 6.30E-02 | 1.04E+00 |
| ENSG000001 | SLC25A25 | 2.74E-01 | 6.28E-01 | 1.08E+00 | 1.10E-01 | 1.08E+00 |
| ENSG00000C | MGAT4A | 2.74E-01 | 6.28E-01 | 8.98E-01 | -1.56E-01 | -1.11E+00 |
| ENSG000001 | TCF7L2 | 2.74E-01 | 6.28E-01 | 9.37E-01 | -9.38E-02 | -1.07E+00 |
| ENSG000001 | DGLUCY | 2.74E-01 | 6.28E-01 | 9.40E-01 | -8.88E-02 | -1.06E+00 |
| ENSG000001 | PRMT7 | 2.74E-01 | 6.28E-01 | 1.06E+00 | 7.76E-02 | 1.06E+00 |
| ENSG000001 | LIMS1 | 2.74E-01 | 6.28E-01 | 9.64E-01 | -5.36E-02 | -1.04E+00 |
| ENSG000002 | FAM133B | 2.74E-01 | 6.28E-01 | 9.54E-01 | -6.83E-02 | -1.05E+00 |
| ENSG000001 | AVIL | 2.74E-01 | 6.28E-01 | 8.12E-01 | -3.01E-01 | -1.23E+00 |
| ENSG000001 | TMA16 | 2.75E-01 | 6.28E-01 | 9.54E-01 | -6.83E-02 | -1.05E+00 |
| ENSG000002 | --- | 2.75E-01 | ? | 7.59E-01 | -3.97E-01 | -1.32E+00 |
| ENSG000002 | --- | 2.75E-01 | ? | 6.99E-01 | -5.17E-01 | -1.43E+00 |
| ENSG000002 | METTL21EP | 2.75E-01 | ? | 5.52E-01 | -8.57E-01 | -1.81E+00 |
| ENSG000002 | HIF1A-AS3 | 2.75E-01 | 6.29E-01 | 7.83E-01 | -3.52E-01 | -1.28E+00 |

| | | | | | | |
|---|---|---|---|---|---|---|
| ENSG000001 | SFMBT2 | 2.75E-01 | ? | 8.05E-01 | -3.13E-01 | -1.24E+00 |
| ENSG000002 | --- | 2.75E-01 | ? | 6.08E-01 | -7.18E-01 | -1.65E+00 |
| ENSG000002 | GPRIN2 | 2.75E-01 | 6.29E-01 | 1.20E+00 | 2.58E-01 | 1.20E+00 |
| ENSG000002 | LINC00649 | 2.75E-01 | 6.29E-01 | 1.09E+00 | 1.23E-01 | 1.09E+00 |
| ENSG000002 | --- | 2.75E-01 | ? | 6.32E-01 | -6.62E-01 | -1.58E+00 |
| ENSG000002 | DENND6B | 2.75E-01 | 6.29E-01 | 1.10E+00 | 1.41E-01 | 1.10E+00 |
| ENSG000001 | CARD10 | 2.75E-01 | 6.29E-01 | 1.07E+00 | 9.86E-02 | 1.07E+00 |
| ENSG000001 | TBCD | 2.75E-01 | 6.29E-01 | 1.05E+00 | 6.86E-02 | 1.05E+00 |
| ENSG000001 | VPS33A | 2.75E-01 | 6.29E-01 | 9.66E-01 | -4.97E-02 | -1.04E+00 |
| ENSG000002 | --- | 2.75E-01 | ? | 6.89E-01 | -5.38E-01 | -1.45E+00 |
| ENSG000002 | --- | 2.75E-01 | ? | 1.52E+00 | 6.07E-01 | 1.52E+00 |
| ENSG000001 | TLN2 | 2.76E-01 | 6.30E-01 | 1.10E+00 | 1.33E-01 | 1.10E+00 |
| ENSG000001 | MRPS31 | 2.76E-01 | 6.30E-01 | 9.51E-01 | -7.25E-02 | -1.05E+00 |
| ENSG000001 | GGT6 | 2.76E-01 | 6.30E-01 | 1.18E+00 | 2.36E-01 | 1.18E+00 |
| ENSG000002 | HS1BP3-IT1 | 2.76E-01 | ? | 1.55E+00 | 6.34E-01 | 1.55E+00 |
| ENSG000002 | NBPF15 | 2.76E-01 | 6.30E-01 | 9.57E-01 | -6.38E-02 | -1.05E+00 |
| ENSG000002 | CD63-AS1 | 2.76E-01 | 6.30E-01 | 1.10E+00 | 1.36E-01 | 1.10E+00 |
| ENSG000002 | --- | 2.76E-01 | ? | 7.27E-01 | -4.60E-01 | -1.38E+00 |
| ENSG000002 | --- | 2.76E-01 | ? | 6.70E-01 | -5.79E-01 | -1.49E+00 |
| ENSG000001 | XPNPEP1 | 2.76E-01 | 6.30E-01 | 1.03E+00 | 4.88E-02 | 1.03E+00 |
| ENSG000001 | BET1L | 2.76E-01 | 6.30E-01 | 9.53E-01 | -6.91E-02 | -1.05E+00 |
| ENSG000001 | AKR1B10 | 2.76E-01 | 6.30E-01 | 8.28E-01 | -2.72E-01 | -1.21E+00 |
| ENSG000001 | TASOR2 | 2.76E-01 | 6.30E-01 | 1.03E+00 | 4.82E-02 | 1.03E+00 |
| ENSG000001 | UBL4A | 2.76E-01 | 6.30E-01 | 1.08E+00 | 1.07E-01 | 1.08E+00 |
| ENSG000001 | CNP | 2.76E-01 | 6.30E-01 | 1.05E+00 | 6.91E-02 | 1.05E+00 |
| ENSG000002 | ACACA | 2.76E-01 | 6.30E-01 | 1.05E+00 | 7.36E-02 | 1.05E+00 |
| ENSG000001 | EVI2A | 2.76E-01 | ? | 6.85E-01 | -5.46E-01 | -1.46E+00 |
| ENSG000002 | --- | 2.76E-01 | 6.30E-01 | 8.69E-01 | -2.03E-01 | -1.15E+00 |
| ENSG000002 | RPS24P19 | 2.76E-01 | 6.30E-01 | 1.18E+00 | 2.39E-01 | 1.18E+00 |
| ENSG000001 | PDXK | 2.77E-01 | 6.30E-01 | 1.05E+00 | 7.41E-02 | 1.05E+00 |
| ENSG000002 | --- | 2.77E-01 | ? | 7.85E-01 | -3.49E-01 | -1.27E+00 |
| ENSG000001 | DDX54 | 2.77E-01 | 6.30E-01 | 1.05E+00 | 6.60E-02 | 1.05E+00 |
| ENSG000001 | SUGP1 | 2.77E-01 | 6.30E-01 | 9.58E-01 | -6.21E-02 | -1.04E+00 |
| ENSG000002 | --- | 2.77E-01 | 6.31E-01 | 8.47E-01 | -2.40E-01 | -1.18E+00 |
| ENSG000002 | --- | 2.77E-01 | 6.31E-01 | 5.65E-01 | -8.23E-01 | -1.77E+00 |
| ENSG000001 | ARMC9 | 2.77E-01 | 6.31E-01 | 9.56E-01 | -6.46E-02 | -1.05E+00 |
| ENSG000002 | LINC02562 | 2.77E-01 | ? | 1.26E+00 | 3.34E-01 | 1.26E+00 |
| ENSG000001 | SYCE2 | 2.77E-01 | ? | 6.03E-01 | -7.30E-01 | -1.66E+00 |
| ENSG00000C | GDI2 | 2.77E-01 | 6.31E-01 | 9.72E-01 | -4.11E-02 | -1.03E+00 |
| ENSG000001 | CRELD1 | 2.77E-01 | 6.31E-01 | 9.49E-01 | -7.62E-02 | -1.05E+00 |
| ENSG000002 | --- | 2.77E-01 | ? | 6.83E-01 | -5.51E-01 | -1.46E+00 |
| ENSG000002 | --- | 2.77E-01 | 6.31E-01 | 9.38E-01 | -9.30E-02 | -1.07E+00 |
| ENSG000002 | --- | 2.77E-01 | 6.31E-01 | 1.20E+00 | 2.62E-01 | 1.20E+00 |
| ENSG000002 | LINC01415 | 2.77E-01 | ? | 1.36E+00 | 4.41E-01 | 1.36E+00 |
| ENSG000001 | TAF5 | 2.77E-01 | 6.31E-01 | 9.37E-01 | -9.32E-02 | -1.07E+00 |

| | | | | | | |
|---|---|---|---|---|---|---|
| ENSG000001 | TSPO | 2.78E-01 | 6.32E-01 | 1.19E+00 | 2.51E-01 | 1.19E+00 |
| ENSG000002 | EXOSC10-AS | 2.78E-01 | ? | 1.42E+00 | 5.07E-01 | 1.42E+00 |
| ENSG000002 | NEDD8-MDP | 2.78E-01 | ? | 7.68E-01 | -3.81E-01 | -1.30E+00 |
| ENSG000001 | GCSH | 2.78E-01 | 6.32E-01 | 1.06E+00 | 8.10E-02 | 1.06E+00 |
| ENSG000002 | FRMD3-AS1 | 2.78E-01 | ? | 1.66E+00 | 7.34E-01 | 1.66E+00 |
| ENSG000001 | MOB3B | 2.78E-01 | 6.32E-01 | 1.06E+00 | 7.88E-02 | 1.06E+00 |
| ENSG000001 | RELA | 2.78E-01 | 6.32E-01 | 9.67E-01 | -4.82E-02 | -1.03E+00 |
| ENSG000002 | --- | 2.78E-01 | 6.32E-01 | 8.36E-01 | -2.58E-01 | -1.20E+00 |
| ENSG000001 | WDR26 | 2.78E-01 | 6.32E-01 | 9.71E-01 | -4.27E-02 | -1.03E+00 |
| ENSG00000C | MYLIP | 2.78E-01 | 6.32E-01 | 1.11E+00 | 1.46E-01 | 1.11E+00 |
| ENSG000001 | SPECC1L | 2.78E-01 | 6.32E-01 | 9.71E-01 | -4.23E-02 | -1.03E+00 |
| ENSG000001 | ZNF628 | 2.78E-01 | 6.32E-01 | 1.13E+00 | 1.79E-01 | 1.13E+00 |
| ENSG000002 | GOLGA8S | 2.78E-01 | ? | 1.38E+00 | 4.62E-01 | 1.38E+00 |
| ENSG000001 | ZNF148 | 2.78E-01 | 6.32E-01 | 9.59E-01 | -6.10E-02 | -1.04E+00 |
| ENSG000002 | --- | 2.78E-01 | ? | 5.48E-01 | -8.67E-01 | -1.82E+00 |
| ENSG000002 | --- | 2.78E-01 | ? | 1.23E+00 | 2.94E-01 | 1.23E+00 |
| ENSG000001 | GNG2 | 2.78E-01 | 6.33E-01 | 8.52E-01 | -2.31E-01 | -1.17E+00 |
| ENSG000002 | --- | 2.78E-01 | ? | 6.22E-01 | -6.85E-01 | -1.61E+00 |
| ENSG000002 | CRPPA | 2.79E-01 | 6.33E-01 | 1.09E+00 | 1.26E-01 | 1.09E+00 |
| ENSG000001 | ZNF496 | 2.79E-01 | 6.33E-01 | 1.05E+00 | 7.12E-02 | 1.05E+00 |
| ENSG000001 | CCDC24 | 2.79E-01 | 6.33E-01 | 1.10E+00 | 1.34E-01 | 1.10E+00 |
| ENSG000001 | PINK1 | 2.79E-01 | 6.33E-01 | 1.05E+00 | 7.37E-02 | 1.05E+00 |
| ENSG000002 | --- | 2.79E-01 | ? | 6.14E-01 | -7.03E-01 | -1.63E+00 |
| ENSG000002 | WDR35-DT | 2.79E-01 | ? | 1.27E+00 | 3.43E-01 | 1.27E+00 |
| ENSG000001 | CCT2 | 2.79E-01 | 6.33E-01 | 9.72E-01 | -4.14E-02 | -1.03E+00 |
| ENSG000001 | SLCO5A1 | 2.79E-01 | ? | 1.52E+00 | 6.08E-01 | 1.52E+00 |
| ENSG000001 | MMP17 | 2.79E-01 | 6.33E-01 | 1.11E+00 | 1.47E-01 | 1.11E+00 |
| ENSG000002 | HULC | 2.79E-01 | 6.33E-01 | 1.17E+00 | 2.32E-01 | 1.17E+00 |
| ENSG000002 | SYT15B | 2.79E-01 | 6.33E-01 | 8.96E-01 | -1.59E-01 | -1.12E+00 |
| ENSG000001 | ZFAS1 | 2.79E-01 | 6.33E-01 | 1.04E+00 | 4.98E-02 | 1.04E+00 |
| ENSG000001 | LRRCC1 | 2.79E-01 | 6.33E-01 | 8.81E-01 | -1.83E-01 | -1.14E+00 |
| ENSG000001 | TEX10 | 2.79E-01 | 6.33E-01 | 1.03E+00 | 4.96E-02 | 1.03E+00 |
| ENSG000001 | C1D | 2.79E-01 | 6.33E-01 | 9.50E-01 | -7.45E-02 | -1.05E+00 |
| ENSG000001 | SULT1E1 | 2.79E-01 | ? | 7.74E-01 | -3.69E-01 | -1.29E+00 |
| ENSG000001 | USP39 | 2.79E-01 | 6.33E-01 | 9.65E-01 | -5.08E-02 | -1.04E+00 |
| ENSG000002 | --- | 2.79E-01 | 6.33E-01 | 1.13E+00 | 1.76E-01 | 1.13E+00 |
| ENSG000001 | CLCF1 | 2.79E-01 | 6.33E-01 | 9.15E-01 | -1.28E-01 | -1.09E+00 |
| ENSG000001 | CHID1 | 2.80E-01 | 6.33E-01 | 1.06E+00 | 9.03E-02 | 1.06E+00 |
| ENSG000001 | CDK5RAP2 | 2.80E-01 | 6.33E-01 | 9.59E-01 | -5.99E-02 | -1.04E+00 |
| ENSG000002 | LINC01806 | 2.80E-01 | 6.33E-01 | 1.10E+00 | 1.31E-01 | 1.10E+00 |
| ENSG000001 | IVNS1ABP | 2.80E-01 | 6.33E-01 | 9.51E-01 | -7.22E-02 | -1.05E+00 |
| ENSG00000C | BCAR1 | 2.80E-01 | 6.33E-01 | 1.07E+00 | 1.01E-01 | 1.07E+00 |
| ENSG000001 | LRFN4 | 2.80E-01 | 6.33E-01 | 1.10E+00 | 1.35E-01 | 1.10E+00 |
| ENSG00000C | GOPC | 2.80E-01 | 6.34E-01 | 9.61E-01 | -5.78E-02 | -1.04E+00 |
| ENSG000002 | --- | 2.80E-01 | ? | 5.26E-01 | -9.28E-01 | -1.90E+00 |

| | | | | | | |
|---|---|---|---|---|---|---|
| ENSG000001 | LRRC2 | 2.80E-01 | ? | 7.53E-01 | -4.09E-01 | -1.33E+00 |
| ENSG000001 | OPRD1 | 2.80E-01 | ? | 7.07E-01 | -5.00E-01 | -1.41E+00 |
| ENSG000001 | SIGIRR | 2.80E-01 | 6.34E-01 | 1.15E+00 | 1.97E-01 | 1.15E+00 |
| ENSG000001 | NXT2 | 2.80E-01 | 6.34E-01 | 1.06E+00 | 7.84E-02 | 1.06E+00 |
| ENSG000002 | --- | 2.80E-01 | ? | 6.33E-01 | -6.61E-01 | -1.58E+00 |
| ENSG000001 | ZNF223 | 2.80E-01 | 6.34E-01 | 8.94E-01 | -1.62E-01 | -1.12E+00 |
| ENSG000002 | PCDHA4 | 2.80E-01 | ? | 8.01E-01 | -3.20E-01 | -1.25E+00 |
| ENSG000001 | SART1 | 2.80E-01 | 6.34E-01 | 9.63E-01 | -5.45E-02 | -1.04E+00 |
| ENSG000001 | SAA1 | 2.80E-01 | 6.34E-01 | 1.13E+00 | 1.77E-01 | 1.13E+00 |
| ENSG000002 | --- | 2.80E-01 | ? | 6.63E-01 | -5.93E-01 | -1.51E+00 |
| ENSG000002 | DNM1P41 | 2.80E-01 | ? | 7.55E-01 | -4.06E-01 | -1.32E+00 |
| ENSG000001 | ATP5MC3 | 2.80E-01 | 6.34E-01 | 1.05E+00 | 7.29E-02 | 1.05E+00 |
| ENSG000001 | MRPL55 | 2.80E-01 | 6.34E-01 | 1.09E+00 | 1.25E-01 | 1.09E+00 |
| ENSG000002 | --- | 2.80E-01 | ? | 1.61E+00 | 6.87E-01 | 1.61E+00 |
| ENSG000002 | GTF2IP1 | 2.81E-01 | 6.34E-01 | 8.16E-01 | -2.94E-01 | -1.23E+00 |
| ENSG000001 | CENATAC | 2.81E-01 | 6.34E-01 | 9.09E-01 | -1.38E-01 | -1.10E+00 |
| ENSG000001 | JUN | 2.81E-01 | 6.34E-01 | 9.53E-01 | -6.99E-02 | -1.05E+00 |
| ENSG000001 | ZNF875 | 2.81E-01 | 6.34E-01 | 9.44E-01 | -8.34E-02 | -1.06E+00 |
| ENSG000001 | UBL3 | 2.81E-01 | 6.34E-01 | 9.55E-01 | -6.63E-02 | -1.05E+00 |
| ENSG00000C | SH3GLB1 | 2.81E-01 | 6.34E-01 | 9.63E-01 | -5.40E-02 | -1.04E+00 |
| ENSG000001 | SFTPD | 2.81E-01 | ? | 1.34E+00 | 4.18E-01 | 1.34E+00 |
| ENSG000001 | CDCA8 | 2.81E-01 | 6.35E-01 | 9.37E-01 | -9.40E-02 | -1.07E+00 |
| ENSG000001 | KLK10 | 2.81E-01 | 6.35E-01 | 8.58E-01 | -2.22E-01 | -1.17E+00 |
| ENSG000002 | --- | 2.81E-01 | ? | 5.08E-01 | -9.77E-01 | -1.97E+00 |
| ENSG000002 | --- | 2.81E-01 | 6.35E-01 | 1.16E+00 | 2.17E-01 | 1.16E+00 |
| ENSG000002 | --- | 2.81E-01 | ? | 5.68E-01 | -8.16E-01 | -1.76E+00 |
| ENSG000002 | MTCH1P2 | 2.81E-01 | ? | 1.50E+00 | 5.87E-01 | 1.50E+00 |
| ENSG000001 | EPHA10 | 2.81E-01 | ? | 1.60E+00 | 6.77E-01 | 1.60E+00 |
| ENSG000001 | NPR3 | 2.81E-01 | ? | 1.26E+00 | 3.37E-01 | 1.26E+00 |
| ENSG000001 | RHPN1 | 2.81E-01 | 6.35E-01 | 1.15E+00 | 2.04E-01 | 1.15E+00 |
| ENSG00000C | GCN1 | 2.81E-01 | 6.35E-01 | 1.04E+00 | 5.12E-02 | 1.04E+00 |
| ENSG000002 | SNX2 | 2.81E-01 | 6.35E-01 | 1.04E+00 | 6.03E-02 | 1.04E+00 |
| ENSG000001 | TIMM9 | 2.81E-01 | 6.35E-01 | 9.56E-01 | -6.52E-02 | -1.05E+00 |
| ENSG000002 | --- | 2.81E-01 | 6.35E-01 | 8.38E-01 | -2.55E-01 | -1.19E+00 |
| ENSG000001 | SLC2A4RG | 2.81E-01 | 6.35E-01 | 1.08E+00 | 1.13E-01 | 1.08E+00 |
| ENSG000002 | ZNF844 | 2.81E-01 | 6.35E-01 | 1.08E+00 | 1.05E-01 | 1.08E+00 |
| ENSG000001 | FZR1 | 2.82E-01 | 6.35E-01 | 1.04E+00 | 6.31E-02 | 1.04E+00 |
| ENSG000001 | PYGB | 2.82E-01 | 6.35E-01 | 1.05E+00 | 6.98E-02 | 1.05E+00 |
| ENSG000001 | PTGES3 | 2.82E-01 | 6.35E-01 | 9.80E-01 | -2.87E-02 | -1.02E+00 |
| ENSG000001 | NRF1 | 2.82E-01 | 6.35E-01 | 1.06E+00 | 8.27E-02 | 1.06E+00 |
| ENSG000001 | PRSS36 | 2.82E-01 | ? | 5.85E-01 | -7.74E-01 | -1.71E+00 |
| ENSG000001 | USP7 | 2.82E-01 | 6.35E-01 | 9.63E-01 | -5.48E-02 | -1.04E+00 |
| ENSG000001 | IL37 | 2.82E-01 | ? | 6.56E-01 | -6.09E-01 | -1.53E+00 |
| ENSG000001 | NAPA | 2.82E-01 | 6.35E-01 | 1.04E+00 | 4.96E-02 | 1.04E+00 |
| ENSG00000C | MAP3K1 | 2.82E-01 | 6.35E-01 | 1.05E+00 | 7.10E-02 | 1.05E+00 |

| | | | | | | |
|---|---|---|---|---|---|---|
| ENSG000000 | BTAF1 | 2.82E-01 | 6.35E-01 | 9.55E-01 | -6.64E-02 | -1.05E+00 |
| ENSG000000 | HEATR5B | 2.82E-01 | 6.35E-01 | 9.71E-01 | -4.18E-02 | -1.03E+00 |
| ENSG000002 | --- | 2.82E-01 | 6.35E-01 | 1.09E+00 | 1.19E-01 | 1.09E+00 |
| ENSG000001 | ARMT1 | 2.82E-01 | 6.35E-01 | 1.04E+00 | 5.84E-02 | 1.04E+00 |
| ENSG000001 | UPP1 | 2.82E-01 | 6.35E-01 | 9.55E-01 | -6.58E-02 | -1.05E+00 |
| ENSG000002 | ZNF582-DT | 2.82E-01 | 6.35E-01 | 8.77E-01 | -1.90E-01 | -1.14E+00 |
| ENSG000001 | BATF2 | 2.82E-01 | ? | 1.26E+00 | 3.32E-01 | 1.26E+00 |
| ENSG000001 | CCDC88A | 2.82E-01 | 6.35E-01 | 9.49E-01 | -7.50E-02 | -1.05E+00 |
| ENSG000001 | CALB2 | 2.82E-01 | ? | 1.39E+00 | 4.70E-01 | 1.39E+00 |
| ENSG000002 | --- | 2.82E-01 | 6.35E-01 | 1.15E+00 | 1.96E-01 | 1.15E+00 |
| ENSG000002 | --- | 2.82E-01 | ? | 1.29E+00 | 3.69E-01 | 1.29E+00 |
| ENSG000001 | NR1I3 | 2.82E-01 | ? | 7.85E-01 | -3.50E-01 | -1.27E+00 |
| ENSG000002 | AKR1B10P1 | 2.82E-01 | ? | 6.33E-01 | -6.59E-01 | -1.58E+00 |
| ENSG000001 | ACO2 | 2.82E-01 | 6.35E-01 | 1.03E+00 | 4.50E-02 | 1.03E+00 |
| ENSG000001 | KCNJ15 | 2.83E-01 | 6.35E-01 | 9.38E-01 | -9.17E-02 | -1.07E+00 |
| ENSG000001 | PRR5 | 2.83E-01 | 6.35E-01 | 1.12E+00 | 1.62E-01 | 1.12E+00 |
| ENSG000000 | SLC9A3R2 | 2.83E-01 | 6.35E-01 | 9.43E-01 | -8.48E-02 | -1.06E+00 |
| ENSG000001 | EVA1B | 2.83E-01 | ? | 1.38E+00 | 4.62E-01 | 1.38E+00 |
| ENSG000000 | IDI1 | 2.83E-01 | 6.36E-01 | 9.56E-01 | -6.43E-02 | -1.05E+00 |
| ENSG000001 | MYRFL | 2.83E-01 | ? | 1.43E+00 | 5.20E-01 | 1.43E+00 |
| ENSG000002 | NUDT3 | 2.83E-01 | 6.36E-01 | 9.57E-01 | -6.41E-02 | -1.05E+00 |
| ENSG000001 | PLCB3 | 2.83E-01 | 6.36E-01 | 1.04E+00 | 5.46E-02 | 1.04E+00 |
| ENSG000001 | JAG2 | 2.83E-01 | 6.36E-01 | 1.09E+00 | 1.26E-01 | 1.09E+00 |
| ENSG000001 | MITF | 2.83E-01 | 6.36E-01 | 1.06E+00 | 8.11E-02 | 1.06E+00 |
| ENSG000001 | NPIPB15 | 2.83E-01 | 6.36E-01 | 9.26E-01 | -1.11E-01 | -1.08E+00 |
| ENSG000001 | LRCH1 | 2.83E-01 | 6.36E-01 | 1.05E+00 | 6.83E-02 | 1.05E+00 |
| ENSG000002 | UXT-AS1 | 2.83E-01 | ? | 1.27E+00 | 3.45E-01 | 1.27E+00 |
| ENSG000000 | RIPOR3 | 2.83E-01 | ? | 7.65E-01 | -3.86E-01 | -1.31E+00 |
| ENSG000001 | LIAS | 2.83E-01 | 6.37E-01 | 9.50E-01 | -7.40E-02 | -1.05E+00 |
| ENSG000001 | PLG | 2.84E-01 | ? | 1.41E+00 | 4.91E-01 | 1.41E+00 |
| ENSG000000 | APLP2 | 2.84E-01 | 6.37E-01 | 9.62E-01 | -5.65E-02 | -1.04E+00 |
| ENSG000000 | BARX2 | 2.84E-01 | 6.37E-01 | 9.04E-01 | -1.45E-01 | -1.11E+00 |
| ENSG000001 | PSMD6 | 2.84E-01 | 6.37E-01 | 9.73E-01 | -4.02E-02 | -1.03E+00 |
| ENSG000001 | AOPEP | 2.84E-01 | 6.37E-01 | 1.05E+00 | 7.41E-02 | 1.05E+00 |
| ENSG000002 | LIPK | 2.84E-01 | ? | 1.65E+00 | 7.21E-01 | 1.65E+00 |
| ENSG000001 | PLK2 | 2.84E-01 | 6.37E-01 | 1.05E+00 | 7.59E-02 | 1.05E+00 |
| ENSG000001 | SLC29A3 | 2.84E-01 | 6.37E-01 | 1.07E+00 | 1.01E-01 | 1.07E+00 |
| ENSG000002 | ST13P6 | 2.84E-01 | ? | 7.83E-01 | -3.54E-01 | -1.28E+00 |
| ENSG000001 | FOLR1 | 2.84E-01 | 6.37E-01 | 1.12E+00 | 1.65E-01 | 1.12E+00 |
| ENSG000002 | --- | 2.84E-01 | ? | 7.16E-01 | -4.83E-01 | -1.40E+00 |
| ENSG000002 | FOXP4-AS1 | 2.84E-01 | ? | 6.26E-01 | -6.75E-01 | -1.60E+00 |
| ENSG000001 | BMP1 | 2.84E-01 | 6.38E-01 | 1.05E+00 | 7.43E-02 | 1.05E+00 |
| ENSG000001 | PSTPIP1 | 2.84E-01 | 6.38E-01 | 8.49E-01 | -2.37E-01 | -1.18E+00 |
| ENSG000001 | FAM120C | 2.84E-01 | 6.38E-01 | 8.97E-01 | -1.57E-01 | -1.11E+00 |
| ENSG000001 | HDHD2 | 2.84E-01 | 6.38E-01 | 1.06E+00 | 8.56E-02 | 1.06E+00 |

| | | | | | | |
|---|---|---|---|---|---|---|
| ENSG000002 | --- | 2.85E-01 | 6.38E-01 | 8.53E-01 | -2.29E-01 | -1.17E+00 |
| ENSG000002 | PRPF31-AS1 | 2.85E-01 | ? | 8.45E-01 | -2.44E-01 | -1.18E+00 |
| ENSG000002 | ZNF473CR | 2.85E-01 | ? | 1.29E+00 | 3.71E-01 | 1.29E+00 |
| ENSG00000C | TPD52 | 2.85E-01 | 6.38E-01 | 9.62E-01 | -5.52E-02 | -1.04E+00 |
| ENSG000001 | GPC6 | 2.85E-01 | 6.38E-01 | 8.99E-01 | -1.54E-01 | -1.11E+00 |
| ENSG000001 | SF3B1 | 2.85E-01 | 6.38E-01 | 9.73E-01 | -4.02E-02 | -1.03E+00 |
| ENSG000001 | MPST | 2.85E-01 | 6.38E-01 | 1.08E+00 | 1.11E-01 | 1.08E+00 |
| ENSG000001 | DKK2 | 2.85E-01 | 6.38E-01 | 8.81E-01 | -1.82E-01 | -1.13E+00 |
| ENSG000001 | NMRAL1 | 2.85E-01 | 6.38E-01 | 1.06E+00 | 7.91E-02 | 1.06E+00 |
| ENSG000001 | ATRIP | 2.85E-01 | 6.38E-01 | 1.06E+00 | 8.65E-02 | 1.06E+00 |
| ENSG000002 | RALY-AS1 | 2.85E-01 | 6.38E-01 | 1.12E+00 | 1.60E-01 | 1.12E+00 |
| ENSG000002 | MFF-DT | 2.85E-01 | 6.38E-01 | 8.77E-01 | -1.89E-01 | -1.14E+00 |
| ENSG000001 | RIPK2 | 2.85E-01 | 6.38E-01 | 9.52E-01 | -7.11E-02 | -1.05E+00 |
| ENSG000002 | CD27-AS1 | 2.85E-01 | 6.38E-01 | 1.11E+00 | 1.54E-01 | 1.11E+00 |
| ENSG000002 | LINC00310 | 2.85E-01 | ? | 5.17E-01 | -9.51E-01 | -1.93E+00 |
| ENSG000001 | RAE1 | 2.85E-01 | 6.39E-01 | 1.04E+00 | 5.19E-02 | 1.04E+00 |
| ENSG000001 | RWDD1 | 2.85E-01 | 6.39E-01 | 9.51E-01 | -7.19E-02 | -1.05E+00 |
| ENSG00000C | MARCHF2 | 2.86E-01 | 6.39E-01 | 1.07E+00 | 9.10E-02 | 1.07E+00 |
| ENSG000001 | TIMELESS | 2.86E-01 | 6.39E-01 | 1.06E+00 | 7.80E-02 | 1.06E+00 |
| ENSG00000C | PTPN4 | 2.86E-01 | 6.39E-01 | 9.56E-01 | -6.55E-02 | -1.05E+00 |
| ENSG000001 | SYT17 | 2.86E-01 | 6.39E-01 | 1.15E+00 | 2.02E-01 | 1.15E+00 |
| ENSG000001 | STMN3 | 2.86E-01 | 6.39E-01 | 1.17E+00 | 2.22E-01 | 1.17E+00 |
| ENSG00000C | UQCRC1 | 2.86E-01 | 6.39E-01 | 1.05E+00 | 6.38E-02 | 1.05E+00 |
| ENSG00000C | TNFRSF9 | 2.86E-01 | ? | 1.40E+00 | 4.90E-01 | 1.40E+00 |
| ENSG000001 | KLK1 | 2.86E-01 | ? | 4.96E-01 | -1.01E+00 | -2.02E+00 |
| ENSG000002 | ARPIN-AP3S: | 2.86E-01 | 6.39E-01 | 9.38E-01 | -9.18E-02 | -1.07E+00 |
| ENSG000002 | ACTN1-DT | 2.86E-01 | ? | 5.63E-01 | -8.29E-01 | -1.78E+00 |
| ENSG000002 | GNA15-DT | 2.86E-01 | ? | 6.73E-01 | -5.72E-01 | -1.49E+00 |
| ENSG000001 | IL1R2 | 2.86E-01 | 6.39E-01 | 8.26E-01 | -2.75E-01 | -1.21E+00 |
| ENSG00000C | SNAP23 | 2.86E-01 | 6.39E-01 | 9.59E-01 | -6.03E-02 | -1.04E+00 |
| ENSG000001 | FLRT2 | 2.86E-01 | 6.40E-01 | 9.54E-01 | -6.81E-02 | -1.05E+00 |
| ENSG000001 | ATG9B | 2.86E-01 | ? | 7.84E-01 | -3.50E-01 | -1.27E+00 |
| ENSG000001 | SLC35C1 | 2.86E-01 | 6.40E-01 | 1.05E+00 | 7.32E-02 | 1.05E+00 |
| ENSG000002 | --- | 2.86E-01 | ? | 8.22E-01 | -2.82E-01 | -1.22E+00 |
| ENSG000001 | NPIPA9 | 2.87E-01 | 6.40E-01 | 9.43E-01 | -8.53E-02 | -1.06E+00 |
| ENSG000002 | --- | 2.87E-01 | ? | 1.64E+00 | 7.10E-01 | 1.64E+00 |
| ENSG000001 | POLR1G | 2.87E-01 | 6.40E-01 | 1.05E+00 | 7.10E-02 | 1.05E+00 |
| ENSG000001 | ADAMTS10 | 2.87E-01 | ? | 1.81E+00 | 8.54E-01 | 1.81E+00 |
| ENSG000001 | CHKA | 2.87E-01 | 6.40E-01 | 1.07E+00 | 1.04E-01 | 1.07E+00 |
| ENSG00000C | TBPL1 | 2.87E-01 | 6.40E-01 | 1.05E+00 | 7.38E-02 | 1.05E+00 |
| ENSG000001 | ZNF696 | 2.87E-01 | 6.40E-01 | 9.41E-01 | -8.79E-02 | -1.06E+00 |
| ENSG000002 | RRAS2P1 | 2.87E-01 | ? | 7.81E-01 | -3.57E-01 | -1.28E+00 |
| ENSG000002 | --- | 2.87E-01 | 6.40E-01 | 1.26E+00 | 3.34E-01 | 1.26E+00 |
| ENSG000002 | SOGA3 | 2.87E-01 | ? | 7.82E-01 | -3.55E-01 | -1.28E+00 |
| ENSG000002 | --- | 2.87E-01 | ? | 7.63E-01 | -3.89E-01 | -1.31E+00 |

| | | | | | | | |
|---|---|---|---|---|---|---|---|
| ENSG000002 | --- | 2.87E-01 | ? | | 5.98E-01 | -7.42E-01 | -1.67E+00 |
| ENSG000001 | CNNM4 | 2.87E-01 | | 6.40E-01 | 1.05E+00 | 6.79E-02 | 1.05E+00 |
| ENSG000001 | SNX30 | 2.87E-01 | | 6.40E-01 | 1.05E+00 | 7.72E-02 | 1.05E+00 |
| ENSG000001 | CD81 | 2.87E-01 | | 6.40E-01 | 1.06E+00 | 8.97E-02 | 1.06E+00 |
| ENSG000001 | SPSB4 | 2.87E-01 | ? | | 1.40E+00 | 4.84E-01 | 1.40E+00 |
| ENSG000002 | --- | 2.87E-01 | | 6.40E-01 | 9.45E-01 | -8.18E-02 | -1.06E+00 |
| ENSG000001 | SLIRP | 2.87E-01 | | 6.40E-01 | 1.06E+00 | 8.95E-02 | 1.06E+00 |
| ENSG000002 | MAP4K3-DT | 2.87E-01 | | 6.40E-01 | 9.20E-01 | -1.21E-01 | -1.09E+00 |
| ENSG000002 | --- | 2.87E-01 | ? | | 1.35E+00 | 4.30E-01 | 1.35E+00 |
| ENSG000001 | GPSM1 | 2.87E-01 | | 6.41E-01 | 1.09E+00 | 1.20E-01 | 1.09E+00 |
| ENSG000001 | JRKL | 2.88E-01 | | 6.41E-01 | 9.47E-01 | -7.82E-02 | -1.06E+00 |
| ENSG000002 | LINC02912 | 2.88E-01 | ? | | 1.66E+00 | 7.33E-01 | 1.66E+00 |
| ENSG000001 | THNSL1 | 2.88E-01 | | 6.41E-01 | 1.07E+00 | 9.14E-02 | 1.07E+00 |
| ENSG000001 | GAS2 | 2.88E-01 | ? | | 8.06E-01 | -3.11E-01 | -1.24E+00 |
| ENSG000001 | TUBB | 2.88E-01 | | 6.41E-01 | 1.04E+00 | 5.05E-02 | 1.04E+00 |
| ENSG000001 | API5 | 2.88E-01 | | 6.41E-01 | 9.72E-01 | -4.09E-02 | -1.03E+00 |
| ENSG000001 | MKLN1 | 2.88E-01 | | 6.41E-01 | 9.60E-01 | -5.84E-02 | -1.04E+00 |
| ENSG000001 | SNRPD3 | 2.88E-01 | | 6.41E-01 | 1.03E+00 | 4.88E-02 | 1.03E+00 |
| ENSG000001 | ILDR1 | 2.88E-01 | ? | | 1.26E+00 | 3.37E-01 | 1.26E+00 |
| ENSG000002 | MYBPHL | 2.88E-01 | ? | | 1.40E+00 | 4.86E-01 | 1.40E+00 |
| ENSG000001 | POP1 | 2.88E-01 | | 6.42E-01 | 9.41E-01 | -8.71E-02 | -1.06E+00 |
| ENSG000001 | PER1 | 2.88E-01 | | 6.42E-01 | 1.08E+00 | 1.08E-01 | 1.08E+00 |
| ENSG000002 | --- | 2.88E-01 | ? | | 1.22E+00 | 2.91E-01 | 1.22E+00 |
| ENSG000001 | ZNF433 | 2.89E-01 | | 6.42E-01 | 1.08E+00 | 1.12E-01 | 1.08E+00 |
| ENSG000002 | --- | 2.89E-01 | ? | | 1.37E+00 | 4.50E-01 | 1.37E+00 |
| ENSG000001 | MXRA7 | 2.89E-01 | | 6.42E-01 | 1.03E+00 | 4.38E-02 | 1.03E+00 |
| ENSG000001 | MAST1 | 2.89E-01 | ? | | 7.81E-01 | -3.57E-01 | -1.28E+00 |
| ENSG000002 | SPTY2D1OS | 2.89E-01 | ? | | 6.22E-01 | -6.85E-01 | -1.61E+00 |
| ENSG000002 | CD24 | 2.89E-01 | | 6.42E-01 | 1.05E+00 | 6.62E-02 | 1.05E+00 |
| ENSG000002 | MEX3A | 2.89E-01 | | 6.42E-01 | 9.26E-01 | -1.11E-01 | -1.08E+00 |
| ENSG00000C | PLEKHG6 | 2.89E-01 | | 6.42E-01 | 1.18E+00 | 2.34E-01 | 1.18E+00 |
| ENSG000002 | PTPN23-DT | 2.89E-01 | | 6.42E-01 | 1.20E+00 | 2.57E-01 | 1.20E+00 |
| ENSG000001 | TREML1 | 2.89E-01 | ? | | 1.26E+00 | 3.38E-01 | 1.26E+00 |
| ENSG000001 | MGAT4B | 2.89E-01 | | 6.42E-01 | 1.06E+00 | 8.85E-02 | 1.06E+00 |
| ENSG000001 | NELL2 | 2.89E-01 | ? | | 1.58E+00 | 6.64E-01 | 1.58E+00 |
| ENSG000001 | NLRC3 | 2.89E-01 | ? | | 8.17E-01 | -2.91E-01 | -1.22E+00 |
| ENSG000001 | ENPP1 | 2.89E-01 | | 6.42E-01 | 1.18E+00 | 2.39E-01 | 1.18E+00 |
| ENSG000002 | ZNF850 | 2.89E-01 | | 6.43E-01 | 1.08E+00 | 1.11E-01 | 1.08E+00 |
| ENSG000001 | ARNT2 | 2.89E-01 | | 6.43E-01 | 9.11E-01 | -1.34E-01 | -1.10E+00 |
| ENSG000002 | --- | 2.89E-01 | ? | | 1.64E+00 | 7.15E-01 | 1.64E+00 |
| ENSG000002 | YBX1P1 | 2.89E-01 | | 6.43E-01 | 9.23E-01 | -1.16E-01 | -1.08E+00 |
| ENSG000001 | DPH6 | 2.89E-01 | | 6.43E-01 | 1.08E+00 | 1.13E-01 | 1.08E+00 |
| ENSG000002 | --- | 2.89E-01 | ? | | 7.82E-01 | -3.54E-01 | -1.28E+00 |
| ENSG000001 | SPRR3 | 2.89E-01 | | 6.43E-01 | 7.91E-01 | -3.39E-01 | -1.26E+00 |
| ENSG000001 | SLC27A4 | 2.89E-01 | | 6.43E-01 | 1.07E+00 | 9.21E-02 | 1.07E+00 |

| | | | | | | |
|---|---|---|---|---|---|---|
| ENSG000002 | LRRC37A9P | 2.89E-01 | ? | 1.66E+00 | 7.30E-01 | 1.66E+00 |
| ENSG000001 | SCRN1 | 2.90E-01 | 6.43E-01 | 1.03E+00 | 4.00E-02 | 1.03E+00 |
| ENSG000001 | DSG3 | 2.90E-01 | 6.43E-01 | 1.05E+00 | 6.69E-02 | 1.05E+00 |
| ENSG000001 | DNAH6 | 2.90E-01 | ? | 1.32E+00 | 4.03E-01 | 1.32E+00 |
| ENSG000002 | CYP2T1P | 2.90E-01 | ? | 1.42E+00 | 5.07E-01 | 1.42E+00 |
| ENSG000002 | LINC01569 | 2.90E-01 | ? | 1.25E+00 | 3.21E-01 | 1.25E+00 |
| ENSG000002 | TCAF1P1 | 2.90E-01 | 6.43E-01 | 1.13E+00 | 1.78E-01 | 1.13E+00 |
| ENSG000001 | NAIF1 | 2.90E-01 | 6.43E-01 | 1.06E+00 | 9.05E-02 | 1.06E+00 |
| ENSG000001 | ROCK2 | 2.90E-01 | 6.43E-01 | 9.54E-01 | -6.82E-02 | -1.05E+00 |
| ENSG000001 | AKT2 | 2.90E-01 | 6.43E-01 | 1.04E+00 | 5.44E-02 | 1.04E+00 |
| ENSG000002 | LINC00992 | 2.90E-01 | ? | 1.30E+00 | 3.73E-01 | 1.30E+00 |
| ENSG000001 | CHADL | 2.90E-01 | ? | 1.30E+00 | 3.76E-01 | 1.30E+00 |
| ENSG00000C | YAF2 | 2.90E-01 | 6.43E-01 | 9.38E-01 | -9.21E-02 | -1.07E+00 |
| ENSG000001 | PDLIM5 | 2.90E-01 | 6.43E-01 | 1.03E+00 | 4.58E-02 | 1.03E+00 |
| ENSG000002 | --- | 2.90E-01 | ? | 5.66E-01 | -8.22E-01 | -1.77E+00 |
| ENSG000002 | RBMXL1 | 2.90E-01 | 6.44E-01 | 1.06E+00 | 9.08E-02 | 1.06E+00 |
| ENSG000001 | CD83 | 2.90E-01 | 6.44E-01 | 1.12E+00 | 1.58E-01 | 1.12E+00 |
| ENSG000001 | SLC22A18 | 2.90E-01 | 6.44E-01 | 1.15E+00 | 2.00E-01 | 1.15E+00 |
| ENSG000001 | AGO4 | 2.90E-01 | 6.44E-01 | 9.52E-01 | -7.10E-02 | -1.05E+00 |
| ENSG000001 | RAP1B | 2.91E-01 | 6.44E-01 | 1.05E+00 | 7.09E-02 | 1.05E+00 |
| ENSG000001 | PBXIP1 | 2.91E-01 | 6.44E-01 | 1.05E+00 | 7.48E-02 | 1.05E+00 |
| ENSG000001 | ENG | 2.91E-01 | 6.44E-01 | 1.11E+00 | 1.45E-01 | 1.11E+00 |
| ENSG000002 | --- | 2.91E-01 | ? | 1.72E+00 | 7.86E-01 | 1.72E+00 |
| ENSG00000C | STOML1 | 2.91E-01 | 6.44E-01 | 1.07E+00 | 9.66E-02 | 1.07E+00 |
| ENSG000001 | ZNF250 | 2.91E-01 | 6.44E-01 | 1.06E+00 | 7.73E-02 | 1.06E+00 |
| ENSG000002 | C12orf73 | 2.91E-01 | 6.45E-01 | 9.21E-01 | -1.18E-01 | -1.09E+00 |
| ENSG000001 | SAMD15 | 2.91E-01 | 6.45E-01 | 1.14E+00 | 1.89E-01 | 1.14E+00 |
| ENSG000002 | AGPAT1 | 2.91E-01 | 6.45E-01 | 9.63E-01 | -5.37E-02 | -1.04E+00 |
| ENSG000002 | TNFRSF14-A | 2.91E-01 | 6.45E-01 | 8.24E-01 | -2.79E-01 | -1.21E+00 |
| ENSG000002 | BRPF3-AS1 | 2.92E-01 | ? | 7.63E-01 | -3.90E-01 | -1.31E+00 |
| ENSG000002 | NEXN-AS1 | 2.92E-01 | ? | 1.30E+00 | 3.80E-01 | 1.30E+00 |
| ENSG000001 | SNX21 | 2.92E-01 | 6.46E-01 | 1.09E+00 | 1.18E-01 | 1.09E+00 |
| ENSG000001 | NUP35 | 2.92E-01 | 6.46E-01 | 1.05E+00 | 7.14E-02 | 1.05E+00 |
| ENSG000001 | IGFBP6 | 2.92E-01 | 6.46E-01 | 8.54E-01 | -2.28E-01 | -1.17E+00 |
| ENSG000001 | KLHL18 | 2.92E-01 | 6.46E-01 | 1.04E+00 | 5.14E-02 | 1.04E+00 |
| ENSG000001 | PALS2 | 2.92E-01 | 6.46E-01 | 9.59E-01 | -6.10E-02 | -1.04E+00 |
| ENSG000002 | --- | 2.92E-01 | ? | 7.08E-01 | -4.98E-01 | -1.41E+00 |
| ENSG000001 | RBM39 | 2.92E-01 | 6.46E-01 | 9.61E-01 | -5.75E-02 | -1.04E+00 |
| ENSG000001 | PLTP | 2.92E-01 | 6.46E-01 | 1.04E+00 | 6.12E-02 | 1.04E+00 |
| ENSG000001 | PILRB | 2.92E-01 | 6.46E-01 | 9.03E-01 | -1.47E-01 | -1.11E+00 |
| ENSG000001 | PARS2 | 2.92E-01 | 6.46E-01 | 1.07E+00 | 9.74E-02 | 1.07E+00 |
| ENSG00000C | GYG2 | 2.92E-01 | ? | 1.70E+00 | 7.67E-01 | 1.70E+00 |
| ENSG000001 | PSMD14 | 2.92E-01 | 6.46E-01 | 9.76E-01 | -3.55E-02 | -1.02E+00 |
| ENSG000002 | --- | 2.92E-01 | ? | 1.37E+00 | 4.56E-01 | 1.37E+00 |
| ENSG000002 | RPS27AP16 | 2.92E-01 | 6.46E-01 | 1.11E+00 | 1.50E-01 | 1.11E+00 |

| | | | | | | |
|---|---|---|---|---|---|---|
| ENSG000001 | ABHD3 | 2.92E-01 | 6.46E-01 | 9.42E-01 | -8.64E-02 | -1.06E+00 |
| ENSG000001 | POLR1C | 2.93E-01 | 6.47E-01 | 1.04E+00 | 6.26E-02 | 1.04E+00 |
| ENSG000002 | --- | 2.93E-01 | ? | 7.22E-01 | -4.71E-01 | -1.39E+00 |
| ENSG000002 | AKR1C7P | 2.93E-01 | ? | 6.26E-01 | -6.75E-01 | -1.60E+00 |
| ENSG000001 | FLVCR1 | 2.93E-01 | 6.47E-01 | 9.44E-01 | -8.35E-02 | -1.06E+00 |
| ENSG000001 | VAMP1 | 2.93E-01 | 6.47E-01 | 8.96E-01 | -1.58E-01 | -1.12E+00 |
| ENSG000002 | GTF2I-AS1 | 2.93E-01 | ? | 7.21E-01 | -4.71E-01 | -1.39E+00 |
| ENSG000002 | COL15A1 | 2.93E-01 | ? | 6.85E-01 | -5.46E-01 | -1.46E+00 |
| ENSG00000C | NLE1 | 2.93E-01 | 6.47E-01 | 1.06E+00 | 7.89E-02 | 1.06E+00 |
| ENSG000001 | SPTBN1 | 2.93E-01 | 6.47E-01 | 9.68E-01 | -4.69E-02 | -1.03E+00 |
| ENSG000001 | LMNA | 2.93E-01 | 6.47E-01 | 1.05E+00 | 7.26E-02 | 1.05E+00 |
| ENSG000001 | FAM210B | 2.93E-01 | 6.47E-01 | 1.04E+00 | 5.42E-02 | 1.04E+00 |
| ENSG000001 | CDC42EP4 | 2.93E-01 | 6.47E-01 | 1.06E+00 | 8.04E-02 | 1.06E+00 |
| ENSG000002 | --- | 2.93E-01 | ? | 8.17E-01 | -2.92E-01 | -1.22E+00 |
| ENSG000002 | --- | 2.93E-01 | ? | 8.23E-01 | -2.81E-01 | -1.21E+00 |
| ENSG000001 | FBXW8 | 2.94E-01 | 6.48E-01 | 1.05E+00 | 6.51E-02 | 1.05E+00 |
| ENSG000002 | --- | 2.94E-01 | ? | 6.94E-01 | -5.28E-01 | -1.44E+00 |
| ENSG000001 | MSH3 | 2.94E-01 | 6.48E-01 | 9.52E-01 | -7.11E-02 | -1.05E+00 |
| ENSG000002 | --- | 2.94E-01 | 6.48E-01 | 1.16E+00 | 2.13E-01 | 1.16E+00 |
| ENSG000002 | --- | 2.94E-01 | ? | 1.25E+00 | 3.25E-01 | 1.25E+00 |
| ENSG00000C | EPN2 | 2.94E-01 | 6.48E-01 | 1.04E+00 | 5.08E-02 | 1.04E+00 |
| ENSG000002 | MCTS1 | 2.94E-01 | 6.48E-01 | 9.70E-01 | -4.43E-02 | -1.03E+00 |
| ENSG000001 | RAB20 | 2.94E-01 | 6.48E-01 | 1.10E+00 | 1.39E-01 | 1.10E+00 |
| ENSG000001 | WNK4 | 2.94E-01 | 6.48E-01 | 1.08E+00 | 1.10E-01 | 1.08E+00 |
| ENSG000002 | --- | 2.94E-01 | 6.48E-01 | 8.49E-01 | -2.35E-01 | -1.18E+00 |
| ENSG000002 | RDM1 | 2.94E-01 | ? | 1.56E+00 | 6.37E-01 | 1.56E+00 |
| ENSG000002 | TRIM7-AS1 | 2.94E-01 | ? | 6.89E-01 | -5.37E-01 | -1.45E+00 |
| ENSG000002 | --- | 2.94E-01 | ? | 6.13E-01 | -7.05E-01 | -1.63E+00 |
| ENSG000002 | MKLN1-AS | 2.94E-01 | 6.48E-01 | 1.08E+00 | 1.08E-01 | 1.08E+00 |
| ENSG000001 | ENDOG | 2.94E-01 | 6.49E-01 | 1.10E+00 | 1.42E-01 | 1.10E+00 |
| ENSG000002 | PCDHA7 | 2.94E-01 | ? | 7.14E-01 | -4.87E-01 | -1.40E+00 |
| ENSG000001 | KCNK12 | 2.94E-01 | ? | 6.70E-01 | -5.77E-01 | -1.49E+00 |
| ENSG000001 | GABRA5 | 2.94E-01 | ? | 1.31E+00 | 3.92E-01 | 1.31E+00 |
| ENSG000001 | TUBGCP6 | 2.94E-01 | 6.49E-01 | 1.07E+00 | 9.67E-02 | 1.07E+00 |
| ENSG000002 | --- | 2.94E-01 | ? | 6.78E-01 | -5.61E-01 | -1.48E+00 |
| ENSG000001 | CENPJ | 2.95E-01 | 6.49E-01 | 9.26E-01 | -1.11E-01 | -1.08E+00 |
| ENSG000001 | CLIC5 | 2.95E-01 | ? | 1.26E+00 | 3.39E-01 | 1.26E+00 |
| ENSG000001 | UVRAG | 2.95E-01 | 6.49E-01 | 9.54E-01 | -6.84E-02 | -1.05E+00 |
| ENSG000001 | TFDP2 | 2.95E-01 | 6.49E-01 | 1.04E+00 | 6.14E-02 | 1.04E+00 |
| ENSG000001 | RHOB | 2.95E-01 | 6.49E-01 | 9.56E-01 | -6.49E-02 | -1.05E+00 |
| ENSG000001 | DHFRP1 | 2.95E-01 | ? | 1.52E+00 | 6.06E-01 | 1.52E+00 |
| ENSG000002 | --- | 2.95E-01 | ? | 8.17E-01 | -2.92E-01 | -1.22E+00 |
| ENSG000002 | LINC01358 | 2.95E-01 | ? | 7.09E-01 | -4.96E-01 | -1.41E+00 |
| ENSG00000C | SS18L2 | 2.95E-01 | 6.50E-01 | 1.06E+00 | 8.75E-02 | 1.06E+00 |
| ENSG000001 | MT-ATP6 | 2.95E-01 | 6.50E-01 | 8.93E-01 | -1.63E-01 | -1.12E+00 |

| | | | | | | |
|---|---|---|---|---|---|---|
| ENSG000001 | MORN1 | 2.95E-01 | 6.50E-01 | 1.11E+00 | 1.53E-01 | 1.11E+00 |
| ENSG000002 | --- | 2.95E-01 | ? | 1.50E+00 | 5.83E-01 | 1.50E+00 |
| ENSG000002 | ENTPD1-AS1 | 2.95E-01 | 6.50E-01 | 9.44E-01 | -8.33E-02 | -1.06E+00 |
| ENSG00000C | AAMDC | 2.95E-01 | 6.50E-01 | 9.37E-01 | -9.45E-02 | -1.07E+00 |
| ENSG000002 | --- | 2.95E-01 | ? | 6.09E-01 | -7.15E-01 | -1.64E+00 |
| ENSG000002 | SH3D21 | 2.96E-01 | 6.50E-01 | 9.27E-01 | -1.10E-01 | -1.08E+00 |
| ENSG000001 | IPP | 2.96E-01 | 6.50E-01 | 1.04E+00 | 5.96E-02 | 1.04E+00 |
| ENSG000001 | CDK1 | 2.96E-01 | 6.50E-01 | 8.77E-01 | -1.90E-01 | -1.14E+00 |
| ENSG000002 | --- | 2.96E-01 | ? | 1.39E+00 | 4.78E-01 | 1.39E+00 |
| ENSG00000C | ARCN1 | 2.96E-01 | 6.50E-01 | 9.71E-01 | -4.21E-02 | -1.03E+00 |
| ENSG00000C | PCSK5 | 2.96E-01 | 6.50E-01 | 9.35E-01 | -9.76E-02 | -1.07E+00 |
| ENSG000001 | MCEE | 2.96E-01 | 6.50E-01 | 1.15E+00 | 2.04E-01 | 1.15E+00 |
| ENSG000001 | CENPV | 2.96E-01 | 6.50E-01 | 1.12E+00 | 1.67E-01 | 1.12E+00 |
| ENSG000001 | OMA1 | 2.96E-01 | 6.50E-01 | 1.05E+00 | 6.72E-02 | 1.05E+00 |
| ENSG000002 | --- | 2.96E-01 | ? | 1.18E+00 | 2.39E-01 | 1.18E+00 |
| ENSG00000C | ARHGEF10L | 2.96E-01 | 6.51E-01 | 1.07E+00 | 9.77E-02 | 1.07E+00 |
| ENSG00000C | NUFIP1 | 2.96E-01 | 6.51E-01 | 9.49E-01 | -7.56E-02 | -1.05E+00 |
| ENSG000001 | KLF8 | 2.96E-01 | 6.51E-01 | 9.39E-01 | -9.06E-02 | -1.06E+00 |
| ENSG000001 | IL23A | 2.96E-01 | ? | 1.19E+00 | 2.50E-01 | 1.19E+00 |
| ENSG000001 | NUF2 | 2.96E-01 | 6.51E-01 | 8.67E-01 | -2.06E-01 | -1.15E+00 |
| ENSG000001 | UBE2J1 | 2.96E-01 | 6.51E-01 | 9.71E-01 | -4.20E-02 | -1.03E+00 |
| ENSG000002 | --- | 2.97E-01 | 6.51E-01 | 1.12E+00 | 1.69E-01 | 1.12E+00 |
| ENSG000001 | AMPD3 | 2.97E-01 | 6.51E-01 | 1.04E+00 | 5.51E-02 | 1.04E+00 |
| ENSG000002 | PICART1 | 2.97E-01 | 6.51E-01 | 9.21E-01 | -1.19E-01 | -1.09E+00 |
| ENSG000002 | --- | 2.97E-01 | 6.51E-01 | 9.21E-01 | -1.18E-01 | -1.09E+00 |
| ENSG000001 | KIAA0513 | 2.97E-01 | 6.51E-01 | 1.07E+00 | 9.61E-02 | 1.07E+00 |
| ENSG000002 | MRPL12 | 2.97E-01 | 6.51E-01 | 1.08E+00 | 1.08E-01 | 1.08E+00 |
| ENSG000001 | PRPF38A | 2.97E-01 | 6.51E-01 | 9.60E-01 | -5.84E-02 | -1.04E+00 |
| ENSG000001 | SOX6 | 2.97E-01 | 6.51E-01 | 1.08E+00 | 1.17E-01 | 1.08E+00 |
| ENSG000002 | --- | 2.97E-01 | ? | 1.27E+00 | 3.42E-01 | 1.27E+00 |
| ENSG000001 | THAP6 | 2.97E-01 | 6.51E-01 | 9.58E-01 | -6.23E-02 | -1.04E+00 |
| ENSG000002 | --- | 2.97E-01 | ? | 1.64E+00 | 7.11E-01 | 1.64E+00 |
| ENSG000001 | --- | 2.97E-01 | ? | 1.60E+00 | 6.75E-01 | 1.60E+00 |
| ENSG000001 | AASDH | 2.97E-01 | 6.52E-01 | 1.05E+00 | 7.25E-02 | 1.05E+00 |
| ENSG000002 | RPSAP12 | 2.97E-01 | ? | 1.33E+00 | 4.15E-01 | 1.33E+00 |
| ENSG000001 | ZNF85 | 2.97E-01 | 6.52E-01 | 9.33E-01 | -1.01E-01 | -1.07E+00 |
| ENSG000002 | PRRC2B | 2.97E-01 | 6.52E-01 | 1.03E+00 | 4.60E-02 | 1.03E+00 |
| ENSG000001 | KAZN | 2.97E-01 | 6.52E-01 | 1.04E+00 | 6.14E-02 | 1.04E+00 |
| ENSG000002 | LINC01554 | 2.97E-01 | ? | 5.72E-01 | -8.06E-01 | -1.75E+00 |
| ENSG000002 | --- | 2.97E-01 | 6.52E-01 | 9.08E-01 | -1.40E-01 | -1.10E+00 |
| ENSG000001 | MPDU1 | 2.98E-01 | 6.52E-01 | 1.04E+00 | 5.74E-02 | 1.04E+00 |
| ENSG000002 | --- | 2.98E-01 | ? | 1.56E+00 | 6.38E-01 | 1.56E+00 |
| ENSG000001 | CEND1 | 2.98E-01 | ? | 6.85E-01 | -5.46E-01 | -1.46E+00 |
| ENSG000001 | MORN2 | 2.98E-01 | 6.52E-01 | 1.07E+00 | 1.02E-01 | 1.07E+00 |
| ENSG000002 | SLC25A53 | 2.98E-01 | 6.52E-01 | 1.12E+00 | 1.63E-01 | 1.12E+00 |

| | | | | | | |
|---|---|---|---|---|---|---|
| ENSG000001 | FXR2 | 2.98E-01 | 6.52E-01 | 1.03E+00 | 4.37E-02 | 1.03E+00 |
| ENSG000001 | ANKRD37 | 2.98E-01 | 6.52E-01 | 9.00E-01 | -1.52E-01 | -1.11E+00 |
| ENSG000002 | SZT2-AS1 | 2.98E-01 | ? | 7.77E-01 | -3.63E-01 | -1.29E+00 |
| ENSG000001 | IFT140 | 2.98E-01 | 6.52E-01 | 1.07E+00 | 9.62E-02 | 1.07E+00 |
| ENSG000002 | TMEM44-AS | 2.98E-01 | 6.52E-01 | 8.66E-01 | -2.07E-01 | -1.15E+00 |
| ENSG000002 | SLC6A14 | 2.98E-01 | 6.52E-01 | 8.57E-01 | -2.23E-01 | -1.17E+00 |
| ENSG000002 | --- | 2.98E-01 | 6.52E-01 | 1.19E+00 | 2.55E-01 | 1.19E+00 |
| ENSG000001 | PNP | 2.98E-01 | 6.52E-01 | 1.04E+00 | 5.61E-02 | 1.04E+00 |
| ENSG000002 | PACERR | 2.98E-01 | 6.52E-01 | 8.80E-01 | -1.84E-01 | -1.14E+00 |
| ENSG000002 | --- | 2.98E-01 | 6.52E-01 | 1.11E+00 | 1.50E-01 | 1.11E+00 |
| ENSG000001 | IL22RA1 | 2.98E-01 | 6.52E-01 | 1.23E+00 | 2.94E-01 | 1.23E+00 |
| ENSG000001 | TCTN2 | 2.98E-01 | 6.52E-01 | 9.33E-01 | -9.96E-02 | -1.07E+00 |
| ENSG000002 | RAB33B-AS1 | 2.98E-01 | 6.52E-01 | 8.75E-01 | -1.93E-01 | -1.14E+00 |
| ENSG000001 | JOSD2 | 2.98E-01 | 6.52E-01 | 1.14E+00 | 1.91E-01 | 1.14E+00 |
| ENSG000001 | STING1 | 2.98E-01 | 6.52E-01 | 1.05E+00 | 6.58E-02 | 1.05E+00 |
| ENSG00000C | ZNF263 | 2.99E-01 | 6.52E-01 | 9.66E-01 | -4.95E-02 | -1.03E+00 |
| ENSG000001 | MACIR | 2.99E-01 | 6.52E-01 | 9.56E-01 | -6.52E-02 | -1.05E+00 |
| ENSG00000C | BLNK | 2.99E-01 | 6.53E-01 | 1.10E+00 | 1.40E-01 | 1.10E+00 |
| ENSG000002 | --- | 2.99E-01 | ? | 1.51E+00 | 5.91E-01 | 1.51E+00 |
| ENSG000001 | IGFL1 | 2.99E-01 | ? | 1.23E+00 | 3.04E-01 | 1.23E+00 |
| ENSG000002 | CIRBP-AS1 | 2.99E-01 | 6.53E-01 | 8.72E-01 | -1.97E-01 | -1.15E+00 |
| ENSG000002 | --- | 2.99E-01 | ? | 1.45E+00 | 5.37E-01 | 1.45E+00 |
| ENSG000001 | DGCR6 | 2.99E-01 | 6.53E-01 | 1.10E+00 | 1.42E-01 | 1.10E+00 |
| ENSG000002 | HOMEZ | 2.99E-01 | 6.53E-01 | 1.04E+00 | 5.50E-02 | 1.04E+00 |
| ENSG000001 | CTNND1 | 2.99E-01 | 6.53E-01 | 1.02E+00 | 3.19E-02 | 1.02E+00 |
| ENSG000001 | DLG2 | 2.99E-01 | ? | 1.32E+00 | 4.02E-01 | 1.32E+00 |
| ENSG000002 | USP38-DT | 2.99E-01 | ? | 6.77E-01 | -5.64E-01 | -1.48E+00 |
| ENSG000001 | OLA1 | 2.99E-01 | 6.54E-01 | 9.58E-01 | -6.14E-02 | -1.04E+00 |
| ENSG000001 | ZNF559 | 2.99E-01 | 6.54E-01 | 1.06E+00 | 8.79E-02 | 1.06E+00 |
| ENSG000002 | LINC00562 | 2.99E-01 | ? | 7.87E-01 | -3.45E-01 | -1.27E+00 |
| ENSG000002 | EMBP1 | 3.00E-01 | ? | 1.21E+00 | 2.69E-01 | 1.21E+00 |
| ENSG000001 | RAMAC | 3.00E-01 | 6.54E-01 | 1.03E+00 | 4.79E-02 | 1.03E+00 |
| ENSG000001 | RAB6B | 3.00E-01 | 6.54E-01 | 1.07E+00 | 1.00E-01 | 1.07E+00 |
| ENSG000002 | --- | 3.00E-01 | ? | 1.60E+00 | 6.80E-01 | 1.60E+00 |
| ENSG00000C | DIP2B | 3.00E-01 | 6.55E-01 | 1.04E+00 | 5.08E-02 | 1.04E+00 |
| ENSG000001 | CDH26 | 3.00E-01 | 6.55E-01 | 8.86E-01 | -1.75E-01 | -1.13E+00 |
| ENSG000002 | --- | 3.00E-01 | ? | 6.66E-01 | -5.85E-01 | -1.50E+00 |
| ENSG000002 | --- | 3.00E-01 | 6.55E-01 | 9.18E-01 | -1.23E-01 | -1.09E+00 |
| ENSG000001 | RPL36AL | 3.00E-01 | 6.55E-01 | 1.04E+00 | 5.62E-02 | 1.04E+00 |
| ENSG00000C | PAFAH1B1 | 3.00E-01 | 6.55E-01 | 9.81E-01 | -2.70E-02 | -1.02E+00 |
| ENSG000002 | THEMIS3P | 3.00E-01 | ? | 1.37E+00 | 4.50E-01 | 1.37E+00 |
| ENSG000002 | CSPG4P5 | 3.00E-01 | ? | 7.13E-01 | -4.89E-01 | -1.40E+00 |
| ENSG000002 | --- | 3.00E-01 | ? | 1.31E+00 | 3.86E-01 | 1.31E+00 |
| ENSG000001 | GAA | 3.01E-01 | 6.55E-01 | 1.08E+00 | 1.16E-01 | 1.08E+00 |
| ENSG000002 | --- | 3.01E-01 | ? | 7.23E-01 | -4.67E-01 | -1.38E+00 |

| | | | | | | |
|---|---|---|---|---|---|---|
| ENSG000001 | CDH4 | 3.01E-01 | 6.55E-01 | 9.01E-01 | -1.51E-01 | -1.11E+00 |
| ENSG000001 | INTS7 | 3.01E-01 | 6.55E-01 | 9.60E-01 | -5.89E-02 | -1.04E+00 |
| ENSG000002 | --- | 3.01E-01 | ? | 1.45E+00 | 5.33E-01 | 1.45E+00 |
| ENSG000001 | PIGT | 3.01E-01 | 6.55E-01 | 9.63E-01 | -5.46E-02 | -1.04E+00 |
| ENSG000001 | UBXN4 | 3.01E-01 | 6.55E-01 | 9.73E-01 | -3.97E-02 | -1.03E+00 |
| ENSG000001 | METTL18 | 3.01E-01 | 6.55E-01 | 1.09E+00 | 1.24E-01 | 1.09E+00 |
| ENSG000001 | KLC1 | 3.01E-01 | 6.55E-01 | 9.68E-01 | -4.68E-02 | -1.03E+00 |
| ENSG000001 | KLK7 | 3.01E-01 | 6.55E-01 | 8.01E-01 | -3.20E-01 | -1.25E+00 |
| ENSG00000C | ZBTB25 | 3.01E-01 | 6.56E-01 | 1.07E+00 | 9.52E-02 | 1.07E+00 |
| ENSG00000C | PDK2 | 3.01E-01 | 6.56E-01 | 9.29E-01 | -1.06E-01 | -1.08E+00 |
| ENSG000002 | CDRT4 | 3.01E-01 | ? | 1.98E+00 | 9.82E-01 | 1.98E+00 |
| ENSG000002 | MUC20P1 | 3.01E-01 | ? | 7.78E-01 | -3.62E-01 | -1.28E+00 |
| ENSG000002 | --- | 3.01E-01 | ? | 7.19E-01 | -4.77E-01 | -1.39E+00 |
| ENSG000001 | ZNF420 | 3.01E-01 | 6.56E-01 | 9.34E-01 | -9.92E-02 | -1.07E+00 |
| ENSG00000C | SPTB | 3.01E-01 | 6.56E-01 | 1.17E+00 | 2.21E-01 | 1.17E+00 |
| ENSG00000C | TRMT11 | 3.01E-01 | 6.56E-01 | 1.05E+00 | 7.59E-02 | 1.05E+00 |
| ENSG000001 | PAICS | 3.01E-01 | 6.56E-01 | 1.04E+00 | 5.96E-02 | 1.04E+00 |
| ENSG000001 | GOT1 | 3.01E-01 | 6.56E-01 | 1.04E+00 | 6.08E-02 | 1.04E+00 |
| ENSG000001 | AUTS2 | 3.01E-01 | 6.56E-01 | 9.61E-01 | -5.79E-02 | -1.04E+00 |
| ENSG000001 | DOCK2 | 3.02E-01 | ? | 7.85E-01 | -3.49E-01 | -1.27E+00 |
| ENSG000001 | A2ML1 | 3.02E-01 | 6.56E-01 | 1.09E+00 | 1.26E-01 | 1.09E+00 |
| ENSG000001 | DACT2 | 3.02E-01 | ? | 1.39E+00 | 4.77E-01 | 1.39E+00 |
| ENSG000002 | RPL6P10 | 3.02E-01 | 6.56E-01 | 1.07E+00 | 1.02E-01 | 1.07E+00 |
| ENSG000002 | --- | 3.02E-01 | ? | 8.23E-01 | -2.82E-01 | -1.22E+00 |
| ENSG000001 | PLLP | 3.02E-01 | 6.56E-01 | 1.06E+00 | 9.05E-02 | 1.06E+00 |
| ENSG000002 | --- | 3.02E-01 | ? | 8.12E-01 | -3.00E-01 | -1.23E+00 |
| ENSG000001 | ZNF146 | 3.02E-01 | 6.56E-01 | 9.67E-01 | -4.87E-02 | -1.03E+00 |
| ENSG000002 | --- | 3.02E-01 | 6.56E-01 | 8.83E-01 | -1.80E-01 | -1.13E+00 |
| ENSG000002 | PARD6G-AS1 | 3.02E-01 | 6.57E-01 | 1.17E+00 | 2.29E-01 | 1.17E+00 |
| ENSG000001 | IRS4 | 3.02E-01 | ? | 1.72E+00 | 7.80E-01 | 1.72E+00 |
| ENSG00000C | GOLGA5 | 3.02E-01 | 6.57E-01 | 9.52E-01 | -7.06E-02 | -1.05E+00 |
| ENSG000001 | OTUD7A | 3.02E-01 | ? | 1.34E+00 | 4.22E-01 | 1.34E+00 |
| ENSG000001 | NENF | 3.02E-01 | 6.57E-01 | 1.08E+00 | 1.09E-01 | 1.08E+00 |
| ENSG000001 | SBDS | 3.02E-01 | 6.57E-01 | 9.65E-01 | -5.19E-02 | -1.04E+00 |
| ENSG000002 | HLA-F-AS1 | 3.02E-01 | 6.57E-01 | 9.06E-01 | -1.43E-01 | -1.10E+00 |
| ENSG000002 | TOP1P1 | 3.03E-01 | ? | 7.02E-01 | -5.11E-01 | -1.42E+00 |
| ENSG000002 | --- | 3.03E-01 | ? | 1.56E+00 | 6.39E-01 | 1.56E+00 |
| ENSG000001 | DNAI4 | 3.03E-01 | 6.57E-01 | 1.09E+00 | 1.22E-01 | 1.09E+00 |
| ENSG000001 | SOCS3 | 3.03E-01 | 6.57E-01 | 8.94E-01 | -1.62E-01 | -1.12E+00 |
| ENSG000001 | BHLHA15 | 3.03E-01 | ? | 6.92E-01 | -5.30E-01 | -1.44E+00 |
| ENSG000001 | CD151 | 3.03E-01 | 6.57E-01 | 1.08E+00 | 1.04E-01 | 1.08E+00 |
| ENSG000002 | --- | 3.03E-01 | ? | 1.58E+00 | 6.58E-01 | 1.58E+00 |
| ENSG000001 | SERINC5 | 3.03E-01 | 6.58E-01 | 1.02E+00 | 3.28E-02 | 1.02E+00 |
| ENSG000002 | --- | 3.03E-01 | ? | 6.73E-01 | -5.71E-01 | -1.49E+00 |
| ENSG000002 | CCDC85C | 3.03E-01 | 6.58E-01 | 1.05E+00 | 7.37E-02 | 1.05E+00 |

| | | | | | | |
|---|---|---|---|---|---|---|
| ENSG000001 | TSPAN33 | 3.03E-01 | 6.58E-01 | 1.10E+00 | 1.43E-01 | 1.10E+00 |
| ENSG000002 | ST20-AS1 | 3.03E-01 | 6.58E-01 | 1.07E+00 | 1.03E-01 | 1.07E+00 |
| ENSG000001 | C10orf88 | 3.03E-01 | 6.58E-01 | 9.53E-01 | -6.89E-02 | -1.05E+00 |
| ENSG000002 | LIN52 | 3.03E-01 | 6.58E-01 | 1.07E+00 | 1.02E-01 | 1.07E+00 |
| ENSG000001 | ILKAP | 3.03E-01 | 6.58E-01 | 1.04E+00 | 5.48E-02 | 1.04E+00 |
| ENSG000002 | CFAP298-TCF | 3.03E-01 | 6.58E-01 | 8.50E-01 | -2.34E-01 | -1.18E+00 |
| ENSG000002 | ZNF559-ZNF | 3.03E-01 | ? | 7.04E-01 | -5.06E-01 | -1.42E+00 |
| ENSG000001 | EFHD2 | 3.03E-01 | 6.58E-01 | 1.04E+00 | 5.59E-02 | 1.04E+00 |
| ENSG000002 | BMPR2 | 3.03E-01 | 6.58E-01 | 9.54E-01 | -6.87E-02 | -1.05E+00 |
| ENSG000001 | F12 | 3.03E-01 | 6.58E-01 | 1.15E+00 | 1.96E-01 | 1.15E+00 |
| ENSG000001 | CCDC87 | 3.04E-01 | ? | 1.29E+00 | 3.67E-01 | 1.29E+00 |
| ENSG000002 | --- | 3.04E-01 | ? | 1.39E+00 | 4.79E-01 | 1.39E+00 |
| ENSG000001 | TOX3 | 3.04E-01 | ? | 7.30E-01 | -4.55E-01 | -1.37E+00 |
| ENSG000001 | UCK1 | 3.04E-01 | 6.58E-01 | 1.06E+00 | 8.71E-02 | 1.06E+00 |
| ENSG000002 | DDX12P | 3.04E-01 | ? | 8.15E-01 | -2.96E-01 | -1.23E+00 |
| ENSG000002 | --- | 3.04E-01 | ? | 1.33E+00 | 4.16E-01 | 1.33E+00 |
| ENSG000002 | JARID2-DT | 3.04E-01 | ? | 6.75E-01 | -5.67E-01 | -1.48E+00 |
| ENSG000001 | ANXA2R-OT1 | 3.04E-01 | 6.58E-01 | 9.23E-01 | -1.16E-01 | -1.08E+00 |
| ENSG000000 | AK2 | 3.04E-01 | 6.58E-01 | 1.03E+00 | 4.14E-02 | 1.03E+00 |
| ENSG000002 | --- | 3.04E-01 | 6.59E-01 | 9.00E-01 | -1.51E-01 | -1.11E+00 |
| ENSG000001 | ZNF106 | 3.04E-01 | 6.59E-01 | 9.53E-01 | -6.99E-02 | -1.05E+00 |
| ENSG000002 | --- | 3.04E-01 | 6.59E-01 | 1.17E+00 | 2.22E-01 | 1.17E+00 |
| ENSG000001 | HSFX1 | 3.04E-01 | ? | 3.68E+00 | 1.88E+00 | 3.68E+00 |
| ENSG000001 | C11orf65 | 3.04E-01 | ? | 8.38E-01 | -2.55E-01 | -1.19E+00 |
| ENSG000002 | OR5BA1P | 3.04E-01 | ? | 1.35E+00 | 4.31E-01 | 1.35E+00 |
| ENSG000001 | ZNF251 | 3.05E-01 | 6.59E-01 | 1.07E+00 | 9.32E-02 | 1.07E+00 |
| ENSG000001 | IL1RL2 | 3.05E-01 | 6.59E-01 | 9.08E-01 | -1.39E-01 | -1.10E+00 |
| ENSG000001 | S100A12 | 3.05E-01 | 6.59E-01 | 1.19E+00 | 2.48E-01 | 1.19E+00 |
| ENSG000002 | --- | 3.05E-01 | ? | 1.33E+00 | 4.08E-01 | 1.33E+00 |
| ENSG000002 | ZNF878 | 3.05E-01 | ? | 7.43E-01 | -4.28E-01 | -1.34E+00 |
| ENSG000001 | NSMF | 3.05E-01 | 6.59E-01 | 1.05E+00 | 6.39E-02 | 1.05E+00 |
| ENSG000002 | --- | 3.05E-01 | ? | 5.75E-01 | -8.00E-01 | -1.74E+00 |
| ENSG000002 | --- | 3.05E-01 | ? | 6.14E-01 | -7.04E-01 | -1.63E+00 |
| ENSG000002 | RPL21P93 | 3.05E-01 | ? | 1.44E+00 | 5.27E-01 | 1.44E+00 |
| ENSG000002 | MIR646HG | 3.05E-01 | 6.59E-01 | 8.59E-01 | -2.19E-01 | -1.16E+00 |
| ENSG000001 | BUB3 | 3.05E-01 | 6.59E-01 | 1.02E+00 | 3.39E-02 | 1.02E+00 |
| ENSG000001 | POLE4 | 3.05E-01 | 6.59E-01 | 1.08E+00 | 1.06E-01 | 1.08E+00 |
| ENSG000001 | SPATA5L1 | 3.05E-01 | 6.59E-01 | 9.57E-01 | -6.32E-02 | -1.04E+00 |
| ENSG000001 | INPPL1 | 3.05E-01 | 6.59E-01 | 1.06E+00 | 7.74E-02 | 1.06E+00 |
| ENSG000002 | CSNK2B | 3.05E-01 | 6.59E-01 | 1.02E+00 | 3.27E-02 | 1.02E+00 |
| ENSG000001 | PHF12 | 3.05E-01 | 6.59E-01 | 9.63E-01 | -5.39E-02 | -1.04E+00 |
| ENSG000002 | --- | 3.05E-01 | 6.59E-01 | 9.20E-01 | -1.20E-01 | -1.09E+00 |
| ENSG000001 | HIBADH | 3.05E-01 | 6.59E-01 | 1.04E+00 | 6.31E-02 | 1.04E+00 |
| ENSG000002 | LINC00460 | 3.05E-01 | ? | 7.95E-01 | -3.32E-01 | -1.26E+00 |
| ENSG000001 | ZBTB7A | 3.05E-01 | 6.59E-01 | 9.69E-01 | -4.60E-02 | -1.03E+00 |

| | | | | | | |
|---|---|---|---|---|---|---|
| ENSG000001 | PGGHG | 3.05E-01 | 6.59E-01 | 1.09E+00 | 1.26E-01 | 1.09E+00 |
| ENSG000001 | UBE2B | 3.05E-01 | 6.59E-01 | 1.04E+00 | 5.07E-02 | 1.04E+00 |
| ENSG000001 | ITPR1 | 3.05E-01 | 6.60E-01 | 9.47E-01 | -7.82E-02 | -1.06E+00 |
| ENSG000002 | --- | 3.05E-01 | ? | 1.23E+00 | 3.02E-01 | 1.23E+00 |
| ENSG000001 | ZNF347 | 3.06E-01 | 6.60E-01 | 1.07E+00 | 9.12E-02 | 1.07E+00 |
| ENSG000001 | PMEPA1 | 3.06E-01 | 6.60E-01 | 9.56E-01 | -6.47E-02 | -1.05E+00 |
| ENSG000001 | NCBP2 | 3.06E-01 | 6.60E-01 | 1.03E+00 | 4.31E-02 | 1.03E+00 |
| ENSG000001 | FBXL7 | 3.06E-01 | 6.60E-01 | 9.41E-01 | -8.77E-02 | -1.06E+00 |
| ENSG000002 | --- | 3.06E-01 | ? | 8.46E-01 | -2.41E-01 | -1.18E+00 |
| ENSG000001 | KLHL2 | 3.06E-01 | 6.60E-01 | 1.04E+00 | 5.97E-02 | 1.04E+00 |
| ENSG000001 | ROBO2 | 3.06E-01 | 6.60E-01 | 8.96E-01 | -1.58E-01 | -1.12E+00 |
| ENSG000002 | --- | 3.06E-01 | 6.60E-01 | 1.09E+00 | 1.18E-01 | 1.09E+00 |
| ENSG00000C | AFP | 3.06E-01 | ? | 1.31E+00 | 3.93E-01 | 1.31E+00 |
| ENSG000001 | ZBTB7B | 3.06E-01 | 6.60E-01 | 1.06E+00 | 8.84E-02 | 1.06E+00 |
| ENSG000002 | --- | 3.06E-01 | 6.60E-01 | 1.17E+00 | 2.23E-01 | 1.17E+00 |
| ENSG000001 | FAM149A | 3.06E-01 | ? | 1.21E+00 | 2.77E-01 | 1.21E+00 |
| ENSG000001 | MFSD12 | 3.06E-01 | 6.60E-01 | 1.06E+00 | 8.77E-02 | 1.06E+00 |
| ENSG000001 | ANKRD7 | 3.06E-01 | ? | 7.29E-01 | -4.57E-01 | -1.37E+00 |
| ENSG00000C | DDX20 | 3.06E-01 | 6.60E-01 | 1.03E+00 | 4.40E-02 | 1.03E+00 |
| ENSG000002 | TBC1D3L | 3.06E-01 | 6.60E-01 | 8.68E-01 | -2.05E-01 | -1.15E+00 |
| ENSG00000C | TSPAN15 | 3.06E-01 | 6.60E-01 | 1.07E+00 | 9.78E-02 | 1.07E+00 |
| ENSG000002 | --- | 3.06E-01 | 6.60E-01 | 1.19E+00 | 2.47E-01 | 1.19E+00 |
| ENSG000001 | DHDDS | 3.06E-01 | 6.60E-01 | 9.73E-01 | -3.95E-02 | -1.03E+00 |
| ENSG000001 | MTERF2 | 3.06E-01 | 6.60E-01 | 9.35E-01 | -9.65E-02 | -1.07E+00 |
| ENSG000002 | --- | 3.06E-01 | 6.60E-01 | 9.04E-01 | -1.46E-01 | -1.11E+00 |
| ENSG000001 | ATF7 | 3.07E-01 | 6.60E-01 | 9.63E-01 | -5.47E-02 | -1.04E+00 |
| ENSG000001 | TOMM70 | 3.07E-01 | 6.60E-01 | 1.03E+00 | 3.93E-02 | 1.03E+00 |
| ENSG000001 | PPP3CA | 3.07E-01 | 6.60E-01 | 9.65E-01 | -5.17E-02 | -1.04E+00 |
| ENSG000001 | NOXRED1 | 3.07E-01 | ? | 8.27E-01 | -2.74E-01 | -1.21E+00 |
| ENSG000002 | --- | 3.07E-01 | ? | 1.59E+00 | 6.66E-01 | 1.59E+00 |
| ENSG000002 | --- | 3.07E-01 | ? | 7.60E-01 | -3.96E-01 | -1.32E+00 |
| ENSG000001 | GSTO1 | 3.07E-01 | 6.61E-01 | 1.05E+00 | 7.57E-02 | 1.05E+00 |
| ENSG000001 | PIFO | 3.07E-01 | 6.61E-01 | 1.14E+00 | 1.87E-01 | 1.14E+00 |
| ENSG000002 | --- | 3.07E-01 | 6.61E-01 | 1.23E+00 | 3.00E-01 | 1.23E+00 |
| ENSG00000C | NOX4 | 3.07E-01 | ? | 8.14E-01 | -2.96E-01 | -1.23E+00 |
| ENSG000001 | MFNG | 3.07E-01 | 6.61E-01 | 1.11E+00 | 1.51E-01 | 1.11E+00 |
| ENSG000001 | PODNL1 | 3.07E-01 | 6.61E-01 | 9.12E-01 | -1.33E-01 | -1.10E+00 |
| ENSG000001 | KBTBD2 | 3.07E-01 | 6.61E-01 | 9.70E-01 | -4.37E-02 | -1.03E+00 |
| ENSG000001 | GK4P | 3.07E-01 | ? | 1.49E+00 | 5.78E-01 | 1.49E+00 |
| ENSG000001 | RNFT2 | 3.07E-01 | ? | 1.24E+00 | 3.16E-01 | 1.24E+00 |
| ENSG000001 | NCL | 3.07E-01 | 6.61E-01 | 9.59E-01 | -6.03E-02 | -1.04E+00 |
| ENSG000002 | --- | 3.07E-01 | ? | 7.98E-01 | -3.26E-01 | -1.25E+00 |
| ENSG00000C | ZFAND6 | 3.07E-01 | 6.61E-01 | 9.68E-01 | -4.63E-02 | -1.03E+00 |
| ENSG000002 | CAMSAP1-DT | 3.07E-01 | ? | 1.27E+00 | 3.42E-01 | 1.27E+00 |
| ENSG000002 | HYPK | 3.07E-01 | 6.61E-01 | 9.37E-01 | -9.33E-02 | -1.07E+00 |

| | | | | | | |
|---|---|---|---|---|---|---|
| ENSG000001 | ZNF287 | 3.07E-01 | 6.61E-01 | 1.07E+00 | 1.01E-01 | 1.07E+00 |
| ENSG000002 | --- | 3.08E-01 | 6.61E-01 | 8.50E-01 | -2.35E-01 | -1.18E+00 |
| ENSG000002 | --- | 3.08E-01 | 6.61E-01 | 8.97E-01 | -1.58E-01 | -1.12E+00 |
| ENSG000002 | --- | 3.08E-01 | ? | 1.33E+00 | 4.17E-01 | 1.33E+00 |
| ENSG000001 | MAN1B1 | 3.08E-01 | 6.61E-01 | 1.05E+00 | 7.15E-02 | 1.05E+00 |
| ENSG000001 | RANBP2 | 3.08E-01 | 6.61E-01 | 9.53E-01 | -6.91E-02 | -1.05E+00 |
| ENSG000001 | FRMD5 | 3.08E-01 | 6.61E-01 | 1.08E+00 | 1.17E-01 | 1.08E+00 |
| ENSG000001 | CXCL6 | 3.08E-01 | 6.61E-01 | 8.36E-01 | -2.59E-01 | -1.20E+00 |
| ENSG000001 | CASC3 | 3.08E-01 | 6.61E-01 | 9.70E-01 | -4.34E-02 | -1.03E+00 |
| ENSG000001 | METTL2B | 3.08E-01 | 6.61E-01 | 9.64E-01 | -5.30E-02 | -1.04E+00 |
| ENSG000001 | TMEM132B | 3.08E-01 | 6.61E-01 | 8.69E-01 | -2.02E-01 | -1.15E+00 |
| ENSG000002 | TAS2R15P | 3.08E-01 | ? | 1.64E+00 | 7.17E-01 | 1.64E+00 |
| ENSG000001 | ZWILCH | 3.08E-01 | 6.61E-01 | 1.04E+00 | 5.56E-02 | 1.04E+00 |
| ENSG000002 | CHD1-DT | 3.08E-01 | ? | 1.19E+00 | 2.55E-01 | 1.19E+00 |
| ENSG00000C | PALS1 | 3.08E-01 | 6.61E-01 | 9.55E-01 | -6.65E-02 | -1.05E+00 |
| ENSG000002 | NFATC2IP-AS | 3.08E-01 | 6.61E-01 | 8.93E-01 | -1.63E-01 | -1.12E+00 |
| ENSG000002 | PIN4P1 | 3.08E-01 | ? | 5.24E-01 | -9.31E-01 | -1.91E+00 |
| ENSG000001 | RIDA | 3.08E-01 | 6.61E-01 | 1.06E+00 | 8.37E-02 | 1.06E+00 |
| ENSG000001 | GLO1 | 3.08E-01 | 6.61E-01 | 1.02E+00 | 3.46E-02 | 1.02E+00 |
| ENSG000001 | TMPRSS2 | 3.08E-01 | ? | 7.82E-01 | -3.55E-01 | -1.28E+00 |
| ENSG000001 | SRC | 3.08E-01 | 6.61E-01 | 1.05E+00 | 6.96E-02 | 1.05E+00 |
| ENSG00000C | CDC34 | 3.08E-01 | 6.62E-01 | 1.05E+00 | 7.12E-02 | 1.05E+00 |
| ENSG000002 | PET117 | 3.09E-01 | 6.62E-01 | 8.85E-01 | -1.76E-01 | -1.13E+00 |
| ENSG000002 | --- | 3.09E-01 | ? | 1.47E+00 | 5.54E-01 | 1.47E+00 |
| ENSG000001 | SERPINA3 | 3.09E-01 | 6.62E-01 | 8.11E-01 | -3.02E-01 | -1.23E+00 |
| ENSG000001 | ZSWIM5 | 3.09E-01 | ? | 1.34E+00 | 4.21E-01 | 1.34E+00 |
| ENSG000001 | PARM1 | 3.09E-01 | ? | 1.21E+00 | 2.72E-01 | 1.21E+00 |
| ENSG000001 | EFS | 3.09E-01 | 6.62E-01 | 1.14E+00 | 1.92E-01 | 1.14E+00 |
| ENSG000002 | --- | 3.09E-01 | ? | 7.83E-01 | -3.53E-01 | -1.28E+00 |
| ENSG000002 | --- | 3.09E-01 | 6.62E-01 | 9.18E-01 | -1.24E-01 | -1.09E+00 |
| ENSG000002 | --- | 3.09E-01 | ? | 1.53E+00 | 6.16E-01 | 1.53E+00 |
| ENSG000001 | MUTYH | 3.09E-01 | 6.62E-01 | 1.11E+00 | 1.46E-01 | 1.11E+00 |
| ENSG000001 | NPHP3 | 3.09E-01 | 6.62E-01 | 9.55E-01 | -6.72E-02 | -1.05E+00 |
| ENSG000002 | PSENEN | 3.09E-01 | 6.62E-01 | 1.07E+00 | 9.71E-02 | 1.07E+00 |
| ENSG000001 | PAK1 | 3.09E-01 | 6.62E-01 | 1.03E+00 | 4.11E-02 | 1.03E+00 |
| ENSG000002 | LINC00240 | 3.09E-01 | 6.62E-01 | 1.13E+00 | 1.72E-01 | 1.13E+00 |
| ENSG000002 | --- | 3.09E-01 | ? | 1.33E+00 | 4.15E-01 | 1.33E+00 |
| ENSG000001 | LZTFL1 | 3.09E-01 | 6.62E-01 | 9.44E-01 | -8.27E-02 | -1.06E+00 |
| ENSG000001 | CAND2 | 3.09E-01 | ? | 1.26E+00 | 3.32E-01 | 1.26E+00 |
| ENSG000002 | SMIM6 | 3.09E-01 | ? | 6.56E-01 | -6.09E-01 | -1.53E+00 |
| ENSG000001 | CHSY1 | 3.09E-01 | 6.62E-01 | 9.48E-01 | -7.76E-02 | -1.06E+00 |
| ENSG000001 | TMPRSS9 | 3.09E-01 | ? | 1.32E+00 | 4.02E-01 | 1.32E+00 |
| ENSG000001 | DNAJB7 | 3.09E-01 | ? | 6.51E-01 | -6.18E-01 | -1.54E+00 |
| ENSG00000C | FUZ | 3.09E-01 | 6.62E-01 | 1.17E+00 | 2.26E-01 | 1.17E+00 |
| ENSG000001 | PRC1 | 3.09E-01 | 6.62E-01 | 9.07E-01 | -1.41E-01 | -1.10E+00 |

| | | | | | | | |
|---|---|---|---|---|---|---|---|
| ENSG000001 | GOLM2 | 3.09E-01 | | 6.62E-01 | 9.52E-01 | -7.12E-02 | -1.05E+00 |
| ENSG000001 | GASK1A | 3.10E-01 | ? | | 6.50E-01 | -6.22E-01 | -1.54E+00 |
| ENSG000002 | --- | 3.10E-01 | ? | | 7.90E-01 | -3.40E-01 | -1.27E+00 |
| ENSG000001 | NPY1R | 3.10E-01 | ? | | 6.21E-01 | -6.88E-01 | -1.61E+00 |
| ENSG000001 | RAB11FIP1 | 3.10E-01 | | 6.62E-01 | 1.07E+00 | 1.02E-01 | 1.07E+00 |
| ENSG000001 | SELENOM | 3.10E-01 | | 6.62E-01 | 1.11E+00 | 1.48E-01 | 1.11E+00 |
| ENSG000001 | APRT | 3.10E-01 | | 6.62E-01 | 1.07E+00 | 9.92E-02 | 1.07E+00 |
| ENSG000001 | PPIL1 | 3.10E-01 | | 6.62E-01 | 9.57E-01 | -6.32E-02 | -1.04E+00 |
| ENSG000002 | --- | 3.10E-01 | ? | | 6.31E-01 | -6.64E-01 | -1.58E+00 |
| ENSG00000C | SLC9A1 | 3.10E-01 | | 6.62E-01 | 1.03E+00 | 3.65E-02 | 1.03E+00 |
| ENSG000001 | TRIAP1 | 3.10E-01 | | 6.63E-01 | 1.04E+00 | 6.23E-02 | 1.04E+00 |
| ENSG000002 | LINC02928 | 3.10E-01 | ? | | 5.30E-01 | -9.16E-01 | -1.89E+00 |
| ENSG000001 | MECR | 3.10E-01 | | 6.63E-01 | 1.06E+00 | 8.71E-02 | 1.06E+00 |
| ENSG000002 | MIRLET7A1H | 3.10E-01 | | 6.63E-01 | 8.74E-01 | -1.94E-01 | -1.14E+00 |
| ENSG000001 | MCMBP | 3.10E-01 | | 6.63E-01 | 9.65E-01 | -5.12E-02 | -1.04E+00 |
| ENSG000001 | FLOT2 | 3.10E-01 | | 6.63E-01 | 9.59E-01 | -6.09E-02 | -1.04E+00 |
| ENSG000002 | RPL36AP44 | 3.10E-01 | ? | | 7.75E-01 | -3.68E-01 | -1.29E+00 |
| ENSG000002 | --- | 3.10E-01 | ? | | 6.41E-01 | -6.41E-01 | -1.56E+00 |
| ENSG000002 | LINC00954 | 3.10E-01 | ? | | 6.11E-01 | -7.11E-01 | -1.64E+00 |
| ENSG000001 | SOX17 | 3.10E-01 | ? | | 6.05E-01 | -7.26E-01 | -1.65E+00 |
| ENSG000001 | ARHGAP25 | 3.10E-01 | | 6.63E-01 | 9.54E-01 | -6.76E-02 | -1.05E+00 |
| ENSG000002 | --- | 3.10E-01 | | 6.63E-01 | 8.97E-01 | -1.57E-01 | -1.11E+00 |
| ENSG000001 | RANBP6 | 3.10E-01 | | 6.63E-01 | 1.03E+00 | 4.44E-02 | 1.03E+00 |
| ENSG000001 | GPAA1 | 3.10E-01 | | 6.63E-01 | 1.09E+00 | 1.30E-01 | 1.09E+00 |
| ENSG000002 | SNHG19 | 3.10E-01 | | 6.63E-01 | 1.17E+00 | 2.28E-01 | 1.17E+00 |
| ENSG000001 | ANKRD53 | 3.10E-01 | ? | | 6.89E-01 | -5.38E-01 | -1.45E+00 |
| ENSG000001 | AIFM1 | 3.11E-01 | | 6.63E-01 | 9.66E-01 | -4.95E-02 | -1.03E+00 |
| ENSG00000C | AP2B1 | 3.11E-01 | | 6.63E-01 | 1.02E+00 | 3.06E-02 | 1.02E+00 |
| ENSG000002 | HAGLR | 3.11E-01 | ? | | 7.35E-01 | -4.44E-01 | -1.36E+00 |
| ENSG000001 | SCAMP2 | 3.11E-01 | | 6.63E-01 | 1.04E+00 | 5.94E-02 | 1.04E+00 |
| ENSG000001 | VPS28 | 3.11E-01 | | 6.63E-01 | 1.06E+00 | 7.77E-02 | 1.06E+00 |
| ENSG000001 | ARSB | 3.11E-01 | | 6.63E-01 | 1.05E+00 | 6.75E-02 | 1.05E+00 |
| ENSG000001 | NUDT9 | 3.11E-01 | | 6.63E-01 | 9.73E-01 | -3.97E-02 | -1.03E+00 |
| ENSG000001 | SLC2A13 | 3.11E-01 | | 6.63E-01 | 9.37E-01 | -9.39E-02 | -1.07E+00 |
| ENSG000002 | OVCH1-AS1 | 3.11E-01 | ? | | 7.77E-01 | -3.64E-01 | -1.29E+00 |
| ENSG000002 | LINC01011 | 3.11E-01 | ? | | 1.23E+00 | 2.99E-01 | 1.23E+00 |
| ENSG000001 | MDFI | 3.11E-01 | | 6.63E-01 | 1.07E+00 | 9.14E-02 | 1.07E+00 |
| ENSG000001 | ZSCAN9 | 3.11E-01 | | 6.63E-01 | 9.47E-01 | -7.93E-02 | -1.06E+00 |
| ENSG000002 | --- | 3.11E-01 | ? | | 5.86E-01 | -7.71E-01 | -1.71E+00 |
| ENSG000001 | C1orf50 | 3.11E-01 | | 6.63E-01 | 1.06E+00 | 8.37E-02 | 1.06E+00 |
| ENSG00000C | ARHGAP6 | 3.11E-01 | ? | | 1.28E+00 | 3.61E-01 | 1.28E+00 |
| ENSG000001 | CACNA2D1 | 3.11E-01 | | 6.64E-01 | 9.02E-01 | -1.49E-01 | -1.11E+00 |
| ENSG00000C | SNRNP40 | 3.11E-01 | | 6.64E-01 | 1.03E+00 | 4.70E-02 | 1.03E+00 |
| ENSG000001 | MYC | 3.11E-01 | | 6.64E-01 | 1.06E+00 | 7.75E-02 | 1.06E+00 |
| ENSG000002 | SMIM20 | 3.11E-01 | | 6.64E-01 | 1.04E+00 | 5.71E-02 | 1.04E+00 |

| | | | | | | |
|---|---|---|---|---|---|---|
| ENSG000001 | BRICD5 | 3.12E-01 | ? | 8.05E-01 | -3.12E-01 | -1.24E+00 |
| ENSG000001 | CFAP157 | 3.12E-01 | 6.64E-01 | 1.12E+00 | 1.60E-01 | 1.12E+00 |
| ENSG000001 | H4C16 | 3.12E-01 | 6.64E-01 | 8.65E-01 | -2.10E-01 | -1.16E+00 |
| ENSG000001 | ATF1 | 3.12E-01 | 6.64E-01 | 9.51E-01 | -7.25E-02 | -1.05E+00 |
| ENSG000001 | VRK1 | 3.12E-01 | 6.64E-01 | 1.05E+00 | 7.53E-02 | 1.05E+00 |
| ENSG000001 | ZNF212 | 3.12E-01 | 6.64E-01 | 1.08E+00 | 1.10E-01 | 1.08E+00 |
| ENSG000002 | --- | 3.12E-01 | 6.64E-01 | 1.24E+00 | 3.12E-01 | 1.24E+00 |
| ENSG000001 | TMEM79 | 3.12E-01 | 6.64E-01 | 1.08E+00 | 1.17E-01 | 1.08E+00 |
| ENSG000001 | KICS2 | 3.12E-01 | 6.64E-01 | 1.06E+00 | 8.37E-02 | 1.06E+00 |
| ENSG000001 | THRA | 3.12E-01 | 6.64E-01 | 9.41E-01 | -8.72E-02 | -1.06E+00 |
| ENSG000001 | CSNK1G2 | 3.12E-01 | 6.64E-01 | 1.05E+00 | 6.43E-02 | 1.05E+00 |
| ENSG000001 | POLD4 | 3.12E-01 | 6.64E-01 | 1.10E+00 | 1.35E-01 | 1.10E+00 |
| ENSG000001 | MYH3 | 3.12E-01 | ? | 8.39E-01 | -2.54E-01 | -1.19E+00 |
| ENSG000002 | --- | 3.12E-01 | ? | 1.30E+00 | 3.77E-01 | 1.30E+00 |
| ENSG000002 | --- | 3.12E-01 | ? | 1.67E+00 | 7.38E-01 | 1.67E+00 |
| ENSG000001 | CEP85 | 3.12E-01 | 6.64E-01 | 9.52E-01 | -7.15E-02 | -1.05E+00 |
| ENSG000002 | DSTNP2 | 3.12E-01 | 6.64E-01 | 8.57E-01 | -2.23E-01 | -1.17E+00 |
| ENSG000001 | UBAC2 | 3.12E-01 | 6.64E-01 | 1.03E+00 | 4.51E-02 | 1.03E+00 |
| ENSG000002 | MCPH1-DT | 3.12E-01 | 6.64E-01 | 9.13E-01 | -1.32E-01 | -1.10E+00 |
| ENSG00000C | FOXC1 | 3.12E-01 | 6.64E-01 | 9.37E-01 | -9.37E-02 | -1.07E+00 |
| ENSG000001 | SMNDC1 | 3.12E-01 | 6.64E-01 | 9.65E-01 | -5.21E-02 | -1.04E+00 |
| ENSG00000C | PAFAH1B3 | 3.12E-01 | 6.64E-01 | 1.05E+00 | 7.68E-02 | 1.05E+00 |
| ENSG000001 | TMEM232 | 3.12E-01 | ? | 1.34E+00 | 4.22E-01 | 1.34E+00 |
| ENSG000001 | HASPIN | 3.13E-01 | 6.64E-01 | 1.11E+00 | 1.47E-01 | 1.11E+00 |
| ENSG000002 | --- | 3.13E-01 | ? | 1.21E+00 | 2.75E-01 | 1.21E+00 |
| ENSG000001 | SNX27 | 3.13E-01 | 6.64E-01 | 9.70E-01 | -4.47E-02 | -1.03E+00 |
| ENSG000001 | HP1BP3 | 3.13E-01 | 6.64E-01 | 9.59E-01 | -6.04E-02 | -1.04E+00 |
| ENSG000001 | ZNF846 | 3.13E-01 | 6.64E-01 | 9.24E-01 | -1.14E-01 | -1.08E+00 |
| ENSG000001 | MAPK14 | 3.13E-01 | 6.64E-01 | 9.68E-01 | -4.74E-02 | -1.03E+00 |
| ENSG000001 | SMCO2 | 3.13E-01 | ? | 7.40E-01 | -4.34E-01 | -1.35E+00 |
| ENSG000001 | SPEF2 | 3.13E-01 | 6.64E-01 | 1.15E+00 | 2.03E-01 | 1.15E+00 |
| ENSG000001 | SMPDL3B | 3.13E-01 | 6.64E-01 | 1.13E+00 | 1.82E-01 | 1.13E+00 |
| ENSG000001 | L3MBTL3 | 3.13E-01 | 6.64E-01 | 9.65E-01 | -5.12E-02 | -1.04E+00 |
| ENSG000001 | CARS1 | 3.13E-01 | 6.64E-01 | 9.72E-01 | -4.12E-02 | -1.03E+00 |
| ENSG000002 | ZNF286A-TB | 3.13E-01 | ? | 6.45E-01 | -6.32E-01 | -1.55E+00 |
| ENSG000002 | --- | 3.13E-01 | 6.64E-01 | 8.94E-01 | -1.61E-01 | -1.12E+00 |
| ENSG000001 | NR1H2 | 3.13E-01 | 6.64E-01 | 1.07E+00 | 9.77E-02 | 1.07E+00 |
| ENSG000001 | MX2 | 3.13E-01 | ? | 1.38E+00 | 4.65E-01 | 1.38E+00 |
| ENSG00000C | P3H2 | 3.13E-01 | 6.65E-01 | 1.07E+00 | 9.35E-02 | 1.07E+00 |
| ENSG000001 | TMEM91 | 3.13E-01 | 6.65E-01 | 1.14E+00 | 1.92E-01 | 1.14E+00 |
| ENSG000001 | SCLT1 | 3.13E-01 | 6.65E-01 | 9.35E-01 | -9.73E-02 | -1.07E+00 |
| ENSG00000C | TYRO3 | 3.14E-01 | 6.65E-01 | 1.08E+00 | 1.14E-01 | 1.08E+00 |
| ENSG000002 | --- | 3.14E-01 | 6.65E-01 | 7.88E-01 | -3.45E-01 | -1.27E+00 |
| ENSG000002 | ROCK1P1 | 3.14E-01 | ? | 7.84E-01 | -3.52E-01 | -1.28E+00 |
| ENSG000001 | TGFB1 | 3.14E-01 | 6.65E-01 | 9.54E-01 | -6.84E-02 | -1.05E+00 |

| | | | | | | |
|---|---|---|---|---|---|---|
| ENSG000001 | SS18 | 3.14E-01 | 6.65E-01 | 9.67E-01 | -4.89E-02 | -1.03E+00 |
| ENSG000001 | FRZB | 3.14E-01 | ? | 6.05E-01 | -7.25E-01 | -1.65E+00 |
| ENSG000001 | CSTF1 | 3.14E-01 | 6.65E-01 | 1.04E+00 | 5.18E-02 | 1.04E+00 |
| ENSG000001 | DACT1 | 3.14E-01 | ? | 1.27E+00 | 3.41E-01 | 1.27E+00 |
| ENSG00000C | TNIP3 | 3.14E-01 | ? | 1.25E+00 | 3.24E-01 | 1.25E+00 |
| ENSG000002 | CAPN8 | 3.14E-01 | ? | 1.34E+00 | 4.20E-01 | 1.34E+00 |
| ENSG000001 | VPS45 | 3.14E-01 | 6.65E-01 | 1.03E+00 | 4.81E-02 | 1.03E+00 |
| ENSG000001 | IFNAR2 | 3.14E-01 | 6.65E-01 | 1.04E+00 | 5.40E-02 | 1.04E+00 |
| ENSG000001 | MPLKIP | 3.14E-01 | 6.65E-01 | 1.06E+00 | 8.55E-02 | 1.06E+00 |
| ENSG000001 | NDUFC2 | 3.14E-01 | 6.65E-01 | 1.03E+00 | 4.64E-02 | 1.03E+00 |
| ENSG000002 | --- | 3.14E-01 | 6.65E-01 | 1.14E+00 | 1.94E-01 | 1.14E+00 |
| ENSG000001 | TEPSIN | 3.14E-01 | 6.65E-01 | 1.09E+00 | 1.26E-01 | 1.09E+00 |
| ENSG000001 | HSPB2 | 3.14E-01 | ? | 1.68E+00 | 7.47E-01 | 1.68E+00 |
| ENSG000001 | ARPC2 | 3.14E-01 | 6.65E-01 | 1.02E+00 | 3.00E-02 | 1.02E+00 |
| ENSG000001 | C4orf33 | 3.14E-01 | 6.65E-01 | 9.43E-01 | -8.50E-02 | -1.06E+00 |
| ENSG000001 | RIN1 | 3.14E-01 | 6.65E-01 | 1.07E+00 | 1.02E-01 | 1.07E+00 |
| ENSG000001 | FGF2 | 3.14E-01 | 6.65E-01 | 8.50E-01 | -2.34E-01 | -1.18E+00 |
| ENSG000002 | WBP1LP2 | 3.14E-01 | ? | 1.20E+00 | 2.64E-01 | 1.20E+00 |
| ENSG000002 | --- | 3.14E-01 | 6.65E-01 | 9.07E-01 | -1.41E-01 | -1.10E+00 |
| ENSG000001 | BRSK1 | 3.14E-01 | 6.65E-01 | 1.15E+00 | 2.06E-01 | 1.15E+00 |
| ENSG000001 | DYNC2H1 | 3.15E-01 | 6.66E-01 | 1.07E+00 | 9.11E-02 | 1.07E+00 |
| ENSG000002 | SPRY4-AS1 | 3.15E-01 | ? | 1.24E+00 | 3.11E-01 | 1.24E+00 |
| ENSG000001 | TWSG1 | 3.15E-01 | 6.66E-01 | 9.46E-01 | -7.97E-02 | -1.06E+00 |
| ENSG000001 | TNPO2 | 3.15E-01 | 6.66E-01 | 1.04E+00 | 5.40E-02 | 1.04E+00 |
| ENSG000002 | NDUFB2-AS1 | 3.15E-01 | 6.66E-01 | 8.83E-01 | -1.79E-01 | -1.13E+00 |
| ENSG000001 | ATP13A1 | 3.15E-01 | 6.66E-01 | 1.04E+00 | 6.22E-02 | 1.04E+00 |
| ENSG000002 | --- | 3.15E-01 | ? | 1.53E+00 | 6.15E-01 | 1.53E+00 |
| ENSG000002 | --- | 3.15E-01 | 6.66E-01 | 8.77E-01 | -1.89E-01 | -1.14E+00 |
| ENSG000001 | AK1 | 3.15E-01 | 6.66E-01 | 1.06E+00 | 8.07E-02 | 1.06E+00 |
| ENSG000002 | --- | 3.15E-01 | ? | 8.19E-01 | -2.88E-01 | -1.22E+00 |
| ENSG000002 | --- | 3.15E-01 | ? | 6.06E-01 | -7.22E-01 | -1.65E+00 |
| ENSG000001 | ZNF768 | 3.15E-01 | 6.66E-01 | 1.05E+00 | 7.32E-02 | 1.05E+00 |
| ENSG00000C | ACOT7 | 3.15E-01 | 6.66E-01 | 1.04E+00 | 6.25E-02 | 1.04E+00 |
| ENSG000002 | TMEFF1 | 3.16E-01 | ? | 5.66E-01 | -8.22E-01 | -1.77E+00 |
| ENSG000001 | INHA | 3.16E-01 | ? | 6.38E-01 | -6.49E-01 | -1.57E+00 |
| ENSG000001 | POLR1B | 3.16E-01 | 6.67E-01 | 1.04E+00 | 5.24E-02 | 1.04E+00 |
| ENSG000001 | STIP1 | 3.16E-01 | 6.67E-01 | 9.74E-01 | -3.82E-02 | -1.03E+00 |
| ENSG000001 | PLCE1 | 3.16E-01 | 6.67E-01 | 9.29E-01 | -1.07E-01 | -1.08E+00 |
| ENSG000002 | HSP90AB2P | 3.16E-01 | ? | 1.38E+00 | 4.60E-01 | 1.38E+00 |
| ENSG000001 | TMEM33 | 3.16E-01 | 6.67E-01 | 9.59E-01 | -5.97E-02 | -1.04E+00 |
| ENSG000001 | IQCK | 3.16E-01 | 6.68E-01 | 9.42E-01 | -8.62E-02 | -1.06E+00 |
| ENSG000002 | --- | 3.16E-01 | ? | 1.89E+00 | 9.22E-01 | 1.89E+00 |
| ENSG000002 | RPL37P12 | 3.17E-01 | ? | 1.61E+00 | 6.91E-01 | 1.61E+00 |
| ENSG000001 | FSD1L | 3.17E-01 | 6.68E-01 | 9.31E-01 | -1.03E-01 | -1.07E+00 |
| ENSG00000C | MYLK | 3.17E-01 | 6.68E-01 | 1.12E+00 | 1.69E-01 | 1.12E+00 |

| | | | | | | |
|---|---|---|---|---|---|---|
| ENSG000001 | SYNGR2 | 3.17E-01 | 6.68E-01 | 9.43E-01 | -8.49E-02 | -1.06E+00 |
| ENSG000001 | PTK2B | 3.17E-01 | 6.68E-01 | 9.28E-01 | -1.08E-01 | -1.08E+00 |
| ENSG000002 | ZNF8-DT | 3.17E-01 | 6.68E-01 | 8.91E-01 | -1.66E-01 | -1.12E+00 |
| ENSG000001 | UBQLN1 | 3.17E-01 | 6.68E-01 | 9.74E-01 | -3.75E-02 | -1.03E+00 |
| ENSG000002 | SCDP1 | 3.17E-01 | ? | 6.79E-01 | -5.59E-01 | -1.47E+00 |
| ENSG000001 | MYPOP | 3.17E-01 | 6.68E-01 | 1.09E+00 | 1.21E-01 | 1.09E+00 |
| ENSG000002 | EPHA2-AS1 | 3.17E-01 | ? | 7.99E-01 | -3.25E-01 | -1.25E+00 |
| ENSG000002 | --- | 3.17E-01 | ? | 1.49E+00 | 5.78E-01 | 1.49E+00 |
| ENSG000002 | KANSL1L-AS1 | 3.17E-01 | ? | 7.16E-01 | -4.82E-01 | -1.40E+00 |
| ENSG000001 | NOTCH2 | 3.17E-01 | 6.68E-01 | 1.06E+00 | 8.29E-02 | 1.06E+00 |
| ENSG000001 | GMPPB | 3.17E-01 | 6.68E-01 | 9.50E-01 | -7.36E-02 | -1.05E+00 |
| ENSG000001 | RAB11FIP2 | 3.17E-01 | 6.68E-01 | 1.05E+00 | 7.57E-02 | 1.05E+00 |
| ENSG000001 | HTR1B | 3.17E-01 | 6.68E-01 | 9.00E-01 | -1.53E-01 | -1.11E+00 |
| ENSG000001 | WDR48 | 3.17E-01 | 6.68E-01 | 9.66E-01 | -5.03E-02 | -1.04E+00 |
| ENSG000001 | CCND1 | 3.17E-01 | 6.68E-01 | 9.65E-01 | -5.09E-02 | -1.04E+00 |
| ENSG000000 | CHFR | 3.17E-01 | 6.68E-01 | 9.62E-01 | -5.64E-02 | -1.04E+00 |
| ENSG000001 | POLG | 3.17E-01 | 6.68E-01 | 1.04E+00 | 5.62E-02 | 1.04E+00 |
| ENSG000001 | HDX | 3.17E-01 | 6.68E-01 | 9.36E-01 | -9.48E-02 | -1.07E+00 |
| ENSG000001 | UBP1 | 3.17E-01 | 6.68E-01 | 9.75E-01 | -3.59E-02 | -1.03E+00 |
| ENSG000001 | GAD1 | 3.17E-01 | 6.68E-01 | 1.20E+00 | 2.62E-01 | 1.20E+00 |
| ENSG000002 | SOCS7 | 3.17E-01 | 6.68E-01 | 1.04E+00 | 5.61E-02 | 1.04E+00 |
| ENSG000002 | NBPF19 | 3.17E-01 | 6.68E-01 | 9.39E-01 | -9.03E-02 | -1.06E+00 |
| ENSG000002 | RPSAP54 | 3.17E-01 | ? | 6.70E-01 | -5.78E-01 | -1.49E+00 |
| ENSG000002 | MAN1B1-DT | 3.18E-01 | ? | 1.18E+00 | 2.35E-01 | 1.18E+00 |
| ENSG000002 | UBE2SP1 | 3.18E-01 | ? | 7.58E-01 | -4.00E-01 | -1.32E+00 |
| ENSG000002 | SMIM22 | 3.18E-01 | 6.68E-01 | 1.18E+00 | 2.39E-01 | 1.18E+00 |
| ENSG000001 | FAT4 | 3.18E-01 | 6.68E-01 | 1.13E+00 | 1.76E-01 | 1.13E+00 |
| ENSG000002 | TPT1P4 | 3.18E-01 | ? | 6.47E-01 | -6.29E-01 | -1.55E+00 |
| ENSG000002 | --- | 3.18E-01 | ? | 6.72E-01 | -5.74E-01 | -1.49E+00 |
| ENSG000001 | GSPT2 | 3.18E-01 | 6.68E-01 | 9.42E-01 | -8.58E-02 | -1.06E+00 |
| ENSG000002 | TRIM16L | 3.18E-01 | 6.68E-01 | 9.18E-01 | -1.23E-01 | -1.09E+00 |
| ENSG000001 | ACYP2 | 3.18E-01 | 6.68E-01 | 1.05E+00 | 7.57E-02 | 1.05E+00 |
| ENSG000002 | IPO5P1 | 3.18E-01 | 6.68E-01 | 1.08E+00 | 1.11E-01 | 1.08E+00 |
| ENSG000002 | ZDHHC12-DT | 3.18E-01 | 6.68E-01 | 1.17E+00 | 2.29E-01 | 1.17E+00 |
| ENSG000001 | PREPL | 3.18E-01 | 6.68E-01 | 9.55E-01 | -6.65E-02 | -1.05E+00 |
| ENSG000000 | ACTR6 | 3.18E-01 | 6.68E-01 | 1.03E+00 | 4.81E-02 | 1.03E+00 |
| ENSG000000 | FOXP3 | 3.18E-01 | ? | 1.49E+00 | 5.73E-01 | 1.49E+00 |
| ENSG000002 | TRAC | 3.18E-01 | ? | 1.27E+00 | 3.46E-01 | 1.27E+00 |
| ENSG000002 | RDUR | 3.18E-01 | 6.68E-01 | 1.55E+00 | 6.36E-01 | 1.55E+00 |
| ENSG000001 | ADAM23 | 3.18E-01 | 6.69E-01 | 1.09E+00 | 1.18E-01 | 1.09E+00 |
| ENSG000000 | SLC4A4 | 3.18E-01 | 6.69E-01 | 9.04E-01 | -1.46E-01 | -1.11E+00 |
| ENSG000000 | CNOT4 | 3.18E-01 | 6.69E-01 | 9.49E-01 | -7.48E-02 | -1.05E+00 |
| ENSG000002 | ERVV-2 | 3.18E-01 | ? | 6.63E-01 | -5.94E-01 | -1.51E+00 |
| ENSG000002 | --- | 3.18E-01 | 6.69E-01 | 1.08E+00 | 1.16E-01 | 1.08E+00 |
| ENSG000001 | HSF1 | 3.19E-01 | 6.69E-01 | 1.05E+00 | 6.87E-02 | 1.05E+00 |

| | | | | | | |
|---|---|---|---|---|---|---|
| ENSG000001 | CAAP1 | 3.19E-01 | 6.69E-01 | 9.53E-01 | -7.02E-02 | -1.05E+00 |
| ENSG000001 | ZNF668 | 3.19E-01 | 6.69E-01 | 1.10E+00 | 1.38E-01 | 1.10E+00 |
| ENSG000002 | SYS1 | 3.19E-01 | 6.69E-01 | 1.05E+00 | 6.85E-02 | 1.05E+00 |
| ENSG000002 | --- | 3.19E-01 | ? | 6.18E-01 | -6.94E-01 | -1.62E+00 |
| ENSG000002 | --- | 3.19E-01 | 6.69E-01 | 9.26E-01 | -1.11E-01 | -1.08E+00 |
| ENSG000001 | FAAP100 | 3.19E-01 | 6.69E-01 | 1.08E+00 | 1.06E-01 | 1.08E+00 |
| ENSG000001 | MRPS9 | 3.19E-01 | 6.70E-01 | 9.56E-01 | -6.51E-02 | -1.05E+00 |
| ENSG000002 | KCTD21-AS1 | 3.19E-01 | 6.70E-01 | 8.44E-01 | -2.45E-01 | -1.18E+00 |
| ENSG00000C | TMEM40 | 3.19E-01 | 6.70E-01 | 1.06E+00 | 8.14E-02 | 1.06E+00 |
| ENSG000002 | --- | 3.19E-01 | ? | 7.80E-01 | -3.59E-01 | -1.28E+00 |
| ENSG00000C | RORA | 3.19E-01 | 6.70E-01 | 1.08E+00 | 1.05E-01 | 1.08E+00 |
| ENSG000001 | TMEM150A | 3.19E-01 | 6.70E-01 | 1.06E+00 | 8.73E-02 | 1.06E+00 |
| ENSG00000C | IL12RB2 | 3.19E-01 | 6.70E-01 | 1.11E+00 | 1.51E-01 | 1.11E+00 |
| ENSG000002 | --- | 3.19E-01 | ? | 1.32E+00 | 4.01E-01 | 1.32E+00 |
| ENSG000001 | FEZ2 | 3.19E-01 | 6.70E-01 | 1.03E+00 | 4.15E-02 | 1.03E+00 |
| ENSG000001 | TMEM61 | 3.20E-01 | 6.70E-01 | 1.15E+00 | 2.02E-01 | 1.15E+00 |
| ENSG000002 | LINC01521 | 3.20E-01 | ? | 7.98E-01 | -3.26E-01 | -1.25E+00 |
| ENSG000002 | GDI2P2 | 3.20E-01 | ? | 1.52E+00 | 6.04E-01 | 1.52E+00 |
| ENSG000002 | C2CD4D | 3.20E-01 | ? | 6.40E-01 | -6.44E-01 | -1.56E+00 |
| ENSG000002 | --- | 3.20E-01 | ? | 1.31E+00 | 3.87E-01 | 1.31E+00 |
| ENSG000002 | SNHG26 | 3.20E-01 | 6.70E-01 | 9.27E-01 | -1.09E-01 | -1.08E+00 |
| ENSG000001 | ARHGAP5 | 3.20E-01 | 6.70E-01 | 1.04E+00 | 5.84E-02 | 1.04E+00 |
| ENSG00000C | HECW1 | 3.20E-01 | ? | 7.50E-01 | -4.15E-01 | -1.33E+00 |
| ENSG000002 | --- | 3.20E-01 | 6.70E-01 | 8.37E-01 | -2.56E-01 | -1.19E+00 |
| ENSG00000C | POLRMT | 3.20E-01 | 6.70E-01 | 1.07E+00 | 9.79E-02 | 1.07E+00 |
| ENSG00000C | OTUD5 | 3.20E-01 | 6.70E-01 | 9.70E-01 | -4.44E-02 | -1.03E+00 |
| ENSG000002 | --- | 3.20E-01 | ? | 1.37E+00 | 4.56E-01 | 1.37E+00 |
| ENSG000002 | CCDC169 | 3.20E-01 | ? | 1.19E+00 | 2.51E-01 | 1.19E+00 |
| ENSG000001 | FEM1A | 3.20E-01 | 6.70E-01 | 1.08E+00 | 1.06E-01 | 1.08E+00 |
| ENSG000001 | RTN3 | 3.20E-01 | 6.70E-01 | 1.03E+00 | 3.81E-02 | 1.03E+00 |
| ENSG000001 | FLVCR1-DT | 3.20E-01 | ? | 1.25E+00 | 3.18E-01 | 1.25E+00 |
| ENSG000001 | RBM33 | 3.20E-01 | 6.70E-01 | 9.63E-01 | -5.40E-02 | -1.04E+00 |
| ENSG000001 | FAHD2A | 3.20E-01 | 6.70E-01 | 1.04E+00 | 6.19E-02 | 1.04E+00 |
| ENSG000001 | PSRC1 | 3.20E-01 | 6.70E-01 | 1.12E+00 | 1.59E-01 | 1.12E+00 |
| ENSG000001 | ITGA5 | 3.20E-01 | 6.70E-01 | 1.04E+00 | 5.79E-02 | 1.04E+00 |
| ENSG000001 | RBMX | 3.20E-01 | 6.70E-01 | 1.03E+00 | 4.10E-02 | 1.03E+00 |
| ENSG000001 | MAMSTR | 3.20E-01 | ? | 1.29E+00 | 3.72E-01 | 1.29E+00 |
| ENSG00000C | SDK2 | 3.20E-01 | 6.70E-01 | 1.05E+00 | 6.79E-02 | 1.05E+00 |
| ENSG000002 | ZNF32-AS2 | 3.20E-01 | ? | 7.46E-01 | -4.23E-01 | -1.34E+00 |
| ENSG000001 | MEPCE | 3.20E-01 | 6.70E-01 | 1.05E+00 | 7.00E-02 | 1.05E+00 |
| ENSG000002 | CLCA3P | 3.20E-01 | ? | 5.15E-01 | -9.57E-01 | -1.94E+00 |
| ENSG000001 | MAML2 | 3.21E-01 | 6.70E-01 | 9.49E-01 | -7.59E-02 | -1.05E+00 |
| ENSG000002 | C1DP1 | 3.21E-01 | ? | 1.28E+00 | 3.57E-01 | 1.28E+00 |
| ENSG000001 | CYB561D2 | 3.21E-01 | 6.70E-01 | 1.13E+00 | 1.75E-01 | 1.13E+00 |
| ENSG000001 | APOD | 3.21E-01 | 6.70E-01 | 9.45E-01 | -8.11E-02 | -1.06E+00 |

| | | | | | | |
|---|---|---|---|---|---|---|
| ENSG000001 | DARS1 | 3.21E-01 | 6.70E-01 | 9.75E-01 | -3.64E-02 | -1.03E+00 |
| ENSG000001 | PHF3 | 3.21E-01 | 6.70E-01 | 9.54E-01 | -6.74E-02 | -1.05E+00 |
| ENSG000001 | TMEM139 | 3.21E-01 | 6.70E-01 | 9.31E-01 | -1.03E-01 | -1.07E+00 |
| ENSG000002 | LINC02739 | 3.21E-01 ? | | 6.55E-01 | -6.11E-01 | -1.53E+00 |
| ENSG000001 | C2orf68 | 3.21E-01 | 6.70E-01 | 9.45E-01 | -8.17E-02 | -1.06E+00 |
| ENSG000001 | IRS1 | 3.21E-01 | 6.70E-01 | 9.68E-01 | -4.76E-02 | -1.03E+00 |
| ENSG000001 | AK4 | 3.21E-01 | 6.70E-01 | 9.64E-01 | -5.33E-02 | -1.04E+00 |
| ENSG000001 | DYNC2LI1 | 3.21E-01 | 6.70E-01 | 1.05E+00 | 7.45E-02 | 1.05E+00 |
| ENSG00000C | RAD52 | 3.21E-01 | 6.70E-01 | 9.11E-01 | -1.34E-01 | -1.10E+00 |
| ENSG000002 | --- | 3.21E-01 ? | | 6.98E-01 | -5.18E-01 | -1.43E+00 |
| ENSG00000C | ERMP1 | 3.21E-01 | 6.70E-01 | 1.03E+00 | 4.80E-02 | 1.03E+00 |
| ENSG000002 | --- | 3.21E-01 ? | | 1.20E+00 | 2.63E-01 | 1.20E+00 |
| ENSG000001 | --- | 3.21E-01 | 6.70E-01 | 1.08E+00 | 1.17E-01 | 1.08E+00 |
| ENSG000001 | PSMC1 | 3.21E-01 | 6.70E-01 | 9.80E-01 | -2.95E-02 | -1.02E+00 |
| ENSG000002 | RPSAP41 | 3.21E-01 ? | | 8.15E-01 | -2.95E-01 | -1.23E+00 |
| ENSG000001 | YIPF4 | 3.21E-01 | 6.70E-01 | 9.73E-01 | -3.98E-02 | -1.03E+00 |
| ENSG000001 | MATN1-AS1 | 3.21E-01 ? | | 1.21E+00 | 2.74E-01 | 1.21E+00 |
| ENSG000001 | TAF6L | 3.21E-01 | 6.70E-01 | 1.08E+00 | 1.17E-01 | 1.08E+00 |
| ENSG00000C | FGFR3 | 3.21E-01 | 6.70E-01 | 9.11E-01 | -1.34E-01 | -1.10E+00 |
| ENSG000001 | RERE | 3.21E-01 | 6.70E-01 | 9.63E-01 | -5.38E-02 | -1.04E+00 |
| ENSG00000C | CTSA | 3.21E-01 | 6.70E-01 | 1.04E+00 | 5.05E-02 | 1.04E+00 |
| ENSG00000C | DDHD2 | 3.21E-01 | 6.70E-01 | 1.03E+00 | 3.72E-02 | 1.03E+00 |
| ENSG000002 | --- | 3.21E-01 ? | | 1.41E+00 | 4.95E-01 | 1.41E+00 |
| ENSG000001 | CFD | 3.21E-01 ? | | 1.27E+00 | 3.48E-01 | 1.27E+00 |
| ENSG00000C | SYT7 | 3.21E-01 | 6.70E-01 | 9.12E-01 | -1.33E-01 | -1.10E+00 |
| ENSG000002 | --- | 3.22E-01 ? | | 6.45E-01 | -6.33E-01 | -1.55E+00 |
| ENSG000002 | --- | 3.22E-01 ? | | 7.83E-01 | -3.53E-01 | -1.28E+00 |
| ENSG000001 | PPRC1 | 3.22E-01 | 6.70E-01 | 1.03E+00 | 3.73E-02 | 1.03E+00 |
| ENSG000001 | STYXL1 | 3.22E-01 | 6.70E-01 | 1.05E+00 | 6.73E-02 | 1.05E+00 |
| ENSG000001 | SPTBN2 | 3.22E-01 | 6.70E-01 | 1.03E+00 | 4.50E-02 | 1.03E+00 |
| ENSG000002 | PEG13 | 3.22E-01 ? | | 6.48E-01 | -6.26E-01 | -1.54E+00 |
| ENSG00000C | ADRB1 | 3.22E-01 ? | | 7.02E-01 | -5.11E-01 | -1.42E+00 |
| ENSG000001 | ERICH5 | 3.22E-01 | 6.71E-01 | 8.87E-01 | -1.73E-01 | -1.13E+00 |
| ENSG000002 | --- | 3.22E-01 ? | | 1.54E+00 | 6.25E-01 | 1.54E+00 |
| ENSG000002 | ALOX12P2 | 3.22E-01 | 6.71E-01 | 8.69E-01 | -2.02E-01 | -1.15E+00 |
| ENSG000001 | ESYT3 | 3.22E-01 | 6.71E-01 | 1.16E+00 | 2.18E-01 | 1.16E+00 |
| ENSG000001 | ZFP36L1 | 3.22E-01 | 6.71E-01 | 9.59E-01 | -6.02E-02 | -1.04E+00 |
| ENSG000001 | MTFMT | 3.22E-01 | 6.71E-01 | 1.04E+00 | 5.65E-02 | 1.04E+00 |
| ENSG000002 | C17orf100 | 3.22E-01 | 6.71E-01 | 1.09E+00 | 1.29E-01 | 1.09E+00 |
| ENSG000002 | DHX16 | 3.22E-01 | 6.71E-01 | 9.68E-01 | -4.68E-02 | -1.03E+00 |
| ENSG000002 | DUXAP9 | 3.22E-01 | 6.71E-01 | 1.15E+00 | 2.01E-01 | 1.15E+00 |
| ENSG000001 | MCTS2 | 3.22E-01 ? | | 8.16E-01 | -2.93E-01 | -1.22E+00 |
| ENSG000001 | --- | 3.22E-01 ? | | 1.56E+00 | 6.40E-01 | 1.56E+00 |
| ENSG000002 | THRIL | 3.22E-01 ? | | 1.79E+00 | 8.41E-01 | 1.79E+00 |
| ENSG000001 | B3GALNT1 | 3.22E-01 | 6.71E-01 | 9.37E-01 | -9.42E-02 | -1.07E+00 |

| | | | | | | |
|---|---|---|---|---|---|---|
| ENSG000001 | LSM10 | 3.22E-01 | 6.71E-01 | 9.35E-01 | -9.62E-02 | -1.07E+00 |
| ENSG000001 | LRRC19 | 3.22E-01 | ? | 1.40E+00 | 4.86E-01 | 1.40E+00 |
| ENSG000001 | UHMK1 | 3.22E-01 | 6.71E-01 | 1.03E+00 | 4.68E-02 | 1.03E+00 |
| ENSG000002 | --- | 3.22E-01 | ? | 1.57E+00 | 6.55E-01 | 1.57E+00 |
| ENSG000002 | --- | 3.23E-01 | 6.71E-01 | 7.55E-01 | -4.06E-01 | -1.33E+00 |
| ENSG000002 | ATAD3C | 3.23E-01 | 6.71E-01 | 1.12E+00 | 1.61E-01 | 1.12E+00 |
| ENSG000002 | --- | 3.23E-01 | ? | 1.37E+00 | 4.49E-01 | 1.37E+00 |
| ENSG000002 | ZBTB22 | 3.23E-01 | 6.71E-01 | 1.06E+00 | 8.63E-02 | 1.06E+00 |
| ENSG000001 | HNMT | 3.23E-01 | 6.71E-01 | 1.13E+00 | 1.72E-01 | 1.13E+00 |
| ENSG000002 | FAM216A | 3.23E-01 | 6.71E-01 | 1.07E+00 | 9.92E-02 | 1.07E+00 |
| ENSG000002 | --- | 3.23E-01 | ? | 7.97E-01 | -3.28E-01 | -1.25E+00 |
| ENSG000001 | CHST7 | 3.23E-01 | 6.72E-01 | 1.13E+00 | 1.81E-01 | 1.13E+00 |
| ENSG000002 | LPAL2 | 3.23E-01 | ? | 1.50E+00 | 5.82E-01 | 1.50E+00 |
| ENSG000001 | TXNDC17 | 3.23E-01 | 6.72E-01 | 1.06E+00 | 7.75E-02 | 1.06E+00 |
| ENSG000001 | LARP4 | 3.23E-01 | 6.72E-01 | 9.60E-01 | -5.86E-02 | -1.04E+00 |
| ENSG000002 | --- | 3.23E-01 | ? | 1.33E+00 | 4.06E-01 | 1.33E+00 |
| ENSG000001 | PDCD10 | 3.23E-01 | 6.72E-01 | 9.60E-01 | -5.85E-02 | -1.04E+00 |
| ENSG00000C | DEPDC1B | 3.24E-01 | 6.72E-01 | 9.24E-01 | -1.14E-01 | -1.08E+00 |
| ENSG000001 | TLX2 | 3.24E-01 | ? | 1.61E+00 | 6.89E-01 | 1.61E+00 |
| ENSG000001 | RNF34 | 3.24E-01 | 6.72E-01 | 1.03E+00 | 3.84E-02 | 1.03E+00 |
| ENSG00000C | NRCAM | 3.24E-01 | 6.72E-01 | 9.35E-01 | -9.64E-02 | -1.07E+00 |
| ENSG000001 | SRSF9 | 3.24E-01 | 6.72E-01 | 1.03E+00 | 4.72E-02 | 1.03E+00 |
| ENSG000002 | --- | 3.24E-01 | 6.72E-01 | 9.26E-01 | -1.11E-01 | -1.08E+00 |
| ENSG000001 | ZNF467 | 3.24E-01 | ? | 8.00E-01 | -3.22E-01 | -1.25E+00 |
| ENSG000002 | --- | 3.24E-01 | ? | 7.58E-01 | -3.99E-01 | -1.32E+00 |
| ENSG000002 | LINC00920 | 3.24E-01 | 6.73E-01 | 1.16E+00 | 2.11E-01 | 1.16E+00 |
| ENSG00000C | TUBE1 | 3.24E-01 | 6.73E-01 | 9.30E-01 | -1.05E-01 | -1.08E+00 |
| ENSG000001 | ENDOV | 3.24E-01 | 6.73E-01 | 1.09E+00 | 1.19E-01 | 1.09E+00 |
| ENSG000001 | MRPL41 | 3.24E-01 | 6.73E-01 | 1.08E+00 | 1.14E-01 | 1.08E+00 |
| ENSG000001 | TCEANC2 | 3.24E-01 | 6.73E-01 | 1.05E+00 | 7.57E-02 | 1.05E+00 |
| ENSG000001 | MRTFA | 3.24E-01 | 6.73E-01 | 1.04E+00 | 5.08E-02 | 1.04E+00 |
| ENSG000002 | FAM174C | 3.24E-01 | 6.73E-01 | 1.08E+00 | 1.14E-01 | 1.08E+00 |
| ENSG00000C | LLGL2 | 3.24E-01 | 6.73E-01 | 1.08E+00 | 1.12E-01 | 1.08E+00 |
| ENSG000001 | WDR36 | 3.24E-01 | 6.73E-01 | 1.03E+00 | 4.58E-02 | 1.03E+00 |
| ENSG000001 | GATA3 | 3.25E-01 | ? | 1.23E+00 | 3.01E-01 | 1.23E+00 |
| ENSG00000C | MASP2 | 3.25E-01 | 6.73E-01 | 8.97E-01 | -1.56E-01 | -1.11E+00 |
| ENSG000002 | GLIPR1-AS1 | 3.25E-01 | 6.73E-01 | 8.83E-01 | -1.79E-01 | -1.13E+00 |
| ENSG00000C | ALPK1 | 3.25E-01 | 6.73E-01 | 9.54E-01 | -6.82E-02 | -1.05E+00 |
| ENSG000001 | BEX1 | 3.25E-01 | ? | 1.39E+00 | 4.80E-01 | 1.39E+00 |
| ENSG00000C | LAS1L | 3.25E-01 | 6.73E-01 | 1.03E+00 | 4.38E-02 | 1.03E+00 |
| ENSG000002 | RPL7P57 | 3.25E-01 | ? | 1.59E+00 | 6.68E-01 | 1.59E+00 |
| ENSG00000C | TP53INP2 | 3.25E-01 | 6.73E-01 | 9.34E-01 | -9.87E-02 | -1.07E+00 |
| ENSG000001 | SMAD2 | 3.25E-01 | 6.73E-01 | 1.02E+00 | 3.53E-02 | 1.02E+00 |
| ENSG000001 | GADD45GIP1 | 3.25E-01 | 6.73E-01 | 9.54E-01 | -6.85E-02 | -1.05E+00 |
| ENSG000002 | MIR205HG | 3.25E-01 | 6.73E-01 | 1.07E+00 | 9.62E-02 | 1.07E+00 |

| | | | | | | |
|---|---|---|---|---|---|---|
| ENSG000001 | LRRC4C | 3.25E-01 | ? | 1.46E+00 | 5.42E-01 | 1.46E+00 |
| ENSG000001 | P4HB | 3.25E-01 | 6.73E-01 | 9.56E-01 | -6.45E-02 | -1.05E+00 |
| ENSG000001 | IDNK | 3.25E-01 | ? | 1.19E+00 | 2.52E-01 | 1.19E+00 |
| ENSG000001 | RINL | 3.25E-01 | 6.73E-01 | 1.07E+00 | 9.77E-02 | 1.07E+00 |
| ENSG000001 | SPTLC3 | 3.25E-01 | 6.73E-01 | 1.06E+00 | 7.98E-02 | 1.06E+00 |
| ENSG000001 | TOM1L2 | 3.25E-01 | 6.73E-01 | 1.04E+00 | 5.53E-02 | 1.04E+00 |
| ENSG00000C | TDRD1 | 3.25E-01 | ? | 1.29E+00 | 3.63E-01 | 1.29E+00 |
| ENSG00000C | TIMM13 | 3.25E-01 | 6.73E-01 | 1.08E+00 | 1.13E-01 | 1.08E+00 |
| ENSG00000C | DVL2 | 3.25E-01 | 6.73E-01 | 1.07E+00 | 9.23E-02 | 1.07E+00 |
| ENSG000002 | INAFM2 | 3.25E-01 | 6.73E-01 | 1.04E+00 | 5.36E-02 | 1.04E+00 |
| ENSG000002 | COL11A2 | 3.25E-01 | 6.73E-01 | 1.16E+00 | 2.12E-01 | 1.16E+00 |
| ENSG000002 | --- | 3.25E-01 | ? | 6.38E-01 | -6.48E-01 | -1.57E+00 |
| ENSG000001 | RIT1 | 3.26E-01 | 6.73E-01 | 1.04E+00 | 4.99E-02 | 1.04E+00 |
| ENSG000002 | --- | 3.26E-01 | ? | 5.76E-01 | -7.96E-01 | -1.74E+00 |
| ENSG000001 | TOR4A | 3.26E-01 | 6.74E-01 | 9.32E-01 | -1.01E-01 | -1.07E+00 |
| ENSG000002 | --- | 3.26E-01 | ? | 7.28E-01 | -4.59E-01 | -1.37E+00 |
| ENSG000001 | ZNF221 | 3.26E-01 | 6.74E-01 | 9.10E-01 | -1.37E-01 | -1.10E+00 |
| ENSG000001 | KLHL3 | 3.26E-01 | 6.74E-01 | 9.19E-01 | -1.22E-01 | -1.09E+00 |
| ENSG000002 | --- | 3.26E-01 | 6.74E-01 | 8.64E-01 | -2.11E-01 | -1.16E+00 |
| ENSG00000C | RNF31 | 3.26E-01 | 6.74E-01 | 1.05E+00 | 7.31E-02 | 1.05E+00 |
| ENSG000002 | NSMCE1-DT | 3.26E-01 | ? | 6.97E-01 | -5.20E-01 | -1.43E+00 |
| ENSG000002 | KMT2E-AS1 | 3.26E-01 | ? | 1.22E+00 | 2.92E-01 | 1.22E+00 |
| ENSG00000C | PITPNM2 | 3.26E-01 | 6.74E-01 | 9.50E-01 | -7.38E-02 | -1.05E+00 |
| ENSG000001 | PPP1R3B | 3.26E-01 | 6.74E-01 | 9.53E-01 | -6.92E-02 | -1.05E+00 |
| ENSG000001 | PNPLA7 | 3.26E-01 | 6.74E-01 | 8.63E-01 | -2.13E-01 | -1.16E+00 |
| ENSG000002 | --- | 3.26E-01 | 6.74E-01 | 9.41E-01 | -8.83E-02 | -1.06E+00 |
| ENSG000001 | CBR4 | 3.26E-01 | 6.74E-01 | 9.39E-01 | -9.01E-02 | -1.06E+00 |
| ENSG000002 | SNX18P7 | 3.26E-01 | ? | 7.91E-01 | -3.39E-01 | -1.26E+00 |
| ENSG000001 | NFIC | 3.26E-01 | 6.74E-01 | 9.68E-01 | -4.64E-02 | -1.03E+00 |
| ENSG000001 | LSM4 | 3.26E-01 | 6.74E-01 | 1.06E+00 | 8.43E-02 | 1.06E+00 |
| ENSG000001 | SLC25A21 | 3.26E-01 | 6.74E-01 | 8.49E-01 | -2.37E-01 | -1.18E+00 |
| ENSG000002 | RPL15P3 | 3.26E-01 | 6.74E-01 | 9.11E-01 | -1.34E-01 | -1.10E+00 |
| ENSG000002 | --- | 3.26E-01 | ? | 6.33E-01 | -6.59E-01 | -1.58E+00 |
| ENSG00000C | TBC1D22B | 3.26E-01 | 6.74E-01 | 1.05E+00 | 6.38E-02 | 1.05E+00 |
| ENSG000001 | TFB2M | 3.26E-01 | 6.74E-01 | 1.04E+00 | 6.02E-02 | 1.04E+00 |
| ENSG000001 | KDM1B | 3.26E-01 | 6.74E-01 | 1.03E+00 | 4.51E-02 | 1.03E+00 |
| ENSG000001 | KRT75 | 3.27E-01 | 6.74E-01 | 1.19E+00 | 2.55E-01 | 1.19E+00 |
| ENSG000001 | PI3 | 3.27E-01 | 6.74E-01 | 1.19E+00 | 2.49E-01 | 1.19E+00 |
| ENSG000001 | CARNS1 | 3.27E-01 | ? | 6.50E-01 | -6.22E-01 | -1.54E+00 |
| ENSG000001 | BRD1 | 3.27E-01 | 6.74E-01 | 1.04E+00 | 5.21E-02 | 1.04E+00 |
| ENSG000002 | --- | 3.27E-01 | ? | 6.94E-01 | -5.27E-01 | -1.44E+00 |
| ENSG000002 | --- | 3.27E-01 | ? | 7.49E-01 | -4.17E-01 | -1.33E+00 |
| ENSG000001 | CDC40 | 3.27E-01 | 6.75E-01 | 1.03E+00 | 4.32E-02 | 1.03E+00 |
| ENSG000001 | CLU | 3.27E-01 | 6.75E-01 | 9.43E-01 | -8.43E-02 | -1.06E+00 |
| ENSG000002 | --- | 3.27E-01 | ? | 7.67E-01 | -3.83E-01 | -1.30E+00 |

| | | | | | | | |
|---|---|---|---|---|---|---|---|
| ENSG000002 | --- | 3.27E-01 | | 6.75E-01 | 1.12E+00 | 1.65E-01 | 1.12E+00 |
| ENSG000000 | LHX5 | 3.27E-01 | ? | | 1.25E+00 | 3.27E-01 | 1.25E+00 |
| ENSG000001 | EXPH5 | 3.27E-01 | | 6.75E-01 | 1.09E+00 | 1.30E-01 | 1.09E+00 |
| ENSG000001 | NFATC2 | 3.27E-01 | ? | | 6.97E-01 | -5.21E-01 | -1.43E+00 |
| ENSG000001 | KCTD9P2 | 3.27E-01 | ? | | 6.07E-01 | -7.20E-01 | -1.65E+00 |
| ENSG000001 | TMUB2 | 3.27E-01 | | 6.75E-01 | 1.05E+00 | 6.83E-02 | 1.05E+00 |
| ENSG000002 | --- | 3.28E-01 | | 6.75E-01 | 1.22E+00 | 2.84E-01 | 1.22E+00 |
| ENSG000001 | TBC1D2B | 3.28E-01 | | 6.75E-01 | 1.03E+00 | 3.83E-02 | 1.03E+00 |
| ENSG000001 | ARHGEF3 | 3.28E-01 | | 6.75E-01 | 1.03E+00 | 4.09E-02 | 1.03E+00 |
| ENSG000001 | ZNF581 | 3.28E-01 | | 6.75E-01 | 1.09E+00 | 1.21E-01 | 1.09E+00 |
| ENSG000001 | NMI | 3.28E-01 | | 6.75E-01 | 1.07E+00 | 9.74E-02 | 1.07E+00 |
| ENSG000001 | PACSIN3 | 3.28E-01 | | 6.75E-01 | 1.04E+00 | 5.90E-02 | 1.04E+00 |
| ENSG000001 | COMTD1 | 3.28E-01 | | 6.75E-01 | 1.13E+00 | 1.71E-01 | 1.13E+00 |
| ENSG000002 | --- | 3.28E-01 | | 6.75E-01 | 9.28E-01 | -1.08E-01 | -1.08E+00 |
| ENSG000002 | A2M-AS1 | 3.28E-01 | ? | | 6.15E-01 | -7.00E-01 | -1.62E+00 |
| ENSG000000 | HDAC6 | 3.28E-01 | | 6.75E-01 | 1.04E+00 | 5.61E-02 | 1.04E+00 |
| ENSG000001 | ILF2 | 3.28E-01 | | 6.75E-01 | 1.03E+00 | 3.74E-02 | 1.03E+00 |
| ENSG000001 | DOK1 | 3.28E-01 | | 6.75E-01 | 1.13E+00 | 1.82E-01 | 1.13E+00 |
| ENSG000001 | CYP27A1 | 3.28E-01 | ? | | 1.28E+00 | 3.59E-01 | 1.28E+00 |
| ENSG000002 | --- | 3.28E-01 | | 6.75E-01 | 8.18E-01 | -2.90E-01 | -1.22E+00 |
| ENSG000002 | --- | 3.28E-01 | ? | | 1.30E+00 | 3.78E-01 | 1.30E+00 |
| ENSG000002 | --- | 3.28E-01 | ? | | 1.30E+00 | 3.78E-01 | 1.30E+00 |
| ENSG000001 | DPH5 | 3.28E-01 | | 6.75E-01 | 1.04E+00 | 5.29E-02 | 1.04E+00 |
| ENSG000002 | C8orf88 | 3.28E-01 | ? | | 1.35E+00 | 4.32E-01 | 1.35E+00 |
| ENSG000002 | LINC00607 | 3.28E-01 | ? | | 1.19E+00 | 2.50E-01 | 1.19E+00 |
| ENSG000001 | ABCA1 | 3.28E-01 | | 6.75E-01 | 9.29E-01 | -1.07E-01 | -1.08E+00 |
| ENSG000002 | --- | 3.28E-01 | | 6.75E-01 | 8.84E-01 | -1.78E-01 | -1.13E+00 |
| ENSG000002 | PLCG1-AS1 | 3.28E-01 | ? | | 8.37E-01 | -2.56E-01 | -1.19E+00 |
| ENSG000001 | C15orf62 | 3.28E-01 | | 6.75E-01 | 8.68E-01 | -2.04E-01 | -1.15E+00 |
| ENSG000000 | CACNA2D2 | 3.28E-01 | ? | | 6.72E-01 | -5.74E-01 | -1.49E+00 |
| ENSG000001 | RB1 | 3.28E-01 | | 6.75E-01 | 9.65E-01 | -5.08E-02 | -1.04E+00 |
| ENSG000002 | GUSBP5 | 3.29E-01 | ? | | 7.76E-01 | -3.65E-01 | -1.29E+00 |
| ENSG000001 | TLE3 | 3.29E-01 | | 6.76E-01 | 9.63E-01 | -5.44E-02 | -1.04E+00 |
| ENSG000002 | --- | 3.29E-01 | ? | | 6.77E-01 | -5.63E-01 | -1.48E+00 |
| ENSG000002 | LINC02175 | 3.29E-01 | | 6.76E-01 | 1.18E+00 | 2.38E-01 | 1.18E+00 |
| ENSG000002 | --- | 3.29E-01 | ? | | 1.21E+00 | 2.74E-01 | 1.21E+00 |
| ENSG000002 | STARD4-AS1 | 3.29E-01 | | 6.76E-01 | 9.07E-01 | -1.41E-01 | -1.10E+00 |
| ENSG000001 | ANO10 | 3.29E-01 | | 6.76E-01 | 1.03E+00 | 4.63E-02 | 1.03E+00 |
| ENSG000000 | CDC42 | 3.29E-01 | | 6.76E-01 | 1.05E+00 | 7.14E-02 | 1.05E+00 |
| ENSG000001 | HOPX | 3.29E-01 | | 6.76E-01 | 1.17E+00 | 2.31E-01 | 1.17E+00 |
| ENSG000002 | --- | 3.29E-01 | ? | | 7.66E-01 | -3.85E-01 | -1.31E+00 |
| ENSG000002 | HCG20 | 3.29E-01 | ? | | 7.03E-01 | -5.08E-01 | -1.42E+00 |
| ENSG000002 | MROH7-TTC4 | 3.29E-01 | ? | | 6.08E-01 | -7.18E-01 | -1.65E+00 |
| ENSG000001 | MVK | 3.29E-01 | | 6.77E-01 | 1.06E+00 | 8.08E-02 | 1.06E+00 |
| ENSG000002 | DSP-AS1 | 3.30E-01 | | 6.77E-01 | 1.11E+00 | 1.44E-01 | 1.11E+00 |

| | | | | | | | |
|---|---|---|---|---|---|---|---|
| ENSG000001 | ZKSCAN4 | 3.30E-01 | | 6.77E-01 | 1.06E+00 | 8.74E-02 | 1.06E+00 |
| ENSG000001 | UTP25 | 3.30E-01 | | 6.78E-01 | 9.58E-01 | -6.20E-02 | -1.04E+00 |
| ENSG000002 | H2AC8 | 3.30E-01 | | 6.78E-01 | 1.12E+00 | 1.65E-01 | 1.12E+00 |
| ENSG000002 | --- | 3.30E-01 | ? | | 7.06E-01 | -5.02E-01 | -1.42E+00 |
| ENSG000001 | CEP78 | 3.30E-01 | | 6.78E-01 | 9.58E-01 | -6.14E-02 | -1.04E+00 |
| ENSG000002 | FTLP14 | 3.30E-01 | ? | | 7.24E-01 | -4.66E-01 | -1.38E+00 |
| ENSG000001 | DHRS1 | 3.30E-01 | | 6.78E-01 | 1.05E+00 | 6.46E-02 | 1.05E+00 |
| ENSG000001 | MUC3A | 3.30E-01 | | 6.78E-01 | 9.11E-01 | -1.35E-01 | -1.10E+00 |
| ENSG000001 | NT5DC1 | 3.30E-01 | | 6.78E-01 | 1.04E+00 | 5.65E-02 | 1.04E+00 |
| ENSG000001 | ZNF574 | 3.30E-01 | | 6.78E-01 | 1.05E+00 | 7.67E-02 | 1.05E+00 |
| ENSG000001 | CHAD | 3.31E-01 | ? | | 1.17E+00 | 2.26E-01 | 1.17E+00 |
| ENSG00000C | HDAC4 | 3.31E-01 | | 6.78E-01 | 1.05E+00 | 6.51E-02 | 1.05E+00 |
| ENSG00000C | ADAM22 | 3.31E-01 | ? | | 1.27E+00 | 3.41E-01 | 1.27E+00 |
| ENSG000001 | ARL6IP6 | 3.31E-01 | | 6.78E-01 | 9.60E-01 | -5.94E-02 | -1.04E+00 |
| ENSG000001 | LYRM1 | 3.31E-01 | | 6.78E-01 | 1.05E+00 | 7.43E-02 | 1.05E+00 |
| ENSG000002 | --- | 3.31E-01 | ? | | 1.93E+00 | 9.48E-01 | 1.93E+00 |
| ENSG000001 | ZC3H8 | 3.31E-01 | | 6.78E-01 | 9.50E-01 | -7.43E-02 | -1.05E+00 |
| ENSG000001 | RLN2 | 3.31E-01 | ? | | 8.02E-01 | -3.19E-01 | -1.25E+00 |
| ENSG000001 | DAPK3 | 3.31E-01 | | 6.78E-01 | 1.05E+00 | 7.40E-02 | 1.05E+00 |
| ENSG000002 | DYNLL2 | 3.31E-01 | | 6.78E-01 | 9.68E-01 | -4.69E-02 | -1.03E+00 |
| ENSG000002 | --- | 3.31E-01 | ? | | 1.27E+00 | 3.47E-01 | 1.27E+00 |
| ENSG000001 | ALOXE3 | 3.31E-01 | | 6.78E-01 | 1.18E+00 | 2.40E-01 | 1.18E+00 |
| ENSG000002 | --- | 3.31E-01 | ? | | 1.24E+00 | 3.11E-01 | 1.24E+00 |
| ENSG00000C | VMP1 | 3.31E-01 | | 6.78E-01 | 9.57E-01 | -6.33E-02 | -1.04E+00 |
| ENSG000001 | RNF20 | 3.31E-01 | | 6.78E-01 | 1.04E+00 | 5.38E-02 | 1.04E+00 |
| ENSG000001 | ALOX5AP | 3.31E-01 | ? | | 1.18E+00 | 2.38E-01 | 1.18E+00 |
| ENSG000002 | TMEM185A | 3.31E-01 | | 6.79E-01 | 1.05E+00 | 7.26E-02 | 1.05E+00 |
| ENSG000001 | CEP126 | 3.31E-01 | | 6.79E-01 | 9.38E-01 | -9.18E-02 | -1.07E+00 |
| ENSG000001 | CCIN | 3.31E-01 | ? | | 7.89E-01 | -3.41E-01 | -1.27E+00 |
| ENSG000001 | EFNA3 | 3.31E-01 | | 6.79E-01 | 1.14E+00 | 1.90E-01 | 1.14E+00 |
| ENSG000001 | ABHD8 | 3.31E-01 | | 6.79E-01 | 1.11E+00 | 1.54E-01 | 1.11E+00 |
| ENSG000001 | INSC | 3.32E-01 | | 6.79E-01 | 1.13E+00 | 1.72E-01 | 1.13E+00 |
| ENSG000002 | P2RX5-TAX1I | 3.32E-01 | | 6.79E-01 | 8.26E-01 | -2.75E-01 | -1.21E+00 |
| ENSG000001 | HTR1D | 3.32E-01 | ? | | 1.29E+00 | 3.68E-01 | 1.29E+00 |
| ENSG000001 | JAM2 | 3.32E-01 | ? | | 1.41E+00 | 5.01E-01 | 1.41E+00 |
| ENSG00000C | EPB41L2 | 3.32E-01 | | 6.79E-01 | 9.42E-01 | -8.59E-02 | -1.06E+00 |
| ENSG000002 | KDM4A-AS1 | 3.32E-01 | | 6.79E-01 | 8.90E-01 | -1.69E-01 | -1.12E+00 |
| ENSG000001 | ZNF182 | 3.32E-01 | | 6.79E-01 | 9.51E-01 | -7.18E-02 | -1.05E+00 |
| ENSG000001 | IL1B | 3.32E-01 | | 6.79E-01 | 9.40E-01 | -8.92E-02 | -1.06E+00 |
| ENSG000001 | ING5 | 3.32E-01 | | 6.79E-01 | 1.05E+00 | 7.65E-02 | 1.05E+00 |
| ENSG000002 | --- | 3.32E-01 | | 6.79E-01 | 8.33E-01 | -2.63E-01 | -1.20E+00 |
| ENSG000001 | TRMT5 | 3.32E-01 | | 6.79E-01 | 1.03E+00 | 4.86E-02 | 1.03E+00 |
| ENSG000001 | TNFAIP2 | 3.32E-01 | | 6.79E-01 | 8.76E-01 | -1.91E-01 | -1.14E+00 |
| ENSG000001 | PTPN6 | 3.32E-01 | | 6.79E-01 | 1.04E+00 | 6.26E-02 | 1.04E+00 |
| ENSG000001 | ARRDC4 | 3.32E-01 | | 6.79E-01 | 9.68E-01 | -4.63E-02 | -1.03E+00 |

| | | | | | | |
|---|---|---|---|---|---|---|
| ENSG000001 | DMAP1 | 3.32E-01 | 6.79E-01 | 9.33E-01 | -9.95E-02 | -1.07E+00 |
| ENSG000002 | --- | 3.32E-01 | ? | 7.95E-01 | -3.30E-01 | -1.26E+00 |
| ENSG00000C | TCF7 | 3.32E-01 | 6.79E-01 | 8.99E-01 | -1.53E-01 | -1.11E+00 |
| ENSG000001 | TSC1 | 3.33E-01 | 6.79E-01 | 1.04E+00 | 5.45E-02 | 1.04E+00 |
| ENSG000002 | --- | 3.33E-01 | ? | 6.43E-01 | -6.37E-01 | -1.55E+00 |
| ENSG000001 | ZNF276 | 3.33E-01 | 6.80E-01 | 9.45E-01 | -8.13E-02 | -1.06E+00 |
| ENSG000002 | --- | 3.33E-01 | ? | 7.00E-01 | -5.14E-01 | -1.43E+00 |
| ENSG000001 | RAC3 | 3.33E-01 | 6.80E-01 | 1.09E+00 | 1.21E-01 | 1.09E+00 |
| ENSG000001 | TRABD | 3.33E-01 | 6.80E-01 | 1.09E+00 | 1.21E-01 | 1.09E+00 |
| ENSG000001 | POLR3C | 3.33E-01 | 6.80E-01 | 9.63E-01 | -5.41E-02 | -1.04E+00 |
| ENSG00000C | TMED1 | 3.33E-01 | 6.80E-01 | 1.07E+00 | 1.01E-01 | 1.07E+00 |
| ENSG000001 | WTAP | 3.33E-01 | 6.80E-01 | 9.75E-01 | -3.70E-02 | -1.03E+00 |
| ENSG000002 | --- | 3.33E-01 | ? | 8.18E-01 | -2.90E-01 | -1.22E+00 |
| ENSG000001 | POFUT2 | 3.33E-01 | 6.80E-01 | 9.62E-01 | -5.64E-02 | -1.04E+00 |
| ENSG000001 | CEP89 | 3.33E-01 | 6.80E-01 | 9.71E-01 | -4.25E-02 | -1.03E+00 |
| ENSG000001 | CHST1 | 3.33E-01 | ? | 1.32E+00 | 4.03E-01 | 1.32E+00 |
| ENSG000002 | --- | 3.33E-01 | ? | 7.74E-01 | -3.69E-01 | -1.29E+00 |
| ENSG000001 | LGALS9 | 3.33E-01 | ? | 7.65E-01 | -3.87E-01 | -1.31E+00 |
| ENSG00000C | POLB | 3.33E-01 | 6.80E-01 | 9.58E-01 | -6.14E-02 | -1.04E+00 |
| ENSG000002 | --- | 3.33E-01 | 6.80E-01 | 8.90E-01 | -1.68E-01 | -1.12E+00 |
| ENSG000001 | NHP2 | 3.33E-01 | 6.80E-01 | 1.03E+00 | 4.27E-02 | 1.03E+00 |
| ENSG000002 | LINC00504 | 3.33E-01 | ? | 1.31E+00 | 3.92E-01 | 1.31E+00 |
| ENSG000002 | LSP1P5 | 3.34E-01 | 6.80E-01 | 1.07E+00 | 9.76E-02 | 1.07E+00 |
| ENSG000001 | PMP22 | 3.34E-01 | 6.80E-01 | 8.61E-01 | -2.15E-01 | -1.16E+00 |
| ENSG000001 | RIOK3 | 3.34E-01 | 6.80E-01 | 9.69E-01 | -4.56E-02 | -1.03E+00 |
| ENSG000002 | --- | 3.34E-01 | 6.80E-01 | 8.92E-01 | -1.65E-01 | -1.12E+00 |
| ENSG000001 | PARL | 3.34E-01 | 6.80E-01 | 1.03E+00 | 4.90E-02 | 1.03E+00 |
| ENSG000001 | LZIC | 3.34E-01 | 6.80E-01 | 9.74E-01 | -3.78E-02 | -1.03E+00 |
| ENSG000002 | --- | 3.34E-01 | ? | 1.31E+00 | 3.89E-01 | 1.31E+00 |
| ENSG000002 | --- | 3.34E-01 | ? | 6.86E-01 | -5.43E-01 | -1.46E+00 |
| ENSG000001 | TMCO6 | 3.34E-01 | 6.81E-01 | 1.08E+00 | 1.07E-01 | 1.08E+00 |
| ENSG000001 | PHF11 | 3.34E-01 | 6.81E-01 | 9.53E-01 | -6.87E-02 | -1.05E+00 |
| ENSG000001 | LIN9 | 3.34E-01 | 6.81E-01 | 9.28E-01 | -1.09E-01 | -1.08E+00 |
| ENSG00000C | NUCKS1 | 3.34E-01 | 6.81E-01 | 1.03E+00 | 4.62E-02 | 1.03E+00 |
| ENSG000001 | SLC35E1 | 3.34E-01 | 6.81E-01 | 1.03E+00 | 4.41E-02 | 1.03E+00 |
| ENSG000001 | PLCZ1 | 3.34E-01 | ? | 5.58E-01 | -8.41E-01 | -1.79E+00 |
| ENSG000001 | NCF1 | 3.35E-01 | ? | 1.48E+00 | 5.69E-01 | 1.48E+00 |
| ENSG000001 | INPP5F | 3.35E-01 | 6.81E-01 | 9.69E-01 | -4.58E-02 | -1.03E+00 |
| ENSG000001 | GLYCTK | 3.35E-01 | 6.81E-01 | 1.12E+00 | 1.60E-01 | 1.12E+00 |
| ENSG000001 | SRGN | 3.35E-01 | 6.81E-01 | 8.14E-01 | -2.96E-01 | -1.23E+00 |
| ENSG000001 | SVIL | 3.35E-01 | 6.81E-01 | 1.03E+00 | 4.10E-02 | 1.03E+00 |
| ENSG000002 | LINC01173 | 3.35E-01 | ? | 6.40E-01 | -6.45E-01 | -1.56E+00 |
| ENSG000002 | EID1 | 3.35E-01 | 6.81E-01 | 1.03E+00 | 3.75E-02 | 1.03E+00 |
| ENSG000002 | --- | 3.35E-01 | 6.81E-01 | 1.07E+00 | 1.01E-01 | 1.07E+00 |
| ENSG000001 | ITGB4 | 3.35E-01 | 6.81E-01 | 9.54E-01 | -6.77E-02 | -1.05E+00 |

| | | | | | | |
|---|---|---|---|---|---|---|
| ENSG000001 | EOMES | 3.35E-01 | ? | 8.39E-01 | -2.53E-01 | -1.19E+00 |
| ENSG000001 | GPRC5B | 3.35E-01 | ? | 1.60E+00 | 6.82E-01 | 1.60E+00 |
| ENSG00000C | NCKAP1 | 3.35E-01 | 6.82E-01 | 9.77E-01 | -3.37E-02 | -1.02E+00 |
| ENSG000001 | CUTC | 3.35E-01 | 6.82E-01 | 1.05E+00 | 6.82E-02 | 1.05E+00 |
| ENSG000001 | SLC39A6 | 3.35E-01 | 6.82E-01 | 9.71E-01 | -4.25E-02 | -1.03E+00 |
| ENSG000001 | SNX8 | 3.35E-01 | 6.82E-01 | 1.08E+00 | 1.05E-01 | 1.08E+00 |
| ENSG000001 | ATP5F1A | 3.35E-01 | 6.82E-01 | 1.04E+00 | 5.46E-02 | 1.04E+00 |
| ENSG000002 | DNAJC19 | 3.35E-01 | 6.82E-01 | 1.06E+00 | 7.82E-02 | 1.06E+00 |
| ENSG000001 | PAK4 | 3.35E-01 | 6.82E-01 | 1.06E+00 | 8.51E-02 | 1.06E+00 |
| ENSG000002 | --- | 3.35E-01 | 6.82E-01 | 1.13E+00 | 1.74E-01 | 1.13E+00 |
| ENSG000001 | ZC2HC1C | 3.35E-01 | 6.82E-01 | 9.04E-01 | -1.46E-01 | -1.11E+00 |
| ENSG000001 | CCDC142 | 3.35E-01 | 6.82E-01 | 1.06E+00 | 8.19E-02 | 1.06E+00 |
| ENSG000002 | --- | 3.35E-01 | ? | 7.61E-01 | -3.93E-01 | -1.31E+00 |
| ENSG000001 | LYPD3 | 3.36E-01 | 6.82E-01 | 1.06E+00 | 8.35E-02 | 1.06E+00 |
| ENSG000002 | --- | 3.36E-01 | ? | 6.10E-01 | -7.12E-01 | -1.64E+00 |
| ENSG000001 | ZNF430 | 3.36E-01 | 6.82E-01 | 9.40E-01 | -8.93E-02 | -1.06E+00 |
| ENSG000001 | TRMT10C | 3.36E-01 | 6.82E-01 | 9.66E-01 | -5.01E-02 | -1.04E+00 |
| ENSG000001 | PHF14 | 3.36E-01 | 6.82E-01 | 9.58E-01 | -6.24E-02 | -1.04E+00 |
| ENSG000002 | --- | 3.36E-01 | 6.82E-01 | 1.16E+00 | 2.12E-01 | 1.16E+00 |
| ENSG000002 | RELA-DT | 3.36E-01 | ? | 1.16E+00 | 2.16E-01 | 1.16E+00 |
| ENSG000001 | C5orf24 | 3.36E-01 | 6.82E-01 | 9.54E-01 | -6.81E-02 | -1.05E+00 |
| ENSG000001 | RET | 3.36E-01 | 6.82E-01 | 1.08E+00 | 1.08E-01 | 1.08E+00 |
| ENSG000002 | RNASEH2B-A | 3.36E-01 | ? | 1.64E+00 | 7.15E-01 | 1.64E+00 |
| ENSG000001 | AGPAT2 | 3.36E-01 | 6.82E-01 | 1.07E+00 | 1.01E-01 | 1.07E+00 |
| ENSG000001 | PROX2 | 3.36E-01 | ? | 8.13E-01 | -2.99E-01 | -1.23E+00 |
| ENSG000002 | --- | 3.36E-01 | ? | 5.05E-01 | -9.87E-01 | -1.98E+00 |
| ENSG000001 | WDR19 | 3.36E-01 | 6.82E-01 | 9.56E-01 | -6.55E-02 | -1.05E+00 |
| ENSG000002 | PRAF2 | 3.36E-01 | 6.82E-01 | 1.05E+00 | 7.26E-02 | 1.05E+00 |
| ENSG000002 | --- | 3.36E-01 | 6.82E-01 | 1.09E+00 | 1.29E-01 | 1.09E+00 |
| ENSG00000C | SCAMP1 | 3.36E-01 | 6.82E-01 | 9.63E-01 | -5.49E-02 | -1.04E+00 |
| ENSG000002 | ZNF550 | 3.36E-01 | 6.82E-01 | 9.35E-01 | -9.63E-02 | -1.07E+00 |
| ENSG000002 | --- | 3.36E-01 | ? | 4.10E-01 | -1.29E+00 | -2.44E+00 |
| ENSG00000C | KARS1 | 3.36E-01 | 6.82E-01 | 9.76E-01 | -3.49E-02 | -1.02E+00 |
| ENSG000002 | CBLL1-AS1 | 3.36E-01 | 6.82E-01 | 8.93E-01 | -1.64E-01 | -1.12E+00 |
| ENSG000002 | TMEM249 | 3.36E-01 | ? | 7.95E-01 | -3.32E-01 | -1.26E+00 |
| ENSG000001 | EIF1AX | 3.36E-01 | 6.82E-01 | 9.77E-01 | -3.43E-02 | -1.02E+00 |
| ENSG000002 | MKRN2OS | 3.36E-01 | ? | 1.18E+00 | 2.35E-01 | 1.18E+00 |
| ENSG000001 | ANAPC13 | 3.36E-01 | 6.82E-01 | 9.72E-01 | -4.07E-02 | -1.03E+00 |
| ENSG000001 | BMP2K | 3.36E-01 | 6.82E-01 | 9.47E-01 | -7.78E-02 | -1.06E+00 |
| ENSG000001 | MAPKAPK3 | 3.37E-01 | 6.82E-01 | 1.04E+00 | 5.50E-02 | 1.04E+00 |
| ENSG000002 | ANKRD28 | 3.37E-01 | 6.82E-01 | 9.65E-01 | -5.11E-02 | -1.04E+00 |
| ENSG000001 | EXD3 | 3.37E-01 | 6.82E-01 | 1.05E+00 | 7.14E-02 | 1.05E+00 |
| ENSG000001 | ACVR1 | 3.37E-01 | 6.82E-01 | 1.03E+00 | 4.78E-02 | 1.03E+00 |
| ENSG000001 | GPD2 | 3.37E-01 | 6.82E-01 | 1.04E+00 | 5.47E-02 | 1.04E+00 |
| ENSG000001 | CHSY3 | 3.37E-01 | 6.82E-01 | 1.11E+00 | 1.47E-01 | 1.11E+00 |

| | | | | | | |
|---|---|---|---|---|---|---|
| ENSG000002 | --- | 3.37E-01 | ? | 7.26E-01 | -4.62E-01 | -1.38E+00 |
| ENSG000002 | RRP7BP | 3.37E-01 | 6.82E-01 | 9.28E-01 | -1.08E-01 | -1.08E+00 |
| ENSG000001 | SNRPB | 3.37E-01 | 6.82E-01 | 1.06E+00 | 8.75E-02 | 1.06E+00 |
| ENSG000002 | --- | 3.37E-01 | ? | 6.27E-01 | -6.73E-01 | -1.59E+00 |
| ENSG000001 | PXDNL | 3.37E-01 | ? | 7.36E-01 | -4.42E-01 | -1.36E+00 |
| ENSG000001 | NXT1 | 3.37E-01 | 6.82E-01 | 1.07E+00 | 9.69E-02 | 1.07E+00 |
| ENSG000002 | --- | 3.37E-01 | ? | 1.35E+00 | 4.28E-01 | 1.35E+00 |
| ENSG000002 | NPIPB11 | 3.37E-01 | 6.82E-01 | 9.03E-01 | -1.47E-01 | -1.11E+00 |
| ENSG00000C | SUSD2 | 3.37E-01 | ? | 8.25E-01 | -2.78E-01 | -1.21E+00 |
| ENSG000002 | JMJD1C-AS1 | 3.37E-01 | ? | 1.52E+00 | 6.09E-01 | 1.52E+00 |
| ENSG000002 | --- | 3.37E-01 | ? | 8.18E-01 | -2.90E-01 | -1.22E+00 |
| ENSG000001 | HSF4 | 3.37E-01 | 6.82E-01 | 1.16E+00 | 2.20E-01 | 1.16E+00 |
| ENSG000002 | --- | 3.37E-01 | ? | 8.37E-01 | -2.57E-01 | -1.20E+00 |
| ENSG00000C | CDC45 | 3.37E-01 | 6.82E-01 | 8.81E-01 | -1.83E-01 | -1.14E+00 |
| ENSG000001 | TOP2A | 3.37E-01 | 6.82E-01 | 8.73E-01 | -1.96E-01 | -1.15E+00 |
| ENSG000001 | RPS26 | 3.37E-01 | 6.82E-01 | 1.03E+00 | 4.54E-02 | 1.03E+00 |
| ENSG000002 | RPEP4 | 3.38E-01 | ? | 6.08E-01 | -7.18E-01 | -1.65E+00 |
| ENSG000002 | ANKRD18B | 3.38E-01 | 6.82E-01 | 9.40E-01 | -8.88E-02 | -1.06E+00 |
| ENSG000001 | TMC5 | 3.38E-01 | 6.82E-01 | 8.79E-01 | -1.85E-01 | -1.14E+00 |
| ENSG000002 | --- | 3.38E-01 | 6.82E-01 | 7.85E-01 | -3.48E-01 | -1.27E+00 |
| ENSG000001 | SYTL4 | 3.38E-01 | 6.82E-01 | 9.14E-01 | -1.29E-01 | -1.09E+00 |
| ENSG000001 | CIAO2B | 3.38E-01 | 6.82E-01 | 1.06E+00 | 9.01E-02 | 1.06E+00 |
| ENSG000001 | LGR5 | 3.38E-01 | ? | 1.25E+00 | 3.28E-01 | 1.25E+00 |
| ENSG000001 | GPR85 | 3.38E-01 | 6.82E-01 | 1.10E+00 | 1.31E-01 | 1.10E+00 |
| ENSG000002 | SH3BP5-AS1 | 3.38E-01 | 6.82E-01 | 9.16E-01 | -1.26E-01 | -1.09E+00 |
| ENSG000001 | RAET1L | 3.38E-01 | 6.82E-01 | 9.14E-01 | -1.29E-01 | -1.09E+00 |
| ENSG000001 | RNGTT | 3.38E-01 | 6.82E-01 | 9.68E-01 | -4.68E-02 | -1.03E+00 |
| ENSG000001 | ZNF114 | 3.38E-01 | ? | 8.07E-01 | -3.10E-01 | -1.24E+00 |
| ENSG000001 | TAGLN3 | 3.38E-01 | 6.82E-01 | 9.15E-01 | -1.29E-01 | -1.09E+00 |
| ENSG000001 | WNT5B | 3.38E-01 | 6.82E-01 | 1.09E+00 | 1.29E-01 | 1.09E+00 |
| ENSG000002 | HSPE1-MOB4 | 3.38E-01 | 6.82E-01 | 1.09E+00 | 1.21E-01 | 1.09E+00 |
| ENSG00000C | KIF26A | 3.38E-01 | ? | 8.39E-01 | -2.53E-01 | -1.19E+00 |
| ENSG000002 | --- | 3.38E-01 | 6.82E-01 | 8.74E-01 | -1.94E-01 | -1.14E+00 |
| ENSG000001 | ENO2 | 3.38E-01 | 6.82E-01 | 9.36E-01 | -9.60E-02 | -1.07E+00 |
| ENSG000002 | --- | 3.38E-01 | 6.83E-01 | 9.00E-01 | -1.53E-01 | -1.11E+00 |
| ENSG000001 | ADAMTS9 | 3.38E-01 | ? | 8.50E-01 | -2.34E-01 | -1.18E+00 |
| ENSG000002 | --- | 3.38E-01 | 6.83E-01 | 8.52E-01 | -2.32E-01 | -1.17E+00 |
| ENSG000001 | STK26 | 3.38E-01 | 6.83E-01 | 1.04E+00 | 5.24E-02 | 1.04E+00 |
| ENSG000002 | PEDS1 | 3.38E-01 | 6.83E-01 | 1.03E+00 | 3.97E-02 | 1.03E+00 |
| ENSG000001 | TUBG1 | 3.38E-01 | 6.83E-01 | 1.03E+00 | 4.36E-02 | 1.03E+00 |
| ENSG000002 | C10orf143 | 3.38E-01 | ? | 1.19E+00 | 2.48E-01 | 1.19E+00 |
| ENSG00000C | PKN2 | 3.38E-01 | 6.83E-01 | 9.72E-01 | -4.08E-02 | -1.03E+00 |
| ENSG000001 | ADARB1 | 3.39E-01 | 6.83E-01 | 1.05E+00 | 6.36E-02 | 1.05E+00 |
| ENSG000001 | SH3BP1 | 3.39E-01 | 6.83E-01 | 1.06E+00 | 7.94E-02 | 1.06E+00 |
| ENSG000001 | IRX1 | 3.39E-01 | ? | 1.17E+00 | 2.32E-01 | 1.17E+00 |

| | | | | | | |
|---|---|---|---|---|---|---|
| ENSG000002 | PCDHGB9P | 3.39E-01 | ? | 1.53E+00 | 6.16E-01 | 1.53E+00 |
| ENSG000001 | TSC22D4 | 3.39E-01 | 6.83E-01 | 1.08E+00 | 1.13E-01 | 1.08E+00 |
| ENSG000002 | URAHP | 3.39E-01 | ? | 1.24E+00 | 3.16E-01 | 1.24E+00 |
| ENSG000001 | ABCF3 | 3.39E-01 | 6.83E-01 | 1.03E+00 | 4.72E-02 | 1.03E+00 |
| ENSG00000C | CLEC2D | 3.39E-01 | 6.83E-01 | 8.47E-01 | -2.39E-01 | -1.18E+00 |
| ENSG000002 | MYG1-AS1 | 3.39E-01 | ? | 1.21E+00 | 2.72E-01 | 1.21E+00 |
| ENSG000001 | URB2 | 3.39E-01 | 6.83E-01 | 1.04E+00 | 5.43E-02 | 1.04E+00 |
| ENSG000001 | NUTF2 | 3.39E-01 | 6.83E-01 | 9.67E-01 | -4.87E-02 | -1.03E+00 |
| ENSG00000C | SPAG5 | 3.39E-01 | 6.83E-01 | 9.27E-01 | -1.10E-01 | -1.08E+00 |
| ENSG000001 | NIT1 | 3.39E-01 | 6.83E-01 | 1.05E+00 | 7.13E-02 | 1.05E+00 |
| ENSG00000C | PAF1 | 3.39E-01 | 6.83E-01 | 9.62E-01 | -5.53E-02 | -1.04E+00 |
| ENSG00000C | RETSAT | 3.39E-01 | 6.84E-01 | 1.04E+00 | 5.60E-02 | 1.04E+00 |
| ENSG000002 | --- | 3.39E-01 | ? | 6.56E-01 | -6.08E-01 | -1.52E+00 |
| ENSG000001 | SNX19 | 3.39E-01 | 6.84E-01 | 9.75E-01 | -3.68E-02 | -1.03E+00 |
| ENSG000002 | --- | 3.40E-01 | ? | 6.35E-01 | -6.56E-01 | -1.58E+00 |
| ENSG000001 | GPRC5C | 3.40E-01 | 6.84E-01 | 1.08E+00 | 1.06E-01 | 1.08E+00 |
| ENSG000002 | PWP2 | 3.40E-01 | 6.84E-01 | 1.05E+00 | 7.18E-02 | 1.05E+00 |
| ENSG000002 | CTAGE6 | 3.40E-01 | ? | 7.54E-01 | -4.08E-01 | -1.33E+00 |
| ENSG000001 | ZFP62 | 3.40E-01 | 6.84E-01 | 1.04E+00 | 5.63E-02 | 1.04E+00 |
| ENSG000001 | CERK | 3.40E-01 | 6.84E-01 | 1.04E+00 | 5.71E-02 | 1.04E+00 |
| ENSG000001 | KHDRBS1 | 3.40E-01 | 6.84E-01 | 1.02E+00 | 2.73E-02 | 1.02E+00 |
| ENSG000002 | LINC02983 | 3.40E-01 | ? | 7.38E-01 | -4.39E-01 | -1.36E+00 |
| ENSG000002 | --- | 3.40E-01 | ? | 7.91E-01 | -3.38E-01 | -1.26E+00 |
| ENSG000002 | DNAJC9-AS1 | 3.40E-01 | ? | 1.26E+00 | 3.29E-01 | 1.26E+00 |
| ENSG000002 | --- | 3.40E-01 | ? | 1.33E+00 | 4.06E-01 | 1.33E+00 |
| ENSG000001 | ZNF48 | 3.40E-01 | 6.85E-01 | 1.06E+00 | 8.41E-02 | 1.06E+00 |
| ENSG000001 | S100A11 | 3.40E-01 | 6.85E-01 | 1.04E+00 | 5.34E-02 | 1.04E+00 |
| ENSG000001 | KLHL15 | 3.40E-01 | 6.85E-01 | 1.06E+00 | 8.56E-02 | 1.06E+00 |
| ENSG000002 | PHACTR2P1 | 3.40E-01 | ? | 6.78E-01 | -5.61E-01 | -1.48E+00 |
| ENSG000002 | --- | 3.40E-01 | ? | 7.96E-01 | -3.29E-01 | -1.26E+00 |
| ENSG000001 | MTG1 | 3.41E-01 | 6.85E-01 | 1.07E+00 | 9.18E-02 | 1.07E+00 |
| ENSG000002 | TMEM183BP | 3.41E-01 | 6.85E-01 | 1.17E+00 | 2.32E-01 | 1.17E+00 |
| ENSG000002 | --- | 3.41E-01 | ? | 1.44E+00 | 5.22E-01 | 1.44E+00 |
| ENSG000001 | PAXX | 3.41E-01 | 6.85E-01 | 1.13E+00 | 1.80E-01 | 1.13E+00 |
| ENSG000001 | PUS7L | 3.41E-01 | 6.85E-01 | 9.48E-01 | -7.72E-02 | -1.05E+00 |
| ENSG000001 | DOCK8 | 3.41E-01 | 6.85E-01 | 1.06E+00 | 8.27E-02 | 1.06E+00 |
| ENSG000001 | CWC27 | 3.41E-01 | 6.85E-01 | 9.52E-01 | -7.04E-02 | -1.05E+00 |
| ENSG000001 | ATP8B3 | 3.41E-01 | 6.85E-01 | 9.26E-01 | -1.12E-01 | -1.08E+00 |
| ENSG00000C | TGM1 | 3.41E-01 | 6.85E-01 | 1.08E+00 | 1.12E-01 | 1.08E+00 |
| ENSG000001 | CCZ1B | 3.41E-01 | 6.85E-01 | 9.71E-01 | -4.28E-02 | -1.03E+00 |
| ENSG000002 | --- | 3.41E-01 | ? | 1.38E+00 | 4.66E-01 | 1.38E+00 |
| ENSG000001 | TSSC4 | 3.41E-01 | 6.85E-01 | 1.09E+00 | 1.29E-01 | 1.09E+00 |
| ENSG000001 | QSOX2 | 3.41E-01 | 6.85E-01 | 1.05E+00 | 7.06E-02 | 1.05E+00 |
| ENSG000001 | NSMCE2 | 3.41E-01 | 6.85E-01 | 9.55E-01 | -6.67E-02 | -1.05E+00 |
| ENSG000001 | APC | 3.41E-01 | 6.85E-01 | 9.51E-01 | -7.28E-02 | -1.05E+00 |

| | | | | | | |
|---|---|---|---|---|---|---|
| ENSG000001 | MRPL13 | 3.41E-01 | 6.85E-01 | 1.04E+00 | 5.64E-02 | 1.04E+00 |
| ENSG000002 | PLS3-AS1 | 3.41E-01 | ? | 1.35E+00 | 4.34E-01 | 1.35E+00 |
| ENSG000002 | --- | 3.41E-01 | ? | 1.72E+00 | 7.84E-01 | 1.72E+00 |
| ENSG000002 | LINC01226 | 3.41E-01 | ? | 1.57E+00 | 6.54E-01 | 1.57E+00 |
| ENSG000002 | MARS2 | 3.41E-01 | 6.85E-01 | 1.06E+00 | 8.20E-02 | 1.06E+00 |
| ENSG000002 | --- | 3.41E-01 | ? | 5.88E-01 | -7.66E-01 | -1.70E+00 |
| ENSG00000C | USP36 | 3.42E-01 | 6.85E-01 | 9.76E-01 | -3.52E-02 | -1.02E+00 |
| ENSG000001 | RHD | 3.42E-01 | ? | 1.51E+00 | 5.96E-01 | 1.51E+00 |
| ENSG000002 | EFCAB6-DT | 3.42E-01 | ? | 1.44E+00 | 5.23E-01 | 1.44E+00 |
| ENSG000001 | FBLN7 | 3.42E-01 | 6.85E-01 | 1.14E+00 | 1.87E-01 | 1.14E+00 |
| ENSG000001 | PTPRB | 3.42E-01 | 6.86E-01 | 9.09E-01 | -1.37E-01 | -1.10E+00 |
| ENSG000002 | YWHAZP10 | 3.42E-01 | ? | 1.36E+00 | 4.45E-01 | 1.36E+00 |
| ENSG000001 | CLBA1 | 3.42E-01 | 6.86E-01 | 1.07E+00 | 1.00E-01 | 1.07E+00 |
| ENSG000002 | --- | 3.42E-01 | ? | 8.31E-01 | -2.67E-01 | -1.20E+00 |
| ENSG000002 | RBM26-AS1 | 3.42E-01 | 6.86E-01 | 1.09E+00 | 1.26E-01 | 1.09E+00 |
| ENSG000002 | --- | 3.42E-01 | ? | 1.21E+00 | 2.71E-01 | 1.21E+00 |
| ENSG000001 | ZNF514 | 3.42E-01 | 6.86E-01 | 9.36E-01 | -9.48E-02 | -1.07E+00 |
| ENSG000001 | C11orf58 | 3.42E-01 | 6.86E-01 | 9.75E-01 | -3.66E-02 | -1.03E+00 |
| ENSG000002 | --- | 3.42E-01 | ? | 7.08E-01 | -4.98E-01 | -1.41E+00 |
| ENSG000001 | ZNF429 | 3.42E-01 | 6.86E-01 | 9.49E-01 | -7.62E-02 | -1.05E+00 |
| ENSG000001 | PCSK7 | 3.42E-01 | 6.86E-01 | 9.65E-01 | -5.20E-02 | -1.04E+00 |
| ENSG000001 | ZSWIM4 | 3.43E-01 | 6.86E-01 | 9.18E-01 | -1.23E-01 | -1.09E+00 |
| ENSG000001 | ZSWIM9 | 3.43E-01 | 6.86E-01 | 9.30E-01 | -1.04E-01 | -1.08E+00 |
| ENSG000001 | SHMT2 | 3.43E-01 | 6.87E-01 | 1.05E+00 | 6.71E-02 | 1.05E+00 |
| ENSG000001 | PSG9 | 3.43E-01 | ? | 7.32E-01 | -4.49E-01 | -1.37E+00 |
| ENSG000001 | MZT2B | 3.43E-01 | 6.87E-01 | 1.13E+00 | 1.75E-01 | 1.13E+00 |
| ENSG000001 | CYP20A1 | 3.43E-01 | 6.87E-01 | 1.04E+00 | 5.92E-02 | 1.04E+00 |
| ENSG000001 | NAP1L2 | 3.43E-01 | 6.87E-01 | 1.07E+00 | 1.02E-01 | 1.07E+00 |
| ENSG000002 | LINC01303 | 3.43E-01 | 6.87E-01 | 8.73E-01 | -1.96E-01 | -1.15E+00 |
| ENSG000002 | DPY19L1P1 | 3.43E-01 | ? | 1.40E+00 | 4.88E-01 | 1.40E+00 |
| ENSG000001 | KLB | 3.43E-01 | ? | 1.28E+00 | 3.53E-01 | 1.28E+00 |
| ENSG000001 | C2CD2 | 3.43E-01 | 6.87E-01 | 1.04E+00 | 5.48E-02 | 1.04E+00 |
| ENSG000001 | BNIP1 | 3.43E-01 | 6.87E-01 | 1.05E+00 | 7.57E-02 | 1.05E+00 |
| ENSG000002 | FTH1P23 | 3.43E-01 | ? | 1.42E+00 | 5.05E-01 | 1.42E+00 |
| ENSG000001 | C2CD4A | 3.43E-01 | ? | 7.10E-01 | -4.95E-01 | -1.41E+00 |
| ENSG000002 | L3MBTL2-AS | 3.43E-01 | 6.88E-01 | 8.93E-01 | -1.64E-01 | -1.12E+00 |
| ENSG00000C | EFNA2 | 3.44E-01 | ? | 6.30E-01 | -6.66E-01 | -1.59E+00 |
| ENSG000002 | KRT8P14 | 3.44E-01 | ? | 8.14E-01 | -2.97E-01 | -1.23E+00 |
| ENSG00000C | NDUFAB1 | 3.44E-01 | 6.88E-01 | 1.05E+00 | 6.51E-02 | 1.05E+00 |
| ENSG000001 | RABL2A | 3.44E-01 | 6.88E-01 | 1.09E+00 | 1.26E-01 | 1.09E+00 |
| ENSG000001 | SHC1 | 3.44E-01 | 6.88E-01 | 1.04E+00 | 6.07E-02 | 1.04E+00 |
| ENSG000002 | PGAM5 | 3.44E-01 | 6.88E-01 | 1.05E+00 | 6.52E-02 | 1.05E+00 |
| ENSG000002 | --- | 3.44E-01 | 6.88E-01 | 9.15E-01 | -1.29E-01 | -1.09E+00 |
| ENSG000001 | RPS19BP1 | 3.44E-01 | 6.88E-01 | 1.07E+00 | 9.38E-02 | 1.07E+00 |
| ENSG000001 | PTGER2 | 3.44E-01 | ? | 1.35E+00 | 4.30E-01 | 1.35E+00 |

| | | | | | | |
|---|---|---|---|---|---|---|
| ENSG000001 | CCNB1IP1 | 3.44E-01 | 6.88E-01 | 1.04E+00 | 6.13E-02 | 1.04E+00 |
| ENSG000001 | PHKB | 3.44E-01 | 6.88E-01 | 1.03E+00 | 4.18E-02 | 1.03E+00 |
| ENSG000002 | FGF14-AS2 | 3.44E-01 | ? | 1.40E+00 | 4.88E-01 | 1.40E+00 |
| ENSG00000C | TMEM260 | 3.44E-01 | 6.88E-01 | 1.03E+00 | 4.84E-02 | 1.03E+00 |
| ENSG000001 | NEMF | 3.44E-01 | 6.88E-01 | 9.61E-01 | -5.68E-02 | -1.04E+00 |
| ENSG000002 | IDI2-AS1 | 3.44E-01 | 6.88E-01 | 8.51E-01 | -2.33E-01 | -1.17E+00 |
| ENSG000001 | ANKAR | 3.44E-01 | 6.88E-01 | 8.65E-01 | -2.10E-01 | -1.16E+00 |
| ENSG000002 | FCHO2-DT | 3.44E-01 | ? | 1.39E+00 | 4.75E-01 | 1.39E+00 |
| ENSG000002 | CCNI2 | 3.44E-01 | ? | 1.20E+00 | 2.63E-01 | 1.20E+00 |
| ENSG000001 | TUBBP5 | 3.44E-01 | ? | 2.87E-01 | -1.80E+00 | -3.49E+00 |
| ENSG000001 | ALDH1A3 | 3.45E-01 | 6.88E-01 | 9.47E-01 | -7.90E-02 | -1.06E+00 |
| ENSG000001 | UNC5C | 3.45E-01 | ? | 7.73E-01 | -3.72E-01 | -1.29E+00 |
| ENSG000002 | ZNF595 | 3.45E-01 | 6.89E-01 | 1.06E+00 | 8.82E-02 | 1.06E+00 |
| ENSG000001 | HMGB1 | 3.45E-01 | 6.89E-01 | 9.55E-01 | -6.63E-02 | -1.05E+00 |
| ENSG000001 | BVES | 3.45E-01 | 6.89E-01 | 1.06E+00 | 8.68E-02 | 1.06E+00 |
| ENSG000001 | F2 | 3.45E-01 | ? | 7.49E-01 | -4.17E-01 | -1.33E+00 |
| ENSG000001 | TTC23 | 3.45E-01 | 6.89E-01 | 9.52E-01 | -7.16E-02 | -1.05E+00 |
| ENSG000001 | ELOF1 | 3.45E-01 | 6.89E-01 | 1.05E+00 | 7.44E-02 | 1.05E+00 |
| ENSG000002 | --- | 3.45E-01 | ? | 1.44E+00 | 5.27E-01 | 1.44E+00 |
| ENSG000002 | HHIP-AS1 | 3.45E-01 | ? | 6.57E-01 | -6.07E-01 | -1.52E+00 |
| ENSG000001 | C1GALT1C1 | 3.45E-01 | 6.89E-01 | 1.04E+00 | 5.71E-02 | 1.04E+00 |
| ENSG000001 | FIGN | 3.45E-01 | 6.89E-01 | 9.40E-01 | -8.95E-02 | -1.06E+00 |
| ENSG000002 | --- | 3.45E-01 | ? | 1.48E+00 | 5.65E-01 | 1.48E+00 |
| ENSG000002 | RAB44 | 3.45E-01 | ? | 7.22E-01 | -4.71E-01 | -1.39E+00 |
| ENSG000001 | HDLBP | 3.45E-01 | 6.89E-01 | 9.78E-01 | -3.28E-02 | -1.02E+00 |
| ENSG000001 | MRPL21 | 3.45E-01 | 6.89E-01 | 1.04E+00 | 5.67E-02 | 1.04E+00 |
| ENSG000002 | --- | 3.45E-01 | ? | 8.07E-01 | -3.09E-01 | -1.24E+00 |
| ENSG000001 | TMEM266 | 3.45E-01 | ? | 1.33E+00 | 4.12E-01 | 1.33E+00 |
| ENSG000001 | CARD8 | 3.45E-01 | 6.89E-01 | 9.47E-01 | -7.92E-02 | -1.06E+00 |
| ENSG00000C | HFE | 3.46E-01 | 6.89E-01 | 9.44E-01 | -8.27E-02 | -1.06E+00 |
| ENSG000001 | XPOT | 3.46E-01 | 6.89E-01 | 9.66E-01 | -5.04E-02 | -1.04E+00 |
| ENSG000001 | CHCHD6 | 3.46E-01 | 6.89E-01 | 1.07E+00 | 9.28E-02 | 1.07E+00 |
| ENSG000001 | PLEKHF2 | 3.46E-01 | 6.89E-01 | 1.04E+00 | 6.21E-02 | 1.04E+00 |
| ENSG000002 | --- | 3.46E-01 | ? | 6.68E-01 | -5.81E-01 | -1.50E+00 |
| ENSG000002 | PELATON | 3.46E-01 | ? | 5.93E-01 | -7.54E-01 | -1.69E+00 |
| ENSG000001 | ZNF599 | 3.46E-01 | 6.90E-01 | 9.09E-01 | -1.37E-01 | -1.10E+00 |
| ENSG00000C | PRKACA | 3.46E-01 | 6.90E-01 | 1.03E+00 | 3.82E-02 | 1.03E+00 |
| ENSG000001 | ATG4C | 3.46E-01 | 6.90E-01 | 9.47E-01 | -7.82E-02 | -1.06E+00 |
| ENSG000002 | --- | 3.46E-01 | ? | 1.26E+00 | 3.38E-01 | 1.26E+00 |
| ENSG000002 | RN7SL689P | 3.46E-01 | ? | 1.50E+00 | 5.86E-01 | 1.50E+00 |
| ENSG000002 | --- | 3.46E-01 | ? | 1.50E+00 | 5.87E-01 | 1.50E+00 |
| ENSG000001 | SPATA6L | 3.46E-01 | 6.90E-01 | 8.73E-01 | -1.96E-01 | -1.15E+00 |
| ENSG00000C | DGAT2 | 3.46E-01 | 6.90E-01 | 1.07E+00 | 9.71E-02 | 1.07E+00 |
| ENSG000001 | KDM4D | 3.46E-01 | 6.90E-01 | 8.70E-01 | -2.01E-01 | -1.15E+00 |
| ENSG000002 | --- | 3.46E-01 | ? | 1.45E+00 | 5.36E-01 | 1.45E+00 |

| | | | | | | |
|---|---|---|---|---|---|---|
| ENSG000001 | HK2 | 3.47E-01 | 6.90E-01 | 1.03E+00 | 3.87E-02 | 1.03E+00 |
| ENSG000001 | BLOC1S1 | 3.47E-01 | 6.90E-01 | 1.06E+00 | 7.77E-02 | 1.06E+00 |
| ENSG000001 | MISP3 | 3.47E-01 | 6.90E-01 | 1.14E+00 | 1.86E-01 | 1.14E+00 |
| ENSG000002 | --- | 3.47E-01 | ? | 8.12E-01 | -3.01E-01 | -1.23E+00 |
| ENSG000001 | OGA | 3.47E-01 | 6.90E-01 | 9.69E-01 | -4.60E-02 | -1.03E+00 |
| ENSG00000C | EZR | 3.47E-01 | 6.90E-01 | 1.03E+00 | 4.72E-02 | 1.03E+00 |
| ENSG000002 | PRC1-AS1 | 3.47E-01 | ? | 7.82E-01 | -3.54E-01 | -1.28E+00 |
| ENSG000001 | MSTO1 | 3.47E-01 | 6.90E-01 | 9.58E-01 | -6.12E-02 | -1.04E+00 |
| ENSG000001 | TMSB15B | 3.47E-01 | 6.90E-01 | 1.13E+00 | 1.79E-01 | 1.13E+00 |
| ENSG000001 | PTX3 | 3.47E-01 | 6.90E-01 | 1.18E+00 | 2.43E-01 | 1.18E+00 |
| ENSG000002 | NUTM2E | 3.47E-01 | 6.90E-01 | 1.10E+00 | 1.36E-01 | 1.10E+00 |
| ENSG000002 | LDHAP4 | 3.47E-01 | 6.90E-01 | 1.06E+00 | 8.00E-02 | 1.06E+00 |
| ENSG000001 | BPTF | 3.47E-01 | 6.90E-01 | 9.54E-01 | -6.74E-02 | -1.05E+00 |
| ENSG000001 | RAB8A | 3.47E-01 | 6.90E-01 | 9.61E-01 | -5.76E-02 | -1.04E+00 |
| ENSG000001 | BMAL1 | 3.47E-01 | 6.90E-01 | 1.05E+00 | 7.25E-02 | 1.05E+00 |
| ENSG000001 | GTF2H1 | 3.47E-01 | 6.90E-01 | 1.03E+00 | 4.25E-02 | 1.03E+00 |
| ENSG000001 | NHLH2 | 3.47E-01 | 6.90E-01 | 9.30E-01 | -1.05E-01 | -1.08E+00 |
| ENSG000002 | --- | 3.47E-01 | ? | 1.69E+00 | 7.53E-01 | 1.69E+00 |
| ENSG000001 | ZNF772 | 3.47E-01 | 6.90E-01 | 1.04E+00 | 5.87E-02 | 1.04E+00 |
| ENSG000001 | CHRM3 | 3.47E-01 | ? | 7.85E-01 | -3.50E-01 | -1.27E+00 |
| ENSG000002 | DNAAF3-AS1 | 3.47E-01 | 6.90E-01 | 8.81E-01 | -1.83E-01 | -1.14E+00 |
| ENSG000001 | CAMK2G | 3.47E-01 | 6.90E-01 | 1.04E+00 | 5.28E-02 | 1.04E+00 |
| ENSG000001 | APC2 | 3.47E-01 | 6.90E-01 | 8.83E-01 | -1.79E-01 | -1.13E+00 |
| ENSG000002 | ATP6V0E2-A: | 3.47E-01 | ? | 1.23E+00 | 2.95E-01 | 1.23E+00 |
| ENSG000001 | CNTN3 | 3.48E-01 | 6.91E-01 | 1.08E+00 | 1.10E-01 | 1.08E+00 |
| ENSG000002 | ZHX1-C8orf7( | 3.48E-01 | ? | 1.29E+00 | 3.70E-01 | 1.29E+00 |
| ENSG000001 | PTS | 3.48E-01 | 6.91E-01 | 9.62E-01 | -5.54E-02 | -1.04E+00 |
| ENSG000001 | TFR2 | 3.48E-01 | ? | 1.47E+00 | 5.52E-01 | 1.47E+00 |
| ENSG00000C | TINF2 | 3.48E-01 | 6.91E-01 | 1.05E+00 | 6.42E-02 | 1.05E+00 |
| ENSG000001 | KANK1 | 3.48E-01 | 6.91E-01 | 1.03E+00 | 3.75E-02 | 1.03E+00 |
| ENSG00000C | MAP3K4 | 3.48E-01 | 6.91E-01 | 9.64E-01 | -5.29E-02 | -1.04E+00 |
| ENSG000001 | DYRK1A | 3.48E-01 | 6.91E-01 | 9.63E-01 | -5.44E-02 | -1.04E+00 |
| ENSG000001 | ZDHHC3 | 3.48E-01 | 6.91E-01 | 1.02E+00 | 2.88E-02 | 1.02E+00 |
| ENSG000001 | FAM87B | 3.48E-01 | ? | 1.46E+00 | 5.45E-01 | 1.46E+00 |
| ENSG000001 | DLX4 | 3.48E-01 | ? | 1.38E+00 | 4.67E-01 | 1.38E+00 |
| ENSG000001 | SOD3 | 3.48E-01 | ? | 6.41E-01 | -6.43E-01 | -1.56E+00 |
| ENSG00000C | DHX33 | 3.48E-01 | 6.91E-01 | 1.03E+00 | 3.80E-02 | 1.03E+00 |
| ENSG000002 | --- | 3.48E-01 | ? | 7.86E-01 | -3.48E-01 | -1.27E+00 |
| ENSG000001 | GARIN5A | 3.48E-01 | 6.91E-01 | 1.17E+00 | 2.23E-01 | 1.17E+00 |
| ENSG000001 | RPP25 | 3.48E-01 | 6.91E-01 | 1.08E+00 | 1.06E-01 | 1.08E+00 |
| ENSG000001 | ANKRD42 | 3.48E-01 | 6.91E-01 | 9.61E-01 | -5.74E-02 | -1.04E+00 |
| ENSG00000C | TSPAN9 | 3.49E-01 | 6.91E-01 | 1.06E+00 | 8.05E-02 | 1.06E+00 |
| ENSG000001 | TIMP1 | 3.49E-01 | 6.91E-01 | 1.06E+00 | 8.99E-02 | 1.06E+00 |
| ENSG000001 | PTPRZ1 | 3.49E-01 | 6.91E-01 | 8.77E-01 | -1.89E-01 | -1.14E+00 |
| ENSG000001 | FANCI | 3.49E-01 | 6.91E-01 | 9.49E-01 | -7.61E-02 | -1.05E+00 |

| | | | | | | |
|---|---|---|---|---|---|---|
| ENSG000002 | --- | 3.49E-01 | ? | 5.90E-01 | -7.61E-01 | -1.70E+00 |
| ENSG000002 | --- | 3.49E-01 | ? | 5.73E-01 | -8.04E-01 | -1.75E+00 |
| ENSG000000 | WAS | 3.49E-01 | ? | 6.94E-01 | -5.27E-01 | -1.44E+00 |
| ENSG000002 | PPIEL | 3.49E-01 | 6.92E-01 | 9.04E-01 | -1.45E-01 | -1.11E+00 |
| ENSG000001 | GPR156 | 3.49E-01 | ? | 1.27E+00 | 3.49E-01 | 1.27E+00 |
| ENSG000001 | NFIL3 | 3.49E-01 | 6.92E-01 | 1.06E+00 | 8.41E-02 | 1.06E+00 |
| ENSG000000 | DOCK3 | 3.49E-01 | ? | 1.16E+00 | 2.13E-01 | 1.16E+00 |
| ENSG000002 | --- | 3.49E-01 | ? | 6.04E-01 | -7.27E-01 | -1.65E+00 |
| ENSG000001 | CCS | 3.49E-01 | 6.93E-01 | 1.06E+00 | 8.27E-02 | 1.06E+00 |
| ENSG000002 | --- | 3.49E-01 | ? | 1.51E+00 | 5.95E-01 | 1.51E+00 |
| ENSG000002 | --- | 3.49E-01 | ? | 7.96E-01 | -3.30E-01 | -1.26E+00 |
| ENSG000002 | EIF4EBP3 | 3.50E-01 | ? | 1.34E+00 | 4.18E-01 | 1.34E+00 |
| ENSG000002 | YBX3P1 | 3.50E-01 | ? | 1.59E+00 | 6.65E-01 | 1.59E+00 |
| ENSG000001 | ZNF354A | 3.50E-01 | 6.93E-01 | 9.52E-01 | -7.10E-02 | -1.05E+00 |
| ENSG000001 | FBXO39 | 3.50E-01 | ? | 8.02E-01 | -3.18E-01 | -1.25E+00 |
| ENSG000001 | RND3 | 3.50E-01 | 6.93E-01 | 1.04E+00 | 6.07E-02 | 1.04E+00 |
| ENSG000002 | LINC02938 | 3.50E-01 | ? | 6.86E-01 | -5.44E-01 | -1.46E+00 |
| ENSG000001 | TMEM131L | 3.50E-01 | 6.93E-01 | 1.04E+00 | 5.16E-02 | 1.04E+00 |
| ENSG000001 | EAPP | 3.50E-01 | 6.93E-01 | 9.60E-01 | -5.86E-02 | -1.04E+00 |
| ENSG000000 | B4GALT1 | 3.50E-01 | 6.93E-01 | 9.27E-01 | -1.09E-01 | -1.08E+00 |
| ENSG000001 | AP3S1 | 3.50E-01 | 6.93E-01 | 9.70E-01 | -4.34E-02 | -1.03E+00 |
| ENSG000002 | SPIRE2 | 3.50E-01 | 6.93E-01 | 1.16E+00 | 2.17E-01 | 1.16E+00 |
| ENSG000001 | SMIM12 | 3.50E-01 | 6.93E-01 | 1.02E+00 | 3.44E-02 | 1.02E+00 |
| ENSG000001 | NEIL2 | 3.50E-01 | 6.93E-01 | 1.06E+00 | 8.49E-02 | 1.06E+00 |
| ENSG000002 | ZNF815P | 3.50E-01 | ? | 1.22E+00 | 2.83E-01 | 1.22E+00 |
| ENSG000001 | CGRRF1 | 3.50E-01 | 6.93E-01 | 9.47E-01 | -7.91E-02 | -1.06E+00 |
| ENSG000001 | CSRP2 | 3.50E-01 | 6.93E-01 | 1.08E+00 | 1.11E-01 | 1.08E+00 |
| ENSG000001 | DNAJC14 | 3.50E-01 | 6.93E-01 | 1.04E+00 | 5.36E-02 | 1.04E+00 |
| ENSG000001 | KCNK5 | 3.50E-01 | ? | 1.34E+00 | 4.17E-01 | 1.34E+00 |
| ENSG000001 | SLC24A2 | 3.50E-01 | ? | 7.78E-01 | -3.62E-01 | -1.29E+00 |
| ENSG000001 | RNF44 | 3.50E-01 | 6.93E-01 | 1.04E+00 | 5.86E-02 | 1.04E+00 |
| ENSG000001 | METTL23 | 3.50E-01 | 6.93E-01 | 1.05E+00 | 6.71E-02 | 1.05E+00 |
| ENSG000001 | SSNA1 | 3.50E-01 | 6.93E-01 | 1.07E+00 | 9.19E-02 | 1.07E+00 |
| ENSG000002 | TRAF3IP1 | 3.50E-01 | 6.93E-01 | 9.54E-01 | -6.85E-02 | -1.05E+00 |
| ENSG000001 | ZNF771 | 3.50E-01 | 6.93E-01 | 1.13E+00 | 1.71E-01 | 1.13E+00 |
| ENSG000001 | DTX2P1 | 3.51E-01 | ? | 7.59E-01 | -3.97E-01 | -1.32E+00 |
| ENSG000001 | SETD7 | 3.51E-01 | 6.93E-01 | 9.57E-01 | -6.31E-02 | -1.04E+00 |
| ENSG000002 | --- | 3.51E-01 | 6.93E-01 | 1.18E+00 | 2.39E-01 | 1.18E+00 |
| ENSG000001 | MAK16 | 3.51E-01 | 6.93E-01 | 1.04E+00 | 5.76E-02 | 1.04E+00 |
| ENSG000002 | MGC16275 | 3.51E-01 | ? | 1.48E+00 | 5.63E-01 | 1.48E+00 |
| ENSG000002 | --- | 3.51E-01 | 6.93E-01 | 8.62E-01 | -2.14E-01 | -1.16E+00 |
| ENSG000001 | HNRNPA1 | 3.51E-01 | 6.93E-01 | 1.03E+00 | 4.73E-02 | 1.03E+00 |
| ENSG000002 | EHMT2 | 3.51E-01 | 6.93E-01 | 1.05E+00 | 6.79E-02 | 1.05E+00 |
| ENSG000002 | --- | 3.51E-01 | ? | 1.72E+00 | 7.81E-01 | 1.72E+00 |
| ENSG000001 | ANKRD16 | 3.51E-01 | 6.93E-01 | 9.11E-01 | -1.34E-01 | -1.10E+00 |

| | | | | | | |
|---|---|---|---|---|---|---|
| ENSG000001 | ANKRD18A | 3.51E-01 | 6.93E-01 | 9.38E-01 | -9.23E-02 | -1.07E+00 |
| ENSG000002 | LINC00881 | 3.51E-01 | ? | 7.31E-01 | -4.51E-01 | -1.37E+00 |
| ENSG000001 | THOC5 | 3.51E-01 | 6.93E-01 | 1.03E+00 | 4.23E-02 | 1.03E+00 |
| ENSG000001 | SCO1 | 3.51E-01 | 6.93E-01 | 9.76E-01 | -3.47E-02 | -1.02E+00 |
| ENSG000001 | RHBDL1 | 3.51E-01 | 6.93E-01 | 1.18E+00 | 2.37E-01 | 1.18E+00 |
| ENSG000001 | NME6 | 3.51E-01 | 6.93E-01 | 9.58E-01 | -6.12E-02 | -1.04E+00 |
| ENSG000002 | SLC25A36P1 | 3.51E-01 | ? | 6.12E-01 | -7.08E-01 | -1.63E+00 |
| ENSG000002 | NUTM2B-AS | 3.51E-01 | 6.93E-01 | 9.63E-01 | -5.41E-02 | -1.04E+00 |
| ENSG000002 | --- | 3.51E-01 | ? | 1.67E+00 | 7.37E-01 | 1.67E+00 |
| ENSG00000C | TMEM38A | 3.51E-01 | 6.93E-01 | 1.13E+00 | 1.77E-01 | 1.13E+00 |
| ENSG000001 | ZFP2 | 3.51E-01 | 6.93E-01 | 1.15E+00 | 2.00E-01 | 1.15E+00 |
| ENSG000002 | DDX59-AS1 | 3.51E-01 | ? | 1.29E+00 | 3.72E-01 | 1.29E+00 |
| ENSG000001 | CEP295 | 3.52E-01 | 6.94E-01 | 1.05E+00 | 7.07E-02 | 1.05E+00 |
| ENSG000002 | --- | 3.52E-01 | ? | 1.47E+00 | 5.53E-01 | 1.47E+00 |
| ENSG000002 | --- | 3.52E-01 | ? | 1.26E+00 | 3.37E-01 | 1.26E+00 |
| ENSG000001 | SLC51A | 3.52E-01 | 6.94E-01 | 1.08E+00 | 1.13E-01 | 1.08E+00 |
| ENSG000002 | BCYRN1 | 3.52E-01 | 6.94E-01 | 9.33E-01 | -9.97E-02 | -1.07E+00 |
| ENSG000001 | MCF2 | 3.52E-01 | ? | 1.62E+00 | 6.99E-01 | 1.62E+00 |
| ENSG000001 | NAALADL2 | 3.52E-01 | 6.94E-01 | 9.09E-01 | -1.37E-01 | -1.10E+00 |
| ENSG000001 | DNAAF1 | 3.52E-01 | ? | 1.38E+00 | 4.66E-01 | 1.38E+00 |
| ENSG000001 | RAD9B | 3.52E-01 | ? | 8.29E-01 | -2.70E-01 | -1.21E+00 |
| ENSG000001 | ATR | 3.52E-01 | 6.94E-01 | 9.64E-01 | -5.26E-02 | -1.04E+00 |
| ENSG000001 | MGRN1 | 3.52E-01 | 6.94E-01 | 1.04E+00 | 6.06E-02 | 1.04E+00 |
| ENSG000001 | SKA2 | 3.52E-01 | 6.94E-01 | 9.57E-01 | -6.36E-02 | -1.05E+00 |
| ENSG000002 | --- | 3.52E-01 | ? | 1.37E+00 | 4.49E-01 | 1.37E+00 |
| ENSG000001 | ZNF780B | 3.52E-01 | 6.94E-01 | 1.06E+00 | 8.66E-02 | 1.06E+00 |
| ENSG000002 | --- | 3.52E-01 | ? | 7.67E-01 | -3.84E-01 | -1.30E+00 |
| ENSG000001 | DSCAM | 3.52E-01 | 6.94E-01 | 1.08E+00 | 1.04E-01 | 1.08E+00 |
| ENSG000002 | --- | 3.52E-01 | ? | 1.74E+00 | 7.99E-01 | 1.74E+00 |
| ENSG000001 | MBNL2 | 3.52E-01 | 6.94E-01 | 1.04E+00 | 5.90E-02 | 1.04E+00 |
| ENSG000001 | TRIM9 | 3.52E-01 | ? | 1.17E+00 | 2.24E-01 | 1.17E+00 |
| ENSG000001 | CREBRF | 3.53E-01 | 6.95E-01 | 1.08E+00 | 1.09E-01 | 1.08E+00 |
| ENSG000002 | EIF1B-AS1 | 3.53E-01 | ? | 1.18E+00 | 2.33E-01 | 1.18E+00 |
| ENSG000001 | COPA | 3.53E-01 | 6.95E-01 | 9.76E-01 | -3.53E-02 | -1.02E+00 |
| ENSG000002 | --- | 3.53E-01 | ? | 1.39E+00 | 4.71E-01 | 1.39E+00 |
| ENSG000001 | SEC23A | 3.53E-01 | 6.95E-01 | 1.03E+00 | 4.70E-02 | 1.03E+00 |
| ENSG000001 | ZNF774 | 3.53E-01 | 6.95E-01 | 1.07E+00 | 9.09E-02 | 1.07E+00 |
| ENSG000002 | BUD13P1 | 3.53E-01 | ? | 1.75E+00 | 8.07E-01 | 1.75E+00 |
| ENSG000002 | NAA80 | 3.53E-01 | 6.95E-01 | 9.31E-01 | -1.02E-01 | -1.07E+00 |
| ENSG000002 | --- | 3.53E-01 | 6.95E-01 | 1.11E+00 | 1.50E-01 | 1.11E+00 |
| ENSG000001 | MREG | 3.53E-01 | 6.95E-01 | 1.04E+00 | 6.34E-02 | 1.04E+00 |
| ENSG000002 | --- | 3.53E-01 | ? | 1.34E+00 | 4.24E-01 | 1.34E+00 |
| ENSG000002 | --- | 3.53E-01 | ? | 1.29E+00 | 3.65E-01 | 1.29E+00 |
| ENSG000001 | KLC2 | 3.53E-01 | 6.95E-01 | 9.65E-01 | -5.16E-02 | -1.04E+00 |
| ENSG000002 | LINC00589 | 3.53E-01 | 6.95E-01 | 8.91E-01 | -1.67E-01 | -1.12E+00 |

| | | | | | | |
|---|---|---|---|---|---|---|
| ENSG000002 | ACAD9-DT | 3.53E-01 | ? | 1.18E+00 | 2.36E-01 | 1.18E+00 |
| ENSG00000C | GLI2 | 3.53E-01 | 6.95E-01 | 8.60E-01 | -2.17E-01 | -1.16E+00 |
| ENSG000002 | HERC2P7 | 3.53E-01 | ? | 1.30E+00 | 3.77E-01 | 1.30E+00 |
| ENSG000001 | DIPK2A | 3.53E-01 | 6.95E-01 | 1.04E+00 | 5.99E-02 | 1.04E+00 |
| ENSG000001 | RBFA | 3.53E-01 | 6.95E-01 | 9.60E-01 | -5.95E-02 | -1.04E+00 |
| ENSG000001 | OBSCN-AS1 | 3.53E-01 | ? | 1.21E+00 | 2.77E-01 | 1.21E+00 |
| ENSG000001 | FAM88B | 3.53E-01 | ? | 8.62E-01 | -2.14E-01 | -1.16E+00 |
| ENSG000002 | --- | 3.53E-01 | 6.95E-01 | 1.12E+00 | 1.61E-01 | 1.12E+00 |
| ENSG000002 | NPEPL1 | 3.53E-01 | 6.95E-01 | 1.06E+00 | 8.48E-02 | 1.06E+00 |
| ENSG000001 | MMP13 | 3.53E-01 | ? | 6.94E-01 | -5.27E-01 | -1.44E+00 |
| ENSG000001 | TANC1 | 3.54E-01 | 6.95E-01 | 9.78E-01 | -3.24E-02 | -1.02E+00 |
| ENSG000001 | PPP1R12C | 3.54E-01 | 6.96E-01 | 1.05E+00 | 6.58E-02 | 1.05E+00 |
| ENSG000001 | H2AC13 | 3.54E-01 | ? | 1.28E+00 | 3.57E-01 | 1.28E+00 |
| ENSG000002 | BTF3P10 | 3.54E-01 | ? | 1.58E+00 | 6.63E-01 | 1.58E+00 |
| ENSG000001 | COG1 | 3.54E-01 | 6.96E-01 | 9.69E-01 | -4.55E-02 | -1.03E+00 |
| ENSG000001 | EPHB3 | 3.54E-01 | 6.96E-01 | 1.12E+00 | 1.58E-01 | 1.12E+00 |
| ENSG000001 | RYR2 | 3.54E-01 | ? | 8.19E-01 | -2.88E-01 | -1.22E+00 |
| ENSG000001 | ZNF383 | 3.54E-01 | 6.96E-01 | 1.06E+00 | 8.59E-02 | 1.06E+00 |
| ENSG000001 | RBM15 | 3.54E-01 | 6.96E-01 | 9.63E-01 | -5.42E-02 | -1.04E+00 |
| ENSG000001 | OTUD4P1 | 3.54E-01 | ? | 6.80E-01 | -5.56E-01 | -1.47E+00 |
| ENSG000002 | --- | 3.54E-01 | 6.96E-01 | 8.98E-01 | -1.55E-01 | -1.11E+00 |
| ENSG000002 | RETREG1-AS | 3.54E-01 | ? | 8.17E-01 | -2.92E-01 | -1.22E+00 |
| ENSG000001 | SMURF1 | 3.54E-01 | 6.96E-01 | 9.76E-01 | -3.53E-02 | -1.02E+00 |
| ENSG000002 | HMGN1P4 | 3.54E-01 | ? | 7.45E-01 | -4.25E-01 | -1.34E+00 |
| ENSG000002 | --- | 3.54E-01 | ? | 7.91E-01 | -3.38E-01 | -1.26E+00 |
| ENSG000002 | LINC00662 | 3.54E-01 | 6.96E-01 | 9.54E-01 | -6.80E-02 | -1.05E+00 |
| ENSG000001 | RWDD3 | 3.54E-01 | 6.96E-01 | 1.06E+00 | 7.99E-02 | 1.06E+00 |
| ENSG00000C | CYP26B1 | 3.54E-01 | 6.96E-01 | 9.31E-01 | -1.03E-01 | -1.07E+00 |
| ENSG000001 | GNG12 | 3.55E-01 | 6.96E-01 | 1.03E+00 | 4.34E-02 | 1.03E+00 |
| ENSG000002 | --- | 3.55E-01 | ? | 1.35E+00 | 4.30E-01 | 1.35E+00 |
| ENSG000001 | RUSF1 | 3.55E-01 | 6.97E-01 | 1.04E+00 | 6.25E-02 | 1.04E+00 |
| ENSG000001 | RGS19 | 3.55E-01 | 6.97E-01 | 9.24E-01 | -1.14E-01 | -1.08E+00 |
| ENSG000002 | --- | 3.55E-01 | ? | 7.79E-01 | -3.60E-01 | -1.28E+00 |
| ENSG000001 | LRRC9 | 3.55E-01 | ? | 1.35E+00 | 4.31E-01 | 1.35E+00 |
| ENSG00000C | IFT88 | 3.55E-01 | 6.97E-01 | 9.52E-01 | -7.05E-02 | -1.05E+00 |
| ENSG00000C | SCIN | 3.55E-01 | ? | 1.53E+00 | 6.16E-01 | 1.53E+00 |
| ENSG000002 | KLRC3 | 3.55E-01 | ? | 1.60E+00 | 6.82E-01 | 1.60E+00 |
| ENSG000001 | NDFIP2 | 3.55E-01 | 6.97E-01 | 1.03E+00 | 4.25E-02 | 1.03E+00 |
| ENSG000001 | PCBP3 | 3.56E-01 | ? | 7.30E-01 | -4.53E-01 | -1.37E+00 |
| ENSG000001 | SCAF4 | 3.56E-01 | 6.98E-01 | 9.56E-01 | -6.48E-02 | -1.05E+00 |
| ENSG000001 | LENG8 | 3.56E-01 | 6.98E-01 | 9.12E-01 | -1.33E-01 | -1.10E+00 |
| ENSG000002 | --- | 3.56E-01 | 6.98E-01 | 8.82E-01 | -1.82E-01 | -1.13E+00 |
| ENSG000001 | RCHY1 | 3.56E-01 | 6.98E-01 | 9.62E-01 | -5.58E-02 | -1.04E+00 |
| ENSG000001 | SLC25A4 | 3.56E-01 | 6.98E-01 | 9.64E-01 | -5.25E-02 | -1.04E+00 |
| ENSG000001 | FER | 3.56E-01 | 6.98E-01 | 9.52E-01 | -7.08E-02 | -1.05E+00 |

| | | | | | | | |
|---|---|---|---|---|---|---|---|
| ENSG000002 | --- | 3.56E-01 | | 6.98E-01 | 1.16E+00 | 2.09E-01 | 1.16E+00 |
| ENSG000002 | ZNF805 | 3.56E-01 | | 6.98E-01 | 9.48E-01 | -7.70E-02 | -1.05E+00 |
| ENSG000002 | BTBD3-AS1 | 3.56E-01 | ? | | 7.14E-01 | -4.85E-01 | -1.40E+00 |
| ENSG000001 | CDC73 | 3.56E-01 | | 6.98E-01 | 9.66E-01 | -4.95E-02 | -1.03E+00 |
| ENSG000001 | GGACT | 3.56E-01 | | 6.98E-01 | 1.07E+00 | 9.25E-02 | 1.07E+00 |
| ENSG000001 | CPSF2 | 3.56E-01 | | 6.98E-01 | 1.03E+00 | 3.89E-02 | 1.03E+00 |
| ENSG000002 | --- | 3.56E-01 | ? | | 7.10E-01 | -4.93E-01 | -1.41E+00 |
| ENSG000002 | DNM1P35 | 3.56E-01 | ? | | 7.28E-01 | -4.59E-01 | -1.37E+00 |
| ENSG000001 | PUSL1 | 3.56E-01 | | 6.98E-01 | 1.08E+00 | 1.15E-01 | 1.08E+00 |
| ENSG000001 | RASL11B | 3.56E-01 | | 6.98E-01 | 8.76E-01 | -1.91E-01 | -1.14E+00 |
| ENSG000002 | LINC01347 | 3.57E-01 | | 6.98E-01 | 9.31E-01 | -1.03E-01 | -1.07E+00 |
| ENSG000001 | SRGAP2C | 3.57E-01 | | 6.99E-01 | 9.49E-01 | -7.57E-02 | -1.05E+00 |
| ENSG000001 | NSD2 | 3.57E-01 | | 6.99E-01 | 9.72E-01 | -4.10E-02 | -1.03E+00 |
| ENSG000001 | SHPRH | 3.57E-01 | | 6.99E-01 | 1.04E+00 | 6.06E-02 | 1.04E+00 |
| ENSG000001 | VAV3 | 3.57E-01 | | 6.99E-01 | 1.07E+00 | 1.02E-01 | 1.07E+00 |
| ENSG000001 | THAP7 | 3.57E-01 | | 6.99E-01 | 1.07E+00 | 9.91E-02 | 1.07E+00 |
| ENSG000002 | --- | 3.57E-01 | ? | | 8.03E-01 | -3.17E-01 | -1.25E+00 |
| ENSG000001 | PLA2G4A | 3.57E-01 | | 6.99E-01 | 1.06E+00 | 7.83E-02 | 1.06E+00 |
| ENSG000001 | DUSP4 | 3.57E-01 | | 6.99E-01 | 9.09E-01 | -1.38E-01 | -1.10E+00 |
| ENSG00000C | ZFYVE16 | 3.57E-01 | | 6.99E-01 | 1.03E+00 | 4.07E-02 | 1.03E+00 |
| ENSG000001 | RSRC1 | 3.57E-01 | | 6.99E-01 | 9.61E-01 | -5.76E-02 | -1.04E+00 |
| ENSG000001 | SUFU | 3.58E-01 | | 6.99E-01 | 1.04E+00 | 5.26E-02 | 1.04E+00 |
| ENSG000002 | --- | 3.58E-01 | ? | | 8.22E-01 | -2.83E-01 | -1.22E+00 |
| ENSG000001 | RNLS | 3.58E-01 | | 6.99E-01 | 9.23E-01 | -1.15E-01 | -1.08E+00 |
| ENSG000001 | WDR53 | 3.58E-01 | | 6.99E-01 | 9.49E-01 | -7.55E-02 | -1.05E+00 |
| ENSG000002 | --- | 3.58E-01 | | 6.99E-01 | 1.09E+00 | 1.21E-01 | 1.09E+00 |
| ENSG000002 | CTBP2P4 | 3.58E-01 | ? | | 7.49E-01 | -4.17E-01 | -1.34E+00 |
| ENSG000002 | SPCS2P4 | 3.58E-01 | | 7.00E-01 | 1.17E+00 | 2.21E-01 | 1.17E+00 |
| ENSG00000C | ZFR | 3.58E-01 | | 7.00E-01 | 9.65E-01 | -5.20E-02 | -1.04E+00 |
| ENSG000002 | --- | 3.58E-01 | ? | | 7.61E-01 | -3.93E-01 | -1.31E+00 |
| ENSG000002 | NATD1 | 3.58E-01 | | 7.00E-01 | 1.08E+00 | 1.11E-01 | 1.08E+00 |
| ENSG000001 | ZSWIM3 | 3.59E-01 | | 7.01E-01 | 9.20E-01 | -1.21E-01 | -1.09E+00 |
| ENSG000002 | --- | 3.59E-01 | ? | | 8.21E-01 | -2.85E-01 | -1.22E+00 |
| ENSG000001 | GIPC2 | 3.59E-01 | ? | | 1.33E+00 | 4.09E-01 | 1.33E+00 |
| ENSG000001 | MED12L | 3.59E-01 | ? | | 1.16E+00 | 2.09E-01 | 1.16E+00 |
| ENSG000002 | --- | 3.59E-01 | ? | | 1.72E+00 | 7.80E-01 | 1.72E+00 |
| ENSG000001 | JAZF1 | 3.59E-01 | | 7.01E-01 | 9.49E-01 | -7.53E-02 | -1.05E+00 |
| ENSG00000C | HEBP1 | 3.59E-01 | | 7.01E-01 | 1.04E+00 | 5.88E-02 | 1.04E+00 |
| ENSG000002 | PTGES3P1 | 3.59E-01 | ? | | 8.04E-01 | -3.15E-01 | -1.24E+00 |
| ENSG000002 | ARL2 | 3.59E-01 | | 7.01E-01 | 1.06E+00 | 8.31E-02 | 1.06E+00 |
| ENSG000002 | --- | 3.59E-01 | | 7.01E-01 | 1.11E+00 | 1.48E-01 | 1.11E+00 |
| ENSG000002 | EFNA4 | 3.59E-01 | | 7.01E-01 | 1.07E+00 | 9.20E-02 | 1.07E+00 |
| ENSG000002 | --- | 3.59E-01 | | 7.01E-01 | 1.07E+00 | 9.34E-02 | 1.07E+00 |
| ENSG000001 | HSPA4L | 3.59E-01 | | 7.01E-01 | 1.10E+00 | 1.36E-01 | 1.10E+00 |
| ENSG000002 | --- | 3.59E-01 | ? | | 7.86E-01 | -3.47E-01 | -1.27E+00 |

| | | | | | | |
|---|---|---|---|---|---|---|
| ENSG000002 | --- | 3.59E-01 | 7.01E-01 | 9.19E-01 | -1.22E-01 | -1.09E+00 |
| ENSG000001 | STIM2 | 3.59E-01 | 7.01E-01 | 1.04E+00 | 5.19E-02 | 1.04E+00 |
| ENSG000001 | CASP6 | 3.59E-01 | 7.01E-01 | 9.61E-01 | -5.80E-02 | -1.04E+00 |
| ENSG000001 | ATP5F1B | 3.60E-01 | 7.01E-01 | 1.04E+00 | 5.61E-02 | 1.04E+00 |
| ENSG000001 | NCAM2 | 3.60E-01 | ? | 7.21E-01 | -4.73E-01 | -1.39E+00 |
| ENSG000001 | MT-CO1 | 3.60E-01 | 7.01E-01 | 9.27E-01 | -1.09E-01 | -1.08E+00 |
| ENSG00000C | SRBD1 | 3.60E-01 | 7.02E-01 | 9.56E-01 | -6.44E-02 | -1.05E+00 |
| ENSG00000C | LIPE | 3.60E-01 | 7.02E-01 | 1.08E+00 | 1.10E-01 | 1.08E+00 |
| ENSG00000C | RPS6KA6 | 3.60E-01 | 7.02E-01 | 1.07E+00 | 9.31E-02 | 1.07E+00 |
| ENSG000002 | MT-TC | 3.60E-01 | ? | 1.48E+00 | 5.69E-01 | 1.48E+00 |
| ENSG000001 | TBC1D20 | 3.60E-01 | 7.02E-01 | 1.03E+00 | 3.76E-02 | 1.03E+00 |
| ENSG000002 | --- | 3.60E-01 | ? | 7.69E-01 | -3.78E-01 | -1.30E+00 |
| ENSG000001 | ARHGAP40 | 3.60E-01 | 7.02E-01 | 1.09E+00 | 1.22E-01 | 1.09E+00 |
| ENSG000001 | TEX30 | 3.60E-01 | 7.02E-01 | 9.33E-01 | -1.00E-01 | -1.07E+00 |
| ENSG00000C | LIMS2 | 3.60E-01 | ? | 1.27E+00 | 3.48E-01 | 1.27E+00 |
| ENSG000001 | RO60 | 3.60E-01 | 7.02E-01 | 1.04E+00 | 6.15E-02 | 1.04E+00 |
| ENSG00000C | CCDC124 | 3.60E-01 | 7.02E-01 | 1.04E+00 | 5.66E-02 | 1.04E+00 |
| ENSG00000C | MCAM | 3.60E-01 | 7.02E-01 | 9.47E-01 | -7.79E-02 | -1.06E+00 |
| ENSG000002 | --- | 3.61E-01 | 7.02E-01 | 7.65E-01 | -3.87E-01 | -1.31E+00 |
| ENSG00000C | FOSL2 | 3.61E-01 | 7.02E-01 | 1.04E+00 | 5.01E-02 | 1.04E+00 |
| ENSG000002 | RPLP0P2 | 3.61E-01 | 7.02E-01 | 8.39E-01 | -2.54E-01 | -1.19E+00 |
| ENSG000001 | PIM1 | 3.61E-01 | 7.02E-01 | 1.06E+00 | 8.65E-02 | 1.06E+00 |
| ENSG000001 | MT-CO2 | 3.61E-01 | 7.02E-01 | 9.31E-01 | -1.04E-01 | -1.07E+00 |
| ENSG000002 | ERICD | 3.61E-01 | ? | 8.46E-01 | -2.41E-01 | -1.18E+00 |
| ENSG000002 | SPANXA2-OT | 3.61E-01 | ? | 8.03E-01 | -3.16E-01 | -1.25E+00 |
| ENSG000001 | GPN3 | 3.61E-01 | 7.03E-01 | 1.04E+00 | 6.24E-02 | 1.04E+00 |
| ENSG000001 | LIX1 | 3.61E-01 | ? | 1.66E+00 | 7.34E-01 | 1.66E+00 |
| ENSG000002 | --- | 3.61E-01 | 7.03E-01 | 9.26E-01 | -1.11E-01 | -1.08E+00 |
| ENSG000001 | FRMD8 | 3.61E-01 | 7.03E-01 | 1.07E+00 | 9.13E-02 | 1.07E+00 |
| ENSG000001 | TAB1 | 3.61E-01 | 7.03E-01 | 1.04E+00 | 6.16E-02 | 1.04E+00 |
| ENSG000001 | RRAD | 3.61E-01 | 7.03E-01 | 1.17E+00 | 2.32E-01 | 1.17E+00 |
| ENSG000002 | RTL9 | 3.61E-01 | ? | 1.16E+00 | 2.17E-01 | 1.16E+00 |
| ENSG000002 | GABPB1-IT1 | 3.61E-01 | 7.03E-01 | 1.08E+00 | 1.10E-01 | 1.08E+00 |
| ENSG000001 | TFDP1 | 3.61E-01 | 7.03E-01 | 1.03E+00 | 4.32E-02 | 1.03E+00 |
| ENSG00000C | PSEN1 | 3.61E-01 | 7.03E-01 | 9.74E-01 | -3.83E-02 | -1.03E+00 |
| ENSG000001 | KIAA1549L | 3.62E-01 | 7.03E-01 | 1.05E+00 | 6.37E-02 | 1.05E+00 |
| ENSG00000C | ALDH3B1 | 3.62E-01 | 7.03E-01 | 1.10E+00 | 1.42E-01 | 1.10E+00 |
| ENSG000001 | EPHB6 | 3.62E-01 | ? | 8.32E-01 | -2.66E-01 | -1.20E+00 |
| ENSG000001 | HS3ST2 | 3.62E-01 | 7.04E-01 | 8.59E-01 | -2.19E-01 | -1.16E+00 |
| ENSG000002 | --- | 3.62E-01 | ? | 7.68E-01 | -3.81E-01 | -1.30E+00 |
| ENSG000001 | SRP54 | 3.62E-01 | 7.04E-01 | 9.67E-01 | -4.86E-02 | -1.03E+00 |
| ENSG000001 | CATSPER2 | 3.62E-01 | 7.04E-01 | 8.77E-01 | -1.89E-01 | -1.14E+00 |
| ENSG000001 | TAOK1 | 3.62E-01 | 7.04E-01 | 9.53E-01 | -6.92E-02 | -1.05E+00 |
| ENSG00000C | MYCBP2 | 3.62E-01 | 7.04E-01 | 9.36E-01 | -9.51E-02 | -1.07E+00 |
| ENSG000001 | FARSB | 3.62E-01 | 7.04E-01 | 9.70E-01 | -4.37E-02 | -1.03E+00 |

| | | | | | | |
|---|---|---|---|---|---|---|
| ENSG000001 | GPR61 | 3.62E-01 | ? | 1.49E+00 | 5.76E-01 | 1.49E+00 |
| ENSG000001 | SRPRB | 3.62E-01 | 7.04E-01 | 1.03E+00 | 4.00E-02 | 1.03E+00 |
| ENSG000002 | LINC01208 | 3.62E-01 | ? | 1.63E+00 | 7.03E-01 | 1.63E+00 |
| ENSG000001 | RPP30 | 3.62E-01 | 7.04E-01 | 1.03E+00 | 4.66E-02 | 1.03E+00 |
| ENSG000001 | INVS | 3.62E-01 | 7.04E-01 | 1.03E+00 | 4.67E-02 | 1.03E+00 |
| ENSG000001 | TTK | 3.62E-01 | 7.04E-01 | 9.04E-01 | -1.46E-01 | -1.11E+00 |
| ENSG000001 | C1QTNF12 | 3.62E-01 | ? | 1.41E+00 | 4.99E-01 | 1.41E+00 |
| ENSG000001 | NUP214 | 3.62E-01 | 7.04E-01 | 1.03E+00 | 3.65E-02 | 1.03E+00 |
| ENSG000001 | ARMH4 | 3.63E-01 | 7.04E-01 | 1.11E+00 | 1.44E-01 | 1.11E+00 |
| ENSG000001 | SCRIB | 3.63E-01 | 7.04E-01 | 1.08E+00 | 1.13E-01 | 1.08E+00 |
| ENSG000001 | RFC4 | 3.63E-01 | 7.04E-01 | 9.45E-01 | -8.12E-02 | -1.06E+00 |
| ENSG000002 | LASTR | 3.63E-01 | 7.04E-01 | 9.52E-01 | -7.03E-02 | -1.05E+00 |
| ENSG000002 | --- | 3.63E-01 | 7.04E-01 | 1.19E+00 | 2.48E-01 | 1.19E+00 |
| ENSG000001 | DCTN3 | 3.63E-01 | 7.04E-01 | 1.05E+00 | 6.86E-02 | 1.05E+00 |
| ENSG000001 | MRPS25 | 3.63E-01 | 7.04E-01 | 9.56E-01 | -6.45E-02 | -1.05E+00 |
| ENSG000001 | CELF2-AS1 | 3.63E-01 | ? | 1.26E+00 | 3.38E-01 | 1.26E+00 |
| ENSG000002 | --- | 3.63E-01 | 7.04E-01 | 9.26E-01 | -1.11E-01 | -1.08E+00 |
| ENSG000001 | EPHB4 | 3.63E-01 | 7.04E-01 | 1.05E+00 | 6.66E-02 | 1.05E+00 |
| ENSG000002 | IFI30 | 3.63E-01 | 7.04E-01 | 1.09E+00 | 1.20E-01 | 1.09E+00 |
| ENSG000002 | CYB5RL | 3.63E-01 | 7.04E-01 | 1.11E+00 | 1.50E-01 | 1.11E+00 |
| ENSG000001 | KAT2A | 3.63E-01 | 7.04E-01 | 1.06E+00 | 8.08E-02 | 1.06E+00 |
| ENSG000002 | --- | 3.63E-01 | ? | 1.50E+00 | 5.81E-01 | 1.50E+00 |
| ENSG000002 | --- | 3.63E-01 | 7.04E-01 | 8.58E-01 | -2.21E-01 | -1.17E+00 |
| ENSG000001 | ZNF750 | 3.63E-01 | 7.04E-01 | 1.12E+00 | 1.69E-01 | 1.12E+00 |
| ENSG000001 | CSTB | 3.63E-01 | 7.04E-01 | 1.06E+00 | 7.74E-02 | 1.06E+00 |
| ENSG000002 | MT-TN | 3.63E-01 | 7.04E-01 | 8.27E-01 | -2.75E-01 | -1.21E+00 |
| ENSG000002 | HLA-DMA | 3.63E-01 | ? | 8.37E-01 | -2.57E-01 | -1.20E+00 |
| ENSG00000C | NFATC3 | 3.64E-01 | 7.04E-01 | 1.04E+00 | 5.92E-02 | 1.04E+00 |
| ENSG000001 | CENPH | 3.64E-01 | 7.04E-01 | 1.08E+00 | 1.15E-01 | 1.08E+00 |
| ENSG000002 | --- | 3.64E-01 | 7.04E-01 | 8.93E-01 | -1.63E-01 | -1.12E+00 |
| ENSG000002 | SCIRT | 3.64E-01 | ? | 8.35E-01 | -2.60E-01 | -1.20E+00 |
| ENSG000001 | PDHB | 3.64E-01 | 7.04E-01 | 9.74E-01 | -3.76E-02 | -1.03E+00 |
| ENSG000002 | --- | 3.64E-01 | ? | 1.34E+00 | 4.27E-01 | 1.34E+00 |
| ENSG000001 | CCDC168 | 3.64E-01 | ? | 7.29E-01 | -4.55E-01 | -1.37E+00 |
| ENSG000002 | --- | 3.64E-01 | ? | 6.09E-01 | -7.16E-01 | -1.64E+00 |
| ENSG000002 | LINC01291 | 3.64E-01 | ? | 1.45E+00 | 5.39E-01 | 1.45E+00 |
| ENSG000001 | KCND3 | 3.64E-01 | ? | 8.17E-01 | -2.91E-01 | -1.22E+00 |
| ENSG000002 | --- | 3.64E-01 | 7.05E-01 | 8.83E-01 | -1.80E-01 | -1.13E+00 |
| ENSG000002 | --- | 3.64E-01 | ? | 6.74E-01 | -5.69E-01 | -1.48E+00 |
| ENSG000002 | MRC1 | 3.64E-01 | ? | 6.60E-01 | -5.99E-01 | -1.52E+00 |
| ENSG000002 | --- | 3.64E-01 | ? | 1.29E+00 | 3.63E-01 | 1.29E+00 |
| ENSG000002 | ABCA11P | 3.64E-01 | 7.05E-01 | 1.09E+00 | 1.23E-01 | 1.09E+00 |
| ENSG000001 | KCNJ12 | 3.64E-01 | 7.05E-01 | 1.13E+00 | 1.77E-01 | 1.13E+00 |
| ENSG000002 | MT-TA | 3.64E-01 | 7.05E-01 | 8.51E-01 | -2.33E-01 | -1.18E+00 |
| ENSG000001 | TWNK | 3.64E-01 | 7.05E-01 | 1.04E+00 | 5.99E-02 | 1.04E+00 |

| | | | | | | |
|---|---|---|---|---|---|---|
| ENSG000001 | C16orf74 | 3.64E-01 | 7.05E-01 | 1.08E+00 | 1.05E-01 | 1.08E+00 |
| ENSG000001 | ARHGAP29 | 3.64E-01 | 7.05E-01 | 9.45E-01 | -8.11E-02 | -1.06E+00 |
| ENSG000001 | PKIA | 3.64E-01 | 7.05E-01 | 1.07E+00 | 1.04E-01 | 1.07E+00 |
| ENSG000002 | FAM169BP | 3.64E-01 | ? | 6.76E-01 | -5.64E-01 | -1.48E+00 |
| ENSG000001 | NDUFB11 | 3.64E-01 | 7.05E-01 | 1.06E+00 | 8.32E-02 | 1.06E+00 |
| ENSG000002 | --- | 3.64E-01 | ? | 6.97E-01 | -5.21E-01 | -1.43E+00 |
| ENSG000001 | NPTN | 3.64E-01 | 7.05E-01 | 1.02E+00 | 2.93E-02 | 1.02E+00 |
| ENSG000001 | ADRA2C | 3.64E-01 | ? | 8.15E-01 | -2.96E-01 | -1.23E+00 |
| ENSG000001 | GTF2B | 3.64E-01 | 7.05E-01 | 1.03E+00 | 4.49E-02 | 1.03E+00 |
| ENSG000002 | CLEC5A | 3.65E-01 | ? | 6.90E-01 | -5.36E-01 | -1.45E+00 |
| ENSG000001 | SIK1 | 3.65E-01 | 7.05E-01 | 1.08E+00 | 1.05E-01 | 1.08E+00 |
| ENSG000001 | SH2D3A | 3.65E-01 | 7.05E-01 | 9.66E-01 | -4.96E-02 | -1.03E+00 |
| ENSG000001 | GTF3C3 | 3.65E-01 | 7.05E-01 | 9.75E-01 | -3.64E-02 | -1.03E+00 |
| ENSG000001 | DCAF15 | 3.65E-01 | 7.05E-01 | 1.06E+00 | 8.77E-02 | 1.06E+00 |
| ENSG000001 | TNFSF13 | 3.65E-01 | 7.06E-01 | 9.47E-01 | -7.90E-02 | -1.06E+00 |
| ENSG000001 | CD2BP2 | 3.65E-01 | 7.06E-01 | 1.03E+00 | 4.75E-02 | 1.03E+00 |
| ENSG000002 | RCN1P2 | 3.66E-01 | 7.06E-01 | 8.71E-01 | -1.99E-01 | -1.15E+00 |
| ENSG000001 | JTB | 3.66E-01 | 7.06E-01 | 1.06E+00 | 8.70E-02 | 1.06E+00 |
| ENSG000001 | RRAGC | 3.66E-01 | 7.06E-01 | 1.04E+00 | 5.81E-02 | 1.04E+00 |
| ENSG000001 | TEDC1 | 3.66E-01 | 7.06E-01 | 1.11E+00 | 1.56E-01 | 1.11E+00 |
| ENSG000002 | --- | 3.66E-01 | ? | 6.95E-01 | -5.25E-01 | -1.44E+00 |
| ENSG000001 | TRAPPC8 | 3.66E-01 | 7.06E-01 | 1.02E+00 | 3.28E-02 | 1.02E+00 |
| ENSG000001 | IL18BP | 3.66E-01 | 7.06E-01 | 9.34E-01 | -9.77E-02 | -1.07E+00 |
| ENSG000002 | DCTN1 | 3.66E-01 | 7.06E-01 | 9.77E-01 | -3.31E-02 | -1.02E+00 |
| ENSG000002 | --- | 3.66E-01 | 7.06E-01 | 9.03E-01 | -1.47E-01 | -1.11E+00 |
| ENSG000001 | SPDYE1 | 3.66E-01 | ? | 7.99E-01 | -3.23E-01 | -1.25E+00 |
| ENSG000002 | --- | 3.66E-01 | 7.06E-01 | 1.15E+00 | 2.04E-01 | 1.15E+00 |
| ENSG000001 | SBK1 | 3.66E-01 | ? | 1.23E+00 | 2.94E-01 | 1.23E+00 |
| ENSG000002 | LINC00578 | 3.66E-01 | ? | 7.46E-01 | -4.23E-01 | -1.34E+00 |
| ENSG000001 | NAP1L1 | 3.66E-01 | 7.07E-01 | 1.03E+00 | 4.73E-02 | 1.03E+00 |
| ENSG000002 | CKLF-CMTM1 | 3.66E-01 | ? | 1.39E+00 | 4.78E-01 | 1.39E+00 |
| ENSG000001 | FBXO32 | 3.66E-01 | 7.07E-01 | 8.88E-01 | -1.71E-01 | -1.13E+00 |
| ENSG000001 | CDKAL1 | 3.66E-01 | 7.07E-01 | 1.04E+00 | 5.48E-02 | 1.04E+00 |
| ENSG000001 | TMTC1 | 3.66E-01 | 7.07E-01 | 1.11E+00 | 1.45E-01 | 1.11E+00 |
| ENSG000001 | ARMCX2 | 3.67E-01 | 7.07E-01 | 1.03E+00 | 4.19E-02 | 1.03E+00 |
| ENSG000001 | CWC22 | 3.67E-01 | 7.07E-01 | 9.62E-01 | -5.62E-02 | -1.04E+00 |
| ENSG000000 | RANBP1 | 3.67E-01 | 7.07E-01 | 1.05E+00 | 6.47E-02 | 1.05E+00 |
| ENSG000001 | SULT4A1 | 3.67E-01 | 7.07E-01 | 1.10E+00 | 1.39E-01 | 1.10E+00 |
| ENSG000001 | GNRH2 | 3.67E-01 | ? | 1.48E+00 | 5.64E-01 | 1.48E+00 |
| ENSG000001 | IRF7 | 3.67E-01 | 7.07E-01 | 8.92E-01 | -1.64E-01 | -1.12E+00 |
| ENSG000002 | --- | 3.67E-01 | 7.07E-01 | 1.09E+00 | 1.24E-01 | 1.09E+00 |
| ENSG000002 | --- | 3.67E-01 | 7.07E-01 | 8.75E-01 | -1.92E-01 | -1.14E+00 |
| ENSG000002 | --- | 3.67E-01 | 7.07E-01 | 9.18E-01 | -1.24E-01 | -1.09E+00 |
| ENSG000001 | SLC35A3 | 3.67E-01 | 7.07E-01 | 1.03E+00 | 4.58E-02 | 1.03E+00 |
| ENSG000002 | PCDHGA3 | 3.67E-01 | ? | 7.79E-01 | -3.60E-01 | -1.28E+00 |

| | | | | | | |
|---|---|---|---|---|---|---|
| ENSG000001 | SURF6 | 3.67E-01 | 7.07E-01 | 1.03E+00 | 4.03E-02 | 1.03E+00 |
| ENSG000002 | --- | 3.67E-01 | ? | 8.28E-01 | -2.72E-01 | -1.21E+00 |
| ENSG000001 | DIPK1B | 3.67E-01 | ? | 1.21E+00 | 2.72E-01 | 1.21E+00 |
| ENSG000002 | ARRDC3-AS1 | 3.67E-01 | 7.07E-01 | 1.15E+00 | 2.05E-01 | 1.15E+00 |
| ENSG000002 | --- | 3.67E-01 | ? | 1.19E+00 | 2.49E-01 | 1.19E+00 |
| ENSG000001 | KIF27 | 3.67E-01 | 7.07E-01 | 9.52E-01 | -7.16E-02 | -1.05E+00 |
| ENSG000001 | CABP1 | 3.68E-01 | ? | 7.37E-01 | -4.39E-01 | -1.36E+00 |
| ENSG000002 | --- | 3.68E-01 | ? | 1.38E+00 | 4.67E-01 | 1.38E+00 |
| ENSG000002 | C17orf107 | 3.68E-01 | 7.08E-01 | 8.76E-01 | -1.90E-01 | -1.14E+00 |
| ENSG000001 | PKD1P3 | 3.68E-01 | 7.08E-01 | 8.34E-01 | -2.62E-01 | -1.20E+00 |
| ENSG000002 | LINC00205 | 3.68E-01 | 7.08E-01 | 1.06E+00 | 8.82E-02 | 1.06E+00 |
| ENSG000002 | --- | 3.68E-01 | ? | 7.82E-01 | -3.56E-01 | -1.28E+00 |
| ENSG000002 | --- | 3.68E-01 | 7.08E-01 | 1.14E+00 | 1.95E-01 | 1.14E+00 |
| ENSG000002 | --- | 3.68E-01 | ? | 8.29E-01 | -2.71E-01 | -1.21E+00 |
| ENSG000001 | USP14 | 3.68E-01 | 7.08E-01 | 9.74E-01 | -3.84E-02 | -1.03E+00 |
| ENSG000001 | NUAK2 | 3.68E-01 | 7.08E-01 | 1.05E+00 | 7.43E-02 | 1.05E+00 |
| ENSG000002 | --- | 3.68E-01 | 7.08E-01 | 1.07E+00 | 1.01E-01 | 1.07E+00 |
| ENSG000002 | --- | 3.68E-01 | ? | 8.27E-01 | -2.74E-01 | -1.21E+00 |
| ENSG000001 | FOXA2 | 3.68E-01 | 7.08E-01 | 9.54E-01 | -6.82E-02 | -1.05E+00 |
| ENSG000002 | AKAP1-DT | 3.68E-01 | ? | 1.20E+00 | 2.68E-01 | 1.20E+00 |
| ENSG000002 | --- | 3.68E-01 | ? | 1.17E+00 | 2.30E-01 | 1.17E+00 |
| ENSG000001 | PDXDC1 | 3.69E-01 | 7.09E-01 | 1.03E+00 | 3.89E-02 | 1.03E+00 |
| ENSG000001 | PLEK2 | 3.69E-01 | 7.09E-01 | 9.66E-01 | -4.97E-02 | -1.04E+00 |
| ENSG000001 | DHX58 | 3.69E-01 | 7.09E-01 | 1.11E+00 | 1.53E-01 | 1.11E+00 |
| ENSG000002 | KLK9 | 3.69E-01 | ? | 6.66E-01 | -5.87E-01 | -1.50E+00 |
| ENSG00000C | IL17RB | 3.69E-01 | 7.09E-01 | 1.11E+00 | 1.50E-01 | 1.11E+00 |
| ENSG00000C | KITLG | 3.69E-01 | 7.09E-01 | 1.05E+00 | 6.50E-02 | 1.05E+00 |
| ENSG000002 | KLRK1-AS1 | 3.69E-01 | 7.09E-01 | 1.07E+00 | 9.97E-02 | 1.07E+00 |
| ENSG000002 | LCAL1 | 3.69E-01 | 7.09E-01 | 9.52E-01 | -7.17E-02 | -1.05E+00 |
| ENSG000002 | GABPB1-AS1 | 3.69E-01 | 7.09E-01 | 9.04E-01 | -1.45E-01 | -1.11E+00 |
| ENSG000002 | --- | 3.69E-01 | ? | 1.24E+00 | 3.09E-01 | 1.24E+00 |
| ENSG000001 | TLE6 | 3.69E-01 | ? | 7.49E-01 | -4.17E-01 | -1.33E+00 |
| ENSG000001 | PSMG2 | 3.69E-01 | 7.09E-01 | 9.74E-01 | -3.73E-02 | -1.03E+00 |
| ENSG000002 | TMEM147-A: | 3.69E-01 | 7.09E-01 | 1.06E+00 | 9.06E-02 | 1.06E+00 |
| ENSG000001 | NABP2 | 3.69E-01 | 7.09E-01 | 1.03E+00 | 4.76E-02 | 1.03E+00 |
| ENSG000001 | PCYT1A | 3.69E-01 | 7.09E-01 | 1.02E+00 | 2.79E-02 | 1.02E+00 |
| ENSG000001 | MUC6 | 3.69E-01 | ? | 8.19E-01 | -2.87E-01 | -1.22E+00 |
| ENSG000002 | --- | 3.69E-01 | 7.09E-01 | 1.24E+00 | 3.11E-01 | 1.24E+00 |
| ENSG00000C | FSTL3 | 3.70E-01 | 7.09E-01 | 9.55E-01 | -6.72E-02 | -1.05E+00 |
| ENSG000001 | ZNF485 | 3.70E-01 | 7.10E-01 | 1.06E+00 | 8.13E-02 | 1.06E+00 |
| ENSG000002 | HNRNPA3P5 | 3.70E-01 | ? | 1.50E+00 | 5.84E-01 | 1.50E+00 |
| ENSG000001 | ZFYVE27 | 3.70E-01 | 7.10E-01 | 9.49E-01 | -7.55E-02 | -1.05E+00 |
| ENSG00000C | TRPM3 | 3.70E-01 | ? | 1.76E+00 | 8.17E-01 | 1.76E+00 |
| ENSG00000C | TBC1D1 | 3.70E-01 | 7.10E-01 | 9.62E-01 | -5.52E-02 | -1.04E+00 |
| ENSG000001 | SLC26A9 | 3.70E-01 | ? | 1.62E+00 | 6.93E-01 | 1.62E+00 |

| | | | | | | | |
|---|---|---|---|---|---|---|---|
| ENSG000001 | CORO6 | 3.70E-01 | | 7.10E-01 | 1.11E+00 | 1.55E-01 | 1.11E+00 |
| ENSG000001 | STRN | 3.70E-01 | | 7.10E-01 | 9.63E-01 | -5.49E-02 | -1.04E+00 |
| ENSG000002 | --- | 3.70E-01 | | 7.10E-01 | 1.13E+00 | 1.75E-01 | 1.13E+00 |
| ENSG000002 | --- | 3.70E-01 | ? | | 8.13E-01 | -2.99E-01 | -1.23E+00 |
| ENSG000001 | USP1 | 3.70E-01 | | 7.10E-01 | 9.58E-01 | -6.23E-02 | -1.04E+00 |
| ENSG000002 | --- | 3.70E-01 | ? | | 1.19E+00 | 2.52E-01 | 1.19E+00 |
| ENSG000001 | RPS7 | 3.70E-01 | | 7.10E-01 | 1.03E+00 | 3.66E-02 | 1.03E+00 |
| ENSG000002 | --- | 3.70E-01 | ? | | 8.02E-01 | -3.18E-01 | -1.25E+00 |
| ENSG000002 | --- | 3.70E-01 | ? | | 6.30E-01 | -6.67E-01 | -1.59E+00 |
| ENSG000002 | LINC01977 | 3.70E-01 | ? | | 6.51E-01 | -6.19E-01 | -1.54E+00 |
| ENSG000002 | PXN-AS1 | 3.70E-01 | | 7.10E-01 | 9.21E-01 | -1.18E-01 | -1.09E+00 |
| ENSG000002 | ACOXL-AS1 | 3.70E-01 | ? | | 1.46E+00 | 5.48E-01 | 1.46E+00 |
| ENSG000001 | B4GALNT4 | 3.70E-01 | | 7.10E-01 | 1.07E+00 | 1.01E-01 | 1.07E+00 |
| ENSG000001 | KIDINS220 | 3.70E-01 | | 7.10E-01 | 9.69E-01 | -4.48E-02 | -1.03E+00 |
| ENSG000001 | BDKRB1 | 3.70E-01 | | 7.10E-01 | 1.14E+00 | 1.85E-01 | 1.14E+00 |
| ENSG000001 | ID2 | 3.71E-01 | ? | | 8.17E-01 | -2.92E-01 | -1.22E+00 |
| ENSG000002 | RUNDC3A-AS | 3.71E-01 | | 7.10E-01 | 1.06E+00 | 8.59E-02 | 1.06E+00 |
| ENSG000001 | NOLC1 | 3.71E-01 | | 7.10E-01 | 9.69E-01 | -4.57E-02 | -1.03E+00 |
| ENSG000001 | TNXB | 3.71E-01 | ? | | 1.20E+00 | 2.66E-01 | 1.20E+00 |
| ENSG000002 | GET4 | 3.71E-01 | | 7.10E-01 | 1.05E+00 | 6.70E-02 | 1.05E+00 |
| ENSG000001 | ABHD18 | 3.71E-01 | | 7.10E-01 | 9.53E-01 | -6.94E-02 | -1.05E+00 |
| ENSG000001 | CALM2 | 3.71E-01 | | 7.10E-01 | 9.74E-01 | -3.77E-02 | -1.03E+00 |
| ENSG000002 | CTTN-DT | 3.71E-01 | ? | | 7.59E-01 | -3.98E-01 | -1.32E+00 |
| ENSG000001 | RPS24 | 3.71E-01 | | 7.10E-01 | 1.04E+00 | 5.64E-02 | 1.04E+00 |
| ENSG000001 | POC1A | 3.71E-01 | | 7.10E-01 | 1.08E+00 | 1.05E-01 | 1.08E+00 |
| ENSG000002 | --- | 3.71E-01 | ? | | 1.20E+00 | 2.69E-01 | 1.20E+00 |
| ENSG000001 | ZMYND15 | 3.71E-01 | ? | | 8.32E-01 | -2.65E-01 | -1.20E+00 |
| ENSG000001 | PLAC8L1 | 3.71E-01 | ? | | 7.80E-01 | -3.59E-01 | -1.28E+00 |
| ENSG000001 | ESCO2 | 3.71E-01 | | 7.11E-01 | 8.90E-01 | -1.67E-01 | -1.12E+00 |
| ENSG000002 | --- | 3.71E-01 | ? | | 1.61E+00 | 6.88E-01 | 1.61E+00 |
| ENSG000001 | PPP4R1 | 3.71E-01 | | 7.11E-01 | 9.80E-01 | -2.90E-02 | -1.02E+00 |
| ENSG00000C | RARB | 3.71E-01 | ? | | 1.35E+00 | 4.28E-01 | 1.35E+00 |
| ENSG000002 | --- | 3.72E-01 | ? | | 7.66E-01 | -3.86E-01 | -1.31E+00 |
| ENSG000002 | --- | 3.72E-01 | ? | | 1.38E+00 | 4.70E-01 | 1.38E+00 |
| ENSG000001 | ELMO3 | 3.72E-01 | | 7.11E-01 | 1.10E+00 | 1.34E-01 | 1.10E+00 |
| ENSG000001 | NR2F6 | 3.72E-01 | | 7.12E-01 | 1.08E+00 | 1.10E-01 | 1.08E+00 |
| ENSG00000C | ZDHHC6 | 3.72E-01 | | 7.12E-01 | 9.52E-01 | -7.13E-02 | -1.05E+00 |
| ENSG000001 | KIF1C | 3.72E-01 | | 7.12E-01 | 1.02E+00 | 2.76E-02 | 1.02E+00 |
| ENSG000001 | SETD3 | 3.72E-01 | | 7.12E-01 | 1.03E+00 | 3.92E-02 | 1.03E+00 |
| ENSG00000C | TUBG2 | 3.72E-01 | | 7.12E-01 | 1.05E+00 | 7.64E-02 | 1.05E+00 |
| ENSG000001 | B3GALT6 | 3.72E-01 | | 7.12E-01 | 1.05E+00 | 6.56E-02 | 1.05E+00 |
| ENSG000001 | DRC3 | 3.72E-01 | | 7.12E-01 | 8.93E-01 | -1.64E-01 | -1.12E+00 |
| ENSG000002 | MICB-DT | 3.72E-01 | ? | | 1.22E+00 | 2.84E-01 | 1.22E+00 |
| ENSG000002 | CUTALP | 3.72E-01 | | 7.12E-01 | 1.06E+00 | 8.63E-02 | 1.06E+00 |
| ENSG000001 | SYNC | 3.72E-01 | | 7.12E-01 | 1.04E+00 | 5.30E-02 | 1.04E+00 |

| | | | | | | |
|---|---|---|---|---|---|---|
| ENSG000001 | SUSD4 | 3.72E-01 | 7.12E-01 | 1.08E+00 | 1.16E-01 | 1.08E+00 |
| ENSG00000C | PTGS1 | 3.72E-01 | 7.12E-01 | 9.49E-01 | -7.55E-02 | -1.05E+00 |
| ENSG000002 | --- | 3.73E-01 | ? | 1.46E+00 | 5.47E-01 | 1.46E+00 |
| ENSG000001 | SYAP1 | 3.73E-01 | 7.12E-01 | 1.03E+00 | 4.25E-02 | 1.03E+00 |
| ENSG000002 | BNIP3P4 | 3.73E-01 | ? | 1.62E+00 | 6.96E-01 | 1.62E+00 |
| ENSG00000C | BCL3 | 3.73E-01 | 7.12E-01 | 9.35E-01 | -9.66E-02 | -1.07E+00 |
| ENSG000001 | CCDC9B | 3.73E-01 | 7.12E-01 | 9.49E-01 | -7.60E-02 | -1.05E+00 |
| ENSG000001 | FUBP1 | 3.73E-01 | 7.12E-01 | 9.64E-01 | -5.26E-02 | -1.04E+00 |
| ENSG000001 | LPAR1 | 3.73E-01 | 7.12E-01 | 1.04E+00 | 5.30E-02 | 1.04E+00 |
| ENSG000001 | BUB1 | 3.73E-01 | 7.12E-01 | 9.22E-01 | -1.18E-01 | -1.08E+00 |
| ENSG000002 | HLA-B | 3.73E-01 | 7.12E-01 | 1.07E+00 | 9.94E-02 | 1.07E+00 |
| ENSG000002 | ZNF551 | 3.73E-01 | 7.12E-01 | 1.04E+00 | 5.64E-02 | 1.04E+00 |
| ENSG000001 | TMEM182 | 3.73E-01 | 7.12E-01 | 9.28E-01 | -1.07E-01 | -1.08E+00 |
| ENSG000001 | REC8 | 3.73E-01 | 7.12E-01 | 8.96E-01 | -1.59E-01 | -1.12E+00 |
| ENSG000002 | MICALL2-DT | 3.73E-01 | ? | 1.47E+00 | 5.60E-01 | 1.47E+00 |
| ENSG000001 | TP53I13 | 3.73E-01 | 7.12E-01 | 1.07E+00 | 9.25E-02 | 1.07E+00 |
| ENSG000002 | SEC63P1 | 3.73E-01 | ? | 8.32E-01 | -2.66E-01 | -1.20E+00 |
| ENSG000001 | POLR2K | 3.73E-01 | 7.12E-01 | 9.68E-01 | -4.67E-02 | -1.03E+00 |
| ENSG00000C | ABCB11 | 3.73E-01 | ? | 8.20E-01 | -2.87E-01 | -1.22E+00 |
| ENSG000001 | NACAD | 3.73E-01 | 7.12E-01 | 9.50E-01 | -7.44E-02 | -1.05E+00 |
| ENSG000002 | --- | 3.73E-01 | 7.12E-01 | 8.57E-01 | -2.23E-01 | -1.17E+00 |
| ENSG000002 | NKAPP1 | 3.73E-01 | ? | 7.91E-01 | -3.37E-01 | -1.26E+00 |
| ENSG000001 | ZC3H12D | 3.73E-01 | ? | 6.50E-01 | -6.21E-01 | -1.54E+00 |
| ENSG000002 | HCG18 | 3.73E-01 | 7.12E-01 | 9.68E-01 | -4.73E-02 | -1.03E+00 |
| ENSG000001 | RCBTB1 | 3.74E-01 | 7.12E-01 | 1.04E+00 | 5.01E-02 | 1.04E+00 |
| ENSG000001 | ARHGAP24 | 3.74E-01 | 7.12E-01 | 9.22E-01 | -1.16E-01 | -1.08E+00 |
| ENSG000001 | TEKTIP1 | 3.74E-01 | ? | 1.38E+00 | 4.60E-01 | 1.38E+00 |
| ENSG000001 | AARS2 | 3.74E-01 | 7.13E-01 | 1.04E+00 | 6.10E-02 | 1.04E+00 |
| ENSG000002 | ZNF818P | 3.74E-01 | 7.13E-01 | 1.07E+00 | 9.51E-02 | 1.07E+00 |
| ENSG000001 | EIF4A3 | 3.74E-01 | 7.13E-01 | 1.04E+00 | 5.37E-02 | 1.04E+00 |
| ENSG000001 | ZNF677 | 3.74E-01 | 7.13E-01 | 1.05E+00 | 6.53E-02 | 1.05E+00 |
| ENSG000002 | --- | 3.74E-01 | ? | 7.11E-01 | -4.92E-01 | -1.41E+00 |
| ENSG000001 | DNAAF11 | 3.74E-01 | 7.13E-01 | 8.77E-01 | -1.89E-01 | -1.14E+00 |
| ENSG000001 | MFAP3L | 3.74E-01 | 7.13E-01 | 9.42E-01 | -8.64E-02 | -1.06E+00 |
| ENSG000002 | ACTL10 | 3.74E-01 | ? | 1.24E+00 | 3.14E-01 | 1.24E+00 |
| ENSG000001 | DEPDC5 | 3.74E-01 | 7.13E-01 | 1.04E+00 | 6.29E-02 | 1.04E+00 |
| ENSG000001 | CTBP1 | 3.74E-01 | 7.13E-01 | 1.05E+00 | 6.83E-02 | 1.05E+00 |
| ENSG000002 | --- | 3.74E-01 | ? | 7.35E-01 | -4.43E-01 | -1.36E+00 |
| ENSG00000C | UFL1 | 3.74E-01 | 7.13E-01 | 9.62E-01 | -5.52E-02 | -1.04E+00 |
| ENSG000001 | ACTBP7 | 3.74E-01 | ? | 1.67E+00 | 7.39E-01 | 1.67E+00 |
| ENSG000002 | CENPS-CORT | 3.74E-01 | ? | 8.04E-01 | -3.14E-01 | -1.24E+00 |
| ENSG000001 | LRIT3 | 3.74E-01 | ? | 7.01E-01 | -5.11E-01 | -1.43E+00 |
| ENSG000002 | GPANK1 | 3.74E-01 | 7.13E-01 | 1.06E+00 | 7.95E-02 | 1.06E+00 |
| ENSG000001 | PPP1R37 | 3.74E-01 | 7.13E-01 | 1.08E+00 | 1.09E-01 | 1.08E+00 |
| ENSG000001 | LPP | 3.74E-01 | 7.13E-01 | 9.45E-01 | -8.11E-02 | -1.06E+00 |

| | | | | | | |
|---|---|---|---|---|---|---|
| ENSG000002 | --- | 3.74E-01 | 7.13E-01 | 1.12E+00 | 1.69E-01 | 1.12E+00 |
| ENSG000001 | SERF1A | 3.75E-01 | 7.13E-01 | 1.05E+00 | 6.61E-02 | 1.05E+00 |
| ENSG000002 | SNHG22 | 3.75E-01 | 7.13E-01 | 8.85E-01 | -1.76E-01 | -1.13E+00 |
| ENSG000001 | CCDC47 | 3.75E-01 | 7.13E-01 | 9.74E-01 | -3.78E-02 | -1.03E+00 |
| ENSG000001 | ZNF792 | 3.75E-01 | 7.13E-01 | 1.07E+00 | 9.40E-02 | 1.07E+00 |
| ENSG000002 | --- | 3.75E-01 | ? | 1.74E+00 | 8.02E-01 | 1.74E+00 |
| ENSG000001 | ZNF511 | 3.75E-01 | 7.13E-01 | 9.45E-01 | -8.21E-02 | -1.06E+00 |
| ENSG000001 | CD160 | 3.75E-01 | ? | 8.46E-01 | -2.41E-01 | -1.18E+00 |
| ENSG000001 | UBAP2L | 3.75E-01 | 7.13E-01 | 9.71E-01 | -4.22E-02 | -1.03E+00 |
| ENSG000001 | SETMAR | 3.75E-01 | 7.13E-01 | 1.06E+00 | 7.78E-02 | 1.06E+00 |
| ENSG000002 | --- | 3.75E-01 | ? | 7.67E-01 | -3.83E-01 | -1.30E+00 |
| ENSG000001 | HRH1 | 3.75E-01 | 7.13E-01 | 9.70E-01 | -4.44E-02 | -1.03E+00 |
| ENSG000001 | MATCAP2 | 3.75E-01 | 7.13E-01 | 9.48E-01 | -7.76E-02 | -1.06E+00 |
| ENSG000001 | DCHS2 | 3.75E-01 | ? | 8.02E-01 | -3.18E-01 | -1.25E+00 |
| ENSG000001 | ODAPH | 3.75E-01 | ? | 8.31E-01 | -2.67E-01 | -1.20E+00 |
| ENSG000002 | --- | 3.75E-01 | ? | 7.84E-01 | -3.52E-01 | -1.28E+00 |
| ENSG00000C | VRK2 | 3.75E-01 | 7.14E-01 | 9.60E-01 | -5.93E-02 | -1.04E+00 |
| ENSG000002 | PAICSP4 | 3.75E-01 | ? | 1.49E+00 | 5.77E-01 | 1.49E+00 |
| ENSG00000C | PSD | 3.76E-01 | ? | 1.27E+00 | 3.39E-01 | 1.27E+00 |
| ENSG000002 | ABHD16A | 3.76E-01 | 7.14E-01 | 1.06E+00 | 8.65E-02 | 1.06E+00 |
| ENSG000001 | PAIP1 | 3.76E-01 | 7.14E-01 | 9.67E-01 | -4.86E-02 | -1.03E+00 |
| ENSG000001 | TELO2 | 3.76E-01 | 7.14E-01 | 1.11E+00 | 1.49E-01 | 1.11E+00 |
| ENSG000002 | --- | 3.76E-01 | ? | 1.43E+00 | 5.17E-01 | 1.43E+00 |
| ENSG00000C | MRE11 | 3.76E-01 | 7.14E-01 | 9.64E-01 | -5.29E-02 | -1.04E+00 |
| ENSG000001 | KATNA1 | 3.76E-01 | 7.14E-01 | 1.04E+00 | 5.17E-02 | 1.04E+00 |
| ENSG000001 | HS6ST2 | 3.76E-01 | 7.14E-01 | 1.06E+00 | 8.06E-02 | 1.06E+00 |
| ENSG00000C | CRMP1 | 3.76E-01 | 7.14E-01 | 9.23E-01 | -1.16E-01 | -1.08E+00 |
| ENSG000002 | --- | 3.76E-01 | ? | 7.75E-01 | -3.68E-01 | -1.29E+00 |
| ENSG000002 | --- | 3.76E-01 | ? | 8.27E-01 | -2.74E-01 | -1.21E+00 |
| ENSG000002 | PCDHAC2 | 3.76E-01 | 7.14E-01 | 1.06E+00 | 8.46E-02 | 1.06E+00 |
| ENSG000002 | BRK1 | 3.76E-01 | 7.14E-01 | 1.02E+00 | 3.10E-02 | 1.02E+00 |
| ENSG000001 | IKZF3 | 3.76E-01 | ? | 1.18E+00 | 2.43E-01 | 1.18E+00 |
| ENSG000002 | GDI1 | 3.76E-01 | 7.14E-01 | 9.71E-01 | -4.18E-02 | -1.03E+00 |
| ENSG000002 | LMCD1-AS1 | 3.76E-01 | ? | 1.35E+00 | 4.28E-01 | 1.35E+00 |
| ENSG000001 | USP20 | 3.76E-01 | 7.14E-01 | 1.04E+00 | 5.61E-02 | 1.04E+00 |
| ENSG000001 | TMEM201 | 3.76E-01 | 7.14E-01 | 1.06E+00 | 8.83E-02 | 1.06E+00 |
| ENSG000001 | PTPA | 3.76E-01 | 7.14E-01 | 1.05E+00 | 7.27E-02 | 1.05E+00 |
| ENSG000001 | ARID5A | 3.76E-01 | 7.14E-01 | 1.09E+00 | 1.30E-01 | 1.09E+00 |
| ENSG000001 | CEP19 | 3.76E-01 | 7.14E-01 | 1.09E+00 | 1.20E-01 | 1.09E+00 |
| ENSG000001 | PNN | 3.76E-01 | 7.14E-01 | 9.45E-01 | -8.10E-02 | -1.06E+00 |
| ENSG000001 | RCCD1 | 3.76E-01 | 7.14E-01 | 1.05E+00 | 7.38E-02 | 1.05E+00 |
| ENSG000002 | LINC02701 | 3.77E-01 | ? | 1.36E+00 | 4.47E-01 | 1.36E+00 |
| ENSG00000C | HEXB | 3.77E-01 | 7.14E-01 | 1.04E+00 | 5.40E-02 | 1.04E+00 |
| ENSG000001 | OPRL1 | 3.77E-01 | ? | 7.81E-01 | -3.57E-01 | -1.28E+00 |
| ENSG000001 | RNF128 | 3.77E-01 | 7.14E-01 | 1.05E+00 | 7.40E-02 | 1.05E+00 |

| | | | | | | |
|---|---|---|---|---|---|---|
| ENSG000001 | CLMP | 3.77E-01 | 7.14E-01 | 9.28E-01 | -1.08E-01 | -1.08E+00 |
| ENSG000001 | NCOR2 | 3.77E-01 | 7.14E-01 | 9.61E-01 | -5.78E-02 | -1.04E+00 |
| ENSG000002 | --- | 3.77E-01 | ? | 7.59E-01 | -3.98E-01 | -1.32E+00 |
| ENSG000001 | PRKRIP1 | 3.77E-01 | 7.14E-01 | 9.57E-01 | -6.39E-02 | -1.05E+00 |
| ENSG000001 | RAB3D | 3.77E-01 | 7.14E-01 | 1.05E+00 | 7.06E-02 | 1.05E+00 |
| ENSG000001 | ATP5F1C | 3.77E-01 | 7.14E-01 | 1.02E+00 | 3.04E-02 | 1.02E+00 |
| ENSG000001 | HMGXB3 | 3.77E-01 | 7.14E-01 | 1.03E+00 | 3.91E-02 | 1.03E+00 |
| ENSG000002 | RPS7P1 | 3.77E-01 | 7.14E-01 | 1.05E+00 | 7.20E-02 | 1.05E+00 |
| ENSG000002 | --- | 3.77E-01 | ? | 7.47E-01 | -4.20E-01 | -1.34E+00 |
| ENSG000001 | RBBP8 | 3.77E-01 | 7.14E-01 | 1.03E+00 | 4.52E-02 | 1.03E+00 |
| ENSG000002 | --- | 3.77E-01 | 7.14E-01 | 8.94E-01 | -1.61E-01 | -1.12E+00 |
| ENSG000002 | ISY1 | 3.77E-01 | 7.14E-01 | 1.04E+00 | 6.08E-02 | 1.04E+00 |
| ENSG000001 | COQ10B | 3.77E-01 | 7.14E-01 | 1.03E+00 | 4.29E-02 | 1.03E+00 |
| ENSG000001 | IDUA | 3.77E-01 | 7.14E-01 | 1.08E+00 | 1.10E-01 | 1.08E+00 |
| ENSG000002 | CCDC71L | 3.77E-01 | 7.14E-01 | 1.05E+00 | 6.79E-02 | 1.05E+00 |
| ENSG000001 | CHRNA5 | 3.77E-01 | 7.14E-01 | 8.66E-01 | -2.08E-01 | -1.16E+00 |
| ENSG000001 | CLDN1 | 3.78E-01 | 7.14E-01 | 1.04E+00 | 6.26E-02 | 1.04E+00 |
| ENSG000001 | CCDC154 | 3.78E-01 | ? | 7.19E-01 | -4.76E-01 | -1.39E+00 |
| ENSG000001 | LBHD1 | 3.78E-01 | 7.15E-01 | 9.19E-01 | -1.22E-01 | -1.09E+00 |
| ENSG000001 | FSIP1 | 3.78E-01 | 7.15E-01 | 1.16E+00 | 2.13E-01 | 1.16E+00 |
| ENSG000002 | CHUK | 3.78E-01 | 7.15E-01 | 9.69E-01 | -4.50E-02 | -1.03E+00 |
| ENSG000001 | LHFPL6 | 3.78E-01 | 7.15E-01 | 1.04E+00 | 6.25E-02 | 1.04E+00 |
| ENSG000001 | MBD2 | 3.78E-01 | 7.15E-01 | 9.65E-01 | -5.14E-02 | -1.04E+00 |
| ENSG000002 | WASH8P | 3.78E-01 | 7.15E-01 | 9.43E-01 | -8.53E-02 | -1.06E+00 |
| ENSG000002 | --- | 3.78E-01 | ? | 7.78E-01 | -3.62E-01 | -1.28E+00 |
| ENSG000001 | LENG1 | 3.78E-01 | 7.15E-01 | 9.37E-01 | -9.38E-02 | -1.07E+00 |
| ENSG000001 | CRNN | 3.78E-01 | ? | 1.59E+00 | 6.65E-01 | 1.59E+00 |
| ENSG000002 | MIR210HG | 3.78E-01 | 7.15E-01 | 1.08E+00 | 1.05E-01 | 1.08E+00 |
| ENSG000002 | LINC02454 | 3.78E-01 | 7.15E-01 | 1.06E+00 | 8.04E-02 | 1.06E+00 |
| ENSG000001 | ITGA4 | 3.78E-01 | 7.15E-01 | 1.07E+00 | 9.98E-02 | 1.07E+00 |
| ENSG000001 | GPAT4 | 3.78E-01 | 7.15E-01 | 9.79E-01 | -3.00E-02 | -1.02E+00 |
| ENSG000002 | APTR | 3.78E-01 | 7.15E-01 | 1.09E+00 | 1.23E-01 | 1.09E+00 |
| ENSG000002 | HSP90AA3P | 3.78E-01 | ? | 1.27E+00 | 3.45E-01 | 1.27E+00 |
| ENSG000002 | PNISR-AS1 | 3.78E-01 | 7.15E-01 | 1.09E+00 | 1.31E-01 | 1.09E+00 |
| ENSG000001 | IGFLR1 | 3.78E-01 | ? | 1.54E+00 | 6.22E-01 | 1.54E+00 |
| ENSG000001 | WNT10A | 3.79E-01 | ? | 1.46E+00 | 5.41E-01 | 1.46E+00 |
| ENSG000001 | RPL10AP2 | 3.79E-01 | ? | 1.70E+00 | 7.67E-01 | 1.70E+00 |
| ENSG00000C | SERPIND1 | 3.79E-01 | ? | 7.71E-01 | -3.76E-01 | -1.30E+00 |
| ENSG000001 | MOSPD1 | 3.79E-01 | 7.15E-01 | 1.03E+00 | 4.67E-02 | 1.03E+00 |
| ENSG00000C | WDR3 | 3.79E-01 | 7.15E-01 | 1.03E+00 | 4.74E-02 | 1.03E+00 |
| ENSG00000C | CELSR1 | 3.79E-01 | 7.15E-01 | 1.04E+00 | 5.17E-02 | 1.04E+00 |
| ENSG000001 | IQCH | 3.79E-01 | 7.15E-01 | 1.11E+00 | 1.51E-01 | 1.11E+00 |
| ENSG000002 | --- | 3.79E-01 | 7.15E-01 | 9.31E-01 | -1.03E-01 | -1.07E+00 |
| ENSG000002 | TAF15 | 3.79E-01 | 7.15E-01 | 9.69E-01 | -4.58E-02 | -1.03E+00 |
| ENSG000002 | HNRNPCP2 | 3.79E-01 | 7.15E-01 | 8.80E-01 | -1.85E-01 | -1.14E+00 |

| | | | | | | |
|---|---|---|---|---|---|---|
| ENSG000001 | ATP7A | 3.79E-01 | 7.15E-01 | 9.59E-01 | -5.98E-02 | -1.04E+00 |
| ENSG000002 | CHIC1 | 3.79E-01 | 7.15E-01 | 1.05E+00 | 7.47E-02 | 1.05E+00 |
| ENSG000001 | BAZ2B | 3.79E-01 | 7.15E-01 | 9.58E-01 | -6.26E-02 | -1.04E+00 |
| ENSG000002 | --- | 3.79E-01 | 7.15E-01 | 7.96E-01 | -3.30E-01 | -1.26E+00 |
| ENSG000002 | --- | 3.79E-01 | ? | 1.39E+00 | 4.76E-01 | 1.39E+00 |
| ENSG000002 | --- | 3.79E-01 | 7.15E-01 | 9.11E-01 | -1.35E-01 | -1.10E+00 |
| ENSG000001 | FAM83E | 3.79E-01 | 7.15E-01 | 8.94E-01 | -1.61E-01 | -1.12E+00 |
| ENSG000002 | C2CD4D-AS1 | 3.79E-01 | ? | 7.70E-01 | -3.78E-01 | -1.30E+00 |
| ENSG000001 | ZSCAN29 | 3.79E-01 | 7.15E-01 | 9.71E-01 | -4.21E-02 | -1.03E+00 |
| ENSG000001 | STX18 | 3.79E-01 | 7.15E-01 | 9.68E-01 | -4.67E-02 | -1.03E+00 |
| ENSG000001 | SLC22A5 | 3.80E-01 | 7.16E-01 | 9.60E-01 | -5.86E-02 | -1.04E+00 |
| ENSG000001 | TUBB4A | 3.80E-01 | ? | 5.45E-01 | -8.76E-01 | -1.84E+00 |
| ENSG000001 | FBXO34 | 3.80E-01 | 7.16E-01 | 1.03E+00 | 4.89E-02 | 1.03E+00 |
| ENSG00000C | RIF1 | 3.80E-01 | 7.16E-01 | 9.53E-01 | -7.01E-02 | -1.05E+00 |
| ENSG000002 | MUC12-AS1 | 3.80E-01 | ? | 1.48E+00 | 5.67E-01 | 1.48E+00 |
| ENSG000001 | ATAD3B | 3.80E-01 | 7.16E-01 | 9.47E-01 | -7.84E-02 | -1.06E+00 |
| ENSG000001 | GOLIM4 | 3.80E-01 | 7.16E-01 | 9.63E-01 | -5.48E-02 | -1.04E+00 |
| ENSG00000C | LNX1 | 3.80E-01 | ? | 8.27E-01 | -2.75E-01 | -1.21E+00 |
| ENSG000001 | PLA2G12A | 3.80E-01 | 7.16E-01 | 9.68E-01 | -4.72E-02 | -1.03E+00 |
| ENSG000002 | ZNF726 | 3.80E-01 | ? | 8.17E-01 | -2.91E-01 | -1.22E+00 |
| ENSG000001 | LRFN3 | 3.80E-01 | 7.16E-01 | 1.07E+00 | 9.11E-02 | 1.07E+00 |
| ENSG000001 | XRCC6 | 3.80E-01 | 7.16E-01 | 1.02E+00 | 2.75E-02 | 1.02E+00 |
| ENSG000001 | PTGES3L-AA | 3.80E-01 | ? | 7.18E-01 | -4.78E-01 | -1.39E+00 |
| ENSG000001 | U2AF1 | 3.80E-01 | 7.16E-01 | 9.52E-01 | -7.12E-02 | -1.05E+00 |
| ENSG000002 | CROCCP2 | 3.80E-01 | ? | 6.90E-01 | -5.35E-01 | -1.45E+00 |
| ENSG000001 | PTK6 | 3.80E-01 | 7.16E-01 | 9.34E-01 | -9.91E-02 | -1.07E+00 |
| ENSG000001 | FBXO30 | 3.80E-01 | 7.16E-01 | 9.65E-01 | -5.21E-02 | -1.04E+00 |
| ENSG000002 | --- | 3.80E-01 | ? | 1.53E+00 | 6.16E-01 | 1.53E+00 |
| ENSG000001 | ZNF841 | 3.80E-01 | 7.16E-01 | 9.47E-01 | -7.85E-02 | -1.06E+00 |
| ENSG000001 | THEM4 | 3.80E-01 | 7.16E-01 | 1.05E+00 | 6.58E-02 | 1.05E+00 |
| ENSG000001 | CBS | 3.80E-01 | 7.16E-01 | 1.22E+00 | 2.87E-01 | 1.22E+00 |
| ENSG000002 | MSANTD2-A: | 3.81E-01 | 7.16E-01 | 1.11E+00 | 1.47E-01 | 1.11E+00 |
| ENSG000001 | PAX8-AS1 | 3.81E-01 | 7.16E-01 | 1.05E+00 | 7.22E-02 | 1.05E+00 |
| ENSG000001 | MT-CYB | 3.81E-01 | 7.16E-01 | 9.16E-01 | -1.26E-01 | -1.09E+00 |
| ENSG000001 | CCSAP | 3.81E-01 | 7.16E-01 | 1.05E+00 | 6.98E-02 | 1.05E+00 |
| ENSG000002 | SNHG14 | 3.81E-01 | 7.16E-01 | 9.66E-01 | -5.06E-02 | -1.04E+00 |
| ENSG000001 | CNNM3 | 3.81E-01 | 7.16E-01 | 9.51E-01 | -7.30E-02 | -1.05E+00 |
| ENSG000002 | SDCBP2-AS1 | 3.81E-01 | 7.16E-01 | 9.02E-01 | -1.48E-01 | -1.11E+00 |
| ENSG000001 | SGF29 | 3.81E-01 | 7.16E-01 | 1.08E+00 | 1.10E-01 | 1.08E+00 |
| ENSG00000C | PLXNA2 | 3.81E-01 | 7.16E-01 | 9.71E-01 | -4.19E-02 | -1.03E+00 |
| ENSG000001 | NAV1 | 3.81E-01 | 7.16E-01 | 9.69E-01 | -4.50E-02 | -1.03E+00 |
| ENSG000001 | COL7A1 | 3.81E-01 | 7.16E-01 | 9.37E-01 | -9.40E-02 | -1.07E+00 |
| ENSG000002 | PINLYP | 3.81E-01 | 7.16E-01 | 8.64E-01 | -2.10E-01 | -1.16E+00 |
| ENSG00000C | GPBP1 | 3.81E-01 | 7.16E-01 | 9.71E-01 | -4.32E-02 | -1.03E+00 |
| ENSG000001 | NADSYN1 | 3.81E-01 | 7.16E-01 | 1.04E+00 | 5.78E-02 | 1.04E+00 |

| | | | | | | |
|---|---|---|---|---|---|---|
| ENSG000002 | KIAA1143P1 | 3.81E-01 | ? | 7.38E-01 | -4.38E-01 | -1.35E+00 |
| ENSG000002 | --- | 3.81E-01 | ? | 6.88E-01 | -5.39E-01 | -1.45E+00 |
| ENSG000001 | RAPGEF1 | 3.81E-01 | 7.16E-01 | 1.03E+00 | 4.13E-02 | 1.03E+00 |
| ENSG000001 | OXCT2 | 3.81E-01 | ? | 8.37E-01 | -2.57E-01 | -1.19E+00 |
| ENSG000002 | --- | 3.81E-01 | 7.16E-01 | 8.98E-01 | -1.55E-01 | -1.11E+00 |
| ENSG000002 | --- | 3.81E-01 | ? | 5.84E-01 | -7.76E-01 | -1.71E+00 |
| ENSG000001 | TMEM248 | 3.81E-01 | 7.16E-01 | 9.78E-01 | -3.18E-02 | -1.02E+00 |
| ENSG000002 | ZRANB2-DT | 3.82E-01 | ? | 7.96E-01 | -3.29E-01 | -1.26E+00 |
| ENSG000001 | ZNF266 | 3.82E-01 | 7.16E-01 | 9.63E-01 | -5.39E-02 | -1.04E+00 |
| ENSG000001 | LCA5 | 3.82E-01 | 7.16E-01 | 9.53E-01 | -7.02E-02 | -1.05E+00 |
| ENSG000001 | AGPAT3 | 3.82E-01 | 7.16E-01 | 1.03E+00 | 4.60E-02 | 1.03E+00 |
| ENSG000001 | SLC9A9 | 3.82E-01 | 7.16E-01 | 1.12E+00 | 1.59E-01 | 1.12E+00 |
| ENSG000001 | HIBCH | 3.82E-01 | 7.16E-01 | 1.04E+00 | 5.67E-02 | 1.04E+00 |
| ENSG000001 | PLPPR1 | 3.82E-01 | ? | 7.93E-01 | -3.35E-01 | -1.26E+00 |
| ENSG000001 | NEUROD2 | 3.82E-01 | ? | 1.22E+00 | 2.87E-01 | 1.22E+00 |
| ENSG00000C | DNAH5 | 3.82E-01 | 7.16E-01 | 9.42E-01 | -8.55E-02 | -1.06E+00 |
| ENSG000002 | LINC02742 | 3.82E-01 | 7.16E-01 | 8.78E-01 | -1.87E-01 | -1.14E+00 |
| ENSG000001 | PSMD2 | 3.82E-01 | 7.16E-01 | 1.03E+00 | 3.91E-02 | 1.03E+00 |
| ENSG000001 | ACAT2 | 3.82E-01 | 7.16E-01 | 1.04E+00 | 5.98E-02 | 1.04E+00 |
| ENSG000001 | TH | 3.82E-01 | ? | 8.17E-01 | -2.91E-01 | -1.22E+00 |
| ENSG000001 | ADAT2 | 3.82E-01 | 7.17E-01 | 1.05E+00 | 7.66E-02 | 1.05E+00 |
| ENSG000002 | AGAP7P | 3.82E-01 | ? | 6.75E-01 | -5.67E-01 | -1.48E+00 |
| ENSG000002 | AK3P3 | 3.82E-01 | ? | 1.39E+00 | 4.76E-01 | 1.39E+00 |
| ENSG000002 | H3C14 | 3.82E-01 | ? | 1.61E+00 | 6.83E-01 | 1.61E+00 |
| ENSG000002 | PPP5D1P | 3.82E-01 | ? | 1.16E+00 | 2.12E-01 | 1.16E+00 |
| ENSG00000C | POLR2B | 3.82E-01 | 7.17E-01 | 9.82E-01 | -2.66E-02 | -1.02E+00 |
| ENSG000002 | RPL18AP3 | 3.82E-01 | 7.17E-01 | 1.07E+00 | 9.17E-02 | 1.07E+00 |
| ENSG000001 | PPWD1 | 3.82E-01 | 7.17E-01 | 9.61E-01 | -5.78E-02 | -1.04E+00 |
| ENSG000002 | ELOCP19 | 3.82E-01 | ? | 1.27E+00 | 3.46E-01 | 1.27E+00 |
| ENSG00000C | RNF19A | 3.82E-01 | 7.17E-01 | 9.62E-01 | -5.62E-02 | -1.04E+00 |
| ENSG000001 | TIGD5 | 3.82E-01 | 7.17E-01 | 1.09E+00 | 1.24E-01 | 1.09E+00 |
| ENSG000001 | TRIM55 | 3.82E-01 | ? | 1.26E+00 | 3.32E-01 | 1.26E+00 |
| ENSG000001 | OXA1L | 3.82E-01 | 7.17E-01 | 1.02E+00 | 3.26E-02 | 1.02E+00 |
| ENSG000002 | LINC00452 | 3.83E-01 | ? | 8.08E-01 | -3.08E-01 | -1.24E+00 |
| ENSG000002 | --- | 3.83E-01 | 7.17E-01 | 9.19E-01 | -1.21E-01 | -1.09E+00 |
| ENSG000001 | ALKBH8 | 3.83E-01 | 7.17E-01 | 9.53E-01 | -6.96E-02 | -1.05E+00 |
| ENSG000002 | --- | 3.83E-01 | 7.17E-01 | 8.74E-01 | -1.94E-01 | -1.14E+00 |
| ENSG000002 | ISL1-DT | 3.83E-01 | ? | 8.20E-01 | -2.87E-01 | -1.22E+00 |
| ENSG000001 | ZKSCAN2 | 3.83E-01 | 7.17E-01 | 9.58E-01 | -6.14E-02 | -1.04E+00 |
| ENSG000001 | DHX40 | 3.83E-01 | 7.17E-01 | 1.03E+00 | 4.08E-02 | 1.03E+00 |
| ENSG00000C | INPP4A | 3.83E-01 | 7.18E-01 | 1.03E+00 | 3.77E-02 | 1.03E+00 |
| ENSG000001 | TOX | 3.83E-01 | ? | 7.40E-01 | -4.34E-01 | -1.35E+00 |
| ENSG00000C | SMG6 | 3.83E-01 | 7.18E-01 | 1.03E+00 | 3.64E-02 | 1.03E+00 |
| ENSG000002 | ZNF747-DT | 3.83E-01 | ? | 1.15E+00 | 2.03E-01 | 1.15E+00 |
| ENSG000002 | HSPA1B | 3.83E-01 | 7.18E-01 | 1.04E+00 | 6.22E-02 | 1.04E+00 |

| | | | | | | |
|---|---|---|---|---|---|---|
| ENSG000001 | ARF1 | 3.83E-01 | 7.18E-01 | 1.03E+00 | 4.25E-02 | 1.03E+00 |
| ENSG000001 | UNC119 | 3.84E-01 | 7.18E-01 | 1.04E+00 | 5.39E-02 | 1.04E+00 |
| ENSG000001 | PLAAT1 | 3.84E-01 | ? | 1.17E+00 | 2.23E-01 | 1.17E+00 |
| ENSG000002 | FAM72C | 3.84E-01 | 7.18E-01 | 8.11E-01 | -3.02E-01 | -1.23E+00 |
| ENSG000001 | TMEM205 | 3.84E-01 | 7.18E-01 | 1.06E+00 | 7.87E-02 | 1.06E+00 |
| ENSG000001 | MRPL9 | 3.84E-01 | 7.18E-01 | 9.72E-01 | -4.09E-02 | -1.03E+00 |
| ENSG000001 | PLEKHA3 | 3.84E-01 | 7.18E-01 | 1.04E+00 | 6.21E-02 | 1.04E+00 |
| ENSG000002 | --- | 3.84E-01 | ? | 1.20E+00 | 2.62E-01 | 1.20E+00 |
| ENSG000001 | CCT6A | 3.84E-01 | 7.18E-01 | 9.77E-01 | -3.43E-02 | -1.02E+00 |
| ENSG000001 | PMPCA | 3.84E-01 | 7.18E-01 | 1.03E+00 | 4.14E-02 | 1.03E+00 |
| ENSG000002 | --- | 3.84E-01 | ? | 4.69E-01 | -1.09E+00 | -2.13E+00 |
| ENSG000002 | --- | 3.84E-01 | ? | 6.86E-01 | -5.43E-01 | -1.46E+00 |
| ENSG000001 | ATAD2 | 3.84E-01 | 7.18E-01 | 9.44E-01 | -8.38E-02 | -1.06E+00 |
| ENSG000001 | KANK2 | 3.84E-01 | 7.18E-01 | 9.62E-01 | -5.54E-02 | -1.04E+00 |
| ENSG000002 | ARID3C | 3.84E-01 | ? | 1.43E+00 | 5.21E-01 | 1.43E+00 |
| ENSG000001 | CCDC69 | 3.84E-01 | 7.18E-01 | 9.43E-01 | -8.47E-02 | -1.06E+00 |
| ENSG000001 | MSANTD2 | 3.84E-01 | 7.18E-01 | 9.40E-01 | -9.00E-02 | -1.06E+00 |
| ENSG000001 | TMEM128 | 3.84E-01 | 7.18E-01 | 1.04E+00 | 6.04E-02 | 1.04E+00 |
| ENSG000001 | PPM1A | 3.84E-01 | 7.18E-01 | 9.74E-01 | -3.84E-02 | -1.03E+00 |
| ENSG000001 | PEX14 | 3.84E-01 | 7.18E-01 | 9.64E-01 | -5.29E-02 | -1.04E+00 |
| ENSG000001 | ETFDH | 3.84E-01 | 7.18E-01 | 1.05E+00 | 6.67E-02 | 1.05E+00 |
| ENSG000002 | IRAIN | 3.85E-01 | ? | 8.09E-01 | -3.06E-01 | -1.24E+00 |
| ENSG000001 | MMP21 | 3.85E-01 | ? | 7.50E-01 | -4.16E-01 | -1.33E+00 |
| ENSG000001 | RALGAPB | 3.85E-01 | 7.19E-01 | 9.75E-01 | -3.60E-02 | -1.03E+00 |
| ENSG000001 | ZFPM1 | 3.85E-01 | 7.19E-01 | 9.25E-01 | -1.13E-01 | -1.08E+00 |
| ENSG000002 | SDHAP4 | 3.85E-01 | 7.19E-01 | 9.33E-01 | -1.00E-01 | -1.07E+00 |
| ENSG000002 | MIDEAS-AS1 | 3.85E-01 | ? | 1.35E+00 | 4.34E-01 | 1.35E+00 |
| ENSG000002 | SARNP | 3.85E-01 | 7.19E-01 | 1.03E+00 | 4.47E-02 | 1.03E+00 |
| ENSG000001 | PSMA7 | 3.85E-01 | 7.19E-01 | 1.05E+00 | 7.37E-02 | 1.05E+00 |
| ENSG000002 | --- | 3.85E-01 | ? | 1.46E+00 | 5.41E-01 | 1.46E+00 |
| ENSG000001 | CSRNP3 | 3.85E-01 | 7.19E-01 | 8.95E-01 | -1.60E-01 | -1.12E+00 |
| ENSG000001 | LHX6 | 3.85E-01 | ? | 1.18E+00 | 2.42E-01 | 1.18E+00 |
| ENSG000002 | SIPA1 | 3.85E-01 | 7.19E-01 | 9.21E-01 | -1.19E-01 | -1.09E+00 |
| ENSG000001 | B3GNT8 | 3.85E-01 | 7.19E-01 | 1.10E+00 | 1.42E-01 | 1.10E+00 |
| ENSG000001 | PLD6 | 3.85E-01 | 7.19E-01 | 1.06E+00 | 8.92E-02 | 1.06E+00 |
| ENSG000001 | DOCK6 | 3.85E-01 | 7.19E-01 | 1.03E+00 | 4.32E-02 | 1.03E+00 |
| ENSG000001 | SETDB1 | 3.85E-01 | 7.19E-01 | 1.03E+00 | 4.51E-02 | 1.03E+00 |
| ENSG00000C | CDH1 | 3.85E-01 | 7.19E-01 | 9.81E-01 | -2.76E-02 | -1.02E+00 |
| ENSG000002 | --- | 3.86E-01 | 7.19E-01 | 9.00E-01 | -1.52E-01 | -1.11E+00 |
| ENSG000002 | --- | 3.86E-01 | ? | 1.43E+00 | 5.19E-01 | 1.43E+00 |
| ENSG000002 | --- | 3.86E-01 | ? | 1.46E+00 | 5.43E-01 | 1.46E+00 |
| ENSG00000C | PHLPP2 | 3.86E-01 | 7.19E-01 | 1.04E+00 | 5.74E-02 | 1.04E+00 |
| ENSG000001 | SPON2 | 3.86E-01 | 7.19E-01 | 8.63E-01 | -2.12E-01 | -1.16E+00 |
| ENSG00000C | RAB27A | 3.86E-01 | 7.19E-01 | 9.60E-01 | -5.86E-02 | -1.04E+00 |
| ENSG000001 | DET1 | 3.86E-01 | 7.19E-01 | 1.09E+00 | 1.25E-01 | 1.09E+00 |

| | | | | | | | |
|---|---|---|---|---|---|---|---|
| ENSG000001 | ZNF823 | 3.86E-01 | | 7.19E-01 | 1.07E+00 | 9.23E-02 | 1.07E+00 |
| ENSG000001 | INTS6L | 3.86E-01 | | 7.19E-01 | 1.05E+00 | 6.74E-02 | 1.05E+00 |
| ENSG000002 | MCPH1-AS1 | 3.86E-01 | | 7.19E-01 | 8.82E-01 | -1.81E-01 | -1.13E+00 |
| ENSG000001 | C11orf87 | 3.86E-01 | ? | | 1.49E+00 | 5.74E-01 | 1.49E+00 |
| ENSG000002 | --- | 3.86E-01 | | 7.19E-01 | 1.16E+00 | 2.15E-01 | 1.16E+00 |
| ENSG000001 | GLIS3 | 3.86E-01 | | 7.19E-01 | 9.13E-01 | -1.31E-01 | -1.09E+00 |
| ENSG000001 | USP6NL | 3.86E-01 | | 7.19E-01 | 1.04E+00 | 5.62E-02 | 1.04E+00 |
| ENSG000002 | NBAT1 | 3.86E-01 | ? | | 8.07E-01 | -3.09E-01 | -1.24E+00 |
| ENSG000001 | LSM1 | 3.86E-01 | | 7.19E-01 | 9.65E-01 | -5.21E-02 | -1.04E+00 |
| ENSG000001 | PDLIM2 | 3.86E-01 | | 7.19E-01 | 9.29E-01 | -1.06E-01 | -1.08E+00 |
| ENSG000002 | PCAT1 | 3.86E-01 | | 7.19E-01 | 1.14E+00 | 1.86E-01 | 1.14E+00 |
| ENSG000002 | --- | 3.87E-01 | ? | | 5.85E-01 | -7.73E-01 | -1.71E+00 |
| ENSG000002 | --- | 3.87E-01 | ? | | 6.83E-01 | -5.50E-01 | -1.46E+00 |
| ENSG000001 | QPCT | 3.87E-01 | | 7.20E-01 | 1.07E+00 | 9.85E-02 | 1.07E+00 |
| ENSG000002 | --- | 3.87E-01 | ? | | 8.18E-01 | -2.89E-01 | -1.22E+00 |
| ENSG000001 | PBX4 | 3.87E-01 | ? | | 1.39E+00 | 4.75E-01 | 1.39E+00 |
| ENSG000002 | --- | 3.87E-01 | ? | | 7.58E-01 | -3.99E-01 | -1.32E+00 |
| ENSG000001 | SEMA3D | 3.87E-01 | ? | | 7.95E-01 | -3.31E-01 | -1.26E+00 |
| ENSG000002 | C3orf36 | 3.87E-01 | ? | | 7.50E-01 | -4.14E-01 | -1.33E+00 |
| ENSG000001 | GLCCI1 | 3.87E-01 | | 7.20E-01 | 9.41E-01 | -8.77E-02 | -1.06E+00 |
| ENSG000002 | CKLF | 3.87E-01 | | 7.20E-01 | 1.08E+00 | 1.07E-01 | 1.08E+00 |
| ENSG000002 | TCEA1P2 | 3.87E-01 | | 7.20E-01 | 8.79E-01 | -1.87E-01 | -1.14E+00 |
| ENSG000001 | MPG | 3.87E-01 | | 7.20E-01 | 1.07E+00 | 9.34E-02 | 1.07E+00 |
| ENSG00000C | CD99 | 3.87E-01 | | 7.20E-01 | 9.72E-01 | -4.13E-02 | -1.03E+00 |
| ENSG000001 | KAT8 | 3.87E-01 | | 7.20E-01 | 1.03E+00 | 4.66E-02 | 1.03E+00 |
| ENSG000001 | XPR1 | 3.87E-01 | | 7.20E-01 | 1.03E+00 | 4.38E-02 | 1.03E+00 |
| ENSG000002 | --- | 3.87E-01 | ? | | 1.28E+00 | 3.60E-01 | 1.28E+00 |
| ENSG000001 | FAM131B | 3.87E-01 | ? | | 1.16E+00 | 2.19E-01 | 1.16E+00 |
| ENSG000001 | MYO1B | 3.87E-01 | | 7.20E-01 | 1.03E+00 | 3.67E-02 | 1.03E+00 |
| ENSG000001 | RALGAPA1 | 3.87E-01 | | 7.20E-01 | 1.03E+00 | 4.48E-02 | 1.03E+00 |
| ENSG000001 | UBXN11 | 3.87E-01 | | 7.20E-01 | 1.07E+00 | 9.38E-02 | 1.07E+00 |
| ENSG000001 | RBM19 | 3.87E-01 | | 7.20E-01 | 1.03E+00 | 4.65E-02 | 1.03E+00 |
| ENSG000002 | --- | 3.87E-01 | ? | | 7.58E-01 | -4.01E-01 | -1.32E+00 |
| ENSG000001 | FITM1 | 3.88E-01 | ? | | 1.35E+00 | 4.32E-01 | 1.35E+00 |
| ENSG000001 | LMBR1 | 3.88E-01 | | 7.20E-01 | 1.02E+00 | 3.55E-02 | 1.02E+00 |
| ENSG000002 | --- | 3.88E-01 | ? | | 1.35E+00 | 4.36E-01 | 1.35E+00 |
| ENSG000001 | CD28 | 3.88E-01 | ? | | 6.63E-01 | -5.93E-01 | -1.51E+00 |
| ENSG000002 | ZNF286B | 3.88E-01 | | 7.20E-01 | 1.07E+00 | 1.02E-01 | 1.07E+00 |
| ENSG000001 | SKIC3 | 3.88E-01 | | 7.20E-01 | 9.77E-01 | -3.30E-02 | -1.02E+00 |
| ENSG000001 | NUDT4B | 3.88E-01 | | 7.20E-01 | 9.54E-01 | -6.84E-02 | -1.05E+00 |
| ENSG000001 | LUC7L2 | 3.88E-01 | | 7.21E-01 | 9.70E-01 | -4.32E-02 | -1.03E+00 |
| ENSG000002 | --- | 3.88E-01 | ? | | 7.12E-01 | -4.91E-01 | -1.41E+00 |
| ENSG000002 | FAM187A | 3.88E-01 | ? | | 1.33E+00 | 4.16E-01 | 1.33E+00 |
| ENSG000002 | --- | 3.88E-01 | ? | | 7.30E-01 | -4.55E-01 | -1.37E+00 |
| ENSG000001 | ZNF710 | 3.88E-01 | | 7.21E-01 | 9.65E-01 | -5.13E-02 | -1.04E+00 |

| | | | | | | |
|---|---|---|---|---|---|---|
| ENSG000001 | KRT6B | 3.88E-01 | 7.21E-01 | 1.08E+00 | 1.06E-01 | 1.08E+00 |
| ENSG000001 | ZBTB44-DT | 3.88E-01 | ? | 6.94E-01 | -5.26E-01 | -1.44E+00 |
| ENSG000002 | TUG1 | 3.88E-01 | 7.21E-01 | 9.72E-01 | -4.07E-02 | -1.03E+00 |
| ENSG000002 | --- | 3.88E-01 | ? | 5.81E-01 | -7.84E-01 | -1.72E+00 |
| ENSG000001 | NR5A2 | 3.88E-01 | ? | 6.78E-01 | -5.60E-01 | -1.47E+00 |
| ENSG000001 | KIAA2013 | 3.88E-01 | 7.21E-01 | 1.05E+00 | 6.72E-02 | 1.05E+00 |
| ENSG000002 | HCG22 | 3.89E-01 | ? | 1.30E+00 | 3.83E-01 | 1.30E+00 |
| ENSG000002 | KLF7-IT1 | 3.89E-01 | ? | 1.32E+00 | 3.99E-01 | 1.32E+00 |
| ENSG000001 | FAM43B | 3.89E-01 | ? | 1.54E+00 | 6.27E-01 | 1.54E+00 |
| ENSG000002 | MSNP1 | 3.89E-01 | ? | 1.22E+00 | 2.88E-01 | 1.22E+00 |
| ENSG000002 | --- | 3.89E-01 | ? | 8.54E-01 | -2.27E-01 | -1.17E+00 |
| ENSG000002 | ZNF470-DT | 3.89E-01 | ? | 1.18E+00 | 2.33E-01 | 1.18E+00 |
| ENSG00000C | NDC80 | 3.89E-01 | 7.22E-01 | 8.81E-01 | -1.83E-01 | -1.14E+00 |
| ENSG000002 | --- | 3.89E-01 | ? | 1.30E+00 | 3.81E-01 | 1.30E+00 |
| ENSG000001 | CHIC2 | 3.90E-01 | 7.23E-01 | 9.54E-01 | -6.75E-02 | -1.05E+00 |
| ENSG000002 | ANXA2P2 | 3.90E-01 | 7.23E-01 | 9.68E-01 | -4.71E-02 | -1.03E+00 |
| ENSG00000C | JMJD6 | 3.90E-01 | 7.23E-01 | 9.62E-01 | -5.64E-02 | -1.04E+00 |
| ENSG000002 | CENATAC-DT | 3.90E-01 | ? | 1.22E+00 | 2.92E-01 | 1.22E+00 |
| ENSG000001 | SERPINA10 | 3.90E-01 | ? | 8.32E-01 | -2.65E-01 | -1.20E+00 |
| ENSG000001 | SCIMP | 3.90E-01 | ? | 1.56E+00 | 6.45E-01 | 1.56E+00 |
| ENSG000002 | --- | 3.90E-01 | ? | 1.55E+00 | 6.29E-01 | 1.55E+00 |
| ENSG000001 | CDC20 | 3.90E-01 | 7.23E-01 | 8.79E-01 | -1.86E-01 | -1.14E+00 |
| ENSG000001 | ZNF540 | 3.90E-01 | ? | 1.19E+00 | 2.54E-01 | 1.19E+00 |
| ENSG000002 | --- | 3.90E-01 | ? | 7.71E-01 | -3.76E-01 | -1.30E+00 |
| ENSG000002 | CHML | 3.90E-01 | 7.23E-01 | 1.05E+00 | 7.71E-02 | 1.05E+00 |
| ENSG000001 | IGF2R | 3.90E-01 | 7.23E-01 | 1.03E+00 | 4.29E-02 | 1.03E+00 |
| ENSG000001 | MLEC | 3.90E-01 | 7.23E-01 | 1.04E+00 | 5.03E-02 | 1.04E+00 |
| ENSG000002 | DGUOK-AS1 | 3.90E-01 | 7.23E-01 | 1.11E+00 | 1.56E-01 | 1.11E+00 |
| ENSG000002 | RPS15AP1 | 3.90E-01 | ? | 6.70E-01 | -5.78E-01 | -1.49E+00 |
| ENSG000002 | POLRMTP1 | 3.90E-01 | 7.24E-01 | 8.73E-01 | -1.96E-01 | -1.15E+00 |
| ENSG000002 | RPL23AP81 | 3.91E-01 | ? | 7.06E-01 | -5.03E-01 | -1.42E+00 |
| ENSG000001 | HDGFL2 | 3.91E-01 | 7.24E-01 | 9.67E-01 | -4.89E-02 | -1.03E+00 |
| ENSG000001 | ABCC4 | 3.91E-01 | 7.24E-01 | 1.04E+00 | 5.23E-02 | 1.04E+00 |
| ENSG000001 | TPD52L2 | 3.91E-01 | 7.24E-01 | 1.02E+00 | 3.53E-02 | 1.02E+00 |
| ENSG000001 | PPP2R3B | 3.91E-01 | 7.24E-01 | 1.07E+00 | 9.31E-02 | 1.07E+00 |
| ENSG000001 | UBE3D | 3.91E-01 | 7.24E-01 | 9.31E-01 | -1.03E-01 | -1.07E+00 |
| ENSG000002 | TPM3P5 | 3.91E-01 | ? | 8.16E-01 | -2.93E-01 | -1.22E+00 |
| ENSG000001 | COPRS | 3.91E-01 | 7.24E-01 | 1.04E+00 | 5.79E-02 | 1.04E+00 |
| ENSG000002 | DNAH10OS | 3.91E-01 | 7.24E-01 | 8.65E-01 | -2.10E-01 | -1.16E+00 |
| ENSG000002 | --- | 3.91E-01 | ? | 7.60E-01 | -3.96E-01 | -1.32E+00 |
| ENSG000001 | TNKS | 3.91E-01 | 7.24E-01 | 9.70E-01 | -4.34E-02 | -1.03E+00 |
| ENSG000001 | LRRC20 | 3.91E-01 | 7.24E-01 | 9.43E-01 | -8.41E-02 | -1.06E+00 |
| ENSG000001 | PYCR3 | 3.91E-01 | 7.24E-01 | 1.09E+00 | 1.18E-01 | 1.09E+00 |
| ENSG000001 | NMT2 | 3.91E-01 | 7.24E-01 | 1.04E+00 | 5.30E-02 | 1.04E+00 |
| ENSG000001 | RPA2 | 3.91E-01 | 7.24E-01 | 1.03E+00 | 4.88E-02 | 1.03E+00 |

| | | | | | | |
|---|---|---|---|---|---|---|
| ENSG000001 | LMNTD2 | 3.91E-01 | ? | 1.18E+00 | 2.42E-01 | 1.18E+00 |
| ENSG000001 | MIR4435-2H( | 3.91E-01 | 7.24E-01 | 9.49E-01 | -7.50E-02 | -1.05E+00 |
| ENSG000002 | CTBP1-AS | 3.91E-01 | 7.24E-01 | 8.83E-01 | -1.80E-01 | -1.13E+00 |
| ENSG000001 | IMMP2L | 3.91E-01 | 7.24E-01 | 1.05E+00 | 6.97E-02 | 1.05E+00 |
| ENSG000002 | --- | 3.91E-01 | 7.24E-01 | 8.98E-01 | -1.55E-01 | -1.11E+00 |
| ENSG000001 | PAN3 | 3.91E-01 | 7.24E-01 | 9.71E-01 | -4.28E-02 | -1.03E+00 |
| ENSG000001 | COP1 | 3.91E-01 | 7.24E-01 | 1.02E+00 | 3.39E-02 | 1.02E+00 |
| ENSG000001 | FAM83G | 3.92E-01 | 7.24E-01 | 1.04E+00 | 5.01E-02 | 1.04E+00 |
| ENSG00000C | PIK3C2A | 3.92E-01 | 7.24E-01 | 9.65E-01 | -5.10E-02 | -1.04E+00 |
| ENSG000001 | BOK | 3.92E-01 | 7.24E-01 | 1.06E+00 | 8.45E-02 | 1.06E+00 |
| ENSG000001 | ECM1 | 3.92E-01 | 7.24E-01 | 9.35E-01 | -9.62E-02 | -1.07E+00 |
| ENSG000001 | NES | 3.92E-01 | ? | 8.42E-01 | -2.47E-01 | -1.19E+00 |
| ENSG000002 | --- | 3.92E-01 | ? | 7.48E-01 | -4.20E-01 | -1.34E+00 |
| ENSG000001 | CDCP1 | 3.92E-01 | 7.24E-01 | 9.67E-01 | -4.78E-02 | -1.03E+00 |
| ENSG000002 | RPL37P23 | 3.92E-01 | ? | 5.70E-01 | -8.11E-01 | -1.75E+00 |
| ENSG000002 | --- | 3.92E-01 | ? | 1.22E+00 | 2.83E-01 | 1.22E+00 |
| ENSG000001 | CTHRC1 | 3.92E-01 | ? | 6.95E-01 | -5.25E-01 | -1.44E+00 |
| ENSG000002 | B4GALT1-AS | 3.92E-01 | 7.24E-01 | 8.58E-01 | -2.22E-01 | -1.17E+00 |
| ENSG00000C | MYOM2 | 3.92E-01 | ? | 1.23E+00 | 3.02E-01 | 1.23E+00 |
| ENSG000001 | PTDSS2 | 3.92E-01 | 7.24E-01 | 1.05E+00 | 6.44E-02 | 1.05E+00 |
| ENSG000001 | ZSCAN4 | 3.92E-01 | 7.24E-01 | 1.16E+00 | 2.09E-01 | 1.16E+00 |
| ENSG000001 | RPGR | 3.92E-01 | 7.24E-01 | 9.38E-01 | -9.25E-02 | -1.07E+00 |
| ENSG000001 | LAMB2 | 3.92E-01 | 7.24E-01 | 9.62E-01 | -5.65E-02 | -1.04E+00 |
| ENSG000002 | --- | 3.92E-01 | ? | 6.73E-01 | -5.71E-01 | -1.49E+00 |
| ENSG000001 | XPO6 | 3.92E-01 | 7.24E-01 | 1.02E+00 | 2.98E-02 | 1.02E+00 |
| ENSG000002 | MANEA-DT | 3.92E-01 | ? | 7.25E-01 | -4.63E-01 | -1.38E+00 |
| ENSG000001 | PWWP2B | 3.92E-01 | 7.24E-01 | 1.10E+00 | 1.37E-01 | 1.10E+00 |
| ENSG00000C | NUDCD3 | 3.92E-01 | 7.24E-01 | 1.03E+00 | 4.18E-02 | 1.03E+00 |
| ENSG000001 | PIGK | 3.92E-01 | 7.24E-01 | 9.60E-01 | -5.86E-02 | -1.04E+00 |
| ENSG000001 | ADAM10 | 3.92E-01 | 7.24E-01 | 1.03E+00 | 3.93E-02 | 1.03E+00 |
| ENSG000001 | DNAJA3 | 3.92E-01 | 7.24E-01 | 1.02E+00 | 3.47E-02 | 1.02E+00 |
| ENSG000001 | ZNF319 | 3.92E-01 | 7.24E-01 | 1.05E+00 | 6.90E-02 | 1.05E+00 |
| ENSG000002 | SEPTIN10P1 | 3.93E-01 | ? | 8.01E-01 | -3.20E-01 | -1.25E+00 |
| ENSG000002 | --- | 3.93E-01 | 7.24E-01 | 1.14E+00 | 1.95E-01 | 1.14E+00 |
| ENSG000001 | PDE3B | 3.93E-01 | 7.25E-01 | 1.09E+00 | 1.25E-01 | 1.09E+00 |
| ENSG000002 | ST3GAL5-AS | 3.93E-01 | ? | 1.33E+00 | 4.13E-01 | 1.33E+00 |
| ENSG000001 | ZKSCAN1 | 3.93E-01 | 7.25E-01 | 9.77E-01 | -3.39E-02 | -1.02E+00 |
| ENSG000001 | SURF4 | 3.93E-01 | 7.25E-01 | 1.03E+00 | 4.19E-02 | 1.03E+00 |
| ENSG000002 | JDP2-AS1 | 3.93E-01 | ? | 1.39E+00 | 4.75E-01 | 1.39E+00 |
| ENSG000002 | --- | 3.93E-01 | ? | 6.65E-01 | -5.88E-01 | -1.50E+00 |
| ENSG000001 | MAPK8IP3 | 3.93E-01 | 7.25E-01 | 9.32E-01 | -1.01E-01 | -1.07E+00 |
| ENSG000001 | TGIF1 | 3.93E-01 | 7.25E-01 | 1.03E+00 | 4.63E-02 | 1.03E+00 |
| ENSG000001 | TMPRSS6 | 3.93E-01 | ? | 6.90E-01 | -5.35E-01 | -1.45E+00 |
| ENSG000001 | PFDN5 | 3.93E-01 | 7.25E-01 | 1.03E+00 | 3.58E-02 | 1.03E+00 |
| ENSG00000C | KLHL13 | 3.93E-01 | 7.25E-01 | 1.06E+00 | 7.93E-02 | 1.06E+00 |

| | | | | | | |
|---|---|---|---|---|---|---|
| ENSG000002 | AATBC | 3.93E-01 | 7.25E-01 | 1.08E+00 | 1.10E-01 | 1.08E+00 |
| ENSG000002 | --- | 3.93E-01 | 7.25E-01 | 1.18E+00 | 2.35E-01 | 1.18E+00 |
| ENSG000001 | CD79A | 3.93E-01 | ? | 1.46E+00 | 5.51E-01 | 1.46E+00 |
| ENSG000001 | SH3BGRL | 3.93E-01 | 7.25E-01 | 1.03E+00 | 4.90E-02 | 1.03E+00 |
| ENSG000001 | RPL39L | 3.94E-01 | 7.25E-01 | 1.09E+00 | 1.22E-01 | 1.09E+00 |
| ENSG000001 | SENP2 | 3.94E-01 | 7.25E-01 | 9.74E-01 | -3.83E-02 | -1.03E+00 |
| ENSG000001 | DCHS1 | 3.94E-01 | ? | 1.18E+00 | 2.38E-01 | 1.18E+00 |
| ENSG000002 | --- | 3.94E-01 | ? | 6.84E-01 | -5.48E-01 | -1.46E+00 |
| ENSG000002 | UGT1A1 | 3.94E-01 | 7.25E-01 | 9.29E-01 | -1.06E-01 | -1.08E+00 |
| ENSG000001 | VPS29 | 3.94E-01 | 7.25E-01 | 9.73E-01 | -4.01E-02 | -1.03E+00 |
| ENSG000001 | SYT11 | 3.94E-01 | 7.25E-01 | 1.13E+00 | 1.79E-01 | 1.13E+00 |
| ENSG000001 | CD14 | 3.94E-01 | ? | 1.41E+00 | 4.94E-01 | 1.41E+00 |
| ENSG000001 | CBLN3 | 3.94E-01 | 7.25E-01 | 9.35E-01 | -9.67E-02 | -1.07E+00 |
| ENSG000001 | CYP2E1 | 3.94E-01 | ? | 8.35E-01 | -2.61E-01 | -1.20E+00 |
| ENSG000001 | SEC22A | 3.94E-01 | 7.25E-01 | 1.04E+00 | 5.11E-02 | 1.04E+00 |
| ENSG00000C | LRRFIP2 | 3.94E-01 | 7.25E-01 | 1.04E+00 | 6.03E-02 | 1.04E+00 |
| ENSG000002 | --- | 3.94E-01 | 7.25E-01 | 9.07E-01 | -1.41E-01 | -1.10E+00 |
| ENSG000001 | THBD | 3.94E-01 | 7.25E-01 | 9.55E-01 | -6.70E-02 | -1.05E+00 |
| ENSG000001 | DNAJC21 | 3.94E-01 | 7.25E-01 | 9.71E-01 | -4.21E-02 | -1.03E+00 |
| ENSG00000C | SMC1A | 3.94E-01 | 7.25E-01 | 9.72E-01 | -4.17E-02 | -1.03E+00 |
| ENSG000002 | --- | 3.94E-01 | 7.25E-01 | 1.17E+00 | 2.25E-01 | 1.17E+00 |
| ENSG000001 | PPP2R5E | 3.94E-01 | 7.25E-01 | 9.78E-01 | -3.17E-02 | -1.02E+00 |
| ENSG000001 | AXIN2 | 3.94E-01 | ? | 1.26E+00 | 3.32E-01 | 1.26E+00 |
| ENSG000002 | --- | 3.94E-01 | ? | 1.37E+00 | 4.55E-01 | 1.37E+00 |
| ENSG000001 | GKAP1 | 3.94E-01 | 7.25E-01 | 8.90E-01 | -1.68E-01 | -1.12E+00 |
| ENSG000002 | CROCCP3 | 3.94E-01 | 7.25E-01 | 8.94E-01 | -1.61E-01 | -1.12E+00 |
| ENSG000001 | IFIT2 | 3.94E-01 | 7.25E-01 | 9.15E-01 | -1.27E-01 | -1.09E+00 |
| ENSG000002 | --- | 3.94E-01 | ? | 1.52E+00 | 6.05E-01 | 1.52E+00 |
| ENSG000001 | PAQR7 | 3.95E-01 | 7.25E-01 | 1.05E+00 | 6.72E-02 | 1.05E+00 |
| ENSG000002 | --- | 3.95E-01 | ? | 1.19E+00 | 2.50E-01 | 1.19E+00 |
| ENSG000002 | --- | 3.95E-01 | ? | 8.03E-01 | -3.17E-01 | -1.25E+00 |
| ENSG000002 | --- | 3.95E-01 | 7.25E-01 | 8.66E-01 | -2.07E-01 | -1.15E+00 |
| ENSG000002 | --- | 3.95E-01 | ? | 1.27E+00 | 3.47E-01 | 1.27E+00 |
| ENSG000001 | SPOPL | 3.95E-01 | 7.25E-01 | 9.63E-01 | -5.38E-02 | -1.04E+00 |
| ENSG000002 | SELENOTP1 | 3.95E-01 | ? | 1.33E+00 | 4.16E-01 | 1.33E+00 |
| ENSG000001 | SLC52A1 | 3.95E-01 | ? | 7.16E-01 | -4.83E-01 | -1.40E+00 |
| ENSG000002 | --- | 3.95E-01 | ? | 1.18E+00 | 2.33E-01 | 1.18E+00 |
| ENSG000002 | --- | 3.95E-01 | ? | 8.48E-01 | -2.39E-01 | -1.18E+00 |
| ENSG000001 | EDF1 | 3.95E-01 | 7.26E-01 | 1.03E+00 | 3.60E-02 | 1.03E+00 |
| ENSG000002 | NDUFC2-KCT | 3.95E-01 | 7.26E-01 | 1.17E+00 | 2.32E-01 | 1.17E+00 |
| ENSG000002 | --- | 3.95E-01 | ? | 7.75E-01 | -3.68E-01 | -1.29E+00 |
| ENSG000001 | RAB6A | 3.95E-01 | 7.26E-01 | 9.60E-01 | -5.88E-02 | -1.04E+00 |
| ENSG000001 | ELF1 | 3.95E-01 | 7.26E-01 | 1.03E+00 | 4.12E-02 | 1.03E+00 |
| ENSG000001 | DUSP2 | 3.95E-01 | ? | 1.26E+00 | 3.33E-01 | 1.26E+00 |
| ENSG000002 | MAP1LC3B2 | 3.95E-01 | 7.26E-01 | 9.07E-01 | -1.41E-01 | -1.10E+00 |

| | | | | | | |
|---|---|---|---|---|---|---|
| ENSG000001 | NAP1L3 | 3.95E-01 | ? | 1.17E+00 | 2.30E-01 | 1.17E+00 |
| ENSG000002 | SRRM5 | 3.95E-01 | 7.26E-01 | 8.64E-01 | -2.12E-01 | -1.16E+00 |
| ENSG000001 | PEX7 | 3.95E-01 | 7.26E-01 | 1.06E+00 | 8.77E-02 | 1.06E+00 |
| ENSG000001 | EID2B | 3.95E-01 | 7.26E-01 | 1.15E+00 | 2.04E-01 | 1.15E+00 |
| ENSG000001 | FARSA | 3.95E-01 | 7.26E-01 | 1.04E+00 | 5.00E-02 | 1.04E+00 |
| ENSG000002 | --- | 3.96E-01 | ? | 7.98E-01 | -3.26E-01 | -1.25E+00 |
| ENSG000002 | --- | 3.96E-01 | 7.26E-01 | 9.27E-01 | -1.09E-01 | -1.08E+00 |
| ENSG000002 | EXTL3-AS1 | 3.96E-01 | 7.26E-01 | 1.15E+00 | 1.96E-01 | 1.15E+00 |
| ENSG000001 | SYT12 | 3.96E-01 | 7.27E-01 | 9.18E-01 | -1.24E-01 | -1.09E+00 |
| ENSG000001 | FAM117A | 3.96E-01 | 7.27E-01 | 1.07E+00 | 1.00E-01 | 1.07E+00 |
| ENSG000001 | ARHGEF39 | 3.96E-01 | 7.27E-01 | 8.99E-01 | -1.53E-01 | -1.11E+00 |
| ENSG000001 | MINDY3 | 3.96E-01 | 7.27E-01 | 9.70E-01 | -4.37E-02 | -1.03E+00 |
| ENSG000001 | BLM | 3.96E-01 | 7.27E-01 | 9.16E-01 | -1.27E-01 | -1.09E+00 |
| ENSG000002 | --- | 3.96E-01 | ? | 1.47E+00 | 5.53E-01 | 1.47E+00 |
| ENSG000002 | CFL1P5 | 3.96E-01 | ? | 7.07E-01 | -5.00E-01 | -1.41E+00 |
| ENSG000001 | ALKAL1 | 3.96E-01 | ? | 7.86E-01 | -3.47E-01 | -1.27E+00 |
| ENSG000002 | --- | 3.97E-01 | ? | 6.72E-01 | -5.75E-01 | -1.49E+00 |
| ENSG000001 | SPIN1 | 3.97E-01 | 7.28E-01 | 9.80E-01 | -2.89E-02 | -1.02E+00 |
| ENSG000001 | PACSIN2 | 3.97E-01 | 7.28E-01 | 1.02E+00 | 3.52E-02 | 1.02E+00 |
| ENSG000002 | --- | 3.97E-01 | ? | 8.60E-01 | -2.17E-01 | -1.16E+00 |
| ENSG000001 | ITGA9 | 3.97E-01 | ? | 1.41E+00 | 5.00E-01 | 1.41E+00 |
| ENSG00000C | PLEKHG2 | 3.97E-01 | 7.28E-01 | 1.04E+00 | 5.02E-02 | 1.04E+00 |
| ENSG000002 | EPM2A-DT | 3.97E-01 | 7.28E-01 | 1.14E+00 | 1.95E-01 | 1.14E+00 |
| ENSG000001 | OTUB1 | 3.97E-01 | 7.28E-01 | 1.03E+00 | 4.95E-02 | 1.03E+00 |
| ENSG00000C | EVI5 | 3.97E-01 | 7.28E-01 | 9.68E-01 | -4.69E-02 | -1.03E+00 |
| ENSG000002 | --- | 3.97E-01 | 7.28E-01 | 1.09E+00 | 1.23E-01 | 1.09E+00 |
| ENSG000001 | MGAT2 | 3.97E-01 | 7.28E-01 | 1.07E+00 | 9.44E-02 | 1.07E+00 |
| ENSG000001 | TMEM14B | 3.97E-01 | 7.28E-01 | 1.04E+00 | 5.24E-02 | 1.04E+00 |
| ENSG000001 | RSPH1 | 3.97E-01 | 7.28E-01 | 1.13E+00 | 1.72E-01 | 1.13E+00 |
| ENSG000001 | AGL | 3.97E-01 | 7.28E-01 | 9.61E-01 | -5.74E-02 | -1.04E+00 |
| ENSG00000C | RDH11 | 3.97E-01 | 7.28E-01 | 1.02E+00 | 3.18E-02 | 1.02E+00 |
| ENSG000002 | --- | 3.97E-01 | 7.28E-01 | 8.83E-01 | -1.79E-01 | -1.13E+00 |
| ENSG000001 | ZNF711 | 3.98E-01 | 7.28E-01 | 1.10E+00 | 1.39E-01 | 1.10E+00 |
| ENSG000001 | DIS3L | 3.98E-01 | 7.28E-01 | 9.58E-01 | -6.14E-02 | -1.04E+00 |
| ENSG000001 | CYTH2 | 3.98E-01 | 7.28E-01 | 9.48E-01 | -7.69E-02 | -1.05E+00 |
| ENSG000001 | RNF212 | 3.98E-01 | ? | 1.34E+00 | 4.26E-01 | 1.34E+00 |
| ENSG000002 | YBX1P2 | 3.98E-01 | 7.28E-01 | 8.93E-01 | -1.63E-01 | -1.12E+00 |
| ENSG00000C | TMSB10 | 3.98E-01 | 7.28E-01 | 1.03E+00 | 4.51E-02 | 1.03E+00 |
| ENSG000002 | --- | 3.98E-01 | ? | 6.81E-01 | -5.54E-01 | -1.47E+00 |
| ENSG000001 | DRAP1 | 3.98E-01 | 7.28E-01 | 9.65E-01 | -5.18E-02 | -1.04E+00 |
| ENSG000001 | EIF2AK4 | 3.98E-01 | 7.28E-01 | 1.03E+00 | 4.18E-02 | 1.03E+00 |
| ENSG000002 | SYCE1L | 3.98E-01 | ? | 7.42E-01 | -4.31E-01 | -1.35E+00 |
| ENSG000001 | ATP6V0A2 | 3.98E-01 | 7.28E-01 | 9.66E-01 | -5.05E-02 | -1.04E+00 |
| ENSG000001 | FLNC | 3.98E-01 | 7.28E-01 | 8.78E-01 | -1.88E-01 | -1.14E+00 |
| ENSG000001 | ZDHHC24 | 3.98E-01 | 7.28E-01 | 1.04E+00 | 5.42E-02 | 1.04E+00 |

| | | | | | | |
|---|---|---|---|---|---|---|
| ENSG000002 | COMMD3-BN | 3.98E-01 | 7.28E-01 | 1.46E+00 | 5.41E-01 | 1.46E+00 |
| ENSG000001 | TMEM115 | 3.98E-01 | 7.28E-01 | 1.04E+00 | 5.80E-02 | 1.04E+00 |
| ENSG000001 | UBN2 | 3.98E-01 | 7.28E-01 | 9.46E-01 | -8.00E-02 | -1.06E+00 |
| ENSG000001 | KPNA1 | 3.98E-01 | 7.28E-01 | 1.02E+00 | 3.26E-02 | 1.02E+00 |
| ENSG000001 | EDNRA | 3.98E-01 | 7.28E-01 | 1.08E+00 | 1.08E-01 | 1.08E+00 |
| ENSG000002 | --- | 3.98E-01 | ? | 1.22E+00 | 2.83E-01 | 1.22E+00 |
| ENSG00000C | LXN | 3.98E-01 | 7.28E-01 | 9.26E-01 | -1.10E-01 | -1.08E+00 |
| ENSG000002 | LINC00933 | 3.98E-01 | ? | 1.21E+00 | 2.81E-01 | 1.21E+00 |
| ENSG000002 | RPL10P9 | 3.99E-01 | ? | 1.27E+00 | 3.40E-01 | 1.27E+00 |
| ENSG000001 | SMIM8 | 3.99E-01 | 7.28E-01 | 1.07E+00 | 9.91E-02 | 1.07E+00 |
| ENSG000002 | MSH5-SAPCD | 3.99E-01 | 7.28E-01 | 8.70E-01 | -2.01E-01 | -1.15E+00 |
| ENSG000002 | --- | 3.99E-01 | ? | 1.22E+00 | 2.90E-01 | 1.22E+00 |
| ENSG000002 | --- | 3.99E-01 | ? | 8.04E-01 | -3.15E-01 | -1.24E+00 |
| ENSG000001 | TMEM9 | 3.99E-01 | 7.28E-01 | 1.06E+00 | 8.20E-02 | 1.06E+00 |
| ENSG000002 | UPK3BL1 | 3.99E-01 | 7.28E-01 | 1.11E+00 | 1.49E-01 | 1.11E+00 |
| ENSG000001 | CNOT7 | 3.99E-01 | 7.28E-01 | 9.75E-01 | -3.68E-02 | -1.03E+00 |
| ENSG000002 | --- | 3.99E-01 | ? | 1.45E+00 | 5.36E-01 | 1.45E+00 |
| ENSG000002 | LINC02826 | 3.99E-01 | ? | 8.20E-01 | -2.85E-01 | -1.22E+00 |
| ENSG000002 | LINC02960 | 3.99E-01 | ? | 1.15E+00 | 2.06E-01 | 1.15E+00 |
| ENSG000002 | NFKBIL1 | 3.99E-01 | 7.28E-01 | 1.07E+00 | 1.02E-01 | 1.07E+00 |
| ENSG000002 | LINC02036 | 3.99E-01 | ? | 1.26E+00 | 3.33E-01 | 1.26E+00 |
| ENSG000001 | RNMT | 3.99E-01 | 7.28E-01 | 1.03E+00 | 4.39E-02 | 1.03E+00 |
| ENSG000002 | --- | 3.99E-01 | ? | 1.24E+00 | 3.14E-01 | 1.24E+00 |
| ENSG000001 | EFCAB14 | 3.99E-01 | 7.28E-01 | 9.78E-01 | -3.14E-02 | -1.02E+00 |
| ENSG000001 | NAT9 | 3.99E-01 | 7.28E-01 | 1.05E+00 | 6.90E-02 | 1.05E+00 |
| ENSG000001 | CACNB3 | 3.99E-01 | 7.28E-01 | 1.04E+00 | 5.61E-02 | 1.04E+00 |
| ENSG000001 | C3orf52 | 3.99E-01 | 7.28E-01 | 9.45E-01 | -8.09E-02 | -1.06E+00 |
| ENSG000002 | PRAG1 | 3.99E-01 | 7.28E-01 | 1.06E+00 | 7.73E-02 | 1.06E+00 |
| ENSG000001 | PDGFC | 3.99E-01 | 7.28E-01 | 1.04E+00 | 5.53E-02 | 1.04E+00 |
| ENSG000002 | --- | 3.99E-01 | ? | 7.93E-01 | -3.35E-01 | -1.26E+00 |
| ENSG000002 | PKD1P4 | 3.99E-01 | 7.28E-01 | 8.38E-01 | -2.56E-01 | -1.19E+00 |
| ENSG00000C | CDKL5 | 3.99E-01 | 7.28E-01 | 1.05E+00 | 6.54E-02 | 1.05E+00 |
| ENSG000002 | --- | 3.99E-01 | 7.28E-01 | 8.55E-01 | -2.26E-01 | -1.17E+00 |
| ENSG00000C | ATP2C2 | 3.99E-01 | 7.28E-01 | 8.70E-01 | -2.01E-01 | -1.15E+00 |
| ENSG000002 | RPL3P4 | 4.00E-01 | 7.28E-01 | 1.08E+00 | 1.09E-01 | 1.08E+00 |
| ENSG000001 | SLC29A1 | 4.00E-01 | 7.28E-01 | 1.05E+00 | 6.98E-02 | 1.05E+00 |
| ENSG000002 | --- | 4.00E-01 | 7.28E-01 | 1.22E+00 | 2.83E-01 | 1.22E+00 |
| ENSG000001 | PLEKHA7 | 4.00E-01 | 7.28E-01 | 1.10E+00 | 1.44E-01 | 1.10E+00 |
| ENSG000002 | --- | 4.00E-01 | 7.28E-01 | 9.29E-01 | -1.06E-01 | -1.08E+00 |
| ENSG00000C | CDV3 | 4.00E-01 | 7.28E-01 | 1.02E+00 | 2.84E-02 | 1.02E+00 |
| ENSG000001 | ACTBL2 | 4.00E-01 | 7.28E-01 | 1.09E+00 | 1.24E-01 | 1.09E+00 |
| ENSG000002 | --- | 4.00E-01 | 7.28E-01 | 9.19E-01 | -1.22E-01 | -1.09E+00 |
| ENSG00000C | YTHDC2 | 4.00E-01 | 7.28E-01 | 9.74E-01 | -3.85E-02 | -1.03E+00 |
| ENSG000001 | ALG13 | 4.00E-01 | 7.28E-01 | 9.57E-01 | -6.30E-02 | -1.04E+00 |
| ENSG000002 | MIR34AHG | 4.00E-01 | 7.28E-01 | 1.10E+00 | 1.34E-01 | 1.10E+00 |

| | | | | | | |
|---|---|---|---|---|---|---|
| ENSG000002 | GABBR1 | 4.00E-01 | 7.28E-01 | 9.25E-01 | -1.12E-01 | -1.08E+00 |
| ENSG000001 | AMPD2 | 4.00E-01 | 7.29E-01 | 1.04E+00 | 5.41E-02 | 1.04E+00 |
| ENSG000002 | MEMO1P1 | 4.00E-01 | ? | 1.24E+00 | 3.09E-01 | 1.24E+00 |
| ENSG000001 | ABCD1 | 4.00E-01 | 7.29E-01 | 1.06E+00 | 7.74E-02 | 1.06E+00 |
| ENSG000002 | --- | 4.01E-01 | ? | 6.00E-01 | -7.37E-01 | -1.67E+00 |
| ENSG000001 | SCRN2 | 4.01E-01 | 7.29E-01 | 1.08E+00 | 1.06E-01 | 1.08E+00 |
| ENSG000001 | CARD19 | 4.01E-01 | 7.29E-01 | 1.07E+00 | 1.03E-01 | 1.07E+00 |
| ENSG000002 | --- | 4.01E-01 | ? | 1.14E+00 | 1.91E-01 | 1.14E+00 |
| ENSG000002 | SMIM13 | 4.01E-01 | 7.29E-01 | 9.67E-01 | -4.79E-02 | -1.03E+00 |
| ENSG000002 | RPL22P24 | 4.01E-01 | ? | 1.40E+00 | 4.86E-01 | 1.40E+00 |
| ENSG000001 | DLX5 | 4.01E-01 | ? | 1.23E+00 | 3.04E-01 | 1.23E+00 |
| ENSG000001 | RAD51B | 4.01E-01 | 7.29E-01 | 9.36E-01 | -9.55E-02 | -1.07E+00 |
| ENSG000002 | LINC00526 | 4.01E-01 | ? | 1.18E+00 | 2.39E-01 | 1.18E+00 |
| ENSG000002 | --- | 4.01E-01 | ? | 6.32E-01 | -6.62E-01 | -1.58E+00 |
| ENSG000002 | OR2A7 | 4.01E-01 | 7.29E-01 | 9.33E-01 | -1.01E-01 | -1.07E+00 |
| ENSG000001 | TTLL4 | 4.01E-01 | 7.29E-01 | 1.03E+00 | 4.35E-02 | 1.03E+00 |
| ENSG000001 | ANO9 | 4.01E-01 | 7.29E-01 | 1.07E+00 | 9.75E-02 | 1.07E+00 |
| ENSG000001 | LAGE3 | 4.01E-01 | 7.29E-01 | 1.11E+00 | 1.56E-01 | 1.11E+00 |
| ENSG000002 | --- | 4.01E-01 | ? | 8.31E-01 | -2.67E-01 | -1.20E+00 |
| ENSG00000C | PPP2R5A | 4.01E-01 | 7.30E-01 | 9.71E-01 | -4.26E-02 | -1.03E+00 |
| ENSG000001 | ACRBP | 4.01E-01 | ? | 7.52E-01 | -4.11E-01 | -1.33E+00 |
| ENSG000001 | GIN1 | 4.01E-01 | 7.30E-01 | 9.52E-01 | -7.08E-02 | -1.05E+00 |
| ENSG000001 | SLC2A11 | 4.01E-01 | 7.30E-01 | 9.32E-01 | -1.01E-01 | -1.07E+00 |
| ENSG000001 | GLDC | 4.01E-01 | ? | 1.15E+00 | 1.96E-01 | 1.15E+00 |
| ENSG000001 | SLC37A3 | 4.01E-01 | 7.30E-01 | 9.69E-01 | -4.60E-02 | -1.03E+00 |
| ENSG000001 | HEMK1 | 4.02E-01 | 7.30E-01 | 1.06E+00 | 7.77E-02 | 1.06E+00 |
| ENSG000001 | C20orf144 | 4.02E-01 | ? | 1.45E+00 | 5.34E-01 | 1.45E+00 |
| ENSG00000C | MZF1 | 4.02E-01 | 7.30E-01 | 9.33E-01 | -1.01E-01 | -1.07E+00 |
| ENSG00000C | RSBN1 | 4.02E-01 | 7.30E-01 | 9.71E-01 | -4.25E-02 | -1.03E+00 |
| ENSG000002 | UQCC3 | 4.02E-01 | 7.30E-01 | 1.08E+00 | 1.16E-01 | 1.08E+00 |
| ENSG000001 | DNAH11 | 4.02E-01 | 7.30E-01 | 9.51E-01 | -7.29E-02 | -1.05E+00 |
| ENSG000001 | CDIPT | 4.02E-01 | 7.30E-01 | 1.06E+00 | 8.31E-02 | 1.06E+00 |
| ENSG000002 | --- | 4.02E-01 | 7.30E-01 | 1.09E+00 | 1.24E-01 | 1.09E+00 |
| ENSG000001 | ZNF699 | 4.02E-01 | 7.30E-01 | 9.64E-01 | -5.26E-02 | -1.04E+00 |
| ENSG000001 | KBTBD11 | 4.02E-01 | 7.30E-01 | 1.07E+00 | 1.02E-01 | 1.07E+00 |
| ENSG000002 | --- | 4.02E-01 | ? | 1.22E+00 | 2.92E-01 | 1.22E+00 |
| ENSG000002 | LYN | 4.02E-01 | 7.30E-01 | 1.03E+00 | 3.92E-02 | 1.03E+00 |
| ENSG000002 | RPL41P2 | 4.02E-01 | 7.30E-01 | 1.09E+00 | 1.22E-01 | 1.09E+00 |
| ENSG000001 | PNPLA2 | 4.02E-01 | 7.30E-01 | 1.05E+00 | 7.22E-02 | 1.05E+00 |
| ENSG000001 | GCNT3 | 4.02E-01 | 7.30E-01 | 8.89E-01 | -1.69E-01 | -1.12E+00 |
| ENSG000001 | HSPA14 | 4.03E-01 | 7.30E-01 | 1.04E+00 | 6.04E-02 | 1.04E+00 |
| ENSG000001 | C11orf24 | 4.03E-01 | 7.30E-01 | 9.45E-01 | -8.14E-02 | -1.06E+00 |
| ENSG000001 | NMU | 4.03E-01 | ? | 8.11E-01 | -3.02E-01 | -1.23E+00 |
| ENSG000001 | SLC29A4 | 4.03E-01 | 7.30E-01 | 1.11E+00 | 1.50E-01 | 1.11E+00 |
| ENSG000001 | PRPSAP2 | 4.03E-01 | 7.30E-01 | 9.66E-01 | -5.03E-02 | -1.04E+00 |

| | | | | | | |
|---|---|---|---|---|---|---|
| ENSG000001 | FRMD4A | 4.03E-01 | 7.30E-01 | 9.67E-01 | -4.90E-02 | -1.03E+00 |
| ENSG000002 | --- | 4.03E-01 | ? | 7.23E-01 | -4.69E-01 | -1.38E+00 |
| ENSG000002 | GOLGA8H | 4.03E-01 | ? | 8.69E-01 | -2.02E-01 | -1.15E+00 |
| ENSG000001 | RIC1 | 4.03E-01 | 7.31E-01 | 1.02E+00 | 3.41E-02 | 1.02E+00 |
| ENSG000001 | ACOT8 | 4.03E-01 | 7.31E-01 | 1.04E+00 | 6.02E-02 | 1.04E+00 |
| ENSG000002 | KLHL7-DT | 4.03E-01 | ? | 1.20E+00 | 2.65E-01 | 1.20E+00 |
| ENSG000001 | JDP2 | 4.03E-01 | 7.31E-01 | 1.08E+00 | 1.06E-01 | 1.08E+00 |
| ENSG000001 | WIPI2 | 4.03E-01 | 7.31E-01 | 1.03E+00 | 3.66E-02 | 1.03E+00 |
| ENSG000001 | GDF7 | 4.03E-01 | ? | 6.80E-01 | -5.57E-01 | -1.47E+00 |
| ENSG000001 | EEF1AKMT1 | 4.03E-01 | 7.31E-01 | 1.07E+00 | 9.87E-02 | 1.07E+00 |
| ENSG000001 | HLA-DQA1 | 4.03E-01 | ? | 1.48E+00 | 5.69E-01 | 1.48E+00 |
| ENSG000002 | --- | 4.03E-01 | ? | 1.31E+00 | 3.93E-01 | 1.31E+00 |
| ENSG000002 | FAM157C | 4.03E-01 | 7.31E-01 | 9.34E-01 | -9.82E-02 | -1.07E+00 |
| ENSG000001 | NPRL2 | 4.03E-01 | 7.31E-01 | 1.06E+00 | 8.45E-02 | 1.06E+00 |
| ENSG000001 | PHF21A | 4.03E-01 | 7.31E-01 | 9.57E-01 | -6.30E-02 | -1.04E+00 |
| ENSG000001 | UBASH3B | 4.04E-01 | 7.31E-01 | 9.67E-01 | -4.86E-02 | -1.03E+00 |
| ENSG000002 | ZNF260 | 4.04E-01 | 7.31E-01 | 9.60E-01 | -5.86E-02 | -1.04E+00 |
| ENSG000001 | FGF5 | 4.04E-01 | 7.31E-01 | 9.01E-01 | -1.51E-01 | -1.11E+00 |
| ENSG000001 | PRPS1 | 4.04E-01 | 7.31E-01 | 1.03E+00 | 3.92E-02 | 1.03E+00 |
| ENSG000001 | CUTA | 4.04E-01 | 7.31E-01 | 1.04E+00 | 5.61E-02 | 1.04E+00 |
| ENSG00000C | ANO2 | 4.04E-01 | ? | 1.22E+00 | 2.82E-01 | 1.22E+00 |
| ENSG000001 | PPP1R14BP3 | 4.04E-01 | 7.31E-01 | 8.70E-01 | -2.01E-01 | -1.15E+00 |
| ENSG000002 | ZNF407 | 4.04E-01 | 7.31E-01 | 1.03E+00 | 4.53E-02 | 1.03E+00 |
| ENSG000001 | MCCC2 | 4.04E-01 | 7.31E-01 | 1.03E+00 | 3.85E-02 | 1.03E+00 |
| ENSG000001 | HMGB1P5 | 4.04E-01 | 7.31E-01 | 9.19E-01 | -1.22E-01 | -1.09E+00 |
| ENSG000001 | IL6 | 4.04E-01 | 7.31E-01 | 8.05E-01 | -3.14E-01 | -1.24E+00 |
| ENSG000002 | --- | 4.04E-01 | ? | 6.81E-01 | -5.54E-01 | -1.47E+00 |
| ENSG000001 | GUF1 | 4.04E-01 | 7.31E-01 | 9.67E-01 | -4.78E-02 | -1.03E+00 |
| ENSG000002 | --- | 4.04E-01 | 7.31E-01 | 8.95E-01 | -1.60E-01 | -1.12E+00 |
| ENSG000001 | HCFC1R1 | 4.05E-01 | 7.32E-01 | 1.06E+00 | 9.06E-02 | 1.06E+00 |
| ENSG000001 | CHST4 | 4.05E-01 | ? | 1.31E+00 | 3.88E-01 | 1.31E+00 |
| ENSG000002 | ARPC3P1 | 4.05E-01 | ? | 1.45E+00 | 5.39E-01 | 1.45E+00 |
| ENSG000001 | FOXD4 | 4.05E-01 | ? | 8.27E-01 | -2.74E-01 | -1.21E+00 |
| ENSG000002 | --- | 4.05E-01 | ? | 1.46E+00 | 5.43E-01 | 1.46E+00 |
| ENSG000001 | HRH4 | 4.05E-01 | ? | 1.33E+00 | 4.12E-01 | 1.33E+00 |
| ENSG000002 | SUMO2P17 | 4.05E-01 | ? | 7.42E-01 | -4.31E-01 | -1.35E+00 |
| ENSG000001 | JMJD7-PLA2( | 4.05E-01 | 7.32E-01 | 9.12E-01 | -1.34E-01 | -1.10E+00 |
| ENSG000001 | TMEM164 | 4.05E-01 | 7.32E-01 | 1.03E+00 | 4.75E-02 | 1.03E+00 |
| ENSG000001 | BATF | 4.05E-01 | ? | 7.83E-01 | -3.54E-01 | -1.28E+00 |
| ENSG000002 | --- | 4.05E-01 | ? | 1.35E+00 | 4.31E-01 | 1.35E+00 |
| ENSG000001 | KCTD3 | 4.05E-01 | 7.33E-01 | 9.71E-01 | -4.27E-02 | -1.03E+00 |
| ENSG000002 | --- | 4.06E-01 | ? | 1.42E+00 | 5.10E-01 | 1.42E+00 |
| ENSG000002 | HSPE1P2 | 4.06E-01 | ? | 1.45E+00 | 5.32E-01 | 1.45E+00 |
| ENSG000001 | GTF2IRD2 | 4.06E-01 | 7.33E-01 | 8.64E-01 | -2.12E-01 | -1.16E+00 |
| ENSG000002 | --- | 4.06E-01 | 7.33E-01 | 9.38E-01 | -9.17E-02 | -1.07E+00 |

| | | | | | | |
|---|---|---|---|---|---|---|
| ENSG000002 | SDAD1P1 | 4.06E-01 | 7.33E-01 | 1.09E+00 | 1.27E-01 | 1.09E+00 |
| ENSG000001 | VIPR1 | 4.06E-01 | 7.33E-01 | 1.17E+00 | 2.22E-01 | 1.17E+00 |
| ENSG000002 | LINC00479 | 4.06E-01 | ? | 6.25E-01 | -6.77E-01 | -1.60E+00 |
| ENSG000001 | LRRC8D | 4.06E-01 | 7.33E-01 | 1.03E+00 | 3.83E-02 | 1.03E+00 |
| ENSG000002 | EBLN2 | 4.06E-01 | 7.33E-01 | 8.65E-01 | -2.10E-01 | -1.16E+00 |
| ENSG000001 | ALPK3 | 4.06E-01 | 7.33E-01 | 8.91E-01 | -1.66E-01 | -1.12E+00 |
| ENSG000002 | NARF-AS2 | 4.06E-01 | 7.33E-01 | 8.78E-01 | -1.88E-01 | -1.14E+00 |
| ENSG000001 | ZNF808 | 4.06E-01 | 7.33E-01 | 1.06E+00 | 7.97E-02 | 1.06E+00 |
| ENSG000001 | CDKN2D | 4.06E-01 | 7.33E-01 | 1.07E+00 | 9.95E-02 | 1.07E+00 |
| ENSG00000C | TMPRSS11E | 4.06E-01 | 7.33E-01 | 1.06E+00 | 8.24E-02 | 1.06E+00 |
| ENSG000002 | --- | 4.06E-01 | ? | 1.16E+00 | 2.19E-01 | 1.16E+00 |
| ENSG000001 | UTP4 | 4.06E-01 | 7.33E-01 | 1.03E+00 | 4.41E-02 | 1.03E+00 |
| ENSG000001 | CCDC74B | 4.06E-01 | ? | 1.28E+00 | 3.53E-01 | 1.28E+00 |
| ENSG000002 | FTH1P20 | 4.06E-01 | ? | 8.57E-01 | -2.23E-01 | -1.17E+00 |
| ENSG000001 | KCNJ1 | 4.06E-01 | ? | 7.02E-01 | -5.11E-01 | -1.42E+00 |
| ENSG000002 | BOLA3-DT | 4.06E-01 | 7.33E-01 | 9.23E-01 | -1.16E-01 | -1.08E+00 |
| ENSG000001 | OSGIN2 | 4.06E-01 | 7.33E-01 | 9.60E-01 | -5.88E-02 | -1.04E+00 |
| ENSG000001 | C3orf14 | 4.06E-01 | 7.33E-01 | 9.57E-01 | -6.34E-02 | -1.04E+00 |
| ENSG000001 | PLCL2 | 4.06E-01 | 7.33E-01 | 9.05E-01 | -1.44E-01 | -1.11E+00 |
| ENSG000001 | KLHL28 | 4.06E-01 | 7.33E-01 | 9.66E-01 | -4.92E-02 | -1.03E+00 |
| ENSG00000C | NDUFB7 | 4.07E-01 | 7.33E-01 | 1.06E+00 | 7.74E-02 | 1.06E+00 |
| ENSG000001 | COMMD7 | 4.07E-01 | 7.33E-01 | 1.03E+00 | 4.19E-02 | 1.03E+00 |
| ENSG000001 | NPIPA1 | 4.07E-01 | 7.33E-01 | 9.55E-01 | -6.63E-02 | -1.05E+00 |
| ENSG000001 | SLC16A6 | 4.07E-01 | ? | 1.35E+00 | 4.37E-01 | 1.35E+00 |
| ENSG000001 | PLA2G7 | 4.07E-01 | ? | 1.18E+00 | 2.44E-01 | 1.18E+00 |
| ENSG000001 | SMARCA4 | 4.07E-01 | 7.33E-01 | 9.81E-01 | -2.72E-02 | -1.02E+00 |
| ENSG000001 | KLHL36 | 4.07E-01 | 7.33E-01 | 1.03E+00 | 4.57E-02 | 1.03E+00 |
| ENSG000002 | --- | 4.07E-01 | ? | 6.32E-01 | -6.62E-01 | -1.58E+00 |
| ENSG00000C | KDM4A | 4.07E-01 | 7.33E-01 | 1.02E+00 | 3.30E-02 | 1.02E+00 |
| ENSG000002 | --- | 4.07E-01 | ? | 7.57E-01 | -4.02E-01 | -1.32E+00 |
| ENSG000001 | YIF1B | 4.07E-01 | 7.33E-01 | 1.06E+00 | 8.10E-02 | 1.06E+00 |
| ENSG000001 | DMBT1 | 4.07E-01 | ? | 1.32E+00 | 3.99E-01 | 1.32E+00 |
| ENSG000001 | SLC27A6 | 4.07E-01 | 7.33E-01 | 9.13E-01 | -1.32E-01 | -1.10E+00 |
| ENSG000001 | CXCL3 | 4.07E-01 | 7.33E-01 | 8.15E-01 | -2.95E-01 | -1.23E+00 |
| ENSG000002 | --- | 4.07E-01 | 7.33E-01 | 9.00E-01 | -1.51E-01 | -1.11E+00 |
| ENSG000001 | WSB2 | 4.07E-01 | 7.33E-01 | 1.03E+00 | 3.82E-02 | 1.03E+00 |
| ENSG000002 | --- | 4.07E-01 | ? | 7.31E-01 | -4.51E-01 | -1.37E+00 |
| ENSG000002 | --- | 4.07E-01 | ? | 6.28E-01 | -6.71E-01 | -1.59E+00 |
| ENSG000002 | BAALC-AS1 | 4.07E-01 | 7.34E-01 | 1.07E+00 | 1.00E-01 | 1.07E+00 |
| ENSG000002 | --- | 4.07E-01 | ? | 4.64E-01 | -1.11E+00 | -2.16E+00 |
| ENSG000001 | CPNE2 | 4.07E-01 | 7.34E-01 | 1.04E+00 | 6.03E-02 | 1.04E+00 |
| ENSG000001 | CEACAM19 | 4.07E-01 | 7.34E-01 | 9.38E-01 | -9.21E-02 | -1.07E+00 |
| ENSG000001 | MARF1 | 4.07E-01 | 7.34E-01 | 9.71E-01 | -4.22E-02 | -1.03E+00 |
| ENSG000001 | KCNN4 | 4.07E-01 | 7.34E-01 | 9.34E-01 | -9.79E-02 | -1.07E+00 |
| ENSG000001 | RAB34 | 4.07E-01 | 7.34E-01 | 1.05E+00 | 7.00E-02 | 1.05E+00 |

| | | | | | | | |
|---|---|---|---|---|---|---|---|
| ENSG000001 | NBL1 | 4.08E-01 | | 7.34E-01 | 1.06E+00 | 8.33E-02 | 1.06E+00 |
| ENSG000002 | --- | 4.08E-01 | ? | | 7.99E-01 | -3.24E-01 | -1.25E+00 |
| ENSG000002 | --- | 4.08E-01 | ? | | 1.26E+00 | 3.31E-01 | 1.26E+00 |
| ENSG000001 | CREB1 | 4.08E-01 | | 7.34E-01 | 9.69E-01 | -4.54E-02 | -1.03E+00 |
| ENSG000002 | --- | 4.08E-01 | ? | | 1.16E+00 | 2.17E-01 | 1.16E+00 |
| ENSG000002 | --- | 4.08E-01 | ? | | 1.28E+00 | 3.57E-01 | 1.28E+00 |
| ENSG000001 | KRT6C | 4.08E-01 | | 7.34E-01 | 1.13E+00 | 1.71E-01 | 1.13E+00 |
| ENSG000001 | NCOA6 | 4.08E-01 | | 7.34E-01 | 9.63E-01 | -5.41E-02 | -1.04E+00 |
| ENSG000001 | CD40 | 4.08E-01 | | 7.34E-01 | 1.13E+00 | 1.73E-01 | 1.13E+00 |
| ENSG000001 | CCNJ | 4.08E-01 | | 7.34E-01 | 9.56E-01 | -6.43E-02 | -1.05E+00 |
| ENSG00000C | TRAF4 | 4.08E-01 | | 7.34E-01 | 1.05E+00 | 6.48E-02 | 1.05E+00 |
| ENSG000001 | KCNJ11 | 4.08E-01 | ? | | 1.50E+00 | 5.88E-01 | 1.50E+00 |
| ENSG00000C | TRMT2A | 4.08E-01 | | 7.34E-01 | 1.06E+00 | 8.33E-02 | 1.06E+00 |
| ENSG000001 | SLC25A15 | 4.08E-01 | | 7.34E-01 | 1.05E+00 | 6.69E-02 | 1.05E+00 |
| ENSG000001 | RINT1 | 4.08E-01 | | 7.34E-01 | 9.69E-01 | -4.47E-02 | -1.03E+00 |
| ENSG000001 | ADTRP | 4.08E-01 | | 7.34E-01 | 9.63E-01 | -5.50E-02 | -1.04E+00 |
| ENSG000002 | ATP2C2-AS1 | 4.08E-01 | | 7.34E-01 | 9.10E-01 | -1.36E-01 | -1.10E+00 |
| ENSG000002 | --- | 4.08E-01 | ? | | 1.34E+00 | 4.22E-01 | 1.34E+00 |
| ENSG000001 | ZNF517 | 4.08E-01 | | 7.34E-01 | 9.15E-01 | -1.29E-01 | -1.09E+00 |
| ENSG000001 | OCIAD1 | 4.09E-01 | | 7.34E-01 | 9.77E-01 | -3.42E-02 | -1.02E+00 |
| ENSG000002 | ST13P18 | 4.09E-01 | ? | | 1.51E+00 | 5.99E-01 | 1.51E+00 |
| ENSG000001 | ATF2 | 4.09E-01 | | 7.34E-01 | 9.70E-01 | -4.44E-02 | -1.03E+00 |
| ENSG000002 | SULT1A4 | 4.09E-01 | | 7.34E-01 | 1.07E+00 | 1.02E-01 | 1.07E+00 |
| ENSG000002 | --- | 4.09E-01 | | 7.34E-01 | 9.13E-01 | -1.31E-01 | -1.09E+00 |
| ENSG000002 | IQCH-AS1 | 4.09E-01 | | 7.34E-01 | 1.07E+00 | 9.66E-02 | 1.07E+00 |
| ENSG00000C | BAZ1B | 4.09E-01 | | 7.35E-01 | 1.03E+00 | 4.33E-02 | 1.03E+00 |
| ENSG000002 | B3GNT9 | 4.09E-01 | | 7.35E-01 | 1.06E+00 | 8.34E-02 | 1.06E+00 |
| ENSG000002 | MIR17HG | 4.09E-01 | | 7.35E-01 | 1.14E+00 | 1.89E-01 | 1.14E+00 |
| ENSG000001 | EHHADH | 4.09E-01 | | 7.35E-01 | 1.03E+00 | 4.75E-02 | 1.03E+00 |
| ENSG000001 | LAD1 | 4.09E-01 | | 7.35E-01 | 1.03E+00 | 3.69E-02 | 1.03E+00 |
| ENSG000001 | ECI2 | 4.09E-01 | | 7.35E-01 | 9.71E-01 | -4.26E-02 | -1.03E+00 |
| ENSG000002 | NPIPB8 | 4.09E-01 | ? | | 6.89E-01 | -5.38E-01 | -1.45E+00 |
| ENSG000001 | DAP | 4.10E-01 | | 7.35E-01 | 9.74E-01 | -3.77E-02 | -1.03E+00 |
| ENSG000001 | ZNF780A | 4.10E-01 | | 7.35E-01 | 1.04E+00 | 5.49E-02 | 1.04E+00 |
| ENSG000001 | SSBP4 | 4.10E-01 | | 7.35E-01 | 1.09E+00 | 1.20E-01 | 1.09E+00 |
| ENSG000002 | --- | 4.10E-01 | | 7.35E-01 | 8.48E-01 | -2.37E-01 | -1.18E+00 |
| ENSG000001 | SUPT6H | 4.10E-01 | | 7.35E-01 | 9.82E-01 | -2.60E-02 | -1.02E+00 |
| ENSG000002 | --- | 4.10E-01 | ? | | 1.77E+00 | 8.28E-01 | 1.77E+00 |
| ENSG000001 | C1orf131 | 4.10E-01 | | 7.35E-01 | 1.04E+00 | 5.89E-02 | 1.04E+00 |
| ENSG000001 | ERH | 4.10E-01 | | 7.35E-01 | 9.74E-01 | -3.87E-02 | -1.03E+00 |
| ENSG000002 | --- | 4.10E-01 | | 7.35E-01 | 9.23E-01 | -1.16E-01 | -1.08E+00 |
| ENSG00000C | KDM5A | 4.10E-01 | | 7.35E-01 | 9.72E-01 | -4.11E-02 | -1.03E+00 |
| ENSG000002 | --- | 4.10E-01 | ? | | 8.06E-01 | -3.11E-01 | -1.24E+00 |
| ENSG000002 | --- | 4.10E-01 | ? | | 1.49E+00 | 5.78E-01 | 1.49E+00 |
| ENSG000002 | DHRS11 | 4.10E-01 | | 7.35E-01 | 9.41E-01 | -8.79E-02 | -1.06E+00 |

| | | | | | | |
|---|---|---|---|---|---|---|
| ENSG000001 | GCNA | 4.10E-01 | 7.35E-01 | 8.93E-01 | -1.63E-01 | -1.12E+00 |
| ENSG00000C | ZCWPW1 | 4.10E-01 | 7.35E-01 | 9.26E-01 | -1.10E-01 | -1.08E+00 |
| ENSG000002 | --- | 4.10E-01 | ? | 1.36E+00 | 4.44E-01 | 1.36E+00 |
| ENSG000001 | KPNA7 | 4.11E-01 | 7.36E-01 | 8.87E-01 | -1.72E-01 | -1.13E+00 |
| ENSG000001 | MYL6B | 4.11E-01 | 7.36E-01 | 1.04E+00 | 6.10E-02 | 1.04E+00 |
| ENSG000001 | SERPING1 | 4.11E-01 | 7.36E-01 | 9.37E-01 | -9.39E-02 | -1.07E+00 |
| ENSG000001 | TEX9 | 4.11E-01 | 7.36E-01 | 9.06E-01 | -1.43E-01 | -1.10E+00 |
| ENSG000001 | RAD9A | 4.11E-01 | 7.36E-01 | 1.09E+00 | 1.18E-01 | 1.09E+00 |
| ENSG000001 | NCAPG2 | 4.11E-01 | 7.36E-01 | 9.52E-01 | -7.14E-02 | -1.05E+00 |
| ENSG000001 | ZNF513 | 4.11E-01 | 7.36E-01 | 1.05E+00 | 6.69E-02 | 1.05E+00 |
| ENSG000001 | CNBD2 | 4.11E-01 | ? | 6.23E-01 | -6.82E-01 | -1.60E+00 |
| ENSG000001 | TRPT1 | 4.11E-01 | 7.36E-01 | 1.08E+00 | 1.13E-01 | 1.08E+00 |
| ENSG000001 | TMEM198B | 4.11E-01 | 7.36E-01 | 9.25E-01 | -1.13E-01 | -1.08E+00 |
| ENSG000001 | PARVA | 4.11E-01 | 7.36E-01 | 1.02E+00 | 3.16E-02 | 1.02E+00 |
| ENSG000002 | --- | 4.11E-01 | ? | 1.46E+00 | 5.44E-01 | 1.46E+00 |
| ENSG00000C | MSMO1 | 4.11E-01 | 7.36E-01 | 1.03E+00 | 3.86E-02 | 1.03E+00 |
| ENSG000002 | --- | 4.11E-01 | 7.36E-01 | 9.04E-01 | -1.45E-01 | -1.11E+00 |
| ENSG000001 | SLC16A7 | 4.11E-01 | 7.36E-01 | 1.12E+00 | 1.61E-01 | 1.12E+00 |
| ENSG000001 | SRGAP1 | 4.11E-01 | 7.36E-01 | 1.04E+00 | 6.28E-02 | 1.04E+00 |
| ENSG000001 | RETREG1 | 4.11E-01 | 7.36E-01 | 9.30E-01 | -1.05E-01 | -1.08E+00 |
| ENSG000002 | --- | 4.11E-01 | ? | 8.66E-01 | -2.08E-01 | -1.16E+00 |
| ENSG000002 | THBS4-AS1 | 4.12E-01 | ? | 1.39E+00 | 4.72E-01 | 1.39E+00 |
| ENSG000002 | --- | 4.12E-01 | ? | 8.45E-01 | -2.43E-01 | -1.18E+00 |
| ENSG000001 | ZNF782 | 4.12E-01 | 7.36E-01 | 9.28E-01 | -1.08E-01 | -1.08E+00 |
| ENSG000001 | SBF1 | 4.12E-01 | 7.36E-01 | 1.06E+00 | 8.68E-02 | 1.06E+00 |
| ENSG000001 | MLLT3 | 4.12E-01 | 7.36E-01 | 1.04E+00 | 5.80E-02 | 1.04E+00 |
| ENSG000001 | HYCC1 | 4.12E-01 | 7.36E-01 | 1.03E+00 | 4.43E-02 | 1.03E+00 |
| ENSG000002 | --- | 4.12E-01 | ? | 7.91E-01 | -3.38E-01 | -1.26E+00 |
| ENSG000001 | CERS5 | 4.12E-01 | 7.37E-01 | 1.03E+00 | 3.72E-02 | 1.03E+00 |
| ENSG000002 | --- | 4.12E-01 | ? | 1.28E+00 | 3.53E-01 | 1.28E+00 |
| ENSG000002 | PVT1 | 4.12E-01 | 7.37E-01 | 9.45E-01 | -8.18E-02 | -1.06E+00 |
| ENSG000002 | STX18-AS1 | 4.12E-01 | 7.37E-01 | 1.07E+00 | 9.86E-02 | 1.07E+00 |
| ENSG000001 | ZNF169 | 4.12E-01 | 7.37E-01 | 9.18E-01 | -1.23E-01 | -1.09E+00 |
| ENSG000002 | --- | 4.12E-01 | 7.37E-01 | 1.08E+00 | 1.10E-01 | 1.08E+00 |
| ENSG000002 | LINC02916 | 4.12E-01 | ? | 7.88E-01 | -3.44E-01 | -1.27E+00 |
| ENSG000001 | NPIPB12 | 4.13E-01 | 7.37E-01 | 8.81E-01 | -1.82E-01 | -1.13E+00 |
| ENSG000001 | IFT43 | 4.13E-01 | 7.37E-01 | 1.06E+00 | 7.85E-02 | 1.06E+00 |
| ENSG000002 | --- | 4.13E-01 | ? | 5.84E-01 | -7.75E-01 | -1.71E+00 |
| ENSG000001 | SMPD3 | 4.13E-01 | ? | 8.14E-01 | -2.97E-01 | -1.23E+00 |
| ENSG000002 | --- | 4.13E-01 | 7.37E-01 | 1.17E+00 | 2.30E-01 | 1.17E+00 |
| ENSG000002 | --- | 4.13E-01 | ? | 7.23E-01 | -4.69E-01 | -1.38E+00 |
| ENSG000001 | YWHAH-AS1 | 4.13E-01 | ? | 7.91E-01 | -3.39E-01 | -1.26E+00 |
| ENSG000002 | --- | 4.13E-01 | ? | 7.00E-01 | -5.14E-01 | -1.43E+00 |
| ENSG000001 | SHB | 4.13E-01 | 7.37E-01 | 9.76E-01 | -3.54E-02 | -1.02E+00 |
| ENSG00000C | ATG16L1 | 4.13E-01 | 7.37E-01 | 1.03E+00 | 4.11E-02 | 1.03E+00 |

| | | | | | | |
|---|---|---|---|---|---|---|
| ENSG000001 | BAZ1A | 4.13E-01 | 7.37E-01 | 9.71E-01 | -4.19E-02 | -1.03E+00 |
| ENSG000002 | TMEM256-PI | 4.13E-01 | ? | 1.34E+00 | 4.22E-01 | 1.34E+00 |
| ENSG000002 | --- | 4.13E-01 | ? | 1.15E+00 | 2.07E-01 | 1.15E+00 |
| ENSG000001 | NINL | 4.13E-01 | 7.37E-01 | 9.37E-01 | -9.42E-02 | -1.07E+00 |
| ENSG000001 | RASA3 | 4.13E-01 | 7.37E-01 | 9.69E-01 | -4.55E-02 | -1.03E+00 |
| ENSG000002 | MED14OS | 4.13E-01 | ? | 1.22E+00 | 2.88E-01 | 1.22E+00 |
| ENSG000001 | DAG1 | 4.13E-01 | 7.37E-01 | 1.02E+00 | 3.49E-02 | 1.02E+00 |
| ENSG000001 | DTYMK | 4.13E-01 | 7.37E-01 | 1.07E+00 | 9.32E-02 | 1.07E+00 |
| ENSG000002 | GIHCG | 4.13E-01 | 7.37E-01 | 9.00E-01 | -1.52E-01 | -1.11E+00 |
| ENSG000001 | SH2D5 | 4.13E-01 | 7.37E-01 | 9.55E-01 | -6.70E-02 | -1.05E+00 |
| ENSG000002 | MYL12BP1 | 4.13E-01 | ? | 1.40E+00 | 4.90E-01 | 1.40E+00 |
| ENSG000001 | TANC2 | 4.13E-01 | 7.37E-01 | 1.03E+00 | 4.70E-02 | 1.03E+00 |
| ENSG000002 | CDK11B | 4.13E-01 | 7.37E-01 | 9.74E-01 | -3.83E-02 | -1.03E+00 |
| ENSG000002 | CCND2-AS1 | 4.13E-01 | ? | 7.71E-01 | -3.76E-01 | -1.30E+00 |
| ENSG000002 | SLC2A1-DT | 4.13E-01 | 7.37E-01 | 1.11E+00 | 1.47E-01 | 1.11E+00 |
| ENSG00000C | ZBTB11 | 4.13E-01 | 7.37E-01 | 9.68E-01 | -4.70E-02 | -1.03E+00 |
| ENSG000001 | HSPB8 | 4.13E-01 | 7.37E-01 | 1.07E+00 | 9.35E-02 | 1.07E+00 |
| ENSG000002 | ZSWIM8 | 4.14E-01 | 7.37E-01 | 1.04E+00 | 5.02E-02 | 1.04E+00 |
| ENSG000001 | POMZP3 | 4.14E-01 | 7.37E-01 | 1.07E+00 | 9.92E-02 | 1.07E+00 |
| ENSG00000C | PPP2R3A | 4.14E-01 | 7.37E-01 | 9.57E-01 | -6.33E-02 | -1.04E+00 |
| ENSG000002 | ANAPC1P2 | 4.14E-01 | 7.37E-01 | 6.98E-01 | -5.18E-01 | -1.43E+00 |
| ENSG000002 | NDUFA6-DT | 4.14E-01 | 7.37E-01 | 1.10E+00 | 1.42E-01 | 1.10E+00 |
| ENSG000001 | LGALS8 | 4.14E-01 | 7.37E-01 | 9.76E-01 | -3.53E-02 | -1.02E+00 |
| ENSG00000C | GPATCH2L | 4.14E-01 | 7.37E-01 | 9.69E-01 | -4.51E-02 | -1.03E+00 |
| ENSG000001 | CAPN10 | 4.14E-01 | 7.37E-01 | 1.06E+00 | 8.62E-02 | 1.06E+00 |
| ENSG000001 | ADCY5 | 4.14E-01 | ? | 8.14E-01 | -2.98E-01 | -1.23E+00 |
| ENSG000001 | NOD2 | 4.14E-01 | ? | 7.94E-01 | -3.32E-01 | -1.26E+00 |
| ENSG000001 | ABL2 | 4.14E-01 | 7.38E-01 | 9.50E-01 | -7.42E-02 | -1.05E+00 |
| ENSG000001 | SRP14 | 4.14E-01 | 7.38E-01 | 9.68E-01 | -4.68E-02 | -1.03E+00 |
| ENSG000001 | CREB3L4 | 4.14E-01 | 7.38E-01 | 1.07E+00 | 1.04E-01 | 1.07E+00 |
| ENSG000002 | --- | 4.15E-01 | ? | 7.56E-01 | -4.03E-01 | -1.32E+00 |
| ENSG000001 | ABHD14A-AC | 4.15E-01 | 7.38E-01 | 1.15E+00 | 2.00E-01 | 1.15E+00 |
| ENSG000001 | ZNF408 | 4.15E-01 | 7.38E-01 | 1.06E+00 | 8.15E-02 | 1.06E+00 |
| ENSG000002 | OR7E128P | 4.15E-01 | ? | 1.46E+00 | 5.43E-01 | 1.46E+00 |
| ENSG000001 | NWD1 | 4.15E-01 | 7.38E-01 | 1.09E+00 | 1.26E-01 | 1.09E+00 |
| ENSG000001 | CPLANE2 | 4.15E-01 | 7.38E-01 | 1.09E+00 | 1.29E-01 | 1.09E+00 |
| ENSG000001 | ILVBL | 4.15E-01 | 7.38E-01 | 1.08E+00 | 1.06E-01 | 1.08E+00 |
| ENSG000002 | --- | 4.15E-01 | 7.38E-01 | 8.87E-01 | -1.72E-01 | -1.13E+00 |
| ENSG000001 | MMD | 4.15E-01 | 7.38E-01 | 1.05E+00 | 7.15E-02 | 1.05E+00 |
| ENSG000001 | KRTCAP2 | 4.15E-01 | 7.38E-01 | 1.04E+00 | 5.07E-02 | 1.04E+00 |
| ENSG000002 | --- | 4.15E-01 | ? | 1.18E+00 | 2.36E-01 | 1.18E+00 |
| ENSG000001 | SGTA | 4.15E-01 | 7.38E-01 | 9.72E-01 | -4.09E-02 | -1.03E+00 |
| ENSG000002 | H2AC19 | 4.15E-01 | 7.38E-01 | 1.11E+00 | 1.44E-01 | 1.11E+00 |
| ENSG000002 | --- | 4.15E-01 | ? | 6.73E-01 | -5.72E-01 | -1.49E+00 |
| ENSG000002 | C22orf39 | 4.15E-01 | 7.38E-01 | 1.04E+00 | 5.97E-02 | 1.04E+00 |

| | | | | | | |
|---|---|---|---|---|---|---|
| ENSG000002 | --- | 4.15E-01 | 7.38E-01 | 9.19E-01 | -1.21E-01 | -1.09E+00 |
| ENSG000001 | NDUFV1-DT | 4.15E-01 | ? | 6.24E-01 | -6.80E-01 | -1.60E+00 |
| ENSG000002 | LINC01127 | 4.15E-01 | ? | 8.30E-01 | -2.69E-01 | -1.21E+00 |
| ENSG000002 | --- | 4.15E-01 | ? | 7.29E-01 | -4.56E-01 | -1.37E+00 |
| ENSG000002 | --- | 4.16E-01 | 7.39E-01 | 9.27E-01 | -1.09E-01 | -1.08E+00 |
| ENSG000001 | ACTA2-AS1 | 4.16E-01 | ? | 7.60E-01 | -3.95E-01 | -1.32E+00 |
| ENSG000002 | --- | 4.16E-01 | ? | 1.37E+00 | 4.52E-01 | 1.37E+00 |
| ENSG000001 | FHL2 | 4.16E-01 | 7.39E-01 | 1.04E+00 | 6.23E-02 | 1.04E+00 |
| ENSG000001 | BEND6 | 4.16E-01 | 7.39E-01 | 1.15E+00 | 1.99E-01 | 1.15E+00 |
| ENSG000002 | PPP3R1 | 4.16E-01 | 7.39E-01 | 9.79E-01 | -2.99E-02 | -1.02E+00 |
| ENSG000001 | TRIM45 | 4.16E-01 | 7.39E-01 | 1.07E+00 | 1.01E-01 | 1.07E+00 |
| ENSG000001 | SPSB3 | 4.16E-01 | 7.39E-01 | 1.07E+00 | 9.97E-02 | 1.07E+00 |
| ENSG000002 | --- | 4.16E-01 | 7.39E-01 | 9.16E-01 | -1.26E-01 | -1.09E+00 |
| ENSG000002 | CYP51A1P1 | 4.16E-01 | ? | 1.33E+00 | 4.12E-01 | 1.33E+00 |
| ENSG000001 | ZNF655 | 4.16E-01 | 7.39E-01 | 9.59E-01 | -5.97E-02 | -1.04E+00 |
| ENSG000002 | --- | 4.16E-01 | 7.39E-01 | 1.17E+00 | 2.25E-01 | 1.17E+00 |
| ENSG000002 | ZFP57 | 4.16E-01 | 7.39E-01 | 1.06E+00 | 8.76E-02 | 1.06E+00 |
| ENSG000001 | H3-5 | 4.16E-01 | ? | 1.53E+00 | 6.16E-01 | 1.53E+00 |
| ENSG000001 | PUM1 | 4.16E-01 | 7.39E-01 | 9.80E-01 | -2.93E-02 | -1.02E+00 |
| ENSG000001 | LMTK2 | 4.16E-01 | 7.39E-01 | 9.73E-01 | -3.99E-02 | -1.03E+00 |
| ENSG000001 | SCUBE3 | 4.16E-01 | ? | 1.23E+00 | 3.00E-01 | 1.23E+00 |
| ENSG00000C | MAMLD1 | 4.16E-01 | 7.39E-01 | 9.22E-01 | -1.17E-01 | -1.08E+00 |
| ENSG00000C | CDC6 | 4.16E-01 | 7.39E-01 | 9.27E-01 | -1.09E-01 | -1.08E+00 |
| ENSG000001 | MAP3K7 | 4.17E-01 | 7.39E-01 | 9.71E-01 | -4.22E-02 | -1.03E+00 |
| ENSG000001 | CDKL1 | 4.17E-01 | 7.39E-01 | 9.44E-01 | -8.31E-02 | -1.06E+00 |
| ENSG000001 | CFL1 | 4.17E-01 | 7.39E-01 | 1.03E+00 | 3.83E-02 | 1.03E+00 |
| ENSG000002 | --- | 4.17E-01 | ? | 1.47E+00 | 5.60E-01 | 1.47E+00 |
| ENSG000001 | NDUFAF2 | 4.17E-01 | 7.39E-01 | 9.52E-01 | -7.02E-02 | -1.05E+00 |
| ENSG000001 | ALDH16A1 | 4.17E-01 | 7.39E-01 | 1.09E+00 | 1.20E-01 | 1.09E+00 |
| ENSG000002 | H2AC15 | 4.17E-01 | ? | 1.28E+00 | 3.58E-01 | 1.28E+00 |
| ENSG000001 | MFSD1 | 4.17E-01 | 7.39E-01 | 1.03E+00 | 3.69E-02 | 1.03E+00 |
| ENSG00000C | GNAI3 | 4.17E-01 | 7.39E-01 | 1.02E+00 | 2.77E-02 | 1.02E+00 |
| ENSG00000C | NUBP2 | 4.17E-01 | 7.39E-01 | 1.08E+00 | 1.14E-01 | 1.08E+00 |
| ENSG000001 | GDF9 | 4.17E-01 | ? | 7.57E-01 | -4.01E-01 | -1.32E+00 |
| ENSG000002 | BTF3L4P2 | 4.17E-01 | ? | 8.10E-01 | -3.04E-01 | -1.23E+00 |
| ENSG000001 | BARD1 | 4.17E-01 | 7.39E-01 | 9.34E-01 | -9.92E-02 | -1.07E+00 |
| ENSG000001 | RPL7AP28 | 4.17E-01 | ? | 1.52E+00 | 6.06E-01 | 1.52E+00 |
| ENSG000001 | SPDYE2B | 4.17E-01 | ? | 7.20E-01 | -4.73E-01 | -1.39E+00 |
| ENSG000001 | C22orf23 | 4.17E-01 | 7.39E-01 | 8.69E-01 | -2.03E-01 | -1.15E+00 |
| ENSG000002 | --- | 4.17E-01 | 7.39E-01 | 1.08E+00 | 1.14E-01 | 1.08E+00 |
| ENSG000001 | HDHD3 | 4.17E-01 | 7.39E-01 | 1.07E+00 | 9.71E-02 | 1.07E+00 |
| ENSG000002 | --- | 4.17E-01 | ? | 1.14E+00 | 1.91E-01 | 1.14E+00 |
| ENSG000001 | ZNF397 | 4.17E-01 | 7.39E-01 | 1.04E+00 | 5.03E-02 | 1.04E+00 |
| ENSG000001 | SEMA5A | 4.17E-01 | 7.39E-01 | 8.95E-01 | -1.60E-01 | -1.12E+00 |
| ENSG000001 | HELZ2 | 4.17E-01 | 7.39E-01 | 1.07E+00 | 9.18E-02 | 1.07E+00 |

| | | | | | | |
|---|---|---|---|---|---|---|
| ENSG000002 | PIGY | 4.17E-01 | 7.39E-01 | 1.04E+00 | 5.12E-02 | 1.04E+00 |
| ENSG000001 | PYURF | 4.17E-01 | 7.39E-01 | 1.04E+00 | 5.12E-02 | 1.04E+00 |
| ENSG000002 | --- | 4.17E-01 | ? | 1.27E+00 | 3.39E-01 | 1.27E+00 |
| ENSG000002 | KMT5AP2 | 4.17E-01 | ? | 1.47E+00 | 5.57E-01 | 1.47E+00 |
| ENSG000001 | DTX3L | 4.17E-01 | 7.39E-01 | 9.65E-01 | -5.20E-02 | -1.04E+00 |
| ENSG000002 | --- | 4.17E-01 | ? | 1.30E+00 | 3.79E-01 | 1.30E+00 |
| ENSG000002 | SCN1A-AS1 | 4.17E-01 | 7.39E-01 | 9.26E-01 | -1.10E-01 | -1.08E+00 |
| ENSG00000C | CNOT3 | 4.17E-01 | 7.39E-01 | 9.71E-01 | -4.19E-02 | -1.03E+00 |
| ENSG000001 | MED11 | 4.18E-01 | 7.39E-01 | 9.53E-01 | -6.90E-02 | -1.05E+00 |
| ENSG000002 | EHBP1-AS1 | 4.18E-01 | 7.40E-01 | 8.96E-01 | -1.58E-01 | -1.12E+00 |
| ENSG000001 | ABCB7 | 4.18E-01 | 7.40E-01 | 1.03E+00 | 4.05E-02 | 1.03E+00 |
| ENSG000001 | RXYLT1 | 4.18E-01 | 7.40E-01 | 9.69E-01 | -4.53E-02 | -1.03E+00 |
| ENSG000002 | SELENOH | 4.18E-01 | 7.40E-01 | 1.07E+00 | 9.42E-02 | 1.07E+00 |
| ENSG000002 | --- | 4.18E-01 | ? | 8.53E-01 | -2.30E-01 | -1.17E+00 |
| ENSG000001 | IQUB | 4.18E-01 | ? | 8.02E-01 | -3.18E-01 | -1.25E+00 |
| ENSG000002 | BOP1 | 4.18E-01 | 7.40E-01 | 1.04E+00 | 6.34E-02 | 1.04E+00 |
| ENSG000001 | UBXN6 | 4.18E-01 | 7.40E-01 | 1.06E+00 | 7.78E-02 | 1.06E+00 |
| ENSG000002 | RPL5P34 | 4.18E-01 | 7.40E-01 | 1.13E+00 | 1.70E-01 | 1.13E+00 |
| ENSG000001 | ZNF594 | 4.18E-01 | 7.40E-01 | 9.24E-01 | -1.14E-01 | -1.08E+00 |
| ENSG000002 | --- | 4.18E-01 | 7.40E-01 | 1.12E+00 | 1.59E-01 | 1.12E+00 |
| ENSG000001 | DHRS3 | 4.18E-01 | 7.40E-01 | 1.10E+00 | 1.32E-01 | 1.10E+00 |
| ENSG000001 | TBC1D15 | 4.18E-01 | 7.40E-01 | 9.64E-01 | -5.29E-02 | -1.04E+00 |
| ENSG000002 | MAP3K14-AS | 4.18E-01 | 7.40E-01 | 9.35E-01 | -9.70E-02 | -1.07E+00 |
| ENSG00000C | ABCC2 | 4.18E-01 | 7.40E-01 | 1.08E+00 | 1.05E-01 | 1.08E+00 |
| ENSG000002 | --- | 4.19E-01 | ? | 1.43E+00 | 5.13E-01 | 1.43E+00 |
| ENSG000002 | --- | 4.19E-01 | ? | 7.82E-01 | -3.54E-01 | -1.28E+00 |
| ENSG000001 | IL15RA | 4.19E-01 | 7.40E-01 | 9.31E-01 | -1.03E-01 | -1.07E+00 |
| ENSG000002 | --- | 4.19E-01 | ? | 7.89E-01 | -3.41E-01 | -1.27E+00 |
| ENSG000001 | ZNF443 | 4.19E-01 | 7.40E-01 | 9.26E-01 | -1.10E-01 | -1.08E+00 |
| ENSG000001 | PHTF1 | 4.19E-01 | 7.40E-01 | 9.64E-01 | -5.32E-02 | -1.04E+00 |
| ENSG000002 | RNF217-AS1 | 4.19E-01 | 7.40E-01 | 1.09E+00 | 1.26E-01 | 1.09E+00 |
| ENSG000001 | C1QTNF1 | 4.19E-01 | 7.40E-01 | 1.07E+00 | 9.91E-02 | 1.07E+00 |
| ENSG000002 | --- | 4.19E-01 | ? | 7.52E-01 | -4.12E-01 | -1.33E+00 |
| ENSG000001 | PPP2CA | 4.19E-01 | 7.40E-01 | 1.01E+00 | 2.06E-02 | 1.01E+00 |
| ENSG000001 | NDUFS7 | 4.19E-01 | 7.40E-01 | 1.07E+00 | 9.48E-02 | 1.07E+00 |
| ENSG00000C | IP6K2 | 4.19E-01 | 7.40E-01 | 1.03E+00 | 3.90E-02 | 1.03E+00 |
| ENSG000001 | IL6-AS1 | 4.19E-01 | ? | 1.29E+00 | 3.70E-01 | 1.29E+00 |
| ENSG000001 | ENOSF1 | 4.19E-01 | 7.40E-01 | 9.62E-01 | -5.65E-02 | -1.04E+00 |
| ENSG000002 | LINC00629 | 4.19E-01 | ? | 1.38E+00 | 4.69E-01 | 1.38E+00 |
| ENSG000001 | ULK3 | 4.19E-01 | 7.40E-01 | 1.05E+00 | 6.95E-02 | 1.05E+00 |
| ENSG000001 | GRAMD1C | 4.19E-01 | 7.40E-01 | 9.03E-01 | -1.47E-01 | -1.11E+00 |
| ENSG000001 | SMAD6 | 4.19E-01 | ? | 1.56E+00 | 6.37E-01 | 1.56E+00 |
| ENSG000001 | PAK5 | 4.19E-01 | ? | 1.15E+00 | 1.98E-01 | 1.15E+00 |
| ENSG000001 | SEC23IP | 4.20E-01 | 7.40E-01 | 9.77E-01 | -3.42E-02 | -1.02E+00 |
| ENSG000002 | --- | 4.20E-01 | 7.40E-01 | 8.87E-01 | -1.73E-01 | -1.13E+00 |

| | | | | | | |
|---|---|---|---|---|---|---|
| ENSG000002 | --- | 4.20E-01 | 7.40E-01 | 1.08E+00 | 1.07E-01 | 1.08E+00 |
| ENSG00000C | MYO9A | 4.20E-01 | 7.40E-01 | 9.60E-01 | -5.93E-02 | -1.04E+00 |
| ENSG00000C | UBR2 | 4.20E-01 | 7.40E-01 | 9.75E-01 | -3.61E-02 | -1.03E+00 |
| ENSG000002 | --- | 4.20E-01 ? | | 8.21E-01 | -2.85E-01 | -1.22E+00 |
| ENSG000001 | IL17RE | 4.20E-01 | 7.40E-01 | 1.06E+00 | 7.84E-02 | 1.06E+00 |
| ENSG000002 | SMG1P7 | 4.20E-01 ? | | 8.29E-01 | -2.70E-01 | -1.21E+00 |
| ENSG000002 | LINC00536 | 4.20E-01 ? | | 1.55E+00 | 6.36E-01 | 1.55E+00 |
| ENSG000001 | RHOU | 4.20E-01 ? | | 8.71E-01 | -1.99E-01 | -1.15E+00 |
| ENSG00000C | TCF3 | 4.20E-01 | 7.40E-01 | 9.64E-01 | -5.25E-02 | -1.04E+00 |
| ENSG000001 | WARS2 | 4.20E-01 | 7.40E-01 | 9.66E-01 | -5.05E-02 | -1.04E+00 |
| ENSG000001 | SEMA6C | 4.20E-01 | 7.41E-01 | 1.09E+00 | 1.22E-01 | 1.09E+00 |
| ENSG000002 | --- | 4.20E-01 ? | | 1.20E+00 | 2.61E-01 | 1.20E+00 |
| ENSG000001 | ZNF227 | 4.20E-01 | 7.41E-01 | 9.54E-01 | -6.83E-02 | -1.05E+00 |
| ENSG00000C | CASP8 | 4.20E-01 | 7.41E-01 | 1.03E+00 | 4.07E-02 | 1.03E+00 |
| ENSG000002 | ISCA1P1 | 4.20E-01 ? | | 1.27E+00 | 3.43E-01 | 1.27E+00 |
| ENSG000001 | TMEM127 | 4.20E-01 | 7.41E-01 | 1.02E+00 | 3.17E-02 | 1.02E+00 |
| ENSG000001 | ALYREF | 4.21E-01 | 7.41E-01 | 9.64E-01 | -5.23E-02 | -1.04E+00 |
| ENSG000001 | LHPP | 4.21E-01 | 7.41E-01 | 1.08E+00 | 1.11E-01 | 1.08E+00 |
| ENSG000001 | PLIN3 | 4.21E-01 | 7.41E-01 | 9.61E-01 | -5.75E-02 | -1.04E+00 |
| ENSG000002 | MIR29B2CHG | 4.21E-01 | 7.41E-01 | 9.27E-01 | -1.09E-01 | -1.08E+00 |
| ENSG000002 | --- | 4.21E-01 ? | | 1.23E+00 | 2.96E-01 | 1.23E+00 |
| ENSG000002 | DPP3 | 4.21E-01 | 7.41E-01 | 1.04E+00 | 5.82E-02 | 1.04E+00 |
| ENSG000001 | CEP76 | 4.21E-01 | 7.41E-01 | 1.05E+00 | 6.59E-02 | 1.05E+00 |
| ENSG000001 | NUDCD2 | 4.21E-01 | 7.41E-01 | 9.71E-01 | -4.18E-02 | -1.03E+00 |
| ENSG000002 | --- | 4.21E-01 ? | | 6.82E-01 | -5.52E-01 | -1.47E+00 |
| ENSG000001 | GALNT10 | 4.21E-01 | 7.41E-01 | 1.03E+00 | 3.76E-02 | 1.03E+00 |
| ENSG000002 | TWF2 | 4.21E-01 | 7.41E-01 | 1.04E+00 | 5.82E-02 | 1.04E+00 |
| ENSG000001 | MYOZ3 | 4.21E-01 ? | | 8.39E-01 | -2.53E-01 | -1.19E+00 |
| ENSG000002 | LINC01829 | 4.21E-01 ? | | 7.04E-01 | -5.07E-01 | -1.42E+00 |
| ENSG000001 | COPS3 | 4.21E-01 | 7.41E-01 | 1.02E+00 | 2.94E-02 | 1.02E+00 |
| ENSG000002 | SUGT1P4-ST | 4.21E-01 | 7.41E-01 | 1.11E+00 | 1.54E-01 | 1.11E+00 |
| ENSG000001 | TRMT61B | 4.21E-01 | 7.41E-01 | 1.04E+00 | 5.35E-02 | 1.04E+00 |
| ENSG000001 | NCAPD3 | 4.21E-01 | 7.41E-01 | 1.03E+00 | 3.98E-02 | 1.03E+00 |
| ENSG000001 | ERCC6L2 | 4.21E-01 | 7.41E-01 | 1.04E+00 | 5.33E-02 | 1.04E+00 |
| ENSG000002 | RAMACL | 4.21E-01 ? | | 1.20E+00 | 2.61E-01 | 1.20E+00 |
| ENSG000001 | IER5L | 4.21E-01 | 7.41E-01 | 9.16E-01 | -1.27E-01 | -1.09E+00 |
| ENSG000002 | RPP21 | 4.21E-01 | 7.41E-01 | 1.06E+00 | 8.59E-02 | 1.06E+00 |
| ENSG000001 | TRIM66 | 4.22E-01 | 7.41E-01 | 9.42E-01 | -8.70E-02 | -1.06E+00 |
| ENSG00000C | WWTR1 | 4.22E-01 | 7.41E-01 | 9.59E-01 | -6.08E-02 | -1.04E+00 |
| ENSG000002 | --- | 4.22E-01 ? | | 1.35E+00 | 4.28E-01 | 1.35E+00 |
| ENSG000001 | PCDHB10 | 4.22E-01 ? | | 6.71E-01 | -5.77E-01 | -1.49E+00 |
| ENSG00000C | ISOC2 | 4.22E-01 | 7.42E-01 | 1.07E+00 | 9.89E-02 | 1.07E+00 |
| ENSG00000C | ZNF506 | 4.22E-01 | 7.42E-01 | 1.05E+00 | 7.56E-02 | 1.05E+00 |
| ENSG000001 | ZNF605 | 4.22E-01 | 7.42E-01 | 9.60E-01 | -5.94E-02 | -1.04E+00 |
| ENSG000001 | SLC35A5 | 4.22E-01 | 7.42E-01 | 9.73E-01 | -3.93E-02 | -1.03E+00 |

| | | | | | | |
|---|---|---|---|---|---|---|
| ENSG000002 | ACVR2B-AS1 | 4.22E-01 | ? | 7.22E-01 | -4.71E-01 | -1.39E+00 |
| ENSG000001 | TAS2R5 | 4.22E-01 | ? | 8.79E-01 | -1.87E-01 | -1.14E+00 |
| ENSG000002 | TRIM39 | 4.22E-01 | 7.42E-01 | 9.69E-01 | -4.48E-02 | -1.03E+00 |
| ENSG000002 | CCDC153 | 4.23E-01 | ? | 1.24E+00 | 3.07E-01 | 1.24E+00 |
| ENSG000001 | TMEM51 | 4.23E-01 | 7.42E-01 | 1.03E+00 | 4.47E-02 | 1.03E+00 |
| ENSG000001 | CHRM4 | 4.23E-01 | ? | 7.70E-01 | -3.77E-01 | -1.30E+00 |
| ENSG000001 | RNASEH2B | 4.23E-01 | 7.42E-01 | 9.63E-01 | -5.42E-02 | -1.04E+00 |
| ENSG000002 | HEXD-IT1 | 4.23E-01 | ? | 1.21E+00 | 2.77E-01 | 1.21E+00 |
| ENSG000001 | VANGL2 | 4.23E-01 | 7.43E-01 | 1.04E+00 | 5.40E-02 | 1.04E+00 |
| ENSG000001 | CALCOCO2 | 4.23E-01 | 7.43E-01 | 9.70E-01 | -4.41E-02 | -1.03E+00 |
| ENSG000002 | --- | 4.23E-01 | 7.43E-01 | 1.10E+00 | 1.44E-01 | 1.10E+00 |
| ENSG000001 | HTRA1 | 4.23E-01 | 7.43E-01 | 1.09E+00 | 1.29E-01 | 1.09E+00 |
| ENSG00000C | MARK4 | 4.23E-01 | 7.43E-01 | 9.78E-01 | -3.14E-02 | -1.02E+00 |
| ENSG000001 | PPM1F | 4.23E-01 | 7.43E-01 | 1.04E+00 | 5.73E-02 | 1.04E+00 |
| ENSG000001 | SERPINA9 | 4.23E-01 | 7.43E-01 | 8.25E-01 | -2.78E-01 | -1.21E+00 |
| ENSG00000C | EDN1 | 4.23E-01 | 7.43E-01 | 9.54E-01 | -6.84E-02 | -1.05E+00 |
| ENSG00000C | ARHGAP33 | 4.23E-01 | 7.43E-01 | 1.12E+00 | 1.64E-01 | 1.12E+00 |
| ENSG000001 | SYNGAP1 | 4.24E-01 | 7.43E-01 | 9.63E-01 | -5.46E-02 | -1.04E+00 |
| ENSG000001 | ACTG1 | 4.24E-01 | 7.43E-01 | 9.68E-01 | -4.71E-02 | -1.03E+00 |
| ENSG000001 | NOL8 | 4.24E-01 | 7.43E-01 | 9.69E-01 | -4.58E-02 | -1.03E+00 |
| ENSG000002 | --- | 4.24E-01 | ? | 1.87E+00 | 9.06E-01 | 1.87E+00 |
| ENSG000001 | MTNR1A | 4.24E-01 | ? | 1.31E+00 | 3.88E-01 | 1.31E+00 |
| ENSG000001 | TPST1 | 4.24E-01 | 7.44E-01 | 1.04E+00 | 5.99E-02 | 1.04E+00 |
| ENSG000001 | OIT3 | 4.24E-01 | ? | 1.64E+00 | 7.16E-01 | 1.64E+00 |
| ENSG000002 | FABP5P7 | 4.24E-01 | 7.44E-01 | 8.83E-01 | -1.79E-01 | -1.13E+00 |
| ENSG000002 | HTD2 | 4.24E-01 | 7.44E-01 | 9.06E-01 | -1.43E-01 | -1.10E+00 |
| ENSG000002 | --- | 4.24E-01 | 7.44E-01 | 1.11E+00 | 1.49E-01 | 1.11E+00 |
| ENSG000001 | DNAJC24 | 4.24E-01 | 7.44E-01 | 9.65E-01 | -5.16E-02 | -1.04E+00 |
| ENSG000002 | --- | 4.24E-01 | ? | 7.10E-01 | -4.95E-01 | -1.41E+00 |
| ENSG000002 | ZNRF2P2 | 4.24E-01 | ? | 7.55E-01 | -4.05E-01 | -1.32E+00 |
| ENSG000001 | WFS1 | 4.24E-01 | 7.44E-01 | 1.05E+00 | 6.60E-02 | 1.05E+00 |
| ENSG000001 | CSGALNACT1 | 4.25E-01 | ? | 1.28E+00 | 3.57E-01 | 1.28E+00 |
| ENSG000002 | RPL5P1 | 4.25E-01 | ? | 1.17E+00 | 2.26E-01 | 1.17E+00 |
| ENSG000001 | HCFC2 | 4.25E-01 | 7.44E-01 | 9.63E-01 | -5.40E-02 | -1.04E+00 |
| ENSG000001 | RHOF | 4.25E-01 | 7.45E-01 | 9.52E-01 | -7.16E-02 | -1.05E+00 |
| ENSG000001 | ZFPM2 | 4.25E-01 | ? | 7.97E-01 | -3.27E-01 | -1.25E+00 |
| ENSG000001 | ZSCAN32 | 4.25E-01 | 7.45E-01 | 9.65E-01 | -5.20E-02 | -1.04E+00 |
| ENSG000002 | RPS3AP38 | 4.25E-01 | ? | 8.13E-01 | -2.99E-01 | -1.23E+00 |
| ENSG000002 | --- | 4.25E-01 | 7.45E-01 | 1.56E+00 | 6.45E-01 | 1.56E+00 |
| ENSG000001 | WTIP | 4.25E-01 | 7.45E-01 | 1.06E+00 | 8.78E-02 | 1.06E+00 |
| ENSG000001 | MMAA | 4.25E-01 | 7.45E-01 | 9.49E-01 | -7.61E-02 | -1.05E+00 |
| ENSG000002 | TAF9 | 4.25E-01 | 7.45E-01 | 9.75E-01 | -3.58E-02 | -1.03E+00 |
| ENSG000001 | SCML2 | 4.25E-01 | ? | 8.36E-01 | -2.58E-01 | -1.20E+00 |
| ENSG000002 | ITGB1P1 | 4.25E-01 | 7.45E-01 | 1.04E+00 | 6.33E-02 | 1.04E+00 |
| ENSG000001 | ZBTB21 | 4.25E-01 | 7.45E-01 | 9.62E-01 | -5.61E-02 | -1.04E+00 |

| | | | | | | |
|---|---|---|---|---|---|---|
| ENSG000002 | --- | 4.25E-01 | ? | 7.31E-01 | -4.52E-01 | -1.37E+00 |
| ENSG000001 | HIRA | 4.25E-01 | 7.45E-01 | 1.03E+00 | 3.82E-02 | 1.03E+00 |
| ENSG000002 | LINC02290 | 4.25E-01 | ? | 7.42E-01 | -4.30E-01 | -1.35E+00 |
| ENSG000002 | CCHCR1 | 4.25E-01 | 7.45E-01 | 9.31E-01 | -1.04E-01 | -1.07E+00 |
| ENSG000002 | THUMPD3-A: | 4.26E-01 | 7.45E-01 | 9.34E-01 | -9.86E-02 | -1.07E+00 |
| ENSG000001 | NACA3P | 4.26E-01 | ? | 7.43E-01 | -4.29E-01 | -1.35E+00 |
| ENSG000002 | --- | 4.26E-01 | ? | 1.31E+00 | 3.94E-01 | 1.31E+00 |
| ENSG000002 | MT1H | 4.26E-01 | ? | 7.33E-01 | -4.49E-01 | -1.36E+00 |
| ENSG000001 | SYNPO2L | 4.26E-01 | ? | 1.48E+00 | 5.61E-01 | 1.48E+00 |
| ENSG000001 | SLC12A9 | 4.26E-01 | 7.45E-01 | 1.05E+00 | 7.14E-02 | 1.05E+00 |
| ENSG000002 | --- | 4.26E-01 | 7.45E-01 | 9.21E-01 | -1.19E-01 | -1.09E+00 |
| ENSG000001 | SNRPD1 | 4.26E-01 | 7.45E-01 | 1.03E+00 | 4.38E-02 | 1.03E+00 |
| ENSG000001 | ARSJ | 4.26E-01 | 7.45E-01 | 9.57E-01 | -6.30E-02 | -1.04E+00 |
| ENSG000002 | --- | 4.26E-01 | ? | 8.38E-01 | -2.55E-01 | -1.19E+00 |
| ENSG000001 | TARS1 | 4.26E-01 | 7.45E-01 | 9.76E-01 | -3.49E-02 | -1.02E+00 |
| ENSG00000C | MAPK9 | 4.26E-01 | 7.45E-01 | 1.03E+00 | 3.93E-02 | 1.03E+00 |
| ENSG000002 | SRGAP2 | 4.26E-01 | 7.45E-01 | 1.03E+00 | 4.49E-02 | 1.03E+00 |
| ENSG000002 | LNP1 | 4.26E-01 | ? | 1.17E+00 | 2.21E-01 | 1.17E+00 |
| ENSG000002 | --- | 4.26E-01 | ? | 7.29E-01 | -4.56E-01 | -1.37E+00 |
| ENSG000001 | ARSG | 4.26E-01 | 7.45E-01 | 1.06E+00 | 8.67E-02 | 1.06E+00 |
| ENSG000002 | --- | 4.26E-01 | ? | 7.19E-01 | -4.76E-01 | -1.39E+00 |
| ENSG000001 | SKI | 4.26E-01 | 7.45E-01 | 9.76E-01 | -3.46E-02 | -1.02E+00 |
| ENSG000001 | RASA4B | 4.26E-01 | 7.45E-01 | 9.21E-01 | -1.19E-01 | -1.09E+00 |
| ENSG000001 | SPART | 4.26E-01 | 7.45E-01 | 1.02E+00 | 3.48E-02 | 1.02E+00 |
| ENSG000001 | CYP2R1 | 4.26E-01 | 7.45E-01 | 1.04E+00 | 5.84E-02 | 1.04E+00 |
| ENSG000002 | --- | 4.26E-01 | ? | 7.69E-01 | -3.78E-01 | -1.30E+00 |
| ENSG000001 | CCL20 | 4.27E-01 | 7.45E-01 | 8.99E-01 | -1.54E-01 | -1.11E+00 |
| ENSG000001 | PDAP1 | 4.27E-01 | 7.45E-01 | 1.02E+00 | 2.97E-02 | 1.02E+00 |
| ENSG000002 | HERC3 | 4.27E-01 | 7.45E-01 | 1.23E+00 | 2.98E-01 | 1.23E+00 |
| ENSG000001 | ZFP41 | 4.27E-01 | 7.45E-01 | 1.05E+00 | 7.18E-02 | 1.05E+00 |
| ENSG000001 | ECE1 | 4.27E-01 | 7.45E-01 | 9.78E-01 | -3.27E-02 | -1.02E+00 |
| ENSG000002 | --- | 4.27E-01 | ? | 7.69E-01 | -3.79E-01 | -1.30E+00 |
| ENSG000002 | --- | 4.27E-01 | 7.45E-01 | 9.40E-01 | -8.88E-02 | -1.06E+00 |
| ENSG000001 | LPIN3 | 4.27E-01 | 7.45E-01 | 1.04E+00 | 5.95E-02 | 1.04E+00 |
| ENSG000002 | --- | 4.27E-01 | ? | 1.22E+00 | 2.89E-01 | 1.22E+00 |
| ENSG000002 | ATP10A | 4.27E-01 | 7.45E-01 | 8.97E-01 | -1.56E-01 | -1.11E+00 |
| ENSG000002 | DSTNP1 | 4.27E-01 | 7.45E-01 | 8.94E-01 | -1.61E-01 | -1.12E+00 |
| ENSG000002 | PAIP1P1 | 4.27E-01 | ? | 1.28E+00 | 3.55E-01 | 1.28E+00 |
| ENSG000002 | --- | 4.27E-01 | ? | 7.03E-01 | -5.08E-01 | -1.42E+00 |
| ENSG000002 | SPRR2E | 4.27E-01 | 7.46E-01 | 1.24E+00 | 3.12E-01 | 1.24E+00 |
| ENSG000001 | CTCF | 4.27E-01 | 7.46E-01 | 9.69E-01 | -4.56E-02 | -1.03E+00 |
| ENSG000002 | --- | 4.27E-01 | ? | 1.14E+00 | 1.87E-01 | 1.14E+00 |
| ENSG000001 | FAM182A | 4.28E-01 | 7.46E-01 | 8.89E-01 | -1.70E-01 | -1.12E+00 |
| ENSG000001 | NOTCH1 | 4.28E-01 | 7.46E-01 | 1.06E+00 | 8.35E-02 | 1.06E+00 |
| ENSG000001 | ZNF326 | 4.28E-01 | 7.46E-01 | 9.51E-01 | -7.25E-02 | -1.05E+00 |

| | | | | | | |
|---|---|---|---|---|---|---|
| ENSG000001 | MYG1 | 4.28E-01 | 7.46E-01 | 1.12E+00 | 1.62E-01 | 1.12E+00 |
| ENSG000001 | TRIM61 | 4.28E-01 | 7.46E-01 | 8.88E-01 | -1.71E-01 | -1.13E+00 |
| ENSG000002 | TMEM253 | 4.28E-01 | 7.46E-01 | 1.19E+00 | 2.55E-01 | 1.19E+00 |
| ENSG000000 | REXO5 | 4.28E-01 | 7.46E-01 | 1.11E+00 | 1.45E-01 | 1.11E+00 |
| ENSG000001 | PFDN4 | 4.28E-01 | 7.46E-01 | 9.50E-01 | -7.44E-02 | -1.05E+00 |
| ENSG000001 | ESS2 | 4.28E-01 | 7.46E-01 | 1.03E+00 | 4.64E-02 | 1.03E+00 |
| ENSG000001 | PTRHD1 | 4.28E-01 | 7.46E-01 | 1.06E+00 | 7.86E-02 | 1.06E+00 |
| ENSG000001 | MRFAP1L1 | 4.28E-01 | 7.46E-01 | 9.74E-01 | -3.75E-02 | -1.03E+00 |
| ENSG000001 | SLC25A11 | 4.28E-01 | 7.47E-01 | 1.05E+00 | 7.06E-02 | 1.05E+00 |
| ENSG000001 | NUDT17 | 4.28E-01 | 7.47E-01 | 9.03E-01 | -1.48E-01 | -1.11E+00 |
| ENSG000002 | --- | 4.28E-01 | ? | 8.28E-01 | -2.73E-01 | -1.21E+00 |
| ENSG000002 | YPEL3-DT | 4.29E-01 | ? | 1.44E+00 | 5.22E-01 | 1.44E+00 |
| ENSG000001 | MRPL32 | 4.29E-01 | 7.47E-01 | 9.67E-01 | -4.80E-02 | -1.03E+00 |
| ENSG000001 | SRPK2 | 4.29E-01 | 7.47E-01 | 1.02E+00 | 3.41E-02 | 1.02E+00 |
| ENSG000001 | DIP2A | 4.29E-01 | 7.47E-01 | 1.03E+00 | 3.78E-02 | 1.03E+00 |
| ENSG000000 | CDON | 4.29E-01 | 7.47E-01 | 9.37E-01 | -9.44E-02 | -1.07E+00 |
| ENSG000002 | FTH1P10 | 4.29E-01 | 7.47E-01 | 9.02E-01 | -1.49E-01 | -1.11E+00 |
| ENSG000001 | TRIM4 | 4.29E-01 | 7.47E-01 | 1.03E+00 | 4.21E-02 | 1.03E+00 |
| ENSG000001 | LRP5 | 4.29E-01 | 7.47E-01 | 1.04E+00 | 6.23E-02 | 1.04E+00 |
| ENSG000002 | NFAM1 | 4.29E-01 | ? | 1.38E+00 | 4.65E-01 | 1.38E+00 |
| ENSG000001 | ZNF154 | 4.29E-01 | 7.47E-01 | 1.09E+00 | 1.28E-01 | 1.09E+00 |
| ENSG000002 | FTH1P8 | 4.29E-01 | 7.47E-01 | 8.83E-01 | -1.79E-01 | -1.13E+00 |
| ENSG000001 | GPX1P1 | 4.29E-01 | ? | 1.19E+00 | 2.56E-01 | 1.19E+00 |
| ENSG000002 | LINC02887 | 4.29E-01 | 7.47E-01 | 9.03E-01 | -1.46E-01 | -1.11E+00 |
| ENSG000001 | ETV4 | 4.29E-01 | 7.47E-01 | 1.03E+00 | 4.69E-02 | 1.03E+00 |
| ENSG000001 | ACTR5 | 4.29E-01 | 7.47E-01 | 1.05E+00 | 7.34E-02 | 1.05E+00 |
| ENSG000000 | NCDN | 4.29E-01 | 7.47E-01 | 1.05E+00 | 6.77E-02 | 1.05E+00 |
| ENSG000001 | STX2 | 4.29E-01 | 7.47E-01 | 9.74E-01 | -3.79E-02 | -1.03E+00 |
| ENSG000002 | --- | 4.29E-01 | ? | 7.38E-01 | -4.37E-01 | -1.35E+00 |
| ENSG000001 | PRRT2 | 4.29E-01 | ? | 1.16E+00 | 2.18E-01 | 1.16E+00 |
| ENSG000001 | --- | 4.29E-01 | 7.47E-01 | 1.20E+00 | 2.60E-01 | 1.20E+00 |
| ENSG000001 | GPX7 | 4.29E-01 | ? | 1.18E+00 | 2.35E-01 | 1.18E+00 |
| ENSG000001 | SORL1 | 4.30E-01 | 7.47E-01 | 1.06E+00 | 8.28E-02 | 1.06E+00 |
| ENSG000001 | TNFSF18 | 4.30E-01 | ? | 7.79E-01 | -3.61E-01 | -1.28E+00 |
| ENSG000001 | DTL | 4.30E-01 | 7.47E-01 | 8.91E-01 | -1.67E-01 | -1.12E+00 |
| ENSG000001 | NUTM2F | 4.30E-01 | ? | 1.39E+00 | 4.79E-01 | 1.39E+00 |
| ENSG000001 | IER2 | 4.30E-01 | 7.47E-01 | 9.42E-01 | -8.60E-02 | -1.06E+00 |
| ENSG000001 | SYCP2L | 4.30E-01 | 7.47E-01 | 8.84E-01 | -1.78E-01 | -1.13E+00 |
| ENSG000001 | SMPD4BP | 4.30E-01 | 7.47E-01 | 9.40E-01 | -8.87E-02 | -1.06E+00 |
| ENSG000001 | LRPPRC | 4.30E-01 | 7.47E-01 | 1.02E+00 | 3.00E-02 | 1.02E+00 |
| ENSG000001 | BDH1 | 4.30E-01 | 7.47E-01 | 9.52E-01 | -7.14E-02 | -1.05E+00 |
| ENSG000001 | HERC2 | 4.30E-01 | 7.47E-01 | 9.61E-01 | -5.78E-02 | -1.04E+00 |
| ENSG000001 | FAM241B | 4.30E-01 | 7.47E-01 | 9.34E-01 | -9.87E-02 | -1.07E+00 |
| ENSG000002 | SLC35B4 | 4.30E-01 | 7.47E-01 | 9.74E-01 | -3.75E-02 | -1.03E+00 |
| ENSG000002 | RARA-AS1 | 4.30E-01 | ? | 7.48E-01 | -4.18E-01 | -1.34E+00 |

| | | | | | | |
|---|---|---|---|---|---|---|
| ENSG000002 | --- | 4.30E-01 | 7.48E-01 | 1.10E+00 | 1.43E-01 | 1.10E+00 |
| ENSG000002 | HSPA8P9 | 4.30E-01 | ? | 6.94E-01 | -5.26E-01 | -1.44E+00 |
| ENSG000001 | TRMT1 | 4.30E-01 | 7.48E-01 | 1.04E+00 | 5.59E-02 | 1.04E+00 |
| ENSG000001 | ZNF575 | 4.30E-01 | ? | 8.22E-01 | -2.83E-01 | -1.22E+00 |
| ENSG000001 | TBC1D9B | 4.30E-01 | 7.48E-01 | 1.03E+00 | 4.10E-02 | 1.03E+00 |
| ENSG000001 | CERT1 | 4.31E-01 | 7.48E-01 | 1.02E+00 | 3.40E-02 | 1.02E+00 |
| ENSG000001 | GUK1 | 4.31E-01 | 7.48E-01 | 1.04E+00 | 5.16E-02 | 1.04E+00 |
| ENSG000001 | ZNF627 | 4.31E-01 | 7.48E-01 | 1.04E+00 | 5.96E-02 | 1.04E+00 |
| ENSG000001 | SCNM1 | 4.31E-01 | 7.48E-01 | 9.64E-01 | -5.32E-02 | -1.04E+00 |
| ENSG000002 | --- | 4.31E-01 | ? | 6.40E-01 | -6.44E-01 | -1.56E+00 |
| ENSG000001 | ZNF135 | 4.31E-01 | 7.48E-01 | 9.45E-01 | -8.17E-02 | -1.06E+00 |
| ENSG000001 | CINP | 4.31E-01 | 7.48E-01 | 1.03E+00 | 4.40E-02 | 1.03E+00 |
| ENSG000001 | NEPRO | 4.31E-01 | 7.48E-01 | 9.80E-01 | -2.97E-02 | -1.02E+00 |
| ENSG00000C | ZBTB32 | 4.31E-01 | ? | 7.56E-01 | -4.04E-01 | -1.32E+00 |
| ENSG000002 | KRT8P39 | 4.31E-01 | 7.48E-01 | 8.66E-01 | -2.08E-01 | -1.16E+00 |
| ENSG000001 | ZNF500 | 4.31E-01 | 7.48E-01 | 1.06E+00 | 8.52E-02 | 1.06E+00 |
| ENSG000002 | --- | 4.31E-01 | ? | 1.45E+00 | 5.32E-01 | 1.45E+00 |
| ENSG000001 | PCYOX1 | 4.31E-01 | 7.48E-01 | 1.03E+00 | 4.14E-02 | 1.03E+00 |
| ENSG000001 | TUBBP1 | 4.31E-01 | 7.48E-01 | 1.12E+00 | 1.67E-01 | 1.12E+00 |
| ENSG000002 | TENM3 | 4.31E-01 | 7.48E-01 | 8.90E-01 | -1.67E-01 | -1.12E+00 |
| ENSG000002 | SMN2 | 4.31E-01 | 7.48E-01 | 9.48E-01 | -7.72E-02 | -1.05E+00 |
| ENSG000001 | TMCO4 | 4.31E-01 | 7.48E-01 | 1.08E+00 | 1.14E-01 | 1.08E+00 |
| ENSG000002 | MICA | 4.31E-01 | 7.48E-01 | 1.02E+00 | 2.97E-02 | 1.02E+00 |
| ENSG000002 | ZNF234 | 4.31E-01 | 7.48E-01 | 9.58E-01 | -6.12E-02 | -1.04E+00 |
| ENSG000002 | MRPL33 | 4.32E-01 | 7.48E-01 | 1.03E+00 | 4.84E-02 | 1.03E+00 |
| ENSG000001 | FGG | 4.32E-01 | ? | 6.99E-01 | -5.17E-01 | -1.43E+00 |
| ENSG000001 | DUS1L | 4.32E-01 | 7.48E-01 | 1.04E+00 | 5.77E-02 | 1.04E+00 |
| ENSG000001 | ZNF286A | 4.32E-01 | 7.48E-01 | 1.04E+00 | 5.33E-02 | 1.04E+00 |
| ENSG000001 | SEPTIN6 | 4.32E-01 | 7.48E-01 | 9.52E-01 | -7.17E-02 | -1.05E+00 |
| ENSG000001 | OGG1 | 4.32E-01 | 7.48E-01 | 1.05E+00 | 6.50E-02 | 1.05E+00 |
| ENSG000001 | RGPD2 | 4.32E-01 | ? | 7.56E-01 | -4.04E-01 | -1.32E+00 |
| ENSG000001 | CA3 | 4.32E-01 | ? | 1.28E+00 | 3.56E-01 | 1.28E+00 |
| ENSG000002 | --- | 4.32E-01 | ? | 1.58E+00 | 6.58E-01 | 1.58E+00 |
| ENSG000002 | GANC | 4.32E-01 | 7.49E-01 | 1.03E+00 | 4.05E-02 | 1.03E+00 |
| ENSG00000C | ANGEL1 | 4.32E-01 | 7.49E-01 | 1.03E+00 | 4.89E-02 | 1.03E+00 |
| ENSG000001 | ZNF197 | 4.32E-01 | 7.49E-01 | 9.72E-01 | -4.04E-02 | -1.03E+00 |
| ENSG000002 | --- | 4.32E-01 | ? | 7.91E-01 | -3.38E-01 | -1.26E+00 |
| ENSG000002 | --- | 4.32E-01 | ? | 1.57E+00 | 6.55E-01 | 1.57E+00 |
| ENSG000001 | CEP63 | 4.32E-01 | 7.49E-01 | 9.57E-01 | -6.34E-02 | -1.04E+00 |
| ENSG000001 | TDP2 | 4.32E-01 | 7.49E-01 | 1.02E+00 | 2.98E-02 | 1.02E+00 |
| ENSG000001 | OTUD4 | 4.33E-01 | 7.49E-01 | 9.79E-01 | -3.03E-02 | -1.02E+00 |
| ENSG000002 | SMG7-AS1 | 4.33E-01 | 7.49E-01 | 1.12E+00 | 1.62E-01 | 1.12E+00 |
| ENSG000002 | --- | 4.33E-01 | ? | 1.19E+00 | 2.50E-01 | 1.19E+00 |
| ENSG000002 | --- | 4.33E-01 | ? | 7.91E-01 | -3.38E-01 | -1.26E+00 |
| ENSG000002 | PRICKLE4 | 4.33E-01 | ? | 1.40E+00 | 4.86E-01 | 1.40E+00 |

| | | | | | | |
|---|---|---|---|---|---|---|
| ENSG000002 | --- | 4.33E-01 | ? | 8.62E-01 | -2.15E-01 | -1.16E+00 |
| ENSG000001 | MT1E | 4.33E-01 | 7.49E-01 | 9.59E-01 | -6.08E-02 | -1.04E+00 |
| ENSG000001 | SNX29P1 | 4.33E-01 | ? | 1.44E+00 | 5.22E-01 | 1.44E+00 |
| ENSG000001 | IMPG1 | 4.33E-01 | ? | 1.34E+00 | 4.18E-01 | 1.34E+00 |
| ENSG000001 | SLU7 | 4.33E-01 | 7.49E-01 | 9.71E-01 | -4.28E-02 | -1.03E+00 |
| ENSG000001 | DHODH | 4.33E-01 | 7.49E-01 | 1.05E+00 | 7.23E-02 | 1.05E+00 |
| ENSG00000C | ATP9A | 4.33E-01 | 7.49E-01 | 1.04E+00 | 5.12E-02 | 1.04E+00 |
| ENSG000002 | --- | 4.33E-01 | 7.49E-01 | 9.20E-01 | -1.20E-01 | -1.09E+00 |
| ENSG000002 | A1BG-AS1 | 4.33E-01 | 7.49E-01 | 1.06E+00 | 8.38E-02 | 1.06E+00 |
| ENSG000002 | C17orf49 | 4.33E-01 | 7.49E-01 | 1.05E+00 | 6.78E-02 | 1.05E+00 |
| ENSG000001 | STK16 | 4.33E-01 | 7.49E-01 | 1.04E+00 | 5.75E-02 | 1.04E+00 |
| ENSG000001 | FAM72B | 4.33E-01 | 7.49E-01 | 8.83E-01 | -1.80E-01 | -1.13E+00 |
| ENSG000002 | N4BP2L2-IT2 | 4.34E-01 | ? | 8.63E-01 | -2.13E-01 | -1.16E+00 |
| ENSG000002 | --- | 4.34E-01 | 7.50E-01 | 1.14E+00 | 1.91E-01 | 1.14E+00 |
| ENSG000001 | SLC6A6 | 4.34E-01 | 7.50E-01 | 1.04E+00 | 5.01E-02 | 1.04E+00 |
| ENSG000001 | ZNF626 | 4.34E-01 | 7.50E-01 | 9.57E-01 | -6.34E-02 | -1.04E+00 |
| ENSG000002 | LINC02542 | 4.34E-01 | ? | 8.04E-01 | -3.14E-01 | -1.24E+00 |
| ENSG000002 | RPL21P16 | 4.34E-01 | 7.50E-01 | 1.05E+00 | 6.90E-02 | 1.05E+00 |
| ENSG000001 | NOS1AP | 4.34E-01 | ? | 7.72E-01 | -3.73E-01 | -1.30E+00 |
| ENSG000002 | CYFIP1 | 4.34E-01 | 7.50E-01 | 1.03E+00 | 4.89E-02 | 1.03E+00 |
| ENSG000001 | YIF1A | 4.34E-01 | 7.50E-01 | 1.05E+00 | 7.21E-02 | 1.05E+00 |
| ENSG000001 | KIF7 | 4.34E-01 | 7.50E-01 | 1.10E+00 | 1.33E-01 | 1.10E+00 |
| ENSG000002 | --- | 4.34E-01 | ? | 8.59E-01 | -2.19E-01 | -1.16E+00 |
| ENSG000001 | EEF2K | 4.34E-01 | 7.50E-01 | 1.03E+00 | 4.17E-02 | 1.03E+00 |
| ENSG000001 | PCDH19 | 4.34E-01 | 7.50E-01 | 1.26E+00 | 3.29E-01 | 1.26E+00 |
| ENSG000002 | LIN37 | 4.34E-01 | 7.50E-01 | 9.54E-01 | -6.74E-02 | -1.05E+00 |
| ENSG000001 | DHX38 | 4.34E-01 | 7.50E-01 | 9.82E-01 | -2.63E-02 | -1.02E+00 |
| ENSG000001 | DAGLB | 4.34E-01 | 7.50E-01 | 1.03E+00 | 4.70E-02 | 1.03E+00 |
| ENSG000001 | OGFOD3 | 4.34E-01 | 7.50E-01 | 1.05E+00 | 6.45E-02 | 1.05E+00 |
| ENSG000002 | --- | 4.35E-01 | ? | 6.89E-01 | -5.37E-01 | -1.45E+00 |
| ENSG000002 | TBC1D3C | 4.35E-01 | 7.50E-01 | 1.17E+00 | 2.25E-01 | 1.17E+00 |
| ENSG000001 | GDF15 | 4.35E-01 | 7.50E-01 | 9.29E-01 | -1.06E-01 | -1.08E+00 |
| ENSG000002 | --- | 4.35E-01 | ? | 8.80E-01 | -1.84E-01 | -1.14E+00 |
| ENSG000001 | ZFAND3 | 4.35E-01 | 7.50E-01 | 1.02E+00 | 2.57E-02 | 1.02E+00 |
| ENSG000001 | ORMDL1 | 4.35E-01 | 7.50E-01 | 9.69E-01 | -4.58E-02 | -1.03E+00 |
| ENSG000001 | MLYCD | 4.35E-01 | 7.50E-01 | 9.38E-01 | -9.17E-02 | -1.07E+00 |
| ENSG000001 | CCDC12 | 4.35E-01 | 7.50E-01 | 1.04E+00 | 5.45E-02 | 1.04E+00 |
| ENSG000002 | --- | 4.35E-01 | ? | 5.69E-01 | -8.14E-01 | -1.76E+00 |
| ENSG00000C | DERL2 | 4.35E-01 | 7.50E-01 | 9.78E-01 | -3.17E-02 | -1.02E+00 |
| ENSG000001 | AHNAK2 | 4.35E-01 | 7.50E-01 | 1.05E+00 | 6.98E-02 | 1.05E+00 |
| ENSG000001 | AK3 | 4.35E-01 | 7.50E-01 | 9.68E-01 | -4.65E-02 | -1.03E+00 |
| ENSG000001 | SYT9 | 4.35E-01 | ? | 1.16E+00 | 2.12E-01 | 1.16E+00 |
| ENSG000001 | NRBF2 | 4.36E-01 | 7.51E-01 | 1.02E+00 | 3.35E-02 | 1.02E+00 |
| ENSG000001 | SLC24A4 | 4.36E-01 | ? | 1.14E+00 | 1.90E-01 | 1.14E+00 |
| ENSG000001 | ENTPD4 | 4.36E-01 | 7.51E-01 | 1.02E+00 | 3.44E-02 | 1.02E+00 |

| | | | | | | |
|---|---|---|---|---|---|---|
| ENSG000001 | TBC1D31 | 4.36E-01 | 7.51E-01 | 1.06E+00 | 7.91E-02 | 1.06E+00 |
| ENSG000002 | LNCSRLR | 4.36E-01 | 7.51E-01 | 8.78E-01 | -1.87E-01 | -1.14E+00 |
| ENSG000002 | ERVK13-1 | 4.36E-01 | 7.51E-01 | 9.36E-01 | -9.55E-02 | -1.07E+00 |
| ENSG000002 | --- | 4.36E-01 | 7.51E-01 | 1.10E+00 | 1.40E-01 | 1.10E+00 |
| ENSG000001 | ABCC10 | 4.36E-01 | 7.51E-01 | 1.05E+00 | 7.02E-02 | 1.05E+00 |
| ENSG000002 | --- | 4.36E-01 | 7.51E-01 | 1.09E+00 | 1.25E-01 | 1.09E+00 |
| ENSG000001 | SLC4A2 | 4.36E-01 | 7.51E-01 | 1.04E+00 | 5.82E-02 | 1.04E+00 |
| ENSG000001 | MSS51 | 4.36E-01 | 7.51E-01 | 8.79E-01 | -1.87E-01 | -1.14E+00 |
| ENSG000001 | FN1 | 4.36E-01 | 7.51E-01 | 9.56E-01 | -6.45E-02 | -1.05E+00 |
| ENSG000001 | PRX | 4.36E-01 | ? | 8.62E-01 | -2.14E-01 | -1.16E+00 |
| ENSG000002 | --- | 4.36E-01 | 7.51E-01 | 1.07E+00 | 9.30E-02 | 1.07E+00 |
| ENSG000001 | RBM12B | 4.36E-01 | 7.51E-01 | 9.73E-01 | -3.90E-02 | -1.03E+00 |
| ENSG000002 | LYPLA2P1 | 4.36E-01 | ? | 1.51E+00 | 5.90E-01 | 1.51E+00 |
| ENSG000002 | --- | 4.37E-01 | ? | 1.53E+00 | 6.17E-01 | 1.53E+00 |
| ENSG00000C | SNCB | 4.37E-01 | ? | 7.98E-01 | -3.25E-01 | -1.25E+00 |
| ENSG000001 | ZNF107 | 4.37E-01 | 7.51E-01 | 9.48E-01 | -7.67E-02 | -1.05E+00 |
| ENSG000001 | GRIN1 | 4.37E-01 | ? | 8.24E-01 | -2.79E-01 | -1.21E+00 |
| ENSG000002 | MT-TE | 4.37E-01 | ? | 7.67E-01 | -3.82E-01 | -1.30E+00 |
| ENSG000001 | PRELID3B | 4.37E-01 | 7.52E-01 | 9.65E-01 | -5.08E-02 | -1.04E+00 |
| ENSG000001 | PRRG2 | 4.37E-01 | 7.52E-01 | 1.07E+00 | 9.84E-02 | 1.07E+00 |
| ENSG000002 | --- | 4.37E-01 | ? | 1.18E+00 | 2.43E-01 | 1.18E+00 |
| ENSG000001 | UBXN2A | 4.37E-01 | 7.52E-01 | 9.76E-01 | -3.46E-02 | -1.02E+00 |
| ENSG000002 | LINC00242 | 4.37E-01 | ? | 1.47E+00 | 5.52E-01 | 1.47E+00 |
| ENSG000001 | MED16 | 4.37E-01 | 7.52E-01 | 1.04E+00 | 5.71E-02 | 1.04E+00 |
| ENSG000001 | MRPS11 | 4.37E-01 | 7.52E-01 | 9.68E-01 | -4.72E-02 | -1.03E+00 |
| ENSG00000C | NAALAD2 | 4.37E-01 | ? | 8.74E-01 | -1.94E-01 | -1.14E+00 |
| ENSG000002 | --- | 4.37E-01 | ? | 1.23E+00 | 2.96E-01 | 1.23E+00 |
| ENSG000002 | TMEM167AP | 4.37E-01 | ? | 1.15E+00 | 2.04E-01 | 1.15E+00 |
| ENSG000002 | CCT7P1 | 4.37E-01 | ? | 1.38E+00 | 4.66E-01 | 1.38E+00 |
| ENSG000002 | --- | 4.37E-01 | ? | 6.56E-01 | -6.08E-01 | -1.52E+00 |
| ENSG000002 | CNPY2 | 4.37E-01 | 7.52E-01 | 9.67E-01 | -4.81E-02 | -1.03E+00 |
| ENSG000001 | DMGDH | 4.37E-01 | ? | 6.40E-01 | -6.44E-01 | -1.56E+00 |
| ENSG000002 | RENO1 | 4.37E-01 | 7.52E-01 | 8.82E-01 | -1.82E-01 | -1.13E+00 |
| ENSG000002 | --- | 4.37E-01 | 7.52E-01 | 9.40E-01 | -8.86E-02 | -1.06E+00 |
| ENSG000001 | GPR143 | 4.37E-01 | ? | 1.22E+00 | 2.82E-01 | 1.22E+00 |
| ENSG000002 | C4orf48 | 4.37E-01 | ? | 1.36E+00 | 4.45E-01 | 1.36E+00 |
| ENSG000001 | RRM1 | 4.37E-01 | 7.52E-01 | 1.03E+00 | 4.62E-02 | 1.03E+00 |
| ENSG000002 | CKMT2-AS1 | 4.37E-01 | 7.52E-01 | 1.08E+00 | 1.13E-01 | 1.08E+00 |
| ENSG000002 | --- | 4.37E-01 | ? | 1.35E+00 | 4.31E-01 | 1.35E+00 |
| ENSG000002 | SMIM1 | 4.37E-01 | ? | 1.40E+00 | 4.84E-01 | 1.40E+00 |
| ENSG000001 | SLC35D1 | 4.38E-01 | 7.52E-01 | 9.69E-01 | -4.51E-02 | -1.03E+00 |
| ENSG000001 | KALRN | 4.38E-01 | 7.52E-01 | 9.08E-01 | -1.38E-01 | -1.10E+00 |
| ENSG00000C | HPF1 | 4.38E-01 | 7.52E-01 | 9.52E-01 | -7.03E-02 | -1.05E+00 |
| ENSG000002 | RASAL2-AS1 | 4.38E-01 | 7.52E-01 | 1.13E+00 | 1.77E-01 | 1.13E+00 |
| ENSG000001 | DDX19B | 4.38E-01 | 7.52E-01 | 1.03E+00 | 4.18E-02 | 1.03E+00 |

| | | | | | | |
|---|---|---|---|---|---|---|
| ENSG000001 | CYSTM1 | 4.38E-01 | 7.53E-01 | 1.04E+00 | 5.49E-02 | 1.04E+00 |
| ENSG000001 | IAH1 | 4.38E-01 | 7.53E-01 | 1.03E+00 | 4.55E-02 | 1.03E+00 |
| ENSG000002 | POU2F2-AS1 | 4.38E-01 | ? | 7.74E-01 | -3.69E-01 | -1.29E+00 |
| ENSG00000C | FKBP5 | 4.39E-01 | 7.53E-01 | 1.03E+00 | 4.66E-02 | 1.03E+00 |
| ENSG000001 | PPP1R3C | 4.39E-01 | 7.54E-01 | 9.23E-01 | -1.15E-01 | -1.08E+00 |
| ENSG000001 | OAZ1 | 4.39E-01 | 7.54E-01 | 9.58E-01 | -6.13E-02 | -1.04E+00 |
| ENSG000001 | KCNQ5 | 4.39E-01 | 7.54E-01 | 9.56E-01 | -6.44E-02 | -1.05E+00 |
| ENSG000002 | RNA5-8SN3 | 4.39E-01 | ? | 1.43E+00 | 5.18E-01 | 1.43E+00 |
| ENSG000002 | RNA5-8SN1 | 4.39E-01 | ? | 1.43E+00 | 5.18E-01 | 1.43E+00 |
| ENSG000002 | RNA5-8SN2 | 4.39E-01 | ? | 1.43E+00 | 5.18E-01 | 1.43E+00 |
| ENSG000002 | 5_8S_rRNA | 4.39E-01 | ? | 1.43E+00 | 5.18E-01 | 1.43E+00 |
| ENSG000002 | --- | 4.39E-01 | 7.54E-01 | 8.93E-01 | -1.63E-01 | -1.12E+00 |
| ENSG00000C | APBA3 | 4.39E-01 | 7.54E-01 | 1.06E+00 | 8.00E-02 | 1.06E+00 |
| ENSG000001 | POTEE | 4.39E-01 | ? | 1.26E+00 | 3.30E-01 | 1.26E+00 |
| ENSG000002 | --- | 4.39E-01 | ? | 8.34E-01 | -2.62E-01 | -1.20E+00 |
| ENSG00000C | XYLB | 4.39E-01 | 7.54E-01 | 1.06E+00 | 8.27E-02 | 1.06E+00 |
| ENSG000001 | SNRPE | 4.39E-01 | 7.54E-01 | 1.03E+00 | 3.97E-02 | 1.03E+00 |
| ENSG000002 | --- | 4.39E-01 | 7.54E-01 | 1.05E+00 | 7.68E-02 | 1.05E+00 |
| ENSG000001 | SLC41A1 | 4.39E-01 | 7.54E-01 | 9.75E-01 | -3.65E-02 | -1.03E+00 |
| ENSG000002 | --- | 4.39E-01 | ? | 1.16E+00 | 2.12E-01 | 1.16E+00 |
| ENSG000002 | PCGF2 | 4.39E-01 | 7.54E-01 | 1.04E+00 | 5.42E-02 | 1.04E+00 |
| ENSG00000C | MRPS35 | 4.40E-01 | 7.54E-01 | 9.75E-01 | -3.71E-02 | -1.03E+00 |
| ENSG000002 | LINC01589 | 4.40E-01 | ? | 1.43E+00 | 5.16E-01 | 1.43E+00 |
| ENSG000001 | LMO4 | 4.40E-01 | 7.54E-01 | 1.04E+00 | 6.10E-02 | 1.04E+00 |
| ENSG000002 | --- | 4.40E-01 | 7.54E-01 | 8.92E-01 | -1.65E-01 | -1.12E+00 |
| ENSG00000C | OAT | 4.40E-01 | 7.54E-01 | 1.03E+00 | 4.58E-02 | 1.03E+00 |
| ENSG000002 | LINC03016 | 4.40E-01 | ? | 1.21E+00 | 2.75E-01 | 1.21E+00 |
| ENSG000001 | TNRC6B | 4.40E-01 | 7.54E-01 | 9.53E-01 | -6.92E-02 | -1.05E+00 |
| ENSG000001 | YTHDF1 | 4.40E-01 | 7.54E-01 | 9.80E-01 | -2.86E-02 | -1.02E+00 |
| ENSG000002 | AMZ2P1 | 4.40E-01 | 7.55E-01 | 9.20E-01 | -1.20E-01 | -1.09E+00 |
| ENSG000001 | PCDH7 | 4.40E-01 | 7.55E-01 | 1.04E+00 | 5.36E-02 | 1.04E+00 |
| ENSG000001 | LHFPL2 | 4.40E-01 | 7.55E-01 | 9.48E-01 | -7.73E-02 | -1.06E+00 |
| ENSG000001 | PSMD13 | 4.40E-01 | 7.55E-01 | 1.02E+00 | 2.95E-02 | 1.02E+00 |
| ENSG000001 | SLC22A13 | 4.40E-01 | ? | 1.35E+00 | 4.34E-01 | 1.35E+00 |
| ENSG000001 | SMC6 | 4.40E-01 | 7.55E-01 | 9.66E-01 | -5.06E-02 | -1.04E+00 |
| ENSG000001 | FBRSL1 | 4.41E-01 | 7.55E-01 | 9.58E-01 | -6.15E-02 | -1.04E+00 |
| ENSG000001 | ABCC5 | 4.41E-01 | 7.55E-01 | 9.61E-01 | -5.70E-02 | -1.04E+00 |
| ENSG000001 | AZIN2 | 4.41E-01 | 7.55E-01 | 9.13E-01 | -1.31E-01 | -1.09E+00 |
| ENSG000001 | SYNGR3 | 4.41E-01 | 7.55E-01 | 1.13E+00 | 1.81E-01 | 1.13E+00 |
| ENSG000001 | AMT | 4.41E-01 | 7.55E-01 | 8.45E-01 | -2.43E-01 | -1.18E+00 |
| ENSG000001 | SMIM10L2B | 4.41E-01 | ? | 8.39E-01 | -2.53E-01 | -1.19E+00 |
| ENSG000002 | --- | 4.41E-01 | ? | 8.56E-01 | -2.24E-01 | -1.17E+00 |
| ENSG000001 | TLR1 | 4.41E-01 | ? | 7.86E-01 | -3.48E-01 | -1.27E+00 |
| ENSG000001 | BPNT2 | 4.41E-01 | 7.55E-01 | 9.68E-01 | -4.68E-02 | -1.03E+00 |
| ENSG000002 | --- | 4.41E-01 | ? | 8.29E-01 | -2.71E-01 | -1.21E+00 |

| | | | | | | | |
|---|---|---|---|---|---|---|---|
| ENSG000002 | --- | 4.41E-01 | | 7.55E-01 | 1.05E+00 | 6.78E-02 | 1.05E+00 |
| ENSG000002 | ASB14 | 4.41E-01 | ? | | 8.58E-01 | -2.22E-01 | -1.17E+00 |
| ENSG000001 | SLCO3A1 | 4.41E-01 | | 7.55E-01 | 1.04E+00 | 6.27E-02 | 1.04E+00 |
| ENSG000002 | MUC20-OT1 | 4.41E-01 | | 7.55E-01 | 9.46E-01 | -8.06E-02 | -1.06E+00 |
| ENSG000001 | PSMG4 | 4.41E-01 | | 7.55E-01 | 1.05E+00 | 6.41E-02 | 1.05E+00 |
| ENSG000002 | --- | 4.41E-01 | | 7.55E-01 | 8.69E-01 | -2.02E-01 | -1.15E+00 |
| ENSG000001 | ZYG11B | 4.41E-01 | | 7.55E-01 | 9.74E-01 | -3.81E-02 | -1.03E+00 |
| ENSG000001 | FCHSD1 | 4.41E-01 | | 7.55E-01 | 9.47E-01 | -7.89E-02 | -1.06E+00 |
| ENSG000001 | CIT | 4.41E-01 | | 7.55E-01 | 9.30E-01 | -1.05E-01 | -1.08E+00 |
| ENSG000002 | PTGES3P3 | 4.42E-01 | ? | | 7.65E-01 | -3.87E-01 | -1.31E+00 |
| ENSG000002 | LINC02298 | 4.42E-01 | | 7.55E-01 | 9.09E-01 | -1.38E-01 | -1.10E+00 |
| ENSG000002 | ZNF793-AS1 | 4.42E-01 | ? | | 1.15E+00 | 2.05E-01 | 1.15E+00 |
| ENSG000001 | ANKK1 | 4.42E-01 | | 7.55E-01 | 1.12E+00 | 1.65E-01 | 1.12E+00 |
| ENSG000001 | PAIP1P2 | 4.42E-01 | | 7.55E-01 | 8.98E-01 | -1.56E-01 | -1.11E+00 |
| ENSG00000C | BUD23 | 4.42E-01 | | 7.55E-01 | 1.03E+00 | 4.35E-02 | 1.03E+00 |
| ENSG000001 | PRDM16-DT | 4.42E-01 | ? | | 1.35E+00 | 4.36E-01 | 1.35E+00 |
| ENSG000001 | PTPN22 | 4.42E-01 | ? | | 1.29E+00 | 3.63E-01 | 1.29E+00 |
| ENSG000002 | --- | 4.42E-01 | | 7.56E-01 | 8.59E-01 | -2.19E-01 | -1.16E+00 |
| ENSG000001 | KLHL6 | 4.42E-01 | ? | | 1.18E+00 | 2.44E-01 | 1.18E+00 |
| ENSG000002 | NPM1P24 | 4.42E-01 | ? | | 7.80E-01 | -3.58E-01 | -1.28E+00 |
| ENSG00000C | MAT2B | 4.42E-01 | | 7.56E-01 | 1.02E+00 | 2.89E-02 | 1.02E+00 |
| ENSG000001 | RIMKLB | 4.42E-01 | | 7.56E-01 | 1.06E+00 | 8.53E-02 | 1.06E+00 |
| ENSG000002 | --- | 4.42E-01 | ? | | 6.66E-01 | -5.87E-01 | -1.50E+00 |
| ENSG000002 | SCARNA7 | 4.42E-01 | | 7.56E-01 | 8.84E-01 | -1.78E-01 | -1.13E+00 |
| ENSG000001 | VPS11 | 4.42E-01 | | 7.56E-01 | 1.02E+00 | 3.40E-02 | 1.02E+00 |
| ENSG000001 | PNMA1 | 4.42E-01 | | 7.56E-01 | 9.76E-01 | -3.44E-02 | -1.02E+00 |
| ENSG000002 | SNRPGP10 | 4.43E-01 | ? | | 8.60E-01 | -2.18E-01 | -1.16E+00 |
| ENSG000001 | GGA1 | 4.43E-01 | | 7.56E-01 | 1.04E+00 | 5.60E-02 | 1.04E+00 |
| ENSG000001 | TEFM | 4.43E-01 | | 7.56E-01 | 1.04E+00 | 6.13E-02 | 1.04E+00 |
| ENSG000002 | PDCD6IP-DT | 4.43E-01 | | 7.56E-01 | 9.62E-01 | -5.58E-02 | -1.04E+00 |
| ENSG000001 | RAB22A | 4.43E-01 | | 7.56E-01 | 9.71E-01 | -4.22E-02 | -1.03E+00 |
| ENSG000001 | EXTL2 | 4.43E-01 | | 7.56E-01 | 9.52E-01 | -7.11E-02 | -1.05E+00 |
| ENSG000002 | LINC03032 | 4.43E-01 | ? | | 8.20E-01 | -2.86E-01 | -1.22E+00 |
| ENSG00000C | MOK | 4.43E-01 | | 7.56E-01 | 9.46E-01 | -8.08E-02 | -1.06E+00 |
| ENSG000002 | ZNF630 | 4.43E-01 | | 7.56E-01 | 9.33E-01 | -1.00E-01 | -1.07E+00 |
| ENSG000001 | FAM120AOS | 4.43E-01 | | 7.56E-01 | 9.66E-01 | -5.05E-02 | -1.04E+00 |
| ENSG00000C | AKR1B1 | 4.43E-01 | | 7.56E-01 | 1.05E+00 | 6.43E-02 | 1.05E+00 |
| ENSG000002 | LINC01151 | 4.43E-01 | ? | | 1.23E+00 | 2.93E-01 | 1.23E+00 |
| ENSG000001 | RGCC | 4.43E-01 | | 7.56E-01 | 8.85E-01 | -1.77E-01 | -1.13E+00 |
| ENSG000002 | HSP90B2P | 4.43E-01 | ? | | 7.59E-01 | -3.98E-01 | -1.32E+00 |
| ENSG000002 | --- | 4.43E-01 | ? | | 7.04E-01 | -5.07E-01 | -1.42E+00 |
| ENSG000001 | TMEM126B | 4.43E-01 | | 7.56E-01 | 1.05E+00 | 6.82E-02 | 1.05E+00 |
| ENSG000002 | SMG1P6 | 4.43E-01 | ? | | 1.32E+00 | 3.99E-01 | 1.32E+00 |
| ENSG000001 | NUDCD1 | 4.43E-01 | | 7.56E-01 | 1.03E+00 | 3.58E-02 | 1.03E+00 |
| ENSG000001 | TMEM219 | 4.43E-01 | | 7.56E-01 | 1.05E+00 | 7.39E-02 | 1.05E+00 |

| | | | | | | |
|---|---|---|---|---|---|---|
| ENSG000002 | C2orf27A | 4.43E-01 | 7.56E-01 | 1.04E+00 | 5.97E-02 | 1.04E+00 |
| ENSG00000C | ZNF195 | 4.43E-01 | 7.56E-01 | 9.46E-01 | -7.96E-02 | -1.06E+00 |
| ENSG000001 | PLGRKT | 4.43E-01 | 7.56E-01 | 9.70E-01 | -4.40E-02 | -1.03E+00 |
| ENSG000001 | PDZD7 | 4.43E-01 | 7.56E-01 | 9.22E-01 | -1.18E-01 | -1.09E+00 |
| ENSG00000C | ITGA3 | 4.44E-01 | 7.56E-01 | 1.03E+00 | 3.92E-02 | 1.03E+00 |
| ENSG000002 | PA2G4P5 | 4.44E-01 | ? | 1.32E+00 | 4.02E-01 | 1.32E+00 |
| ENSG000001 | KDM5B | 4.44E-01 | 7.56E-01 | 1.02E+00 | 3.06E-02 | 1.02E+00 |
| ENSG000001 | CRISPLD2 | 4.44E-01 | 7.56E-01 | 8.87E-01 | -1.73E-01 | -1.13E+00 |
| ENSG000001 | GTF2E2 | 4.44E-01 | 7.56E-01 | 9.69E-01 | -4.54E-02 | -1.03E+00 |
| ENSG000001 | SPIRE1 | 4.44E-01 | 7.56E-01 | 1.03E+00 | 4.40E-02 | 1.03E+00 |
| ENSG000002 | --- | 4.44E-01 | ? | 8.17E-01 | -2.92E-01 | -1.22E+00 |
| ENSG000002 | DYNC1I2P1 | 4.44E-01 | 7.56E-01 | 1.10E+00 | 1.31E-01 | 1.10E+00 |
| ENSG00000C | ERP44 | 4.44E-01 | 7.56E-01 | 9.77E-01 | -3.31E-02 | -1.02E+00 |
| ENSG000001 | EVC2 | 4.44E-01 | 7.56E-01 | 9.54E-01 | -6.86E-02 | -1.05E+00 |
| ENSG000001 | CACFD1 | 4.44E-01 | 7.57E-01 | 1.09E+00 | 1.29E-01 | 1.09E+00 |
| ENSG000001 | NSUN5 | 4.44E-01 | 7.57E-01 | 9.55E-01 | -6.68E-02 | -1.05E+00 |
| ENSG00000C | FMO4 | 4.44E-01 | 7.57E-01 | 9.22E-01 | -1.18E-01 | -1.08E+00 |
| ENSG000001 | EIF5B | 4.44E-01 | 7.57E-01 | 9.72E-01 | -4.17E-02 | -1.03E+00 |
| ENSG000002 | SLC9A7P1 | 4.45E-01 | 7.57E-01 | 1.13E+00 | 1.79E-01 | 1.13E+00 |
| ENSG000002 | PSME2P2 | 4.45E-01 | ? | 1.23E+00 | 3.01E-01 | 1.23E+00 |
| ENSG000002 | SLC16A1-AS1 | 4.45E-01 | 7.57E-01 | 9.54E-01 | -6.82E-02 | -1.05E+00 |
| ENSG000001 | PPIA | 4.45E-01 | 7.57E-01 | 1.01E+00 | 2.12E-02 | 1.01E+00 |
| ENSG00000C | MSH2 | 4.45E-01 | 7.57E-01 | 9.69E-01 | -4.62E-02 | -1.03E+00 |
| ENSG000001 | TIAM2 | 4.45E-01 | 7.57E-01 | 9.35E-01 | -9.71E-02 | -1.07E+00 |
| ENSG000002 | --- | 4.45E-01 | ? | 1.48E+00 | 5.69E-01 | 1.48E+00 |
| ENSG000001 | MTRR | 4.45E-01 | 7.57E-01 | 9.74E-01 | -3.77E-02 | -1.03E+00 |
| ENSG000002 | --- | 4.45E-01 | ? | 1.42E+00 | 5.04E-01 | 1.42E+00 |
| ENSG000001 | PAXIP1 | 4.45E-01 | 7.57E-01 | 9.69E-01 | -4.58E-02 | -1.03E+00 |
| ENSG000001 | MTRF1 | 4.45E-01 | 7.57E-01 | 9.53E-01 | -6.97E-02 | -1.05E+00 |
| ENSG000002 | --- | 4.45E-01 | 7.57E-01 | 1.04E+00 | 5.99E-02 | 1.04E+00 |
| ENSG000001 | RGS12 | 4.45E-01 | 7.57E-01 | 1.04E+00 | 5.26E-02 | 1.04E+00 |
| ENSG000002 | LINC01002 | 4.45E-01 | 7.57E-01 | 9.27E-01 | -1.09E-01 | -1.08E+00 |
| ENSG000002 | --- | 4.45E-01 | 7.57E-01 | 1.04E+00 | 5.07E-02 | 1.04E+00 |
| ENSG000001 | CADM2 | 4.45E-01 | ? | 7.57E-01 | -4.03E-01 | -1.32E+00 |
| ENSG000002 | --- | 4.46E-01 | ? | 8.20E-01 | -2.86E-01 | -1.22E+00 |
| ENSG00000C | DLX3 | 4.46E-01 | 7.57E-01 | 1.09E+00 | 1.31E-01 | 1.09E+00 |
| ENSG000001 | ZNF354C | 4.46E-01 | 7.57E-01 | 9.51E-01 | -7.23E-02 | -1.05E+00 |
| ENSG000002 | --- | 4.46E-01 | ? | 1.32E+00 | 3.98E-01 | 1.32E+00 |
| ENSG000001 | CDCA7 | 4.46E-01 | 7.57E-01 | 1.06E+00 | 8.40E-02 | 1.06E+00 |
| ENSG000001 | LACC1 | 4.46E-01 | 7.57E-01 | 1.04E+00 | 6.12E-02 | 1.04E+00 |
| ENSG00000C | ERCC8 | 4.46E-01 | 7.57E-01 | 9.62E-01 | -5.52E-02 | -1.04E+00 |
| ENSG000001 | CCR10 | 4.46E-01 | ? | 1.44E+00 | 5.28E-01 | 1.44E+00 |
| ENSG000001 | SLC6A12 | 4.46E-01 | ? | 1.32E+00 | 3.98E-01 | 1.32E+00 |
| ENSG000002 | --- | 4.46E-01 | ? | 7.26E-01 | -4.61E-01 | -1.38E+00 |
| ENSG000001 | ODF2 | 4.46E-01 | 7.57E-01 | 9.64E-01 | -5.30E-02 | -1.04E+00 |

| | | | | | | |
|---|---|---|---|---|---|---|
| ENSG000002 | RPL7P21 | 4.46E-01 | ? | 8.00E-01 | -3.22E-01 | -1.25E+00 |
| ENSG000001 | CENPX | 4.46E-01 | 7.57E-01 | 1.07E+00 | 9.18E-02 | 1.07E+00 |
| ENSG000001 | KMT2D | 4.46E-01 | 7.57E-01 | 9.59E-01 | -6.11E-02 | -1.04E+00 |
| ENSG000001 | ICE1 | 4.46E-01 | 7.57E-01 | 9.69E-01 | -4.57E-02 | -1.03E+00 |
| ENSG000001 | SMIM10 | 4.46E-01 | ? | 1.14E+00 | 1.87E-01 | 1.14E+00 |
| ENSG000002 | --- | 4.46E-01 | ? | 1.25E+00 | 3.19E-01 | 1.25E+00 |
| ENSG000001 | COMMD9 | 4.46E-01 | 7.57E-01 | 1.04E+00 | 5.40E-02 | 1.04E+00 |
| ENSG000001 | PTAFR | 4.46E-01 | 7.57E-01 | 1.03E+00 | 3.74E-02 | 1.03E+00 |
| ENSG000001 | CYP4B1 | 4.46E-01 | ? | 1.24E+00 | 3.12E-01 | 1.24E+00 |
| ENSG000001 | BUB1B | 4.46E-01 | 7.58E-01 | 9.08E-01 | -1.39E-01 | -1.10E+00 |
| ENSG000002 | TSTD3 | 4.46E-01 | 7.58E-01 | 9.29E-01 | -1.07E-01 | -1.08E+00 |
| ENSG000002 | --- | 4.46E-01 | ? | 7.62E-01 | -3.93E-01 | -1.31E+00 |
| ENSG000001 | EGFL6 | 4.46E-01 | ? | 1.19E+00 | 2.48E-01 | 1.19E+00 |
| ENSG000002 | --- | 4.46E-01 | 7.58E-01 | 8.86E-01 | -1.75E-01 | -1.13E+00 |
| ENSG000001 | ZNF549 | 4.46E-01 | 7.58E-01 | 9.67E-01 | -4.78E-02 | -1.03E+00 |
| ENSG000002 | --- | 4.47E-01 | 7.58E-01 | 9.10E-01 | -1.36E-01 | -1.10E+00 |
| ENSG000002 | VIRMA-DT | 4.47E-01 | ? | 8.26E-01 | -2.77E-01 | -1.21E+00 |
| ENSG000001 | VILL | 4.47E-01 | 7.58E-01 | 1.11E+00 | 1.53E-01 | 1.11E+00 |
| ENSG000001 | AZIN1 | 4.47E-01 | 7.58E-01 | 1.02E+00 | 2.98E-02 | 1.02E+00 |
| ENSG000001 | UPK1A | 4.47E-01 | ? | 1.44E+00 | 5.30E-01 | 1.44E+00 |
| ENSG00000C | TMEM230 | 4.47E-01 | 7.58E-01 | 1.02E+00 | 2.97E-02 | 1.02E+00 |
| ENSG000002 | --- | 4.47E-01 | ? | 1.44E+00 | 5.29E-01 | 1.44E+00 |
| ENSG000001 | HDAC11 | 4.47E-01 | 7.58E-01 | 1.05E+00 | 6.36E-02 | 1.05E+00 |
| ENSG000002 | --- | 4.47E-01 | 7.58E-01 | 1.11E+00 | 1.56E-01 | 1.11E+00 |
| ENSG000001 | NBPF3 | 4.47E-01 | 7.58E-01 | 1.05E+00 | 6.97E-02 | 1.05E+00 |
| ENSG000002 | --- | 4.47E-01 | ? | 8.13E-01 | -2.98E-01 | -1.23E+00 |
| ENSG000001 | PLCG1 | 4.47E-01 | 7.58E-01 | 1.02E+00 | 3.25E-02 | 1.02E+00 |
| ENSG000001 | DENND10 | 4.47E-01 | 7.58E-01 | 1.03E+00 | 3.73E-02 | 1.03E+00 |
| ENSG000002 | PKD1P1 | 4.47E-01 | 7.58E-01 | 8.74E-01 | -1.94E-01 | -1.14E+00 |
| ENSG000001 | C9orf78 | 4.47E-01 | 7.58E-01 | 9.76E-01 | -3.46E-02 | -1.02E+00 |
| ENSG000001 | LYPLA1 | 4.47E-01 | 7.58E-01 | 9.67E-01 | -4.86E-02 | -1.03E+00 |
| ENSG000001 | SFI1 | 4.47E-01 | 7.58E-01 | 9.54E-01 | -6.79E-02 | -1.05E+00 |
| ENSG000001 | C2CD3 | 4.47E-01 | 7.58E-01 | 1.03E+00 | 3.99E-02 | 1.03E+00 |
| ENSG000001 | TOMM22 | 4.48E-01 | 7.58E-01 | 1.02E+00 | 3.28E-02 | 1.02E+00 |
| ENSG00000C | RFFL | 4.48E-01 | 7.58E-01 | 1.03E+00 | 4.00E-02 | 1.03E+00 |
| ENSG000001 | SH2B3 | 4.48E-01 | 7.58E-01 | 9.73E-01 | -3.90E-02 | -1.03E+00 |
| ENSG00000C | NUP50 | 4.48E-01 | 7.58E-01 | 1.02E+00 | 3.32E-02 | 1.02E+00 |
| ENSG000001 | HSD17B1 | 4.48E-01 | 7.58E-01 | 1.08E+00 | 1.14E-01 | 1.08E+00 |
| ENSG000001 | BMPR1B | 4.48E-01 | 7.58E-01 | 9.61E-01 | -5.69E-02 | -1.04E+00 |
| ENSG000001 | GRK4 | 4.48E-01 | 7.58E-01 | 9.40E-01 | -8.94E-02 | -1.06E+00 |
| ENSG000001 | IK | 4.48E-01 | 7.58E-01 | 9.62E-01 | -5.57E-02 | -1.04E+00 |
| ENSG000002 | OR9A4 | 4.48E-01 | ? | 1.54E+00 | 6.22E-01 | 1.54E+00 |
| ENSG000001 | BTBD10 | 4.48E-01 | 7.58E-01 | 1.02E+00 | 3.17E-02 | 1.02E+00 |
| ENSG00000C | RFX3 | 4.48E-01 | 7.58E-01 | 1.05E+00 | 6.64E-02 | 1.05E+00 |
| ENSG000001 | FES | 4.48E-01 | ? | 7.43E-01 | -4.29E-01 | -1.35E+00 |

| | | | | | | |
|---|---|---|---|---|---|---|
| ENSG000001 | CNIH2 | 4.48E-01 | ? | 1.29E+00 | 3.66E-01 | 1.29E+00 |
| ENSG000001 | SPNS2 | 4.48E-01 | 7.58E-01 | 8.50E-01 | -2.34E-01 | -1.18E+00 |
| ENSG000000 | MTMR2 | 4.48E-01 | 7.58E-01 | 9.77E-01 | -3.40E-02 | -1.02E+00 |
| ENSG000000 | POLR1A | 4.48E-01 | 7.58E-01 | 1.03E+00 | 4.07E-02 | 1.03E+00 |
| ENSG000001 | TAF1 | 4.48E-01 | 7.58E-01 | 9.68E-01 | -4.65E-02 | -1.03E+00 |
| ENSG000001 | FCGR1A | 4.48E-01 | ? | 7.13E-01 | -4.88E-01 | -1.40E+00 |
| ENSG000002 | DGCR11 | 4.48E-01 | 7.58E-01 | 1.07E+00 | 9.18E-02 | 1.07E+00 |
| ENSG000001 | CCDC17 | 4.48E-01 | 7.58E-01 | 8.83E-01 | -1.80E-01 | -1.13E+00 |
| ENSG000001 | PGLS | 4.48E-01 | 7.58E-01 | 1.04E+00 | 6.15E-02 | 1.04E+00 |
| ENSG000001 | MRNIP | 4.49E-01 | 7.58E-01 | 9.19E-01 | -1.21E-01 | -1.09E+00 |
| ENSG000002 | --- | 4.49E-01 | ? | 1.32E+00 | 4.03E-01 | 1.32E+00 |
| ENSG000002 | --- | 4.49E-01 | ? | 7.17E-01 | -4.80E-01 | -1.39E+00 |
| ENSG000002 | SNRPEP4 | 4.49E-01 | ? | 8.42E-01 | -2.47E-01 | -1.19E+00 |
| ENSG000002 | --- | 4.49E-01 | ? | 5.85E-01 | -7.74E-01 | -1.71E+00 |
| ENSG000001 | GABRB3 | 4.49E-01 | 7.58E-01 | 1.06E+00 | 8.13E-02 | 1.06E+00 |
| ENSG000001 | GCC1 | 4.49E-01 | 7.58E-01 | 9.69E-01 | -4.59E-02 | -1.03E+00 |
| ENSG000001 | PIEZO1 | 4.49E-01 | 7.58E-01 | 1.04E+00 | 5.43E-02 | 1.04E+00 |
| ENSG000002 | --- | 4.49E-01 | 7.58E-01 | 1.13E+00 | 1.77E-01 | 1.13E+00 |
| ENSG000001 | PFKFB1 | 4.49E-01 | ? | 7.95E-01 | -3.30E-01 | -1.26E+00 |
| ENSG000001 | JAK3 | 4.49E-01 | 7.59E-01 | 1.07E+00 | 1.03E-01 | 1.07E+00 |
| ENSG000002 | LINC01725 | 4.49E-01 | ? | 1.50E+00 | 5.83E-01 | 1.50E+00 |
| ENSG000002 | --- | 4.49E-01 | ? | 1.42E+00 | 5.08E-01 | 1.42E+00 |
| ENSG000001 | NAB1 | 4.49E-01 | 7.59E-01 | 9.65E-01 | -5.08E-02 | -1.04E+00 |
| ENSG000002 | DNASE1 | 4.49E-01 | 7.59E-01 | 1.05E+00 | 6.54E-02 | 1.05E+00 |
| ENSG000002 | DPY19L2P3 | 4.49E-01 | ? | 7.02E-01 | -5.11E-01 | -1.43E+00 |
| ENSG000002 | --- | 4.49E-01 | ? | 1.16E+00 | 2.14E-01 | 1.16E+00 |
| ENSG000002 | --- | 4.49E-01 | ? | 6.37E-01 | -6.50E-01 | -1.57E+00 |
| ENSG000002 | KLF3-AS1 | 4.49E-01 | ? | 7.16E-01 | -4.82E-01 | -1.40E+00 |
| ENSG000001 | TMEM154 | 4.49E-01 | 7.59E-01 | 9.67E-01 | -4.82E-02 | -1.03E+00 |
| ENSG000002 | FCGR1BP | 4.50E-01 | ? | 8.22E-01 | -2.82E-01 | -1.22E+00 |
| ENSG000001 | LCTL | 4.50E-01 | ? | 8.18E-01 | -2.90E-01 | -1.22E+00 |
| ENSG000002 | EEF1AKMT2 | 4.50E-01 | 7.59E-01 | 1.04E+00 | 5.68E-02 | 1.04E+00 |
| ENSG000002 | CSNK1E | 4.50E-01 | 7.59E-01 | 1.04E+00 | 5.71E-02 | 1.04E+00 |
| ENSG000002 | --- | 4.50E-01 | ? | 8.51E-01 | -2.32E-01 | -1.17E+00 |
| ENSG000002 | --- | 4.50E-01 | ? | 7.89E-01 | -3.41E-01 | -1.27E+00 |
| ENSG000002 | SCGB2B2 | 4.50E-01 | ? | 8.49E-01 | -2.36E-01 | -1.18E+00 |
| ENSG000002 | --- | 4.50E-01 | ? | 1.31E+00 | 3.87E-01 | 1.31E+00 |
| ENSG000002 | PTOV1-AS1 | 4.50E-01 | ? | 7.43E-01 | -4.29E-01 | -1.35E+00 |
| ENSG000002 | --- | 4.50E-01 | ? | 1.32E+00 | 3.99E-01 | 1.32E+00 |
| ENSG000002 | --- | 4.50E-01 | 7.60E-01 | 1.15E+00 | 1.96E-01 | 1.15E+00 |
| ENSG000002 | --- | 4.50E-01 | ? | 8.00E-01 | -3.22E-01 | -1.25E+00 |
| ENSG000001 | STRIP1 | 4.50E-01 | 7.60E-01 | 9.69E-01 | -4.51E-02 | -1.03E+00 |
| ENSG000001 | LINC01560 | 4.50E-01 | ? | 1.16E+00 | 2.17E-01 | 1.16E+00 |
| ENSG000001 | LRRC1 | 4.50E-01 | 7.60E-01 | 1.03E+00 | 4.85E-02 | 1.03E+00 |
| ENSG000001 | SLC22A3 | 4.50E-01 | 7.60E-01 | 9.55E-01 | -6.71E-02 | -1.05E+00 |

| | | | | | | |
|---|---|---|---|---|---|---|
| ENSG000002 | --- | 4.50E-01 | ? | 7.38E-01 | -4.38E-01 | -1.35E+00 |
| ENSG000001 | SAP30BP | 4.50E-01 | 7.60E-01 | 9.80E-01 | -2.89E-02 | -1.02E+00 |
| ENSG000002 | LINC01772 | 4.50E-01 | 7.60E-01 | 9.29E-01 | -1.06E-01 | -1.08E+00 |
| ENSG000001 | DDRGK1 | 4.50E-01 | 7.60E-01 | 9.73E-01 | -3.97E-02 | -1.03E+00 |
| ENSG000002 | POLR2J4 | 4.50E-01 | 7.60E-01 | 9.02E-01 | -1.49E-01 | -1.11E+00 |
| ENSG000001 | CRACDL | 4.50E-01 | 7.60E-01 | 1.08E+00 | 1.17E-01 | 1.08E+00 |
| ENSG000001 | ZNF527 | 4.51E-01 | 7.60E-01 | 9.49E-01 | -7.60E-02 | -1.05E+00 |
| ENSG000002 | DACT3-AS1 | 4.51E-01 | ? | 1.18E+00 | 2.37E-01 | 1.18E+00 |
| ENSG000001 | RHOT2 | 4.51E-01 | 7.60E-01 | 1.05E+00 | 7.60E-02 | 1.05E+00 |
| ENSG000001 | PHC1 | 4.51E-01 | 7.60E-01 | 1.04E+00 | 5.86E-02 | 1.04E+00 |
| ENSG000001 | CAPNS1 | 4.51E-01 | 7.60E-01 | 1.02E+00 | 2.55E-02 | 1.02E+00 |
| ENSG000002 | STX17-DT | 4.51E-01 | ? | 7.75E-01 | -3.68E-01 | -1.29E+00 |
| ENSG000001 | CDKN2C | 4.51E-01 | ? | 8.53E-01 | -2.29E-01 | -1.17E+00 |
| ENSG000002 | --- | 4.51E-01 | ? | 8.57E-01 | -2.23E-01 | -1.17E+00 |
| ENSG000002 | --- | 4.51E-01 | ? | 1.21E+00 | 2.75E-01 | 1.21E+00 |
| ENSG00000C | SLC45A4 | 4.51E-01 | 7.60E-01 | 9.51E-01 | -7.22E-02 | -1.05E+00 |
| ENSG000002 | SMIM2-AS1 | 4.51E-01 | 7.60E-01 | 1.12E+00 | 1.57E-01 | 1.12E+00 |
| ENSG000001 | HTR7P1 | 4.51E-01 | 7.60E-01 | 1.07E+00 | 1.01E-01 | 1.07E+00 |
| ENSG000001 | SYDE1 | 4.51E-01 | 7.60E-01 | 1.06E+00 | 8.67E-02 | 1.06E+00 |
| ENSG000002 | --- | 4.51E-01 | 7.60E-01 | 9.22E-01 | -1.17E-01 | -1.08E+00 |
| ENSG000002 | --- | 4.51E-01 | ? | 1.37E+00 | 4.59E-01 | 1.37E+00 |
| ENSG00000C | HHAT | 4.52E-01 | 7.61E-01 | 1.06E+00 | 7.77E-02 | 1.06E+00 |
| ENSG000001 | PDGFA | 4.52E-01 | 7.61E-01 | 9.78E-01 | -3.21E-02 | -1.02E+00 |
| ENSG000001 | SIPA1L1 | 4.52E-01 | 7.61E-01 | 9.40E-01 | -8.90E-02 | -1.06E+00 |
| ENSG000002 | CPNE1 | 4.52E-01 | 7.61E-01 | 1.04E+00 | 6.18E-02 | 1.04E+00 |
| ENSG000002 | --- | 4.52E-01 | ? | 1.51E+00 | 5.92E-01 | 1.51E+00 |
| ENSG000001 | CPLX1 | 4.52E-01 | ? | 1.26E+00 | 3.39E-01 | 1.26E+00 |
| ENSG000002 | LINC02972 | 4.52E-01 | ? | 1.24E+00 | 3.13E-01 | 1.24E+00 |
| ENSG000002 | ZNF204P | 4.52E-01 | ? | 7.04E-01 | -5.06E-01 | -1.42E+00 |
| ENSG000001 | TC2N | 4.52E-01 | 7.61E-01 | 9.66E-01 | -5.01E-02 | -1.04E+00 |
| ENSG000002 | CCNYL6 | 4.52E-01 | ? | 1.41E+00 | 4.92E-01 | 1.41E+00 |
| ENSG000001 | ZNF93 | 4.52E-01 | 7.61E-01 | 9.46E-01 | -8.04E-02 | -1.06E+00 |
| ENSG000001 | MEMO1 | 4.52E-01 | 7.61E-01 | 1.02E+00 | 3.33E-02 | 1.02E+00 |
| ENSG000001 | SLC9A5 | 4.52E-01 | 7.61E-01 | 1.09E+00 | 1.24E-01 | 1.09E+00 |
| ENSG000001 | NRIP3 | 4.52E-01 | 7.61E-01 | 9.40E-01 | -8.85E-02 | -1.06E+00 |
| ENSG000001 | DDN | 4.52E-01 | ? | 1.22E+00 | 2.82E-01 | 1.22E+00 |
| ENSG000001 | TFIP11 | 4.52E-01 | 7.61E-01 | 1.02E+00 | 2.74E-02 | 1.02E+00 |
| ENSG000002 | ZNF579 | 4.52E-01 | 7.61E-01 | 1.08E+00 | 1.14E-01 | 1.08E+00 |
| ENSG000001 | ZNF445 | 4.52E-01 | 7.61E-01 | 1.03E+00 | 4.41E-02 | 1.03E+00 |
| ENSG00000C | KRIT1 | 4.52E-01 | 7.61E-01 | 9.70E-01 | -4.45E-02 | -1.03E+00 |
| ENSG00000C | KCNK6 | 4.52E-01 | 7.61E-01 | 1.02E+00 | 3.05E-02 | 1.02E+00 |
| ENSG00000C | ANK1 | 4.52E-01 | 7.61E-01 | 1.12E+00 | 1.66E-01 | 1.12E+00 |
| ENSG000001 | MAL2 | 4.52E-01 | 7.61E-01 | 1.05E+00 | 7.00E-02 | 1.05E+00 |
| ENSG000002 | --- | 4.52E-01 | ? | 8.12E-01 | -3.01E-01 | -1.23E+00 |
| ENSG000001 | ALX1 | 4.52E-01 | ? | 7.19E-01 | -4.76E-01 | -1.39E+00 |

| | | | | | | | |
|---|---|---|---|---|---|---|---|
| ENSG000002 | --- | 4.52E-01 | | 7.61E-01 | 1.10E+00 | 1.40E-01 | 1.10E+00 |
| ENSG000002 | --- | 4.52E-01 | | 7.61E-01 | 1.09E+00 | 1.27E-01 | 1.09E+00 |
| ENSG000001 | ALKBH1 | 4.52E-01 | | 7.61E-01 | 9.67E-01 | -4.80E-02 | -1.03E+00 |
| ENSG000002 | AP1G2 | 4.53E-01 | | 7.61E-01 | 9.62E-01 | -5.55E-02 | -1.04E+00 |
| ENSG000001 | UBTD1 | 4.53E-01 | | 7.61E-01 | 1.06E+00 | 8.96E-02 | 1.06E+00 |
| ENSG000002 | --- | 4.53E-01 | | 7.61E-01 | 9.01E-01 | -1.50E-01 | -1.11E+00 |
| ENSG000001 | NPSR1 | 4.53E-01 | ? | | 1.41E+00 | 4.94E-01 | 1.41E+00 |
| ENSG000001 | CTSE | 4.53E-01 | ? | | 8.25E-01 | -2.77E-01 | -1.21E+00 |
| ENSG000001 | SRP9 | 4.53E-01 | | 7.61E-01 | 1.02E+00 | 2.62E-02 | 1.02E+00 |
| ENSG000002 | PMS2P4 | 4.53E-01 | | 7.61E-01 | 1.08E+00 | 1.13E-01 | 1.08E+00 |
| ENSG00000C | MARK2 | 4.53E-01 | | 7.61E-01 | 1.02E+00 | 3.36E-02 | 1.02E+00 |
| ENSG00000C | MAP2 | 4.53E-01 | | 7.61E-01 | 9.54E-01 | -6.79E-02 | -1.05E+00 |
| ENSG00000C | PCNP | 4.53E-01 | | 7.61E-01 | 9.73E-01 | -3.95E-02 | -1.03E+00 |
| ENSG000002 | TSPAN4 | 4.53E-01 | | 7.61E-01 | 1.07E+00 | 1.02E-01 | 1.07E+00 |
| ENSG000002 | EPPK1 | 4.54E-01 | | 7.62E-01 | 9.45E-01 | -8.18E-02 | -1.06E+00 |
| ENSG000001 | FAM110A | 4.54E-01 | | 7.62E-01 | 1.07E+00 | 1.01E-01 | 1.07E+00 |
| ENSG000001 | ZBTB39 | 4.54E-01 | | 7.62E-01 | 1.03E+00 | 4.03E-02 | 1.03E+00 |
| ENSG000001 | EIF4A1 | 4.54E-01 | | 7.62E-01 | 1.02E+00 | 3.01E-02 | 1.02E+00 |
| ENSG000001 | C19orf25 | 4.54E-01 | | 7.62E-01 | 1.06E+00 | 7.78E-02 | 1.06E+00 |
| ENSG000002 | COLQ | 4.54E-01 | | 7.62E-01 | 8.71E-01 | -2.00E-01 | -1.15E+00 |
| ENSG000002 | --- | 4.54E-01 | ? | | 1.53E+00 | 6.10E-01 | 1.53E+00 |
| ENSG000002 | --- | 4.54E-01 | ? | | 1.22E+00 | 2.93E-01 | 1.22E+00 |
| ENSG000002 | JRK | 4.54E-01 | | 7.62E-01 | 1.04E+00 | 5.76E-02 | 1.04E+00 |
| ENSG00000C | SNRPA | 4.54E-01 | | 7.62E-01 | 1.03E+00 | 3.87E-02 | 1.03E+00 |
| ENSG000001 | HAUS8 | 4.54E-01 | | 7.62E-01 | 9.31E-01 | -1.03E-01 | -1.07E+00 |
| ENSG000001 | DBP | 4.54E-01 | ? | | 1.17E+00 | 2.26E-01 | 1.17E+00 |
| ENSG000002 | API5P1 | 4.54E-01 | ? | | 8.26E-01 | -2.75E-01 | -1.21E+00 |
| ENSG000002 | CHMP4A | 4.54E-01 | | 7.62E-01 | 1.04E+00 | 5.22E-02 | 1.04E+00 |
| ENSG000001 | HSPE1 | 4.54E-01 | | 7.62E-01 | 1.04E+00 | 5.04E-02 | 1.04E+00 |
| ENSG000001 | SPATA18 | 4.54E-01 | | 7.62E-01 | 1.03E+00 | 4.82E-02 | 1.03E+00 |
| ENSG00000C | BRD9 | 4.54E-01 | | 7.62E-01 | 9.79E-01 | -3.09E-02 | -1.02E+00 |
| ENSG000002 | LINC02605 | 4.54E-01 | ? | | 6.91E-01 | -5.34E-01 | -1.45E+00 |
| ENSG00000C | BTBD1 | 4.55E-01 | | 7.62E-01 | 1.02E+00 | 3.39E-02 | 1.02E+00 |
| ENSG00000C | RASGRF1 | 4.55E-01 | ? | | 1.16E+00 | 2.19E-01 | 1.16E+00 |
| ENSG000002 | --- | 4.55E-01 | ? | | 1.33E+00 | 4.11E-01 | 1.33E+00 |
| ENSG000001 | YY1 | 4.55E-01 | | 7.62E-01 | 9.80E-01 | -2.90E-02 | -1.02E+00 |
| ENSG000002 | --- | 4.55E-01 | ? | | 7.77E-01 | -3.65E-01 | -1.29E+00 |
| ENSG000002 | --- | 4.55E-01 | ? | | 8.79E-01 | -1.87E-01 | -1.14E+00 |
| ENSG00000C | DUSP12 | 4.55E-01 | | 7.62E-01 | 9.62E-01 | -5.52E-02 | -1.04E+00 |
| ENSG000002 | RUSC1-AS1 | 4.55E-01 | | 7.62E-01 | 8.96E-01 | -1.58E-01 | -1.12E+00 |
| ENSG000001 | PLCH1 | 4.55E-01 | ? | | 8.02E-01 | -3.19E-01 | -1.25E+00 |
| ENSG000001 | DOCK7 | 4.55E-01 | | 7.62E-01 | 9.81E-01 | -2.72E-02 | -1.02E+00 |
| ENSG000002 | --- | 4.55E-01 | ? | | 8.38E-01 | -2.55E-01 | -1.19E+00 |
| ENSG000001 | DCTD | 4.55E-01 | | 7.62E-01 | 1.03E+00 | 3.71E-02 | 1.03E+00 |
| ENSG000001 | CD72 | 4.55E-01 | | 7.62E-01 | 1.15E+00 | 1.99E-01 | 1.15E+00 |

| | | | | | | |
|---|---|---|---|---|---|---|
| ENSG000001 | RABAC1 | 4.55E-01 | 7.62E-01 | 1.11E+00 | 1.47E-01 | 1.11E+00 |
| ENSG000001 | B9D2 | 4.55E-01 | 7.62E-01 | 1.08E+00 | 1.11E-01 | 1.08E+00 |
| ENSG000001 | TAF4B | 4.55E-01 | 7.62E-01 | 1.04E+00 | 5.58E-02 | 1.04E+00 |
| ENSG000001 | MAN2A2 | 4.55E-01 | 7.62E-01 | 9.71E-01 | -4.31E-02 | -1.03E+00 |
| ENSG000001 | FBXL2 | 4.55E-01 | 7.62E-01 | 1.03E+00 | 4.76E-02 | 1.03E+00 |
| ENSG000002 | ZNF674-AS1 | 4.56E-01 | 7.62E-01 | 8.95E-01 | -1.60E-01 | -1.12E+00 |
| ENSG000001 | MLX | 4.56E-01 | 7.62E-01 | 9.74E-01 | -3.76E-02 | -1.03E+00 |
| ENSG000001 | PKMYT1 | 4.56E-01 | 7.62E-01 | 1.11E+00 | 1.48E-01 | 1.11E+00 |
| ENSG000002 | --- | 4.56E-01 | ? | 7.41E-01 | -4.33E-01 | -1.35E+00 |
| ENSG000002 | FCGBP | 4.56E-01 | 7.62E-01 | 1.10E+00 | 1.35E-01 | 1.10E+00 |
| ENSG000001 | FBXL9P | 4.56E-01 | ? | 1.16E+00 | 2.10E-01 | 1.16E+00 |
| ENSG000001 | SKIL | 4.56E-01 | 7.62E-01 | 9.56E-01 | -6.55E-02 | -1.05E+00 |
| ENSG000001 | URGCP | 4.56E-01 | 7.62E-01 | 1.02E+00 | 2.73E-02 | 1.02E+00 |
| ENSG000002 | LINC01354 | 4.56E-01 | 7.62E-01 | 1.11E+00 | 1.47E-01 | 1.11E+00 |
| ENSG00000C | TMEM131 | 4.56E-01 | 7.62E-01 | 1.03E+00 | 3.64E-02 | 1.03E+00 |
| ENSG000001 | GORAB | 4.56E-01 | 7.62E-01 | 9.63E-01 | -5.37E-02 | -1.04E+00 |
| ENSG000001 | CTSC | 4.56E-01 | 7.62E-01 | 9.72E-01 | -4.07E-02 | -1.03E+00 |
| ENSG000002 | PCDHGA4 | 4.56E-01 | 7.62E-01 | 1.10E+00 | 1.35E-01 | 1.10E+00 |
| ENSG000002 | --- | 4.56E-01 | ? | 1.57E+00 | 6.47E-01 | 1.57E+00 |
| ENSG000002 | --- | 4.56E-01 | ? | 7.60E-01 | -3.96E-01 | -1.32E+00 |
| ENSG000001 | MEA1 | 4.56E-01 | 7.63E-01 | 1.02E+00 | 3.41E-02 | 1.02E+00 |
| ENSG000001 | INPP5E | 4.56E-01 | 7.63E-01 | 9.60E-01 | -5.84E-02 | -1.04E+00 |
| ENSG000002 | HMGA1P4 | 4.56E-01 | ? | 1.35E+00 | 4.37E-01 | 1.35E+00 |
| ENSG000001 | KANSL3 | 4.56E-01 | 7.63E-01 | 1.02E+00 | 3.40E-02 | 1.02E+00 |
| ENSG000001 | NEU3 | 4.56E-01 | 7.63E-01 | 1.03E+00 | 4.77E-02 | 1.03E+00 |
| ENSG000002 | LINC01134 | 4.56E-01 | ? | 8.62E-01 | -2.14E-01 | -1.16E+00 |
| ENSG000001 | THAP5 | 4.56E-01 | 7.63E-01 | 9.64E-01 | -5.31E-02 | -1.04E+00 |
| ENSG00000C | BCLAF1 | 4.56E-01 | 7.63E-01 | 9.70E-01 | -4.42E-02 | -1.03E+00 |
| ENSG000001 | ADAM9 | 4.56E-01 | 7.63E-01 | 1.02E+00 | 3.33E-02 | 1.02E+00 |
| ENSG000001 | PIM2 | 4.57E-01 | 7.63E-01 | 1.05E+00 | 6.57E-02 | 1.05E+00 |
| ENSG000002 | --- | 4.57E-01 | 7.63E-01 | 9.48E-01 | -7.75E-02 | -1.06E+00 |
| ENSG000002 | EMSLR | 4.57E-01 | 7.63E-01 | 9.41E-01 | -8.80E-02 | -1.06E+00 |
| ENSG000001 | AFDN-DT | 4.57E-01 | ? | 8.68E-01 | -2.04E-01 | -1.15E+00 |
| ENSG000001 | KCND1 | 4.57E-01 | 7.63E-01 | 9.17E-01 | -1.25E-01 | -1.09E+00 |
| ENSG000002 | LINC01215 | 4.57E-01 | ? | 6.45E-01 | -6.33E-01 | -1.55E+00 |
| ENSG000001 | DOCK5 | 4.57E-01 | 7.63E-01 | 9.65E-01 | -5.14E-02 | -1.04E+00 |
| ENSG000002 | LINC01882 | 4.57E-01 | ? | 7.29E-01 | -4.55E-01 | -1.37E+00 |
| ENSG000002 | ATE1OSP | 4.57E-01 | ? | 8.10E-01 | -3.04E-01 | -1.23E+00 |
| ENSG000002 | --- | 4.57E-01 | 7.63E-01 | 9.03E-01 | -1.47E-01 | -1.11E+00 |
| ENSG000001 | KIAA1586 | 4.57E-01 | 7.63E-01 | 9.44E-01 | -8.36E-02 | -1.06E+00 |
| ENSG000001 | TCIRG1 | 4.57E-01 | 7.63E-01 | 1.06E+00 | 8.76E-02 | 1.06E+00 |
| ENSG000002 | --- | 4.57E-01 | ? | 8.39E-01 | -2.53E-01 | -1.19E+00 |
| ENSG000001 | CAPS | 4.58E-01 | 7.64E-01 | 1.07E+00 | 9.97E-02 | 1.07E+00 |
| ENSG000001 | CCDC22 | 4.58E-01 | 7.64E-01 | 1.06E+00 | 8.35E-02 | 1.06E+00 |
| ENSG000002 | TFAP2A-AS2 | 4.58E-01 | 7.64E-01 | 1.09E+00 | 1.19E-01 | 1.09E+00 |

| | | | | | | |
|---|---|---|---|---|---|---|
| ENSG000001 | SLC24A3 | 4.58E-01 | ? | 8.14E-01 | -2.97E-01 | -1.23E+00 |
| ENSG000001 | FBXL20 | 4.58E-01 | 7.64E-01 | 1.04E+00 | 5.94E-02 | 1.04E+00 |
| ENSG000002 | --- | 4.58E-01 | 7.64E-01 | 9.45E-01 | -8.16E-02 | -1.06E+00 |
| ENSG000001 | RAB30 | 4.58E-01 | 7.64E-01 | 1.05E+00 | 7.57E-02 | 1.05E+00 |
| ENSG000001 | GET1 | 4.58E-01 | 7.64E-01 | 1.03E+00 | 3.65E-02 | 1.03E+00 |
| ENSG000001 | FAM89A | 4.58E-01 | 7.64E-01 | 1.07E+00 | 9.82E-02 | 1.07E+00 |
| ENSG00000C | PUM2 | 4.58E-01 | 7.64E-01 | 1.02E+00 | 2.42E-02 | 1.02E+00 |
| ENSG000001 | KIAA1217 | 4.58E-01 | 7.64E-01 | 9.67E-01 | -4.84E-02 | -1.03E+00 |
| ENSG00000C | PSMB1 | 4.58E-01 | 7.64E-01 | 1.02E+00 | 2.45E-02 | 1.02E+00 |
| ENSG000001 | ZNF548 | 4.58E-01 | 7.64E-01 | 9.56E-01 | -6.47E-02 | -1.05E+00 |
| ENSG000002 | --- | 4.58E-01 | ? | 8.27E-01 | -2.73E-01 | -1.21E+00 |
| ENSG000001 | PLCXD1 | 4.58E-01 | 7.64E-01 | 9.57E-01 | -6.29E-02 | -1.04E+00 |
| ENSG00000C | E2F2 | 4.58E-01 | ? | 6.54E-01 | -6.14E-01 | -1.53E+00 |
| ENSG000001 | ITPR2 | 4.58E-01 | 7.64E-01 | 9.57E-01 | -6.40E-02 | -1.05E+00 |
| ENSG000002 | --- | 4.58E-01 | ? | 1.22E+00 | 2.82E-01 | 1.22E+00 |
| ENSG000001 | ANKRD36C | 4.58E-01 | 7.64E-01 | 9.49E-01 | -7.60E-02 | -1.05E+00 |
| ENSG000001 | HECTD3 | 4.58E-01 | 7.64E-01 | 1.03E+00 | 3.61E-02 | 1.03E+00 |
| ENSG000002 | LMO7-AS1 | 4.58E-01 | ? | 1.23E+00 | 2.95E-01 | 1.23E+00 |
| ENSG000001 | NASP | 4.58E-01 | 7.64E-01 | 9.59E-01 | -6.03E-02 | -1.04E+00 |
| ENSG000002 | --- | 4.59E-01 | 7.64E-01 | 1.08E+00 | 1.16E-01 | 1.08E+00 |
| ENSG00000C | HIVEP2 | 4.59E-01 | 7.64E-01 | 1.03E+00 | 4.89E-02 | 1.03E+00 |
| ENSG000001 | DENND3 | 4.59E-01 | 7.64E-01 | 9.64E-01 | -5.24E-02 | -1.04E+00 |
| ENSG000001 | METAP2 | 4.59E-01 | 7.64E-01 | 9.73E-01 | -4.00E-02 | -1.03E+00 |
| ENSG000002 | LINC01698 | 4.59E-01 | ? | 1.25E+00 | 3.24E-01 | 1.25E+00 |
| ENSG000001 | NT5C1A | 4.59E-01 | ? | 1.35E+00 | 4.34E-01 | 1.35E+00 |
| ENSG000002 | --- | 4.59E-01 | ? | 1.19E+00 | 2.53E-01 | 1.19E+00 |
| ENSG00000C | FBXL19 | 4.59E-01 | 7.65E-01 | 1.05E+00 | 7.33E-02 | 1.05E+00 |
| ENSG000002 | --- | 4.59E-01 | ? | 1.14E+00 | 1.89E-01 | 1.14E+00 |
| ENSG000002 | --- | 4.59E-01 | ? | 7.38E-01 | -4.39E-01 | -1.36E+00 |
| ENSG000001 | --- | 4.59E-01 | ? | 1.20E+00 | 2.64E-01 | 1.20E+00 |
| ENSG000002 | ATP5MF-PTC | 4.59E-01 | 7.65E-01 | 1.05E+00 | 7.15E-02 | 1.05E+00 |
| ENSG000001 | SNHG28 | 4.59E-01 | ? | 8.21E-01 | -2.84E-01 | -1.22E+00 |
| ENSG000001 | --- | 4.59E-01 | 7.65E-01 | 9.44E-01 | -8.34E-02 | -1.06E+00 |
| ENSG000001 | STK33 | 4.60E-01 | 7.65E-01 | 1.11E+00 | 1.51E-01 | 1.11E+00 |
| ENSG000001 | TRIM47 | 4.60E-01 | 7.65E-01 | 1.05E+00 | 6.91E-02 | 1.05E+00 |
| ENSG000002 | BAHCC1 | 4.60E-01 | 7.65E-01 | 9.47E-01 | -7.84E-02 | -1.06E+00 |
| ENSG000001 | GTF2A2 | 4.60E-01 | 7.65E-01 | 9.73E-01 | -4.02E-02 | -1.03E+00 |
| ENSG000002 | C2orf92 | 4.60E-01 | 7.65E-01 | 1.09E+00 | 1.25E-01 | 1.09E+00 |
| ENSG000002 | RPS4XP11 | 4.60E-01 | ? | 7.39E-01 | -4.37E-01 | -1.35E+00 |
| ENSG000002 | MIR1915HG | 4.60E-01 | 7.65E-01 | 1.07E+00 | 9.72E-02 | 1.07E+00 |
| ENSG000001 | ASCL2 | 4.60E-01 | ? | 1.46E+00 | 5.47E-01 | 1.46E+00 |
| ENSG000001 | CISH | 4.60E-01 | 7.65E-01 | 9.26E-01 | -1.11E-01 | -1.08E+00 |
| ENSG000002 | ZNF432 | 4.60E-01 | 7.65E-01 | 9.43E-01 | -8.51E-02 | -1.06E+00 |
| ENSG000001 | ZNF461 | 4.60E-01 | 7.65E-01 | 1.04E+00 | 6.28E-02 | 1.04E+00 |
| ENSG000002 | B3GALT4 | 4.60E-01 | 7.65E-01 | 1.13E+00 | 1.72E-01 | 1.13E+00 |

| | | | | | | |
|---|---|---|---|---|---|---|
| ENSG000002 | ARHGAP27P1 | 4.60E-01 | 7.65E-01 | 9.40E-01 | -8.89E-02 | -1.06E+00 |
| ENSG000002 | TMEM263-DT | 4.60E-01 | ? | 1.20E+00 | 2.68E-01 | 1.20E+00 |
| ENSG000001 | HEPACAM2 | 4.60E-01 | ? | 1.47E+00 | 5.55E-01 | 1.47E+00 |
| ENSG000001 | DELEC1 | 4.60E-01 | ? | 8.54E-01 | -2.28E-01 | -1.17E+00 |
| ENSG000001 | OGDH | 4.60E-01 | 7.65E-01 | 1.02E+00 | 2.61E-02 | 1.02E+00 |
| ENSG000002 | ANKRD11P2 | 4.60E-01 | ? | 1.35E+00 | 4.36E-01 | 1.35E+00 |
| ENSG000002 | ZNF492 | 4.61E-01 | ? | 1.14E+00 | 1.94E-01 | 1.14E+00 |
| ENSG000001 | CBR1 | 4.61E-01 | 7.66E-01 | 1.03E+00 | 3.96E-02 | 1.03E+00 |
| ENSG000002 | --- | 4.61E-01 | 7.66E-01 | 1.35E+00 | 4.31E-01 | 1.35E+00 |
| ENSG000002 | --- | 4.61E-01 | ? | 1.36E+00 | 4.42E-01 | 1.36E+00 |
| ENSG000002 | ZNF888 | 4.61E-01 | 7.66E-01 | 9.64E-01 | -5.27E-02 | -1.04E+00 |
| ENSG000001 | STK11IP | 4.61E-01 | 7.66E-01 | 1.05E+00 | 7.51E-02 | 1.05E+00 |
| ENSG000002 | --- | 4.61E-01 | ? | 1.38E+00 | 4.68E-01 | 1.38E+00 |
| ENSG000002 | SAMD12-AS1 | 4.61E-01 | ? | 1.32E+00 | 4.01E-01 | 1.32E+00 |
| ENSG000001 | FBXL22 | 4.61E-01 | ? | 1.41E+00 | 5.01E-01 | 1.41E+00 |
| ENSG000002 | --- | 4.61E-01 | ? | 7.03E-01 | -5.09E-01 | -1.42E+00 |
| ENSG000002 | --- | 4.61E-01 | 7.66E-01 | 9.41E-01 | -8.84E-02 | -1.06E+00 |
| ENSG000001 | EFCAB5 | 4.61E-01 | ? | 1.13E+00 | 1.82E-01 | 1.13E+00 |
| ENSG000001 | NOSIP | 4.62E-01 | 7.67E-01 | 1.05E+00 | 6.41E-02 | 1.05E+00 |
| ENSG000001 | USP46 | 4.62E-01 | 7.67E-01 | 1.02E+00 | 3.25E-02 | 1.02E+00 |
| ENSG000001 | KRAS | 4.62E-01 | 7.67E-01 | 1.02E+00 | 3.53E-02 | 1.02E+00 |
| ENSG000001 | RSRP1 | 4.62E-01 | 7.67E-01 | 9.48E-01 | -7.64E-02 | -1.05E+00 |
| ENSG000000 | TGDS | 4.62E-01 | 7.67E-01 | 1.05E+00 | 6.72E-02 | 1.05E+00 |
| ENSG000001 | PSMD10 | 4.62E-01 | 7.67E-01 | 9.80E-01 | -2.98E-02 | -1.02E+00 |
| ENSG000001 | NECTIN1 | 4.62E-01 | 7.67E-01 | 1.04E+00 | 5.81E-02 | 1.04E+00 |
| ENSG000001 | GNB3 | 4.62E-01 | 7.67E-01 | 9.01E-01 | -1.51E-01 | -1.11E+00 |
| ENSG000001 | ELOVL6 | 4.62E-01 | 7.67E-01 | 9.72E-01 | -4.09E-02 | -1.03E+00 |
| ENSG000001 | MPP7 | 4.62E-01 | 7.67E-01 | 9.34E-01 | -9.79E-02 | -1.07E+00 |
| ENSG000001 | COQ10A | 4.62E-01 | 7.67E-01 | 9.41E-01 | -8.75E-02 | -1.06E+00 |
| ENSG000002 | PDCL3P4 | 4.62E-01 | 7.67E-01 | 9.07E-01 | -1.41E-01 | -1.10E+00 |
| ENSG000002 | --- | 4.62E-01 | ? | 1.14E+00 | 1.95E-01 | 1.14E+00 |
| ENSG000001 | CES4A | 4.62E-01 | 7.67E-01 | 1.08E+00 | 1.08E-01 | 1.08E+00 |
| ENSG000002 | --- | 4.62E-01 | ? | 1.43E+00 | 5.13E-01 | 1.43E+00 |
| ENSG000002 | CEP164P1 | 4.62E-01 | ? | 1.32E+00 | 3.98E-01 | 1.32E+00 |
| ENSG000002 | PRSS23-AS1 | 4.62E-01 | ? | 1.38E+00 | 4.61E-01 | 1.38E+00 |
| ENSG000002 | --- | 4.62E-01 | ? | 7.08E-01 | -4.98E-01 | -1.41E+00 |
| ENSG000001 | CFAP54 | 4.62E-01 | 7.67E-01 | 8.94E-01 | -1.61E-01 | -1.12E+00 |
| ENSG000002 | HECW2-AS1 | 4.63E-01 | ? | 1.16E+00 | 2.14E-01 | 1.16E+00 |
| ENSG000001 | BIRC6 | 4.63E-01 | 7.67E-01 | 9.57E-01 | -6.37E-02 | -1.05E+00 |
| ENSG000001 | PTDSS1 | 4.63E-01 | 7.67E-01 | 1.02E+00 | 3.25E-02 | 1.02E+00 |
| ENSG000001 | BAG3 | 4.63E-01 | 7.67E-01 | 9.81E-01 | -2.82E-02 | -1.02E+00 |
| ENSG000000 | ARHGAP31 | 4.63E-01 | 7.67E-01 | 9.53E-01 | -7.00E-02 | -1.05E+00 |
| ENSG000001 | PABPC1L | 4.63E-01 | 7.67E-01 | 9.12E-01 | -1.33E-01 | -1.10E+00 |
| ENSG000002 | DCTN6-DT | 4.63E-01 | ? | 1.23E+00 | 3.03E-01 | 1.23E+00 |
| ENSG000002 | --- | 4.63E-01 | ? | 1.86E+00 | 8.95E-01 | 1.86E+00 |

| | | | | | | |
|---|---|---|---|---|---|---|
| ENSG000002 | ZC3H12A-DT | 4.63E-01 | 7.67E-01 | 1.09E+00 | 1.23E-01 | 1.09E+00 |
| ENSG000001 | ZNF257 | 4.63E-01 | 7.67E-01 | 9.08E-01 | -1.40E-01 | -1.10E+00 |
| ENSG000002 | --- | 4.63E-01 | 7.67E-01 | 8.93E-01 | -1.63E-01 | -1.12E+00 |
| ENSG000001 | CYB5R2 | 4.63E-01 | 7.67E-01 | 1.06E+00 | 9.08E-02 | 1.06E+00 |
| ENSG000001 | ANKRA2 | 4.63E-01 | 7.67E-01 | 1.03E+00 | 4.89E-02 | 1.03E+00 |
| ENSG000001 | DIXDC1 | 4.63E-01 | 7.67E-01 | 1.03E+00 | 4.94E-02 | 1.03E+00 |
| ENSG000001 | SYT14 | 4.63E-01 | 7.68E-01 | 8.81E-01 | -1.82E-01 | -1.13E+00 |
| ENSG000001 | FAHD1 | 4.64E-01 | 7.68E-01 | 1.03E+00 | 4.53E-02 | 1.03E+00 |
| ENSG000001 | HBS1L | 4.64E-01 | 7.68E-01 | 9.77E-01 | -3.29E-02 | -1.02E+00 |
| ENSG000002 | NSUN6 | 4.64E-01 | 7.68E-01 | 9.58E-01 | -6.27E-02 | -1.04E+00 |
| ENSG000001 | NUP54 | 4.64E-01 | 7.68E-01 | 9.74E-01 | -3.87E-02 | -1.03E+00 |
| ENSG000001 | TRIM69 | 4.64E-01 | 7.68E-01 | 9.60E-01 | -5.82E-02 | -1.04E+00 |
| ENSG000001 | CYB5A | 4.64E-01 | 7.68E-01 | 1.03E+00 | 3.70E-02 | 1.03E+00 |
| ENSG000001 | TPM1 | 4.64E-01 | 7.68E-01 | 1.04E+00 | 5.37E-02 | 1.04E+00 |
| ENSG000001 | STX19 | 4.64E-01 | ? | 8.41E-01 | -2.49E-01 | -1.19E+00 |
| ENSG000002 | --- | 4.64E-01 | ? | 7.70E-01 | -3.78E-01 | -1.30E+00 |
| ENSG000001 | MOSPD3 | 4.64E-01 | 7.68E-01 | 1.07E+00 | 9.42E-02 | 1.07E+00 |
| ENSG000001 | SVBP | 4.64E-01 | 7.68E-01 | 9.53E-01 | -6.89E-02 | -1.05E+00 |
| ENSG000002 | SHLD3 | 4.64E-01 | 7.68E-01 | 9.28E-01 | -1.08E-01 | -1.08E+00 |
| ENSG000001 | PPIL2 | 4.64E-01 | 7.68E-01 | 1.02E+00 | 3.10E-02 | 1.02E+00 |
| ENSG000001 | PLEKHN1 | 4.64E-01 | 7.68E-01 | 1.05E+00 | 6.40E-02 | 1.05E+00 |
| ENSG000001 | GFER | 4.64E-01 | 7.68E-01 | 1.05E+00 | 7.34E-02 | 1.05E+00 |
| ENSG00000C | HS3ST1 | 4.64E-01 | 7.68E-01 | 9.35E-01 | -9.68E-02 | -1.07E+00 |
| ENSG000002 | --- | 4.65E-01 | ? | 1.56E+00 | 6.41E-01 | 1.56E+00 |
| ENSG000002 | ARHGAP11A- | 4.65E-01 | ? | 1.24E+00 | 3.05E-01 | 1.24E+00 |
| ENSG000002 | --- | 4.65E-01 | ? | 7.54E-01 | -4.07E-01 | -1.33E+00 |
| ENSG000001 | TRERF1 | 4.65E-01 | 7.68E-01 | 1.02E+00 | 3.07E-02 | 1.02E+00 |
| ENSG000002 | LINC00685 | 4.65E-01 | ? | 8.06E-01 | -3.11E-01 | -1.24E+00 |
| ENSG000001 | DISP2 | 4.65E-01 | 7.68E-01 | 1.05E+00 | 7.09E-02 | 1.05E+00 |
| ENSG000001 | OR7E13P | 4.65E-01 | ? | 1.39E+00 | 4.80E-01 | 1.39E+00 |
| ENSG000002 | GATA6-AS1 | 4.65E-01 | ? | 1.20E+00 | 2.60E-01 | 1.20E+00 |
| ENSG000002 | --- | 4.65E-01 | ? | 1.22E+00 | 2.84E-01 | 1.22E+00 |
| ENSG00000C | CA12 | 4.65E-01 | 7.69E-01 | 1.09E+00 | 1.26E-01 | 1.09E+00 |
| ENSG000002 | --- | 4.65E-01 | 7.69E-01 | 9.52E-01 | -7.05E-02 | -1.05E+00 |
| ENSG000002 | --- | 4.65E-01 | 7.69E-01 | 1.09E+00 | 1.30E-01 | 1.09E+00 |
| ENSG000001 | TRAF3 | 4.65E-01 | 7.69E-01 | 1.02E+00 | 3.02E-02 | 1.02E+00 |
| ENSG00000C | ATP2B1 | 4.65E-01 | 7.69E-01 | 1.03E+00 | 4.38E-02 | 1.03E+00 |
| ENSG000001 | NT5C3B | 4.66E-01 | 7.69E-01 | 1.03E+00 | 4.05E-02 | 1.03E+00 |
| ENSG000002 | TWIST2 | 4.66E-01 | 7.69E-01 | 9.25E-01 | -1.12E-01 | -1.08E+00 |
| ENSG000002 | --- | 4.66E-01 | 7.69E-01 | 1.12E+00 | 1.67E-01 | 1.12E+00 |
| ENSG000002 | --- | 4.66E-01 | ? | 6.80E-01 | -5.56E-01 | -1.47E+00 |
| ENSG000002 | ZNF736 | 4.66E-01 | 7.69E-01 | 9.61E-01 | -5.76E-02 | -1.04E+00 |
| ENSG000001 | CSE1L | 4.66E-01 | 7.69E-01 | 1.02E+00 | 2.95E-02 | 1.02E+00 |
| ENSG000002 | --- | 4.66E-01 | ? | 8.40E-01 | -2.52E-01 | -1.19E+00 |
| ENSG000001 | OPA3 | 4.66E-01 | 7.69E-01 | 1.02E+00 | 2.83E-02 | 1.02E+00 |

| | | | | | | |
|---|---|---|---|---|---|---|
| ENSG000001 | ARHGAP45 | 4.66E-01 | 7.69E-01 | 1.08E+00 | 1.14E-01 | 1.08E+00 |
| ENSG000001 | TRDMT1 | 4.66E-01 | 7.69E-01 | 9.50E-01 | -7.36E-02 | -1.05E+00 |
| ENSG000002 | --- | 4.66E-01 | ? | 7.83E-01 | -3.53E-01 | -1.28E+00 |
| ENSG000002 | --- | 4.66E-01 | ? | 1.49E+00 | 5.80E-01 | 1.49E+00 |
| ENSG000001 | IL1RN | 4.66E-01 | 7.69E-01 | 1.04E+00 | 5.44E-02 | 1.04E+00 |
| ENSG000001 | --- | 4.66E-01 | 7.69E-01 | 9.48E-01 | -7.67E-02 | -1.05E+00 |
| ENSG000001 | FBXO28 | 4.66E-01 | 7.69E-01 | 9.82E-01 | -2.61E-02 | -1.02E+00 |
| ENSG000002 | --- | 4.66E-01 | 7.69E-01 | 1.10E+00 | 1.36E-01 | 1.10E+00 |
| ENSG000001 | GPR132 | 4.66E-01 | 7.69E-01 | 8.71E-01 | -1.98E-01 | -1.15E+00 |
| ENSG000002 | LINC00842 | 4.66E-01 | 7.69E-01 | 1.05E+00 | 7.05E-02 | 1.05E+00 |
| ENSG000001 | PPFIBP1 | 4.66E-01 | 7.69E-01 | 1.02E+00 | 3.50E-02 | 1.02E+00 |
| ENSG000001 | STOX2 | 4.66E-01 | 7.69E-01 | 1.05E+00 | 7.44E-02 | 1.05E+00 |
| ENSG000002 | --- | 4.67E-01 | ? | 7.95E-01 | -3.31E-01 | -1.26E+00 |
| ENSG000002 | --- | 4.67E-01 | ? | 8.55E-01 | -2.27E-01 | -1.17E+00 |
| ENSG000001 | TMEM102 | 4.67E-01 | 7.69E-01 | 1.05E+00 | 6.92E-02 | 1.05E+00 |
| ENSG000002 | ZNF788P | 4.67E-01 | ? | 8.58E-01 | -2.21E-01 | -1.17E+00 |
| ENSG000002 | --- | 4.67E-01 | ? | 1.46E+00 | 5.41E-01 | 1.46E+00 |
| ENSG000001 | CCDC144CP | 4.67E-01 | ? | 1.15E+00 | 1.98E-01 | 1.15E+00 |
| ENSG000002 | RHPN1-AS1 | 4.67E-01 | ? | 7.77E-01 | -3.63E-01 | -1.29E+00 |
| ENSG00000C | NCK2 | 4.67E-01 | 7.70E-01 | 9.74E-01 | -3.82E-02 | -1.03E+00 |
| ENSG000001 | RCE1 | 4.67E-01 | 7.70E-01 | 1.07E+00 | 9.40E-02 | 1.07E+00 |
| ENSG000001 | SPAG17 | 4.67E-01 | ? | 1.15E+00 | 1.98E-01 | 1.15E+00 |
| ENSG000001 | CSRNP1 | 4.67E-01 | 7.70E-01 | 9.56E-01 | -6.50E-02 | -1.05E+00 |
| ENSG000001 | MYPN | 4.67E-01 | 7.70E-01 | 8.93E-01 | -1.64E-01 | -1.12E+00 |
| ENSG000001 | TUBB4B | 4.67E-01 | 7.70E-01 | 9.77E-01 | -3.41E-02 | -1.02E+00 |
| ENSG000001 | GLMN | 4.67E-01 | 7.70E-01 | 9.61E-01 | -5.78E-02 | -1.04E+00 |
| ENSG000002 | --- | 4.67E-01 | ? | 8.89E-01 | -1.70E-01 | -1.13E+00 |
| ENSG00000C | UPF1 | 4.67E-01 | 7.70E-01 | 1.03E+00 | 4.52E-02 | 1.03E+00 |
| ENSG000001 | NDUFA8 | 4.67E-01 | 7.70E-01 | 1.03E+00 | 4.18E-02 | 1.03E+00 |
| ENSG000001 | KRBA2 | 4.67E-01 | 7.70E-01 | 9.39E-01 | -9.05E-02 | -1.06E+00 |
| ENSG000001 | CHST14 | 4.68E-01 | 7.70E-01 | 1.05E+00 | 6.87E-02 | 1.05E+00 |
| ENSG000001 | ZNF335 | 4.68E-01 | 7.70E-01 | 1.03E+00 | 4.73E-02 | 1.03E+00 |
| ENSG000002 | --- | 4.68E-01 | ? | 7.55E-01 | -4.06E-01 | -1.32E+00 |
| ENSG000002 | DYNC1LI2-DT | 4.68E-01 | ? | 1.27E+00 | 3.45E-01 | 1.27E+00 |
| ENSG000001 | USB1 | 4.68E-01 | 7.70E-01 | 1.02E+00 | 3.16E-02 | 1.02E+00 |
| ENSG000002 | HNRNPD-DT | 4.68E-01 | ? | 1.31E+00 | 3.89E-01 | 1.31E+00 |
| ENSG000001 | ARL11 | 4.68E-01 | ? | 1.33E+00 | 4.13E-01 | 1.33E+00 |
| ENSG000001 | IFITM3 | 4.68E-01 | 7.70E-01 | 1.03E+00 | 4.79E-02 | 1.03E+00 |
| ENSG000002 | --- | 4.68E-01 | ? | 1.22E+00 | 2.88E-01 | 1.22E+00 |
| ENSG000001 | PAQR6 | 4.68E-01 | 7.70E-01 | 8.99E-01 | -1.54E-01 | -1.11E+00 |
| ENSG000001 | SPATC1L | 4.68E-01 | 7.70E-01 | 1.06E+00 | 8.70E-02 | 1.06E+00 |
| ENSG000001 | SLC25A45 | 4.68E-01 | ? | 1.15E+00 | 2.03E-01 | 1.15E+00 |
| ENSG000002 | KRT16P5 | 4.68E-01 | ? | 1.18E+00 | 2.40E-01 | 1.18E+00 |
| ENSG000001 | CPSF3 | 4.68E-01 | 7.70E-01 | 9.76E-01 | -3.58E-02 | -1.03E+00 |
| ENSG000001 | ZNF837 | 4.68E-01 | 7.70E-01 | 1.15E+00 | 1.99E-01 | 1.15E+00 |

| | | | | | | |
|---|---|---|---|---|---|---|
| ENSG000002 | --- | 4.68E-01 | ? | 1.38E+00 | 4.59E-01 | 1.38E+00 |
| ENSG00000C | CDC14B | 4.68E-01 | 7.71E-01 | 9.68E-01 | -4.69E-02 | -1.03E+00 |
| ENSG000001 | NRG4 | 4.68E-01 | ? | 1.17E+00 | 2.26E-01 | 1.17E+00 |
| ENSG000001 | MCRIP2 | 4.68E-01 | 7.71E-01 | 1.08E+00 | 1.13E-01 | 1.08E+00 |
| ENSG000001 | ZNF311 | 4.68E-01 | 7.71E-01 | 1.06E+00 | 8.97E-02 | 1.06E+00 |
| ENSG000002 | --- | 4.68E-01 | 7.71E-01 | 9.26E-01 | -1.10E-01 | -1.08E+00 |
| ENSG000002 | --- | 4.68E-01 | 7.71E-01 | 9.03E-01 | -1.48E-01 | -1.11E+00 |
| ENSG000001 | SLC9A2 | 4.68E-01 | ? | 8.86E-01 | -1.74E-01 | -1.13E+00 |
| ENSG000001 | GATD3 | 4.69E-01 | 7.71E-01 | 1.04E+00 | 5.69E-02 | 1.04E+00 |
| ENSG000002 | --- | 4.69E-01 | 7.71E-01 | 9.13E-01 | -1.31E-01 | -1.09E+00 |
| ENSG000002 | --- | 4.69E-01 | ? | 1.42E+00 | 5.07E-01 | 1.42E+00 |
| ENSG000001 | ZNF560 | 4.69E-01 | ? | 8.56E-01 | -2.24E-01 | -1.17E+00 |
| ENSG000002 | --- | 4.69E-01 | ? | 1.36E+00 | 4.42E-01 | 1.36E+00 |
| ENSG000001 | HMGB1P14 | 4.69E-01 | ? | 1.49E+00 | 5.79E-01 | 1.49E+00 |
| ENSG00000C | SLC38A5 | 4.69E-01 | 7.71E-01 | 1.03E+00 | 4.39E-02 | 1.03E+00 |
| ENSG000002 | --- | 4.69E-01 | 7.71E-01 | 9.40E-01 | -8.88E-02 | -1.06E+00 |
| ENSG000002 | --- | 4.69E-01 | ? | 1.20E+00 | 2.68E-01 | 1.20E+00 |
| ENSG000001 | --- | 4.69E-01 | ? | 8.42E-01 | -2.48E-01 | -1.19E+00 |
| ENSG000002 | --- | 4.69E-01 | 7.71E-01 | 1.30E+00 | 3.80E-01 | 1.30E+00 |
| ENSG000001 | ACP2 | 4.69E-01 | 7.71E-01 | 1.03E+00 | 4.90E-02 | 1.03E+00 |
| ENSG000001 | TTC28 | 4.69E-01 | 7.72E-01 | 1.05E+00 | 7.04E-02 | 1.05E+00 |
| ENSG000001 | ZCCHC17 | 4.70E-01 | 7.72E-01 | 9.76E-01 | -3.44E-02 | -1.02E+00 |
| ENSG000001 | IPO8 | 4.70E-01 | 7.72E-01 | 1.02E+00 | 3.47E-02 | 1.02E+00 |
| ENSG000001 | SDHAP1 | 4.70E-01 | 7.72E-01 | 8.53E-01 | -2.29E-01 | -1.17E+00 |
| ENSG000001 | PHC1P1 | 4.70E-01 | 7.72E-01 | 1.11E+00 | 1.47E-01 | 1.11E+00 |
| ENSG000002 | --- | 4.70E-01 | ? | 8.33E-01 | -2.64E-01 | -1.20E+00 |
| ENSG00000C | HCN2 | 4.70E-01 | 7.72E-01 | 1.07E+00 | 1.04E-01 | 1.07E+00 |
| ENSG000001 | KLF9 | 4.70E-01 | 7.72E-01 | 1.04E+00 | 5.37E-02 | 1.04E+00 |
| ENSG000001 | NDUFB10 | 4.70E-01 | 7.72E-01 | 1.02E+00 | 3.31E-02 | 1.02E+00 |
| ENSG000001 | BIN1 | 4.70E-01 | 7.72E-01 | 1.05E+00 | 6.45E-02 | 1.05E+00 |
| ENSG000001 | NOCT | 4.70E-01 | 7.72E-01 | 1.04E+00 | 5.27E-02 | 1.04E+00 |
| ENSG000002 | LINC01357 | 4.70E-01 | ? | 8.93E-01 | -1.64E-01 | -1.12E+00 |
| ENSG000002 | --- | 4.70E-01 | 7.72E-01 | 8.95E-01 | -1.60E-01 | -1.12E+00 |
| ENSG000002 | APOM | 4.70E-01 | ? | 8.03E-01 | -3.17E-01 | -1.25E+00 |
| ENSG000002 | --- | 4.70E-01 | ? | 7.91E-01 | -3.38E-01 | -1.26E+00 |
| ENSG000001 | TMEM216 | 4.70E-01 | 7.72E-01 | 1.06E+00 | 8.34E-02 | 1.06E+00 |
| ENSG000001 | PTTG1 | 4.70E-01 | 7.72E-01 | 1.03E+00 | 4.24E-02 | 1.03E+00 |
| ENSG000001 | AMIGO1 | 4.70E-01 | 7.72E-01 | 1.10E+00 | 1.43E-01 | 1.10E+00 |
| ENSG00000C | PIK3C3 | 4.70E-01 | 7.72E-01 | 9.76E-01 | -3.47E-02 | -1.02E+00 |
| ENSG000001 | USF1 | 4.70E-01 | 7.72E-01 | 1.03E+00 | 4.15E-02 | 1.03E+00 |
| ENSG000002 | --- | 4.70E-01 | ? | 8.55E-01 | -2.26E-01 | -1.17E+00 |
| ENSG000002 | --- | 4.70E-01 | 7.72E-01 | 1.15E+00 | 2.08E-01 | 1.15E+00 |
| ENSG000001 | GTPBP2 | 4.71E-01 | 7.72E-01 | 9.66E-01 | -4.93E-02 | -1.03E+00 |
| ENSG000001 | POU3F1 | 4.71E-01 | 7.72E-01 | 1.12E+00 | 1.68E-01 | 1.12E+00 |
| ENSG000002 | LINC01133 | 4.71E-01 | 7.72E-01 | 1.06E+00 | 8.98E-02 | 1.06E+00 |

| | | | | | | |
|---|---|---|---|---|---|---|
| ENSG000001 | FOXK2 | 4.71E-01 | 7.72E-01 | 1.02E+00 | 2.19E-02 | 1.02E+00 |
| ENSG000002 | --- | 4.71E-01 | ? | 8.37E-01 | -2.56E-01 | -1.19E+00 |
| ENSG000001 | PWWP3B | 4.71E-01 | 7.72E-01 | 9.16E-01 | -1.26E-01 | -1.09E+00 |
| ENSG000002 | --- | 4.71E-01 | ? | 1.18E+00 | 2.33E-01 | 1.18E+00 |
| ENSG000001 | TSPAN1 | 4.71E-01 | 7.72E-01 | 1.04E+00 | 5.99E-02 | 1.04E+00 |
| ENSG000001 | LRRC8C | 4.71E-01 | 7.72E-01 | 1.02E+00 | 3.31E-02 | 1.02E+00 |
| ENSG000001 | CEP120 | 4.71E-01 | 7.72E-01 | 9.66E-01 | -5.05E-02 | -1.04E+00 |
| ENSG000001 | ABRACL | 4.71E-01 | 7.72E-01 | 9.75E-01 | -3.69E-02 | -1.03E+00 |
| ENSG000002 | --- | 4.71E-01 | 7.72E-01 | 1.08E+00 | 1.05E-01 | 1.08E+00 |
| ENSG000001 | HERC3 | 4.71E-01 | 7.72E-01 | 9.69E-01 | -4.56E-02 | -1.03E+00 |
| ENSG000001 | ANP32A | 4.71E-01 | 7.72E-01 | 9.79E-01 | -3.04E-02 | -1.02E+00 |
| ENSG000001 | FBLN2 | 4.71E-01 | 7.72E-01 | 1.09E+00 | 1.25E-01 | 1.09E+00 |
| ENSG000001 | C15orf61 | 4.71E-01 | 7.72E-01 | 9.19E-01 | -1.22E-01 | -1.09E+00 |
| ENSG000001 | VIRMA | 4.71E-01 | 7.72E-01 | 9.81E-01 | -2.76E-02 | -1.02E+00 |
| ENSG000002 | EEF1B2P3 | 4.71E-01 | 7.72E-01 | 1.11E+00 | 1.46E-01 | 1.11E+00 |
| ENSG000002 | --- | 4.71E-01 | ? | 1.47E+00 | 5.58E-01 | 1.47E+00 |
| ENSG000002 | MT-TP | 4.72E-01 | ? | 7.03E-01 | -5.09E-01 | -1.42E+00 |
| ENSG000002 | ZNF460-AS1 | 4.72E-01 | ? | 7.64E-01 | -3.88E-01 | -1.31E+00 |
| ENSG000002 | --- | 4.72E-01 | ? | 7.76E-01 | -3.65E-01 | -1.29E+00 |
| ENSG000002 | C6orf47 | 4.72E-01 | 7.72E-01 | 1.04E+00 | 5.12E-02 | 1.04E+00 |
| ENSG000002 | --- | 4.72E-01 | ? | 7.75E-01 | -3.68E-01 | -1.29E+00 |
| ENSG000001 | CCDC86 | 4.72E-01 | 7.73E-01 | 9.77E-01 | -3.43E-02 | -1.02E+00 |
| ENSG000001 | TUT4 | 4.72E-01 | 7.73E-01 | 1.04E+00 | 5.55E-02 | 1.04E+00 |
| ENSG000002 | --- | 4.72E-01 | 7.73E-01 | 9.23E-01 | -1.15E-01 | -1.08E+00 |
| ENSG000002 | LINC00598 | 4.72E-01 | ? | 7.31E-01 | -4.52E-01 | -1.37E+00 |
| ENSG000002 | DENND1C | 4.72E-01 | 7.73E-01 | 9.54E-01 | -6.72E-02 | -1.05E+00 |
| ENSG000001 | IRS2 | 4.72E-01 | 7.73E-01 | 1.07E+00 | 9.28E-02 | 1.07E+00 |
| ENSG000001 | FMR1 | 4.72E-01 | 7.73E-01 | 9.76E-01 | -3.47E-02 | -1.02E+00 |
| ENSG000002 | --- | 4.72E-01 | ? | 1.71E+00 | 7.76E-01 | 1.71E+00 |
| ENSG000002 | TECPR1 | 4.72E-01 | 7.73E-01 | 1.05E+00 | 7.16E-02 | 1.05E+00 |
| ENSG000002 | --- | 4.72E-01 | ? | 7.20E-01 | -4.75E-01 | -1.39E+00 |
| ENSG000001 | TRPV2 | 4.72E-01 | ? | 8.53E-01 | -2.29E-01 | -1.17E+00 |
| ENSG000001 | ARFRP1 | 4.72E-01 | 7.73E-01 | 1.04E+00 | 5.82E-02 | 1.04E+00 |
| ENSG00000C | PRSS3 | 4.72E-01 | ? | 8.26E-01 | -2.75E-01 | -1.21E+00 |
| ENSG000001 | USP45 | 4.73E-01 | 7.73E-01 | 9.73E-01 | -4.00E-02 | -1.03E+00 |
| ENSG000001 | CHCHD1 | 4.73E-01 | 7.73E-01 | 1.04E+00 | 5.69E-02 | 1.04E+00 |
| ENSG000001 | NUP85 | 4.73E-01 | 7.73E-01 | 1.03E+00 | 3.66E-02 | 1.03E+00 |
| ENSG000002 | --- | 4.73E-01 | ? | 1.18E+00 | 2.35E-01 | 1.18E+00 |
| ENSG000001 | PPP1R8 | 4.73E-01 | 7.73E-01 | 9.79E-01 | -3.08E-02 | -1.02E+00 |
| ENSG000002 | KLRK1 | 4.73E-01 | ? | 1.17E+00 | 2.23E-01 | 1.17E+00 |
| ENSG000001 | SPDYE5 | 4.73E-01 | ? | 8.81E-01 | -1.82E-01 | -1.13E+00 |
| ENSG000001 | PIK3R1 | 4.73E-01 | 7.73E-01 | 9.71E-01 | -4.19E-02 | -1.03E+00 |
| ENSG000001 | LHFPL5 | 4.73E-01 | 7.73E-01 | 9.08E-01 | -1.39E-01 | -1.10E+00 |
| ENSG000002 | FAM205A | 4.73E-01 | ? | 1.37E+00 | 4.59E-01 | 1.37E+00 |
| ENSG000002 | DNLZ | 4.73E-01 | 7.73E-01 | 1.10E+00 | 1.40E-01 | 1.10E+00 |

| | | | | | | |
|---|---|---|---|---|---|---|
| ENSG000002 | --- | 4.73E-01 | ? | 8.28E-01 | -2.72E-01 | -1.21E+00 |
| ENSG000002 | VN1R83P | 4.73E-01 | ? | 1.24E+00 | 3.14E-01 | 1.24E+00 |
| ENSG000001 | EPM2AIP1 | 4.73E-01 | 7.73E-01 | 1.03E+00 | 4.05E-02 | 1.03E+00 |
| ENSG000001 | CAMSAP1 | 4.73E-01 | 7.73E-01 | 1.02E+00 | 2.51E-02 | 1.02E+00 |
| ENSG000001 | REXO4 | 4.73E-01 | 7.73E-01 | 1.02E+00 | 3.22E-02 | 1.02E+00 |
| ENSG000001 | GNMT | 4.73E-01 | ? | 1.30E+00 | 3.82E-01 | 1.30E+00 |
| ENSG000001 | CC2D1A | 4.73E-01 | 7.73E-01 | 9.62E-01 | -5.53E-02 | -1.04E+00 |
| ENSG00000C | PILRA | 4.73E-01 | ? | 1.33E+00 | 4.11E-01 | 1.33E+00 |
| ENSG000002 | --- | 4.74E-01 | 7.74E-01 | 9.08E-01 | -1.40E-01 | -1.10E+00 |
| ENSG000001 | PLEKHA8P1 | 4.74E-01 | 7.74E-01 | 1.05E+00 | 7.12E-02 | 1.05E+00 |
| ENSG000002 | MIF | 4.74E-01 | 7.74E-01 | 1.09E+00 | 1.22E-01 | 1.09E+00 |
| ENSG00000C | FAM13B | 4.74E-01 | 7.74E-01 | 9.65E-01 | -5.09E-02 | -1.04E+00 |
| ENSG000001 | AGTRAP | 4.74E-01 | 7.74E-01 | 1.05E+00 | 6.83E-02 | 1.05E+00 |
| ENSG000002 | --- | 4.74E-01 | ? | 1.35E+00 | 4.37E-01 | 1.35E+00 |
| ENSG000001 | IL33 | 4.74E-01 | 7.74E-01 | 1.24E+00 | 3.16E-01 | 1.24E+00 |
| ENSG000002 | --- | 4.74E-01 | ? | 8.49E-01 | -2.37E-01 | -1.18E+00 |
| ENSG00000C | DYRK4 | 4.74E-01 | 7.74E-01 | 1.04E+00 | 5.01E-02 | 1.04E+00 |
| ENSG000001 | GOLGA8A | 4.74E-01 | 7.74E-01 | 9.16E-01 | -1.27E-01 | -1.09E+00 |
| ENSG000001 | CCNB2 | 4.74E-01 | 7.74E-01 | 9.32E-01 | -1.02E-01 | -1.07E+00 |
| ENSG000002 | --- | 4.74E-01 | ? | 8.38E-01 | -2.55E-01 | -1.19E+00 |
| ENSG000002 | NQO1-DT | 4.74E-01 | ? | 1.29E+00 | 3.63E-01 | 1.29E+00 |
| ENSG000002 | --- | 4.74E-01 | ? | 7.96E-01 | -3.28E-01 | -1.26E+00 |
| ENSG000002 | --- | 4.74E-01 | ? | 1.49E+00 | 5.74E-01 | 1.49E+00 |
| ENSG000001 | USP34 | 4.74E-01 | 7.74E-01 | 9.66E-01 | -5.01E-02 | -1.04E+00 |
| ENSG000001 | CLIC3 | 4.74E-01 | 7.74E-01 | 1.13E+00 | 1.72E-01 | 1.13E+00 |
| ENSG000001 | METTL14 | 4.75E-01 | 7.74E-01 | 9.74E-01 | -3.77E-02 | -1.03E+00 |
| ENSG000002 | --- | 4.75E-01 | 7.74E-01 | 9.29E-01 | -1.07E-01 | -1.08E+00 |
| ENSG000002 | PCDHGB2 | 4.75E-01 | ? | 1.16E+00 | 2.14E-01 | 1.16E+00 |
| ENSG000001 | EXO5 | 4.75E-01 | 7.74E-01 | 1.05E+00 | 6.57E-02 | 1.05E+00 |
| ENSG000001 | SORD | 4.75E-01 | 7.74E-01 | 1.03E+00 | 4.57E-02 | 1.03E+00 |
| ENSG000002 | RNPC3-DT | 4.75E-01 | ? | 7.12E-01 | -4.91E-01 | -1.41E+00 |
| ENSG000001 | LANCL3 | 4.75E-01 | 7.74E-01 | 1.10E+00 | 1.43E-01 | 1.10E+00 |
| ENSG000002 | TMEM44-AS: | 4.75E-01 | 7.74E-01 | 8.96E-01 | -1.59E-01 | -1.12E+00 |
| ENSG000001 | HMGCLL1 | 4.75E-01 | 7.74E-01 | 8.62E-01 | -2.14E-01 | -1.16E+00 |
| ENSG00000C | CLSPN | 4.75E-01 | 7.74E-01 | 9.08E-01 | -1.39E-01 | -1.10E+00 |
| ENSG00000C | PITX1 | 4.75E-01 | 7.74E-01 | 1.05E+00 | 6.76E-02 | 1.05E+00 |
| ENSG000001 | RORC | 4.75E-01 | ? | 1.33E+00 | 4.08E-01 | 1.33E+00 |
| ENSG000002 | TPM3P9 | 4.75E-01 | 7.75E-01 | 9.41E-01 | -8.73E-02 | -1.06E+00 |
| ENSG000001 | PDZRN3 | 4.75E-01 | ? | 1.18E+00 | 2.34E-01 | 1.18E+00 |
| ENSG000001 | CPA5 | 4.75E-01 | ? | 1.25E+00 | 3.24E-01 | 1.25E+00 |
| ENSG000001 | GPR37 | 4.75E-01 | 7.75E-01 | 1.06E+00 | 7.82E-02 | 1.06E+00 |
| ENSG000002 | MAP3K9-DT | 4.75E-01 | 7.75E-01 | 9.55E-01 | -6.68E-02 | -1.05E+00 |
| ENSG000001 | DRP2 | 4.76E-01 | ? | 1.26E+00 | 3.34E-01 | 1.26E+00 |
| ENSG000001 | POLI | 4.76E-01 | 7.75E-01 | 9.69E-01 | -4.57E-02 | -1.03E+00 |
| ENSG000001 | TMTC3 | 4.76E-01 | 7.75E-01 | 9.70E-01 | -4.37E-02 | -1.03E+00 |

| | | | | | | |
|---|---|---|---|---|---|---|
| ENSG000002 | NIPBL-DT | 4.76E-01 | 7.75E-01 | 1.06E+00 | 8.26E-02 | 1.06E+00 |
| ENSG000002 | C8orf44 | 4.76E-01 | 7.75E-01 | 1.12E+00 | 1.65E-01 | 1.12E+00 |
| ENSG000001 | ADO | 4.76E-01 | 7.75E-01 | 1.02E+00 | 3.25E-02 | 1.02E+00 |
| ENSG000001 | TRIP12 | 4.76E-01 | 7.75E-01 | 9.81E-01 | -2.78E-02 | -1.02E+00 |
| ENSG000001 | CAGE1 | 4.76E-01 | ? | 6.82E-01 | -5.53E-01 | -1.47E+00 |
| ENSG000002 | TMEM123-D | 4.76E-01 | ? | 1.37E+00 | 4.50E-01 | 1.37E+00 |
| ENSG000001 | IGFBP2 | 4.76E-01 | 7.75E-01 | 1.04E+00 | 6.30E-02 | 1.04E+00 |
| ENSG000001 | SIPA1L3 | 4.76E-01 | 7.75E-01 | 1.03E+00 | 3.71E-02 | 1.03E+00 |
| ENSG000001 | GBA1 | 4.76E-01 | 7.75E-01 | 9.74E-01 | -3.77E-02 | -1.03E+00 |
| ENSG000001 | SLC43A1 | 4.76E-01 | ? | 7.81E-01 | -3.57E-01 | -1.28E+00 |
| ENSG000001 | FEM1C | 4.76E-01 | 7.75E-01 | 9.66E-01 | -4.93E-02 | -1.03E+00 |
| ENSG000002 | --- | 4.76E-01 | ? | 1.37E+00 | 4.51E-01 | 1.37E+00 |
| ENSG000001 | SPX | 4.76E-01 | ? | 8.98E-01 | -1.55E-01 | -1.11E+00 |
| ENSG000001 | NKAPL | 4.76E-01 | ? | 1.21E+00 | 2.73E-01 | 1.21E+00 |
| ENSG000000 | HIPK2 | 4.77E-01 | 7.76E-01 | 1.05E+00 | 6.46E-02 | 1.05E+00 |
| ENSG000001 | KCTD21 | 4.77E-01 | 7.76E-01 | 1.03E+00 | 4.91E-02 | 1.03E+00 |
| ENSG000001 | GART | 4.77E-01 | 7.76E-01 | 1.02E+00 | 3.17E-02 | 1.02E+00 |
| ENSG000001 | RFTN2 | 4.77E-01 | ? | 8.10E-01 | -3.05E-01 | -1.24E+00 |
| ENSG000001 | H2AC20 | 4.77E-01 | ? | 1.37E+00 | 4.50E-01 | 1.37E+00 |
| ENSG000001 | RHEB | 4.77E-01 | 7.76E-01 | 9.73E-01 | -3.88E-02 | -1.03E+00 |
| ENSG000002 | PPM1AP1 | 4.77E-01 | ? | 1.29E+00 | 3.63E-01 | 1.29E+00 |
| ENSG000001 | MARCHF1 | 4.77E-01 | 7.76E-01 | 9.14E-01 | -1.30E-01 | -1.09E+00 |
| ENSG000001 | BRD7 | 4.77E-01 | 7.76E-01 | 9.77E-01 | -3.41E-02 | -1.02E+00 |
| ENSG000001 | PHF8 | 4.77E-01 | 7.76E-01 | 1.02E+00 | 3.00E-02 | 1.02E+00 |
| ENSG000001 | PRDM16 | 4.77E-01 | 7.76E-01 | 9.43E-01 | -8.45E-02 | -1.06E+00 |
| ENSG000002 | L1TD1 | 4.77E-01 | ? | 8.39E-01 | -2.54E-01 | -1.19E+00 |
| ENSG000002 | --- | 4.77E-01 | ? | 7.24E-01 | -4.67E-01 | -1.38E+00 |
| ENSG000002 | --- | 4.77E-01 | ? | 1.29E+00 | 3.67E-01 | 1.29E+00 |
| ENSG000001 | ZNF471 | 4.77E-01 | 7.76E-01 | 9.59E-01 | -6.00E-02 | -1.04E+00 |
| ENSG000002 | ZFPM2-AS1 | 4.77E-01 | 7.76E-01 | 9.15E-01 | -1.29E-01 | -1.09E+00 |
| ENSG000002 | --- | 4.77E-01 | ? | 7.66E-01 | -3.84E-01 | -1.31E+00 |
| ENSG000002 | --- | 4.77E-01 | ? | 7.50E-01 | -4.15E-01 | -1.33E+00 |
| ENSG000002 | --- | 4.77E-01 | 7.76E-01 | 1.12E+00 | 1.66E-01 | 1.12E+00 |
| ENSG000002 | PCCA-DT | 4.78E-01 | 7.76E-01 | 1.06E+00 | 8.19E-02 | 1.06E+00 |
| ENSG000002 | --- | 4.78E-01 | ? | 8.71E-01 | -1.99E-01 | -1.15E+00 |
| ENSG000001 | ZNF576 | 4.78E-01 | 7.76E-01 | 1.04E+00 | 5.91E-02 | 1.04E+00 |
| ENSG000001 | CKAP2L | 4.78E-01 | 7.76E-01 | 9.11E-01 | -1.35E-01 | -1.10E+00 |
| ENSG000002 | HNRNPA3P9 | 4.78E-01 | ? | 8.52E-01 | -2.31E-01 | -1.17E+00 |
| ENSG000001 | FRA10AC1 | 4.78E-01 | 7.76E-01 | 9.67E-01 | -4.78E-02 | -1.03E+00 |
| ENSG000001 | DDX50 | 4.78E-01 | 7.76E-01 | 1.02E+00 | 2.82E-02 | 1.02E+00 |
| ENSG000000 | GLRX2 | 4.78E-01 | 7.76E-01 | 9.72E-01 | -4.06E-02 | -1.03E+00 |
| ENSG000002 | DCST1-AS1 | 4.78E-01 | ? | 7.80E-01 | -3.58E-01 | -1.28E+00 |
| ENSG000002 | HAUS6P1 | 4.78E-01 | ? | 7.42E-01 | -4.30E-01 | -1.35E+00 |
| ENSG000001 | ZFP90 | 4.78E-01 | 7.77E-01 | 1.03E+00 | 4.03E-02 | 1.03E+00 |
| ENSG000001 | IRF5 | 4.78E-01 | 7.77E-01 | 1.05E+00 | 7.21E-02 | 1.05E+00 |

| | | | | | | |
|---|---|---|---|---|---|---|
| ENSG000001 | MT-CO3 | 4.78E-01 | 7.77E-01 | 9.48E-01 | -7.63E-02 | -1.05E+00 |
| ENSG000001 | CITED1 | 4.78E-01 | ? | 1.47E+00 | 5.54E-01 | 1.47E+00 |
| ENSG000002 | RPL31P46 | 4.79E-01 | ? | 7.05E-01 | -5.04E-01 | -1.42E+00 |
| ENSG00000C | SNX1 | 4.79E-01 | 7.77E-01 | 1.02E+00 | 2.86E-02 | 1.02E+00 |
| ENSG000002 | --- | 4.79E-01 | ? | 8.05E-01 | -3.13E-01 | -1.24E+00 |
| ENSG000001 | SRFBP1 | 4.79E-01 | 7.78E-01 | 1.03E+00 | 4.94E-02 | 1.03E+00 |
| ENSG000001 | GDNF | 4.79E-01 | ? | 1.28E+00 | 3.53E-01 | 1.28E+00 |
| ENSG000001 | ATL2 | 4.79E-01 | 7.78E-01 | 9.79E-01 | -3.11E-02 | -1.02E+00 |
| ENSG000001 | EXOSC2 | 4.79E-01 | 7.78E-01 | 1.03E+00 | 4.76E-02 | 1.03E+00 |
| ENSG00000C | TM9SF3 | 4.79E-01 | 7.78E-01 | 9.82E-01 | -2.68E-02 | -1.02E+00 |
| ENSG000002 | --- | 4.79E-01 | ? | 1.45E+00 | 5.37E-01 | 1.45E+00 |
| ENSG00000C | ISL1 | 4.79E-01 | 7.78E-01 | 9.02E-01 | -1.49E-01 | -1.11E+00 |
| ENSG000001 | CNGA4 | 4.79E-01 | ? | 8.47E-01 | -2.40E-01 | -1.18E+00 |
| ENSG000001 | PACRGL | 4.79E-01 | 7.78E-01 | 9.65E-01 | -5.15E-02 | -1.04E+00 |
| ENSG000002 | ZPLD2P | 4.80E-01 | ? | 1.49E+00 | 5.72E-01 | 1.49E+00 |
| ENSG000002 | --- | 4.80E-01 | ? | 8.63E-01 | -2.12E-01 | -1.16E+00 |
| ENSG000001 | SLC12A5 | 4.80E-01 | ? | 1.40E+00 | 4.87E-01 | 1.40E+00 |
| ENSG000001 | HYAL1 | 4.80E-01 | 7.78E-01 | 9.14E-01 | -1.29E-01 | -1.09E+00 |
| ENSG000002 | --- | 4.80E-01 | ? | 5.55E-01 | -8.49E-01 | -1.80E+00 |
| ENSG000001 | C17orf97 | 4.80E-01 | 7.78E-01 | 1.10E+00 | 1.35E-01 | 1.10E+00 |
| ENSG000001 | PMS2 | 4.80E-01 | 7.78E-01 | 9.74E-01 | -3.78E-02 | -1.03E+00 |
| ENSG000001 | CHMP3 | 4.80E-01 | 7.78E-01 | 9.81E-01 | -2.81E-02 | -1.02E+00 |
| ENSG000002 | NSA2P2 | 4.80E-01 | ? | 1.41E+00 | 4.98E-01 | 1.41E+00 |
| ENSG000002 | PPIAP31 | 4.80E-01 | ? | 1.23E+00 | 3.04E-01 | 1.23E+00 |
| ENSG000001 | RNF213 | 4.80E-01 | 7.78E-01 | 1.05E+00 | 6.44E-02 | 1.05E+00 |
| ENSG000001 | AP2A2 | 4.80E-01 | 7.78E-01 | 1.02E+00 | 2.76E-02 | 1.02E+00 |
| ENSG000002 | --- | 4.80E-01 | ? | 7.77E-01 | -3.64E-01 | -1.29E+00 |
| ENSG000001 | TMEM165 | 4.80E-01 | 7.79E-01 | 9.83E-01 | -2.42E-02 | -1.02E+00 |
| ENSG000002 | HSPA1L | 4.80E-01 | 7.79E-01 | 1.08E+00 | 1.11E-01 | 1.08E+00 |
| ENSG000001 | FRMD3 | 4.80E-01 | ? | 8.86E-01 | -1.75E-01 | -1.13E+00 |
| ENSG000001 | ECHDC3 | 4.80E-01 | 7.79E-01 | 1.09E+00 | 1.23E-01 | 1.09E+00 |
| ENSG000001 | SNHG11 | 4.80E-01 | 7.79E-01 | 9.40E-01 | -8.88E-02 | -1.06E+00 |
| ENSG000002 | ILRUN-AS1 | 4.81E-01 | ? | 1.17E+00 | 2.21E-01 | 1.17E+00 |
| ENSG000001 | PTRH2 | 4.81E-01 | 7.79E-01 | 1.02E+00 | 3.34E-02 | 1.02E+00 |
| ENSG000001 | CDK2AP2 | 4.81E-01 | 7.79E-01 | 1.04E+00 | 5.38E-02 | 1.04E+00 |
| ENSG000002 | --- | 4.81E-01 | 7.79E-01 | 9.48E-01 | -7.65E-02 | -1.05E+00 |
| ENSG000001 | SLC39A4 | 4.81E-01 | 7.79E-01 | 1.08E+00 | 1.09E-01 | 1.08E+00 |
| ENSG000002 | CNPY2-AS1 | 4.81E-01 | 7.79E-01 | 9.10E-01 | -1.36E-01 | -1.10E+00 |
| ENSG000001 | WBP2 | 4.81E-01 | 7.79E-01 | 9.65E-01 | -5.09E-02 | -1.04E+00 |
| ENSG000001 | ENTREP3 | 4.81E-01 | 7.79E-01 | 1.04E+00 | 5.36E-02 | 1.04E+00 |
| ENSG000002 | POC1B-GALN | 4.81E-01 | 7.79E-01 | 7.60E-01 | -3.96E-01 | -1.32E+00 |
| ENSG000001 | PAK2 | 4.81E-01 | 7.79E-01 | 9.77E-01 | -3.37E-02 | -1.02E+00 |
| ENSG00000C | PRKCZ | 4.81E-01 | 7.79E-01 | 1.03E+00 | 4.45E-02 | 1.03E+00 |
| ENSG000001 | ITFG2 | 4.81E-01 | 7.79E-01 | 9.52E-01 | -7.09E-02 | -1.05E+00 |
| ENSG000002 | --- | 4.81E-01 | ? | 5.42E-01 | -8.83E-01 | -1.84E+00 |

| | | | | | | |
|---|---|---|---|---|---|---|
| ENSG000001 | C5 | 4.82E-01 | 7.79E-01 | 1.12E+00 | 1.66E-01 | 1.12E+00 |
| ENSG000002 | RTL8A | 4.82E-01 | 7.79E-01 | 1.04E+00 | 5.31E-02 | 1.04E+00 |
| ENSG000002 | ARAP1-AS2 | 4.82E-01 | ? | 8.71E-01 | -1.99E-01 | -1.15E+00 |
| ENSG000001 | CDKN2AIP | 4.82E-01 | 7.79E-01 | 9.63E-01 | -5.37E-02 | -1.04E+00 |
| ENSG000002 | ZNF778-DT | 4.82E-01 | 7.79E-01 | 1.09E+00 | 1.26E-01 | 1.09E+00 |
| ENSG000001 | TMEM229B | 4.82E-01 | 7.79E-01 | 1.06E+00 | 8.46E-02 | 1.06E+00 |
| ENSG000001 | HNRNPLL | 4.82E-01 | 7.79E-01 | 1.02E+00 | 3.31E-02 | 1.02E+00 |
| ENSG000002 | H2AZ1-DT | 4.82E-01 | ? | 7.92E-01 | -3.37E-01 | -1.26E+00 |
| ENSG000002 | --- | 4.82E-01 | ? | 1.19E+00 | 2.52E-01 | 1.19E+00 |
| ENSG000002 | ANKRD42-DT | 4.82E-01 | 7.80E-01 | 1.11E+00 | 1.50E-01 | 1.11E+00 |
| ENSG000001 | TUBB8B | 4.82E-01 | ? | 8.46E-01 | -2.41E-01 | -1.18E+00 |
| ENSG000001 | NAPSA | 4.82E-01 | 7.80E-01 | 1.12E+00 | 1.62E-01 | 1.12E+00 |
| ENSG00000C | PDE8A | 4.82E-01 | 7.80E-01 | 9.67E-01 | -4.78E-02 | -1.03E+00 |
| ENSG000002 | ZNF887P | 4.82E-01 | ? | 1.20E+00 | 2.61E-01 | 1.20E+00 |
| ENSG000001 | GSTA4 | 4.82E-01 | 7.80E-01 | 1.04E+00 | 5.39E-02 | 1.04E+00 |
| ENSG000002 | LINC01819 | 4.82E-01 | ? | 1.27E+00 | 3.49E-01 | 1.27E+00 |
| ENSG000002 | --- | 4.82E-01 | ? | 1.15E+00 | 2.04E-01 | 1.15E+00 |
| ENSG000002 | --- | 4.82E-01 | ? | 1.19E+00 | 2.51E-01 | 1.19E+00 |
| ENSG000002 | SBF2-AS1 | 4.82E-01 | 7.80E-01 | 1.08E+00 | 1.07E-01 | 1.08E+00 |
| ENSG00000C | NLRP1 | 4.82E-01 | 7.80E-01 | 1.03E+00 | 4.18E-02 | 1.03E+00 |
| ENSG000002 | --- | 4.83E-01 | ? | 7.28E-01 | -4.57E-01 | -1.37E+00 |
| ENSG000002 | GAS2L2 | 4.83E-01 | ? | 1.24E+00 | 3.12E-01 | 1.24E+00 |
| ENSG00000C | SYPL1 | 4.83E-01 | 7.80E-01 | 9.80E-01 | -2.97E-02 | -1.02E+00 |
| ENSG000002 | CT62 | 4.83E-01 | 7.80E-01 | 1.05E+00 | 6.89E-02 | 1.05E+00 |
| ENSG000001 | CCDC34 | 4.83E-01 | 7.80E-01 | 9.46E-01 | -7.95E-02 | -1.06E+00 |
| ENSG00000C | XK | 4.83E-01 | ? | 1.15E+00 | 2.06E-01 | 1.15E+00 |
| ENSG000002 | STK4-DT | 4.83E-01 | ? | 8.36E-01 | -2.59E-01 | -1.20E+00 |
| ENSG000001 | HMCES | 4.83E-01 | 7.80E-01 | 1.03E+00 | 3.75E-02 | 1.03E+00 |
| ENSG000001 | LTBR | 4.83E-01 | 7.80E-01 | 1.02E+00 | 3.30E-02 | 1.02E+00 |
| ENSG000002 | --- | 4.83E-01 | ? | 1.51E+00 | 5.91E-01 | 1.51E+00 |
| ENSG000001 | DNAL1 | 4.83E-01 | 7.80E-01 | 9.57E-01 | -6.28E-02 | -1.04E+00 |
| ENSG000002 | --- | 4.83E-01 | ? | 1.31E+00 | 3.91E-01 | 1.31E+00 |
| ENSG000001 | WIPF3 | 4.83E-01 | 7.80E-01 | 1.12E+00 | 1.63E-01 | 1.12E+00 |
| ENSG000001 | UBA3 | 4.83E-01 | 7.80E-01 | 9.73E-01 | -3.96E-02 | -1.03E+00 |
| ENSG000001 | FCSK | 4.83E-01 | 7.80E-01 | 1.06E+00 | 8.64E-02 | 1.06E+00 |
| ENSG000002 | SPDYE2 | 4.83E-01 | 7.80E-01 | 1.15E+00 | 1.98E-01 | 1.15E+00 |
| ENSG000001 | SLC10A6 | 4.83E-01 | ? | 1.15E+00 | 1.99E-01 | 1.15E+00 |
| ENSG000002 | PARGP1 | 4.83E-01 | 7.80E-01 | 1.11E+00 | 1.55E-01 | 1.11E+00 |
| ENSG000001 | PSME3IP1 | 4.83E-01 | 7.81E-01 | 9.74E-01 | -3.78E-02 | -1.03E+00 |
| ENSG000001 | GSTT2B | 4.84E-01 | ? | 8.65E-01 | -2.09E-01 | -1.16E+00 |
| ENSG000001 | IST1 | 4.84E-01 | 7.81E-01 | 1.02E+00 | 2.98E-02 | 1.02E+00 |
| ENSG000001 | DLEU2L | 4.84E-01 | ? | 1.15E+00 | 2.00E-01 | 1.15E+00 |
| ENSG000001 | JMY | 4.84E-01 | 7.81E-01 | 1.04E+00 | 5.50E-02 | 1.04E+00 |
| ENSG000001 | FKBP10 | 4.84E-01 | 7.81E-01 | 9.68E-01 | -4.67E-02 | -1.03E+00 |
| ENSG000001 | CEBPZ | 4.84E-01 | 7.81E-01 | 9.80E-01 | -2.92E-02 | -1.02E+00 |

| | | | | | | |
|---|---|---|---|---|---|---|
| ENSG000002 | --- | 4.84E-01 | ? | 1.36E+00 | 4.43E-01 | 1.36E+00 |
| ENSG000002 | --- | 4.84E-01 | ? | 6.75E-01 | -5.68E-01 | -1.48E+00 |
| ENSG000001 | GPRC5D | 4.84E-01 | ? | 8.04E-01 | -3.16E-01 | -1.24E+00 |
| ENSG000002 | GPC1-AS1 | 4.84E-01 | ? | 8.77E-01 | -1.89E-01 | -1.14E+00 |
| ENSG000001 | MPZL1 | 4.84E-01 | 7.81E-01 | 1.02E+00 | 2.32E-02 | 1.02E+00 |
| ENSG000002 | HILPDA-AS1 | 4.84E-01 | ? | 7.65E-01 | -3.86E-01 | -1.31E+00 |
| ENSG000001 | PRELP | 4.84E-01 | ? | 8.03E-01 | -3.17E-01 | -1.25E+00 |
| ENSG000002 | INF2 | 4.85E-01 | 7.82E-01 | 1.04E+00 | 5.96E-02 | 1.04E+00 |
| ENSG000002 | --- | 4.85E-01 | ? | 1.37E+00 | 4.49E-01 | 1.37E+00 |
| ENSG000001 | TIGD4 | 4.85E-01 | ? | 8.41E-01 | -2.51E-01 | -1.19E+00 |
| ENSG000001 | LRRC59 | 4.85E-01 | 7.82E-01 | 1.02E+00 | 2.23E-02 | 1.02E+00 |
| ENSG000001 | ARMC5 | 4.85E-01 | 7.82E-01 | 9.50E-01 | -7.41E-02 | -1.05E+00 |
| ENSG000002 | DNAJB6P1 | 4.85E-01 | 7.82E-01 | 1.10E+00 | 1.37E-01 | 1.10E+00 |
| ENSG000002 | --- | 4.85E-01 | 7.82E-01 | 8.75E-01 | -1.92E-01 | -1.14E+00 |
| ENSG000001 | SMIM19 | 4.85E-01 | 7.82E-01 | 1.05E+00 | 6.37E-02 | 1.05E+00 |
| ENSG000001 | CGGBP1 | 4.85E-01 | 7.82E-01 | 1.02E+00 | 2.98E-02 | 1.02E+00 |
| ENSG000001 | POM121L2 | 4.85E-01 | ? | 7.48E-01 | -4.18E-01 | -1.34E+00 |
| ENSG000001 | LCE1B | 4.85E-01 | ? | 1.34E+00 | 4.18E-01 | 1.34E+00 |
| ENSG000002 | RPS7P11 | 4.85E-01 | ? | 8.62E-01 | -2.14E-01 | -1.16E+00 |
| ENSG000001 | ANO7 | 4.85E-01 | 7.82E-01 | 1.07E+00 | 1.04E-01 | 1.07E+00 |
| ENSG00000C | CHRNA3 | 4.85E-01 | ? | 1.30E+00 | 3.81E-01 | 1.30E+00 |
| ENSG00000C | RCN1 | 4.85E-01 | 7.82E-01 | 9.79E-01 | -3.12E-02 | -1.02E+00 |
| ENSG000002 | TNK2-AS1 | 4.85E-01 | ? | 7.46E-01 | -4.23E-01 | -1.34E+00 |
| ENSG000002 | SGSM2-AS1 | 4.85E-01 | ? | 1.17E+00 | 2.22E-01 | 1.17E+00 |
| ENSG000002 | EEF1A1P11 | 4.85E-01 | ? | 7.79E-01 | -3.60E-01 | -1.28E+00 |
| ENSG00000C | DLG3 | 4.86E-01 | 7.83E-01 | 9.60E-01 | -5.96E-02 | -1.04E+00 |
| ENSG000001 | STK25 | 4.86E-01 | 7.83E-01 | 1.03E+00 | 3.96E-02 | 1.03E+00 |
| ENSG000001 | MRPL18 | 4.86E-01 | 7.83E-01 | 1.02E+00 | 3.09E-02 | 1.02E+00 |
| ENSG000001 | ADORA1 | 4.86E-01 | ? | 8.84E-01 | -1.78E-01 | -1.13E+00 |
| ENSG000002 | SLC38A4-AS1 | 4.86E-01 | 7.83E-01 | 1.11E+00 | 1.47E-01 | 1.11E+00 |
| ENSG000001 | OSBPL2 | 4.86E-01 | 7.83E-01 | 9.81E-01 | -2.75E-02 | -1.02E+00 |
| ENSG000001 | INO80C | 4.86E-01 | 7.83E-01 | 1.03E+00 | 4.87E-02 | 1.03E+00 |
| ENSG000001 | NAGPA | 4.86E-01 | 7.83E-01 | 9.59E-01 | -6.06E-02 | -1.04E+00 |
| ENSG000001 | IGSF6 | 4.86E-01 | ? | 7.36E-01 | -4.42E-01 | -1.36E+00 |
| ENSG000001 | PLD3 | 4.86E-01 | 7.83E-01 | 1.04E+00 | 5.28E-02 | 1.04E+00 |
| ENSG000001 | DCXR | 4.86E-01 | 7.83E-01 | 1.06E+00 | 8.29E-02 | 1.06E+00 |
| ENSG000001 | FIG4 | 4.86E-01 | 7.83E-01 | 1.03E+00 | 4.49E-02 | 1.03E+00 |
| ENSG000001 | SLC6A5 | 4.86E-01 | ? | 7.71E-01 | -3.75E-01 | -1.30E+00 |
| ENSG000002 | NDUFAF2P1 | 4.87E-01 | ? | 8.57E-01 | -2.22E-01 | -1.17E+00 |
| ENSG000001 | TMEM59 | 4.87E-01 | 7.83E-01 | 1.03E+00 | 4.19E-02 | 1.03E+00 |
| ENSG000002 | LACTB2-AS1 | 4.87E-01 | ? | 8.19E-01 | -2.89E-01 | -1.22E+00 |
| ENSG000002 | C2orf15 | 4.87E-01 | ? | 1.15E+00 | 1.96E-01 | 1.15E+00 |
| ENSG000001 | ADAMTS8 | 4.87E-01 | ? | 1.36E+00 | 4.43E-01 | 1.36E+00 |
| ENSG000002 | --- | 4.87E-01 | ? | 1.15E+00 | 2.02E-01 | 1.15E+00 |
| ENSG00000C | NUP133 | 4.87E-01 | 7.84E-01 | 1.02E+00 | 3.17E-02 | 1.02E+00 |

| | | | | | | |
|---|---|---|---|---|---|---|
| ENSG000001 | NIPSNAP3B | 4.87E-01 | 7.84E-01 | 1.10E+00 | 1.44E-01 | 1.10E+00 |
| ENSG000001 | CEP350 | 4.87E-01 | 7.84E-01 | 9.61E-01 | -5.69E-02 | -1.04E+00 |
| ENSG000001 | TRABD2A | 4.87E-01 | 7.84E-01 | 9.22E-01 | -1.16E-01 | -1.08E+00 |
| ENSG000002 | --- | 4.87E-01 | 7.84E-01 | 1.10E+00 | 1.33E-01 | 1.10E+00 |
| ENSG000001 | CEP131 | 4.87E-01 | 7.84E-01 | 1.06E+00 | 8.44E-02 | 1.06E+00 |
| ENSG000002 | MIR9-3HG | 4.87E-01 | ? | 1.31E+00 | 3.88E-01 | 1.31E+00 |
| ENSG000000 | GABRA3 | 4.87E-01 | ? | 1.33E+00 | 4.12E-01 | 1.33E+00 |
| ENSG000000 | XAB2 | 4.87E-01 | 7.84E-01 | 1.03E+00 | 4.70E-02 | 1.03E+00 |
| ENSG000001 | TTC30B | 4.88E-01 | 7.84E-01 | 1.06E+00 | 8.51E-02 | 1.06E+00 |
| ENSG000001 | CALM3 | 4.88E-01 | 7.84E-01 | 9.84E-01 | -2.26E-02 | -1.02E+00 |
| ENSG000002 | --- | 4.88E-01 | ? | 1.20E+00 | 2.63E-01 | 1.20E+00 |
| ENSG000000 | ICA1 | 4.88E-01 | 7.84E-01 | 1.03E+00 | 4.18E-02 | 1.03E+00 |
| ENSG000001 | PGGT1B | 4.88E-01 | 7.84E-01 | 9.66E-01 | -4.95E-02 | -1.03E+00 |
| ENSG000002 | CASC11 | 4.88E-01 | ? | 7.43E-01 | -4.29E-01 | -1.35E+00 |
| ENSG000001 | CPLANE1 | 4.88E-01 | 7.85E-01 | 1.05E+00 | 6.61E-02 | 1.05E+00 |
| ENSG000002 | ZNF436-AS1 | 4.88E-01 | 7.85E-01 | 9.17E-01 | -1.25E-01 | -1.09E+00 |
| ENSG000002 | TUBB7P | 4.88E-01 | ? | 1.34E+00 | 4.22E-01 | 1.34E+00 |
| ENSG000001 | MAP3K6 | 4.88E-01 | 7.85E-01 | 1.04E+00 | 5.55E-02 | 1.04E+00 |
| ENSG000002 | --- | 4.88E-01 | ? | 1.41E+00 | 4.96E-01 | 1.41E+00 |
| ENSG000001 | SPRYD3 | 4.88E-01 | 7.85E-01 | 1.03E+00 | 4.30E-02 | 1.03E+00 |
| ENSG000001 | NVL | 4.88E-01 | 7.85E-01 | 9.73E-01 | -4.01E-02 | -1.03E+00 |
| ENSG000001 | SNRPA1 | 4.89E-01 | 7.85E-01 | 9.76E-01 | -3.58E-02 | -1.03E+00 |
| ENSG000002 | RPL7AP10 | 4.89E-01 | ? | 8.21E-01 | -2.85E-01 | -1.22E+00 |
| ENSG000001 | KYAT1 | 4.89E-01 | 7.85E-01 | 1.07E+00 | 1.00E-01 | 1.07E+00 |
| ENSG000001 | PQBP1 | 4.89E-01 | 7.85E-01 | 9.72E-01 | -4.10E-02 | -1.03E+00 |
| ENSG000001 | FERMT3 | 4.89E-01 | 7.85E-01 | 8.99E-01 | -1.54E-01 | -1.11E+00 |
| ENSG000001 | LEAP2 | 4.89E-01 | ? | 7.74E-01 | -3.69E-01 | -1.29E+00 |
| ENSG000001 | PLK3 | 4.89E-01 | 7.85E-01 | 9.58E-01 | -6.18E-02 | -1.04E+00 |
| ENSG000001 | FUT1 | 4.89E-01 | 7.85E-01 | 1.04E+00 | 5.46E-02 | 1.04E+00 |
| ENSG000001 | LAMTOR5 | 4.89E-01 | 7.85E-01 | 1.02E+00 | 3.22E-02 | 1.02E+00 |
| ENSG000001 | ZNHIT1 | 4.89E-01 | 7.85E-01 | 1.04E+00 | 5.77E-02 | 1.04E+00 |
| ENSG000001 | MNX1 | 4.89E-01 | ? | 1.40E+00 | 4.86E-01 | 1.40E+00 |
| ENSG000001 | SULT1A1 | 4.89E-01 | 7.85E-01 | 9.31E-01 | -1.03E-01 | -1.07E+00 |
| ENSG000001 | MOB2 | 4.89E-01 | 7.85E-01 | 1.04E+00 | 6.23E-02 | 1.04E+00 |
| ENSG000002 | RPL30-AS1 | 4.89E-01 | 7.85E-01 | 9.26E-01 | -1.11E-01 | -1.08E+00 |
| ENSG000001 | FKBP9P1 | 4.89E-01 | 7.85E-01 | 9.07E-01 | -1.41E-01 | -1.10E+00 |
| ENSG000002 | FUT8-AS1 | 4.89E-01 | ? | 1.43E+00 | 5.12E-01 | 1.43E+00 |
| ENSG000001 | IBA57 | 4.89E-01 | 7.85E-01 | 1.05E+00 | 6.86E-02 | 1.05E+00 |
| ENSG000002 | --- | 4.89E-01 | ? | 1.14E+00 | 1.94E-01 | 1.14E+00 |
| ENSG000002 | --- | 4.90E-01 | ? | 1.37E+00 | 4.49E-01 | 1.37E+00 |
| ENSG000001 | BLTP3B | 4.90E-01 | 7.86E-01 | 1.03E+00 | 4.29E-02 | 1.03E+00 |
| ENSG000001 | MNS1 | 4.90E-01 | ? | 8.11E-01 | -3.03E-01 | -1.23E+00 |
| ENSG000002 | CROCCP2 | 4.90E-01 | 7.86E-01 | 9.53E-01 | -7.00E-02 | -1.05E+00 |
| ENSG000002 | --- | 4.90E-01 | 7.86E-01 | 8.68E-01 | -2.04E-01 | -1.15E+00 |
| ENSG000001 | CSTF3 | 4.90E-01 | 7.86E-01 | 1.02E+00 | 3.48E-02 | 1.02E+00 |

| | | | | | | |
|---|---|---|---|---|---|---|
| ENSG000002 | ANKRD20A11 | 4.90E-01 | ? | 7.73E-01 | -3.71E-01 | -1.29E+00 |
| ENSG000001 | SLC22A2 | 4.90E-01 | ? | 7.10E-01 | -4.94E-01 | -1.41E+00 |
| ENSG000001 | DSEL | 4.90E-01 | 7.86E-01 | 1.05E+00 | 7.67E-02 | 1.05E+00 |
| ENSG000002 | --- | 4.90E-01 | ? | 1.37E+00 | 4.58E-01 | 1.37E+00 |
| ENSG000002 | PCDHGB3 | 4.90E-01 | ? | 1.13E+00 | 1.74E-01 | 1.13E+00 |
| ENSG000001 | EIF3C | 4.90E-01 | 7.86E-01 | 9.85E-01 | -2.22E-02 | -1.02E+00 |
| ENSG000002 | --- | 4.90E-01 | ? | 1.12E+00 | 1.63E-01 | 1.12E+00 |
| ENSG000002 | EML6 | 4.90E-01 | ? | 1.11E+00 | 1.51E-01 | 1.11E+00 |
| ENSG000000 | NDUFB2 | 4.90E-01 | 7.86E-01 | 1.04E+00 | 5.71E-02 | 1.04E+00 |
| ENSG000002 | EIF4BP7 | 4.90E-01 | 7.86E-01 | 1.12E+00 | 1.60E-01 | 1.12E+00 |
| ENSG000002 | --- | 4.90E-01 | ? | 8.73E-01 | -1.97E-01 | -1.15E+00 |
| ENSG000002 | PRKCQ-AS1 | 4.90E-01 | 7.86E-01 | 9.55E-01 | -6.60E-02 | -1.05E+00 |
| ENSG000002 | WWP1-AS1 | 4.90E-01 | ? | 8.54E-01 | -2.28E-01 | -1.17E+00 |
| ENSG000001 | SUCLG1 | 4.90E-01 | 7.86E-01 | 1.02E+00 | 3.35E-02 | 1.02E+00 |
| ENSG000002 | --- | 4.90E-01 | ? | 1.53E+00 | 6.11E-01 | 1.53E+00 |
| ENSG000002 | --- | 4.90E-01 | ? | 1.37E+00 | 4.50E-01 | 1.37E+00 |
| ENSG000001 | ASF1B | 4.91E-01 | 7.86E-01 | 8.97E-01 | -1.57E-01 | -1.12E+00 |
| ENSG000001 | USP47 | 4.91E-01 | 7.86E-01 | 9.78E-01 | -3.18E-02 | -1.02E+00 |
| ENSG000001 | SERINC1 | 4.91E-01 | 7.86E-01 | 9.84E-01 | -2.31E-02 | -1.02E+00 |
| ENSG000001 | MALSU1 | 4.91E-01 | 7.86E-01 | 9.60E-01 | -5.90E-02 | -1.04E+00 |
| ENSG000001 | HCG4 | 4.91E-01 | 7.86E-01 | 9.06E-01 | -1.43E-01 | -1.10E+00 |
| ENSG000001 | TPMT | 4.91E-01 | 7.86E-01 | 1.04E+00 | 5.34E-02 | 1.04E+00 |
| ENSG000001 | CPT1C | 4.91E-01 | ? | 8.53E-01 | -2.30E-01 | -1.17E+00 |
| ENSG000002 | EGLN2 | 4.91E-01 | 7.87E-01 | 1.04E+00 | 5.85E-02 | 1.04E+00 |
| ENSG000002 | ZBTB45P1 | 4.91E-01 | ? | 7.54E-01 | -4.07E-01 | -1.33E+00 |
| ENSG000001 | CAP2 | 4.91E-01 | 7.87E-01 | 1.03E+00 | 4.69E-02 | 1.03E+00 |
| ENSG000000 | CYBA | 4.91E-01 | 7.87E-01 | 9.46E-01 | -7.99E-02 | -1.06E+00 |
| ENSG000002 | --- | 4.91E-01 | ? | 8.47E-01 | -2.39E-01 | -1.18E+00 |
| ENSG000001 | ZNF598 | 4.91E-01 | 7.87E-01 | 1.04E+00 | 6.27E-02 | 1.04E+00 |
| ENSG000002 | --- | 4.92E-01 | ? | 1.39E+00 | 4.78E-01 | 1.39E+00 |
| ENSG000001 | NUDT14 | 4.92E-01 | 7.87E-01 | 1.06E+00 | 8.03E-02 | 1.06E+00 |
| ENSG000002 | --- | 4.92E-01 | ? | 7.79E-01 | -3.59E-01 | -1.28E+00 |
| ENSG000001 | ANKRD54 | 4.92E-01 | 7.87E-01 | 9.73E-01 | -3.89E-02 | -1.03E+00 |
| ENSG000002 | --- | 4.92E-01 | 7.87E-01 | 9.27E-01 | -1.10E-01 | -1.08E+00 |
| ENSG000001 | PPP1R16A | 4.92E-01 | 7.87E-01 | 9.44E-01 | -8.27E-02 | -1.06E+00 |
| ENSG000002 | ZDBF2 | 4.92E-01 | 7.87E-01 | 1.03E+00 | 4.74E-02 | 1.03E+00 |
| ENSG000001 | LDLR | 4.92E-01 | 7.87E-01 | 9.76E-01 | -3.48E-02 | -1.02E+00 |
| ENSG000001 | DHX30 | 4.92E-01 | 7.87E-01 | 1.03E+00 | 4.21E-02 | 1.03E+00 |
| ENSG000001 | FGD5 | 4.92E-01 | ? | 7.55E-01 | -4.06E-01 | -1.33E+00 |
| ENSG000002 | --- | 4.92E-01 | ? | 8.68E-01 | -2.05E-01 | -1.15E+00 |
| ENSG000001 | MFSD4A | 4.92E-01 | ? | 1.17E+00 | 2.27E-01 | 1.17E+00 |
| ENSG000001 | CTU2 | 4.92E-01 | 7.88E-01 | 1.05E+00 | 7.11E-02 | 1.05E+00 |
| ENSG000001 | POLR3G | 4.93E-01 | 7.88E-01 | 1.06E+00 | 8.62E-02 | 1.06E+00 |
| ENSG000000 | UBR7 | 4.93E-01 | 7.88E-01 | 9.70E-01 | -4.44E-02 | -1.03E+00 |
| ENSG000002 | --- | 4.93E-01 | ? | 1.39E+00 | 4.76E-01 | 1.39E+00 |

| | | | | | | |
|---|---|---|---|---|---|---|
| ENSG000002 | METTL6 | 4.93E-01 | 7.88E-01 | 1.03E+00 | 4.55E-02 | 1.03E+00 |
| ENSG000001 | POLE3 | 4.93E-01 | 7.88E-01 | 9.76E-01 | -3.55E-02 | -1.02E+00 |
| ENSG000001 | NIT2 | 4.93E-01 | 7.88E-01 | 1.03E+00 | 4.20E-02 | 1.03E+00 |
| ENSG000001 | TMEM121B | 4.93E-01 | ? | 1.35E+00 | 4.29E-01 | 1.35E+00 |
| ENSG000002 | RXRB | 4.93E-01 | 7.88E-01 | 1.05E+00 | 6.72E-02 | 1.05E+00 |
| ENSG000002 | TSTD1 | 4.93E-01 | 7.88E-01 | 1.05E+00 | 7.14E-02 | 1.05E+00 |
| ENSG000001 | FBXO9 | 4.93E-01 | 7.88E-01 | 9.76E-01 | -3.51E-02 | -1.02E+00 |
| ENSG000002 | RPS3AP26 | 4.93E-01 | 7.88E-01 | 9.27E-01 | -1.10E-01 | -1.08E+00 |
| ENSG000002 | --- | 4.93E-01 | ? | 1.21E+00 | 2.72E-01 | 1.21E+00 |
| ENSG000002 | CYTOR | 4.93E-01 | 7.88E-01 | 9.62E-01 | -5.61E-02 | -1.04E+00 |
| ENSG000001 | ULK4 | 4.93E-01 | 7.88E-01 | 1.07E+00 | 1.02E-01 | 1.07E+00 |
| ENSG000001 | RNF181 | 4.94E-01 | 7.88E-01 | 1.03E+00 | 4.74E-02 | 1.03E+00 |
| ENSG000002 | --- | 4.94E-01 | ? | 6.96E-01 | -5.22E-01 | -1.44E+00 |
| ENSG000001 | PHACTR2 | 4.94E-01 | 7.88E-01 | 1.05E+00 | 6.39E-02 | 1.05E+00 |
| ENSG000002 | --- | 4.94E-01 | 7.88E-01 | 9.11E-01 | -1.34E-01 | -1.10E+00 |
| ENSG000001 | TGOLN2 | 4.94E-01 | 7.88E-01 | 1.02E+00 | 2.74E-02 | 1.02E+00 |
| ENSG000001 | BLOC1S5 | 4.94E-01 | 7.88E-01 | 1.04E+00 | 5.39E-02 | 1.04E+00 |
| ENSG000001 | RAPGEF5 | 4.94E-01 | 7.88E-01 | 1.06E+00 | 8.92E-02 | 1.06E+00 |
| ENSG000001 | C9orf24 | 4.94E-01 | ? | 1.45E+00 | 5.36E-01 | 1.45E+00 |
| ENSG000002 | CASTOR3P | 4.94E-01 | 7.89E-01 | 1.06E+00 | 8.14E-02 | 1.06E+00 |
| ENSG000001 | ZNF19 | 4.94E-01 | 7.89E-01 | 9.21E-01 | -1.19E-01 | -1.09E+00 |
| ENSG000001 | TP53I11 | 4.94E-01 | 7.89E-01 | 1.06E+00 | 8.91E-02 | 1.06E+00 |
| ENSG000001 | SEPTIN2 | 4.94E-01 | 7.89E-01 | 9.79E-01 | -3.07E-02 | -1.02E+00 |
| ENSG000002 | PRDX3P1 | 4.94E-01 | ? | 1.37E+00 | 4.58E-01 | 1.37E+00 |
| ENSG00000C | PANX2 | 4.95E-01 | 7.89E-01 | 9.48E-01 | -7.75E-02 | -1.06E+00 |
| ENSG000001 | MROH1 | 4.95E-01 | 7.89E-01 | 1.05E+00 | 6.92E-02 | 1.05E+00 |
| ENSG000002 | --- | 4.95E-01 | 7.89E-01 | 1.02E+00 | 3.12E-02 | 1.02E+00 |
| ENSG00000C | ALG1 | 4.95E-01 | 7.89E-01 | 1.03E+00 | 4.56E-02 | 1.03E+00 |
| ENSG000001 | CNOT8 | 4.95E-01 | 7.89E-01 | 1.02E+00 | 3.08E-02 | 1.02E+00 |
| ENSG000002 | --- | 4.95E-01 | ? | 1.21E+00 | 2.79E-01 | 1.21E+00 |
| ENSG000001 | SETX | 4.95E-01 | 7.89E-01 | 9.70E-01 | -4.37E-02 | -1.03E+00 |
| ENSG00000C | KIZ | 4.95E-01 | 7.89E-01 | 1.03E+00 | 4.88E-02 | 1.03E+00 |
| ENSG000001 | TK2 | 4.95E-01 | 7.89E-01 | 1.03E+00 | 4.88E-02 | 1.03E+00 |
| ENSG000002 | GEMIN7-AS1 | 4.95E-01 | 7.89E-01 | 1.11E+00 | 1.54E-01 | 1.11E+00 |
| ENSG000002 | EDRF1-DT | 4.95E-01 | ? | 1.13E+00 | 1.76E-01 | 1.13E+00 |
| ENSG000001 | SPEN-AS1 | 4.95E-01 | 7.89E-01 | 8.97E-01 | -1.57E-01 | -1.12E+00 |
| ENSG000002 | XKR4 | 4.95E-01 | ? | 8.79E-01 | -1.86E-01 | -1.14E+00 |
| ENSG000002 | AP4M1 | 4.95E-01 | 7.89E-01 | 1.05E+00 | 7.46E-02 | 1.05E+00 |
| ENSG000001 | ENOX1 | 4.95E-01 | 7.89E-01 | 9.01E-01 | -1.50E-01 | -1.11E+00 |
| ENSG00000C | WHRN | 4.95E-01 | 7.89E-01 | 1.08E+00 | 1.08E-01 | 1.08E+00 |
| ENSG000002 | --- | 4.95E-01 | ? | 7.45E-01 | -4.24E-01 | -1.34E+00 |
| ENSG000001 | FANCA | 4.95E-01 | 7.89E-01 | 9.40E-01 | -8.97E-02 | -1.06E+00 |
| ENSG000001 | FCGRT | 4.95E-01 | 7.89E-01 | 1.08E+00 | 1.14E-01 | 1.08E+00 |
| ENSG000001 | BOLA1 | 4.95E-01 | 7.89E-01 | 1.09E+00 | 1.28E-01 | 1.09E+00 |
| ENSG000002 | HERC2P9 | 4.96E-01 | 7.90E-01 | 9.44E-01 | -8.38E-02 | -1.06E+00 |

| | | | | | | |
|---|---|---|---|---|---|---|
| ENSG000001 | PHF19 | 4.96E-01 | 7.90E-01 | 1.04E+00 | 6.21E-02 | 1.04E+00 |
| ENSG000001 | PPP1R36 | 4.96E-01 | ? | 8.28E-01 | -2.72E-01 | -1.21E+00 |
| ENSG00000C | RGPD5 | 4.96E-01 | 7.90E-01 | 1.07E+00 | 1.02E-01 | 1.07E+00 |
| ENSG000001 | ZNF175 | 4.96E-01 | 7.90E-01 | 9.61E-01 | -5.70E-02 | -1.04E+00 |
| ENSG000001 | LIPG | 4.96E-01 | 7.90E-01 | 1.02E+00 | 3.39E-02 | 1.02E+00 |
| ENSG00000C | CBX5 | 4.96E-01 | 7.90E-01 | 1.03E+00 | 4.64E-02 | 1.03E+00 |
| ENSG000001 | ANKRD49 | 4.96E-01 | 7.90E-01 | 1.04E+00 | 5.05E-02 | 1.04E+00 |
| ENSG000002 | LINC00113 | 4.96E-01 | ? | 1.37E+00 | 4.53E-01 | 1.37E+00 |
| ENSG000001 | RALGAPA2 | 4.96E-01 | 7.90E-01 | 9.76E-01 | -3.47E-02 | -1.02E+00 |
| ENSG000001 | RBP5 | 4.96E-01 | 7.90E-01 | 9.08E-01 | -1.40E-01 | -1.10E+00 |
| ENSG000002 | --- | 4.96E-01 | ? | 1.41E+00 | 4.95E-01 | 1.41E+00 |
| ENSG000002 | --- | 4.96E-01 | ? | 8.33E-01 | -2.63E-01 | -1.20E+00 |
| ENSG000001 | USF2 | 4.96E-01 | 7.90E-01 | 9.73E-01 | -3.92E-02 | -1.03E+00 |
| ENSG000002 | --- | 4.97E-01 | ? | 1.26E+00 | 3.39E-01 | 1.26E+00 |
| ENSG000002 | --- | 4.97E-01 | 7.90E-01 | 1.10E+00 | 1.39E-01 | 1.10E+00 |
| ENSG000001 | TBX4 | 4.97E-01 | ? | 1.41E+00 | 4.92E-01 | 1.41E+00 |
| ENSG000001 | GTF3C6 | 4.97E-01 | 7.90E-01 | 9.75E-01 | -3.59E-02 | -1.03E+00 |
| ENSG000002 | --- | 4.97E-01 | ? | 1.28E+00 | 3.57E-01 | 1.28E+00 |
| ENSG000002 | --- | 4.97E-01 | ? | 1.52E+00 | 6.02E-01 | 1.52E+00 |
| ENSG000001 | ACTR10 | 4.97E-01 | 7.90E-01 | 9.76E-01 | -3.45E-02 | -1.02E+00 |
| ENSG000001 | OTOGL | 4.97E-01 | ? | 8.04E-01 | -3.14E-01 | -1.24E+00 |
| ENSG000001 | LACTB2 | 4.97E-01 | 7.91E-01 | 9.58E-01 | -6.19E-02 | -1.04E+00 |
| ENSG000001 | P4HA3 | 4.97E-01 | ? | 1.17E+00 | 2.25E-01 | 1.17E+00 |
| ENSG000001 | ZNF438 | 4.97E-01 | 7.91E-01 | 9.53E-01 | -6.93E-02 | -1.05E+00 |
| ENSG000001 | CYCS | 4.97E-01 | 7.91E-01 | 1.02E+00 | 3.19E-02 | 1.02E+00 |
| ENSG000001 | KRBOX4 | 4.97E-01 | 7.91E-01 | 9.63E-01 | -5.46E-02 | -1.04E+00 |
| ENSG000002 | --- | 4.97E-01 | ? | 1.40E+00 | 4.88E-01 | 1.40E+00 |
| ENSG000002 | RPL36AP37 | 4.97E-01 | 7.91E-01 | 8.82E-01 | -1.81E-01 | -1.13E+00 |
| ENSG000001 | ZNF507 | 4.97E-01 | 7.91E-01 | 9.67E-01 | -4.90E-02 | -1.03E+00 |
| ENSG000001 | LOXL3 | 4.97E-01 | ? | 1.13E+00 | 1.74E-01 | 1.13E+00 |
| ENSG000002 | --- | 4.97E-01 | ? | 8.90E-01 | -1.68E-01 | -1.12E+00 |
| ENSG000001 | MCM4 | 4.98E-01 | 7.91E-01 | 9.66E-01 | -4.95E-02 | -1.03E+00 |
| ENSG000001 | TMEM51-AS | 4.98E-01 | 7.91E-01 | 9.48E-01 | -7.76E-02 | -1.06E+00 |
| ENSG000001 | C15orf48 | 4.98E-01 | 7.91E-01 | 8.98E-01 | -1.55E-01 | -1.11E+00 |
| ENSG00000C | PUS7 | 4.98E-01 | 7.91E-01 | 1.03E+00 | 3.79E-02 | 1.03E+00 |
| ENSG000001 | SHLD2P3 | 4.98E-01 | ? | 1.29E+00 | 3.64E-01 | 1.29E+00 |
| ENSG000001 | ERCC6L | 4.98E-01 | 7.91E-01 | 9.14E-01 | -1.30E-01 | -1.09E+00 |
| ENSG000001 | NUP107 | 4.98E-01 | 7.91E-01 | 1.02E+00 | 3.16E-02 | 1.02E+00 |
| ENSG000001 | PAK1IP1 | 4.98E-01 | 7.91E-01 | 1.03E+00 | 3.73E-02 | 1.03E+00 |
| ENSG000001 | CCDC15 | 4.98E-01 | 7.91E-01 | 9.29E-01 | -1.07E-01 | -1.08E+00 |
| ENSG000001 | RAI2 | 4.98E-01 | ? | 8.05E-01 | -3.13E-01 | -1.24E+00 |
| ENSG000001 | TM4SF19 | 4.98E-01 | ? | 7.98E-01 | -3.26E-01 | -1.25E+00 |
| ENSG000001 | PPIAP22 | 4.98E-01 | 7.91E-01 | 1.03E+00 | 4.65E-02 | 1.03E+00 |
| ENSG00000C | TGFB2 | 4.98E-01 | 7.91E-01 | 9.25E-01 | -1.13E-01 | -1.08E+00 |
| ENSG000002 | LYPLA1P3 | 4.98E-01 | ? | 1.22E+00 | 2.90E-01 | 1.22E+00 |

| | | | | | | |
|---|---|---|---|---|---|---|
| ENSG000001 | AGR2 | 4.98E-01 | 7.91E-01 | 9.63E-01 | -5.49E-02 | -1.04E+00 |
| ENSG000001 | EIF3J-DT | 4.99E-01 | 7.91E-01 | 1.06E+00 | 8.00E-02 | 1.06E+00 |
| ENSG000002 | --- | 4.99E-01 | 7.91E-01 | 9.58E-01 | -6.17E-02 | -1.04E+00 |
| ENSG000002 | NPIPB2 | 4.99E-01 | 7.91E-01 | 1.08E+00 | 1.08E-01 | 1.08E+00 |
| ENSG000001 | RNF183 | 4.99E-01 ? | | 7.80E-01 | -3.58E-01 | -1.28E+00 |
| ENSG00000C | TDRD3 | 4.99E-01 | 7.91E-01 | 9.61E-01 | -5.74E-02 | -1.04E+00 |
| ENSG000001 | HGFAC | 4.99E-01 ? | | 1.35E+00 | 4.37E-01 | 1.35E+00 |
| ENSG000001 | TSPAN14 | 4.99E-01 | 7.91E-01 | 9.82E-01 | -2.58E-02 | -1.02E+00 |
| ENSG000001 | FAM227B | 4.99E-01 | 7.91E-01 | 1.07E+00 | 9.48E-02 | 1.07E+00 |
| ENSG000001 | SEC24B | 4.99E-01 | 7.91E-01 | 9.76E-01 | -3.57E-02 | -1.03E+00 |
| ENSG000001 | LMO2 | 4.99E-01 ? | | 1.13E+00 | 1.81E-01 | 1.13E+00 |
| ENSG000002 | --- | 4.99E-01 ? | | 7.67E-01 | -3.82E-01 | -1.30E+00 |
| ENSG000001 | GTPBP8 | 4.99E-01 | 7.91E-01 | 1.03E+00 | 3.85E-02 | 1.03E+00 |
| ENSG000002 | THAP7-AS1 | 4.99E-01 | 7.92E-01 | 1.06E+00 | 9.08E-02 | 1.06E+00 |
| ENSG000001 | TRAIP | 4.99E-01 | 7.92E-01 | 1.08E+00 | 1.13E-01 | 1.08E+00 |
| ENSG000002 | --- | 4.99E-01 | 7.92E-01 | 9.10E-01 | -1.36E-01 | -1.10E+00 |
| ENSG000002 | --- | 4.99E-01 | 7.92E-01 | 9.49E-01 | -7.62E-02 | -1.05E+00 |
| ENSG000002 | TEKT4P2 | 5.00E-01 | 7.92E-01 | 1.03E+00 | 3.64E-02 | 1.03E+00 |
| ENSG000001 | CCDC106 | 5.00E-01 | 7.92E-01 | 1.07E+00 | 1.02E-01 | 1.07E+00 |
| ENSG00000C | RBMS2 | 5.00E-01 | 7.92E-01 | 9.73E-01 | -4.02E-02 | -1.03E+00 |
| ENSG000001 | CASC2 | 5.00E-01 ? | | 1.18E+00 | 2.34E-01 | 1.18E+00 |
| ENSG000001 | SLC44A3 | 5.00E-01 | 7.92E-01 | 1.03E+00 | 4.59E-02 | 1.03E+00 |
| ENSG000001 | NOL6 | 5.00E-01 | 7.92E-01 | 1.03E+00 | 4.82E-02 | 1.03E+00 |
| ENSG000001 | CCDC96 | 5.00E-01 | 7.92E-01 | 9.31E-01 | -1.04E-01 | -1.07E+00 |
| ENSG000001 | CFL2 | 5.00E-01 | 7.92E-01 | 9.51E-01 | -7.22E-02 | -1.05E+00 |
| ENSG000001 | BCCIP | 5.00E-01 | 7.92E-01 | 9.70E-01 | -4.34E-02 | -1.03E+00 |
| ENSG000002 | OLA1P1 | 5.00E-01 ? | | 8.59E-01 | -2.19E-01 | -1.16E+00 |
| ENSG000002 | --- | 5.00E-01 ? | | 7.76E-01 | -3.66E-01 | -1.29E+00 |
| ENSG000001 | CXCR2 | 5.00E-01 ? | | 1.13E+00 | 1.73E-01 | 1.13E+00 |
| ENSG00000C | ZNF275 | 5.00E-01 | 7.93E-01 | 9.70E-01 | -4.44E-02 | -1.03E+00 |
| ENSG000001 | BCL2L1 | 5.00E-01 | 7.93E-01 | 9.76E-01 | -3.44E-02 | -1.02E+00 |
| ENSG000002 | --- | 5.00E-01 ? | | 8.46E-01 | -2.42E-01 | -1.18E+00 |
| ENSG000001 | DIRAS1 | 5.01E-01 ? | | 1.27E+00 | 3.50E-01 | 1.27E+00 |
| ENSG000002 | --- | 5.01E-01 ? | | 8.25E-01 | -2.77E-01 | -1.21E+00 |
| ENSG000001 | EVA1C | 5.01E-01 | 7.93E-01 | 1.08E+00 | 1.05E-01 | 1.08E+00 |
| ENSG000002 | LINC02967 | 5.01E-01 | 7.93E-01 | 1.06E+00 | 8.69E-02 | 1.06E+00 |
| ENSG000002 | --- | 5.01E-01 | 7.93E-01 | 1.10E+00 | 1.37E-01 | 1.10E+00 |
| ENSG000002 | RCC1L | 5.01E-01 | 7.93E-01 | 1.03E+00 | 4.84E-02 | 1.03E+00 |
| ENSG000001 | FAT3 | 5.01E-01 ? | | 1.40E+00 | 4.83E-01 | 1.40E+00 |
| ENSG000001 | POLR2I | 5.01E-01 | 7.93E-01 | 9.51E-01 | -7.23E-02 | -1.05E+00 |
| ENSG000001 | RIC8B | 5.01E-01 | 7.93E-01 | 9.68E-01 | -4.70E-02 | -1.03E+00 |
| ENSG000001 | MBLAC2 | 5.01E-01 | 7.93E-01 | 1.05E+00 | 7.37E-02 | 1.05E+00 |
| ENSG000002 | TNPO1P3 | 5.01E-01 | 7.93E-01 | 9.02E-01 | -1.49E-01 | -1.11E+00 |
| ENSG000001 | CIP2A | 5.01E-01 | 7.93E-01 | 9.59E-01 | -6.08E-02 | -1.04E+00 |
| ENSG00000C | TNK2 | 5.01E-01 | 7.93E-01 | 1.05E+00 | 6.86E-02 | 1.05E+00 |

| | | | | | | |
|---|---|---|---|---|---|---|
| ENSG000002 | --- | 5.01E-01 | ? | 8.95E-01 | -1.61E-01 | -1.12E+00 |
| ENSG000001 | SCAPER | 5.01E-01 | 7.93E-01 | 9.69E-01 | -4.48E-02 | -1.03E+00 |
| ENSG00000C | YBX1 | 5.01E-01 | 7.93E-01 | 9.70E-01 | -4.46E-02 | -1.03E+00 |
| ENSG000001 | ARRDC3 | 5.01E-01 | 7.93E-01 | 1.04E+00 | 5.90E-02 | 1.04E+00 |
| ENSG000001 | VAC14 | 5.01E-01 | 7.93E-01 | 1.03E+00 | 4.63E-02 | 1.03E+00 |
| ENSG00000C | MOCOS | 5.02E-01 | 7.93E-01 | 1.04E+00 | 5.21E-02 | 1.04E+00 |
| ENSG000001 | NELFB | 5.02E-01 | 7.93E-01 | 1.04E+00 | 6.15E-02 | 1.04E+00 |
| ENSG000002 | SRSF6P2 | 5.02E-01 | ? | 7.49E-01 | -4.18E-01 | -1.34E+00 |
| ENSG000002 | MROH3P | 5.02E-01 | ? | 7.63E-01 | -3.90E-01 | -1.31E+00 |
| ENSG000002 | --- | 5.02E-01 | ? | 1.20E+00 | 2.67E-01 | 1.20E+00 |
| ENSG000001 | SYTL3 | 5.02E-01 | 7.93E-01 | 1.08E+00 | 1.07E-01 | 1.08E+00 |
| ENSG000001 | FADS1 | 5.02E-01 | 7.93E-01 | 1.03E+00 | 4.15E-02 | 1.03E+00 |
| ENSG00000C | MEF2A | 5.02E-01 | 7.93E-01 | 9.65E-01 | -5.11E-02 | -1.04E+00 |
| ENSG000001 | LONRF2 | 5.02E-01 | ? | 1.19E+00 | 2.54E-01 | 1.19E+00 |
| ENSG000001 | LRIG2 | 5.02E-01 | 7.93E-01 | 9.72E-01 | -4.09E-02 | -1.03E+00 |
| ENSG000001 | ZNF862 | 5.02E-01 | 7.93E-01 | 1.04E+00 | 5.70E-02 | 1.04E+00 |
| ENSG000002 | ZNF781 | 5.02E-01 | ? | 1.28E+00 | 3.59E-01 | 1.28E+00 |
| ENSG000002 | --- | 5.02E-01 | ? | 1.38E+00 | 4.65E-01 | 1.38E+00 |
| ENSG000002 | EEF1B2P6 | 5.02E-01 | ? | 8.05E-01 | -3.14E-01 | -1.24E+00 |
| ENSG000001 | TSGA10 | 5.02E-01 | 7.93E-01 | 1.06E+00 | 7.88E-02 | 1.06E+00 |
| ENSG000002 | --- | 5.02E-01 | ? | 7.55E-01 | -4.06E-01 | -1.33E+00 |
| ENSG000001 | RGMB | 5.02E-01 | 7.93E-01 | 9.72E-01 | -4.14E-02 | -1.03E+00 |
| ENSG000001 | BNC2 | 5.02E-01 | ? | 1.33E+00 | 4.11E-01 | 1.33E+00 |
| ENSG000001 | COL4A6 | 5.02E-01 | 7.94E-01 | 1.04E+00 | 5.64E-02 | 1.04E+00 |
| ENSG000001 | CEACAM5 | 5.02E-01 | ? | 8.49E-01 | -2.36E-01 | -1.18E+00 |
| ENSG000001 | OXNAD1 | 5.02E-01 | 7.94E-01 | 1.04E+00 | 5.74E-02 | 1.04E+00 |
| ENSG000001 | CLMN | 5.03E-01 | 7.94E-01 | 1.03E+00 | 4.60E-02 | 1.03E+00 |
| ENSG000001 | PDE6D | 5.03E-01 | 7.94E-01 | 1.04E+00 | 5.62E-02 | 1.04E+00 |
| ENSG000002 | --- | 5.03E-01 | ? | 1.36E+00 | 4.45E-01 | 1.36E+00 |
| ENSG000001 | SPDYE12 | 5.03E-01 | ? | 1.17E+00 | 2.28E-01 | 1.17E+00 |
| ENSG000001 | ITGB1 | 5.03E-01 | 7.94E-01 | 9.69E-01 | -4.49E-02 | -1.03E+00 |
| ENSG000001 | UBE2I | 5.03E-01 | 7.94E-01 | 9.81E-01 | -2.78E-02 | -1.02E+00 |
| ENSG000002 | CRNDE | 5.03E-01 | 7.94E-01 | 9.48E-01 | -7.78E-02 | -1.06E+00 |
| ENSG00000C | GRAMD1B | 5.03E-01 | 7.94E-01 | 8.21E-01 | -2.84E-01 | -1.22E+00 |
| ENSG000002 | TRIM27 | 5.03E-01 | 7.94E-01 | 9.76E-01 | -3.48E-02 | -1.02E+00 |
| ENSG000001 | MFHAS1 | 5.03E-01 | 7.94E-01 | 1.03E+00 | 3.87E-02 | 1.03E+00 |
| ENSG000002 | POLH-AS1 | 5.03E-01 | ? | 1.17E+00 | 2.22E-01 | 1.17E+00 |
| ENSG000001 | ZNF346 | 5.03E-01 | 7.94E-01 | 1.03E+00 | 4.16E-02 | 1.03E+00 |
| ENSG000002 | UGDH-AS1 | 5.03E-01 | 7.94E-01 | 9.28E-01 | -1.08E-01 | -1.08E+00 |
| ENSG000002 | CDC42-AS1 | 5.03E-01 | ? | 1.17E+00 | 2.28E-01 | 1.17E+00 |
| ENSG000001 | KCP | 5.03E-01 | 7.94E-01 | 1.09E+00 | 1.30E-01 | 1.09E+00 |
| ENSG000001 | BTN3A3 | 5.03E-01 | 7.94E-01 | 1.04E+00 | 6.11E-02 | 1.04E+00 |
| ENSG000001 | CST6 | 5.03E-01 | 7.94E-01 | 1.14E+00 | 1.94E-01 | 1.14E+00 |
| ENSG000002 | --- | 5.04E-01 | ? | 8.10E-01 | -3.04E-01 | -1.23E+00 |
| ENSG000001 | PARD6B | 5.04E-01 | 7.94E-01 | 9.53E-01 | -7.00E-02 | -1.05E+00 |

| | | | | | | |
|---|---|---|---|---|---|---|
| ENSG000002 | --- | 5.04E-01 | ? | 8.98E-01 | -1.55E-01 | -1.11E+00 |
| ENSG000002 | --- | 5.04E-01 | ? | 1.19E+00 | 2.56E-01 | 1.19E+00 |
| ENSG000002 | PLAC4 | 5.04E-01 | ? | 1.43E+00 | 5.19E-01 | 1.43E+00 |
| ENSG000002 | --- | 5.04E-01 | ? | 6.56E-01 | -6.09E-01 | -1.53E+00 |
| ENSG000001 | PRKAG1 | 5.04E-01 | 7.95E-01 | 1.02E+00 | 2.60E-02 | 1.02E+00 |
| ENSG000002 | LBH | 5.04E-01 | 7.95E-01 | 9.58E-01 | -6.27E-02 | -1.04E+00 |
| ENSG00000C | CUL1 | 5.04E-01 | 7.95E-01 | 1.02E+00 | 3.15E-02 | 1.02E+00 |
| ENSG000001 | COTL1 | 5.04E-01 | 7.95E-01 | 1.02E+00 | 3.42E-02 | 1.02E+00 |
| ENSG000002 | MEG3 | 5.04E-01 | 7.95E-01 | 9.23E-01 | -1.16E-01 | -1.08E+00 |
| ENSG000002 | PKD2L2-DT | 5.04E-01 | ? | 1.16E+00 | 2.19E-01 | 1.16E+00 |
| ENSG000002 | LSM14A | 5.04E-01 | 7.95E-01 | 9.82E-01 | -2.65E-02 | -1.02E+00 |
| ENSG000001 | GNE | 5.04E-01 | 7.95E-01 | 1.03E+00 | 4.01E-02 | 1.03E+00 |
| ENSG00000C | MYNN | 5.05E-01 | 7.95E-01 | 9.63E-01 | -5.50E-02 | -1.04E+00 |
| ENSG000001 | MIDEAS | 5.05E-01 | 7.95E-01 | 1.03E+00 | 3.79E-02 | 1.03E+00 |
| ENSG000001 | DZIP1 | 5.05E-01 | 7.95E-01 | 1.05E+00 | 7.02E-02 | 1.05E+00 |
| ENSG000002 | CCNT2-AS1 | 5.05E-01 | 7.95E-01 | 1.06E+00 | 8.24E-02 | 1.06E+00 |
| ENSG000001 | POLN | 5.05E-01 | 7.95E-01 | 1.15E+00 | 1.98E-01 | 1.15E+00 |
| ENSG000001 | CEPT1 | 5.05E-01 | 7.95E-01 | 1.02E+00 | 2.80E-02 | 1.02E+00 |
| ENSG000002 | --- | 5.05E-01 | 7.95E-01 | 9.22E-01 | -1.18E-01 | -1.08E+00 |
| ENSG000002 | LIPE-AS1 | 5.05E-01 | ? | 1.16E+00 | 2.15E-01 | 1.16E+00 |
| ENSG000002 | PPP1CB-DT | 5.05E-01 | ? | 1.22E+00 | 2.82E-01 | 1.22E+00 |
| ENSG000001 | PFDN1 | 5.05E-01 | 7.95E-01 | 9.81E-01 | -2.76E-02 | -1.02E+00 |
| ENSG000001 | LEP | 5.05E-01 | ? | 1.40E+00 | 4.86E-01 | 1.40E+00 |
| ENSG000002 | ZNF709 | 5.05E-01 | 7.95E-01 | 1.11E+00 | 1.52E-01 | 1.11E+00 |
| ENSG000001 | SLC22A1 | 5.05E-01 | ? | 1.18E+00 | 2.38E-01 | 1.18E+00 |
| ENSG000001 | AUH | 5.05E-01 | 7.95E-01 | 9.44E-01 | -8.34E-02 | -1.06E+00 |
| ENSG000001 | DUSP5 | 5.05E-01 | 7.95E-01 | 1.03E+00 | 3.75E-02 | 1.03E+00 |
| ENSG000002 | --- | 5.05E-01 | ? | 1.15E+00 | 1.97E-01 | 1.15E+00 |
| ENSG000001 | --- | 5.06E-01 | ? | 1.35E+00 | 4.34E-01 | 1.35E+00 |
| ENSG000002 | XPC-AS1 | 5.06E-01 | ? | 8.74E-01 | -1.95E-01 | -1.14E+00 |
| ENSG000002 | --- | 5.06E-01 | ? | 1.28E+00 | 3.56E-01 | 1.28E+00 |
| ENSG000002 | --- | 5.06E-01 | 7.96E-01 | 1.08E+00 | 1.14E-01 | 1.08E+00 |
| ENSG000002 | POLG2 | 5.06E-01 | 7.96E-01 | 9.53E-01 | -6.99E-02 | -1.05E+00 |
| ENSG000001 | SRP9P1 | 5.06E-01 | ? | 1.21E+00 | 2.79E-01 | 1.21E+00 |
| ENSG000001 | INSIG2 | 5.06E-01 | 7.96E-01 | 9.76E-01 | -3.54E-02 | -1.02E+00 |
| ENSG000002 | PROSER2-AS | 5.06E-01 | ? | 8.28E-01 | -2.72E-01 | -1.21E+00 |
| ENSG000002 | --- | 5.06E-01 | 7.96E-01 | 1.10E+00 | 1.41E-01 | 1.10E+00 |
| ENSG000001 | KCNA7 | 5.06E-01 | ? | 1.12E+00 | 1.62E-01 | 1.12E+00 |
| ENSG000001 | ST3GAL4 | 5.06E-01 | 7.96E-01 | 1.04E+00 | 5.45E-02 | 1.04E+00 |
| ENSG000002 | CCDC28A-AS | 5.06E-01 | ? | 8.68E-01 | -2.04E-01 | -1.15E+00 |
| ENSG00000C | DBF4 | 5.06E-01 | 7.96E-01 | 9.61E-01 | -5.71E-02 | -1.04E+00 |
| ENSG000002 | HOMER3-AS | 5.06E-01 | ? | 1.11E+00 | 1.45E-01 | 1.11E+00 |
| ENSG000001 | FUCA1 | 5.06E-01 | 7.96E-01 | 1.05E+00 | 6.78E-02 | 1.05E+00 |
| ENSG000001 | TEC | 5.06E-01 | 7.96E-01 | 9.07E-01 | -1.40E-01 | -1.10E+00 |
| ENSG000001 | NKAIN4 | 5.06E-01 | ? | 7.18E-01 | -4.77E-01 | -1.39E+00 |

| | | | | | | | |
|---|---|---|---|---|---|---|---|
| ENSG000001 | ZKSCAN7 | 5.06E-01 | | 7.96E-01 | 1.06E+00 | 8.15E-02 | 1.06E+00 |
| ENSG000002 | --- | 5.06E-01 | ? | | 1.25E+00 | 3.19E-01 | 1.25E+00 |
| ENSG000000 | PYGM | 5.06E-01 | ? | | 7.80E-01 | -3.58E-01 | -1.28E+00 |
| ENSG000001 | CYP4F11 | 5.06E-01 | ? | | 8.73E-01 | -1.95E-01 | -1.15E+00 |
| ENSG000001 | THOC2 | 5.06E-01 | | 7.96E-01 | 9.74E-01 | -3.77E-02 | -1.03E+00 |
| ENSG000001 | RPUSD2 | 5.07E-01 | | 7.96E-01 | 1.04E+00 | 5.22E-02 | 1.04E+00 |
| ENSG000002 | --- | 5.07E-01 | ? | | 8.51E-01 | -2.32E-01 | -1.17E+00 |
| ENSG000001 | GABPA | 5.07E-01 | | 7.96E-01 | 1.03E+00 | 4.32E-02 | 1.03E+00 |
| ENSG000002 | --- | 5.07E-01 | ? | | 1.16E+00 | 2.20E-01 | 1.16E+00 |
| ENSG000001 | PPBP | 5.07E-01 | | 7.96E-01 | 7.98E-01 | -3.26E-01 | -1.25E+00 |
| ENSG000002 | PDXDC2P | 5.07E-01 | ? | | 1.68E+00 | 7.46E-01 | 1.68E+00 |
| ENSG000001 | MIRLET7BHG | 5.07E-01 | | 7.96E-01 | 9.39E-01 | -9.01E-02 | -1.06E+00 |
| ENSG000001 | TBC1D17 | 5.07E-01 | | 7.96E-01 | 1.04E+00 | 5.15E-02 | 1.04E+00 |
| ENSG000002 | --- | 5.07E-01 | ? | | 1.48E+00 | 5.62E-01 | 1.48E+00 |
| ENSG000001 | EIF2S2 | 5.07E-01 | | 7.96E-01 | 9.84E-01 | -2.36E-02 | -1.02E+00 |
| ENSG000001 | CYB5B | 5.07E-01 | | 7.96E-01 | 1.02E+00 | 2.64E-02 | 1.02E+00 |
| ENSG000002 | --- | 5.07E-01 | | 7.96E-01 | 9.25E-01 | -1.12E-01 | -1.08E+00 |
| ENSG000000 | C1QTNF3 | 5.07E-01 | ? | | 1.25E+00 | 3.22E-01 | 1.25E+00 |
| ENSG000001 | ALG10 | 5.07E-01 | | 7.96E-01 | 1.04E+00 | 6.02E-02 | 1.04E+00 |
| ENSG000002 | BMS1P14 | 5.07E-01 | ? | | 7.51E-01 | -4.13E-01 | -1.33E+00 |
| ENSG000002 | DPRXP4 | 5.07E-01 | ? | | 7.21E-01 | -4.72E-01 | -1.39E+00 |
| ENSG000001 | IL20 | 5.07E-01 | ? | | 1.28E+00 | 3.60E-01 | 1.28E+00 |
| ENSG000002 | --- | 5.07E-01 | ? | | 1.13E+00 | 1.77E-01 | 1.13E+00 |
| ENSG000001 | PCBD2 | 5.07E-01 | | 7.97E-01 | 9.67E-01 | -4.84E-02 | -1.03E+00 |
| ENSG000002 | ZNF891 | 5.07E-01 | | 7.97E-01 | 9.48E-01 | -7.73E-02 | -1.06E+00 |
| ENSG000002 | RNF5P1 | 5.07E-01 | ? | | 7.53E-01 | -4.10E-01 | -1.33E+00 |
| ENSG000002 | --- | 5.08E-01 | ? | | 1.21E+00 | 2.69E-01 | 1.21E+00 |
| ENSG000002 | TOMM40P4 | 5.08E-01 | | 7.97E-01 | 1.06E+00 | 8.88E-02 | 1.06E+00 |
| ENSG000002 | --- | 5.08E-01 | ? | | 7.66E-01 | -3.84E-01 | -1.30E+00 |
| ENSG000001 | DYM | 5.08E-01 | | 7.97E-01 | 1.02E+00 | 2.83E-02 | 1.02E+00 |
| ENSG000001 | CCDC158 | 5.08E-01 | ? | | 8.09E-01 | -3.05E-01 | -1.24E+00 |
| ENSG000002 | --- | 5.08E-01 | | 7.97E-01 | 1.16E+00 | 2.11E-01 | 1.16E+00 |
| ENSG000001 | TMEM68 | 5.08E-01 | | 7.97E-01 | 1.02E+00 | 3.53E-02 | 1.02E+00 |
| ENSG000002 | GOLGA8T | 5.08E-01 | ? | | 8.64E-01 | -2.12E-01 | -1.16E+00 |
| ENSG000001 | HEYL | 5.08E-01 | ? | | 1.22E+00 | 2.89E-01 | 1.22E+00 |
| ENSG000001 | EAF2 | 5.08E-01 | ? | | 8.93E-01 | -1.64E-01 | -1.12E+00 |
| ENSG000002 | PNMA6A | 5.08E-01 | ? | | 1.29E+00 | 3.65E-01 | 1.29E+00 |
| ENSG000001 | LINC01547 | 5.08E-01 | | 7.98E-01 | 1.10E+00 | 1.34E-01 | 1.10E+00 |
| ENSG000002 | --- | 5.08E-01 | ? | | 7.23E-01 | -4.68E-01 | -1.38E+00 |
| ENSG000001 | ZNF799 | 5.08E-01 | | 7.98E-01 | 1.05E+00 | 7.52E-02 | 1.05E+00 |
| ENSG000002 | --- | 5.08E-01 | ? | | 8.10E-01 | -3.03E-01 | -1.23E+00 |
| ENSG000001 | CLEC2B | 5.08E-01 | | 7.98E-01 | 9.36E-01 | -9.47E-02 | -1.07E+00 |
| ENSG000002 | PCDHA11 | 5.09E-01 | | 7.98E-01 | 9.15E-01 | -1.28E-01 | -1.09E+00 |
| ENSG000001 | IRAG2 | 5.09E-01 | | 7.98E-01 | 1.07E+00 | 9.26E-02 | 1.07E+00 |
| ENSG000002 | CAP1P1 | 5.09E-01 | ? | | 8.67E-01 | -2.06E-01 | -1.15E+00 |

| | | | | | | |
|---|---|---|---|---|---|---|
| ENSG000001 | MTA1 | 5.09E-01 | 7.98E-01 | 1.04E+00 | 5.00E-02 | 1.04E+00 |
| ENSG000001 | MB21D2 | 5.09E-01 | 7.98E-01 | 1.03E+00 | 4.91E-02 | 1.03E+00 |
| ENSG000002 | --- | 5.09E-01 | ? | 1.20E+00 | 2.66E-01 | 1.20E+00 |
| ENSG000002 | CRYZL1 | 5.09E-01 | 7.98E-01 | 9.70E-01 | -4.40E-02 | -1.03E+00 |
| ENSG000001 | ARHGAP39 | 5.09E-01 | 7.98E-01 | 9.40E-01 | -8.89E-02 | -1.06E+00 |
| ENSG000001 | ORC2 | 5.09E-01 | 7.98E-01 | 1.03E+00 | 3.58E-02 | 1.03E+00 |
| ENSG000002 | --- | 5.09E-01 | 7.98E-01 | 1.17E+00 | 2.32E-01 | 1.17E+00 |
| ENSG000002 | --- | 5.09E-01 | ? | 1.17E+00 | 2.23E-01 | 1.17E+00 |
| ENSG000001 | C5orf15 | 5.09E-01 | 7.98E-01 | 1.02E+00 | 3.36E-02 | 1.02E+00 |
| ENSG000001 | TCF7L1 | 5.09E-01 | 7.98E-01 | 1.02E+00 | 3.17E-02 | 1.02E+00 |
| ENSG000001 | E4F1 | 5.09E-01 | 7.98E-01 | 1.05E+00 | 6.87E-02 | 1.05E+00 |
| ENSG000002 | RPL7AP43 | 5.09E-01 | ? | 8.09E-01 | -3.06E-01 | -1.24E+00 |
| ENSG000001 | SLC13A5 | 5.09E-01 | ? | 1.27E+00 | 3.45E-01 | 1.27E+00 |
| ENSG000002 | --- | 5.09E-01 | ? | 7.19E-01 | -4.76E-01 | -1.39E+00 |
| ENSG00000C | BCAS1 | 5.09E-01 | ? | 1.15E+00 | 1.96E-01 | 1.15E+00 |
| ENSG000001 | ZNF813 | 5.09E-01 | 7.98E-01 | 9.69E-01 | -4.58E-02 | -1.03E+00 |
| ENSG000001 | NPM2 | 5.09E-01 | ? | 8.19E-01 | -2.89E-01 | -1.22E+00 |
| ENSG000001 | PKHD1 | 5.09E-01 | ? | 8.16E-01 | -2.93E-01 | -1.22E+00 |
| ENSG000001 | HERC5 | 5.10E-01 | ? | 1.16E+00 | 2.08E-01 | 1.16E+00 |
| ENSG000001 | FGF17 | 5.10E-01 | ? | 1.34E+00 | 4.24E-01 | 1.34E+00 |
| ENSG000001 | GMFB | 5.10E-01 | 7.99E-01 | 1.02E+00 | 2.65E-02 | 1.02E+00 |
| ENSG00000C | ALOX5 | 5.10E-01 | ? | 8.90E-01 | -1.68E-01 | -1.12E+00 |
| ENSG000002 | ZBTB10 | 5.10E-01 | 7.99E-01 | 1.03E+00 | 4.35E-02 | 1.03E+00 |
| ENSG000002 | UBXN2B | 5.10E-01 | 7.99E-01 | 9.73E-01 | -4.01E-02 | -1.03E+00 |
| ENSG000001 | DMWD | 5.10E-01 | 7.99E-01 | 1.03E+00 | 4.72E-02 | 1.03E+00 |
| ENSG000001 | MLST8 | 5.10E-01 | 7.99E-01 | 1.03E+00 | 4.68E-02 | 1.03E+00 |
| ENSG000002 | RPL21P75 | 5.10E-01 | 7.99E-01 | 1.29E+00 | 3.67E-01 | 1.29E+00 |
| ENSG000001 | CDKN2A | 5.10E-01 | 7.99E-01 | 1.05E+00 | 6.76E-02 | 1.05E+00 |
| ENSG000001 | ESR2 | 5.10E-01 | ? | 1.15E+00 | 2.05E-01 | 1.15E+00 |
| ENSG000001 | SH2B1 | 5.10E-01 | 7.99E-01 | 9.64E-01 | -5.35E-02 | -1.04E+00 |
| ENSG000001 | MFSD2A | 5.10E-01 | 7.99E-01 | 1.04E+00 | 5.25E-02 | 1.04E+00 |
| ENSG000001 | DLX2 | 5.10E-01 | ? | 8.17E-01 | -2.91E-01 | -1.22E+00 |
| ENSG000001 | PIPSL | 5.11E-01 | ? | 9.05E-01 | -1.45E-01 | -1.11E+00 |
| ENSG000002 | --- | 5.11E-01 | ? | 1.21E+00 | 2.72E-01 | 1.21E+00 |
| ENSG000002 | --- | 5.11E-01 | ? | 1.45E+00 | 5.33E-01 | 1.45E+00 |
| ENSG000001 | F8 | 5.11E-01 | 7.99E-01 | 9.49E-01 | -7.55E-02 | -1.05E+00 |
| ENSG000001 | LYNX1 | 5.11E-01 | 7.99E-01 | 1.09E+00 | 1.23E-01 | 1.09E+00 |
| ENSG000002 | RPL24P8 | 5.11E-01 | ? | 1.28E+00 | 3.57E-01 | 1.28E+00 |
| ENSG000001 | ZNF669 | 5.11E-01 | 7.99E-01 | 9.70E-01 | -4.38E-02 | -1.03E+00 |
| ENSG000002 | FKBPL | 5.11E-01 | 7.99E-01 | 1.08E+00 | 1.09E-01 | 1.08E+00 |
| ENSG000002 | LAT | 5.11E-01 | 7.99E-01 | 9.28E-01 | -1.08E-01 | -1.08E+00 |
| ENSG000001 | PSPH | 5.11E-01 | 7.99E-01 | 1.03E+00 | 3.95E-02 | 1.03E+00 |
| ENSG000001 | IL2RB | 5.11E-01 | 7.99E-01 | 1.09E+00 | 1.22E-01 | 1.09E+00 |
| ENSG000001 | TEAD4 | 5.11E-01 | 8.00E-01 | 9.55E-01 | -6.70E-02 | -1.05E+00 |
| ENSG00000C | SEL1L3 | 5.11E-01 | 8.00E-01 | 1.02E+00 | 3.31E-02 | 1.02E+00 |

| | | | | | | |
|---|---|---|---|---|---|---|
| ENSG000001 | SARS2 | 5.11E-01 | 8.00E-01 | 1.05E+00 | 6.65E-02 | 1.05E+00 |
| ENSG000001 | DNMBP | 5.12E-01 | 8.00E-01 | 9.77E-01 | -3.34E-02 | -1.02E+00 |
| ENSG000001 | RPL26 | 5.12E-01 | 8.00E-01 | 1.02E+00 | 3.32E-02 | 1.02E+00 |
| ENSG000001 | ATG3 | 5.12E-01 | 8.00E-01 | 9.70E-01 | -4.46E-02 | -1.03E+00 |
| ENSG000002 | PIWIL2-DT | 5.12E-01 | ? | 8.37E-01 | -2.56E-01 | -1.19E+00 |
| ENSG000002 | UBE2MP1 | 5.12E-01 | ? | 1.24E+00 | 3.06E-01 | 1.24E+00 |
| ENSG000002 | --- | 5.12E-01 | ? | 7.24E-01 | -4.66E-01 | -1.38E+00 |
| ENSG000002 | --- | 5.12E-01 | ? | 8.18E-01 | -2.90E-01 | -1.22E+00 |
| ENSG000001 | TMPRSS11B | 5.12E-01 | ? | 7.32E-01 | -4.51E-01 | -1.37E+00 |
| ENSG000002 | RAB1C | 5.12E-01 | ? | 1.47E+00 | 5.57E-01 | 1.47E+00 |
| ENSG000002 | --- | 5.12E-01 | ? | 7.14E-01 | -4.87E-01 | -1.40E+00 |
| ENSG000001 | FGFRL1 | 5.12E-01 | 8.00E-01 | 9.50E-01 | -7.43E-02 | -1.05E+00 |
| ENSG000001 | CYP2J2 | 5.12E-01 | 8.00E-01 | 9.38E-01 | -9.18E-02 | -1.07E+00 |
| ENSG000002 | --- | 5.12E-01 | 8.00E-01 | 8.26E-01 | -2.77E-01 | -1.21E+00 |
| ENSG000001 | PRKD2 | 5.12E-01 | 8.00E-01 | 1.03E+00 | 4.06E-02 | 1.03E+00 |
| ENSG000001 | METTL8 | 5.12E-01 | 8.00E-01 | 1.02E+00 | 2.89E-02 | 1.02E+00 |
| ENSG000002 | --- | 5.12E-01 | 8.00E-01 | 1.09E+00 | 1.24E-01 | 1.09E+00 |
| ENSG00000C | VCL | 5.12E-01 | 8.00E-01 | 1.03E+00 | 3.82E-02 | 1.03E+00 |
| ENSG000001 | KIAA1958 | 5.13E-01 | 8.00E-01 | 9.49E-01 | -7.60E-02 | -1.05E+00 |
| ENSG000002 | ODC1-DT | 5.13E-01 | ? | 8.95E-01 | -1.59E-01 | -1.12E+00 |
| ENSG000002 | ZNF670 | 5.13E-01 | 8.00E-01 | 9.50E-01 | -7.35E-02 | -1.05E+00 |
| ENSG000002 | --- | 5.13E-01 | ? | 1.23E+00 | 3.00E-01 | 1.23E+00 |
| ENSG000002 | --- | 5.13E-01 | ? | 7.17E-01 | -4.81E-01 | -1.40E+00 |
| ENSG000002 | --- | 5.13E-01 | ? | 8.41E-01 | -2.50E-01 | -1.19E+00 |
| ENSG000002 | HLA-DPA1 | 5.13E-01 | ? | 7.23E-01 | -4.68E-01 | -1.38E+00 |
| ENSG00000C | RCOR1 | 5.13E-01 | 8.00E-01 | 9.82E-01 | -2.56E-02 | -1.02E+00 |
| ENSG000001 | MAGEE1 | 5.13E-01 | 8.00E-01 | 1.04E+00 | 5.85E-02 | 1.04E+00 |
| ENSG000001 | RBM4B | 5.13E-01 | 8.00E-01 | 1.03E+00 | 3.91E-02 | 1.03E+00 |
| ENSG000001 | KRCC1 | 5.13E-01 | 8.00E-01 | 9.69E-01 | -4.49E-02 | -1.03E+00 |
| ENSG000001 | FOXN2 | 5.13E-01 | 8.00E-01 | 9.67E-01 | -4.87E-02 | -1.03E+00 |
| ENSG00000C | DOCK9 | 5.13E-01 | 8.00E-01 | 1.03E+00 | 4.65E-02 | 1.03E+00 |
| ENSG000001 | DNMT1 | 5.13E-01 | 8.00E-01 | 9.66E-01 | -4.93E-02 | -1.03E+00 |
| ENSG000001 | ALOX12B | 5.13E-01 | ? | 6.94E-01 | -5.28E-01 | -1.44E+00 |
| ENSG000002 | --- | 5.13E-01 | 8.00E-01 | 1.07E+00 | 1.00E-01 | 1.07E+00 |
| ENSG00000C | IGF2BP2 | 5.13E-01 | 8.00E-01 | 1.03E+00 | 3.61E-02 | 1.03E+00 |
| ENSG000002 | --- | 5.13E-01 | 8.00E-01 | 9.11E-01 | -1.35E-01 | -1.10E+00 |
| ENSG000002 | RNASEK | 5.13E-01 | 8.00E-01 | 9.74E-01 | -3.82E-02 | -1.03E+00 |
| ENSG000001 | COQ4 | 5.13E-01 | 8.00E-01 | 9.52E-01 | -7.04E-02 | -1.05E+00 |
| ENSG000002 | LINC00511 | 5.13E-01 | 8.00E-01 | 9.71E-01 | -4.19E-02 | -1.03E+00 |
| ENSG00000C | ERP29 | 5.13E-01 | 8.00E-01 | 1.02E+00 | 2.83E-02 | 1.02E+00 |
| ENSG00000C | TMCC3 | 5.13E-01 | 8.00E-01 | 9.68E-01 | -4.62E-02 | -1.03E+00 |
| ENSG000001 | ASRGL1 | 5.13E-01 | ? | 7.98E-01 | -3.26E-01 | -1.25E+00 |
| ENSG000001 | HPX | 5.13E-01 | 8.00E-01 | 1.11E+00 | 1.54E-01 | 1.11E+00 |
| ENSG000002 | --- | 5.13E-01 | ? | 1.24E+00 | 3.11E-01 | 1.24E+00 |
| ENSG00000C | PNKP | 5.13E-01 | 8.00E-01 | 1.04E+00 | 5.03E-02 | 1.04E+00 |

| | | | | | | |
|---|---|---|---|---|---|---|
| ENSG000001 | SKP1 | 5.13E-01 | 8.00E-01 | 9.75E-01 | -3.72E-02 | -1.03E+00 |
| ENSG000002 | --- | 5.13E-01 | ? | 1.15E+00 | 2.03E-01 | 1.15E+00 |
| ENSG000001 | RNF32 | 5.13E-01 | 8.00E-01 | 1.10E+00 | 1.40E-01 | 1.10E+00 |
| ENSG000001 | PPP6C | 5.13E-01 | 8.00E-01 | 9.79E-01 | -3.08E-02 | -1.02E+00 |
| ENSG000002 | --- | 5.13E-01 | ? | 1.12E+00 | 1.64E-01 | 1.12E+00 |
| ENSG00000C | GRAMD1A | 5.14E-01 | 8.00E-01 | 1.03E+00 | 4.87E-02 | 1.03E+00 |
| ENSG000001 | ADGRL2 | 5.14E-01 | 8.00E-01 | 1.03E+00 | 3.65E-02 | 1.03E+00 |
| ENSG000001 | BBS4 | 5.14E-01 | 8.00E-01 | 9.66E-01 | -4.97E-02 | -1.04E+00 |
| ENSG00000C | MCM10 | 5.14E-01 | 8.00E-01 | 8.88E-01 | -1.71E-01 | -1.13E+00 |
| ENSG000002 | RPL17P36 | 5.14E-01 | ? | 6.91E-01 | -5.33E-01 | -1.45E+00 |
| ENSG00000C | PNPLA6 | 5.14E-01 | 8.00E-01 | 1.04E+00 | 5.93E-02 | 1.04E+00 |
| ENSG000001 | STX4 | 5.14E-01 | 8.00E-01 | 9.76E-01 | -3.48E-02 | -1.02E+00 |
| ENSG00000C | RSF1 | 5.14E-01 | 8.00E-01 | 9.68E-01 | -4.70E-02 | -1.03E+00 |
| ENSG000001 | CDKN2B | 5.14E-01 | 8.00E-01 | 1.04E+00 | 5.83E-02 | 1.04E+00 |
| ENSG000002 | --- | 5.14E-01 | ? | 1.53E+00 | 6.16E-01 | 1.53E+00 |
| ENSG000001 | HYDIN | 5.14E-01 | 8.00E-01 | 1.12E+00 | 1.60E-01 | 1.12E+00 |
| ENSG000002 | STK38L | 5.14E-01 | 8.00E-01 | 1.03E+00 | 3.78E-02 | 1.03E+00 |
| ENSG000001 | ARL4D | 5.14E-01 | 8.00E-01 | 9.53E-01 | -6.89E-02 | -1.05E+00 |
| ENSG000001 | LINC00518 | 5.14E-01 | ? | 1.24E+00 | 3.13E-01 | 1.24E+00 |
| ENSG000001 | GRIK5 | 5.14E-01 | ? | 1.17E+00 | 2.27E-01 | 1.17E+00 |
| ENSG000002 | TRAM2-AS1 | 5.14E-01 | 8.01E-01 | 1.06E+00 | 8.73E-02 | 1.06E+00 |
| ENSG000001 | MGP | 5.14E-01 | ? | 7.50E-01 | -4.16E-01 | -1.33E+00 |
| ENSG000002 | TMSB4X | 5.15E-01 | 8.01E-01 | 9.79E-01 | -3.04E-02 | -1.02E+00 |
| ENSG000002 | SERF1B | 5.15E-01 | 8.01E-01 | 1.03E+00 | 4.54E-02 | 1.03E+00 |
| ENSG000001 | LINC00471 | 5.15E-01 | ? | 8.11E-01 | -3.03E-01 | -1.23E+00 |
| ENSG000001 | WASF4P | 5.15E-01 | ? | 8.20E-01 | -2.86E-01 | -1.22E+00 |
| ENSG000002 | EPHA1-AS1 | 5.15E-01 | 8.01E-01 | 1.06E+00 | 8.97E-02 | 1.06E+00 |
| ENSG000002 | --- | 5.15E-01 | ? | 1.36E+00 | 4.43E-01 | 1.36E+00 |
| ENSG000001 | H1-4 | 5.15E-01 | ? | 8.18E-01 | -2.90E-01 | -1.22E+00 |
| ENSG000002 | SYPL1P2 | 5.15E-01 | ? | 1.33E+00 | 4.12E-01 | 1.33E+00 |
| ENSG000001 | ERI2 | 5.15E-01 | 8.01E-01 | 1.03E+00 | 3.61E-02 | 1.03E+00 |
| ENSG000001 | SLC25A29 | 5.15E-01 | 8.01E-01 | 1.05E+00 | 7.61E-02 | 1.05E+00 |
| ENSG000001 | MED22 | 5.15E-01 | 8.01E-01 | 1.03E+00 | 4.50E-02 | 1.03E+00 |
| ENSG000002 | NBPF26 | 5.15E-01 | 8.01E-01 | 9.18E-01 | -1.23E-01 | -1.09E+00 |
| ENSG000002 | --- | 5.15E-01 | 8.01E-01 | 9.58E-01 | -6.23E-02 | -1.04E+00 |
| ENSG000002 | --- | 5.15E-01 | ? | 1.22E+00 | 2.92E-01 | 1.22E+00 |
| ENSG000001 | CLP1 | 5.15E-01 | 8.01E-01 | 1.03E+00 | 4.32E-02 | 1.03E+00 |
| ENSG000001 | CYRIB | 5.16E-01 | 8.01E-01 | 9.75E-01 | -3.59E-02 | -1.03E+00 |
| ENSG000001 | FAM83H | 5.16E-01 | 8.01E-01 | 1.04E+00 | 5.24E-02 | 1.04E+00 |
| ENSG000002 | --- | 5.16E-01 | ? | 8.64E-01 | -2.10E-01 | -1.16E+00 |
| ENSG000002 | AATF | 5.16E-01 | 8.01E-01 | 9.80E-01 | -2.97E-02 | -1.02E+00 |
| ENSG000002 | HNRNPA1P4! | 5.16E-01 | 8.02E-01 | 9.00E-01 | -1.52E-01 | -1.11E+00 |
| ENSG000001 | ICAM5 | 5.16E-01 | 8.02E-01 | 9.20E-01 | -1.20E-01 | -1.09E+00 |
| ENSG000002 | LINC02606 | 5.16E-01 | ? | 1.25E+00 | 3.27E-01 | 1.25E+00 |
| ENSG00000C | OSBPL5 | 5.16E-01 | 8.02E-01 | 1.04E+00 | 5.48E-02 | 1.04E+00 |

| | | | | | | |
|---|---|---|---|---|---|---|
| ENSG000002 | C10orf95-AS | 5.16E-01 | 8.02E-01 | 1.06E+00 | 8.45E-02 | 1.06E+00 |
| ENSG000002 | LINC01271 | 5.16E-01 | ? | 8.67E-01 | -2.06E-01 | -1.15E+00 |
| ENSG000001 | ZDHHC8BP | 5.16E-01 | ? | 8.31E-01 | -2.68E-01 | -1.20E+00 |
| ENSG000001 | SPTLC2 | 5.16E-01 | 8.02E-01 | 1.03E+00 | 4.29E-02 | 1.03E+00 |
| ENSG000002 | PRPS1P2 | 5.16E-01 | ? | 8.80E-01 | -1.84E-01 | -1.14E+00 |
| ENSG000001 | IL17RD | 5.16E-01 | 8.02E-01 | 1.04E+00 | 5.30E-02 | 1.04E+00 |
| ENSG000001 | JAK1 | 5.16E-01 | 8.02E-01 | 1.02E+00 | 2.56E-02 | 1.02E+00 |
| ENSG00000C | MXD1 | 5.16E-01 | 8.02E-01 | 9.63E-01 | -5.40E-02 | -1.04E+00 |
| ENSG000002 | ERICH2-DT | 5.17E-01 | ? | 1.33E+00 | 4.12E-01 | 1.33E+00 |
| ENSG000001 | PHEX | 5.17E-01 | 8.02E-01 | 1.03E+00 | 4.56E-02 | 1.03E+00 |
| ENSG000002 | --- | 5.17E-01 | 8.02E-01 | 8.34E-01 | -2.63E-01 | -1.20E+00 |
| ENSG000002 | --- | 5.17E-01 | ? | 1.48E+00 | 5.69E-01 | 1.48E+00 |
| ENSG000001 | FAM219B | 5.17E-01 | 8.02E-01 | 1.03E+00 | 4.22E-02 | 1.03E+00 |
| ENSG000001 | ACSL5 | 5.17E-01 | ? | 1.11E+00 | 1.48E-01 | 1.11E+00 |
| ENSG000002 | FRMD6-AS2 | 5.17E-01 | 8.02E-01 | 1.07E+00 | 1.02E-01 | 1.07E+00 |
| ENSG000001 | RABGGTA | 5.17E-01 | 8.02E-01 | 1.04E+00 | 5.37E-02 | 1.04E+00 |
| ENSG000001 | LTBP3 | 5.17E-01 | 8.02E-01 | 1.04E+00 | 6.19E-02 | 1.04E+00 |
| ENSG000001 | QPRT | 5.17E-01 | ? | 8.36E-01 | -2.59E-01 | -1.20E+00 |
| ENSG000001 | OAS2 | 5.17E-01 | 8.02E-01 | 1.03E+00 | 4.51E-02 | 1.03E+00 |
| ENSG000002 | --- | 5.17E-01 | ? | 1.41E+00 | 4.95E-01 | 1.41E+00 |
| ENSG00000C | ZNF800 | 5.17E-01 | 8.02E-01 | 9.75E-01 | -3.70E-02 | -1.03E+00 |
| ENSG000001 | CTC1 | 5.17E-01 | 8.02E-01 | 1.03E+00 | 4.66E-02 | 1.03E+00 |
| ENSG000002 | --- | 5.17E-01 | 8.02E-01 | 1.05E+00 | 7.70E-02 | 1.05E+00 |
| ENSG000001 | ZBTB42 | 5.17E-01 | 8.02E-01 | 9.26E-01 | -1.11E-01 | -1.08E+00 |
| ENSG000001 | DEPTOR | 5.18E-01 | ? | 1.14E+00 | 1.84E-01 | 1.14E+00 |
| ENSG000002 | --- | 5.18E-01 | ? | 7.60E-01 | -3.97E-01 | -1.32E+00 |
| ENSG000001 | LCA5L | 5.18E-01 | 8.03E-01 | 9.27E-01 | -1.09E-01 | -1.08E+00 |
| ENSG000002 | DPY19L3-DT | 5.18E-01 | ? | 1.36E+00 | 4.42E-01 | 1.36E+00 |
| ENSG000002 | --- | 5.18E-01 | ? | 1.42E+00 | 5.01E-01 | 1.42E+00 |
| ENSG000001 | DPAGT1 | 5.18E-01 | 8.03E-01 | 9.77E-01 | -3.36E-02 | -1.02E+00 |
| ENSG000001 | TIMM29 | 5.18E-01 | 8.03E-01 | 1.05E+00 | 6.52E-02 | 1.05E+00 |
| ENSG000002 | --- | 5.18E-01 | ? | 8.90E-01 | -1.68E-01 | -1.12E+00 |
| ENSG000002 | --- | 5.18E-01 | ? | 8.36E-01 | -2.59E-01 | -1.20E+00 |
| ENSG000001 | CDK18 | 5.18E-01 | 8.03E-01 | 1.03E+00 | 4.07E-02 | 1.03E+00 |
| ENSG000001 | TMEM126A | 5.18E-01 | 8.03E-01 | 1.04E+00 | 5.08E-02 | 1.04E+00 |
| ENSG000001 | ANKRD61 | 5.18E-01 | ? | 8.40E-01 | -2.51E-01 | -1.19E+00 |
| ENSG000002 | --- | 5.18E-01 | ? | 8.89E-01 | -1.70E-01 | -1.13E+00 |
| ENSG000001 | SLAIN2 | 5.18E-01 | 8.03E-01 | 1.03E+00 | 4.31E-02 | 1.03E+00 |
| ENSG000002 | --- | 5.18E-01 | ? | 1.26E+00 | 3.37E-01 | 1.26E+00 |
| ENSG000001 | SDC4 | 5.18E-01 | 8.03E-01 | 1.03E+00 | 3.90E-02 | 1.03E+00 |
| ENSG000001 | RETREG3 | 5.18E-01 | 8.03E-01 | 1.03E+00 | 4.31E-02 | 1.03E+00 |
| ENSG000001 | KLC4 | 5.18E-01 | 8.03E-01 | 1.04E+00 | 5.70E-02 | 1.04E+00 |
| ENSG000002 | NDUFS3 | 5.18E-01 | 8.03E-01 | 1.04E+00 | 5.47E-02 | 1.04E+00 |
| ENSG000001 | IPO4 | 5.18E-01 | 8.03E-01 | 1.03E+00 | 3.90E-02 | 1.03E+00 |
| ENSG000001 | ICOSLG | 5.18E-01 | 8.03E-01 | 1.07E+00 | 9.62E-02 | 1.07E+00 |

| | | | | | | |
|---|---|---|---|---|---|---|
| ENSG000002 | RPL23AP47 | 5.18E-01 | ? | 6.36E-01 | -6.54E-01 | -1.57E+00 |
| ENSG00000C | LAMB4 | 5.19E-01 | ? | 8.52E-01 | -2.32E-01 | -1.17E+00 |
| ENSG000002 | --- | 5.19E-01 | ? | 1.43E+00 | 5.19E-01 | 1.43E+00 |
| ENSG000001 | --- | 5.19E-01 | 8.03E-01 | 9.31E-01 | -1.03E-01 | -1.07E+00 |
| ENSG000001 | CIITA | 5.19E-01 | 8.03E-01 | 1.08E+00 | 1.07E-01 | 1.08E+00 |
| ENSG000002 | --- | 5.19E-01 | 8.03E-01 | 1.10E+00 | 1.35E-01 | 1.10E+00 |
| ENSG000001 | ACOT13 | 5.19E-01 | 8.03E-01 | 1.03E+00 | 3.58E-02 | 1.03E+00 |
| ENSG000002 | --- | 5.19E-01 | ? | 1.16E+00 | 2.14E-01 | 1.16E+00 |
| ENSG000001 | INCA1 | 5.19E-01 | ? | 1.14E+00 | 1.89E-01 | 1.14E+00 |
| ENSG000001 | PAQR4 | 5.19E-01 | 8.03E-01 | 1.04E+00 | 5.61E-02 | 1.04E+00 |
| ENSG000001 | PICK1 | 5.19E-01 | 8.03E-01 | 9.72E-01 | -4.03E-02 | -1.03E+00 |
| ENSG000002 | --- | 5.19E-01 | ? | 8.17E-01 | -2.91E-01 | -1.22E+00 |
| ENSG00000C | RHBDF1 | 5.19E-01 | 8.03E-01 | 9.70E-01 | -4.42E-02 | -1.03E+00 |
| ENSG000001 | GADD45G | 5.19E-01 | ? | 7.32E-01 | -4.50E-01 | -1.37E+00 |
| ENSG000001 | SLC4A3 | 5.19E-01 | 8.03E-01 | 9.59E-01 | -6.05E-02 | -1.04E+00 |
| ENSG000001 | RIBC1 | 5.19E-01 | ? | 1.23E+00 | 3.00E-01 | 1.23E+00 |
| ENSG000002 | TNFRSF10A- | 5.19E-01 | 8.03E-01 | 1.05E+00 | 7.08E-02 | 1.05E+00 |
| ENSG000002 | PET100 | 5.19E-01 | 8.03E-01 | 1.03E+00 | 4.38E-02 | 1.03E+00 |
| ENSG000001 | AFAP1L1 | 5.19E-01 | 8.03E-01 | 1.03E+00 | 3.93E-02 | 1.03E+00 |
| ENSG000001 | AEBP2 | 5.19E-01 | 8.03E-01 | 1.03E+00 | 4.13E-02 | 1.03E+00 |
| ENSG000001 | RAPGEF6 | 5.19E-01 | 8.03E-01 | 9.79E-01 | -3.13E-02 | -1.02E+00 |
| ENSG000001 | COL27A1 | 5.19E-01 | 8.03E-01 | 9.73E-01 | -3.92E-02 | -1.03E+00 |
| ENSG000001 | ARL3 | 5.19E-01 | 8.03E-01 | 1.03E+00 | 4.57E-02 | 1.03E+00 |
| ENSG000001 | SMAD9 | 5.20E-01 | ? | 1.13E+00 | 1.74E-01 | 1.13E+00 |
| ENSG000002 | SCARNA9 | 5.20E-01 | ? | 1.19E+00 | 2.51E-01 | 1.19E+00 |
| ENSG00000C | TFPI | 5.20E-01 | 8.04E-01 | 1.04E+00 | 5.45E-02 | 1.04E+00 |
| ENSG000002 | --- | 5.20E-01 | ? | 1.24E+00 | 3.12E-01 | 1.24E+00 |
| ENSG000001 | MRPS18C | 5.20E-01 | 8.04E-01 | 9.65E-01 | -5.14E-02 | -1.04E+00 |
| ENSG000001 | OLFM2 | 5.20E-01 | 8.04E-01 | 1.08E+00 | 1.12E-01 | 1.08E+00 |
| ENSG000001 | WARS1 | 5.20E-01 | 8.04E-01 | 9.80E-01 | -2.90E-02 | -1.02E+00 |
| ENSG000001 | COL8A1 | 5.20E-01 | 8.04E-01 | 1.04E+00 | 5.55E-02 | 1.04E+00 |
| ENSG000002 | --- | 5.20E-01 | ? | 7.86E-01 | -3.48E-01 | -1.27E+00 |
| ENSG000001 | ZNF681 | 5.20E-01 | 8.04E-01 | 9.43E-01 | -8.53E-02 | -1.06E+00 |
| ENSG000001 | RPTOR | 5.20E-01 | 8.04E-01 | 1.03E+00 | 3.97E-02 | 1.03E+00 |
| ENSG000001 | MAMDC2 | 5.20E-01 | 8.04E-01 | 9.49E-01 | -7.55E-02 | -1.05E+00 |
| ENSG000001 | GPATCH4 | 5.20E-01 | 8.04E-01 | 1.03E+00 | 3.68E-02 | 1.03E+00 |
| ENSG000001 | HMGXB4 | 5.21E-01 | 8.04E-01 | 9.73E-01 | -3.99E-02 | -1.03E+00 |
| ENSG000001 | TAPT1 | 5.21E-01 | 8.04E-01 | 1.02E+00 | 3.50E-02 | 1.02E+00 |
| ENSG000001 | NNT | 5.21E-01 | 8.04E-01 | 1.02E+00 | 3.05E-02 | 1.02E+00 |
| ENSG000002 | --- | 5.21E-01 | ? | 8.54E-01 | -2.28E-01 | -1.17E+00 |
| ENSG000001 | PGBD4 | 5.21E-01 | 8.04E-01 | 1.04E+00 | 5.97E-02 | 1.04E+00 |
| ENSG000002 | C2orf16 | 5.21E-01 | 8.04E-01 | 9.41E-01 | -8.73E-02 | -1.06E+00 |
| ENSG000002 | --- | 5.21E-01 | ? | 1.34E+00 | 4.21E-01 | 1.34E+00 |
| ENSG000001 | ADAM17 | 5.21E-01 | 8.04E-01 | 1.02E+00 | 2.21E-02 | 1.02E+00 |
| ENSG000001 | LONP1 | 5.21E-01 | 8.04E-01 | 1.03E+00 | 4.06E-02 | 1.03E+00 |

| | | | | | | |
|---|---|---|---|---|---|---|
| ENSG000002 | --- | 5.21E-01 | ? | 1.15E+00 | 2.01E-01 | 1.15E+00 |
| ENSG000001 | TANGO2 | 5.21E-01 | 8.04E-01 | 9.72E-01 | -4.11E-02 | -1.03E+00 |
| ENSG000001 | SNX16 | 5.21E-01 | 8.04E-01 | 9.69E-01 | -4.47E-02 | -1.03E+00 |
| ENSG000002 | --- | 5.21E-01 | ? | 7.03E-01 | -5.08E-01 | -1.42E+00 |
| ENSG000001 | TNFAIP8L2 | 5.21E-01 | ? | 1.36E+00 | 4.44E-01 | 1.36E+00 |
| ENSG000002 | --- | 5.21E-01 | ? | 8.07E-01 | -3.09E-01 | -1.24E+00 |
| ENSG000002 | LINC01311 | 5.21E-01 | ? | 1.16E+00 | 2.20E-01 | 1.16E+00 |
| ENSG00000C | PDPR | 5.22E-01 | 8.04E-01 | 1.02E+00 | 2.96E-02 | 1.02E+00 |
| ENSG000001 | BORCS6 | 5.22E-01 | 8.04E-01 | 1.06E+00 | 8.50E-02 | 1.06E+00 |
| ENSG000002 | --- | 5.22E-01 | 8.04E-01 | 1.06E+00 | 8.20E-02 | 1.06E+00 |
| ENSG000001 | RXRA | 5.22E-01 | 8.04E-01 | 9.63E-01 | -5.38E-02 | -1.04E+00 |
| ENSG000001 | ADGRB2 | 5.22E-01 | 8.04E-01 | 9.14E-01 | -1.30E-01 | -1.09E+00 |
| ENSG000001 | GTF2F1 | 5.22E-01 | 8.04E-01 | 9.80E-01 | -2.95E-02 | -1.02E+00 |
| ENSG000001 | NCF2 | 5.22E-01 | 8.04E-01 | 1.04E+00 | 5.89E-02 | 1.04E+00 |
| ENSG000001 | ZNF518A | 5.22E-01 | 8.04E-01 | 9.70E-01 | -4.39E-02 | -1.03E+00 |
| ENSG000001 | SNX3 | 5.22E-01 | 8.05E-01 | 9.76E-01 | -3.56E-02 | -1.02E+00 |
| ENSG000001 | TRPV1 | 5.22E-01 | 8.05E-01 | 9.39E-01 | -9.02E-02 | -1.06E+00 |
| ENSG000002 | --- | 5.22E-01 | ? | 8.72E-01 | -1.97E-01 | -1.15E+00 |
| ENSG000002 | --- | 5.22E-01 | ? | 8.26E-01 | -2.75E-01 | -1.21E+00 |
| ENSG000001 | VASP | 5.22E-01 | 8.05E-01 | 1.03E+00 | 3.71E-02 | 1.03E+00 |
| ENSG000001 | RIPOR2 | 5.22E-01 | ? | 8.44E-01 | -2.44E-01 | -1.18E+00 |
| ENSG000002 | PDE4DIPP2 | 5.23E-01 | ? | 1.13E+00 | 1.74E-01 | 1.13E+00 |
| ENSG000001 | ATPAF1 | 5.23E-01 | 8.05E-01 | 1.03E+00 | 3.61E-02 | 1.03E+00 |
| ENSG000001 | RNPEPL1 | 5.23E-01 | 8.05E-01 | 1.06E+00 | 8.31E-02 | 1.06E+00 |
| ENSG000001 | SLC52A2 | 5.23E-01 | 8.06E-01 | 1.06E+00 | 8.34E-02 | 1.06E+00 |
| ENSG000002 | --- | 5.23E-01 | ? | 1.23E+00 | 2.97E-01 | 1.23E+00 |
| ENSG000002 | --- | 5.23E-01 | ? | 7.66E-01 | -3.84E-01 | -1.30E+00 |
| ENSG000001 | PABPN1 | 5.23E-01 | 8.06E-01 | 1.03E+00 | 4.06E-02 | 1.03E+00 |
| ENSG000001 | UQCC1 | 5.23E-01 | 8.06E-01 | 9.73E-01 | -3.90E-02 | -1.03E+00 |
| ENSG000001 | SLC35E3 | 5.23E-01 | 8.06E-01 | 1.02E+00 | 3.44E-02 | 1.02E+00 |
| ENSG000002 | ATP5MC1P6 | 5.23E-01 | ? | 7.19E-01 | -4.77E-01 | -1.39E+00 |
| ENSG000002 | TMEM11-DT | 5.23E-01 | ? | 1.17E+00 | 2.24E-01 | 1.17E+00 |
| ENSG000002 | --- | 5.23E-01 | ? | 1.35E+00 | 4.38E-01 | 1.35E+00 |
| ENSG000001 | MAX | 5.23E-01 | 8.06E-01 | 9.78E-01 | -3.21E-02 | -1.02E+00 |
| ENSG000001 | DYNC1H1 | 5.23E-01 | 8.06E-01 | 9.66E-01 | -4.95E-02 | -1.03E+00 |
| ENSG000001 | NFKBIB | 5.23E-01 | 8.06E-01 | 1.03E+00 | 4.64E-02 | 1.03E+00 |
| ENSG000002 | PSMB3 | 5.23E-01 | 8.06E-01 | 1.03E+00 | 4.19E-02 | 1.03E+00 |
| ENSG000001 | E2F7 | 5.23E-01 | 8.06E-01 | 9.63E-01 | -5.42E-02 | -1.04E+00 |
| ENSG000002 | PRKACB-DT | 5.23E-01 | ? | 7.91E-01 | -3.38E-01 | -1.26E+00 |
| ENSG000002 | CBX3P2 | 5.23E-01 | ? | 1.32E+00 | 4.05E-01 | 1.32E+00 |
| ENSG00000C | FSCN1 | 5.23E-01 | 8.06E-01 | 1.04E+00 | 6.35E-02 | 1.04E+00 |
| ENSG000001 | HINFP | 5.23E-01 | 8.06E-01 | 9.73E-01 | -3.92E-02 | -1.03E+00 |
| ENSG000001 | GCH1 | 5.23E-01 | 8.06E-01 | 1.03E+00 | 3.91E-02 | 1.03E+00 |
| ENSG000001 | OLFML2A | 5.24E-01 | 8.06E-01 | 1.06E+00 | 8.75E-02 | 1.06E+00 |
| ENSG000002 | FBXW11P1 | 5.24E-01 | ? | 1.29E+00 | 3.67E-01 | 1.29E+00 |

| | | | | | | | |
|---|---|---|---|---|---|---|---|
| ENSG000002 | CARD18 | 5.24E-01 | ? | | 7.92E-01 | -3.36E-01 | -1.26E+00 |
| ENSG000002 | TMEM191B | 5.24E-01 | ? | | 1.18E+00 | 2.38E-01 | 1.18E+00 |
| ENSG000002 | --- | 5.24E-01 | ? | | 1.70E+00 | 7.65E-01 | 1.70E+00 |
| ENSG000001 | PRMT5 | 5.24E-01 | | 8.06E-01 | 1.02E+00 | 2.66E-02 | 1.02E+00 |
| ENSG000001 | REEP4 | 5.24E-01 | | 8.06E-01 | 1.05E+00 | 6.74E-02 | 1.05E+00 |
| ENSG000001 | COQ3 | 5.24E-01 | | 8.06E-01 | 9.60E-01 | -5.82E-02 | -1.04E+00 |
| ENSG000002 | --- | 5.24E-01 | ? | | 8.50E-01 | -2.35E-01 | -1.18E+00 |
| ENSG000001 | FGF18 | 5.24E-01 | ? | | 7.65E-01 | -3.87E-01 | -1.31E+00 |
| ENSG000001 | PJA1 | 5.24E-01 | | 8.06E-01 | 1.02E+00 | 3.14E-02 | 1.02E+00 |
| ENSG000002 | CDK6-AS1 | 5.24E-01 | ? | | 1.24E+00 | 3.16E-01 | 1.24E+00 |
| ENSG000001 | ABCA9 | 5.24E-01 | ? | | 8.86E-01 | -1.75E-01 | -1.13E+00 |
| ENSG000001 | ZBTB14 | 5.24E-01 | | 8.06E-01 | 9.69E-01 | -4.54E-02 | -1.03E+00 |
| ENSG000002 | --- | 5.24E-01 | | 8.06E-01 | 1.37E+00 | 4.50E-01 | 1.37E+00 |
| ENSG000001 | HORMAD1 | 5.24E-01 | ? | | 1.24E+00 | 3.07E-01 | 1.24E+00 |
| ENSG00000C | ORC6 | 5.24E-01 | | 8.07E-01 | 9.40E-01 | -8.87E-02 | -1.06E+00 |
| ENSG000002 | TONSL-AS1 | 5.25E-01 | ? | | 1.37E+00 | 4.51E-01 | 1.37E+00 |
| ENSG000002 | --- | 5.25E-01 | ? | | 7.32E-01 | -4.49E-01 | -1.37E+00 |
| ENSG000002 | INO80D-AS2 | 5.25E-01 | | 8.07E-01 | 1.09E+00 | 1.26E-01 | 1.09E+00 |
| ENSG000002 | --- | 5.25E-01 | ? | | 1.38E+00 | 4.68E-01 | 1.38E+00 |
| ENSG000002 | --- | 5.25E-01 | ? | | 1.14E+00 | 1.89E-01 | 1.14E+00 |
| ENSG000002 | LINC01094 | 5.25E-01 | | 8.07E-01 | 1.07E+00 | 1.01E-01 | 1.07E+00 |
| ENSG000002 | PRDX6-AS1 | 5.25E-01 | ? | | 8.96E-01 | -1.59E-01 | -1.12E+00 |
| ENSG000001 | PTPRE | 5.25E-01 | | 8.07E-01 | 1.02E+00 | 2.62E-02 | 1.02E+00 |
| ENSG000001 | SLC36A1 | 5.25E-01 | | 8.07E-01 | 1.03E+00 | 4.17E-02 | 1.03E+00 |
| ENSG000002 | PCDHGB4 | 5.25E-01 | | 8.07E-01 | 9.60E-01 | -5.90E-02 | -1.04E+00 |
| ENSG000001 | MED1 | 5.25E-01 | | 8.07E-01 | 9.81E-01 | -2.77E-02 | -1.02E+00 |
| ENSG000002 | --- | 5.25E-01 | ? | | 1.23E+00 | 2.93E-01 | 1.23E+00 |
| ENSG000002 | --- | 5.25E-01 | | 8.07E-01 | 9.32E-01 | -1.02E-01 | -1.07E+00 |
| ENSG000002 | TMEM254-A: | 5.25E-01 | ? | | 1.18E+00 | 2.38E-01 | 1.18E+00 |
| ENSG000001 | DMKN | 5.25E-01 | | 8.07E-01 | 1.02E+00 | 2.83E-02 | 1.02E+00 |
| ENSG000001 | GRPR | 5.25E-01 | ? | | 1.34E+00 | 4.25E-01 | 1.34E+00 |
| ENSG000002 | --- | 5.25E-01 | ? | | 7.73E-01 | -3.71E-01 | -1.29E+00 |
| ENSG000002 | --- | 5.25E-01 | ? | | 8.63E-01 | -2.12E-01 | -1.16E+00 |
| ENSG000001 | MXRA8 | 5.26E-01 | | 8.08E-01 | 9.42E-01 | -8.55E-02 | -1.06E+00 |
| ENSG000001 | C19orf48 | 5.26E-01 | | 8.08E-01 | 1.05E+00 | 7.18E-02 | 1.05E+00 |
| ENSG000002 | --- | 5.26E-01 | ? | | 1.21E+00 | 2.70E-01 | 1.21E+00 |
| ENSG00000C | RABEP1 | 5.26E-01 | | 8.08E-01 | 9.77E-01 | -3.33E-02 | -1.02E+00 |
| ENSG000002 | --- | 5.26E-01 | | 8.08E-01 | 9.01E-01 | -1.50E-01 | -1.11E+00 |
| ENSG000002 | URGCP-MRP | 5.26E-01 | ? | | 1.26E+00 | 3.33E-01 | 1.26E+00 |
| ENSG000002 | --- | 5.26E-01 | ? | | 8.48E-01 | -2.38E-01 | -1.18E+00 |
| ENSG000001 | LRRN1 | 5.26E-01 | | 8.08E-01 | 1.06E+00 | 7.95E-02 | 1.06E+00 |
| ENSG000002 | --- | 5.26E-01 | ? | | 8.02E-01 | -3.19E-01 | -1.25E+00 |
| ENSG000001 | MANSC1 | 5.26E-01 | | 8.08E-01 | 9.60E-01 | -5.87E-02 | -1.04E+00 |
| ENSG000002 | --- | 5.26E-01 | ? | | 1.25E+00 | 3.19E-01 | 1.25E+00 |
| ENSG000002 | --- | 5.26E-01 | ? | | 1.41E+00 | 5.00E-01 | 1.41E+00 |

| | | | | | | |
|---|---|---|---|---|---|---|
| ENSG000001 | ARHGEF4 | 5.26E-01 | 8.08E-01 | 1.03E+00 | 4.67E-02 | 1.03E+00 |
| ENSG000002 | BTNL12P | 5.26E-01 | ? | 1.24E+00 | 3.12E-01 | 1.24E+00 |
| ENSG000001 | NDUFA10 | 5.26E-01 | 8.08E-01 | 9.80E-01 | -2.91E-02 | -1.02E+00 |
| ENSG000002 | --- | 5.26E-01 | ? | 8.55E-01 | -2.26E-01 | -1.17E+00 |
| ENSG000001 | DDX51 | 5.26E-01 | 8.08E-01 | 1.03E+00 | 4.55E-02 | 1.03E+00 |
| ENSG000001 | GABBR2 | 5.27E-01 | 8.08E-01 | 9.15E-01 | -1.29E-01 | -1.09E+00 |
| ENSG000001 | JAKMIP3 | 5.27E-01 | 8.08E-01 | 1.09E+00 | 1.24E-01 | 1.09E+00 |
| ENSG000001 | LRFN1 | 5.27E-01 | ? | 1.37E+00 | 4.53E-01 | 1.37E+00 |
| ENSG000002 | --- | 5.27E-01 | 8.08E-01 | 1.04E+00 | 5.87E-02 | 1.04E+00 |
| ENSG000002 | DSTNP3 | 5.27E-01 | ? | 1.22E+00 | 2.87E-01 | 1.22E+00 |
| ENSG000001 | SRPX | 5.27E-01 | 8.08E-01 | 1.02E+00 | 3.24E-02 | 1.02E+00 |
| ENSG000002 | UBE2Q2P2 | 5.27E-01 | ? | 8.97E-01 | -1.58E-01 | -1.12E+00 |
| ENSG000001 | NAT1 | 5.27E-01 | 8.09E-01 | 9.56E-01 | -6.46E-02 | -1.05E+00 |
| ENSG000001 | SERPINF1 | 5.27E-01 | 8.09E-01 | 1.03E+00 | 3.56E-02 | 1.03E+00 |
| ENSG000002 | --- | 5.27E-01 | ? | 1.42E+00 | 5.04E-01 | 1.42E+00 |
| ENSG000001 | PRRT3 | 5.27E-01 | 8.09E-01 | 1.10E+00 | 1.37E-01 | 1.10E+00 |
| ENSG000001 | PHYHD1 | 5.27E-01 | ? | 8.45E-01 | -2.44E-01 | -1.18E+00 |
| ENSG000001 | GRINA | 5.27E-01 | 8.09E-01 | 1.03E+00 | 3.95E-02 | 1.03E+00 |
| ENSG000001 | SPATA25 | 5.27E-01 | ? | 1.25E+00 | 3.20E-01 | 1.25E+00 |
| ENSG000001 | METTL3 | 5.27E-01 | 8.09E-01 | 9.74E-01 | -3.76E-02 | -1.03E+00 |
| ENSG000001 | RPSAP19 | 5.27E-01 | ? | 1.35E+00 | 4.38E-01 | 1.35E+00 |
| ENSG000002 | --- | 5.27E-01 | 8.09E-01 | 9.32E-01 | -1.02E-01 | -1.07E+00 |
| ENSG000001 | PHKG2 | 5.27E-01 | 8.09E-01 | 9.53E-01 | -7.01E-02 | -1.05E+00 |
| ENSG000002 | COX7CP1 | 5.28E-01 | ? | 6.36E-01 | -6.52E-01 | -1.57E+00 |
| ENSG000002 | FLJ20021 | 5.28E-01 | 8.09E-01 | 1.06E+00 | 8.39E-02 | 1.06E+00 |
| ENSG00000C | GPATCH1 | 5.28E-01 | 8.09E-01 | 9.64E-01 | -5.23E-02 | -1.04E+00 |
| ENSG000001 | CD101 | 5.28E-01 | 8.09E-01 | 9.22E-01 | -1.18E-01 | -1.09E+00 |
| ENSG000002 | OXCT1-AS1 | 5.28E-01 | ? | 1.31E+00 | 3.88E-01 | 1.31E+00 |
| ENSG000001 | LPCAT4 | 5.28E-01 | 8.09E-01 | 1.03E+00 | 3.77E-02 | 1.03E+00 |
| ENSG000002 | --- | 5.28E-01 | ? | 7.21E-01 | -4.72E-01 | -1.39E+00 |
| ENSG000001 | KLHL26 | 5.28E-01 | 8.10E-01 | 1.08E+00 | 1.14E-01 | 1.08E+00 |
| ENSG00000C | HSF2 | 5.28E-01 | 8.10E-01 | 1.04E+00 | 5.91E-02 | 1.04E+00 |
| ENSG000001 | HIP1 | 5.29E-01 | 8.10E-01 | 1.03E+00 | 4.00E-02 | 1.03E+00 |
| ENSG000001 | MAP3K7CL | 5.29E-01 | 8.10E-01 | 9.47E-01 | -7.92E-02 | -1.06E+00 |
| ENSG000001 | SENP7 | 5.29E-01 | 8.10E-01 | 9.64E-01 | -5.27E-02 | -1.04E+00 |
| ENSG000001 | METTL26 | 5.29E-01 | 8.10E-01 | 9.54E-01 | -6.81E-02 | -1.05E+00 |
| ENSG000001 | PPP1R9A | 5.29E-01 | ? | 1.17E+00 | 2.26E-01 | 1.17E+00 |
| ENSG000001 | AHDC1 | 5.29E-01 | 8.10E-01 | 1.04E+00 | 5.89E-02 | 1.04E+00 |
| ENSG000001 | PCED1A | 5.29E-01 | 8.10E-01 | 1.05E+00 | 7.35E-02 | 1.05E+00 |
| ENSG000002 | FOXP1-DT | 5.29E-01 | ? | 1.32E+00 | 4.05E-01 | 1.32E+00 |
| ENSG000001 | ZNF778 | 5.29E-01 | 8.10E-01 | 1.04E+00 | 5.06E-02 | 1.04E+00 |
| ENSG00000C | PI4K2B | 5.29E-01 | 8.10E-01 | 1.02E+00 | 3.30E-02 | 1.02E+00 |
| ENSG000001 | S100A9 | 5.29E-01 | 8.10E-01 | 1.06E+00 | 8.96E-02 | 1.06E+00 |
| ENSG000002 | --- | 5.29E-01 | ? | 8.31E-01 | -2.68E-01 | -1.20E+00 |
| ENSG000002 | --- | 5.29E-01 | ? | 8.58E-01 | -2.22E-01 | -1.17E+00 |

| | | | | | | | |
|---|---|---|---|---|---|---|---|
| ENSG00000C | CRAMP1 | 5.29E-01 | | 8.10E-01 | 9.73E-01 | -3.92E-02 | -1.03E+00 |
| ENSG00000C | GEMIN2 | 5.29E-01 | | 8.10E-01 | 9.61E-01 | -5.73E-02 | -1.04E+00 |
| ENSG000002 | ARHGEF2-AS | 5.29E-01 | ? | | 8.65E-01 | -2.09E-01 | -1.16E+00 |
| ENSG000002 | DRG1P1 | 5.29E-01 | ? | | 7.61E-01 | -3.93E-01 | -1.31E+00 |
| ENSG000001 | TTL | 5.29E-01 | | 8.10E-01 | 9.83E-01 | -2.53E-02 | -1.02E+00 |
| ENSG000002 | LINC01186 | 5.30E-01 | ? | | 9.03E-01 | -1.47E-01 | -1.11E+00 |
| ENSG000001 | ADGRF3 | 5.30E-01 | ? | | 8.81E-01 | -1.82E-01 | -1.13E+00 |
| ENSG00000C | THRAP3 | 5.30E-01 | | 8.10E-01 | 9.84E-01 | -2.26E-02 | -1.02E+00 |
| ENSG000001 | MXI1 | 5.30E-01 | | 8.10E-01 | 9.64E-01 | -5.26E-02 | -1.04E+00 |
| ENSG000001 | PARD3 | 5.30E-01 | | 8.10E-01 | 1.02E+00 | 2.39E-02 | 1.02E+00 |
| ENSG000001 | ECD | 5.30E-01 | | 8.10E-01 | 9.74E-01 | -3.82E-02 | -1.03E+00 |
| ENSG000002 | IFITM10 | 5.30E-01 | ? | | 1.22E+00 | 2.82E-01 | 1.22E+00 |
| ENSG000002 | --- | 5.30E-01 | ? | | 7.80E-01 | -3.59E-01 | -1.28E+00 |
| ENSG000001 | R3HCC1L | 5.30E-01 | | 8.11E-01 | 1.02E+00 | 3.55E-02 | 1.02E+00 |
| ENSG000002 | --- | 5.30E-01 | ? | | 8.11E-01 | -3.03E-01 | -1.23E+00 |
| ENSG000001 | MOB3C | 5.30E-01 | | 8.11E-01 | 9.72E-01 | -4.03E-02 | -1.03E+00 |
| ENSG000002 | --- | 5.30E-01 | ? | | 8.25E-01 | -2.78E-01 | -1.21E+00 |
| ENSG000001 | GDAP1 | 5.30E-01 | | 8.11E-01 | 1.03E+00 | 4.86E-02 | 1.03E+00 |
| ENSG000001 | STX16 | 5.30E-01 | | 8.11E-01 | 9.63E-01 | -5.46E-02 | -1.04E+00 |
| ENSG000001 | ACVR1C | 5.30E-01 | ? | | 8.81E-01 | -1.82E-01 | -1.13E+00 |
| ENSG000001 | HES1 | 5.30E-01 | | 8.11E-01 | 9.70E-01 | -4.40E-02 | -1.03E+00 |
| ENSG000001 | TP53BP2 | 5.31E-01 | | 8.11E-01 | 9.72E-01 | -4.15E-02 | -1.03E+00 |
| ENSG000002 | EOLA2-DT | 5.31E-01 | | 8.11E-01 | 1.07E+00 | 9.73E-02 | 1.07E+00 |
| ENSG000001 | CEL | 5.31E-01 | | 8.11E-01 | 9.48E-01 | -7.75E-02 | -1.06E+00 |
| ENSG000002 | VWA7 | 5.31E-01 | | 8.11E-01 | 1.11E+00 | 1.52E-01 | 1.11E+00 |
| ENSG000002 | --- | 5.31E-01 | ? | | 1.35E+00 | 4.38E-01 | 1.35E+00 |
| ENSG00000C | TF | 5.31E-01 | ? | | 8.32E-01 | -2.66E-01 | -1.20E+00 |
| ENSG00000C | MCOLN3 | 5.31E-01 | | 8.11E-01 | 1.05E+00 | 7.66E-02 | 1.05E+00 |
| ENSG000001 | MMP3 | 5.31E-01 | | 8.11E-01 | 9.13E-01 | -1.31E-01 | -1.09E+00 |
| ENSG000002 | --- | 5.31E-01 | ? | | 1.42E+00 | 5.09E-01 | 1.42E+00 |
| ENSG000002 | ZNF497-AS1 | 5.31E-01 | ? | | 8.23E-01 | -2.81E-01 | -1.22E+00 |
| ENSG000001 | AR | 5.31E-01 | ? | | 1.23E+00 | 2.97E-01 | 1.23E+00 |
| ENSG000002 | LINC01144 | 5.31E-01 | ? | | 1.30E+00 | 3.83E-01 | 1.30E+00 |
| ENSG000001 | TMEM186 | 5.31E-01 | | 8.11E-01 | 1.04E+00 | 5.71E-02 | 1.04E+00 |
| ENSG000001 | SCN5A | 5.31E-01 | ? | | 1.44E+00 | 5.26E-01 | 1.44E+00 |
| ENSG000001 | MINDY1 | 5.31E-01 | | 8.11E-01 | 9.63E-01 | -5.46E-02 | -1.04E+00 |
| ENSG000001 | TONSL | 5.31E-01 | | 8.11E-01 | 9.48E-01 | -7.64E-02 | -1.05E+00 |
| ENSG000002 | PSMB9 | 5.31E-01 | ? | | 9.00E-01 | -1.52E-01 | -1.11E+00 |
| ENSG000002 | --- | 5.31E-01 | | 8.12E-01 | 9.23E-01 | -1.16E-01 | -1.08E+00 |
| ENSG000001 | TARBP2 | 5.32E-01 | | 8.12E-01 | 1.05E+00 | 6.49E-02 | 1.05E+00 |
| ENSG00000C | ABHD5 | 5.32E-01 | | 8.12E-01 | 1.02E+00 | 3.43E-02 | 1.02E+00 |
| ENSG000001 | ACTG1P1 | 5.32E-01 | ? | | 1.28E+00 | 3.52E-01 | 1.28E+00 |
| ENSG000001 | TRIM33 | 5.32E-01 | | 8.12E-01 | 9.72E-01 | -4.05E-02 | -1.03E+00 |
| ENSG000001 | ZNF460 | 5.32E-01 | | 8.12E-01 | 1.08E+00 | 1.15E-01 | 1.08E+00 |
| ENSG000001 | PRKAA1 | 5.32E-01 | | 8.12E-01 | 9.74E-01 | -3.81E-02 | -1.03E+00 |

| | | | | | | |
|---|---|---|---|---|---|---|
| ENSG000001 | METTL16 | 5.32E-01 | 8.12E-01 | 1.02E+00 | 2.76E-02 | 1.02E+00 |
| ENSG000002 | SELENOO-AS | 5.32E-01 | ? | 1.25E+00 | 3.18E-01 | 1.25E+00 |
| ENSG000002 | ZNF687-AS1 | 5.32E-01 | ? | 1.29E+00 | 3.69E-01 | 1.29E+00 |
| ENSG000001 | FADS2 | 5.32E-01 | 8.12E-01 | 1.03E+00 | 4.14E-02 | 1.03E+00 |
| ENSG000002 | --- | 5.32E-01 | ? | 1.31E+00 | 3.93E-01 | 1.31E+00 |
| ENSG000001 | KNDC1 | 5.32E-01 | ? | 1.17E+00 | 2.29E-01 | 1.17E+00 |
| ENSG000001 | CEBPB | 5.32E-01 | 8.12E-01 | 9.69E-01 | -4.54E-02 | -1.03E+00 |
| ENSG000002 | NPIPA2 | 5.33E-01 | ? | 7.93E-01 | -3.34E-01 | -1.26E+00 |
| ENSG000001 | NPIPP1 | 5.33E-01 | 8.12E-01 | 9.10E-01 | -1.37E-01 | -1.10E+00 |
| ENSG00000C | PSME4 | 5.33E-01 | 8.12E-01 | 1.03E+00 | 3.87E-02 | 1.03E+00 |
| ENSG000002 | PHB2 | 5.33E-01 | 8.12E-01 | 1.03E+00 | 4.48E-02 | 1.03E+00 |
| ENSG000001 | UGT8 | 5.33E-01 | 8.12E-01 | 9.17E-01 | -1.25E-01 | -1.09E+00 |
| ENSG000001 | PLSCR1 | 5.33E-01 | 8.12E-01 | 1.03E+00 | 4.46E-02 | 1.03E+00 |
| ENSG00000C | SP100 | 5.33E-01 | 8.12E-01 | 9.76E-01 | -3.54E-02 | -1.02E+00 |
| ENSG000001 | PIGH | 5.33E-01 | 8.12E-01 | 1.03E+00 | 4.51E-02 | 1.03E+00 |
| ENSG000002 | --- | 5.33E-01 | ? | 1.48E+00 | 5.62E-01 | 1.48E+00 |
| ENSG000002 | --- | 5.33E-01 | 8.12E-01 | 1.08E+00 | 1.09E-01 | 1.08E+00 |
| ENSG000001 | KIF21A | 5.33E-01 | 8.13E-01 | 1.02E+00 | 3.36E-02 | 1.02E+00 |
| ENSG000002 | FTH1P16 | 5.33E-01 | ? | 8.18E-01 | -2.89E-01 | -1.22E+00 |
| ENSG000002 | --- | 5.33E-01 | 8.13E-01 | 9.14E-01 | -1.29E-01 | -1.09E+00 |
| ENSG000001 | TAF5L | 5.33E-01 | 8.13E-01 | 9.77E-01 | -3.36E-02 | -1.02E+00 |
| ENSG000002 | --- | 5.33E-01 | 8.13E-01 | 8.85E-01 | -1.76E-01 | -1.13E+00 |
| ENSG000002 | ZNF566-AS1 | 5.34E-01 | 8.13E-01 | 1.07E+00 | 9.92E-02 | 1.07E+00 |
| ENSG000002 | --- | 5.34E-01 | 8.13E-01 | 9.26E-01 | -1.10E-01 | -1.08E+00 |
| ENSG000001 | VN1R1 | 5.34E-01 | ? | 1.16E+00 | 2.14E-01 | 1.16E+00 |
| ENSG000002 | --- | 5.34E-01 | ? | 1.34E+00 | 4.22E-01 | 1.34E+00 |
| ENSG000001 | MBD6 | 5.34E-01 | 8.14E-01 | 1.02E+00 | 3.34E-02 | 1.02E+00 |
| ENSG000002 | NRADDP | 5.34E-01 | ? | 8.05E-01 | -3.13E-01 | -1.24E+00 |
| ENSG000001 | NECAB3 | 5.34E-01 | 8.14E-01 | 1.08E+00 | 1.14E-01 | 1.08E+00 |
| ENSG000001 | RPS6KA3 | 5.34E-01 | 8.14E-01 | 1.02E+00 | 3.23E-02 | 1.02E+00 |
| ENSG000001 | BMF | 5.34E-01 | ? | 8.62E-01 | -2.14E-01 | -1.16E+00 |
| ENSG000002 | --- | 5.34E-01 | ? | 7.64E-01 | -3.89E-01 | -1.31E+00 |
| ENSG000001 | BTC | 5.34E-01 | 8.14E-01 | 9.24E-01 | -1.14E-01 | -1.08E+00 |
| ENSG000001 | GRWD1 | 5.34E-01 | 8.14E-01 | 1.03E+00 | 4.47E-02 | 1.03E+00 |
| ENSG00000C | NRP1 | 5.35E-01 | 8.14E-01 | 9.78E-01 | -3.28E-02 | -1.02E+00 |
| ENSG000002 | MCRIP1 | 5.35E-01 | 8.14E-01 | 1.05E+00 | 7.35E-02 | 1.05E+00 |
| ENSG000001 | CCDC78 | 5.35E-01 | 8.14E-01 | 9.34E-01 | -9.87E-02 | -1.07E+00 |
| ENSG000001 | ZNF724 | 5.35E-01 | 8.14E-01 | 1.08E+00 | 1.05E-01 | 1.08E+00 |
| ENSG000001 | APOBEC3B | 5.35E-01 | ? | 1.20E+00 | 2.65E-01 | 1.20E+00 |
| ENSG000001 | TPD52L1 | 5.35E-01 | 8.14E-01 | 1.03E+00 | 3.91E-02 | 1.03E+00 |
| ENSG000002 | --- | 5.35E-01 | 8.14E-01 | 7.39E-01 | -4.36E-01 | -1.35E+00 |
| ENSG000001 | DPM2 | 5.35E-01 | 8.14E-01 | 1.03E+00 | 4.39E-02 | 1.03E+00 |
| ENSG000002 | ARPC4 | 5.35E-01 | 8.14E-01 | 1.03E+00 | 4.82E-02 | 1.03E+00 |
| ENSG000001 | SULT1C2 | 5.35E-01 | ? | 1.26E+00 | 3.28E-01 | 1.26E+00 |
| ENSG000001 | CISD2 | 5.36E-01 | 8.15E-01 | 9.70E-01 | -4.38E-02 | -1.03E+00 |

| | | | | | | |
|---|---|---|---|---|---|---|
| ENSG000002 | VPS33B-DT | 5.36E-01 | ? | 1.31E+00 | 3.91E-01 | 1.31E+00 |
| ENSG000001 | TMUB1 | 5.36E-01 | 8.15E-01 | 1.04E+00 | 5.72E-02 | 1.04E+00 |
| ENSG000002 | SNX10-AS1 | 5.36E-01 | ? | 1.37E+00 | 4.56E-01 | 1.37E+00 |
| ENSG000002 | --- | 5.36E-01 | ? | 6.85E-01 | -5.47E-01 | -1.46E+00 |
| ENSG000002 | --- | 5.36E-01 | ? | 1.16E+00 | 2.16E-01 | 1.16E+00 |
| ENSG000001 | SLC35F3 | 5.36E-01 | 8.15E-01 | 9.60E-01 | -5.92E-02 | -1.04E+00 |
| ENSG000001 | CCDC65 | 5.36E-01 | ? | 8.75E-01 | -1.93E-01 | -1.14E+00 |
| ENSG000001 | ZNF202 | 5.36E-01 | 8.15E-01 | 9.73E-01 | -4.00E-02 | -1.03E+00 |
| ENSG000002 | --- | 5.36E-01 | 8.15E-01 | 1.08E+00 | 1.09E-01 | 1.08E+00 |
| ENSG00000C | SCYL3 | 5.36E-01 | 8.15E-01 | 9.70E-01 | -4.46E-02 | -1.03E+00 |
| ENSG000002 | EVPLL | 5.36E-01 | 8.16E-01 | 9.21E-01 | -1.19E-01 | -1.09E+00 |
| ENSG000002 | --- | 5.37E-01 | ? | 8.87E-01 | -1.74E-01 | -1.13E+00 |
| ENSG000001 | POFUT1 | 5.37E-01 | 8.16E-01 | 9.74E-01 | -3.74E-02 | -1.03E+00 |
| ENSG000001 | PTOV1 | 5.37E-01 | 8.16E-01 | 1.04E+00 | 5.35E-02 | 1.04E+00 |
| ENSG000001 | ARL13B | 5.37E-01 | 8.16E-01 | 9.66E-01 | -5.01E-02 | -1.04E+00 |
| ENSG000002 | UBL7-DT | 5.37E-01 | 8.16E-01 | 9.35E-01 | -9.66E-02 | -1.07E+00 |
| ENSG000002 | --- | 5.37E-01 | ? | 7.65E-01 | -3.86E-01 | -1.31E+00 |
| ENSG000001 | SIRPB1 | 5.37E-01 | 8.16E-01 | 1.04E+00 | 6.17E-02 | 1.04E+00 |
| ENSG000001 | VCPKMT | 5.37E-01 | 8.16E-01 | 9.59E-01 | -6.08E-02 | -1.04E+00 |
| ENSG000001 | SPCS2 | 5.37E-01 | 8.16E-01 | 1.02E+00 | 2.53E-02 | 1.02E+00 |
| ENSG000002 | ZNF814 | 5.37E-01 | 8.16E-01 | 9.64E-01 | -5.35E-02 | -1.04E+00 |
| ENSG000001 | NPFF | 5.37E-01 | 8.16E-01 | 9.00E-01 | -1.52E-01 | -1.11E+00 |
| ENSG000001 | RUSC2 | 5.37E-01 | 8.16E-01 | 1.02E+00 | 2.98E-02 | 1.02E+00 |
| ENSG000001 | SLC13A4 | 5.37E-01 | ? | 8.70E-01 | -2.01E-01 | -1.15E+00 |
| ENSG000001 | WDR83 | 5.37E-01 | 8.16E-01 | 9.56E-01 | -6.49E-02 | -1.05E+00 |
| ENSG000001 | CDR2 | 5.37E-01 | 8.16E-01 | 9.82E-01 | -2.64E-02 | -1.02E+00 |
| ENSG000001 | ADCY9 | 5.38E-01 | 8.16E-01 | 1.03E+00 | 4.45E-02 | 1.03E+00 |
| ENSG000001 | TMEM198 | 5.38E-01 | 8.16E-01 | 8.99E-01 | -1.53E-01 | -1.11E+00 |
| ENSG000001 | MELTF | 5.38E-01 | 8.16E-01 | 1.05E+00 | 7.25E-02 | 1.05E+00 |
| ENSG000001 | NSMCE3 | 5.38E-01 | 8.16E-01 | 9.76E-01 | -3.52E-02 | -1.02E+00 |
| ENSG000002 | --- | 5.38E-01 | ? | 8.59E-01 | -2.19E-01 | -1.16E+00 |
| ENSG000001 | SNX14 | 5.38E-01 | 8.16E-01 | 9.78E-01 | -3.27E-02 | -1.02E+00 |
| ENSG000002 | CENPVL3 | 5.38E-01 | ? | 1.28E+00 | 3.52E-01 | 1.28E+00 |
| ENSG000001 | BNIPL | 5.38E-01 | 8.16E-01 | 1.08E+00 | 1.13E-01 | 1.08E+00 |
| ENSG000001 | BTD | 5.38E-01 | 8.16E-01 | 9.65E-01 | -5.09E-02 | -1.04E+00 |
| ENSG000001 | C1orf115 | 5.38E-01 | 8.16E-01 | 1.05E+00 | 7.31E-02 | 1.05E+00 |
| ENSG000002 | 7SK | 5.38E-01 | ? | 8.56E-01 | -2.24E-01 | -1.17E+00 |
| ENSG000001 | B3GALT5 | 5.38E-01 | ? | 8.65E-01 | -2.09E-01 | -1.16E+00 |
| ENSG000001 | INAVA | 5.38E-01 | 8.16E-01 | 1.03E+00 | 3.81E-02 | 1.03E+00 |
| ENSG000002 | SMIM11 | 5.38E-01 | 8.16E-01 | 1.10E+00 | 1.41E-01 | 1.10E+00 |
| ENSG000001 | TRMT61A | 5.38E-01 | 8.16E-01 | 1.06E+00 | 8.57E-02 | 1.06E+00 |
| ENSG000002 | CYLD-AS1 | 5.38E-01 | ? | 8.61E-01 | -2.16E-01 | -1.16E+00 |
| ENSG000002 | KRT17P6 | 5.38E-01 | ? | 1.34E+00 | 4.18E-01 | 1.34E+00 |
| ENSG000001 | PPP2R2B | 5.38E-01 | 8.16E-01 | 9.47E-01 | -7.92E-02 | -1.06E+00 |
| ENSG00000C | OAS1 | 5.38E-01 | 8.16E-01 | 1.08E+00 | 1.08E-01 | 1.08E+00 |

| | | | | | | |
|---|---|---|---|---|---|---|
| ENSG000001 | SLC4A1AP | 5.38E-01 | 8.16E-01 | 9.78E-01 | -3.26E-02 | -1.02E+00 |
| ENSG000001 | GPC5 | 5.38E-01 | ? | 1.22E+00 | 2.89E-01 | 1.22E+00 |
| ENSG000002 | --- | 5.38E-01 | ? | 1.28E+00 | 3.56E-01 | 1.28E+00 |
| ENSG000002 | --- | 5.39E-01 | 8.16E-01 | 9.29E-01 | -1.06E-01 | -1.08E+00 |
| ENSG000002 | --- | 5.39E-01 | ? | 1.37E+00 | 4.58E-01 | 1.37E+00 |
| ENSG000002 | --- | 5.39E-01 | ? | 8.92E-01 | -1.65E-01 | -1.12E+00 |
| ENSG000002 | --- | 5.39E-01 | 8.17E-01 | 1.11E+00 | 1.56E-01 | 1.11E+00 |
| ENSG000002 | --- | 5.39E-01 | ? | 8.26E-01 | -2.76E-01 | -1.21E+00 |
| ENSG000001 | ADCY6 | 5.39E-01 | 8.17E-01 | 1.03E+00 | 3.82E-02 | 1.03E+00 |
| ENSG000002 | SLC4A1APP1 | 5.39E-01 | ? | 7.27E-01 | -4.60E-01 | -1.38E+00 |
| ENSG000001 | ATG101 | 5.39E-01 | 8.17E-01 | 9.72E-01 | -4.14E-02 | -1.03E+00 |
| ENSG000001 | DIABLO | 5.39E-01 | 8.17E-01 | 1.02E+00 | 2.72E-02 | 1.02E+00 |
| ENSG000001 | EMG1 | 5.39E-01 | 8.17E-01 | 1.03E+00 | 4.53E-02 | 1.03E+00 |
| ENSG000001 | WDR89 | 5.39E-01 | 8.17E-01 | 9.76E-01 | -3.49E-02 | -1.02E+00 |
| ENSG000002 | H3C10 | 5.39E-01 | 8.17E-01 | 1.07E+00 | 1.04E-01 | 1.07E+00 |
| ENSG000001 | NCKAP5L | 5.39E-01 | 8.17E-01 | 1.04E+00 | 5.29E-02 | 1.04E+00 |
| ENSG000001 | PLA2G15 | 5.39E-01 | 8.17E-01 | 1.04E+00 | 5.88E-02 | 1.04E+00 |
| ENSG000001 | CCDC97 | 5.39E-01 | 8.17E-01 | 9.80E-01 | -2.92E-02 | -1.02E+00 |
| ENSG000001 | CLIC2 | 5.39E-01 | ? | 1.14E+00 | 1.89E-01 | 1.14E+00 |
| ENSG000002 | --- | 5.39E-01 | ? | 8.96E-01 | -1.58E-01 | -1.12E+00 |
| ENSG000001 | PTGFR | 5.39E-01 | ? | 1.20E+00 | 2.68E-01 | 1.20E+00 |
| ENSG000002 | RASSF8-AS1 | 5.39E-01 | 8.17E-01 | 9.48E-01 | -7.78E-02 | -1.06E+00 |
| ENSG000001 | SRF | 5.39E-01 | 8.17E-01 | 9.78E-01 | -3.25E-02 | -1.02E+00 |
| ENSG000001 | NUP88 | 5.40E-01 | 8.17E-01 | 9.81E-01 | -2.72E-02 | -1.02E+00 |
| ENSG000001 | POLR2C | 5.40E-01 | 8.17E-01 | 1.03E+00 | 4.01E-02 | 1.03E+00 |
| ENSG000001 | CACUL1 | 5.40E-01 | 8.17E-01 | 9.82E-01 | -2.55E-02 | -1.02E+00 |
| ENSG000002 | LINC01023 | 5.40E-01 | ? | 7.73E-01 | -3.71E-01 | -1.29E+00 |
| ENSG000001 | PPP1R35 | 5.40E-01 | 8.17E-01 | 1.06E+00 | 8.56E-02 | 1.06E+00 |
| ENSG000001 | ZNF646 | 5.40E-01 | 8.17E-01 | 1.03E+00 | 3.75E-02 | 1.03E+00 |
| ENSG000001 | PDIA3 | 5.40E-01 | 8.17E-01 | 9.80E-01 | -2.93E-02 | -1.02E+00 |
| ENSG000001 | CPNE8 | 5.40E-01 | 8.17E-01 | 9.72E-01 | -4.04E-02 | -1.03E+00 |
| ENSG000001 | ZRSR2 | 5.40E-01 | 8.17E-01 | 9.72E-01 | -4.13E-02 | -1.03E+00 |
| ENSG000002 | GEMIN8P4 | 5.40E-01 | ? | 1.32E+00 | 3.98E-01 | 1.32E+00 |
| ENSG000002 | --- | 5.40E-01 | ? | 8.83E-01 | -1.80E-01 | -1.13E+00 |
| ENSG000002 | --- | 5.40E-01 | ? | 7.80E-01 | -3.58E-01 | -1.28E+00 |
| ENSG000001 | HDGF | 5.40E-01 | 8.17E-01 | 9.82E-01 | -2.57E-02 | -1.02E+00 |
| ENSG000002 | --- | 5.40E-01 | 8.17E-01 | 9.25E-01 | -1.12E-01 | -1.08E+00 |
| ENSG000001 | C9orf40 | 5.40E-01 | 8.17E-01 | 1.04E+00 | 5.89E-02 | 1.04E+00 |
| ENSG000002 | CCDC162P | 5.40E-01 | 8.17E-01 | 9.25E-01 | -1.12E-01 | -1.08E+00 |
| ENSG00000C | PRKCH | 5.40E-01 | 8.17E-01 | 9.75E-01 | -3.69E-02 | -1.03E+00 |
| ENSG00000C | HSD17B14 | 5.40E-01 | ? | 8.75E-01 | -1.93E-01 | -1.14E+00 |
| ENSG000001 | LDHAL6A | 5.40E-01 | ? | 1.32E+00 | 4.01E-01 | 1.32E+00 |
| ENSG000001 | SFXN4 | 5.41E-01 | 8.17E-01 | 1.04E+00 | 5.66E-02 | 1.04E+00 |
| ENSG000002 | --- | 5.41E-01 | ? | 1.17E+00 | 2.30E-01 | 1.17E+00 |
| ENSG000002 | --- | 5.41E-01 | ? | 7.69E-01 | -3.79E-01 | -1.30E+00 |

| | | | | | | |
|---|---|---|---|---|---|---|
| ENSG00000C | MCF2L2 | 5.41E-01 | 8.17E-01 | 1.09E+00 | 1.21E-01 | 1.09E+00 |
| ENSG000002 | --- | 5.41E-01 | 8.17E-01 | 1.08E+00 | 1.06E-01 | 1.08E+00 |
| ENSG000001 | SIX5 | 5.41E-01 | 8.18E-01 | 9.35E-01 | -9.76E-02 | -1.07E+00 |
| ENSG000002 | --- | 5.41E-01 | ? | 1.20E+00 | 2.68E-01 | 1.20E+00 |
| ENSG000002 | --- | 5.41E-01 | 8.18E-01 | 1.07E+00 | 9.69E-02 | 1.07E+00 |
| ENSG000002 | LINC02897 | 5.41E-01 | ? | 1.21E+00 | 2.74E-01 | 1.21E+00 |
| ENSG000001 | ZNF57 | 5.41E-01 | 8.18E-01 | 1.03E+00 | 4.83E-02 | 1.03E+00 |
| ENSG000001 | ALOX15B | 5.41E-01 | 8.18E-01 | 1.05E+00 | 6.71E-02 | 1.05E+00 |
| ENSG000001 | TTC5 | 5.41E-01 | 8.18E-01 | 1.03E+00 | 4.77E-02 | 1.03E+00 |
| ENSG000002 | --- | 5.41E-01 | ? | 7.47E-01 | -4.21E-01 | -1.34E+00 |
| ENSG000001 | CRACR2A | 5.41E-01 | 8.18E-01 | 9.34E-01 | -9.90E-02 | -1.07E+00 |
| ENSG000002 | PCDHGB5 | 5.41E-01 | 8.18E-01 | 9.73E-01 | -4.01E-02 | -1.03E+00 |
| ENSG000001 | MAP2K2 | 5.42E-01 | 8.18E-01 | 1.04E+00 | 5.00E-02 | 1.04E+00 |
| ENSG000002 | --- | 5.42E-01 | ? | 8.24E-01 | -2.79E-01 | -1.21E+00 |
| ENSG000002 | --- | 5.42E-01 | ? | 1.38E+00 | 4.60E-01 | 1.38E+00 |
| ENSG000001 | ECM2 | 5.42E-01 | ? | 8.45E-01 | -2.42E-01 | -1.18E+00 |
| ENSG000002 | --- | 5.42E-01 | 8.18E-01 | 1.36E+00 | 4.47E-01 | 1.36E+00 |
| ENSG000002 | CNTNAP3C | 5.42E-01 | ? | 1.31E+00 | 3.95E-01 | 1.31E+00 |
| ENSG000001 | SPIN2B | 5.42E-01 | 8.18E-01 | 1.05E+00 | 6.99E-02 | 1.05E+00 |
| ENSG000001 | LDHAL6B | 5.42E-01 | ? | 1.27E+00 | 3.48E-01 | 1.27E+00 |
| ENSG00000C | AQR | 5.42E-01 | 8.18E-01 | 1.02E+00 | 2.64E-02 | 1.02E+00 |
| ENSG000002 | RAD21-AS1 | 5.42E-01 | ? | 1.17E+00 | 2.23E-01 | 1.17E+00 |
| ENSG000002 | SALRNA1 | 5.42E-01 | ? | 7.09E-01 | -4.96E-01 | -1.41E+00 |
| ENSG000001 | TSKU | 5.42E-01 | 8.18E-01 | 1.04E+00 | 5.13E-02 | 1.04E+00 |
| ENSG000001 | NDNF | 5.42E-01 | 8.18E-01 | 9.16E-01 | -1.27E-01 | -1.09E+00 |
| ENSG000001 | DR1 | 5.42E-01 | 8.18E-01 | 1.02E+00 | 2.94E-02 | 1.02E+00 |
| ENSG000001 | FBXO7 | 5.42E-01 | 8.18E-01 | 1.01E+00 | 2.04E-02 | 1.01E+00 |
| ENSG000001 | TGM3 | 5.42E-01 | ? | 7.45E-01 | -4.25E-01 | -1.34E+00 |
| ENSG000002 | OSGEPL1-AS | 5.43E-01 | ? | 7.49E-01 | -4.16E-01 | -1.33E+00 |
| ENSG000002 | ZNF688 | 5.43E-01 | 8.19E-01 | 1.06E+00 | 8.36E-02 | 1.06E+00 |
| ENSG000001 | ARHGEF19 | 5.43E-01 | 8.19E-01 | 1.09E+00 | 1.26E-01 | 1.09E+00 |
| ENSG000001 | PLXDC2 | 5.43E-01 | 8.19E-01 | 9.70E-01 | -4.37E-02 | -1.03E+00 |
| ENSG000001 | MN1 | 5.43E-01 | 8.19E-01 | 1.03E+00 | 4.80E-02 | 1.03E+00 |
| ENSG000001 | ZFP30 | 5.43E-01 | 8.19E-01 | 1.03E+00 | 3.85E-02 | 1.03E+00 |
| ENSG000001 | CCNP | 5.43E-01 | ? | 7.69E-01 | -3.79E-01 | -1.30E+00 |
| ENSG000002 | --- | 5.43E-01 | ? | 8.12E-01 | -3.01E-01 | -1.23E+00 |
| ENSG000001 | BACE1 | 5.43E-01 | 8.19E-01 | 1.02E+00 | 3.33E-02 | 1.02E+00 |
| ENSG000002 | MXD3 | 5.43E-01 | 8.19E-01 | 9.49E-01 | -7.62E-02 | -1.05E+00 |
| ENSG000001 | CTH | 5.43E-01 | 8.19E-01 | 1.09E+00 | 1.21E-01 | 1.09E+00 |
| ENSG000001 | TMEM63A | 5.43E-01 | 8.19E-01 | 9.66E-01 | -4.99E-02 | -1.04E+00 |
| ENSG00000C | RAB10 | 5.43E-01 | 8.19E-01 | 9.83E-01 | -2.54E-02 | -1.02E+00 |
| ENSG000001 | NOVA1 | 5.43E-01 | ? | 1.16E+00 | 2.12E-01 | 1.16E+00 |
| ENSG000002 | --- | 5.43E-01 | ? | 1.26E+00 | 3.34E-01 | 1.26E+00 |
| ENSG000002 | --- | 5.43E-01 | 8.19E-01 | 9.17E-01 | -1.25E-01 | -1.09E+00 |
| ENSG000002 | POLR2J3 | 5.43E-01 | 8.19E-01 | 9.38E-01 | -9.20E-02 | -1.07E+00 |

| | | | | | | |
|---|---|---|---|---|---|---|
| ENSG000001 | PHKG1 | 5.43E-01 | ? | 1.17E+00 | 2.20E-01 | 1.17E+00 |
| ENSG000001 | GPR83 | 5.43E-01 | ? | 1.40E+00 | 4.86E-01 | 1.40E+00 |
| ENSG000001 | RABEP2 | 5.43E-01 | 8.19E-01 | 1.05E+00 | 7.12E-02 | 1.05E+00 |
| ENSG000002 | UBBP4 | 5.44E-01 | ? | 1.36E+00 | 4.41E-01 | 1.36E+00 |
| ENSG000001 | COQ5 | 5.44E-01 | 8.19E-01 | 1.03E+00 | 3.60E-02 | 1.03E+00 |
| ENSG000001 | PLXNB2 | 5.44E-01 | 8.19E-01 | 1.03E+00 | 3.94E-02 | 1.03E+00 |
| ENSG000001 | CDK2 | 5.44E-01 | 8.19E-01 | 1.03E+00 | 3.57E-02 | 1.03E+00 |
| ENSG000001 | C1orf174 | 5.44E-01 | 8.19E-01 | 1.02E+00 | 3.50E-02 | 1.02E+00 |
| ENSG000001 | FKBP15 | 5.44E-01 | 8.19E-01 | 1.02E+00 | 2.64E-02 | 1.02E+00 |
| ENSG000001 | CCDC61 | 5.44E-01 | 8.19E-01 | 1.07E+00 | 9.18E-02 | 1.07E+00 |
| ENSG000002 | PNLIPRP3 | 5.44E-01 | 8.19E-01 | 9.27E-01 | -1.09E-01 | -1.08E+00 |
| ENSG000002 | OR10A6 | 5.44E-01 | ? | 8.87E-01 | -1.73E-01 | -1.13E+00 |
| ENSG000001 | PTMAP2 | 5.44E-01 | ? | 8.67E-01 | -2.06E-01 | -1.15E+00 |
| ENSG000002 | ZNF155 | 5.44E-01 | 8.19E-01 | 1.04E+00 | 5.83E-02 | 1.04E+00 |
| ENSG000001 | THEMIS2 | 5.44E-01 | 8.19E-01 | 1.08E+00 | 1.15E-01 | 1.08E+00 |
| ENSG000001 | MIGA1 | 5.44E-01 | 8.19E-01 | 9.80E-01 | -2.99E-02 | -1.02E+00 |
| ENSG000001 | KANSL1 | 5.44E-01 | 8.19E-01 | 9.79E-01 | -3.09E-02 | -1.02E+00 |
| ENSG000002 | GAPDHP43 | 5.44E-01 | ? | 1.24E+00 | 3.13E-01 | 1.24E+00 |
| ENSG000001 | DLL1 | 5.45E-01 | 8.20E-01 | 1.05E+00 | 7.07E-02 | 1.05E+00 |
| ENSG000001 | TOR1AIP2 | 5.45E-01 | 8.20E-01 | 9.83E-01 | -2.42E-02 | -1.02E+00 |
| ENSG000002 | MYLK-AS1 | 5.45E-01 | 8.20E-01 | 1.07E+00 | 1.03E-01 | 1.07E+00 |
| ENSG000001 | MCMDC2 | 5.45E-01 | ? | 1.15E+00 | 2.01E-01 | 1.15E+00 |
| ENSG000002 | --- | 5.45E-01 | ? | 1.28E+00 | 3.53E-01 | 1.28E+00 |
| ENSG000001 | QSOX1 | 5.45E-01 | 8.20E-01 | 1.03E+00 | 4.47E-02 | 1.03E+00 |
| ENSG000001 | IQCE | 5.45E-01 | 8.20E-01 | 9.77E-01 | -3.42E-02 | -1.02E+00 |
| ENSG000002 | --- | 5.45E-01 | ? | 1.44E+00 | 5.23E-01 | 1.44E+00 |
| ENSG000001 | CACNA1D | 5.45E-01 | ? | 1.18E+00 | 2.37E-01 | 1.18E+00 |
| ENSG000001 | NCOR1 | 5.45E-01 | 8.20E-01 | 9.79E-01 | -3.09E-02 | -1.02E+00 |
| ENSG000001 | SYT5 | 5.45E-01 | ? | 1.28E+00 | 3.61E-01 | 1.28E+00 |
| ENSG000002 | HSFX2 | 5.45E-01 | ? | 8.82E-01 | -1.82E-01 | -1.13E+00 |
| ENSG000002 | INTS9-AS1 | 5.45E-01 | ? | 7.67E-01 | -3.83E-01 | -1.30E+00 |
| ENSG000002 | --- | 5.45E-01 | ? | 8.25E-01 | -2.77E-01 | -1.21E+00 |
| ENSG000001 | PLA2G4E | 5.45E-01 | ? | 1.25E+00 | 3.28E-01 | 1.25E+00 |
| ENSG000002 | RAP1BL | 5.46E-01 | 8.21E-01 | 1.09E+00 | 1.26E-01 | 1.09E+00 |
| ENSG000002 | --- | 5.46E-01 | ? | 8.52E-01 | -2.31E-01 | -1.17E+00 |
| ENSG000001 | SHROOM1 | 5.46E-01 | 8.21E-01 | 9.53E-01 | -6.87E-02 | -1.05E+00 |
| ENSG000001 | KAT7 | 5.46E-01 | 8.21E-01 | 1.02E+00 | 2.67E-02 | 1.02E+00 |
| ENSG000002 | ADNP-AS1 | 5.46E-01 | ? | 1.16E+00 | 2.13E-01 | 1.16E+00 |
| ENSG000001 | POR | 5.46E-01 | 8.21E-01 | 1.03E+00 | 4.08E-02 | 1.03E+00 |
| ENSG000001 | BCL2L10 | 5.46E-01 | ? | 1.13E+00 | 1.70E-01 | 1.13E+00 |
| ENSG000002 | DLGAP1-AS2 | 5.46E-01 | 8.21E-01 | 9.48E-01 | -7.68E-02 | -1.05E+00 |
| ENSG000001 | C2orf76 | 5.46E-01 | 8.21E-01 | 9.64E-01 | -5.23E-02 | -1.04E+00 |
| ENSG000002 | LINC01679 | 5.46E-01 | ? | 1.16E+00 | 2.09E-01 | 1.16E+00 |
| ENSG000001 | LRRC32 | 5.46E-01 | ? | 1.30E+00 | 3.80E-01 | 1.30E+00 |
| ENSG000001 | CASP8AP2 | 5.46E-01 | 8.21E-01 | 9.69E-01 | -4.49E-02 | -1.03E+00 |

| | | | | | | |
|---|---|---|---|---|---|---|
| ENSG000001 | TMED5 | 5.46E-01 | 8.21E-01 | 9.77E-01 | -3.31E-02 | -1.02E+00 |
| ENSG000002 | --- | 5.46E-01 | 8.21E-01 | 1.08E+00 | 1.09E-01 | 1.08E+00 |
| ENSG000002 | --- | 5.46E-01 | ? | 1.37E+00 | 4.53E-01 | 1.37E+00 |
| ENSG000001 | XRCC2 | 5.46E-01 | 8.21E-01 | 9.32E-01 | -1.02E-01 | -1.07E+00 |
| ENSG000001 | NME3 | 5.46E-01 | 8.21E-01 | 1.09E+00 | 1.29E-01 | 1.09E+00 |
| ENSG000002 | --- | 5.46E-01 | ? | 1.35E+00 | 4.29E-01 | 1.35E+00 |
| ENSG000001 | ZBTB26 | 5.46E-01 | 8.21E-01 | 9.58E-01 | -6.22E-02 | -1.04E+00 |
| ENSG000002 | --- | 5.47E-01 | ? | 8.41E-01 | -2.50E-01 | -1.19E+00 |
| ENSG000002 | DBNDD2 | 5.47E-01 | 8.21E-01 | 1.03E+00 | 4.24E-02 | 1.03E+00 |
| ENSG000001 | GUSBP13 | 5.47E-01 | ? | 8.26E-01 | -2.75E-01 | -1.21E+00 |
| ENSG000002 | TNS2-AS1 | 5.47E-01 | 8.21E-01 | 1.06E+00 | 8.72E-02 | 1.06E+00 |
| ENSG000002 | --- | 5.47E-01 | ? | 1.21E+00 | 2.78E-01 | 1.21E+00 |
| ENSG000001 | DPH1 | 5.47E-01 | 8.21E-01 | 1.04E+00 | 5.13E-02 | 1.04E+00 |
| ENSG000001 | LAMA2 | 5.47E-01 | 8.21E-01 | 9.46E-01 | -8.03E-02 | -1.06E+00 |
| ENSG000002 | ZNF883 | 5.47E-01 | ? | 1.16E+00 | 2.13E-01 | 1.16E+00 |
| ENSG000002 | CNN3-DT | 5.47E-01 | 8.21E-01 | 9.29E-01 | -1.07E-01 | -1.08E+00 |
| ENSG000002 | MAD2L1-DT | 5.47E-01 | ? | 1.28E+00 | 3.54E-01 | 1.28E+00 |
| ENSG000002 | VIM-AS1 | 5.47E-01 | 8.21E-01 | 1.07E+00 | 9.32E-02 | 1.07E+00 |
| ENSG000001 | EFEMP2 | 5.47E-01 | 8.21E-01 | 1.05E+00 | 6.75E-02 | 1.05E+00 |
| ENSG000001 | PSME3 | 5.47E-01 | 8.21E-01 | 9.88E-01 | -1.76E-02 | -1.01E+00 |
| ENSG000001 | NAF1 | 5.47E-01 | 8.21E-01 | 9.72E-01 | -4.15E-02 | -1.03E+00 |
| ENSG000002 | GP6-AS1 | 5.47E-01 | ? | 1.15E+00 | 1.96E-01 | 1.15E+00 |
| ENSG000002 | --- | 5.47E-01 | ? | 1.33E+00 | 4.10E-01 | 1.33E+00 |
| ENSG000001 | QRSL1 | 5.47E-01 | 8.21E-01 | 9.77E-01 | -3.32E-02 | -1.02E+00 |
| ENSG000001 | PDE6A | 5.47E-01 | ? | 1.22E+00 | 2.92E-01 | 1.22E+00 |
| ENSG000002 | --- | 5.47E-01 | ? | 8.21E-01 | -2.84E-01 | -1.22E+00 |
| ENSG000001 | NUP155 | 5.47E-01 | 8.21E-01 | 1.02E+00 | 3.39E-02 | 1.02E+00 |
| ENSG00000C | MAP4K3 | 5.47E-01 | 8.21E-01 | 9.77E-01 | -3.41E-02 | -1.02E+00 |
| ENSG000001 | MVD | 5.47E-01 | 8.21E-01 | 1.04E+00 | 6.21E-02 | 1.04E+00 |
| ENSG000002 | --- | 5.48E-01 | 8.21E-01 | 9.40E-01 | -8.86E-02 | -1.06E+00 |
| ENSG000001 | CETN3 | 5.48E-01 | 8.21E-01 | 1.03E+00 | 4.48E-02 | 1.03E+00 |
| ENSG000002 | PSMB8 | 5.48E-01 | 8.21E-01 | 1.04E+00 | 5.22E-02 | 1.04E+00 |
| ENSG000001 | MRPS26 | 5.48E-01 | 8.21E-01 | 1.05E+00 | 6.35E-02 | 1.05E+00 |
| ENSG000001 | FRMD4B | 5.48E-01 | 8.21E-01 | 9.78E-01 | -3.25E-02 | -1.02E+00 |
| ENSG000001 | VEGFD | 5.48E-01 | ? | 6.86E-01 | -5.43E-01 | -1.46E+00 |
| ENSG00000C | PPP1R13B | 5.48E-01 | 8.21E-01 | 9.74E-01 | -3.78E-02 | -1.03E+00 |
| ENSG000001 | DOLPP1 | 5.48E-01 | 8.21E-01 | 1.03E+00 | 4.61E-02 | 1.03E+00 |
| ENSG000001 | DUS3L | 5.48E-01 | 8.21E-01 | 1.05E+00 | 6.60E-02 | 1.05E+00 |
| ENSG000001 | SCAF11 | 5.48E-01 | 8.21E-01 | 1.02E+00 | 3.31E-02 | 1.02E+00 |
| ENSG000002 | --- | 5.48E-01 | ? | 8.10E-01 | -3.03E-01 | -1.23E+00 |
| ENSG000001 | SPDEF | 5.48E-01 | 8.22E-01 | 9.58E-01 | -6.13E-02 | -1.04E+00 |
| ENSG00000C | LAP3 | 5.48E-01 | 8.22E-01 | 1.02E+00 | 2.29E-02 | 1.02E+00 |
| ENSG000002 | --- | 5.48E-01 | ? | 1.20E+00 | 2.58E-01 | 1.20E+00 |
| ENSG000001 | L3MBTL1 | 5.48E-01 | 8.22E-01 | 1.06E+00 | 8.31E-02 | 1.06E+00 |
| ENSG000002 | --- | 5.48E-01 | ? | 1.20E+00 | 2.69E-01 | 1.20E+00 |

| | | | | | | |
|---|---|---|---|---|---|---|
| ENSG000001 | ASB3 | 5.48E-01 | 8.22E-01 | 9.64E-01 | -5.28E-02 | -1.04E+00 |
| ENSG000001 | ZNF211 | 5.49E-01 | 8.22E-01 | 9.65E-01 | -5.09E-02 | -1.04E+00 |
| ENSG000001 | ZNF607 | 5.49E-01 | 8.22E-01 | 9.74E-01 | -3.78E-02 | -1.03E+00 |
| ENSG000002 | LINC02068 | 5.49E-01 | ? | 7.43E-01 | -4.29E-01 | -1.35E+00 |
| ENSG000001 | ZBTB6 | 5.49E-01 | 8.22E-01 | 9.75E-01 | -3.66E-02 | -1.03E+00 |
| ENSG000001 | GPR3 | 5.49E-01 | 8.22E-01 | 1.09E+00 | 1.22E-01 | 1.09E+00 |
| ENSG000002 | --- | 5.49E-01 | 8.22E-01 | 9.18E-01 | -1.23E-01 | -1.09E+00 |
| ENSG000002 | FBXO48 | 5.49E-01 | 8.22E-01 | 9.58E-01 | -6.21E-02 | -1.04E+00 |
| ENSG000001 | POLR3F | 5.49E-01 | 8.22E-01 | 1.03E+00 | 3.76E-02 | 1.03E+00 |
| ENSG000002 | --- | 5.49E-01 | ? | 9.04E-01 | -1.46E-01 | -1.11E+00 |
| ENSG000001 | SLC15A4 | 5.49E-01 | 8.22E-01 | 9.82E-01 | -2.66E-02 | -1.02E+00 |
| ENSG000001 | CCNE2 | 5.49E-01 | 8.22E-01 | 8.86E-01 | -1.74E-01 | -1.13E+00 |
| ENSG000001 | ITPRID2 | 5.49E-01 | 8.22E-01 | 1.02E+00 | 3.21E-02 | 1.02E+00 |
| ENSG000002 | --- | 5.49E-01 | ? | 1.42E+00 | 5.07E-01 | 1.42E+00 |
| ENSG000001 | RAD23A | 5.49E-01 | 8.22E-01 | 9.72E-01 | -4.10E-02 | -1.03E+00 |
| ENSG000001 | GATAD2A | 5.49E-01 | 8.22E-01 | 9.87E-01 | -1.93E-02 | -1.01E+00 |
| ENSG000001 | CYP1B1 | 5.49E-01 | 8.22E-01 | 1.05E+00 | 7.52E-02 | 1.05E+00 |
| ENSG000001 | B3GNT5 | 5.49E-01 | 8.22E-01 | 9.79E-01 | -3.11E-02 | -1.02E+00 |
| ENSG000002 | --- | 5.49E-01 | ? | 1.12E+00 | 1.69E-01 | 1.12E+00 |
| ENSG000002 | TMEM185B | 5.49E-01 | 8.22E-01 | 9.82E-01 | -2.63E-02 | -1.02E+00 |
| ENSG000001 | WFDC3 | 5.49E-01 | 8.22E-01 | 8.74E-01 | -1.94E-01 | -1.14E+00 |
| ENSG000001 | SNHG12 | 5.50E-01 | 8.22E-01 | 9.55E-01 | -6.62E-02 | -1.05E+00 |
| ENSG000001 | RAP1GDS1 | 5.50E-01 | 8.22E-01 | 9.87E-01 | -1.85E-02 | -1.01E+00 |
| ENSG000002 | --- | 5.50E-01 | ? | 1.10E+00 | 1.38E-01 | 1.10E+00 |
| ENSG000002 | --- | 5.50E-01 | ? | 7.58E-01 | -4.01E-01 | -1.32E+00 |
| ENSG00000C | SF3B2 | 5.50E-01 | 8.22E-01 | 9.88E-01 | -1.68E-02 | -1.01E+00 |
| ENSG000001 | PLPPR5 | 5.50E-01 | 8.22E-01 | 9.20E-01 | -1.21E-01 | -1.09E+00 |
| ENSG000001 | HRAS | 5.50E-01 | 8.22E-01 | 1.03E+00 | 3.78E-02 | 1.03E+00 |
| ENSG000001 | STK31 | 5.50E-01 | ? | 8.98E-01 | -1.56E-01 | -1.11E+00 |
| ENSG00000C | MRPS10 | 5.50E-01 | 8.22E-01 | 9.84E-01 | -2.39E-02 | -1.02E+00 |
| ENSG000002 | GPC2 | 5.50E-01 | ? | 8.76E-01 | -1.91E-01 | -1.14E+00 |
| ENSG000001 | ELL3 | 5.50E-01 | 8.22E-01 | 1.03E+00 | 4.84E-02 | 1.03E+00 |
| ENSG000002 | HNRNPKP2 | 5.50E-01 | ? | 1.22E+00 | 2.89E-01 | 1.22E+00 |
| ENSG00000C | KIF3C | 5.50E-01 | 8.22E-01 | 9.83E-01 | -2.50E-02 | -1.02E+00 |
| ENSG000001 | LDLRAD2 | 5.50E-01 | 8.22E-01 | 9.09E-01 | -1.37E-01 | -1.10E+00 |
| ENSG000001 | RUNDC3B | 5.50E-01 | ? | 1.17E+00 | 2.24E-01 | 1.17E+00 |
| ENSG00000C | PAK3 | 5.50E-01 | 8.22E-01 | 1.07E+00 | 9.47E-02 | 1.07E+00 |
| ENSG000001 | KIF5B | 5.50E-01 | 8.22E-01 | 9.76E-01 | -3.44E-02 | -1.02E+00 |
| ENSG000002 | --- | 5.50E-01 | ? | 1.19E+00 | 2.56E-01 | 1.19E+00 |
| ENSG000002 | --- | 5.51E-01 | ? | 1.24E+00 | 3.07E-01 | 1.24E+00 |
| ENSG000001 | IGSF8 | 5.51E-01 | 8.22E-01 | 1.05E+00 | 6.46E-02 | 1.05E+00 |
| ENSG000001 | PRR7 | 5.51E-01 | 8.22E-01 | 1.11E+00 | 1.52E-01 | 1.11E+00 |
| ENSG000002 | LCMT1-AS1 | 5.51E-01 | ? | 8.24E-01 | -2.79E-01 | -1.21E+00 |
| ENSG000002 | NUS1P2 | 5.51E-01 | ? | 8.17E-01 | -2.92E-01 | -1.22E+00 |
| ENSG000001 | MED27 | 5.51E-01 | 8.22E-01 | 9.75E-01 | -3.64E-02 | -1.03E+00 |

| | | | | | | | |
|---|---|---|---|---|---|---|---|
| ENSG000001 | PCID2 | 5.51E-01 | | 8.23E-01 | 9.69E-01 | -4.59E-02 | -1.03E+00 |
| ENSG000001 | MATN3 | 5.51E-01 | | 8.23E-01 | 9.52E-01 | -7.09E-02 | -1.05E+00 |
| ENSG000001 | PDZD11 | 5.51E-01 | | 8.23E-01 | 1.02E+00 | 3.06E-02 | 1.02E+00 |
| ENSG000001 | KLRC1 | 5.51E-01 | ? | | 1.21E+00 | 2.78E-01 | 1.21E+00 |
| ENSG000001 | EXOSC4 | 5.51E-01 | | 8.23E-01 | 1.06E+00 | 7.97E-02 | 1.06E+00 |
| ENSG000001 | MAB21L3 | 5.51E-01 | | 8.23E-01 | 1.05E+00 | 7.62E-02 | 1.05E+00 |
| ENSG000002 | LINC01948 | 5.51E-01 | ? | | 7.82E-01 | -3.54E-01 | -1.28E+00 |
| ENSG000002 | --- | 5.51E-01 | ? | | 7.89E-01 | -3.42E-01 | -1.27E+00 |
| ENSG000001 | XIRP2 | 5.51E-01 | ? | | 1.14E+00 | 1.95E-01 | 1.14E+00 |
| ENSG000001 | ADAMTS3 | 5.51E-01 | ? | | 9.09E-01 | -1.38E-01 | -1.10E+00 |
| ENSG000002 | TRIM31 | 5.51E-01 | ? | | 1.21E+00 | 2.77E-01 | 1.21E+00 |
| ENSG000000 | MTFR1 | 5.51E-01 | | 8.23E-01 | 1.02E+00 | 2.84E-02 | 1.02E+00 |
| ENSG000001 | ZNF569 | 5.52E-01 | | 8.23E-01 | 9.68E-01 | -4.67E-02 | -1.03E+00 |
| ENSG000002 | MRPS9-AS1 | 5.52E-01 | ? | | 1.18E+00 | 2.40E-01 | 1.18E+00 |
| ENSG000001 | ITM2C | 5.52E-01 | | 8.23E-01 | 9.68E-01 | -4.67E-02 | -1.03E+00 |
| ENSG000001 | NDUFA9 | 5.52E-01 | | 8.23E-01 | 9.72E-01 | -4.06E-02 | -1.03E+00 |
| ENSG000001 | PDIA3P1 | 5.52E-01 | | 8.23E-01 | 9.67E-01 | -4.88E-02 | -1.03E+00 |
| ENSG000002 | --- | 5.52E-01 | ? | | 8.06E-01 | -3.11E-01 | -1.24E+00 |
| ENSG000001 | C17orf75 | 5.52E-01 | | 8.23E-01 | 9.75E-01 | -3.70E-02 | -1.03E+00 |
| ENSG000001 | NME4 | 5.52E-01 | | 8.23E-01 | 1.02E+00 | 2.66E-02 | 1.02E+00 |
| ENSG000002 | KRT19P1 | 5.52E-01 | ? | | 8.81E-01 | -1.83E-01 | -1.13E+00 |
| ENSG000001 | CYTH1 | 5.52E-01 | | 8.24E-01 | 1.02E+00 | 3.14E-02 | 1.02E+00 |
| ENSG000001 | BBS5 | 5.52E-01 | | 8.24E-01 | 1.04E+00 | 6.34E-02 | 1.04E+00 |
| ENSG000001 | TM2D1 | 5.52E-01 | | 8.24E-01 | 9.78E-01 | -3.15E-02 | -1.02E+00 |
| ENSG000002 | GPR179 | 5.52E-01 | ? | | 1.22E+00 | 2.81E-01 | 1.22E+00 |
| ENSG000002 | RPS27AP5 | 5.52E-01 | ? | | 1.12E+00 | 1.59E-01 | 1.12E+00 |
| ENSG000001 | GLUD1 | 5.52E-01 | | 8.24E-01 | 9.85E-01 | -2.17E-02 | -1.02E+00 |
| ENSG000001 | FAM72A | 5.52E-01 | | 8.24E-01 | 1.08E+00 | 1.15E-01 | 1.08E+00 |
| ENSG000002 | PAXIP1-AS2 | 5.52E-01 | | 8.24E-01 | 9.60E-01 | -5.85E-02 | -1.04E+00 |
| ENSG000002 | --- | 5.52E-01 | ? | | 1.51E+00 | 5.97E-01 | 1.51E+00 |
| ENSG000002 | LINC01191 | 5.52E-01 | ? | | 8.42E-01 | -2.49E-01 | -1.19E+00 |
| ENSG000001 | APOO | 5.53E-01 | | 8.24E-01 | 1.05E+00 | 6.67E-02 | 1.05E+00 |
| ENSG000002 | --- | 5.53E-01 | ? | | 1.65E+00 | 7.19E-01 | 1.65E+00 |
| ENSG000000 | TXLNG | 5.53E-01 | | 8.24E-01 | 1.02E+00 | 3.36E-02 | 1.02E+00 |
| ENSG000001 | GLIPR1L2 | 5.53E-01 | ? | | 8.18E-01 | -2.90E-01 | -1.22E+00 |
| ENSG000002 | GET1-SH3BG | 5.53E-01 | ? | | 1.40E+00 | 4.87E-01 | 1.40E+00 |
| ENSG000002 | --- | 5.53E-01 | ? | | 7.34E-01 | -4.47E-01 | -1.36E+00 |
| ENSG000002 | ZNF350-AS1 | 5.53E-01 | ? | | 1.23E+00 | 3.04E-01 | 1.23E+00 |
| ENSG000001 | RNF149 | 5.53E-01 | | 8.25E-01 | 9.80E-01 | -2.92E-02 | -1.02E+00 |
| ENSG000002 | ZNF853 | 5.53E-01 | ? | | 8.51E-01 | -2.33E-01 | -1.18E+00 |
| ENSG000002 | NFYC-AS1 | 5.54E-01 | ? | | 8.63E-01 | -2.13E-01 | -1.16E+00 |
| ENSG000001 | HAX1 | 5.54E-01 | | 8.25E-01 | 1.02E+00 | 3.22E-02 | 1.02E+00 |
| ENSG000001 | PPHLN1 | 5.54E-01 | | 8.25E-01 | 9.82E-01 | -2.59E-02 | -1.02E+00 |
| ENSG000001 | UGCG | 5.54E-01 | | 8.25E-01 | 1.03E+00 | 3.90E-02 | 1.03E+00 |
| ENSG000002 | OR7E7P | 5.54E-01 | ? | | 1.24E+00 | 3.09E-01 | 1.24E+00 |

| | | | | | | |
|---|---|---|---|---|---|---|
| ENSG000002 | --- | 5.54E-01 | ? | 8.85E-01 | -1.77E-01 | -1.13E+00 |
| ENSG000001 | PPM1L | 5.54E-01 | 8.25E-01 | 9.58E-01 | -6.20E-02 | -1.04E+00 |
| ENSG000002 | --- | 5.54E-01 | ? | 1.17E+00 | 2.21E-01 | 1.17E+00 |
| ENSG000001 | ARHGAP32 | 5.54E-01 | 8.25E-01 | 9.79E-01 | -3.12E-02 | -1.02E+00 |
| ENSG000002 | --- | 5.54E-01 | 8.25E-01 | 1.05E+00 | 6.60E-02 | 1.05E+00 |
| ENSG000002 | APPBP2-DT | 5.54E-01 | ? | 8.18E-01 | -2.90E-01 | -1.22E+00 |
| ENSG000002 | DAXX | 5.54E-01 | 8.25E-01 | 1.01E+00 | 2.12E-02 | 1.01E+00 |
| ENSG000001 | ALDH8A1 | 5.54E-01 | ? | 8.07E-01 | -3.09E-01 | -1.24E+00 |
| ENSG000001 | PCDHB13 | 5.54E-01 | ? | 1.15E+00 | 1.99E-01 | 1.15E+00 |
| ENSG000001 | KTN1-AS1 | 5.54E-01 | 8.25E-01 | 1.06E+00 | 8.07E-02 | 1.06E+00 |
| ENSG000002 | --- | 5.55E-01 | ? | 7.88E-01 | -3.43E-01 | -1.27E+00 |
| ENSG00000C | HMMR | 5.55E-01 | 8.25E-01 | 9.26E-01 | -1.10E-01 | -1.08E+00 |
| ENSG000001 | A4GALT | 5.55E-01 | 8.25E-01 | 9.60E-01 | -5.96E-02 | -1.04E+00 |
| ENSG000001 | ZNF654 | 5.55E-01 | 8.25E-01 | 9.78E-01 | -3.15E-02 | -1.02E+00 |
| ENSG000002 | EFCAB7 | 5.55E-01 | 8.25E-01 | 9.50E-01 | -7.46E-02 | -1.05E+00 |
| ENSG000002 | CASTOR2 | 5.55E-01 | 8.25E-01 | 1.03E+00 | 4.68E-02 | 1.03E+00 |
| ENSG00000C | ZFHX4 | 5.55E-01 | 8.25E-01 | 9.20E-01 | -1.20E-01 | -1.09E+00 |
| ENSG000001 | TXNL4A | 5.55E-01 | 8.25E-01 | 1.02E+00 | 2.51E-02 | 1.02E+00 |
| ENSG000001 | TCP11L2 | 5.55E-01 | 8.25E-01 | 1.05E+00 | 6.72E-02 | 1.05E+00 |
| ENSG00000C | FOXJ2 | 5.55E-01 | 8.25E-01 | 9.79E-01 | -3.05E-02 | -1.02E+00 |
| ENSG000001 | MAPK11 | 5.55E-01 | 8.25E-01 | 1.05E+00 | 6.44E-02 | 1.05E+00 |
| ENSG000001 | ANTXR1 | 5.55E-01 | 8.25E-01 | 1.02E+00 | 3.55E-02 | 1.02E+00 |
| ENSG000002 | --- | 5.55E-01 | ? | 8.49E-01 | -2.36E-01 | -1.18E+00 |
| ENSG000001 | NUBP1 | 5.55E-01 | 8.25E-01 | 9.78E-01 | -3.27E-02 | -1.02E+00 |
| ENSG000001 | VAPB | 5.55E-01 | 8.25E-01 | 9.82E-01 | -2.61E-02 | -1.02E+00 |
| ENSG000002 | --- | 5.55E-01 | ? | 1.15E+00 | 2.07E-01 | 1.15E+00 |
| ENSG000002 | FAM86JP | 5.55E-01 | 8.25E-01 | 1.05E+00 | 7.39E-02 | 1.05E+00 |
| ENSG000001 | BTNL9 | 5.55E-01 | 8.25E-01 | 1.10E+00 | 1.39E-01 | 1.10E+00 |
| ENSG000002 | OVCA2 | 5.55E-01 | 8.25E-01 | 1.03E+00 | 4.66E-02 | 1.03E+00 |
| ENSG000001 | LYPD5 | 5.55E-01 | ? | 1.12E+00 | 1.67E-01 | 1.12E+00 |
| ENSG000002 | KMT2B | 5.55E-01 | 8.25E-01 | 9.77E-01 | -3.40E-02 | -1.02E+00 |
| ENSG000001 | DNASE1L2 | 5.55E-01 | ? | 8.53E-01 | -2.29E-01 | -1.17E+00 |
| ENSG00000C | LTBP4 | 5.55E-01 | 8.25E-01 | 1.04E+00 | 5.28E-02 | 1.04E+00 |
| ENSG000001 | RAP2B | 5.55E-01 | 8.25E-01 | 1.02E+00 | 2.46E-02 | 1.02E+00 |
| ENSG000002 | ARPIN | 5.55E-01 | 8.25E-01 | 1.02E+00 | 3.51E-02 | 1.02E+00 |
| ENSG000002 | --- | 5.56E-01 | ? | 1.32E+00 | 3.99E-01 | 1.32E+00 |
| ENSG000001 | PEX2 | 5.56E-01 | 8.26E-01 | 1.02E+00 | 3.38E-02 | 1.02E+00 |
| ENSG000001 | IFT122 | 5.56E-01 | 8.26E-01 | 1.05E+00 | 7.33E-02 | 1.05E+00 |
| ENSG000001 | FBXO36 | 5.56E-01 | 8.26E-01 | 1.06E+00 | 8.42E-02 | 1.06E+00 |
| ENSG000002 | --- | 5.56E-01 | ? | 1.30E+00 | 3.82E-01 | 1.30E+00 |
| ENSG00000C | LRRC7 | 5.56E-01 | ? | 1.15E+00 | 2.01E-01 | 1.15E+00 |
| ENSG000002 | SUMO2P1 | 5.56E-01 | ? | 1.14E+00 | 1.85E-01 | 1.14E+00 |
| ENSG000001 | CLHC1 | 5.56E-01 | 8.26E-01 | 9.57E-01 | -6.39E-02 | -1.05E+00 |
| ENSG000002 | --- | 5.56E-01 | ? | 1.39E+00 | 4.71E-01 | 1.39E+00 |
| ENSG000001 | CDH18 | 5.56E-01 | ? | 1.28E+00 | 3.59E-01 | 1.28E+00 |

| | | | | | | |
|---|---|---|---|---|---|---|
| ENSG000001 | NCMAP | 5.56E-01 | ? | 1.30E+00 | 3.74E-01 | 1.30E+00 |
| ENSG000001 | FBXO27 | 5.56E-01 | 8.26E-01 | 1.04E+00 | 5.52E-02 | 1.04E+00 |
| ENSG000001 | TFPI2 | 5.56E-01 | 8.26E-01 | 1.12E+00 | 1.59E-01 | 1.12E+00 |
| ENSG000002 | --- | 5.56E-01 | ? | 8.06E-01 | -3.11E-01 | -1.24E+00 |
| ENSG000001 | PTPRJ | 5.56E-01 | 8.26E-01 | 1.04E+00 | 5.06E-02 | 1.04E+00 |
| ENSG000002 | --- | 5.57E-01 | 8.26E-01 | 9.17E-01 | -1.24E-01 | -1.09E+00 |
| ENSG000002 | MTA1-DT | 5.57E-01 | 8.26E-01 | 9.23E-01 | -1.16E-01 | -1.08E+00 |
| ENSG000001 | C21orf58 | 5.57E-01 | 8.26E-01 | 1.07E+00 | 1.02E-01 | 1.07E+00 |
| ENSG000002 | EIF4EP1 | 5.57E-01 | ? | 7.36E-01 | -4.43E-01 | -1.36E+00 |
| ENSG000002 | RPL18AP7 | 5.57E-01 | ? | 7.76E-01 | -3.66E-01 | -1.29E+00 |
| ENSG000001 | GEN1 | 5.57E-01 | 8.26E-01 | 9.70E-01 | -4.43E-02 | -1.03E+00 |
| ENSG000002 | ZNF516-AS1 | 5.57E-01 | ? | 1.13E+00 | 1.76E-01 | 1.13E+00 |
| ENSG000002 | BAZ1A-AS1 | 5.57E-01 | 8.26E-01 | 1.10E+00 | 1.35E-01 | 1.10E+00 |
| ENSG000002 | MIR503HG | 5.57E-01 | 8.26E-01 | 9.25E-01 | -1.13E-01 | -1.08E+00 |
| ENSG000002 | MIR181A2HG | 5.57E-01 | ? | 9.13E-01 | -1.31E-01 | -1.10E+00 |
| ENSG000001 | TMEM52 | 5.57E-01 | ? | 1.32E+00 | 4.03E-01 | 1.32E+00 |
| ENSG000002 | GSEC | 5.57E-01 | 8.26E-01 | 1.06E+00 | 7.99E-02 | 1.06E+00 |
| ENSG000002 | SNX18P12 | 5.57E-01 | ? | 1.14E+00 | 1.93E-01 | 1.14E+00 |
| ENSG000001 | FDX1 | 5.57E-01 | 8.26E-01 | 9.73E-01 | -3.91E-02 | -1.03E+00 |
| ENSG000001 | TBX6 | 5.57E-01 | ? | 8.44E-01 | -2.45E-01 | -1.19E+00 |
| ENSG000002 | --- | 5.57E-01 | 8.26E-01 | 9.14E-01 | -1.29E-01 | -1.09E+00 |
| ENSG000001 | IDH2 | 5.57E-01 | 8.26E-01 | 1.05E+00 | 6.96E-02 | 1.05E+00 |
| ENSG000001 | CTNNA3 | 5.57E-01 | ? | 1.12E+00 | 1.57E-01 | 1.12E+00 |
| ENSG000000 | SMAP2 | 5.57E-01 | 8.26E-01 | 1.03E+00 | 4.06E-02 | 1.03E+00 |
| ENSG000002 | --- | 5.57E-01 | ? | 1.16E+00 | 2.13E-01 | 1.16E+00 |
| ENSG000001 | CLCN2 | 5.57E-01 | 8.26E-01 | 9.53E-01 | -6.97E-02 | -1.05E+00 |
| ENSG000000 | KDM1A | 5.57E-01 | 8.26E-01 | 1.01E+00 | 1.93E-02 | 1.01E+00 |
| ENSG000000 | C3orf18 | 5.57E-01 | 8.26E-01 | 9.39E-01 | -9.03E-02 | -1.06E+00 |
| ENSG000001 | MAFK | 5.58E-01 | 8.27E-01 | 9.68E-01 | -4.63E-02 | -1.03E+00 |
| ENSG000001 | RAD51AP1 | 5.58E-01 | 8.27E-01 | 9.29E-01 | -1.06E-01 | -1.08E+00 |
| ENSG000002 | LINC01179 | 5.58E-01 | ? | 8.53E-01 | -2.29E-01 | -1.17E+00 |
| ENSG000002 | TMSB4XP4 | 5.58E-01 | ? | 8.55E-01 | -2.25E-01 | -1.17E+00 |
| ENSG000002 | --- | 5.58E-01 | ? | 8.34E-01 | -2.62E-01 | -1.20E+00 |
| ENSG000002 | PLBD1-AS1 | 5.58E-01 | ? | 1.30E+00 | 3.81E-01 | 1.30E+00 |
| ENSG000001 | RMDN2 | 5.58E-01 | 8.27E-01 | 1.03E+00 | 4.93E-02 | 1.03E+00 |
| ENSG000002 | LRRC37A16P | 5.58E-01 | 8.27E-01 | 9.48E-01 | -7.69E-02 | -1.05E+00 |
| ENSG000001 | SLC26A1 | 5.58E-01 | ? | 8.69E-01 | -2.03E-01 | -1.15E+00 |
| ENSG000000 | ANAPC4 | 5.58E-01 | 8.27E-01 | 9.77E-01 | -3.40E-02 | -1.02E+00 |
| ENSG000001 | ANKRD13D | 5.58E-01 | 8.27E-01 | 9.59E-01 | -5.99E-02 | -1.04E+00 |
| ENSG000001 | PLEKHA8 | 5.58E-01 | 8.27E-01 | 9.77E-01 | -3.33E-02 | -1.02E+00 |
| ENSG000001 | MAP3K10 | 5.58E-01 | 8.27E-01 | 1.05E+00 | 6.92E-02 | 1.05E+00 |
| ENSG000001 | NEMP1 | 5.58E-01 | 8.27E-01 | 1.02E+00 | 2.69E-02 | 1.02E+00 |
| ENSG000001 | COPG2 | 5.58E-01 | 8.27E-01 | 1.02E+00 | 3.03E-02 | 1.02E+00 |
| ENSG000002 | RBM20 | 5.58E-01 | ? | 8.47E-01 | -2.40E-01 | -1.18E+00 |
| ENSG000001 | NOP9 | 5.58E-01 | 8.27E-01 | 1.02E+00 | 2.76E-02 | 1.02E+00 |

| | | | | | | |
|---|---|---|---|---|---|---|
| ENSG000002 | --- | 5.58E-01 | ? | 1.11E+00 | 1.50E-01 | 1.11E+00 |
| ENSG000001 | CCND3 | 5.58E-01 | 8.27E-01 | 1.04E+00 | 4.97E-02 | 1.04E+00 |
| ENSG000001 | EDA2R | 5.58E-01 | 8.27E-01 | 1.04E+00 | 5.29E-02 | 1.04E+00 |
| ENSG000002 | --- | 5.58E-01 | 8.27E-01 | 9.16E-01 | -1.27E-01 | -1.09E+00 |
| ENSG000001 | NELFA | 5.58E-01 | 8.27E-01 | 1.03E+00 | 4.38E-02 | 1.03E+00 |
| ENSG000002 | --- | 5.59E-01 | ? | 1.34E+00 | 4.23E-01 | 1.34E+00 |
| ENSG000002 | --- | 5.59E-01 | ? | 1.44E+00 | 5.28E-01 | 1.44E+00 |
| ENSG000002 | ARMCX5-GPI | 5.59E-01 | 8.27E-01 | 1.06E+00 | 8.98E-02 | 1.06E+00 |
| ENSG000002 | RPL7AP70 | 5.59E-01 | ? | 1.33E+00 | 4.15E-01 | 1.33E+00 |
| ENSG000001 | ZNF30 | 5.59E-01 | 8.27E-01 | 9.57E-01 | -6.41E-02 | -1.05E+00 |
| ENSG000001 | GREB1 | 5.59E-01 | 8.27E-01 | 1.10E+00 | 1.39E-01 | 1.10E+00 |
| ENSG000002 | PFDN6 | 5.59E-01 | 8.27E-01 | 1.02E+00 | 2.89E-02 | 1.02E+00 |
| ENSG000001 | SWI5 | 5.59E-01 | 8.27E-01 | 9.71E-01 | -4.27E-02 | -1.03E+00 |
| ENSG000001 | FAAP20 | 5.59E-01 | 8.27E-01 | 1.05E+00 | 6.67E-02 | 1.05E+00 |
| ENSG00000C | TEAD3 | 5.59E-01 | 8.27E-01 | 1.02E+00 | 3.33E-02 | 1.02E+00 |
| ENSG000002 | TBCE | 5.59E-01 | 8.27E-01 | 1.04E+00 | 5.63E-02 | 1.04E+00 |
| ENSG000001 | LRRC8A | 5.59E-01 | 8.27E-01 | 1.02E+00 | 2.53E-02 | 1.02E+00 |
| ENSG000001 | PCLO | 5.59E-01 | 8.27E-01 | 9.56E-01 | -6.47E-02 | -1.05E+00 |
| ENSG000001 | SGO1 | 5.59E-01 | 8.27E-01 | 9.21E-01 | -1.18E-01 | -1.09E+00 |
| ENSG000002 | SUV39H2-DT | 5.59E-01 | ? | 1.19E+00 | 2.47E-01 | 1.19E+00 |
| ENSG000001 | MRPL30 | 5.59E-01 | 8.27E-01 | 1.02E+00 | 2.95E-02 | 1.02E+00 |
| ENSG000002 | USP34-DT | 5.60E-01 | 8.27E-01 | 9.38E-01 | -9.22E-02 | -1.07E+00 |
| ENSG00000C | GPRC5A | 5.60E-01 | 8.27E-01 | 9.77E-01 | -3.39E-02 | -1.02E+00 |
| ENSG000001 | COQ7 | 5.60E-01 | 8.27E-01 | 9.75E-01 | -3.73E-02 | -1.03E+00 |
| ENSG000001 | POPDC2 | 5.60E-01 | ? | 1.14E+00 | 1.91E-01 | 1.14E+00 |
| ENSG000002 | LIMS4 | 5.60E-01 | 8.28E-01 | 1.12E+00 | 1.66E-01 | 1.12E+00 |
| ENSG000001 | KRT79 | 5.60E-01 | ? | 8.01E-01 | -3.20E-01 | -1.25E+00 |
| ENSG000001 | ARSA | 5.60E-01 | 8.28E-01 | 1.05E+00 | 6.38E-02 | 1.05E+00 |
| ENSG000002 | MIA2-AS1 | 5.60E-01 | ? | 1.26E+00 | 3.33E-01 | 1.26E+00 |
| ENSG000002 | GAS5-AS1 | 5.60E-01 | ? | 8.46E-01 | -2.42E-01 | -1.18E+00 |
| ENSG000001 | PRPF40B | 5.60E-01 | 8.28E-01 | 1.03E+00 | 4.01E-02 | 1.03E+00 |
| ENSG000001 | MECP2 | 5.60E-01 | 8.28E-01 | 9.82E-01 | -2.60E-02 | -1.02E+00 |
| ENSG000001 | ZNF775 | 5.60E-01 | 8.28E-01 | 1.08E+00 | 1.13E-01 | 1.08E+00 |
| ENSG00000C | GCLC | 5.60E-01 | 8.28E-01 | 9.71E-01 | -4.23E-02 | -1.03E+00 |
| ENSG000002 | SLC26A6 | 5.60E-01 | 8.28E-01 | 9.67E-01 | -4.85E-02 | -1.03E+00 |
| ENSG00000C | TGFBR3 | 5.60E-01 | 8.28E-01 | 9.54E-01 | -6.83E-02 | -1.05E+00 |
| ENSG000001 | ANAPC2 | 5.60E-01 | 8.28E-01 | 1.05E+00 | 6.49E-02 | 1.05E+00 |
| ENSG000002 | --- | 5.61E-01 | ? | 9.01E-01 | -1.50E-01 | -1.11E+00 |
| ENSG000002 | NPY4R | 5.61E-01 | ? | 8.56E-01 | -2.25E-01 | -1.17E+00 |
| ENSG000002 | ANXA2R-AS1 | 5.61E-01 | ? | 8.89E-01 | -1.69E-01 | -1.12E+00 |
| ENSG000001 | ABTB3 | 5.61E-01 | 8.28E-01 | 9.68E-01 | -4.64E-02 | -1.03E+00 |
| ENSG000001 | VSTM2L | 5.61E-01 | 8.28E-01 | 9.50E-01 | -7.36E-02 | -1.05E+00 |
| ENSG000001 | GPATCH8 | 5.61E-01 | 8.28E-01 | 9.81E-01 | -2.70E-02 | -1.02E+00 |
| ENSG000001 | KIAA1328 | 5.61E-01 | 8.28E-01 | 9.48E-01 | -7.69E-02 | -1.05E+00 |
| ENSG000002 | --- | 5.61E-01 | ? | 8.02E-01 | -3.19E-01 | -1.25E+00 |

| | | | | | | |
|---|---|---|---|---|---|---|
| ENSG000002 | PPT2 | 5.61E-01 | 8.28E-01 | 1.04E+00 | 6.29E-02 | 1.04E+00 |
| ENSG000001 | NME7 | 5.61E-01 | 8.28E-01 | 1.03E+00 | 4.61E-02 | 1.03E+00 |
| ENSG000001 | ASAP1 | 5.61E-01 | 8.28E-01 | 1.03E+00 | 3.94E-02 | 1.03E+00 |
| ENSG000001 | CSAD | 5.61E-01 | 8.28E-01 | 9.23E-01 | -1.16E-01 | -1.08E+00 |
| ENSG000001 | MARCHF5 | 5.61E-01 | 8.28E-01 | 1.02E+00 | 2.48E-02 | 1.02E+00 |
| ENSG000001 | WDR4 | 5.61E-01 | 8.28E-01 | 1.04E+00 | 5.22E-02 | 1.04E+00 |
| ENSG000002 | ACTG1P20 | 5.61E-01 | ? | 7.94E-01 | -3.33E-01 | -1.26E+00 |
| ENSG000002 | GNG12-AS1 | 5.61E-01 | ? | 8.00E-01 | -3.22E-01 | -1.25E+00 |
| ENSG000001 | CD200R1 | 5.62E-01 | ? | 1.25E+00 | 3.24E-01 | 1.25E+00 |
| ENSG000002 | NSUN5P1 | 5.62E-01 | 8.28E-01 | 9.53E-01 | -6.99E-02 | -1.05E+00 |
| ENSG000001 | PUS1 | 5.62E-01 | 8.28E-01 | 1.03E+00 | 4.59E-02 | 1.03E+00 |
| ENSG000002 | CEMP1 | 5.62E-01 | ? | 1.61E+00 | 6.84E-01 | 1.61E+00 |
| ENSG000002 | BHLHE40-AS | 5.62E-01 | ? | 1.16E+00 | 2.19E-01 | 1.16E+00 |
| ENSG00000C | BRAP | 5.62E-01 | 8.28E-01 | 1.02E+00 | 2.85E-02 | 1.02E+00 |
| ENSG000001 | SLC30A7 | 5.62E-01 | 8.28E-01 | 9.79E-01 | -3.04E-02 | -1.02E+00 |
| ENSG000001 | MFSD5 | 5.62E-01 | 8.28E-01 | 1.04E+00 | 5.75E-02 | 1.04E+00 |
| ENSG000001 | IDH3B | 5.62E-01 | 8.28E-01 | 9.80E-01 | -2.87E-02 | -1.02E+00 |
| ENSG000002 | CD101-AS1 | 5.62E-01 | ? | 1.24E+00 | 3.11E-01 | 1.24E+00 |
| ENSG000001 | SLC6A20 | 5.62E-01 | ? | 1.23E+00 | 2.97E-01 | 1.23E+00 |
| ENSG000001 | TBL3 | 5.62E-01 | 8.28E-01 | 1.04E+00 | 5.48E-02 | 1.04E+00 |
| ENSG00000C | XRCC5 | 5.62E-01 | 8.28E-01 | 9.86E-01 | -2.10E-02 | -1.01E+00 |
| ENSG000002 | --- | 5.62E-01 | ? | 7.64E-01 | -3.87E-01 | -1.31E+00 |
| ENSG000001 | CBX2 | 5.62E-01 | 8.28E-01 | 1.05E+00 | 7.63E-02 | 1.05E+00 |
| ENSG000001 | DUSP10 | 5.62E-01 | 8.28E-01 | 1.03E+00 | 4.67E-02 | 1.03E+00 |
| ENSG000002 | --- | 5.62E-01 | ? | 8.51E-01 | -2.33E-01 | -1.18E+00 |
| ENSG000001 | IRF2BPL | 5.62E-01 | 8.28E-01 | 1.04E+00 | 6.06E-02 | 1.04E+00 |
| ENSG00000C | CLK1 | 5.62E-01 | 8.28E-01 | 9.63E-01 | -5.41E-02 | -1.04E+00 |
| ENSG00000C | MYDGF | 5.62E-01 | 8.28E-01 | 1.03E+00 | 4.46E-02 | 1.03E+00 |
| ENSG000001 | EIF3J | 5.62E-01 | 8.28E-01 | 9.74E-01 | -3.74E-02 | -1.03E+00 |
| ENSG000001 | RECQL4 | 5.62E-01 | 8.28E-01 | 1.09E+00 | 1.30E-01 | 1.09E+00 |
| ENSG000002 | MAGIX | 5.62E-01 | ? | 1.13E+00 | 1.78E-01 | 1.13E+00 |
| ENSG000001 | NSMCE4A | 5.63E-01 | 8.28E-01 | 1.03E+00 | 3.67E-02 | 1.03E+00 |
| ENSG000002 | --- | 5.63E-01 | ? | 1.21E+00 | 2.78E-01 | 1.21E+00 |
| ENSG000001 | GK | 5.63E-01 | 8.28E-01 | 9.52E-01 | -7.03E-02 | -1.05E+00 |
| ENSG000001 | EGF | 5.63E-01 | ? | 8.66E-01 | -2.07E-01 | -1.15E+00 |
| ENSG000002 | PFN1P1 | 5.63E-01 | ? | 1.29E+00 | 3.72E-01 | 1.29E+00 |
| ENSG00000C | PHRF1 | 5.63E-01 | 8.29E-01 | 9.82E-01 | -2.59E-02 | -1.02E+00 |
| ENSG000001 | TUFT1 | 5.63E-01 | 8.29E-01 | 9.85E-01 | -2.13E-02 | -1.01E+00 |
| ENSG000001 | ADGRL3 | 5.63E-01 | ? | 1.19E+00 | 2.54E-01 | 1.19E+00 |
| ENSG000001 | CFAP43 | 5.63E-01 | ? | 9.12E-01 | -1.33E-01 | -1.10E+00 |
| ENSG000001 | OGT | 5.63E-01 | 8.29E-01 | 9.67E-01 | -4.89E-02 | -1.03E+00 |
| ENSG000001 | MEIS1 | 5.63E-01 | 8.29E-01 | 1.04E+00 | 5.45E-02 | 1.04E+00 |
| ENSG000001 | RAMP1 | 5.63E-01 | ? | 1.14E+00 | 1.84E-01 | 1.14E+00 |
| ENSG000001 | SV2C | 5.63E-01 | ? | 8.90E-01 | -1.69E-01 | -1.12E+00 |
| ENSG000002 | TNKS2-DT | 5.64E-01 | ? | 1.26E+00 | 3.30E-01 | 1.26E+00 |

| | | | | | | |
|---|---|---|---|---|---|---|
| ENSG000001 | CCNA1 | 5.64E-01 | 8.30E-01 | 1.08E+00 | 1.16E-01 | 1.08E+00 |
| ENSG000002 | --- | 5.64E-01 | ? | 1.33E+00 | 4.12E-01 | 1.33E+00 |
| ENSG000001 | NLN | 5.64E-01 | 8.30E-01 | 9.78E-01 | -3.20E-02 | -1.02E+00 |
| ENSG000002 | TNPO1P2 | 5.64E-01 | ? | 1.16E+00 | 2.08E-01 | 1.16E+00 |
| ENSG000001 | KRT13 | 5.64E-01 | 8.30E-01 | 1.08E+00 | 1.16E-01 | 1.08E+00 |
| ENSG00000C | THUMPD1 | 5.64E-01 | 8.30E-01 | 1.02E+00 | 2.97E-02 | 1.02E+00 |
| ENSG000002 | EMP2 | 5.64E-01 | 8.30E-01 | 9.71E-01 | -4.32E-02 | -1.03E+00 |
| ENSG000002 | --- | 5.64E-01 | ? | 1.20E+00 | 2.67E-01 | 1.20E+00 |
| ENSG000001 | RGS4 | 5.64E-01 | ? | 1.15E+00 | 2.07E-01 | 1.15E+00 |
| ENSG000001 | SPOCK3 | 5.64E-01 | ? | 1.13E+00 | 1.78E-01 | 1.13E+00 |
| ENSG000001 | BEX4 | 5.64E-01 | 8.30E-01 | 9.75E-01 | -3.61E-02 | -1.03E+00 |
| ENSG000001 | ZNF682 | 5.64E-01 | 8.30E-01 | 9.47E-01 | -7.88E-02 | -1.06E+00 |
| ENSG000001 | ZNF142 | 5.64E-01 | 8.30E-01 | 9.79E-01 | -3.05E-02 | -1.02E+00 |
| ENSG000001 | KCNE1 | 5.64E-01 | ? | 7.09E-01 | -4.97E-01 | -1.41E+00 |
| ENSG00000C | PKD1 | 5.65E-01 | 8.30E-01 | 1.04E+00 | 6.30E-02 | 1.04E+00 |
| ENSG000002 | --- | 5.65E-01 | 8.30E-01 | 9.10E-01 | -1.35E-01 | -1.10E+00 |
| ENSG000002 | HNRNPKP1 | 5.65E-01 | ? | 1.39E+00 | 4.79E-01 | 1.39E+00 |
| ENSG000002 | FNBP1P1 | 5.65E-01 | 8.30E-01 | 1.09E+00 | 1.26E-01 | 1.09E+00 |
| ENSG00000C | SEC61A2 | 5.65E-01 | 8.30E-01 | 9.64E-01 | -5.28E-02 | -1.04E+00 |
| ENSG00000C | ZXDC | 5.65E-01 | 8.30E-01 | 1.02E+00 | 3.20E-02 | 1.02E+00 |
| ENSG000001 | FAM167A | 5.65E-01 | 8.30E-01 | 1.12E+00 | 1.66E-01 | 1.12E+00 |
| ENSG000002 | --- | 5.65E-01 | ? | 1.33E+00 | 4.13E-01 | 1.33E+00 |
| ENSG000002 | --- | 5.65E-01 | 8.30E-01 | 1.11E+00 | 1.56E-01 | 1.11E+00 |
| ENSG00000C | RASGRP2 | 5.65E-01 | ? | 7.69E-01 | -3.80E-01 | -1.30E+00 |
| ENSG000001 | SLC1A1 | 5.65E-01 | 8.30E-01 | 9.66E-01 | -4.92E-02 | -1.03E+00 |
| ENSG000001 | ZNF641 | 5.65E-01 | 8.30E-01 | 1.03E+00 | 3.93E-02 | 1.03E+00 |
| ENSG000001 | DOP1B | 5.65E-01 | 8.30E-01 | 1.02E+00 | 3.54E-02 | 1.02E+00 |
| ENSG000001 | SND1 | 5.65E-01 | 8.30E-01 | 9.83E-01 | -2.49E-02 | -1.02E+00 |
| ENSG000001 | LARP6 | 5.65E-01 | 8.30E-01 | 1.04E+00 | 5.75E-02 | 1.04E+00 |
| ENSG000002 | --- | 5.65E-01 | ? | 1.22E+00 | 2.83E-01 | 1.22E+00 |
| ENSG000002 | --- | 5.65E-01 | ? | 8.20E-01 | -2.86E-01 | -1.22E+00 |
| ENSG000001 | PELP1 | 5.65E-01 | 8.30E-01 | 1.03E+00 | 3.63E-02 | 1.03E+00 |
| ENSG000001 | BABAM2 | 5.65E-01 | 8.30E-01 | 1.03E+00 | 3.75E-02 | 1.03E+00 |
| ENSG00000C | IRAK3 | 5.65E-01 | 8.30E-01 | 1.05E+00 | 6.67E-02 | 1.05E+00 |
| ENSG000002 | --- | 5.65E-01 | 8.30E-01 | 1.31E+00 | 3.86E-01 | 1.31E+00 |
| ENSG000001 | FAIM | 5.66E-01 | 8.30E-01 | 9.60E-01 | -5.88E-02 | -1.04E+00 |
| ENSG000001 | CORIN | 5.66E-01 | ? | 9.01E-01 | -1.51E-01 | -1.11E+00 |
| ENSG000002 | YWHAZP3 | 5.66E-01 | 8.30E-01 | 1.10E+00 | 1.36E-01 | 1.10E+00 |
| ENSG000001 | KIAA0408 | 5.66E-01 | ? | 1.21E+00 | 2.71E-01 | 1.21E+00 |
| ENSG000001 | TNFSF13B | 5.66E-01 | ? | 9.11E-01 | -1.35E-01 | -1.10E+00 |
| ENSG000001 | ZNF236 | 5.66E-01 | 8.30E-01 | 9.76E-01 | -3.55E-02 | -1.02E+00 |
| ENSG000001 | MTF2 | 5.66E-01 | 8.30E-01 | 9.75E-01 | -3.61E-02 | -1.03E+00 |
| ENSG000001 | PLAC1 | 5.66E-01 | ? | 8.75E-01 | -1.93E-01 | -1.14E+00 |
| ENSG000001 | SLPI | 5.66E-01 | 8.30E-01 | 1.07E+00 | 9.99E-02 | 1.07E+00 |
| ENSG000001 | KMT2A | 5.66E-01 | 8.30E-01 | 9.61E-01 | -5.76E-02 | -1.04E+00 |

| | | | | | | |
|---|---|---|---|---|---|---|
| ENSG000001 | BCL2L15 | 5.66E-01 | ? | 8.02E-01 | -3.18E-01 | -1.25E+00 |
| ENSG000001 | MORC3 | 5.66E-01 | 8.30E-01 | 1.03E+00 | 3.71E-02 | 1.03E+00 |
| ENSG000001 | SMG5 | 5.66E-01 | 8.30E-01 | 1.02E+00 | 3.37E-02 | 1.02E+00 |
| ENSG000002 | EEF1A1P12 | 5.66E-01 | ? | 7.75E-01 | -3.67E-01 | -1.29E+00 |
| ENSG000001 | FAM172BP | 5.66E-01 | 8.30E-01 | 9.47E-01 | -7.85E-02 | -1.06E+00 |
| ENSG000001 | PPM1B | 5.66E-01 | 8.30E-01 | 1.03E+00 | 3.57E-02 | 1.03E+00 |
| ENSG000001 | CMAHP | 5.66E-01 | ? | 8.80E-01 | -1.84E-01 | -1.14E+00 |
| ENSG000001 | ENO3 | 5.66E-01 | 8.30E-01 | 1.09E+00 | 1.27E-01 | 1.09E+00 |
| ENSG000001 | SUV39H2 | 5.66E-01 | 8.30E-01 | 1.03E+00 | 4.10E-02 | 1.03E+00 |
| ENSG000001 | NTAN1 | 5.66E-01 | 8.30E-01 | 9.75E-01 | -3.61E-02 | -1.03E+00 |
| ENSG00000C | SMARCD1 | 5.66E-01 | 8.30E-01 | 9.86E-01 | -2.02E-02 | -1.01E+00 |
| ENSG000002 | --- | 5.66E-01 | ? | 7.68E-01 | -3.81E-01 | -1.30E+00 |
| ENSG000001 | VPS13A | 5.66E-01 | 8.30E-01 | 1.03E+00 | 4.69E-02 | 1.03E+00 |
| ENSG000001 | EEF1E1 | 5.66E-01 | 8.30E-01 | 9.73E-01 | -3.89E-02 | -1.03E+00 |
| ENSG00000C | IPO11 | 5.66E-01 | 8.30E-01 | 9.85E-01 | -2.14E-02 | -1.01E+00 |
| ENSG000001 | MSRB3 | 5.66E-01 | 8.30E-01 | 1.03E+00 | 3.82E-02 | 1.03E+00 |
| ENSG00000C | SDF4 | 5.67E-01 | 8.30E-01 | 1.02E+00 | 2.53E-02 | 1.02E+00 |
| ENSG000001 | CEP41 | 5.67E-01 | 8.30E-01 | 9.73E-01 | -3.88E-02 | -1.03E+00 |
| ENSG00000C | MAP3K13 | 5.67E-01 | 8.30E-01 | 9.80E-01 | -2.91E-02 | -1.02E+00 |
| ENSG00000C | OPN3 | 5.67E-01 | 8.30E-01 | 9.72E-01 | -4.08E-02 | -1.03E+00 |
| ENSG000002 | --- | 5.67E-01 | 8.30E-01 | 9.31E-01 | -1.03E-01 | -1.07E+00 |
| ENSG000001 | SNX22 | 5.67E-01 | 8.30E-01 | 1.05E+00 | 6.42E-02 | 1.05E+00 |
| ENSG000002 | KCTD7 | 5.67E-01 | 8.30E-01 | 1.04E+00 | 5.47E-02 | 1.04E+00 |
| ENSG000001 | MTOR | 5.67E-01 | 8.30E-01 | 9.80E-01 | -2.89E-02 | -1.02E+00 |
| ENSG000002 | LINC01132 | 5.67E-01 | ? | 8.61E-01 | -2.16E-01 | -1.16E+00 |
| ENSG000002 | KRT8P33 | 5.67E-01 | ? | 8.94E-01 | -1.61E-01 | -1.12E+00 |
| ENSG000001 | FAM193B | 5.67E-01 | 8.30E-01 | 1.03E+00 | 4.80E-02 | 1.03E+00 |
| ENSG000001 | FRG1 | 5.67E-01 | 8.30E-01 | 9.72E-01 | -4.13E-02 | -1.03E+00 |
| ENSG000002 | RPL39P3 | 5.67E-01 | 8.30E-01 | 1.07E+00 | 1.02E-01 | 1.07E+00 |
| ENSG000001 | RANGAP1 | 5.67E-01 | 8.30E-01 | 1.03E+00 | 3.85E-02 | 1.03E+00 |
| ENSG000001 | TP53I3 | 5.67E-01 | 8.31E-01 | 9.73E-01 | -3.88E-02 | -1.03E+00 |
| ENSG000002 | TSNAX-DISC1 | 5.68E-01 | ? | 7.79E-01 | -3.61E-01 | -1.28E+00 |
| ENSG000001 | KIF11 | 5.68E-01 | 8.31E-01 | 9.51E-01 | -7.25E-02 | -1.05E+00 |
| ENSG00000C | NTHL1 | 5.68E-01 | 8.31E-01 | 1.05E+00 | 7.09E-02 | 1.05E+00 |
| ENSG000002 | --- | 5.68E-01 | ? | 1.37E+00 | 4.57E-01 | 1.37E+00 |
| ENSG000002 | --- | 5.68E-01 | ? | 1.19E+00 | 2.45E-01 | 1.19E+00 |
| ENSG000002 | --- | 5.68E-01 | ? | 8.47E-01 | -2.40E-01 | -1.18E+00 |
| ENSG000001 | ALDH2 | 5.68E-01 | 8.31E-01 | 1.03E+00 | 4.33E-02 | 1.03E+00 |
| ENSG000001 | ZNF230 | 5.68E-01 | 8.31E-01 | 9.58E-01 | -6.18E-02 | -1.04E+00 |
| ENSG00000C | KDM2B | 5.68E-01 | 8.31E-01 | 1.02E+00 | 2.93E-02 | 1.02E+00 |
| ENSG000001 | DYNLT2 | 5.68E-01 | 8.31E-01 | 1.10E+00 | 1.35E-01 | 1.10E+00 |
| ENSG000002 | --- | 5.68E-01 | ? | 8.51E-01 | -2.32E-01 | -1.17E+00 |
| ENSG000002 | --- | 5.68E-01 | 8.31E-01 | 1.04E+00 | 6.12E-02 | 1.04E+00 |
| ENSG000002 | FTH1P7 | 5.69E-01 | 8.31E-01 | 1.06E+00 | 7.77E-02 | 1.06E+00 |
| ENSG000002 | WASH9P | 5.69E-01 | ? | 8.69E-01 | -2.02E-01 | -1.15E+00 |

| | | | | | | |
|---|---|---|---|---|---|---|
| ENSG000002 | SLX1B-SULT1 | 5.69E-01 | 8.32E-01 | 9.45E-01 | -8.09E-02 | -1.06E+00 |
| ENSG000001 | NIBAN3 | 5.69E-01 | ? | 1.12E+00 | 1.65E-01 | 1.12E+00 |
| ENSG000001 | MAPK1 | 5.69E-01 | 8.32E-01 | 1.01E+00 | 1.70E-02 | 1.01E+00 |
| ENSG00000C | SAMD4A | 5.69E-01 | 8.32E-01 | 1.02E+00 | 2.42E-02 | 1.02E+00 |
| ENSG000001 | UBA1 | 5.69E-01 | 8.32E-01 | 1.02E+00 | 2.74E-02 | 1.02E+00 |
| ENSG000002 | --- | 5.69E-01 | 8.32E-01 | 1.06E+00 | 8.11E-02 | 1.06E+00 |
| ENSG000001 | ZFHX2 | 5.69E-01 | 8.32E-01 | 9.50E-01 | -7.42E-02 | -1.05E+00 |
| ENSG000002 | HLA-C | 5.69E-01 | 8.32E-01 | 1.04E+00 | 5.13E-02 | 1.04E+00 |
| ENSG000001 | MAGI2 | 5.69E-01 | 8.32E-01 | 9.16E-01 | -1.27E-01 | -1.09E+00 |
| ENSG000001 | SLFNL1 | 5.69E-01 | 8.32E-01 | 9.18E-01 | -1.23E-01 | -1.09E+00 |
| ENSG000002 | FNIP1 | 5.69E-01 | 8.32E-01 | 9.61E-01 | -5.80E-02 | -1.04E+00 |
| ENSG000002 | TBC1D8-AS1 | 5.69E-01 | ? | 1.17E+00 | 2.31E-01 | 1.17E+00 |
| ENSG000002 | LINC02965 | 5.69E-01 | ? | 1.22E+00 | 2.84E-01 | 1.22E+00 |
| ENSG000001 | SSBP3 | 5.69E-01 | 8.32E-01 | 1.02E+00 | 2.67E-02 | 1.02E+00 |
| ENSG000001 | ZBTB33 | 5.69E-01 | 8.32E-01 | 9.78E-01 | -3.21E-02 | -1.02E+00 |
| ENSG000002 | FBXO17 | 5.69E-01 | 8.32E-01 | 1.05E+00 | 6.50E-02 | 1.05E+00 |
| ENSG000001 | KLHDC7A | 5.69E-01 | ? | 1.24E+00 | 3.07E-01 | 1.24E+00 |
| ENSG000002 | PCDHA10 | 5.70E-01 | 8.32E-01 | 9.27E-01 | -1.09E-01 | -1.08E+00 |
| ENSG000002 | DECR2 | 5.70E-01 | 8.32E-01 | 1.05E+00 | 7.63E-02 | 1.05E+00 |
| ENSG000002 | --- | 5.70E-01 | 8.32E-01 | 9.37E-01 | -9.32E-02 | -1.07E+00 |
| ENSG000002 | LINC02595 | 5.70E-01 | ? | 9.08E-01 | -1.39E-01 | -1.10E+00 |
| ENSG000002 | --- | 5.70E-01 | 8.32E-01 | 1.04E+00 | 5.62E-02 | 1.04E+00 |
| ENSG000002 | FAM157D | 5.70E-01 | ? | 8.03E-01 | -3.17E-01 | -1.25E+00 |
| ENSG000002 | --- | 5.70E-01 | ? | 1.13E+00 | 1.82E-01 | 1.13E+00 |
| ENSG000002 | LINC00938 | 5.70E-01 | 8.32E-01 | 9.58E-01 | -6.13E-02 | -1.04E+00 |
| ENSG000001 | TMCC1 | 5.70E-01 | 8.32E-01 | 9.83E-01 | -2.49E-02 | -1.02E+00 |
| ENSG000001 | SPTBN5 | 5.70E-01 | 8.32E-01 | 9.39E-01 | -9.03E-02 | -1.06E+00 |
| ENSG000001 | NR6A1 | 5.70E-01 | 8.32E-01 | 1.05E+00 | 7.36E-02 | 1.05E+00 |
| ENSG000001 | MRM3 | 5.70E-01 | 8.32E-01 | 1.03E+00 | 4.73E-02 | 1.03E+00 |
| ENSG000001 | GALM | 5.70E-01 | 8.32E-01 | 9.41E-01 | -8.80E-02 | -1.06E+00 |
| ENSG000002 | --- | 5.70E-01 | 8.32E-01 | 8.70E-01 | -2.00E-01 | -1.15E+00 |
| ENSG000001 | RNF26 | 5.70E-01 | 8.32E-01 | 9.71E-01 | -4.20E-02 | -1.03E+00 |
| ENSG000002 | --- | 5.70E-01 | ? | 8.04E-01 | -3.14E-01 | -1.24E+00 |
| ENSG000001 | CENPP | 5.70E-01 | 8.32E-01 | 9.57E-01 | -6.28E-02 | -1.04E+00 |
| ENSG000002 | LINC01204 | 5.70E-01 | 8.32E-01 | 9.47E-01 | -7.81E-02 | -1.06E+00 |
| ENSG000001 | MACF1 | 5.70E-01 | 8.32E-01 | 9.59E-01 | -6.03E-02 | -1.04E+00 |
| ENSG000001 | F2RL2 | 5.71E-01 | ? | 1.18E+00 | 2.38E-01 | 1.18E+00 |
| ENSG000002 | LINC02889 | 5.71E-01 | 8.32E-01 | 1.11E+00 | 1.44E-01 | 1.11E+00 |
| ENSG000002 | COPS8-DT | 5.71E-01 | 8.32E-01 | 9.30E-01 | -1.04E-01 | -1.07E+00 |
| ENSG000002 | PFN1P8 | 5.71E-01 | ? | 1.20E+00 | 2.63E-01 | 1.20E+00 |
| ENSG000002 | --- | 5.71E-01 | ? | 1.26E+00 | 3.30E-01 | 1.26E+00 |
| ENSG000002 | CRIP1 | 5.71E-01 | ? | 8.96E-01 | -1.59E-01 | -1.12E+00 |
| ENSG000001 | MRPS5 | 5.71E-01 | 8.33E-01 | 9.81E-01 | -2.82E-02 | -1.02E+00 |
| ENSG000001 | DPM3 | 5.71E-01 | 8.33E-01 | 9.52E-01 | -7.17E-02 | -1.05E+00 |
| ENSG00000C | SPDL1 | 5.71E-01 | 8.33E-01 | 9.68E-01 | -4.72E-02 | -1.03E+00 |

| | | | | | | |
|---|---|---|---|---|---|---|
| ENSG000001 | TIAL1 | 5.71E-01 | 8.33E-01 | 9.84E-01 | -2.34E-02 | -1.02E+00 |
| ENSG000002 | --- | 5.72E-01 | ? | 1.45E+00 | 5.35E-01 | 1.45E+00 |
| ENSG000002 | --- | 5.72E-01 | ? | 1.45E+00 | 5.35E-01 | 1.45E+00 |
| ENSG000001 | TTC8 | 5.72E-01 | 8.33E-01 | 1.02E+00 | 3.43E-02 | 1.02E+00 |
| ENSG000001 | ITPA | 5.72E-01 | 8.33E-01 | 1.03E+00 | 4.37E-02 | 1.03E+00 |
| ENSG000002 | LINC02256 | 5.72E-01 | ? | 1.24E+00 | 3.13E-01 | 1.24E+00 |
| ENSG000001 | ALKBH2 | 5.72E-01 | 8.33E-01 | 9.63E-01 | -5.41E-02 | -1.04E+00 |
| ENSG000001 | TPBG | 5.72E-01 | 8.33E-01 | 9.72E-01 | -4.14E-02 | -1.03E+00 |
| ENSG000001 | PDCL | 5.72E-01 | 8.33E-01 | 9.75E-01 | -3.61E-02 | -1.03E+00 |
| ENSG000001 | UBE2Q1 | 5.72E-01 | 8.33E-01 | 1.02E+00 | 3.18E-02 | 1.02E+00 |
| ENSG000002 | --- | 5.72E-01 | ? | 9.14E-01 | -1.30E-01 | -1.09E+00 |
| ENSG000002 | PRKG1-AS1 | 5.72E-01 | ? | 1.20E+00 | 2.61E-01 | 1.20E+00 |
| ENSG000001 | FUT6 | 5.72E-01 | ? | 1.26E+00 | 3.31E-01 | 1.26E+00 |
| ENSG000001 | PIN1 | 5.72E-01 | 8.33E-01 | 1.04E+00 | 6.07E-02 | 1.04E+00 |
| ENSG000001 | SLC3A2 | 5.72E-01 | 8.33E-01 | 1.04E+00 | 5.57E-02 | 1.04E+00 |
| ENSG000002 | MFSD2B | 5.72E-01 | ? | 7.69E-01 | -3.79E-01 | -1.30E+00 |
| ENSG00000C | UIMC1 | 5.72E-01 | 8.33E-01 | 1.02E+00 | 2.88E-02 | 1.02E+00 |
| ENSG000001 | MAPK7 | 5.72E-01 | 8.33E-01 | 1.03E+00 | 4.45E-02 | 1.03E+00 |
| ENSG000001 | PRPF31 | 5.72E-01 | 8.33E-01 | 1.02E+00 | 2.28E-02 | 1.02E+00 |
| ENSG000002 | CASC9 | 5.73E-01 | ? | 1.32E+00 | 4.03E-01 | 1.32E+00 |
| ENSG000002 | --- | 5.73E-01 | ? | 1.14E+00 | 1.94E-01 | 1.14E+00 |
| ENSG000001 | IRF2 | 5.73E-01 | 8.33E-01 | 1.03E+00 | 4.04E-02 | 1.03E+00 |
| ENSG000001 | KCTD4 | 5.73E-01 | 8.33E-01 | 1.11E+00 | 1.56E-01 | 1.11E+00 |
| ENSG000002 | --- | 5.73E-01 | ? | 8.51E-01 | -2.33E-01 | -1.18E+00 |
| ENSG000002 | ANP32BP1 | 5.73E-01 | ? | 7.41E-01 | -4.32E-01 | -1.35E+00 |
| ENSG000001 | DDX59 | 5.73E-01 | 8.34E-01 | 9.76E-01 | -3.57E-02 | -1.03E+00 |
| ENSG000002 | RNF208 | 5.73E-01 | ? | 8.36E-01 | -2.58E-01 | -1.20E+00 |
| ENSG000002 | CD177 | 5.73E-01 | ? | 9.09E-01 | -1.37E-01 | -1.10E+00 |
| ENSG000001 | MAPK1IP1L | 5.73E-01 | 8.34E-01 | 9.84E-01 | -2.29E-02 | -1.02E+00 |
| ENSG000002 | --- | 5.73E-01 | ? | 7.41E-01 | -4.33E-01 | -1.35E+00 |
| ENSG00000C | CWF19L1 | 5.73E-01 | 8.34E-01 | 1.02E+00 | 3.45E-02 | 1.02E+00 |
| ENSG000002 | --- | 5.73E-01 | ? | 1.19E+00 | 2.48E-01 | 1.19E+00 |
| ENSG000002 | --- | 5.73E-01 | ? | 1.25E+00 | 3.19E-01 | 1.25E+00 |
| ENSG000001 | RIPK3 | 5.73E-01 | 8.34E-01 | 9.68E-01 | -4.62E-02 | -1.03E+00 |
| ENSG000001 | TRPA1 | 5.73E-01 | 8.34E-01 | 9.30E-01 | -1.05E-01 | -1.08E+00 |
| ENSG000001 | RNF207 | 5.74E-01 | 8.34E-01 | 9.44E-01 | -8.33E-02 | -1.06E+00 |
| ENSG00000C | RUNX1T1 | 5.74E-01 | ? | 1.13E+00 | 1.77E-01 | 1.13E+00 |
| ENSG000002 | --- | 5.74E-01 | ? | 1.25E+00 | 3.19E-01 | 1.25E+00 |
| ENSG000001 | ZNF609 | 5.74E-01 | 8.34E-01 | 1.01E+00 | 1.71E-02 | 1.01E+00 |
| ENSG000001 | MDM1 | 5.74E-01 | 8.34E-01 | 9.63E-01 | -5.37E-02 | -1.04E+00 |
| ENSG000001 | RP9 | 5.74E-01 | 8.34E-01 | 1.03E+00 | 3.92E-02 | 1.03E+00 |
| ENSG000002 | --- | 5.74E-01 | ? | 8.21E-01 | -2.85E-01 | -1.22E+00 |
| ENSG000002 | --- | 5.74E-01 | ? | 1.12E+00 | 1.61E-01 | 1.12E+00 |
| ENSG000002 | ASIC3 | 5.74E-01 | 8.34E-01 | 9.22E-01 | -1.17E-01 | -1.08E+00 |
| ENSG000001 | FAM174B | 5.74E-01 | 8.35E-01 | 9.65E-01 | -5.20E-02 | -1.04E+00 |

| | | | | | | |
|---|---|---|---|---|---|---|
| ENSG000002 | --- | 5.74E-01 | ? | 8.10E-01 | -3.04E-01 | -1.23E+00 |
| ENSG000002 | --- | 5.74E-01 | 8.35E-01 | 9.57E-01 | -6.36E-02 | -1.05E+00 |
| ENSG000002 | --- | 5.74E-01 | 8.35E-01 | 9.14E-01 | -1.31E-01 | -1.09E+00 |
| ENSG000002 | --- | 5.74E-01 | 8.35E-01 | 9.13E-01 | -1.31E-01 | -1.09E+00 |
| ENSG000002 | LINC01932 | 5.74E-01 | ? | 8.94E-01 | -1.61E-01 | -1.12E+00 |
| ENSG000001 | STK17A | 5.74E-01 | 8.35E-01 | 1.02E+00 | 3.28E-02 | 1.02E+00 |
| ENSG000002 | PIGBOS1 | 5.75E-01 | 8.35E-01 | 1.06E+00 | 8.42E-02 | 1.06E+00 |
| ENSG000001 | WNK2 | 5.75E-01 | ? | 1.23E+00 | 2.96E-01 | 1.23E+00 |
| ENSG000001 | CLYBL | 5.75E-01 | 8.35E-01 | 9.31E-01 | -1.03E-01 | -1.07E+00 |
| ENSG000001 | CTDNEP1 | 5.75E-01 | 8.35E-01 | 1.02E+00 | 3.06E-02 | 1.02E+00 |
| ENSG000002 | LINC01483 | 5.75E-01 | ? | 1.23E+00 | 3.03E-01 | 1.23E+00 |
| ENSG000001 | FARS2 | 5.75E-01 | 8.35E-01 | 1.04E+00 | 6.08E-02 | 1.04E+00 |
| ENSG000001 | METTL25 | 5.75E-01 | 8.35E-01 | 1.04E+00 | 5.13E-02 | 1.04E+00 |
| ENSG000001 | IRAK1 | 5.75E-01 | 8.35E-01 | 1.03E+00 | 4.17E-02 | 1.03E+00 |
| ENSG000002 | --- | 5.75E-01 | ? | 1.19E+00 | 2.47E-01 | 1.19E+00 |
| ENSG000001 | CABYR | 5.75E-01 | 8.35E-01 | 1.04E+00 | 6.05E-02 | 1.04E+00 |
| ENSG000002 | TSPAN5-DT | 5.75E-01 | 8.35E-01 | 9.10E-01 | -1.35E-01 | -1.10E+00 |
| ENSG000001 | RPRD2 | 5.75E-01 | 8.35E-01 | 9.81E-01 | -2.70E-02 | -1.02E+00 |
| ENSG000002 | --- | 5.75E-01 | ? | 8.61E-01 | -2.16E-01 | -1.16E+00 |
| ENSG00000C | PSMD5 | 5.75E-01 | 8.35E-01 | 1.01E+00 | 1.85E-02 | 1.01E+00 |
| ENSG000001 | GSAP | 5.75E-01 | 8.35E-01 | 9.53E-01 | -6.90E-02 | -1.05E+00 |
| ENSG000001 | DBR1 | 5.75E-01 | 8.35E-01 | 1.03E+00 | 3.80E-02 | 1.03E+00 |
| ENSG000001 | SEC24C | 5.75E-01 | 8.35E-01 | 1.02E+00 | 2.16E-02 | 1.02E+00 |
| ENSG00000C | ACOX3 | 5.75E-01 | 8.35E-01 | 9.61E-01 | -5.74E-02 | -1.04E+00 |
| ENSG000002 | MORF4L2-AS | 5.75E-01 | ? | 8.08E-01 | -3.07E-01 | -1.24E+00 |
| ENSG000002 | --- | 5.75E-01 | ? | 7.33E-01 | -4.49E-01 | -1.36E+00 |
| ENSG000001 | MYL2 | 5.75E-01 | ? | 1.31E+00 | 3.87E-01 | 1.31E+00 |
| ENSG000002 | --- | 5.76E-01 | ? | 8.48E-01 | -2.38E-01 | -1.18E+00 |
| ENSG00000C | SULT2B1 | 5.76E-01 | 8.35E-01 | 9.07E-01 | -1.40E-01 | -1.10E+00 |
| ENSG000002 | SLC44A4 | 5.76E-01 | ? | 8.37E-01 | -2.57E-01 | -1.20E+00 |
| ENSG000001 | CCP110 | 5.76E-01 | 8.35E-01 | 9.71E-01 | -4.32E-02 | -1.03E+00 |
| ENSG000002 | --- | 5.76E-01 | ? | 8.85E-01 | -1.76E-01 | -1.13E+00 |
| ENSG00000C | COL4A4 | 5.76E-01 | 8.35E-01 | 9.44E-01 | -8.35E-02 | -1.06E+00 |
| ENSG00000C | ARHGEF1 | 5.76E-01 | 8.35E-01 | 1.03E+00 | 4.57E-02 | 1.03E+00 |
| ENSG000002 | --- | 5.76E-01 | ? | 1.26E+00 | 3.34E-01 | 1.26E+00 |
| ENSG000001 | TMEM179B | 5.76E-01 | 8.35E-01 | 1.02E+00 | 3.24E-02 | 1.02E+00 |
| ENSG000002 | ATF6-DT | 5.76E-01 | ? | 1.15E+00 | 2.02E-01 | 1.15E+00 |
| ENSG00000C | OVGP1 | 5.76E-01 | 8.35E-01 | 1.09E+00 | 1.18E-01 | 1.09E+00 |
| ENSG00000C | FLT4 | 5.76E-01 | ? | 1.16E+00 | 2.16E-01 | 1.16E+00 |
| ENSG000002 | PIGHP1 | 5.76E-01 | ? | 7.56E-01 | -4.03E-01 | -1.32E+00 |
| ENSG000002 | LIF-AS2 | 5.76E-01 | ? | 1.24E+00 | 3.09E-01 | 1.24E+00 |
| ENSG000002 | PCDHA3 | 5.76E-01 | ? | 7.41E-01 | -4.32E-01 | -1.35E+00 |
| ENSG000001 | MERTK | 5.76E-01 | ? | 1.13E+00 | 1.77E-01 | 1.13E+00 |
| ENSG000002 | --- | 5.76E-01 | ? | 8.95E-01 | -1.60E-01 | -1.12E+00 |
| ENSG000001 | IMPDH1 | 5.76E-01 | 8.36E-01 | 1.03E+00 | 3.84E-02 | 1.03E+00 |

| | | | | | | |
|---|---|---|---|---|---|---|
| ENSG000002 | WASH2P | 5.76E-01 | 8.36E-01 | 1.03E+00 | 4.78E-02 | 1.03E+00 |
| ENSG000001 | TNFAIP8L3 | 5.77E-01 | ? | 1.12E+00 | 1.63E-01 | 1.12E+00 |
| ENSG000002 | --- | 5.77E-01 | ? | 1.42E+00 | 5.04E-01 | 1.42E+00 |
| ENSG00000C | ARID4A | 5.77E-01 | 8.36E-01 | 1.03E+00 | 4.67E-02 | 1.03E+00 |
| ENSG000001 | SSBP3-AS1 | 5.77E-01 | ? | 8.57E-01 | -2.22E-01 | -1.17E+00 |
| ENSG000001 | FADD | 5.77E-01 | 8.36E-01 | 1.02E+00 | 3.26E-02 | 1.02E+00 |
| ENSG000001 | NEB | 5.77E-01 | 8.36E-01 | 1.06E+00 | 8.57E-02 | 1.06E+00 |
| ENSG000001 | SPG11 | 5.77E-01 | 8.36E-01 | 1.02E+00 | 2.97E-02 | 1.02E+00 |
| ENSG00000C | RHBDD2 | 5.77E-01 | 8.36E-01 | 1.03E+00 | 4.62E-02 | 1.03E+00 |
| ENSG000001 | MTHFSD | 5.77E-01 | 8.36E-01 | 9.75E-01 | -3.69E-02 | -1.03E+00 |
| ENSG000002 | --- | 5.77E-01 | ? | 1.16E+00 | 2.16E-01 | 1.16E+00 |
| ENSG000001 | STBD1 | 5.77E-01 | 8.36E-01 | 1.02E+00 | 2.68E-02 | 1.02E+00 |
| ENSG000002 | FAM95B1 | 5.77E-01 | ? | 1.32E+00 | 4.03E-01 | 1.32E+00 |
| ENSG00000C | CSDE1 | 5.77E-01 | 8.36E-01 | 9.83E-01 | -2.43E-02 | -1.02E+00 |
| ENSG000002 | UBQLN4P1 | 5.77E-01 | ? | 1.15E+00 | 1.96E-01 | 1.15E+00 |
| ENSG000002 | --- | 5.77E-01 | ? | 1.13E+00 | 1.80E-01 | 1.13E+00 |
| ENSG000002 | FOXO3B | 5.77E-01 | 8.36E-01 | 1.05E+00 | 6.36E-02 | 1.05E+00 |
| ENSG000002 | --- | 5.77E-01 | ? | 8.78E-01 | -1.87E-01 | -1.14E+00 |
| ENSG000002 | LOXL1-AS1 | 5.77E-01 | 8.36E-01 | 1.05E+00 | 7.49E-02 | 1.05E+00 |
| ENSG000001 | CNOT6L | 5.77E-01 | 8.36E-01 | 1.02E+00 | 3.09E-02 | 1.02E+00 |
| ENSG000002 | --- | 5.77E-01 | 8.36E-01 | 1.04E+00 | 5.55E-02 | 1.04E+00 |
| ENSG000001 | RHCE | 5.77E-01 | ? | 7.20E-01 | -4.73E-01 | -1.39E+00 |
| ENSG000001 | MCM8 | 5.77E-01 | 8.36E-01 | 9.64E-01 | -5.35E-02 | -1.04E+00 |
| ENSG000002 | RPEP6 | 5.77E-01 | ? | 1.16E+00 | 2.14E-01 | 1.16E+00 |
| ENSG000001 | DCAF13 | 5.77E-01 | 8.36E-01 | 9.85E-01 | -2.21E-02 | -1.02E+00 |
| ENSG000002 | ASS1P2 | 5.78E-01 | ? | 7.32E-01 | -4.49E-01 | -1.37E+00 |
| ENSG000001 | NONO | 5.78E-01 | 8.36E-01 | 9.82E-01 | -2.69E-02 | -1.02E+00 |
| ENSG000002 | ARPC3P3 | 5.78E-01 | ? | 1.15E+00 | 2.01E-01 | 1.15E+00 |
| ENSG000002 | BVES-AS1 | 5.78E-01 | ? | 8.30E-01 | -2.68E-01 | -1.20E+00 |
| ENSG000002 | --- | 5.78E-01 | 8.36E-01 | 1.09E+00 | 1.29E-01 | 1.09E+00 |
| ENSG000002 | --- | 5.78E-01 | ? | 7.23E-01 | -4.68E-01 | -1.38E+00 |
| ENSG000001 | PDZK1IP1 | 5.78E-01 | 8.36E-01 | 1.06E+00 | 8.61E-02 | 1.06E+00 |
| ENSG000002 | EEF1A1P9 | 5.78E-01 | ? | 8.18E-01 | -2.90E-01 | -1.22E+00 |
| ENSG000002 | TPI1P1 | 5.78E-01 | 8.36E-01 | 9.34E-01 | -9.84E-02 | -1.07E+00 |
| ENSG000001 | ZNF620 | 5.78E-01 | 8.36E-01 | 9.17E-01 | -1.26E-01 | -1.09E+00 |
| ENSG000001 | TROAP | 5.78E-01 | 8.36E-01 | 9.19E-01 | -1.22E-01 | -1.09E+00 |
| ENSG000001 | ZC3H7A | 5.78E-01 | 8.36E-01 | 1.02E+00 | 2.67E-02 | 1.02E+00 |
| ENSG000001 | WDR91 | 5.78E-01 | 8.36E-01 | 1.03E+00 | 3.85E-02 | 1.03E+00 |
| ENSG000002 | XKR7 | 5.78E-01 | ? | 6.97E-01 | -5.21E-01 | -1.43E+00 |
| ENSG000001 | ST6GAL2 | 5.78E-01 | ? | 8.78E-01 | -1.87E-01 | -1.14E+00 |
| ENSG000001 | RNF19B | 5.78E-01 | 8.36E-01 | 1.02E+00 | 2.66E-02 | 1.02E+00 |
| ENSG000001 | UTP23 | 5.78E-01 | 8.36E-01 | 9.80E-01 | -2.99E-02 | -1.02E+00 |
| ENSG000001 | ZNF296 | 5.78E-01 | 8.36E-01 | 1.07E+00 | 9.66E-02 | 1.07E+00 |
| ENSG000001 | EIF2D | 5.78E-01 | 8.36E-01 | 1.02E+00 | 2.50E-02 | 1.02E+00 |
| ENSG000002 | PPIL3 | 5.78E-01 | 8.36E-01 | 9.76E-01 | -3.56E-02 | -1.03E+00 |

| | | | | | | |
|---|---|---|---|---|---|---|
| ENSG000001 | ZCCHC24 | 5.78E-01 | 8.36E-01 | 1.04E+00 | 6.12E-02 | 1.04E+00 |
| ENSG000001 | TUBGCP4 | 5.78E-01 | 8.36E-01 | 9.83E-01 | -2.40E-02 | -1.02E+00 |
| ENSG000001 | TRIM65 | 5.78E-01 | 8.36E-01 | 1.03E+00 | 4.52E-02 | 1.03E+00 |
| ENSG000001 | JPH3 | 5.79E-01 | ? | 1.19E+00 | 2.54E-01 | 1.19E+00 |
| ENSG000002 | --- | 5.79E-01 | ? | 1.23E+00 | 3.01E-01 | 1.23E+00 |
| ENSG00000C | ATRX | 5.79E-01 | 8.36E-01 | 9.70E-01 | -4.35E-02 | -1.03E+00 |
| ENSG000002 | ZNF253 | 5.79E-01 | 8.36E-01 | 9.68E-01 | -4.65E-02 | -1.03E+00 |
| ENSG00000C | NOP58 | 5.79E-01 | 8.36E-01 | 9.76E-01 | -3.51E-02 | -1.02E+00 |
| ENSG000001 | SCAF1 | 5.79E-01 | 8.36E-01 | 1.03E+00 | 4.94E-02 | 1.03E+00 |
| ENSG000001 | GRIA1 | 5.79E-01 | ? | 1.19E+00 | 2.50E-01 | 1.19E+00 |
| ENSG000001 | SPRR2D | 5.79E-01 | 8.36E-01 | 1.11E+00 | 1.49E-01 | 1.11E+00 |
| ENSG000001 | HDAC2 | 5.79E-01 | 8.36E-01 | 9.82E-01 | -2.68E-02 | -1.02E+00 |
| ENSG000002 | RRN3P3 | 5.79E-01 | 8.36E-01 | 9.54E-01 | -6.77E-02 | -1.05E+00 |
| ENSG000002 | PCDHGA11 | 5.79E-01 | 8.36E-01 | 1.04E+00 | 5.16E-02 | 1.04E+00 |
| ENSG000001 | KCNC4 | 5.79E-01 | 8.36E-01 | 1.03E+00 | 4.25E-02 | 1.03E+00 |
| ENSG00000C | ZC3HC1 | 5.79E-01 | 8.36E-01 | 1.03E+00 | 3.84E-02 | 1.03E+00 |
| ENSG00000C | TRNT1 | 5.79E-01 | 8.36E-01 | 9.76E-01 | -3.45E-02 | -1.02E+00 |
| ENSG00000C | ZNF184 | 5.79E-01 | 8.36E-01 | 9.60E-01 | -5.90E-02 | -1.04E+00 |
| ENSG000002 | BCL10-AS1 | 5.79E-01 | ? | 1.29E+00 | 3.70E-01 | 1.29E+00 |
| ENSG00000C | SCD | 5.79E-01 | 8.36E-01 | 9.75E-01 | -3.71E-02 | -1.03E+00 |
| ENSG000001 | ADCY7 | 5.80E-01 | 8.36E-01 | 1.04E+00 | 5.22E-02 | 1.04E+00 |
| ENSG000001 | ERICH1 | 5.80E-01 | 8.36E-01 | 9.74E-01 | -3.82E-02 | -1.03E+00 |
| ENSG000001 | CCDC159 | 5.80E-01 | 8.36E-01 | 9.36E-01 | -9.57E-02 | -1.07E+00 |
| ENSG000002 | --- | 5.80E-01 | ? | 7.51E-01 | -4.14E-01 | -1.33E+00 |
| ENSG000001 | NANOS3 | 5.80E-01 | ? | 7.72E-01 | -3.73E-01 | -1.29E+00 |
| ENSG00000C | CYB561 | 5.80E-01 | 8.37E-01 | 9.69E-01 | -4.49E-02 | -1.03E+00 |
| ENSG000001 | SERPINA1 | 5.80E-01 | 8.37E-01 | 1.03E+00 | 4.52E-02 | 1.03E+00 |
| ENSG000002 | FRG1JP | 5.80E-01 | ? | 1.22E+00 | 2.90E-01 | 1.22E+00 |
| ENSG000002 | --- | 5.80E-01 | ? | 8.44E-01 | -2.45E-01 | -1.18E+00 |
| ENSG000002 | --- | 5.80E-01 | 8.37E-01 | 8.99E-01 | -1.54E-01 | -1.11E+00 |
| ENSG000001 | SLFN13 | 5.80E-01 | 8.37E-01 | 9.10E-01 | -1.36E-01 | -1.10E+00 |
| ENSG000001 | AMBRA1 | 5.80E-01 | 8.37E-01 | 1.02E+00 | 3.21E-02 | 1.02E+00 |
| ENSG000002 | --- | 5.80E-01 | ? | 7.50E-01 | -4.16E-01 | -1.33E+00 |
| ENSG000002 | NR2E3 | 5.80E-01 | ? | 7.91E-01 | -3.38E-01 | -1.26E+00 |
| ENSG000001 | AGO3 | 5.80E-01 | 8.37E-01 | 1.04E+00 | 5.29E-02 | 1.04E+00 |
| ENSG000002 | PRRT3-AS1 | 5.80E-01 | ? | 1.12E+00 | 1.66E-01 | 1.12E+00 |
| ENSG000001 | TTC21B | 5.81E-01 | 8.37E-01 | 9.75E-01 | -3.64E-02 | -1.03E+00 |
| ENSG00000C | HSD17B6 | 5.81E-01 | ? | 1.33E+00 | 4.06E-01 | 1.33E+00 |
| ENSG000001 | TNNT1 | 5.81E-01 | 8.37E-01 | 9.63E-01 | -5.51E-02 | -1.04E+00 |
| ENSG000002 | --- | 5.81E-01 | ? | 7.43E-01 | -4.28E-01 | -1.35E+00 |
| ENSG000001 | MPPE1 | 5.81E-01 | 8.37E-01 | 1.04E+00 | 5.33E-02 | 1.04E+00 |
| ENSG000002 | ANKRD20A8I | 5.81E-01 | ? | 8.09E-01 | -3.06E-01 | -1.24E+00 |
| ENSG000002 | ZNF134 | 5.81E-01 | 8.37E-01 | 1.02E+00 | 2.33E-02 | 1.02E+00 |
| ENSG000002 | --- | 5.81E-01 | ? | 8.77E-01 | -1.89E-01 | -1.14E+00 |
| ENSG000001 | BTF3 | 5.81E-01 | 8.37E-01 | 1.02E+00 | 2.62E-02 | 1.02E+00 |

| | | | | | | |
|---|---|---|---|---|---|---|
| ENSG000001 | ITGB6 | 5.81E-01 | 8.37E-01 | 1.03E+00 | 4.69E-02 | 1.03E+00 |
| ENSG000002 | CA3-AS1 | 5.81E-01 | 8.37E-01 | 9.10E-01 | -1.36E-01 | -1.10E+00 |
| ENSG000001 | RAD23B | 5.81E-01 | 8.37E-01 | 1.01E+00 | 1.51E-02 | 1.01E+00 |
| ENSG000001 | GLRA4 | 5.81E-01 | ? | 1.11E+00 | 1.45E-01 | 1.11E+00 |
| ENSG00000C | RBBP9 | 5.81E-01 | 8.38E-01 | 1.03E+00 | 3.93E-02 | 1.03E+00 |
| ENSG000001 | PCSK1 | 5.81E-01 | ? | 1.20E+00 | 2.67E-01 | 1.20E+00 |
| ENSG00000C | ENPP4 | 5.81E-01 | 8.38E-01 | 9.34E-01 | -9.83E-02 | -1.07E+00 |
| ENSG000001 | RPAP2 | 5.81E-01 | 8.38E-01 | 9.69E-01 | -4.58E-02 | -1.03E+00 |
| ENSG000001 | OSGEPL1 | 5.82E-01 | 8.38E-01 | 9.59E-01 | -5.98E-02 | -1.04E+00 |
| ENSG000001 | TTPAL | 5.82E-01 | 8.38E-01 | 1.02E+00 | 2.52E-02 | 1.02E+00 |
| ENSG000001 | DMTN | 5.82E-01 | 8.38E-01 | 9.70E-01 | -4.33E-02 | -1.03E+00 |
| ENSG000001 | ACTR3 | 5.82E-01 | 8.38E-01 | 1.01E+00 | 1.82E-02 | 1.01E+00 |
| ENSG000001 | CALML4 | 5.82E-01 | 8.38E-01 | 9.34E-01 | -9.91E-02 | -1.07E+00 |
| ENSG000002 | FKBP14-AS1 | 5.82E-01 | ? | 1.14E+00 | 1.93E-01 | 1.14E+00 |
| ENSG000001 | GJA4 | 5.82E-01 | ? | 1.29E+00 | 3.65E-01 | 1.29E+00 |
| ENSG000001 | ZNF398 | 5.82E-01 | 8.38E-01 | 1.02E+00 | 3.16E-02 | 1.02E+00 |
| ENSG000002 | DSG2-AS1 | 5.82E-01 | ? | 1.11E+00 | 1.45E-01 | 1.11E+00 |
| ENSG000001 | DHRS7B | 5.82E-01 | 8.38E-01 | 9.67E-01 | -4.78E-02 | -1.03E+00 |
| ENSG000001 | PRCC | 5.82E-01 | 8.38E-01 | 1.02E+00 | 2.16E-02 | 1.02E+00 |
| ENSG000001 | CNTRL | 5.82E-01 | 8.38E-01 | 9.69E-01 | -4.54E-02 | -1.03E+00 |
| ENSG00000C | MCUB | 5.82E-01 | 8.38E-01 | 1.03E+00 | 3.93E-02 | 1.03E+00 |
| ENSG000002 | ST13P7 | 5.82E-01 | ? | 8.63E-01 | -2.13E-01 | -1.16E+00 |
| ENSG00000C | ERLEC1 | 5.82E-01 | 8.38E-01 | 9.78E-01 | -3.27E-02 | -1.02E+00 |
| ENSG00000C | RTRAF | 5.83E-01 | 8.38E-01 | 9.85E-01 | -2.20E-02 | -1.02E+00 |
| ENSG000002 | LCMT1 | 5.83E-01 | 8.38E-01 | 9.77E-01 | -3.36E-02 | -1.02E+00 |
| ENSG000001 | DGKZP1 | 5.83E-01 | 8.38E-01 | 1.07E+00 | 9.75E-02 | 1.07E+00 |
| ENSG000001 | NUP205 | 5.83E-01 | 8.38E-01 | 9.83E-01 | -2.54E-02 | -1.02E+00 |
| ENSG000001 | MST1R | 5.83E-01 | 8.38E-01 | 1.03E+00 | 4.79E-02 | 1.03E+00 |
| ENSG000002 | --- | 5.83E-01 | ? | 8.28E-01 | -2.72E-01 | -1.21E+00 |
| ENSG000002 | --- | 5.83E-01 | ? | 7.73E-01 | -3.71E-01 | -1.29E+00 |
| ENSG000002 | --- | 5.83E-01 | 8.38E-01 | 9.29E-01 | -1.06E-01 | -1.08E+00 |
| ENSG000002 | --- | 5.83E-01 | ? | 8.92E-01 | -1.64E-01 | -1.12E+00 |
| ENSG000001 | MSN | 5.83E-01 | 8.38E-01 | 9.82E-01 | -2.67E-02 | -1.02E+00 |
| ENSG000001 | KRTAP19-1 | 5.83E-01 | ? | 1.27E+00 | 3.49E-01 | 1.27E+00 |
| ENSG000001 | SMYD3 | 5.83E-01 | 8.38E-01 | 1.03E+00 | 4.26E-02 | 1.03E+00 |
| ENSG000001 | LRRC27 | 5.83E-01 | 8.39E-01 | 1.04E+00 | 5.71E-02 | 1.04E+00 |
| ENSG000001 | ZSWIM1 | 5.83E-01 | 8.39E-01 | 9.69E-01 | -4.61E-02 | -1.03E+00 |
| ENSG000001 | PCDHB15 | 5.83E-01 | ? | 8.52E-01 | -2.32E-01 | -1.17E+00 |
| ENSG000002 | --- | 5.83E-01 | ? | 1.20E+00 | 2.62E-01 | 1.20E+00 |
| ENSG000001 | TXNDC2 | 5.83E-01 | ? | 1.29E+00 | 3.68E-01 | 1.29E+00 |
| ENSG000001 | ZNF653 | 5.83E-01 | 8.39E-01 | 9.58E-01 | -6.22E-02 | -1.04E+00 |
| ENSG00000C | QPCTL | 5.83E-01 | 8.39E-01 | 1.05E+00 | 7.18E-02 | 1.05E+00 |
| ENSG000001 | PIF1 | 5.84E-01 | 8.39E-01 | 8.97E-01 | -1.56E-01 | -1.11E+00 |
| ENSG000001 | RPE | 5.84E-01 | 8.39E-01 | 1.01E+00 | 2.01E-02 | 1.01E+00 |
| ENSG000001 | C20orf96 | 5.84E-01 | 8.39E-01 | 9.36E-01 | -9.51E-02 | -1.07E+00 |

| | | | | | | |
|---|---|---|---|---|---|---|
| ENSG000002 | RPL24P2 | 5.84E-01 | ? | 1.30E+00 | 3.83E-01 | 1.30E+00 |
| ENSG000002 | VAC14-AS1 | 5.84E-01 | ? | 1.14E+00 | 1.93E-01 | 1.14E+00 |
| ENSG000001 | TTBK2 | 5.84E-01 | 8.39E-01 | 1.03E+00 | 4.64E-02 | 1.03E+00 |
| ENSG000002 | --- | 5.84E-01 | ? | 8.92E-01 | -1.65E-01 | -1.12E+00 |
| ENSG000002 | --- | 5.84E-01 | ? | 8.39E-01 | -2.53E-01 | -1.19E+00 |
| ENSG000002 | BAG6 | 5.84E-01 | 8.39E-01 | 9.83E-01 | -2.42E-02 | -1.02E+00 |
| ENSG000002 | --- | 5.84E-01 | ? | 8.13E-01 | -2.98E-01 | -1.23E+00 |
| ENSG000001 | EFEMP1 | 5.84E-01 | 8.39E-01 | 1.05E+00 | 6.68E-02 | 1.05E+00 |
| ENSG000001 | LGALS3BP | 5.84E-01 | 8.39E-01 | 1.03E+00 | 3.90E-02 | 1.03E+00 |
| ENSG000002 | SLC12A2-DT | 5.84E-01 | 8.39E-01 | 1.03E+00 | 3.79E-02 | 1.03E+00 |
| ENSG000002 | RTL8B | 5.84E-01 | 8.39E-01 | 1.03E+00 | 4.94E-02 | 1.03E+00 |
| ENSG000001 | BRINP2 | 5.85E-01 | ? | 1.16E+00 | 2.09E-01 | 1.16E+00 |
| ENSG000001 | BRD7P2 | 5.85E-01 | ? | 1.13E+00 | 1.77E-01 | 1.13E+00 |
| ENSG000001 | ITGA7 | 5.85E-01 | ? | 1.14E+00 | 1.94E-01 | 1.14E+00 |
| ENSG000001 | SHISA2 | 5.85E-01 | ? | 1.14E+00 | 1.88E-01 | 1.14E+00 |
| ENSG000001 | MCM3AP | 5.85E-01 | 8.40E-01 | 9.85E-01 | -2.22E-02 | -1.02E+00 |
| ENSG000002 | RRN3P3 | 5.85E-01 | ? | 7.49E-01 | -4.18E-01 | -1.34E+00 |
| ENSG000002 | RPL23AP2 | 5.85E-01 | ? | 1.26E+00 | 3.36E-01 | 1.26E+00 |
| ENSG000001 | GBF1 | 5.85E-01 | 8.40E-01 | 9.89E-01 | -1.65E-02 | -1.01E+00 |
| ENSG000001 | TAF7 | 5.85E-01 | 8.40E-01 | 1.01E+00 | 2.07E-02 | 1.01E+00 |
| ENSG000002 | KANTR | 5.86E-01 | 8.40E-01 | 1.06E+00 | 8.51E-02 | 1.06E+00 |
| ENSG000002 | LINC01515 | 5.86E-01 | 8.40E-01 | 9.21E-01 | -1.19E-01 | -1.09E+00 |
| ENSG000002 | TUBB2BP1 | 5.86E-01 | ? | 7.51E-01 | -4.14E-01 | -1.33E+00 |
| ENSG000001 | CCNO | 5.86E-01 | 8.40E-01 | 9.15E-01 | -1.28E-01 | -1.09E+00 |
| ENSG000001 | LEKR1 | 5.86E-01 | ? | 1.24E+00 | 3.12E-01 | 1.24E+00 |
| ENSG000001 | MRPL16 | 5.86E-01 | 8.40E-01 | 1.03E+00 | 4.48E-02 | 1.03E+00 |
| ENSG000001 | KCTD6 | 5.86E-01 | 8.40E-01 | 9.61E-01 | -5.73E-02 | -1.04E+00 |
| ENSG000001 | MRPL49 | 5.86E-01 | 8.40E-01 | 9.79E-01 | -3.00E-02 | -1.02E+00 |
| ENSG00000C | USE1 | 5.86E-01 | 8.40E-01 | 9.64E-01 | -5.35E-02 | -1.04E+00 |
| ENSG000001 | SOX15 | 5.86E-01 | 8.40E-01 | 9.73E-01 | -3.89E-02 | -1.03E+00 |
| ENSG000001 | COL5A1 | 5.86E-01 | 8.40E-01 | 1.04E+00 | 5.09E-02 | 1.04E+00 |
| ENSG00000C | VDAC3 | 5.86E-01 | 8.40E-01 | 9.83E-01 | -2.47E-02 | -1.02E+00 |
| ENSG000001 | NTRK1 | 5.86E-01 | ? | 1.36E+00 | 4.49E-01 | 1.36E+00 |
| ENSG00000C | ATOSA | 5.86E-01 | 8.40E-01 | 1.03E+00 | 3.79E-02 | 1.03E+00 |
| ENSG000001 | LRRC37A3 | 5.86E-01 | 8.40E-01 | 1.04E+00 | 5.56E-02 | 1.04E+00 |
| ENSG000002 | --- | 5.86E-01 | ? | 8.44E-01 | -2.44E-01 | -1.18E+00 |
| ENSG000001 | PPL | 5.86E-01 | 8.40E-01 | 9.36E-01 | -9.57E-02 | -1.07E+00 |
| ENSG000001 | SNAPC3 | 5.86E-01 | 8.40E-01 | 9.85E-01 | -2.16E-02 | -1.02E+00 |
| ENSG000001 | CSF1R | 5.86E-01 | ? | 8.84E-01 | -1.78E-01 | -1.13E+00 |
| ENSG000002 | ALOX12-AS1 | 5.86E-01 | 8.40E-01 | 9.09E-01 | -1.38E-01 | -1.10E+00 |
| ENSG000002 | --- | 5.86E-01 | ? | 8.68E-01 | -2.05E-01 | -1.15E+00 |
| ENSG000001 | IL36B | 5.86E-01 | ? | 1.23E+00 | 2.99E-01 | 1.23E+00 |
| ENSG000001 | NMNAT3 | 5.86E-01 | ? | 1.10E+00 | 1.41E-01 | 1.10E+00 |
| ENSG000001 | YWHAH | 5.86E-01 | 8.40E-01 | 1.02E+00 | 2.16E-02 | 1.02E+00 |
| ENSG000002 | ADM5 | 5.86E-01 | ? | 8.34E-01 | -2.62E-01 | -1.20E+00 |

| | | | | | | |
|---|---|---|---|---|---|---|
| ENSG00000C | ITGAE | 5.87E-01 | 8.40E-01 | 9.58E-01 | -6.23E-02 | -1.04E+00 |
| ENSG000002 | --- | 5.87E-01 | 8.40E-01 | 9.25E-01 | -1.13E-01 | -1.08E+00 |
| ENSG000002 | --- | 5.87E-01 | ? | 1.12E+00 | 1.69E-01 | 1.12E+00 |
| ENSG000001 | IFI16 | 5.87E-01 | 8.40E-01 | 9.78E-01 | -3.24E-02 | -1.02E+00 |
| ENSG000001 | INTS1 | 5.87E-01 | 8.40E-01 | 1.03E+00 | 4.31E-02 | 1.03E+00 |
| ENSG000002 | --- | 5.87E-01 | ? | 8.73E-01 | -1.96E-01 | -1.15E+00 |
| ENSG000002 | RNF139-DT | 5.87E-01 | 8.40E-01 | 9.23E-01 | -1.15E-01 | -1.08E+00 |
| ENSG000001 | DGKH | 5.87E-01 | 8.40E-01 | 9.63E-01 | -5.44E-02 | -1.04E+00 |
| ENSG00000C | AP3M2 | 5.87E-01 | 8.40E-01 | 1.02E+00 | 2.84E-02 | 1.02E+00 |
| ENSG000002 | --- | 5.87E-01 | 8.40E-01 | 9.36E-01 | -9.48E-02 | -1.07E+00 |
| ENSG000002 | GJD3-AS1 | 5.87E-01 | 8.40E-01 | 9.49E-01 | -7.48E-02 | -1.05E+00 |
| ENSG000002 | --- | 5.87E-01 | ? | 9.00E-01 | -1.52E-01 | -1.11E+00 |
| ENSG00000C | RAB11FIP3 | 5.87E-01 | 8.40E-01 | 1.03E+00 | 4.11E-02 | 1.03E+00 |
| ENSG000001 | DMRT1 | 5.87E-01 | ? | 8.15E-01 | -2.95E-01 | -1.23E+00 |
| ENSG000001 | GLOD4 | 5.87E-01 | 8.40E-01 | 9.84E-01 | -2.33E-02 | -1.02E+00 |
| ENSG00000C | WNK1 | 5.87E-01 | 8.40E-01 | 9.78E-01 | -3.20E-02 | -1.02E+00 |
| ENSG000002 | MUC12 | 5.87E-01 | ? | 7.61E-01 | -3.93E-01 | -1.31E+00 |
| ENSG000001 | ABHD11 | 5.87E-01 | 8.40E-01 | 9.67E-01 | -4.89E-02 | -1.03E+00 |
| ENSG000001 | SLC26A7 | 5.87E-01 | 8.40E-01 | 9.30E-01 | -1.04E-01 | -1.07E+00 |
| ENSG000002 | PCP4L1 | 5.87E-01 | ? | 1.15E+00 | 1.96E-01 | 1.15E+00 |
| ENSG000002 | RPAP3-DT | 5.87E-01 | ? | 8.98E-01 | -1.55E-01 | -1.11E+00 |
| ENSG000001 | POLD2 | 5.87E-01 | 8.41E-01 | 1.02E+00 | 2.59E-02 | 1.02E+00 |
| ENSG000002 | DCDC2C | 5.87E-01 | ? | 1.27E+00 | 3.41E-01 | 1.27E+00 |
| ENSG000002 | PCBP2P2 | 5.87E-01 | ? | 8.92E-01 | -1.65E-01 | -1.12E+00 |
| ENSG000001 | AGMAT | 5.87E-01 | 8.41E-01 | 1.06E+00 | 8.03E-02 | 1.06E+00 |
| ENSG000002 | TSC22D1-AS: | 5.88E-01 | ? | 8.72E-01 | -1.98E-01 | -1.15E+00 |
| ENSG000001 | RARS1 | 5.88E-01 | 8.41E-01 | 9.85E-01 | -2.19E-02 | -1.02E+00 |
| ENSG000001 | SH3TC2 | 5.88E-01 | 8.41E-01 | 9.73E-01 | -3.95E-02 | -1.03E+00 |
| ENSG00000C | ATXN3 | 5.88E-01 | 8.41E-01 | 1.02E+00 | 3.47E-02 | 1.02E+00 |
| ENSG000002 | --- | 5.88E-01 | ? | 1.17E+00 | 2.31E-01 | 1.17E+00 |
| ENSG000001 | C7orf50 | 5.88E-01 | 8.41E-01 | 1.04E+00 | 5.74E-02 | 1.04E+00 |
| ENSG000002 | --- | 5.88E-01 | ? | 8.60E-01 | -2.17E-01 | -1.16E+00 |
| ENSG000001 | PAN2 | 5.88E-01 | 8.41E-01 | 1.03E+00 | 4.71E-02 | 1.03E+00 |
| ENSG000001 | BCKDK | 5.88E-01 | 8.41E-01 | 1.04E+00 | 4.99E-02 | 1.04E+00 |
| ENSG000001 | TMEM67 | 5.88E-01 | 8.41E-01 | 9.67E-01 | -4.90E-02 | -1.03E+00 |
| ENSG000002 | LINC01119 | 5.88E-01 | ? | 8.13E-01 | -2.98E-01 | -1.23E+00 |
| ENSG000002 | MAP2K5-DT | 5.88E-01 | ? | 1.16E+00 | 2.09E-01 | 1.16E+00 |
| ENSG000001 | BAG1 | 5.88E-01 | 8.41E-01 | 9.68E-01 | -4.71E-02 | -1.03E+00 |
| ENSG000002 | HNRNPA1P1( | 5.88E-01 | ? | 8.80E-01 | -1.84E-01 | -1.14E+00 |
| ENSG000002 | GNG10 | 5.88E-01 | 8.41E-01 | 9.78E-01 | -3.23E-02 | -1.02E+00 |
| ENSG000001 | ACTR8 | 5.88E-01 | 8.41E-01 | 9.81E-01 | -2.78E-02 | -1.02E+00 |
| ENSG000001 | C9 | 5.88E-01 | ? | 1.21E+00 | 2.75E-01 | 1.21E+00 |
| ENSG000001 | ZNF543 | 5.88E-01 | 8.41E-01 | 1.04E+00 | 5.01E-02 | 1.04E+00 |
| ENSG000001 | RTKN2 | 5.89E-01 | 8.41E-01 | 1.08E+00 | 1.09E-01 | 1.08E+00 |
| ENSG000001 | NALF2 | 5.89E-01 | 8.41E-01 | 9.68E-01 | -4.71E-02 | -1.03E+00 |

| | | | | | | |
|---|---|---|---|---|---|---|
| ENSG000002 | SYT9-AS1 | 5.89E-01 | 8.41E-01 | 9.37E-01 | -9.41E-02 | -1.07E+00 |
| ENSG000001 | SALL2 | 5.89E-01 | 8.41E-01 | 1.09E+00 | 1.22E-01 | 1.09E+00 |
| ENSG000002 | --- | 5.89E-01 | ? | 8.55E-01 | -2.26E-01 | -1.17E+00 |
| ENSG000001 | ULBP3 | 5.89E-01 | 8.41E-01 | 1.02E+00 | 3.26E-02 | 1.02E+00 |
| ENSG000002 | TMEM92-AS | 5.89E-01 | ? | 8.06E-01 | -3.11E-01 | -1.24E+00 |
| ENSG000001 | SECISBP2L | 5.89E-01 | 8.41E-01 | 9.79E-01 | -3.07E-02 | -1.02E+00 |
| ENSG000002 | --- | 5.89E-01 | ? | 1.18E+00 | 2.38E-01 | 1.18E+00 |
| ENSG000001 | ABI2 | 5.89E-01 | 8.41E-01 | 9.75E-01 | -3.64E-02 | -1.03E+00 |
| ENSG000001 | POLR3D | 5.89E-01 | 8.41E-01 | 9.81E-01 | -2.77E-02 | -1.02E+00 |
| ENSG000001 | PAPSS2 | 5.89E-01 | 8.41E-01 | 9.67E-01 | -4.84E-02 | -1.03E+00 |
| ENSG000001 | UFSP2 | 5.89E-01 | 8.41E-01 | 1.02E+00 | 3.01E-02 | 1.02E+00 |
| ENSG000001 | CHAF1B | 5.89E-01 | 8.41E-01 | 9.49E-01 | -7.52E-02 | -1.05E+00 |
| ENSG000002 | --- | 5.89E-01 | ? | 8.82E-01 | -1.81E-01 | -1.13E+00 |
| ENSG00000C | CASS4 | 5.89E-01 | ? | 8.96E-01 | -1.59E-01 | -1.12E+00 |
| ENSG000002 | --- | 5.90E-01 | ? | 7.96E-01 | -3.30E-01 | -1.26E+00 |
| ENSG000002 | SLC35E1P1 | 5.90E-01 | ? | 1.22E+00 | 2.84E-01 | 1.22E+00 |
| ENSG000002 | NHSL1-AS1 | 5.90E-01 | 8.42E-01 | 1.06E+00 | 9.07E-02 | 1.06E+00 |
| ENSG000002 | --- | 5.90E-01 | ? | 1.14E+00 | 1.90E-01 | 1.14E+00 |
| ENSG000001 | MAGED1 | 5.90E-01 | 8.42E-01 | 9.75E-01 | -3.69E-02 | -1.03E+00 |
| ENSG000001 | EPHA4 | 5.90E-01 | 8.42E-01 | 1.03E+00 | 4.88E-02 | 1.03E+00 |
| ENSG000001 | CAMP | 5.90E-01 | ? | 1.14E+00 | 1.84E-01 | 1.14E+00 |
| ENSG000002 | --- | 5.90E-01 | ? | 8.08E-01 | -3.07E-01 | -1.24E+00 |
| ENSG000001 | AP1S2 | 5.90E-01 | 8.43E-01 | 9.72E-01 | -4.07E-02 | -1.03E+00 |
| ENSG000002 | SLX1A-SULT1 | 5.90E-01 | 8.43E-01 | 1.04E+00 | 6.03E-02 | 1.04E+00 |
| ENSG000001 | OTUD3 | 5.91E-01 | 8.43E-01 | 1.03E+00 | 4.31E-02 | 1.03E+00 |
| ENSG000002 | --- | 5.91E-01 | ? | 8.26E-01 | -2.76E-01 | -1.21E+00 |
| ENSG000002 | TTC23L-AS1 | 5.91E-01 | ? | 1.24E+00 | 3.12E-01 | 1.24E+00 |
| ENSG00000C | CROCC | 5.91E-01 | 8.43E-01 | 1.05E+00 | 6.98E-02 | 1.05E+00 |
| ENSG000002 | PRDM8-AS1 | 5.91E-01 | ? | 1.27E+00 | 3.44E-01 | 1.27E+00 |
| ENSG000001 | JUP | 5.91E-01 | 8.43E-01 | 1.06E+00 | 8.49E-02 | 1.06E+00 |
| ENSG000001 | ZNF592 | 5.91E-01 | 8.43E-01 | 9.82E-01 | -2.69E-02 | -1.02E+00 |
| ENSG000002 | ZBTB48 | 5.91E-01 | 8.43E-01 | 1.04E+00 | 5.07E-02 | 1.04E+00 |
| ENSG00000C | TFE3 | 5.91E-01 | 8.43E-01 | 1.02E+00 | 2.32E-02 | 1.02E+00 |
| ENSG000001 | LINC02908 | 5.91E-01 | ? | 8.34E-01 | -2.62E-01 | -1.20E+00 |
| ENSG000002 | ZNRD2-DT | 5.91E-01 | ? | 1.11E+00 | 1.55E-01 | 1.11E+00 |
| ENSG000002 | FTH1P11 | 5.91E-01 | ? | 1.10E+00 | 1.38E-01 | 1.10E+00 |
| ENSG000002 | PRR29 | 5.92E-01 | ? | 1.28E+00 | 3.54E-01 | 1.28E+00 |
| ENSG000001 | ZNF687 | 5.92E-01 | 8.44E-01 | 1.03E+00 | 3.74E-02 | 1.03E+00 |
| ENSG000001 | DDX21 | 5.92E-01 | 8.44E-01 | 1.02E+00 | 2.60E-02 | 1.02E+00 |
| ENSG000002 | MICAL3 | 5.92E-01 | 8.44E-01 | 1.02E+00 | 2.47E-02 | 1.02E+00 |
| ENSG000002 | --- | 5.92E-01 | ? | 8.42E-01 | -2.47E-01 | -1.19E+00 |
| ENSG000001 | LRCH2 | 5.92E-01 | ? | 1.18E+00 | 2.43E-01 | 1.18E+00 |
| ENSG000002 | --- | 5.92E-01 | ? | 1.26E+00 | 3.36E-01 | 1.26E+00 |
| ENSG000001 | HNRNPD | 5.92E-01 | 8.44E-01 | 9.79E-01 | -3.01E-02 | -1.02E+00 |
| ENSG000002 | --- | 5.92E-01 | 8.44E-01 | 9.00E-01 | -1.51E-01 | -1.11E+00 |

| | | | | | | |
|---|---|---|---|---|---|---|
| ENSG000001 | SLC25A28 | 5.92E-01 | 8.44E-01 | 1.03E+00 | 3.93E-02 | 1.03E+00 |
| ENSG000002 | CERS1 | 5.92E-01 | 8.44E-01 | 9.11E-01 | -1.34E-01 | -1.10E+00 |
| ENSG000001 | STAG3L3 | 5.92E-01 | ? | 1.73E+00 | 7.88E-01 | 1.73E+00 |
| ENSG000001 | TAF12 | 5.92E-01 | 8.44E-01 | 9.74E-01 | -3.83E-02 | -1.03E+00 |
| ENSG000002 | TFAP2A-AS1 | 5.92E-01 | ? | 9.21E-01 | -1.19E-01 | -1.09E+00 |
| ENSG000002 | RPL9P7 | 5.92E-01 | ? | 1.35E+00 | 4.36E-01 | 1.35E+00 |
| ENSG000002 | NT5C3AP1 | 5.92E-01 | ? | 7.80E-01 | -3.58E-01 | -1.28E+00 |
| ENSG000001 | SH3PXD2B | 5.92E-01 | 8.44E-01 | 9.82E-01 | -2.69E-02 | -1.02E+00 |
| ENSG000001 | EPM2A | 5.92E-01 | 8.44E-01 | 9.61E-01 | -5.78E-02 | -1.04E+00 |
| ENSG000001 | MED7 | 5.93E-01 | 8.44E-01 | 9.72E-01 | -4.14E-02 | -1.03E+00 |
| ENSG000002 | --- | 5.93E-01 | ? | 7.90E-01 | -3.40E-01 | -1.27E+00 |
| ENSG000001 | GMPPA | 5.93E-01 | 8.44E-01 | 9.74E-01 | -3.83E-02 | -1.03E+00 |
| ENSG000001 | CYP39A1 | 5.93E-01 | ? | 1.09E+00 | 1.31E-01 | 1.09E+00 |
| ENSG000001 | LGALSL | 5.93E-01 | 8.44E-01 | 1.04E+00 | 5.40E-02 | 1.04E+00 |
| ENSG000002 | --- | 5.93E-01 | ? | 8.25E-01 | -2.78E-01 | -1.21E+00 |
| ENSG00000( | GTF3C1 | 5.93E-01 | 8.44E-01 | 1.01E+00 | 1.54E-02 | 1.01E+00 |
| ENSG000002 | NUP62 | 5.93E-01 | 8.44E-01 | 1.01E+00 | 2.05E-02 | 1.01E+00 |
| ENSG000002 | ECE1-AS1 | 5.93E-01 | ? | 8.21E-01 | -2.85E-01 | -1.22E+00 |
| ENSG000002 | STEAP3-AS1 | 5.93E-01 | ? | 1.15E+00 | 2.05E-01 | 1.15E+00 |
| ENSG000002 | --- | 5.93E-01 | ? | 9.16E-01 | -1.27E-01 | -1.09E+00 |
| ENSG000001 | FPR1 | 5.93E-01 | ? | 1.26E+00 | 3.37E-01 | 1.26E+00 |
| ENSG00000( | NCBP3 | 5.93E-01 | 8.45E-01 | 9.85E-01 | -2.15E-02 | -1.02E+00 |
| ENSG000001 | DDX55 | 5.93E-01 | 8.45E-01 | 9.72E-01 | -4.15E-02 | -1.03E+00 |
| ENSG000002 | --- | 5.93E-01 | ? | 8.78E-01 | -1.88E-01 | -1.14E+00 |
| ENSG000002 | --- | 5.93E-01 | ? | 7.83E-01 | -3.52E-01 | -1.28E+00 |
| ENSG000001 | CFAP52 | 5.93E-01 | ? | 7.46E-01 | -4.23E-01 | -1.34E+00 |
| ENSG000001 | PTMS | 5.94E-01 | 8.45E-01 | 9.79E-01 | -3.12E-02 | -1.02E+00 |
| ENSG000002 | --- | 5.94E-01 | 8.45E-01 | 1.07E+00 | 9.19E-02 | 1.07E+00 |
| ENSG000001 | RSPRY1 | 5.94E-01 | 8.45E-01 | 9.85E-01 | -2.15E-02 | -1.02E+00 |
| ENSG000001 | DGKZ | 5.94E-01 | 8.45E-01 | 1.03E+00 | 3.69E-02 | 1.03E+00 |
| ENSG000001 | ZNF79 | 5.94E-01 | 8.45E-01 | 1.03E+00 | 4.45E-02 | 1.03E+00 |
| ENSG000001 | ADIPOR1 | 5.94E-01 | 8.45E-01 | 1.02E+00 | 2.52E-02 | 1.02E+00 |
| ENSG000002 | --- | 5.94E-01 | ? | 8.46E-01 | -2.40E-01 | -1.18E+00 |
| ENSG000001 | SLC25A12 | 5.94E-01 | 8.45E-01 | 1.02E+00 | 2.87E-02 | 1.02E+00 |
| ENSG000002 | INKA2-AS1 | 5.94E-01 | ? | 8.35E-01 | -2.60E-01 | -1.20E+00 |
| ENSG000001 | SAC3D1 | 5.94E-01 | 8.45E-01 | 1.04E+00 | 5.36E-02 | 1.04E+00 |
| ENSG000002 | RPL23AP53 | 5.94E-01 | 8.45E-01 | 1.06E+00 | 8.67E-02 | 1.06E+00 |
| ENSG000002 | AGGF1P2 | 5.94E-01 | ? | 1.31E+00 | 3.85E-01 | 1.31E+00 |
| ENSG000001 | FNDC4 | 5.94E-01 | 8.45E-01 | 1.04E+00 | 5.74E-02 | 1.04E+00 |
| ENSG000002 | --- | 5.94E-01 | ? | 8.79E-01 | -1.86E-01 | -1.14E+00 |
| ENSG000002 | LOH12CR2 | 5.94E-01 | ? | 1.23E+00 | 3.04E-01 | 1.23E+00 |
| ENSG000001 | TMEM19 | 5.94E-01 | 8.45E-01 | 1.02E+00 | 2.45E-02 | 1.02E+00 |
| ENSG000001 | ARFGEF3 | 5.94E-01 | 8.45E-01 | 9.69E-01 | -4.48E-02 | -1.03E+00 |
| ENSG000001 | XRCC4 | 5.94E-01 | 8.45E-01 | 9.69E-01 | -4.60E-02 | -1.03E+00 |
| ENSG00000( | ERC1 | 5.94E-01 | 8.45E-01 | 1.02E+00 | 3.01E-02 | 1.02E+00 |

| | | | | | | |
|---|---|---|---|---|---|---|
| ENSG000001 | RBP4 | 5.95E-01 | ? | 1.26E+00 | 3.39E-01 | 1.26E+00 |
| ENSG000002 | LINC01176 | 5.95E-01 | 8.45E-01 | 1.10E+00 | 1.33E-01 | 1.10E+00 |
| ENSG000002 | ZNF316 | 5.95E-01 | 8.45E-01 | 9.66E-01 | -4.96E-02 | -1.03E+00 |
| ENSG000001 | ZFTRAF1 | 5.95E-01 | 8.45E-01 | 1.04E+00 | 5.86E-02 | 1.04E+00 |
| ENSG00000C | LRCH4 | 5.95E-01 | 8.45E-01 | 9.73E-01 | -4.00E-02 | -1.03E+00 |
| ENSG000002 | --- | 5.95E-01 | ? | 1.43E+00 | 5.19E-01 | 1.43E+00 |
| ENSG000002 | --- | 5.95E-01 | ? | 8.06E-01 | -3.11E-01 | -1.24E+00 |
| ENSG000002 | AS3MT | 5.95E-01 | ? | 1.39E+00 | 4.79E-01 | 1.39E+00 |
| ENSG000001 | DRICH1 | 5.95E-01 | ? | 1.30E+00 | 3.78E-01 | 1.30E+00 |
| ENSG000002 | POLR1HASP | 5.95E-01 | 8.46E-01 | 9.55E-01 | -6.66E-02 | -1.05E+00 |
| ENSG000001 | GPR155 | 5.95E-01 | 8.46E-01 | 9.79E-01 | -3.08E-02 | -1.02E+00 |
| ENSG00000C | EYA2 | 5.95E-01 | 8.46E-01 | 1.03E+00 | 3.62E-02 | 1.03E+00 |
| ENSG000002 | PHB1P9 | 5.95E-01 | ? | 8.37E-01 | -2.56E-01 | -1.19E+00 |
| ENSG000001 | LMBRD2 | 5.95E-01 | 8.46E-01 | 9.73E-01 | -3.97E-02 | -1.03E+00 |
| ENSG000001 | DCTN6 | 5.96E-01 | 8.46E-01 | 9.71E-01 | -4.18E-02 | -1.03E+00 |
| ENSG000001 | NFRKB | 5.96E-01 | 8.46E-01 | 9.79E-01 | -3.08E-02 | -1.02E+00 |
| ENSG000001 | CRELD2 | 5.96E-01 | 8.46E-01 | 1.03E+00 | 4.14E-02 | 1.03E+00 |
| ENSG000002 | C6orf163 | 5.96E-01 | ? | 9.03E-01 | -1.47E-01 | -1.11E+00 |
| ENSG000001 | ZDHHC23 | 5.96E-01 | 8.46E-01 | 1.04E+00 | 5.35E-02 | 1.04E+00 |
| ENSG000001 | TCEAL9 | 5.96E-01 | 8.46E-01 | 9.79E-01 | -3.01E-02 | -1.02E+00 |
| ENSG000002 | C13orf46 | 5.96E-01 | ? | 8.82E-01 | -1.81E-01 | -1.13E+00 |
| ENSG000001 | SELENOI | 5.96E-01 | 8.46E-01 | 1.02E+00 | 3.00E-02 | 1.02E+00 |
| ENSG000001 | APRG1 | 5.96E-01 | ? | 1.11E+00 | 1.47E-01 | 1.11E+00 |
| ENSG000002 | --- | 5.96E-01 | ? | 1.22E+00 | 2.82E-01 | 1.22E+00 |
| ENSG000001 | ZC3H7B | 5.96E-01 | 8.46E-01 | 1.02E+00 | 2.66E-02 | 1.02E+00 |
| ENSG000001 | LRCH3 | 5.96E-01 | 8.46E-01 | 1.02E+00 | 2.46E-02 | 1.02E+00 |
| ENSG000002 | --- | 5.96E-01 | ? | 8.77E-01 | -1.89E-01 | -1.14E+00 |
| ENSG000001 | BET1 | 5.96E-01 | 8.47E-01 | 1.02E+00 | 3.26E-02 | 1.02E+00 |
| ENSG000002 | --- | 5.97E-01 | ? | 7.93E-01 | -3.35E-01 | -1.26E+00 |
| ENSG000002 | --- | 5.97E-01 | ? | 8.62E-01 | -2.14E-01 | -1.16E+00 |
| ENSG000001 | NMB | 5.97E-01 | 8.47E-01 | 1.08E+00 | 1.13E-01 | 1.08E+00 |
| ENSG000002 | --- | 5.97E-01 | ? | 8.66E-01 | -2.07E-01 | -1.15E+00 |
| ENSG000002 | --- | 5.97E-01 | ? | 1.34E+00 | 4.22E-01 | 1.34E+00 |
| ENSG00000C | PKP2 | 5.97E-01 | 8.47E-01 | 9.76E-01 | -3.58E-02 | -1.03E+00 |
| ENSG000002 | --- | 5.97E-01 | ? | 9.08E-01 | -1.39E-01 | -1.10E+00 |
| ENSG000002 | --- | 5.97E-01 | ? | 8.68E-01 | -2.04E-01 | -1.15E+00 |
| ENSG000001 | CERS3 | 5.97E-01 | ? | 1.20E+00 | 2.68E-01 | 1.20E+00 |
| ENSG000001 | MKKS | 5.97E-01 | 8.47E-01 | 9.77E-01 | -3.29E-02 | -1.02E+00 |
| ENSG000001 | TDRD12 | 5.97E-01 | 8.47E-01 | 9.54E-01 | -6.78E-02 | -1.05E+00 |
| ENSG000002 | --- | 5.97E-01 | 8.47E-01 | 9.13E-01 | -1.31E-01 | -1.09E+00 |
| ENSG000001 | COMP | 5.97E-01 | ? | 1.22E+00 | 2.87E-01 | 1.22E+00 |
| ENSG000001 | RCOR3 | 5.97E-01 | 8.47E-01 | 9.75E-01 | -3.73E-02 | -1.03E+00 |
| ENSG000001 | CATSPER3 | 5.97E-01 | ? | 8.18E-01 | -2.91E-01 | -1.22E+00 |
| ENSG000002 | RSL24D1P11 | 5.97E-01 | ? | 6.91E-01 | -5.33E-01 | -1.45E+00 |
| ENSG000002 | --- | 5.97E-01 | ? | 1.12E+00 | 1.67E-01 | 1.12E+00 |

| | | | | | | |
|---|---|---|---|---|---|---|
| ENSG000001 | UFM1 | 5.98E-01 | 8.47E-01 | 9.83E-01 | -2.48E-02 | -1.02E+00 |
| ENSG000001 | B4GALT4 | 5.98E-01 | 8.47E-01 | 9.78E-01 | -3.24E-02 | -1.02E+00 |
| ENSG000001 | MINDY4 | 5.98E-01 | 8.47E-01 | 9.59E-01 | -6.01E-02 | -1.04E+00 |
| ENSG000002 | --- | 5.98E-01 | 8.47E-01 | 1.06E+00 | 7.76E-02 | 1.06E+00 |
| ENSG000001 | SAMD10 | 5.98E-01 | 8.47E-01 | 1.06E+00 | 8.60E-02 | 1.06E+00 |
| ENSG000001 | IFTAP | 5.98E-01 | 8.47E-01 | 9.69E-01 | -4.57E-02 | -1.03E+00 |
| ENSG000001 | LYAR | 5.98E-01 | 8.47E-01 | 9.71E-01 | -4.29E-02 | -1.03E+00 |
| ENSG000001 | NR3C2 | 5.98E-01 | 8.47E-01 | 1.11E+00 | 1.54E-01 | 1.11E+00 |
| ENSG000002 | PCDHGA7 | 5.98E-01 | 8.47E-01 | 1.08E+00 | 1.06E-01 | 1.08E+00 |
| ENSG000002 | --- | 5.98E-01 | ? | 1.13E+00 | 1.73E-01 | 1.13E+00 |
| ENSG00000C | MATR3 | 5.98E-01 | 8.47E-01 | 9.72E-01 | -4.07E-02 | -1.03E+00 |
| ENSG000001 | CLUH | 5.98E-01 | 8.47E-01 | 9.74E-01 | -3.84E-02 | -1.03E+00 |
| ENSG000001 | INTS12 | 5.98E-01 | 8.47E-01 | 9.80E-01 | -2.92E-02 | -1.02E+00 |
| ENSG000002 | --- | 5.98E-01 | ? | 1.15E+00 | 2.03E-01 | 1.15E+00 |
| ENSG000002 | ATP5PDP4 | 5.98E-01 | ? | 1.24E+00 | 3.12E-01 | 1.24E+00 |
| ENSG000001 | HSH2D | 5.98E-01 | ? | 1.25E+00 | 3.26E-01 | 1.25E+00 |
| ENSG000001 | CUBN | 5.98E-01 | 8.47E-01 | 9.10E-01 | -1.36E-01 | -1.10E+00 |
| ENSG000001 | ZC3H12B | 5.98E-01 | ? | 9.05E-01 | -1.44E-01 | -1.10E+00 |
| ENSG000002 | RPL23AP7 | 5.98E-01 | 8.47E-01 | 1.03E+00 | 4.17E-02 | 1.03E+00 |
| ENSG000001 | ITM2B | 5.98E-01 | 8.47E-01 | 9.84E-01 | -2.36E-02 | -1.02E+00 |
| ENSG000001 | REEP5 | 5.98E-01 | 8.47E-01 | 9.87E-01 | -1.90E-02 | -1.01E+00 |
| ENSG000002 | LINC01128 | 5.99E-01 | 8.47E-01 | 9.80E-01 | -2.93E-02 | -1.02E+00 |
| ENSG000001 | NUDT22 | 5.99E-01 | 8.47E-01 | 1.04E+00 | 6.04E-02 | 1.04E+00 |
| ENSG000002 | --- | 5.99E-01 | ? | 8.40E-01 | -2.52E-01 | -1.19E+00 |
| ENSG00000C | FAT1 | 5.99E-01 | 8.47E-01 | 1.03E+00 | 4.87E-02 | 1.03E+00 |
| ENSG000001 | HIPK3 | 5.99E-01 | 8.47E-01 | 1.02E+00 | 2.61E-02 | 1.02E+00 |
| ENSG00000C | HDAC7 | 5.99E-01 | 8.47E-01 | 1.03E+00 | 3.76E-02 | 1.03E+00 |
| ENSG000002 | PTGES3L | 5.99E-01 | ? | 1.15E+00 | 2.02E-01 | 1.15E+00 |
| ENSG000001 | CCDC107 | 5.99E-01 | 8.47E-01 | 9.67E-01 | -4.79E-02 | -1.03E+00 |
| ENSG000001 | TIMM10B | 5.99E-01 | 8.47E-01 | 9.76E-01 | -3.45E-02 | -1.02E+00 |
| ENSG00000C | NR1H3 | 5.99E-01 | 8.47E-01 | 9.47E-01 | -7.92E-02 | -1.06E+00 |
| ENSG000002 | SNORD3A | 5.99E-01 | ? | 8.49E-01 | -2.36E-01 | -1.18E+00 |
| ENSG000002 | GAS5 | 5.99E-01 | 8.47E-01 | 1.02E+00 | 3.03E-02 | 1.02E+00 |
| ENSG000002 | CKMT1B | 5.99E-01 | 8.47E-01 | 1.02E+00 | 3.30E-02 | 1.02E+00 |
| ENSG000001 | SRRM1 | 5.99E-01 | 8.47E-01 | 9.78E-01 | -3.19E-02 | -1.02E+00 |
| ENSG000002 | ZNF337-AS1 | 5.99E-01 | 8.47E-01 | 9.34E-01 | -9.90E-02 | -1.07E+00 |
| ENSG000001 | GDPD1 | 5.99E-01 | 8.47E-01 | 9.49E-01 | -7.49E-02 | -1.05E+00 |
| ENSG000002 | --- | 5.99E-01 | ? | 8.34E-01 | -2.62E-01 | -1.20E+00 |
| ENSG000002 | TMEM191C | 5.99E-01 | ? | 9.04E-01 | -1.46E-01 | -1.11E+00 |
| ENSG000001 | C14orf28 | 5.99E-01 | 8.48E-01 | 1.04E+00 | 6.03E-02 | 1.04E+00 |
| ENSG000001 | UBQLNL | 5.99E-01 | ? | 1.20E+00 | 2.62E-01 | 1.20E+00 |
| ENSG000001 | RGPD3 | 5.99E-01 | ? | 1.17E+00 | 2.26E-01 | 1.17E+00 |
| ENSG000001 | LSP1 | 5.99E-01 | ? | 1.33E+00 | 4.11E-01 | 1.33E+00 |
| ENSG000002 | --- | 5.99E-01 | 8.48E-01 | 1.10E+00 | 1.42E-01 | 1.10E+00 |
| ENSG000002 | --- | 6.00E-01 | ? | 1.26E+00 | 3.34E-01 | 1.26E+00 |

| | | | | | | |
|---|---|---|---|---|---|---|
| ENSG000002 | PCDHAC1 | 6.00E-01 | 8.48E-01 | 9.47E-01 | -7.86E-02 | -1.06E+00 |
| ENSG000002 | LINC01719 | 6.00E-01 | 8.48E-01 | 9.60E-01 | -5.96E-02 | -1.04E+00 |
| ENSG000002 | PMS2P7 | 6.00E-01 | ? | 9.00E-01 | -1.52E-01 | -1.11E+00 |
| ENSG000002 | --- | 6.00E-01 | ? | 1.12E+00 | 1.59E-01 | 1.12E+00 |
| ENSG000002 | SUCLG2-DT | 6.00E-01 | 8.48E-01 | 1.05E+00 | 6.36E-02 | 1.05E+00 |
| ENSG00000C | MTMR11 | 6.00E-01 | 8.48E-01 | 9.66E-01 | -5.02E-02 | -1.04E+00 |
| ENSG000002 | --- | 6.00E-01 | 8.48E-01 | 9.31E-01 | -1.04E-01 | -1.07E+00 |
| ENSG00000C | FAM107B | 6.00E-01 | 8.48E-01 | 1.03E+00 | 3.67E-02 | 1.03E+00 |
| ENSG000002 | LINC02018 | 6.00E-01 | ? | 1.18E+00 | 2.43E-01 | 1.18E+00 |
| ENSG000001 | RP1L1 | 6.00E-01 | ? | 8.07E-01 | -3.10E-01 | -1.24E+00 |
| ENSG000002 | HEXA-AS1 | 6.00E-01 | ? | 1.11E+00 | 1.48E-01 | 1.11E+00 |
| ENSG000001 | RAB11A | 6.00E-01 | 8.48E-01 | 1.01E+00 | 1.79E-02 | 1.01E+00 |
| ENSG000002 | NDUFV2-AS1 | 6.00E-01 | ? | 1.21E+00 | 2.76E-01 | 1.21E+00 |
| ENSG000002 | NUS1P1 | 6.00E-01 | 8.49E-01 | 9.33E-01 | -9.97E-02 | -1.07E+00 |
| ENSG000001 | GRPEL1 | 6.01E-01 | 8.49E-01 | 1.02E+00 | 2.49E-02 | 1.02E+00 |
| ENSG000002 | --- | 6.01E-01 | ? | 1.18E+00 | 2.44E-01 | 1.18E+00 |
| ENSG000001 | ADAMTS5 | 6.01E-01 | 8.49E-01 | 1.07E+00 | 9.19E-02 | 1.07E+00 |
| ENSG000001 | --- | 6.01E-01 | 8.49E-01 | 9.16E-01 | -1.27E-01 | -1.09E+00 |
| ENSG000001 | NPHP4 | 6.01E-01 | 8.49E-01 | 1.03E+00 | 4.57E-02 | 1.03E+00 |
| ENSG000001 | ATP8A2 | 6.01E-01 | 8.49E-01 | 9.51E-01 | -7.20E-02 | -1.05E+00 |
| ENSG000002 | NEMP2-DT | 6.01E-01 | ? | 8.26E-01 | -2.75E-01 | -1.21E+00 |
| ENSG000002 | ZNF816-ZNF | 6.01E-01 | ? | 8.81E-01 | -1.83E-01 | -1.13E+00 |
| ENSG000002 | MT-TQ | 6.01E-01 | ? | 8.08E-01 | -3.08E-01 | -1.24E+00 |
| ENSG000001 | CYB5R3 | 6.01E-01 | 8.49E-01 | 9.83E-01 | -2.54E-02 | -1.02E+00 |
| ENSG000001 | NAGA | 6.01E-01 | 8.49E-01 | 1.03E+00 | 3.83E-02 | 1.03E+00 |
| ENSG000001 | ABTB2 | 6.01E-01 | 8.49E-01 | 9.72E-01 | -4.09E-02 | -1.03E+00 |
| ENSG000002 | RBFADN | 6.01E-01 | ? | 8.45E-01 | -2.42E-01 | -1.18E+00 |
| ENSG000001 | NUP210 | 6.01E-01 | ? | 8.92E-01 | -1.66E-01 | -1.12E+00 |
| ENSG000002 | RPL35P1 | 6.01E-01 | ? | 7.63E-01 | -3.90E-01 | -1.31E+00 |
| ENSG000001 | NDRG2 | 6.01E-01 | 8.49E-01 | 1.09E+00 | 1.19E-01 | 1.09E+00 |
| ENSG000002 | LINC01424 | 6.01E-01 | ? | 8.59E-01 | -2.19E-01 | -1.16E+00 |
| ENSG000001 | CHURC1-FNT | 6.01E-01 | ? | 6.51E-01 | -6.20E-01 | -1.54E+00 |
| ENSG000002 | ING2-DT | 6.01E-01 | ? | 1.21E+00 | 2.71E-01 | 1.21E+00 |
| ENSG000001 | CCT6B | 6.01E-01 | ? | 1.10E+00 | 1.41E-01 | 1.10E+00 |
| ENSG000002 | --- | 6.01E-01 | ? | 8.92E-01 | -1.64E-01 | -1.12E+00 |
| ENSG000002 | SFTA3 | 6.01E-01 | 8.49E-01 | 1.09E+00 | 1.20E-01 | 1.09E+00 |
| ENSG000002 | LINC02846 | 6.01E-01 | ? | 1.21E+00 | 2.79E-01 | 1.21E+00 |
| ENSG000001 | SERPINF2 | 6.01E-01 | 8.49E-01 | 1.06E+00 | 8.49E-02 | 1.06E+00 |
| ENSG000001 | PRR12 | 6.01E-01 | 8.49E-01 | 9.69E-01 | -4.60E-02 | -1.03E+00 |
| ENSG000002 | ZNF324B | 6.01E-01 | 8.49E-01 | 1.04E+00 | 5.74E-02 | 1.04E+00 |
| ENSG000001 | ANKRD31 | 6.02E-01 | ? | 1.17E+00 | 2.31E-01 | 1.17E+00 |
| ENSG000001 | TMEM44 | 6.02E-01 | 8.49E-01 | 9.76E-01 | -3.56E-02 | -1.03E+00 |
| ENSG000001 | ANP32B | 6.02E-01 | 8.49E-01 | 1.02E+00 | 3.04E-02 | 1.02E+00 |
| ENSG000002 | NALT1 | 6.02E-01 | ? | 1.13E+00 | 1.79E-01 | 1.13E+00 |
| ENSG000001 | DMRT2 | 6.02E-01 | 8.49E-01 | 9.11E-01 | -1.34E-01 | -1.10E+00 |

| | | | | | | |
|---|---|---|---|---|---|---|
| ENSG000001 | HSD11B2 | 6.02E-01 | 8.49E-01 | 1.08E+00 | 1.08E-01 | 1.08E+00 |
| ENSG000001 | TULP4 | 6.02E-01 | 8.49E-01 | 9.74E-01 | -3.81E-02 | -1.03E+00 |
| ENSG000002 | GUSBP14 | 6.02E-01 | ? | 1.10E+00 | 1.34E-01 | 1.10E+00 |
| ENSG000002 | --- | 6.02E-01 | ? | 8.18E-01 | -2.90E-01 | -1.22E+00 |
| ENSG000002 | ANKRD65 | 6.02E-01 | 8.49E-01 | 9.66E-01 | -4.98E-02 | -1.04E+00 |
| ENSG000001 | OSR1 | 6.02E-01 | ? | 1.12E+00 | 1.69E-01 | 1.12E+00 |
| ENSG000001 | RSPH4A | 6.02E-01 | ? | 8.16E-01 | -2.93E-01 | -1.22E+00 |
| ENSG000001 | MUC15 | 6.02E-01 | ? | 1.16E+00 | 2.10E-01 | 1.16E+00 |
| ENSG000002 | --- | 6.02E-01 | ? | 1.14E+00 | 1.84E-01 | 1.14E+00 |
| ENSG000002 | SPRN | 6.02E-01 | 8.49E-01 | 9.24E-01 | -1.14E-01 | -1.08E+00 |
| ENSG000001 | TMEM268 | 6.02E-01 | 8.49E-01 | 1.03E+00 | 4.53E-02 | 1.03E+00 |
| ENSG000002 | HNRNPA1P1( | 6.02E-01 | ? | 8.84E-01 | -1.78E-01 | -1.13E+00 |
| ENSG000001 | N4BP2L1 | 6.02E-01 | ? | 1.10E+00 | 1.36E-01 | 1.10E+00 |
| ENSG00000C | RASSF7 | 6.02E-01 | 8.49E-01 | 1.06E+00 | 7.76E-02 | 1.06E+00 |
| ENSG000001 | SERPINE2 | 6.03E-01 | 8.49E-01 | 9.77E-01 | -3.33E-02 | -1.02E+00 |
| ENSG000001 | H2BC4 | 6.03E-01 | 8.49E-01 | 9.71E-01 | -4.29E-02 | -1.03E+00 |
| ENSG000001 | ELK4 | 6.03E-01 | 8.49E-01 | 9.78E-01 | -3.24E-02 | -1.02E+00 |
| ENSG000001 | RPL32P18 | 6.03E-01 | ? | 1.09E+00 | 1.18E-01 | 1.09E+00 |
| ENSG000001 | SCUBE2 | 6.03E-01 | ? | 7.77E-01 | -3.64E-01 | -1.29E+00 |
| ENSG000002 | --- | 6.03E-01 | 8.50E-01 | 9.63E-01 | -5.42E-02 | -1.04E+00 |
| ENSG000001 | PLSCR4 | 6.03E-01 | 8.50E-01 | 1.05E+00 | 7.47E-02 | 1.05E+00 |
| ENSG000001 | LRRC4B | 6.03E-01 | ? | 7.43E-01 | -4.28E-01 | -1.35E+00 |
| ENSG000002 | DDX11-AS1 | 6.03E-01 | ? | 1.15E+00 | 2.02E-01 | 1.15E+00 |
| ENSG000002 | --- | 6.03E-01 | ? | 8.85E-01 | -1.76E-01 | -1.13E+00 |
| ENSG000002 | --- | 6.03E-01 | ? | 1.10E+00 | 1.33E-01 | 1.10E+00 |
| ENSG000002 | MARCKS | 6.03E-01 | 8.50E-01 | 1.02E+00 | 3.15E-02 | 1.02E+00 |
| ENSG000001 | CEMIP | 6.03E-01 | 8.50E-01 | 8.74E-01 | -1.94E-01 | -1.14E+00 |
| ENSG00000C | PCDHA6 | 6.03E-01 | ? | 1.11E+00 | 1.50E-01 | 1.11E+00 |
| ENSG000001 | PBK | 6.03E-01 | 8.50E-01 | 9.23E-01 | -1.15E-01 | -1.08E+00 |
| ENSG00000C | UNC13D | 6.04E-01 | 8.50E-01 | 1.05E+00 | 6.93E-02 | 1.05E+00 |
| ENSG000001 | BBS2 | 6.04E-01 | 8.50E-01 | 1.02E+00 | 2.66E-02 | 1.02E+00 |
| ENSG000002 | --- | 6.04E-01 | ? | 1.12E+00 | 1.58E-01 | 1.12E+00 |
| ENSG00000C | VCAN | 6.04E-01 | 8.50E-01 | 1.04E+00 | 5.21E-02 | 1.04E+00 |
| ENSG000002 | NOTCH4 | 6.04E-01 | 8.50E-01 | 1.07E+00 | 9.94E-02 | 1.07E+00 |
| ENSG000001 | FUBP3 | 6.04E-01 | 8.50E-01 | 1.02E+00 | 2.57E-02 | 1.02E+00 |
| ENSG000002 | --- | 6.04E-01 | ? | 1.20E+00 | 2.61E-01 | 1.20E+00 |
| ENSG000001 | NEK8 | 6.04E-01 | 8.50E-01 | 1.03E+00 | 4.72E-02 | 1.03E+00 |
| ENSG000001 | CFAP69 | 6.04E-01 | 8.50E-01 | 1.07E+00 | 9.76E-02 | 1.07E+00 |
| ENSG000002 | --- | 6.04E-01 | ? | 1.23E+00 | 2.96E-01 | 1.23E+00 |
| ENSG000002 | H4C15 | 6.04E-01 | 8.50E-01 | 1.05E+00 | 7.09E-02 | 1.05E+00 |
| ENSG000002 | MRPS6 | 6.04E-01 | 8.50E-01 | 1.02E+00 | 2.80E-02 | 1.02E+00 |
| ENSG000002 | MED8-AS1 | 6.04E-01 | ? | 8.03E-01 | -3.17E-01 | -1.25E+00 |
| ENSG00000C | GABARAPL2 | 6.04E-01 | 8.50E-01 | 1.02E+00 | 2.93E-02 | 1.02E+00 |
| ENSG000002 | RING1 | 6.04E-01 | 8.51E-01 | 9.81E-01 | -2.81E-02 | -1.02E+00 |
| ENSG000001 | NOS3 | 6.04E-01 | ? | 1.22E+00 | 2.81E-01 | 1.22E+00 |

| | | | | | | | |
|---|---|---|---|---|---|---|---|
| ENSG000001 | CHD2 | 6.05E-01 | | 8.51E-01 | 9.79E-01 | -3.08E-02 | -1.02E+00 |
| ENSG000002 | ITGA1 | 6.05E-01 | | 8.51E-01 | 9.67E-01 | -4.81E-02 | -1.03E+00 |
| ENSG00000C | PTCD2 | 6.05E-01 | | 8.51E-01 | 1.02E+00 | 3.35E-02 | 1.02E+00 |
| ENSG000002 | SEC24B-AS1 | 6.05E-01 | ? | | 1.12E+00 | 1.59E-01 | 1.12E+00 |
| ENSG000002 | --- | 6.05E-01 | ? | | 7.92E-01 | -3.36E-01 | -1.26E+00 |
| ENSG000001 | FIBIN | 6.05E-01 | ? | | 1.16E+00 | 2.11E-01 | 1.16E+00 |
| ENSG000001 | COPS2 | 6.05E-01 | | 8.51E-01 | 9.80E-01 | -2.89E-02 | -1.02E+00 |
| ENSG000001 | KLK8 | 6.05E-01 | ? | | 8.99E-01 | -1.54E-01 | -1.11E+00 |
| ENSG00000C | DDX18 | 6.05E-01 | | 8.51E-01 | 9.86E-01 | -2.07E-02 | -1.01E+00 |
| ENSG000002 | UBR5-DT | 6.05E-01 | ? | | 1.15E+00 | 2.03E-01 | 1.15E+00 |
| ENSG000002 | RBM33-DT | 6.05E-01 | | 8.51E-01 | 1.07E+00 | 9.75E-02 | 1.07E+00 |
| ENSG000002 | PPP1R3G | 6.05E-01 | ? | | 1.12E+00 | 1.65E-01 | 1.12E+00 |
| ENSG000001 | TMEM259 | 6.05E-01 | | 8.51E-01 | 1.03E+00 | 4.48E-02 | 1.03E+00 |
| ENSG000001 | TFPT | 6.05E-01 | | 8.51E-01 | 9.73E-01 | -4.00E-02 | -1.03E+00 |
| ENSG000001 | CNPY4 | 6.05E-01 | | 8.51E-01 | 1.04E+00 | 6.13E-02 | 1.04E+00 |
| ENSG000001 | PAPPA | 6.05E-01 | ? | | 9.11E-01 | -1.35E-01 | -1.10E+00 |
| ENSG000002 | SPON1 | 6.05E-01 | | 8.51E-01 | 9.26E-01 | -1.10E-01 | -1.08E+00 |
| ENSG000001 | PPP4R3A | 6.06E-01 | | 8.51E-01 | 9.85E-01 | -2.12E-02 | -1.01E+00 |
| ENSG000001 | AKT1 | 6.06E-01 | | 8.51E-01 | 9.78E-01 | -3.23E-02 | -1.02E+00 |
| ENSG000002 | TTLL3 | 6.06E-01 | | 8.51E-01 | 9.37E-01 | -9.42E-02 | -1.07E+00 |
| ENSG000001 | SLC25A22 | 6.06E-01 | | 8.51E-01 | 1.04E+00 | 5.74E-02 | 1.04E+00 |
| ENSG000002 | TTC3P1 | 6.06E-01 | | 8.51E-01 | 9.62E-01 | -5.56E-02 | -1.04E+00 |
| ENSG000002 | --- | 6.06E-01 | | 8.51E-01 | 9.32E-01 | -1.01E-01 | -1.07E+00 |
| ENSG000001 | TSPAN10 | 6.06E-01 | ? | | 1.13E+00 | 1.74E-01 | 1.13E+00 |
| ENSG000001 | LRIG1 | 6.06E-01 | | 8.51E-01 | 1.04E+00 | 5.30E-02 | 1.04E+00 |
| ENSG000002 | MIR3142HG | 6.06E-01 | ? | | 1.25E+00 | 3.24E-01 | 1.25E+00 |
| ENSG000001 | RSBN1L | 6.06E-01 | | 8.51E-01 | 1.02E+00 | 3.26E-02 | 1.02E+00 |
| ENSG000001 | SRPX2 | 6.06E-01 | | 8.52E-01 | 1.03E+00 | 3.60E-02 | 1.03E+00 |
| ENSG00000C | CDC7 | 6.06E-01 | | 8.52E-01 | 9.49E-01 | -7.51E-02 | -1.05E+00 |
| ENSG000002 | SUPT16HP1 | 6.06E-01 | | 8.52E-01 | 1.08E+00 | 1.05E-01 | 1.08E+00 |
| ENSG000002 | FAM246C | 6.07E-01 | ? | | 1.21E+00 | 2.77E-01 | 1.21E+00 |
| ENSG000001 | TBKBP1 | 6.07E-01 | | 8.52E-01 | 1.03E+00 | 4.61E-02 | 1.03E+00 |
| ENSG000002 | --- | 6.07E-01 | ? | | 7.52E-01 | -4.11E-01 | -1.33E+00 |
| ENSG000002 | LINC00640 | 6.07E-01 | ? | | 8.52E-01 | -2.31E-01 | -1.17E+00 |
| ENSG000001 | ATF7IP | 6.07E-01 | | 8.52E-01 | 9.77E-01 | -3.33E-02 | -1.02E+00 |
| ENSG00000C | PHLPP1 | 6.07E-01 | | 8.52E-01 | 1.03E+00 | 4.32E-02 | 1.03E+00 |
| ENSG000002 | NAV2-AS2 | 6.07E-01 | ? | | 8.26E-01 | -2.76E-01 | -1.21E+00 |
| ENSG000001 | KRT10 | 6.07E-01 | | 8.52E-01 | 1.03E+00 | 4.45E-02 | 1.03E+00 |
| ENSG000002 | --- | 6.07E-01 | | 8.52E-01 | 8.72E-01 | -1.97E-01 | -1.15E+00 |
| ENSG000002 | KCNMA1-AS1 | 6.07E-01 | ? | | 7.85E-01 | -3.48E-01 | -1.27E+00 |
| ENSG00000C | ARHGAP28 | 6.07E-01 | ? | | 1.11E+00 | 1.51E-01 | 1.11E+00 |
| ENSG000001 | MTHFD1 | 6.07E-01 | | 8.52E-01 | 9.81E-01 | -2.72E-02 | -1.02E+00 |
| ENSG00000C | GEMIN5 | 6.07E-01 | | 8.52E-01 | 9.81E-01 | -2.74E-02 | -1.02E+00 |
| ENSG000002 | --- | 6.07E-01 | | 8.52E-01 | 1.05E+00 | 7.12E-02 | 1.05E+00 |
| ENSG000002 | --- | 6.07E-01 | ? | | 1.22E+00 | 2.82E-01 | 1.22E+00 |

| | | | | | | | |
|---|---|---|---|---|---|---|---|
| ENSG000001 | EBNA1BP2 | 6.07E-01 | | 8.52E-01 | 9.82E-01 | -2.56E-02 | -1.02E+00 |
| ENSG000002 | LINC00923 | 6.08E-01 | | 8.52E-01 | 1.09E+00 | 1.22E-01 | 1.09E+00 |
| ENSG000001 | CGB7 | 6.08E-01 | | 8.53E-01 | 1.05E+00 | 6.60E-02 | 1.05E+00 |
| ENSG000002 | --- | 6.08E-01 | ? | | 1.31E+00 | 3.92E-01 | 1.31E+00 |
| ENSG00000C | QSER1 | 6.08E-01 | | 8.53E-01 | 1.02E+00 | 3.36E-02 | 1.02E+00 |
| ENSG000001 | PCNX4 | 6.08E-01 | | 8.53E-01 | 9.83E-01 | -2.51E-02 | -1.02E+00 |
| ENSG000001 | EPC1 | 6.08E-01 | | 8.53E-01 | 1.02E+00 | 3.15E-02 | 1.02E+00 |
| ENSG000002 | --- | 6.08E-01 | ? | | 1.26E+00 | 3.39E-01 | 1.26E+00 |
| ENSG000002 | ITGA9-AS1 | 6.08E-01 | | 8.53E-01 | 1.09E+00 | 1.21E-01 | 1.09E+00 |
| ENSG000002 | --- | 6.08E-01 | ? | | 8.40E-01 | -2.52E-01 | -1.19E+00 |
| ENSG00000C | CD74 | 6.08E-01 | | 8.53E-01 | 9.72E-01 | -4.14E-02 | -1.03E+00 |
| ENSG000002 | --- | 6.09E-01 | ? | | 1.21E+00 | 2.72E-01 | 1.21E+00 |
| ENSG000001 | TMEM53 | 6.09E-01 | | 8.53E-01 | 1.05E+00 | 6.76E-02 | 1.05E+00 |
| ENSG00000C | SYNE2 | 6.09E-01 | | 8.53E-01 | 1.06E+00 | 8.03E-02 | 1.06E+00 |
| ENSG000002 | NBPF9 | 6.09E-01 | | 8.53E-01 | 1.02E+00 | 2.66E-02 | 1.02E+00 |
| ENSG000001 | PCNPP5 | 6.09E-01 | ? | | 7.73E-01 | -3.72E-01 | -1.29E+00 |
| ENSG000002 | FAM91A3P | 6.09E-01 | ? | | 1.24E+00 | 3.13E-01 | 1.24E+00 |
| ENSG000001 | CNFN | 6.09E-01 | ? | | 8.35E-01 | -2.60E-01 | -1.20E+00 |
| ENSG000001 | NDUFS5 | 6.09E-01 | | 8.54E-01 | 1.01E+00 | 2.12E-02 | 1.01E+00 |
| ENSG000002 | RNF115 | 6.09E-01 | | 8.54E-01 | 9.76E-01 | -3.48E-02 | -1.02E+00 |
| ENSG000002 | PFN1P2 | 6.09E-01 | | 8.54E-01 | 9.52E-01 | -7.04E-02 | -1.05E+00 |
| ENSG000002 | LTB4R2 | 6.09E-01 | | 8.54E-01 | 1.04E+00 | 5.93E-02 | 1.04E+00 |
| ENSG000001 | CLDN15 | 6.09E-01 | | 8.54E-01 | 9.47E-01 | -7.81E-02 | -1.06E+00 |
| ENSG000001 | BDH2 | 6.09E-01 | | 8.54E-01 | 1.03E+00 | 4.54E-02 | 1.03E+00 |
| ENSG000001 | SWAP70 | 6.09E-01 | | 8.54E-01 | 1.02E+00 | 2.66E-02 | 1.02E+00 |
| ENSG000001 | SMOC2 | 6.09E-01 | ? | | 1.09E+00 | 1.23E-01 | 1.09E+00 |
| ENSG000002 | --- | 6.09E-01 | | 8.54E-01 | 7.43E-01 | -4.29E-01 | -1.35E+00 |
| ENSG000001 | ZFAND5 | 6.09E-01 | | 8.54E-01 | 1.01E+00 | 1.83E-02 | 1.01E+00 |
| ENSG000001 | RBM24 | 6.10E-01 | | 8.54E-01 | 9.27E-01 | -1.09E-01 | -1.08E+00 |
| ENSG000001 | TMEM106A | 6.10E-01 | | 8.54E-01 | 9.60E-01 | -5.81E-02 | -1.04E+00 |
| ENSG000002 | LINC01356 | 6.10E-01 | ? | | 1.14E+00 | 1.87E-01 | 1.14E+00 |
| ENSG000001 | TCEAL8 | 6.10E-01 | | 8.54E-01 | 1.02E+00 | 3.16E-02 | 1.02E+00 |
| ENSG000002 | MIRLET7IHG | 6.10E-01 | | 8.54E-01 | 9.30E-01 | -1.04E-01 | -1.07E+00 |
| ENSG000001 | MBIP | 6.10E-01 | | 8.54E-01 | 9.71E-01 | -4.23E-02 | -1.03E+00 |
| ENSG000002 | EIF3EP1 | 6.10E-01 | | 8.54E-01 | 9.06E-01 | -1.42E-01 | -1.10E+00 |
| ENSG000001 | THAP11 | 6.10E-01 | | 8.54E-01 | 1.04E+00 | 4.98E-02 | 1.04E+00 |
| ENSG000001 | IKBKB | 6.10E-01 | | 8.54E-01 | 9.81E-01 | -2.72E-02 | -1.02E+00 |
| ENSG000002 | BLACAT1 | 6.10E-01 | | 8.54E-01 | 1.07E+00 | 1.01E-01 | 1.07E+00 |
| ENSG000001 | NAALADL1 | 6.10E-01 | | 8.54E-01 | 1.08E+00 | 1.12E-01 | 1.08E+00 |
| ENSG000002 | H2BC8 | 6.10E-01 | | 8.54E-01 | 1.05E+00 | 7.50E-02 | 1.05E+00 |
| ENSG000001 | RYR3 | 6.10E-01 | | 8.54E-01 | 9.38E-01 | -9.25E-02 | -1.07E+00 |
| ENSG000002 | --- | 6.11E-01 | ? | | 8.59E-01 | -2.20E-01 | -1.16E+00 |
| ENSG000001 | IL18R1 | 6.11E-01 | | 8.54E-01 | 9.56E-01 | -6.49E-02 | -1.05E+00 |
| ENSG000001 | RICTOR | 6.11E-01 | | 8.54E-01 | 1.03E+00 | 3.57E-02 | 1.03E+00 |
| ENSG000002 | LINC01315 | 6.11E-01 | ? | | 1.29E+00 | 3.70E-01 | 1.29E+00 |

| | | | | | | |
|---|---|---|---|---|---|---|
| ENSG000001 | GPT2 | 6.11E-01 | 8.54E-01 | 9.75E-01 | -3.66E-02 | -1.03E+00 |
| ENSG000001 | STKLD1 | 6.11E-01 | ? | 1.26E+00 | 3.36E-01 | 1.26E+00 |
| ENSG000001 | COQ6 | 6.11E-01 | 8.54E-01 | 1.03E+00 | 3.94E-02 | 1.03E+00 |
| ENSG000001 | PADI3 | 6.11E-01 | 8.55E-01 | 9.54E-01 | -6.84E-02 | -1.05E+00 |
| ENSG000001 | --- | 6.11E-01 | ? | 9.07E-01 | -1.41E-01 | -1.10E+00 |
| ENSG000001 | ELK1 | 6.11E-01 | 8.55E-01 | 1.03E+00 | 3.69E-02 | 1.03E+00 |
| ENSG000002 | AIDAP1 | 6.11E-01 | ? | 8.38E-01 | -2.55E-01 | -1.19E+00 |
| ENSG000002 | --- | 6.11E-01 | ? | 1.11E+00 | 1.53E-01 | 1.11E+00 |
| ENSG000002 | EPCAM-DT | 6.11E-01 | ? | 8.53E-01 | -2.30E-01 | -1.17E+00 |
| ENSG000001 | RCN3 | 6.11E-01 | ? | 8.28E-01 | -2.73E-01 | -1.21E+00 |
| ENSG000002 | TNFRSF25 | 6.12E-01 | 8.55E-01 | 9.57E-01 | -6.41E-02 | -1.05E+00 |
| ENSG000002 | SEPSECS-AS1 | 6.12E-01 | ? | 1.19E+00 | 2.53E-01 | 1.19E+00 |
| ENSG000001 | CYBC1 | 6.12E-01 | 8.55E-01 | 9.70E-01 | -4.34E-02 | -1.03E+00 |
| ENSG000001 | RFX1 | 6.12E-01 | 8.55E-01 | 1.04E+00 | 6.04E-02 | 1.04E+00 |
| ENSG000001 | RPL23AP82 | 6.12E-01 | ? | 5.84E-01 | -7.77E-01 | -1.71E+00 |
| ENSG000001 | DEF8 | 6.12E-01 | 8.55E-01 | 1.03E+00 | 4.32E-02 | 1.03E+00 |
| ENSG000001 | FGD2 | 6.12E-01 | ? | 7.92E-01 | -3.37E-01 | -1.26E+00 |
| ENSG000002 | --- | 6.12E-01 | ? | 8.45E-01 | -2.43E-01 | -1.18E+00 |
| ENSG000001 | RAB11B | 6.12E-01 | 8.55E-01 | 1.02E+00 | 3.44E-02 | 1.02E+00 |
| ENSG000001 | CDC42EP1 | 6.12E-01 | 8.55E-01 | 1.03E+00 | 3.76E-02 | 1.03E+00 |
| ENSG000001 | SLX4 | 6.12E-01 | 8.55E-01 | 9.82E-01 | -2.55E-02 | -1.02E+00 |
| ENSG00000C | CPSF1 | 6.12E-01 | 8.55E-01 | 1.03E+00 | 4.26E-02 | 1.03E+00 |
| ENSG000002 | --- | 6.12E-01 | ? | 1.13E+00 | 1.80E-01 | 1.13E+00 |
| ENSG000002 | LINC02551 | 6.12E-01 | 8.55E-01 | 1.05E+00 | 7.32E-02 | 1.05E+00 |
| ENSG00000C | ADD1 | 6.12E-01 | 8.55E-01 | 9.91E-01 | -1.27E-02 | -1.01E+00 |
| ENSG000001 | CGB5 | 6.12E-01 | ? | 1.27E+00 | 3.42E-01 | 1.27E+00 |
| ENSG000002 | --- | 6.12E-01 | ? | 8.59E-01 | -2.18E-01 | -1.16E+00 |
| ENSG000002 | EMC1-AS1 | 6.12E-01 | ? | 8.27E-01 | -2.73E-01 | -1.21E+00 |
| ENSG000002 | LINC02028 | 6.12E-01 | ? | 7.96E-01 | -3.30E-01 | -1.26E+00 |
| ENSG00000C | LIG3 | 6.12E-01 | 8.55E-01 | 9.79E-01 | -3.00E-02 | -1.02E+00 |
| ENSG000001 | PEX1 | 6.12E-01 | 8.55E-01 | 9.75E-01 | -3.62E-02 | -1.03E+00 |
| ENSG00000C | NDUFS1 | 6.12E-01 | 8.55E-01 | 9.86E-01 | -1.99E-02 | -1.01E+00 |
| ENSG000001 | SFRP1 | 6.13E-01 | 8.55E-01 | 1.15E+00 | 2.01E-01 | 1.15E+00 |
| ENSG000002 | --- | 6.13E-01 | ? | 7.50E-01 | -4.15E-01 | -1.33E+00 |
| ENSG000001 | RBM34 | 6.13E-01 | 8.55E-01 | 9.71E-01 | -4.26E-02 | -1.03E+00 |
| ENSG000002 | --- | 6.13E-01 | ? | 8.69E-01 | -2.03E-01 | -1.15E+00 |
| ENSG000001 | CCNJL | 6.13E-01 | 8.55E-01 | 1.02E+00 | 2.72E-02 | 1.02E+00 |
| ENSG000001 | SMIM14 | 6.13E-01 | 8.55E-01 | 1.02E+00 | 2.45E-02 | 1.02E+00 |
| ENSG000001 | CLSTN3 | 6.13E-01 | 8.55E-01 | 1.02E+00 | 3.11E-02 | 1.02E+00 |
| ENSG00000C | EPDR1 | 6.13E-01 | 8.55E-01 | 9.57E-01 | -6.37E-02 | -1.05E+00 |
| ENSG000002 | ZKSCAN2-DT | 6.13E-01 | 8.55E-01 | 1.08E+00 | 1.06E-01 | 1.08E+00 |
| ENSG000002 | --- | 6.13E-01 | ? | 1.30E+00 | 3.77E-01 | 1.30E+00 |
| ENSG00000C | ME2 | 6.13E-01 | 8.56E-01 | 9.83E-01 | -2.53E-02 | -1.02E+00 |
| ENSG000002 | GTF2IP1 | 6.13E-01 | 8.56E-01 | 1.03E+00 | 4.81E-02 | 1.03E+00 |
| ENSG000002 | SLC25A21-AS | 6.13E-01 | ? | 1.13E+00 | 1.71E-01 | 1.13E+00 |

| | | | | | | |
|---|---|---|---|---|---|---|
| ENSG000002 | --- | 6.13E-01 | ? | 8.33E-01 | -2.64E-01 | -1.20E+00 |
| ENSG000001 | SETD1B | 6.13E-01 | 8.56E-01 | 9.80E-01 | -2.94E-02 | -1.02E+00 |
| ENSG000002 | --- | 6.14E-01 | ? | 1.15E+00 | 2.03E-01 | 1.15E+00 |
| ENSG000001 | CD53 | 6.14E-01 | ? | 7.63E-01 | -3.90E-01 | -1.31E+00 |
| ENSG000001 | NDUFB5 | 6.14E-01 | 8.56E-01 | 1.01E+00 | 2.15E-02 | 1.01E+00 |
| ENSG000001 | NACC1 | 6.14E-01 | 8.56E-01 | 1.02E+00 | 3.34E-02 | 1.02E+00 |
| ENSG00000C | RTF2 | 6.14E-01 | 8.56E-01 | 1.01E+00 | 1.76E-02 | 1.01E+00 |
| ENSG000001 | EMC6 | 6.14E-01 | 8.56E-01 | 1.03E+00 | 3.90E-02 | 1.03E+00 |
| ENSG000002 | RAB12 | 6.14E-01 | 8.56E-01 | 9.83E-01 | -2.54E-02 | -1.02E+00 |
| ENSG000001 | EIF4EBP2 | 6.14E-01 | 8.56E-01 | 9.85E-01 | -2.18E-02 | -1.02E+00 |
| ENSG000001 | STX17 | 6.14E-01 | 8.56E-01 | 9.82E-01 | -2.64E-02 | -1.02E+00 |
| ENSG00000C | SORBS1 | 6.14E-01 | 8.56E-01 | 1.06E+00 | 8.47E-02 | 1.06E+00 |
| ENSG000002 | --- | 6.14E-01 | ? | 8.86E-01 | -1.74E-01 | -1.13E+00 |
| ENSG000001 | SPCS3 | 6.14E-01 | 8.56E-01 | 1.02E+00 | 2.28E-02 | 1.02E+00 |
| ENSG000002 | RP9P | 6.14E-01 | 8.56E-01 | 9.57E-01 | -6.29E-02 | -1.04E+00 |
| ENSG000001 | RAI1 | 6.14E-01 | 8.56E-01 | 9.72E-01 | -4.14E-02 | -1.03E+00 |
| ENSG000001 | CLEC18B | 6.14E-01 | ? | 8.86E-01 | -1.74E-01 | -1.13E+00 |
| ENSG000002 | IPO8P1 | 6.15E-01 | ? | 7.92E-01 | -3.36E-01 | -1.26E+00 |
| ENSG000002 | C19orf38 | 6.15E-01 | ? | 1.13E+00 | 1.74E-01 | 1.13E+00 |
| ENSG000002 | --- | 6.15E-01 | ? | 1.47E+00 | 5.59E-01 | 1.47E+00 |
| ENSG000001 | CD37 | 6.15E-01 | ? | 9.16E-01 | -1.27E-01 | -1.09E+00 |
| ENSG000001 | RHBDD1 | 6.15E-01 | 8.57E-01 | 1.01E+00 | 2.13E-02 | 1.01E+00 |
| ENSG000002 | --- | 6.15E-01 | ? | 1.27E+00 | 3.41E-01 | 1.27E+00 |
| ENSG000002 | CAP2P1 | 6.15E-01 | ? | 1.28E+00 | 3.51E-01 | 1.28E+00 |
| ENSG00000C | BOD1L1 | 6.15E-01 | 8.57E-01 | 9.74E-01 | -3.81E-02 | -1.03E+00 |
| ENSG000001 | SLC5A2 | 6.15E-01 | ? | 1.25E+00 | 3.16E-01 | 1.25E+00 |
| ENSG000002 | --- | 6.15E-01 | ? | 1.14E+00 | 1.83E-01 | 1.14E+00 |
| ENSG00000C | GPC1 | 6.15E-01 | 8.57E-01 | 1.03E+00 | 4.83E-02 | 1.03E+00 |
| ENSG000001 | RTL8C | 6.15E-01 | 8.57E-01 | 9.72E-01 | -4.07E-02 | -1.03E+00 |
| ENSG000002 | COSMOC | 6.15E-01 | ? | 8.53E-01 | -2.29E-01 | -1.17E+00 |
| ENSG00000C | ABLIM1 | 6.16E-01 | 8.57E-01 | 1.03E+00 | 3.84E-02 | 1.03E+00 |
| ENSG000002 | --- | 6.16E-01 | ? | 1.10E+00 | 1.44E-01 | 1.10E+00 |
| ENSG000002 | MRPS31P5 | 6.16E-01 | 8.57E-01 | 9.60E-01 | -5.87E-02 | -1.04E+00 |
| ENSG000002 | GOLGA8O | 6.16E-01 | 8.57E-01 | 9.56E-01 | -6.51E-02 | -1.05E+00 |
| ENSG000002 | PRDM4-AS1 | 6.16E-01 | ? | 8.87E-01 | -1.73E-01 | -1.13E+00 |
| ENSG000001 | DNAAF2 | 6.16E-01 | 8.57E-01 | 9.77E-01 | -3.42E-02 | -1.02E+00 |
| ENSG000001 | MCF2L | 6.16E-01 | ? | 1.36E+00 | 4.47E-01 | 1.36E+00 |
| ENSG000001 | MTRES1 | 6.16E-01 | 8.57E-01 | 1.02E+00 | 3.36E-02 | 1.02E+00 |
| ENSG000001 | PIH1D1 | 6.16E-01 | 8.57E-01 | 9.79E-01 | -3.07E-02 | -1.02E+00 |
| ENSG000001 | ATP5PF | 6.16E-01 | 8.57E-01 | 1.02E+00 | 2.19E-02 | 1.02E+00 |
| ENSG000002 | MIR130AHG | 6.16E-01 | ? | 8.95E-01 | -1.60E-01 | -1.12E+00 |
| ENSG000001 | CCDC28B | 6.16E-01 | 8.57E-01 | 9.44E-01 | -8.28E-02 | -1.06E+00 |
| ENSG000002 | ZNF584-DT | 6.16E-01 | 8.57E-01 | 1.03E+00 | 4.46E-02 | 1.03E+00 |
| ENSG000002 | --- | 6.16E-01 | 8.57E-01 | 1.05E+00 | 6.44E-02 | 1.05E+00 |
| ENSG000002 | --- | 6.16E-01 | ? | 1.22E+00 | 2.84E-01 | 1.22E+00 |

| | | | | | | |
|---|---|---|---|---|---|---|
| ENSG000002 | --- | 6.16E-01 | ? | 9.15E-01 | -1.28E-01 | -1.09E+00 |
| ENSG00000C | DCBLD2 | 6.16E-01 | 8.57E-01 | 1.02E+00 | 2.39E-02 | 1.02E+00 |
| ENSG000001 | GPR153 | 6.16E-01 | 8.58E-01 | 9.60E-01 | -5.90E-02 | -1.04E+00 |
| ENSG000001 | PURB | 6.16E-01 | 8.58E-01 | 9.80E-01 | -2.91E-02 | -1.02E+00 |
| ENSG000001 | MED12 | 6.16E-01 | 8.58E-01 | 1.02E+00 | 2.62E-02 | 1.02E+00 |
| ENSG000001 | AKAP17A | 6.17E-01 | 8.58E-01 | 9.80E-01 | -2.98E-02 | -1.02E+00 |
| ENSG000002 | ERICH2 | 6.17E-01 | 8.58E-01 | 1.08E+00 | 1.17E-01 | 1.08E+00 |
| ENSG000002 | ZDHHC11B | 6.17E-01 | ? | 1.10E+00 | 1.34E-01 | 1.10E+00 |
| ENSG000002 | --- | 6.17E-01 | ? | 1.12E+00 | 1.57E-01 | 1.12E+00 |
| ENSG000002 | GTF2A1-AS1 | 6.17E-01 | ? | 1.17E+00 | 2.24E-01 | 1.17E+00 |
| ENSG000001 | GRIP2 | 6.17E-01 | ? | 1.26E+00 | 3.32E-01 | 1.26E+00 |
| ENSG000002 | CLASP1-AS1 | 6.17E-01 | ? | 1.19E+00 | 2.53E-01 | 1.19E+00 |
| ENSG000002 | TMEM30A-D | 6.17E-01 | ? | 1.12E+00 | 1.67E-01 | 1.12E+00 |
| ENSG000001 | CA14 | 6.17E-01 | ? | 1.13E+00 | 1.74E-01 | 1.13E+00 |
| ENSG000002 | SNHG7 | 6.17E-01 | 8.58E-01 | 9.60E-01 | -5.85E-02 | -1.04E+00 |
| ENSG000002 | LINC00449 | 6.17E-01 | ? | 8.13E-01 | -2.98E-01 | -1.23E+00 |
| ENSG000002 | IPO7 | 6.17E-01 | 8.58E-01 | 1.02E+00 | 2.49E-02 | 1.02E+00 |
| ENSG000001 | CRYL1 | 6.17E-01 | 8.58E-01 | 1.03E+00 | 4.30E-02 | 1.03E+00 |
| ENSG000002 | --- | 6.18E-01 | ? | 1.21E+00 | 2.75E-01 | 1.21E+00 |
| ENSG000001 | CXorf38 | 6.18E-01 | 8.59E-01 | 9.81E-01 | -2.70E-02 | -1.02E+00 |
| ENSG00000C | MNT | 6.18E-01 | 8.59E-01 | 9.78E-01 | -3.25E-02 | -1.02E+00 |
| ENSG000002 | CHMP3-AS1 | 6.18E-01 | ? | 8.67E-01 | -2.06E-01 | -1.15E+00 |
| ENSG000001 | FOXP2 | 6.18E-01 | 8.59E-01 | 9.22E-01 | -1.17E-01 | -1.08E+00 |
| ENSG000002 | --- | 6.18E-01 | ? | 7.97E-01 | -3.28E-01 | -1.25E+00 |
| ENSG000001 | ADGRV1 | 6.18E-01 | ? | 8.48E-01 | -2.38E-01 | -1.18E+00 |
| ENSG000001 | WASF2 | 6.18E-01 | 8.59E-01 | 9.88E-01 | -1.68E-02 | -1.01E+00 |
| ENSG000001 | H2AZ1 | 6.18E-01 | 8.59E-01 | 1.03E+00 | 3.72E-02 | 1.03E+00 |
| ENSG000002 | ACTR3-AS1 | 6.18E-01 | ? | 1.20E+00 | 2.69E-01 | 1.20E+00 |
| ENSG000002 | HERPUD2-AS | 6.18E-01 | 8.59E-01 | 9.78E-01 | -3.27E-02 | -1.02E+00 |
| ENSG000002 | PCDHGC4 | 6.18E-01 | ? | 1.22E+00 | 2.89E-01 | 1.22E+00 |
| ENSG000001 | ART3 | 6.18E-01 | ? | 1.24E+00 | 3.05E-01 | 1.24E+00 |
| ENSG000001 | PCDH1 | 6.18E-01 | 8.59E-01 | 9.80E-01 | -2.92E-02 | -1.02E+00 |
| ENSG000001 | ARMC12 | 6.18E-01 | ? | 8.55E-01 | -2.26E-01 | -1.17E+00 |
| ENSG000001 | SPCS1 | 6.18E-01 | 8.59E-01 | 1.02E+00 | 2.67E-02 | 1.02E+00 |
| ENSG000001 | PLAU | 6.18E-01 | 8.59E-01 | 9.81E-01 | -2.74E-02 | -1.02E+00 |
| ENSG000001 | SEM1 | 6.18E-01 | 8.59E-01 | 1.02E+00 | 2.48E-02 | 1.02E+00 |
| ENSG000001 | UQCRC2 | 6.18E-01 | 8.59E-01 | 1.01E+00 | 1.99E-02 | 1.01E+00 |
| ENSG00000C | SIRT6 | 6.19E-01 | 8.59E-01 | 1.04E+00 | 6.05E-02 | 1.04E+00 |
| ENSG000002 | --- | 6.19E-01 | ? | 8.51E-01 | -2.33E-01 | -1.17E+00 |
| ENSG000001 | METTL5 | 6.19E-01 | 8.59E-01 | 1.02E+00 | 2.71E-02 | 1.02E+00 |
| ENSG000002 | EFCAB8 | 6.19E-01 | ? | 1.21E+00 | 2.73E-01 | 1.21E+00 |
| ENSG000001 | MAF1 | 6.19E-01 | 8.59E-01 | 9.79E-01 | -2.99E-02 | -1.02E+00 |
| ENSG000001 | XDH | 6.19E-01 | 8.59E-01 | 1.03E+00 | 3.88E-02 | 1.03E+00 |
| ENSG000002 | --- | 6.19E-01 | ? | 1.13E+00 | 1.76E-01 | 1.13E+00 |
| ENSG000001 | PPFIA4 | 6.19E-01 | 8.59E-01 | 9.56E-01 | -6.44E-02 | -1.05E+00 |

| | | | | | | |
|---|---|---|---|---|---|---|
| ENSG000002 | DICER1-AS1 | 6.19E-01 | 8.59E-01 | 9.30E-01 | -1.05E-01 | -1.08E+00 |
| ENSG000002 | LINC01920 | 6.19E-01 | ? | 1.27E+00 | 3.46E-01 | 1.27E+00 |
| ENSG000002 | HDAC4-AS1 | 6.19E-01 | ? | 1.20E+00 | 2.68E-01 | 1.20E+00 |
| ENSG000002 | --- | 6.19E-01 | ? | 8.05E-01 | -3.12E-01 | -1.24E+00 |
| ENSG00000C | CLDN11 | 6.19E-01 | ? | 8.92E-01 | -1.66E-01 | -1.12E+00 |
| ENSG00000C | SLC4A8 | 6.19E-01 | 8.60E-01 | 9.60E-01 | -5.90E-02 | -1.04E+00 |
| ENSG000001 | FGD3 | 6.19E-01 | ? | 9.02E-01 | -1.49E-01 | -1.11E+00 |
| ENSG000002 | OR2A1-AS1 | 6.20E-01 | ? | 8.81E-01 | -1.84E-01 | -1.14E+00 |
| ENSG000002 | MRPS30-DT | 6.20E-01 | ? | 1.08E+00 | 1.09E-01 | 1.08E+00 |
| ENSG000001 | LEMD3 | 6.20E-01 | 8.60E-01 | 9.86E-01 | -1.99E-02 | -1.01E+00 |
| ENSG00000C | ITGB5 | 6.20E-01 | 8.60E-01 | 1.02E+00 | 3.28E-02 | 1.02E+00 |
| ENSG000001 | L1CAM | 6.20E-01 | 8.60E-01 | 1.02E+00 | 2.91E-02 | 1.02E+00 |
| ENSG000001 | ISG20 | 6.20E-01 | 8.60E-01 | 1.05E+00 | 7.14E-02 | 1.05E+00 |
| ENSG000002 | CHMP1B-AS1 | 6.20E-01 | 8.60E-01 | 1.31E+00 | 3.89E-01 | 1.31E+00 |
| ENSG000002 | --- | 6.20E-01 | ? | 1.20E+00 | 2.60E-01 | 1.20E+00 |
| ENSG000002 | ETF1P2 | 6.20E-01 | ? | 1.28E+00 | 3.54E-01 | 1.28E+00 |
| ENSG000002 | --- | 6.20E-01 | ? | 1.14E+00 | 1.84E-01 | 1.14E+00 |
| ENSG000002 | --- | 6.20E-01 | ? | 1.17E+00 | 2.23E-01 | 1.17E+00 |
| ENSG000001 | FMNL2 | 6.20E-01 | 8.60E-01 | 9.83E-01 | -2.50E-02 | -1.02E+00 |
| ENSG000002 | RNASE4 | 6.20E-01 | 8.60E-01 | 1.15E+00 | 2.03E-01 | 1.15E+00 |
| ENSG000002 | RPL5P4 | 6.20E-01 | 8.60E-01 | 1.06E+00 | 8.59E-02 | 1.06E+00 |
| ENSG000001 | PROS1 | 6.20E-01 | 8.60E-01 | 9.62E-01 | -5.54E-02 | -1.04E+00 |
| ENSG000001 | OR7D2 | 6.20E-01 | 8.60E-01 | 9.34E-01 | -9.89E-02 | -1.07E+00 |
| ENSG000001 | RBMS3 | 6.20E-01 | 8.60E-01 | 9.59E-01 | -6.01E-02 | -1.04E+00 |
| ENSG000002 | --- | 6.20E-01 | 8.60E-01 | 9.44E-01 | -8.39E-02 | -1.06E+00 |
| ENSG000001 | RILP | 6.20E-01 | 8.60E-01 | 1.05E+00 | 6.49E-02 | 1.05E+00 |
| ENSG000001 | TCAF2P1 | 6.21E-01 | ? | 1.21E+00 | 2.73E-01 | 1.21E+00 |
| ENSG000001 | MATCAP1 | 6.21E-01 | 8.60E-01 | 1.04E+00 | 5.12E-02 | 1.04E+00 |
| ENSG000001 | TRIM68 | 6.21E-01 | 8.60E-01 | 1.02E+00 | 2.98E-02 | 1.02E+00 |
| ENSG000002 | --- | 6.21E-01 | ? | 8.69E-01 | -2.02E-01 | -1.15E+00 |
| ENSG000001 | RENBP | 6.21E-01 | ? | 1.15E+00 | 2.04E-01 | 1.15E+00 |
| ENSG000002 | HNRNPA1P3 | 6.21E-01 | 8.60E-01 | 1.03E+00 | 4.44E-02 | 1.03E+00 |
| ENSG000002 | NELFE | 6.21E-01 | 8.60E-01 | 1.01E+00 | 2.00E-02 | 1.01E+00 |
| ENSG000002 | --- | 6.21E-01 | ? | 1.18E+00 | 2.34E-01 | 1.18E+00 |
| ENSG000002 | ATP6AP1L | 6.21E-01 | 8.60E-01 | 9.43E-01 | -8.42E-02 | -1.06E+00 |
| ENSG000001 | PIMREG | 6.21E-01 | 8.60E-01 | 9.02E-01 | -1.49E-01 | -1.11E+00 |
| ENSG000001 | BNC1 | 6.21E-01 | 8.60E-01 | 9.86E-01 | -2.11E-02 | -1.01E+00 |
| ENSG000002 | --- | 6.21E-01 | 8.60E-01 | 9.32E-01 | -1.02E-01 | -1.07E+00 |
| ENSG000002 | RPL12P16 | 6.21E-01 | 8.60E-01 | 9.47E-01 | -7.86E-02 | -1.06E+00 |
| ENSG000001 | S100A1 | 6.21E-01 | 8.60E-01 | 1.07E+00 | 9.68E-02 | 1.07E+00 |
| ENSG000001 | NDST2 | 6.21E-01 | 8.60E-01 | 1.02E+00 | 3.23E-02 | 1.02E+00 |
| ENSG000002 | C10orf55 | 6.21E-01 | 8.60E-01 | 9.50E-01 | -7.44E-02 | -1.05E+00 |
| ENSG000001 | FUNDC2 | 6.21E-01 | 8.60E-01 | 1.02E+00 | 2.21E-02 | 1.02E+00 |
| ENSG000001 | TIPARP | 6.21E-01 | 8.60E-01 | 9.85E-01 | -2.14E-02 | -1.01E+00 |
| ENSG000001 | S1PR4 | 6.21E-01 | ? | 1.27E+00 | 3.50E-01 | 1.27E+00 |

| | | | | | | |
|---|---|---|---|---|---|---|
| ENSG000001 | GOLGA8G | 6.22E-01 | ? | 8.65E-01 | -2.10E-01 | -1.16E+00 |
| ENSG000001 | LARP4B | 6.22E-01 | 8.60E-01 | 9.87E-01 | -1.91E-02 | -1.01E+00 |
| ENSG000001 | DVL3 | 6.22E-01 | 8.60E-01 | 9.86E-01 | -2.11E-02 | -1.01E+00 |
| ENSG000001 | COX16 | 6.22E-01 | 8.61E-01 | 1.02E+00 | 3.17E-02 | 1.02E+00 |
| ENSG000001 | SLC34A1 | 6.22E-01 | ? | 8.50E-01 | -2.35E-01 | -1.18E+00 |
| ENSG000002 | RPL32P16 | 6.22E-01 | ? | 8.38E-01 | -2.55E-01 | -1.19E+00 |
| ENSG000001 | C1orf21 | 6.22E-01 | 8.61E-01 | 1.03E+00 | 3.69E-02 | 1.03E+00 |
| ENSG000002 | TRNP1 | 6.22E-01 | 8.61E-01 | 1.03E+00 | 3.79E-02 | 1.03E+00 |
| ENSG000001 | AP3B2 | 6.22E-01 | ? | 1.10E+00 | 1.36E-01 | 1.10E+00 |
| ENSG000002 | C8orf82 | 6.22E-01 | 8.61E-01 | 9.51E-01 | -7.31E-02 | -1.05E+00 |
| ENSG000001 | FABP6 | 6.22E-01 | ? | 1.12E+00 | 1.70E-01 | 1.12E+00 |
| ENSG000001 | ITGBL1 | 6.22E-01 | ? | 1.10E+00 | 1.41E-01 | 1.10E+00 |
| ENSG000001 | SLMAP | 6.22E-01 | 8.61E-01 | 9.84E-01 | -2.34E-02 | -1.02E+00 |
| ENSG000002 | PDCD5P1 | 6.22E-01 | ? | 7.70E-01 | -3.77E-01 | -1.30E+00 |
| ENSG00000C | MRPL22 | 6.23E-01 | 8.61E-01 | 9.81E-01 | -2.76E-02 | -1.02E+00 |
| ENSG000001 | ENTPD3 | 6.23E-01 | 8.61E-01 | 9.59E-01 | -5.97E-02 | -1.04E+00 |
| ENSG000002 | STX5-DT | 6.23E-01 | ? | 8.22E-01 | -2.82E-01 | -1.22E+00 |
| ENSG000001 | FAM171A2 | 6.23E-01 | 8.61E-01 | 9.35E-01 | -9.73E-02 | -1.07E+00 |
| ENSG000001 | ANO5 | 6.23E-01 | ? | 1.29E+00 | 3.70E-01 | 1.29E+00 |
| ENSG000001 | DECR1 | 6.23E-01 | 8.61E-01 | 1.02E+00 | 2.43E-02 | 1.02E+00 |
| ENSG000001 | RADIL | 6.23E-01 | ? | 1.22E+00 | 2.88E-01 | 1.22E+00 |
| ENSG00000C | MRPL43 | 6.23E-01 | 8.62E-01 | 1.02E+00 | 2.53E-02 | 1.02E+00 |
| ENSG000002 | --- | 6.23E-01 | 8.62E-01 | 9.21E-01 | -1.19E-01 | -1.09E+00 |
| ENSG00000C | CEP170B | 6.23E-01 | 8.62E-01 | 1.03E+00 | 4.65E-02 | 1.03E+00 |
| ENSG000002 | --- | 6.23E-01 | ? | 1.19E+00 | 2.56E-01 | 1.19E+00 |
| ENSG000001 | RALB | 6.24E-01 | 8.62E-01 | 1.01E+00 | 1.89E-02 | 1.01E+00 |
| ENSG000001 | CCDC122 | 6.24E-01 | 8.62E-01 | 1.03E+00 | 4.76E-02 | 1.03E+00 |
| ENSG000001 | GPR150 | 6.24E-01 | ? | 1.26E+00 | 3.28E-01 | 1.26E+00 |
| ENSG000002 | TMCC1-DT | 6.24E-01 | 8.62E-01 | 1.07E+00 | 9.72E-02 | 1.07E+00 |
| ENSG000002 | --- | 6.24E-01 | ? | 1.15E+00 | 2.03E-01 | 1.15E+00 |
| ENSG000001 | POF1B | 6.24E-01 | 8.62E-01 | 9.73E-01 | -3.91E-02 | -1.03E+00 |
| ENSG000001 | SLC12A4 | 6.24E-01 | 8.62E-01 | 1.03E+00 | 3.78E-02 | 1.03E+00 |
| ENSG000002 | SLC25A38P1 | 6.24E-01 | ? | 7.26E-01 | -4.61E-01 | -1.38E+00 |
| ENSG000001 | YWHAG | 6.24E-01 | 8.63E-01 | 9.85E-01 | -2.17E-02 | -1.02E+00 |
| ENSG000002 | ARHGAP8 | 6.24E-01 | 8.63E-01 | 1.02E+00 | 3.25E-02 | 1.02E+00 |
| ENSG000001 | CAPN9 | 6.24E-01 | ? | 1.20E+00 | 2.67E-01 | 1.20E+00 |
| ENSG000002 | CHKB-CPT1B | 6.25E-01 | 8.63E-01 | 9.25E-01 | -1.13E-01 | -1.08E+00 |
| ENSG00000C | IFI35 | 6.25E-01 | 8.63E-01 | 9.35E-01 | -9.66E-02 | -1.07E+00 |
| ENSG000001 | STK39 | 6.25E-01 | 8.63E-01 | 1.02E+00 | 3.12E-02 | 1.02E+00 |
| ENSG000002 | SAP30-DT | 6.25E-01 | ? | 9.02E-01 | -1.49E-01 | -1.11E+00 |
| ENSG000001 | SNRPG | 6.25E-01 | 8.63E-01 | 1.02E+00 | 3.18E-02 | 1.02E+00 |
| ENSG000001 | EIF3G | 6.25E-01 | 8.63E-01 | 9.88E-01 | -1.80E-02 | -1.01E+00 |
| ENSG000001 | RCSD1 | 6.25E-01 | ? | 7.78E-01 | -3.62E-01 | -1.29E+00 |
| ENSG000002 | LINC01786 | 6.25E-01 | ? | 7.71E-01 | -3.75E-01 | -1.30E+00 |
| ENSG000002 | PSORS1C1 | 6.25E-01 | 8.63E-01 | 1.04E+00 | 6.32E-02 | 1.04E+00 |

| | | | | | | |
|---|---|---|---|---|---|---|
| ENSG000001 | ACOT4 | 6.25E-01 | ? | 8.08E-01 | -3.08E-01 | -1.24E+00 |
| ENSG000001 | ZSCAN30 | 6.25E-01 | 8.63E-01 | 9.68E-01 | -4.69E-02 | -1.03E+00 |
| ENSG000001 | CCZ1 | 6.25E-01 | 8.63E-01 | 9.83E-01 | -2.43E-02 | -1.02E+00 |
| ENSG000001 | SULF1 | 6.25E-01 | 8.63E-01 | 9.29E-01 | -1.07E-01 | -1.08E+00 |
| ENSG000001 | CDCA7L | 6.25E-01 | 8.63E-01 | 1.03E+00 | 3.63E-02 | 1.03E+00 |
| ENSG000001 | CPEB3 | 6.25E-01 | 8.63E-01 | 1.05E+00 | 7.11E-02 | 1.05E+00 |
| ENSG000001 | MDH1B | 6.25E-01 | ? | 1.16E+00 | 2.15E-01 | 1.16E+00 |
| ENSG000002 | --- | 6.26E-01 | ? | 8.19E-01 | -2.88E-01 | -1.22E+00 |
| ENSG000002 | --- | 6.26E-01 | 8.63E-01 | 9.38E-01 | -9.18E-02 | -1.07E+00 |
| ENSG00000C | UPRT | 6.26E-01 | 8.63E-01 | 9.77E-01 | -3.34E-02 | -1.02E+00 |
| ENSG000001 | TIMM50 | 6.26E-01 | 8.63E-01 | 1.02E+00 | 3.24E-02 | 1.02E+00 |
| ENSG00000C | ZNF839 | 6.26E-01 | 8.63E-01 | 1.03E+00 | 3.67E-02 | 1.03E+00 |
| ENSG00000C | KIF2A | 6.26E-01 | 8.63E-01 | 9.85E-01 | -2.19E-02 | -1.02E+00 |
| ENSG000001 | CAVIN4 | 6.26E-01 | ? | 8.79E-01 | -1.86E-01 | -1.14E+00 |
| ENSG000001 | EMILIN2 | 6.26E-01 | ? | 8.84E-01 | -1.77E-01 | -1.13E+00 |
| ENSG000001 | RSPH9 | 6.26E-01 | 8.63E-01 | 1.10E+00 | 1.35E-01 | 1.10E+00 |
| ENSG000002 | --- | 6.26E-01 | ? | 1.26E+00 | 3.31E-01 | 1.26E+00 |
| ENSG000001 | STON2 | 6.26E-01 | 8.63E-01 | 9.74E-01 | -3.79E-02 | -1.03E+00 |
| ENSG000002 | PFN1P4 | 6.26E-01 | ? | 1.30E+00 | 3.76E-01 | 1.30E+00 |
| ENSG00000C | THAP3 | 6.26E-01 | 8.64E-01 | 1.04E+00 | 5.25E-02 | 1.04E+00 |
| ENSG000001 | DAZAP2 | 6.26E-01 | 8.64E-01 | 1.02E+00 | 2.96E-02 | 1.02E+00 |
| ENSG000001 | FOXD4L1 | 6.27E-01 | ? | 8.86E-01 | -1.75E-01 | -1.13E+00 |
| ENSG000002 | --- | 6.27E-01 | ? | 8.99E-01 | -1.53E-01 | -1.11E+00 |
| ENSG000002 | --- | 6.27E-01 | ? | 1.26E+00 | 3.38E-01 | 1.26E+00 |
| ENSG000002 | KRT16P2 | 6.27E-01 | ? | 1.12E+00 | 1.59E-01 | 1.12E+00 |
| ENSG000001 | CHMP1A | 6.27E-01 | 8.64E-01 | 1.02E+00 | 3.22E-02 | 1.02E+00 |
| ENSG000001 | PARPBP | 6.27E-01 | 8.64E-01 | 1.04E+00 | 5.94E-02 | 1.04E+00 |
| ENSG000001 | RAVER1 | 6.27E-01 | 8.64E-01 | 1.04E+00 | 5.81E-02 | 1.04E+00 |
| ENSG000001 | BAIAP2L2 | 6.27E-01 | 8.64E-01 | 9.36E-01 | -9.61E-02 | -1.07E+00 |
| ENSG000001 | NDUFA13 | 6.27E-01 | 8.64E-01 | 1.02E+00 | 2.83E-02 | 1.02E+00 |
| ENSG000001 | GIMAP2 | 6.27E-01 | ? | 1.21E+00 | 2.74E-01 | 1.21E+00 |
| ENSG00000C | SEMA4G | 6.27E-01 | 8.64E-01 | 9.52E-01 | -7.09E-02 | -1.05E+00 |
| ENSG000001 | VWCE | 6.27E-01 | 8.64E-01 | 1.09E+00 | 1.24E-01 | 1.09E+00 |
| ENSG000001 | SC5D | 6.27E-01 | 8.64E-01 | 9.81E-01 | -2.69E-02 | -1.02E+00 |
| ENSG00000C | CADPS2 | 6.27E-01 | 8.64E-01 | 1.05E+00 | 7.36E-02 | 1.05E+00 |
| ENSG000002 | --- | 6.27E-01 | ? | 1.26E+00 | 3.39E-01 | 1.26E+00 |
| ENSG000002 | --- | 6.27E-01 | ? | 1.14E+00 | 1.95E-01 | 1.14E+00 |
| ENSG00000C | EHD3 | 6.27E-01 | 8.64E-01 | 1.02E+00 | 2.78E-02 | 1.02E+00 |
| ENSG00000C | SPEG | 6.28E-01 | 8.64E-01 | 1.04E+00 | 5.20E-02 | 1.04E+00 |
| ENSG000001 | YPEL2 | 6.28E-01 | 8.64E-01 | 1.03E+00 | 3.67E-02 | 1.03E+00 |
| ENSG000001 | --- | 6.28E-01 | ? | 1.20E+00 | 2.65E-01 | 1.20E+00 |
| ENSG000001 | TMEM106B | 6.28E-01 | 8.64E-01 | 9.76E-01 | -3.44E-02 | -1.02E+00 |
| ENSG000002 | ALMS1P1 | 6.28E-01 | ? | 1.18E+00 | 2.35E-01 | 1.18E+00 |
| ENSG000002 | EIF4A1P10 | 6.28E-01 | 8.64E-01 | 9.52E-01 | -7.12E-02 | -1.05E+00 |
| ENSG00000C | TOMM34 | 6.28E-01 | 8.64E-01 | 1.02E+00 | 3.21E-02 | 1.02E+00 |

| | | | | | | | |
|---|---|---|---|---|---|---|---|
| ENSG000001 | VKORC1 | 6.28E-01 | | 8.64E-01 | 1.02E+00 | 3.40E-02 | 1.02E+00 |
| ENSG000001 | VSIG1 | 6.28E-01 | | 8.64E-01 | 9.55E-01 | -6.57E-02 | -1.05E+00 |
| ENSG000002 | ST3GAL1-DT | 6.28E-01 | ? | | 1.15E+00 | 1.96E-01 | 1.15E+00 |
| ENSG000001 | TMLHE | 6.28E-01 | | 8.64E-01 | 1.03E+00 | 4.08E-02 | 1.03E+00 |
| ENSG00000C | PHTF2 | 6.28E-01 | | 8.64E-01 | 9.77E-01 | -3.35E-02 | -1.02E+00 |
| ENSG000002 | --- | 6.28E-01 | ? | | 9.13E-01 | -1.31E-01 | -1.10E+00 |
| ENSG000001 | CDS1 | 6.28E-01 | | 8.64E-01 | 1.01E+00 | 1.94E-02 | 1.01E+00 |
| ENSG000002 | --- | 6.28E-01 | | 8.64E-01 | 9.39E-01 | -9.15E-02 | -1.07E+00 |
| ENSG000001 | USP3 | 6.28E-01 | | 8.64E-01 | 9.81E-01 | -2.76E-02 | -1.02E+00 |
| ENSG000001 | CDNF | 6.28E-01 | ? | | 8.92E-01 | -1.65E-01 | -1.12E+00 |
| ENSG000001 | IQSEC1 | 6.28E-01 | | 8.64E-01 | 1.02E+00 | 2.94E-02 | 1.02E+00 |
| ENSG000001 | ZNF81 | 6.28E-01 | | 8.64E-01 | 9.78E-01 | -3.23E-02 | -1.02E+00 |
| ENSG000001 | HEATR5A | 6.28E-01 | | 8.64E-01 | 9.81E-01 | -2.80E-02 | -1.02E+00 |
| ENSG000001 | KRT23 | 6.28E-01 | ? | | 1.09E+00 | 1.25E-01 | 1.09E+00 |
| ENSG000002 | --- | 6.29E-01 | ? | | 8.80E-01 | -1.85E-01 | -1.14E+00 |
| ENSG000001 | PLAC9 | 6.29E-01 | ? | | 8.50E-01 | -2.35E-01 | -1.18E+00 |
| ENSG000002 | --- | 6.29E-01 | ? | | 7.72E-01 | -3.74E-01 | -1.30E+00 |
| ENSG000001 | HPS3 | 6.29E-01 | | 8.65E-01 | 1.02E+00 | 2.61E-02 | 1.02E+00 |
| ENSG00000C | GTF2IRD1 | 6.29E-01 | | 8.65E-01 | 1.02E+00 | 3.29E-02 | 1.02E+00 |
| ENSG000002 | LINC02362 | 6.29E-01 | ? | | 8.14E-01 | -2.96E-01 | -1.23E+00 |
| ENSG000002 | THTPA | 6.29E-01 | | 8.65E-01 | 1.03E+00 | 4.85E-02 | 1.03E+00 |
| ENSG00000C | MFAP3 | 6.29E-01 | | 8.65E-01 | 1.01E+00 | 2.13E-02 | 1.01E+00 |
| ENSG000002 | --- | 6.29E-01 | ? | | 8.11E-01 | -3.03E-01 | -1.23E+00 |
| ENSG000002 | THOC1-DT | 6.29E-01 | ? | | 8.83E-01 | -1.79E-01 | -1.13E+00 |
| ENSG000001 | SHBG | 6.29E-01 | ? | | 1.13E+00 | 1.81E-01 | 1.13E+00 |
| ENSG000002 | --- | 6.29E-01 | ? | | 1.26E+00 | 3.35E-01 | 1.26E+00 |
| ENSG000001 | CTDSP1 | 6.29E-01 | | 8.65E-01 | 1.03E+00 | 4.16E-02 | 1.03E+00 |
| ENSG000001 | KLF3 | 6.29E-01 | | 8.65E-01 | 9.82E-01 | -2.62E-02 | -1.02E+00 |
| ENSG000002 | GPSM3 | 6.29E-01 | | 8.65E-01 | 9.72E-01 | -4.07E-02 | -1.03E+00 |
| ENSG000001 | SNRNP48 | 6.29E-01 | | 8.65E-01 | 9.79E-01 | -3.02E-02 | -1.02E+00 |
| ENSG000002 | WDR11-DT | 6.29E-01 | ? | | 1.38E+00 | 4.66E-01 | 1.38E+00 |
| ENSG00000C | TTC19 | 6.30E-01 | | 8.65E-01 | 9.84E-01 | -2.37E-02 | -1.02E+00 |
| ENSG000001 | TAOK2 | 6.30E-01 | | 8.65E-01 | 1.02E+00 | 2.92E-02 | 1.02E+00 |
| ENSG000001 | TMEM169 | 6.30E-01 | ? | | 1.25E+00 | 3.18E-01 | 1.25E+00 |
| ENSG000001 | PDE4D | 6.30E-01 | | 8.65E-01 | 9.49E-01 | -7.58E-02 | -1.05E+00 |
| ENSG000002 | --- | 6.30E-01 | ? | | 8.07E-01 | -3.10E-01 | -1.24E+00 |
| ENSG000001 | --- | 6.30E-01 | ? | | 8.63E-01 | -2.12E-01 | -1.16E+00 |
| ENSG000001 | EPS8 | 6.30E-01 | | 8.65E-01 | 9.69E-01 | -4.50E-02 | -1.03E+00 |
| ENSG000002 | --- | 6.30E-01 | ? | | 1.19E+00 | 2.57E-01 | 1.19E+00 |
| ENSG000001 | ADAL | 6.30E-01 | | 8.65E-01 | 9.61E-01 | -5.70E-02 | -1.04E+00 |
| ENSG000002 | FAM193B-DT | 6.30E-01 | ? | | 1.27E+00 | 3.46E-01 | 1.27E+00 |
| ENSG000002 | GOLGA2P10 | 6.30E-01 | | 8.65E-01 | 1.22E+00 | 2.83E-01 | 1.22E+00 |
| ENSG000001 | TM6SF1 | 6.30E-01 | ? | | 8.48E-01 | -2.39E-01 | -1.18E+00 |
| ENSG000002 | GVINP1 | 6.30E-01 | ? | | 1.54E+00 | 6.22E-01 | 1.54E+00 |
| ENSG000001 | DNAJB4 | 6.30E-01 | | 8.65E-01 | 1.02E+00 | 2.95E-02 | 1.02E+00 |

| | | | | | | |
|---|---|---|---|---|---|---|
| ENSG00000C | WIZ | 6.30E-01 | 8.65E-01 | 1.02E+00 | 3.13E-02 | 1.02E+00 |
| ENSG000002 | MRPS18AP1 | 6.30E-01 | ? | 1.22E+00 | 2.82E-01 | 1.22E+00 |
| ENSG000002 | HMGB1P6 | 6.30E-01 | 8.65E-01 | 1.04E+00 | 5.82E-02 | 1.04E+00 |
| ENSG000001 | RMI1 | 6.30E-01 | 8.65E-01 | 9.71E-01 | -4.32E-02 | -1.03E+00 |
| ENSG000001 | HAT1 | 6.30E-01 | 8.65E-01 | 9.84E-01 | -2.28E-02 | -1.02E+00 |
| ENSG00000C | PPP5C | 6.30E-01 | 8.65E-01 | 1.02E+00 | 2.19E-02 | 1.02E+00 |
| ENSG000002 | CAHM | 6.30E-01 | ? | 8.37E-01 | -2.57E-01 | -1.20E+00 |
| ENSG000001 | TMEM192 | 6.30E-01 | 8.65E-01 | 1.02E+00 | 2.24E-02 | 1.02E+00 |
| ENSG000002 | --- | 6.30E-01 | 8.65E-01 | 9.63E-01 | -5.38E-02 | -1.04E+00 |
| ENSG000001 | ATP5MG | 6.31E-01 | 8.65E-01 | 1.02E+00 | 2.58E-02 | 1.02E+00 |
| ENSG000001 | COMMD10 | 6.31E-01 | 8.65E-01 | 9.77E-01 | -3.39E-02 | -1.02E+00 |
| ENSG000002 | --- | 6.31E-01 | ? | 8.86E-01 | -1.75E-01 | -1.13E+00 |
| ENSG000001 | NOP2 | 6.31E-01 | 8.65E-01 | 1.02E+00 | 2.92E-02 | 1.02E+00 |
| ENSG000001 | TOM1L1 | 6.31E-01 | 8.65E-01 | 9.83E-01 | -2.51E-02 | -1.02E+00 |
| ENSG000001 | ARL6IP5 | 6.31E-01 | 8.65E-01 | 9.86E-01 | -2.08E-02 | -1.01E+00 |
| ENSG000001 | DRD1 | 6.31E-01 | ? | 1.18E+00 | 2.39E-01 | 1.18E+00 |
| ENSG000001 | WDR74 | 6.31E-01 | 8.65E-01 | 1.02E+00 | 3.22E-02 | 1.02E+00 |
| ENSG000002 | --- | 6.31E-01 | 8.65E-01 | 9.59E-01 | -6.07E-02 | -1.04E+00 |
| ENSG000002 | LNCTAM34A | 6.31E-01 | 8.65E-01 | 1.07E+00 | 9.24E-02 | 1.07E+00 |
| ENSG000001 | TFF3 | 6.31E-01 | ? | 1.18E+00 | 2.34E-01 | 1.18E+00 |
| ENSG000001 | CCDC25 | 6.31E-01 | 8.65E-01 | 1.01E+00 | 2.06E-02 | 1.01E+00 |
| ENSG000002 | LGALS7 | 6.31E-01 | 8.65E-01 | 1.06E+00 | 7.82E-02 | 1.06E+00 |
| ENSG000001 | KRT14 | 6.31E-01 | 8.65E-01 | 1.03E+00 | 4.38E-02 | 1.03E+00 |
| ENSG000001 | FAM149B1 | 6.31E-01 | 8.65E-01 | 9.77E-01 | -3.29E-02 | -1.02E+00 |
| ENSG000001 | CARD14 | 6.31E-01 | 8.65E-01 | 1.08E+00 | 1.08E-01 | 1.08E+00 |
| ENSG000001 | HNF4A | 6.31E-01 | ? | 7.84E-01 | -3.51E-01 | -1.28E+00 |
| ENSG000001 | ARRDC2 | 6.31E-01 | 8.65E-01 | 9.67E-01 | -4.81E-02 | -1.03E+00 |
| ENSG000002 | --- | 6.31E-01 | ? | 7.57E-01 | -4.02E-01 | -1.32E+00 |
| ENSG000001 | TNFRSF11B | 6.31E-01 | ? | 1.13E+00 | 1.80E-01 | 1.13E+00 |
| ENSG000002 | --- | 6.32E-01 | ? | 8.24E-01 | -2.79E-01 | -1.21E+00 |
| ENSG000002 | --- | 6.32E-01 | ? | 1.14E+00 | 1.86E-01 | 1.14E+00 |
| ENSG000001 | EXOC4 | 6.32E-01 | 8.66E-01 | 1.01E+00 | 1.94E-02 | 1.01E+00 |
| ENSG000002 | BTG3-AS1 | 6.32E-01 | ? | 1.09E+00 | 1.26E-01 | 1.09E+00 |
| ENSG000001 | WDR12 | 6.32E-01 | 8.66E-01 | 9.82E-01 | -2.59E-02 | -1.02E+00 |
| ENSG00000C | AKAP8L | 6.32E-01 | 8.66E-01 | 1.01E+00 | 2.01E-02 | 1.01E+00 |
| ENSG000001 | CYP7B1 | 6.32E-01 | 8.66E-01 | 9.56E-01 | -6.47E-02 | -1.05E+00 |
| ENSG000001 | GDPD5 | 6.32E-01 | 8.66E-01 | 9.64E-01 | -5.23E-02 | -1.04E+00 |
| ENSG000001 | NUP153 | 6.32E-01 | 8.66E-01 | 1.02E+00 | 2.29E-02 | 1.02E+00 |
| ENSG000001 | CNRIP1 | 6.32E-01 | 8.66E-01 | 1.07E+00 | 1.01E-01 | 1.07E+00 |
| ENSG000001 | CAST | 6.32E-01 | 8.66E-01 | 1.02E+00 | 2.51E-02 | 1.02E+00 |
| ENSG000002 | --- | 6.32E-01 | ? | 1.23E+00 | 3.01E-01 | 1.23E+00 |
| ENSG000001 | TMEM17 | 6.32E-01 | 8.66E-01 | 9.62E-01 | -5.62E-02 | -1.04E+00 |
| ENSG000001 | PLXNA1 | 6.32E-01 | 8.66E-01 | 1.02E+00 | 2.99E-02 | 1.02E+00 |
| ENSG000001 | CCSER1 | 6.32E-01 | ? | 1.11E+00 | 1.52E-01 | 1.11E+00 |
| ENSG000001 | PDCL3 | 6.32E-01 | 8.66E-01 | 9.83E-01 | -2.48E-02 | -1.02E+00 |

| | | | | | | |
|---|---|---|---|---|---|---|
| ENSG000002 | NPM1P7 | 6.32E-01 | 8.66E-01 | 9.27E-01 | -1.09E-01 | -1.08E+00 |
| ENSG00000C | HAGH | 6.33E-01 | 8.66E-01 | 1.03E+00 | 3.70E-02 | 1.03E+00 |
| ENSG000002 | RPS15P4 | 6.33E-01 | ? | 1.15E+00 | 2.01E-01 | 1.15E+00 |
| ENSG000002 | --- | 6.33E-01 | ? | 1.12E+00 | 1.69E-01 | 1.12E+00 |
| ENSG000001 | DGKQ | 6.33E-01 | 8.66E-01 | 1.04E+00 | 5.19E-02 | 1.04E+00 |
| ENSG000002 | --- | 6.33E-01 | ? | 1.31E+00 | 3.86E-01 | 1.31E+00 |
| ENSG000002 | SNHG3 | 6.33E-01 | 8.66E-01 | 9.59E-01 | -6.10E-02 | -1.04E+00 |
| ENSG000001 | IL3RA | 6.33E-01 | ? | 8.67E-01 | -2.05E-01 | -1.15E+00 |
| ENSG000002 | --- | 6.33E-01 | ? | 1.10E+00 | 1.42E-01 | 1.10E+00 |
| ENSG00000C | ZFP64 | 6.33E-01 | 8.66E-01 | 1.01E+00 | 2.11E-02 | 1.01E+00 |
| ENSG000001 | SLC38A6 | 6.33E-01 | 8.66E-01 | 9.72E-01 | -4.14E-02 | -1.03E+00 |
| ENSG000001 | CTSV | 6.33E-01 | 8.66E-01 | 1.03E+00 | 3.86E-02 | 1.03E+00 |
| ENSG000001 | CHRNB1 | 6.33E-01 | 8.66E-01 | 1.04E+00 | 5.13E-02 | 1.04E+00 |
| ENSG00000C | COL11A1 | 6.33E-01 | 8.66E-01 | 9.32E-01 | -1.01E-01 | -1.07E+00 |
| ENSG000001 | SSH3 | 6.33E-01 | 8.66E-01 | 1.03E+00 | 4.04E-02 | 1.03E+00 |
| ENSG000001 | EMC1 | 6.33E-01 | 8.66E-01 | 9.87E-01 | -1.83E-02 | -1.01E+00 |
| ENSG00000C | NDUFAF7 | 6.33E-01 | 8.66E-01 | 9.78E-01 | -3.15E-02 | -1.02E+00 |
| ENSG000002 | GDNF-AS1 | 6.33E-01 | ? | 1.22E+00 | 2.82E-01 | 1.22E+00 |
| ENSG000001 | DNAJC6 | 6.33E-01 | 8.66E-01 | 9.68E-01 | -4.75E-02 | -1.03E+00 |
| ENSG000002 | ELF3-AS1 | 6.33E-01 | 8.66E-01 | 1.07E+00 | 9.44E-02 | 1.07E+00 |
| ENSG000001 | SLX1A | 6.33E-01 | 8.66E-01 | 9.40E-01 | -8.98E-02 | -1.06E+00 |
| ENSG000002 | BTG1-DT | 6.33E-01 | ? | 8.41E-01 | -2.51E-01 | -1.19E+00 |
| ENSG000002 | CEP43 | 6.33E-01 | 8.66E-01 | 1.03E+00 | 3.99E-02 | 1.03E+00 |
| ENSG000001 | TRPM1 | 6.33E-01 | ? | 1.23E+00 | 3.02E-01 | 1.23E+00 |
| ENSG000001 | UQCC2 | 6.34E-01 | 8.66E-01 | 1.03E+00 | 4.21E-02 | 1.03E+00 |
| ENSG000001 | NLRP3 | 6.34E-01 | 8.66E-01 | 9.40E-01 | -8.90E-02 | -1.06E+00 |
| ENSG000001 | PIKFYVE | 6.34E-01 | 8.66E-01 | 1.02E+00 | 2.78E-02 | 1.02E+00 |
| ENSG000002 | --- | 6.34E-01 | ? | 1.11E+00 | 1.50E-01 | 1.11E+00 |
| ENSG000001 | ARMH3 | 6.34E-01 | 8.66E-01 | 1.02E+00 | 2.32E-02 | 1.02E+00 |
| ENSG000002 | --- | 6.34E-01 | 8.66E-01 | 9.47E-01 | -7.87E-02 | -1.06E+00 |
| ENSG000001 | CARS2 | 6.34E-01 | 8.66E-01 | 1.02E+00 | 2.81E-02 | 1.02E+00 |
| ENSG000001 | ARL14EP | 6.34E-01 | 8.66E-01 | 1.02E+00 | 2.68E-02 | 1.02E+00 |
| ENSG000001 | PSMB6 | 6.34E-01 | 8.66E-01 | 9.77E-01 | -3.39E-02 | -1.02E+00 |
| ENSG000001 | H1-0 | 6.34E-01 | 8.66E-01 | 1.03E+00 | 3.91E-02 | 1.03E+00 |
| ENSG000002 | --- | 6.34E-01 | ? | 7.77E-01 | -3.64E-01 | -1.29E+00 |
| ENSG000001 | SASH1 | 6.34E-01 | 8.66E-01 | 1.02E+00 | 3.43E-02 | 1.02E+00 |
| ENSG000002 | --- | 6.34E-01 | ? | 1.20E+00 | 2.63E-01 | 1.20E+00 |
| ENSG000002 | RHOQ-AS1 | 6.35E-01 | ? | 8.27E-01 | -2.75E-01 | -1.21E+00 |
| ENSG000001 | ASPHD1 | 6.35E-01 | 8.67E-01 | 9.40E-01 | -8.89E-02 | -1.06E+00 |
| ENSG000002 | UBE2F-SCLY | 6.35E-01 | ? | 8.10E-01 | -3.03E-01 | -1.23E+00 |
| ENSG000002 | F8A2 | 6.35E-01 | 8.67E-01 | 9.34E-01 | -9.87E-02 | -1.07E+00 |
| ENSG000001 | MICU1 | 6.35E-01 | 8.67E-01 | 1.02E+00 | 2.98E-02 | 1.02E+00 |
| ENSG000002 | ZSCAN5A-AS | 6.35E-01 | ? | 1.26E+00 | 3.28E-01 | 1.26E+00 |
| ENSG000002 | --- | 6.35E-01 | ? | 7.87E-01 | -3.46E-01 | -1.27E+00 |
| ENSG000001 | GOLPH3 | 6.35E-01 | 8.67E-01 | 9.88E-01 | -1.70E-02 | -1.01E+00 |

| | | | | | | |
|---|---|---|---|---|---|---|
| ENSG000001 | SLC25A18 | 6.35E-01 | ? | 1.09E+00 | 1.22E-01 | 1.09E+00 |
| ENSG000001 | ZNF555 | 6.35E-01 | 8.67E-01 | 9.68E-01 | -4.64E-02 | -1.03E+00 |
| ENSG000002 | ZNF468 | 6.35E-01 | 8.67E-01 | 1.02E+00 | 3.27E-02 | 1.02E+00 |
| ENSG000001 | CMC1 | 6.35E-01 | 8.67E-01 | 9.73E-01 | -3.94E-02 | -1.03E+00 |
| ENSG00000C | LZTS1 | 6.35E-01 | ? | 1.18E+00 | 2.33E-01 | 1.18E+00 |
| ENSG000001 | CHPF | 6.35E-01 | 8.67E-01 | 1.05E+00 | 6.95E-02 | 1.05E+00 |
| ENSG000001 | SEC16A | 6.35E-01 | 8.67E-01 | 9.85E-01 | -2.20E-02 | -1.02E+00 |
| ENSG000002 | --- | 6.35E-01 | ? | 1.18E+00 | 2.44E-01 | 1.18E+00 |
| ENSG000002 | --- | 6.36E-01 | ? | 8.17E-01 | -2.91E-01 | -1.22E+00 |
| ENSG00000C | MRI1 | 6.36E-01 | 8.67E-01 | 1.03E+00 | 3.67E-02 | 1.03E+00 |
| ENSG000001 | SLC39A14 | 6.36E-01 | 8.67E-01 | 1.02E+00 | 2.92E-02 | 1.02E+00 |
| ENSG000002 | NBPF25P | 6.36E-01 | 8.67E-01 | 9.30E-01 | -1.05E-01 | -1.08E+00 |
| ENSG000001 | GRIN3B | 6.36E-01 | 8.67E-01 | 9.34E-01 | -9.83E-02 | -1.07E+00 |
| ENSG000001 | C14orf93 | 6.36E-01 | 8.67E-01 | 1.04E+00 | 5.90E-02 | 1.04E+00 |
| ENSG000002 | --- | 6.36E-01 | ? | 1.24E+00 | 3.11E-01 | 1.24E+00 |
| ENSG000001 | ZEB2 | 6.36E-01 | ? | 1.24E+00 | 3.14E-01 | 1.24E+00 |
| ENSG000002 | MYO1B-AS1 | 6.36E-01 | ? | 8.60E-01 | -2.18E-01 | -1.16E+00 |
| ENSG000001 | POLR2J3 | 6.36E-01 | 8.67E-01 | 1.03E+00 | 3.81E-02 | 1.03E+00 |
| ENSG000001 | HMBOX1 | 6.36E-01 | 8.67E-01 | 9.72E-01 | -4.06E-02 | -1.03E+00 |
| ENSG000001 | GAB3 | 6.36E-01 | ? | 9.16E-01 | -1.27E-01 | -1.09E+00 |
| ENSG000002 | ZNF300P1 | 6.36E-01 | ? | 8.09E-01 | -3.06E-01 | -1.24E+00 |
| ENSG000001 | TAPBPL | 6.36E-01 | 8.67E-01 | 1.05E+00 | 6.50E-02 | 1.05E+00 |
| ENSG000001 | PANK2 | 6.36E-01 | 8.67E-01 | 1.02E+00 | 2.23E-02 | 1.02E+00 |
| ENSG000002 | --- | 6.36E-01 | ? | 8.05E-01 | -3.13E-01 | -1.24E+00 |
| ENSG000001 | SOCS5 | 6.36E-01 | 8.67E-01 | 1.02E+00 | 3.46E-02 | 1.02E+00 |
| ENSG000001 | CNTNAP2 | 6.36E-01 | ? | 8.99E-01 | -1.54E-01 | -1.11E+00 |
| ENSG00000C | NUP37 | 6.36E-01 | 8.67E-01 | 1.02E+00 | 2.61E-02 | 1.02E+00 |
| ENSG000001 | ARHGAP42 | 6.36E-01 | 8.67E-01 | 9.69E-01 | -4.54E-02 | -1.03E+00 |
| ENSG000002 | ICMT-DT | 6.36E-01 | ? | 7.70E-01 | -3.77E-01 | -1.30E+00 |
| ENSG000002 | H4C5 | 6.36E-01 | ? | 1.14E+00 | 1.87E-01 | 1.14E+00 |
| ENSG000001 | TPGS1 | 6.37E-01 | 8.67E-01 | 9.48E-01 | -7.73E-02 | -1.06E+00 |
| ENSG000001 | MESD | 6.37E-01 | 8.67E-01 | 1.02E+00 | 2.24E-02 | 1.02E+00 |
| ENSG000001 | TBX19 | 6.37E-01 | 8.67E-01 | 1.06E+00 | 7.97E-02 | 1.06E+00 |
| ENSG00000C | CNGB1 | 6.37E-01 | 8.67E-01 | 1.04E+00 | 5.02E-02 | 1.04E+00 |
| ENSG00000C | CAMKK1 | 6.37E-01 | 8.67E-01 | 9.64E-01 | -5.22E-02 | -1.04E+00 |
| ENSG000001 | RABEPK | 6.37E-01 | 8.67E-01 | 1.02E+00 | 3.51E-02 | 1.02E+00 |
| ENSG000001 | TENT2 | 6.37E-01 | 8.67E-01 | 9.81E-01 | -2.75E-02 | -1.02E+00 |
| ENSG000002 | RPL7P22 | 6.37E-01 | ? | 9.19E-01 | -1.22E-01 | -1.09E+00 |
| ENSG000002 | ZNF8-ERVK3 | 6.37E-01 | 8.67E-01 | 1.06E+00 | 8.75E-02 | 1.06E+00 |
| ENSG000002 | POT1-AS1 | 6.37E-01 | ? | 9.19E-01 | -1.22E-01 | -1.09E+00 |
| ENSG000001 | RASA1 | 6.37E-01 | 8.67E-01 | 9.80E-01 | -2.89E-02 | -1.02E+00 |
| ENSG000002 | RFPL3S | 6.37E-01 | ? | 8.46E-01 | -2.42E-01 | -1.18E+00 |
| ENSG000001 | CUZD1 | 6.37E-01 | ? | 1.12E+00 | 1.57E-01 | 1.12E+00 |
| ENSG000001 | TEX15 | 6.37E-01 | ? | 1.12E+00 | 1.69E-01 | 1.12E+00 |
| ENSG000002 | --- | 6.37E-01 | ? | 1.14E+00 | 1.86E-01 | 1.14E+00 |

| | | | | | | |
|---|---|---|---|---|---|---|
| ENSG000001 | CHST6 | 6.37E-01 | ? | 9.12E-01 | -1.33E-01 | -1.10E+00 |
| ENSG000002 | --- | 6.37E-01 | 8.68E-01 | 9.31E-01 | -1.03E-01 | -1.07E+00 |
| ENSG000001 | DGKE | 6.37E-01 | 8.68E-01 | 1.03E+00 | 4.68E-02 | 1.03E+00 |
| ENSG000001 | HNRNPR | 6.37E-01 | 8.68E-01 | 9.80E-01 | -2.87E-02 | -1.02E+00 |
| ENSG000002 | --- | 6.37E-01 | 8.68E-01 | 9.56E-01 | -6.54E-02 | -1.05E+00 |
| ENSG000002 | IRF2-DT | 6.37E-01 | ? | 1.12E+00 | 1.65E-01 | 1.12E+00 |
| ENSG000001 | PCOLCE2 | 6.37E-01 | 8.68E-01 | 1.06E+00 | 8.88E-02 | 1.06E+00 |
| ENSG000002 | --- | 6.37E-01 | ? | 8.86E-01 | -1.74E-01 | -1.13E+00 |
| ENSG000001 | NUP43 | 6.38E-01 | 8.68E-01 | 9.84E-01 | -2.28E-02 | -1.02E+00 |
| ENSG000002 | ARIH2OS | 6.38E-01 | ? | 1.19E+00 | 2.52E-01 | 1.19E+00 |
| ENSG00000C | MRPS33 | 6.38E-01 | 8.68E-01 | 1.02E+00 | 2.65E-02 | 1.02E+00 |
| ENSG000001 | GNA13 | 6.38E-01 | 8.68E-01 | 1.01E+00 | 2.12E-02 | 1.01E+00 |
| ENSG000001 | FAM181B | 6.38E-01 | ? | 9.02E-01 | -1.50E-01 | -1.11E+00 |
| ENSG000001 | MFAP1 | 6.38E-01 | 8.68E-01 | 1.02E+00 | 2.81E-02 | 1.02E+00 |
| ENSG000001 | PTPRA | 6.38E-01 | 8.68E-01 | 9.85E-01 | -2.17E-02 | -1.02E+00 |
| ENSG00000C | TARBP1 | 6.38E-01 | 8.68E-01 | 1.02E+00 | 3.43E-02 | 1.02E+00 |
| ENSG000001 | TENT5C | 6.38E-01 | 8.68E-01 | 9.28E-01 | -1.08E-01 | -1.08E+00 |
| ENSG00000C | MAST3 | 6.38E-01 | 8.68E-01 | 9.72E-01 | -4.12E-02 | -1.03E+00 |
| ENSG000002 | ZNF578 | 6.38E-01 | 8.68E-01 | 9.76E-01 | -3.53E-02 | -1.02E+00 |
| ENSG000001 | SDHC | 6.38E-01 | 8.68E-01 | 9.80E-01 | -2.91E-02 | -1.02E+00 |
| ENSG000001 | PUS10 | 6.38E-01 | 8.68E-01 | 1.03E+00 | 4.53E-02 | 1.03E+00 |
| ENSG000002 | TUBGCP5 | 6.38E-01 | 8.68E-01 | 9.84E-01 | -2.37E-02 | -1.02E+00 |
| ENSG000002 | LIMS3 | 6.38E-01 | 8.68E-01 | 9.09E-01 | -1.38E-01 | -1.10E+00 |
| ENSG000001 | ALDH5A1 | 6.38E-01 | 8.68E-01 | 1.03E+00 | 3.79E-02 | 1.03E+00 |
| ENSG000001 | ASH2L | 6.38E-01 | 8.68E-01 | 1.01E+00 | 2.10E-02 | 1.01E+00 |
| ENSG000001 | CCDC120 | 6.38E-01 | 8.68E-01 | 9.74E-01 | -3.83E-02 | -1.03E+00 |
| ENSG000001 | SGSM1 | 6.38E-01 | ? | 7.70E-01 | -3.76E-01 | -1.30E+00 |
| ENSG000001 | APOBEC3H | 6.38E-01 | ? | 8.29E-01 | -2.71E-01 | -1.21E+00 |
| ENSG00000C | RAD51 | 6.38E-01 | 8.68E-01 | 1.06E+00 | 7.92E-02 | 1.06E+00 |
| ENSG000001 | HSPA12A | 6.39E-01 | 8.68E-01 | 1.03E+00 | 3.81E-02 | 1.03E+00 |
| ENSG00000C | CIRBP | 6.39E-01 | 8.68E-01 | 1.02E+00 | 3.45E-02 | 1.02E+00 |
| ENSG000002 | SVIL-AS1 | 6.39E-01 | 8.68E-01 | 1.02E+00 | 2.37E-02 | 1.02E+00 |
| ENSG000001 | LINC00839 | 6.39E-01 | 8.68E-01 | 1.02E+00 | 3.53E-02 | 1.02E+00 |
| ENSG000001 | UBOX5 | 6.39E-01 | 8.68E-01 | 9.66E-01 | -4.96E-02 | -1.03E+00 |
| ENSG000001 | ENTR1 | 6.39E-01 | 8.68E-01 | 9.82E-01 | -2.61E-02 | -1.02E+00 |
| ENSG000001 | NCSTN | 6.39E-01 | 8.68E-01 | 1.01E+00 | 1.86E-02 | 1.01E+00 |
| ENSG000002 | TSIX | 6.39E-01 | ? | 8.86E-01 | -1.74E-01 | -1.13E+00 |
| ENSG000002 | IER5L-AS1 | 6.39E-01 | ? | 1.24E+00 | 3.09E-01 | 1.24E+00 |
| ENSG000001 | FRMD6 | 6.39E-01 | 8.68E-01 | 1.01E+00 | 1.84E-02 | 1.01E+00 |
| ENSG000002 | CICP25 | 6.39E-01 | ? | 7.74E-01 | -3.71E-01 | -1.29E+00 |
| ENSG000002 | --- | 6.39E-01 | ? | 1.20E+00 | 2.68E-01 | 1.20E+00 |
| ENSG000002 | --- | 6.39E-01 | ? | 8.44E-01 | -2.45E-01 | -1.19E+00 |
| ENSG000001 | STK40 | 6.39E-01 | 8.68E-01 | 9.78E-01 | -3.28E-02 | -1.02E+00 |
| ENSG000001 | CLIP4 | 6.39E-01 | 8.68E-01 | 1.02E+00 | 2.63E-02 | 1.02E+00 |
| ENSG000002 | LINC02331 | 6.39E-01 | ? | 9.05E-01 | -1.44E-01 | -1.10E+00 |

| | | | | | | |
|---|---|---|---|---|---|---|
| ENSG000001 | ZNF385B | 6.40E-01 | ? | 8.27E-01 | -2.73E-01 | -1.21E+00 |
| ENSG000001 | AUP1 | 6.40E-01 | 8.68E-01 | 1.02E+00 | 2.17E-02 | 1.02E+00 |
| ENSG000002 | --- | 6.40E-01 | 8.68E-01 | 1.05E+00 | 7.49E-02 | 1.05E+00 |
| ENSG000002 | AKR7A2P1 | 6.40E-01 | ? | 1.21E+00 | 2.75E-01 | 1.21E+00 |
| ENSG000002 | C10orf88B | 6.40E-01 | 8.69E-01 | 9.43E-01 | -8.40E-02 | -1.06E+00 |
| ENSG000001 | SAMD11 | 6.40E-01 | ? | 1.21E+00 | 2.76E-01 | 1.21E+00 |
| ENSG000001 | DISP1 | 6.40E-01 | 8.69E-01 | 1.04E+00 | 5.05E-02 | 1.04E+00 |
| ENSG000002 | H4C12 | 6.40E-01 | ? | 8.60E-01 | -2.18E-01 | -1.16E+00 |
| ENSG000002 | ACBD3-AS1 | 6.40E-01 | ? | 7.67E-01 | -3.83E-01 | -1.30E+00 |
| ENSG000001 | KCNMB3 | 6.40E-01 | ? | 9.22E-01 | -1.17E-01 | -1.08E+00 |
| ENSG000002 | XRCC6P2 | 6.40E-01 | ? | 8.85E-01 | -1.77E-01 | -1.13E+00 |
| ENSG000001 | NIPBL | 6.40E-01 | 8.69E-01 | 9.75E-01 | -3.64E-02 | -1.03E+00 |
| ENSG000002 | MHENCR | 6.40E-01 | 8.69E-01 | 9.53E-01 | -6.91E-02 | -1.05E+00 |
| ENSG000001 | GALNS | 6.40E-01 | 8.69E-01 | 9.73E-01 | -3.94E-02 | -1.03E+00 |
| ENSG000001 | ADAM32 | 6.40E-01 | ? | 8.92E-01 | -1.64E-01 | -1.12E+00 |
| ENSG000001 | GARS1-DT | 6.40E-01 | 8.69E-01 | 9.64E-01 | -5.30E-02 | -1.04E+00 |
| ENSG000001 | EYA3 | 6.40E-01 | 8.69E-01 | 9.87E-01 | -1.89E-02 | -1.01E+00 |
| ENSG000002 | --- | 6.41E-01 | 8.69E-01 | 9.65E-01 | -5.14E-02 | -1.04E+00 |
| ENSG000002 | VPS9D1-AS1 | 6.41E-01 | 8.69E-01 | 1.03E+00 | 4.89E-02 | 1.03E+00 |
| ENSG00000C | PIGB | 6.41E-01 | 8.69E-01 | 1.02E+00 | 3.37E-02 | 1.02E+00 |
| ENSG000001 | BTG4 | 6.41E-01 | ? | 1.14E+00 | 1.89E-01 | 1.14E+00 |
| ENSG000001 | PAIP2 | 6.41E-01 | 8.69E-01 | 1.02E+00 | 2.49E-02 | 1.02E+00 |
| ENSG000001 | ZNF24 | 6.41E-01 | 8.69E-01 | 9.85E-01 | -2.21E-02 | -1.02E+00 |
| ENSG000002 | --- | 6.41E-01 | ? | 7.96E-01 | -3.29E-01 | -1.26E+00 |
| ENSG000001 | CFAP97 | 6.41E-01 | 8.69E-01 | 1.02E+00 | 2.91E-02 | 1.02E+00 |
| ENSG000002 | MALINC1 | 6.41E-01 | 8.69E-01 | 9.38E-01 | -9.30E-02 | -1.07E+00 |
| ENSG00000C | FTSJ1 | 6.41E-01 | 8.69E-01 | 9.88E-01 | -1.70E-02 | -1.01E+00 |
| ENSG000001 | NSA2 | 6.41E-01 | 8.69E-01 | 1.02E+00 | 2.44E-02 | 1.02E+00 |
| ENSG00000C | HSPA5 | 6.41E-01 | 8.69E-01 | 9.77E-01 | -3.31E-02 | -1.02E+00 |
| ENSG000002 | YY2 | 6.41E-01 | 8.69E-01 | 9.43E-01 | -8.54E-02 | -1.06E+00 |
| ENSG000001 | CHI3L1 | 6.41E-01 | ? | 1.14E+00 | 1.88E-01 | 1.14E+00 |
| ENSG000001 | ATXN7L2 | 6.41E-01 | 8.69E-01 | 9.56E-01 | -6.49E-02 | -1.05E+00 |
| ENSG000002 | MIR4458HG | 6.41E-01 | 8.69E-01 | 1.06E+00 | 7.83E-02 | 1.06E+00 |
| ENSG000001 | IL17D | 6.41E-01 | 8.69E-01 | 9.62E-01 | -5.54E-02 | -1.04E+00 |
| ENSG000001 | COL2A1 | 6.41E-01 | ? | 1.17E+00 | 2.25E-01 | 1.17E+00 |
| ENSG000001 | FHIT | 6.41E-01 | 8.69E-01 | 9.53E-01 | -6.96E-02 | -1.05E+00 |
| ENSG000002 | PCGF3-AS1 | 6.41E-01 | 8.69E-01 | 9.51E-01 | -7.28E-02 | -1.05E+00 |
| ENSG000002 | STAG3L4 | 6.42E-01 | 8.69E-01 | 9.71E-01 | -4.24E-02 | -1.03E+00 |
| ENSG000001 | ZNF7 | 6.42E-01 | 8.69E-01 | 1.02E+00 | 2.97E-02 | 1.02E+00 |
| ENSG000001 | PRPH2 | 6.42E-01 | ? | 8.09E-01 | -3.06E-01 | -1.24E+00 |
| ENSG000002 | ZBED8 | 6.42E-01 | 8.69E-01 | 1.04E+00 | 6.13E-02 | 1.04E+00 |
| ENSG000001 | SLC35F5 | 6.42E-01 | 8.69E-01 | 1.02E+00 | 2.74E-02 | 1.02E+00 |
| ENSG000001 | TOPBP1 | 6.42E-01 | 8.69E-01 | 9.81E-01 | -2.70E-02 | -1.02E+00 |
| ENSG000002 | --- | 6.42E-01 | ? | 1.20E+00 | 2.66E-01 | 1.20E+00 |
| ENSG000002 | PCDHGA9 | 6.42E-01 | ? | 1.09E+00 | 1.31E-01 | 1.09E+00 |

| | | | | | | |
|---|---|---|---|---|---|---|
| ENSG000001 | ESPN | 6.42E-01 | 8.69E-01 | 9.70E-01 | -4.43E-02 | -1.03E+00 |
| ENSG00000C | LTBP1 | 6.42E-01 | 8.69E-01 | 9.37E-01 | -9.36E-02 | -1.07E+00 |
| ENSG00000C | CCDC28A | 6.42E-01 | 8.69E-01 | 1.03E+00 | 4.82E-02 | 1.03E+00 |
| ENSG000001 | ADAM15 | 6.43E-01 | 8.69E-01 | 1.03E+00 | 4.90E-02 | 1.03E+00 |
| ENSG000002 | RBM38-AS1 | 6.43E-01 | ? | 8.54E-01 | -2.28E-01 | -1.17E+00 |
| ENSG000002 | --- | 6.43E-01 | ? | 1.18E+00 | 2.44E-01 | 1.18E+00 |
| ENSG000002 | --- | 6.43E-01 | 8.69E-01 | 9.49E-01 | -7.56E-02 | -1.05E+00 |
| ENSG000002 | PAPOLA-DT | 6.43E-01 | ? | 1.14E+00 | 1.85E-01 | 1.14E+00 |
| ENSG000001 | HTATSF1 | 6.43E-01 | 8.69E-01 | 9.84E-01 | -2.35E-02 | -1.02E+00 |
| ENSG000001 | MTO1 | 6.43E-01 | 8.69E-01 | 1.02E+00 | 2.90E-02 | 1.02E+00 |
| ENSG000001 | RPIA | 6.43E-01 | 8.69E-01 | 1.03E+00 | 3.94E-02 | 1.03E+00 |
| ENSG00000C | CAMK2B | 6.43E-01 | ? | 8.67E-01 | -2.07E-01 | -1.15E+00 |
| ENSG000001 | CHD5 | 6.43E-01 | ? | 1.16E+00 | 2.17E-01 | 1.16E+00 |
| ENSG000001 | BPGM | 6.43E-01 | 8.69E-01 | 9.79E-01 | -3.00E-02 | -1.02E+00 |
| ENSG000002 | TMEM88B | 6.43E-01 | ? | 8.23E-01 | -2.80E-01 | -1.21E+00 |
| ENSG000001 | KRT9 | 6.43E-01 | 8.69E-01 | 1.05E+00 | 7.49E-02 | 1.05E+00 |
| ENSG000002 | TWSG1-DT | 6.43E-01 | ? | 1.20E+00 | 2.61E-01 | 1.20E+00 |
| ENSG000002 | --- | 6.43E-01 | ? | 8.60E-01 | -2.18E-01 | -1.16E+00 |
| ENSG000001 | LRRN2 | 6.43E-01 | ? | 1.11E+00 | 1.49E-01 | 1.11E+00 |
| ENSG000001 | KRT10-AS1 | 6.43E-01 | 8.70E-01 | 1.03E+00 | 4.46E-02 | 1.03E+00 |
| ENSG000002 | FAUP1 | 6.43E-01 | ? | 1.20E+00 | 2.64E-01 | 1.20E+00 |
| ENSG000001 | THAP12 | 6.43E-01 | 8.70E-01 | 9.85E-01 | -2.23E-02 | -1.02E+00 |
| ENSG000001 | PAX8 | 6.43E-01 | 8.70E-01 | 1.06E+00 | 7.87E-02 | 1.06E+00 |
| ENSG000002 | --- | 6.43E-01 | 8.70E-01 | 9.64E-01 | -5.28E-02 | -1.04E+00 |
| ENSG000001 | LRG1 | 6.43E-01 | 8.70E-01 | 9.57E-01 | -6.34E-02 | -1.04E+00 |
| ENSG000001 | MUC20 | 6.43E-01 | 8.70E-01 | 1.07E+00 | 9.88E-02 | 1.07E+00 |
| ENSG00000C | DNAJC25 | 6.43E-01 | 8.70E-01 | 9.74E-01 | -3.86E-02 | -1.03E+00 |
| ENSG000001 | FURIN | 6.44E-01 | 8.70E-01 | 1.02E+00 | 3.49E-02 | 1.02E+00 |
| ENSG000001 | BAAT | 6.44E-01 | ? | 1.18E+00 | 2.34E-01 | 1.18E+00 |
| ENSG000001 | IDE | 6.44E-01 | 8.70E-01 | 9.81E-01 | -2.73E-02 | -1.02E+00 |
| ENSG000001 | FAM13A | 6.44E-01 | 8.70E-01 | 9.72E-01 | -4.03E-02 | -1.03E+00 |
| ENSG000001 | ELP5 | 6.44E-01 | 8.70E-01 | 1.02E+00 | 3.12E-02 | 1.02E+00 |
| ENSG000001 | SETD5 | 6.44E-01 | 8.70E-01 | 9.84E-01 | -2.33E-02 | -1.02E+00 |
| ENSG000001 | EGFR | 6.44E-01 | 8.70E-01 | 9.80E-01 | -2.87E-02 | -1.02E+00 |
| ENSG000002 | SLC44A3-AS1 | 6.44E-01 | 8.70E-01 | 9.57E-01 | -6.28E-02 | -1.04E+00 |
| ENSG000002 | ERICH6-AS1 | 6.44E-01 | ? | 1.08E+00 | 1.08E-01 | 1.08E+00 |
| ENSG000001 | SCOC-AS1 | 6.44E-01 | ? | 1.08E+00 | 1.15E-01 | 1.08E+00 |
| ENSG00000C | SEMA3F | 6.44E-01 | 8.70E-01 | 1.03E+00 | 4.16E-02 | 1.03E+00 |
| ENSG000002 | CLN8-AS1 | 6.44E-01 | 8.70E-01 | 1.04E+00 | 5.77E-02 | 1.04E+00 |
| ENSG000002 | METTL14-DT | 6.44E-01 | ? | 1.14E+00 | 1.87E-01 | 1.14E+00 |
| ENSG000002 | SNHG25 | 6.44E-01 | ? | 8.29E-01 | -2.71E-01 | -1.21E+00 |
| ENSG000001 | RILPL1 | 6.44E-01 | 8.70E-01 | 9.81E-01 | -2.75E-02 | -1.02E+00 |
| ENSG000001 | WRAP53 | 6.44E-01 | 8.70E-01 | 9.74E-01 | -3.78E-02 | -1.03E+00 |
| ENSG000001 | FOXP4 | 6.44E-01 | 8.70E-01 | 9.84E-01 | -2.34E-02 | -1.02E+00 |
| ENSG000001 | TET3 | 6.44E-01 | 8.70E-01 | 9.79E-01 | -3.11E-02 | -1.02E+00 |

| | | | | | | |
|---|---|---|---|---|---|---|
| ENSG000001 | USP24 | 6.44E-01 | 8.70E-01 | 9.82E-01 | -2.69E-02 | -1.02E+00 |
| ENSG000001 | RBM28 | 6.44E-01 | 8.70E-01 | 1.02E+00 | 2.23E-02 | 1.02E+00 |
| ENSG000001 | PLCG2 | 6.45E-01 | 8.70E-01 | 1.05E+00 | 7.64E-02 | 1.05E+00 |
| ENSG000002 | --- | 6.45E-01 | ? | 1.10E+00 | 1.42E-01 | 1.10E+00 |
| ENSG000002 | SSBP3P1 | 6.45E-01 | ? | 8.61E-01 | -2.16E-01 | -1.16E+00 |
| ENSG000002 | ARL2-SNX15 | 6.45E-01 | 8.70E-01 | 1.23E+00 | 2.97E-01 | 1.23E+00 |
| ENSG000002 | --- | 6.45E-01 | 8.70E-01 | 1.02E+00 | 2.48E-02 | 1.02E+00 |
| ENSG000002 | --- | 6.45E-01 | ? | 1.13E+00 | 1.82E-01 | 1.13E+00 |
| ENSG000002 | ZNF295-AS1 | 6.45E-01 | ? | 8.34E-01 | -2.62E-01 | -1.20E+00 |
| ENSG000001 | CCDC40 | 6.45E-01 | 8.70E-01 | 1.06E+00 | 8.47E-02 | 1.06E+00 |
| ENSG000001 | MED13L | 6.45E-01 | 8.70E-01 | 9.80E-01 | -2.87E-02 | -1.02E+00 |
| ENSG000001 | ARHGDIB | 6.45E-01 | 8.70E-01 | 9.75E-01 | -3.67E-02 | -1.03E+00 |
| ENSG000001 | SNRPC | 6.45E-01 | 8.70E-01 | 1.02E+00 | 2.23E-02 | 1.02E+00 |
| ENSG000002 | --- | 6.45E-01 | 8.70E-01 | 9.19E-01 | -1.23E-01 | -1.09E+00 |
| ENSG00000C | USP2 | 6.45E-01 | 8.70E-01 | 1.04E+00 | 6.02E-02 | 1.04E+00 |
| ENSG00000C | FH | 6.45E-01 | 8.70E-01 | 1.02E+00 | 2.72E-02 | 1.02E+00 |
| ENSG000001 | CRYZ | 6.45E-01 | 8.70E-01 | 9.89E-01 | -1.61E-02 | -1.01E+00 |
| ENSG000002 | ICAM4-AS1 | 6.46E-01 | ? | 8.25E-01 | -2.77E-01 | -1.21E+00 |
| ENSG000001 | HTATIP2 | 6.46E-01 | 8.70E-01 | 1.02E+00 | 2.20E-02 | 1.02E+00 |
| ENSG000001 | SPIN2A | 6.46E-01 | ? | 8.80E-01 | -1.85E-01 | -1.14E+00 |
| ENSG000001 | MRPS16 | 6.46E-01 | 8.71E-01 | 1.02E+00 | 2.63E-02 | 1.02E+00 |
| ENSG000001 | KLF16 | 6.46E-01 | 8.71E-01 | 1.04E+00 | 5.11E-02 | 1.04E+00 |
| ENSG000001 | MAGOHB | 6.46E-01 | 8.71E-01 | 1.03E+00 | 3.60E-02 | 1.03E+00 |
| ENSG000001 | SOD1 | 6.46E-01 | 8.71E-01 | 9.83E-01 | -2.44E-02 | -1.02E+00 |
| ENSG000001 | EPHB1 | 6.46E-01 | ? | 1.12E+00 | 1.67E-01 | 1.12E+00 |
| ENSG000001 | SFXN5 | 6.46E-01 | 8.71E-01 | 1.03E+00 | 3.84E-02 | 1.03E+00 |
| ENSG000001 | KIF3A | 6.46E-01 | 8.71E-01 | 9.76E-01 | -3.52E-02 | -1.02E+00 |
| ENSG000001 | ABHD17AP1 | 6.46E-01 | ? | 1.12E+00 | 1.58E-01 | 1.12E+00 |
| ENSG000002 | MTCO2P12 | 6.46E-01 | ? | 1.12E+00 | 1.66E-01 | 1.12E+00 |
| ENSG00000C | TNPO3 | 6.46E-01 | 8.71E-01 | 9.88E-01 | -1.78E-02 | -1.01E+00 |
| ENSG000002 | --- | 6.46E-01 | ? | 1.24E+00 | 3.15E-01 | 1.24E+00 |
| ENSG000002 | APP-DT | 6.46E-01 | ? | 1.15E+00 | 2.07E-01 | 1.15E+00 |
| ENSG000001 | SLC25A19 | 6.46E-01 | 8.71E-01 | 1.03E+00 | 3.75E-02 | 1.03E+00 |
| ENSG000001 | PNPLA3 | 6.46E-01 | 8.71E-01 | 9.67E-01 | -4.80E-02 | -1.03E+00 |
| ENSG000001 | GP1BA | 6.47E-01 | 8.71E-01 | 9.44E-01 | -8.32E-02 | -1.06E+00 |
| ENSG000001 | ADH6 | 6.47E-01 | ? | 9.01E-01 | -1.50E-01 | -1.11E+00 |
| ENSG000002 | --- | 6.47E-01 | ? | 1.22E+00 | 2.82E-01 | 1.22E+00 |
| ENSG000002 | PDXP | 6.47E-01 | 8.71E-01 | 8.39E-01 | -2.54E-01 | -1.19E+00 |
| ENSG000002 | LINC00200 | 6.47E-01 | ? | 8.79E-01 | -1.86E-01 | -1.14E+00 |
| ENSG000002 | --- | 6.47E-01 | ? | 1.15E+00 | 2.02E-01 | 1.15E+00 |
| ENSG000002 | --- | 6.47E-01 | ? | 8.48E-01 | -2.38E-01 | -1.18E+00 |
| ENSG000002 | --- | 6.47E-01 | ? | 1.31E+00 | 3.86E-01 | 1.31E+00 |
| ENSG000002 | ZNF8 | 6.47E-01 | 8.71E-01 | 1.02E+00 | 3.25E-02 | 1.02E+00 |
| ENSG000002 | --- | 6.47E-01 | ? | 1.30E+00 | 3.77E-01 | 1.30E+00 |
| ENSG000001 | ASB7 | 6.47E-01 | 8.71E-01 | 9.80E-01 | -2.91E-02 | -1.02E+00 |

| | | | | | | |
|---|---|---|---|---|---|---|
| ENSG000001 | TMEM100 | 6.47E-01 | ? | 1.14E+00 | 1.90E-01 | 1.14E+00 |
| ENSG000001 | MRPL52 | 6.47E-01 | 8.71E-01 | 1.02E+00 | 3.18E-02 | 1.02E+00 |
| ENSG000002 | --- | 6.47E-01 | ? | 1.18E+00 | 2.38E-01 | 1.18E+00 |
| ENSG000001 | IFT81 | 6.47E-01 | 8.71E-01 | 9.77E-01 | -3.43E-02 | -1.02E+00 |
| ENSG000002 | --- | 6.47E-01 | 8.71E-01 | 9.22E-01 | -1.17E-01 | -1.08E+00 |
| ENSG000002 | --- | 6.47E-01 | ? | 1.09E+00 | 1.27E-01 | 1.09E+00 |
| ENSG000001 | CCDC150 | 6.47E-01 | ? | 9.16E-01 | -1.26E-01 | -1.09E+00 |
| ENSG000001 | CD58 | 6.47E-01 | 8.72E-01 | 9.79E-01 | -3.12E-02 | -1.02E+00 |
| ENSG000001 | ALS2CL | 6.48E-01 | 8.72E-01 | 9.69E-01 | -4.50E-02 | -1.03E+00 |
| ENSG000001 | KHDRBS3 | 6.48E-01 | 8.72E-01 | 1.04E+00 | 5.11E-02 | 1.04E+00 |
| ENSG000002 | TIMM23 | 6.48E-01 | 8.72E-01 | 1.01E+00 | 1.79E-02 | 1.01E+00 |
| ENSG000001 | ETV6 | 6.48E-01 | 8.72E-01 | 1.02E+00 | 2.31E-02 | 1.02E+00 |
| ENSG000002 | --- | 6.48E-01 | ? | 8.93E-01 | -1.64E-01 | -1.12E+00 |
| ENSG000001 | B3GAT2 | 6.48E-01 | 8.72E-01 | 9.33E-01 | -1.00E-01 | -1.07E+00 |
| ENSG000001 | SLFN5 | 6.48E-01 | 8.72E-01 | 1.02E+00 | 2.76E-02 | 1.02E+00 |
| ENSG000001 | C15orf39 | 6.48E-01 | 8.72E-01 | 1.02E+00 | 3.53E-02 | 1.02E+00 |
| ENSG000001 | PFAS | 6.48E-01 | 8.72E-01 | 1.02E+00 | 3.31E-02 | 1.02E+00 |
| ENSG000002 | MSANTD5 | 6.48E-01 | ? | 1.13E+00 | 1.77E-01 | 1.13E+00 |
| ENSG000001 | CATSPERE | 6.48E-01 | 8.72E-01 | 1.08E+00 | 1.17E-01 | 1.08E+00 |
| ENSG000001 | ONECUT2 | 6.48E-01 | ? | 1.10E+00 | 1.38E-01 | 1.10E+00 |
| ENSG000001 | RABGGTB | 6.48E-01 | 8.72E-01 | 9.78E-01 | -3.16E-02 | -1.02E+00 |
| ENSG000002 | --- | 6.48E-01 | ? | 8.05E-01 | -3.13E-01 | -1.24E+00 |
| ENSG000001 | LCN2 | 6.48E-01 | 8.72E-01 | 1.09E+00 | 1.29E-01 | 1.09E+00 |
| ENSG000001 | ZBED4 | 6.48E-01 | 8.72E-01 | 9.87E-01 | -1.83E-02 | -1.01E+00 |
| ENSG000002 | TMEM191A | 6.49E-01 | 8.72E-01 | 9.22E-01 | -1.17E-01 | -1.08E+00 |
| ENSG000001 | AKAP7 | 6.49E-01 | 8.72E-01 | 9.49E-01 | -7.58E-02 | -1.05E+00 |
| ENSG00000C | STEEP1 | 6.49E-01 | 8.72E-01 | 1.02E+00 | 2.98E-02 | 1.02E+00 |
| ENSG000002 | --- | 6.49E-01 | ? | 8.25E-01 | -2.77E-01 | -1.21E+00 |
| ENSG000001 | CCDC178 | 6.49E-01 | ? | 8.39E-01 | -2.54E-01 | -1.19E+00 |
| ENSG000002 | SAP25 | 6.49E-01 | ? | 9.17E-01 | -1.26E-01 | -1.09E+00 |
| ENSG000001 | DUSP5P1 | 6.49E-01 | ? | 1.20E+00 | 2.62E-01 | 1.20E+00 |
| ENSG000001 | ADM | 6.49E-01 | 8.72E-01 | 9.71E-01 | -4.27E-02 | -1.03E+00 |
| ENSG00000C | STAG3 | 6.49E-01 | 8.72E-01 | 9.46E-01 | -8.06E-02 | -1.06E+00 |
| ENSG000002 | HMGN3-AS1 | 6.49E-01 | 8.72E-01 | 9.45E-01 | -8.24E-02 | -1.06E+00 |
| ENSG000002 | TICAM2 | 6.49E-01 | ? | 1.10E+00 | 1.39E-01 | 1.10E+00 |
| ENSG000001 | HMGN2P46 | 6.49E-01 | ? | 1.31E+00 | 3.88E-01 | 1.31E+00 |
| ENSG000001 | IFITM2 | 6.49E-01 | 8.72E-01 | 9.78E-01 | -3.26E-02 | -1.02E+00 |
| ENSG000001 | CALB1 | 6.49E-01 | ? | 1.19E+00 | 2.49E-01 | 1.19E+00 |
| ENSG000001 | COL6A2 | 6.49E-01 | 8.72E-01 | 1.09E+00 | 1.28E-01 | 1.09E+00 |
| ENSG000001 | RASGRP3 | 6.49E-01 | ? | 1.09E+00 | 1.26E-01 | 1.09E+00 |
| ENSG000001 | ANKRD35 | 6.49E-01 | ? | 1.12E+00 | 1.58E-01 | 1.12E+00 |
| ENSG000002 | BMS1P20 | 6.49E-01 | 8.72E-01 | 9.64E-01 | -5.29E-02 | -1.04E+00 |
| ENSG000001 | VEGFB | 6.49E-01 | 8.72E-01 | 1.03E+00 | 4.52E-02 | 1.03E+00 |
| ENSG000002 | --- | 6.49E-01 | ? | 1.17E+00 | 2.32E-01 | 1.17E+00 |
| ENSG000001 | PCNX2 | 6.49E-01 | 8.72E-01 | 1.02E+00 | 2.42E-02 | 1.02E+00 |

| | | | | | | |
|---|---|---|---|---|---|---|
| ENSG000002 | NUP50-DT | 6.49E-01 | 8.72E-01 | 1.04E+00 | 5.48E-02 | 1.04E+00 |
| ENSG000001 | STAT5B | 6.49E-01 | 8.72E-01 | 1.02E+00 | 2.45E-02 | 1.02E+00 |
| ENSG000002 | --- | 6.49E-01 | ? | 8.49E-01 | -2.36E-01 | -1.18E+00 |
| ENSG000001 | HLCS | 6.49E-01 | 8.72E-01 | 1.02E+00 | 3.01E-02 | 1.02E+00 |
| ENSG000001 | PHF24 | 6.50E-01 | ? | 8.99E-01 | -1.54E-01 | -1.11E+00 |
| ENSG000001 | CAV1 | 6.50E-01 | 8.72E-01 | 1.02E+00 | 2.52E-02 | 1.02E+00 |
| ENSG000001 | DCLK2 | 6.50E-01 | ? | 1.14E+00 | 1.88E-01 | 1.14E+00 |
| ENSG00000C | TPR | 6.50E-01 | 8.72E-01 | 9.81E-01 | -2.82E-02 | -1.02E+00 |
| ENSG000001 | NCOA5 | 6.50E-01 | 8.72E-01 | 1.01E+00 | 2.00E-02 | 1.01E+00 |
| ENSG000001 | LRP10 | 6.50E-01 | 8.72E-01 | 1.02E+00 | 3.14E-02 | 1.02E+00 |
| ENSG000001 | INTS2 | 6.50E-01 | 8.72E-01 | 9.82E-01 | -2.64E-02 | -1.02E+00 |
| ENSG000002 | --- | 6.50E-01 | ? | 1.29E+00 | 3.66E-01 | 1.29E+00 |
| ENSG000001 | HNRNPA2B1 | 6.50E-01 | 8.72E-01 | 9.86E-01 | -1.97E-02 | -1.01E+00 |
| ENSG000001 | YIPF2 | 6.50E-01 | 8.72E-01 | 9.73E-01 | -3.97E-02 | -1.03E+00 |
| ENSG000002 | --- | 6.50E-01 | ? | 8.06E-01 | -3.12E-01 | -1.24E+00 |
| ENSG000001 | SUN1 | 6.50E-01 | 8.72E-01 | 1.01E+00 | 1.64E-02 | 1.01E+00 |
| ENSG000002 | VWA8-AS1 | 6.50E-01 | ? | 8.40E-01 | -2.52E-01 | -1.19E+00 |
| ENSG000001 | DNAJC5 | 6.50E-01 | 8.72E-01 | 1.02E+00 | 3.10E-02 | 1.02E+00 |
| ENSG000002 | ELFN1 | 6.50E-01 | ? | 1.09E+00 | 1.20E-01 | 1.09E+00 |
| ENSG000002 | PPME1 | 6.50E-01 | 8.72E-01 | 1.01E+00 | 2.01E-02 | 1.01E+00 |
| ENSG000001 | WSCD1 | 6.50E-01 | ? | 9.18E-01 | -1.23E-01 | -1.09E+00 |
| ENSG000001 | BRD8 | 6.50E-01 | 8.72E-01 | 1.02E+00 | 3.00E-02 | 1.02E+00 |
| ENSG00000C | PHACTR3 | 6.50E-01 | 8.72E-01 | 1.04E+00 | 6.17E-02 | 1.04E+00 |
| ENSG000002 | --- | 6.50E-01 | 8.72E-01 | 1.07E+00 | 9.83E-02 | 1.07E+00 |
| ENSG000001 | OR5P2 | 6.50E-01 | ? | 7.92E-01 | -3.36E-01 | -1.26E+00 |
| ENSG000001 | TUBGCP2 | 6.51E-01 | 8.72E-01 | 9.84E-01 | -2.28E-02 | -1.02E+00 |
| ENSG000002 | LINC01376 | 6.51E-01 | ? | 8.48E-01 | -2.38E-01 | -1.18E+00 |
| ENSG00000C | POLD3 | 6.51E-01 | 8.72E-01 | 1.03E+00 | 4.14E-02 | 1.03E+00 |
| ENSG000001 | AKR1C3 | 6.51E-01 | 8.72E-01 | 9.74E-01 | -3.77E-02 | -1.03E+00 |
| ENSG000002 | --- | 6.51E-01 | 8.72E-01 | 7.90E-01 | -3.39E-01 | -1.27E+00 |
| ENSG000002 | CRYGS | 6.51E-01 | 8.72E-01 | 9.20E-01 | -1.20E-01 | -1.09E+00 |
| ENSG000001 | EMC2 | 6.51E-01 | 8.72E-01 | 1.02E+00 | 2.62E-02 | 1.02E+00 |
| ENSG000002 | INHBA-AS1 | 6.51E-01 | ? | 1.24E+00 | 3.15E-01 | 1.24E+00 |
| ENSG000001 | MIR22HG | 6.51E-01 | 8.72E-01 | 1.03E+00 | 3.69E-02 | 1.03E+00 |
| ENSG000001 | ISG20L2 | 6.51E-01 | 8.73E-01 | 1.01E+00 | 1.93E-02 | 1.01E+00 |
| ENSG00000C | SEC61A1 | 6.51E-01 | 8.73E-01 | 1.01E+00 | 1.82E-02 | 1.01E+00 |
| ENSG000001 | LPIN1 | 6.51E-01 | 8.73E-01 | 9.80E-01 | -2.99E-02 | -1.02E+00 |
| ENSG000002 | PSMA6P1 | 6.51E-01 | ? | 1.19E+00 | 2.50E-01 | 1.19E+00 |
| ENSG000001 | PRR15 | 6.51E-01 | ? | 1.11E+00 | 1.49E-01 | 1.11E+00 |
| ENSG000001 | ANXA10 | 6.51E-01 | ? | 7.93E-01 | -3.34E-01 | -1.26E+00 |
| ENSG000002 | --- | 6.51E-01 | ? | 1.20E+00 | 2.66E-01 | 1.20E+00 |
| ENSG000001 | WDHD1 | 6.51E-01 | 8.73E-01 | 9.73E-01 | -3.99E-02 | -1.03E+00 |
| ENSG000002 | DIRC3-AS1 | 6.51E-01 | ? | 1.17E+00 | 2.23E-01 | 1.17E+00 |
| ENSG000002 | RMRP | 6.51E-01 | ? | 1.18E+00 | 2.42E-01 | 1.18E+00 |
| ENSG00000C | ZIC2 | 6.52E-01 | ? | 1.21E+00 | 2.74E-01 | 1.21E+00 |

| | | | | | | |
|---|---|---|---|---|---|---|
| ENSG00000C | BIRC3 | 6.52E-01 | 8.73E-01 | 1.05E+00 | 6.91E-02 | 1.05E+00 |
| ENSG000001 | ADAMTS14 | 6.52E-01 | ? | 8.15E-01 | -2.95E-01 | -1.23E+00 |
| ENSG00000C | CTTN | 6.52E-01 | 8.73E-01 | 9.91E-01 | -1.37E-02 | -1.01E+00 |
| ENSG000002 | AIDAP2 | 6.52E-01 | ? | 9.02E-01 | -1.48E-01 | -1.11E+00 |
| ENSG000001 | KPNA3 | 6.52E-01 | 8.73E-01 | 9.86E-01 | -2.08E-02 | -1.01E+00 |
| ENSG000001 | IQSEC2 | 6.52E-01 | 8.73E-01 | 1.03E+00 | 3.73E-02 | 1.03E+00 |
| ENSG000002 | --- | 6.52E-01 | 8.73E-01 | 9.38E-01 | -9.18E-02 | -1.07E+00 |
| ENSG000002 | --- | 6.52E-01 | ? | 1.17E+00 | 2.21E-01 | 1.17E+00 |
| ENSG000002 | --- | 6.52E-01 | ? | 8.88E-01 | -1.71E-01 | -1.13E+00 |
| ENSG000002 | HEXIM2-AS1 | 6.52E-01 | ? | 1.10E+00 | 1.43E-01 | 1.10E+00 |
| ENSG000001 | HMGB2 | 6.52E-01 | 8.73E-01 | 9.65E-01 | -5.16E-02 | -1.04E+00 |
| ENSG000002 | LINC00882 | 6.52E-01 | ? | 8.88E-01 | -1.71E-01 | -1.13E+00 |
| ENSG000001 | BBS7 | 6.52E-01 | 8.73E-01 | 9.79E-01 | -3.10E-02 | -1.02E+00 |
| ENSG000001 | ACSF3 | 6.52E-01 | 8.73E-01 | 1.02E+00 | 3.04E-02 | 1.02E+00 |
| ENSG000002 | --- | 6.52E-01 | ? | 7.75E-01 | -3.67E-01 | -1.29E+00 |
| ENSG000002 | MAILR | 6.52E-01 | ? | 8.75E-01 | -1.92E-01 | -1.14E+00 |
| ENSG000001 | DPH7 | 6.52E-01 | 8.73E-01 | 1.02E+00 | 2.97E-02 | 1.02E+00 |
| ENSG000001 | BBS12 | 6.52E-01 | 8.73E-01 | 9.75E-01 | -3.72E-02 | -1.03E+00 |
| ENSG000002 | --- | 6.52E-01 | ? | 1.29E+00 | 3.72E-01 | 1.29E+00 |
| ENSG000001 | NCCRP1 | 6.53E-01 | 8.73E-01 | 9.35E-01 | -9.77E-02 | -1.07E+00 |
| ENSG000001 | ARHGAP26 | 6.53E-01 | 8.74E-01 | 9.78E-01 | -3.22E-02 | -1.02E+00 |
| ENSG000001 | MARCHF10 | 6.53E-01 | ? | 8.61E-01 | -2.16E-01 | -1.16E+00 |
| ENSG000002 | --- | 6.53E-01 | ? | 8.97E-01 | -1.56E-01 | -1.11E+00 |
| ENSG000001 | GRM6 | 6.53E-01 | ? | 1.14E+00 | 1.89E-01 | 1.14E+00 |
| ENSG00000C | TIPIN | 6.53E-01 | 8.74E-01 | 1.04E+00 | 5.58E-02 | 1.04E+00 |
| ENSG000002 | ALG11 | 6.53E-01 | 8.74E-01 | 1.01E+00 | 1.86E-02 | 1.01E+00 |
| ENSG000001 | ZNF684 | 6.53E-01 | 8.74E-01 | 9.39E-01 | -9.15E-02 | -1.07E+00 |
| ENSG000001 | HRNR | 6.53E-01 | 8.74E-01 | 9.29E-01 | -1.07E-01 | -1.08E+00 |
| ENSG000001 | NDOR1 | 6.53E-01 | 8.74E-01 | 9.61E-01 | -5.74E-02 | -1.04E+00 |
| ENSG000001 | ZNG1B | 6.53E-01 | 8.74E-01 | 9.76E-01 | -3.48E-02 | -1.02E+00 |
| ENSG000001 | KLHL21 | 6.53E-01 | 8.74E-01 | 1.03E+00 | 3.81E-02 | 1.03E+00 |
| ENSG000002 | --- | 6.53E-01 | ? | 8.83E-01 | -1.80E-01 | -1.13E+00 |
| ENSG000002 | --- | 6.53E-01 | 8.74E-01 | 9.78E-01 | -3.17E-02 | -1.02E+00 |
| ENSG000001 | MYRF | 6.54E-01 | 8.74E-01 | 9.39E-01 | -9.15E-02 | -1.07E+00 |
| ENSG000001 | AFTPH | 6.54E-01 | 8.74E-01 | 9.85E-01 | -2.12E-02 | -1.01E+00 |
| ENSG000001 | ATP6V1E1 | 6.54E-01 | 8.74E-01 | 9.89E-01 | -1.63E-02 | -1.01E+00 |
| ENSG000001 | IL17RC | 6.54E-01 | 8.74E-01 | 1.04E+00 | 5.19E-02 | 1.04E+00 |
| ENSG000001 | ZNF334 | 6.54E-01 | 8.74E-01 | 9.60E-01 | -5.90E-02 | -1.04E+00 |
| ENSG000001 | NRK | 6.54E-01 | 8.74E-01 | 1.07E+00 | 9.52E-02 | 1.07E+00 |
| ENSG000002 | --- | 6.54E-01 | ? | 8.75E-01 | -1.93E-01 | -1.14E+00 |
| ENSG000002 | --- | 6.54E-01 | 8.74E-01 | 1.05E+00 | 6.67E-02 | 1.05E+00 |
| ENSG000001 | AGBL3 | 6.54E-01 | ? | 1.08E+00 | 1.11E-01 | 1.08E+00 |
| ENSG000001 | RNFT1 | 6.54E-01 | 8.74E-01 | 9.81E-01 | -2.78E-02 | -1.02E+00 |
| ENSG00000C | WAPL | 6.54E-01 | 8.74E-01 | 9.86E-01 | -2.05E-02 | -1.01E+00 |
| ENSG000002 | ARHGAP11B | 6.54E-01 | 8.74E-01 | 9.10E-01 | -1.36E-01 | -1.10E+00 |

| | | | | | | |
|---|---|---|---|---|---|---|
| ENSG000001 | ADK | 6.54E-01 | 8.74E-01 | 9.86E-01 | -1.98E-02 | -1.01E+00 |
| ENSG000001 | FSCN3 | 6.54E-01 | ? | 8.45E-01 | -2.43E-01 | -1.18E+00 |
| ENSG000001 | KAT2B | 6.54E-01 | 8.74E-01 | 9.75E-01 | -3.67E-02 | -1.03E+00 |
| ENSG000001 | MIEN1 | 6.54E-01 | 8.74E-01 | 9.78E-01 | -3.17E-02 | -1.02E+00 |
| ENSG000001 | SEMA3E | 6.54E-01 | 8.74E-01 | 1.05E+00 | 7.69E-02 | 1.05E+00 |
| ENSG000002 | --- | 6.54E-01 | ? | 8.16E-01 | -2.94E-01 | -1.23E+00 |
| ENSG000002 | GPX1 | 6.54E-01 | 8.74E-01 | 1.03E+00 | 3.95E-02 | 1.03E+00 |
| ENSG000001 | FAM104B | 6.54E-01 | 8.74E-01 | 1.02E+00 | 3.12E-02 | 1.02E+00 |
| ENSG000001 | TNFRSF14 | 6.55E-01 | 8.74E-01 | 1.05E+00 | 6.71E-02 | 1.05E+00 |
| ENSG000001 | SNIP1 | 6.55E-01 | 8.74E-01 | 9.86E-01 | -2.07E-02 | -1.01E+00 |
| ENSG000001 | C9orf131 | 6.55E-01 | ? | 1.25E+00 | 3.19E-01 | 1.25E+00 |
| ENSG000001 | DENND4B | 6.55E-01 | 8.74E-01 | 1.02E+00 | 3.54E-02 | 1.02E+00 |
| ENSG00000C | TBC1D22A | 6.55E-01 | 8.74E-01 | 9.78E-01 | -3.24E-02 | -1.02E+00 |
| ENSG000001 | SEPTIN9 | 6.55E-01 | 8.74E-01 | 9.77E-01 | -3.34E-02 | -1.02E+00 |
| ENSG000002 | MMP25-AS1 | 6.55E-01 | 8.74E-01 | 9.53E-01 | -6.99E-02 | -1.05E+00 |
| ENSG000002 | --- | 6.55E-01 | 8.74E-01 | 1.09E+00 | 1.25E-01 | 1.09E+00 |
| ENSG000001 | BSCL2 | 6.55E-01 | 8.74E-01 | 1.05E+00 | 6.86E-02 | 1.05E+00 |
| ENSG000001 | U2SURP | 6.55E-01 | 8.74E-01 | 9.82E-01 | -2.62E-02 | -1.02E+00 |
| ENSG00000C | SMARCA2 | 6.55E-01 | 8.74E-01 | 9.85E-01 | -2.22E-02 | -1.02E+00 |
| ENSG000002 | --- | 6.55E-01 | ? | 8.31E-01 | -2.67E-01 | -1.20E+00 |
| ENSG000002 | MAGEL2 | 6.55E-01 | ? | 7.48E-01 | -4.19E-01 | -1.34E+00 |
| ENSG000001 | SLC5A1 | 6.55E-01 | ? | 8.19E-01 | -2.88E-01 | -1.22E+00 |
| ENSG000001 | NAT8L | 6.55E-01 | ? | 1.11E+00 | 1.44E-01 | 1.11E+00 |
| ENSG000001 | C4orf19 | 6.55E-01 | 8.74E-01 | 9.42E-01 | -8.62E-02 | -1.06E+00 |
| ENSG000002 | --- | 6.55E-01 | 8.74E-01 | 1.07E+00 | 9.25E-02 | 1.07E+00 |
| ENSG000002 | --- | 6.55E-01 | ? | 1.17E+00 | 2.28E-01 | 1.17E+00 |
| ENSG000002 | --- | 6.55E-01 | ? | 8.55E-01 | -2.27E-01 | -1.17E+00 |
| ENSG000001 | KCNS1 | 6.55E-01 | ? | 1.10E+00 | 1.36E-01 | 1.10E+00 |
| ENSG00000C | ST6GALNAC1 | 6.55E-01 | 8.74E-01 | 1.03E+00 | 4.81E-02 | 1.03E+00 |
| ENSG000001 | TERF1 | 6.55E-01 | 8.74E-01 | 9.83E-01 | -2.49E-02 | -1.02E+00 |
| ENSG000001 | MFSD10 | 6.55E-01 | 8.74E-01 | 1.06E+00 | 8.00E-02 | 1.06E+00 |
| ENSG00000C | PTBP1 | 6.56E-01 | 8.74E-01 | 9.84E-01 | -2.27E-02 | -1.02E+00 |
| ENSG000001 | POC5 | 6.56E-01 | 8.74E-01 | 9.73E-01 | -3.99E-02 | -1.03E+00 |
| ENSG000001 | PPAN | 6.56E-01 | 8.74E-01 | 9.82E-01 | -2.65E-02 | -1.02E+00 |
| ENSG000002 | GHET1 | 6.56E-01 | 8.74E-01 | 1.07E+00 | 9.80E-02 | 1.07E+00 |
| ENSG000001 | IQGAP1 | 6.56E-01 | 8.74E-01 | 9.84E-01 | -2.40E-02 | -1.02E+00 |
| ENSG000001 | IMMP1L | 6.56E-01 | 8.74E-01 | 9.69E-01 | -4.48E-02 | -1.03E+00 |
| ENSG000001 | TRAPPC12 | 6.56E-01 | 8.74E-01 | 9.83E-01 | -2.46E-02 | -1.02E+00 |
| ENSG000002 | ZKSCAN8P1 | 6.56E-01 | ? | 9.17E-01 | -1.24E-01 | -1.09E+00 |
| ENSG000001 | METTL21A | 6.56E-01 | 8.74E-01 | 1.03E+00 | 4.57E-02 | 1.03E+00 |
| ENSG000001 | GORASP1 | 6.56E-01 | 8.74E-01 | 1.02E+00 | 2.34E-02 | 1.02E+00 |
| ENSG000001 | EXOC3L1 | 6.56E-01 | ? | 7.72E-01 | -3.74E-01 | -1.30E+00 |
| ENSG000001 | P3H3 | 6.56E-01 | 8.74E-01 | 1.04E+00 | 5.05E-02 | 1.04E+00 |
| ENSG000001 | ECHDC2 | 6.56E-01 | 8.74E-01 | 1.04E+00 | 5.12E-02 | 1.04E+00 |
| ENSG000001 | AVEN | 6.56E-01 | 8.74E-01 | 9.82E-01 | -2.65E-02 | -1.02E+00 |

| | | | | | | |
|---|---|---|---|---|---|---|
| ENSG00000C | RAB7A | 6.56E-01 | 8.74E-01 | 9.83E-01 | -2.45E-02 | -1.02E+00 |
| ENSG000001 | REP15 | 6.56E-01 | 8.74E-01 | 1.06E+00 | 8.59E-02 | 1.06E+00 |
| ENSG000001 | ELMOD3 | 6.56E-01 | 8.74E-01 | 1.02E+00 | 2.63E-02 | 1.02E+00 |
| ENSG000002 | --- | 6.57E-01 | ? | 8.33E-01 | -2.64E-01 | -1.20E+00 |
| ENSG000002 | --- | 6.57E-01 | 8.74E-01 | 1.13E+00 | 1.73E-01 | 1.13E+00 |
| ENSG000002 | --- | 6.57E-01 | ? | 8.24E-01 | -2.80E-01 | -1.21E+00 |
| ENSG000001 | EIF4A2 | 6.57E-01 | 8.74E-01 | 9.78E-01 | -3.19E-02 | -1.02E+00 |
| ENSG000002 | --- | 6.57E-01 | ? | 8.88E-01 | -1.72E-01 | -1.13E+00 |
| ENSG000002 | --- | 6.57E-01 | ? | 8.87E-01 | -1.73E-01 | -1.13E+00 |
| ENSG000002 | MED4-AS1 | 6.57E-01 | ? | 1.16E+00 | 2.10E-01 | 1.16E+00 |
| ENSG000001 | SAV1 | 6.57E-01 | 8.75E-01 | 9.79E-01 | -3.09E-02 | -1.02E+00 |
| ENSG000001 | XPO7 | 6.57E-01 | 8.75E-01 | 1.01E+00 | 1.47E-02 | 1.01E+00 |
| ENSG000001 | SNRPN | 6.57E-01 | 8.75E-01 | 9.82E-01 | -2.66E-02 | -1.02E+00 |
| ENSG000001 | MANF | 6.57E-01 | 8.75E-01 | 1.02E+00 | 2.72E-02 | 1.02E+00 |
| ENSG000002 | --- | 6.57E-01 | ? | 1.28E+00 | 3.58E-01 | 1.28E+00 |
| ENSG000002 | --- | 6.57E-01 | ? | 7.43E-01 | -4.28E-01 | -1.35E+00 |
| ENSG000002 | --- | 6.57E-01 | 8.75E-01 | 1.08E+00 | 1.10E-01 | 1.08E+00 |
| ENSG000001 | BCAR3 | 6.57E-01 | 8.75E-01 | 1.02E+00 | 3.25E-02 | 1.02E+00 |
| ENSG000001 | ASPSCR1 | 6.57E-01 | 8.75E-01 | 1.04E+00 | 5.01E-02 | 1.04E+00 |
| ENSG000001 | PRDX4 | 6.57E-01 | 8.75E-01 | 1.02E+00 | 2.83E-02 | 1.02E+00 |
| ENSG000001 | C1orf147 | 6.58E-01 | ? | 1.12E+00 | 1.66E-01 | 1.12E+00 |
| ENSG000002 | IGSF22-AS1 | 6.58E-01 | ? | 8.22E-01 | -2.84E-01 | -1.22E+00 |
| ENSG000002 | --- | 6.58E-01 | 8.75E-01 | 9.46E-01 | -8.01E-02 | -1.06E+00 |
| ENSG000002 | C21orf62-AS | 6.58E-01 | ? | 1.09E+00 | 1.27E-01 | 1.09E+00 |
| ENSG000001 | USP32P1 | 6.58E-01 | ? | 1.34E+00 | 4.19E-01 | 1.34E+00 |
| ENSG000002 | --- | 6.58E-01 | ? | 1.10E+00 | 1.43E-01 | 1.10E+00 |
| ENSG000001 | MEGF6 | 6.58E-01 | 8.75E-01 | 9.50E-01 | -7.45E-02 | -1.05E+00 |
| ENSG000001 | MYO5A | 6.58E-01 | 8.75E-01 | 9.81E-01 | -2.78E-02 | -1.02E+00 |
| ENSG000002 | ZNF611 | 6.58E-01 | 8.75E-01 | 1.03E+00 | 4.74E-02 | 1.03E+00 |
| ENSG000001 | SPATA2L | 6.58E-01 | 8.75E-01 | 1.06E+00 | 8.51E-02 | 1.06E+00 |
| ENSG000002 | PJVK | 6.58E-01 | ? | 9.30E-01 | -1.05E-01 | -1.08E+00 |
| ENSG000001 | ZNF232 | 6.58E-01 | 8.75E-01 | 1.04E+00 | 6.22E-02 | 1.04E+00 |
| ENSG000002 | PPP4R3B | 6.58E-01 | 8.75E-01 | 1.01E+00 | 1.90E-02 | 1.01E+00 |
| ENSG000001 | DDX56 | 6.58E-01 | 8.75E-01 | 1.02E+00 | 2.92E-02 | 1.02E+00 |
| ENSG000001 | DDX39A | 6.58E-01 | 8.75E-01 | 1.03E+00 | 3.70E-02 | 1.03E+00 |
| ENSG000001 | PGP | 6.58E-01 | 8.75E-01 | 9.76E-01 | -3.55E-02 | -1.02E+00 |
| ENSG000002 | --- | 6.58E-01 | ? | 1.28E+00 | 3.54E-01 | 1.28E+00 |
| ENSG000002 | TMSB15B | 6.59E-01 | 8.75E-01 | 9.37E-01 | -9.41E-02 | -1.07E+00 |
| ENSG00000C | SPRTN | 6.59E-01 | 8.75E-01 | 9.81E-01 | -2.81E-02 | -1.02E+00 |
| ENSG000002 | --- | 6.59E-01 | ? | 1.21E+00 | 2.76E-01 | 1.21E+00 |
| ENSG000002 | --- | 6.59E-01 | ? | 8.97E-01 | -1.58E-01 | -1.12E+00 |
| ENSG000002 | RIMBP3B | 6.59E-01 | ? | 1.31E+00 | 3.94E-01 | 1.31E+00 |
| ENSG000002 | --- | 6.59E-01 | 8.75E-01 | 1.07E+00 | 9.19E-02 | 1.07E+00 |
| ENSG000002 | --- | 6.59E-01 | ? | 7.88E-01 | -3.44E-01 | -1.27E+00 |
| ENSG000002 | PCDHGA12 | 6.59E-01 | 8.75E-01 | 9.60E-01 | -5.95E-02 | -1.04E+00 |

| | | | | | | |
|---|---|---|---|---|---|---|
| ENSG000002 | STARD7-AS1 | 6.59E-01 | 8.75E-01 | 9.67E-01 | -4.86E-02 | -1.03E+00 |
| ENSG000002 | --- | 6.59E-01 | ? | 1.12E+00 | 1.65E-01 | 1.12E+00 |
| ENSG000001 | STS | 6.59E-01 | 8.75E-01 | 9.83E-01 | -2.45E-02 | -1.02E+00 |
| ENSG000002 | --- | 6.59E-01 | ? | 1.14E+00 | 1.84E-01 | 1.14E+00 |
| ENSG000001 | FDPS | 6.59E-01 | 8.75E-01 | 1.02E+00 | 2.99E-02 | 1.02E+00 |
| ENSG000002 | INSL3 | 6.59E-01 | ? | 1.10E+00 | 1.32E-01 | 1.10E+00 |
| ENSG000001 | CDH5 | 6.59E-01 | 8.75E-01 | 1.06E+00 | 8.96E-02 | 1.06E+00 |
| ENSG000001 | ARL14 | 6.59E-01 | 8.75E-01 | 1.04E+00 | 5.87E-02 | 1.04E+00 |
| ENSG000002 | HMGB1P24 | 6.59E-01 | ? | 1.32E+00 | 3.98E-01 | 1.32E+00 |
| ENSG000002 | KPNA2P3 | 6.59E-01 | ? | 1.22E+00 | 2.93E-01 | 1.22E+00 |
| ENSG000002 | --- | 6.59E-01 | ? | 9.34E-01 | -9.77E-02 | -1.07E+00 |
| ENSG000001 | ZNF394 | 6.60E-01 | 8.76E-01 | 9.83E-01 | -2.48E-02 | -1.02E+00 |
| ENSG000001 | TRIM38 | 6.60E-01 | 8.76E-01 | 9.83E-01 | -2.46E-02 | -1.02E+00 |
| ENSG000002 | LINC02532 | 6.60E-01 | ? | 1.26E+00 | 3.31E-01 | 1.26E+00 |
| ENSG000001 | SNX6 | 6.60E-01 | 8.76E-01 | 9.88E-01 | -1.75E-02 | -1.01E+00 |
| ENSG000002 | GAPDHP60 | 6.60E-01 | ? | 7.98E-01 | -3.26E-01 | -1.25E+00 |
| ENSG000001 | CYP4X1 | 6.60E-01 | ? | 1.11E+00 | 1.45E-01 | 1.11E+00 |
| ENSG000001 | TUBAL3 | 6.60E-01 | ? | 1.10E+00 | 1.36E-01 | 1.10E+00 |
| ENSG00000C | IFNGR1 | 6.60E-01 | 8.76E-01 | 9.87E-01 | -1.94E-02 | -1.01E+00 |
| ENSG00000C | PTPN23 | 6.60E-01 | 8.76E-01 | 9.82E-01 | -2.63E-02 | -1.02E+00 |
| ENSG00000C | MAVS | 6.60E-01 | 8.76E-01 | 9.91E-01 | -1.35E-02 | -1.01E+00 |
| ENSG000002 | --- | 6.60E-01 | ? | 8.85E-01 | -1.76E-01 | -1.13E+00 |
| ENSG000001 | DAD1 | 6.60E-01 | 8.76E-01 | 9.87E-01 | -1.96E-02 | -1.01E+00 |
| ENSG000001 | C10orf95 | 6.60E-01 | 8.76E-01 | 1.11E+00 | 1.54E-01 | 1.11E+00 |
| ENSG000001 | RSAD2 | 6.60E-01 | ? | 1.19E+00 | 2.55E-01 | 1.19E+00 |
| ENSG00000C | TG | 6.60E-01 | 8.76E-01 | 1.05E+00 | 6.86E-02 | 1.05E+00 |
| ENSG000002 | --- | 6.60E-01 | ? | 9.10E-01 | -1.36E-01 | -1.10E+00 |
| ENSG000001 | ALDH1L2 | 6.60E-01 | 8.76E-01 | 9.65E-01 | -5.10E-02 | -1.04E+00 |
| ENSG000001 | SOBP | 6.60E-01 | 8.76E-01 | 1.05E+00 | 7.03E-02 | 1.05E+00 |
| ENSG000001 | C1orf35 | 6.60E-01 | 8.76E-01 | 1.02E+00 | 3.29E-02 | 1.02E+00 |
| ENSG000001 | TRIB3 | 6.61E-01 | 8.76E-01 | 9.70E-01 | -4.39E-02 | -1.03E+00 |
| ENSG000002 | AFAP1-AS1 | 6.61E-01 | 8.76E-01 | 9.22E-01 | -1.17E-01 | -1.08E+00 |
| ENSG000001 | SYT8 | 6.61E-01 | ? | 9.20E-01 | -1.21E-01 | -1.09E+00 |
| ENSG000001 | ZNF619 | 6.61E-01 | 8.76E-01 | 9.73E-01 | -3.94E-02 | -1.03E+00 |
| ENSG000001 | HYOU1 | 6.61E-01 | 8.76E-01 | 9.88E-01 | -1.79E-02 | -1.01E+00 |
| ENSG000001 | OCRL | 6.61E-01 | 8.76E-01 | 1.01E+00 | 2.04E-02 | 1.01E+00 |
| ENSG000001 | NUMA1 | 6.61E-01 | 8.76E-01 | 1.01E+00 | 1.86E-02 | 1.01E+00 |
| ENSG000002 | --- | 6.61E-01 | ? | 9.00E-01 | -1.52E-01 | -1.11E+00 |
| ENSG000002 | RNF14P3 | 6.61E-01 | ? | 8.16E-01 | -2.93E-01 | -1.23E+00 |
| ENSG00000C | PIAS2 | 6.61E-01 | 8.76E-01 | 9.84E-01 | -2.39E-02 | -1.02E+00 |
| ENSG000001 | MGME1 | 6.61E-01 | 8.76E-01 | 1.02E+00 | 2.39E-02 | 1.02E+00 |
| ENSG000001 | BIK | 6.61E-01 | 8.76E-01 | 1.05E+00 | 6.44E-02 | 1.05E+00 |
| ENSG00000C | TULP3 | 6.61E-01 | 8.76E-01 | 9.81E-01 | -2.78E-02 | -1.02E+00 |
| ENSG00000C | FHL1 | 6.61E-01 | 8.76E-01 | 1.02E+00 | 2.60E-02 | 1.02E+00 |
| ENSG000002 | --- | 6.61E-01 | ? | 8.92E-01 | -1.66E-01 | -1.12E+00 |

| | | | | | | |
|---|---|---|---|---|---|---|
| ENSG000001 | CCL24 | 6.61E-01 | ? | 8.16E-01 | -2.93E-01 | -1.23E+00 |
| ENSG000001 | NMNAT1 | 6.62E-01 | 8.77E-01 | 9.84E-01 | -2.34E-02 | -1.02E+00 |
| ENSG000002 | --- | 6.62E-01 | ? | 8.62E-01 | -2.14E-01 | -1.16E+00 |
| ENSG000001 | CEP162 | 6.62E-01 | 8.77E-01 | 9.63E-01 | -5.39E-02 | -1.04E+00 |
| ENSG000002 | CTXND1 | 6.62E-01 | ? | 8.17E-01 | -2.91E-01 | -1.22E+00 |
| ENSG000002 | LINC01451 | 6.62E-01 | 8.77E-01 | 9.26E-01 | -1.10E-01 | -1.08E+00 |
| ENSG000001 | DCAF17 | 6.62E-01 | 8.77E-01 | 9.83E-01 | -2.41E-02 | -1.02E+00 |
| ENSG000001 | NINJ2-AS1 | 6.62E-01 | 8.77E-01 | 9.46E-01 | -8.03E-02 | -1.06E+00 |
| ENSG000001 | CWF19L2 | 6.62E-01 | 8.77E-01 | 1.02E+00 | 2.65E-02 | 1.02E+00 |
| ENSG000001 | ZNF69 | 6.62E-01 | 8.77E-01 | 9.65E-01 | -5.15E-02 | -1.04E+00 |
| ENSG000001 | RNF111 | 6.62E-01 | 8.77E-01 | 9.85E-01 | -2.23E-02 | -1.02E+00 |
| ENSG000001 | ARL8B | 6.62E-01 | 8.77E-01 | 1.02E+00 | 2.22E-02 | 1.02E+00 |
| ENSG00000C | TRIT1 | 6.62E-01 | 8.77E-01 | 1.02E+00 | 2.46E-02 | 1.02E+00 |
| ENSG000001 | ABR | 6.62E-01 | 8.77E-01 | 9.88E-01 | -1.80E-02 | -1.01E+00 |
| ENSG000002 | --- | 6.62E-01 | ? | 9.17E-01 | -1.26E-01 | -1.09E+00 |
| ENSG000001 | MVB12A | 6.62E-01 | 8.77E-01 | 1.04E+00 | 5.27E-02 | 1.04E+00 |
| ENSG000001 | LINC00313 | 6.62E-01 | 8.77E-01 | 1.06E+00 | 8.44E-02 | 1.06E+00 |
| ENSG000001 | C18orf21 | 6.62E-01 | 8.77E-01 | 9.78E-01 | -3.27E-02 | -1.02E+00 |
| ENSG00000C | DEF6 | 6.63E-01 | 8.77E-01 | 1.03E+00 | 3.78E-02 | 1.03E+00 |
| ENSG000002 | --- | 6.63E-01 | 8.77E-01 | 1.07E+00 | 9.35E-02 | 1.07E+00 |
| ENSG000001 | APOBEC2 | 6.63E-01 | ? | 1.14E+00 | 1.83E-01 | 1.14E+00 |
| ENSG000001 | STT3B | 6.63E-01 | 8.77E-01 | 9.87E-01 | -1.84E-02 | -1.01E+00 |
| ENSG000001 | CHTOP | 6.63E-01 | 8.77E-01 | 9.85E-01 | -2.24E-02 | -1.02E+00 |
| ENSG000001 | SGSH | 6.63E-01 | 8.77E-01 | 1.03E+00 | 4.74E-02 | 1.03E+00 |
| ENSG000002 | LINC02615 | 6.63E-01 | 8.77E-01 | 1.05E+00 | 7.42E-02 | 1.05E+00 |
| ENSG000001 | THAP10 | 6.63E-01 | 8.77E-01 | 1.03E+00 | 4.59E-02 | 1.03E+00 |
| ENSG000001 | IGFBP5 | 6.63E-01 | 8.77E-01 | 1.09E+00 | 1.22E-01 | 1.09E+00 |
| ENSG000001 | CREG1 | 6.63E-01 | 8.77E-01 | 1.02E+00 | 3.15E-02 | 1.02E+00 |
| ENSG000001 | CARD6 | 6.63E-01 | 8.77E-01 | 9.85E-01 | -2.15E-02 | -1.02E+00 |
| ENSG000001 | SNX18 | 6.63E-01 | 8.77E-01 | 1.01E+00 | 1.95E-02 | 1.01E+00 |
| ENSG000002 | RTCA-AS1 | 6.63E-01 | 8.77E-01 | 1.05E+00 | 6.68E-02 | 1.05E+00 |
| ENSG000001 | HEATR3 | 6.63E-01 | 8.77E-01 | 1.02E+00 | 2.59E-02 | 1.02E+00 |
| ENSG000001 | MANEA | 6.63E-01 | 8.77E-01 | 9.79E-01 | -2.99E-02 | -1.02E+00 |
| ENSG000001 | ZNF283 | 6.63E-01 | 8.77E-01 | 1.03E+00 | 3.93E-02 | 1.03E+00 |
| ENSG000001 | SAR1B | 6.63E-01 | 8.77E-01 | 9.86E-01 | -2.06E-02 | -1.01E+00 |
| ENSG000001 | CASQ2 | 6.63E-01 | ? | 8.26E-01 | -2.75E-01 | -1.21E+00 |
| ENSG000001 | ADPRM | 6.63E-01 | 8.77E-01 | 9.65E-01 | -5.17E-02 | -1.04E+00 |
| ENSG000002 | --- | 6.64E-01 | ? | 1.13E+00 | 1.76E-01 | 1.13E+00 |
| ENSG00000C | TUBA3D | 6.64E-01 | ? | 1.18E+00 | 2.35E-01 | 1.18E+00 |
| ENSG000001 | ZDHHC13 | 6.64E-01 | 8.77E-01 | 1.02E+00 | 2.61E-02 | 1.02E+00 |
| ENSG000002 | IL6R-AS1 | 6.64E-01 | ? | 1.16E+00 | 2.16E-01 | 1.16E+00 |
| ENSG000002 | LINGO3 | 6.64E-01 | ? | 8.23E-01 | -2.81E-01 | -1.22E+00 |
| ENSG000001 | MRPS7 | 6.64E-01 | 8.77E-01 | 1.02E+00 | 2.75E-02 | 1.02E+00 |
| ENSG000002 | LRRC73 | 6.64E-01 | ? | 1.19E+00 | 2.49E-01 | 1.19E+00 |
| ENSG000002 | --- | 6.64E-01 | 8.77E-01 | 9.83E-01 | -2.51E-02 | -1.02E+00 |

| | | | | | | |
|---|---|---|---|---|---|---|
| ENSG000002 | --- | 6.64E-01 | ? | 8.40E-01 | -2.51E-01 | -1.19E+00 |
| ENSG000001 | EMB | 6.64E-01 | 8.77E-01 | 9.83E-01 | -2.51E-02 | -1.02E+00 |
| ENSG000002 | --- | 6.64E-01 | 8.77E-01 | 9.72E-01 | -4.14E-02 | -1.03E+00 |
| ENSG000002 | --- | 6.64E-01 | ? | 9.21E-01 | -1.18E-01 | -1.09E+00 |
| ENSG000002 | SETP14 | 6.64E-01 | 8.77E-01 | 1.06E+00 | 8.92E-02 | 1.06E+00 |
| ENSG000002 | LINC02278 | 6.64E-01 | ? | 1.20E+00 | 2.62E-01 | 1.20E+00 |
| ENSG000002 | --- | 6.64E-01 | 8.77E-01 | 1.06E+00 | 8.87E-02 | 1.06E+00 |
| ENSG000001 | GRIK2 | 6.64E-01 | ? | 1.10E+00 | 1.41E-01 | 1.10E+00 |
| ENSG000002 | RAD51-AS1 | 6.64E-01 | 8.77E-01 | 1.06E+00 | 8.55E-02 | 1.06E+00 |
| ENSG000002 | --- | 6.64E-01 | 8.77E-01 | 9.20E-01 | -1.20E-01 | -1.09E+00 |
| ENSG000001 | TMCO1 | 6.65E-01 | 8.77E-01 | 1.01E+00 | 1.67E-02 | 1.01E+00 |
| ENSG000002 | EIF2AK3-DT | 6.65E-01 | 8.77E-01 | 1.06E+00 | 7.93E-02 | 1.06E+00 |
| ENSG000001 | ATF3 | 6.65E-01 | 8.77E-01 | 1.07E+00 | 9.98E-02 | 1.07E+00 |
| ENSG00000C | GOLGA3 | 6.65E-01 | 8.77E-01 | 9.86E-01 | -1.97E-02 | -1.01E+00 |
| ENSG000001 | CCDC136 | 6.65E-01 | ? | 9.02E-01 | -1.48E-01 | -1.11E+00 |
| ENSG000001 | NICN1 | 6.65E-01 | 8.78E-01 | 9.49E-01 | -7.60E-02 | -1.05E+00 |
| ENSG000001 | ACSM1 | 6.65E-01 | ? | 1.17E+00 | 2.31E-01 | 1.17E+00 |
| ENSG00000C | SUCO | 6.65E-01 | 8.78E-01 | 1.01E+00 | 2.09E-02 | 1.01E+00 |
| ENSG000001 | ZNF747 | 6.65E-01 | 8.78E-01 | 9.66E-01 | -4.94E-02 | -1.03E+00 |
| ENSG000002 | LINC01588 | 6.65E-01 | 8.78E-01 | 9.36E-01 | -9.47E-02 | -1.07E+00 |
| ENSG000002 | KHDC1-AS1 | 6.65E-01 | 8.78E-01 | 1.03E+00 | 3.98E-02 | 1.03E+00 |
| ENSG000001 | ANAPC10 | 6.66E-01 | 8.78E-01 | 9.76E-01 | -3.56E-02 | -1.02E+00 |
| ENSG000002 | --- | 6.66E-01 | ? | 1.18E+00 | 2.34E-01 | 1.18E+00 |
| ENSG000002 | RANP1 | 6.66E-01 | ? | 9.14E-01 | -1.30E-01 | -1.09E+00 |
| ENSG000002 | --- | 6.66E-01 | 8.78E-01 | 9.64E-01 | -5.28E-02 | -1.04E+00 |
| ENSG000001 | LYSMD4 | 6.66E-01 | 8.78E-01 | 9.70E-01 | -4.38E-02 | -1.03E+00 |
| ENSG000002 | HIVEP2-DT | 6.66E-01 | ? | 1.13E+00 | 1.76E-01 | 1.13E+00 |
| ENSG000001 | ZBTB38 | 6.66E-01 | 8.78E-01 | 9.83E-01 | -2.47E-02 | -1.02E+00 |
| ENSG000001 | SLC25A17 | 6.66E-01 | 8.78E-01 | 9.80E-01 | -2.89E-02 | -1.02E+00 |
| ENSG000001 | IGF1R | 6.66E-01 | 8.78E-01 | 9.83E-01 | -2.42E-02 | -1.02E+00 |
| ENSG00000C | GSK3B | 6.66E-01 | 8.78E-01 | 1.01E+00 | 1.92E-02 | 1.01E+00 |
| ENSG000001 | SLC35B3 | 6.66E-01 | 8.78E-01 | 1.01E+00 | 2.00E-02 | 1.01E+00 |
| ENSG000001 | TSR2 | 6.66E-01 | 8.78E-01 | 1.01E+00 | 1.82E-02 | 1.01E+00 |
| ENSG000002 | YWHAZP2 | 6.66E-01 | ? | 1.10E+00 | 1.35E-01 | 1.10E+00 |
| ENSG000002 | RPS10P7 | 6.66E-01 | 8.78E-01 | 1.07E+00 | 9.65E-02 | 1.07E+00 |
| ENSG000001 | CRK | 6.66E-01 | 8.78E-01 | 1.01E+00 | 1.44E-02 | 1.01E+00 |
| ENSG000001 | USP53 | 6.66E-01 | 8.78E-01 | 1.02E+00 | 3.28E-02 | 1.02E+00 |
| ENSG000002 | AGAP1-IT1 | 6.66E-01 | ? | 7.88E-01 | -3.44E-01 | -1.27E+00 |
| ENSG000001 | BRSK2 | 6.66E-01 | ? | 1.15E+00 | 1.98E-01 | 1.15E+00 |
| ENSG000002 | GHRLOS | 6.67E-01 | 8.78E-01 | 9.48E-01 | -7.70E-02 | -1.05E+00 |
| ENSG000001 | CCDC90B | 6.67E-01 | 8.78E-01 | 1.02E+00 | 2.51E-02 | 1.02E+00 |
| ENSG000001 | FRS2 | 6.67E-01 | 8.78E-01 | 1.01E+00 | 2.03E-02 | 1.01E+00 |
| ENSG000002 | DBIL5P | 6.67E-01 | ? | 1.12E+00 | 1.63E-01 | 1.12E+00 |
| ENSG000001 | AP1G1 | 6.67E-01 | 8.78E-01 | 9.89E-01 | -1.57E-02 | -1.01E+00 |
| ENSG000001 | ELAPOR2 | 6.67E-01 | 8.78E-01 | 1.03E+00 | 4.45E-02 | 1.03E+00 |

| | | | | | | |
|---|---|---|---|---|---|---|
| ENSG000001 | FBXO8 | 6.67E-01 | 8.78E-01 | 9.78E-01 | -3.24E-02 | -1.02E+00 |
| ENSG000002 | LINC02767 | 6.67E-01 | ? | 8.56E-01 | -2.24E-01 | -1.17E+00 |
| ENSG000002 | --- | 6.67E-01 | 8.78E-01 | 9.24E-01 | -1.13E-01 | -1.08E+00 |
| ENSG000001 | ILDR2 | 6.67E-01 | ? | 1.20E+00 | 2.64E-01 | 1.20E+00 |
| ENSG00000C | LMO3 | 6.67E-01 | 8.78E-01 | 1.02E+00 | 2.87E-02 | 1.02E+00 |
| ENSG000001 | OR7E38P | 6.67E-01 | 8.78E-01 | 1.06E+00 | 8.38E-02 | 1.06E+00 |
| ENSG000001 | EIF3B | 6.67E-01 | 8.78E-01 | 9.91E-01 | -1.24E-02 | -1.01E+00 |
| ENSG000001 | TMED6 | 6.67E-01 | ? | 1.09E+00 | 1.26E-01 | 1.09E+00 |
| ENSG000002 | RPS20P14 | 6.67E-01 | ? | 8.85E-01 | -1.76E-01 | -1.13E+00 |
| ENSG00000C | SLC2A3 | 6.67E-01 | 8.78E-01 | 1.02E+00 | 3.38E-02 | 1.02E+00 |
| ENSG000001 | DESI2 | 6.67E-01 | 8.78E-01 | 1.01E+00 | 1.79E-02 | 1.01E+00 |
| ENSG000002 | --- | 6.67E-01 | 8.78E-01 | 9.43E-01 | -8.53E-02 | -1.06E+00 |
| ENSG000002 | --- | 6.68E-01 | ? | 1.15E+00 | 2.00E-01 | 1.15E+00 |
| ENSG000002 | LINC02057 | 6.68E-01 | 8.79E-01 | 9.55E-01 | -6.70E-02 | -1.05E+00 |
| ENSG000002 | --- | 6.68E-01 | 8.79E-01 | 1.12E+00 | 1.65E-01 | 1.12E+00 |
| ENSG000001 | POMT1 | 6.68E-01 | 8.79E-01 | 1.02E+00 | 2.80E-02 | 1.02E+00 |
| ENSG000002 | --- | 6.68E-01 | ? | 8.59E-01 | -2.19E-01 | -1.16E+00 |
| ENSG000001 | MID1IP1 | 6.68E-01 | 8.79E-01 | 1.02E+00 | 3.01E-02 | 1.02E+00 |
| ENSG000002 | LINC01605 | 6.68E-01 | 8.79E-01 | 9.62E-01 | -5.63E-02 | -1.04E+00 |
| ENSG000001 | VPS37C | 6.68E-01 | 8.79E-01 | 1.02E+00 | 3.26E-02 | 1.02E+00 |
| ENSG000001 | POLL | 6.68E-01 | 8.79E-01 | 1.03E+00 | 4.32E-02 | 1.03E+00 |
| ENSG000002 | --- | 6.68E-01 | ? | 1.09E+00 | 1.30E-01 | 1.09E+00 |
| ENSG00000C | SPAG7 | 6.68E-01 | 8.79E-01 | 1.02E+00 | 2.19E-02 | 1.02E+00 |
| ENSG000002 | --- | 6.68E-01 | ? | 9.09E-01 | -1.38E-01 | -1.10E+00 |
| ENSG000002 | --- | 6.68E-01 | ? | 1.19E+00 | 2.56E-01 | 1.19E+00 |
| ENSG000002 | TMC3-AS1 | 6.68E-01 | ? | 8.39E-01 | -2.54E-01 | -1.19E+00 |
| ENSG000001 | NLGN2 | 6.68E-01 | 8.79E-01 | 1.03E+00 | 3.87E-02 | 1.03E+00 |
| ENSG000001 | ANKEF1 | 6.68E-01 | 8.79E-01 | 9.80E-01 | -2.85E-02 | -1.02E+00 |
| ENSG000001 | KPNA2 | 6.68E-01 | 8.79E-01 | 9.84E-01 | -2.33E-02 | -1.02E+00 |
| ENSG000002 | --- | 6.69E-01 | ? | 9.16E-01 | -1.27E-01 | -1.09E+00 |
| ENSG000002 | LINC00184 | 6.69E-01 | ? | 9.16E-01 | -1.27E-01 | -1.09E+00 |
| ENSG000001 | TMBIM1 | 6.69E-01 | 8.79E-01 | 1.02E+00 | 2.42E-02 | 1.02E+00 |
| ENSG000002 | --- | 6.69E-01 | ? | 8.30E-01 | -2.68E-01 | -1.20E+00 |
| ENSG000001 | SAMD1 | 6.69E-01 | 8.79E-01 | 9.79E-01 | -3.10E-02 | -1.02E+00 |
| ENSG000002 | --- | 6.69E-01 | ? | 8.98E-01 | -1.55E-01 | -1.11E+00 |
| ENSG000001 | RPL23P2 | 6.69E-01 | ? | 1.20E+00 | 2.67E-01 | 1.20E+00 |
| ENSG000001 | PPP1R13L | 6.69E-01 | 8.79E-01 | 1.02E+00 | 3.52E-02 | 1.02E+00 |
| ENSG000002 | ZNF2 | 6.69E-01 | 8.79E-01 | 1.03E+00 | 4.15E-02 | 1.03E+00 |
| ENSG000002 | --- | 6.69E-01 | ? | 1.13E+00 | 1.76E-01 | 1.13E+00 |
| ENSG000002 | --- | 6.69E-01 | 8.79E-01 | 8.85E-01 | -1.76E-01 | -1.13E+00 |
| ENSG000001 | ZNF491 | 6.69E-01 | 8.79E-01 | 9.51E-01 | -7.27E-02 | -1.05E+00 |
| ENSG000001 | BEAN1 | 6.69E-01 | 8.79E-01 | 1.07E+00 | 1.00E-01 | 1.07E+00 |
| ENSG00000C | FUNDC1 | 6.69E-01 | 8.79E-01 | 9.76E-01 | -3.58E-02 | -1.03E+00 |
| ENSG000001 | IFI27L1 | 6.69E-01 | 8.79E-01 | 1.04E+00 | 5.46E-02 | 1.04E+00 |
| ENSG000001 | ELF3 | 6.69E-01 | 8.79E-01 | 9.43E-01 | -8.46E-02 | -1.06E+00 |

| | | | | | | |
|---|---|---|---|---|---|---|
| ENSG000001 | MDK | 6.69E-01 | 8.79E-01 | 9.68E-01 | -4.65E-02 | -1.03E+00 |
| ENSG000002 | --- | 6.69E-01 | ? | 1.15E+00 | 2.05E-01 | 1.15E+00 |
| ENSG000001 | NDUFAF6 | 6.70E-01 | 8.79E-01 | 1.02E+00 | 2.85E-02 | 1.02E+00 |
| ENSG000001 | BRPF1 | 6.70E-01 | 8.79E-01 | 1.01E+00 | 2.13E-02 | 1.01E+00 |
| ENSG000001 | POU6F1 | 6.70E-01 | 8.79E-01 | 1.07E+00 | 9.80E-02 | 1.07E+00 |
| ENSG000001 | MICALL2 | 6.70E-01 | 8.79E-01 | 1.03E+00 | 4.74E-02 | 1.03E+00 |
| ENSG000002 | --- | 6.70E-01 | ? | 8.26E-01 | -2.75E-01 | -1.21E+00 |
| ENSG000002 | SAMD5 | 6.70E-01 | 8.79E-01 | 9.80E-01 | -2.97E-02 | -1.02E+00 |
| ENSG000002 | MINCR | 6.70E-01 | 8.79E-01 | 1.07E+00 | 9.51E-02 | 1.07E+00 |
| ENSG000002 | --- | 6.70E-01 | ? | 8.48E-01 | -2.38E-01 | -1.18E+00 |
| ENSG000001 | RFLNB | 6.70E-01 | 8.80E-01 | 1.03E+00 | 4.34E-02 | 1.03E+00 |
| ENSG000001 | DUSP9 | 6.70E-01 | ? | 1.10E+00 | 1.41E-01 | 1.10E+00 |
| ENSG000001 | CLDN7 | 6.70E-01 | 8.80E-01 | 9.66E-01 | -5.01E-02 | -1.04E+00 |
| ENSG000001 | CITED2 | 6.71E-01 | 8.80E-01 | 9.80E-01 | -2.91E-02 | -1.02E+00 |
| ENSG000001 | ZNF777 | 6.71E-01 | 8.80E-01 | 9.84E-01 | -2.36E-02 | -1.02E+00 |
| ENSG000002 | --- | 6.71E-01 | 8.80E-01 | 9.07E-01 | -1.41E-01 | -1.10E+00 |
| ENSG000002 | --- | 6.71E-01 | ? | 8.75E-01 | -1.93E-01 | -1.14E+00 |
| ENSG000001 | DDX46 | 6.71E-01 | 8.80E-01 | 1.01E+00 | 2.14E-02 | 1.01E+00 |
| ENSG000001 | LIMD2 | 6.71E-01 | 8.80E-01 | 9.59E-01 | -6.03E-02 | -1.04E+00 |
| ENSG000001 | DDA1 | 6.71E-01 | 8.80E-01 | 1.02E+00 | 2.18E-02 | 1.02E+00 |
| ENSG000001 | KIAA0753 | 6.71E-01 | 8.80E-01 | 1.01E+00 | 2.01E-02 | 1.01E+00 |
| ENSG000001 | NR3C1 | 6.71E-01 | 8.80E-01 | 1.02E+00 | 2.77E-02 | 1.02E+00 |
| ENSG000001 | CCK | 6.71E-01 | 8.80E-01 | 9.45E-01 | -8.19E-02 | -1.06E+00 |
| ENSG000001 | RESF1 | 6.71E-01 | 8.80E-01 | 9.72E-01 | -4.07E-02 | -1.03E+00 |
| ENSG000002 | --- | 6.71E-01 | ? | 8.94E-01 | -1.61E-01 | -1.12E+00 |
| ENSG000001 | LCP1 | 6.71E-01 | 8.80E-01 | 1.08E+00 | 1.08E-01 | 1.08E+00 |
| ENSG000002 | BCLAF1P2 | 6.71E-01 | ? | 9.21E-01 | -1.19E-01 | -1.09E+00 |
| ENSG000002 | SYNJ2BP-CO | 6.71E-01 | 8.80E-01 | 8.64E-01 | -2.11E-01 | -1.16E+00 |
| ENSG000002 | AKR7L | 6.71E-01 | ? | 8.59E-01 | -2.20E-01 | -1.16E+00 |
| ENSG000002 | --- | 6.71E-01 | 8.80E-01 | 7.83E-01 | -3.53E-01 | -1.28E+00 |
| ENSG000001 | HTR4 | 6.71E-01 | ? | 1.10E+00 | 1.32E-01 | 1.10E+00 |
| ENSG000001 | SNRNP27 | 6.71E-01 | 8.80E-01 | 1.02E+00 | 3.11E-02 | 1.02E+00 |
| ENSG000002 | --- | 6.71E-01 | ? | 1.09E+00 | 1.24E-01 | 1.09E+00 |
| ENSG000002 | --- | 6.71E-01 | 8.80E-01 | 9.53E-01 | -6.88E-02 | -1.05E+00 |
| ENSG000001 | DCPS | 6.71E-01 | 8.80E-01 | 1.02E+00 | 2.95E-02 | 1.02E+00 |
| ENSG000001 | H2AC11 | 6.71E-01 | ? | 8.90E-01 | -1.69E-01 | -1.12E+00 |
| ENSG00000C | HOOK2 | 6.71E-01 | 8.80E-01 | 9.73E-01 | -4.02E-02 | -1.03E+00 |
| ENSG000001 | PTPN14 | 6.72E-01 | 8.80E-01 | 1.02E+00 | 2.74E-02 | 1.02E+00 |
| ENSG000001 | VANGL1 | 6.72E-01 | 8.80E-01 | 1.01E+00 | 1.75E-02 | 1.01E+00 |
| ENSG000002 | --- | 6.72E-01 | ? | 1.17E+00 | 2.25E-01 | 1.17E+00 |
| ENSG000002 | ADAM20P1 | 6.72E-01 | ? | 7.98E-01 | -3.25E-01 | -1.25E+00 |
| ENSG000002 | --- | 6.72E-01 | 8.80E-01 | 1.04E+00 | 5.63E-02 | 1.04E+00 |
| ENSG000002 | --- | 6.72E-01 | ? | 7.80E-01 | -3.59E-01 | -1.28E+00 |
| ENSG000001 | HES7 | 6.72E-01 | ? | 8.78E-01 | -1.88E-01 | -1.14E+00 |
| ENSG000002 | --- | 6.72E-01 | 8.80E-01 | 1.04E+00 | 5.26E-02 | 1.04E+00 |

| | | | | | | | |
|---|---|---|---|---|---|---|---|
| ENSG000002 | --- | 6.72E-01 | | 8.80E-01 | 9.35E-01 | -9.76E-02 | -1.07E+00 |
| ENSG000002 | --- | 6.72E-01 | ? | | 1.19E+00 | 2.53E-01 | 1.19E+00 |
| ENSG00000C | CAMK1G | 6.72E-01 | ? | | 1.21E+00 | 2.75E-01 | 1.21E+00 |
| ENSG000001 | NBPF20 | 6.72E-01 | | 8.80E-01 | 9.59E-01 | -6.08E-02 | -1.04E+00 |
| ENSG000001 | CCT8 | 6.72E-01 | | 8.80E-01 | 1.01E+00 | 1.32E-02 | 1.01E+00 |
| ENSG000002 | APOBEC3D | 6.72E-01 | ? | | 1.22E+00 | 2.82E-01 | 1.22E+00 |
| ENSG00000C | HES2 | 6.73E-01 | | 8.80E-01 | 1.01E+00 | 1.74E-02 | 1.01E+00 |
| ENSG000001 | FAM151B | 6.73E-01 | ? | | 9.30E-01 | -1.05E-01 | -1.08E+00 |
| ENSG000001 | SMAD3 | 6.73E-01 | | 8.80E-01 | 9.86E-01 | -2.00E-02 | -1.01E+00 |
| ENSG00000C | TSR3 | 6.73E-01 | | 8.80E-01 | 1.03E+00 | 3.73E-02 | 1.03E+00 |
| ENSG000001 | RAB33B | 6.73E-01 | | 8.80E-01 | 9.80E-01 | -2.96E-02 | -1.02E+00 |
| ENSG000001 | VAV2 | 6.73E-01 | | 8.80E-01 | 1.01E+00 | 1.85E-02 | 1.01E+00 |
| ENSG000001 | ZNF623 | 6.73E-01 | | 8.80E-01 | 9.81E-01 | -2.71E-02 | -1.02E+00 |
| ENSG000001 | DNAH1 | 6.73E-01 | | 8.80E-01 | 1.04E+00 | 5.41E-02 | 1.04E+00 |
| ENSG000001 | AHI1 | 6.73E-01 | | 8.80E-01 | 9.77E-01 | -3.32E-02 | -1.02E+00 |
| ENSG00000C | SLC24A1 | 6.73E-01 | | 8.80E-01 | 9.81E-01 | -2.74E-02 | -1.02E+00 |
| ENSG000001 | C2CD6 | 6.73E-01 | ? | | 1.20E+00 | 2.66E-01 | 1.20E+00 |
| ENSG000001 | VPS26A | 6.73E-01 | | 8.80E-01 | 1.01E+00 | 2.01E-02 | 1.01E+00 |
| ENSG000001 | DENND2C | 6.73E-01 | | 8.80E-01 | 9.81E-01 | -2.81E-02 | -1.02E+00 |
| ENSG000001 | LRRC43 | 6.73E-01 | ? | | 1.09E+00 | 1.27E-01 | 1.09E+00 |
| ENSG000002 | --- | 6.73E-01 | ? | | 9.02E-01 | -1.49E-01 | -1.11E+00 |
| ENSG000002 | --- | 6.73E-01 | ? | | 8.11E-01 | -3.01E-01 | -1.23E+00 |
| ENSG000002 | CPHL1P | 6.73E-01 | ? | | 1.20E+00 | 2.66E-01 | 1.20E+00 |
| ENSG000002 | ANKHD1-DT | 6.73E-01 | ? | | 1.17E+00 | 2.29E-01 | 1.17E+00 |
| ENSG000001 | NFKBID | 6.73E-01 | | 8.81E-01 | 9.53E-01 | -6.95E-02 | -1.05E+00 |
| ENSG000002 | --- | 6.73E-01 | ? | | 1.19E+00 | 2.56E-01 | 1.19E+00 |
| ENSG000001 | SYNE1 | 6.73E-01 | | 8.81E-01 | 9.74E-01 | -3.80E-02 | -1.03E+00 |
| ENSG000002 | ICA1-AS1 | 6.73E-01 | ? | | 1.27E+00 | 3.45E-01 | 1.27E+00 |
| ENSG000001 | TVP23C | 6.73E-01 | | 8.81E-01 | 9.64E-01 | -5.24E-02 | -1.04E+00 |
| ENSG000002 | PLPP4 | 6.73E-01 | | 8.81E-01 | 1.06E+00 | 8.67E-02 | 1.06E+00 |
| ENSG000001 | NT5DC3 | 6.73E-01 | | 8.81E-01 | 9.86E-01 | -2.01E-02 | -1.01E+00 |
| ENSG000001 | SLC26A2 | 6.74E-01 | | 8.81E-01 | 9.78E-01 | -3.15E-02 | -1.02E+00 |
| ENSG000001 | ODF3B | 6.74E-01 | ? | | 1.08E+00 | 1.12E-01 | 1.08E+00 |
| ENSG00000C | VIM | 6.74E-01 | | 8.81E-01 | 9.81E-01 | -2.80E-02 | -1.02E+00 |
| ENSG000002 | --- | 6.74E-01 | ? | | 1.09E+00 | 1.28E-01 | 1.09E+00 |
| ENSG00000C | LASP1 | 6.74E-01 | | 8.81E-01 | 1.01E+00 | 1.60E-02 | 1.01E+00 |
| ENSG000001 | PDE5A | 6.74E-01 | | 8.81E-01 | 1.03E+00 | 4.08E-02 | 1.03E+00 |
| ENSG000002 | RAB11FIP1P1 | 6.74E-01 | ? | | 8.93E-01 | -1.64E-01 | -1.12E+00 |
| ENSG000001 | RPGRIP1L | 6.74E-01 | | 8.81E-01 | 9.77E-01 | -3.37E-02 | -1.02E+00 |
| ENSG000001 | UBE4B | 6.74E-01 | | 8.81E-01 | 9.88E-01 | -1.81E-02 | -1.01E+00 |
| ENSG000002 | --- | 6.74E-01 | ? | | 8.98E-01 | -1.55E-01 | -1.11E+00 |
| ENSG000001 | RYK | 6.74E-01 | | 8.81E-01 | 1.02E+00 | 2.45E-02 | 1.02E+00 |
| ENSG000001 | ZFHX3 | 6.74E-01 | | 8.81E-01 | 9.73E-01 | -4.01E-02 | -1.03E+00 |
| ENSG000001 | WWP1 | 6.74E-01 | | 8.81E-01 | 1.01E+00 | 1.80E-02 | 1.01E+00 |
| ENSG000001 | BLVRA | 6.74E-01 | | 8.81E-01 | 9.83E-01 | -2.52E-02 | -1.02E+00 |

| | | | | | | |
|---|---|---|---|---|---|---|
| ENSG000000 | ADAM28 | 6.74E-01 | 8.81E-01 | 9.27E-01 | -1.09E-01 | -1.08E+00 |
| ENSG000000 | CALCOCO1 | 6.75E-01 | 8.81E-01 | 9.82E-01 | -2.57E-02 | -1.02E+00 |
| ENSG000000 | ASNS | 6.75E-01 | 8.81E-01 | 1.03E+00 | 3.57E-02 | 1.03E+00 |
| ENSG000001 | MRPS27 | 6.75E-01 | 8.81E-01 | 9.87E-01 | -1.84E-02 | -1.01E+00 |
| ENSG000001 | TRIM21 | 6.75E-01 | 8.81E-01 | 9.84E-01 | -2.32E-02 | -1.02E+00 |
| ENSG000001 | USP25 | 6.75E-01 | 8.81E-01 | 1.02E+00 | 2.42E-02 | 1.02E+00 |
| ENSG000001 | CDC42SE2 | 6.75E-01 | 8.81E-01 | 1.01E+00 | 1.54E-02 | 1.01E+00 |
| ENSG000000 | SEH1L | 6.75E-01 | 8.81E-01 | 9.84E-01 | -2.28E-02 | -1.02E+00 |
| ENSG000001 | C8orf48 | 6.75E-01 | 8.81E-01 | 9.56E-01 | -6.50E-02 | -1.05E+00 |
| ENSG000002 | IKBKB-DT | 6.75E-01 | 8.81E-01 | 9.29E-01 | -1.07E-01 | -1.08E+00 |
| ENSG000002 | ZNF571-AS1 | 6.75E-01 | 8.81E-01 | 9.50E-01 | -7.37E-02 | -1.05E+00 |
| ENSG000002 | --- | 6.75E-01 | 8.81E-01 | 9.28E-01 | -1.07E-01 | -1.08E+00 |
| ENSG000002 | --- | 6.75E-01 | ? | 8.33E-01 | -2.64E-01 | -1.20E+00 |
| ENSG000001 | ZNF784 | 6.75E-01 | 8.81E-01 | 1.05E+00 | 6.50E-02 | 1.05E+00 |
| ENSG000002 | --- | 6.75E-01 | 8.81E-01 | 9.62E-01 | -5.52E-02 | -1.04E+00 |
| ENSG000002 | C5orf60 | 6.75E-01 | ? | 1.25E+00 | 3.23E-01 | 1.25E+00 |
| ENSG000001 | GCC2 | 6.75E-01 | 8.81E-01 | 9.77E-01 | -3.36E-02 | -1.02E+00 |
| ENSG000002 | RPS2P7 | 6.75E-01 | 8.81E-01 | 9.51E-01 | -7.24E-02 | -1.05E+00 |
| ENSG000001 | FAM98B | 6.75E-01 | 8.81E-01 | 1.01E+00 | 2.10E-02 | 1.01E+00 |
| ENSG000002 | FAM222A-AS | 6.75E-01 | ? | 8.60E-01 | -2.18E-01 | -1.16E+00 |
| ENSG000001 | PTCH2 | 6.75E-01 | ? | 1.10E+00 | 1.32E-01 | 1.10E+00 |
| ENSG000001 | IMP4 | 6.76E-01 | 8.81E-01 | 1.02E+00 | 2.47E-02 | 1.02E+00 |
| ENSG000002 | --- | 6.76E-01 | ? | 7.18E-01 | -4.78E-01 | -1.39E+00 |
| ENSG000001 | RPRD1B | 6.76E-01 | 8.81E-01 | 1.01E+00 | 1.81E-02 | 1.01E+00 |
| ENSG000001 | FAM50B | 6.76E-01 | 8.81E-01 | 9.57E-01 | -6.31E-02 | -1.04E+00 |
| ENSG000001 | ZNF749 | 6.76E-01 | 8.81E-01 | 9.77E-01 | -3.41E-02 | -1.02E+00 |
| ENSG000001 | SMG1P5 | 6.76E-01 | 8.81E-01 | 1.08E+00 | 1.16E-01 | 1.08E+00 |
| ENSG000001 | PGBD1 | 6.76E-01 | 8.81E-01 | 9.71E-01 | -4.22E-02 | -1.03E+00 |
| ENSG000002 | --- | 6.76E-01 | ? | 8.50E-01 | -2.34E-01 | -1.18E+00 |
| ENSG000001 | DVL1 | 6.76E-01 | 8.81E-01 | 9.72E-01 | -4.11E-02 | -1.03E+00 |
| ENSG000002 | --- | 6.76E-01 | ? | 1.21E+00 | 2.76E-01 | 1.21E+00 |
| ENSG000002 | ENO1P4 | 6.76E-01 | ? | 7.08E-01 | -4.99E-01 | -1.41E+00 |
| ENSG000002 | SRSF8 | 6.76E-01 | 8.81E-01 | 1.01E+00 | 1.90E-02 | 1.01E+00 |
| ENSG000002 | --- | 6.76E-01 | ? | 1.18E+00 | 2.41E-01 | 1.18E+00 |
| ENSG000001 | CENPI | 6.76E-01 | 8.81E-01 | 9.67E-01 | -4.77E-02 | -1.03E+00 |
| ENSG000001 | PPP2CB | 6.76E-01 | 8.81E-01 | 1.01E+00 | 1.54E-02 | 1.01E+00 |
| ENSG000001 | CD164 | 6.76E-01 | 8.81E-01 | 9.89E-01 | -1.59E-02 | -1.01E+00 |
| ENSG000001 | ABCA15P | 6.76E-01 | ? | 8.94E-01 | -1.61E-01 | -1.12E+00 |
| ENSG000002 | C11orf94 | 6.76E-01 | ? | 9.02E-01 | -1.48E-01 | -1.11E+00 |
| ENSG000000 | SYN1 | 6.76E-01 | ? | 1.13E+00 | 1.73E-01 | 1.13E+00 |
| ENSG000002 | --- | 6.76E-01 | ? | 1.16E+00 | 2.13E-01 | 1.16E+00 |
| ENSG000002 | EIF3FP3 | 6.76E-01 | 8.81E-01 | 9.46E-01 | -8.01E-02 | -1.06E+00 |
| ENSG000001 | SIK3 | 6.76E-01 | 8.81E-01 | 1.01E+00 | 1.83E-02 | 1.01E+00 |
| ENSG000002 | --- | 6.76E-01 | ? | 8.22E-01 | -2.82E-01 | -1.22E+00 |
| ENSG000001 | EPSTI1 | 6.77E-01 | 8.81E-01 | 1.05E+00 | 6.84E-02 | 1.05E+00 |

| | | | | | | |
|---|---|---|---|---|---|---|
| ENSG000001 | NCLN | 6.77E-01 | 8.82E-01 | 1.03E+00 | 4.11E-02 | 1.03E+00 |
| ENSG000001 | PROSER3 | 6.77E-01 | 8.82E-01 | 1.02E+00 | 2.22E-02 | 1.02E+00 |
| ENSG000002 | HERC2P2 | 6.77E-01 | 8.82E-01 | 1.05E+00 | 6.61E-02 | 1.05E+00 |
| ENSG000001 | PPP6R1 | 6.77E-01 | 8.82E-01 | 9.86E-01 | -1.97E-02 | -1.01E+00 |
| ENSG000002 | --- | 6.77E-01 | ? | 6.91E-01 | -5.34E-01 | -1.45E+00 |
| ENSG00000C | PTPRU | 6.77E-01 | 8.82E-01 | 1.02E+00 | 3.15E-02 | 1.02E+00 |
| ENSG000002 | --- | 6.77E-01 | ? | 8.39E-01 | -2.53E-01 | -1.19E+00 |
| ENSG000001 | IFIT1 | 6.77E-01 | ? | 1.10E+00 | 1.40E-01 | 1.10E+00 |
| ENSG00000C | TFAP4 | 6.78E-01 | 8.82E-01 | 9.68E-01 | -4.67E-02 | -1.03E+00 |
| ENSG000002 | --- | 6.78E-01 | ? | 8.81E-01 | -1.83E-01 | -1.13E+00 |
| ENSG000001 | OCIAD2 | 6.78E-01 | 8.82E-01 | 1.01E+00 | 1.61E-02 | 1.01E+00 |
| ENSG00000C | FAM3A | 6.78E-01 | 8.82E-01 | 9.76E-01 | -3.52E-02 | -1.02E+00 |
| ENSG000001 | RTTN | 6.78E-01 | 8.82E-01 | 1.02E+00 | 2.17E-02 | 1.02E+00 |
| ENSG00000C | TLL1 | 6.78E-01 | ? | 1.12E+00 | 1.58E-01 | 1.12E+00 |
| ENSG000001 | ZNF519 | 6.78E-01 | 8.82E-01 | 1.06E+00 | 8.29E-02 | 1.06E+00 |
| ENSG000001 | UBB | 6.78E-01 | 8.82E-01 | 9.89E-01 | -1.58E-02 | -1.01E+00 |
| ENSG000001 | RPAIN | 6.78E-01 | 8.82E-01 | 9.73E-01 | -3.92E-02 | -1.03E+00 |
| ENSG000001 | ANKRD2 | 6.78E-01 | 8.82E-01 | 9.38E-01 | -9.16E-02 | -1.07E+00 |
| ENSG000001 | ADRA1B | 6.78E-01 | ? | 1.13E+00 | 1.73E-01 | 1.13E+00 |
| ENSG000002 | --- | 6.78E-01 | ? | 1.16E+00 | 2.14E-01 | 1.16E+00 |
| ENSG000001 | SPIDR | 6.78E-01 | 8.82E-01 | 1.02E+00 | 2.80E-02 | 1.02E+00 |
| ENSG000001 | CHMP4C | 6.78E-01 | 8.82E-01 | 1.02E+00 | 2.24E-02 | 1.02E+00 |
| ENSG000001 | CLCN7 | 6.78E-01 | 8.82E-01 | 1.03E+00 | 4.21E-02 | 1.03E+00 |
| ENSG000002 | FAM86B3P | 6.78E-01 | 8.82E-01 | 1.03E+00 | 4.55E-02 | 1.03E+00 |
| ENSG000001 | CHST2 | 6.79E-01 | 8.83E-01 | 1.03E+00 | 4.55E-02 | 1.03E+00 |
| ENSG00000C | CLCN6 | 6.79E-01 | 8.83E-01 | 9.77E-01 | -3.29E-02 | -1.02E+00 |
| ENSG000001 | EMILIN1 | 6.79E-01 | ? | 1.16E+00 | 2.18E-01 | 1.16E+00 |
| ENSG000002 | SPDYE3 | 6.79E-01 | ? | 8.62E-01 | -2.14E-01 | -1.16E+00 |
| ENSG000001 | HAPSTR1 | 6.79E-01 | 8.83E-01 | 9.84E-01 | -2.32E-02 | -1.02E+00 |
| ENSG000001 | INPP5J | 6.79E-01 | ? | 9.12E-01 | -1.33E-01 | -1.10E+00 |
| ENSG000001 | RAN | 6.79E-01 | 8.83E-01 | 1.01E+00 | 1.45E-02 | 1.01E+00 |
| ENSG000002 | SERTAD4-AS | 6.79E-01 | 8.83E-01 | 1.07E+00 | 1.03E-01 | 1.07E+00 |
| ENSG000001 | SHLD2P1 | 6.79E-01 | ? | 8.91E-01 | -1.67E-01 | -1.12E+00 |
| ENSG000001 | TBCB | 6.79E-01 | 8.83E-01 | 1.02E+00 | 2.57E-02 | 1.02E+00 |
| ENSG000001 | ZNF552 | 6.79E-01 | 8.83E-01 | 1.02E+00 | 3.37E-02 | 1.02E+00 |
| ENSG000001 | F5 | 6.79E-01 | ? | 8.72E-01 | -1.98E-01 | -1.15E+00 |
| ENSG000001 | FOXRED2 | 6.79E-01 | 8.83E-01 | 9.65E-01 | -5.08E-02 | -1.04E+00 |
| ENSG000001 | NECTIN3 | 6.80E-01 | 8.83E-01 | 9.81E-01 | -2.78E-02 | -1.02E+00 |
| ENSG000002 | --- | 6.80E-01 | ? | 1.20E+00 | 2.64E-01 | 1.20E+00 |
| ENSG000002 | LRRFIP1P1 | 6.80E-01 | 8.83E-01 | 9.44E-01 | -8.39E-02 | -1.06E+00 |
| ENSG000002 | MIR2117HG | 6.80E-01 | ? | 8.84E-01 | -1.77E-01 | -1.13E+00 |
| ENSG00000C | CCDC88C | 6.80E-01 | 8.83E-01 | 1.02E+00 | 3.52E-02 | 1.02E+00 |
| ENSG000001 | SPATA13 | 6.80E-01 | 8.84E-01 | 1.04E+00 | 5.37E-02 | 1.04E+00 |
| ENSG000001 | LIN7C | 6.80E-01 | 8.84E-01 | 9.83E-01 | -2.49E-02 | -1.02E+00 |
| ENSG000002 | SOD2-OT1 | 6.80E-01 | ? | 1.13E+00 | 1.72E-01 | 1.13E+00 |

| | | | | | | |
|---|---|---|---|---|---|---|
| ENSG000002 | --- | 6.80E-01 | ? | 8.31E-01 | -2.68E-01 | -1.20E+00 |
| ENSG000001 | F2RL1 | 6.80E-01 | 8.84E-01 | 1.02E+00 | 2.70E-02 | 1.02E+00 |
| ENSG000002 | UGT1A3 | 6.80E-01 | 8.84E-01 | 9.39E-01 | -9.04E-02 | -1.06E+00 |
| ENSG000002 | SAPCD1 | 6.80E-01 | ? | 1.15E+00 | 2.00E-01 | 1.15E+00 |
| ENSG000001 | EHD4 | 6.80E-01 | 8.84E-01 | 9.83E-01 | -2.43E-02 | -1.02E+00 |
| ENSG000002 | NAXD | 6.81E-01 | 8.84E-01 | 1.03E+00 | 3.60E-02 | 1.03E+00 |
| ENSG00000C | PALB2 | 6.81E-01 | 8.84E-01 | 1.02E+00 | 2.50E-02 | 1.02E+00 |
| ENSG000002 | IPO7P2 | 6.81E-01 | ? | 1.14E+00 | 1.93E-01 | 1.14E+00 |
| ENSG000002 | MEG9 | 6.81E-01 | 8.84E-01 | 1.08E+00 | 1.12E-01 | 1.08E+00 |
| ENSG000002 | ST6GALNAC6 | 6.81E-01 | 8.84E-01 | 9.59E-01 | -5.99E-02 | -1.04E+00 |
| ENSG000002 | RN7SKP11 | 6.81E-01 | ? | 1.11E+00 | 1.56E-01 | 1.11E+00 |
| ENSG000001 | CBFA2T3 | 6.81E-01 | 8.84E-01 | 9.56E-01 | -6.42E-02 | -1.05E+00 |
| ENSG000002 | PCBP2-OT1 | 6.81E-01 | ? | 1.17E+00 | 2.31E-01 | 1.17E+00 |
| ENSG00000C | ZDHHC8 | 6.81E-01 | 8.84E-01 | 1.03E+00 | 4.28E-02 | 1.03E+00 |
| ENSG00000C | STARD7 | 6.81E-01 | 8.84E-01 | 1.01E+00 | 2.13E-02 | 1.01E+00 |
| ENSG000001 | ACTRT3 | 6.81E-01 | 8.84E-01 | 1.03E+00 | 3.58E-02 | 1.03E+00 |
| ENSG000001 | AP4B1 | 6.81E-01 | 8.84E-01 | 9.81E-01 | -2.73E-02 | -1.02E+00 |
| ENSG000001 | FAM25A | 6.81E-01 | ? | 1.27E+00 | 3.47E-01 | 1.27E+00 |
| ENSG000001 | FAM53C | 6.82E-01 | 8.84E-01 | 1.01E+00 | 2.04E-02 | 1.01E+00 |
| ENSG000001 | DYNAP | 6.82E-01 | ? | 8.27E-01 | -2.74E-01 | -1.21E+00 |
| ENSG00000C | RC3H2 | 6.82E-01 | 8.84E-01 | 9.86E-01 | -2.07E-02 | -1.01E+00 |
| ENSG000002 | FPGT | 6.82E-01 | 8.84E-01 | 1.02E+00 | 2.29E-02 | 1.02E+00 |
| ENSG000002 | --- | 6.82E-01 | ? | 8.50E-01 | -2.35E-01 | -1.18E+00 |
| ENSG00000C | NEXMIF | 6.82E-01 | ? | 1.17E+00 | 2.32E-01 | 1.17E+00 |
| ENSG000001 | HK1 | 6.82E-01 | 8.84E-01 | 1.01E+00 | 1.03E-02 | 1.01E+00 |
| ENSG000001 | SHFL | 6.82E-01 | 8.84E-01 | 9.78E-01 | -3.21E-02 | -1.02E+00 |
| ENSG00000C | ITIH4 | 6.82E-01 | 8.84E-01 | 9.36E-01 | -9.56E-02 | -1.07E+00 |
| ENSG000002 | --- | 6.82E-01 | ? | 8.49E-01 | -2.36E-01 | -1.18E+00 |
| ENSG000001 | CCSER2 | 6.82E-01 | 8.84E-01 | 9.83E-01 | -2.54E-02 | -1.02E+00 |
| ENSG000002 | MIR762HG | 6.82E-01 | ? | 1.09E+00 | 1.24E-01 | 1.09E+00 |
| ENSG000001 | KCTD15 | 6.82E-01 | 8.84E-01 | 1.02E+00 | 2.65E-02 | 1.02E+00 |
| ENSG000001 | PLA2G4D | 6.82E-01 | ? | 8.79E-01 | -1.86E-01 | -1.14E+00 |
| ENSG000001 | VTI1B | 6.82E-01 | 8.84E-01 | 9.90E-01 | -1.48E-02 | -1.01E+00 |
| ENSG000002 | --- | 6.82E-01 | 8.84E-01 | 9.34E-01 | -9.84E-02 | -1.07E+00 |
| ENSG000001 | RNF169 | 6.82E-01 | 8.84E-01 | 1.02E+00 | 2.84E-02 | 1.02E+00 |
| ENSG000001 | MTUS2 | 6.82E-01 | ? | 1.24E+00 | 3.06E-01 | 1.24E+00 |
| ENSG000001 | CTXN1 | 6.82E-01 | 8.85E-01 | 1.04E+00 | 5.65E-02 | 1.04E+00 |
| ENSG000002 | PHKA1P1 | 6.82E-01 | ? | 8.69E-01 | -2.03E-01 | -1.15E+00 |
| ENSG000002 | RCCD1-AS1 | 6.82E-01 | ? | 8.86E-01 | -1.75E-01 | -1.13E+00 |
| ENSG000001 | ISM1 | 6.82E-01 | ? | 1.08E+00 | 1.13E-01 | 1.08E+00 |
| ENSG000001 | LIFR | 6.83E-01 | 8.85E-01 | 9.71E-01 | -4.18E-02 | -1.03E+00 |
| ENSG000001 | CDHR3 | 6.83E-01 | ? | 1.10E+00 | 1.31E-01 | 1.10E+00 |
| ENSG000001 | RHCG | 6.83E-01 | 8.85E-01 | 9.58E-01 | -6.21E-02 | -1.04E+00 |
| ENSG000002 | ORAI1 | 6.83E-01 | 8.85E-01 | 1.03E+00 | 3.76E-02 | 1.03E+00 |
| ENSG000002 | --- | 6.83E-01 | ? | 1.12E+00 | 1.69E-01 | 1.12E+00 |

| | | | | | | |
|---|---|---|---|---|---|---|
| ENSG000002 | --- | 6.83E-01 | ? | 1.17E+00 | 2.25E-01 | 1.17E+00 |
| ENSG000002 | EIF5-DT | 6.83E-01 | ? | 8.33E-01 | -2.63E-01 | -1.20E+00 |
| ENSG000001 | ATP5MK | 6.83E-01 | 8.85E-01 | 1.02E+00 | 2.60E-02 | 1.02E+00 |
| ENSG000001 | FAM78A | 6.83E-01 | ? | 1.08E+00 | 1.09E-01 | 1.08E+00 |
| ENSG000001 | PCDHB2 | 6.83E-01 | 8.85E-01 | 9.09E-01 | -1.37E-01 | -1.10E+00 |
| ENSG000002 | --- | 6.83E-01 | 8.85E-01 | 9.14E-01 | -1.29E-01 | -1.09E+00 |
| ENSG00000C | SLC7A9 | 6.83E-01 | ? | 1.14E+00 | 1.95E-01 | 1.14E+00 |
| ENSG000001 | LHFPL1 | 6.83E-01 | ? | 8.82E-01 | -1.81E-01 | -1.13E+00 |
| ENSG00000C | ANKRD44 | 6.83E-01 | 8.85E-01 | 9.56E-01 | -6.51E-02 | -1.05E+00 |
| ENSG000002 | RPS4XP16 | 6.83E-01 | ? | 1.08E+00 | 1.15E-01 | 1.08E+00 |
| ENSG000002 | --- | 6.83E-01 | ? | 1.09E+00 | 1.22E-01 | 1.09E+00 |
| ENSG000002 | --- | 6.83E-01 | ? | 8.55E-01 | -2.27E-01 | -1.17E+00 |
| ENSG000001 | DNA2 | 6.83E-01 | 8.85E-01 | 1.04E+00 | 4.96E-02 | 1.04E+00 |
| ENSG000001 | KANSL2 | 6.83E-01 | 8.85E-01 | 1.02E+00 | 2.94E-02 | 1.02E+00 |
| ENSG000002 | --- | 6.84E-01 | ? | 1.09E+00 | 1.25E-01 | 1.09E+00 |
| ENSG000001 | CA9 | 6.84E-01 | 8.85E-01 | 1.04E+00 | 5.51E-02 | 1.04E+00 |
| ENSG000002 | --- | 6.84E-01 | ? | 1.23E+00 | 2.96E-01 | 1.23E+00 |
| ENSG000001 | PABIR3 | 6.84E-01 | ? | 1.07E+00 | 9.31E-02 | 1.07E+00 |
| ENSG00000C | POMGNT1 | 6.84E-01 | 8.86E-01 | 1.02E+00 | 2.46E-02 | 1.02E+00 |
| ENSG000002 | FAM223A | 6.84E-01 | ? | 8.53E-01 | -2.30E-01 | -1.17E+00 |
| ENSG000002 | GIGYF2 | 6.84E-01 | 8.86E-01 | 9.87E-01 | -1.94E-02 | -1.01E+00 |
| ENSG000001 | DUSP28 | 6.84E-01 | 8.86E-01 | 1.05E+00 | 7.08E-02 | 1.05E+00 |
| ENSG000002 | --- | 6.84E-01 | ? | 9.08E-01 | -1.39E-01 | -1.10E+00 |
| ENSG000001 | WDR45B | 6.84E-01 | 8.86E-01 | 9.88E-01 | -1.74E-02 | -1.01E+00 |
| ENSG000001 | LRSAM1 | 6.84E-01 | 8.86E-01 | 9.82E-01 | -2.69E-02 | -1.02E+00 |
| ENSG000002 | --- | 6.84E-01 | ? | 9.03E-01 | -1.48E-01 | -1.11E+00 |
| ENSG000002 | MRPS35-DT | 6.84E-01 | ? | 1.22E+00 | 2.81E-01 | 1.22E+00 |
| ENSG000002 | --- | 6.84E-01 | ? | 1.11E+00 | 1.50E-01 | 1.11E+00 |
| ENSG00000C | REXO1 | 6.85E-01 | 8.86E-01 | 1.03E+00 | 3.86E-02 | 1.03E+00 |
| ENSG000001 | BEX2 | 6.85E-01 | 8.86E-01 | 1.05E+00 | 7.18E-02 | 1.05E+00 |
| ENSG000001 | MLLT1 | 6.85E-01 | 8.86E-01 | 1.02E+00 | 2.83E-02 | 1.02E+00 |
| ENSG000001 | MFSD6 | 6.85E-01 | 8.86E-01 | 9.83E-01 | -2.46E-02 | -1.02E+00 |
| ENSG000002 | BMPR1AP2 | 6.85E-01 | ? | 1.16E+00 | 2.11E-01 | 1.16E+00 |
| ENSG000001 | CYSRT1 | 6.85E-01 | ? | 8.86E-01 | -1.74E-01 | -1.13E+00 |
| ENSG000001 | TTC13 | 6.85E-01 | 8.86E-01 | 9.74E-01 | -3.84E-02 | -1.03E+00 |
| ENSG000002 | ISY1-RAB43 | 6.85E-01 | 8.86E-01 | 9.82E-01 | -2.67E-02 | -1.02E+00 |
| ENSG000001 | GATAD1 | 6.85E-01 | 8.86E-01 | 9.84E-01 | -2.26E-02 | -1.02E+00 |
| ENSG00000C | OSGEP | 6.85E-01 | 8.86E-01 | 1.02E+00 | 3.25E-02 | 1.02E+00 |
| ENSG000001 | CKAP2 | 6.85E-01 | 8.86E-01 | 9.80E-01 | -2.90E-02 | -1.02E+00 |
| ENSG000001 | NCOA3 | 6.85E-01 | 8.86E-01 | 1.01E+00 | 2.00E-02 | 1.01E+00 |
| ENSG000001 | TCEAL3 | 6.85E-01 | 8.86E-01 | 1.03E+00 | 3.96E-02 | 1.03E+00 |
| ENSG000002 | BICDL3P | 6.86E-01 | ? | 8.36E-01 | -2.59E-01 | -1.20E+00 |
| ENSG00000C | ATP1B3 | 6.86E-01 | 8.87E-01 | 1.01E+00 | 1.98E-02 | 1.01E+00 |
| ENSG000002 | FAM238C | 6.86E-01 | ? | 9.15E-01 | -1.28E-01 | -1.09E+00 |
| ENSG000002 | YWHAZP5 | 6.86E-01 | ? | 9.08E-01 | -1.39E-01 | -1.10E+00 |

| | | | | | | |
|---|---|---|---|---|---|---|
| ENSG000002 | --- | 6.86E-01 | ? | 8.78E-01 | -1.87E-01 | -1.14E+00 |
| ENSG000001 | B3GNTL1 | 6.86E-01 | 8.87E-01 | 1.06E+00 | 8.18E-02 | 1.06E+00 |
| ENSG000001 | LINC02873 | 6.86E-01 | ? | 1.16E+00 | 2.19E-01 | 1.16E+00 |
| ENSG000002 | HNRNPCP7 | 6.86E-01 | ? | 1.13E+00 | 1.72E-01 | 1.13E+00 |
| ENSG000002 | --- | 6.86E-01 | 8.87E-01 | 9.42E-01 | -8.55E-02 | -1.06E+00 |
| ENSG00000C | MAP2K7 | 6.86E-01 | 8.87E-01 | 9.78E-01 | -3.24E-02 | -1.02E+00 |
| ENSG00000C | ZNF343 | 6.86E-01 | 8.87E-01 | 9.85E-01 | -2.23E-02 | -1.02E+00 |
| ENSG000002 | CCDC163 | 6.86E-01 | ? | 9.37E-01 | -9.46E-02 | -1.07E+00 |
| ENSG000002 | MRPL46 | 6.86E-01 | 8.87E-01 | 1.02E+00 | 2.92E-02 | 1.02E+00 |
| ENSG000002 | --- | 6.86E-01 | 8.87E-01 | 9.69E-01 | -4.62E-02 | -1.03E+00 |
| ENSG00000C | RAD54L | 6.86E-01 | 8.87E-01 | 9.18E-01 | -1.24E-01 | -1.09E+00 |
| ENSG000001 | ZC3H12C | 6.87E-01 | 8.87E-01 | 9.77E-01 | -3.39E-02 | -1.02E+00 |
| ENSG000001 | FKRP | 6.87E-01 | 8.87E-01 | 9.78E-01 | -3.18E-02 | -1.02E+00 |
| ENSG000001 | LRBA | 6.87E-01 | 8.87E-01 | 9.82E-01 | -2.64E-02 | -1.02E+00 |
| ENSG000001 | CSF1 | 6.87E-01 | 8.87E-01 | 1.02E+00 | 3.25E-02 | 1.02E+00 |
| ENSG000001 | UPK2 | 6.87E-01 | ? | 1.11E+00 | 1.52E-01 | 1.11E+00 |
| ENSG000002 | --- | 6.87E-01 | ? | 9.07E-01 | -1.41E-01 | -1.10E+00 |
| ENSG000002 | --- | 6.87E-01 | ? | 1.08E+00 | 1.15E-01 | 1.08E+00 |
| ENSG000001 | STRBP | 6.87E-01 | 8.87E-01 | 9.82E-01 | -2.68E-02 | -1.02E+00 |
| ENSG000001 | SEMA4D | 6.87E-01 | 8.87E-01 | 9.67E-01 | -4.78E-02 | -1.03E+00 |
| ENSG000001 | FRMPD4 | 6.87E-01 | 8.87E-01 | 1.05E+00 | 6.98E-02 | 1.05E+00 |
| ENSG000001 | CCDC85B | 6.87E-01 | 8.87E-01 | 1.06E+00 | 8.89E-02 | 1.06E+00 |
| ENSG000001 | CFDP1 | 6.87E-01 | 8.87E-01 | 9.83E-01 | -2.51E-02 | -1.02E+00 |
| ENSG000002 | --- | 6.87E-01 | ? | 1.13E+00 | 1.71E-01 | 1.13E+00 |
| ENSG000001 | ALKBH4 | 6.87E-01 | 8.88E-01 | 9.68E-01 | -4.64E-02 | -1.03E+00 |
| ENSG000001 | CSPG5 | 6.88E-01 | ? | 9.07E-01 | -1.41E-01 | -1.10E+00 |
| ENSG000002 | TTC41P | 6.88E-01 | 8.88E-01 | 9.40E-01 | -8.95E-02 | -1.06E+00 |
| ENSG000001 | GLIPR2 | 6.88E-01 | 8.88E-01 | 9.51E-01 | -7.26E-02 | -1.05E+00 |
| ENSG000001 | ROR1 | 6.88E-01 | 8.88E-01 | 9.65E-01 | -5.13E-02 | -1.04E+00 |
| ENSG00000C | MAP2K4 | 6.88E-01 | 8.88E-01 | 1.01E+00 | 1.52E-02 | 1.01E+00 |
| ENSG000001 | PDE4B | 6.88E-01 | ? | 8.55E-01 | -2.25E-01 | -1.17E+00 |
| ENSG000002 | SLC9A3-AS1 | 6.88E-01 | 8.88E-01 | 1.05E+00 | 6.92E-02 | 1.05E+00 |
| ENSG000001 | NOMO1 | 6.88E-01 | 8.88E-01 | 1.01E+00 | 1.88E-02 | 1.01E+00 |
| ENSG000001 | PXMP2 | 6.88E-01 | 8.88E-01 | 1.06E+00 | 8.16E-02 | 1.06E+00 |
| ENSG000001 | NOPCHAP1 | 6.88E-01 | 8.88E-01 | 1.04E+00 | 5.15E-02 | 1.04E+00 |
| ENSG00000C | VEZT | 6.88E-01 | 8.88E-01 | 9.83E-01 | -2.48E-02 | -1.02E+00 |
| ENSG000001 | OAZ2 | 6.88E-01 | 8.88E-01 | 1.02E+00 | 2.52E-02 | 1.02E+00 |
| ENSG000001 | CEP170P1 | 6.88E-01 | ? | 1.21E+00 | 2.78E-01 | 1.21E+00 |
| ENSG000001 | ZFP36L2 | 6.88E-01 | 8.88E-01 | 9.80E-01 | -2.97E-02 | -1.02E+00 |
| ENSG000001 | ZMYM1 | 6.88E-01 | 8.88E-01 | 9.81E-01 | -2.83E-02 | -1.02E+00 |
| ENSG00000C | ZNF280C | 6.88E-01 | 8.88E-01 | 9.74E-01 | -3.77E-02 | -1.03E+00 |
| ENSG00000C | RPUSD1 | 6.88E-01 | 8.88E-01 | 1.03E+00 | 4.92E-02 | 1.03E+00 |
| ENSG000002 | SMC3P1 | 6.89E-01 | ? | 1.16E+00 | 2.15E-01 | 1.16E+00 |
| ENSG000002 | --- | 6.89E-01 | 8.88E-01 | 1.03E+00 | 4.68E-02 | 1.03E+00 |
| ENSG00000C | SPAST | 6.89E-01 | 8.88E-01 | 9.79E-01 | -3.03E-02 | -1.02E+00 |

| | | | | | | |
|---|---|---|---|---|---|---|
| ENSG000001 | FCRLB | 6.89E-01 | ? | 1.09E+00 | 1.22E-01 | 1.09E+00 |
| ENSG00000C | ZNF112 | 6.89E-01 | 8.88E-01 | 9.74E-01 | -3.80E-02 | -1.03E+00 |
| ENSG000001 | CHRM5 | 6.89E-01 | ? | 8.33E-01 | -2.64E-01 | -1.20E+00 |
| ENSG00000C | TP53BP1 | 6.89E-01 | 8.89E-01 | 9.89E-01 | -1.62E-02 | -1.01E+00 |
| ENSG000002 | --- | 6.89E-01 | ? | 1.24E+00 | 3.05E-01 | 1.24E+00 |
| ENSG000002 | --- | 6.89E-01 | ? | 1.09E+00 | 1.27E-01 | 1.09E+00 |
| ENSG000002 | TNFRSF10A- | 6.90E-01 | 8.89E-01 | 9.72E-01 | -4.05E-02 | -1.03E+00 |
| ENSG000001 | LMX1B | 6.90E-01 | ? | 1.20E+00 | 2.66E-01 | 1.20E+00 |
| ENSG000001 | CADM4 | 6.90E-01 | 8.89E-01 | 9.66E-01 | -4.95E-02 | -1.03E+00 |
| ENSG000001 | PCDHB3 | 6.90E-01 | ? | 1.17E+00 | 2.29E-01 | 1.17E+00 |
| ENSG000001 | UBE2G1 | 6.90E-01 | 8.89E-01 | 9.87E-01 | -1.89E-02 | -1.01E+00 |
| ENSG000001 | MEIS3P2 | 6.90E-01 | ? | 1.11E+00 | 1.55E-01 | 1.11E+00 |
| ENSG000001 | STARD9 | 6.90E-01 | 8.89E-01 | 9.70E-01 | -4.44E-02 | -1.03E+00 |
| ENSG000002 | SMIM30 | 6.90E-01 | 8.89E-01 | 9.82E-01 | -2.56E-02 | -1.02E+00 |
| ENSG000001 | PDE9A | 6.90E-01 | 8.89E-01 | 9.46E-01 | -8.08E-02 | -1.06E+00 |
| ENSG000002 | --- | 6.90E-01 | ? | 9.12E-01 | -1.32E-01 | -1.10E+00 |
| ENSG000001 | GALNT3 | 6.90E-01 | 8.89E-01 | 1.02E+00 | 2.48E-02 | 1.02E+00 |
| ENSG000001 | CERS2 | 6.90E-01 | 8.89E-01 | 9.86E-01 | -2.07E-02 | -1.01E+00 |
| ENSG000002 | TP53TG3D | 6.90E-01 | ? | 8.53E-01 | -2.30E-01 | -1.17E+00 |
| ENSG000001 | DPY30 | 6.90E-01 | 8.89E-01 | 9.82E-01 | -2.55E-02 | -1.02E+00 |
| ENSG000001 | CHCHD7 | 6.90E-01 | 8.89E-01 | 9.86E-01 | -2.10E-02 | -1.01E+00 |
| ENSG000002 | MIAT | 6.90E-01 | ? | 1.09E+00 | 1.24E-01 | 1.09E+00 |
| ENSG000001 | SLC2A8 | 6.90E-01 | 8.89E-01 | 1.04E+00 | 5.60E-02 | 1.04E+00 |
| ENSG000002 | NPM1P6 | 6.90E-01 | ? | 1.19E+00 | 2.56E-01 | 1.19E+00 |
| ENSG000001 | RBPMS | 6.90E-01 | 8.89E-01 | 9.82E-01 | -2.60E-02 | -1.02E+00 |
| ENSG000001 | OVCH2 | 6.90E-01 | 8.89E-01 | 9.46E-01 | -8.04E-02 | -1.06E+00 |
| ENSG00000C | RABL2B | 6.90E-01 | 8.89E-01 | 9.85E-01 | -2.21E-02 | -1.02E+00 |
| ENSG000001 | PRXL2B | 6.90E-01 | 8.89E-01 | 9.79E-01 | -3.03E-02 | -1.02E+00 |
| ENSG000002 | WAPL-DT | 6.91E-01 | ? | 9.35E-01 | -9.71E-02 | -1.07E+00 |
| ENSG000001 | YY1AP1 | 6.91E-01 | 8.89E-01 | 9.88E-01 | -1.79E-02 | -1.01E+00 |
| ENSG000002 | SMIM31 | 6.91E-01 | ? | 7.92E-01 | -3.37E-01 | -1.26E+00 |
| ENSG000001 | CATSPERB | 6.91E-01 | 8.89E-01 | 1.05E+00 | 7.66E-02 | 1.05E+00 |
| ENSG000001 | CDKL2 | 6.91E-01 | 8.89E-01 | 9.65E-01 | -5.15E-02 | -1.04E+00 |
| ENSG000002 | --- | 6.91E-01 | ? | 8.01E-01 | -3.20E-01 | -1.25E+00 |
| ENSG000001 | THBS4 | 6.91E-01 | ? | 8.85E-01 | -1.76E-01 | -1.13E+00 |
| ENSG000002 | --- | 6.91E-01 | ? | 8.95E-01 | -1.60E-01 | -1.12E+00 |
| ENSG000001 | CARHSP1 | 6.91E-01 | 8.89E-01 | 9.78E-01 | -3.19E-02 | -1.02E+00 |
| ENSG000001 | LRP1 | 6.91E-01 | 8.89E-01 | 9.79E-01 | -3.11E-02 | -1.02E+00 |
| ENSG000001 | BST1 | 6.91E-01 | ? | 9.16E-01 | -1.26E-01 | -1.09E+00 |
| ENSG000002 | --- | 6.91E-01 | ? | 1.15E+00 | 1.99E-01 | 1.15E+00 |
| ENSG00000C | ZNF213 | 6.91E-01 | 8.89E-01 | 1.03E+00 | 4.30E-02 | 1.03E+00 |
| ENSG000001 | DNAJC22 | 6.91E-01 | ? | 9.20E-01 | -1.20E-01 | -1.09E+00 |
| ENSG000001 | PDE11A | 6.91E-01 | ? | 1.08E+00 | 1.12E-01 | 1.08E+00 |
| ENSG000002 | ERVE-1 | 6.91E-01 | ? | 8.80E-01 | -1.84E-01 | -1.14E+00 |
| ENSG000002 | --- | 6.91E-01 | ? | 1.10E+00 | 1.42E-01 | 1.10E+00 |

| | | | | | | |
|---|---|---|---|---|---|---|
| ENSG000001 | DRAXIN | 6.91E-01 | ? | 1.13E+00 | 1.83E-01 | 1.13E+00 |
| ENSG000001 | UNC93B1 | 6.92E-01 | 8.90E-01 | 1.03E+00 | 4.00E-02 | 1.03E+00 |
| ENSG000002 | AACSP1 | 6.92E-01 | ? | 1.19E+00 | 2.49E-01 | 1.19E+00 |
| ENSG000001 | TMEM143 | 6.92E-01 | 8.90E-01 | 9.57E-01 | -6.36E-02 | -1.05E+00 |
| ENSG000001 | AHSA2P | 6.92E-01 | 8.90E-01 | 9.66E-01 | -4.93E-02 | -1.03E+00 |
| ENSG000001 | RGPD4 | 6.92E-01 | ? | 1.11E+00 | 1.48E-01 | 1.11E+00 |
| ENSG000002 | LINC02240 | 6.92E-01 | ? | 1.20E+00 | 2.65E-01 | 1.20E+00 |
| ENSG000002 | --- | 6.92E-01 | ? | 8.26E-01 | -2.75E-01 | -1.21E+00 |
| ENSG000001 | RPL28 | 6.92E-01 | 8.90E-01 | 1.02E+00 | 2.79E-02 | 1.02E+00 |
| ENSG000001 | APPL1 | 6.92E-01 | 8.90E-01 | 1.01E+00 | 1.83E-02 | 1.01E+00 |
| ENSG000001 | KIAA1671 | 6.92E-01 | 8.90E-01 | 9.87E-01 | -1.89E-02 | -1.01E+00 |
| ENSG000002 | --- | 6.92E-01 | ? | 8.68E-01 | -2.04E-01 | -1.15E+00 |
| ENSG000002 | --- | 6.92E-01 | ? | 9.21E-01 | -1.18E-01 | -1.09E+00 |
| ENSG000002 | WTAPP1 | 6.92E-01 | ? | 8.96E-01 | -1.59E-01 | -1.12E+00 |
| ENSG000002 | NOL12 | 6.92E-01 | 8.90E-01 | 1.02E+00 | 3.51E-02 | 1.02E+00 |
| ENSG000002 | LINC02989 | 6.92E-01 | ? | 1.09E+00 | 1.23E-01 | 1.09E+00 |
| ENSG000001 | UGT2B17 | 6.92E-01 | ? | 8.25E-01 | -2.78E-01 | -1.21E+00 |
| ENSG000001 | C7orf57 | 6.92E-01 | ? | 1.19E+00 | 2.56E-01 | 1.19E+00 |
| ENSG00000C | COASY | 6.93E-01 | 8.90E-01 | 9.83E-01 | -2.45E-02 | -1.02E+00 |
| ENSG000002 | --- | 6.93E-01 | ? | 8.79E-01 | -1.87E-01 | -1.14E+00 |
| ENSG000002 | FNTB | 6.93E-01 | 8.91E-01 | 1.02E+00 | 3.24E-02 | 1.02E+00 |
| ENSG000002 | --- | 6.93E-01 | 8.91E-01 | 9.58E-01 | -6.22E-02 | -1.04E+00 |
| ENSG000001 | BTG1 | 6.93E-01 | 8.91E-01 | 9.82E-01 | -2.60E-02 | -1.02E+00 |
| ENSG000001 | LINC02878 | 6.93E-01 | ? | 1.09E+00 | 1.19E-01 | 1.09E+00 |
| ENSG000001 | KIF13B | 6.93E-01 | 8.91E-01 | 1.02E+00 | 2.73E-02 | 1.02E+00 |
| ENSG000001 | FRAT2 | 6.93E-01 | 8.91E-01 | 1.03E+00 | 3.82E-02 | 1.03E+00 |
| ENSG000002 | --- | 6.93E-01 | 8.91E-01 | 9.50E-01 | -7.35E-02 | -1.05E+00 |
| ENSG000001 | RPL12P4 | 6.93E-01 | ? | 1.08E+00 | 1.16E-01 | 1.08E+00 |
| ENSG00000C | RNF216 | 6.93E-01 | 8.91E-01 | 1.01E+00 | 1.68E-02 | 1.01E+00 |
| ENSG000002 | ULK4P1 | 6.93E-01 | ? | 1.17E+00 | 2.24E-01 | 1.17E+00 |
| ENSG000001 | TATDN1 | 6.93E-01 | 8.91E-01 | 1.01E+00 | 2.12E-02 | 1.01E+00 |
| ENSG000002 | FOXN3-AS1 | 6.93E-01 | 8.91E-01 | 1.08E+00 | 1.13E-01 | 1.08E+00 |
| ENSG000001 | SIK2 | 6.93E-01 | 8.91E-01 | 9.87E-01 | -1.96E-02 | -1.01E+00 |
| ENSG000002 | RPL23AP42 | 6.93E-01 | 8.91E-01 | 1.02E+00 | 3.48E-02 | 1.02E+00 |
| ENSG000001 | CXCL5 | 6.94E-01 | 8.91E-01 | 9.24E-01 | -1.15E-01 | -1.08E+00 |
| ENSG000002 | --- | 6.94E-01 | ? | 8.54E-01 | -2.27E-01 | -1.17E+00 |
| ENSG00000C | MAPKAPK5 | 6.94E-01 | 8.91E-01 | 9.87E-01 | -1.91E-02 | -1.01E+00 |
| ENSG000002 | NEAT1 | 6.94E-01 | 8.91E-01 | 9.55E-01 | -6.60E-02 | -1.05E+00 |
| ENSG00000C | ABCA7 | 6.94E-01 | 8.91E-01 | 9.72E-01 | -4.14E-02 | -1.03E+00 |
| ENSG000001 | MAP1S | 6.94E-01 | 8.91E-01 | 9.66E-01 | -5.06E-02 | -1.04E+00 |
| ENSG000001 | ARL6IP4 | 6.94E-01 | 8.91E-01 | 9.89E-01 | -1.64E-02 | -1.01E+00 |
| ENSG000001 | UFD1-AS1 | 6.94E-01 | ? | 9.31E-01 | -1.03E-01 | -1.07E+00 |
| ENSG000002 | KLC1-AS1 | 6.94E-01 | 8.91E-01 | 9.68E-01 | -4.63E-02 | -1.03E+00 |
| ENSG000002 | GOLGA8K | 6.94E-01 | ? | 9.09E-01 | -1.37E-01 | -1.10E+00 |
| ENSG000001 | KDM3A | 6.94E-01 | 8.91E-01 | 1.02E+00 | 2.25E-02 | 1.02E+00 |

| | | | | | | | |
|---|---|---|---|---|---|---|---|
| ENSG000001 | TXNL4B | 6.94E-01 | | 8.91E-01 | 1.02E+00 | 2.98E-02 | 1.02E+00 |
| ENSG000002 | --- | 6.94E-01 | ? | | 1.08E+00 | 1.13E-01 | 1.08E+00 |
| ENSG000001 | SEPSECS | 6.94E-01 | | 8.91E-01 | 1.02E+00 | 2.97E-02 | 1.02E+00 |
| ENSG00000C | CACNG4 | 6.95E-01 | ? | | 1.16E+00 | 2.18E-01 | 1.16E+00 |
| ENSG000001 | FLYWCH2 | 6.95E-01 | | 8.91E-01 | 1.03E+00 | 4.34E-02 | 1.03E+00 |
| ENSG000001 | PFN4 | 6.95E-01 | ? | | 8.93E-01 | -1.63E-01 | -1.12E+00 |
| ENSG000001 | TM9SF1 | 6.95E-01 | | 8.91E-01 | 1.03E+00 | 3.73E-02 | 1.03E+00 |
| ENSG000001 | CIB1 | 6.95E-01 | | 8.91E-01 | 1.03E+00 | 3.86E-02 | 1.03E+00 |
| ENSG000002 | --- | 6.95E-01 | ? | | 1.11E+00 | 1.52E-01 | 1.11E+00 |
| ENSG000002 | LINC00911 | 6.95E-01 | ? | | 1.20E+00 | 2.65E-01 | 1.20E+00 |
| ENSG000001 | MSRA | 6.95E-01 | | 8.91E-01 | 1.03E+00 | 4.21E-02 | 1.03E+00 |
| ENSG000001 | KLHDC4 | 6.95E-01 | | 8.91E-01 | 1.02E+00 | 2.49E-02 | 1.02E+00 |
| ENSG000001 | GLB1L3 | 6.95E-01 | | 8.91E-01 | 9.69E-01 | -4.53E-02 | -1.03E+00 |
| ENSG000001 | EGFL7 | 6.95E-01 | | 8.91E-01 | 1.04E+00 | 5.74E-02 | 1.04E+00 |
| ENSG000001 | MGMT | 6.95E-01 | | 8.91E-01 | 1.04E+00 | 5.81E-02 | 1.04E+00 |
| ENSG000002 | AURKAP1 | 6.95E-01 | ? | | 8.12E-01 | -3.00E-01 | -1.23E+00 |
| ENSG000001 | SELENOK | 6.95E-01 | | 8.91E-01 | 9.83E-01 | -2.43E-02 | -1.02E+00 |
| ENSG000002 | --- | 6.95E-01 | ? | | 8.42E-01 | -2.48E-01 | -1.19E+00 |
| ENSG000001 | DNALI1 | 6.95E-01 | | 8.91E-01 | 9.46E-01 | -7.97E-02 | -1.06E+00 |
| ENSG00000C | MID2 | 6.95E-01 | | 8.91E-01 | 9.83E-01 | -2.43E-02 | -1.02E+00 |
| ENSG000001 | CLDN4 | 6.95E-01 | | 8.91E-01 | 1.06E+00 | 8.14E-02 | 1.06E+00 |
| ENSG000001 | ZNF132 | 6.95E-01 | | 8.91E-01 | 1.03E+00 | 4.46E-02 | 1.03E+00 |
| ENSG00000C | RASSF1 | 6.95E-01 | | 8.91E-01 | 1.02E+00 | 3.15E-02 | 1.02E+00 |
| ENSG000001 | ATP6V1B2 | 6.95E-01 | | 8.91E-01 | 1.02E+00 | 2.52E-02 | 1.02E+00 |
| ENSG000002 | --- | 6.95E-01 | ? | | 8.79E-01 | -1.85E-01 | -1.14E+00 |
| ENSG000002 | ADAM20P1 | 6.95E-01 | ? | | 7.96E-01 | -3.30E-01 | -1.26E+00 |
| ENSG000002 | --- | 6.95E-01 | ? | | 1.17E+00 | 2.31E-01 | 1.17E+00 |
| ENSG000002 | --- | 6.95E-01 | ? | | 9.35E-01 | -9.64E-02 | -1.07E+00 |
| ENSG000001 | HNRNPAB | 6.95E-01 | | 8.91E-01 | 1.01E+00 | 1.65E-02 | 1.01E+00 |
| ENSG000001 | --- | 6.96E-01 | ? | | 1.19E+00 | 2.45E-01 | 1.19E+00 |
| ENSG000001 | RAB5IF | 6.96E-01 | | 8.91E-01 | 9.80E-01 | -2.95E-02 | -1.02E+00 |
| ENSG000001 | LYL1 | 6.96E-01 | ? | | 1.17E+00 | 2.22E-01 | 1.17E+00 |
| ENSG000001 | TRIM5 | 6.96E-01 | | 8.91E-01 | 1.01E+00 | 2.04E-02 | 1.01E+00 |
| ENSG000002 | HMGN1P38 | 6.96E-01 | | 8.91E-01 | 9.49E-01 | -7.59E-02 | -1.05E+00 |
| ENSG000001 | MIER2 | 6.96E-01 | | 8.91E-01 | 1.02E+00 | 2.45E-02 | 1.02E+00 |
| ENSG000001 | PRKAB1 | 6.96E-01 | | 8.91E-01 | 1.01E+00 | 2.02E-02 | 1.01E+00 |
| ENSG000002 | --- | 6.96E-01 | | 8.91E-01 | 9.59E-01 | -6.01E-02 | -1.04E+00 |
| ENSG000001 | RABGAP1L | 6.96E-01 | | 8.91E-01 | 9.83E-01 | -2.46E-02 | -1.02E+00 |
| ENSG000001 | TSLP | 6.96E-01 | | 8.92E-01 | 1.05E+00 | 7.27E-02 | 1.05E+00 |
| ENSG000002 | RPS26P6 | 6.96E-01 | ? | | 1.23E+00 | 2.93E-01 | 1.23E+00 |
| ENSG000001 | VPS26C | 6.96E-01 | | 8.92E-01 | 1.01E+00 | 1.96E-02 | 1.01E+00 |
| ENSG000002 | --- | 6.96E-01 | ? | | 1.10E+00 | 1.32E-01 | 1.10E+00 |
| ENSG000001 | NAA15 | 6.96E-01 | | 8.92E-01 | 9.89E-01 | -1.60E-02 | -1.01E+00 |
| ENSG000002 | ETV5 | 6.97E-01 | | 8.92E-01 | 1.01E+00 | 1.70E-02 | 1.01E+00 |
| ENSG000002 | ENTPD3-AS1 | 6.97E-01 | | 8.92E-01 | 1.06E+00 | 8.12E-02 | 1.06E+00 |

| | | | | | | |
|---|---|---|---|---|---|---|
| ENSG000001 | SMG1 | 6.97E-01 | 8.92E-01 | 9.74E-01 | -3.82E-02 | -1.03E+00 |
| ENSG000001 | AXIN1 | 6.97E-01 | 8.92E-01 | 9.83E-01 | -2.42E-02 | -1.02E+00 |
| ENSG000002 | MIR34BHG | 6.97E-01 | ? | 1.16E+00 | 2.14E-01 | 1.16E+00 |
| ENSG000002 | CSRP1-AS1 | 6.97E-01 | ? | 9.11E-01 | -1.34E-01 | -1.10E+00 |
| ENSG00000C | RTN4R | 6.97E-01 | 8.92E-01 | 1.05E+00 | 7.07E-02 | 1.05E+00 |
| ENSG000001 | ANKLE1 | 6.97E-01 | ? | 9.26E-01 | -1.11E-01 | -1.08E+00 |
| ENSG000001 | LIF | 6.97E-01 | 8.92E-01 | 1.03E+00 | 4.83E-02 | 1.03E+00 |
| ENSG000002 | BCDIN3D-AS | 6.97E-01 | 8.92E-01 | 1.06E+00 | 8.53E-02 | 1.06E+00 |
| ENSG000002 | --- | 6.97E-01 | 8.92E-01 | 1.06E+00 | 8.61E-02 | 1.06E+00 |
| ENSG000001 | TERF2 | 6.97E-01 | 8.92E-01 | 9.87E-01 | -1.94E-02 | -1.01E+00 |
| ENSG000001 | JMJD8 | 6.97E-01 | 8.92E-01 | 9.78E-01 | -3.20E-02 | -1.02E+00 |
| ENSG000002 | RPL37A-DT | 6.97E-01 | ? | 1.09E+00 | 1.26E-01 | 1.09E+00 |
| ENSG000002 | --- | 6.97E-01 | ? | 7.76E-01 | -3.65E-01 | -1.29E+00 |
| ENSG000002 | --- | 6.97E-01 | ? | 1.12E+00 | 1.67E-01 | 1.12E+00 |
| ENSG00000C | MIPEP | 6.98E-01 | 8.92E-01 | 1.02E+00 | 3.15E-02 | 1.02E+00 |
| ENSG000002 | --- | 6.98E-01 | 8.92E-01 | 9.45E-01 | -8.18E-02 | -1.06E+00 |
| ENSG000001 | RWDD4P2 | 6.98E-01 | 8.92E-01 | 9.53E-01 | -6.88E-02 | -1.05E+00 |
| ENSG000002 | H4C4 | 6.98E-01 | ? | 8.84E-01 | -1.78E-01 | -1.13E+00 |
| ENSG00000C | SNX10 | 6.98E-01 | 8.92E-01 | 9.74E-01 | -3.74E-02 | -1.03E+00 |
| ENSG000002 | --- | 6.98E-01 | 8.92E-01 | 9.35E-01 | -9.68E-02 | -1.07E+00 |
| ENSG000001 | LDLRAP1 | 6.98E-01 | 8.92E-01 | 1.01E+00 | 2.03E-02 | 1.01E+00 |
| ENSG000001 | SFXN2 | 6.98E-01 | 8.92E-01 | 9.57E-01 | -6.32E-02 | -1.04E+00 |
| ENSG000002 | --- | 6.98E-01 | 8.92E-01 | 9.68E-01 | -4.65E-02 | -1.03E+00 |
| ENSG000002 | --- | 6.98E-01 | ? | 1.19E+00 | 2.52E-01 | 1.19E+00 |
| ENSG000002 | ZNF503-AS2 | 6.98E-01 | 8.92E-01 | 1.04E+00 | 6.26E-02 | 1.04E+00 |
| ENSG000002 | MIR30DHG | 6.98E-01 | 8.92E-01 | 1.05E+00 | 6.59E-02 | 1.05E+00 |
| ENSG000001 | CEP70 | 6.98E-01 | 8.92E-01 | 1.02E+00 | 3.52E-02 | 1.02E+00 |
| ENSG000001 | EXOC2 | 6.98E-01 | 8.92E-01 | 1.01E+00 | 1.25E-02 | 1.01E+00 |
| ENSG000001 | EGLN3 | 6.99E-01 | 8.92E-01 | 1.02E+00 | 3.10E-02 | 1.02E+00 |
| ENSG000001 | RTCB | 6.99E-01 | 8.92E-01 | 1.01E+00 | 1.54E-02 | 1.01E+00 |
| ENSG000002 | --- | 6.99E-01 | ? | 1.12E+00 | 1.66E-01 | 1.12E+00 |
| ENSG000001 | ASB8 | 6.99E-01 | 8.92E-01 | 9.85E-01 | -2.19E-02 | -1.02E+00 |
| ENSG000002 | CERNA1 | 6.99E-01 | ? | 1.07E+00 | 9.19E-02 | 1.07E+00 |
| ENSG000001 | DNER | 6.99E-01 | ? | 9.24E-01 | -1.13E-01 | -1.08E+00 |
| ENSG000001 | AMDHD1 | 6.99E-01 | ? | 8.58E-01 | -2.21E-01 | -1.17E+00 |
| ENSG000001 | CFAP298 | 6.99E-01 | 8.93E-01 | 9.84E-01 | -2.33E-02 | -1.02E+00 |
| ENSG000001 | ALOX12 | 6.99E-01 | ? | 1.07E+00 | 1.00E-01 | 1.07E+00 |
| ENSG000002 | RPS7P10 | 6.99E-01 | ? | 1.13E+00 | 1.70E-01 | 1.13E+00 |
| ENSG000001 | TCEAL4 | 6.99E-01 | 8.93E-01 | 1.02E+00 | 2.93E-02 | 1.02E+00 |
| ENSG00000C | ADD2 | 6.99E-01 | ? | 1.16E+00 | 2.19E-01 | 1.16E+00 |
| ENSG000001 | PTER | 6.99E-01 | 8.93E-01 | 9.81E-01 | -2.77E-02 | -1.02E+00 |
| ENSG000001 | TRPC6 | 6.99E-01 | 8.93E-01 | 1.07E+00 | 1.03E-01 | 1.07E+00 |
| ENSG000001 | SGPP1 | 6.99E-01 | 8.93E-01 | 9.85E-01 | -2.20E-02 | -1.02E+00 |
| ENSG000002 | GCC2-AS1 | 6.99E-01 | ? | 1.20E+00 | 2.58E-01 | 1.20E+00 |
| ENSG000001 | TLR5 | 6.99E-01 | ? | 9.26E-01 | -1.11E-01 | -1.08E+00 |

| | | | | | | |
|---|---|---|---|---|---|---|
| ENSG000001 | DRD2 | 6.99E-01 | ? | 8.63E-01 | -2.13E-01 | -1.16E+00 |
| ENSG000002 | GOLGA6L10 | 6.99E-01 | 8.93E-01 | 9.50E-01 | -7.47E-02 | -1.05E+00 |
| ENSG000002 | HCG15 | 6.99E-01 | ? | 1.19E+00 | 2.48E-01 | 1.19E+00 |
| ENSG000001 | IPO9 | 6.99E-01 | 8.93E-01 | 1.01E+00 | 1.34E-02 | 1.01E+00 |
| ENSG000001 | TNFSF15 | 6.99E-01 | 8.93E-01 | 9.74E-01 | -3.82E-02 | -1.03E+00 |
| ENSG000002 | --- | 6.99E-01 | ? | 1.11E+00 | 1.47E-01 | 1.11E+00 |
| ENSG000001 | PDE6B | 7.00E-01 | ? | 1.20E+00 | 2.63E-01 | 1.20E+00 |
| ENSG000002 | CRYZL2P | 7.00E-01 | 8.93E-01 | 9.73E-01 | -3.97E-02 | -1.03E+00 |
| ENSG000001 | EMC4 | 7.00E-01 | 8.93E-01 | 1.01E+00 | 1.76E-02 | 1.01E+00 |
| ENSG000002 | --- | 7.00E-01 | ? | 9.34E-01 | -9.93E-02 | -1.07E+00 |
| ENSG000001 | CLTA | 7.00E-01 | 8.93E-01 | 1.01E+00 | 2.00E-02 | 1.01E+00 |
| ENSG000001 | HDAC5 | 7.00E-01 | 8.93E-01 | 1.04E+00 | 5.65E-02 | 1.04E+00 |
| ENSG000001 | SCAI | 7.00E-01 | 8.93E-01 | 1.03E+00 | 3.78E-02 | 1.03E+00 |
| ENSG000002 | GJA5 | 7.00E-01 | ? | 1.12E+00 | 1.63E-01 | 1.12E+00 |
| ENSG000002 | LRRC37A6P | 7.00E-01 | ? | 8.29E-01 | -2.71E-01 | -1.21E+00 |
| ENSG000002 | RPL23AP74 | 7.00E-01 | ? | 1.20E+00 | 2.65E-01 | 1.20E+00 |
| ENSG000002 | FSCN1P1 | 7.00E-01 | ? | 8.44E-01 | -2.45E-01 | -1.19E+00 |
| ENSG000001 | SYMPK | 7.00E-01 | 8.93E-01 | 9.88E-01 | -1.79E-02 | -1.01E+00 |
| ENSG000001 | H2AC6 | 7.00E-01 | 8.93E-01 | 9.87E-01 | -1.82E-02 | -1.01E+00 |
| ENSG000002 | ZNF451-AS1 | 7.00E-01 | ? | 8.94E-01 | -1.61E-01 | -1.12E+00 |
| ENSG000001 | MAP3K8 | 7.00E-01 | ? | 9.27E-01 | -1.09E-01 | -1.08E+00 |
| ENSG000001 | CREB5 | 7.00E-01 | 8.93E-01 | 9.74E-01 | -3.85E-02 | -1.03E+00 |
| ENSG000002 | --- | 7.00E-01 | ? | 8.75E-01 | -1.93E-01 | -1.14E+00 |
| ENSG000001 | SHC2 | 7.01E-01 | 8.93E-01 | 9.49E-01 | -7.55E-02 | -1.05E+00 |
| ENSG000002 | --- | 7.01E-01 | ? | 7.73E-01 | -3.72E-01 | -1.29E+00 |
| ENSG000001 | ZDHHC11 | 7.01E-01 | 8.93E-01 | 1.04E+00 | 5.32E-02 | 1.04E+00 |
| ENSG000001 | COQ8B | 7.01E-01 | 8.93E-01 | 1.03E+00 | 3.67E-02 | 1.03E+00 |
| ENSG000002 | GNAS-AS1 | 7.01E-01 | ? | 1.20E+00 | 2.66E-01 | 1.20E+00 |
| ENSG00000C | TSPAN6 | 7.01E-01 | 8.93E-01 | 9.75E-01 | -3.62E-02 | -1.03E+00 |
| ENSG000001 | LRP12 | 7.01E-01 | 8.93E-01 | 1.02E+00 | 2.59E-02 | 1.02E+00 |
| ENSG000001 | FHOD3 | 7.01E-01 | 8.93E-01 | 9.55E-01 | -6.65E-02 | -1.05E+00 |
| ENSG000001 | PRKD3 | 7.01E-01 | 8.93E-01 | 9.87E-01 | -1.84E-02 | -1.01E+00 |
| ENSG000002 | LINC01607 | 7.01E-01 | ? | 1.16E+00 | 2.19E-01 | 1.16E+00 |
| ENSG000001 | CCDC74A | 7.01E-01 | 8.93E-01 | 1.07E+00 | 9.35E-02 | 1.07E+00 |
| ENSG000002 | C4orf46 | 7.01E-01 | 8.93E-01 | 1.02E+00 | 2.98E-02 | 1.02E+00 |
| ENSG000002 | PCDHGB6 | 7.01E-01 | 8.93E-01 | 1.05E+00 | 7.56E-02 | 1.05E+00 |
| ENSG000001 | SPINT1 | 7.01E-01 | 8.93E-01 | 1.02E+00 | 3.10E-02 | 1.02E+00 |
| ENSG000002 | FAM229A | 7.01E-01 | 8.93E-01 | 9.42E-01 | -8.68E-02 | -1.06E+00 |
| ENSG000001 | MBD4 | 7.01E-01 | 8.93E-01 | 9.84E-01 | -2.38E-02 | -1.02E+00 |
| ENSG000002 | MIR181A1HC | 7.01E-01 | ? | 8.43E-01 | -2.46E-01 | -1.19E+00 |
| ENSG000002 | LINC02894 | 7.01E-01 | ? | 8.95E-01 | -1.59E-01 | -1.12E+00 |
| ENSG000001 | NSUN3 | 7.01E-01 | 8.94E-01 | 9.76E-01 | -3.49E-02 | -1.02E+00 |
| ENSG000001 | RPN1 | 7.02E-01 | 8.94E-01 | 1.01E+00 | 1.65E-02 | 1.01E+00 |
| ENSG000001 | ZNF667-AS1 | 7.02E-01 | 8.94E-01 | 9.76E-01 | -3.53E-02 | -1.02E+00 |
| ENSG00000C | TRAPPC3 | 7.02E-01 | 8.94E-01 | 1.01E+00 | 1.43E-02 | 1.01E+00 |

| | | | | | | |
|---|---|---|---|---|---|---|
| ENSG000002 | LINC02076 | 7.02E-01 | ? | 1.10E+00 | 1.38E-01 | 1.10E+00 |
| ENSG000002 | HSPA8P8 | 7.02E-01 | ? | 8.54E-01 | -2.27E-01 | -1.17E+00 |
| ENSG000002 | --- | 7.02E-01 | ? | 9.22E-01 | -1.17E-01 | -1.08E+00 |
| ENSG00000C | POLR3E | 7.02E-01 | 8.94E-01 | 1.01E+00 | 1.82E-02 | 1.01E+00 |
| ENSG000001 | RNF40 | 7.02E-01 | 8.94E-01 | 9.90E-01 | -1.39E-02 | -1.01E+00 |
| ENSG000001 | ESAM | 7.02E-01 | ? | 8.71E-01 | -1.99E-01 | -1.15E+00 |
| ENSG000001 | ATRNL1 | 7.02E-01 | ? | 1.21E+00 | 2.71E-01 | 1.21E+00 |
| ENSG000001 | SLC16A5 | 7.02E-01 | 8.94E-01 | 1.04E+00 | 5.57E-02 | 1.04E+00 |
| ENSG000002 | PRANCR | 7.02E-01 | ? | 9.31E-01 | -1.03E-01 | -1.07E+00 |
| ENSG000001 | TAMM41 | 7.02E-01 | 8.94E-01 | 1.02E+00 | 2.62E-02 | 1.02E+00 |
| ENSG000001 | TBC1D30 | 7.03E-01 | 8.94E-01 | 1.05E+00 | 6.37E-02 | 1.05E+00 |
| ENSG000002 | --- | 7.03E-01 | ? | 1.14E+00 | 1.94E-01 | 1.14E+00 |
| ENSG00000C | DERL3 | 7.03E-01 | ? | 1.14E+00 | 1.88E-01 | 1.14E+00 |
| ENSG000001 | PDIK1L | 7.03E-01 | 8.94E-01 | 9.71E-01 | -4.29E-02 | -1.03E+00 |
| ENSG000002 | MRPL23 | 7.03E-01 | 8.94E-01 | 1.03E+00 | 3.71E-02 | 1.03E+00 |
| ENSG000001 | DNAJC12 | 7.03E-01 | ? | 8.56E-01 | -2.24E-01 | -1.17E+00 |
| ENSG000002 | --- | 7.03E-01 | ? | 1.18E+00 | 2.38E-01 | 1.18E+00 |
| ENSG000002 | VASH1-AS1 | 7.03E-01 | 8.94E-01 | 1.04E+00 | 5.49E-02 | 1.04E+00 |
| ENSG000001 | TSPAN2 | 7.03E-01 | 8.94E-01 | 9.50E-01 | -7.34E-02 | -1.05E+00 |
| ENSG000001 | UTP6 | 7.03E-01 | 8.94E-01 | 9.89E-01 | -1.57E-02 | -1.01E+00 |
| ENSG000001 | ELOVL4 | 7.03E-01 | 8.94E-01 | 1.02E+00 | 3.29E-02 | 1.02E+00 |
| ENSG000001 | LTA4H | 7.03E-01 | 8.94E-01 | 1.01E+00 | 1.45E-02 | 1.01E+00 |
| ENSG000002 | COX10-DT | 7.03E-01 | 8.94E-01 | 9.77E-01 | -3.43E-02 | -1.02E+00 |
| ENSG000001 | IL31RA | 7.03E-01 | 8.94E-01 | 1.03E+00 | 4.09E-02 | 1.03E+00 |
| ENSG000001 | ATP10B | 7.03E-01 | 8.95E-01 | 9.49E-01 | -7.54E-02 | -1.05E+00 |
| ENSG000002 | --- | 7.03E-01 | ? | 1.20E+00 | 2.59E-01 | 1.20E+00 |
| ENSG000001 | PTGR2 | 7.03E-01 | 8.95E-01 | 1.03E+00 | 4.37E-02 | 1.03E+00 |
| ENSG000001 | AQP7P1 | 7.03E-01 | ? | 7.94E-01 | -3.34E-01 | -1.26E+00 |
| ENSG000001 | PIGN | 7.03E-01 | 8.95E-01 | 1.01E+00 | 2.06E-02 | 1.01E+00 |
| ENSG000001 | NOXA1 | 7.03E-01 | ? | 1.18E+00 | 2.40E-01 | 1.18E+00 |
| ENSG000001 | CLPP | 7.04E-01 | 8.95E-01 | 1.02E+00 | 2.62E-02 | 1.02E+00 |
| ENSG000002 | --- | 7.04E-01 | ? | 1.16E+00 | 2.14E-01 | 1.16E+00 |
| ENSG000002 | AARSD1 | 7.04E-01 | 8.95E-01 | 1.02E+00 | 2.93E-02 | 1.02E+00 |
| ENSG000001 | DDHD1 | 7.04E-01 | 8.95E-01 | 9.81E-01 | -2.71E-02 | -1.02E+00 |
| ENSG000002 | TGFBR3L | 7.04E-01 | ? | 1.09E+00 | 1.22E-01 | 1.09E+00 |
| ENSG000002 | SUZ12P1 | 7.04E-01 | ? | 7.95E-01 | -3.31E-01 | -1.26E+00 |
| ENSG000001 | FAM111A | 7.04E-01 | 8.95E-01 | 1.02E+00 | 3.40E-02 | 1.02E+00 |
| ENSG000001 | RBM17 | 7.04E-01 | 8.95E-01 | 9.87E-01 | -1.85E-02 | -1.01E+00 |
| ENSG000002 | --- | 7.04E-01 | 8.95E-01 | 9.29E-01 | -1.06E-01 | -1.08E+00 |
| ENSG000001 | XRRA1 | 7.04E-01 | 8.95E-01 | 1.02E+00 | 2.53E-02 | 1.02E+00 |
| ENSG000002 | --- | 7.04E-01 | ? | 1.15E+00 | 1.99E-01 | 1.15E+00 |
| ENSG000001 | ZGPAT | 7.04E-01 | 8.95E-01 | 1.02E+00 | 2.96E-02 | 1.02E+00 |
| ENSG000002 | --- | 7.04E-01 | ? | 1.09E+00 | 1.21E-01 | 1.09E+00 |
| ENSG000001 | BLZF1 | 7.04E-01 | 8.95E-01 | 9.90E-01 | -1.45E-02 | -1.01E+00 |
| ENSG000001 | SIX4 | 7.04E-01 | 8.95E-01 | 1.02E+00 | 3.19E-02 | 1.02E+00 |

| | | | | | | |
|---|---|---|---|---|---|---|
| ENSG000002 | NAP1L4 | 7.04E-01 | 8.95E-01 | 1.01E+00 | 1.39E-02 | 1.01E+00 |
| ENSG000001 | CIBAR1P1 | 7.04E-01 | 8.95E-01 | 1.06E+00 | 8.06E-02 | 1.06E+00 |
| ENSG000002 | SF1-DT | 7.04E-01 | ? | 8.32E-01 | -2.66E-01 | -1.20E+00 |
| ENSG000002 | --- | 7.04E-01 | 8.95E-01 | 9.41E-01 | -8.76E-02 | -1.06E+00 |
| ENSG000001 | LYPD6 | 7.04E-01 | 8.95E-01 | 9.50E-01 | -7.39E-02 | -1.05E+00 |
| ENSG000002 | STARD13-AS | 7.04E-01 | ? | 1.18E+00 | 2.39E-01 | 1.18E+00 |
| ENSG000001 | TCAP | 7.04E-01 | ? | 8.49E-01 | -2.37E-01 | -1.18E+00 |
| ENSG000001 | STX10 | 7.04E-01 | 8.95E-01 | 1.02E+00 | 2.73E-02 | 1.02E+00 |
| ENSG000002 | --- | 7.05E-01 | ? | 1.14E+00 | 1.83E-01 | 1.14E+00 |
| ENSG000001 | ZNF558 | 7.05E-01 | 8.95E-01 | 9.80E-01 | -2.97E-02 | -1.02E+00 |
| ENSG00000C | ADSS2 | 7.05E-01 | 8.95E-01 | 1.01E+00 | 1.81E-02 | 1.01E+00 |
| ENSG00000C | ALG9 | 7.05E-01 | 8.95E-01 | 9.89E-01 | -1.54E-02 | -1.01E+00 |
| ENSG000002 | BEND3P3 | 7.05E-01 | 8.95E-01 | 1.06E+00 | 7.81E-02 | 1.06E+00 |
| ENSG000001 | SH3BGRL3 | 7.05E-01 | 8.95E-01 | 1.01E+00 | 2.10E-02 | 1.01E+00 |
| ENSG000002 | LINC01618 | 7.05E-01 | ? | 1.06E+00 | 9.01E-02 | 1.06E+00 |
| ENSG000001 | MACROH2A1 | 7.05E-01 | 8.95E-01 | 1.01E+00 | 1.14E-02 | 1.01E+00 |
| ENSG000001 | SRPRA | 7.05E-01 | 8.95E-01 | 9.88E-01 | -1.70E-02 | -1.01E+00 |
| ENSG00000C | EPS15 | 7.05E-01 | 8.95E-01 | 9.88E-01 | -1.74E-02 | -1.01E+00 |
| ENSG000002 | --- | 7.05E-01 | ? | 1.08E+00 | 1.14E-01 | 1.08E+00 |
| ENSG000001 | AGTPBP1 | 7.05E-01 | 8.95E-01 | 1.02E+00 | 2.45E-02 | 1.02E+00 |
| ENSG000002 | MIR600HG | 7.05E-01 | 8.95E-01 | 1.05E+00 | 6.57E-02 | 1.05E+00 |
| ENSG000002 | H4C14 | 7.05E-01 | 8.95E-01 | 9.30E-01 | -1.05E-01 | -1.08E+00 |
| ENSG000002 | AGAP14P | 7.06E-01 | ? | 9.19E-01 | -1.21E-01 | -1.09E+00 |
| ENSG000001 | SH3TC1 | 7.06E-01 | 8.95E-01 | 1.03E+00 | 3.75E-02 | 1.03E+00 |
| ENSG000001 | EXD1 | 7.06E-01 | ? | 7.94E-01 | -3.33E-01 | -1.26E+00 |
| ENSG00000C | MCUR1 | 7.06E-01 | 8.95E-01 | 9.84E-01 | -2.39E-02 | -1.02E+00 |
| ENSG000001 | PMF1 | 7.06E-01 | 8.95E-01 | 1.02E+00 | 2.29E-02 | 1.02E+00 |
| ENSG000001 | PDIA4 | 7.06E-01 | 8.95E-01 | 9.82E-01 | -2.66E-02 | -1.02E+00 |
| ENSG000001 | ADPRS | 7.06E-01 | 8.95E-01 | 1.02E+00 | 2.76E-02 | 1.02E+00 |
| ENSG000001 | HDDC3 | 7.06E-01 | 8.95E-01 | 1.03E+00 | 3.74E-02 | 1.03E+00 |
| ENSG000002 | --- | 7.06E-01 | ? | 8.61E-01 | -2.16E-01 | -1.16E+00 |
| ENSG00000C | GLG1 | 7.06E-01 | 8.95E-01 | 1.01E+00 | 1.23E-02 | 1.01E+00 |
| ENSG000001 | LIPT2 | 7.06E-01 | ? | 9.35E-01 | -9.67E-02 | -1.07E+00 |
| ENSG000001 | DCAF4 | 7.06E-01 | 8.95E-01 | 9.84E-01 | -2.25E-02 | -1.02E+00 |
| ENSG000001 | HERC1 | 7.06E-01 | 8.95E-01 | 9.75E-01 | -3.65E-02 | -1.03E+00 |
| ENSG000001 | RSU1 | 7.06E-01 | 8.95E-01 | 1.01E+00 | 1.84E-02 | 1.01E+00 |
| ENSG000001 | KLF12 | 7.06E-01 | 8.95E-01 | 9.82E-01 | -2.66E-02 | -1.02E+00 |
| ENSG00000C | WIPI1 | 7.06E-01 | 8.95E-01 | 1.02E+00 | 2.58E-02 | 1.02E+00 |
| ENSG000002 | GAPDHP35 | 7.06E-01 | ? | 8.41E-01 | -2.50E-01 | -1.19E+00 |
| ENSG000001 | TIRAP | 7.06E-01 | 8.95E-01 | 9.81E-01 | -2.83E-02 | -1.02E+00 |
| ENSG000001 | MPHOSPH6 | 7.06E-01 | 8.95E-01 | 1.02E+00 | 2.84E-02 | 1.02E+00 |
| ENSG000002 | BORCS7-ASN | 7.06E-01 | ? | 1.40E+00 | 4.87E-01 | 1.40E+00 |
| ENSG000002 | TICAM2-AS1 | 7.06E-01 | ? | 9.09E-01 | -1.37E-01 | -1.10E+00 |
| ENSG000002 | DCP1A | 7.06E-01 | 8.95E-01 | 9.89E-01 | -1.56E-02 | -1.01E+00 |
| ENSG000001 | TMEM241 | 7.06E-01 | 8.95E-01 | 1.03E+00 | 3.75E-02 | 1.03E+00 |

| | | | | | | |
|---|---|---|---|---|---|---|
| ENSG000002 | FAM223B | 7.07E-01 | ? | 8.53E-01 | -2.30E-01 | -1.17E+00 |
| ENSG000001 | ZSCAN18 | 7.07E-01 | 8.95E-01 | 1.02E+00 | 3.43E-02 | 1.02E+00 |
| ENSG000001 | VPS4B | 7.07E-01 | 8.95E-01 | 9.89E-01 | -1.59E-02 | -1.01E+00 |
| ENSG00000C | SLC7A4 | 7.07E-01 | 8.95E-01 | 1.07E+00 | 9.26E-02 | 1.07E+00 |
| ENSG000001 | PCYT2 | 7.07E-01 | 8.95E-01 | 1.02E+00 | 3.32E-02 | 1.02E+00 |
| ENSG000001 | CAB39 | 7.07E-01 | 8.95E-01 | 9.87E-01 | -1.82E-02 | -1.01E+00 |
| ENSG00000C | POLR3B | 7.07E-01 | 8.95E-01 | 9.83E-01 | -2.47E-02 | -1.02E+00 |
| ENSG000001 | FAM219A | 7.07E-01 | 8.95E-01 | 9.86E-01 | -2.03E-02 | -1.01E+00 |
| ENSG000001 | CABLES2 | 7.07E-01 | 8.95E-01 | 9.72E-01 | -4.10E-02 | -1.03E+00 |
| ENSG000001 | CHAC1 | 7.07E-01 | 8.95E-01 | 9.44E-01 | -8.29E-02 | -1.06E+00 |
| ENSG000001 | NUDT16L1 | 7.07E-01 | 8.95E-01 | 1.03E+00 | 4.87E-02 | 1.03E+00 |
| ENSG000001 | LDB3 | 7.07E-01 | ? | 8.59E-01 | -2.20E-01 | -1.16E+00 |
| ENSG00000C | ST3GAL6 | 7.07E-01 | 8.95E-01 | 9.61E-01 | -5.68E-02 | -1.04E+00 |
| ENSG000001 | KLHL8 | 7.07E-01 | 8.95E-01 | 9.84E-01 | -2.26E-02 | -1.02E+00 |
| ENSG000001 | SH2B2 | 7.07E-01 | 8.95E-01 | 9.46E-01 | -8.01E-02 | -1.06E+00 |
| ENSG000002 | --- | 7.07E-01 | ? | 8.83E-01 | -1.80E-01 | -1.13E+00 |
| ENSG000001 | MXRA5 | 7.07E-01 | 8.95E-01 | 1.03E+00 | 3.58E-02 | 1.03E+00 |
| ENSG000001 | NPDC1 | 7.07E-01 | 8.95E-01 | 1.03E+00 | 4.93E-02 | 1.03E+00 |
| ENSG000001 | IRAK2 | 7.08E-01 | 8.95E-01 | 1.03E+00 | 3.60E-02 | 1.03E+00 |
| ENSG000001 | CAMK4 | 7.08E-01 | 8.95E-01 | 9.62E-01 | -5.61E-02 | -1.04E+00 |
| ENSG000001 | SPRED2 | 7.08E-01 | 8.95E-01 | 9.90E-01 | -1.51E-02 | -1.01E+00 |
| ENSG00000C | SCARF1 | 7.08E-01 | ? | 9.38E-01 | -9.17E-02 | -1.07E+00 |
| ENSG00000C | FKBP1A | 7.08E-01 | 8.95E-01 | 1.01E+00 | 1.43E-02 | 1.01E+00 |
| ENSG000001 | ABHD6 | 7.08E-01 | 8.95E-01 | 9.78E-01 | -3.26E-02 | -1.02E+00 |
| ENSG000002 | --- | 7.08E-01 | ? | 1.09E+00 | 1.21E-01 | 1.09E+00 |
| ENSG000001 | OS9 | 7.08E-01 | 8.95E-01 | 9.91E-01 | -1.33E-02 | -1.01E+00 |
| ENSG000002 | --- | 7.08E-01 | ? | 1.14E+00 | 1.86E-01 | 1.14E+00 |
| ENSG000001 | CNST | 7.08E-01 | 8.95E-01 | 9.85E-01 | -2.23E-02 | -1.02E+00 |
| ENSG000002 | C17orf67 | 7.08E-01 | ? | 1.09E+00 | 1.23E-01 | 1.09E+00 |
| ENSG000001 | NDUFS8 | 7.08E-01 | 8.95E-01 | 1.03E+00 | 3.60E-02 | 1.03E+00 |
| ENSG000001 | CDC25C | 7.08E-01 | ? | 9.40E-01 | -8.93E-02 | -1.06E+00 |
| ENSG00000C | TYMP | 7.08E-01 | 8.95E-01 | 9.66E-01 | -5.03E-02 | -1.04E+00 |
| ENSG000002 | BCAR3-AS1 | 7.08E-01 | ? | 1.13E+00 | 1.78E-01 | 1.13E+00 |
| ENSG000002 | SFTA1P | 7.08E-01 | 8.95E-01 | 9.72E-01 | -4.03E-02 | -1.03E+00 |
| ENSG000002 | --- | 7.08E-01 | ? | 9.32E-01 | -1.02E-01 | -1.07E+00 |
| ENSG000002 | --- | 7.08E-01 | ? | 9.04E-01 | -1.46E-01 | -1.11E+00 |
| ENSG000002 | NSUN5P1 | 7.09E-01 | 8.96E-01 | 1.07E+00 | 1.01E-01 | 1.07E+00 |
| ENSG000002 | --- | 7.09E-01 | ? | 1.20E+00 | 2.58E-01 | 1.20E+00 |
| ENSG000001 | ZBTB18 | 7.09E-01 | 8.96E-01 | 1.02E+00 | 2.45E-02 | 1.02E+00 |
| ENSG000001 | UTP14A | 7.09E-01 | 8.96E-01 | 9.84E-01 | -2.29E-02 | -1.02E+00 |
| ENSG000002 | --- | 7.09E-01 | 8.96E-01 | 9.47E-01 | -7.81E-02 | -1.06E+00 |
| ENSG000001 | GDF6 | 7.09E-01 | ? | 8.44E-01 | -2.44E-01 | -1.18E+00 |
| ENSG000001 | EDA | 7.09E-01 | 8.96E-01 | 1.04E+00 | 5.25E-02 | 1.04E+00 |
| ENSG000001 | FABP3 | 7.09E-01 | ? | 1.07E+00 | 9.79E-02 | 1.07E+00 |
| ENSG000001 | UBTF | 7.09E-01 | 8.96E-01 | 1.01E+00 | 1.46E-02 | 1.01E+00 |

| | | | | | | | |
|---|---|---|---|---|---|---|---|
| ENSG000001 | BCO2 | 7.09E-01 | ? | | 9.10E-01 | -1.36E-01 | -1.10E+00 |
| ENSG000001 | CALHM4 | 7.09E-01 | ? | | 8.05E-01 | -3.13E-01 | -1.24E+00 |
| ENSG000001 | SACS | 7.09E-01 | | 8.96E-01 | 9.73E-01 | -4.00E-02 | -1.03E+00 |
| ENSG000001 | MCOLN2 | 7.09E-01 | | 8.96E-01 | 1.04E+00 | 5.23E-02 | 1.04E+00 |
| ENSG00000C | PDE4A | 7.09E-01 | | 8.96E-01 | 1.03E+00 | 3.97E-02 | 1.03E+00 |
| ENSG000001 | C11orf1 | 7.09E-01 | | 8.96E-01 | 9.76E-01 | -3.57E-02 | -1.03E+00 |
| ENSG000002 | --- | 7.09E-01 | | 8.96E-01 | 1.10E+00 | 1.44E-01 | 1.10E+00 |
| ENSG000002 | --- | 7.09E-01 | | 8.96E-01 | 1.05E+00 | 6.42E-02 | 1.05E+00 |
| ENSG000001 | ADAMTS7 | 7.09E-01 | | 8.96E-01 | 1.04E+00 | 5.80E-02 | 1.04E+00 |
| ENSG000001 | SNRPB2 | 7.09E-01 | | 8.96E-01 | 1.01E+00 | 2.06E-02 | 1.01E+00 |
| ENSG000001 | MGA | 7.09E-01 | | 8.96E-01 | 9.80E-01 | -2.98E-02 | -1.02E+00 |
| ENSG000002 | --- | 7.09E-01 | ? | | 1.11E+00 | 1.55E-01 | 1.11E+00 |
| ENSG000002 | --- | 7.10E-01 | | 8.96E-01 | 1.18E+00 | 2.36E-01 | 1.18E+00 |
| ENSG000001 | MTCO3P12 | 7.10E-01 | ? | | 1.11E+00 | 1.45E-01 | 1.11E+00 |
| ENSG000001 | RBP7 | 7.10E-01 | ? | | 1.13E+00 | 1.76E-01 | 1.13E+00 |
| ENSG000002 | STX16-NPEPI | 7.10E-01 | ? | | 9.10E-01 | -1.36E-01 | -1.10E+00 |
| ENSG000001 | S100A8 | 7.10E-01 | | 8.96E-01 | 1.07E+00 | 9.69E-02 | 1.07E+00 |
| ENSG000001 | AGRN | 7.10E-01 | | 8.96E-01 | 1.04E+00 | 5.34E-02 | 1.04E+00 |
| ENSG000002 | AP4B1-AS1 | 7.10E-01 | ? | | 9.21E-01 | -1.19E-01 | -1.09E+00 |
| ENSG000002 | --- | 7.10E-01 | ? | | 8.86E-01 | -1.75E-01 | -1.13E+00 |
| ENSG000001 | PTPRR | 7.10E-01 | | 8.96E-01 | 9.42E-01 | -8.68E-02 | -1.06E+00 |
| ENSG000002 | SBDSP1 | 7.10E-01 | | 8.96E-01 | 8.89E-01 | -1.70E-01 | -1.13E+00 |
| ENSG000002 | --- | 7.10E-01 | ? | | 8.99E-01 | -1.54E-01 | -1.11E+00 |
| ENSG000002 | MIR302CHG | 7.10E-01 | ? | | 9.16E-01 | -1.27E-01 | -1.09E+00 |
| ENSG00000C | UNKL | 7.10E-01 | | 8.96E-01 | 9.67E-01 | -4.91E-02 | -1.03E+00 |
| ENSG000002 | RAB43P1 | 7.10E-01 | | 8.96E-01 | 1.07E+00 | 1.04E-01 | 1.07E+00 |
| ENSG000001 | IFNE | 7.10E-01 | | 8.96E-01 | 9.44E-01 | -8.31E-02 | -1.06E+00 |
| ENSG000002 | RBM8A | 7.10E-01 | | 8.96E-01 | 9.89E-01 | -1.53E-02 | -1.01E+00 |
| ENSG000001 | FILIP1 | 7.10E-01 | ? | | 8.52E-01 | -2.31E-01 | -1.17E+00 |
| ENSG000001 | USP30 | 7.10E-01 | | 8.96E-01 | 9.78E-01 | -3.26E-02 | -1.02E+00 |
| ENSG000002 | --- | 7.10E-01 | ? | | 8.69E-01 | -2.03E-01 | -1.15E+00 |
| ENSG000001 | PAQR8 | 7.10E-01 | | 8.96E-01 | 1.03E+00 | 4.51E-02 | 1.03E+00 |
| ENSG000001 | MORC2 | 7.11E-01 | | 8.96E-01 | 1.01E+00 | 1.44E-02 | 1.01E+00 |
| ENSG000001 | ZC3HAV1 | 7.11E-01 | | 8.96E-01 | 9.84E-01 | -2.28E-02 | -1.02E+00 |
| ENSG000002 | LINC01138 | 7.11E-01 | | 8.96E-01 | 9.56E-01 | -6.49E-02 | -1.05E+00 |
| ENSG00000C | PSMC4 | 7.11E-01 | | 8.96E-01 | 9.90E-01 | -1.50E-02 | -1.01E+00 |
| ENSG00000C | MCCC1 | 7.11E-01 | | 8.96E-01 | 1.02E+00 | 3.21E-02 | 1.02E+00 |
| ENSG000001 | ZNF71 | 7.11E-01 | | 8.96E-01 | 1.02E+00 | 2.84E-02 | 1.02E+00 |
| ENSG00000C | ROGDI | 7.11E-01 | | 8.96E-01 | 9.80E-01 | -2.86E-02 | -1.02E+00 |
| ENSG000001 | AFF2 | 7.11E-01 | ? | | 1.08E+00 | 1.16E-01 | 1.08E+00 |
| ENSG000001 | BSN | 7.11E-01 | | 8.96E-01 | 9.51E-01 | -7.28E-02 | -1.05E+00 |
| ENSG000002 | --- | 7.11E-01 | ? | | 1.15E+00 | 2.08E-01 | 1.15E+00 |
| ENSG00000C | AAAS | 7.11E-01 | | 8.96E-01 | 1.02E+00 | 2.19E-02 | 1.02E+00 |
| ENSG000002 | FLG-AS1 | 7.11E-01 | | 8.96E-01 | 9.68E-01 | -4.73E-02 | -1.03E+00 |
| ENSG000001 | DPH2 | 7.11E-01 | | 8.96E-01 | 1.02E+00 | 3.17E-02 | 1.02E+00 |

| | | | | | | |
|---|---|---|---|---|---|---|
| ENSG000001 | BAG4 | 7.11E-01 | 8.96E-01 | 1.02E+00 | 2.59E-02 | 1.02E+00 |
| ENSG000002 | --- | 7.11E-01 | ? | 1.12E+00 | 1.58E-01 | 1.12E+00 |
| ENSG000001 | SLC12A7 | 7.11E-01 | 8.96E-01 | 9.82E-01 | -2.61E-02 | -1.02E+00 |
| ENSG000001 | DHTKD1 | 7.12E-01 | 8.96E-01 | 1.02E+00 | 2.74E-02 | 1.02E+00 |
| ENSG000001 | DNAJC8 | 7.12E-01 | 8.96E-01 | 9.85E-01 | -2.21E-02 | -1.02E+00 |
| ENSG000001 | BAK1P1 | 7.12E-01 | ? | 8.49E-01 | -2.37E-01 | -1.18E+00 |
| ENSG000001 | DBN1 | 7.12E-01 | 8.96E-01 | 9.85E-01 | -2.25E-02 | -1.02E+00 |
| ENSG000002 | --- | 7.12E-01 | 8.97E-01 | 9.50E-01 | -7.38E-02 | -1.05E+00 |
| ENSG000002 | --- | 7.12E-01 | ? | 9.23E-01 | -1.16E-01 | -1.08E+00 |
| ENSG000002 | AP1AR-DT | 7.12E-01 | ? | 1.09E+00 | 1.25E-01 | 1.09E+00 |
| ENSG00000C | DCN | 7.12E-01 | ? | 1.15E+00 | 2.04E-01 | 1.15E+00 |
| ENSG000002 | MMADHC-DT | 7.12E-01 | ? | 8.67E-01 | -2.06E-01 | -1.15E+00 |
| ENSG000002 | DDX18P1 | 7.12E-01 | ? | 1.17E+00 | 2.27E-01 | 1.17E+00 |
| ENSG000002 | SFT2D2 | 7.12E-01 | 8.97E-01 | 1.02E+00 | 2.49E-02 | 1.02E+00 |
| ENSG000002 | RPS23P8 | 7.12E-01 | 8.97E-01 | 1.05E+00 | 6.98E-02 | 1.05E+00 |
| ENSG000002 | --- | 7.12E-01 | ? | 1.07E+00 | 9.53E-02 | 1.07E+00 |
| ENSG000001 | PCLAF | 7.13E-01 | 8.97E-01 | 9.44E-01 | -8.27E-02 | -1.06E+00 |
| ENSG000002 | TIPARP-AS1 | 7.13E-01 | ? | 1.17E+00 | 2.24E-01 | 1.17E+00 |
| ENSG000002 | RPS2P55 | 7.13E-01 | ? | 1.11E+00 | 1.51E-01 | 1.11E+00 |
| ENSG000001 | PRKAR1A | 7.13E-01 | 8.97E-01 | 9.82E-01 | -2.58E-02 | -1.02E+00 |
| ENSG000001 | MX1 | 7.13E-01 | 8.97E-01 | 9.69E-01 | -4.48E-02 | -1.03E+00 |
| ENSG000001 | SYNCRIP | 7.13E-01 | 8.97E-01 | 9.88E-01 | -1.80E-02 | -1.01E+00 |
| ENSG000001 | WFDC2 | 7.13E-01 | ? | 8.91E-01 | -1.66E-01 | -1.12E+00 |
| ENSG000002 | LINC02009 | 7.13E-01 | 8.97E-01 | 1.06E+00 | 8.99E-02 | 1.06E+00 |
| ENSG000001 | NEIL1 | 7.13E-01 | 8.97E-01 | 1.04E+00 | 5.03E-02 | 1.04E+00 |
| ENSG000002 | CASC15 | 7.13E-01 | 8.97E-01 | 9.51E-01 | -7.28E-02 | -1.05E+00 |
| ENSG000001 | MIS12 | 7.13E-01 | 8.97E-01 | 1.02E+00 | 2.69E-02 | 1.02E+00 |
| ENSG000002 | KCNQ5-DT | 7.13E-01 | ? | 8.75E-01 | -1.93E-01 | -1.14E+00 |
| ENSG000002 | FMNL1-AS1 | 7.13E-01 | ? | 1.10E+00 | 1.41E-01 | 1.10E+00 |
| ENSG000002 | LIVAR | 7.13E-01 | 8.97E-01 | 1.06E+00 | 8.72E-02 | 1.06E+00 |
| ENSG000002 | MKNK1-AS1 | 7.14E-01 | ? | 9.18E-01 | -1.23E-01 | -1.09E+00 |
| ENSG000001 | WDR43 | 7.14E-01 | 8.98E-01 | 9.84E-01 | -2.36E-02 | -1.02E+00 |
| ENSG000001 | CD248 | 7.14E-01 | ? | 8.05E-01 | -3.12E-01 | -1.24E+00 |
| ENSG000001 | CRTC3 | 7.14E-01 | 8.98E-01 | 1.02E+00 | 2.26E-02 | 1.02E+00 |
| ENSG00000C | SCARB1 | 7.14E-01 | 8.98E-01 | 1.02E+00 | 2.56E-02 | 1.02E+00 |
| ENSG000001 | TBC1D12 | 7.14E-01 | 8.98E-01 | 9.83E-01 | -2.45E-02 | -1.02E+00 |
| ENSG000001 | HELQ | 7.14E-01 | 8.98E-01 | 1.02E+00 | 2.49E-02 | 1.02E+00 |
| ENSG000001 | FBXW10 | 7.14E-01 | ? | 9.16E-01 | -1.26E-01 | -1.09E+00 |
| ENSG000001 | S100A7 | 7.14E-01 | 8.98E-01 | 1.05E+00 | 7.42E-02 | 1.05E+00 |
| ENSG000001 | BICDL1 | 7.14E-01 | 8.98E-01 | 1.03E+00 | 4.58E-02 | 1.03E+00 |
| ENSG000001 | COX7B | 7.14E-01 | 8.98E-01 | 9.86E-01 | -1.98E-02 | -1.01E+00 |
| ENSG000002 | AGER | 7.14E-01 | 8.98E-01 | 9.65E-01 | -5.19E-02 | -1.04E+00 |
| ENSG000001 | CHD6 | 7.14E-01 | 8.98E-01 | 1.01E+00 | 2.06E-02 | 1.01E+00 |
| ENSG000002 | --- | 7.14E-01 | 8.98E-01 | 1.04E+00 | 5.90E-02 | 1.04E+00 |
| ENSG000001 | HES4 | 7.14E-01 | 8.98E-01 | 1.05E+00 | 7.12E-02 | 1.05E+00 |

| | | | | | | |
|---|---|---|---|---|---|---|
| ENSG00000C | LAPTM4A | 7.15E-01 | 8.98E-01 | 1.01E+00 | 1.34E-02 | 1.01E+00 |
| ENSG000002 | --- | 7.15E-01 | ? | 1.08E+00 | 1.12E-01 | 1.08E+00 |
| ENSG000002 | --- | 7.15E-01 | ? | 1.08E+00 | 1.09E-01 | 1.08E+00 |
| ENSG000001 | EMC3 | 7.15E-01 | 8.98E-01 | 1.01E+00 | 1.61E-02 | 1.01E+00 |
| ENSG000002 | --- | 7.15E-01 | ? | 9.04E-01 | -1.46E-01 | -1.11E+00 |
| ENSG000001 | FMO5 | 7.15E-01 | 8.98E-01 | 9.51E-01 | -7.28E-02 | -1.05E+00 |
| ENSG000001 | CSTPP1 | 7.15E-01 | 8.98E-01 | 9.76E-01 | -3.45E-02 | -1.02E+00 |
| ENSG000002 | MEF2B | 7.15E-01 | 8.98E-01 | 1.09E+00 | 1.28E-01 | 1.09E+00 |
| ENSG000002 | --- | 7.15E-01 | ? | 1.18E+00 | 2.44E-01 | 1.18E+00 |
| ENSG000001 | GDPD2 | 7.15E-01 | ? | 1.12E+00 | 1.61E-01 | 1.12E+00 |
| ENSG000002 | YAE1 | 7.15E-01 | 8.98E-01 | 9.75E-01 | -3.64E-02 | -1.03E+00 |
| ENSG000002 | TMEM199 | 7.15E-01 | 8.98E-01 | 9.89E-01 | -1.57E-02 | -1.01E+00 |
| ENSG000002 | PKD1P6-NPIF | 7.15E-01 | 8.98E-01 | 1.05E+00 | 7.51E-02 | 1.05E+00 |
| ENSG000002 | --- | 7.15E-01 | ? | 8.41E-01 | -2.49E-01 | -1.19E+00 |
| ENSG000001 | CAV2 | 7.15E-01 | 8.98E-01 | 9.86E-01 | -2.06E-02 | -1.01E+00 |
| ENSG000001 | WDR24 | 7.15E-01 | 8.98E-01 | 1.03E+00 | 4.22E-02 | 1.03E+00 |
| ENSG000002 | CISD3 | 7.15E-01 | 8.98E-01 | 1.02E+00 | 2.19E-02 | 1.02E+00 |
| ENSG000001 | SHC3 | 7.15E-01 | 8.98E-01 | 1.05E+00 | 6.56E-02 | 1.05E+00 |
| ENSG000001 | CCT7 | 7.15E-01 | 8.98E-01 | 1.01E+00 | 1.43E-02 | 1.01E+00 |
| ENSG000002 | ATP2B1-AS1 | 7.15E-01 | ? | 1.13E+00 | 1.70E-01 | 1.13E+00 |
| ENSG000002 | CA5BP1 | 7.15E-01 | 8.98E-01 | 9.81E-01 | -2.83E-02 | -1.02E+00 |
| ENSG000001 | C1orf162 | 7.15E-01 | ? | 1.11E+00 | 1.53E-01 | 1.11E+00 |
| ENSG000001 | MUS81 | 7.15E-01 | 8.98E-01 | 1.02E+00 | 2.62E-02 | 1.02E+00 |
| ENSG000001 | ST20 | 7.15E-01 | 8.98E-01 | 9.63E-01 | -5.50E-02 | -1.04E+00 |
| ENSG000001 | ANO4 | 7.15E-01 | 8.98E-01 | 9.69E-01 | -4.58E-02 | -1.03E+00 |
| ENSG000002 | VPS13B-DT | 7.15E-01 | ? | 9.17E-01 | -1.24E-01 | -1.09E+00 |
| ENSG000002 | --- | 7.15E-01 | 8.98E-01 | 1.04E+00 | 5.49E-02 | 1.04E+00 |
| ENSG000001 | --- | 7.15E-01 | ? | 1.12E+00 | 1.60E-01 | 1.12E+00 |
| ENSG000001 | S100P | 7.16E-01 | 8.98E-01 | 9.24E-01 | -1.14E-01 | -1.08E+00 |
| ENSG000001 | ZNF281 | 7.16E-01 | 8.98E-01 | 1.01E+00 | 1.97E-02 | 1.01E+00 |
| ENSG000001 | RNF2 | 7.16E-01 | 8.98E-01 | 9.81E-01 | -2.81E-02 | -1.02E+00 |
| ENSG000001 | ADGRF1 | 7.16E-01 | 8.98E-01 | 1.04E+00 | 5.50E-02 | 1.04E+00 |
| ENSG000001 | KDSR | 7.16E-01 | 8.98E-01 | 9.90E-01 | -1.38E-02 | -1.01E+00 |
| ENSG000001 | ZNF526 | 7.16E-01 | 8.98E-01 | 1.02E+00 | 2.43E-02 | 1.02E+00 |
| ENSG000001 | ZNF557 | 7.16E-01 | 8.98E-01 | 9.82E-01 | -2.67E-02 | -1.02E+00 |
| ENSG00000C | ZNF324 | 7.16E-01 | 8.98E-01 | 1.02E+00 | 3.03E-02 | 1.02E+00 |
| ENSG000001 | FAM227A | 7.16E-01 | 8.98E-01 | 1.06E+00 | 7.87E-02 | 1.06E+00 |
| ENSG00000C | PSMD8 | 7.16E-01 | 8.98E-01 | 1.01E+00 | 1.49E-02 | 1.01E+00 |
| ENSG000001 | MT1F | 7.16E-01 | ? | 1.07E+00 | 1.02E-01 | 1.07E+00 |
| ENSG000001 | CD164L2 | 7.16E-01 | ? | 1.07E+00 | 1.00E-01 | 1.07E+00 |
| ENSG000001 | PLGLB1 | 7.16E-01 | 8.98E-01 | 1.07E+00 | 1.03E-01 | 1.07E+00 |
| ENSG000002 | KIF23-AS1 | 7.16E-01 | ? | 1.09E+00 | 1.22E-01 | 1.09E+00 |
| ENSG000002 | --- | 7.17E-01 | 8.98E-01 | 1.04E+00 | 5.76E-02 | 1.04E+00 |
| ENSG000001 | F2R | 7.17E-01 | 8.98E-01 | 1.01E+00 | 2.05E-02 | 1.01E+00 |
| ENSG000002 | MAP3K4-AS1 | 7.17E-01 | ? | 9.23E-01 | -1.15E-01 | -1.08E+00 |

| | | | | | | |
|---|---|---|---|---|---|---|
| ENSG000001 | KCNT1 | 7.17E-01 | ? | 8.72E-01 | -1.97E-01 | -1.15E+00 |
| ENSG000002 | --- | 7.17E-01 | 8.98E-01 | 1.03E+00 | 4.09E-02 | 1.03E+00 |
| ENSG000002 | C4orf54 | 7.17E-01 | ? | 1.13E+00 | 1.73E-01 | 1.13E+00 |
| ENSG000001 | ZNF787 | 7.17E-01 | 8.98E-01 | 1.02E+00 | 3.30E-02 | 1.02E+00 |
| ENSG000001 | APOC1 | 7.17E-01 | ? | 9.28E-01 | -1.07E-01 | -1.08E+00 |
| ENSG000001 | TPRKB | 7.17E-01 | 8.98E-01 | 9.79E-01 | -3.10E-02 | -1.02E+00 |
| ENSG000001 | EDN2 | 7.17E-01 | ? | 1.13E+00 | 1.82E-01 | 1.13E+00 |
| ENSG00000C | LAT2 | 7.17E-01 | 8.98E-01 | 1.05E+00 | 6.42E-02 | 1.05E+00 |
| ENSG000002 | TRIM8-DT | 7.17E-01 | 8.98E-01 | 9.52E-01 | -7.04E-02 | -1.05E+00 |
| ENSG000001 | ZNF474 | 7.17E-01 | ? | 8.43E-01 | -2.47E-01 | -1.19E+00 |
| ENSG000001 | MBOAT7 | 7.17E-01 | 8.98E-01 | 9.87E-01 | -1.93E-02 | -1.01E+00 |
| ENSG000001 | ARF6 | 7.17E-01 | 8.98E-01 | 9.92E-01 | -1.15E-02 | -1.01E+00 |
| ENSG000001 | GPR137 | 7.17E-01 | 8.98E-01 | 1.03E+00 | 4.14E-02 | 1.03E+00 |
| ENSG000001 | MRPS14 | 7.17E-01 | 8.98E-01 | 1.02E+00 | 2.19E-02 | 1.02E+00 |
| ENSG000002 | LINC02918 | 7.17E-01 | ? | 1.08E+00 | 1.13E-01 | 1.08E+00 |
| ENSG000002 | RBPMS-AS1 | 7.17E-01 | ? | 8.76E-01 | -1.90E-01 | -1.14E+00 |
| ENSG000002 | --- | 7.17E-01 | ? | 1.13E+00 | 1.81E-01 | 1.13E+00 |
| ENSG000002 | PLD5P1 | 7.17E-01 | 8.98E-01 | 1.06E+00 | 7.98E-02 | 1.06E+00 |
| ENSG000002 | PPP2CA-DT | 7.18E-01 | ? | 1.12E+00 | 1.59E-01 | 1.12E+00 |
| ENSG000002 | --- | 7.18E-01 | 8.99E-01 | 8.41E-01 | -2.49E-01 | -1.19E+00 |
| ENSG000001 | MAPK4 | 7.18E-01 | ? | 1.14E+00 | 1.95E-01 | 1.14E+00 |
| ENSG000002 | --- | 7.18E-01 | 8.99E-01 | 9.39E-01 | -9.05E-02 | -1.06E+00 |
| ENSG000002 | --- | 7.18E-01 | ? | 1.10E+00 | 1.33E-01 | 1.10E+00 |
| ENSG000002 | --- | 7.18E-01 | 8.99E-01 | 1.05E+00 | 7.40E-02 | 1.05E+00 |
| ENSG000002 | ITPR1-DT | 7.18E-01 | ? | 1.09E+00 | 1.21E-01 | 1.09E+00 |
| ENSG000002 | --- | 7.18E-01 | ? | 8.93E-01 | -1.63E-01 | -1.12E+00 |
| ENSG000002 | --- | 7.18E-01 | ? | 1.14E+00 | 1.86E-01 | 1.14E+00 |
| ENSG000001 | CFAP126 | 7.18E-01 | 8.99E-01 | 1.05E+00 | 7.42E-02 | 1.05E+00 |
| ENSG000001 | RAB3IP | 7.18E-01 | 8.99E-01 | 9.83E-01 | -2.49E-02 | -1.02E+00 |
| ENSG000001 | C9orf163 | 7.18E-01 | ? | 8.96E-01 | -1.59E-01 | -1.12E+00 |
| ENSG000001 | UNC13A | 7.18E-01 | ? | 8.59E-01 | -2.19E-01 | -1.16E+00 |
| ENSG000002 | --- | 7.18E-01 | ? | 1.14E+00 | 1.91E-01 | 1.14E+00 |
| ENSG000001 | LMAN2 | 7.18E-01 | 8.99E-01 | 1.01E+00 | 1.94E-02 | 1.01E+00 |
| ENSG000002 | --- | 7.18E-01 | ? | 8.22E-01 | -2.82E-01 | -1.22E+00 |
| ENSG000002 | ELOA-AS1 | 7.18E-01 | ? | 9.40E-01 | -8.96E-02 | -1.06E+00 |
| ENSG000001 | WDR55 | 7.19E-01 | 8.99E-01 | 1.02E+00 | 2.26E-02 | 1.02E+00 |
| ENSG000002 | LINC01473 | 7.19E-01 | ? | 1.13E+00 | 1.75E-01 | 1.13E+00 |
| ENSG000001 | ZNF565 | 7.19E-01 | 8.99E-01 | 9.65E-01 | -5.11E-02 | -1.04E+00 |
| ENSG000002 | PCDH20 | 7.19E-01 | 8.99E-01 | 1.05E+00 | 7.11E-02 | 1.05E+00 |
| ENSG000001 | ANKDD1B | 7.19E-01 | ? | 1.06E+00 | 8.67E-02 | 1.06E+00 |
| ENSG000001 | EMX1 | 7.19E-01 | ? | 1.19E+00 | 2.47E-01 | 1.19E+00 |
| ENSG000001 | PRR4 | 7.19E-01 | ? | 8.76E-01 | -1.90E-01 | -1.14E+00 |
| ENSG000001 | MEGF11 | 7.19E-01 | ? | 1.17E+00 | 2.24E-01 | 1.17E+00 |
| ENSG000001 | ADGRG4 | 7.19E-01 | ? | 1.15E+00 | 1.97E-01 | 1.15E+00 |
| ENSG000002 | TOMM6 | 7.19E-01 | 8.99E-01 | 9.88E-01 | -1.74E-02 | -1.01E+00 |

| | | | | | | |
|---|---|---|---|---|---|---|
| ENSG000002 | --- | 7.19E-01 | ? | 8.89E-01 | -1.70E-01 | -1.13E+00 |
| ENSG000002 | --- | 7.19E-01 | ? | 1.09E+00 | 1.20E-01 | 1.09E+00 |
| ENSG000001 | CIAO2A | 7.19E-01 | 8.99E-01 | 1.01E+00 | 1.80E-02 | 1.01E+00 |
| ENSG000001 | TBC1D24 | 7.19E-01 | 8.99E-01 | 9.80E-01 | -2.92E-02 | -1.02E+00 |
| ENSG000002 | RPL15P14 | 7.19E-01 | ? | 8.12E-01 | -3.01E-01 | -1.23E+00 |
| ENSG000002 | FTH1P2 | 7.19E-01 | 9.00E-01 | 9.60E-01 | -5.86E-02 | -1.04E+00 |
| ENSG000002 | LINC02777 | 7.19E-01 | 9.00E-01 | 9.39E-01 | -9.06E-02 | -1.06E+00 |
| ENSG00000C | POLR2J | 7.19E-01 | 9.00E-01 | 1.01E+00 | 1.98E-02 | 1.01E+00 |
| ENSG000001 | EML2 | 7.19E-01 | 9.00E-01 | 9.80E-01 | -2.98E-02 | -1.02E+00 |
| ENSG000002 | DYNLRB2-AS | 7.19E-01 | ? | 1.09E+00 | 1.21E-01 | 1.09E+00 |
| ENSG000001 | NAA60 | 7.20E-01 | 9.00E-01 | 1.01E+00 | 2.07E-02 | 1.01E+00 |
| ENSG00000C | MON2 | 7.20E-01 | 9.00E-01 | 1.01E+00 | 1.65E-02 | 1.01E+00 |
| ENSG00000C | MED17 | 7.20E-01 | 9.00E-01 | 9.87E-01 | -1.89E-02 | -1.01E+00 |
| ENSG000002 | C6orf136 | 7.20E-01 | 9.00E-01 | 9.80E-01 | -2.88E-02 | -1.02E+00 |
| ENSG000002 | HMGB3P24 | 7.20E-01 | ? | 8.68E-01 | -2.04E-01 | -1.15E+00 |
| ENSG000001 | ANKRD1 | 7.20E-01 | 9.00E-01 | 1.08E+00 | 1.09E-01 | 1.08E+00 |
| ENSG000001 | PPM1G | 7.20E-01 | 9.00E-01 | 1.01E+00 | 2.03E-02 | 1.01E+00 |
| ENSG00000C | KLHL42 | 7.20E-01 | 9.00E-01 | 9.86E-01 | -2.05E-02 | -1.01E+00 |
| ENSG000002 | HSPD1P11 | 7.20E-01 | ? | 1.13E+00 | 1.82E-01 | 1.13E+00 |
| ENSG000002 | MTND4P9 | 7.20E-01 | ? | 8.63E-01 | -2.12E-01 | -1.16E+00 |
| ENSG000001 | ERBIN | 7.20E-01 | 9.00E-01 | 1.01E+00 | 2.04E-02 | 1.01E+00 |
| ENSG000002 | NFE2L1-DT | 7.20E-01 | ? | 1.08E+00 | 1.16E-01 | 1.08E+00 |
| ENSG000002 | --- | 7.20E-01 | 9.00E-01 | 9.73E-01 | -3.97E-02 | -1.03E+00 |
| ENSG000001 | HTRA2 | 7.20E-01 | 9.00E-01 | 1.02E+00 | 2.34E-02 | 1.02E+00 |
| ENSG000001 | LIPT1 | 7.20E-01 | 9.00E-01 | 9.65E-01 | -5.14E-02 | -1.04E+00 |
| ENSG000001 | MCM3 | 7.20E-01 | 9.00E-01 | 1.02E+00 | 2.70E-02 | 1.02E+00 |
| ENSG000002 | --- | 7.20E-01 | ? | 1.11E+00 | 1.45E-01 | 1.11E+00 |
| ENSG000001 | TUBA8 | 7.20E-01 | ? | 8.65E-01 | -2.10E-01 | -1.16E+00 |
| ENSG00000C | IDH3G | 7.20E-01 | 9.00E-01 | 1.01E+00 | 1.82E-02 | 1.01E+00 |
| ENSG000002 | --- | 7.20E-01 | 9.00E-01 | 9.49E-01 | -7.58E-02 | -1.05E+00 |
| ENSG000002 | LRRC37A2 | 7.21E-01 | 9.00E-01 | 9.80E-01 | -2.95E-02 | -1.02E+00 |
| ENSG000002 | LINC01232 | 7.21E-01 | ? | 9.29E-01 | -1.06E-01 | -1.08E+00 |
| ENSG000001 | NIFK | 7.21E-01 | 9.00E-01 | 1.01E+00 | 1.76E-02 | 1.01E+00 |
| ENSG000001 | ATP8A1 | 7.21E-01 | ? | 9.16E-01 | -1.27E-01 | -1.09E+00 |
| ENSG000002 | ZNF433-AS1 | 7.21E-01 | 9.00E-01 | 9.64E-01 | -5.22E-02 | -1.04E+00 |
| ENSG000001 | BRME1 | 7.21E-01 | ? | 8.87E-01 | -1.73E-01 | -1.13E+00 |
| ENSG000002 | --- | 7.21E-01 | 9.00E-01 | 9.29E-01 | -1.06E-01 | -1.08E+00 |
| ENSG000002 | AJUBA-DT | 7.21E-01 | ? | 9.24E-01 | -1.14E-01 | -1.08E+00 |
| ENSG000001 | ATP2A1 | 7.21E-01 | 9.00E-01 | 9.42E-01 | -8.60E-02 | -1.06E+00 |
| ENSG00000C | INTS13 | 7.21E-01 | 9.00E-01 | 1.01E+00 | 1.81E-02 | 1.01E+00 |
| ENSG000001 | ETS2 | 7.21E-01 | 9.00E-01 | 1.03E+00 | 3.69E-02 | 1.03E+00 |
| ENSG000001 | RTF1 | 7.21E-01 | 9.00E-01 | 1.01E+00 | 1.61E-02 | 1.01E+00 |
| ENSG000002 | MSRB3-AS1 | 7.21E-01 | ? | 9.23E-01 | -1.16E-01 | -1.08E+00 |
| ENSG000001 | GTF3C5 | 7.21E-01 | 9.00E-01 | 9.86E-01 | -1.97E-02 | -1.01E+00 |
| ENSG000001 | UBQLN2 | 7.21E-01 | 9.00E-01 | 1.01E+00 | 1.49E-02 | 1.01E+00 |

| | | | | | | | |
|---|---|---|---|---|---|---|---|
| ENSG000001 | HIF1AN | 7.22E-01 | | 9.00E-01 | 9.91E-01 | -1.24E-02 | -1.01E+00 |
| ENSG000002 | TGFB2-AS1 | 7.22E-01 | ? | | 8.42E-01 | -2.48E-01 | -1.19E+00 |
| ENSG000001 | CCR3 | 7.22E-01 | | 9.00E-01 | 9.47E-01 | -7.85E-02 | -1.06E+00 |
| ENSG000002 | PRELID1P1 | 7.22E-01 | ? | | 1.20E+00 | 2.60E-01 | 1.20E+00 |
| ENSG000002 | PDF | 7.22E-01 | | 9.00E-01 | 1.07E+00 | 9.91E-02 | 1.07E+00 |
| ENSG000002 | --- | 7.22E-01 | ? | | 1.07E+00 | 9.54E-02 | 1.07E+00 |
| ENSG000001 | EARS2 | 7.22E-01 | | 9.00E-01 | 1.01E+00 | 1.60E-02 | 1.01E+00 |
| ENSG000002 | CLDN14-AS1 | 7.22E-01 | | 9.00E-01 | 1.05E+00 | 6.69E-02 | 1.05E+00 |
| ENSG00000C | GNB5 | 7.22E-01 | | 9.00E-01 | 9.89E-01 | -1.65E-02 | -1.01E+00 |
| ENSG000002 | KRT8P46 | 7.22E-01 | ? | | 9.07E-01 | -1.41E-01 | -1.10E+00 |
| ENSG000001 | LRRC3 | 7.22E-01 | | 9.01E-01 | 1.02E+00 | 3.13E-02 | 1.02E+00 |
| ENSG000001 | B4GALT3 | 7.22E-01 | | 9.01E-01 | 1.02E+00 | 2.18E-02 | 1.02E+00 |
| ENSG000001 | FEN1 | 7.22E-01 | | 9.01E-01 | 1.02E+00 | 3.00E-02 | 1.02E+00 |
| ENSG000002 | --- | 7.22E-01 | ? | | 8.74E-01 | -1.95E-01 | -1.14E+00 |
| ENSG000002 | --- | 7.22E-01 | ? | | 1.13E+00 | 1.76E-01 | 1.13E+00 |
| ENSG000001 | PLBD1 | 7.23E-01 | ? | | 1.08E+00 | 1.12E-01 | 1.08E+00 |
| ENSG000001 | RASIP1 | 7.23E-01 | | 9.01E-01 | 9.55E-01 | -6.58E-02 | -1.05E+00 |
| ENSG000002 | EIF4A2P1 | 7.23E-01 | ? | | 8.81E-01 | -1.82E-01 | -1.13E+00 |
| ENSG000002 | RPL17P50 | 7.23E-01 | ? | | 1.10E+00 | 1.39E-01 | 1.10E+00 |
| ENSG000002 | --- | 7.23E-01 | | 9.01E-01 | 1.12E+00 | 1.68E-01 | 1.12E+00 |
| ENSG000002 | CAP1P2 | 7.23E-01 | ? | | 1.10E+00 | 1.40E-01 | 1.10E+00 |
| ENSG000001 | ZNF544 | 7.23E-01 | | 9.01E-01 | 9.87E-01 | -1.87E-02 | -1.01E+00 |
| ENSG000001 | DENND4C | 7.23E-01 | | 9.01E-01 | 1.01E+00 | 1.44E-02 | 1.01E+00 |
| ENSG000002 | KRT16P6 | 7.23E-01 | ? | | 1.12E+00 | 1.69E-01 | 1.12E+00 |
| ENSG000001 | MRAS | 7.23E-01 | | 9.01E-01 | 1.02E+00 | 2.94E-02 | 1.02E+00 |
| ENSG000002 | EGFL8 | 7.23E-01 | | 9.01E-01 | 1.06E+00 | 8.46E-02 | 1.06E+00 |
| ENSG000001 | SPINK5 | 7.23E-01 | | 9.01E-01 | 1.04E+00 | 5.96E-02 | 1.04E+00 |
| ENSG000001 | RAB3A | 7.23E-01 | | 9.01E-01 | 9.48E-01 | -7.71E-02 | -1.05E+00 |
| ENSG000001 | USP49 | 7.23E-01 | | 9.01E-01 | 9.79E-01 | -3.01E-02 | -1.02E+00 |
| ENSG000001 | NOMO3 | 7.24E-01 | | 9.01E-01 | 1.01E+00 | 1.96E-02 | 1.01E+00 |
| ENSG000002 | --- | 7.24E-01 | ? | | 1.07E+00 | 9.80E-02 | 1.07E+00 |
| ENSG000001 | ZNF695 | 7.24E-01 | ? | | 9.05E-01 | -1.45E-01 | -1.11E+00 |
| ENSG000002 | LINC01550 | 7.24E-01 | ? | | 1.13E+00 | 1.81E-01 | 1.13E+00 |
| ENSG000001 | FAM20B | 7.24E-01 | | 9.01E-01 | 1.01E+00 | 1.31E-02 | 1.01E+00 |
| ENSG000002 | SLC22A18AS | 7.24E-01 | | 9.01E-01 | 9.51E-01 | -7.24E-02 | -1.05E+00 |
| ENSG000001 | ITPRIP | 7.24E-01 | | 9.01E-01 | 9.76E-01 | -3.46E-02 | -1.02E+00 |
| ENSG000001 | NCBP1 | 7.24E-01 | | 9.01E-01 | 1.01E+00 | 1.68E-02 | 1.01E+00 |
| ENSG000002 | NPM1P27 | 7.24E-01 | | 9.01E-01 | 9.73E-01 | -3.91E-02 | -1.03E+00 |
| ENSG000002 | --- | 7.24E-01 | ? | | 1.10E+00 | 1.35E-01 | 1.10E+00 |
| ENSG000001 | SP1 | 7.24E-01 | | 9.01E-01 | 1.01E+00 | 1.65E-02 | 1.01E+00 |
| ENSG000002 | LINC01630 | 7.24E-01 | | 9.01E-01 | 9.73E-01 | -3.89E-02 | -1.03E+00 |
| ENSG000001 | CFAP119 | 7.24E-01 | ? | | 9.46E-01 | -8.08E-02 | -1.06E+00 |
| ENSG000001 | ENKD1 | 7.24E-01 | | 9.01E-01 | 1.04E+00 | 6.09E-02 | 1.04E+00 |
| ENSG000001 | EGR2 | 7.24E-01 | ? | | 1.20E+00 | 2.60E-01 | 1.20E+00 |
| ENSG000001 | FBXL4 | 7.24E-01 | | 9.01E-01 | 1.01E+00 | 2.13E-02 | 1.01E+00 |

| | | | | | | |
|---|---|---|---|---|---|---|
| ENSG000001 | ICA1L | 7.24E-01 | 9.01E-01 | 1.02E+00 | 3.15E-02 | 1.02E+00 |
| ENSG000002 | LINC00939 | 7.24E-01 | 9.01E-01 | 9.45E-01 | -8.18E-02 | -1.06E+00 |
| ENSG000001 | EPHX4 | 7.25E-01 | 9.01E-01 | 1.04E+00 | 5.10E-02 | 1.04E+00 |
| ENSG000001 | IFI27 | 7.25E-01 | 9.01E-01 | 9.58E-01 | -6.26E-02 | -1.04E+00 |
| ENSG000001 | IL11RA | 7.25E-01 | 9.01E-01 | 9.56E-01 | -6.49E-02 | -1.05E+00 |
| ENSG000001 | GAPDH | 7.25E-01 | 9.01E-01 | 1.01E+00 | 2.01E-02 | 1.01E+00 |
| ENSG000002 | --- | 7.25E-01 | 9.01E-01 | 9.69E-01 | -4.55E-02 | -1.03E+00 |
| ENSG000001 | TCEAL7 | 7.25E-01 | ? | 8.98E-01 | -1.55E-01 | -1.11E+00 |
| ENSG000002 | --- | 7.25E-01 | ? | 8.64E-01 | -2.11E-01 | -1.16E+00 |
| ENSG000001 | MVB12B | 7.25E-01 | 9.01E-01 | 1.03E+00 | 4.15E-02 | 1.03E+00 |
| ENSG000002 | TRAPPC2B | 7.25E-01 | 9.01E-01 | 1.03E+00 | 4.38E-02 | 1.03E+00 |
| ENSG000001 | ZCCHC7 | 7.25E-01 | 9.01E-01 | 1.01E+00 | 2.05E-02 | 1.01E+00 |
| ENSG000002 | LINC01852 | 7.25E-01 | ? | 8.83E-01 | -1.80E-01 | -1.13E+00 |
| ENSG000002 | LINC00235 | 7.25E-01 | ? | 9.24E-01 | -1.14E-01 | -1.08E+00 |
| ENSG000001 | MAMDC4 | 7.25E-01 | 9.01E-01 | 9.49E-01 | -7.58E-02 | -1.05E+00 |
| ENSG000002 | --- | 7.25E-01 | ? | 9.03E-01 | -1.47E-01 | -1.11E+00 |
| ENSG00000C | CRYBG3 | 7.25E-01 | 9.01E-01 | 1.02E+00 | 2.84E-02 | 1.02E+00 |
| ENSG000001 | PCBP1-AS1 | 7.25E-01 | 9.01E-01 | 9.72E-01 | -4.15E-02 | -1.03E+00 |
| ENSG000001 | ARID5B | 7.25E-01 | 9.01E-01 | 1.02E+00 | 3.06E-02 | 1.02E+00 |
| ENSG000001 | SHARPIN | 7.25E-01 | 9.01E-01 | 9.78E-01 | -3.22E-02 | -1.02E+00 |
| ENSG000001 | KLHL35 | 7.25E-01 | 9.01E-01 | 9.47E-01 | -7.90E-02 | -1.06E+00 |
| ENSG000001 | COL25A1 | 7.25E-01 | ? | 1.14E+00 | 1.88E-01 | 1.14E+00 |
| ENSG000001 | RASD2 | 7.25E-01 | 9.02E-01 | 1.04E+00 | 6.19E-02 | 1.04E+00 |
| ENSG000001 | ZNF572 | 7.25E-01 | ? | 1.08E+00 | 1.06E-01 | 1.08E+00 |
| ENSG000001 | POMP | 7.25E-01 | 9.02E-01 | 1.01E+00 | 2.06E-02 | 1.01E+00 |
| ENSG000001 | ZSCAN5A | 7.25E-01 | 9.02E-01 | 1.04E+00 | 5.28E-02 | 1.04E+00 |
| ENSG000002 | --- | 7.26E-01 | ? | 1.18E+00 | 2.36E-01 | 1.18E+00 |
| ENSG000001 | CES2 | 7.26E-01 | 9.02E-01 | 9.88E-01 | -1.70E-02 | -1.01E+00 |
| ENSG000002 | --- | 7.26E-01 | ? | 1.07E+00 | 1.01E-01 | 1.07E+00 |
| ENSG000001 | RABIF | 7.26E-01 | 9.02E-01 | 1.01E+00 | 1.87E-02 | 1.01E+00 |
| ENSG000002 | CEROX1 | 7.26E-01 | ? | 8.61E-01 | -2.16E-01 | -1.16E+00 |
| ENSG000001 | CLEC18A | 7.26E-01 | 9.02E-01 | 9.39E-01 | -9.02E-02 | -1.06E+00 |
| ENSG000002 | RPS27AP2 | 7.26E-01 | ? | 8.20E-01 | -2.86E-01 | -1.22E+00 |
| ENSG000002 | --- | 7.26E-01 | ? | 1.07E+00 | 1.04E-01 | 1.07E+00 |
| ENSG000001 | LEO1 | 7.26E-01 | 9.02E-01 | 9.81E-01 | -2.81E-02 | -1.02E+00 |
| ENSG00000C | BAIAP2L1 | 7.26E-01 | 9.02E-01 | 9.88E-01 | -1.75E-02 | -1.01E+00 |
| ENSG000002 | --- | 7.26E-01 | ? | 8.96E-01 | -1.58E-01 | -1.12E+00 |
| ENSG000001 | SULF2 | 7.26E-01 | 9.02E-01 | 9.89E-01 | -1.66E-02 | -1.01E+00 |
| ENSG00000C | LARS2 | 7.26E-01 | 9.02E-01 | 9.87E-01 | -1.93E-02 | -1.01E+00 |
| ENSG000001 | FANCB | 7.26E-01 | 9.02E-01 | 9.65E-01 | -5.14E-02 | -1.04E+00 |
| ENSG000001 | CASKIN2 | 7.26E-01 | 9.02E-01 | 1.02E+00 | 3.23E-02 | 1.02E+00 |
| ENSG000001 | POLA1 | 7.26E-01 | 9.02E-01 | 9.75E-01 | -3.58E-02 | -1.03E+00 |
| ENSG000001 | BAALC | 7.26E-01 | ? | 1.20E+00 | 2.64E-01 | 1.20E+00 |
| ENSG000001 | MRPS36 | 7.26E-01 | 9.02E-01 | 9.81E-01 | -2.74E-02 | -1.02E+00 |
| ENSG000001 | POLR1F | 7.26E-01 | 9.02E-01 | 1.01E+00 | 1.45E-02 | 1.01E+00 |

| | | | | | | |
|---|---|---|---|---|---|---|
| ENSG000001 | GMEB1 | 7.27E-01 | 9.02E-01 | 9.87E-01 | -1.96E-02 | -1.01E+00 |
| ENSG000001 | SNRNP70 | 7.27E-01 | 9.02E-01 | 1.02E+00 | 2.43E-02 | 1.02E+00 |
| ENSG000001 | RPL10P16 | 7.27E-01 | 9.02E-01 | 1.04E+00 | 5.88E-02 | 1.04E+00 |
| ENSG00000C | AHRR | 7.27E-01 | ? | 8.21E-01 | -2.84E-01 | -1.22E+00 |
| ENSG000002 | --- | 7.27E-01 | ? | 1.19E+00 | 2.50E-01 | 1.19E+00 |
| ENSG000001 | LPGAT1 | 7.27E-01 | 9.02E-01 | 1.01E+00 | 1.75E-02 | 1.01E+00 |
| ENSG00000C | IYD | 7.27E-01 | ? | 9.16E-01 | -1.27E-01 | -1.09E+00 |
| ENSG000002 | BORCS8 | 7.27E-01 | 9.02E-01 | 1.02E+00 | 3.04E-02 | 1.02E+00 |
| ENSG000001 | GPR84 | 7.27E-01 | ? | 8.57E-01 | -2.22E-01 | -1.17E+00 |
| ENSG000001 | DNAJC13 | 7.27E-01 | 9.02E-01 | 9.87E-01 | -1.90E-02 | -1.01E+00 |
| ENSG000002 | HNRNPA3P1: | 7.27E-01 | ? | 8.43E-01 | -2.47E-01 | -1.19E+00 |
| ENSG000001 | KDM4C | 7.27E-01 | 9.02E-01 | 1.01E+00 | 1.86E-02 | 1.01E+00 |
| ENSG000002 | RPL7AP16 | 7.27E-01 | ? | 1.09E+00 | 1.29E-01 | 1.09E+00 |
| ENSG000002 | --- | 7.27E-01 | ? | 1.09E+00 | 1.20E-01 | 1.09E+00 |
| ENSG000001 | KLF11 | 7.27E-01 | 9.02E-01 | 1.02E+00 | 2.77E-02 | 1.02E+00 |
| ENSG000002 | KCTD11 | 7.28E-01 | 9.02E-01 | 9.82E-01 | -2.62E-02 | -1.02E+00 |
| ENSG000001 | MRPS2 | 7.28E-01 | 9.02E-01 | 1.01E+00 | 2.15E-02 | 1.01E+00 |
| ENSG000002 | --- | 7.28E-01 | 9.02E-01 | 9.57E-01 | -6.37E-02 | -1.05E+00 |
| ENSG000002 | --- | 7.28E-01 | 9.03E-01 | 1.04E+00 | 6.28E-02 | 1.04E+00 |
| ENSG000002 | --- | 7.28E-01 | ? | 9.31E-01 | -1.03E-01 | -1.07E+00 |
| ENSG000002 | YBX1P10 | 7.28E-01 | 9.03E-01 | 9.62E-01 | -5.52E-02 | -1.04E+00 |
| ENSG000001 | BCL11A | 7.28E-01 | 9.03E-01 | 9.82E-01 | -2.59E-02 | -1.02E+00 |
| ENSG000001 | HNRNPU | 7.28E-01 | 9.03E-01 | 9.92E-01 | -1.12E-02 | -1.01E+00 |
| ENSG000002 | --- | 7.28E-01 | ? | 1.12E+00 | 1.57E-01 | 1.12E+00 |
| ENSG000002 | --- | 7.28E-01 | 9.03E-01 | 9.52E-01 | -7.08E-02 | -1.05E+00 |
| ENSG000002 | LRIG2-DT | 7.28E-01 | ? | 1.14E+00 | 1.92E-01 | 1.14E+00 |
| ENSG000001 | ZNF676 | 7.28E-01 | ? | 9.22E-01 | -1.17E-01 | -1.08E+00 |
| ENSG000001 | METTL7A | 7.28E-01 | ? | 1.06E+00 | 9.04E-02 | 1.06E+00 |
| ENSG000001 | CDC42BPG | 7.28E-01 | 9.03E-01 | 1.02E+00 | 2.36E-02 | 1.02E+00 |
| ENSG000001 | TRRAP | 7.28E-01 | 9.03E-01 | 9.85E-01 | -2.21E-02 | -1.02E+00 |
| ENSG000001 | KIF18A | 7.28E-01 | 9.03E-01 | 9.69E-01 | -4.52E-02 | -1.03E+00 |
| ENSG000002 | --- | 7.28E-01 | ? | 8.23E-01 | -2.82E-01 | -1.22E+00 |
| ENSG000002 | RUFY2 | 7.28E-01 | 9.03E-01 | 9.84E-01 | -2.30E-02 | -1.02E+00 |
| ENSG000002 | RHOQP1 | 7.29E-01 | ? | 1.08E+00 | 1.12E-01 | 1.08E+00 |
| ENSG000001 | ATP13A4 | 7.29E-01 | ? | 8.96E-01 | -1.59E-01 | -1.12E+00 |
| ENSG000001 | MTFR2 | 7.29E-01 | 9.03E-01 | 9.58E-01 | -6.26E-02 | -1.04E+00 |
| ENSG00000C | IFT74 | 7.29E-01 | 9.03E-01 | 9.82E-01 | -2.55E-02 | -1.02E+00 |
| ENSG000001 | C4orf17 | 7.29E-01 | ? | 8.21E-01 | -2.84E-01 | -1.22E+00 |
| ENSG000001 | AKR1E2 | 7.29E-01 | ? | 1.06E+00 | 9.08E-02 | 1.06E+00 |
| ENSG000001 | CPNE5 | 7.29E-01 | ? | 9.26E-01 | -1.11E-01 | -1.08E+00 |
| ENSG000001 | ZNF493 | 7.29E-01 | 9.03E-01 | 9.75E-01 | -3.71E-02 | -1.03E+00 |
| ENSG000002 | FAM24B | 7.29E-01 | 9.03E-01 | 1.03E+00 | 4.47E-02 | 1.03E+00 |
| ENSG000002 | --- | 7.29E-01 | 9.03E-01 | 9.36E-01 | -9.48E-02 | -1.07E+00 |
| ENSG000001 | C1R | 7.29E-01 | 9.03E-01 | 1.03E+00 | 4.21E-02 | 1.03E+00 |
| ENSG000002 | EIF4BP3 | 7.29E-01 | ? | 1.09E+00 | 1.24E-01 | 1.09E+00 |

| | | | | | | |
|---|---|---|---|---|---|---|
| ENSG000001 | PPIP5K1 | 7.29E-01 | 9.03E-01 | 1.01E+00 | 1.78E-02 | 1.01E+00 |
| ENSG000002 | PDCL3P5 | 7.29E-01 | ? | 1.13E+00 | 1.80E-01 | 1.13E+00 |
| ENSG000002 | --- | 7.29E-01 | ? | 8.95E-01 | -1.61E-01 | -1.12E+00 |
| ENSG000002 | RPL3P7 | 7.29E-01 | ? | 8.60E-01 | -2.17E-01 | -1.16E+00 |
| ENSG00000C | SUGP2 | 7.29E-01 | 9.03E-01 | 1.02E+00 | 2.21E-02 | 1.02E+00 |
| ENSG000001 | MRRF | 7.29E-01 | 9.03E-01 | 9.88E-01 | -1.74E-02 | -1.01E+00 |
| ENSG000001 | CCDC77 | 7.30E-01 | 9.03E-01 | 9.76E-01 | -3.52E-02 | -1.02E+00 |
| ENSG000001 | UGGT1 | 7.30E-01 | 9.03E-01 | 1.02E+00 | 2.19E-02 | 1.02E+00 |
| ENSG00000C | FARP2 | 7.30E-01 | 9.03E-01 | 1.01E+00 | 1.35E-02 | 1.01E+00 |
| ENSG000002 | --- | 7.30E-01 | 9.03E-01 | 9.33E-01 | -9.94E-02 | -1.07E+00 |
| ENSG000002 | --- | 7.30E-01 | ? | 9.04E-01 | -1.46E-01 | -1.11E+00 |
| ENSG000001 | LMTK3 | 7.30E-01 | 9.03E-01 | 9.78E-01 | -3.22E-02 | -1.02E+00 |
| ENSG00000C | OSBPL7 | 7.30E-01 | 9.03E-01 | 9.75E-01 | -3.66E-02 | -1.03E+00 |
| ENSG000001 | CENPA | 7.30E-01 | 9.03E-01 | 9.36E-01 | -9.59E-02 | -1.07E+00 |
| ENSG000002 | ZNF594-DT | 7.30E-01 | ? | 9.23E-01 | -1.16E-01 | -1.08E+00 |
| ENSG000002 | --- | 7.30E-01 | 9.04E-01 | 9.75E-01 | -3.61E-02 | -1.03E+00 |
| ENSG000001 | ATP6AP1-DT | 7.30E-01 | ? | 1.06E+00 | 8.88E-02 | 1.06E+00 |
| ENSG000002 | --- | 7.30E-01 | ? | 1.19E+00 | 2.47E-01 | 1.19E+00 |
| ENSG000001 | SH3BGR | 7.30E-01 | 9.04E-01 | 9.54E-01 | -6.87E-02 | -1.05E+00 |
| ENSG000002 | PRR5-ARHGA | 7.30E-01 | 9.04E-01 | 1.05E+00 | 6.50E-02 | 1.05E+00 |
| ENSG000002 | --- | 7.31E-01 | ? | 8.37E-01 | -2.57E-01 | -1.19E+00 |
| ENSG000001 | BCL2L12 | 7.31E-01 | 9.04E-01 | 1.03E+00 | 4.77E-02 | 1.03E+00 |
| ENSG000002 | --- | 7.31E-01 | 9.04E-01 | 9.67E-01 | -4.90E-02 | -1.03E+00 |
| ENSG000002 | GPRC5D-AS1 | 7.31E-01 | 9.04E-01 | 1.03E+00 | 3.98E-02 | 1.03E+00 |
| ENSG000002 | GBAP1 | 7.31E-01 | 9.04E-01 | 1.03E+00 | 3.90E-02 | 1.03E+00 |
| ENSG000001 | CTSD | 7.31E-01 | 9.04E-01 | 9.77E-01 | -3.40E-02 | -1.02E+00 |
| ENSG000001 | ZSCAN2 | 7.31E-01 | 9.04E-01 | 9.78E-01 | -3.22E-02 | -1.02E+00 |
| ENSG000002 | --- | 7.31E-01 | ? | 1.10E+00 | 1.40E-01 | 1.10E+00 |
| ENSG000002 | TRBV12-4 | 7.31E-01 | ? | 9.15E-01 | -1.27E-01 | -1.09E+00 |
| ENSG000002 | SRA1 | 7.31E-01 | 9.04E-01 | 1.01E+00 | 2.06E-02 | 1.01E+00 |
| ENSG000002 | --- | 7.31E-01 | ? | 1.14E+00 | 1.93E-01 | 1.14E+00 |
| ENSG000001 | LGALS7B | 7.31E-01 | 9.04E-01 | 1.06E+00 | 8.02E-02 | 1.06E+00 |
| ENSG000001 | AMMECR1L | 7.31E-01 | 9.04E-01 | 1.01E+00 | 1.44E-02 | 1.01E+00 |
| ENSG000001 | PRSS53 | 7.31E-01 | 9.04E-01 | 9.69E-01 | -4.54E-02 | -1.03E+00 |
| ENSG000001 | KCNJ14 | 7.31E-01 | 9.04E-01 | 9.69E-01 | -4.50E-02 | -1.03E+00 |
| ENSG000001 | MEI1 | 7.31E-01 | ? | 8.97E-01 | -1.57E-01 | -1.11E+00 |
| ENSG000001 | AP4S1 | 7.31E-01 | 9.04E-01 | 1.02E+00 | 3.47E-02 | 1.02E+00 |
| ENSG000001 | CBLB | 7.32E-01 | 9.04E-01 | 9.85E-01 | -2.25E-02 | -1.02E+00 |
| ENSG000001 | TMEM9B | 7.32E-01 | 9.04E-01 | 1.01E+00 | 1.25E-02 | 1.01E+00 |
| ENSG000002 | --- | 7.32E-01 | ? | 8.72E-01 | -1.97E-01 | -1.15E+00 |
| ENSG000002 | --- | 7.32E-01 | ? | 9.47E-01 | -7.91E-02 | -1.06E+00 |
| ENSG000001 | BMP8B | 7.32E-01 | ? | 1.07E+00 | 1.00E-01 | 1.07E+00 |
| ENSG000001 | SDK1 | 7.32E-01 | 9.04E-01 | 9.62E-01 | -5.66E-02 | -1.04E+00 |
| ENSG000002 | --- | 7.32E-01 | ? | 1.27E+00 | 3.40E-01 | 1.27E+00 |
| ENSG000001 | SVIP | 7.32E-01 | 9.04E-01 | 9.81E-01 | -2.77E-02 | -1.02E+00 |

| | | | | | | | |
|---|---|---|---|---|---|---|---|
| ENSG00000C | TNS1 | 7.32E-01 | | 9.04E-01 | 1.02E+00 | 2.55E-02 | 1.02E+00 |
| ENSG000002 | FAM88F | 7.32E-01 | ? | | 9.24E-01 | -1.15E-01 | -1.08E+00 |
| ENSG000001 | EFTUD2 | 7.32E-01 | | 9.04E-01 | 1.01E+00 | 1.58E-02 | 1.01E+00 |
| ENSG000001 | NBR2 | 7.32E-01 | | 9.04E-01 | 9.72E-01 | -4.04E-02 | -1.03E+00 |
| ENSG000001 | SSR1 | 7.32E-01 | | 9.04E-01 | 1.01E+00 | 1.77E-02 | 1.01E+00 |
| ENSG000002 | --- | 7.32E-01 | ? | | 1.08E+00 | 1.12E-01 | 1.08E+00 |
| ENSG00000C | BTBD7 | 7.32E-01 | | 9.04E-01 | 9.88E-01 | -1.79E-02 | -1.01E+00 |
| ENSG000001 | TDG | 7.32E-01 | | 9.04E-01 | 1.01E+00 | 1.71E-02 | 1.01E+00 |
| ENSG000001 | ABHD10 | 7.32E-01 | | 9.04E-01 | 1.02E+00 | 2.89E-02 | 1.02E+00 |
| ENSG000001 | PHETA1 | 7.33E-01 | | 9.05E-01 | 9.78E-01 | -3.18E-02 | -1.02E+00 |
| ENSG000002 | --- | 7.33E-01 | ? | | 8.16E-01 | -2.93E-01 | -1.23E+00 |
| ENSG000002 | RPL21P89 | 7.33E-01 | ? | | 8.29E-01 | -2.70E-01 | -1.21E+00 |
| ENSG000002 | AMACR | 7.33E-01 | | 9.05E-01 | 1.02E+00 | 2.16E-02 | 1.02E+00 |
| ENSG000002 | SMIM17 | 7.33E-01 | ? | | 8.92E-01 | -1.66E-01 | -1.12E+00 |
| ENSG000001 | MACC1 | 7.33E-01 | | 9.05E-01 | 1.02E+00 | 3.40E-02 | 1.02E+00 |
| ENSG00000C | FECH | 7.33E-01 | | 9.05E-01 | 9.89E-01 | -1.55E-02 | -1.01E+00 |
| ENSG000002 | --- | 7.33E-01 | ? | | 8.74E-01 | -1.95E-01 | -1.14E+00 |
| ENSG000001 | BAG2 | 7.33E-01 | | 9.05E-01 | 1.02E+00 | 2.79E-02 | 1.02E+00 |
| ENSG000001 | NUFIP2 | 7.33E-01 | | 9.05E-01 | 9.87E-01 | -1.89E-02 | -1.01E+00 |
| ENSG000001 | SERPINC1 | 7.33E-01 | ? | | 1.12E+00 | 1.65E-01 | 1.12E+00 |
| ENSG000001 | METRN | 7.33E-01 | | 9.05E-01 | 1.03E+00 | 4.36E-02 | 1.03E+00 |
| ENSG000001 | PLBD2 | 7.33E-01 | | 9.05E-01 | 9.83E-01 | -2.42E-02 | -1.02E+00 |
| ENSG000001 | ZNF22-AS1 | 7.33E-01 | ? | | 9.39E-01 | -9.15E-02 | -1.07E+00 |
| ENSG000002 | SPRR2A | 7.34E-01 | | 9.05E-01 | 9.42E-01 | -8.58E-02 | -1.06E+00 |
| ENSG000002 | --- | 7.34E-01 | ? | | 1.06E+00 | 8.55E-02 | 1.06E+00 |
| ENSG000001 | TEX14 | 7.34E-01 | ? | | 8.63E-01 | -2.13E-01 | -1.16E+00 |
| ENSG000002 | ARPC4-TTLL3 | 7.34E-01 | | 9.05E-01 | 9.47E-01 | -7.90E-02 | -1.06E+00 |
| ENSG000001 | CASP9 | 7.34E-01 | | 9.05E-01 | 1.02E+00 | 2.92E-02 | 1.02E+00 |
| ENSG000002 | --- | 7.34E-01 | ? | | 8.52E-01 | -2.31E-01 | -1.17E+00 |
| ENSG000002 | --- | 7.34E-01 | | 9.05E-01 | 1.05E+00 | 7.69E-02 | 1.05E+00 |
| ENSG000001 | UBR5 | 7.34E-01 | | 9.05E-01 | 1.02E+00 | 2.35E-02 | 1.02E+00 |
| ENSG000001 | GPRASP2 | 7.34E-01 | | 9.05E-01 | 1.02E+00 | 2.62E-02 | 1.02E+00 |
| ENSG000001 | GATA6 | 7.34E-01 | | 9.05E-01 | 9.84E-01 | -2.38E-02 | -1.02E+00 |
| ENSG000002 | HNRNPUL2-E | 7.34E-01 | | 9.05E-01 | 9.88E-01 | -1.79E-02 | -1.01E+00 |
| ENSG000002 | --- | 7.34E-01 | ? | | 8.19E-01 | -2.88E-01 | -1.22E+00 |
| ENSG000002 | --- | 7.34E-01 | | 9.05E-01 | 1.07E+00 | 9.29E-02 | 1.07E+00 |
| ENSG000001 | PRIM2 | 7.34E-01 | | 9.05E-01 | 9.82E-01 | -2.59E-02 | -1.02E+00 |
| ENSG00000C | SYDE2 | 7.34E-01 | | 9.05E-01 | 9.71E-01 | -4.32E-02 | -1.03E+00 |
| ENSG00000C | --- | 7.35E-01 | | 9.05E-01 | 1.10E+00 | 1.38E-01 | 1.10E+00 |
| ENSG000002 | LINC01572 | 7.35E-01 | | 9.05E-01 | 9.50E-01 | -7.39E-02 | -1.05E+00 |
| ENSG000001 | UEVLD | 7.35E-01 | | 9.05E-01 | 9.89E-01 | -1.56E-02 | -1.01E+00 |
| ENSG000001 | NPNT | 7.35E-01 | | 9.05E-01 | 9.66E-01 | -4.97E-02 | -1.04E+00 |
| ENSG00000C | ITCH | 7.35E-01 | | 9.05E-01 | 1.01E+00 | 1.33E-02 | 1.01E+00 |
| ENSG000002 | FMC1-LUC7L | 7.35E-01 | | 9.05E-01 | 9.71E-01 | -4.30E-02 | -1.03E+00 |
| ENSG00000C | CD59 | 7.35E-01 | | 9.05E-01 | 1.01E+00 | 1.52E-02 | 1.01E+00 |

| | | | | | | |
|---|---|---|---|---|---|---|
| ENSG000001 | CCDC60 | 7.35E-01 | ? | 1.08E+00 | 1.08E-01 | 1.08E+00 |
| ENSG000002 | --- | 7.35E-01 | ? | 8.82E-01 | -1.82E-01 | -1.13E+00 |
| ENSG000002 | --- | 7.35E-01 | ? | 8.93E-01 | -1.63E-01 | -1.12E+00 |
| ENSG00000C | DAPK2 | 7.35E-01 | 9.06E-01 | 1.05E+00 | 7.45E-02 | 1.05E+00 |
| ENSG00000C | CBFA2T2 | 7.35E-01 | 9.06E-01 | 1.02E+00 | 2.36E-02 | 1.02E+00 |
| ENSG000002 | --- | 7.35E-01 | ? | 1.09E+00 | 1.30E-01 | 1.09E+00 |
| ENSG000002 | --- | 7.35E-01 | ? | 1.10E+00 | 1.42E-01 | 1.10E+00 |
| ENSG00000C | UTP18 | 7.35E-01 | 9.06E-01 | 9.89E-01 | -1.60E-02 | -1.01E+00 |
| ENSG000002 | DHRS4L1 | 7.36E-01 | ? | 1.19E+00 | 2.53E-01 | 1.19E+00 |
| ENSG000001 | RNASEH2A | 7.36E-01 | 9.06E-01 | 9.71E-01 | -4.30E-02 | -1.03E+00 |
| ENSG000001 | YPEL5 | 7.36E-01 | 9.06E-01 | 1.01E+00 | 1.30E-02 | 1.01E+00 |
| ENSG000002 | SMC5-DT | 7.36E-01 | ? | 1.07E+00 | 9.64E-02 | 1.07E+00 |
| ENSG000001 | GLCE | 7.36E-01 | 9.06E-01 | 9.86E-01 | -2.11E-02 | -1.01E+00 |
| ENSG000002 | ZNF587B | 7.36E-01 | 9.06E-01 | 1.02E+00 | 2.22E-02 | 1.02E+00 |
| ENSG000002 | MRPL20-AS1 | 7.36E-01 | 9.06E-01 | 1.02E+00 | 2.73E-02 | 1.02E+00 |
| ENSG000001 | EP400 | 7.36E-01 | 9.06E-01 | 9.84E-01 | -2.26E-02 | -1.02E+00 |
| ENSG000001 | SNX7 | 7.36E-01 | 9.06E-01 | 9.87E-01 | -1.91E-02 | -1.01E+00 |
| ENSG000002 | GATD1-DT | 7.36E-01 | 9.06E-01 | 1.05E+00 | 6.41E-02 | 1.05E+00 |
| ENSG00000C | ACAP1 | 7.37E-01 | 9.06E-01 | 9.51E-01 | -7.29E-02 | -1.05E+00 |
| ENSG000001 | FLCN | 7.37E-01 | 9.06E-01 | 1.02E+00 | 2.93E-02 | 1.02E+00 |
| ENSG000001 | DMXL1 | 7.37E-01 | 9.06E-01 | 9.82E-01 | -2.61E-02 | -1.02E+00 |
| ENSG000002 | --- | 7.37E-01 | ? | 8.92E-01 | -1.65E-01 | -1.12E+00 |
| ENSG000002 | --- | 7.37E-01 | ? | 9.27E-01 | -1.09E-01 | -1.08E+00 |
| ENSG000002 | HLA-V | 7.37E-01 | 9.06E-01 | 9.69E-01 | -4.50E-02 | -1.03E+00 |
| ENSG000001 | ELMO1 | 7.37E-01 | ? | 1.11E+00 | 1.54E-01 | 1.11E+00 |
| ENSG000001 | LINC00482 | 7.37E-01 | ? | 8.69E-01 | -2.03E-01 | -1.15E+00 |
| ENSG000001 | IFNA1 | 7.37E-01 | ? | 8.51E-01 | -2.32E-01 | -1.17E+00 |
| ENSG00000C | CFLAR | 7.37E-01 | 9.06E-01 | 9.89E-01 | -1.53E-02 | -1.01E+00 |
| ENSG000002 | --- | 7.37E-01 | ? | 8.54E-01 | -2.28E-01 | -1.17E+00 |
| ENSG000002 | MACC1-DT | 7.37E-01 | ? | 9.40E-01 | -8.94E-02 | -1.06E+00 |
| ENSG000001 | ICMT | 7.37E-01 | 9.06E-01 | 1.01E+00 | 1.57E-02 | 1.01E+00 |
| ENSG000001 | ZFYVE19 | 7.37E-01 | 9.06E-01 | 1.02E+00 | 2.42E-02 | 1.02E+00 |
| ENSG000001 | CBX4 | 7.37E-01 | 9.06E-01 | 9.81E-01 | -2.76E-02 | -1.02E+00 |
| ENSG000001 | SLC28A3 | 7.37E-01 | 9.06E-01 | 1.03E+00 | 4.21E-02 | 1.03E+00 |
| ENSG000001 | HOOK3 | 7.37E-01 | 9.06E-01 | 9.82E-01 | -2.60E-02 | -1.02E+00 |
| ENSG000001 | CCDC9 | 7.37E-01 | 9.06E-01 | 9.78E-01 | -3.16E-02 | -1.02E+00 |
| ENSG000002 | --- | 7.37E-01 | ? | 1.07E+00 | 9.84E-02 | 1.07E+00 |
| ENSG000001 | TOP1MT | 7.37E-01 | 9.06E-01 | 9.86E-01 | -2.00E-02 | -1.01E+00 |
| ENSG000002 | HACD2 | 7.37E-01 | 9.06E-01 | 1.01E+00 | 1.48E-02 | 1.01E+00 |
| ENSG000001 | TLK1 | 7.37E-01 | 9.07E-01 | 9.88E-01 | -1.71E-02 | -1.01E+00 |
| ENSG000001 | CALR | 7.38E-01 | 9.07E-01 | 9.86E-01 | -2.02E-02 | -1.01E+00 |
| ENSG000002 | --- | 7.38E-01 | ? | 8.66E-01 | -2.07E-01 | -1.15E+00 |
| ENSG00000C | CD6 | 7.38E-01 | ? | 8.94E-01 | -1.61E-01 | -1.12E+00 |
| ENSG000001 | FBXO33 | 7.38E-01 | 9.07E-01 | 9.88E-01 | -1.78E-02 | -1.01E+00 |
| ENSG000002 | EGLN3-AS1 | 7.38E-01 | ? | 1.10E+00 | 1.38E-01 | 1.10E+00 |

| | | | | | | |
|---|---|---|---|---|---|---|
| ENSG000002 | EGOT | 7.38E-01 | ? | 9.04E-01 | -1.46E-01 | -1.11E+00 |
| ENSG00000C | SARM1 | 7.38E-01 | 9.07E-01 | 1.03E+00 | 4.39E-02 | 1.03E+00 |
| ENSG000001 | LYG1 | 7.38E-01 | ? | 1.09E+00 | 1.25E-01 | 1.09E+00 |
| ENSG000002 | --- | 7.38E-01 | ? | 9.12E-01 | -1.32E-01 | -1.10E+00 |
| ENSG000001 | ARHGEF18 | 7.38E-01 | 9.07E-01 | 9.84E-01 | -2.39E-02 | -1.02E+00 |
| ENSG000002 | LINC02901 | 7.38E-01 | ? | 8.78E-01 | -1.88E-01 | -1.14E+00 |
| ENSG000001 | TEX56P | 7.38E-01 | ? | 1.13E+00 | 1.74E-01 | 1.13E+00 |
| ENSG000002 | --- | 7.38E-01 | ? | 9.40E-01 | -8.96E-02 | -1.06E+00 |
| ENSG000001 | STAB2 | 7.38E-01 | ? | 8.82E-01 | -1.82E-01 | -1.13E+00 |
| ENSG000001 | SNED1 | 7.38E-01 | 9.07E-01 | 9.48E-01 | -7.70E-02 | -1.05E+00 |
| ENSG000002 | BOLA2-SMG | 7.38E-01 | 9.07E-01 | 9.79E-01 | -3.05E-02 | -1.02E+00 |
| ENSG000002 | --- | 7.39E-01 | ? | 9.28E-01 | -1.09E-01 | -1.08E+00 |
| ENSG000002 | --- | 7.39E-01 | ? | 8.62E-01 | -2.15E-01 | -1.16E+00 |
| ENSG000001 | CDK10 | 7.39E-01 | 9.08E-01 | 1.02E+00 | 2.90E-02 | 1.02E+00 |
| ENSG000001 | TMX4 | 7.39E-01 | 9.08E-01 | 9.85E-01 | -2.14E-02 | -1.01E+00 |
| ENSG000001 | SRPK3 | 7.39E-01 | ? | 1.09E+00 | 1.19E-01 | 1.09E+00 |
| ENSG000001 | ENO4 | 7.39E-01 | ? | 1.15E+00 | 2.02E-01 | 1.15E+00 |
| ENSG000001 | SYCP3 | 7.39E-01 | ? | 8.92E-01 | -1.66E-01 | -1.12E+00 |
| ENSG000001 | FBXW5 | 7.39E-01 | 9.08E-01 | 1.02E+00 | 2.96E-02 | 1.02E+00 |
| ENSG000001 | SQSTM1 | 7.39E-01 | 9.08E-01 | 9.83E-01 | -2.40E-02 | -1.02E+00 |
| ENSG000002 | MIF-AS1 | 7.39E-01 | 9.08E-01 | 1.08E+00 | 1.06E-01 | 1.08E+00 |
| ENSG000001 | ZFYVE21 | 7.39E-01 | 9.08E-01 | 9.87E-01 | -1.92E-02 | -1.01E+00 |
| ENSG000002 | --- | 7.39E-01 | 9.08E-01 | 9.77E-01 | -3.39E-02 | -1.02E+00 |
| ENSG000002 | --- | 7.39E-01 | ? | 1.07E+00 | 9.31E-02 | 1.07E+00 |
| ENSG000002 | UBE2Q2P1 | 7.39E-01 | 9.08E-01 | 9.65E-01 | -5.17E-02 | -1.04E+00 |
| ENSG000001 | GPR173 | 7.39E-01 | ? | 9.09E-01 | -1.37E-01 | -1.10E+00 |
| ENSG000001 | EMC10 | 7.39E-01 | 9.08E-01 | 9.84E-01 | -2.34E-02 | -1.02E+00 |
| ENSG000001 | PRSS35 | 7.40E-01 | ? | 1.20E+00 | 2.61E-01 | 1.20E+00 |
| ENSG000002 | NOP56P3 | 7.40E-01 | ? | 1.13E+00 | 1.74E-01 | 1.13E+00 |
| ENSG000001 | SLC16A11 | 7.40E-01 | ? | 1.13E+00 | 1.79E-01 | 1.13E+00 |
| ENSG000002 | NBDY | 7.40E-01 | 9.08E-01 | 1.02E+00 | 2.28E-02 | 1.02E+00 |
| ENSG000001 | ZSCAN16 | 7.40E-01 | ? | 9.51E-01 | -7.21E-02 | -1.05E+00 |
| ENSG000001 | PFDN2 | 7.40E-01 | 9.08E-01 | 9.85E-01 | -2.20E-02 | -1.02E+00 |
| ENSG000002 | --- | 7.40E-01 | ? | 1.08E+00 | 1.14E-01 | 1.08E+00 |
| ENSG000001 | PIEZO2 | 7.40E-01 | ? | 9.24E-01 | -1.14E-01 | -1.08E+00 |
| ENSG000002 | NUP153-AS1 | 7.40E-01 | ? | 1.07E+00 | 1.01E-01 | 1.07E+00 |
| ENSG000001 | ACSS3 | 7.40E-01 | ? | 1.13E+00 | 1.77E-01 | 1.13E+00 |
| ENSG000001 | CENPB | 7.40E-01 | 9.08E-01 | 1.02E+00 | 2.40E-02 | 1.02E+00 |
| ENSG000001 | HDAC8 | 7.40E-01 | 9.08E-01 | 1.01E+00 | 1.94E-02 | 1.01E+00 |
| ENSG000002 | SYNJ2BP | 7.40E-01 | 9.08E-01 | 1.02E+00 | 2.34E-02 | 1.02E+00 |
| ENSG000002 | --- | 7.40E-01 | ? | 8.65E-01 | -2.08E-01 | -1.16E+00 |
| ENSG000001 | ZNF137P | 7.41E-01 | 9.08E-01 | 9.61E-01 | -5.67E-02 | -1.04E+00 |
| ENSG000001 | ODC1 | 7.41E-01 | 9.08E-01 | 9.85E-01 | -2.24E-02 | -1.02E+00 |
| ENSG000001 | PCGF1 | 7.41E-01 | 9.08E-01 | 1.02E+00 | 2.59E-02 | 1.02E+00 |
| ENSG000001 | CDA | 7.41E-01 | 9.08E-01 | 9.82E-01 | -2.56E-02 | -1.02E+00 |

| | | | | | | |
|---|---|---|---|---|---|---|
| ENSG000001 | DNM1 | 7.41E-01 | 9.08E-01 | 1.05E+00 | 6.76E-02 | 1.05E+00 |
| ENSG000002 | --- | 7.41E-01 | 9.08E-01 | 9.48E-01 | -7.71E-02 | -1.05E+00 |
| ENSG000001 | CLK2 | 7.41E-01 | 9.08E-01 | 9.82E-01 | -2.55E-02 | -1.02E+00 |
| ENSG000002 | --- | 7.41E-01 | ? | 8.62E-01 | -2.15E-01 | -1.16E+00 |
| ENSG000001 | OSBP2 | 7.41E-01 | 9.08E-01 | 9.48E-01 | -7.65E-02 | -1.05E+00 |
| ENSG00000C | HNRNPM | 7.41E-01 | 9.08E-01 | 9.93E-01 | -1.03E-02 | -1.01E+00 |
| ENSG000001 | DMAC2 | 7.41E-01 | 9.08E-01 | 9.90E-01 | -1.39E-02 | -1.01E+00 |
| ENSG000001 | ACTR2 | 7.41E-01 | 9.08E-01 | 9.90E-01 | -1.50E-02 | -1.01E+00 |
| ENSG000001 | G3BP2 | 7.41E-01 | 9.08E-01 | 1.01E+00 | 1.39E-02 | 1.01E+00 |
| ENSG00000C | MTAP | 7.41E-01 | 9.08E-01 | 9.87E-01 | -1.87E-02 | -1.01E+00 |
| ENSG000001 | RASA4 | 7.41E-01 | 9.08E-01 | 9.74E-01 | -3.78E-02 | -1.03E+00 |
| ENSG000001 | XPA | 7.41E-01 | 9.08E-01 | 1.02E+00 | 3.20E-02 | 1.02E+00 |
| ENSG000001 | BSG | 7.41E-01 | 9.08E-01 | 1.01E+00 | 1.93E-02 | 1.01E+00 |
| ENSG000001 | RNF167 | 7.41E-01 | 9.08E-01 | 1.01E+00 | 1.54E-02 | 1.01E+00 |
| ENSG000001 | MARS1 | 7.41E-01 | 9.08E-01 | 1.01E+00 | 1.32E-02 | 1.01E+00 |
| ENSG000001 | IMMT | 7.41E-01 | 9.08E-01 | 1.01E+00 | 1.37E-02 | 1.01E+00 |
| ENSG000001 | TRIM3 | 7.41E-01 | 9.08E-01 | 1.02E+00 | 2.49E-02 | 1.02E+00 |
| ENSG000001 | LRRC47 | 7.41E-01 | 9.08E-01 | 1.01E+00 | 1.98E-02 | 1.01E+00 |
| ENSG000002 | --- | 7.41E-01 | ? | 1.22E+00 | 2.93E-01 | 1.22E+00 |
| ENSG000001 | DHRS4 | 7.41E-01 | 9.08E-01 | 1.02E+00 | 2.72E-02 | 1.02E+00 |
| ENSG000001 | STK11 | 7.41E-01 | 9.08E-01 | 1.02E+00 | 2.52E-02 | 1.02E+00 |
| ENSG000001 | MCRS1 | 7.41E-01 | 9.08E-01 | 9.81E-01 | -2.76E-02 | -1.02E+00 |
| ENSG000001 | NDRG3 | 7.41E-01 | 9.08E-01 | 9.89E-01 | -1.61E-02 | -1.01E+00 |
| ENSG000001 | APPL2 | 7.41E-01 | 9.08E-01 | 9.85E-01 | -2.23E-02 | -1.02E+00 |
| ENSG000001 | REPS2 | 7.42E-01 | 9.08E-01 | 9.80E-01 | -2.98E-02 | -1.02E+00 |
| ENSG00000C | FHIP1B | 7.42E-01 | 9.08E-01 | 1.03E+00 | 3.70E-02 | 1.03E+00 |
| ENSG000001 | ACBD7 | 7.42E-01 | ? | 9.36E-01 | -9.52E-02 | -1.07E+00 |
| ENSG00000C | MTREX | 7.42E-01 | 9.08E-01 | 1.01E+00 | 1.13E-02 | 1.01E+00 |
| ENSG000001 | SPATA24 | 7.42E-01 | 9.08E-01 | 9.59E-01 | -6.08E-02 | -1.04E+00 |
| ENSG000002 | --- | 7.42E-01 | ? | 8.72E-01 | -1.98E-01 | -1.15E+00 |
| ENSG000001 | SYNPO | 7.42E-01 | 9.08E-01 | 1.01E+00 | 1.32E-02 | 1.01E+00 |
| ENSG000002 | SMG1P5 | 7.42E-01 | 9.08E-01 | 1.02E+00 | 2.78E-02 | 1.02E+00 |
| ENSG000001 | MSC | 7.42E-01 | ? | 1.06E+00 | 8.96E-02 | 1.06E+00 |
| ENSG000002 | RBM5-AS1 | 7.43E-01 | 9.09E-01 | 9.29E-01 | -1.06E-01 | -1.08E+00 |
| ENSG000001 | RTN2 | 7.43E-01 | 9.09E-01 | 9.80E-01 | -2.95E-02 | -1.02E+00 |
| ENSG000001 | CSTF2 | 7.43E-01 | 9.09E-01 | 1.02E+00 | 2.16E-02 | 1.02E+00 |
| ENSG000002 | TRIM34 | 7.43E-01 | 9.09E-01 | 1.02E+00 | 3.13E-02 | 1.02E+00 |
| ENSG000001 | ASCC3 | 7.43E-01 | 9.09E-01 | 9.88E-01 | -1.69E-02 | -1.01E+00 |
| ENSG000002 | SLC28A2-AS1 | 7.43E-01 | ? | 1.08E+00 | 1.13E-01 | 1.08E+00 |
| ENSG000002 | EXOSC3P1 | 7.43E-01 | ? | 8.98E-01 | -1.55E-01 | -1.11E+00 |
| ENSG000001 | TXN | 7.43E-01 | 9.09E-01 | 9.89E-01 | -1.60E-02 | -1.01E+00 |
| ENSG000001 | PPM1M | 7.43E-01 | 9.09E-01 | 1.03E+00 | 4.54E-02 | 1.03E+00 |
| ENSG000001 | KAT5 | 7.43E-01 | 9.09E-01 | 1.01E+00 | 1.91E-02 | 1.01E+00 |
| ENSG000001 | PPARGC1A | 7.43E-01 | ? | 1.12E+00 | 1.66E-01 | 1.12E+00 |
| ENSG00000C | USP28 | 7.43E-01 | 9.09E-01 | 1.01E+00 | 1.92E-02 | 1.01E+00 |

| | | | | | | |
|---|---|---|---|---|---|---|
| ENSG000001 | RGS16 | 7.43E-01 | 9.09E-01 | 1.04E+00 | 5.97E-02 | 1.04E+00 |
| ENSG000001 | SLC8A1 | 7.43E-01 | 9.09E-01 | 1.04E+00 | 5.34E-02 | 1.04E+00 |
| ENSG000001 | MPV17L | 7.43E-01 | ? | 1.08E+00 | 1.16E-01 | 1.08E+00 |
| ENSG000001 | GJA3 | 7.43E-01 | 9.09E-01 | 1.05E+00 | 6.56E-02 | 1.05E+00 |
| ENSG000001 | NNMT | 7.43E-01 | 9.09E-01 | 9.66E-01 | -4.95E-02 | -1.03E+00 |
| ENSG000001 | INPP1 | 7.43E-01 | 9.09E-01 | 9.86E-01 | -1.98E-02 | -1.01E+00 |
| ENSG000002 | MSANTD7 | 7.43E-01 | 9.09E-01 | 1.01E+00 | 1.70E-02 | 1.01E+00 |
| ENSG000001 | GMNN | 7.43E-01 | 9.09E-01 | 9.79E-01 | -3.01E-02 | -1.02E+00 |
| ENSG000002 | LINC00880 | 7.44E-01 | 9.09E-01 | 9.43E-01 | -8.45E-02 | -1.06E+00 |
| ENSG000001 | CHMP2A | 7.44E-01 | 9.09E-01 | 9.83E-01 | -2.42E-02 | -1.02E+00 |
| ENSG000001 | TBX18 | 7.44E-01 | 9.09E-01 | 1.02E+00 | 2.41E-02 | 1.02E+00 |
| ENSG000001 | --- | 7.44E-01 | ? | 9.24E-01 | -1.13E-01 | -1.08E+00 |
| ENSG000001 | SPRYD4 | 7.44E-01 | 9.09E-01 | 9.80E-01 | -2.86E-02 | -1.02E+00 |
| ENSG00000C | GNA11 | 7.44E-01 | 9.09E-01 | 9.83E-01 | -2.51E-02 | -1.02E+00 |
| ENSG000001 | CNPPD1 | 7.44E-01 | 9.09E-01 | 1.02E+00 | 2.50E-02 | 1.02E+00 |
| ENSG000001 | MANBA | 7.44E-01 | 9.09E-01 | 9.90E-01 | -1.47E-02 | -1.01E+00 |
| ENSG000001 | PTCD3 | 7.44E-01 | 9.09E-01 | 1.01E+00 | 1.56E-02 | 1.01E+00 |
| ENSG000001 | ARMCX5 | 7.44E-01 | 9.09E-01 | 9.86E-01 | -2.05E-02 | -1.01E+00 |
| ENSG000001 | PES1 | 7.44E-01 | 9.09E-01 | 1.01E+00 | 1.66E-02 | 1.01E+00 |
| ENSG000002 | TMEM150C | 7.44E-01 | 9.09E-01 | 9.67E-01 | -4.79E-02 | -1.03E+00 |
| ENSG000002 | DNAJC9 | 7.44E-01 | 9.09E-01 | 9.83E-01 | -2.45E-02 | -1.02E+00 |
| ENSG000001 | NOMO2 | 7.44E-01 | 9.09E-01 | 9.90E-01 | -1.47E-02 | -1.01E+00 |
| ENSG000001 | USP21 | 7.44E-01 | 9.09E-01 | 1.02E+00 | 2.66E-02 | 1.02E+00 |
| ENSG000001 | RNF130 | 7.44E-01 | 9.09E-01 | 1.01E+00 | 1.73E-02 | 1.01E+00 |
| ENSG000001 | PLEKHG3 | 7.44E-01 | 9.09E-01 | 1.01E+00 | 1.89E-02 | 1.01E+00 |
| ENSG000002 | --- | 7.45E-01 | 9.09E-01 | 1.20E+00 | 2.60E-01 | 1.20E+00 |
| ENSG000001 | COL4A3 | 7.45E-01 | ? | 9.37E-01 | -9.42E-02 | -1.07E+00 |
| ENSG000001 | SMG9 | 7.45E-01 | 9.09E-01 | 1.01E+00 | 2.01E-02 | 1.01E+00 |
| ENSG000001 | MBD1 | 7.45E-01 | 9.09E-01 | 9.92E-01 | -1.21E-02 | -1.01E+00 |
| ENSG000001 | PPIL6 | 7.45E-01 | 9.09E-01 | 1.05E+00 | 7.72E-02 | 1.05E+00 |
| ENSG000001 | GRIN3A | 7.45E-01 | ? | 9.20E-01 | -1.21E-01 | -1.09E+00 |
| ENSG000002 | --- | 7.45E-01 | 9.09E-01 | 9.75E-01 | -3.63E-02 | -1.03E+00 |
| ENSG000001 | ALG12 | 7.45E-01 | 9.09E-01 | 1.02E+00 | 3.02E-02 | 1.02E+00 |
| ENSG000001 | C5AR2 | 7.45E-01 | ? | 1.12E+00 | 1.61E-01 | 1.12E+00 |
| ENSG000001 | GCK | 7.45E-01 | ? | 8.68E-01 | -2.04E-01 | -1.15E+00 |
| ENSG000002 | --- | 7.45E-01 | ? | 1.08E+00 | 1.06E-01 | 1.08E+00 |
| ENSG000002 | CDH12P1 | 7.45E-01 | ? | 1.31E+00 | 3.84E-01 | 1.31E+00 |
| ENSG000002 | CDH12P3 | 7.45E-01 | ? | 1.31E+00 | 3.84E-01 | 1.31E+00 |
| ENSG000001 | CHRNB4 | 7.45E-01 | ? | 9.40E-01 | -8.91E-02 | -1.06E+00 |
| ENSG000002 | DOK6 | 7.45E-01 | ? | 1.13E+00 | 1.71E-01 | 1.13E+00 |
| ENSG000001 | JCAD | 7.45E-01 | 9.09E-01 | 9.62E-01 | -5.65E-02 | -1.04E+00 |
| ENSG000002 | RSC1A1 | 7.45E-01 | 9.09E-01 | 1.04E+00 | 6.21E-02 | 1.04E+00 |
| ENSG000001 | AKR7A3 | 7.46E-01 | ? | 8.75E-01 | -1.93E-01 | -1.14E+00 |
| ENSG000001 | C5orf63 | 7.46E-01 | 9.09E-01 | 9.71E-01 | -4.22E-02 | -1.03E+00 |
| ENSG000002 | --- | 7.46E-01 | ? | 1.16E+00 | 2.10E-01 | 1.16E+00 |

| | | | | | | |
|---|---|---|---|---|---|---|
| ENSG000002 | FANK1 | 7.46E-01 | 9.09E-01 | 1.02E+00 | 2.99E-02 | 1.02E+00 |
| ENSG000001 | DTX3 | 7.46E-01 | 9.09E-01 | 1.03E+00 | 3.75E-02 | 1.03E+00 |
| ENSG000002 | --- | 7.46E-01 | 9.09E-01 | 9.49E-01 | -7.60E-02 | -1.05E+00 |
| ENSG000002 | SNURF | 7.46E-01 | 9.09E-01 | 9.53E-01 | -6.99E-02 | -1.05E+00 |
| ENSG000002 | --- | 7.46E-01 | 9.09E-01 | 1.15E+00 | 2.07E-01 | 1.15E+00 |
| ENSG000001 | TNFSF10 | 7.46E-01 | 9.09E-01 | 9.82E-01 | -2.57E-02 | -1.02E+00 |
| ENSG000002 | --- | 7.46E-01 | ? | 9.19E-01 | -1.21E-01 | -1.09E+00 |
| ENSG000002 | LRTOMT | 7.46E-01 | ? | 9.45E-01 | -8.09E-02 | -1.06E+00 |
| ENSG000001 | WDR88 | 7.46E-01 | ? | 8.77E-01 | -1.89E-01 | -1.14E+00 |
| ENSG000001 | SLC9A8 | 7.46E-01 | 9.10E-01 | 9.86E-01 | -2.03E-02 | -1.01E+00 |
| ENSG000002 | --- | 7.47E-01 | ? | 1.17E+00 | 2.26E-01 | 1.17E+00 |
| ENSG000002 | MYO19 | 7.47E-01 | 9.10E-01 | 9.87E-01 | -1.85E-02 | -1.01E+00 |
| ENSG000001 | SYNPO2 | 7.47E-01 | 9.10E-01 | 9.73E-01 | -3.98E-02 | -1.03E+00 |
| ENSG000001 | ZNF773 | 7.47E-01 | 9.10E-01 | 1.02E+00 | 2.96E-02 | 1.02E+00 |
| ENSG000001 | ZNF101 | 7.47E-01 | 9.10E-01 | 9.84E-01 | -2.34E-02 | -1.02E+00 |
| ENSG000001 | PODXL2 | 7.47E-01 | 9.10E-01 | 1.04E+00 | 5.41E-02 | 1.04E+00 |
| ENSG000001 | RC3H1 | 7.47E-01 | 9.10E-01 | 1.02E+00 | 2.58E-02 | 1.02E+00 |
| ENSG000001 | PRKCG | 7.47E-01 | ? | 1.22E+00 | 2.82E-01 | 1.22E+00 |
| ENSG000001 | COX4I1 | 7.47E-01 | 9.10E-01 | 1.01E+00 | 1.28E-02 | 1.01E+00 |
| ENSG000001 | ARHGAP30 | 7.47E-01 | ? | 9.38E-01 | -9.21E-02 | -1.07E+00 |
| ENSG000001 | WRN | 7.47E-01 | 9.10E-01 | 1.01E+00 | 1.60E-02 | 1.01E+00 |
| ENSG000001 | SIX2 | 7.47E-01 | 9.10E-01 | 9.82E-01 | -2.63E-02 | -1.02E+00 |
| ENSG000002 | RNPS1P1 | 7.47E-01 | ? | 8.95E-01 | -1.61E-01 | -1.12E+00 |
| ENSG000002 | SNX18P3 | 7.47E-01 | ? | 8.99E-01 | -1.53E-01 | -1.11E+00 |
| ENSG000001 | SCPEP1 | 7.47E-01 | 9.10E-01 | 9.84E-01 | -2.33E-02 | -1.02E+00 |
| ENSG000002 | --- | 7.47E-01 | ? | 1.08E+00 | 1.14E-01 | 1.08E+00 |
| ENSG000002 | LINC01748 | 7.47E-01 | ? | 8.80E-01 | -1.84E-01 | -1.14E+00 |
| ENSG000002 | RNPS1 | 7.48E-01 | 9.10E-01 | 9.91E-01 | -1.30E-02 | -1.01E+00 |
| ENSG000001 | ACAD9 | 7.48E-01 | 9.10E-01 | 9.90E-01 | -1.51E-02 | -1.01E+00 |
| ENSG000002 | MAST4-AS1 | 7.48E-01 | ? | 1.11E+00 | 1.47E-01 | 1.11E+00 |
| ENSG000002 | PCDHA12 | 7.48E-01 | 9.10E-01 | 1.04E+00 | 5.67E-02 | 1.04E+00 |
| ENSG000001 | FBXO15 | 7.48E-01 | ? | 1.08E+00 | 1.16E-01 | 1.08E+00 |
| ENSG000000 | C1orf112 | 7.48E-01 | 9.10E-01 | 9.80E-01 | -2.94E-02 | -1.02E+00 |
| ENSG000001 | PGBD2 | 7.48E-01 | 9.10E-01 | 1.03E+00 | 3.67E-02 | 1.03E+00 |
| ENSG000001 | PLEKHG5 | 7.48E-01 | 9.10E-01 | 1.02E+00 | 2.55E-02 | 1.02E+00 |
| ENSG000001 | RASAL1 | 7.48E-01 | ? | 1.15E+00 | 2.02E-01 | 1.15E+00 |
| ENSG000002 | LINC00623 | 7.48E-01 | 9.10E-01 | 1.02E+00 | 2.62E-02 | 1.02E+00 |
| ENSG000002 | SULT1A3 | 7.48E-01 | 9.10E-01 | 9.64E-01 | -5.34E-02 | -1.04E+00 |
| ENSG000001 | CMPK1 | 7.48E-01 | 9.10E-01 | 1.01E+00 | 1.49E-02 | 1.01E+00 |
| ENSG000001 | ZNF830 | 7.48E-01 | 9.10E-01 | 1.02E+00 | 2.30E-02 | 1.02E+00 |
| ENSG000001 | SGCB | 7.48E-01 | 9.10E-01 | 1.02E+00 | 2.27E-02 | 1.02E+00 |
| ENSG000002 | KRT42P | 7.48E-01 | ? | 1.11E+00 | 1.44E-01 | 1.11E+00 |
| ENSG000001 | VKORC1L1 | 7.48E-01 | 9.10E-01 | 9.91E-01 | -1.29E-02 | -1.01E+00 |
| ENSG000002 | --- | 7.48E-01 | ? | 8.45E-01 | -2.42E-01 | -1.18E+00 |
| ENSG000001 | MORN3 | 7.48E-01 | 9.10E-01 | 1.05E+00 | 7.52E-02 | 1.05E+00 |

| | | | | | | |
|---|---|---|---|---|---|---|
| ENSG000002 | --- | 7.48E-01 | 9.10E-01 | 1.04E+00 | 5.63E-02 | 1.04E+00 |
| ENSG000001 | RFNG | 7.48E-01 | 9.10E-01 | 9.73E-01 | -3.93E-02 | -1.03E+00 |
| ENSG000002 | BASP1-AS1 | 7.48E-01 | ? | 1.12E+00 | 1.62E-01 | 1.12E+00 |
| ENSG00000C | SF3A1 | 7.48E-01 | 9.10E-01 | 9.94E-01 | -9.08E-03 | -1.01E+00 |
| ENSG000002 | RPSAP26 | 7.48E-01 | ? | 1.21E+00 | 2.80E-01 | 1.21E+00 |
| ENSG000001 | SLC7A6OS | 7.48E-01 | 9.10E-01 | 1.01E+00 | 1.70E-02 | 1.01E+00 |
| ENSG000002 | TMX2 | 7.48E-01 | 9.10E-01 | 1.01E+00 | 1.19E-02 | 1.01E+00 |
| ENSG000001 | ADGRG6 | 7.48E-01 | 9.10E-01 | 9.83E-01 | -2.44E-02 | -1.02E+00 |
| ENSG000001 | RPP38-DT | 7.48E-01 | ? | 1.14E+00 | 1.89E-01 | 1.14E+00 |
| ENSG000002 | --- | 7.48E-01 | ? | 9.02E-01 | -1.48E-01 | -1.11E+00 |
| ENSG000002 | --- | 7.48E-01 | 9.10E-01 | 1.05E+00 | 7.71E-02 | 1.05E+00 |
| ENSG000001 | FAM131C | 7.49E-01 | ? | 9.24E-01 | -1.14E-01 | -1.08E+00 |
| ENSG000002 | --- | 7.49E-01 | ? | 9.20E-01 | -1.20E-01 | -1.09E+00 |
| ENSG000002 | AGGF1P3 | 7.49E-01 | ? | 8.42E-01 | -2.49E-01 | -1.19E+00 |
| ENSG000001 | ETAA1 | 7.49E-01 | 9.10E-01 | 1.01E+00 | 2.15E-02 | 1.01E+00 |
| ENSG000002 | ZNRF2P1 | 7.49E-01 | ? | 1.09E+00 | 1.29E-01 | 1.09E+00 |
| ENSG000001 | STEAP1 | 7.49E-01 | 9.10E-01 | 9.76E-01 | -3.46E-02 | -1.02E+00 |
| ENSG000001 | KCTD14 | 7.49E-01 | 9.10E-01 | 9.75E-01 | -3.72E-02 | -1.03E+00 |
| ENSG000002 | --- | 7.49E-01 | ? | 8.50E-01 | -2.35E-01 | -1.18E+00 |
| ENSG000002 | YWHAEP5 | 7.49E-01 | ? | 1.19E+00 | 2.49E-01 | 1.19E+00 |
| ENSG000001 | PIGF | 7.50E-01 | 9.11E-01 | 1.01E+00 | 2.00E-02 | 1.01E+00 |
| ENSG000002 | TPT1P9 | 7.50E-01 | 9.11E-01 | 1.05E+00 | 7.36E-02 | 1.05E+00 |
| ENSG000002 | --- | 7.50E-01 | ? | 7.47E-01 | -4.20E-01 | -1.34E+00 |
| ENSG000002 | --- | 7.50E-01 | 9.11E-01 | 9.47E-01 | -7.81E-02 | -1.06E+00 |
| ENSG000001 | SPRY1 | 7.50E-01 | 9.11E-01 | 1.03E+00 | 4.69E-02 | 1.03E+00 |
| ENSG000001 | CD86 | 7.50E-01 | ? | 9.21E-01 | -1.19E-01 | -1.09E+00 |
| ENSG000002 | --- | 7.50E-01 | ? | 8.11E-01 | -3.03E-01 | -1.23E+00 |
| ENSG000001 | ZNF28 | 7.50E-01 | 9.11E-01 | 1.01E+00 | 2.02E-02 | 1.01E+00 |
| ENSG000001 | ROPN1L | 7.50E-01 | ? | 8.71E-01 | -1.99E-01 | -1.15E+00 |
| ENSG000002 | --- | 7.50E-01 | ? | 1.14E+00 | 1.95E-01 | 1.14E+00 |
| ENSG000001 | C9orf43 | 7.50E-01 | ? | 9.37E-01 | -9.38E-02 | -1.07E+00 |
| ENSG000001 | XRCC3 | 7.50E-01 | 9.11E-01 | 1.03E+00 | 4.28E-02 | 1.03E+00 |
| ENSG000002 | RNF5 | 7.50E-01 | 9.11E-01 | 1.02E+00 | 2.44E-02 | 1.02E+00 |
| ENSG000002 | LINC01714 | 7.50E-01 | ? | 1.17E+00 | 2.32E-01 | 1.17E+00 |
| ENSG000002 | --- | 7.50E-01 | 9.11E-01 | 1.05E+00 | 7.21E-02 | 1.05E+00 |
| ENSG000002 | --- | 7.50E-01 | ? | 1.08E+00 | 1.09E-01 | 1.08E+00 |
| ENSG000001 | C20orf204 | 7.50E-01 | ? | 9.19E-01 | -1.22E-01 | -1.09E+00 |
| ENSG000001 | TSC22D3 | 7.50E-01 | 9.11E-01 | 1.02E+00 | 3.40E-02 | 1.02E+00 |
| ENSG000002 | ZCWPW2 | 7.50E-01 | ? | 9.49E-01 | -7.62E-02 | -1.05E+00 |
| ENSG000001 | SLC19A3 | 7.50E-01 | ? | 9.24E-01 | -1.14E-01 | -1.08E+00 |
| ENSG000001 | SLC7A7 | 7.50E-01 | ? | 1.07E+00 | 1.01E-01 | 1.07E+00 |
| ENSG000002 | DINOL | 7.50E-01 | ? | 1.11E+00 | 1.50E-01 | 1.11E+00 |
| ENSG000001 | ZFP42 | 7.50E-01 | ? | 1.07E+00 | 9.97E-02 | 1.07E+00 |
| ENSG000001 | TET2 | 7.50E-01 | 9.11E-01 | 9.85E-01 | -2.18E-02 | -1.02E+00 |
| ENSG000002 | CCT6P3 | 7.51E-01 | 9.11E-01 | 9.48E-01 | -7.74E-02 | -1.06E+00 |

| | | | | | | |
|---|---|---|---|---|---|---|
| ENSG000002 | --- | 7.51E-01 | 9.11E-01 | 1.04E+00 | 6.10E-02 | 1.04E+00 |
| ENSG000001 | PMM2 | 7.51E-01 | 9.11E-01 | 9.90E-01 | -1.49E-02 | -1.01E+00 |
| ENSG000002 | CEBPA | 7.51E-01 | 9.11E-01 | 1.06E+00 | 8.65E-02 | 1.06E+00 |
| ENSG000001 | PRKAR1B | 7.51E-01 | 9.11E-01 | 1.02E+00 | 3.02E-02 | 1.02E+00 |
| ENSG000001 | KYNU | 7.51E-01 | 9.11E-01 | 9.74E-01 | -3.78E-02 | -1.03E+00 |
| ENSG000001 | RTL6 | 7.51E-01 | 9.11E-01 | 1.01E+00 | 1.37E-02 | 1.01E+00 |
| ENSG000002 | --- | 7.51E-01 | 9.11E-01 | 9.49E-01 | -7.49E-02 | -1.05E+00 |
| ENSG000001 | BRF2 | 7.51E-01 | 9.11E-01 | 1.02E+00 | 2.91E-02 | 1.02E+00 |
| ENSG000002 | --- | 7.51E-01 | 9.11E-01 | 1.13E+00 | 1.76E-01 | 1.13E+00 |
| ENSG000002 | --- | 7.51E-01 | ? | 8.94E-01 | -1.62E-01 | -1.12E+00 |
| ENSG000001 | CAPN12 | 7.51E-01 | 9.11E-01 | 9.64E-01 | -5.22E-02 | -1.04E+00 |
| ENSG000001 | IGFBPL1 | 7.51E-01 | ? | 1.07E+00 | 9.85E-02 | 1.07E+00 |
| ENSG000002 | LINC02015 | 7.51E-01 | ? | 1.07E+00 | 9.36E-02 | 1.07E+00 |
| ENSG000001 | BCL2L2 | 7.51E-01 | 9.11E-01 | 1.01E+00 | 1.74E-02 | 1.01E+00 |
| ENSG000001 | ZNF181 | 7.51E-01 | 9.11E-01 | 1.02E+00 | 3.06E-02 | 1.02E+00 |
| ENSG000002 | --- | 7.52E-01 | 9.12E-01 | 9.42E-01 | -8.63E-02 | -1.06E+00 |
| ENSG000001 | SPATA17 | 7.52E-01 | ? | 9.29E-01 | -1.06E-01 | -1.08E+00 |
| ENSG000001 | THNSL2 | 7.52E-01 | ? | 9.09E-01 | -1.37E-01 | -1.10E+00 |
| ENSG000001 | HMG20A | 7.52E-01 | 9.12E-01 | 9.89E-01 | -1.60E-02 | -1.01E+00 |
| ENSG000001 | LRRC17 | 7.52E-01 | ? | 9.23E-01 | -1.15E-01 | -1.08E+00 |
| ENSG000001 | ATP11C | 7.52E-01 | 9.12E-01 | 1.01E+00 | 1.87E-02 | 1.01E+00 |
| ENSG000001 | TMOD4 | 7.52E-01 | ? | 1.12E+00 | 1.57E-01 | 1.12E+00 |
| ENSG000001 | GJC2 | 7.52E-01 | ? | 8.78E-01 | -1.87E-01 | -1.14E+00 |
| ENSG000001 | NPAS1 | 7.52E-01 | ? | 8.95E-01 | -1.60E-01 | -1.12E+00 |
| ENSG000002 | MICB | 7.52E-01 | 9.12E-01 | 1.02E+00 | 2.73E-02 | 1.02E+00 |
| ENSG000001 | TTYH3 | 7.52E-01 | 9.12E-01 | 1.02E+00 | 2.43E-02 | 1.02E+00 |
| ENSG000001 | SLC30A10 | 7.52E-01 | ? | 1.14E+00 | 1.86E-01 | 1.14E+00 |
| ENSG000001 | CXCR6 | 7.52E-01 | ? | 9.13E-01 | -1.31E-01 | -1.10E+00 |
| ENSG000002 | --- | 7.52E-01 | ? | 1.18E+00 | 2.39E-01 | 1.18E+00 |
| ENSG000001 | AFAP1 | 7.53E-01 | 9.12E-01 | 1.01E+00 | 1.36E-02 | 1.01E+00 |
| ENSG000002 | --- | 7.53E-01 | 9.12E-01 | 9.56E-01 | -6.56E-02 | -1.05E+00 |
| ENSG000001 | LAMA4 | 7.53E-01 | ? | 8.92E-01 | -1.64E-01 | -1.12E+00 |
| ENSG000001 | SRMS | 7.53E-01 | ? | 8.83E-01 | -1.79E-01 | -1.13E+00 |
| ENSG000002 | MDP1 | 7.53E-01 | 9.13E-01 | 9.55E-01 | -6.71E-02 | -1.05E+00 |
| ENSG000001 | NFU1 | 7.53E-01 | 9.13E-01 | 9.87E-01 | -1.94E-02 | -1.01E+00 |
| ENSG000001 | DUSP19 | 7.53E-01 | 9.13E-01 | 9.53E-01 | -7.01E-02 | -1.05E+00 |
| ENSG000002 | RFX5-AS1 | 7.53E-01 | ? | 9.10E-01 | -1.35E-01 | -1.10E+00 |
| ENSG000002 | --- | 7.53E-01 | ? | 1.08E+00 | 1.16E-01 | 1.08E+00 |
| ENSG000001 | PIGA | 7.53E-01 | 9.13E-01 | 9.90E-01 | -1.52E-02 | -1.01E+00 |
| ENSG000001 | RAB2B | 7.53E-01 | 9.13E-01 | 9.87E-01 | -1.93E-02 | -1.01E+00 |
| ENSG000002 | ZNF486 | 7.53E-01 | 9.13E-01 | 1.02E+00 | 2.25E-02 | 1.02E+00 |
| ENSG000001 | CRCT1 | 7.53E-01 | 9.13E-01 | 1.05E+00 | 7.17E-02 | 1.05E+00 |
| ENSG000002 | --- | 7.53E-01 | ? | 9.06E-01 | -1.43E-01 | -1.10E+00 |
| ENSG000001 | SLC22A20P | 7.53E-01 | 9.13E-01 | 1.04E+00 | 5.90E-02 | 1.04E+00 |
| ENSG000001 | UPK1B | 7.53E-01 | 9.13E-01 | 1.06E+00 | 7.80E-02 | 1.06E+00 |

| | | | | | | |
|---|---|---|---|---|---|---|
| ENSG00000C | IRAG1 | 7.53E-01 | 9.13E-01 | 1.04E+00 | 5.55E-02 | 1.04E+00 |
| ENSG000002 | LENG8-AS1 | 7.54E-01 | 9.13E-01 | 9.37E-01 | -9.34E-02 | -1.07E+00 |
| ENSG000002 | --- | 7.54E-01 | ? | 1.05E+00 | 7.61E-02 | 1.05E+00 |
| ENSG000001 | KIFC3 | 7.54E-01 | 9.13E-01 | 1.02E+00 | 2.39E-02 | 1.02E+00 |
| ENSG000001 | RIPK1 | 7.54E-01 | 9.13E-01 | 1.01E+00 | 1.36E-02 | 1.01E+00 |
| ENSG000002 | --- | 7.54E-01 | ? | 9.03E-01 | -1.47E-01 | -1.11E+00 |
| ENSG000002 | --- | 7.54E-01 | ? | 1.06E+00 | 8.23E-02 | 1.06E+00 |
| ENSG000001 | TJAP1 | 7.54E-01 | 9.13E-01 | 9.85E-01 | -2.18E-02 | -1.02E+00 |
| ENSG00000C | ACACB | 7.54E-01 | ? | 9.39E-01 | -9.14E-02 | -1.07E+00 |
| ENSG00000C | MYO16 | 7.54E-01 | ? | 1.06E+00 | 8.02E-02 | 1.06E+00 |
| ENSG000002 | --- | 7.54E-01 | 9.13E-01 | 9.65E-01 | -5.18E-02 | -1.04E+00 |
| ENSG000001 | AMZ1 | 7.54E-01 | ? | 1.10E+00 | 1.33E-01 | 1.10E+00 |
| ENSG000002 | --- | 7.54E-01 | 9.13E-01 | 1.05E+00 | 7.48E-02 | 1.05E+00 |
| ENSG000001 | SNX32 | 7.54E-01 | 9.13E-01 | 1.04E+00 | 5.20E-02 | 1.04E+00 |
| ENSG00000C | KMT2E | 7.54E-01 | 9.13E-01 | 9.84E-01 | -2.26E-02 | -1.02E+00 |
| ENSG000001 | FLG2 | 7.54E-01 | ? | 1.10E+00 | 1.36E-01 | 1.10E+00 |
| ENSG000001 | DEFB4A | 7.54E-01 | ? | 8.61E-01 | -2.16E-01 | -1.16E+00 |
| ENSG000001 | ADAMTSL3 | 7.54E-01 | ? | 1.13E+00 | 1.76E-01 | 1.13E+00 |
| ENSG000001 | BORCS5 | 7.54E-01 | 9.13E-01 | 9.81E-01 | -2.79E-02 | -1.02E+00 |
| ENSG000002 | --- | 7.54E-01 | ? | 9.18E-01 | -1.24E-01 | -1.09E+00 |
| ENSG000002 | HSD17B7P2 | 7.54E-01 | 9.13E-01 | 1.04E+00 | 5.79E-02 | 1.04E+00 |
| ENSG000002 | UGT1A7 | 7.54E-01 | ? | 8.97E-01 | -1.57E-01 | -1.12E+00 |
| ENSG000002 | --- | 7.55E-01 | 9.13E-01 | 1.05E+00 | 6.61E-02 | 1.05E+00 |
| ENSG000001 | EFHD1 | 7.55E-01 | ? | 1.15E+00 | 1.99E-01 | 1.15E+00 |
| ENSG000001 | TLCD1 | 7.55E-01 | 9.13E-01 | 1.02E+00 | 3.30E-02 | 1.02E+00 |
| ENSG000001 | PYGO1 | 7.55E-01 | 9.13E-01 | 9.74E-01 | -3.87E-02 | -1.03E+00 |
| ENSG000001 | MARCHF3 | 7.55E-01 | ? | 9.22E-01 | -1.17E-01 | -1.08E+00 |
| ENSG000002 | --- | 7.55E-01 | ? | 8.97E-01 | -1.57E-01 | -1.12E+00 |
| ENSG000002 | XNDC1N-ZNI | 7.55E-01 | ? | 1.25E+00 | 3.18E-01 | 1.25E+00 |
| ENSG000001 | POLR3GL | 7.55E-01 | 9.13E-01 | 9.84E-01 | -2.27E-02 | -1.02E+00 |
| ENSG000002 | PDK1-AS1 | 7.55E-01 | 9.13E-01 | 9.72E-01 | -4.03E-02 | -1.03E+00 |
| ENSG000001 | CCL22 | 7.55E-01 | ? | 1.14E+00 | 1.89E-01 | 1.14E+00 |
| ENSG000002 | --- | 7.55E-01 | 9.13E-01 | 9.22E-01 | -1.17E-01 | -1.08E+00 |
| ENSG00000C | DIMT1 | 7.55E-01 | 9.13E-01 | 1.01E+00 | 1.58E-02 | 1.01E+00 |
| ENSG000002 | RN7SK | 7.55E-01 | ? | 1.09E+00 | 1.30E-01 | 1.09E+00 |
| ENSG000001 | AMD1 | 7.55E-01 | 9.13E-01 | 9.92E-01 | -1.22E-02 | -1.01E+00 |
| ENSG000001 | ARPC3 | 7.55E-01 | 9.13E-01 | 9.93E-01 | -1.00E-02 | -1.01E+00 |
| ENSG000002 | HNRNPLP2 | 7.55E-01 | ? | 1.07E+00 | 1.01E-01 | 1.07E+00 |
| ENSG000002 | SOCS2-AS1 | 7.55E-01 | ? | 9.14E-01 | -1.30E-01 | -1.09E+00 |
| ENSG000001 | PEX11G | 7.55E-01 | ? | 1.07E+00 | 9.21E-02 | 1.07E+00 |
| ENSG000002 | FTLP3 | 7.56E-01 | 9.14E-01 | 9.68E-01 | -4.68E-02 | -1.03E+00 |
| ENSG000001 | PIH1D2 | 7.56E-01 | 9.14E-01 | 1.05E+00 | 6.60E-02 | 1.05E+00 |
| ENSG000001 | GLB1L | 7.56E-01 | 9.14E-01 | 9.68E-01 | -4.65E-02 | -1.03E+00 |
| ENSG000001 | GTF3A | 7.56E-01 | 9.14E-01 | 1.01E+00 | 1.77E-02 | 1.01E+00 |
| ENSG000002 | --- | 7.56E-01 | 9.14E-01 | 9.63E-01 | -5.46E-02 | -1.04E+00 |

| | | | | | | |
|---|---|---|---|---|---|---|
| ENSG000002 | --- | 7.56E-01 | ? | 9.50E-01 | -7.42E-02 | -1.05E+00 |
| ENSG000002 | TAF1A-AS1 | 7.56E-01 | ? | 9.46E-01 | -7.98E-02 | -1.06E+00 |
| ENSG000001 | LGALS9B | 7.56E-01 | 9.14E-01 | 9.63E-01 | -5.40E-02 | -1.04E+00 |
| ENSG000002 | --- | 7.56E-01 | ? | 1.14E+00 | 1.93E-01 | 1.14E+00 |
| ENSG000001 | KCNJ6 | 7.56E-01 | ? | 8.56E-01 | -2.25E-01 | -1.17E+00 |
| ENSG000002 | ZNF580 | 7.56E-01 | 9.14E-01 | 9.78E-01 | -3.16E-02 | -1.02E+00 |
| ENSG000002 | ALG3 | 7.56E-01 | 9.14E-01 | 9.88E-01 | -1.81E-02 | -1.01E+00 |
| ENSG000001 | NDUFA11 | 7.56E-01 | 9.14E-01 | 1.01E+00 | 2.05E-02 | 1.01E+00 |
| ENSG000001 | C9orf64 | 7.56E-01 | 9.14E-01 | 1.01E+00 | 2.11E-02 | 1.01E+00 |
| ENSG000001 | H4C3 | 7.56E-01 | ? | 8.66E-01 | -2.08E-01 | -1.15E+00 |
| ENSG000001 | ARL5A | 7.56E-01 | 9.14E-01 | 9.82E-01 | -2.56E-02 | -1.02E+00 |
| ENSG000002 | --- | 7.57E-01 | ? | 1.13E+00 | 1.78E-01 | 1.13E+00 |
| ENSG000002 | --- | 7.57E-01 | ? | 1.08E+00 | 1.06E-01 | 1.08E+00 |
| ENSG000001 | TCAIM | 7.57E-01 | 9.14E-01 | 1.01E+00 | 1.89E-02 | 1.01E+00 |
| ENSG000002 | GAPDHP65 | 7.57E-01 | ? | 9.14E-01 | -1.30E-01 | -1.09E+00 |
| ENSG000002 | LIPT2-AS1 | 7.57E-01 | 9.14E-01 | 9.66E-01 | -4.93E-02 | -1.03E+00 |
| ENSG000001 | PGBD5 | 7.57E-01 | 9.14E-01 | 9.65E-01 | -5.21E-02 | -1.04E+00 |
| ENSG000002 | --- | 7.57E-01 | ? | 9.10E-01 | -1.36E-01 | -1.10E+00 |
| ENSG000001 | MYOM3 | 7.57E-01 | 9.14E-01 | 9.57E-01 | -6.33E-02 | -1.04E+00 |
| ENSG000001 | SCG2 | 7.57E-01 | ? | 9.37E-01 | -9.45E-02 | -1.07E+00 |
| ENSG000001 | PYCR2 | 7.57E-01 | 9.14E-01 | 9.82E-01 | -2.69E-02 | -1.02E+00 |
| ENSG000002 | --- | 7.57E-01 | ? | 1.08E+00 | 1.11E-01 | 1.08E+00 |
| ENSG000001 | C6orf141 | 7.57E-01 | 9.14E-01 | 1.02E+00 | 3.52E-02 | 1.02E+00 |
| ENSG000001 | TBP | 7.57E-01 | 9.14E-01 | 9.86E-01 | -2.01E-02 | -1.01E+00 |
| ENSG00000C | P2RX6 | 7.57E-01 | 9.14E-01 | 1.03E+00 | 4.25E-02 | 1.03E+00 |
| ENSG00000C | SPAG4 | 7.57E-01 | 9.14E-01 | 9.55E-01 | -6.68E-02 | -1.05E+00 |
| ENSG000001 | SIAE | 7.57E-01 | 9.14E-01 | 1.01E+00 | 1.54E-02 | 1.01E+00 |
| ENSG000001 | CTNNAL1 | 7.57E-01 | 9.14E-01 | 9.82E-01 | -2.67E-02 | -1.02E+00 |
| ENSG000001 | DOT1L | 7.57E-01 | 9.14E-01 | 1.02E+00 | 2.34E-02 | 1.02E+00 |
| ENSG000001 | SP6 | 7.57E-01 | 9.14E-01 | 9.76E-01 | -3.43E-02 | -1.02E+00 |
| ENSG000002 | --- | 7.58E-01 | 9.14E-01 | 1.04E+00 | 5.92E-02 | 1.04E+00 |
| ENSG000002 | ZNF888-AS1 | 7.58E-01 | ? | 1.19E+00 | 2.48E-01 | 1.19E+00 |
| ENSG000002 | FAM27E3 | 7.58E-01 | 9.14E-01 | 1.05E+00 | 7.00E-02 | 1.05E+00 |
| ENSG00000C | ANO8 | 7.58E-01 | 9.15E-01 | 9.78E-01 | -3.15E-02 | -1.02E+00 |
| ENSG000002 | --- | 7.58E-01 | ? | 9.21E-01 | -1.19E-01 | -1.09E+00 |
| ENSG000002 | --- | 7.58E-01 | ? | 9.33E-01 | -1.01E-01 | -1.07E+00 |
| ENSG00000C | RBM7 | 7.58E-01 | 9.15E-01 | 1.01E+00 | 2.14E-02 | 1.01E+00 |
| ENSG000001 | P3H1 | 7.58E-01 | 9.15E-01 | 1.01E+00 | 1.87E-02 | 1.01E+00 |
| ENSG000001 | SNHG16 | 7.58E-01 | 9.15E-01 | 1.01E+00 | 1.69E-02 | 1.01E+00 |
| ENSG000001 | SPSB1 | 7.58E-01 | 9.15E-01 | 1.01E+00 | 1.53E-02 | 1.01E+00 |
| ENSG000001 | CBY1 | 7.58E-01 | 9.15E-01 | 1.01E+00 | 1.91E-02 | 1.01E+00 |
| ENSG000002 | BSG-AS1 | 7.58E-01 | ? | 1.16E+00 | 2.11E-01 | 1.16E+00 |
| ENSG000002 | COPG2IT1 | 7.58E-01 | ? | 8.97E-01 | -1.57E-01 | -1.11E+00 |
| ENSG00000C | SIDT1 | 7.59E-01 | 9.15E-01 | 1.03E+00 | 4.43E-02 | 1.03E+00 |
| ENSG000001 | RAMP2-AS1 | 7.59E-01 | ? | 8.76E-01 | -1.91E-01 | -1.14E+00 |

| | | | | | | |
|---|---|---|---|---|---|---|
| ENSG000002 | --- | 7.59E-01 | 9.15E-01 | 1.05E+00 | 7.12E-02 | 1.05E+00 |
| ENSG000001 | TGFB3 | 7.59E-01 | 9.15E-01 | 1.04E+00 | 5.31E-02 | 1.04E+00 |
| ENSG000001 | CHST15 | 7.59E-01 | 9.15E-01 | 9.92E-01 | -1.09E-02 | -1.01E+00 |
| ENSG000001 | LRRC8E | 7.59E-01 | 9.15E-01 | 1.01E+00 | 2.05E-02 | 1.01E+00 |
| ENSG000001 | PP2D1 | 7.59E-01 | ? | 1.13E+00 | 1.75E-01 | 1.13E+00 |
| ENSG000001 | CLDN12 | 7.59E-01 | 9.15E-01 | 1.01E+00 | 1.76E-02 | 1.01E+00 |
| ENSG000001 | CCDC57 | 7.59E-01 | 9.15E-01 | 1.02E+00 | 3.10E-02 | 1.02E+00 |
| ENSG000002 | LINC00327 | 7.60E-01 | ? | 8.99E-01 | -1.54E-01 | -1.11E+00 |
| ENSG000001 | DNTTIP1 | 7.60E-01 | 9.15E-01 | 9.86E-01 | -2.09E-02 | -1.01E+00 |
| ENSG000001 | APCDD1L | 7.60E-01 | 9.15E-01 | 1.02E+00 | 2.84E-02 | 1.02E+00 |
| ENSG00000C | PITHD1 | 7.60E-01 | 9.15E-01 | 1.01E+00 | 1.18E-02 | 1.01E+00 |
| ENSG000002 | STRC | 7.60E-01 | ? | 1.07E+00 | 1.03E-01 | 1.07E+00 |
| ENSG00000C | PLPP1 | 7.60E-01 | 9.15E-01 | 9.85E-01 | -2.22E-02 | -1.02E+00 |
| ENSG000001 | ZNF75D | 7.60E-01 | 9.15E-01 | 1.02E+00 | 2.44E-02 | 1.02E+00 |
| ENSG000002 | --- | 7.60E-01 | ? | 1.17E+00 | 2.32E-01 | 1.17E+00 |
| ENSG000002 | --- | 7.60E-01 | ? | 9.02E-01 | -1.49E-01 | -1.11E+00 |
| ENSG000002 | --- | 7.60E-01 | ? | 9.16E-01 | -1.27E-01 | -1.09E+00 |
| ENSG000001 | GON4L | 7.60E-01 | 9.15E-01 | 9.91E-01 | -1.27E-02 | -1.01E+00 |
| ENSG000001 | ATP13A3 | 7.60E-01 | 9.15E-01 | 1.01E+00 | 1.58E-02 | 1.01E+00 |
| ENSG000001 | MTUS1 | 7.60E-01 | 9.15E-01 | 1.03E+00 | 4.96E-02 | 1.03E+00 |
| ENSG000001 | NMT1 | 7.60E-01 | 9.15E-01 | 1.01E+00 | 1.69E-02 | 1.01E+00 |
| ENSG00000C | SPPL2B | 7.60E-01 | 9.15E-01 | 1.02E+00 | 3.40E-02 | 1.02E+00 |
| ENSG00000C | CFTR | 7.60E-01 | ? | 8.84E-01 | -1.78E-01 | -1.13E+00 |
| ENSG000001 | GARNL3 | 7.60E-01 | 9.15E-01 | 9.76E-01 | -3.45E-02 | -1.02E+00 |
| ENSG00000C | MBTPS2 | 7.60E-01 | 9.15E-01 | 1.01E+00 | 1.86E-02 | 1.01E+00 |
| ENSG000001 | ABHD17B | 7.60E-01 | 9.15E-01 | 1.01E+00 | 1.85E-02 | 1.01E+00 |
| ENSG000002 | RPL4P6 | 7.60E-01 | ? | 9.48E-01 | -7.65E-02 | -1.05E+00 |
| ENSG000001 | IQCC | 7.60E-01 | 9.16E-01 | 1.03E+00 | 4.80E-02 | 1.03E+00 |
| ENSG000002 | --- | 7.60E-01 | ? | 1.08E+00 | 1.14E-01 | 1.08E+00 |
| ENSG000002 | PTPRD-AS1 | 7.61E-01 | ? | 1.12E+00 | 1.64E-01 | 1.12E+00 |
| ENSG000002 | GUSBP1 | 7.61E-01 | 9.16E-01 | 9.72E-01 | -4.04E-02 | -1.03E+00 |
| ENSG000001 | PLA2G3 | 7.61E-01 | ? | 9.10E-01 | -1.36E-01 | -1.10E+00 |
| ENSG000001 | SHKBP1 | 7.61E-01 | 9.16E-01 | 1.02E+00 | 2.73E-02 | 1.02E+00 |
| ENSG00000C | DPF1 | 7.61E-01 | 9.16E-01 | 1.05E+00 | 6.43E-02 | 1.05E+00 |
| ENSG000001 | CYREN | 7.61E-01 | 9.16E-01 | 1.02E+00 | 2.22E-02 | 1.02E+00 |
| ENSG000002 | --- | 7.61E-01 | ? | 7.63E-01 | -3.90E-01 | -1.31E+00 |
| ENSG000001 | PDGFD | 7.61E-01 | 9.16E-01 | 9.72E-01 | -4.10E-02 | -1.03E+00 |
| ENSG000002 | ERVH48-1 | 7.61E-01 | ? | 1.10E+00 | 1.35E-01 | 1.10E+00 |
| ENSG000002 | --- | 7.61E-01 | ? | 1.11E+00 | 1.50E-01 | 1.11E+00 |
| ENSG000002 | ADCY10P1 | 7.61E-01 | 9.16E-01 | 1.05E+00 | 6.82E-02 | 1.05E+00 |
| ENSG000001 | SLC8A3 | 7.61E-01 | ? | 9.28E-01 | -1.09E-01 | -1.08E+00 |
| ENSG000002 | TTC28-AS1 | 7.61E-01 | 9.16E-01 | 1.02E+00 | 3.39E-02 | 1.02E+00 |
| ENSG000002 | CD2AP-DT | 7.61E-01 | ? | 1.11E+00 | 1.46E-01 | 1.11E+00 |
| ENSG000002 | GSN-AS1 | 7.61E-01 | 9.16E-01 | 9.51E-01 | -7.27E-02 | -1.05E+00 |
| ENSG00000C | TNRC6A | 7.61E-01 | 9.16E-01 | 9.89E-01 | -1.54E-02 | -1.01E+00 |

| | | | | | | |
|---|---|---|---|---|---|---|
| ENSG000001 | ANXA9 | 7.61E-01 | 9.16E-01 | 9.55E-01 | -6.63E-02 | -1.05E+00 |
| ENSG000000 | SEC31B | 7.61E-01 | 9.16E-01 | 9.72E-01 | -4.03E-02 | -1.03E+00 |
| ENSG000002 | --- | 7.62E-01 | 9.16E-01 | 1.03E+00 | 3.57E-02 | 1.03E+00 |
| ENSG000001 | EEA1 | 7.62E-01 | 9.16E-01 | 1.02E+00 | 2.59E-02 | 1.02E+00 |
| ENSG000002 | --- | 7.62E-01 | ? | 1.15E+00 | 2.01E-01 | 1.15E+00 |
| ENSG000000 | ZMYND10 | 7.62E-01 | ? | 8.39E-01 | -2.53E-01 | -1.19E+00 |
| ENSG000001 | RNF122 | 7.62E-01 | 9.16E-01 | 9.78E-01 | -3.23E-02 | -1.02E+00 |
| ENSG000001 | THRB | 7.62E-01 | 9.16E-01 | 9.73E-01 | -3.88E-02 | -1.03E+00 |
| ENSG000002 | --- | 7.62E-01 | ? | 1.10E+00 | 1.37E-01 | 1.10E+00 |
| ENSG000001 | TMPRSS7 | 7.62E-01 | ? | 1.17E+00 | 2.22E-01 | 1.17E+00 |
| ENSG000001 | EIF5A | 7.62E-01 | 9.16E-01 | 1.01E+00 | 1.52E-02 | 1.01E+00 |
| ENSG000001 | NUDT16 | 7.62E-01 | 9.16E-01 | 1.01E+00 | 1.66E-02 | 1.01E+00 |
| ENSG000002 | EXOC3L2 | 7.62E-01 | ? | 9.20E-01 | -1.20E-01 | -1.09E+00 |
| ENSG000001 | SGK3 | 7.62E-01 | 9.16E-01 | 1.02E+00 | 2.41E-02 | 1.02E+00 |
| ENSG000001 | TRPM8 | 7.62E-01 | ? | 1.20E+00 | 2.61E-01 | 1.20E+00 |
| ENSG000001 | POT1 | 7.62E-01 | 9.16E-01 | 9.89E-01 | -1.54E-02 | -1.01E+00 |
| ENSG000002 | RNF223 | 7.62E-01 | ? | 9.35E-01 | -9.76E-02 | -1.07E+00 |
| ENSG000002 | --- | 7.62E-01 | ? | 9.32E-01 | -1.02E-01 | -1.07E+00 |
| ENSG000001 | PCSK1N | 7.62E-01 | ? | 1.07E+00 | 9.41E-02 | 1.07E+00 |
| ENSG000001 | NAPRT | 7.62E-01 | 9.16E-01 | 1.03E+00 | 4.02E-02 | 1.03E+00 |
| ENSG000002 | ABCC13 | 7.62E-01 | ? | 8.84E-01 | -1.79E-01 | -1.13E+00 |
| ENSG000001 | PTPRM | 7.62E-01 | 9.16E-01 | 9.89E-01 | -1.65E-02 | -1.01E+00 |
| ENSG000001 | SLC35E2B | 7.62E-01 | 9.16E-01 | 9.89E-01 | -1.56E-02 | -1.01E+00 |
| ENSG000001 | PYROXD2 | 7.62E-01 | 9.16E-01 | 9.74E-01 | -3.77E-02 | -1.03E+00 |
| ENSG000001 | ZNF577 | 7.62E-01 | 9.16E-01 | 9.75E-01 | -3.72E-02 | -1.03E+00 |
| ENSG000002 | --- | 7.62E-01 | ? | 9.27E-01 | -1.10E-01 | -1.08E+00 |
| ENSG000001 | SEC13 | 7.62E-01 | 9.16E-01 | 9.90E-01 | -1.48E-02 | -1.01E+00 |
| ENSG000002 | CFAP97D1 | 7.63E-01 | ? | 1.20E+00 | 2.66E-01 | 1.20E+00 |
| ENSG000001 | DOHH | 7.63E-01 | 9.16E-01 | 1.03E+00 | 4.90E-02 | 1.03E+00 |
| ENSG000001 | PNRC1 | 7.63E-01 | 9.16E-01 | 1.01E+00 | 1.81E-02 | 1.01E+00 |
| ENSG000001 | PDPK1 | 7.63E-01 | 9.16E-01 | 1.01E+00 | 1.00E-02 | 1.01E+00 |
| ENSG000001 | EEPD1 | 7.63E-01 | 9.16E-01 | 9.71E-01 | -4.17E-02 | -1.03E+00 |
| ENSG000001 | EIF1 | 7.63E-01 | 9.16E-01 | 1.01E+00 | 1.54E-02 | 1.01E+00 |
| ENSG000001 | YEATS4 | 7.63E-01 | 9.16E-01 | 9.86E-01 | -2.06E-02 | -1.01E+00 |
| ENSG000002 | MRPL45P2 | 7.63E-01 | ? | 9.44E-01 | -8.35E-02 | -1.06E+00 |
| ENSG000002 | RABGAP1L-D | 7.63E-01 | ? | 1.06E+00 | 8.41E-02 | 1.06E+00 |
| ENSG000001 | LRRC37A | 7.63E-01 | 9.16E-01 | 1.04E+00 | 6.16E-02 | 1.04E+00 |
| ENSG000001 | CPAMD8 | 7.63E-01 | ? | 1.05E+00 | 6.87E-02 | 1.05E+00 |
| ENSG000001 | RNF144A | 7.63E-01 | ? | 1.08E+00 | 1.11E-01 | 1.08E+00 |
| ENSG000001 | DHX9 | 7.63E-01 | 9.16E-01 | 1.01E+00 | 1.17E-02 | 1.01E+00 |
| ENSG000001 | ZNF546 | 7.63E-01 | 9.16E-01 | 9.64E-01 | -5.25E-02 | -1.04E+00 |
| ENSG000000 | NGFR | 7.63E-01 | ? | 1.08E+00 | 1.06E-01 | 1.08E+00 |
| ENSG000002 | MGAM2 | 7.63E-01 | ? | 9.30E-01 | -1.05E-01 | -1.08E+00 |
| ENSG000001 | UBA2 | 7.63E-01 | 9.16E-01 | 9.90E-01 | -1.44E-02 | -1.01E+00 |
| ENSG000000 | CAPN3 | 7.64E-01 | 9.16E-01 | 1.03E+00 | 3.78E-02 | 1.03E+00 |

| | | | | | | |
|---|---|---|---|---|---|---|
| ENSG000002 | --- | 7.64E-01 | ? | 8.90E-01 | -1.68E-01 | -1.12E+00 |
| ENSG000001 | PNRC2 | 7.64E-01 | 9.16E-01 | 1.01E+00 | 1.59E-02 | 1.01E+00 |
| ENSG000001 | HNF4G | 7.64E-01 | ? | 8.45E-01 | -2.43E-01 | -1.18E+00 |
| ENSG000002 | FAM218A | 7.64E-01 | ? | 1.06E+00 | 8.45E-02 | 1.06E+00 |
| ENSG000001 | PPM1E | 7.64E-01 | ? | 8.92E-01 | -1.64E-01 | -1.12E+00 |
| ENSG000001 | VWA2 | 7.64E-01 | 9.17E-01 | 1.02E+00 | 2.55E-02 | 1.02E+00 |
| ENSG000001 | VPS54 | 7.64E-01 | 9.17E-01 | 1.01E+00 | 1.67E-02 | 1.01E+00 |
| ENSG000002 | P2RY11 | 7.64E-01 | 9.17E-01 | 9.76E-01 | -3.53E-02 | -1.02E+00 |
| ENSG000001 | TIGD7 | 7.64E-01 | 9.17E-01 | 9.41E-01 | -8.70E-02 | -1.06E+00 |
| ENSG000001 | ZBTB43 | 7.64E-01 | 9.17E-01 | 1.02E+00 | 2.18E-02 | 1.02E+00 |
| ENSG000001 | CDK19 | 7.64E-01 | 9.17E-01 | 9.86E-01 | -2.00E-02 | -1.01E+00 |
| ENSG000002 | BCAN-AS1 | 7.64E-01 | ? | 9.06E-01 | -1.42E-01 | -1.10E+00 |
| ENSG000001 | FBXW9 | 7.64E-01 | 9.17E-01 | 1.03E+00 | 4.64E-02 | 1.03E+00 |
| ENSG000002 | ACBD6 | 7.64E-01 | 9.17E-01 | 1.01E+00 | 1.44E-02 | 1.01E+00 |
| ENSG000001 | WDPCP | 7.64E-01 | 9.17E-01 | 1.02E+00 | 3.43E-02 | 1.02E+00 |
| ENSG000001 | ATL1 | 7.64E-01 | 9.17E-01 | 9.84E-01 | -2.31E-02 | -1.02E+00 |
| ENSG000001 | FTH1 | 7.64E-01 | 9.17E-01 | 1.01E+00 | 1.88E-02 | 1.01E+00 |
| ENSG000001 | METTL4 | 7.65E-01 | 9.17E-01 | 9.82E-01 | -2.57E-02 | -1.02E+00 |
| ENSG00000C | ASB1 | 7.65E-01 | 9.17E-01 | 1.01E+00 | 1.56E-02 | 1.01E+00 |
| ENSG000001 | ATP6AP2 | 7.65E-01 | 9.17E-01 | 1.01E+00 | 1.28E-02 | 1.01E+00 |
| ENSG000001 | DNMT3A | 7.65E-01 | 9.17E-01 | 1.02E+00 | 2.48E-02 | 1.02E+00 |
| ENSG000002 | UPF3AP1 | 7.65E-01 | ? | 1.11E+00 | 1.48E-01 | 1.11E+00 |
| ENSG000001 | YIPF6 | 7.65E-01 | 9.17E-01 | 1.01E+00 | 1.18E-02 | 1.01E+00 |
| ENSG000002 | CDKN2AIPNL | 7.65E-01 | 9.17E-01 | 9.81E-01 | -2.80E-02 | -1.02E+00 |
| ENSG000002 | RFX3-DT | 7.65E-01 | ? | 8.95E-01 | -1.60E-01 | -1.12E+00 |
| ENSG000002 | --- | 7.65E-01 | ? | 1.11E+00 | 1.57E-01 | 1.11E+00 |
| ENSG000002 | TRAPPC12-A: | 7.65E-01 | ? | 1.05E+00 | 7.00E-02 | 1.05E+00 |
| ENSG000001 | DACH2 | 7.65E-01 | ? | 9.23E-01 | -1.15E-01 | -1.08E+00 |
| ENSG000002 | --- | 7.65E-01 | ? | 8.83E-01 | -1.80E-01 | -1.13E+00 |
| ENSG000001 | OR7C1 | 7.66E-01 | ? | 1.06E+00 | 8.90E-02 | 1.06E+00 |
| ENSG000002 | --- | 7.66E-01 | ? | 1.18E+00 | 2.39E-01 | 1.18E+00 |
| ENSG000001 | RPS6KA5 | 7.66E-01 | 9.17E-01 | 1.03E+00 | 4.46E-02 | 1.03E+00 |
| ENSG000001 | JUNB | 7.66E-01 | 9.17E-01 | 9.67E-01 | -4.86E-02 | -1.03E+00 |
| ENSG000001 | DENND2B | 7.66E-01 | 9.17E-01 | 1.01E+00 | 1.69E-02 | 1.01E+00 |
| ENSG000002 | MAFTRR | 7.66E-01 | ? | 1.12E+00 | 1.68E-01 | 1.12E+00 |
| ENSG000001 | RCL1 | 7.66E-01 | 9.17E-01 | 1.01E+00 | 1.99E-02 | 1.01E+00 |
| ENSG000001 | ABHD17C | 7.66E-01 | 9.17E-01 | 1.02E+00 | 3.00E-02 | 1.02E+00 |
| ENSG000002 | --- | 7.66E-01 | ? | 1.06E+00 | 8.98E-02 | 1.06E+00 |
| ENSG000001 | ATAD5 | 7.66E-01 | 9.17E-01 | 1.03E+00 | 3.61E-02 | 1.03E+00 |
| ENSG000002 | --- | 7.66E-01 | ? | 1.08E+00 | 1.14E-01 | 1.08E+00 |
| ENSG000001 | TLDC2 | 7.66E-01 | ? | 8.97E-01 | -1.57E-01 | -1.12E+00 |
| ENSG000001 | METTL9 | 7.66E-01 | 9.18E-01 | 1.01E+00 | 1.07E-02 | 1.01E+00 |
| ENSG000001 | WWP2 | 7.66E-01 | 9.18E-01 | 9.91E-01 | -1.35E-02 | -1.01E+00 |
| ENSG000002 | --- | 7.66E-01 | ? | 8.90E-01 | -1.68E-01 | -1.12E+00 |
| ENSG000002 | --- | 7.66E-01 | ? | 8.98E-01 | -1.56E-01 | -1.11E+00 |

| | | | | | | |
|---|---|---|---|---|---|---|
| ENSG000001 | PPP1R26 | 7.66E-01 | 9.18E-01 | 1.01E+00 | 2.03E-02 | 1.01E+00 |
| ENSG000002 | MBLAC1 | 7.66E-01 | ? | 1.09E+00 | 1.20E-01 | 1.09E+00 |
| ENSG00000C | NFX1 | 7.66E-01 | 9.18E-01 | 1.01E+00 | 1.22E-02 | 1.01E+00 |
| ENSG000001 | PI4KAP2 | 7.67E-01 | 9.18E-01 | 9.66E-01 | -5.02E-02 | -1.04E+00 |
| ENSG000001 | NRM | 7.67E-01 | 9.18E-01 | 1.03E+00 | 3.93E-02 | 1.03E+00 |
| ENSG000001 | ATAD3A | 7.67E-01 | 9.18E-01 | 9.85E-01 | -2.22E-02 | -1.02E+00 |
| ENSG000001 | ATG16L2 | 7.67E-01 | 9.18E-01 | 1.03E+00 | 4.15E-02 | 1.03E+00 |
| ENSG000002 | LINC01747 | 7.67E-01 | ? | 9.19E-01 | -1.22E-01 | -1.09E+00 |
| ENSG000002 | GAS8-AS1 | 7.67E-01 | ? | 9.20E-01 | -1.21E-01 | -1.09E+00 |
| ENSG000001 | C1orf216 | 7.67E-01 | 9.18E-01 | 1.02E+00 | 2.41E-02 | 1.02E+00 |
| ENSG000001 | ZBED2 | 7.67E-01 | 9.18E-01 | 9.85E-01 | -2.21E-02 | -1.02E+00 |
| ENSG000001 | ADAP1 | 7.67E-01 | 9.18E-01 | 9.61E-01 | -5.74E-02 | -1.04E+00 |
| ENSG000002 | --- | 7.67E-01 | ? | 1.09E+00 | 1.22E-01 | 1.09E+00 |
| ENSG000001 | RTN1 | 7.67E-01 | ? | 1.10E+00 | 1.42E-01 | 1.10E+00 |
| ENSG000001 | P3H4 | 7.67E-01 | 9.18E-01 | 9.83E-01 | -2.52E-02 | -1.02E+00 |
| ENSG00000C | RNF14 | 7.67E-01 | 9.18E-01 | 9.93E-01 | -9.89E-03 | -1.01E+00 |
| ENSG00000C | CELF2 | 7.67E-01 | 9.18E-01 | 9.87E-01 | -1.84E-02 | -1.01E+00 |
| ENSG00000C | GCFC2 | 7.68E-01 | 9.18E-01 | 9.85E-01 | -2.21E-02 | -1.02E+00 |
| ENSG000001 | NDUFB1 | 7.68E-01 | 9.18E-01 | 1.01E+00 | 2.08E-02 | 1.01E+00 |
| ENSG00000C | EXOSC5 | 7.68E-01 | 9.18E-01 | 1.03E+00 | 4.26E-02 | 1.03E+00 |
| ENSG000001 | DCUN1D4 | 7.68E-01 | 9.18E-01 | 1.01E+00 | 1.30E-02 | 1.01E+00 |
| ENSG000002 | --- | 7.68E-01 | 9.18E-01 | 1.03E+00 | 3.75E-02 | 1.03E+00 |
| ENSG000002 | --- | 7.68E-01 | ? | 8.70E-01 | -2.01E-01 | -1.15E+00 |
| ENSG000001 | LIN7B | 7.68E-01 | ? | 1.08E+00 | 1.15E-01 | 1.08E+00 |
| ENSG000002 | --- | 7.68E-01 | ? | 9.00E-01 | -1.52E-01 | -1.11E+00 |
| ENSG000002 | --- | 7.68E-01 | ? | 1.10E+00 | 1.36E-01 | 1.10E+00 |
| ENSG00000C | RFC2 | 7.68E-01 | 9.19E-01 | 1.02E+00 | 2.91E-02 | 1.02E+00 |
| ENSG000001 | BYSL | 7.68E-01 | 9.19E-01 | 1.02E+00 | 2.42E-02 | 1.02E+00 |
| ENSG000001 | FBXW2 | 7.68E-01 | 9.19E-01 | 1.01E+00 | 8.91E-03 | 1.01E+00 |
| ENSG000001 | SHISA5 | 7.68E-01 | 9.19E-01 | 1.01E+00 | 1.39E-02 | 1.01E+00 |
| ENSG000002 | --- | 7.69E-01 | ? | 1.10E+00 | 1.33E-01 | 1.10E+00 |
| ENSG000001 | FAM204A | 7.69E-01 | 9.19E-01 | 9.89E-01 | -1.52E-02 | -1.01E+00 |
| ENSG000001 | ILRUN | 7.69E-01 | 9.19E-01 | 9.89E-01 | -1.65E-02 | -1.01E+00 |
| ENSG000002 | --- | 7.69E-01 | ? | 9.27E-01 | -1.10E-01 | -1.08E+00 |
| ENSG000001 | NEGR1 | 7.69E-01 | 9.19E-01 | 9.63E-01 | -5.49E-02 | -1.04E+00 |
| ENSG000001 | ABITRAM | 7.69E-01 | 9.19E-01 | 9.87E-01 | -1.93E-02 | -1.01E+00 |
| ENSG000002 | RPL7AP50 | 7.69E-01 | ? | 8.72E-01 | -1.98E-01 | -1.15E+00 |
| ENSG000001 | CHCHD5 | 7.69E-01 | 9.19E-01 | 1.03E+00 | 3.65E-02 | 1.03E+00 |
| ENSG000001 | CNTROB | 7.69E-01 | 9.19E-01 | 9.89E-01 | -1.61E-02 | -1.01E+00 |
| ENSG000001 | SYNJ1 | 7.69E-01 | 9.19E-01 | 9.85E-01 | -2.23E-02 | -1.02E+00 |
| ENSG000002 | HRAT17 | 7.69E-01 | ? | 1.14E+00 | 1.91E-01 | 1.14E+00 |
| ENSG000001 | N6AMT1 | 7.69E-01 | 9.19E-01 | 1.01E+00 | 1.85E-02 | 1.01E+00 |
| ENSG000002 | --- | 7.69E-01 | ? | 1.13E+00 | 1.81E-01 | 1.13E+00 |
| ENSG000001 | ARL9 | 7.70E-01 | 9.19E-01 | 1.03E+00 | 3.91E-02 | 1.03E+00 |
| ENSG000001 | PHF5A | 7.70E-01 | 9.19E-01 | 9.88E-01 | -1.72E-02 | -1.01E+00 |

| | | | | | | |
|---|---|---|---|---|---|---|
| ENSG000001 | TMEM63C | 7.70E-01 | 9.19E-01 | 1.02E+00 | 3.43E-02 | 1.02E+00 |
| ENSG00000C | PKP1 | 7.70E-01 | 9.19E-01 | 9.92E-01 | -1.11E-02 | -1.01E+00 |
| ENSG000002 | --- | 7.70E-01 | ? | 1.11E+00 | 1.52E-01 | 1.11E+00 |
| ENSG000002 | --- | 7.70E-01 | 9.19E-01 | 1.05E+00 | 6.41E-02 | 1.05E+00 |
| ENSG000001 | TDRKH | 7.70E-01 | 9.19E-01 | 1.02E+00 | 2.49E-02 | 1.02E+00 |
| ENSG000002 | --- | 7.70E-01 | ? | 9.34E-01 | -9.89E-02 | -1.07E+00 |
| ENSG000001 | GULP1 | 7.70E-01 | 9.19E-01 | 1.04E+00 | 6.07E-02 | 1.04E+00 |
| ENSG000001 | GFUS | 7.70E-01 | 9.19E-01 | 1.01E+00 | 1.79E-02 | 1.01E+00 |
| ENSG000002 | UNGP3 | 7.70E-01 | 9.19E-01 | 1.05E+00 | 6.48E-02 | 1.05E+00 |
| ENSG000002 | --- | 7.70E-01 | ? | 1.07E+00 | 9.28E-02 | 1.07E+00 |
| ENSG000002 | --- | 7.71E-01 | ? | 1.31E+00 | 3.84E-01 | 1.31E+00 |
| ENSG00000C | HSP90AB1 | 7.71E-01 | 9.20E-01 | 9.96E-01 | -5.93E-03 | -1.00E+00 |
| ENSG000002 | --- | 7.71E-01 | ? | 1.07E+00 | 9.70E-02 | 1.07E+00 |
| ENSG000001 | CIBAR1 | 7.71E-01 | 9.20E-01 | 9.81E-01 | -2.84E-02 | -1.02E+00 |
| ENSG000002 | F8A3 | 7.71E-01 | 9.20E-01 | 9.57E-01 | -6.28E-02 | -1.04E+00 |
| ENSG000002 | --- | 7.71E-01 | ? | 1.18E+00 | 2.45E-01 | 1.18E+00 |
| ENSG000001 | ARHGEF11 | 7.71E-01 | 9.20E-01 | 9.91E-01 | -1.32E-02 | -1.01E+00 |
| ENSG000001 | TMBIM4 | 7.71E-01 | 9.20E-01 | 9.89E-01 | -1.58E-02 | -1.01E+00 |
| ENSG000001 | ZNF689 | 7.71E-01 | 9.20E-01 | 1.02E+00 | 3.16E-02 | 1.02E+00 |
| ENSG000002 | CSGALNACT2 | 7.71E-01 | ? | 1.11E+00 | 1.55E-01 | 1.11E+00 |
| ENSG000001 | ECSIT | 7.72E-01 | 9.21E-01 | 1.02E+00 | 2.64E-02 | 1.02E+00 |
| ENSG000001 | RBM11 | 7.72E-01 | ? | 9.34E-01 | -9.86E-02 | -1.07E+00 |
| ENSG000001 | AGAP4 | 7.72E-01 | 9.21E-01 | 9.69E-01 | -4.47E-02 | -1.03E+00 |
| ENSG000001 | GNAI2 | 7.72E-01 | 9.21E-01 | 9.88E-01 | -1.78E-02 | -1.01E+00 |
| ENSG000002 | TPTEP1 | 7.72E-01 | 9.21E-01 | 9.83E-01 | -2.44E-02 | -1.02E+00 |
| ENSG000002 | DPH1-AS1 | 7.72E-01 | ? | 1.14E+00 | 1.88E-01 | 1.14E+00 |
| ENSG000001 | OIP5 | 7.72E-01 | ? | 1.07E+00 | 9.50E-02 | 1.07E+00 |
| ENSG000002 | HNRNPA1P59 | 7.72E-01 | ? | 1.09E+00 | 1.26E-01 | 1.09E+00 |
| ENSG000001 | ANXA1 | 7.72E-01 | 9.21E-01 | 1.01E+00 | 1.19E-02 | 1.01E+00 |
| ENSG000001 | FLNA | 7.72E-01 | 9.21E-01 | 1.01E+00 | 1.80E-02 | 1.01E+00 |
| ENSG000001 | --- | 7.72E-01 | ? | 8.75E-01 | -1.92E-01 | -1.14E+00 |
| ENSG000002 | --- | 7.72E-01 | ? | 1.15E+00 | 2.02E-01 | 1.15E+00 |
| ENSG000001 | ACVR2A | 7.72E-01 | 9.21E-01 | 1.01E+00 | 2.13E-02 | 1.01E+00 |
| ENSG000002 | MINDY2-DT | 7.72E-01 | ? | 1.09E+00 | 1.29E-01 | 1.09E+00 |
| ENSG000002 | --- | 7.72E-01 | ? | 8.54E-01 | -2.28E-01 | -1.17E+00 |
| ENSG000001 | FCMR | 7.72E-01 | 9.21E-01 | 9.69E-01 | -4.60E-02 | -1.03E+00 |
| ENSG000002 | TMEM9B-AS | 7.72E-01 | 9.21E-01 | 1.02E+00 | 3.26E-02 | 1.02E+00 |
| ENSG00000C | EIF2AK1 | 7.72E-01 | 9.21E-01 | 9.90E-01 | -1.39E-02 | -1.01E+00 |
| ENSG000001 | CPD | 7.72E-01 | 9.21E-01 | 1.01E+00 | 1.52E-02 | 1.01E+00 |
| ENSG000001 | CHMP4B | 7.72E-01 | 9.21E-01 | 1.01E+00 | 1.48E-02 | 1.01E+00 |
| ENSG000002 | --- | 7.73E-01 | 9.21E-01 | 9.59E-01 | -6.02E-02 | -1.04E+00 |
| ENSG000001 | TMEM214 | 7.73E-01 | 9.21E-01 | 1.01E+00 | 1.56E-02 | 1.01E+00 |
| ENSG000002 | PRELID1P4 | 7.73E-01 | ? | 8.32E-01 | -2.65E-01 | -1.20E+00 |
| ENSG000001 | RPRD1A | 7.73E-01 | 9.21E-01 | 1.01E+00 | 1.27E-02 | 1.01E+00 |
| ENSG000001 | DDX42 | 7.73E-01 | 9.21E-01 | 1.01E+00 | 8.83E-03 | 1.01E+00 |

| | | | | | | |
|---|---|---|---|---|---|---|
| ENSG000001 | ULBP1 | 7.73E-01 | 9.21E-01 | 1.04E+00 | 5.07E-02 | 1.04E+00 |
| ENSG000001 | BRAT1 | 7.73E-01 | 9.21E-01 | 1.02E+00 | 3.38E-02 | 1.02E+00 |
| ENSG000002 | KRT8P10 | 7.73E-01 | ? | 9.19E-01 | -1.22E-01 | -1.09E+00 |
| ENSG000001 | MYBBP1A | 7.73E-01 | 9.21E-01 | 9.89E-01 | -1.58E-02 | -1.01E+00 |
| ENSG000002 | --- | 7.73E-01 | ? | 1.23E+00 | 3.00E-01 | 1.23E+00 |
| ENSG000002 | LINC02861 | 7.73E-01 | ? | 1.08E+00 | 1.15E-01 | 1.08E+00 |
| ENSG000001 | TMEM50A | 7.73E-01 | 9.21E-01 | 9.93E-01 | -1.06E-02 | -1.01E+00 |
| ENSG00000C | FYB1 | 7.73E-01 | 9.21E-01 | 1.03E+00 | 4.88E-02 | 1.03E+00 |
| ENSG000001 | BIVM | 7.73E-01 | 9.21E-01 | 1.01E+00 | 1.68E-02 | 1.01E+00 |
| ENSG000002 | --- | 7.73E-01 | ? | 9.12E-01 | -1.32E-01 | -1.10E+00 |
| ENSG000001 | SERBP1 | 7.73E-01 | 9.21E-01 | 1.01E+00 | 1.14E-02 | 1.01E+00 |
| ENSG000001 | MAP4K1 | 7.73E-01 | ? | 1.14E+00 | 1.92E-01 | 1.14E+00 |
| ENSG000001 | MAP3K11 | 7.73E-01 | 9.21E-01 | 1.02E+00 | 2.69E-02 | 1.02E+00 |
| ENSG000002 | --- | 7.74E-01 | ? | 8.27E-01 | -2.74E-01 | -1.21E+00 |
| ENSG000002 | CCNYL2 | 7.74E-01 | ? | 1.13E+00 | 1.78E-01 | 1.13E+00 |
| ENSG000002 | --- | 7.74E-01 | ? | 1.07E+00 | 1.04E-01 | 1.07E+00 |
| ENSG000001 | CD70 | 7.74E-01 | ? | 1.14E+00 | 1.91E-01 | 1.14E+00 |
| ENSG000001 | ACD | 7.74E-01 | 9.21E-01 | 1.02E+00 | 2.46E-02 | 1.02E+00 |
| ENSG000001 | EIF3D | 7.74E-01 | 9.21E-01 | 9.93E-01 | -1.02E-02 | -1.01E+00 |
| ENSG000001 | BOLA2 | 7.74E-01 | 9.21E-01 | 9.40E-01 | -8.85E-02 | -1.06E+00 |
| ENSG000002 | BEND3P1 | 7.74E-01 | ? | 1.08E+00 | 1.11E-01 | 1.08E+00 |
| ENSG00000C | OGFOD1 | 7.74E-01 | 9.21E-01 | 1.01E+00 | 1.16E-02 | 1.01E+00 |
| ENSG000001 | COPS7B | 7.74E-01 | 9.21E-01 | 1.01E+00 | 1.32E-02 | 1.01E+00 |
| ENSG000002 | --- | 7.74E-01 | 9.21E-01 | 9.60E-01 | -5.84E-02 | -1.04E+00 |
| ENSG000001 | FASTKD1 | 7.74E-01 | 9.21E-01 | 9.89E-01 | -1.58E-02 | -1.01E+00 |
| ENSG000002 | GALNT7-DT | 7.74E-01 | 9.21E-01 | 9.57E-01 | -6.30E-02 | -1.04E+00 |
| ENSG00000C | SLC25A43 | 7.74E-01 | 9.21E-01 | 9.90E-01 | -1.45E-02 | -1.01E+00 |
| ENSG000001 | PRSS27 | 7.74E-01 | ? | 9.32E-01 | -1.02E-01 | -1.07E+00 |
| ENSG000002 | --- | 7.74E-01 | 9.21E-01 | 1.03E+00 | 3.71E-02 | 1.03E+00 |
| ENSG000002 | LNCAROD | 7.74E-01 | 9.21E-01 | 1.05E+00 | 6.94E-02 | 1.05E+00 |
| ENSG000001 | ANKMY2 | 7.74E-01 | 9.21E-01 | 1.02E+00 | 2.56E-02 | 1.02E+00 |
| ENSG000001 | DSTYK | 7.74E-01 | 9.21E-01 | 1.01E+00 | 1.44E-02 | 1.01E+00 |
| ENSG000001 | VAMP5 | 7.74E-01 | ? | 9.37E-01 | -9.32E-02 | -1.07E+00 |
| ENSG000001 | RPUSD3 | 7.74E-01 | 9.21E-01 | 9.87E-01 | -1.95E-02 | -1.01E+00 |
| ENSG00000C | BAMBI | 7.74E-01 | 9.21E-01 | 1.02E+00 | 2.99E-02 | 1.02E+00 |
| ENSG000001 | GAL3ST4 | 7.74E-01 | 9.21E-01 | 1.04E+00 | 5.14E-02 | 1.04E+00 |
| ENSG000002 | SNHG20 | 7.74E-01 | 9.21E-01 | 9.80E-01 | -2.98E-02 | -1.02E+00 |
| ENSG000001 | RNASE7 | 7.75E-01 | 9.21E-01 | 9.66E-01 | -4.97E-02 | -1.04E+00 |
| ENSG00000C | HSD3B7 | 7.75E-01 | 9.21E-01 | 1.03E+00 | 4.46E-02 | 1.03E+00 |
| ENSG000001 | TMEM30A | 7.75E-01 | 9.21E-01 | 9.92E-01 | -1.14E-02 | -1.01E+00 |
| ENSG000001 | USP15 | 7.75E-01 | 9.21E-01 | 9.89E-01 | -1.60E-02 | -1.01E+00 |
| ENSG000001 | LCLAT1 | 7.75E-01 | 9.21E-01 | 1.01E+00 | 1.34E-02 | 1.01E+00 |
| ENSG000002 | TCEA3 | 7.75E-01 | ? | 1.08E+00 | 1.15E-01 | 1.08E+00 |
| ENSG000001 | TMEM160 | 7.75E-01 | ? | 1.06E+00 | 9.07E-02 | 1.06E+00 |
| ENSG000001 | SENP8 | 7.75E-01 | 9.21E-01 | 9.81E-01 | -2.79E-02 | -1.02E+00 |

| | | | | | | |
|---|---|---|---|---|---|---|
| ENSG000001 | LGMN | 7.75E-01 | 9.21E-01 | 1.01E+00 | 1.31E-02 | 1.01E+00 |
| ENSG000001 | CC2D2B | 7.75E-01 | ? | 1.15E+00 | 1.96E-01 | 1.15E+00 |
| ENSG000002 | --- | 7.75E-01 | 9.21E-01 | 1.04E+00 | 6.03E-02 | 1.04E+00 |
| ENSG000002 | --- | 7.76E-01 | ? | 1.15E+00 | 1.99E-01 | 1.15E+00 |
| ENSG000001 | MRPL42 | 7.76E-01 | 9.21E-01 | 9.92E-01 | -1.23E-02 | -1.01E+00 |
| ENSG000001 | SLC52A3 | 7.76E-01 | 9.21E-01 | 9.69E-01 | -4.61E-02 | -1.03E+00 |
| ENSG000001 | PIP4P2 | 7.76E-01 | 9.21E-01 | 9.86E-01 | -2.09E-02 | -1.01E+00 |
| ENSG000002 | --- | 7.76E-01 | ? | 1.10E+00 | 1.33E-01 | 1.10E+00 |
| ENSG000002 | --- | 7.76E-01 | 9.21E-01 | 9.27E-01 | -1.09E-01 | -1.08E+00 |
| ENSG000002 | --- | 7.76E-01 | ? | 1.07E+00 | 9.62E-02 | 1.07E+00 |
| ENSG000002 | FAM88C | 7.76E-01 | 9.21E-01 | 1.05E+00 | 6.56E-02 | 1.05E+00 |
| ENSG000002 | --- | 7.76E-01 | 9.21E-01 | 9.45E-01 | -8.20E-02 | -1.06E+00 |
| ENSG00000C | BRCA1 | 7.76E-01 | 9.21E-01 | 9.82E-01 | -2.67E-02 | -1.02E+00 |
| ENSG000001 | CPPED1 | 7.76E-01 | 9.21E-01 | 9.88E-01 | -1.70E-02 | -1.01E+00 |
| ENSG000001 | HSD17B11 | 7.76E-01 | 9.21E-01 | 9.82E-01 | -2.60E-02 | -1.02E+00 |
| ENSG000001 | ACAD8 | 7.76E-01 | 9.21E-01 | 9.86E-01 | -2.00E-02 | -1.01E+00 |
| ENSG000001 | ATP6V0D2 | 7.76E-01 | ? | 1.13E+00 | 1.78E-01 | 1.13E+00 |
| ENSG000002 | SPRR2F | 7.76E-01 | ? | 8.98E-01 | -1.56E-01 | -1.11E+00 |
| ENSG000002 | --- | 7.76E-01 | 9.21E-01 | 8.70E-01 | -2.01E-01 | -1.15E+00 |
| ENSG000001 | FAM102A | 7.76E-01 | 9.21E-01 | 1.02E+00 | 2.80E-02 | 1.02E+00 |
| ENSG000001 | DHRSX | 7.76E-01 | 9.21E-01 | 1.02E+00 | 2.17E-02 | 1.02E+00 |
| ENSG000001 | MYO5B | 7.77E-01 | 9.21E-01 | 1.02E+00 | 2.95E-02 | 1.02E+00 |
| ENSG000001 | C1GALT1 | 7.77E-01 | 9.21E-01 | 1.01E+00 | 1.74E-02 | 1.01E+00 |
| ENSG000001 | ZCCHC18 | 7.77E-01 | ? | 8.46E-01 | -2.42E-01 | -1.18E+00 |
| ENSG000001 | HECTD2 | 7.77E-01 | 9.21E-01 | 1.02E+00 | 2.69E-02 | 1.02E+00 |
| ENSG000001 | SUMO1 | 7.77E-01 | 9.21E-01 | 1.01E+00 | 1.24E-02 | 1.01E+00 |
| ENSG00000C | HNRNPC | 7.77E-01 | 9.21E-01 | 1.01E+00 | 1.15E-02 | 1.01E+00 |
| ENSG000002 | --- | 7.77E-01 | ? | 9.38E-01 | -9.27E-02 | -1.07E+00 |
| ENSG000002 | CDH3-AS1 | 7.77E-01 | 9.21E-01 | 9.54E-01 | -6.76E-02 | -1.05E+00 |
| ENSG000001 | ABCG1 | 7.77E-01 | 9.21E-01 | 1.02E+00 | 2.73E-02 | 1.02E+00 |
| ENSG000001 | HARBI1 | 7.77E-01 | 9.21E-01 | 1.02E+00 | 2.98E-02 | 1.02E+00 |
| ENSG000001 | CFAP92 | 7.77E-01 | ? | 1.06E+00 | 8.23E-02 | 1.06E+00 |
| ENSG000001 | SLA2 | 7.77E-01 | ? | 8.80E-01 | -1.84E-01 | -1.14E+00 |
| ENSG000001 | EIF1AD | 7.77E-01 | 9.21E-01 | 9.89E-01 | -1.61E-02 | -1.01E+00 |
| ENSG000001 | SRSF3 | 7.77E-01 | 9.21E-01 | 9.93E-01 | -1.05E-02 | -1.01E+00 |
| ENSG000002 | --- | 7.77E-01 | ? | 7.94E-01 | -3.32E-01 | -1.26E+00 |
| ENSG000002 | --- | 7.77E-01 | ? | 9.08E-01 | -1.39E-01 | -1.10E+00 |
| ENSG00000C | PPIE | 7.77E-01 | 9.21E-01 | 9.88E-01 | -1.71E-02 | -1.01E+00 |
| ENSG000002 | --- | 7.77E-01 | ? | 9.10E-01 | -1.36E-01 | -1.10E+00 |
| ENSG000002 | ZNF407-AS1 | 7.77E-01 | 9.21E-01 | 1.02E+00 | 2.50E-02 | 1.02E+00 |
| ENSG00000C | USP48 | 7.77E-01 | 9.21E-01 | 9.92E-01 | -1.12E-02 | -1.01E+00 |
| ENSG000002 | RPL3P2 | 7.77E-01 | ? | 1.11E+00 | 1.52E-01 | 1.11E+00 |
| ENSG000002 | CRYBB2 | 7.77E-01 | ? | 8.77E-01 | -1.90E-01 | -1.14E+00 |
| ENSG000001 | ELFN2 | 7.77E-01 | 9.22E-01 | 9.82E-01 | -2.69E-02 | -1.02E+00 |
| ENSG000001 | UBALD2 | 7.77E-01 | 9.22E-01 | 9.83E-01 | -2.47E-02 | -1.02E+00 |

| | | | | | | |
|---|---|---|---|---|---|---|
| ENSG000001 | MRPL2 | 7.77E-01 | 9.22E-01 | 1.02E+00 | 2.22E-02 | 1.02E+00 |
| ENSG000002 | MT-RNR2 | 7.78E-01 | 9.22E-01 | 1.02E+00 | 2.84E-02 | 1.02E+00 |
| ENSG000002 | --- | 7.78E-01 | 9.22E-01 | 1.03E+00 | 3.66E-02 | 1.03E+00 |
| ENSG000002 | --- | 7.78E-01 | ? | 1.10E+00 | 1.43E-01 | 1.10E+00 |
| ENSG000002 | C2orf49-DT | 7.78E-01 | 9.22E-01 | 9.65E-01 | -5.12E-02 | -1.04E+00 |
| ENSG000001 | LMAN2L | 7.78E-01 | 9.22E-01 | 9.85E-01 | -2.14E-02 | -1.01E+00 |
| ENSG000002 | --- | 7.78E-01 | ? | 1.13E+00 | 1.75E-01 | 1.13E+00 |
| ENSG000002 | HNRNPA3P1( | 7.78E-01 | ? | 1.14E+00 | 1.88E-01 | 1.14E+00 |
| ENSG000001 | PRUNE1 | 7.78E-01 | 9.22E-01 | 1.02E+00 | 2.29E-02 | 1.02E+00 |
| ENSG000002 | ANG | 7.78E-01 | 9.22E-01 | 9.65E-01 | -5.14E-02 | -1.04E+00 |
| ENSG000001 | SSX2IP | 7.78E-01 | 9.22E-01 | 1.02E+00 | 2.32E-02 | 1.02E+00 |
| ENSG000002 | --- | 7.78E-01 | ? | 1.10E+00 | 1.33E-01 | 1.10E+00 |
| ENSG000001 | MAP4K2 | 7.78E-01 | 9.22E-01 | 1.02E+00 | 2.42E-02 | 1.02E+00 |
| ENSG000002 | --- | 7.79E-01 | ? | 9.18E-01 | -1.24E-01 | -1.09E+00 |
| ENSG00000C | FGR | 7.79E-01 | ? | 1.10E+00 | 1.38E-01 | 1.10E+00 |
| ENSG00000C | FAM136A | 7.79E-01 | 9.22E-01 | 1.01E+00 | 1.49E-02 | 1.01E+00 |
| ENSG000002 | --- | 7.79E-01 | 9.22E-01 | 1.04E+00 | 5.52E-02 | 1.04E+00 |
| ENSG000002 | --- | 7.79E-01 | 9.22E-01 | 9.54E-01 | -6.83E-02 | -1.05E+00 |
| ENSG000001 | PLA2G6 | 7.79E-01 | 9.23E-01 | 1.03E+00 | 3.77E-02 | 1.03E+00 |
| ENSG000002 | --- | 7.79E-01 | 9.23E-01 | 1.04E+00 | 6.18E-02 | 1.04E+00 |
| ENSG000002 | KRT6A | 7.79E-01 | 9.23E-01 | 9.90E-01 | -1.47E-02 | -1.01E+00 |
| ENSG000002 | ABHD14A | 7.79E-01 | 9.23E-01 | 1.03E+00 | 4.73E-02 | 1.03E+00 |
| ENSG000001 | TECTA | 7.79E-01 | ? | 9.24E-01 | -1.15E-01 | -1.08E+00 |
| ENSG000002 | GALT | 7.79E-01 | 9.23E-01 | 1.02E+00 | 2.66E-02 | 1.02E+00 |
| ENSG000001 | CHAF1A | 7.79E-01 | 9.23E-01 | 9.86E-01 | -1.97E-02 | -1.01E+00 |
| ENSG000001 | ZNF285BP | 7.79E-01 | ? | 1.09E+00 | 1.25E-01 | 1.09E+00 |
| ENSG000001 | NR4A2 | 7.80E-01 | 9.23E-01 | 9.56E-01 | -6.44E-02 | -1.05E+00 |
| ENSG000002 | --- | 7.80E-01 | ? | 1.10E+00 | 1.34E-01 | 1.10E+00 |
| ENSG00000C | MAD1L1 | 7.80E-01 | 9.23E-01 | 1.01E+00 | 2.06E-02 | 1.01E+00 |
| ENSG000002 | SOCS5P4 | 7.80E-01 | ? | 9.05E-01 | -1.44E-01 | -1.10E+00 |
| ENSG000001 | ZCRB1 | 7.80E-01 | 9.23E-01 | 9.87E-01 | -1.93E-02 | -1.01E+00 |
| ENSG00000C | CA11 | 7.80E-01 | 9.23E-01 | 1.04E+00 | 6.31E-02 | 1.04E+00 |
| ENSG000002 | LINC00520 | 7.80E-01 | 9.23E-01 | 9.57E-01 | -6.39E-02 | -1.05E+00 |
| ENSG000002 | SNHG4 | 7.80E-01 | 9.23E-01 | 1.03E+00 | 3.97E-02 | 1.03E+00 |
| ENSG000002 | PAXIP1-DT | 7.80E-01 | 9.23E-01 | 9.59E-01 | -6.07E-02 | -1.04E+00 |
| ENSG000002 | ZNF616 | 7.80E-01 | 9.23E-01 | 9.87E-01 | -1.82E-02 | -1.01E+00 |
| ENSG000001 | USP35 | 7.80E-01 | 9.23E-01 | 1.02E+00 | 2.47E-02 | 1.02E+00 |
| ENSG000002 | --- | 7.80E-01 | ? | 8.93E-01 | -1.63E-01 | -1.12E+00 |
| ENSG000001 | ZNF284 | 7.80E-01 | 9.23E-01 | 9.79E-01 | -3.08E-02 | -1.02E+00 |
| ENSG000002 | --- | 7.80E-01 | 9.23E-01 | 1.05E+00 | 6.94E-02 | 1.05E+00 |
| ENSG000002 | --- | 7.80E-01 | ? | 1.14E+00 | 1.93E-01 | 1.14E+00 |
| ENSG000002 | CYP1B1-AS1 | 7.80E-01 | ? | 1.13E+00 | 1.73E-01 | 1.13E+00 |
| ENSG000002 | ZNF469 | 7.80E-01 | 9.23E-01 | 1.04E+00 | 6.11E-02 | 1.04E+00 |
| ENSG000001 | CASKIN1 | 7.80E-01 | 9.23E-01 | 9.47E-01 | -7.86E-02 | -1.06E+00 |
| ENSG000001 | GBP3 | 7.80E-01 | 9.23E-01 | 9.89E-01 | -1.59E-02 | -1.01E+00 |

| | | | | | | |
|---|---|---|---|---|---|---|
| ENSG000001 | TES | 7.80E-01 | 9.23E-01 | 1.01E+00 | 1.18E-02 | 1.01E+00 |
| ENSG000001 | BOLA2B | 7.80E-01 | 9.23E-01 | 9.68E-01 | -4.63E-02 | -1.03E+00 |
| ENSG000001 | GLI1 | 7.80E-01 | 9.23E-01 | 1.04E+00 | 6.17E-02 | 1.04E+00 |
| ENSG000002 | --- | 7.80E-01 | ? | 9.38E-01 | -9.21E-02 | -1.07E+00 |
| ENSG000001 | CCNB1 | 7.80E-01 | 9.23E-01 | 9.79E-01 | -3.04E-02 | -1.02E+00 |
| ENSG000002 | LINC03011 | 7.81E-01 | ? | 9.40E-01 | -8.99E-02 | -1.06E+00 |
| ENSG000001 | RGS20 | 7.81E-01 | 9.23E-01 | 9.87E-01 | -1.87E-02 | -1.01E+00 |
| ENSG000001 | PROC | 7.81E-01 | ? | 1.20E+00 | 2.57E-01 | 1.20E+00 |
| ENSG000001 | DSN1 | 7.81E-01 | 9.23E-01 | 9.82E-01 | -2.62E-02 | -1.02E+00 |
| ENSG000002 | USP3-AS1 | 7.81E-01 | ? | 9.49E-01 | -7.52E-02 | -1.05E+00 |
| ENSG000001 | ANKRD23 | 7.81E-01 | ? | 1.06E+00 | 7.83E-02 | 1.06E+00 |
| ENSG00000C | DLEC1 | 7.81E-01 | ? | 9.39E-01 | -9.07E-02 | -1.06E+00 |
| ENSG000001 | ERCC2 | 7.81E-01 | 9.23E-01 | 1.01E+00 | 1.98E-02 | 1.01E+00 |
| ENSG000002 | LINC02987 | 7.81E-01 | 9.23E-01 | 9.88E-01 | -1.79E-02 | -1.01E+00 |
| ENSG00000C | MEF2C | 7.81E-01 | 9.23E-01 | 9.77E-01 | -3.36E-02 | -1.02E+00 |
| ENSG000002 | --- | 7.81E-01 | ? | 8.75E-01 | -1.93E-01 | -1.14E+00 |
| ENSG000001 | GNG11 | 7.81E-01 | 9.23E-01 | 1.04E+00 | 5.90E-02 | 1.04E+00 |
| ENSG000002 | H2BC9 | 7.81E-01 | ? | 9.24E-01 | -1.13E-01 | -1.08E+00 |
| ENSG000002 | --- | 7.81E-01 | ? | 8.45E-01 | -2.43E-01 | -1.18E+00 |
| ENSG000001 | MTMR14 | 7.81E-01 | 9.23E-01 | 9.89E-01 | -1.58E-02 | -1.01E+00 |
| ENSG000001 | ADRB2 | 7.82E-01 | 9.23E-01 | 9.89E-01 | -1.65E-02 | -1.01E+00 |
| ENSG000002 | --- | 7.82E-01 | ? | 1.09E+00 | 1.18E-01 | 1.09E+00 |
| ENSG000001 | WDR73 | 7.82E-01 | 9.23E-01 | 1.01E+00 | 2.03E-02 | 1.01E+00 |
| ENSG000001 | ZFYVE28 | 7.82E-01 | ? | 1.06E+00 | 7.77E-02 | 1.06E+00 |
| ENSG000001 | ADAMTSL5 | 7.82E-01 | 9.23E-01 | 1.02E+00 | 2.54E-02 | 1.02E+00 |
| ENSG000001 | ZNF570 | 7.82E-01 | 9.23E-01 | 9.86E-01 | -2.10E-02 | -1.01E+00 |
| ENSG000001 | KLK5 | 7.82E-01 | 9.23E-01 | 9.43E-01 | -8.51E-02 | -1.06E+00 |
| ENSG000002 | --- | 7.82E-01 | ? | 1.12E+00 | 1.62E-01 | 1.12E+00 |
| ENSG000001 | CDKN1C | 7.82E-01 | 9.23E-01 | 9.50E-01 | -7.45E-02 | -1.05E+00 |
| ENSG000002 | YTHDF3-DT | 7.82E-01 | ? | 1.14E+00 | 1.85E-01 | 1.14E+00 |
| ENSG000001 | REN | 7.82E-01 | ? | 1.11E+00 | 1.46E-01 | 1.11E+00 |
| ENSG000001 | AADAT | 7.82E-01 | 9.24E-01 | 1.02E+00 | 3.06E-02 | 1.02E+00 |
| ENSG000002 | EIF4A1P2 | 7.82E-01 | ? | 9.36E-01 | -9.47E-02 | -1.07E+00 |
| ENSG000001 | FAM186B | 7.82E-01 | ? | 9.34E-01 | -9.82E-02 | -1.07E+00 |
| ENSG000002 | --- | 7.82E-01 | 9.24E-01 | 9.57E-01 | -6.28E-02 | -1.04E+00 |
| ENSG000001 | PJA2 | 7.83E-01 | 9.24E-01 | 1.01E+00 | 1.27E-02 | 1.01E+00 |
| ENSG000001 | FAM172A | 7.83E-01 | 9.24E-01 | 9.90E-01 | -1.47E-02 | -1.01E+00 |
| ENSG000001 | TRADD | 7.83E-01 | 9.24E-01 | 1.02E+00 | 3.51E-02 | 1.02E+00 |
| ENSG000001 | SERAC1 | 7.83E-01 | 9.24E-01 | 9.86E-01 | -2.00E-02 | -1.01E+00 |
| ENSG000001 | RSPH10B | 7.83E-01 | ? | 1.05E+00 | 7.41E-02 | 1.05E+00 |
| ENSG000002 | --- | 7.83E-01 | ? | 8.85E-01 | -1.76E-01 | -1.13E+00 |
| ENSG000002 | FASTKD5 | 7.83E-01 | 9.24E-01 | 9.89E-01 | -1.59E-02 | -1.01E+00 |
| ENSG000001 | LRMDA | 7.83E-01 | 9.24E-01 | 9.63E-01 | -5.49E-02 | -1.04E+00 |
| ENSG000001 | LONP2 | 7.83E-01 | 9.24E-01 | 9.93E-01 | -1.08E-02 | -1.01E+00 |
| ENSG000002 | --- | 7.83E-01 | ? | 9.32E-01 | -1.02E-01 | -1.07E+00 |

| | | | | | | |
|---|---|---|---|---|---|---|
| ENSG000001 | SPRR1A | 7.83E-01 | 9.24E-01 | 1.10E+00 | 1.43E-01 | 1.10E+00 |
| ENSG000001 | PRKCI | 7.83E-01 | 9.24E-01 | 1.01E+00 | 1.32E-02 | 1.01E+00 |
| ENSG000002 | C1GALT1C1L | 7.84E-01 | ? | 1.14E+00 | 1.90E-01 | 1.14E+00 |
| ENSG000002 | --- | 7.84E-01 | ? | 9.10E-01 | -1.35E-01 | -1.10E+00 |
| ENSG000002 | --- | 7.84E-01 | 9.24E-01 | 1.02E+00 | 3.04E-02 | 1.02E+00 |
| ENSG000001 | MRPL47 | 7.84E-01 | 9.25E-01 | 9.90E-01 | -1.43E-02 | -1.01E+00 |
| ENSG000001 | PRMT2 | 7.84E-01 | 9.25E-01 | 9.92E-01 | -1.09E-02 | -1.01E+00 |
| ENSG000002 | --- | 7.84E-01 | ? | 1.10E+00 | 1.43E-01 | 1.10E+00 |
| ENSG000001 | MON1B | 7.84E-01 | 9.25E-01 | 1.01E+00 | 1.22E-02 | 1.01E+00 |
| ENSG000002 | --- | 7.84E-01 | ? | 1.09E+00 | 1.19E-01 | 1.09E+00 |
| ENSG00000C | ZNF302 | 7.84E-01 | 9.25E-01 | 1.01E+00 | 2.11E-02 | 1.01E+00 |
| ENSG00000C | NRXN3 | 7.84E-01 | ? | 9.52E-01 | -7.04E-02 | -1.05E+00 |
| ENSG000002 | --- | 7.84E-01 | ? | 8.94E-01 | -1.62E-01 | -1.12E+00 |
| ENSG000002 | --- | 7.85E-01 | ? | 1.15E+00 | 2.04E-01 | 1.15E+00 |
| ENSG000002 | PKD1P6 | 7.85E-01 | 9.25E-01 | 9.56E-01 | -6.51E-02 | -1.05E+00 |
| ENSG00000C | N4BP2 | 7.85E-01 | 9.25E-01 | 1.01E+00 | 2.02E-02 | 1.01E+00 |
| ENSG000002 | ZNF783 | 7.85E-01 | 9.25E-01 | 9.81E-01 | -2.81E-02 | -1.02E+00 |
| ENSG000001 | FAM183A | 7.85E-01 | ? | 1.08E+00 | 1.09E-01 | 1.08E+00 |
| ENSG000001 | EIF5A2 | 7.85E-01 | 9.25E-01 | 9.85E-01 | -2.17E-02 | -1.02E+00 |
| ENSG000001 | TMEM277P | 7.85E-01 | ? | 1.12E+00 | 1.60E-01 | 1.12E+00 |
| ENSG00000C | TCOF1 | 7.85E-01 | 9.25E-01 | 1.01E+00 | 1.72E-02 | 1.01E+00 |
| ENSG000001 | SLC5A6 | 7.85E-01 | 9.25E-01 | 9.83E-01 | -2.52E-02 | -1.02E+00 |
| ENSG000002 | --- | 7.85E-01 | ? | 1.18E+00 | 2.38E-01 | 1.18E+00 |
| ENSG000001 | SMARCA5 | 7.85E-01 | 9.25E-01 | 9.90E-01 | -1.42E-02 | -1.01E+00 |
| ENSG00000C | BMAL2 | 7.85E-01 | 9.25E-01 | 9.84E-01 | -2.29E-02 | -1.02E+00 |
| ENSG000002 | --- | 7.85E-01 | ? | 9.34E-01 | -9.83E-02 | -1.07E+00 |
| ENSG000001 | FHOD1 | 7.85E-01 | 9.25E-01 | 1.02E+00 | 2.35E-02 | 1.02E+00 |
| ENSG000001 | FOXF2 | 7.85E-01 | ? | 8.88E-01 | -1.71E-01 | -1.13E+00 |
| ENSG000001 | TMEM184B | 7.85E-01 | 9.25E-01 | 1.01E+00 | 1.80E-02 | 1.01E+00 |
| ENSG000001 | HPSE | 7.85E-01 | 9.25E-01 | 1.03E+00 | 4.69E-02 | 1.03E+00 |
| ENSG000001 | SHF | 7.86E-01 | ? | 8.91E-01 | -1.66E-01 | -1.12E+00 |
| ENSG000001 | SAMD14 | 7.86E-01 | ? | 9.38E-01 | -9.22E-02 | -1.07E+00 |
| ENSG000001 | DEFB1 | 7.86E-01 | 9.25E-01 | 9.46E-01 | -7.97E-02 | -1.06E+00 |
| ENSG000001 | CAMTA2 | 7.86E-01 | 9.25E-01 | 9.89E-01 | -1.55E-02 | -1.01E+00 |
| ENSG000001 | IKZF4 | 7.86E-01 | 9.25E-01 | 1.02E+00 | 2.49E-02 | 1.02E+00 |
| ENSG000002 | EXOC3-AS1 | 7.86E-01 | 9.25E-01 | 1.03E+00 | 4.60E-02 | 1.03E+00 |
| ENSG00000C | RNASET2 | 7.86E-01 | 9.25E-01 | 1.01E+00 | 1.99E-02 | 1.01E+00 |
| ENSG00000C | CYLD | 7.86E-01 | 9.25E-01 | 1.01E+00 | 2.00E-02 | 1.01E+00 |
| ENSG000002 | --- | 7.86E-01 | ? | 1.09E+00 | 1.24E-01 | 1.09E+00 |
| ENSG000001 | PMS2CL | 7.86E-01 | 9.25E-01 | 8.69E-01 | -2.03E-01 | -1.15E+00 |
| ENSG000001 | ASB6 | 7.86E-01 | 9.25E-01 | 1.01E+00 | 1.67E-02 | 1.01E+00 |
| ENSG000002 | NTAN1P2 | 7.86E-01 | ? | 9.13E-01 | -1.31E-01 | -1.10E+00 |
| ENSG000001 | EFCAB11 | 7.86E-01 | 9.25E-01 | 1.02E+00 | 2.41E-02 | 1.02E+00 |
| ENSG000002 | --- | 7.86E-01 | ? | 9.14E-01 | -1.29E-01 | -1.09E+00 |
| ENSG000001 | MMRN2 | 7.86E-01 | 9.25E-01 | 9.65E-01 | -5.13E-02 | -1.04E+00 |

| | | | | | | |
|---|---|---|---|---|---|---|
| ENSG000001 | HYAL3 | 7.86E-01 | 9.25E-01 | 9.80E-01 | -2.92E-02 | -1.02E+00 |
| ENSG000001 | CCDC144A | 7.86E-01 | 9.25E-01 | 1.04E+00 | 6.30E-02 | 1.04E+00 |
| ENSG000001 | CILP2 | 7.86E-01 | ? | 9.42E-01 | -8.55E-02 | -1.06E+00 |
| ENSG000002 | --- | 7.86E-01 | ? | 9.06E-01 | -1.42E-01 | -1.10E+00 |
| ENSG000001 | BLOC1S3 | 7.86E-01 | 9.25E-01 | 9.85E-01 | -2.20E-02 | -1.02E+00 |
| ENSG000001 | LTO1 | 7.86E-01 | 9.25E-01 | 1.01E+00 | 1.85E-02 | 1.01E+00 |
| ENSG000001 | TM2D2 | 7.87E-01 | 9.25E-01 | 1.01E+00 | 1.52E-02 | 1.01E+00 |
| ENSG000001 | ZNF280D | 7.87E-01 | 9.25E-01 | 9.86E-01 | -1.99E-02 | -1.01E+00 |
| ENSG000002 | --- | 7.87E-01 | 9.26E-01 | 9.65E-01 | -5.14E-02 | -1.04E+00 |
| ENSG000002 | GCATP1 | 7.87E-01 | ? | 9.04E-01 | -1.45E-01 | -1.11E+00 |
| ENSG000002 | --- | 7.87E-01 | ? | 1.08E+00 | 1.09E-01 | 1.08E+00 |
| ENSG000001 | CLGN | 7.87E-01 | ? | 9.40E-01 | -8.97E-02 | -1.06E+00 |
| ENSG000002 | LINC00871 | 7.87E-01 | ? | 1.11E+00 | 1.53E-01 | 1.11E+00 |
| ENSG000001 | PCSK6 | 7.87E-01 | 9.26E-01 | 1.03E+00 | 4.05E-02 | 1.03E+00 |
| ENSG000001 | --- | 7.87E-01 | 9.26E-01 | 9.19E-01 | -1.22E-01 | -1.09E+00 |
| ENSG000001 | GSR | 7.87E-01 | 9.26E-01 | 1.01E+00 | 1.88E-02 | 1.01E+00 |
| ENSG000002 | UPF3AP2 | 7.87E-01 | ? | 9.08E-01 | -1.40E-01 | -1.10E+00 |
| ENSG000002 | CDC42P6 | 7.87E-01 | ? | 9.52E-01 | -7.08E-02 | -1.05E+00 |
| ENSG000002 | HSPD1P4 | 7.87E-01 | ? | 9.10E-01 | -1.36E-01 | -1.10E+00 |
| ENSG000001 | SEPTIN7 | 7.87E-01 | 9.26E-01 | 9.91E-01 | -1.31E-02 | -1.01E+00 |
| ENSG000001 | BCL2 | 7.87E-01 | 9.26E-01 | 1.04E+00 | 5.41E-02 | 1.04E+00 |
| ENSG00000C | ZNF510 | 7.87E-01 | 9.26E-01 | 9.87E-01 | -1.87E-02 | -1.01E+00 |
| ENSG000001 | GLIS2 | 7.88E-01 | 9.26E-01 | 1.02E+00 | 2.92E-02 | 1.02E+00 |
| ENSG00000C | SLC6A13 | 7.88E-01 | ? | 9.29E-01 | -1.06E-01 | -1.08E+00 |
| ENSG000001 | NRBP2 | 7.88E-01 | 9.26E-01 | 1.01E+00 | 1.61E-02 | 1.01E+00 |
| ENSG000001 | CENPK | 7.88E-01 | 9.26E-01 | 1.03E+00 | 4.52E-02 | 1.03E+00 |
| ENSG000001 | ZNF404 | 7.88E-01 | 9.26E-01 | 9.78E-01 | -3.15E-02 | -1.02E+00 |
| ENSG000002 | --- | 7.88E-01 | ? | 1.10E+00 | 1.32E-01 | 1.10E+00 |
| ENSG000001 | ATG4A | 7.88E-01 | 9.26E-01 | 9.90E-01 | -1.42E-02 | -1.01E+00 |
| ENSG000001 | RUNX2 | 7.88E-01 | 9.26E-01 | 9.85E-01 | -2.20E-02 | -1.02E+00 |
| ENSG00000C | POLQ | 7.88E-01 | 9.26E-01 | 9.65E-01 | -5.09E-02 | -1.04E+00 |
| ENSG000002 | --- | 7.88E-01 | ? | 8.89E-01 | -1.69E-01 | -1.12E+00 |
| ENSG000002 | TDRKH-AS1 | 7.88E-01 | ? | 8.96E-01 | -1.58E-01 | -1.12E+00 |
| ENSG000001 | FOXL1 | 7.88E-01 | 9.26E-01 | 1.02E+00 | 2.65E-02 | 1.02E+00 |
| ENSG000002 | LINC01934 | 7.88E-01 | ? | 8.89E-01 | -1.69E-01 | -1.12E+00 |
| ENSG000001 | HMGCS1 | 7.88E-01 | 9.26E-01 | 1.01E+00 | 1.65E-02 | 1.01E+00 |
| ENSG000002 | --- | 7.88E-01 | ? | 8.97E-01 | -1.58E-01 | -1.12E+00 |
| ENSG000001 | PDLIM4 | 7.88E-01 | 9.26E-01 | 1.03E+00 | 3.80E-02 | 1.03E+00 |
| ENSG000002 | HMGA1P2 | 7.88E-01 | ? | 8.95E-01 | -1.60E-01 | -1.12E+00 |
| ENSG000001 | FAM102B | 7.88E-01 | 9.26E-01 | 1.02E+00 | 2.55E-02 | 1.02E+00 |
| ENSG000001 | UBN1 | 7.89E-01 | 9.26E-01 | 1.01E+00 | 1.08E-02 | 1.01E+00 |
| ENSG000001 | MAL | 7.89E-01 | ? | 1.15E+00 | 1.96E-01 | 1.15E+00 |
| ENSG000002 | --- | 7.89E-01 | 9.26E-01 | 9.63E-01 | -5.43E-02 | -1.04E+00 |
| ENSG000001 | CRISPLD1 | 7.89E-01 | 9.26E-01 | 1.04E+00 | 5.56E-02 | 1.04E+00 |
| ENSG000001 | TRIM44 | 7.89E-01 | 9.26E-01 | 9.93E-01 | -9.85E-03 | -1.01E+00 |

| | | | | | | | |
|---|---|---|---|---|---|---|---|
| ENSG000001 | RBKS | 7.89E-01 | | 9.26E-01 | 1.04E+00 | 5.03E-02 | 1.04E+00 |
| ENSG000001 | EXOSC1 | 7.89E-01 | | 9.26E-01 | 1.01E+00 | 1.70E-02 | 1.01E+00 |
| ENSG000001 | ENTPD5 | 7.89E-01 | | 9.26E-01 | 9.90E-01 | -1.42E-02 | -1.01E+00 |
| ENSG00000C | ZFX | 7.89E-01 | | 9.26E-01 | 1.01E+00 | 1.14E-02 | 1.01E+00 |
| ENSG000001 | MORC4 | 7.89E-01 | | 9.26E-01 | 9.91E-01 | -1.36E-02 | -1.01E+00 |
| ENSG000002 | HIF1A-AS1 | 7.89E-01 | ? | | 9.19E-01 | -1.22E-01 | -1.09E+00 |
| ENSG000002 | --- | 7.89E-01 | ? | | 9.54E-01 | -6.78E-02 | -1.05E+00 |
| ENSG000001 | C1orf74 | 7.89E-01 | | 9.26E-01 | 1.04E+00 | 5.13E-02 | 1.04E+00 |
| ENSG00000C | ACAT1 | 7.89E-01 | | 9.26E-01 | 9.89E-01 | -1.60E-02 | -1.01E+00 |
| ENSG000002 | --- | 7.89E-01 | | 9.26E-01 | 9.59E-01 | -6.07E-02 | -1.04E+00 |
| ENSG000001 | TM9SF4 | 7.89E-01 | | 9.26E-01 | 1.01E+00 | 1.14E-02 | 1.01E+00 |
| ENSG000001 | ZNRF3 | 7.90E-01 | | 9.26E-01 | 9.88E-01 | -1.77E-02 | -1.01E+00 |
| ENSG000002 | --- | 7.90E-01 | ? | | 8.84E-01 | -1.78E-01 | -1.13E+00 |
| ENSG000001 | HMCN1 | 7.90E-01 | | 9.26E-01 | 1.04E+00 | 6.18E-02 | 1.04E+00 |
| ENSG000001 | ZCCHC10 | 7.90E-01 | | 9.26E-01 | 9.89E-01 | -1.57E-02 | -1.01E+00 |
| ENSG000001 | PABIR2 | 7.90E-01 | | 9.26E-01 | 9.89E-01 | -1.53E-02 | -1.01E+00 |
| ENSG000001 | SLC35E4 | 7.90E-01 | | 9.26E-01 | 9.76E-01 | -3.53E-02 | -1.02E+00 |
| ENSG000002 | --- | 7.90E-01 | ? | | 1.13E+00 | 1.78E-01 | 1.13E+00 |
| ENSG000002 | ARRDC1-AS1 | 7.90E-01 | | 9.26E-01 | 1.02E+00 | 2.66E-02 | 1.02E+00 |
| ENSG000002 | --- | 7.90E-01 | ? | | 1.05E+00 | 7.38E-02 | 1.05E+00 |
| ENSG000001 | C4orf36 | 7.90E-01 | | 9.26E-01 | 1.04E+00 | 5.25E-02 | 1.04E+00 |
| ENSG000002 | ZNF225 | 7.90E-01 | | 9.26E-01 | 9.82E-01 | -2.65E-02 | -1.02E+00 |
| ENSG00000C | AARS1 | 7.90E-01 | | 9.26E-01 | 9.91E-01 | -1.30E-02 | -1.01E+00 |
| ENSG000002 | H3P6 | 7.90E-01 | | 9.26E-01 | 9.66E-01 | -5.06E-02 | -1.04E+00 |
| ENSG000002 | --- | 7.90E-01 | ? | | 9.20E-01 | -1.20E-01 | -1.09E+00 |
| ENSG000001 | ADHFE1 | 7.90E-01 | | 9.26E-01 | 9.64E-01 | -5.28E-02 | -1.04E+00 |
| ENSG000001 | ZER1 | 7.90E-01 | | 9.26E-01 | 9.87E-01 | -1.84E-02 | -1.01E+00 |
| ENSG000001 | LRRC66 | 7.90E-01 | | 9.27E-01 | 9.55E-01 | -6.61E-02 | -1.05E+00 |
| ENSG000001 | FGFR4 | 7.90E-01 | ? | | 9.48E-01 | -7.76E-02 | -1.06E+00 |
| ENSG000001 | TMEM220 | 7.90E-01 | ? | | 1.06E+00 | 7.75E-02 | 1.06E+00 |
| ENSG000002 | --- | 7.90E-01 | ? | | 1.11E+00 | 1.53E-01 | 1.11E+00 |
| ENSG000001 | ZNF239 | 7.90E-01 | | 9.27E-01 | 9.78E-01 | -3.19E-02 | -1.02E+00 |
| ENSG000002 | MRFAP1P1 | 7.91E-01 | ? | | 1.10E+00 | 1.43E-01 | 1.10E+00 |
| ENSG000001 | TRPV4 | 7.91E-01 | | 9.27E-01 | 1.03E+00 | 4.89E-02 | 1.03E+00 |
| ENSG000001 | FAAH2 | 7.91E-01 | | 9.27E-01 | 1.03E+00 | 4.58E-02 | 1.03E+00 |
| ENSG000002 | NKX2-1-AS1 | 7.91E-01 | ? | | 1.15E+00 | 1.98E-01 | 1.15E+00 |
| ENSG000002 | --- | 7.91E-01 | ? | | 9.32E-01 | -1.02E-01 | -1.07E+00 |
| ENSG00000C | TNFRSF12A | 7.91E-01 | | 9.27E-01 | 1.01E+00 | 2.12E-02 | 1.01E+00 |
| ENSG000001 | HCAR1 | 7.91E-01 | | 9.27E-01 | 1.03E+00 | 4.34E-02 | 1.03E+00 |
| ENSG00000C | ADA2 | 7.91E-01 | ? | | 9.32E-01 | -1.01E-01 | -1.07E+00 |
| ENSG000001 | C1QBP | 7.91E-01 | | 9.27E-01 | 1.01E+00 | 1.52E-02 | 1.01E+00 |
| ENSG000001 | PARD6A | 7.91E-01 | ? | | 9.28E-01 | -1.07E-01 | -1.08E+00 |
| ENSG000001 | MTERF3 | 7.91E-01 | | 9.27E-01 | 9.88E-01 | -1.70E-02 | -1.01E+00 |
| ENSG000001 | CILK1 | 7.91E-01 | | 9.27E-01 | 1.01E+00 | 1.63E-02 | 1.01E+00 |
| ENSG000001 | MFSD14B | 7.91E-01 | | 9.27E-01 | 1.01E+00 | 1.12E-02 | 1.01E+00 |

| | | | | | | |
|---|---|---|---|---|---|---|
| ENSG000001 | TIGD6 | 7.92E-01 | 9.27E-01 | 1.01E+00 | 2.11E-02 | 1.01E+00 |
| ENSG000001 | PM20D2 | 7.92E-01 | 9.27E-01 | 1.01E+00 | 1.98E-02 | 1.01E+00 |
| ENSG000000 | PLEKHH3 | 7.92E-01 | 9.27E-01 | 9.80E-01 | -2.90E-02 | -1.02E+00 |
| ENSG000001 | MEIG1 | 7.92E-01 | ? | 1.11E+00 | 1.48E-01 | 1.11E+00 |
| ENSG000001 | DDX28 | 7.92E-01 | 9.27E-01 | 1.02E+00 | 2.90E-02 | 1.02E+00 |
| ENSG000002 | --- | 7.92E-01 | ? | 8.86E-01 | -1.75E-01 | -1.13E+00 |
| ENSG000002 | --- | 7.92E-01 | ? | 1.10E+00 | 1.35E-01 | 1.10E+00 |
| ENSG000000 | MSANTD3 | 7.92E-01 | 9.27E-01 | 9.87E-01 | -1.88E-02 | -1.01E+00 |
| ENSG000001 | EXOSC8 | 7.92E-01 | 9.27E-01 | 9.86E-01 | -1.97E-02 | -1.01E+00 |
| ENSG000001 | DIPK2B | 7.92E-01 | ? | 1.14E+00 | 1.87E-01 | 1.14E+00 |
| ENSG000002 | PRKCIP1 | 7.92E-01 | ? | 1.14E+00 | 1.85E-01 | 1.14E+00 |
| ENSG000001 | ZNF274 | 7.92E-01 | 9.27E-01 | 9.90E-01 | -1.49E-02 | -1.01E+00 |
| ENSG000002 | --- | 7.92E-01 | 9.27E-01 | 9.67E-01 | -4.88E-02 | -1.03E+00 |
| ENSG000001 | TCF25 | 7.92E-01 | 9.27E-01 | 1.01E+00 | 1.79E-02 | 1.01E+00 |
| ENSG000001 | KCNMA1 | 7.92E-01 | 9.27E-01 | 1.01E+00 | 1.68E-02 | 1.01E+00 |
| ENSG000001 | CNKSR1 | 7.92E-01 | 9.27E-01 | 1.01E+00 | 1.93E-02 | 1.01E+00 |
| ENSG000002 | --- | 7.92E-01 | ? | 9.56E-01 | -6.44E-02 | -1.05E+00 |
| ENSG000001 | PIP5K1C | 7.92E-01 | 9.27E-01 | 1.01E+00 | 1.74E-02 | 1.01E+00 |
| ENSG000001 | CCDC103 | 7.93E-01 | 9.28E-01 | 1.04E+00 | 5.45E-02 | 1.04E+00 |
| ENSG000002 | TPI1P2 | 7.93E-01 | ? | 1.05E+00 | 6.62E-02 | 1.05E+00 |
| ENSG000001 | SRSF7 | 7.93E-01 | 9.28E-01 | 1.01E+00 | 1.22E-02 | 1.01E+00 |
| ENSG000001 | GRM1 | 7.93E-01 | ? | 8.98E-01 | -1.55E-01 | -1.11E+00 |
| ENSG000002 | --- | 7.93E-01 | ? | 9.00E-01 | -1.53E-01 | -1.11E+00 |
| ENSG000001 | TRIM2 | 7.93E-01 | 9.28E-01 | 9.88E-01 | -1.73E-02 | -1.01E+00 |
| ENSG000001 | C2orf88 | 7.93E-01 | ? | 1.05E+00 | 7.58E-02 | 1.05E+00 |
| ENSG000001 | CLK4 | 7.93E-01 | 9.28E-01 | 1.02E+00 | 2.30E-02 | 1.02E+00 |
| ENSG000001 | PAX9 | 7.93E-01 | 9.28E-01 | 1.01E+00 | 1.73E-02 | 1.01E+00 |
| ENSG000001 | RDH16 | 7.93E-01 | ? | 1.07E+00 | 1.02E-01 | 1.07E+00 |
| ENSG000001 | TMEM94 | 7.93E-01 | 9.28E-01 | 1.01E+00 | 1.98E-02 | 1.01E+00 |
| ENSG000001 | GPN2 | 7.93E-01 | 9.28E-01 | 1.01E+00 | 1.29E-02 | 1.01E+00 |
| ENSG000001 | SUMO2 | 7.94E-01 | 9.28E-01 | 1.01E+00 | 1.34E-02 | 1.01E+00 |
| ENSG000000 | HOMER3 | 7.94E-01 | 9.28E-01 | 1.01E+00 | 1.70E-02 | 1.01E+00 |
| ENSG000002 | LINC00960 | 7.94E-01 | 9.28E-01 | 1.01E+00 | 1.45E-02 | 1.01E+00 |
| ENSG000002 | --- | 7.94E-01 | ? | 1.17E+00 | 2.31E-01 | 1.17E+00 |
| ENSG000002 | SSR4P1 | 7.94E-01 | ? | 1.07E+00 | 9.22E-02 | 1.07E+00 |
| ENSG000001 | DAPK1 | 7.94E-01 | 9.28E-01 | 1.03E+00 | 3.85E-02 | 1.03E+00 |
| ENSG000001 | AIMP2 | 7.94E-01 | 9.28E-01 | 1.01E+00 | 1.33E-02 | 1.01E+00 |
| ENSG000002 | --- | 7.94E-01 | ? | 8.77E-01 | -1.89E-01 | -1.14E+00 |
| ENSG000002 | --- | 7.94E-01 | 9.28E-01 | 1.05E+00 | 6.69E-02 | 1.05E+00 |
| ENSG000001 | IER5 | 7.94E-01 | 9.28E-01 | 9.87E-01 | -1.82E-02 | -1.01E+00 |
| ENSG000001 | ZNRF2 | 7.94E-01 | 9.28E-01 | 9.86E-01 | -2.00E-02 | -1.01E+00 |
| ENSG000001 | OTULINL | 7.94E-01 | 9.28E-01 | 1.04E+00 | 5.28E-02 | 1.04E+00 |
| ENSG000002 | LINC01792 | 7.94E-01 | ? | 9.57E-01 | -6.39E-02 | -1.05E+00 |
| ENSG000002 | NRSN2-AS1 | 7.94E-01 | 9.28E-01 | 9.78E-01 | -3.23E-02 | -1.02E+00 |
| ENSG000001 | SFXN1 | 7.94E-01 | 9.28E-01 | 9.89E-01 | -1.57E-02 | -1.01E+00 |

| | | | | | | |
|---|---|---|---|---|---|---|
| ENSG000001 | INO80B | 7.94E-01 | 9.28E-01 | 9.70E-01 | -4.37E-02 | -1.03E+00 |
| ENSG000002 | --- | 7.94E-01 | ? | 1.08E+00 | 1.11E-01 | 1.08E+00 |
| ENSG000001 | F13A1 | 7.94E-01 | ? | 9.30E-01 | -1.05E-01 | -1.08E+00 |
| ENSG000001 | ADAP2 | 7.94E-01 | 9.28E-01 | 1.02E+00 | 3.15E-02 | 1.02E+00 |
| ENSG000001 | ASB16 | 7.94E-01 | 9.28E-01 | 9.59E-01 | -6.01E-02 | -1.04E+00 |
| ENSG000001 | MRPS15 | 7.94E-01 | 9.28E-01 | 1.01E+00 | 1.20E-02 | 1.01E+00 |
| ENSG000002 | MRPS18B | 7.94E-01 | 9.28E-01 | 1.01E+00 | 1.61E-02 | 1.01E+00 |
| ENSG000001 | SLC16A12 | 7.94E-01 | 9.28E-01 | 9.68E-01 | -4.71E-02 | -1.03E+00 |
| ENSG000001 | ZNF382 | 7.94E-01 | 9.28E-01 | 1.02E+00 | 2.37E-02 | 1.02E+00 |
| ENSG000002 | --- | 7.94E-01 | ? | 8.73E-01 | -1.97E-01 | -1.15E+00 |
| ENSG000001 | RHNO1 | 7.94E-01 | 9.28E-01 | 9.81E-01 | -2.69E-02 | -1.02E+00 |
| ENSG000001 | MPP1 | 7.95E-01 | 9.28E-01 | 1.02E+00 | 3.16E-02 | 1.02E+00 |
| ENSG000002 | TNFSF12 | 7.95E-01 | 9.28E-01 | 1.03E+00 | 3.88E-02 | 1.03E+00 |
| ENSG000001 | OSBPL10 | 7.95E-01 | 9.28E-01 | 9.91E-01 | -1.33E-02 | -1.01E+00 |
| ENSG000001 | RRP1 | 7.95E-01 | 9.28E-01 | 1.01E+00 | 1.77E-02 | 1.01E+00 |
| ENSG000001 | AFMID | 7.95E-01 | 9.28E-01 | 1.02E+00 | 2.36E-02 | 1.02E+00 |
| ENSG000002 | LINC01182 | 7.95E-01 | ? | 9.20E-01 | -1.21E-01 | -1.09E+00 |
| ENSG000002 | --- | 7.95E-01 | ? | 9.28E-01 | -1.08E-01 | -1.08E+00 |
| ENSG000002 | TBC1D3 | 7.95E-01 | ? | 8.67E-01 | -2.06E-01 | -1.15E+00 |
| ENSG000002 | C4B | 7.95E-01 | ? | 1.07E+00 | 9.57E-02 | 1.07E+00 |
| ENSG000002 | --- | 7.95E-01 | ? | 9.26E-01 | -1.11E-01 | -1.08E+00 |
| ENSG000002 | --- | 7.95E-01 | ? | 1.05E+00 | 6.65E-02 | 1.05E+00 |
| ENSG000001 | C3orf38 | 7.95E-01 | 9.28E-01 | 9.89E-01 | -1.67E-02 | -1.01E+00 |
| ENSG000002 | SERBP1P6 | 7.95E-01 | ? | 1.07E+00 | 9.97E-02 | 1.07E+00 |
| ENSG000002 | PYY2 | 7.95E-01 | ? | 1.08E+00 | 1.07E-01 | 1.08E+00 |
| ENSG000001 | ZNF732 | 7.95E-01 | ? | 1.09E+00 | 1.21E-01 | 1.09E+00 |
| ENSG000002 | SMIM15-AS1 | 7.95E-01 | ? | 9.47E-01 | -7.92E-02 | -1.06E+00 |
| ENSG000002 | HSBP1L1 | 7.95E-01 | 9.28E-01 | 1.01E+00 | 2.14E-02 | 1.01E+00 |
| ENSG000002 | KIF1C-AS1 | 7.96E-01 | ? | 9.18E-01 | -1.23E-01 | -1.09E+00 |
| ENSG000002 | --- | 7.96E-01 | 9.28E-01 | 9.85E-01 | -2.18E-02 | -1.02E+00 |
| ENSG000001 | PPARA | 7.96E-01 | 9.28E-01 | 1.01E+00 | 1.75E-02 | 1.01E+00 |
| ENSG000001 | PALD1 | 7.96E-01 | ? | 1.07E+00 | 9.63E-02 | 1.07E+00 |
| ENSG000001 | ARHGAP1 | 7.96E-01 | 9.28E-01 | 1.01E+00 | 1.45E-02 | 1.01E+00 |
| ENSG000002 | ZNF516-DT | 7.96E-01 | ? | 1.08E+00 | 1.06E-01 | 1.08E+00 |
| ENSG000001 | ANKMY1 | 7.96E-01 | 9.28E-01 | 1.02E+00 | 3.43E-02 | 1.02E+00 |
| ENSG000001 | CUX2 | 7.96E-01 | 9.28E-01 | 9.83E-01 | -2.50E-02 | -1.02E+00 |
| ENSG000002 | --- | 7.96E-01 | ? | 9.29E-01 | -1.06E-01 | -1.08E+00 |
| ENSG00000C | SLC9A3 | 7.96E-01 | 9.28E-01 | 1.03E+00 | 4.03E-02 | 1.03E+00 |
| ENSG000002 | --- | 7.96E-01 | ? | 1.15E+00 | 1.97E-01 | 1.15E+00 |
| ENSG00000C | FTL | 7.96E-01 | 9.28E-01 | 1.01E+00 | 2.11E-02 | 1.01E+00 |
| ENSG000001 | ST8SIA5 | 7.96E-01 | ? | 9.35E-01 | -9.75E-02 | -1.07E+00 |
| ENSG000002 | PTPRG-AS1 | 7.96E-01 | ? | 9.49E-01 | -7.54E-02 | -1.05E+00 |
| ENSG000001 | AHCYL1 | 7.96E-01 | 9.28E-01 | 9.94E-01 | -8.60E-03 | -1.01E+00 |
| ENSG000002 | MCM3AP-AS | 7.96E-01 | 9.28E-01 | 9.83E-01 | -2.54E-02 | -1.02E+00 |
| ENSG000001 | SLC22A15 | 7.96E-01 | 9.28E-01 | 1.01E+00 | 1.61E-02 | 1.01E+00 |

| | | | | | | |
|---|---|---|---|---|---|---|
| ENSG000002 | GSK3B-DT | 7.96E-01 | 9.28E-01 | 1.04E+00 | 5.31E-02 | 1.04E+00 |
| ENSG000001 | FAM184A | 7.96E-01 | ? | 1.08E+00 | 1.06E-01 | 1.08E+00 |
| ENSG000001 | PLPP2 | 7.96E-01 | 9.28E-01 | 1.03E+00 | 3.75E-02 | 1.03E+00 |
| ENSG000001 | ACTR3B | 7.96E-01 | 9.28E-01 | 9.82E-01 | -2.64E-02 | -1.02E+00 |
| ENSG000001 | TMEM50B | 7.96E-01 | 9.28E-01 | 1.01E+00 | 1.49E-02 | 1.01E+00 |
| ENSG000001 | STX5 | 7.96E-01 | 9.28E-01 | 1.01E+00 | 1.49E-02 | 1.01E+00 |
| ENSG000001 | FAM124A | 7.97E-01 | ? | 9.19E-01 | -1.21E-01 | -1.09E+00 |
| ENSG000001 | CLCA3P | 7.97E-01 | ? | 1.25E+00 | 3.24E-01 | 1.25E+00 |
| ENSG000001 | ZNF606 | 7.97E-01 | 9.28E-01 | 1.02E+00 | 2.21E-02 | 1.02E+00 |
| ENSG000001 | ARMCX4 | 7.97E-01 | 9.28E-01 | 9.79E-01 | -3.05E-02 | -1.02E+00 |
| ENSG000001 | DPP9 | 7.97E-01 | 9.28E-01 | 1.01E+00 | 1.57E-02 | 1.01E+00 |
| ENSG000001 | FAM241A | 7.97E-01 | 9.28E-01 | 9.84E-01 | -2.31E-02 | -1.02E+00 |
| ENSG000002 | LINC00899 | 7.97E-01 | 9.28E-01 | 9.76E-01 | -3.51E-02 | -1.02E+00 |
| ENSG000001 | PARP4 | 7.97E-01 | 9.28E-01 | 1.01E+00 | 9.24E-03 | 1.01E+00 |
| ENSG000001 | IQCN | 7.97E-01 | ? | 1.08E+00 | 1.15E-01 | 1.08E+00 |
| ENSG000001 | APTX | 7.97E-01 | 9.28E-01 | 1.01E+00 | 1.16E-02 | 1.01E+00 |
| ENSG00000C | DBNDD1 | 7.97E-01 | 9.28E-01 | 9.83E-01 | -2.48E-02 | -1.02E+00 |
| ENSG000001 | ATP9B | 7.97E-01 | 9.28E-01 | 1.01E+00 | 1.48E-02 | 1.01E+00 |
| ENSG000001 | GUCY2D | 7.97E-01 | ? | 1.13E+00 | 1.72E-01 | 1.13E+00 |
| ENSG000002 | --- | 7.97E-01 | ? | 1.13E+00 | 1.72E-01 | 1.13E+00 |
| ENSG000002 | ZNF229 | 7.97E-01 | 9.29E-01 | 9.90E-01 | -1.51E-02 | -1.01E+00 |
| ENSG00000C | MOV10L1 | 7.97E-01 | ? | 9.45E-01 | -8.12E-02 | -1.06E+00 |
| ENSG000001 | DEK | 7.97E-01 | 9.29E-01 | 1.01E+00 | 1.59E-02 | 1.01E+00 |
| ENSG000001 | SUPV3L1 | 7.97E-01 | 9.29E-01 | 9.90E-01 | -1.42E-02 | -1.01E+00 |
| ENSG000002 | TPRG1LP1 | 7.97E-01 | ? | 9.07E-01 | -1.41E-01 | -1.10E+00 |
| ENSG000001 | RAB15 | 7.98E-01 | 9.29E-01 | 9.75E-01 | -3.63E-02 | -1.03E+00 |
| ENSG000001 | MMADHC | 7.98E-01 | 9.29E-01 | 1.01E+00 | 1.01E-02 | 1.01E+00 |
| ENSG000001 | COPG1 | 7.98E-01 | 9.29E-01 | 9.91E-01 | -1.32E-02 | -1.01E+00 |
| ENSG000001 | PWP1 | 7.98E-01 | 9.29E-01 | 1.01E+00 | 1.13E-02 | 1.01E+00 |
| ENSG000002 | --- | 7.98E-01 | 9.29E-01 | 1.02E+00 | 3.10E-02 | 1.02E+00 |
| ENSG000001 | PMFBP1 | 7.98E-01 | ? | 9.35E-01 | -9.64E-02 | -1.07E+00 |
| ENSG000001 | RAB40A | 7.98E-01 | 9.29E-01 | 9.63E-01 | -5.46E-02 | -1.04E+00 |
| ENSG000002 | --- | 7.98E-01 | ? | 9.44E-01 | -8.34E-02 | -1.06E+00 |
| ENSG000001 | STK35 | 7.98E-01 | 9.29E-01 | 9.91E-01 | -1.23E-02 | -1.01E+00 |
| ENSG000002 | ZNF767P | 7.98E-01 | 9.29E-01 | 9.81E-01 | -2.82E-02 | -1.02E+00 |
| ENSG000001 | SCLY | 7.98E-01 | 9.29E-01 | 1.01E+00 | 1.94E-02 | 1.01E+00 |
| ENSG000001 | PARP6 | 7.98E-01 | 9.29E-01 | 1.01E+00 | 2.04E-02 | 1.01E+00 |
| ENSG000001 | MTHFD1L | 7.98E-01 | 9.29E-01 | 9.92E-01 | -1.18E-02 | -1.01E+00 |
| ENSG000001 | GPI | 7.98E-01 | 9.29E-01 | 1.01E+00 | 1.21E-02 | 1.01E+00 |
| ENSG000002 | --- | 7.99E-01 | ? | 8.70E-01 | -2.01E-01 | -1.15E+00 |
| ENSG000001 | SNX31 | 7.99E-01 | ? | 1.08E+00 | 1.14E-01 | 1.08E+00 |
| ENSG000002 | AP5Z1 | 7.99E-01 | 9.29E-01 | 9.82E-01 | -2.68E-02 | -1.02E+00 |
| ENSG000001 | GALNT13 | 7.99E-01 | ? | 8.97E-01 | -1.56E-01 | -1.11E+00 |
| ENSG000001 | NPM1 | 7.99E-01 | 9.29E-01 | 9.90E-01 | -1.49E-02 | -1.01E+00 |
| ENSG000001 | LRWD1 | 7.99E-01 | 9.29E-01 | 1.01E+00 | 2.08E-02 | 1.01E+00 |

| | | | | | | |
|---|---|---|---|---|---|---|
| ENSG000002 | MSTO2P | 7.99E-01 | ? | 9.56E-01 | -6.42E-02 | -1.05E+00 |
| ENSG000001 | TMEM140 | 7.99E-01 | 9.29E-01 | 1.01E+00 | 1.76E-02 | 1.01E+00 |
| ENSG000002 | --- | 7.99E-01 | ? | 1.09E+00 | 1.26E-01 | 1.09E+00 |
| ENSG000001 | MSANTD4 | 7.99E-01 | 9.29E-01 | 1.01E+00 | 1.52E-02 | 1.01E+00 |
| ENSG000001 | RECQL5 | 7.99E-01 | 9.29E-01 | 1.02E+00 | 2.19E-02 | 1.02E+00 |
| ENSG000001 | FAM118A | 7.99E-01 | 9.29E-01 | 1.01E+00 | 1.63E-02 | 1.01E+00 |
| ENSG000001 | MLH3 | 7.99E-01 | 9.29E-01 | 1.01E+00 | 1.29E-02 | 1.01E+00 |
| ENSG000002 | KRT18P8 | 7.99E-01 | ? | 1.11E+00 | 1.54E-01 | 1.11E+00 |
| ENSG000001 | CD93 | 7.99E-01 | ? | 9.06E-01 | -1.43E-01 | -1.10E+00 |
| ENSG000002 | RTEL1 | 7.99E-01 | 9.29E-01 | 1.02E+00 | 2.71E-02 | 1.02E+00 |
| ENSG000001 | PLEKHM3 | 8.00E-01 | 9.29E-01 | 9.90E-01 | -1.41E-02 | -1.01E+00 |
| ENSG000002 | UNC5B-AS1 | 8.00E-01 | ? | 1.07E+00 | 1.03E-01 | 1.07E+00 |
| ENSG000002 | --- | 8.00E-01 | ? | 8.82E-01 | -1.81E-01 | -1.13E+00 |
| ENSG000002 | EIF3CL | 8.00E-01 | 9.30E-01 | 1.04E+00 | 5.33E-02 | 1.04E+00 |
| ENSG000002 | SLC41A3-AS1 | 8.00E-01 | ? | 1.11E+00 | 1.48E-01 | 1.11E+00 |
| ENSG000001 | GLS2 | 8.00E-01 | ? | 9.57E-01 | -6.41E-02 | -1.05E+00 |
| ENSG000001 | NCOA7 | 8.00E-01 | 9.30E-01 | 1.01E+00 | 1.94E-02 | 1.01E+00 |
| ENSG000001 | GCOM1 | 8.00E-01 | 9.30E-01 | 1.02E+00 | 3.50E-02 | 1.02E+00 |
| ENSG000001 | SIAH1 | 8.00E-01 | 9.30E-01 | 1.01E+00 | 1.61E-02 | 1.01E+00 |
| ENSG000001 | OCLN | 8.00E-01 | 9.30E-01 | 9.77E-01 | -3.32E-02 | -1.02E+00 |
| ENSG000002 | --- | 8.00E-01 | 9.30E-01 | 9.60E-01 | -5.83E-02 | -1.04E+00 |
| ENSG000001 | SIN3B | 8.00E-01 | 9.30E-01 | 1.01E+00 | 1.41E-02 | 1.01E+00 |
| ENSG000002 | VPS52 | 8.00E-01 | 9.30E-01 | 9.91E-01 | -1.28E-02 | -1.01E+00 |
| ENSG000002 | PDCD6 | 8.00E-01 | 9.30E-01 | 1.01E+00 | 1.23E-02 | 1.01E+00 |
| ENSG000002 | RORA-AS1 | 8.00E-01 | ? | 1.11E+00 | 1.54E-01 | 1.11E+00 |
| ENSG000000 | TRIB2 | 8.01E-01 | 9.30E-01 | 9.81E-01 | -2.71E-02 | -1.02E+00 |
| ENSG000001 | SYTL1 | 8.01E-01 | 9.30E-01 | 1.02E+00 | 3.16E-02 | 1.02E+00 |
| ENSG000002 | --- | 8.01E-01 | ? | 1.11E+00 | 1.47E-01 | 1.11E+00 |
| ENSG000002 | --- | 8.01E-01 | 9.30E-01 | 1.04E+00 | 6.01E-02 | 1.04E+00 |
| ENSG000001 | MAPK15 | 8.01E-01 | 9.30E-01 | 1.03E+00 | 4.60E-02 | 1.03E+00 |
| ENSG000002 | --- | 8.01E-01 | ? | 9.60E-01 | -5.93E-02 | -1.04E+00 |
| ENSG000000 | XPO1 | 8.01E-01 | 9.30E-01 | 9.93E-01 | -9.75E-03 | -1.01E+00 |
| ENSG000002 | --- | 8.01E-01 | ? | 9.51E-01 | -7.29E-02 | -1.05E+00 |
| ENSG000001 | ADAMTS12 | 8.01E-01 | 9.30E-01 | 1.05E+00 | 7.53E-02 | 1.05E+00 |
| ENSG000001 | SCNN1D | 8.01E-01 | ? | 1.07E+00 | 9.14E-02 | 1.07E+00 |
| ENSG000001 | TNIP2 | 8.01E-01 | 9.30E-01 | 9.89E-01 | -1.62E-02 | -1.01E+00 |
| ENSG000000 | CEACAM1 | 8.01E-01 | 9.30E-01 | 9.85E-01 | -2.16E-02 | -1.02E+00 |
| ENSG000002 | SPECC1L-ADO | 8.01E-01 | ? | 9.30E-01 | -1.05E-01 | -1.08E+00 |
| ENSG000002 | SLFNL1-AS1 | 8.02E-01 | 9.30E-01 | 1.03E+00 | 4.30E-02 | 1.03E+00 |
| ENSG000001 | ASPHD2 | 8.02E-01 | 9.30E-01 | 1.01E+00 | 1.98E-02 | 1.01E+00 |
| ENSG000002 | ACTG1P17 | 8.02E-01 | ? | 9.13E-01 | -1.31E-01 | -1.09E+00 |
| ENSG000002 | LINC00624 | 8.02E-01 | 9.31E-01 | 9.60E-01 | -5.90E-02 | -1.04E+00 |
| ENSG000002 | --- | 8.02E-01 | 9.31E-01 | 9.62E-01 | -5.63E-02 | -1.04E+00 |
| ENSG000001 | PIGU | 8.02E-01 | 9.31E-01 | 9.90E-01 | -1.46E-02 | -1.01E+00 |
| ENSG000001 | MFGE8 | 8.02E-01 | 9.31E-01 | 9.84E-01 | -2.35E-02 | -1.02E+00 |

| | | | | | | |
|---|---|---|---|---|---|---|
| ENSG000002 | --- | 8.02E-01 | 9.31E-01 | 1.05E+00 | 6.78E-02 | 1.05E+00 |
| ENSG000002 | --- | 8.02E-01 | ? | 9.28E-01 | -1.07E-01 | -1.08E+00 |
| ENSG000001 | FBXO22 | 8.02E-01 | 9.31E-01 | 1.01E+00 | 1.49E-02 | 1.01E+00 |
| ENSG000001 | KBTBD8 | 8.02E-01 | 9.31E-01 | 1.01E+00 | 2.06E-02 | 1.01E+00 |
| ENSG000001 | KLK12 | 8.02E-01 | ? | 8.93E-01 | -1.64E-01 | -1.12E+00 |
| ENSG000002 | LINC00997 | 8.02E-01 | 9.31E-01 | 9.66E-01 | -5.04E-02 | -1.04E+00 |
| ENSG000001 | APOL3 | 8.02E-01 | ? | 9.40E-01 | -8.90E-02 | -1.06E+00 |
| ENSG000002 | --- | 8.02E-01 | ? | 1.09E+00 | 1.26E-01 | 1.09E+00 |
| ENSG000001 | GPR75 | 8.02E-01 | 9.31E-01 | 9.69E-01 | -4.52E-02 | -1.03E+00 |
| ENSG000001 | KCNMB4 | 8.02E-01 | 9.31E-01 | 9.69E-01 | -4.54E-02 | -1.03E+00 |
| ENSG000002 | HAUS7 | 8.02E-01 | 9.31E-01 | 9.82E-01 | -2.64E-02 | -1.02E+00 |
| ENSG000002 | AHCTF1P1 | 8.03E-01 | 9.31E-01 | 9.76E-01 | -3.49E-02 | -1.02E+00 |
| ENSG000001 | CLTRN | 8.03E-01 | 9.31E-01 | 9.68E-01 | -4.70E-02 | -1.03E+00 |
| ENSG000001 | EYA1 | 8.03E-01 | 9.31E-01 | 1.02E+00 | 2.33E-02 | 1.02E+00 |
| ENSG000002 | RPL23AP7 | 8.03E-01 | 9.31E-01 | 1.01E+00 | 1.47E-02 | 1.01E+00 |
| ENSG00000C | RASAL2 | 8.03E-01 | 9.31E-01 | 9.88E-01 | -1.68E-02 | -1.01E+00 |
| ENSG000002 | --- | 8.03E-01 | 9.31E-01 | 8.95E-01 | -1.60E-01 | -1.12E+00 |
| ENSG000001 | B3GNT7 | 8.03E-01 | ? | 1.06E+00 | 8.21E-02 | 1.06E+00 |
| ENSG000002 | --- | 8.03E-01 | ? | 1.13E+00 | 1.79E-01 | 1.13E+00 |
| ENSG000002 | --- | 8.03E-01 | ? | 1.12E+00 | 1.58E-01 | 1.12E+00 |
| ENSG000001 | MAK | 8.03E-01 | ? | 1.10E+00 | 1.40E-01 | 1.10E+00 |
| ENSG000001 | ZNF764 | 8.03E-01 | 9.31E-01 | 1.02E+00 | 3.25E-02 | 1.02E+00 |
| ENSG000001 | AGAP2 | 8.03E-01 | 9.31E-01 | 9.62E-01 | -5.58E-02 | -1.04E+00 |
| ENSG000002 | --- | 8.03E-01 | 9.31E-01 | 1.02E+00 | 2.87E-02 | 1.02E+00 |
| ENSG000001 | COL8A2 | 8.04E-01 | 9.31E-01 | 9.77E-01 | -3.38E-02 | -1.02E+00 |
| ENSG000002 | RPL29P11 | 8.04E-01 | ? | 9.21E-01 | -1.19E-01 | -1.09E+00 |
| ENSG000001 | VMAC | 8.04E-01 | 9.31E-01 | 1.03E+00 | 3.95E-02 | 1.03E+00 |
| ENSG000001 | BORA | 8.04E-01 | 9.31E-01 | 9.84E-01 | -2.40E-02 | -1.02E+00 |
| ENSG000002 | TMEM170B | 8.04E-01 | ? | 9.37E-01 | -9.41E-02 | -1.07E+00 |
| ENSG000002 | LINC00941 | 8.04E-01 | 9.31E-01 | 9.35E-01 | -9.70E-02 | -1.07E+00 |
| ENSG000001 | RAB5C | 8.04E-01 | 9.31E-01 | 9.92E-01 | -1.09E-02 | -1.01E+00 |
| ENSG000002 | --- | 8.04E-01 | ? | 1.05E+00 | 6.48E-02 | 1.05E+00 |
| ENSG000001 | DPF2 | 8.04E-01 | 9.31E-01 | 9.92E-01 | -1.23E-02 | -1.01E+00 |
| ENSG000002 | --- | 8.04E-01 | 9.31E-01 | 9.58E-01 | -6.23E-02 | -1.04E+00 |
| ENSG000002 | ZNF90 | 8.04E-01 | 9.31E-01 | 9.78E-01 | -3.19E-02 | -1.02E+00 |
| ENSG000001 | SPRR2G | 8.04E-01 | ? | 1.09E+00 | 1.21E-01 | 1.09E+00 |
| ENSG000001 | COL21A1 | 8.04E-01 | ? | 9.49E-01 | -7.55E-02 | -1.05E+00 |
| ENSG000002 | --- | 8.04E-01 | ? | 1.10E+00 | 1.33E-01 | 1.10E+00 |
| ENSG00000C | GUCY1B1 | 8.04E-01 | ? | 1.06E+00 | 8.46E-02 | 1.06E+00 |
| ENSG000002 | --- | 8.04E-01 | 9.31E-01 | 1.02E+00 | 2.21E-02 | 1.02E+00 |
| ENSG00000C | ZFYVE26 | 8.04E-01 | 9.31E-01 | 1.01E+00 | 1.51E-02 | 1.01E+00 |
| ENSG000001 | YPEL1 | 8.04E-01 | ? | 9.18E-01 | -1.23E-01 | -1.09E+00 |
| ENSG000002 | ACTBP2 | 8.04E-01 | ? | 9.42E-01 | -8.61E-02 | -1.06E+00 |
| ENSG000001 | NPAT | 8.05E-01 | 9.31E-01 | 9.89E-01 | -1.54E-02 | -1.01E+00 |
| ENSG000002 | --- | 8.05E-01 | ? | 9.45E-01 | -8.14E-02 | -1.06E+00 |

| | | | | | | | |
|---|---|---|---|---|---|---|---|
| ENSG000001 | FBXO25 | 8.05E-01 | | 9.32E-01 | 9.91E-01 | -1.30E-02 | -1.01E+00 |
| ENSG000001 | TEK | 8.05E-01 | ? | | 9.44E-01 | -8.33E-02 | -1.06E+00 |
| ENSG000001 | NETO2 | 8.05E-01 | | 9.32E-01 | 1.02E+00 | 2.27E-02 | 1.02E+00 |
| ENSG000001 | RSL1D1 | 8.05E-01 | | 9.32E-01 | 9.91E-01 | -1.35E-02 | -1.01E+00 |
| ENSG000001 | RBM43 | 8.05E-01 | | 9.32E-01 | 1.01E+00 | 1.90E-02 | 1.01E+00 |
| ENSG000002 | RPL35P5 | 8.05E-01 | ? | | 9.46E-01 | -7.96E-02 | -1.06E+00 |
| ENSG000002 | --- | 8.05E-01 | ? | | 1.13E+00 | 1.81E-01 | 1.13E+00 |
| ENSG000001 | ZNF502 | 8.05E-01 | | 9.32E-01 | 9.84E-01 | -2.32E-02 | -1.02E+00 |
| ENSG000002 | --- | 8.05E-01 | ? | | 1.07E+00 | 9.12E-02 | 1.07E+00 |
| ENSG000002 | --- | 8.05E-01 | ? | | 1.07E+00 | 9.12E-02 | 1.07E+00 |
| ENSG000002 | --- | 8.05E-01 | ? | | 1.07E+00 | 9.12E-02 | 1.07E+00 |
| ENSG000001 | KRT16 | 8.05E-01 | | 9.32E-01 | 1.04E+00 | 6.12E-02 | 1.04E+00 |
| ENSG000001 | PDLIM7 | 8.05E-01 | | 9.32E-01 | 9.88E-01 | -1.73E-02 | -1.01E+00 |
| ENSG000001 | ZNF484 | 8.05E-01 | | 9.32E-01 | 1.02E+00 | 2.52E-02 | 1.02E+00 |
| ENSG000002 | BPTFP1 | 8.05E-01 | ? | | 8.38E-01 | -2.55E-01 | -1.19E+00 |
| ENSG000001 | FZD7 | 8.05E-01 | | 9.32E-01 | 9.73E-01 | -3.99E-02 | -1.03E+00 |
| ENSG000001 | DMXL2 | 8.05E-01 | | 9.32E-01 | 1.01E+00 | 2.04E-02 | 1.01E+00 |
| ENSG000002 | HDAC2-AS2 | 8.06E-01 | ? | | 1.07E+00 | 9.26E-02 | 1.07E+00 |
| ENSG000001 | NEURL1 | 8.06E-01 | ? | | 9.06E-01 | -1.42E-01 | -1.10E+00 |
| ENSG000002 | AJM1 | 8.06E-01 | | 9.32E-01 | 1.04E+00 | 5.57E-02 | 1.04E+00 |
| ENSG000001 | TUFM | 8.06E-01 | | 9.32E-01 | 1.01E+00 | 1.25E-02 | 1.01E+00 |
| ENSG000002 | --- | 8.06E-01 | | 9.32E-01 | 8.51E-01 | -2.32E-01 | -1.17E+00 |
| ENSG000001 | SF3A2 | 8.06E-01 | | 9.32E-01 | 1.01E+00 | 1.77E-02 | 1.01E+00 |
| ENSG00000C | SLC8B1 | 8.06E-01 | | 9.32E-01 | 9.86E-01 | -2.03E-02 | -1.01E+00 |
| ENSG000002 | DNAAF4 | 8.06E-01 | | 9.32E-01 | 9.70E-01 | -4.36E-02 | -1.03E+00 |
| ENSG000002 | EDDM13 | 8.06E-01 | ? | | 8.52E-01 | -2.32E-01 | -1.17E+00 |
| ENSG000001 | IRAK1BP1 | 8.06E-01 | | 9.32E-01 | 9.85E-01 | -2.22E-02 | -1.02E+00 |
| ENSG000002 | PWAR5 | 8.06E-01 | | 9.32E-01 | 9.57E-01 | -6.40E-02 | -1.05E+00 |
| ENSG000002 | LINC02802 | 8.06E-01 | ? | | 9.19E-01 | -1.22E-01 | -1.09E+00 |
| ENSG000001 | ADAMTS17 | 8.07E-01 | ? | | 9.41E-01 | -8.73E-02 | -1.06E+00 |
| ENSG000001 | ZFYVE9 | 8.07E-01 | | 9.32E-01 | 9.93E-01 | -1.02E-02 | -1.01E+00 |
| ENSG000001 | GRM4 | 8.07E-01 | ? | | 9.37E-01 | -9.45E-02 | -1.07E+00 |
| ENSG000001 | FCHO1 | 8.07E-01 | | 9.32E-01 | 9.75E-01 | -3.63E-02 | -1.03E+00 |
| ENSG000002 | BCAN-AS2 | 8.07E-01 | ? | | 9.23E-01 | -1.16E-01 | -1.08E+00 |
| ENSG000001 | OSTC | 8.07E-01 | | 9.32E-01 | 1.01E+00 | 1.09E-02 | 1.01E+00 |
| ENSG000001 | TMEM69 | 8.07E-01 | | 9.32E-01 | 1.01E+00 | 1.45E-02 | 1.01E+00 |
| ENSG000002 | --- | 8.07E-01 | | 9.32E-01 | 9.63E-01 | -5.45E-02 | -1.04E+00 |
| ENSG000001 | DNAJB13 | 8.07E-01 | ? | | 9.45E-01 | -8.20E-02 | -1.06E+00 |
| ENSG000002 | CDK3 | 8.07E-01 | | 9.32E-01 | 1.06E+00 | 8.44E-02 | 1.06E+00 |
| ENSG000002 | --- | 8.07E-01 | ? | | 1.10E+00 | 1.35E-01 | 1.10E+00 |
| ENSG000002 | ZNF658 | 8.07E-01 | | 9.32E-01 | 9.85E-01 | -2.16E-02 | -1.02E+00 |
| ENSG000001 | DOK7 | 8.07E-01 | | 9.32E-01 | 9.60E-01 | -5.88E-02 | -1.04E+00 |
| ENSG000002 | --- | 8.07E-01 | | 9.32E-01 | 9.49E-01 | -7.55E-02 | -1.05E+00 |
| ENSG000001 | PLEKHA4 | 8.07E-01 | | 9.32E-01 | 9.67E-01 | -4.85E-02 | -1.03E+00 |
| ENSG000002 | --- | 8.07E-01 | ? | | 1.07E+00 | 1.04E-01 | 1.07E+00 |

| | | | | | | |
|---|---|---|---|---|---|---|
| ENSG000001 | ZMAT2 | 8.07E-01 | 9.32E-01 | 1.01E+00 | 1.09E-02 | 1.01E+00 |
| ENSG000002 | LINC00243 | 8.07E-01 | ? | 9.53E-01 | -6.94E-02 | -1.05E+00 |
| ENSG000002 | CPEB1 | 8.07E-01 | ? | 1.05E+00 | 7.00E-02 | 1.05E+00 |
| ENSG000001 | HMGN3 | 8.07E-01 | 9.32E-01 | 9.87E-01 | -1.90E-02 | -1.01E+00 |
| ENSG000001 | EPC2 | 8.07E-01 | 9.32E-01 | 9.86E-01 | -1.97E-02 | -1.01E+00 |
| ENSG000001 | TNK1 | 8.08E-01 | 9.33E-01 | 1.01E+00 | 1.76E-02 | 1.01E+00 |
| ENSG000001 | NTMT1 | 8.08E-01 | 9.33E-01 | 1.02E+00 | 2.51E-02 | 1.02E+00 |
| ENSG000002 | TDGP1 | 8.08E-01 | ? | 8.87E-01 | -1.73E-01 | -1.13E+00 |
| ENSG000001 | RPP14 | 8.08E-01 | 9.33E-01 | 1.01E+00 | 1.08E-02 | 1.01E+00 |
| ENSG000002 | LINC01465 | 8.08E-01 | ? | 1.07E+00 | 9.20E-02 | 1.07E+00 |
| ENSG000002 | --- | 8.08E-01 | ? | 1.07E+00 | 9.40E-02 | 1.07E+00 |
| ENSG000002 | NAPA-AS1 | 8.08E-01 | ? | 1.05E+00 | 7.11E-02 | 1.05E+00 |
| ENSG000002 | --- | 8.08E-01 | ? | 9.42E-01 | -8.60E-02 | -1.06E+00 |
| ENSG000002 | --- | 8.08E-01 | ? | 1.14E+00 | 1.85E-01 | 1.14E+00 |
| ENSG000000 | VPS13D | 8.08E-01 | 9.33E-01 | 9.85E-01 | -2.15E-02 | -1.01E+00 |
| ENSG000001 | ZNG1E | 8.08E-01 | 9.33E-01 | 9.86E-01 | -2.05E-02 | -1.01E+00 |
| ENSG000002 | LINC01088 | 8.09E-01 | ? | 1.04E+00 | 6.26E-02 | 1.04E+00 |
| ENSG000001 | BEND7 | 8.09E-01 | 9.33E-01 | 9.84E-01 | -2.30E-02 | -1.02E+00 |
| ENSG000002 | GLI4 | 8.09E-01 | 9.33E-01 | 9.80E-01 | -2.89E-02 | -1.02E+00 |
| ENSG000001 | CAMK1 | 8.09E-01 | ? | 9.53E-01 | -6.98E-02 | -1.05E+00 |
| ENSG000001 | ACSL3 | 8.09E-01 | 9.34E-01 | 9.92E-01 | -1.14E-02 | -1.01E+00 |
| ENSG000001 | XKR8 | 8.09E-01 | 9.34E-01 | 9.82E-01 | -2.60E-02 | -1.02E+00 |
| ENSG000002 | --- | 8.09E-01 | 9.34E-01 | 9.70E-01 | -4.37E-02 | -1.03E+00 |
| ENSG000001 | MRPL17 | 8.09E-01 | 9.34E-01 | 9.85E-01 | -2.13E-02 | -1.01E+00 |
| ENSG000001 | NOL4L | 8.09E-01 | 9.34E-01 | 1.01E+00 | 1.46E-02 | 1.01E+00 |
| ENSG000002 | HMBS | 8.09E-01 | 9.34E-01 | 1.02E+00 | 2.21E-02 | 1.02E+00 |
| ENSG000002 | SEPTIN7-DT | 8.09E-01 | 9.34E-01 | 9.53E-01 | -6.98E-02 | -1.05E+00 |
| ENSG000000 | ADGRF5 | 8.09E-01 | 9.34E-01 | 9.65E-01 | -5.17E-02 | -1.04E+00 |
| ENSG000000 | NUCB2 | 8.09E-01 | 9.34E-01 | 1.01E+00 | 1.84E-02 | 1.01E+00 |
| ENSG000002 | --- | 8.09E-01 | ? | 1.06E+00 | 8.59E-02 | 1.06E+00 |
| ENSG000001 | SMOC1 | 8.09E-01 | 9.34E-01 | 1.11E+00 | 1.45E-01 | 1.11E+00 |
| ENSG000001 | TLE4 | 8.10E-01 | 9.34E-01 | 9.88E-01 | -1.70E-02 | -1.01E+00 |
| ENSG000000 | CEACAM6 | 8.10E-01 | 9.34E-01 | 9.79E-01 | -3.13E-02 | -1.02E+00 |
| ENSG000002 | RN7SL832P | 8.10E-01 | ? | 9.40E-01 | -8.90E-02 | -1.06E+00 |
| ENSG000001 | NPLOC4 | 8.10E-01 | 9.34E-01 | 1.01E+00 | 1.11E-02 | 1.01E+00 |
| ENSG000001 | RNF146 | 8.10E-01 | 9.34E-01 | 9.92E-01 | -1.17E-02 | -1.01E+00 |
| ENSG000001 | FAAP24 | 8.10E-01 | 9.34E-01 | 1.02E+00 | 3.52E-02 | 1.02E+00 |
| ENSG000002 | --- | 8.10E-01 | 9.34E-01 | 9.66E-01 | -4.99E-02 | -1.04E+00 |
| ENSG000001 | RNF113A | 8.10E-01 | 9.34E-01 | 1.01E+00 | 2.04E-02 | 1.01E+00 |
| ENSG000001 | PBX3 | 8.10E-01 | 9.34E-01 | 9.87E-01 | -1.86E-02 | -1.01E+00 |
| ENSG000001 | TPCN2 | 8.10E-01 | 9.34E-01 | 9.84E-01 | -2.28E-02 | -1.02E+00 |
| ENSG000001 | WASHC5 | 8.10E-01 | 9.34E-01 | 9.93E-01 | -1.01E-02 | -1.01E+00 |
| ENSG000001 | SERPINB13 | 8.10E-01 | 9.34E-01 | 9.77E-01 | -3.42E-02 | -1.02E+00 |
| ENSG000002 | --- | 8.10E-01 | ? | 1.06E+00 | 8.74E-02 | 1.06E+00 |
| ENSG000001 | CNOT6 | 8.10E-01 | 9.34E-01 | 1.01E+00 | 8.80E-03 | 1.01E+00 |

| | | | | | | |
|---|---|---|---|---|---|---|
| ENSG00000C | GAL | 8.11E-01 | 9.34E-01 | 1.02E+00 | 3.01E-02 | 1.02E+00 |
| ENSG000002 | NAP1L6P | 8.11E-01 | ? | 9.16E-01 | -1.26E-01 | -1.09E+00 |
| ENSG000001 | TMF1 | 8.11E-01 | 9.34E-01 | 1.01E+00 | 1.46E-02 | 1.01E+00 |
| ENSG000002 | --- | 8.11E-01 | ? | 1.09E+00 | 1.24E-01 | 1.09E+00 |
| ENSG000002 | LINC01670 | 8.11E-01 | ? | 1.07E+00 | 1.00E-01 | 1.07E+00 |
| ENSG000002 | --- | 8.11E-01 | 9.34E-01 | 9.73E-01 | -3.97E-02 | -1.03E+00 |
| ENSG000002 | FBXO38-DT | 8.11E-01 | 9.34E-01 | 9.71E-01 | -4.24E-02 | -1.03E+00 |
| ENSG000001 | P2RY2 | 8.11E-01 | 9.34E-01 | 9.93E-01 | -1.08E-02 | -1.01E+00 |
| ENSG000002 | MPRIPP1 | 8.11E-01 | ? | 9.35E-01 | -9.76E-02 | -1.07E+00 |
| ENSG000001 | MCM5 | 8.11E-01 | 9.34E-01 | 1.02E+00 | 3.55E-02 | 1.02E+00 |
| ENSG000001 | NIPAL1 | 8.11E-01 | 9.34E-01 | 1.01E+00 | 1.35E-02 | 1.01E+00 |
| ENSG000001 | SPARC | 8.11E-01 | 9.34E-01 | 1.01E+00 | 1.61E-02 | 1.01E+00 |
| ENSG000002 | DUX4L50 | 8.11E-01 | 9.34E-01 | 1.03E+00 | 4.00E-02 | 1.03E+00 |
| ENSG000001 | PLEKHA6 | 8.11E-01 | 9.34E-01 | 9.73E-01 | -3.96E-02 | -1.03E+00 |
| ENSG000001 | COL13A1 | 8.11E-01 | 9.34E-01 | 1.03E+00 | 4.07E-02 | 1.03E+00 |
| ENSG000001 | VCPIP1 | 8.11E-01 | 9.34E-01 | 9.91E-01 | -1.27E-02 | -1.01E+00 |
| ENSG000001 | OR10A3 | 8.11E-01 | ? | 9.41E-01 | -8.71E-02 | -1.06E+00 |
| ENSG000002 | ERVW-1 | 8.11E-01 | ? | 1.10E+00 | 1.44E-01 | 1.10E+00 |
| ENSG000002 | ZGLP1 | 8.11E-01 | 9.34E-01 | 9.69E-01 | -4.55E-02 | -1.03E+00 |
| ENSG000001 | ROM1 | 8.11E-01 | 9.34E-01 | 1.04E+00 | 5.85E-02 | 1.04E+00 |
| ENSG000002 | TMEM217B | 8.11E-01 | ? | 1.16E+00 | 2.19E-01 | 1.16E+00 |
| ENSG000001 | RIMS4 | 8.11E-01 | ? | 9.01E-01 | -1.50E-01 | -1.11E+00 |
| ENSG000001 | TMEM25 | 8.11E-01 | 9.34E-01 | 1.01E+00 | 1.85E-02 | 1.01E+00 |
| ENSG000001 | RAB26 | 8.11E-01 | ? | 9.31E-01 | -1.03E-01 | -1.07E+00 |
| ENSG000001 | SLC16A3 | 8.11E-01 | 9.34E-01 | 9.82E-01 | -2.60E-02 | -1.02E+00 |
| ENSG00000C | MKS1 | 8.12E-01 | 9.34E-01 | 1.02E+00 | 2.32E-02 | 1.02E+00 |
| ENSG000001 | MRPL48 | 8.12E-01 | 9.34E-01 | 9.89E-01 | -1.62E-02 | -1.01E+00 |
| ENSG000002 | --- | 8.12E-01 | ? | 1.07E+00 | 9.45E-02 | 1.07E+00 |
| ENSG000002 | HSPA8P5 | 8.12E-01 | ? | 1.09E+00 | 1.20E-01 | 1.09E+00 |
| ENSG000001 | TTC7B | 8.12E-01 | 9.34E-01 | 1.01E+00 | 1.57E-02 | 1.01E+00 |
| ENSG000001 | ARMH1 | 8.12E-01 | ? | 9.11E-01 | -1.35E-01 | -1.10E+00 |
| ENSG000001 | TAFAZZIN | 8.12E-01 | 9.34E-01 | 1.02E+00 | 2.68E-02 | 1.02E+00 |
| ENSG000002 | DCAF8-DT | 8.12E-01 | ? | 9.33E-01 | -1.01E-01 | -1.07E+00 |
| ENSG000002 | PCDHGB1 | 8.12E-01 | ? | 1.09E+00 | 1.23E-01 | 1.09E+00 |
| ENSG000001 | MAGEF1 | 8.12E-01 | 9.34E-01 | 1.01E+00 | 1.85E-02 | 1.01E+00 |
| ENSG000001 | ZFP1 | 8.12E-01 | 9.34E-01 | 9.89E-01 | -1.65E-02 | -1.01E+00 |
| ENSG000002 | --- | 8.12E-01 | ? | 1.09E+00 | 1.22E-01 | 1.09E+00 |
| ENSG000002 | --- | 8.12E-01 | 9.34E-01 | 9.63E-01 | -5.38E-02 | -1.04E+00 |
| ENSG000002 | CAPN14 | 8.12E-01 | 9.34E-01 | 9.72E-01 | -4.11E-02 | -1.03E+00 |
| ENSG000001 | GATM | 8.12E-01 | ? | 1.04E+00 | 6.27E-02 | 1.04E+00 |
| ENSG00000C | SLC6A16 | 8.12E-01 | ? | 1.08E+00 | 1.06E-01 | 1.08E+00 |
| ENSG000002 | PSME2P1 | 8.12E-01 | ? | 9.47E-01 | -7.81E-02 | -1.06E+00 |
| ENSG000001 | ITGAV | 8.12E-01 | 9.34E-01 | 1.01E+00 | 9.97E-03 | 1.01E+00 |
| ENSG000002 | SNORD3B-1 | 8.12E-01 | ? | 8.75E-01 | -1.92E-01 | -1.14E+00 |
| ENSG000001 | DOCK10 | 8.13E-01 | ? | 1.06E+00 | 8.01E-02 | 1.06E+00 |

| | | | | | | |
|---|---|---|---|---|---|---|
| ENSG000001 | ETNK2 | 8.13E-01 | 9.34E-01 | 1.02E+00 | 2.62E-02 | 1.02E+00 |
| ENSG000001 | UCP2 | 8.13E-01 | 9.34E-01 | 1.02E+00 | 3.14E-02 | 1.02E+00 |
| ENSG00000C | ULK2 | 8.13E-01 | 9.34E-01 | 9.91E-01 | -1.35E-02 | -1.01E+00 |
| ENSG00000C | EDC4 | 8.13E-01 | 9.34E-01 | 9.90E-01 | -1.43E-02 | -1.01E+00 |
| ENSG000002 | PPP4R3B-DT | 8.13E-01 | ? | 1.07E+00 | 9.56E-02 | 1.07E+00 |
| ENSG000002 | --- | 8.13E-01 | ? | 8.77E-01 | -1.89E-01 | -1.14E+00 |
| ENSG000002 | RASSF10-DT | 8.13E-01 | ? | 1.05E+00 | 7.05E-02 | 1.05E+00 |
| ENSG000002 | QKILA | 8.13E-01 | ? | 1.08E+00 | 1.17E-01 | 1.08E+00 |
| ENSG000002 | --- | 8.13E-01 | ? | 9.22E-01 | -1.17E-01 | -1.08E+00 |
| ENSG000002 | PABPC4L | 8.13E-01 | ? | 9.41E-01 | -8.75E-02 | -1.06E+00 |
| ENSG000002 | --- | 8.14E-01 | ? | 8.78E-01 | -1.88E-01 | -1.14E+00 |
| ENSG000002 | --- | 8.14E-01 | ? | 9.39E-01 | -9.06E-02 | -1.06E+00 |
| ENSG000001 | KCTD13 | 8.14E-01 | 9.35E-01 | 9.85E-01 | -2.24E-02 | -1.02E+00 |
| ENSG000001 | DEDD2 | 8.14E-01 | 9.35E-01 | 1.01E+00 | 1.30E-02 | 1.01E+00 |
| ENSG000002 | --- | 8.14E-01 | ? | 1.06E+00 | 8.31E-02 | 1.06E+00 |
| ENSG000001 | DLGAP1 | 8.14E-01 | ? | 1.13E+00 | 1.78E-01 | 1.13E+00 |
| ENSG000002 | --- | 8.14E-01 | ? | 9.07E-01 | -1.41E-01 | -1.10E+00 |
| ENSG000001 | MITD1 | 8.14E-01 | 9.36E-01 | 9.87E-01 | -1.92E-02 | -1.01E+00 |
| ENSG000002 | --- | 8.14E-01 | ? | 1.09E+00 | 1.18E-01 | 1.09E+00 |
| ENSG000002 | --- | 8.14E-01 | 9.36E-01 | 1.04E+00 | 5.04E-02 | 1.04E+00 |
| ENSG00000C | TECR | 8.14E-01 | 9.36E-01 | 9.89E-01 | -1.56E-02 | -1.01E+00 |
| ENSG000002 | --- | 8.14E-01 | ? | 9.43E-01 | -8.53E-02 | -1.06E+00 |
| ENSG00000C | CRY1 | 8.14E-01 | 9.36E-01 | 9.91E-01 | -1.32E-02 | -1.01E+00 |
| ENSG000001 | TMED4 | 8.14E-01 | 9.36E-01 | 1.01E+00 | 1.31E-02 | 1.01E+00 |
| ENSG000002 | LINC00622 | 8.15E-01 | ? | 9.47E-01 | -7.82E-02 | -1.06E+00 |
| ENSG000002 | BCKDHA | 8.15E-01 | 9.36E-01 | 9.86E-01 | -1.98E-02 | -1.01E+00 |
| ENSG000001 | BAIAP2 | 8.15E-01 | 9.36E-01 | 1.01E+00 | 1.90E-02 | 1.01E+00 |
| ENSG000002 | --- | 8.15E-01 | ? | 9.26E-01 | -1.11E-01 | -1.08E+00 |
| ENSG000001 | SNRPA1P1 | 8.15E-01 | ? | 9.22E-01 | -1.17E-01 | -1.08E+00 |
| ENSG00000C | MCM2 | 8.15E-01 | 9.36E-01 | 1.02E+00 | 2.41E-02 | 1.02E+00 |
| ENSG000001 | AOC3 | 8.15E-01 | 9.36E-01 | 9.63E-01 | -5.50E-02 | -1.04E+00 |
| ENSG000002 | NLRP2B | 8.15E-01 | ? | 1.12E+00 | 1.63E-01 | 1.12E+00 |
| ENSG000001 | GMEB2 | 8.15E-01 | 9.36E-01 | 9.87E-01 | -1.94E-02 | -1.01E+00 |
| ENSG000002 | RHOQP3 | 8.15E-01 | ? | 1.08E+00 | 1.13E-01 | 1.08E+00 |
| ENSG000002 | DDX52 | 8.15E-01 | 9.36E-01 | 9.92E-01 | -1.10E-02 | -1.01E+00 |
| ENSG000001 | ZNF692 | 8.15E-01 | 9.36E-01 | 1.02E+00 | 2.60E-02 | 1.02E+00 |
| ENSG000001 | MYADM | 8.15E-01 | 9.36E-01 | 1.01E+00 | 1.02E-02 | 1.01E+00 |
| ENSG000001 | ERFE | 8.15E-01 | ? | 9.36E-01 | -9.58E-02 | -1.07E+00 |
| ENSG000002 | CRYZL2P-SEC | 8.15E-01 | 9.36E-01 | 1.04E+00 | 6.09E-02 | 1.04E+00 |
| ENSG000001 | STARD3 | 8.15E-01 | 9.36E-01 | 1.01E+00 | 1.74E-02 | 1.01E+00 |
| ENSG000001 | PDZK1 | 8.16E-01 | 9.36E-01 | 9.72E-01 | -4.15E-02 | -1.03E+00 |
| ENSG000002 | TCEA1P4 | 8.16E-01 | ? | 9.42E-01 | -8.68E-02 | -1.06E+00 |
| ENSG000001 | NEK11 | 8.16E-01 | 9.36E-01 | 9.89E-01 | -1.66E-02 | -1.01E+00 |
| ENSG000001 | MED23 | 8.16E-01 | 9.36E-01 | 1.01E+00 | 1.08E-02 | 1.01E+00 |
| ENSG000001 | AOC2 | 8.16E-01 | 9.36E-01 | 1.02E+00 | 2.89E-02 | 1.02E+00 |

| | | | | | | |
|---|---|---|---|---|---|---|
| ENSG000001 | GLUD2 | 8.16E-01 | ? | 1.04E+00 | 5.18E-02 | 1.04E+00 |
| ENSG000001 | METTL27 | 8.16E-01 | 9.36E-01 | 9.63E-01 | -5.50E-02 | -1.04E+00 |
| ENSG00000C | DSP | 8.16E-01 | 9.36E-01 | 1.01E+00 | 1.60E-02 | 1.01E+00 |
| ENSG000001 | GINS3 | 8.16E-01 | 9.36E-01 | 1.02E+00 | 2.22E-02 | 1.02E+00 |
| ENSG00000C | EPN1 | 8.16E-01 | 9.36E-01 | 9.87E-01 | -1.86E-02 | -1.01E+00 |
| ENSG000001 | EVPL | 8.16E-01 | 9.36E-01 | 9.90E-01 | -1.43E-02 | -1.01E+00 |
| ENSG000001 | TBK1 | 8.16E-01 | 9.36E-01 | 9.92E-01 | -1.22E-02 | -1.01E+00 |
| ENSG000001 | ABCA6 | 8.16E-01 | ? | 1.06E+00 | 8.35E-02 | 1.06E+00 |
| ENSG000001 | ZNF91 | 8.16E-01 | 9.36E-01 | 9.86E-01 | -2.02E-02 | -1.01E+00 |
| ENSG000002 | RAB30-DT | 8.16E-01 | 9.36E-01 | 1.02E+00 | 2.72E-02 | 1.02E+00 |
| ENSG000002 | SLC25A5-AS1 | 8.16E-01 | ? | 1.05E+00 | 6.76E-02 | 1.05E+00 |
| ENSG000002 | LINC01089 | 8.16E-01 | 9.36E-01 | 1.03E+00 | 3.74E-02 | 1.03E+00 |
| ENSG000001 | SNORC | 8.16E-01 | 9.36E-01 | 1.04E+00 | 5.65E-02 | 1.04E+00 |
| ENSG000002 | --- | 8.16E-01 | ? | 1.09E+00 | 1.30E-01 | 1.09E+00 |
| ENSG000001 | NDUFAF1 | 8.16E-01 | 9.36E-01 | 9.90E-01 | -1.38E-02 | -1.01E+00 |
| ENSG000001 | CES3 | 8.16E-01 | 9.36E-01 | 9.61E-01 | -5.73E-02 | -1.04E+00 |
| ENSG000001 | ZFP69 | 8.16E-01 | 9.36E-01 | 9.78E-01 | -3.22E-02 | -1.02E+00 |
| ENSG000001 | CD226 | 8.17E-01 | 9.36E-01 | 1.02E+00 | 3.35E-02 | 1.02E+00 |
| ENSG000001 | AMTN | 8.17E-01 | ? | 1.07E+00 | 1.04E-01 | 1.07E+00 |
| ENSG000002 | --- | 8.17E-01 | ? | 8.28E-01 | -2.72E-01 | -1.21E+00 |
| ENSG000001 | MCAT | 8.17E-01 | 9.36E-01 | 1.01E+00 | 1.97E-02 | 1.01E+00 |
| ENSG000001 | SLC25A27 | 8.17E-01 | 9.36E-01 | 1.03E+00 | 3.71E-02 | 1.03E+00 |
| ENSG000001 | MEF2D | 8.17E-01 | 9.36E-01 | 9.89E-01 | -1.63E-02 | -1.01E+00 |
| ENSG00000C | FGFR1 | 8.17E-01 | 9.37E-01 | 1.01E+00 | 1.99E-02 | 1.01E+00 |
| ENSG000001 | SLC12A6 | 8.18E-01 | 9.37E-01 | 1.01E+00 | 1.21E-02 | 1.01E+00 |
| ENSG000002 | LINC00654 | 8.18E-01 | ? | 9.34E-01 | -9.85E-02 | -1.07E+00 |
| ENSG000001 | GLA | 8.18E-01 | 9.37E-01 | 1.01E+00 | 1.29E-02 | 1.01E+00 |
| ENSG00000C | BID | 8.18E-01 | 9.37E-01 | 1.01E+00 | 1.53E-02 | 1.01E+00 |
| ENSG000002 | LRRC8C-DT | 8.18E-01 | 9.37E-01 | 9.65E-01 | -5.20E-02 | -1.04E+00 |
| ENSG000001 | FOCAD | 8.18E-01 | 9.37E-01 | 1.01E+00 | 1.37E-02 | 1.01E+00 |
| ENSG000001 | DNAH17 | 8.18E-01 | ? | 1.05E+00 | 6.99E-02 | 1.05E+00 |
| ENSG000001 | SPATA5 | 8.18E-01 | 9.37E-01 | 9.83E-01 | -2.44E-02 | -1.02E+00 |
| ENSG000001 | ABHD13 | 8.18E-01 | 9.37E-01 | 9.91E-01 | -1.24E-02 | -1.01E+00 |
| ENSG000002 | CHASERR | 8.18E-01 | 9.37E-01 | 1.01E+00 | 1.80E-02 | 1.01E+00 |
| ENSG000001 | SPATA9 | 8.18E-01 | ? | 1.09E+00 | 1.29E-01 | 1.09E+00 |
| ENSG000001 | MCM7 | 8.18E-01 | 9.37E-01 | 9.86E-01 | -1.97E-02 | -1.01E+00 |
| ENSG000001 | EIF1B | 8.18E-01 | 9.37E-01 | 9.91E-01 | -1.26E-02 | -1.01E+00 |
| ENSG000002 | --- | 8.18E-01 | 9.37E-01 | 1.21E+00 | 2.74E-01 | 1.21E+00 |
| ENSG000001 | UBE2L3 | 8.18E-01 | 9.37E-01 | 1.01E+00 | 9.49E-03 | 1.01E+00 |
| ENSG000001 | TCAF2 | 8.18E-01 | 9.37E-01 | 1.01E+00 | 1.66E-02 | 1.01E+00 |
| ENSG00000C | PACC1 | 8.19E-01 | 9.37E-01 | 9.86E-01 | -2.00E-02 | -1.01E+00 |
| ENSG000001 | KRI1 | 8.19E-01 | 9.37E-01 | 1.01E+00 | 9.98E-03 | 1.01E+00 |
| ENSG000001 | NOM1 | 8.19E-01 | 9.37E-01 | 1.01E+00 | 1.12E-02 | 1.01E+00 |
| ENSG000002 | HERC2P9 | 8.19E-01 | ? | 9.41E-01 | -8.72E-02 | -1.06E+00 |
| ENSG000002 | CPT1B | 8.19E-01 | 9.37E-01 | 9.72E-01 | -4.11E-02 | -1.03E+00 |

| | | | | | | |
|---|---|---|---|---|---|---|
| ENSG000001 | PRDM15 | 8.19E-01 | 9.37E-01 | 9.91E-01 | -1.25E-02 | -1.01E+00 |
| ENSG000001 | H4C11 | 8.19E-01 | ? | 9.41E-01 | -8.79E-02 | -1.06E+00 |
| ENSG000002 | --- | 8.19E-01 | ? | 1.09E+00 | 1.29E-01 | 1.09E+00 |
| ENSG00000C | DHX29 | 8.19E-01 | 9.37E-01 | 9.94E-01 | -8.05E-03 | -1.01E+00 |
| ENSG00000C | KRT31 | 8.19E-01 | 9.37E-01 | 9.73E-01 | -3.99E-02 | -1.03E+00 |
| ENSG00000C | CTTNBP2 | 8.19E-01 | 9.37E-01 | 9.76E-01 | -3.47E-02 | -1.02E+00 |
| ENSG000001 | DHX34 | 8.19E-01 | 9.37E-01 | 9.88E-01 | -1.68E-02 | -1.01E+00 |
| ENSG000001 | GHRL | 8.19E-01 | ? | 1.13E+00 | 1.76E-01 | 1.13E+00 |
| ENSG000002 | RBM25-AS1 | 8.19E-01 | ? | 9.21E-01 | -1.19E-01 | -1.09E+00 |
| ENSG000002 | ARF4-AS1 | 8.19E-01 | ? | 9.28E-01 | -1.08E-01 | -1.08E+00 |
| ENSG000001 | WDR27 | 8.19E-01 | 9.37E-01 | 9.88E-01 | -1.67E-02 | -1.01E+00 |
| ENSG000001 | METAP1D | 8.19E-01 | 9.37E-01 | 1.02E+00 | 2.18E-02 | 1.02E+00 |
| ENSG000001 | NPIPB6 | 8.19E-01 | ? | 1.07E+00 | 9.95E-02 | 1.07E+00 |
| ENSG000001 | TMEM184A | 8.20E-01 | 9.37E-01 | 1.01E+00 | 1.99E-02 | 1.01E+00 |
| ENSG000001 | ZNF248 | 8.20E-01 | 9.37E-01 | 9.86E-01 | -2.00E-02 | -1.01E+00 |
| ENSG000001 | NDUFB9 | 8.20E-01 | 9.37E-01 | 9.91E-01 | -1.28E-02 | -1.01E+00 |
| ENSG000002 | PCDHGB7 | 8.20E-01 | 9.37E-01 | 1.02E+00 | 2.54E-02 | 1.02E+00 |
| ENSG000002 | USP2-AS1 | 8.20E-01 | ? | 9.46E-01 | -7.99E-02 | -1.06E+00 |
| ENSG000001 | DMD | 8.20E-01 | 9.37E-01 | 1.01E+00 | 1.49E-02 | 1.01E+00 |
| ENSG000002 | LINC01409 | 8.20E-01 | ? | 9.53E-01 | -7.00E-02 | -1.05E+00 |
| ENSG000001 | GNPTAB | 8.20E-01 | 9.37E-01 | 1.01E+00 | 1.67E-02 | 1.01E+00 |
| ENSG000001 | ACIN1 | 8.20E-01 | 9.37E-01 | 9.92E-01 | -1.10E-02 | -1.01E+00 |
| ENSG000002 | RPS3AP6 | 8.20E-01 | 9.37E-01 | 9.86E-01 | -2.05E-02 | -1.01E+00 |
| ENSG000002 | HLA-G | 8.20E-01 | 9.37E-01 | 9.76E-01 | -3.57E-02 | -1.03E+00 |
| ENSG000001 | DCK | 8.20E-01 | 9.37E-01 | 1.01E+00 | 1.69E-02 | 1.01E+00 |
| ENSG00000C | RABGAP1 | 8.20E-01 | 9.37E-01 | 1.01E+00 | 9.85E-03 | 1.01E+00 |
| ENSG000002 | SBNO1-AS1 | 8.20E-01 | 9.37E-01 | 9.57E-01 | -6.31E-02 | -1.04E+00 |
| ENSG000002 | SERTAD3-AS | 8.20E-01 | ? | 1.06E+00 | 8.91E-02 | 1.06E+00 |
| ENSG000001 | YDJC | 8.21E-01 | 9.37E-01 | 1.02E+00 | 3.12E-02 | 1.02E+00 |
| ENSG000001 | KLRD1 | 8.21E-01 | ? | 1.10E+00 | 1.37E-01 | 1.10E+00 |
| ENSG000002 | --- | 8.21E-01 | 9.37E-01 | 9.78E-01 | -3.22E-02 | -1.02E+00 |
| ENSG000001 | ABHD17A | 8.21E-01 | 9.37E-01 | 1.02E+00 | 2.27E-02 | 1.02E+00 |
| ENSG000001 | VMA21 | 8.21E-01 | 9.37E-01 | 9.93E-01 | -1.03E-02 | -1.01E+00 |
| ENSG00000C | ACSL4 | 8.21E-01 | 9.37E-01 | 1.01E+00 | 1.07E-02 | 1.01E+00 |
| ENSG000002 | ACAD11 | 8.21E-01 | 9.37E-01 | 9.89E-01 | -1.64E-02 | -1.01E+00 |
| ENSG00000C | GGCT | 8.21E-01 | 9.37E-01 | 1.01E+00 | 1.47E-02 | 1.01E+00 |
| ENSG000002 | --- | 8.21E-01 | 9.37E-01 | 1.04E+00 | 5.30E-02 | 1.04E+00 |
| ENSG000002 | --- | 8.21E-01 | ? | 1.09E+00 | 1.26E-01 | 1.09E+00 |
| ENSG000001 | MAML3 | 8.21E-01 | 9.37E-01 | 1.02E+00 | 3.08E-02 | 1.02E+00 |
| ENSG000002 | --- | 8.21E-01 | 9.37E-01 | 1.03E+00 | 4.94E-02 | 1.03E+00 |
| ENSG000001 | HAGHL | 8.21E-01 | 9.37E-01 | 9.76E-01 | -3.48E-02 | -1.02E+00 |
| ENSG000002 | EEF1A1P6 | 8.21E-01 | 9.37E-01 | 9.63E-01 | -5.42E-02 | -1.04E+00 |
| ENSG000002 | --- | 8.21E-01 | ? | 1.07E+00 | 1.00E-01 | 1.07E+00 |
| ENSG000001 | UBE2L6 | 8.21E-01 | 9.37E-01 | 9.83E-01 | -2.49E-02 | -1.02E+00 |
| ENSG000002 | --- | 8.21E-01 | ? | 9.30E-01 | -1.05E-01 | -1.08E+00 |

| | | | | | | |
|---|---|---|---|---|---|---|
| ENSG000001 | THAP9 | 8.21E-01 | 9.37E-01 | 1.02E+00 | 2.86E-02 | 1.02E+00 |
| ENSG000002 | --- | 8.21E-01 | ? | 1.19E+00 | 2.56E-01 | 1.19E+00 |
| ENSG000001 | LMLN | 8.21E-01 | 9.37E-01 | 1.01E+00 | 1.91E-02 | 1.01E+00 |
| ENSG000002 | PPP1R12A-A: | 8.21E-01 | 9.37E-01 | 1.03E+00 | 4.18E-02 | 1.03E+00 |
| ENSG000002 | DNAJC25-GN | 8.22E-01 | 9.38E-01 | 9.83E-01 | -2.55E-02 | -1.02E+00 |
| ENSG000001 | ARFIP2 | 8.22E-01 | 9.38E-01 | 9.91E-01 | -1.31E-02 | -1.01E+00 |
| ENSG000001 | NQO2 | 8.22E-01 | 9.38E-01 | 1.02E+00 | 3.54E-02 | 1.02E+00 |
| ENSG000002 | --- | 8.22E-01 | ? | 9.55E-01 | -6.67E-02 | -1.05E+00 |
| ENSG000002 | DHRS4-AS1 | 8.22E-01 | 9.38E-01 | 1.01E+00 | 1.52E-02 | 1.01E+00 |
| ENSG000001 | ADPRHL1 | 8.22E-01 | 9.38E-01 | 1.03E+00 | 4.44E-02 | 1.03E+00 |
| ENSG000002 | --- | 8.22E-01 | ? | 1.09E+00 | 1.20E-01 | 1.09E+00 |
| ENSG000002 | --- | 8.22E-01 | ? | 9.20E-01 | -1.20E-01 | -1.09E+00 |
| ENSG000002 | FAM215B | 8.22E-01 | ? | 1.05E+00 | 6.68E-02 | 1.05E+00 |
| ENSG000001 | SEMA4B | 8.22E-01 | 9.38E-01 | 9.84E-01 | -2.36E-02 | -1.02E+00 |
| ENSG000002 | NOTCH2NLC | 8.22E-01 | 9.38E-01 | 1.02E+00 | 3.46E-02 | 1.02E+00 |
| ENSG000002 | DPP3-DT | 8.22E-01 | 9.38E-01 | 1.03E+00 | 4.65E-02 | 1.03E+00 |
| ENSG000001 | STK4 | 8.23E-01 | 9.38E-01 | 9.94E-01 | -8.11E-03 | -1.01E+00 |
| ENSG000001 | FZD9 | 8.23E-01 | ? | 9.05E-01 | -1.44E-01 | -1.10E+00 |
| ENSG000002 | KRT18P17 | 8.23E-01 | ? | 9.26E-01 | -1.11E-01 | -1.08E+00 |
| ENSG000002 | TARS1-DT | 8.23E-01 | ? | 9.65E-01 | -5.16E-02 | -1.04E+00 |
| ENSG000001 | ZNF667 | 8.23E-01 | 9.38E-01 | 1.01E+00 | 1.62E-02 | 1.01E+00 |
| ENSG000002 | --- | 8.23E-01 | ? | 1.04E+00 | 5.64E-02 | 1.04E+00 |
| ENSG000001 | AKIRIN1 | 8.23E-01 | 9.38E-01 | 9.95E-01 | -7.45E-03 | -1.01E+00 |
| ENSG000001 | C1QTNF2 | 8.23E-01 | ? | 9.42E-01 | -8.57E-02 | -1.06E+00 |
| ENSG000002 | PRMT5-DT | 8.23E-01 | ? | 9.49E-01 | -7.61E-02 | -1.05E+00 |
| ENSG000001 | SNAPC4 | 8.23E-01 | 9.38E-01 | 9.88E-01 | -1.80E-02 | -1.01E+00 |
| ENSG000001 | PGRMC2 | 8.23E-01 | 9.38E-01 | 9.93E-01 | -9.57E-03 | -1.01E+00 |
| ENSG000001 | KCTD2 | 8.23E-01 | 9.38E-01 | 1.01E+00 | 1.92E-02 | 1.01E+00 |
| ENSG000002 | NLRP3P1 | 8.23E-01 | 9.38E-01 | 1.04E+00 | 4.99E-02 | 1.04E+00 |
| ENSG000002 | RPS4XP22 | 8.23E-01 | ? | 9.24E-01 | -1.13E-01 | -1.08E+00 |
| ENSG000001 | DISC1 | 8.23E-01 | 9.38E-01 | 1.02E+00 | 2.67E-02 | 1.02E+00 |
| ENSG000001 | CMTR2 | 8.23E-01 | 9.38E-01 | 1.01E+00 | 9.42E-03 | 1.01E+00 |
| ENSG000002 | --- | 8.23E-01 | ? | 9.29E-01 | -1.06E-01 | -1.08E+00 |
| ENSG000001 | UXT | 8.23E-01 | 9.38E-01 | 9.87E-01 | -1.90E-02 | -1.01E+00 |
| ENSG000001 | FUT2 | 8.23E-01 | 9.38E-01 | 1.03E+00 | 3.64E-02 | 1.03E+00 |
| ENSG000002 | RNF39 | 8.24E-01 | 9.38E-01 | 9.83E-01 | -2.50E-02 | -1.02E+00 |
| ENSG000002 | --- | 8.24E-01 | ? | 1.04E+00 | 6.09E-02 | 1.04E+00 |
| ENSG00000C | SLC46A1 | 8.24E-01 | 9.38E-01 | 1.01E+00 | 1.46E-02 | 1.01E+00 |
| ENSG000002 | --- | 8.24E-01 | ? | 1.05E+00 | 7.65E-02 | 1.05E+00 |
| ENSG000001 | ST13 | 8.24E-01 | 9.38E-01 | 1.01E+00 | 1.06E-02 | 1.01E+00 |
| ENSG000001 | SSB | 8.24E-01 | 9.38E-01 | 9.90E-01 | -1.45E-02 | -1.01E+00 |
| ENSG000002 | --- | 8.24E-01 | 9.38E-01 | 1.04E+00 | 5.52E-02 | 1.04E+00 |
| ENSG000001 | INO80E | 8.24E-01 | 9.38E-01 | 1.01E+00 | 1.67E-02 | 1.01E+00 |
| ENSG000001 | ZDHHC14 | 8.24E-01 | 9.38E-01 | 1.01E+00 | 1.50E-02 | 1.01E+00 |
| ENSG000001 | CENPN | 8.24E-01 | 9.38E-01 | 1.01E+00 | 2.05E-02 | 1.01E+00 |

| | | | | | | |
|---|---|---|---|---|---|---|
| ENSG000002 | --- | 8.24E-01 | ? | 9.18E-01 | -1.23E-01 | -1.09E+00 |
| ENSG000002 | FAM169A-AS | 8.24E-01 | ? | 8.99E-01 | -1.53E-01 | -1.11E+00 |
| ENSG000001 | GSKIP | 8.24E-01 | 9.38E-01 | 9.93E-01 | -1.04E-02 | -1.01E+00 |
| ENSG000002 | --- | 8.24E-01 | ? | 1.07E+00 | 9.91E-02 | 1.07E+00 |
| ENSG000001 | NOC2L | 8.24E-01 | 9.38E-01 | 1.01E+00 | 1.18E-02 | 1.01E+00 |
| ENSG000001 | CDK2AP1 | 8.24E-01 | 9.39E-01 | 9.93E-01 | -9.50E-03 | -1.01E+00 |
| ENSG000001 | ZNF596 | 8.25E-01 | 9.39E-01 | 1.02E+00 | 2.67E-02 | 1.02E+00 |
| ENSG000002 | MTCO1P12 | 8.25E-01 | 9.39E-01 | 1.02E+00 | 3.54E-02 | 1.02E+00 |
| ENSG000002 | HLCS-AS1 | 8.25E-01 | ? | 9.45E-01 | -8.21E-02 | -1.06E+00 |
| ENSG000001 | GGN | 8.25E-01 | ? | 9.44E-01 | -8.31E-02 | -1.06E+00 |
| ENSG000002 | --- | 8.25E-01 | ? | 9.32E-01 | -1.01E-01 | -1.07E+00 |
| ENSG000002 | --- | 8.25E-01 | ? | 9.42E-01 | -8.64E-02 | -1.06E+00 |
| ENSG000001 | TUBB2A | 8.25E-01 | 9.39E-01 | 1.01E+00 | 1.34E-02 | 1.01E+00 |
| ENSG000002 | --- | 8.25E-01 | ? | 1.13E+00 | 1.81E-01 | 1.13E+00 |
| ENSG000001 | NAA30 | 8.25E-01 | 9.39E-01 | 1.01E+00 | 8.48E-03 | 1.01E+00 |
| ENSG000001 | CFAP58 | 8.25E-01 | 9.39E-01 | 1.02E+00 | 2.60E-02 | 1.02E+00 |
| ENSG000001 | DHX15 | 8.25E-01 | 9.39E-01 | 1.01E+00 | 8.44E-03 | 1.01E+00 |
| ENSG000001 | AGAP5 | 8.25E-01 | 9.39E-01 | 9.51E-01 | -7.27E-02 | -1.05E+00 |
| ENSG000001 | ATP5IF1 | 8.25E-01 | 9.39E-01 | 9.93E-01 | -9.79E-03 | -1.01E+00 |
| ENSG000000 | ZNF638 | 8.25E-01 | 9.39E-01 | 1.01E+00 | 1.21E-02 | 1.01E+00 |
| ENSG000001 | PID1 | 8.25E-01 | 9.39E-01 | 9.82E-01 | -2.62E-02 | -1.02E+00 |
| ENSG000001 | MDFIC | 8.25E-01 | 9.39E-01 | 9.91E-01 | -1.24E-02 | -1.01E+00 |
| ENSG000002 | SNHG15 | 8.25E-01 | 9.39E-01 | 1.01E+00 | 2.09E-02 | 1.01E+00 |
| ENSG000001 | --- | 8.26E-01 | ? | 1.04E+00 | 5.40E-02 | 1.04E+00 |
| ENSG000002 | TMX2P1 | 8.26E-01 | 9.39E-01 | 1.02E+00 | 2.61E-02 | 1.02E+00 |
| ENSG000001 | POLE2 | 8.26E-01 | ? | 9.66E-01 | -4.97E-02 | -1.04E+00 |
| ENSG000002 | --- | 8.26E-01 | ? | 9.06E-01 | -1.42E-01 | -1.10E+00 |
| ENSG000001 | OXR1 | 8.26E-01 | 9.39E-01 | 9.93E-01 | -9.93E-03 | -1.01E+00 |
| ENSG000002 | --- | 8.26E-01 | ? | 1.06E+00 | 8.78E-02 | 1.06E+00 |
| ENSG000002 | --- | 8.26E-01 | ? | 1.06E+00 | 8.15E-02 | 1.06E+00 |
| ENSG000000 | GCKR | 8.26E-01 | ? | 9.14E-01 | -1.30E-01 | -1.09E+00 |
| ENSG000001 | NUDT6 | 8.26E-01 | 9.39E-01 | 1.02E+00 | 3.23E-02 | 1.02E+00 |
| ENSG000002 | --- | 8.26E-01 | ? | 1.09E+00 | 1.26E-01 | 1.09E+00 |
| ENSG000001 | APLN | 8.26E-01 | 9.39E-01 | 1.02E+00 | 3.33E-02 | 1.02E+00 |
| ENSG000002 | TEX21P | 8.26E-01 | ? | 9.33E-01 | -9.94E-02 | -1.07E+00 |
| ENSG000002 | FRG1CP | 8.26E-01 | ? | 8.53E-01 | -2.30E-01 | -1.17E+00 |
| ENSG000001 | BLTP1 | 8.26E-01 | 9.39E-01 | 9.85E-01 | -2.12E-02 | -1.01E+00 |
| ENSG000001 | IFNK | 8.26E-01 | ? | 8.91E-01 | -1.67E-01 | -1.12E+00 |
| ENSG000002 | --- | 8.26E-01 | ? | 9.12E-01 | -1.33E-01 | -1.10E+00 |
| ENSG000002 | --- | 8.26E-01 | ? | 8.93E-01 | -1.63E-01 | -1.12E+00 |
| ENSG000002 | CDK13-DT | 8.26E-01 | ? | 1.08E+00 | 1.06E-01 | 1.08E+00 |
| ENSG000001 | ZNF292 | 8.26E-01 | 9.39E-01 | 9.86E-01 | -2.05E-02 | -1.01E+00 |
| ENSG000002 | RPS2P32 | 8.27E-01 | ? | 1.10E+00 | 1.36E-01 | 1.10E+00 |
| ENSG000001 | DCAF6 | 8.27E-01 | 9.40E-01 | 1.01E+00 | 1.06E-02 | 1.01E+00 |
| ENSG000001 | GTPBP3 | 8.27E-01 | 9.40E-01 | 1.02E+00 | 2.28E-02 | 1.02E+00 |

| | | | | | | | |
|---|---|---|---|---|---|---|---|
| ENSG000001 | LIN54 | 8.27E-01 | | 9.40E-01 | 9.87E-01 | -1.84E-02 | -1.01E+00 |
| ENSG000002 | STEAP1B-AS | 8.27E-01 | | 9.40E-01 | 9.64E-01 | -5.22E-02 | -1.04E+00 |
| ENSG000001 | COL3A1 | 8.27E-01 | ? | | 9.33E-01 | -9.98E-02 | -1.07E+00 |
| ENSG000002 | --- | 8.27E-01 | ? | | 1.03E+00 | 4.81E-02 | 1.03E+00 |
| ENSG000001 | SIRT7 | 8.27E-01 | | 9.40E-01 | 9.87E-01 | -1.87E-02 | -1.01E+00 |
| ENSG00000C | ST7L | 8.28E-01 | | 9.40E-01 | 9.90E-01 | -1.52E-02 | -1.01E+00 |
| ENSG000001 | RHOT1 | 8.28E-01 | | 9.40E-01 | 9.91E-01 | -1.28E-02 | -1.01E+00 |
| ENSG00000C | DELE1 | 8.28E-01 | | 9.40E-01 | 1.01E+00 | 1.31E-02 | 1.01E+00 |
| ENSG000001 | HYLS1 | 8.28E-01 | | 9.40E-01 | 9.81E-01 | -2.75E-02 | -1.02E+00 |
| ENSG000002 | LINC01431 | 8.28E-01 | ? | | 9.36E-01 | -9.55E-02 | -1.07E+00 |
| ENSG000002 | --- | 8.28E-01 | | 9.40E-01 | 9.61E-01 | -5.73E-02 | -1.04E+00 |
| ENSG000001 | SCAMP3 | 8.28E-01 | | 9.40E-01 | 1.01E+00 | 1.53E-02 | 1.01E+00 |
| ENSG000001 | PHOSPHO2 | 8.28E-01 | | 9.40E-01 | 9.71E-01 | -4.25E-02 | -1.03E+00 |
| ENSG000002 | TEX22 | 8.28E-01 | ? | | 8.92E-01 | -1.65E-01 | -1.12E+00 |
| ENSG000001 | UTRN | 8.28E-01 | | 9.40E-01 | 1.01E+00 | 1.70E-02 | 1.01E+00 |
| ENSG000002 | CTBP2P9 | 8.28E-01 | ? | | 9.42E-01 | -8.69E-02 | -1.06E+00 |
| ENSG000002 | KC6 | 8.28E-01 | ? | | 8.94E-01 | -1.62E-01 | -1.12E+00 |
| ENSG000002 | LINC00702 | 8.28E-01 | ? | | 1.07E+00 | 9.86E-02 | 1.07E+00 |
| ENSG000002 | PKD1P5 | 8.28E-01 | | 9.40E-01 | 1.04E+00 | 5.54E-02 | 1.04E+00 |
| ENSG000001 | CPNE7 | 8.28E-01 | | 9.40E-01 | 9.76E-01 | -3.55E-02 | -1.02E+00 |
| ENSG000001 | MRPL3 | 8.28E-01 | | 9.40E-01 | 9.94E-01 | -9.10E-03 | -1.01E+00 |
| ENSG000001 | ALG8 | 8.28E-01 | | 9.40E-01 | 1.01E+00 | 1.41E-02 | 1.01E+00 |
| ENSG000001 | ARV1 | 8.28E-01 | | 9.40E-01 | 9.89E-01 | -1.58E-02 | -1.01E+00 |
| ENSG000001 | PHC3 | 8.28E-01 | | 9.40E-01 | 9.88E-01 | -1.80E-02 | -1.01E+00 |
| ENSG000001 | KCNJ10 | 8.29E-01 | ? | | 9.17E-01 | -1.25E-01 | -1.09E+00 |
| ENSG00000C | IPCEF1 | 8.29E-01 | ? | | 1.10E+00 | 1.39E-01 | 1.10E+00 |
| ENSG000001 | MYO1E | 8.29E-01 | | 9.41E-01 | 1.01E+00 | 7.20E-03 | 1.01E+00 |
| ENSG000002 | PDC-AS1 | 8.29E-01 | ? | | 9.35E-01 | -9.67E-02 | -1.07E+00 |
| ENSG000001 | SRSF6 | 8.29E-01 | | 9.41E-01 | 1.01E+00 | 8.62E-03 | 1.01E+00 |
| ENSG000001 | SLC6A9 | 8.29E-01 | | 9.41E-01 | 1.03E+00 | 3.82E-02 | 1.03E+00 |
| ENSG000001 | S100A4 | 8.29E-01 | | 9.41E-01 | 9.76E-01 | -3.50E-02 | -1.02E+00 |
| ENSG000002 | MIR99AHG | 8.29E-01 | | 9.41E-01 | 9.69E-01 | -4.56E-02 | -1.03E+00 |
| ENSG000001 | THOC7 | 8.29E-01 | | 9.41E-01 | 9.92E-01 | -1.18E-02 | -1.01E+00 |
| ENSG000002 | --- | 8.29E-01 | ? | | 1.07E+00 | 1.04E-01 | 1.07E+00 |
| ENSG000001 | SYNGR1 | 8.29E-01 | ? | | 9.34E-01 | -9.79E-02 | -1.07E+00 |
| ENSG000001 | ENDOU | 8.29E-01 | ? | | 1.13E+00 | 1.71E-01 | 1.13E+00 |
| ENSG000002 | --- | 8.29E-01 | ? | | 1.07E+00 | 1.00E-01 | 1.07E+00 |
| ENSG000002 | --- | 8.29E-01 | ? | | 1.06E+00 | 7.81E-02 | 1.06E+00 |
| ENSG000002 | --- | 8.29E-01 | ? | | 1.07E+00 | 9.75E-02 | 1.07E+00 |
| ENSG000002 | CHMP1B | 8.29E-01 | | 9.41E-01 | 9.93E-01 | -9.99E-03 | -1.01E+00 |
| ENSG000001 | ZNF222 | 8.29E-01 | | 9.41E-01 | 1.02E+00 | 2.90E-02 | 1.02E+00 |
| ENSG000001 | AQP11 | 8.29E-01 | ? | | 9.42E-01 | -8.65E-02 | -1.06E+00 |
| ENSG000001 | SLC9B2 | 8.30E-01 | | 9.41E-01 | 9.94E-01 | -8.96E-03 | -1.01E+00 |
| ENSG000002 | --- | 8.30E-01 | ? | | 9.17E-01 | -1.25E-01 | -1.09E+00 |
| ENSG000001 | CMBL | 8.30E-01 | | 9.41E-01 | 1.01E+00 | 1.08E-02 | 1.01E+00 |

| | | | | | | |
|---|---|---|---|---|---|---|
| ENSG00000C | HMG20B | 8.30E-01 | 9.41E-01 | 9.90E-01 | -1.47E-02 | -1.01E+00 |
| ENSG000002 | --- | 8.30E-01 | ? | 9.28E-01 | -1.09E-01 | -1.08E+00 |
| ENSG000001 | IGFBP4 | 8.30E-01 | 9.41E-01 | 1.01E+00 | 1.90E-02 | 1.01E+00 |
| ENSG00000C | BCL7C | 8.30E-01 | 9.41E-01 | 1.01E+00 | 1.61E-02 | 1.01E+00 |
| ENSG000001 | WFDC5 | 8.30E-01 | ? | 8.94E-01 | -1.61E-01 | -1.12E+00 |
| ENSG000001 | EPS8L1 | 8.30E-01 | 9.41E-01 | 1.02E+00 | 2.23E-02 | 1.02E+00 |
| ENSG000002 | --- | 8.30E-01 | 9.41E-01 | 1.01E+00 | 1.23E-02 | 1.01E+00 |
| ENSG000002 | --- | 8.30E-01 | ? | 1.07E+00 | 1.02E-01 | 1.07E+00 |
| ENSG000002 | ERRFI1-DT | 8.30E-01 | ? | 9.27E-01 | -1.09E-01 | -1.08E+00 |
| ENSG000001 | ZNF763 | 8.30E-01 | 9.41E-01 | 1.02E+00 | 2.91E-02 | 1.02E+00 |
| ENSG000001 | RBM42 | 8.30E-01 | 9.41E-01 | 1.01E+00 | 2.07E-02 | 1.01E+00 |
| ENSG000001 | TTC31 | 8.30E-01 | 9.41E-01 | 9.91E-01 | -1.30E-02 | -1.01E+00 |
| ENSG000001 | NINJ2 | 8.30E-01 | ? | 1.09E+00 | 1.29E-01 | 1.09E+00 |
| ENSG000001 | CSTF2T | 8.31E-01 | 9.41E-01 | 9.93E-01 | -1.06E-02 | -1.01E+00 |
| ENSG000002 | LINC02236 | 8.31E-01 | ? | 9.49E-01 | -7.55E-02 | -1.05E+00 |
| ENSG000002 | ZSWIM7 | 8.31E-01 | 9.41E-01 | 1.01E+00 | 1.32E-02 | 1.01E+00 |
| ENSG000001 | C16orf86 | 8.31E-01 | ? | 9.17E-01 | -1.25E-01 | -1.09E+00 |
| ENSG000001 | TEP1 | 8.31E-01 | 9.41E-01 | 1.01E+00 | 1.67E-02 | 1.01E+00 |
| ENSG000002 | --- | 8.31E-01 | 9.41E-01 | 9.57E-01 | -6.37E-02 | -1.05E+00 |
| ENSG000002 | STAG3L5P | 8.31E-01 | 9.41E-01 | 9.82E-01 | -2.65E-02 | -1.02E+00 |
| ENSG000002 | --- | 8.31E-01 | ? | 9.10E-01 | -1.35E-01 | -1.10E+00 |
| ENSG000001 | PROX1 | 8.31E-01 | 9.42E-01 | 1.02E+00 | 3.02E-02 | 1.02E+00 |
| ENSG000002 | IMPDH1P5 | 8.31E-01 | ? | 1.07E+00 | 1.04E-01 | 1.07E+00 |
| ENSG000002 | HLA-DOB | 8.31E-01 | 9.42E-01 | 9.76E-01 | -3.48E-02 | -1.02E+00 |
| ENSG000002 | --- | 8.31E-01 | ? | 9.32E-01 | -1.01E-01 | -1.07E+00 |
| ENSG000001 | NUDT1 | 8.32E-01 | 9.42E-01 | 1.02E+00 | 2.90E-02 | 1.02E+00 |
| ENSG000002 | SNHG9 | 8.32E-01 | ? | 1.04E+00 | 5.91E-02 | 1.04E+00 |
| ENSG000001 | NTAQ1 | 8.32E-01 | 9.42E-01 | 9.86E-01 | -2.00E-02 | -1.01E+00 |
| ENSG000001 | SAMHD1 | 8.32E-01 | 9.42E-01 | 9.91E-01 | -1.31E-02 | -1.01E+00 |
| ENSG000001 | DAAM2 | 8.32E-01 | ? | 1.07E+00 | 9.11E-02 | 1.07E+00 |
| ENSG000002 | --- | 8.32E-01 | ? | 9.12E-01 | -1.34E-01 | -1.10E+00 |
| ENSG000002 | --- | 8.32E-01 | 9.42E-01 | 9.53E-01 | -6.95E-02 | -1.05E+00 |
| ENSG000001 | IARS1 | 8.32E-01 | 9.42E-01 | 9.93E-01 | -9.47E-03 | -1.01E+00 |
| ENSG000001 | LIMD1 | 8.32E-01 | 9.42E-01 | 1.01E+00 | 1.23E-02 | 1.01E+00 |
| ENSG000001 | MIIP | 8.32E-01 | 9.42E-01 | 9.81E-01 | -2.75E-02 | -1.02E+00 |
| ENSG00000C | DMRT3 | 8.32E-01 | ? | 1.13E+00 | 1.72E-01 | 1.13E+00 |
| ENSG000001 | PLAC8 | 8.33E-01 | ? | 1.07E+00 | 9.43E-02 | 1.07E+00 |
| ENSG000001 | SYN2 | 8.33E-01 | ? | 9.16E-01 | -1.26E-01 | -1.09E+00 |
| ENSG000001 | PSMA5 | 8.33E-01 | 9.43E-01 | 9.93E-01 | -9.42E-03 | -1.01E+00 |
| ENSG000002 | KRT17P3 | 8.33E-01 | ? | 8.98E-01 | -1.55E-01 | -1.11E+00 |
| ENSG000002 | DLG1-AS1 | 8.33E-01 | ? | 1.05E+00 | 6.54E-02 | 1.05E+00 |
| ENSG000001 | TSHZ2 | 8.33E-01 | ? | 1.04E+00 | 5.50E-02 | 1.04E+00 |
| ENSG000001 | DOCK11 | 8.33E-01 | 9.43E-01 | 1.03E+00 | 4.54E-02 | 1.03E+00 |
| ENSG000002 | LINC00921 | 8.33E-01 | ? | 9.51E-01 | -7.32E-02 | -1.05E+00 |
| ENSG000001 | SIGLEC16 | 8.33E-01 | ? | 9.28E-01 | -1.08E-01 | -1.08E+00 |

| | | | | | | |
|---|---|---|---|---|---|---|
| ENSG000002 | --- | 8.33E-01 | ? | 9.08E-01 | -1.39E-01 | -1.10E+00 |
| ENSG000001 | CDK5RAP3 | 8.33E-01 | 9.43E-01 | 1.01E+00 | 1.72E-02 | 1.01E+00 |
| ENSG000001 | HELLS | 8.33E-01 | 9.43E-01 | 9.79E-01 | -3.07E-02 | -1.02E+00 |
| ENSG000002 | TSBP1-AS1 | 8.33E-01 | ? | 9.45E-01 | -8.14E-02 | -1.06E+00 |
| ENSG00000C | TAF11 | 8.33E-01 | 9.43E-01 | 9.93E-01 | -1.02E-02 | -1.01E+00 |
| ENSG000002 | CKMT1A | 8.33E-01 | 9.43E-01 | 9.92E-01 | -1.14E-02 | -1.01E+00 |
| ENSG00000C | ZNF268 | 8.33E-01 | 9.43E-01 | 9.91E-01 | -1.28E-02 | -1.01E+00 |
| ENSG000002 | PTGES2-AS1 | 8.33E-01 | ? | 9.15E-01 | -1.29E-01 | -1.09E+00 |
| ENSG00000C | VSIG2 | 8.33E-01 | 9.43E-01 | 1.04E+00 | 5.52E-02 | 1.04E+00 |
| ENSG00000C | MRPL28 | 8.33E-01 | 9.43E-01 | 9.89E-01 | -1.66E-02 | -1.01E+00 |
| ENSG000001 | LRRC49 | 8.33E-01 | 9.43E-01 | 9.88E-01 | -1.75E-02 | -1.01E+00 |
| ENSG000001 | THSD7B | 8.34E-01 | ? | 1.07E+00 | 1.03E-01 | 1.07E+00 |
| ENSG000002 | --- | 8.34E-01 | ? | 9.08E-01 | -1.39E-01 | -1.10E+00 |
| ENSG000002 | LINC01239 | 8.34E-01 | 9.43E-01 | 9.68E-01 | -4.71E-02 | -1.03E+00 |
| ENSG000002 | TSPOAP1-AS | 8.34E-01 | 9.43E-01 | 1.03E+00 | 4.58E-02 | 1.03E+00 |
| ENSG000002 | TRD-AS1 | 8.34E-01 | ? | 1.07E+00 | 9.66E-02 | 1.07E+00 |
| ENSG000002 | CUX1 | 8.34E-01 | 9.43E-01 | 9.93E-01 | -1.02E-02 | -1.01E+00 |
| ENSG000001 | MOSMO | 8.34E-01 | 9.43E-01 | 1.01E+00 | 1.76E-02 | 1.01E+00 |
| ENSG000002 | --- | 8.34E-01 | 9.43E-01 | 9.71E-01 | -4.21E-02 | -1.03E+00 |
| ENSG000001 | GRIP1 | 8.34E-01 | 9.43E-01 | 9.91E-01 | -1.34E-02 | -1.01E+00 |
| ENSG000002 | --- | 8.34E-01 | ? | 8.95E-01 | -1.59E-01 | -1.12E+00 |
| ENSG000001 | PRR18 | 8.34E-01 | ? | 1.09E+00 | 1.18E-01 | 1.09E+00 |
| ENSG000002 | CCDC18-AS1 | 8.34E-01 | 9.43E-01 | 1.02E+00 | 2.96E-02 | 1.02E+00 |
| ENSG000001 | C3AR1 | 8.34E-01 | ? | 1.04E+00 | 5.41E-02 | 1.04E+00 |
| ENSG000002 | LINC02422 | 8.34E-01 | ? | 9.11E-01 | -1.35E-01 | -1.10E+00 |
| ENSG000001 | KANK3 | 8.34E-01 | ? | 9.60E-01 | -5.88E-02 | -1.04E+00 |
| ENSG000002 | FBXO16 | 8.34E-01 | 9.43E-01 | 1.02E+00 | 3.45E-02 | 1.02E+00 |
| ENSG000002 | S1PR1-DT | 8.35E-01 | ? | 1.07E+00 | 9.39E-02 | 1.07E+00 |
| ENSG000001 | NLRC5 | 8.35E-01 | 9.43E-01 | 9.80E-01 | -2.92E-02 | -1.02E+00 |
| ENSG000001 | DPY19L2 | 8.35E-01 | 9.43E-01 | 1.03E+00 | 4.59E-02 | 1.03E+00 |
| ENSG000002 | LINC01828 | 8.35E-01 | ? | 1.05E+00 | 6.79E-02 | 1.05E+00 |
| ENSG000001 | WDR11 | 8.35E-01 | 9.43E-01 | 9.95E-01 | -7.44E-03 | -1.01E+00 |
| ENSG000001 | ZNF124 | 8.35E-01 | 9.44E-01 | 9.85E-01 | -2.12E-02 | -1.01E+00 |
| ENSG000002 | SUMO1P3 | 8.35E-01 | ? | 9.15E-01 | -1.29E-01 | -1.09E+00 |
| ENSG000001 | VPS13B | 8.35E-01 | 9.44E-01 | 1.01E+00 | 1.76E-02 | 1.01E+00 |
| ENSG000001 | R3HDM4 | 8.35E-01 | 9.44E-01 | 9.92E-01 | -1.21E-02 | -1.01E+00 |
| ENSG000002 | --- | 8.35E-01 | ? | 1.06E+00 | 7.91E-02 | 1.06E+00 |
| ENSG000002 | --- | 8.35E-01 | ? | 1.06E+00 | 8.71E-02 | 1.06E+00 |
| ENSG000001 | CCDC181 | 8.36E-01 | ? | 9.35E-01 | -9.69E-02 | -1.07E+00 |
| ENSG000001 | SAAL1 | 8.36E-01 | 9.44E-01 | 9.90E-01 | -1.39E-02 | -1.01E+00 |
| ENSG000001 | HYI | 8.36E-01 | 9.44E-01 | 9.89E-01 | -1.64E-02 | -1.01E+00 |
| ENSG000002 | GRID2IP | 8.36E-01 | ? | 8.90E-01 | -1.67E-01 | -1.12E+00 |
| ENSG000001 | PAQR3 | 8.36E-01 | 9.44E-01 | 1.01E+00 | 1.05E-02 | 1.01E+00 |
| ENSG000002 | --- | 8.36E-01 | ? | 9.50E-01 | -7.44E-02 | -1.05E+00 |
| ENSG000001 | GLIPR1 | 8.36E-01 | 9.44E-01 | 1.01E+00 | 1.76E-02 | 1.01E+00 |

| | | | | | | |
|---|---|---|---|---|---|---|
| ENSG000002 | CASP16P | 8.36E-01 | ? | 1.11E+00 | 1.48E-01 | 1.11E+00 |
| ENSG000002 | --- | 8.36E-01 | ? | 9.35E-01 | -9.72E-02 | -1.07E+00 |
| ENSG000001 | EPRS1 | 8.36E-01 | 9.44E-01 | 9.94E-01 | -8.08E-03 | -1.01E+00 |
| ENSG000001 | ZNF300 | 8.36E-01 | 9.44E-01 | 9.83E-01 | -2.41E-02 | -1.02E+00 |
| ENSG000002 | --- | 8.37E-01 | ? | 1.05E+00 | 6.94E-02 | 1.05E+00 |
| ENSG000001 | TRIM22 | 8.37E-01 | 9.45E-01 | 1.01E+00 | 1.66E-02 | 1.01E+00 |
| ENSG000001 | PROZ | 8.37E-01 | ? | 1.10E+00 | 1.36E-01 | 1.10E+00 |
| ENSG000002 | HGH1 | 8.37E-01 | 9.45E-01 | 1.01E+00 | 1.76E-02 | 1.01E+00 |
| ENSG000001 | RACGAP1 | 8.37E-01 | 9.45E-01 | 9.87E-01 | -1.85E-02 | -1.01E+00 |
| ENSG00000C | POMT2 | 8.37E-01 | 9.45E-01 | 1.01E+00 | 1.21E-02 | 1.01E+00 |
| ENSG000002 | --- | 8.37E-01 | ? | 9.31E-01 | -1.03E-01 | -1.07E+00 |
| ENSG000001 | ANKRD50 | 8.37E-01 | 9.45E-01 | 1.01E+00 | 1.24E-02 | 1.01E+00 |
| ENSG000001 | RAD51C | 8.37E-01 | 9.45E-01 | 1.01E+00 | 1.32E-02 | 1.01E+00 |
| ENSG000001 | ISL2 | 8.37E-01 | ? | 1.09E+00 | 1.21E-01 | 1.09E+00 |
| ENSG000001 | WDR35 | 8.38E-01 | 9.45E-01 | 1.01E+00 | 1.55E-02 | 1.01E+00 |
| ENSG000001 | NOC3L | 8.38E-01 | 9.45E-01 | 9.92E-01 | -1.10E-02 | -1.01E+00 |
| ENSG000001 | GHDC | 8.38E-01 | 9.45E-01 | 1.03E+00 | 3.84E-02 | 1.03E+00 |
| ENSG000001 | C12orf57 | 8.38E-01 | 9.45E-01 | 1.01E+00 | 2.02E-02 | 1.01E+00 |
| ENSG000002 | --- | 8.38E-01 | ? | 9.08E-01 | -1.39E-01 | -1.10E+00 |
| ENSG000002 | --- | 8.38E-01 | ? | 9.63E-01 | -5.42E-02 | -1.04E+00 |
| ENSG00000C | YPEL3 | 8.38E-01 | 9.46E-01 | 1.02E+00 | 2.31E-02 | 1.02E+00 |
| ENSG000002 | --- | 8.38E-01 | 9.46E-01 | 1.03E+00 | 3.79E-02 | 1.03E+00 |
| ENSG000002 | MYL5 | 8.38E-01 | 9.46E-01 | 1.03E+00 | 3.71E-02 | 1.03E+00 |
| ENSG000001 | VAV1 | 8.38E-01 | ? | 1.03E+00 | 4.78E-02 | 1.03E+00 |
| ENSG000001 | DGCR8 | 8.38E-01 | 9.46E-01 | 9.95E-01 | -7.41E-03 | -1.01E+00 |
| ENSG00000C | TOP2B | 8.38E-01 | 9.46E-01 | 1.01E+00 | 1.07E-02 | 1.01E+00 |
| ENSG000001 | GSDMA | 8.38E-01 | 9.46E-01 | 1.02E+00 | 3.02E-02 | 1.02E+00 |
| ENSG000001 | LDOC1 | 8.38E-01 | 9.46E-01 | 9.91E-01 | -1.35E-02 | -1.01E+00 |
| ENSG000002 | GLUD1P3 | 8.39E-01 | ? | 1.03E+00 | 4.51E-02 | 1.03E+00 |
| ENSG000002 | --- | 8.39E-01 | ? | 1.08E+00 | 1.07E-01 | 1.08E+00 |
| ENSG000002 | --- | 8.39E-01 | ? | 1.12E+00 | 1.61E-01 | 1.12E+00 |
| ENSG000002 | MAP10 | 8.39E-01 | 9.46E-01 | 1.02E+00 | 2.89E-02 | 1.02E+00 |
| ENSG000002 | SOWAHCP2 | 8.39E-01 | ? | 1.05E+00 | 7.22E-02 | 1.05E+00 |
| ENSG000001 | GPR27 | 8.39E-01 | 9.46E-01 | 9.70E-01 | -4.39E-02 | -1.03E+00 |
| ENSG000001 | YKT6 | 8.39E-01 | 9.46E-01 | 1.01E+00 | 7.40E-03 | 1.01E+00 |
| ENSG000001 | PABIR1 | 8.39E-01 | 9.46E-01 | 9.90E-01 | -1.42E-02 | -1.01E+00 |
| ENSG000001 | NAPB | 8.39E-01 | 9.46E-01 | 1.01E+00 | 1.69E-02 | 1.01E+00 |
| ENSG000001 | HYKK | 8.39E-01 | 9.46E-01 | 9.75E-01 | -3.65E-02 | -1.03E+00 |
| ENSG000002 | --- | 8.39E-01 | 9.46E-01 | 1.02E+00 | 2.81E-02 | 1.02E+00 |
| ENSG000001 | KCNK13 | 8.39E-01 | ? | 8.95E-01 | -1.59E-01 | -1.12E+00 |
| ENSG000001 | DTX4 | 8.39E-01 | 9.46E-01 | 9.91E-01 | -1.34E-02 | -1.01E+00 |
| ENSG000001 | TCEANC | 8.39E-01 | 9.46E-01 | 9.80E-01 | -2.97E-02 | -1.02E+00 |
| ENSG000002 | --- | 8.39E-01 | ? | 1.12E+00 | 1.61E-01 | 1.12E+00 |
| ENSG000001 | ARPC1B | 8.39E-01 | 9.46E-01 | 9.89E-01 | -1.62E-02 | -1.01E+00 |
| ENSG000002 | --- | 8.39E-01 | 9.46E-01 | 1.02E+00 | 2.48E-02 | 1.02E+00 |

| | | | | | | |
|---|---|---|---|---|---|---|
| ENSG000002 | TRMT9B | 8.40E-01 | ? | 9.64E-01 | -5.22E-02 | -1.04E+00 |
| ENSG000001 | DGAT1 | 8.40E-01 | 9.46E-01 | 1.01E+00 | 1.99E-02 | 1.01E+00 |
| ENSG000002 | --- | 8.40E-01 | ? | 9.50E-01 | -7.46E-02 | -1.05E+00 |
| ENSG000001 | ANKRD13B | 8.40E-01 | 9.46E-01 | 9.90E-01 | -1.46E-02 | -1.01E+00 |
| ENSG000002 | AKR1B15 | 8.40E-01 | ? | 9.37E-01 | -9.42E-02 | -1.07E+00 |
| ENSG000001 | TTF2 | 8.40E-01 | 9.46E-01 | 9.92E-01 | -1.13E-02 | -1.01E+00 |
| ENSG000001 | MYLK4 | 8.40E-01 | ? | 1.04E+00 | 5.34E-02 | 1.04E+00 |
| ENSG000001 | CAPN15 | 8.40E-01 | 9.46E-01 | 1.02E+00 | 2.54E-02 | 1.02E+00 |
| ENSG000002 | LINC00857 | 8.40E-01 | 9.46E-01 | 1.02E+00 | 2.89E-02 | 1.02E+00 |
| ENSG000001 | SHLD1 | 8.40E-01 | 9.46E-01 | 1.02E+00 | 2.99E-02 | 1.02E+00 |
| ENSG000002 | --- | 8.40E-01 | ? | 9.47E-01 | -7.81E-02 | -1.06E+00 |
| ENSG000001 | PBRM1 | 8.40E-01 | 9.46E-01 | 9.93E-01 | -1.06E-02 | -1.01E+00 |
| ENSG000001 | DOK4 | 8.40E-01 | 9.46E-01 | 1.01E+00 | 1.70E-02 | 1.01E+00 |
| ENSG000001 | RER1 | 8.40E-01 | 9.46E-01 | 1.01E+00 | 1.16E-02 | 1.01E+00 |
| ENSG000001 | HEXD | 8.40E-01 | 9.46E-01 | 1.02E+00 | 2.40E-02 | 1.02E+00 |
| ENSG000001 | ZNF614 | 8.40E-01 | 9.46E-01 | 1.01E+00 | 1.26E-02 | 1.01E+00 |
| ENSG000002 | GTF2H2B | 8.40E-01 | 9.46E-01 | 1.02E+00 | 2.71E-02 | 1.02E+00 |
| ENSG000002 | LINC01535 | 8.40E-01 | ? | 9.47E-01 | -7.82E-02 | -1.06E+00 |
| ENSG000001 | AHNAK | 8.40E-01 | 9.46E-01 | 9.81E-01 | -2.81E-02 | -1.02E+00 |
| ENSG000002 | --- | 8.40E-01 | ? | 9.37E-01 | -9.36E-02 | -1.07E+00 |
| ENSG000001 | TRIM7 | 8.40E-01 | 9.46E-01 | 9.90E-01 | -1.39E-02 | -1.01E+00 |
| ENSG00000C | PDCD2 | 8.40E-01 | 9.46E-01 | 1.01E+00 | 9.05E-03 | 1.01E+00 |
| ENSG000001 | SLC1A5 | 8.40E-01 | 9.46E-01 | 1.01E+00 | 1.67E-02 | 1.01E+00 |
| ENSG000001 | FANCC | 8.40E-01 | 9.46E-01 | 1.01E+00 | 1.72E-02 | 1.01E+00 |
| ENSG000002 | FOSL2-AS1 | 8.40E-01 | ? | 9.57E-01 | -6.33E-02 | -1.04E+00 |
| ENSG00000C | CHGB | 8.41E-01 | 9.46E-01 | 9.66E-01 | -4.95E-02 | -1.03E+00 |
| ENSG000001 | PLAA | 8.41E-01 | 9.46E-01 | 1.01E+00 | 7.35E-03 | 1.01E+00 |
| ENSG000002 | --- | 8.41E-01 | ? | 1.05E+00 | 6.68E-02 | 1.05E+00 |
| ENSG000001 | VPS39 | 8.41E-01 | 9.46E-01 | 9.94E-01 | -8.29E-03 | -1.01E+00 |
| ENSG000001 | GINS4 | 8.41E-01 | 9.46E-01 | 9.86E-01 | -2.01E-02 | -1.01E+00 |
| ENSG000002 | LINC02936 | 8.41E-01 | ? | 1.06E+00 | 8.45E-02 | 1.06E+00 |
| ENSG000002 | --- | 8.41E-01 | ? | 1.05E+00 | 7.72E-02 | 1.05E+00 |
| ENSG000001 | TAFA2 | 8.41E-01 | ? | 1.05E+00 | 6.91E-02 | 1.05E+00 |
| ENSG000001 | VASN | 8.41E-01 | 9.46E-01 | 9.75E-01 | -3.69E-02 | -1.03E+00 |
| ENSG000001 | ADGRE2 | 8.41E-01 | ? | 1.04E+00 | 5.39E-02 | 1.04E+00 |
| ENSG000002 | NADK2-AS1 | 8.41E-01 | ? | 1.09E+00 | 1.23E-01 | 1.09E+00 |
| ENSG000002 | --- | 8.41E-01 | ? | 1.09E+00 | 1.31E-01 | 1.09E+00 |
| ENSG000001 | ACP1 | 8.41E-01 | 9.46E-01 | 9.95E-01 | -7.26E-03 | -1.01E+00 |
| ENSG000001 | SAP18 | 8.41E-01 | 9.46E-01 | 1.01E+00 | 7.47E-03 | 1.01E+00 |
| ENSG000001 | RUFY1 | 8.41E-01 | 9.46E-01 | 9.92E-01 | -1.11E-02 | -1.01E+00 |
| ENSG000002 | LINC02804 | 8.41E-01 | ? | 9.24E-01 | -1.14E-01 | -1.08E+00 |
| ENSG000001 | DRG2 | 8.41E-01 | 9.46E-01 | 9.91E-01 | -1.29E-02 | -1.01E+00 |
| ENSG000001 | KIAA0586 | 8.41E-01 | 9.46E-01 | 9.91E-01 | -1.33E-02 | -1.01E+00 |
| ENSG000001 | ORC4 | 8.41E-01 | 9.46E-01 | 9.94E-01 | -9.40E-03 | -1.01E+00 |
| ENSG000001 | TLR3 | 8.41E-01 | 9.46E-01 | 1.02E+00 | 2.97E-02 | 1.02E+00 |

| | | | | | | |
|---|---|---|---|---|---|---|
| ENSG000001 | ZNF440 | 8.41E-01 | 9.46E-01 | 9.91E-01 | -1.37E-02 | -1.01E+00 |
| ENSG000001 | BBX | 8.41E-01 | 9.46E-01 | 9.92E-01 | -1.13E-02 | -1.01E+00 |
| ENSG000001 | DNAJB2 | 8.41E-01 | 9.46E-01 | 1.01E+00 | 1.28E-02 | 1.01E+00 |
| ENSG000001 | TBC1D13 | 8.42E-01 | 9.46E-01 | 9.90E-01 | -1.51E-02 | -1.01E+00 |
| ENSG000001 | APLP1 | 8.42E-01 | ? | 9.62E-01 | -5.54E-02 | -1.04E+00 |
| ENSG000001 | CACNB4 | 8.42E-01 | ? | 9.56E-01 | -6.42E-02 | -1.05E+00 |
| ENSG000002 | --- | 8.42E-01 | ? | 1.08E+00 | 1.16E-01 | 1.08E+00 |
| ENSG000001 | KLHL4 | 8.42E-01 | 9.46E-01 | 9.88E-01 | -1.73E-02 | -1.01E+00 |
| ENSG000001 | ZCCHC4 | 8.42E-01 | 9.46E-01 | 1.01E+00 | 1.47E-02 | 1.01E+00 |
| ENSG00000C | PHF7 | 8.42E-01 | 9.46E-01 | 1.02E+00 | 2.30E-02 | 1.02E+00 |
| ENSG000002 | SNHG31 | 8.42E-01 | 9.46E-01 | 1.03E+00 | 4.74E-02 | 1.03E+00 |
| ENSG000002 | --- | 8.42E-01 | ? | 9.54E-01 | -6.73E-02 | -1.05E+00 |
| ENSG000002 | DDN-AS1 | 8.42E-01 | ? | 9.45E-01 | -8.16E-02 | -1.06E+00 |
| ENSG00000C | RANBP3 | 8.42E-01 | 9.46E-01 | 9.93E-01 | -1.03E-02 | -1.01E+00 |
| ENSG000001 | METTL15 | 8.42E-01 | 9.46E-01 | 1.01E+00 | 1.27E-02 | 1.01E+00 |
| ENSG000002 | --- | 8.42E-01 | ? | 9.55E-01 | -6.63E-02 | -1.05E+00 |
| ENSG00000C | PCDHGA2 | 8.42E-01 | 9.46E-01 | 9.71E-01 | -4.23E-02 | -1.03E+00 |
| ENSG000001 | SH2D4B | 8.42E-01 | ? | 9.39E-01 | -9.13E-02 | -1.07E+00 |
| ENSG000001 | NR4A3 | 8.42E-01 | ? | 1.05E+00 | 7.42E-02 | 1.05E+00 |
| ENSG000001 | N4BP3 | 8.42E-01 | 9.46E-01 | 9.78E-01 | -3.25E-02 | -1.02E+00 |
| ENSG00000C | FUCA2 | 8.43E-01 | 9.46E-01 | 1.01E+00 | 7.92E-03 | 1.01E+00 |
| ENSG000002 | --- | 8.43E-01 | ? | 1.09E+00 | 1.18E-01 | 1.09E+00 |
| ENSG000001 | CMC2 | 8.43E-01 | 9.46E-01 | 9.90E-01 | -1.45E-02 | -1.01E+00 |
| ENSG000001 | SLC25A35 | 8.43E-01 | 9.47E-01 | 9.70E-01 | -4.36E-02 | -1.03E+00 |
| ENSG000002 | BRWD1-AS2 | 8.43E-01 | ? | 1.08E+00 | 1.11E-01 | 1.08E+00 |
| ENSG000001 | EHD1 | 8.43E-01 | 9.47E-01 | 1.01E+00 | 1.63E-02 | 1.01E+00 |
| ENSG00000C | UBE2T | 8.43E-01 | 9.47E-01 | 9.84E-01 | -2.26E-02 | -1.02E+00 |
| ENSG000002 | KRT18P34 | 8.43E-01 | ? | 1.03E+00 | 4.63E-02 | 1.03E+00 |
| ENSG000001 | RGS9 | 8.43E-01 | 9.47E-01 | 1.02E+00 | 2.99E-02 | 1.02E+00 |
| ENSG000001 | SELENBP1 | 8.43E-01 | ? | 9.38E-01 | -9.28E-02 | -1.07E+00 |
| ENSG000002 | --- | 8.43E-01 | ? | 9.52E-01 | -7.16E-02 | -1.05E+00 |
| ENSG000002 | --- | 8.43E-01 | ? | 1.05E+00 | 7.24E-02 | 1.05E+00 |
| ENSG000002 | USP27X-DT | 8.43E-01 | ? | 1.04E+00 | 6.18E-02 | 1.04E+00 |
| ENSG000001 | ARHGAP22 | 8.43E-01 | 9.47E-01 | 1.01E+00 | 1.25E-02 | 1.01E+00 |
| ENSG00000C | GPR137B | 8.43E-01 | 9.47E-01 | 9.92E-01 | -1.23E-02 | -1.01E+00 |
| ENSG000001 | NFIB | 8.43E-01 | 9.47E-01 | 9.92E-01 | -1.19E-02 | -1.01E+00 |
| ENSG00000C | RGS11 | 8.44E-01 | ? | 1.05E+00 | 6.48E-02 | 1.05E+00 |
| ENSG000001 | BEST1 | 8.44E-01 | 9.47E-01 | 9.83E-01 | -2.49E-02 | -1.02E+00 |
| ENSG000002 | LINC01152 | 8.44E-01 | ? | 1.07E+00 | 1.03E-01 | 1.07E+00 |
| ENSG000001 | HGS | 8.44E-01 | 9.47E-01 | 1.01E+00 | 1.52E-02 | 1.01E+00 |
| ENSG000001 | CLPTM1 | 8.44E-01 | 9.47E-01 | 9.94E-01 | -8.78E-03 | -1.01E+00 |
| ENSG000001 | NAA40 | 8.44E-01 | 9.47E-01 | 1.01E+00 | 1.11E-02 | 1.01E+00 |
| ENSG000001 | GFI1 | 8.44E-01 | ? | 1.06E+00 | 9.01E-02 | 1.06E+00 |
| ENSG000001 | GPRIN1 | 8.44E-01 | 9.47E-01 | 1.02E+00 | 2.22E-02 | 1.02E+00 |
| ENSG000002 | --- | 8.44E-01 | ? | 1.05E+00 | 7.22E-02 | 1.05E+00 |

| | | | | | | |
|---|---|---|---|---|---|---|
| ENSG000001 | ARFGEF2 | 8.44E-01 | 9.47E-01 | 9.94E-01 | -7.97E-03 | -1.01E+00 |
| ENSG000002 | CSNK2A3 | 8.44E-01 | 9.47E-01 | 1.03E+00 | 4.09E-02 | 1.03E+00 |
| ENSG000001 | ARHGEF16 | 8.44E-01 | 9.47E-01 | 9.86E-01 | -2.01E-02 | -1.01E+00 |
| ENSG000001 | ATAD2B | 8.44E-01 | 9.47E-01 | 9.88E-01 | -1.67E-02 | -1.01E+00 |
| ENSG00000C | MISP | 8.44E-01 | 9.47E-01 | 1.01E+00 | 1.52E-02 | 1.01E+00 |
| ENSG000001 | CCT4 | 8.45E-01 | 9.47E-01 | 9.96E-01 | -5.83E-03 | -1.00E+00 |
| ENSG000001 | SPATA6 | 8.45E-01 | 9.47E-01 | 9.83E-01 | -2.49E-02 | -1.02E+00 |
| ENSG000001 | RNF11 | 8.45E-01 | 9.47E-01 | 1.00E+00 | 7.18E-03 | 1.00E+00 |
| ENSG000001 | MED25 | 8.45E-01 | 9.47E-01 | 1.01E+00 | 1.23E-02 | 1.01E+00 |
| ENSG000001 | C6orf132 | 8.45E-01 | 9.47E-01 | 1.01E+00 | 8.20E-03 | 1.01E+00 |
| ENSG000002 | --- | 8.45E-01 | ? | 1.12E+00 | 1.63E-01 | 1.12E+00 |
| ENSG000002 | SNTG2-AS1 | 8.45E-01 | ? | 1.11E+00 | 1.53E-01 | 1.11E+00 |
| ENSG000001 | SMCO4 | 8.45E-01 | ? | 1.05E+00 | 6.77E-02 | 1.05E+00 |
| ENSG00000C | TTC22 | 8.45E-01 | 9.47E-01 | 1.01E+00 | 1.59E-02 | 1.01E+00 |
| ENSG000002 | --- | 8.45E-01 | 9.47E-01 | 1.03E+00 | 3.73E-02 | 1.03E+00 |
| ENSG000001 | CCDC81 | 8.45E-01 | ? | 9.40E-01 | -8.98E-02 | -1.06E+00 |
| ENSG000001 | NKX3-1 | 8.45E-01 | 9.47E-01 | 9.88E-01 | -1.76E-02 | -1.01E+00 |
| ENSG000001 | DUSP23 | 8.45E-01 | 9.47E-01 | 9.85E-01 | -2.24E-02 | -1.02E+00 |
| ENSG000001 | CNIH1 | 8.45E-01 | 9.47E-01 | 1.01E+00 | 9.84E-03 | 1.01E+00 |
| ENSG000002 | --- | 8.45E-01 | ? | 1.10E+00 | 1.42E-01 | 1.10E+00 |
| ENSG000001 | HIRIP3 | 8.45E-01 | 9.47E-01 | 9.90E-01 | -1.43E-02 | -1.01E+00 |
| ENSG000001 | DUOXA2 | 8.45E-01 | ? | 9.30E-01 | -1.05E-01 | -1.08E+00 |
| ENSG000002 | --- | 8.45E-01 | ? | 1.09E+00 | 1.25E-01 | 1.09E+00 |
| ENSG000001 | TYMS | 8.45E-01 | 9.47E-01 | 1.01E+00 | 1.16E-02 | 1.01E+00 |
| ENSG000002 | FSBP | 8.45E-01 | 9.47E-01 | 9.75E-01 | -3.61E-02 | -1.03E+00 |
| ENSG000002 | GOLGA8N | 8.45E-01 | 9.47E-01 | 9.87E-01 | -1.81E-02 | -1.01E+00 |
| ENSG000002 | DMTF1-AS1 | 8.46E-01 | ? | 1.08E+00 | 1.05E-01 | 1.08E+00 |
| ENSG000002 | --- | 8.46E-01 | ? | 9.55E-01 | -6.68E-02 | -1.05E+00 |
| ENSG000002 | --- | 8.46E-01 | ? | 1.10E+00 | 1.34E-01 | 1.10E+00 |
| ENSG000002 | --- | 8.46E-01 | ? | 1.09E+00 | 1.24E-01 | 1.09E+00 |
| ENSG000002 | --- | 8.46E-01 | 9.47E-01 | 1.01E+00 | 1.87E-02 | 1.01E+00 |
| ENSG000001 | SLC33A1 | 8.46E-01 | 9.47E-01 | 9.93E-01 | -9.94E-03 | -1.01E+00 |
| ENSG000002 | --- | 8.46E-01 | ? | 9.51E-01 | -7.27E-02 | -1.05E+00 |
| ENSG000001 | PHETA2 | 8.46E-01 | ? | 9.62E-01 | -5.63E-02 | -1.04E+00 |
| ENSG000002 | TXNP6 | 8.46E-01 | ? | 1.09E+00 | 1.30E-01 | 1.09E+00 |
| ENSG000002 | --- | 8.46E-01 | ? | 1.09E+00 | 1.30E-01 | 1.09E+00 |
| ENSG000002 | EPB41L4A-AS | 8.46E-01 | 9.47E-01 | 9.85E-01 | -2.22E-02 | -1.02E+00 |
| ENSG000002 | --- | 8.46E-01 | 9.47E-01 | 9.70E-01 | -4.42E-02 | -1.03E+00 |
| ENSG000001 | TPGS2 | 8.46E-01 | 9.47E-01 | 1.01E+00 | 8.78E-03 | 1.01E+00 |
| ENSG000001 | DDX31 | 8.46E-01 | 9.47E-01 | 9.92E-01 | -1.14E-02 | -1.01E+00 |
| ENSG000002 | --- | 8.46E-01 | ? | 1.03E+00 | 4.48E-02 | 1.03E+00 |
| ENSG000001 | OSCAR | 8.46E-01 | ? | 9.17E-01 | -1.25E-01 | -1.09E+00 |
| ENSG000001 | ZBTB45 | 8.46E-01 | 9.48E-01 | 1.01E+00 | 1.78E-02 | 1.01E+00 |
| ENSG000001 | GMDS | 8.46E-01 | 9.48E-01 | 1.01E+00 | 1.57E-02 | 1.01E+00 |
| ENSG000001 | NYAP1 | 8.47E-01 | ? | 9.60E-01 | -5.82E-02 | -1.04E+00 |

| | | | | | | |
|---|---|---|---|---|---|---|
| ENSG00000C | ARHGAP4 | 8.47E-01 | ? | 9.50E-01 | -7.41E-02 | -1.05E+00 |
| ENSG000002 | CRCP | 8.47E-01 | 9.48E-01 | 1.01E+00 | 1.02E-02 | 1.01E+00 |
| ENSG000001 | H2AX | 8.47E-01 | 9.48E-01 | 9.89E-01 | -1.65E-02 | -1.01E+00 |
| ENSG000001 | LUZP2 | 8.47E-01 | ? | 1.08E+00 | 1.08E-01 | 1.08E+00 |
| ENSG000001 | MAP1A | 8.47E-01 | 9.48E-01 | 1.02E+00 | 3.48E-02 | 1.02E+00 |
| ENSG000001 | ZKSCAN3 | 8.47E-01 | 9.48E-01 | 1.01E+00 | 1.91E-02 | 1.01E+00 |
| ENSG000002 | MZF1-AS1 | 8.47E-01 | 9.48E-01 | 1.02E+00 | 3.02E-02 | 1.02E+00 |
| ENSG000002 | DHX9-AS1 | 8.47E-01 | ? | 9.21E-01 | -1.19E-01 | -1.09E+00 |
| ENSG000001 | MEX3D | 8.47E-01 | 9.48E-01 | 1.01E+00 | 1.70E-02 | 1.01E+00 |
| ENSG000002 | --- | 8.47E-01 | ? | 9.51E-01 | -7.26E-02 | -1.05E+00 |
| ENSG000001 | UBL7 | 8.47E-01 | 9.48E-01 | 1.01E+00 | 1.55E-02 | 1.01E+00 |
| ENSG000001 | MTSS2 | 8.47E-01 | 9.48E-01 | 9.91E-01 | -1.33E-02 | -1.01E+00 |
| ENSG000001 | IQCG | 8.47E-01 | 9.48E-01 | 9.84E-01 | -2.28E-02 | -1.02E+00 |
| ENSG000002 | --- | 8.47E-01 | ? | 9.63E-01 | -5.39E-02 | -1.04E+00 |
| ENSG000002 | CTAGE7P | 8.47E-01 | ? | 1.04E+00 | 6.19E-02 | 1.04E+00 |
| ENSG000002 | --- | 8.47E-01 | ? | 1.08E+00 | 1.17E-01 | 1.08E+00 |
| ENSG000001 | RFX7 | 8.47E-01 | 9.48E-01 | 9.92E-01 | -1.12E-02 | -1.01E+00 |
| ENSG000001 | PARP2 | 8.47E-01 | 9.48E-01 | 9.92E-01 | -1.19E-02 | -1.01E+00 |
| ENSG000002 | GAPDHP1 | 8.48E-01 | 9.48E-01 | 1.03E+00 | 3.79E-02 | 1.03E+00 |
| ENSG000002 | --- | 8.48E-01 | ? | 1.07E+00 | 9.89E-02 | 1.07E+00 |
| ENSG000001 | UBE2R2 | 8.48E-01 | 9.48E-01 | 1.00E+00 | 6.03E-03 | 1.00E+00 |
| ENSG000002 | HNF1B | 8.48E-01 | 9.48E-01 | 1.01E+00 | 2.08E-02 | 1.01E+00 |
| ENSG000001 | ETFA | 8.48E-01 | 9.48E-01 | 9.94E-01 | -8.16E-03 | -1.01E+00 |
| ENSG000001 | APAF1 | 8.48E-01 | 9.48E-01 | 1.01E+00 | 8.14E-03 | 1.01E+00 |
| ENSG000002 | --- | 8.48E-01 | ? | 1.07E+00 | 1.01E-01 | 1.07E+00 |
| ENSG000001 | ZNF3 | 8.48E-01 | 9.48E-01 | 9.92E-01 | -1.15E-02 | -1.01E+00 |
| ENSG000001 | NKX2-8 | 8.48E-01 | 9.48E-01 | 1.03E+00 | 4.35E-02 | 1.03E+00 |
| ENSG000001 | SNF8 | 8.48E-01 | 9.48E-01 | 1.01E+00 | 1.03E-02 | 1.01E+00 |
| ENSG000001 | POLR2H | 8.48E-01 | 9.48E-01 | 1.01E+00 | 1.29E-02 | 1.01E+00 |
| ENSG000002 | RPL7AP45 | 8.48E-01 | ? | 9.62E-01 | -5.53E-02 | -1.04E+00 |
| ENSG000001 | TTLL12 | 8.48E-01 | 9.48E-01 | 9.91E-01 | -1.29E-02 | -1.01E+00 |
| ENSG00000C | ELAC2 | 8.48E-01 | 9.48E-01 | 1.01E+00 | 7.95E-03 | 1.01E+00 |
| ENSG000001 | MMP23B | 8.48E-01 | ? | 9.42E-01 | -8.69E-02 | -1.06E+00 |
| ENSG000002 | SNHG10 | 8.48E-01 | 9.48E-01 | 9.81E-01 | -2.72E-02 | -1.02E+00 |
| ENSG000002 | SRRM2-AS1 | 8.48E-01 | ? | 1.04E+00 | 5.80E-02 | 1.04E+00 |
| ENSG000001 | TBC1D19 | 8.48E-01 | 9.48E-01 | 9.86E-01 | -1.97E-02 | -1.01E+00 |
| ENSG000001 | C5orf34 | 8.48E-01 | 9.48E-01 | 1.02E+00 | 3.33E-02 | 1.02E+00 |
| ENSG000002 | --- | 8.48E-01 | ? | 9.00E-01 | -1.52E-01 | -1.11E+00 |
| ENSG000001 | RPL24P4 | 8.48E-01 | 9.48E-01 | 9.75E-01 | -3.72E-02 | -1.03E+00 |
| ENSG000001 | DHX36 | 8.48E-01 | 9.48E-01 | 9.96E-01 | -6.25E-03 | -1.00E+00 |
| ENSG000001 | LLPH | 8.49E-01 | 9.48E-01 | 9.92E-01 | -1.22E-02 | -1.01E+00 |
| ENSG000001 | RRP1B | 8.49E-01 | 9.48E-01 | 1.01E+00 | 7.79E-03 | 1.01E+00 |
| ENSG000001 | ZNF528 | 8.49E-01 | 9.48E-01 | 9.90E-01 | -1.40E-02 | -1.01E+00 |
| ENSG000002 | --- | 8.49E-01 | ? | 9.14E-01 | -1.30E-01 | -1.09E+00 |
| ENSG00000C | GNAS | 8.49E-01 | 9.48E-01 | 9.95E-01 | -7.11E-03 | -1.00E+00 |

| | | | | | | |
|---|---|---|---|---|---|---|
| ENSG000001 | PCMTD1 | 8.49E-01 | 9.48E-01 | 1.01E+00 | 1.12E-02 | 1.01E+00 |
| ENSG000001 | TRIM58 | 8.49E-01 | ? | 9.28E-01 | -1.07E-01 | -1.08E+00 |
| ENSG000002 | LTB4R | 8.49E-01 | 9.48E-01 | 1.02E+00 | 2.43E-02 | 1.02E+00 |
| ENSG000002 | --- | 8.49E-01 | ? | 1.06E+00 | 8.68E-02 | 1.06E+00 |
| ENSG000001 | ST3GAL5 | 8.49E-01 | 9.48E-01 | 9.92E-01 | -1.14E-02 | -1.01E+00 |
| ENSG000001 | SLC6A8 | 8.49E-01 | 9.48E-01 | 9.91E-01 | -1.35E-02 | -1.01E+00 |
| ENSG000002 | --- | 8.49E-01 | ? | 9.45E-01 | -8.12E-02 | -1.06E+00 |
| ENSG000001 | GEM | 8.49E-01 | 9.48E-01 | 9.71E-01 | -4.30E-02 | -1.03E+00 |
| ENSG000002 | HEATR5A-DT | 8.49E-01 | ? | 1.04E+00 | 5.03E-02 | 1.04E+00 |
| ENSG000001 | MDN1 | 8.50E-01 | 9.48E-01 | 9.88E-01 | -1.77E-02 | -1.01E+00 |
| ENSG000002 | --- | 8.50E-01 | ? | 9.26E-01 | -1.11E-01 | -1.08E+00 |
| ENSG000002 | --- | 8.50E-01 | 9.48E-01 | 9.67E-01 | -4.78E-02 | -1.03E+00 |
| ENSG000001 | IFT172 | 8.50E-01 | 9.48E-01 | 1.01E+00 | 2.00E-02 | 1.01E+00 |
| ENSG000002 | --- | 8.50E-01 | 9.48E-01 | 1.02E+00 | 3.32E-02 | 1.02E+00 |
| ENSG000001 | RAB9B | 8.50E-01 | ? | 1.05E+00 | 7.36E-02 | 1.05E+00 |
| ENSG000001 | MRGBP | 8.50E-01 | 9.48E-01 | 1.01E+00 | 1.19E-02 | 1.01E+00 |
| ENSG000002 | --- | 8.50E-01 | ? | 9.11E-01 | -1.35E-01 | -1.10E+00 |
| ENSG000002 | LINC01998 | 8.50E-01 | ? | 9.63E-01 | -5.48E-02 | -1.04E+00 |
| ENSG000001 | PTCD1 | 8.50E-01 | 9.48E-01 | 1.01E+00 | 2.04E-02 | 1.01E+00 |
| ENSG000002 | SATB2-AS1 | 8.50E-01 | 9.48E-01 | 1.03E+00 | 4.03E-02 | 1.03E+00 |
| ENSG000002 | LINC01836 | 8.50E-01 | 9.48E-01 | 1.02E+00 | 3.55E-02 | 1.02E+00 |
| ENSG000001 | YRDC | 8.50E-01 | 9.48E-01 | 1.01E+00 | 1.35E-02 | 1.01E+00 |
| ENSG000001 | AURKB | 8.50E-01 | 9.49E-01 | 9.81E-01 | -2.83E-02 | -1.02E+00 |
| ENSG000001 | DCAF1 | 8.50E-01 | 9.49E-01 | 1.01E+00 | 9.54E-03 | 1.01E+00 |
| ENSG000001 | SNHG17 | 8.51E-01 | 9.49E-01 | 9.84E-01 | -2.31E-02 | -1.02E+00 |
| ENSG000001 | TFAP2E | 8.51E-01 | ? | 9.55E-01 | -6.63E-02 | -1.05E+00 |
| ENSG000001 | UNC79 | 8.51E-01 | ? | 9.48E-01 | -7.65E-02 | -1.05E+00 |
| ENSG000002 | --- | 8.51E-01 | ? | 1.08E+00 | 1.07E-01 | 1.08E+00 |
| ENSG000001 | NFXL1 | 8.51E-01 | 9.49E-01 | 1.01E+00 | 1.27E-02 | 1.01E+00 |
| ENSG000001 | KMO | 8.51E-01 | 9.49E-01 | 1.03E+00 | 4.80E-02 | 1.03E+00 |
| ENSG000002 | RAC1P2 | 8.51E-01 | 9.49E-01 | 1.03E+00 | 4.20E-02 | 1.03E+00 |
| ENSG000002 | ANXA2P1 | 8.51E-01 | ? | 9.17E-01 | -1.24E-01 | -1.09E+00 |
| ENSG000002 | --- | 8.51E-01 | ? | 1.08E+00 | 1.18E-01 | 1.08E+00 |
| ENSG000001 | STEAP1B | 8.51E-01 | 9.49E-01 | 9.75E-01 | -3.66E-02 | -1.03E+00 |
| ENSG000001 | LRRC75A | 8.51E-01 | ? | 9.47E-01 | -7.92E-02 | -1.06E+00 |
| ENSG000002 | CPTP | 8.51E-01 | 9.49E-01 | 9.83E-01 | -2.53E-02 | -1.02E+00 |
| ENSG000001 | PTPDC1 | 8.51E-01 | 9.49E-01 | 9.91E-01 | -1.37E-02 | -1.01E+00 |
| ENSG000001 | SEPTIN3 | 8.51E-01 | ? | 1.05E+00 | 6.58E-02 | 1.05E+00 |
| ENSG000001 | ATP6V1C2 | 8.51E-01 | 9.49E-01 | 9.75E-01 | -3.61E-02 | -1.03E+00 |
| ENSG000002 | LINC01537 | 8.52E-01 | 9.49E-01 | 9.77E-01 | -3.36E-02 | -1.02E+00 |
| ENSG000001 | HROB | 8.52E-01 | 9.49E-01 | 9.77E-01 | -3.32E-02 | -1.02E+00 |
| ENSG000001 | NKIRAS1 | 8.52E-01 | 9.49E-01 | 1.01E+00 | 1.30E-02 | 1.01E+00 |
| ENSG000001 | SDR42E1 | 8.52E-01 | 9.49E-01 | 9.93E-01 | -1.05E-02 | -1.01E+00 |
| ENSG000001 | ABCA4 | 8.52E-01 | ? | 1.04E+00 | 6.18E-02 | 1.04E+00 |
| ENSG000002 | --- | 8.52E-01 | ? | 1.04E+00 | 6.13E-02 | 1.04E+00 |

| | | | | | | |
|---|---|---|---|---|---|---|
| ENSG000002 | GATA2-AS1 | 8.52E-01 | ? | 1.08E+00 | 1.17E-01 | 1.08E+00 |
| ENSG000002 | EHMT2-AS1 | 8.52E-01 | ? | 1.07E+00 | 9.75E-02 | 1.07E+00 |
| ENSG000001 | SLC49A3 | 8.52E-01 | ? | 9.39E-01 | -9.12E-02 | -1.07E+00 |
| ENSG000002 | --- | 8.52E-01 | ? | 9.35E-01 | -9.64E-02 | -1.07E+00 |
| ENSG000001 | SEPTIN5 | 8.52E-01 | 9.49E-01 | 9.87E-01 | -1.85E-02 | -1.01E+00 |
| ENSG000002 | --- | 8.52E-01 | ? | 1.07E+00 | 9.87E-02 | 1.07E+00 |
| ENSG000002 | SLC8A1-AS1 | 8.52E-01 | ? | 9.23E-01 | -1.16E-01 | -1.08E+00 |
| ENSG00000C | PTPN18 | 8.52E-01 | 9.50E-01 | 9.93E-01 | -9.99E-03 | -1.01E+00 |
| ENSG000002 | TBC1D3D | 8.52E-01 | 9.50E-01 | 9.80E-01 | -2.96E-02 | -1.02E+00 |
| ENSG000001 | SLFN12 | 8.52E-01 | 9.50E-01 | 1.01E+00 | 1.14E-02 | 1.01E+00 |
| ENSG000001 | ADA | 8.52E-01 | 9.50E-01 | 9.88E-01 | -1.77E-02 | -1.01E+00 |
| ENSG000001 | CYP1A1 | 8.52E-01 | 9.50E-01 | 9.82E-01 | -2.67E-02 | -1.02E+00 |
| ENSG000002 | --- | 8.53E-01 | ? | 1.10E+00 | 1.43E-01 | 1.10E+00 |
| ENSG000001 | PLCH2 | 8.53E-01 | 9.50E-01 | 1.01E+00 | 1.90E-02 | 1.01E+00 |
| ENSG000001 | IZUMO1 | 8.53E-01 | ? | 9.55E-01 | -6.63E-02 | -1.05E+00 |
| ENSG000002 | WASL-DT | 8.53E-01 | 9.50E-01 | 1.02E+00 | 2.58E-02 | 1.02E+00 |
| ENSG000001 | AATK | 8.53E-01 | 9.50E-01 | 9.80E-01 | -2.90E-02 | -1.02E+00 |
| ENSG00000C | REX1BD | 8.53E-01 | 9.50E-01 | 1.02E+00 | 2.23E-02 | 1.02E+00 |
| ENSG000002 | --- | 8.53E-01 | 9.50E-01 | 1.02E+00 | 2.75E-02 | 1.02E+00 |
| ENSG000002 | PIGCP1 | 8.53E-01 | 9.50E-01 | 1.01E+00 | 2.01E-02 | 1.01E+00 |
| ENSG000001 | ZNF77 | 8.53E-01 | 9.50E-01 | 9.84E-01 | -2.33E-02 | -1.02E+00 |
| ENSG000002 | IL6ST-DT | 8.53E-01 | ? | 1.09E+00 | 1.30E-01 | 1.09E+00 |
| ENSG000002 | GMDS-DT | 8.53E-01 | 9.50E-01 | 9.77E-01 | -3.33E-02 | -1.02E+00 |
| ENSG000002 | --- | 8.53E-01 | ? | 1.10E+00 | 1.43E-01 | 1.10E+00 |
| ENSG00000C | BCORL1 | 8.54E-01 | 9.50E-01 | 1.01E+00 | 1.09E-02 | 1.01E+00 |
| ENSG000002 | C1orf53 | 8.54E-01 | ? | 9.69E-01 | -4.60E-02 | -1.03E+00 |
| ENSG000002 | LINC02980 | 8.54E-01 | ? | 1.03E+00 | 4.76E-02 | 1.03E+00 |
| ENSG000002 | --- | 8.54E-01 | 9.51E-01 | 1.03E+00 | 4.12E-02 | 1.03E+00 |
| ENSG000001 | TRAPPC4 | 8.54E-01 | 9.51E-01 | 1.01E+00 | 1.01E-02 | 1.01E+00 |
| ENSG000001 | MND1 | 8.54E-01 | ? | 1.04E+00 | 6.34E-02 | 1.04E+00 |
| ENSG000001 | NBEAL1 | 8.54E-01 | 9.51E-01 | 1.01E+00 | 1.61E-02 | 1.01E+00 |
| ENSG000001 | ZBTB7C | 8.54E-01 | ? | 1.04E+00 | 5.05E-02 | 1.04E+00 |
| ENSG000001 | TMED3 | 8.54E-01 | 9.51E-01 | 9.90E-01 | -1.44E-02 | -1.01E+00 |
| ENSG000002 | HLA-A | 8.55E-01 | 9.51E-01 | 1.01E+00 | 1.32E-02 | 1.01E+00 |
| ENSG000002 | --- | 8.55E-01 | ? | 1.11E+00 | 1.56E-01 | 1.11E+00 |
| ENSG00000C | CD4 | 8.55E-01 | ? | 9.56E-01 | -6.44E-02 | -1.05E+00 |
| ENSG00000C | PMS1 | 8.55E-01 | 9.51E-01 | 1.01E+00 | 1.50E-02 | 1.01E+00 |
| ENSG00000C | LAMP2 | 8.55E-01 | 9.51E-01 | 1.00E+00 | 5.66E-03 | 1.00E+00 |
| ENSG000001 | SDCBP | 8.55E-01 | 9.51E-01 | 9.95E-01 | -7.76E-03 | -1.01E+00 |
| ENSG000001 | ZNF789 | 8.55E-01 | 9.51E-01 | 9.82E-01 | -2.57E-02 | -1.02E+00 |
| ENSG000001 | DIP2C | 8.55E-01 | 9.51E-01 | 9.92E-01 | -1.22E-02 | -1.01E+00 |
| ENSG000001 | ST6GALNAC5 | 8.55E-01 | 9.51E-01 | 9.81E-01 | -2.72E-02 | -1.02E+00 |
| ENSG000002 | ADH5P4 | 8.55E-01 | ? | 1.07E+00 | 9.67E-02 | 1.07E+00 |
| ENSG000001 | ZNF738 | 8.55E-01 | 9.51E-01 | 9.85E-01 | -2.19E-02 | -1.02E+00 |
| ENSG000001 | RNF150 | 8.55E-01 | ? | 1.08E+00 | 1.04E-01 | 1.08E+00 |

| | | | | | | |
|---|---|---|---|---|---|---|
| ENSG000002 | --- | 8.55E-01 | ? | 1.06E+00 | 8.44E-02 | 1.06E+00 |
| ENSG00000C | KIF22 | 8.56E-01 | 9.51E-01 | 1.01E+00 | 1.82E-02 | 1.01E+00 |
| ENSG000002 | --- | 8.56E-01 | 9.51E-01 | 1.11E+00 | 1.46E-01 | 1.11E+00 |
| ENSG000001 | NUP42 | 8.56E-01 | 9.51E-01 | 1.01E+00 | 1.32E-02 | 1.01E+00 |
| ENSG000001 | CDK5RAP1 | 8.56E-01 | 9.51E-01 | 9.93E-01 | -1.01E-02 | -1.01E+00 |
| ENSG00000C | MLH1 | 8.56E-01 | 9.51E-01 | 1.01E+00 | 1.09E-02 | 1.01E+00 |
| ENSG000002 | LINC02159 | 8.56E-01 | 9.51E-01 | 9.77E-01 | -3.29E-02 | -1.02E+00 |
| ENSG000001 | SUCLG2 | 8.56E-01 | 9.51E-01 | 9.93E-01 | -1.04E-02 | -1.01E+00 |
| ENSG000001 | GPR89A | 8.56E-01 | 9.51E-01 | 1.01E+00 | 1.18E-02 | 1.01E+00 |
| ENSG000001 | FGGY | 8.56E-01 | 9.51E-01 | 1.01E+00 | 1.20E-02 | 1.01E+00 |
| ENSG000002 | --- | 8.56E-01 | ? | 1.09E+00 | 1.29E-01 | 1.09E+00 |
| ENSG000002 | TREX1 | 8.56E-01 | ? | 1.07E+00 | 9.25E-02 | 1.07E+00 |
| ENSG000001 | MAFG | 8.56E-01 | 9.51E-01 | 9.92E-01 | -1.13E-02 | -1.01E+00 |
| ENSG000001 | RMC1 | 8.56E-01 | 9.52E-01 | 9.93E-01 | -1.03E-02 | -1.01E+00 |
| ENSG000001 | PDCD5 | 8.57E-01 | 9.52E-01 | 9.92E-01 | -1.13E-02 | -1.01E+00 |
| ENSG000001 | EOLA1 | 8.57E-01 | 9.52E-01 | 9.89E-01 | -1.54E-02 | -1.01E+00 |
| ENSG000002 | ZNF528-AS1 | 8.57E-01 | 9.52E-01 | 1.01E+00 | 2.01E-02 | 1.01E+00 |
| ENSG000001 | GTF2H3 | 8.57E-01 | 9.52E-01 | 1.01E+00 | 1.02E-02 | 1.01E+00 |
| ENSG000001 | ARHGEF12 | 8.57E-01 | 9.52E-01 | 1.01E+00 | 1.04E-02 | 1.01E+00 |
| ENSG000002 | PKD1-AS1 | 8.57E-01 | ? | 1.09E+00 | 1.30E-01 | 1.09E+00 |
| ENSG000002 | --- | 8.57E-01 | ? | 1.06E+00 | 8.69E-02 | 1.06E+00 |
| ENSG000001 | NRSN2 | 8.57E-01 | 9.52E-01 | 1.02E+00 | 3.18E-02 | 1.02E+00 |
| ENSG000002 | ATP1A1-AS1 | 8.57E-01 | 9.52E-01 | 9.87E-01 | -1.89E-02 | -1.01E+00 |
| ENSG000001 | NTN5 | 8.57E-01 | ? | 1.06E+00 | 8.98E-02 | 1.06E+00 |
| ENSG000002 | FBXW4P1 | 8.57E-01 | ? | 9.65E-01 | -5.11E-02 | -1.04E+00 |
| ENSG000001 | UQCRB | 8.57E-01 | 9.52E-01 | 1.01E+00 | 9.88E-03 | 1.01E+00 |
| ENSG000001 | SFMBT1 | 8.57E-01 | 9.52E-01 | 9.88E-01 | -1.71E-02 | -1.01E+00 |
| ENSG000002 | PSG7 | 8.57E-01 | ? | 9.51E-01 | -7.26E-02 | -1.05E+00 |
| ENSG000002 | LINC02940 | 8.57E-01 | ? | 9.52E-01 | -7.10E-02 | -1.05E+00 |
| ENSG000001 | UBE2E1 | 8.58E-01 | 9.52E-01 | 1.01E+00 | 9.46E-03 | 1.01E+00 |
| ENSG000002 | GLIDR | 8.58E-01 | 9.52E-01 | 1.02E+00 | 2.82E-02 | 1.02E+00 |
| ENSG000001 | WDR86 | 8.58E-01 | ? | 9.16E-01 | -1.26E-01 | -1.09E+00 |
| ENSG000002 | --- | 8.58E-01 | ? | 9.17E-01 | -1.25E-01 | -1.09E+00 |
| ENSG000002 | --- | 8.58E-01 | ? | 9.41E-01 | -8.74E-02 | -1.06E+00 |
| ENSG00000C | MMP25 | 8.58E-01 | 9.52E-01 | 9.79E-01 | -3.12E-02 | -1.02E+00 |
| ENSG000002 | --- | 8.58E-01 | ? | 9.19E-01 | -1.23E-01 | -1.09E+00 |
| ENSG000001 | NPHP1 | 8.58E-01 | 9.52E-01 | 9.85E-01 | -2.18E-02 | -1.02E+00 |
| ENSG000001 | KLRG1 | 8.58E-01 | ? | 9.46E-01 | -7.97E-02 | -1.06E+00 |
| ENSG000001 | UBXN1 | 8.58E-01 | 9.53E-01 | 9.94E-01 | -9.17E-03 | -1.01E+00 |
| ENSG00000C | CXCL2 | 8.58E-01 | 9.53E-01 | 9.46E-01 | -7.96E-02 | -1.06E+00 |
| ENSG000001 | FAM221A | 8.58E-01 | ? | 1.03E+00 | 4.44E-02 | 1.03E+00 |
| ENSG000001 | YIPF3 | 8.59E-01 | 9.53E-01 | 9.93E-01 | -1.00E-02 | -1.01E+00 |
| ENSG000001 | FGD1 | 8.59E-01 | 9.53E-01 | 9.93E-01 | -1.07E-02 | -1.01E+00 |
| ENSG000002 | --- | 8.59E-01 | ? | 9.67E-01 | -4.81E-02 | -1.03E+00 |
| ENSG000001 | COX6B2 | 8.59E-01 | 9.53E-01 | 9.72E-01 | -4.13E-02 | -1.03E+00 |

| | | | | | | |
|---|---|---|---|---|---|---|
| ENSG000002 | RPS18P9 | 8.59E-01 | 9.53E-01 | 1.01E+00 | 1.85E-02 | 1.01E+00 |
| ENSG000002 | LINC02709 | 8.59E-01 | ? | 9.54E-01 | -6.81E-02 | -1.05E+00 |
| ENSG000001 | THSD4 | 8.59E-01 | 9.53E-01 | 1.01E+00 | 1.10E-02 | 1.01E+00 |
| ENSG000001 | DDX23 | 8.59E-01 | 9.53E-01 | 9.96E-01 | -5.58E-03 | -1.00E+00 |
| ENSG000001 | FN3KRP | 8.59E-01 | 9.53E-01 | 9.92E-01 | -1.19E-02 | -1.01E+00 |
| ENSG000001 | ADSS1 | 8.59E-01 | 9.53E-01 | 9.85E-01 | -2.15E-02 | -1.02E+00 |
| ENSG000002 | --- | 8.59E-01 | ? | 1.05E+00 | 7.50E-02 | 1.05E+00 |
| ENSG000002 | PCNPP3 | 8.59E-01 | ? | 1.05E+00 | 7.66E-02 | 1.05E+00 |
| ENSG000002 | --- | 8.59E-01 | ? | 1.05E+00 | 7.12E-02 | 1.05E+00 |
| ENSG000002 | FAM95A | 8.59E-01 | ? | 9.39E-01 | -9.09E-02 | -1.07E+00 |
| ENSG000001 | ATP1A3 | 8.60E-01 | ? | 1.06E+00 | 8.61E-02 | 1.06E+00 |
| ENSG000001 | DBNL | 8.60E-01 | 9.53E-01 | 1.01E+00 | 9.88E-03 | 1.01E+00 |
| ENSG000001 | TYW1 | 8.60E-01 | 9.53E-01 | 1.01E+00 | 9.54E-03 | 1.01E+00 |
| ENSG000001 | CEP250 | 8.60E-01 | 9.53E-01 | 9.93E-01 | -1.05E-02 | -1.01E+00 |
| ENSG000002 | --- | 8.60E-01 | ? | 1.07E+00 | 9.45E-02 | 1.07E+00 |
| ENSG000002 | --- | 8.60E-01 | ? | 1.10E+00 | 1.34E-01 | 1.10E+00 |
| ENSG00000C | NKAIN1 | 8.60E-01 | ? | 1.05E+00 | 7.23E-02 | 1.05E+00 |
| ENSG000001 | GPR68 | 8.60E-01 | 9.54E-01 | 9.81E-01 | -2.70E-02 | -1.02E+00 |
| ENSG000001 | OPTN | 8.60E-01 | 9.54E-01 | 1.01E+00 | 1.02E-02 | 1.01E+00 |
| ENSG000002 | --- | 8.60E-01 | ? | 9.58E-01 | -6.14E-02 | -1.04E+00 |
| ENSG000001 | SPG7 | 8.60E-01 | 9.54E-01 | 9.93E-01 | -9.83E-03 | -1.01E+00 |
| ENSG000001 | NKPD1 | 8.60E-01 | ? | 1.08E+00 | 1.16E-01 | 1.08E+00 |
| ENSG000002 | DNAJC9-AS1 | 8.60E-01 | ? | 1.07E+00 | 9.79E-02 | 1.07E+00 |
| ENSG000002 | PAPLN-AS1 | 8.60E-01 | 9.54E-01 | 1.02E+00 | 2.39E-02 | 1.02E+00 |
| ENSG000001 | ITFG1 | 8.60E-01 | 9.54E-01 | 1.01E+00 | 7.85E-03 | 1.01E+00 |
| ENSG000001 | RPH3AL | 8.60E-01 | 9.54E-01 | 9.85E-01 | -2.11E-02 | -1.01E+00 |
| ENSG000002 | --- | 8.60E-01 | 9.54E-01 | 9.80E-01 | -2.91E-02 | -1.02E+00 |
| ENSG000001 | GPR78 | 8.60E-01 | ? | 1.09E+00 | 1.29E-01 | 1.09E+00 |
| ENSG000001 | GPD1 | 8.60E-01 | ? | 1.05E+00 | 6.83E-02 | 1.05E+00 |
| ENSG000001 | RGPD8 | 8.61E-01 | 9.54E-01 | 9.80E-01 | -2.98E-02 | -1.02E+00 |
| ENSG000002 | TP73-AS1 | 8.61E-01 | 9.54E-01 | 9.90E-01 | -1.52E-02 | -1.01E+00 |
| ENSG000002 | --- | 8.61E-01 | ? | 1.04E+00 | 5.48E-02 | 1.04E+00 |
| ENSG000002 | LGALS8-AS1 | 8.61E-01 | ? | 9.55E-01 | -6.67E-02 | -1.05E+00 |
| ENSG00000C | SREBF1 | 8.61E-01 | 9.54E-01 | 1.01E+00 | 1.75E-02 | 1.01E+00 |
| ENSG00000C | RELT | 8.61E-01 | 9.54E-01 | 1.01E+00 | 1.95E-02 | 1.01E+00 |
| ENSG000002 | ADSL | 8.61E-01 | 9.54E-01 | 9.91E-01 | -1.36E-02 | -1.01E+00 |
| ENSG00000C | HACD3 | 8.61E-01 | 9.54E-01 | 9.93E-01 | -1.01E-02 | -1.01E+00 |
| ENSG000001 | FAN1 | 8.61E-01 | 9.54E-01 | 9.95E-01 | -7.59E-03 | -1.01E+00 |
| ENSG000002 | --- | 8.61E-01 | ? | 1.10E+00 | 1.33E-01 | 1.10E+00 |
| ENSG000002 | --- | 8.61E-01 | ? | 9.51E-01 | -7.26E-02 | -1.05E+00 |
| ENSG000001 | TRIOBP | 8.61E-01 | 9.54E-01 | 1.01E+00 | 1.07E-02 | 1.01E+00 |
| ENSG000001 | YBEY | 8.61E-01 | 9.54E-01 | 1.01E+00 | 1.16E-02 | 1.01E+00 |
| ENSG000001 | BPNT1 | 8.61E-01 | 9.54E-01 | 9.94E-01 | -8.91E-03 | -1.01E+00 |
| ENSG000001 | UNK | 8.61E-01 | 9.54E-01 | 9.91E-01 | -1.24E-02 | -1.01E+00 |
| ENSG000001 | FAM20A | 8.61E-01 | ? | 1.06E+00 | 7.92E-02 | 1.06E+00 |

| | | | | | | |
|---|---|---|---|---|---|---|
| ENSG000002 | FOXD4L5 | 8.61E-01 | ? | 9.42E-01 | -8.66E-02 | -1.06E+00 |
| ENSG000001 | EGR3 | 8.61E-01 | ? | 9.43E-01 | -8.46E-02 | -1.06E+00 |
| ENSG000001 | LOXL4 | 8.62E-01 | ? | 1.06E+00 | 8.15E-02 | 1.06E+00 |
| ENSG000002 | RPS6P25 | 8.62E-01 | ? | 1.05E+00 | 6.56E-02 | 1.05E+00 |
| ENSG000001 | C2orf42 | 8.62E-01 | 9.54E-01 | 1.01E+00 | 1.07E-02 | 1.01E+00 |
| ENSG000001 | DDX5 | 8.62E-01 | 9.54E-01 | 9.93E-01 | -9.73E-03 | -1.01E+00 |
| ENSG000002 | --- | 8.62E-01 | 9.54E-01 | 1.02E+00 | 2.22E-02 | 1.02E+00 |
| ENSG000002 | --- | 8.62E-01 | ? | 9.71E-01 | -4.23E-02 | -1.03E+00 |
| ENSG000001 | KLF7 | 8.62E-01 | 9.54E-01 | 9.94E-01 | -8.24E-03 | -1.01E+00 |
| ENSG00000C | PIR | 8.62E-01 | 9.54E-01 | 1.01E+00 | 1.89E-02 | 1.01E+00 |
| ENSG000002 | DENND10P1 | 8.62E-01 | 9.54E-01 | 9.78E-01 | -3.26E-02 | -1.02E+00 |
| ENSG000002 | --- | 8.62E-01 | ? | 9.23E-01 | -1.15E-01 | -1.08E+00 |
| ENSG000002 | PSMB8-AS1 | 8.62E-01 | ? | 1.03E+00 | 4.84E-02 | 1.03E+00 |
| ENSG00000C | TENM1 | 8.62E-01 | ? | 1.05E+00 | 6.69E-02 | 1.05E+00 |
| ENSG000001 | MMP15 | 8.62E-01 | 9.54E-01 | 1.01E+00 | 1.57E-02 | 1.01E+00 |
| ENSG000001 | ABCA5 | 8.62E-01 | 9.54E-01 | 9.90E-01 | -1.43E-02 | -1.01E+00 |
| ENSG000002 | --- | 8.62E-01 | ? | 1.05E+00 | 7.03E-02 | 1.05E+00 |
| ENSG000002 | --- | 8.62E-01 | ? | 1.08E+00 | 1.15E-01 | 1.08E+00 |
| ENSG000002 | --- | 8.62E-01 | ? | 1.07E+00 | 9.72E-02 | 1.07E+00 |
| ENSG000001 | INO80 | 8.63E-01 | 9.55E-01 | 1.00E+00 | 6.40E-03 | 1.00E+00 |
| ENSG000002 | --- | 8.63E-01 | ? | 1.08E+00 | 1.11E-01 | 1.08E+00 |
| ENSG000001 | ADCK5 | 8.63E-01 | 9.55E-01 | 9.81E-01 | -2.74E-02 | -1.02E+00 |
| ENSG000001 | GDF5 | 8.63E-01 | ? | 9.50E-01 | -7.37E-02 | -1.05E+00 |
| ENSG000001 | RRP9 | 8.63E-01 | 9.55E-01 | 1.01E+00 | 1.55E-02 | 1.01E+00 |
| ENSG000002 | LINC01341 | 8.63E-01 | ? | 9.42E-01 | -8.58E-02 | -1.06E+00 |
| ENSG00000C | MGST1 | 8.63E-01 | 9.55E-01 | 9.96E-01 | -6.17E-03 | -1.00E+00 |
| ENSG000002 | SENCR | 8.63E-01 | 9.55E-01 | 1.03E+00 | 3.64E-02 | 1.03E+00 |
| ENSG000001 | GRAP | 8.63E-01 | ? | 9.42E-01 | -8.57E-02 | -1.06E+00 |
| ENSG000001 | PRKCSH | 8.63E-01 | 9.55E-01 | 1.01E+00 | 8.89E-03 | 1.01E+00 |
| ENSG000001 | MED31 | 8.63E-01 | 9.55E-01 | 1.01E+00 | 1.36E-02 | 1.01E+00 |
| ENSG000002 | MORF4L1P1 | 8.63E-01 | 9.55E-01 | 1.01E+00 | 1.22E-02 | 1.01E+00 |
| ENSG000001 | CCDC6 | 8.64E-01 | 9.55E-01 | 9.94E-01 | -8.26E-03 | -1.01E+00 |
| ENSG000002 | --- | 8.64E-01 | 9.55E-01 | 9.83E-01 | -2.46E-02 | -1.02E+00 |
| ENSG000001 | ARHGDIA | 8.64E-01 | 9.55E-01 | 9.93E-01 | -1.01E-02 | -1.01E+00 |
| ENSG000001 | LRRC61 | 8.64E-01 | 9.55E-01 | 9.75E-01 | -3.72E-02 | -1.03E+00 |
| ENSG000001 | EXOSC3 | 8.64E-01 | 9.55E-01 | 9.92E-01 | -1.13E-02 | -1.01E+00 |
| ENSG000002 | ZEB1-AS1 | 8.64E-01 | 9.55E-01 | 1.01E+00 | 2.08E-02 | 1.01E+00 |
| ENSG000001 | HAND1 | 8.64E-01 | ? | 9.07E-01 | -1.40E-01 | -1.10E+00 |
| ENSG000001 | SPACA9 | 8.64E-01 | 9.55E-01 | 1.01E+00 | 1.48E-02 | 1.01E+00 |
| ENSG000002 | --- | 8.64E-01 | ? | 9.52E-01 | -7.04E-02 | -1.05E+00 |
| ENSG000001 | PPP1R15B | 8.64E-01 | 9.55E-01 | 9.95E-01 | -6.52E-03 | -1.00E+00 |
| ENSG000001 | CSMD2 | 8.64E-01 | ? | 1.07E+00 | 9.68E-02 | 1.07E+00 |
| ENSG000002 | --- | 8.64E-01 | ? | 9.18E-01 | -1.23E-01 | -1.09E+00 |
| ENSG000001 | BICD1 | 8.64E-01 | 9.55E-01 | 1.01E+00 | 1.18E-02 | 1.01E+00 |
| ENSG000001 | TOX-DT | 8.64E-01 | ? | 9.43E-01 | -8.45E-02 | -1.06E+00 |

| | | | | | | |
|---|---|---|---|---|---|---|
| ENSG000002 | --- | 8.65E-01 | 9.56E-01 | 1.02E+00 | 2.70E-02 | 1.02E+00 |
| ENSG000002 | CHCHD2P6 | 8.65E-01 | ? | 9.06E-01 | -1.42E-01 | -1.10E+00 |
| ENSG000001 | GRB2 | 8.65E-01 | 9.56E-01 | 1.00E+00 | 7.18E-03 | 1.00E+00 |
| ENSG000001 | FAM13C | 8.65E-01 | ? | 1.04E+00 | 6.22E-02 | 1.04E+00 |
| ENSG000001 | ZNF417 | 8.65E-01 | 9.56E-01 | 1.01E+00 | 1.55E-02 | 1.01E+00 |
| ENSG000002 | EDRF1-AS1 | 8.65E-01 | ? | 9.33E-01 | -9.93E-02 | -1.07E+00 |
| ENSG000002 | --- | 8.65E-01 | ? | 9.69E-01 | -4.56E-02 | -1.03E+00 |
| ENSG000002 | --- | 8.65E-01 | ? | 9.48E-01 | -7.73E-02 | -1.06E+00 |
| ENSG000002 | UBE2V1 | 8.65E-01 | 9.56E-01 | 9.95E-01 | -7.30E-03 | -1.01E+00 |
| ENSG000002 | C17orf113 | 8.65E-01 | ? | 9.46E-01 | -8.01E-02 | -1.06E+00 |
| ENSG000001 | TNNT2 | 8.65E-01 | 9.56E-01 | 1.03E+00 | 3.98E-02 | 1.03E+00 |
| ENSG000002 | --- | 8.65E-01 | ? | 1.11E+00 | 1.53E-01 | 1.11E+00 |
| ENSG000002 | --- | 8.65E-01 | ? | 1.05E+00 | 6.54E-02 | 1.05E+00 |
| ENSG000002 | --- | 8.66E-01 | ? | 1.07E+00 | 1.02E-01 | 1.07E+00 |
| ENSG00000C | NFYC | 8.66E-01 | 9.56E-01 | 1.01E+00 | 8.20E-03 | 1.01E+00 |
| ENSG000001 | ABT1 | 8.66E-01 | 9.56E-01 | 1.01E+00 | 1.13E-02 | 1.01E+00 |
| ENSG000001 | KRT78 | 8.66E-01 | ? | 9.39E-01 | -9.09E-02 | -1.07E+00 |
| ENSG000001 | PALM2AKAP2 | 8.66E-01 | 9.56E-01 | 9.92E-01 | -1.13E-02 | -1.01E+00 |
| ENSG000001 | C11orf54 | 8.66E-01 | 9.56E-01 | 9.90E-01 | -1.46E-02 | -1.01E+00 |
| ENSG000002 | CTC-338M12 | 8.66E-01 | 9.56E-01 | 9.72E-01 | -4.04E-02 | -1.03E+00 |
| ENSG000001 | SYCP2 | 8.66E-01 | ? | 9.61E-01 | -5.76E-02 | -1.04E+00 |
| ENSG000001 | TEAD1 | 8.66E-01 | 9.56E-01 | 9.94E-01 | -9.29E-03 | -1.01E+00 |
| ENSG000002 | LINC02966 | 8.66E-01 | 9.57E-01 | 1.02E+00 | 3.07E-02 | 1.02E+00 |
| ENSG000001 | TRIM46 | 8.66E-01 | 9.57E-01 | 9.87E-01 | -1.85E-02 | -1.01E+00 |
| ENSG000001 | AIFM3 | 8.66E-01 | 9.57E-01 | 1.02E+00 | 3.12E-02 | 1.02E+00 |
| ENSG000001 | CA5BP1 | 8.67E-01 | ? | 9.35E-01 | -9.76E-02 | -1.07E+00 |
| ENSG00000C | KCNG1 | 8.67E-01 | 9.57E-01 | 1.01E+00 | 1.91E-02 | 1.01E+00 |
| ENSG000002 | RPL22P2 | 8.67E-01 | ? | 9.28E-01 | -1.07E-01 | -1.08E+00 |
| ENSG000001 | TOB1 | 8.67E-01 | 9.57E-01 | 1.01E+00 | 1.03E-02 | 1.01E+00 |
| ENSG000001 | AHSA1 | 8.67E-01 | 9.57E-01 | 9.96E-01 | -5.82E-03 | -1.00E+00 |
| ENSG000001 | APOBR | 8.67E-01 | ? | 9.45E-01 | -8.16E-02 | -1.06E+00 |
| ENSG000001 | PFN1 | 8.67E-01 | 9.57E-01 | 1.01E+00 | 8.15E-03 | 1.01E+00 |
| ENSG000001 | LRRC51 | 8.67E-01 | 9.57E-01 | 9.85E-01 | -2.13E-02 | -1.01E+00 |
| ENSG000002 | --- | 8.67E-01 | ? | 9.59E-01 | -6.09E-02 | -1.04E+00 |
| ENSG000001 | FNDC1 | 8.67E-01 | ? | 1.08E+00 | 1.10E-01 | 1.08E+00 |
| ENSG000001 | ESRP1 | 8.67E-01 | 9.57E-01 | 9.94E-01 | -9.02E-03 | -1.01E+00 |
| ENSG000001 | TOR1A | 8.67E-01 | 9.57E-01 | 9.95E-01 | -7.32E-03 | -1.01E+00 |
| ENSG000001 | CCDC167 | 8.67E-01 | 9.57E-01 | 9.90E-01 | -1.41E-02 | -1.01E+00 |
| ENSG000001 | LINC03040 | 8.67E-01 | 9.57E-01 | 9.83E-01 | -2.41E-02 | -1.02E+00 |
| ENSG00000C | KATNIP | 8.67E-01 | 9.57E-01 | 1.01E+00 | 9.09E-03 | 1.01E+00 |
| ENSG000001 | AGBL2 | 8.67E-01 | 9.57E-01 | 1.02E+00 | 3.47E-02 | 1.02E+00 |
| ENSG000002 | --- | 8.67E-01 | ? | 1.04E+00 | 6.18E-02 | 1.04E+00 |
| ENSG000001 | BRAF | 8.67E-01 | 9.57E-01 | 1.01E+00 | 1.18E-02 | 1.01E+00 |
| ENSG000001 | PLAAT2 | 8.67E-01 | ? | 9.18E-01 | -1.24E-01 | -1.09E+00 |
| ENSG000002 | MPIG6B | 8.68E-01 | ? | 9.38E-01 | -9.22E-02 | -1.07E+00 |

| | | | | | | |
|---|---|---|---|---|---|---|
| ENSG000002 | --- | 8.68E-01 | 9.57E-01 | 1.02E+00 | 2.94E-02 | 1.02E+00 |
| ENSG000001 | ZNF418 | 8.68E-01 | 9.57E-01 | 1.02E+00 | 2.18E-02 | 1.02E+00 |
| ENSG000001 | WDR97 | 8.68E-01 | ? | 9.67E-01 | -4.85E-02 | -1.03E+00 |
| ENSG00000C | APPBP2 | 8.68E-01 | 9.58E-01 | 1.01E+00 | 7.23E-03 | 1.01E+00 |
| ENSG000002 | --- | 8.68E-01 | ? | 9.06E-01 | -1.42E-01 | -1.10E+00 |
| ENSG00000C | ARID1B | 8.68E-01 | 9.58E-01 | 9.94E-01 | -8.35E-03 | -1.01E+00 |
| ENSG000001 | ZNF18 | 8.68E-01 | 9.58E-01 | 9.90E-01 | -1.45E-02 | -1.01E+00 |
| ENSG000001 | POGLUT1 | 8.69E-01 | 9.58E-01 | 9.94E-01 | -8.18E-03 | -1.01E+00 |
| ENSG000001 | MIR31HG | 8.69E-01 | 9.58E-01 | 1.01E+00 | 1.44E-02 | 1.01E+00 |
| ENSG000002 | --- | 8.69E-01 | ? | 9.32E-01 | -1.02E-01 | -1.07E+00 |
| ENSG000001 | KANSL1L | 8.69E-01 | 9.58E-01 | 1.01E+00 | 1.82E-02 | 1.01E+00 |
| ENSG000001 | DPYSL3 | 8.69E-01 | 9.58E-01 | 1.01E+00 | 8.38E-03 | 1.01E+00 |
| ENSG000002 | --- | 8.69E-01 | ? | 1.05E+00 | 6.64E-02 | 1.05E+00 |
| ENSG000001 | PACRG | 8.69E-01 | ? | 1.08E+00 | 1.08E-01 | 1.08E+00 |
| ENSG000002 | --- | 8.69E-01 | ? | 9.36E-01 | -9.50E-02 | -1.07E+00 |
| ENSG00000C | PALM | 8.69E-01 | ? | 1.04E+00 | 5.42E-02 | 1.04E+00 |
| ENSG000002 | --- | 8.69E-01 | ? | 9.71E-01 | -4.22E-02 | -1.03E+00 |
| ENSG000002 | TRIM52-AS1 | 8.69E-01 | 9.58E-01 | 9.81E-01 | -2.71E-02 | -1.02E+00 |
| ENSG000001 | HUS1 | 8.69E-01 | 9.58E-01 | 1.01E+00 | 7.45E-03 | 1.01E+00 |
| ENSG000002 | --- | 8.69E-01 | 9.58E-01 | 9.82E-01 | -2.58E-02 | -1.02E+00 |
| ENSG000001 | SNAPC5 | 8.69E-01 | 9.58E-01 | 9.90E-01 | -1.48E-02 | -1.01E+00 |
| ENSG000001 | ZFP37 | 8.69E-01 | 9.58E-01 | 9.73E-01 | -3.95E-02 | -1.03E+00 |
| ENSG000001 | GNRHR | 8.69E-01 | ? | 1.05E+00 | 7.69E-02 | 1.05E+00 |
| ENSG000001 | KTI12 | 8.69E-01 | 9.58E-01 | 1.01E+00 | 1.43E-02 | 1.01E+00 |
| ENSG000002 | --- | 8.69E-01 | ? | 1.05E+00 | 6.52E-02 | 1.05E+00 |
| ENSG000002 | COX6A1P2 | 8.69E-01 | ? | 9.45E-01 | -8.17E-02 | -1.06E+00 |
| ENSG000002 | --- | 8.69E-01 | ? | 1.05E+00 | 6.41E-02 | 1.05E+00 |
| ENSG000001 | PEX13 | 8.70E-01 | 9.58E-01 | 1.01E+00 | 8.94E-03 | 1.01E+00 |
| ENSG000001 | PROSER2 | 8.70E-01 | 9.58E-01 | 1.01E+00 | 8.43E-03 | 1.01E+00 |
| ENSG000002 | --- | 8.70E-01 | 9.58E-01 | 9.82E-01 | -2.58E-02 | -1.02E+00 |
| ENSG000001 | PIAS4 | 8.70E-01 | 9.58E-01 | 1.01E+00 | 1.53E-02 | 1.01E+00 |
| ENSG000002 | H1-10-AS1 | 8.70E-01 | 9.58E-01 | 1.02E+00 | 2.93E-02 | 1.02E+00 |
| ENSG000001 | TTC21A | 8.70E-01 | 9.58E-01 | 9.80E-01 | -2.97E-02 | -1.02E+00 |
| ENSG000001 | DPH3 | 8.70E-01 | 9.58E-01 | 9.95E-01 | -6.53E-03 | -1.00E+00 |
| ENSG000001 | MSANTD1 | 8.70E-01 | ? | 9.43E-01 | -8.53E-02 | -1.06E+00 |
| ENSG000002 | --- | 8.70E-01 | 9.58E-01 | 9.64E-01 | -5.31E-02 | -1.04E+00 |
| ENSG000002 | --- | 8.70E-01 | ? | 1.09E+00 | 1.25E-01 | 1.09E+00 |
| ENSG000002 | LNCOG | 8.70E-01 | 9.58E-01 | 1.01E+00 | 1.04E-02 | 1.01E+00 |
| ENSG000002 | PCDHGA8 | 8.70E-01 | 9.58E-01 | 1.03E+00 | 3.88E-02 | 1.03E+00 |
| ENSG000001 | ZNF442 | 8.70E-01 | 9.58E-01 | 1.02E+00 | 3.28E-02 | 1.02E+00 |
| ENSG000001 | DAAM1 | 8.70E-01 | 9.58E-01 | 9.93E-01 | -9.43E-03 | -1.01E+00 |
| ENSG000001 | FGFBP3 | 8.70E-01 | ? | 9.49E-01 | -7.48E-02 | -1.05E+00 |
| ENSG000002 | --- | 8.70E-01 | ? | 9.14E-01 | -1.30E-01 | -1.09E+00 |
| ENSG000001 | ZC3H10 | 8.70E-01 | 9.58E-01 | 9.89E-01 | -1.55E-02 | -1.01E+00 |
| ENSG000001 | PCDHB5 | 8.70E-01 | ? | 1.05E+00 | 6.62E-02 | 1.05E+00 |

| | | | | | | | |
|---|---|---|---|---|---|---|---|
| ENSG00000C | TTC27 | 8.70E-01 | | 9.58E-01 | 9.94E-01 | -8.24E-03 | -1.01E+00 |
| ENSG000001 | ZNF22 | 8.71E-01 | | 9.58E-01 | 9.93E-01 | -9.59E-03 | -1.01E+00 |
| ENSG000001 | SMARCAL1 | 8.71E-01 | | 9.58E-01 | 9.94E-01 | -9.27E-03 | -1.01E+00 |
| ENSG000002 | NAIPP4 | 8.71E-01 | ? | | 8.93E-01 | -1.64E-01 | -1.12E+00 |
| ENSG000002 | KRT8P12 | 8.71E-01 | | 9.58E-01 | 1.02E+00 | 2.58E-02 | 1.02E+00 |
| ENSG000002 | --- | 8.71E-01 | ? | | 1.07E+00 | 9.93E-02 | 1.07E+00 |
| ENSG00000C | PLEKHH1 | 8.71E-01 | | 9.58E-01 | 1.01E+00 | 1.32E-02 | 1.01E+00 |
| ENSG000001 | NUP98 | 8.71E-01 | | 9.58E-01 | 1.00E+00 | 4.86E-03 | 1.00E+00 |
| ENSG000001 | CREB3 | 8.71E-01 | | 9.58E-01 | 9.93E-01 | -9.68E-03 | -1.01E+00 |
| ENSG000001 | EXOC8 | 8.71E-01 | | 9.58E-01 | 9.95E-01 | -7.21E-03 | -1.01E+00 |
| ENSG000002 | --- | 8.71E-01 | ? | | 9.14E-01 | -1.30E-01 | -1.09E+00 |
| ENSG00000C | PRSS21 | 8.71E-01 | ? | | 1.03E+00 | 4.81E-02 | 1.03E+00 |
| ENSG00000C | SDCCAG8 | 8.71E-01 | | 9.58E-01 | 1.01E+00 | 1.23E-02 | 1.01E+00 |
| ENSG000002 | --- | 8.71E-01 | ? | | 1.09E+00 | 1.22E-01 | 1.09E+00 |
| ENSG000002 | --- | 8.71E-01 | | 9.58E-01 | 1.02E+00 | 2.56E-02 | 1.02E+00 |
| ENSG00000C | MMP2 | 8.71E-01 | | 9.58E-01 | 9.90E-01 | -1.51E-02 | -1.01E+00 |
| ENSG000001 | ZNF629 | 8.71E-01 | | 9.58E-01 | 1.01E+00 | 7.26E-03 | 1.01E+00 |
| ENSG000001 | SNX15 | 8.71E-01 | | 9.58E-01 | 9.89E-01 | -1.66E-02 | -1.01E+00 |
| ENSG000002 | ZKSCAN7-AS | 8.72E-01 | ? | | 9.22E-01 | -1.17E-01 | -1.08E+00 |
| ENSG000002 | --- | 8.72E-01 | ? | | 1.06E+00 | 8.40E-02 | 1.06E+00 |
| ENSG000001 | PDGFRL | 8.72E-01 | ? | | 9.48E-01 | -7.70E-02 | -1.05E+00 |
| ENSG000001 | TMEM144 | 8.72E-01 | | 9.58E-01 | 9.90E-01 | -1.39E-02 | -1.01E+00 |
| ENSG000001 | FLVCR2 | 8.72E-01 | ? | | 1.04E+00 | 5.53E-02 | 1.04E+00 |
| ENSG000001 | ELL | 8.72E-01 | | 9.58E-01 | 9.90E-01 | -1.51E-02 | -1.01E+00 |
| ENSG000002 | PRRC2A | 8.72E-01 | | 9.58E-01 | 1.01E+00 | 7.92E-03 | 1.01E+00 |
| ENSG000001 | CD55 | 8.72E-01 | | 9.58E-01 | 9.89E-01 | -1.55E-02 | -1.01E+00 |
| ENSG000001 | SMCHD1 | 8.72E-01 | | 9.58E-01 | 1.01E+00 | 9.46E-03 | 1.01E+00 |
| ENSG000002 | RAB6C | 8.72E-01 | ? | | 1.06E+00 | 8.73E-02 | 1.06E+00 |
| ENSG000002 | --- | 8.72E-01 | ? | | 1.07E+00 | 9.35E-02 | 1.07E+00 |
| ENSG000002 | DOCK9-DT | 8.72E-01 | ? | | 1.04E+00 | 5.29E-02 | 1.04E+00 |
| ENSG000001 | THBS3 | 8.72E-01 | | 9.59E-01 | 1.01E+00 | 1.53E-02 | 1.01E+00 |
| ENSG000002 | PCDHGA5 | 8.73E-01 | ? | | 9.42E-01 | -8.54E-02 | -1.06E+00 |
| ENSG000002 | PDE7A-DT | 8.73E-01 | ? | | 1.04E+00 | 6.04E-02 | 1.04E+00 |
| ENSG000001 | ITGB7 | 8.73E-01 | ? | | 9.66E-01 | -4.94E-02 | -1.03E+00 |
| ENSG000001 | ANKRD13C | 8.73E-01 | | 9.59E-01 | 1.01E+00 | 1.06E-02 | 1.01E+00 |
| ENSG000001 | SLC66A1L | 8.73E-01 | | 9.59E-01 | 9.83E-01 | -2.48E-02 | -1.02E+00 |
| ENSG000002 | --- | 8.73E-01 | ? | | 1.06E+00 | 8.77E-02 | 1.06E+00 |
| ENSG000001 | TEX101 | 8.73E-01 | ? | | 1.10E+00 | 1.41E-01 | 1.10E+00 |
| ENSG000002 | TBCEL-TECTA | 8.73E-01 | ? | | 1.05E+00 | 7.60E-02 | 1.05E+00 |
| ENSG000001 | ROBO1 | 8.73E-01 | | 9.59E-01 | 1.01E+00 | 8.40E-03 | 1.01E+00 |
| ENSG000001 | EME1 | 8.73E-01 | | 9.60E-01 | 1.02E+00 | 3.54E-02 | 1.02E+00 |
| ENSG000001 | ADAMTS4 | 8.74E-01 | ? | | 1.08E+00 | 1.09E-01 | 1.08E+00 |
| ENSG000002 | RPL7AP62 | 8.74E-01 | ? | | 9.54E-01 | -6.76E-02 | -1.05E+00 |
| ENSG000002 | ADGRA3P1 | 8.74E-01 | ? | | 9.51E-01 | -7.18E-02 | -1.05E+00 |
| ENSG000001 | CLEC11A | 8.74E-01 | ? | | 1.06E+00 | 7.73E-02 | 1.06E+00 |

| | | | | | | |
|---|---|---|---|---|---|---|
| ENSG000002 | --- | 8.74E-01 | ? | 9.34E-01 | -9.88E-02 | -1.07E+00 |
| ENSG00000C | CLPTM1L | 8.74E-01 | 9.60E-01 | 9.93E-01 | -9.41E-03 | -1.01E+00 |
| ENSG000001 | SYTL5 | 8.74E-01 | ? | 9.32E-01 | -1.02E-01 | -1.07E+00 |
| ENSG000001 | PPDPF | 8.74E-01 | 9.60E-01 | 1.02E+00 | 2.83E-02 | 1.02E+00 |
| ENSG000001 | GPLD1 | 8.74E-01 | 9.60E-01 | 9.77E-01 | -3.41E-02 | -1.02E+00 |
| ENSG000001 | AKAP12 | 8.74E-01 | 9.60E-01 | 1.02E+00 | 3.41E-02 | 1.02E+00 |
| ENSG000002 | NPM1P39 | 8.74E-01 | ? | 9.75E-01 | -3.59E-02 | -1.03E+00 |
| ENSG000001 | ZNF441 | 8.74E-01 | 9.60E-01 | 9.87E-01 | -1.91E-02 | -1.01E+00 |
| ENSG000001 | ZSCAN22 | 8.74E-01 | 9.60E-01 | 1.01E+00 | 1.30E-02 | 1.01E+00 |
| ENSG000001 | FAM161B | 8.75E-01 | 9.60E-01 | 1.01E+00 | 1.50E-02 | 1.01E+00 |
| ENSG000001 | PPP1R32 | 8.75E-01 | ? | 1.03E+00 | 4.30E-02 | 1.03E+00 |
| ENSG000001 | DENND5A | 8.75E-01 | 9.60E-01 | 1.01E+00 | 7.56E-03 | 1.01E+00 |
| ENSG000001 | USP44 | 8.75E-01 | ? | 1.06E+00 | 8.24E-02 | 1.06E+00 |
| ENSG000001 | ACSBG1 | 8.75E-01 | ? | 9.54E-01 | -6.72E-02 | -1.05E+00 |
| ENSG000002 | PPP1R3B-DT | 8.75E-01 | 9.61E-01 | 1.02E+00 | 3.46E-02 | 1.02E+00 |
| ENSG000001 | MAB21L4 | 8.75E-01 | 9.61E-01 | 9.69E-01 | -4.60E-02 | -1.03E+00 |
| ENSG00000C | RBM27 | 8.75E-01 | 9.61E-01 | 9.94E-01 | -8.25E-03 | -1.01E+00 |
| ENSG000002 | --- | 8.75E-01 | ? | 1.04E+00 | 5.26E-02 | 1.04E+00 |
| ENSG00000C | DDX11 | 8.75E-01 | 9.61E-01 | 1.01E+00 | 1.81E-02 | 1.01E+00 |
| ENSG000001 | JMJD1C | 8.76E-01 | 9.61E-01 | 9.92E-01 | -1.17E-02 | -1.01E+00 |
| ENSG000001 | PARN | 8.76E-01 | 9.61E-01 | 9.95E-01 | -7.22E-03 | -1.01E+00 |
| ENSG000002 | ZMPSTE24-D | 8.76E-01 | 9.61E-01 | 9.82E-01 | -2.55E-02 | -1.02E+00 |
| ENSG000002 | --- | 8.76E-01 | 9.61E-01 | 1.02E+00 | 3.33E-02 | 1.02E+00 |
| ENSG000002 | REPIN1 | 8.76E-01 | 9.61E-01 | 1.01E+00 | 1.43E-02 | 1.01E+00 |
| ENSG000001 | CENPT | 8.76E-01 | 9.61E-01 | 1.01E+00 | 1.07E-02 | 1.01E+00 |
| ENSG000001 | --- | 8.76E-01 | ? | 9.35E-01 | -9.77E-02 | -1.07E+00 |
| ENSG000002 | --- | 8.76E-01 | ? | 1.04E+00 | 6.00E-02 | 1.04E+00 |
| ENSG000002 | --- | 8.76E-01 | ? | 1.04E+00 | 5.11E-02 | 1.04E+00 |
| ENSG000001 | MTMR12 | 8.76E-01 | 9.62E-01 | 9.95E-01 | -7.54E-03 | -1.01E+00 |
| ENSG000002 | DUT-AS1 | 8.76E-01 | ? | 9.50E-01 | -7.38E-02 | -1.05E+00 |
| ENSG000002 | LINC01411 | 8.77E-01 | ? | 9.40E-01 | -8.95E-02 | -1.06E+00 |
| ENSG000001 | SMARCAD1 | 8.77E-01 | 9.62E-01 | 1.00E+00 | 6.97E-03 | 1.00E+00 |
| ENSG000001 | CHTF18 | 8.77E-01 | 9.62E-01 | 9.84E-01 | -2.36E-02 | -1.02E+00 |
| ENSG000002 | ZNF567-DT | 8.77E-01 | 9.62E-01 | 1.03E+00 | 3.82E-02 | 1.03E+00 |
| ENSG000001 | NPRL3 | 8.77E-01 | 9.62E-01 | 9.92E-01 | -1.09E-02 | -1.01E+00 |
| ENSG000002 | --- | 8.77E-01 | 9.62E-01 | 1.02E+00 | 2.43E-02 | 1.02E+00 |
| ENSG000001 | YOD1 | 8.77E-01 | 9.62E-01 | 1.01E+00 | 1.01E-02 | 1.01E+00 |
| ENSG000001 | DUSP11 | 8.77E-01 | 9.62E-01 | 9.93E-01 | -9.83E-03 | -1.01E+00 |
| ENSG000002 | --- | 8.77E-01 | ? | 1.04E+00 | 6.13E-02 | 1.04E+00 |
| ENSG000002 | VDAC1P6 | 8.77E-01 | ? | 9.43E-01 | -8.48E-02 | -1.06E+00 |
| ENSG000002 | DPY19L2P2 | 8.77E-01 | ? | 1.05E+00 | 6.61E-02 | 1.05E+00 |
| ENSG000001 | ANAPC1 | 8.77E-01 | 9.62E-01 | 9.95E-01 | -6.98E-03 | -1.00E+00 |
| ENSG000001 | BSPRY | 8.78E-01 | 9.62E-01 | 9.83E-01 | -2.51E-02 | -1.02E+00 |
| ENSG000002 | RPL14P3 | 8.78E-01 | ? | 9.53E-01 | -6.94E-02 | -1.05E+00 |
| ENSG000001 | SPACDR | 8.78E-01 | ? | 9.62E-01 | -5.55E-02 | -1.04E+00 |

| | | | | | | | |
|---|---|---|---|---|---|---|---|
| ENSG000001 | GON7 | 8.78E-01 | | 9.62E-01 | 1.01E+00 | 1.69E-02 | 1.01E+00 |
| ENSG000001 | WNT3 | 8.78E-01 | | 9.62E-01 | 1.01E+00 | 1.57E-02 | 1.01E+00 |
| ENSG00000C | IMPG2 | 8.78E-01 | | 9.62E-01 | 9.75E-01 | -3.72E-02 | -1.03E+00 |
| ENSG000002 | LINC01203 | 8.78E-01 | ? | | 9.67E-01 | -4.78E-02 | -1.03E+00 |
| ENSG00000C | DNM2 | 8.78E-01 | | 9.62E-01 | 9.96E-01 | -6.14E-03 | -1.00E+00 |
| ENSG000001 | PPOX | 8.78E-01 | | 9.62E-01 | 1.01E+00 | 1.69E-02 | 1.01E+00 |
| ENSG000002 | PSPHP1 | 8.78E-01 | ? | | 9.22E-01 | -1.17E-01 | -1.08E+00 |
| ENSG000001 | DMRTA2 | 8.78E-01 | ? | | 9.42E-01 | -8.67E-02 | -1.06E+00 |
| ENSG000001 | SPATA12 | 8.78E-01 | ? | | 9.55E-01 | -6.59E-02 | -1.05E+00 |
| ENSG000002 | --- | 8.78E-01 | ? | | 1.04E+00 | 5.70E-02 | 1.04E+00 |
| ENSG000002 | --- | 8.78E-01 | ? | | 1.05E+00 | 7.70E-02 | 1.05E+00 |
| ENSG000001 | PRIM1 | 8.78E-01 | | 9.62E-01 | 9.78E-01 | -3.25E-02 | -1.02E+00 |
| ENSG000001 | YWHAQ | 8.78E-01 | | 9.62E-01 | 1.00E+00 | 3.95E-03 | 1.00E+00 |
| ENSG000001 | SRSF12 | 8.78E-01 | ? | | 1.03E+00 | 4.92E-02 | 1.03E+00 |
| ENSG000001 | STAR | 8.78E-01 | ? | | 9.62E-01 | -5.63E-02 | -1.04E+00 |
| ENSG000001 | C18orf32 | 8.78E-01 | | 9.62E-01 | 9.87E-01 | -1.87E-02 | -1.01E+00 |
| ENSG000001 | PDE1A | 8.78E-01 | ? | | 1.04E+00 | 5.34E-02 | 1.04E+00 |
| ENSG000001 | PCCA | 8.78E-01 | | 9.62E-01 | 9.91E-01 | -1.28E-02 | -1.01E+00 |
| ENSG000002 | TMEM242 | 8.78E-01 | | 9.62E-01 | 1.01E+00 | 8.45E-03 | 1.01E+00 |
| ENSG000002 | PROB1 | 8.78E-01 | | 9.62E-01 | 9.80E-01 | -2.95E-02 | -1.02E+00 |
| ENSG000002 | GOLPH3-DT | 8.78E-01 | | 9.62E-01 | 9.76E-01 | -3.50E-02 | -1.02E+00 |
| ENSG000001 | ZNF252P | 8.79E-01 | | 9.62E-01 | 1.04E+00 | 5.72E-02 | 1.04E+00 |
| ENSG000002 | PMS2P10 | 8.79E-01 | | 9.62E-01 | 9.77E-01 | -3.39E-02 | -1.02E+00 |
| ENSG000001 | PIWIL2 | 8.79E-01 | ? | | 9.57E-01 | -6.34E-02 | -1.04E+00 |
| ENSG000002 | --- | 8.79E-01 | ? | | 9.20E-01 | -1.20E-01 | -1.09E+00 |
| ENSG000001 | ABCB8 | 8.79E-01 | | 9.62E-01 | 9.91E-01 | -1.36E-02 | -1.01E+00 |
| ENSG000002 | SNX25P1 | 8.79E-01 | ? | | 9.20E-01 | -1.21E-01 | -1.09E+00 |
| ENSG000002 | ERV3-1 | 8.79E-01 | | 9.62E-01 | 9.85E-01 | -2.25E-02 | -1.02E+00 |
| ENSG000002 | --- | 8.79E-01 | ? | | 1.07E+00 | 9.12E-02 | 1.07E+00 |
| ENSG00000C | FYN | 8.79E-01 | | 9.62E-01 | 1.01E+00 | 8.56E-03 | 1.01E+00 |
| ENSG000001 | NUBPL | 8.79E-01 | | 9.62E-01 | 9.93E-01 | -9.60E-03 | -1.01E+00 |
| ENSG000001 | PPARD | 8.79E-01 | | 9.62E-01 | 1.01E+00 | 9.19E-03 | 1.01E+00 |
| ENSG000001 | FHIP2A | 8.79E-01 | | 9.62E-01 | 9.95E-01 | -6.91E-03 | -1.00E+00 |
| ENSG00000C | ATP6V1H | 8.79E-01 | | 9.62E-01 | 1.00E+00 | 5.67E-03 | 1.00E+00 |
| ENSG000001 | SHMT1 | 8.79E-01 | | 9.62E-01 | 9.93E-01 | -1.01E-02 | -1.01E+00 |
| ENSG000001 | PROCA1 | 8.79E-01 | ? | | 9.66E-01 | -5.06E-02 | -1.04E+00 |
| ENSG000001 | SIRPB2 | 8.79E-01 | | 9.62E-01 | 9.88E-01 | -1.71E-02 | -1.01E+00 |
| ENSG000002 | --- | 8.79E-01 | ? | | 1.09E+00 | 1.23E-01 | 1.09E+00 |
| ENSG000001 | CDC42EP2 | 8.80E-01 | | 9.62E-01 | 1.01E+00 | 1.28E-02 | 1.01E+00 |
| ENSG000001 | ENY2 | 8.80E-01 | | 9.62E-01 | 9.94E-01 | -8.15E-03 | -1.01E+00 |
| ENSG000001 | SPATA41 | 8.80E-01 | ? | | 1.05E+00 | 6.60E-02 | 1.05E+00 |
| ENSG000002 | GTF3C2-AS2 | 8.80E-01 | | 9.62E-01 | 9.86E-01 | -1.99E-02 | -1.01E+00 |
| ENSG000002 | --- | 8.80E-01 | ? | | 9.52E-01 | -7.05E-02 | -1.05E+00 |
| ENSG000001 | TADA1 | 8.80E-01 | | 9.62E-01 | 1.01E+00 | 9.77E-03 | 1.01E+00 |
| ENSG000001 | C16orf87 | 8.80E-01 | | 9.62E-01 | 1.01E+00 | 1.29E-02 | 1.01E+00 |

| | | | | | | |
|---|---|---|---|---|---|---|
| ENSG000001 | JHY | 8.80E-01 | 9.62E-01 | 1.01E+00 | 1.26E-02 | 1.01E+00 |
| ENSG000001 | C1orf198 | 8.80E-01 | 9.62E-01 | 1.01E+00 | 8.28E-03 | 1.01E+00 |
| ENSG000001 | USP6 | 8.80E-01 | ? | 1.05E+00 | 7.32E-02 | 1.05E+00 |
| ENSG000001 | ARL4C | 8.80E-01 | 9.62E-01 | 1.01E+00 | 7.56E-03 | 1.01E+00 |
| ENSG000002 | LINC02361 | 8.80E-01 | ? | 1.05E+00 | 7.26E-02 | 1.05E+00 |
| ENSG000002 | LINC02256 | 8.80E-01 | ? | 9.73E-01 | -3.93E-02 | -1.03E+00 |
| ENSG000001 | ACTL6A | 8.80E-01 | 9.63E-01 | 1.00E+00 | 6.27E-03 | 1.00E+00 |
| ENSG000001 | UCP3 | 8.80E-01 | ? | 9.67E-01 | -4.78E-02 | -1.03E+00 |
| ENSG000002 | SP2-DT | 8.80E-01 | ? | 1.03E+00 | 3.71E-02 | 1.03E+00 |
| ENSG000002 | CEBPB-AS1 | 8.80E-01 | ? | 1.05E+00 | 7.52E-02 | 1.05E+00 |
| ENSG000001 | PTPN2 | 8.81E-01 | 9.63E-01 | 9.93E-01 | -1.07E-02 | -1.01E+00 |
| ENSG00000C | FLT3LG | 8.81E-01 | ? | 1.06E+00 | 8.62E-02 | 1.06E+00 |
| ENSG000001 | C8orf34 | 8.81E-01 | ? | 1.02E+00 | 3.31E-02 | 1.02E+00 |
| ENSG000002 | EIF4EP2 | 8.81E-01 | 9.63E-01 | 1.02E+00 | 2.58E-02 | 1.02E+00 |
| ENSG000001 | DNAJC1 | 8.81E-01 | 9.63E-01 | 1.01E+00 | 8.59E-03 | 1.01E+00 |
| ENSG000001 | VPS36 | 8.81E-01 | 9.63E-01 | 9.93E-01 | -1.01E-02 | -1.01E+00 |
| ENSG000001 | CCNF | 8.81E-01 | 9.63E-01 | 1.01E+00 | 1.34E-02 | 1.01E+00 |
| ENSG00000C | FLYWCH1 | 8.81E-01 | 9.63E-01 | 1.01E+00 | 1.30E-02 | 1.01E+00 |
| ENSG000001 | ARL1 | 8.81E-01 | 9.63E-01 | 9.95E-01 | -7.40E-03 | -1.01E+00 |
| ENSG000002 | --- | 8.81E-01 | ? | 1.05E+00 | 6.44E-02 | 1.05E+00 |
| ENSG000001 | NUTM2A | 8.81E-01 | 9.63E-01 | 1.02E+00 | 2.49E-02 | 1.02E+00 |
| ENSG000001 | --- | 8.81E-01 | ? | 9.39E-01 | -9.10E-02 | -1.07E+00 |
| ENSG000002 | NECTIN3-AS1 | 8.81E-01 | ? | 9.31E-01 | -1.04E-01 | -1.07E+00 |
| ENSG000001 | NGRN | 8.81E-01 | 9.63E-01 | 9.96E-01 | -5.39E-03 | -1.00E+00 |
| ENSG000001 | GDA | 8.81E-01 | 9.63E-01 | 1.03E+00 | 4.55E-02 | 1.03E+00 |
| ENSG000002 | --- | 8.81E-01 | ? | 1.04E+00 | 5.08E-02 | 1.04E+00 |
| ENSG000002 | --- | 8.81E-01 | 9.63E-01 | 1.05E+00 | 7.01E-02 | 1.05E+00 |
| ENSG000001 | CFAP410 | 8.82E-01 | 9.63E-01 | 1.02E+00 | 2.22E-02 | 1.02E+00 |
| ENSG000001 | POU2F3 | 8.82E-01 | 9.63E-01 | 1.02E+00 | 2.99E-02 | 1.02E+00 |
| ENSG000001 | CFAP418 | 8.82E-01 | 9.63E-01 | 1.01E+00 | 1.70E-02 | 1.01E+00 |
| ENSG000001 | IFITM1 | 8.82E-01 | 9.63E-01 | 9.83E-01 | -2.54E-02 | -1.02E+00 |
| ENSG000002 | SNHG1 | 8.82E-01 | 9.63E-01 | 9.90E-01 | -1.44E-02 | -1.01E+00 |
| ENSG000001 | MRM2 | 8.82E-01 | 9.63E-01 | 1.01E+00 | 8.20E-03 | 1.01E+00 |
| ENSG000001 | ZMYM3 | 8.82E-01 | 9.63E-01 | 1.01E+00 | 9.71E-03 | 1.01E+00 |
| ENSG000002 | --- | 8.82E-01 | ? | 9.70E-01 | -4.41E-02 | -1.03E+00 |
| ENSG000001 | CEP85L | 8.82E-01 | 9.63E-01 | 9.91E-01 | -1.29E-02 | -1.01E+00 |
| ENSG000001 | PPP1CA | 8.82E-01 | 9.63E-01 | 1.01E+00 | 8.37E-03 | 1.01E+00 |
| ENSG000002 | --- | 8.82E-01 | ? | 9.64E-01 | -5.30E-02 | -1.04E+00 |
| ENSG000001 | LRRC46 | 8.82E-01 | ? | 9.70E-01 | -4.46E-02 | -1.03E+00 |
| ENSG000001 | TSACC | 8.82E-01 | ? | 9.76E-01 | -3.47E-02 | -1.02E+00 |
| ENSG000001 | MMP14 | 8.83E-01 | 9.64E-01 | 9.94E-01 | -8.68E-03 | -1.01E+00 |
| ENSG000002 | --- | 8.83E-01 | 9.64E-01 | 1.02E+00 | 2.67E-02 | 1.02E+00 |
| ENSG000001 | PON2 | 8.83E-01 | 9.64E-01 | 9.95E-01 | -7.20E-03 | -1.01E+00 |
| ENSG000001 | SFTA2 | 8.83E-01 | ? | 1.04E+00 | 5.66E-02 | 1.04E+00 |
| ENSG000002 | HAUS5 | 8.83E-01 | 9.64E-01 | 1.01E+00 | 1.45E-02 | 1.01E+00 |

| | | | | | | |
|---|---|---|---|---|---|---|
| ENSG000001 | GAS2L1 | 8.83E-01 | 9.64E-01 | 1.01E+00 | 1.28E-02 | 1.01E+00 |
| ENSG000002 | --- | 8.83E-01 | ? | 1.04E+00 | 6.07E-02 | 1.04E+00 |
| ENSG000000 | WDR7 | 8.83E-01 | 9.64E-01 | 1.01E+00 | 1.11E-02 | 1.01E+00 |
| ENSG000002 | RPL13AP7 | 8.83E-01 | ? | 9.36E-01 | -9.49E-02 | -1.07E+00 |
| ENSG000002 | --- | 8.83E-01 | 9.64E-01 | 9.77E-01 | -3.35E-02 | -1.02E+00 |
| ENSG000002 | --- | 8.83E-01 | ? | 9.28E-01 | -1.08E-01 | -1.08E+00 |
| ENSG000002 | --- | 8.83E-01 | ? | 1.06E+00 | 8.32E-02 | 1.06E+00 |
| ENSG000001 | WDR5B | 8.83E-01 | 9.64E-01 | 1.01E+00 | 1.55E-02 | 1.01E+00 |
| ENSG000002 | --- | 8.83E-01 | 9.64E-01 | 1.01E+00 | 1.68E-02 | 1.01E+00 |
| ENSG000001 | SIN3A | 8.83E-01 | 9.64E-01 | 9.96E-01 | -6.09E-03 | -1.00E+00 |
| ENSG000002 | --- | 8.83E-01 | ? | 1.03E+00 | 4.92E-02 | 1.03E+00 |
| ENSG000002 | --- | 8.83E-01 | ? | 9.52E-01 | -7.05E-02 | -1.05E+00 |
| ENSG000001 | TMEM74 | 8.83E-01 | ? | 1.05E+00 | 6.82E-02 | 1.05E+00 |
| ENSG000002 | --- | 8.83E-01 | 9.64E-01 | 1.02E+00 | 2.94E-02 | 1.02E+00 |
| ENSG000002 | NUP107-DT | 8.84E-01 | ? | 1.06E+00 | 7.83E-02 | 1.06E+00 |
| ENSG000001 | UFSP1 | 8.84E-01 | ? | 9.60E-01 | -5.93E-02 | -1.04E+00 |
| ENSG000001 | ABCC3 | 8.84E-01 | 9.64E-01 | 9.88E-01 | -1.68E-02 | -1.01E+00 |
| ENSG000002 | --- | 8.84E-01 | ? | 9.25E-01 | -1.12E-01 | -1.08E+00 |
| ENSG000001 | TRMT2B | 8.84E-01 | 9.64E-01 | 1.01E+00 | 1.32E-02 | 1.01E+00 |
| ENSG000001 | MIS18A | 8.84E-01 | 9.64E-01 | 1.01E+00 | 1.24E-02 | 1.01E+00 |
| ENSG000002 | GOLGA4-AS1 | 8.84E-01 | 9.64E-01 | 1.02E+00 | 2.27E-02 | 1.02E+00 |
| ENSG000002 | NARF-IT1 | 8.84E-01 | ? | 9.42E-01 | -8.58E-02 | -1.06E+00 |
| ENSG000001 | STIM1 | 8.84E-01 | 9.64E-01 | 9.95E-01 | -7.22E-03 | -1.01E+00 |
| ENSG000001 | RBL2 | 8.84E-01 | 9.64E-01 | 1.00E+00 | 5.83E-03 | 1.00E+00 |
| ENSG000001 | MTMR8 | 8.84E-01 | ? | 9.70E-01 | -4.45E-02 | -1.03E+00 |
| ENSG000001 | CHRNE | 8.84E-01 | ? | 9.51E-01 | -7.22E-02 | -1.05E+00 |
| ENSG000000 | FAM50A | 8.84E-01 | 9.64E-01 | 9.95E-01 | -7.33E-03 | -1.01E+00 |
| ENSG000001 | XPO4 | 8.84E-01 | 9.64E-01 | 9.95E-01 | -6.98E-03 | -1.00E+00 |
| ENSG000002 | --- | 8.84E-01 | ? | 1.05E+00 | 6.54E-02 | 1.05E+00 |
| ENSG000001 | USP4 | 8.84E-01 | 9.64E-01 | 1.00E+00 | 5.31E-03 | 1.00E+00 |
| ENSG000002 | NEURL1B | 8.84E-01 | 9.64E-01 | 9.79E-01 | -3.05E-02 | -1.02E+00 |
| ENSG000001 | SOGA1 | 8.84E-01 | 9.64E-01 | 9.94E-01 | -8.13E-03 | -1.01E+00 |
| ENSG000002 | --- | 8.85E-01 | ? | 9.52E-01 | -7.04E-02 | -1.05E+00 |
| ENSG000002 | DXO | 8.85E-01 | 9.64E-01 | 1.01E+00 | 1.29E-02 | 1.01E+00 |
| ENSG000001 | MYH9 | 8.85E-01 | 9.64E-01 | 9.97E-01 | -4.95E-03 | -1.00E+00 |
| ENSG000002 | --- | 8.85E-01 | ? | 9.53E-01 | -6.96E-02 | -1.05E+00 |
| ENSG000001 | CDH13 | 8.85E-01 | 9.64E-01 | 1.00E+00 | 4.89E-03 | 1.00E+00 |
| ENSG000000 | CD9 | 8.85E-01 | 9.64E-01 | 9.94E-01 | -8.72E-03 | -1.01E+00 |
| ENSG000001 | CENPS | 8.85E-01 | 9.64E-01 | 9.89E-01 | -1.56E-02 | -1.01E+00 |
| ENSG000001 | ZNF530 | 8.85E-01 | 9.64E-01 | 1.01E+00 | 1.57E-02 | 1.01E+00 |
| ENSG000001 | SEMA6D | 8.85E-01 | ? | 9.75E-01 | -3.70E-02 | -1.03E+00 |
| ENSG000001 | RTN4IP1 | 8.85E-01 | 9.64E-01 | 9.90E-01 | -1.40E-02 | -1.01E+00 |
| ENSG000001 | CRNKL1 | 8.85E-01 | 9.64E-01 | 9.96E-01 | -6.19E-03 | -1.00E+00 |
| ENSG000001 | AGAP1 | 8.85E-01 | 9.64E-01 | 1.01E+00 | 7.51E-03 | 1.01E+00 |
| ENSG000001 | R3HDM2 | 8.85E-01 | 9.64E-01 | 1.01E+00 | 7.78E-03 | 1.01E+00 |

| | | | | | | |
|---|---|---|---|---|---|---|
| ENSG000002 | --- | 8.85E-01 | ? | 9.25E-01 | -1.12E-01 | -1.08E+00 |
| ENSG000000 | APBA2 | 8.85E-01 | 9.64E-01 | 1.01E+00 | 7.95E-03 | 1.01E+00 |
| ENSG000001 | NCKAP5 | 8.85E-01 | 9.64E-01 | 1.01E+00 | 1.23E-02 | 1.01E+00 |
| ENSG000002 | SRD5A3-AS1 | 8.85E-01 | ? | 1.07E+00 | 9.16E-02 | 1.07E+00 |
| ENSG000001 | STAT1 | 8.85E-01 | 9.64E-01 | 9.96E-01 | -5.21E-03 | -1.00E+00 |
| ENSG000001 | GSTM4 | 8.85E-01 | 9.64E-01 | 1.01E+00 | 1.16E-02 | 1.01E+00 |
| ENSG000001 | MAP3K5 | 8.85E-01 | 9.64E-01 | 9.92E-01 | -1.23E-02 | -1.01E+00 |
| ENSG000001 | UGGT2 | 8.86E-01 | 9.64E-01 | 9.92E-01 | -1.15E-02 | -1.01E+00 |
| ENSG000002 | NBPF12 | 8.86E-01 | 9.64E-01 | 9.92E-01 | -1.12E-02 | -1.01E+00 |
| ENSG000002 | SLC12A8 | 8.86E-01 | 9.64E-01 | 9.93E-01 | -9.44E-03 | -1.01E+00 |
| ENSG000000 | SMC1B | 8.86E-01 | ? | 1.08E+00 | 1.09E-01 | 1.08E+00 |
| ENSG000001 | MYO7A | 8.86E-01 | 9.64E-01 | 1.02E+00 | 3.11E-02 | 1.02E+00 |
| ENSG000002 | CCDC15-DT | 8.86E-01 | 9.64E-01 | 1.02E+00 | 3.10E-02 | 1.02E+00 |
| ENSG000001 | PRXL2C | 8.86E-01 | 9.64E-01 | 9.94E-01 | -8.28E-03 | -1.01E+00 |
| ENSG000002 | --- | 8.86E-01 | ? | 9.45E-01 | -8.09E-02 | -1.06E+00 |
| ENSG000001 | SSPN | 8.86E-01 | 9.64E-01 | 1.02E+00 | 2.24E-02 | 1.02E+00 |
| ENSG000002 | FAM228B | 8.86E-01 | 9.64E-01 | 9.85E-01 | -2.11E-02 | -1.01E+00 |
| ENSG000001 | SAT1 | 8.86E-01 | 9.64E-01 | 9.94E-01 | -8.42E-03 | -1.01E+00 |
| ENSG000001 | DNAJC15 | 8.86E-01 | 9.64E-01 | 9.95E-01 | -7.09E-03 | -1.00E+00 |
| ENSG000001 | GGCX | 8.86E-01 | 9.64E-01 | 1.00E+00 | 6.86E-03 | 1.00E+00 |
| ENSG000001 | IL7 | 8.86E-01 | ? | 1.05E+00 | 7.71E-02 | 1.05E+00 |
| ENSG000001 | DENND2A | 8.86E-01 | 9.64E-01 | 1.03E+00 | 4.06E-02 | 1.03E+00 |
| ENSG000002 | FAM200A | 8.87E-01 | 9.64E-01 | 1.01E+00 | 9.98E-03 | 1.01E+00 |
| ENSG000002 | --- | 8.87E-01 | 9.64E-01 | 1.02E+00 | 2.65E-02 | 1.02E+00 |
| ENSG000002 | AGA-DT | 8.87E-01 | ? | 1.03E+00 | 4.09E-02 | 1.03E+00 |
| ENSG000001 | ACAP3 | 8.87E-01 | 9.65E-01 | 1.01E+00 | 1.88E-02 | 1.01E+00 |
| ENSG000002 | ZNF826P | 8.87E-01 | ? | 1.05E+00 | 7.08E-02 | 1.05E+00 |
| ENSG000001 | RAD54L2 | 8.87E-01 | 9.65E-01 | 1.00E+00 | 6.50E-03 | 1.00E+00 |
| ENSG000002 | --- | 8.87E-01 | ? | 9.29E-01 | -1.07E-01 | -1.08E+00 |
| ENSG000001 | MPL | 8.87E-01 | ? | 9.71E-01 | -4.30E-02 | -1.03E+00 |
| ENSG000002 | RASGRF2-AS | 8.87E-01 | ? | 1.05E+00 | 6.94E-02 | 1.05E+00 |
| ENSG000001 | WDR90 | 8.87E-01 | 9.65E-01 | 1.02E+00 | 2.23E-02 | 1.02E+00 |
| ENSG000001 | OSTF1 | 8.87E-01 | 9.65E-01 | 9.95E-01 | -7.57E-03 | -1.01E+00 |
| ENSG000002 | CASC8 | 8.87E-01 | ? | 9.70E-01 | -4.44E-02 | -1.03E+00 |
| ENSG000002 | C21orf140 | 8.88E-01 | ? | 1.05E+00 | 6.78E-02 | 1.05E+00 |
| ENSG000002 | --- | 8.88E-01 | ? | 1.05E+00 | 6.78E-02 | 1.05E+00 |
| ENSG000001 | SUZ12 | 8.88E-01 | 9.65E-01 | 9.95E-01 | -7.53E-03 | -1.01E+00 |
| ENSG000000 | NLRC4 | 8.88E-01 | ? | 9.52E-01 | -7.10E-02 | -1.05E+00 |
| ENSG000001 | DDX39B | 8.88E-01 | 9.65E-01 | 9.91E-01 | -1.24E-02 | -1.01E+00 |
| ENSG000002 | --- | 8.88E-01 | 9.65E-01 | 9.63E-01 | -5.40E-02 | -1.04E+00 |
| ENSG000001 | AKAP13 | 8.88E-01 | 9.65E-01 | 1.01E+00 | 1.10E-02 | 1.01E+00 |
| ENSG000001 | ACKR3 | 8.88E-01 | 9.65E-01 | 1.01E+00 | 1.31E-02 | 1.01E+00 |
| ENSG000002 | MTND4P12 | 8.88E-01 | 9.65E-01 | 9.77E-01 | -3.38E-02 | -1.02E+00 |
| ENSG000002 | HOGA1 | 8.88E-01 | ? | 1.04E+00 | 5.72E-02 | 1.04E+00 |
| ENSG000002 | TMED2-DT | 8.88E-01 | 9.65E-01 | 1.02E+00 | 2.85E-02 | 1.02E+00 |

| | | | | | | |
|---|---|---|---|---|---|---|
| ENSG000002 | C8orf44-SGK | 8.88E-01 | ? | 1.07E+00 | 1.02E-01 | 1.07E+00 |
| ENSG000001 | GCA | 8.88E-01 | 9.65E-01 | 9.91E-01 | -1.31E-02 | -1.01E+00 |
| ENSG000001 | ALG2 | 8.88E-01 | 9.65E-01 | 9.96E-01 | -5.96E-03 | -1.00E+00 |
| ENSG000002 | DPF3 | 8.88E-01 | 9.65E-01 | 1.02E+00 | 2.43E-02 | 1.02E+00 |
| ENSG00000C | PIBF1 | 8.88E-01 | 9.65E-01 | 9.94E-01 | -8.12E-03 | -1.01E+00 |
| ENSG000001 | USP9X | 8.89E-01 | 9.65E-01 | 9.95E-01 | -7.43E-03 | -1.01E+00 |
| ENSG000002 | --- | 8.89E-01 | ? | 1.05E+00 | 6.83E-02 | 1.05E+00 |
| ENSG000002 | FAM72D | 8.89E-01 | 9.65E-01 | 9.76E-01 | -3.51E-02 | -1.02E+00 |
| ENSG000002 | --- | 8.89E-01 | ? | 9.71E-01 | -4.26E-02 | -1.03E+00 |
| ENSG000002 | PANK2-AS1 | 8.89E-01 | ? | 9.76E-01 | -3.52E-02 | -1.02E+00 |
| ENSG000001 | RNF125 | 8.89E-01 | 9.66E-01 | 1.01E+00 | 1.03E-02 | 1.01E+00 |
| ENSG000002 | SEMA6A-AS1 | 8.89E-01 | ? | 1.04E+00 | 6.00E-02 | 1.04E+00 |
| ENSG000001 | FOXO4 | 8.89E-01 | 9.66E-01 | 9.92E-01 | -1.18E-02 | -1.01E+00 |
| ENSG000001 | MTARC2 | 8.89E-01 | 9.66E-01 | 9.82E-01 | -2.69E-02 | -1.02E+00 |
| ENSG000002 | --- | 8.89E-01 | ? | 9.77E-01 | -3.41E-02 | -1.02E+00 |
| ENSG000001 | LSM12 | 8.89E-01 | 9.66E-01 | 9.95E-01 | -7.41E-03 | -1.01E+00 |
| ENSG000001 | SEMA4C | 8.89E-01 | 9.66E-01 | 9.90E-01 | -1.43E-02 | -1.01E+00 |
| ENSG000002 | CDPF1P1 | 8.89E-01 | ? | 1.04E+00 | 5.84E-02 | 1.04E+00 |
| ENSG000001 | SLC15A3 | 8.90E-01 | ? | 1.05E+00 | 6.89E-02 | 1.05E+00 |
| ENSG00000C | AP4E1 | 8.90E-01 | 9.66E-01 | 9.94E-01 | -9.10E-03 | -1.01E+00 |
| ENSG000001 | WNT3A | 8.90E-01 | ? | 1.05E+00 | 6.86E-02 | 1.05E+00 |
| ENSG000002 | MRPL38 | 8.90E-01 | 9.66E-01 | 9.93E-01 | -1.00E-02 | -1.01E+00 |
| ENSG000001 | AOX1 | 8.90E-01 | 9.66E-01 | 9.95E-01 | -7.72E-03 | -1.01E+00 |
| ENSG000002 | --- | 8.90E-01 | ? | 1.02E+00 | 3.51E-02 | 1.02E+00 |
| ENSG000001 | IFT52 | 8.90E-01 | 9.66E-01 | 1.01E+00 | 9.97E-03 | 1.01E+00 |
| ENSG000001 | PRPF8 | 8.90E-01 | 9.66E-01 | 1.00E+00 | 3.67E-03 | 1.00E+00 |
| ENSG000002 | --- | 8.90E-01 | ? | 9.63E-01 | -5.38E-02 | -1.04E+00 |
| ENSG000002 | --- | 8.90E-01 | ? | 9.55E-01 | -6.65E-02 | -1.05E+00 |
| ENSG000001 | TMEM204 | 8.90E-01 | ? | 1.06E+00 | 8.43E-02 | 1.06E+00 |
| ENSG000001 | NKD2 | 8.90E-01 | ? | 1.06E+00 | 8.70E-02 | 1.06E+00 |
| ENSG000001 | FCHO2 | 8.90E-01 | 9.66E-01 | 9.93E-01 | -9.44E-03 | -1.01E+00 |
| ENSG000001 | GSDMD | 8.90E-01 | 9.66E-01 | 9.83E-01 | -2.47E-02 | -1.02E+00 |
| ENSG000001 | PSIP1 | 8.90E-01 | 9.66E-01 | 9.92E-01 | -1.10E-02 | -1.01E+00 |
| ENSG000001 | GHITM | 8.90E-01 | 9.66E-01 | 9.96E-01 | -5.17E-03 | -1.00E+00 |
| ENSG000002 | PRKXP1 | 8.90E-01 | 9.66E-01 | 9.80E-01 | -2.88E-02 | -1.02E+00 |
| ENSG000001 | SLC16A14 | 8.90E-01 | 9.66E-01 | 1.02E+00 | 3.34E-02 | 1.02E+00 |
| ENSG000002 | --- | 8.91E-01 | 9.66E-01 | 9.80E-01 | -2.86E-02 | -1.02E+00 |
| ENSG000002 | BTN2A3P | 8.91E-01 | 9.66E-01 | 1.01E+00 | 2.02E-02 | 1.01E+00 |
| ENSG000001 | FAM168B | 8.91E-01 | 9.66E-01 | 1.00E+00 | 3.33E-03 | 1.00E+00 |
| ENSG000001 | DCUN1D5 | 8.91E-01 | 9.67E-01 | 1.00E+00 | 6.28E-03 | 1.00E+00 |
| ENSG000002 | LINC01564 | 8.91E-01 | ? | 1.04E+00 | 5.43E-02 | 1.04E+00 |
| ENSG000002 | --- | 8.91E-01 | ? | 9.45E-01 | -8.17E-02 | -1.06E+00 |
| ENSG000002 | --- | 8.91E-01 | ? | 9.44E-01 | -8.29E-02 | -1.06E+00 |
| ENSG000002 | HELLPAR | 8.91E-01 | ? | 9.71E-01 | -4.23E-02 | -1.03E+00 |
| ENSG00000C | ADCK1 | 8.91E-01 | 9.67E-01 | 1.01E+00 | 1.27E-02 | 1.01E+00 |

| | | | | | | | |
|---|---|---|---|---|---|---|---|
| ENSG000002 | UBE2D3-AS1 | 8.91E-01 | | 9.67E-01 | 1.02E+00 | 2.64E-02 | 1.02E+00 |
| ENSG000002 | --- | 8.92E-01 | ? | | 9.79E-01 | -3.03E-02 | -1.02E+00 |
| ENSG000001 | ZNF665 | 8.92E-01 | | 9.67E-01 | 9.88E-01 | -1.77E-02 | -1.01E+00 |
| ENSG000001 | ROR2 | 8.92E-01 | ? | | 9.73E-01 | -3.94E-02 | -1.03E+00 |
| ENSG000001 | ZNF226 | 8.92E-01 | | 9.67E-01 | 1.01E+00 | 1.24E-02 | 1.01E+00 |
| ENSG000001 | GNB1L | 8.92E-01 | | 9.67E-01 | 1.03E+00 | 3.65E-02 | 1.03E+00 |
| ENSG000001 | ERP27 | 8.92E-01 | ? | | 9.73E-01 | -3.92E-02 | -1.03E+00 |
| ENSG000001 | ATF7IP2 | 8.92E-01 | | 9.67E-01 | 1.01E+00 | 1.61E-02 | 1.01E+00 |
| ENSG000001 | STAG2 | 8.92E-01 | | 9.67E-01 | 9.96E-01 | -6.47E-03 | -1.00E+00 |
| ENSG000001 | INTS15 | 8.92E-01 | | 9.67E-01 | 1.01E+00 | 8.62E-03 | 1.01E+00 |
| ENSG000001 | STK24 | 8.92E-01 | | 9.67E-01 | 1.00E+00 | 3.78E-03 | 1.00E+00 |
| ENSG000001 | CARMIL2 | 8.92E-01 | ? | | 1.04E+00 | 5.57E-02 | 1.04E+00 |
| ENSG000002 | ZNF879 | 8.92E-01 | | 9.67E-01 | 1.01E+00 | 1.75E-02 | 1.01E+00 |
| ENSG000001 | CLIP2 | 8.92E-01 | | 9.67E-01 | 9.95E-01 | -7.54E-03 | -1.01E+00 |
| ENSG000002 | --- | 8.92E-01 | ? | | 1.09E+00 | 1.21E-01 | 1.09E+00 |
| ENSG000001 | CCDC157 | 8.92E-01 | | 9.67E-01 | 1.02E+00 | 2.40E-02 | 1.02E+00 |
| ENSG000001 | FOXRED1 | 8.92E-01 | | 9.67E-01 | 1.01E+00 | 1.10E-02 | 1.01E+00 |
| ENSG000002 | --- | 8.92E-01 | ? | | 1.04E+00 | 5.58E-02 | 1.04E+00 |
| ENSG000001 | RADX | 8.92E-01 | | 9.67E-01 | 9.85E-01 | -2.19E-02 | -1.02E+00 |
| ENSG000001 | APH1B | 8.93E-01 | | 9.67E-01 | 9.92E-01 | -1.11E-02 | -1.01E+00 |
| ENSG000002 | TUBBP6 | 8.93E-01 | ? | | 9.53E-01 | -6.99E-02 | -1.05E+00 |
| ENSG000002 | AK4P1 | 8.93E-01 | | 9.67E-01 | 1.02E+00 | 2.72E-02 | 1.02E+00 |
| ENSG000002 | HCAR3 | 8.93E-01 | | 9.67E-01 | 1.01E+00 | 1.81E-02 | 1.01E+00 |
| ENSG000001 | EIF4BP6 | 8.93E-01 | | 9.67E-01 | 1.01E+00 | 2.13E-02 | 1.01E+00 |
| ENSG000001 | FAM98C | 8.93E-01 | | 9.67E-01 | 9.86E-01 | -2.06E-02 | -1.01E+00 |
| ENSG000001 | SPRY3 | 8.93E-01 | | 9.67E-01 | 1.01E+00 | 1.93E-02 | 1.01E+00 |
| ENSG000002 | --- | 8.93E-01 | ? | | 1.05E+00 | 7.03E-02 | 1.05E+00 |
| ENSG000001 | ATXN2L | 8.93E-01 | | 9.67E-01 | 9.96E-01 | -6.26E-03 | -1.00E+00 |
| ENSG000002 | TRMT61A-DT | 8.93E-01 | ? | | 9.70E-01 | -4.40E-02 | -1.03E+00 |
| ENSG000001 | MLANA | 8.93E-01 | | 9.67E-01 | 1.02E+00 | 3.28E-02 | 1.02E+00 |
| ENSG000001 | CNOT11 | 8.93E-01 | | 9.67E-01 | 1.00E+00 | 6.35E-03 | 1.00E+00 |
| ENSG000001 | SUB1 | 8.93E-01 | | 9.67E-01 | 1.00E+00 | 6.88E-03 | 1.00E+00 |
| ENSG000001 | REL | 8.93E-01 | | 9.67E-01 | 9.92E-01 | -1.20E-02 | -1.01E+00 |
| ENSG000001 | CORO1A | 8.93E-01 | | 9.67E-01 | 1.01E+00 | 1.61E-02 | 1.01E+00 |
| ENSG000002 | --- | 8.93E-01 | | 9.67E-01 | 1.02E+00 | 2.16E-02 | 1.02E+00 |
| ENSG000000 | FRYL | 8.93E-01 | | 9.67E-01 | 1.01E+00 | 1.05E-02 | 1.01E+00 |
| ENSG000002 | --- | 8.94E-01 | ? | | 9.49E-01 | -7.52E-02 | -1.05E+00 |
| ENSG000001 | LCE1C | 8.94E-01 | ? | | 1.03E+00 | 3.99E-02 | 1.03E+00 |
| ENSG000002 | --- | 8.94E-01 | | 9.68E-01 | 9.90E-01 | -1.40E-02 | -1.01E+00 |
| ENSG000001 | ACMSD | 8.94E-01 | ? | | 9.34E-01 | -9.85E-02 | -1.07E+00 |
| ENSG000001 | TET1 | 8.94E-01 | ? | | 1.03E+00 | 4.50E-02 | 1.03E+00 |
| ENSG000001 | TTC1 | 8.94E-01 | | 9.68E-01 | 1.00E+00 | 5.72E-03 | 1.00E+00 |
| ENSG000002 | --- | 8.94E-01 | ? | | 1.04E+00 | 6.20E-02 | 1.04E+00 |
| ENSG000001 | CAMTA1 | 8.94E-01 | | 9.68E-01 | 1.01E+00 | 8.56E-03 | 1.01E+00 |
| ENSG000001 | CEP44 | 8.94E-01 | | 9.68E-01 | 1.01E+00 | 8.89E-03 | 1.01E+00 |

| | | | | | | |
|---|---|---|---|---|---|---|
| ENSG000002 | HSPA8P1 | 8.94E-01 | ? | 9.60E-01 | -5.94E-02 | -1.04E+00 |
| ENSG000002 | RPL4P4 | 8.95E-01 | 9.68E-01 | 1.01E+00 | 1.55E-02 | 1.01E+00 |
| ENSG000002 | --- | 8.95E-01 | ? | 9.16E-01 | -1.27E-01 | -1.09E+00 |
| ENSG000002 | --- | 8.95E-01 | ? | 1.07E+00 | 9.89E-02 | 1.07E+00 |
| ENSG00000C | METTL22 | 8.95E-01 | 9.68E-01 | 9.93E-01 | -1.01E-02 | -1.01E+00 |
| ENSG000002 | --- | 8.95E-01 | ? | 1.05E+00 | 6.71E-02 | 1.05E+00 |
| ENSG000002 | ZNF865 | 8.95E-01 | 9.68E-01 | 9.92E-01 | -1.19E-02 | -1.01E+00 |
| ENSG000001 | SCFD2 | 8.95E-01 | 9.68E-01 | 1.01E+00 | 8.68E-03 | 1.01E+00 |
| ENSG000001 | IQCB1 | 8.95E-01 | 9.68E-01 | 9.93E-01 | -9.63E-03 | -1.01E+00 |
| ENSG00000C | HACE1 | 8.95E-01 | 9.68E-01 | 9.92E-01 | -1.19E-02 | -1.01E+00 |
| ENSG000001 | ANKRD45 | 8.95E-01 | ? | 9.61E-01 | -5.67E-02 | -1.04E+00 |
| ENSG000002 | --- | 8.95E-01 | ? | 1.03E+00 | 3.66E-02 | 1.03E+00 |
| ENSG000001 | CHEK2 | 8.95E-01 | 9.68E-01 | 9.93E-01 | -1.04E-02 | -1.01E+00 |
| ENSG000001 | BBS10 | 8.95E-01 | 9.68E-01 | 1.01E+00 | 8.74E-03 | 1.01E+00 |
| ENSG000001 | TGIF2 | 8.95E-01 | 9.68E-01 | 9.95E-01 | -7.73E-03 | -1.01E+00 |
| ENSG000001 | TLNRD1 | 8.95E-01 | 9.68E-01 | 9.94E-01 | -8.60E-03 | -1.01E+00 |
| ENSG000001 | PLCD4 | 8.95E-01 | 9.68E-01 | 9.84E-01 | -2.27E-02 | -1.02E+00 |
| ENSG000002 | KRT16P4 | 8.95E-01 | ? | 1.02E+00 | 3.50E-02 | 1.02E+00 |
| ENSG000002 | --- | 8.95E-01 | 9.68E-01 | 1.02E+00 | 2.48E-02 | 1.02E+00 |
| ENSG000001 | RIPK4 | 8.95E-01 | 9.68E-01 | 9.92E-01 | -1.17E-02 | -1.01E+00 |
| ENSG000001 | ADGRG2 | 8.95E-01 | ? | 9.49E-01 | -7.50E-02 | -1.05E+00 |
| ENSG000002 | --- | 8.95E-01 | ? | 9.63E-01 | -5.42E-02 | -1.04E+00 |
| ENSG000002 | --- | 8.95E-01 | 9.68E-01 | 9.94E-01 | -7.99E-03 | -1.01E+00 |
| ENSG000001 | UBE2M | 8.96E-01 | 9.68E-01 | 9.94E-01 | -9.27E-03 | -1.01E+00 |
| ENSG000002 | HSBP1 | 8.96E-01 | 9.68E-01 | 1.00E+00 | 4.65E-03 | 1.00E+00 |
| ENSG000001 | TSC2 | 8.96E-01 | 9.68E-01 | 9.95E-01 | -7.55E-03 | -1.01E+00 |
| ENSG000001 | CD109 | 8.96E-01 | 9.68E-01 | 9.96E-01 | -5.77E-03 | -1.00E+00 |
| ENSG000001 | ZNF556 | 8.96E-01 | 9.68E-01 | 1.01E+00 | 1.94E-02 | 1.01E+00 |
| ENSG000002 | UBE2D3P1 | 8.96E-01 | ? | 9.48E-01 | -7.77E-02 | -1.06E+00 |
| ENSG000001 | ME3 | 8.96E-01 | 9.68E-01 | 1.01E+00 | 2.08E-02 | 1.01E+00 |
| ENSG00000C | EPHA3 | 8.96E-01 | ? | 1.04E+00 | 6.02E-02 | 1.04E+00 |
| ENSG000001 | TFAM | 8.96E-01 | 9.68E-01 | 1.01E+00 | 7.37E-03 | 1.01E+00 |
| ENSG000002 | GGTA1 | 8.96E-01 | ? | 1.06E+00 | 9.04E-02 | 1.06E+00 |
| ENSG000002 | --- | 8.96E-01 | ? | 9.49E-01 | -7.59E-02 | -1.05E+00 |
| ENSG000002 | LINC01752 | 8.96E-01 | ? | 9.51E-01 | -7.29E-02 | -1.05E+00 |
| ENSG000001 | DEPDC7 | 8.96E-01 | 9.68E-01 | 1.01E+00 | 1.61E-02 | 1.01E+00 |
| ENSG000002 | --- | 8.96E-01 | ? | 9.73E-01 | -3.93E-02 | -1.03E+00 |
| ENSG000001 | ANXA5 | 8.96E-01 | 9.68E-01 | 1.01E+00 | 9.48E-03 | 1.01E+00 |
| ENSG000001 | ZNF664 | 8.96E-01 | 9.68E-01 | 9.96E-01 | -5.86E-03 | -1.00E+00 |
| ENSG000002 | --- | 8.96E-01 | ? | 1.07E+00 | 9.21E-02 | 1.07E+00 |
| ENSG000001 | SAP30L | 8.97E-01 | 9.68E-01 | 9.95E-01 | -6.95E-03 | -1.00E+00 |
| ENSG000001 | PRKD1 | 8.97E-01 | 9.68E-01 | 9.82E-01 | -2.58E-02 | -1.02E+00 |
| ENSG000001 | SSBP1 | 8.97E-01 | 9.68E-01 | 1.00E+00 | 5.37E-03 | 1.00E+00 |
| ENSG000001 | NSD1 | 8.97E-01 | 9.68E-01 | 1.01E+00 | 8.33E-03 | 1.01E+00 |
| ENSG00000C | CMTM1 | 8.97E-01 | ? | 1.03E+00 | 4.18E-02 | 1.03E+00 |

| | | | | | | |
|---|---|---|---|---|---|---|
| ENSG000001 | PCGF5 | 8.97E-01 | 9.68E-01 | 9.94E-01 | -8.12E-03 | -1.01E+00 |
| ENSG000002 | PRMT5-AS1 | 8.97E-01 | ? | 1.04E+00 | 5.38E-02 | 1.04E+00 |
| ENSG000001 | ZCCHC14 | 8.97E-01 | 9.68E-01 | 9.94E-01 | -8.60E-03 | -1.01E+00 |
| ENSG000002 | SNRK-AS1 | 8.97E-01 | ? | 1.04E+00 | 5.47E-02 | 1.04E+00 |
| ENSG000001 | RCC1 | 8.97E-01 | 9.68E-01 | 1.01E+00 | 7.33E-03 | 1.01E+00 |
| ENSG000002 | --- | 8.97E-01 | ? | 1.06E+00 | 8.96E-02 | 1.06E+00 |
| ENSG000002 | --- | 8.97E-01 | ? | 1.03E+00 | 4.62E-02 | 1.03E+00 |
| ENSG000001 | MED6 | 8.97E-01 | 9.68E-01 | 9.95E-01 | -6.83E-03 | -1.00E+00 |
| ENSG000001 | DCAF10 | 8.97E-01 | 9.68E-01 | 1.00E+00 | 6.22E-03 | 1.00E+00 |
| ENSG000002 | ANAPC1P2 | 8.97E-01 | 9.68E-01 | 1.02E+00 | 2.39E-02 | 1.02E+00 |
| ENSG000001 | CSNK1D | 8.97E-01 | 9.68E-01 | 9.96E-01 | -6.16E-03 | -1.00E+00 |
| ENSG000002 | --- | 8.97E-01 | ? | 1.07E+00 | 9.47E-02 | 1.07E+00 |
| ENSG000001 | RHEBL1 | 8.98E-01 | ? | 9.60E-01 | -5.89E-02 | -1.04E+00 |
| ENSG000001 | RP1 | 8.98E-01 | 9.69E-01 | 1.01E+00 | 1.98E-02 | 1.01E+00 |
| ENSG000002 | TNF | 8.98E-01 | ? | 9.69E-01 | -4.51E-02 | -1.03E+00 |
| ENSG000001 | MAP6D1 | 8.98E-01 | ? | 1.07E+00 | 9.36E-02 | 1.07E+00 |
| ENSG000001 | PLEKHB2 | 8.98E-01 | 9.69E-01 | 1.00E+00 | 6.06E-03 | 1.00E+00 |
| ENSG000001 | CYP3A5 | 8.98E-01 | 9.69E-01 | 1.01E+00 | 1.12E-02 | 1.01E+00 |
| ENSG000001 | PYGO2 | 8.98E-01 | 9.69E-01 | 1.01E+00 | 1.18E-02 | 1.01E+00 |
| ENSG000001 | STX3 | 8.98E-01 | 9.69E-01 | 1.01E+00 | 1.02E-02 | 1.01E+00 |
| ENSG000001 | ERGIC1 | 8.98E-01 | 9.69E-01 | 1.01E+00 | 8.42E-03 | 1.01E+00 |
| ENSG000001 | ARHGAP21 | 8.98E-01 | 9.69E-01 | 1.01E+00 | 7.51E-03 | 1.01E+00 |
| ENSG000001 | C19orf47 | 8.98E-01 | 9.69E-01 | 1.01E+00 | 8.72E-03 | 1.01E+00 |
| ENSG000001 | SRP19 | 8.98E-01 | 9.69E-01 | 9.96E-01 | -6.27E-03 | -1.00E+00 |
| ENSG000001 | PHAX | 8.98E-01 | 9.69E-01 | 9.96E-01 | -5.86E-03 | -1.00E+00 |
| ENSG000002 | PMS2P2 | 8.98E-01 | ? | 1.05E+00 | 6.62E-02 | 1.05E+00 |
| ENSG000001 | ARL5B | 8.98E-01 | 9.69E-01 | 1.00E+00 | 7.18E-03 | 1.00E+00 |
| ENSG000001 | KNOP1 | 8.99E-01 | 9.69E-01 | 1.00E+00 | 5.14E-03 | 1.00E+00 |
| ENSG000001 | SGPP2 | 8.99E-01 | 9.69E-01 | 1.01E+00 | 1.98E-02 | 1.01E+00 |
| ENSG000001 | FAM47E | 8.99E-01 | ? | 9.75E-01 | -3.70E-02 | -1.03E+00 |
| ENSG000001 | PIM3 | 8.99E-01 | 9.69E-01 | 1.01E+00 | 1.56E-02 | 1.01E+00 |
| ENSG000002 | WWC2-AS2 | 8.99E-01 | ? | 1.04E+00 | 5.40E-02 | 1.04E+00 |
| ENSG000001 | C19orf44 | 8.99E-01 | 9.69E-01 | 9.91E-01 | -1.29E-02 | -1.01E+00 |
| ENSG00000C | ABCB1 | 8.99E-01 | ? | 9.49E-01 | -7.52E-02 | -1.05E+00 |
| ENSG000002 | --- | 8.99E-01 | ? | 9.76E-01 | -3.50E-02 | -1.02E+00 |
| ENSG000002 | --- | 8.99E-01 | ? | 9.66E-01 | -4.92E-02 | -1.03E+00 |
| ENSG000002 | --- | 8.99E-01 | ? | 1.05E+00 | 6.50E-02 | 1.05E+00 |
| ENSG000001 | RPL7L1 | 8.99E-01 | 9.69E-01 | 9.97E-01 | -3.87E-03 | -1.00E+00 |
| ENSG00000C | NCAPH2 | 8.99E-01 | 9.69E-01 | 1.01E+00 | 1.40E-02 | 1.01E+00 |
| ENSG000002 | ST20-MTHFS | 8.99E-01 | 9.69E-01 | 1.04E+00 | 5.08E-02 | 1.04E+00 |
| ENSG000001 | TSTD2 | 8.99E-01 | 9.69E-01 | 9.95E-01 | -7.12E-03 | -1.00E+00 |
| ENSG000001 | UBE2W | 8.99E-01 | 9.69E-01 | 9.95E-01 | -7.16E-03 | -1.00E+00 |
| ENSG000001 | NT5C1B | 8.99E-01 | 9.69E-01 | 9.81E-01 | -2.77E-02 | -1.02E+00 |
| ENSG000002 | C2orf74-DT | 9.00E-01 | 9.69E-01 | 9.94E-01 | -8.82E-03 | -1.01E+00 |
| ENSG000001 | EIF4EBP1 | 9.00E-01 | 9.69E-01 | 9.93E-01 | -9.59E-03 | -1.01E+00 |

| | | | | | | |
|---|---|---|---|---|---|---|
| ENSG000002 | RPS13P2 | 9.00E-01 | ? | 9.60E-01 | -5.96E-02 | -1.04E+00 |
| ENSG000001 | CTNND2 | 9.00E-01 | ? | 1.03E+00 | 4.61E-02 | 1.03E+00 |
| ENSG000001 | COL1A2 | 9.00E-01 | 9.69E-01 | 9.84E-01 | -2.29E-02 | -1.02E+00 |
| ENSG000001 | PPP1R3D | 9.00E-01 | 9.69E-01 | 9.93E-01 | -1.05E-02 | -1.01E+00 |
| ENSG000002 | --- | 9.00E-01 | ? | 1.04E+00 | 6.10E-02 | 1.04E+00 |
| ENSG000001 | HSPA6 | 9.00E-01 | ? | 9.75E-01 | -3.64E-02 | -1.03E+00 |
| ENSG000001 | ADAM19 | 9.00E-01 | 9.69E-01 | 1.01E+00 | 1.04E-02 | 1.01E+00 |
| ENSG000002 | --- | 9.00E-01 | ? | 9.69E-01 | -4.56E-02 | -1.03E+00 |
| ENSG000001 | STAT2 | 9.00E-01 | 9.69E-01 | 1.01E+00 | 8.84E-03 | 1.01E+00 |
| ENSG000001 | SHOC2 | 9.00E-01 | 9.69E-01 | 9.95E-01 | -6.90E-03 | -1.00E+00 |
| ENSG000001 | DCTN4 | 9.00E-01 | 9.69E-01 | 1.00E+00 | 5.43E-03 | 1.00E+00 |
| ENSG000002 | --- | 9.00E-01 | ? | 9.57E-01 | -6.31E-02 | -1.04E+00 |
| ENSG000002 | LY75-CD302 | 9.00E-01 | 9.69E-01 | 9.58E-01 | -6.25E-02 | -1.04E+00 |
| ENSG000002 | --- | 9.00E-01 | 9.69E-01 | 1.02E+00 | 2.34E-02 | 1.02E+00 |
| ENSG000001 | BUD13 | 9.00E-01 | 9.69E-01 | 9.94E-01 | -9.03E-03 | -1.01E+00 |
| ENSG000001 | LSR | 9.00E-01 | 9.69E-01 | 1.01E+00 | 9.47E-03 | 1.01E+00 |
| ENSG000002 | --- | 9.00E-01 | 9.69E-01 | 9.80E-01 | -2.96E-02 | -1.02E+00 |
| ENSG000002 | KRT18P11 | 9.01E-01 | ? | 1.05E+00 | 7.30E-02 | 1.05E+00 |
| ENSG00000C | STXBP2 | 9.01E-01 | 9.69E-01 | 1.01E+00 | 9.49E-03 | 1.01E+00 |
| ENSG000001 | ETFBKMT | 9.01E-01 | 9.69E-01 | 9.85E-01 | -2.15E-02 | -1.01E+00 |
| ENSG000002 | PROX1-AS1 | 9.01E-01 | ? | 1.08E+00 | 1.08E-01 | 1.08E+00 |
| ENSG000001 | HARS1 | 9.01E-01 | 9.69E-01 | 9.96E-01 | -6.03E-03 | -1.00E+00 |
| ENSG000001 | RSAD1 | 9.01E-01 | 9.69E-01 | 1.01E+00 | 1.01E-02 | 1.01E+00 |
| ENSG000001 | RPL13AP25 | 9.01E-01 | ? | 1.04E+00 | 6.21E-02 | 1.04E+00 |
| ENSG000001 | RBP1 | 9.01E-01 | 9.69E-01 | 9.85E-01 | -2.19E-02 | -1.02E+00 |
| ENSG000002 | FLJ46284 | 9.01E-01 | ? | 9.64E-01 | -5.30E-02 | -1.04E+00 |
| ENSG000001 | SRP72 | 9.01E-01 | 9.69E-01 | 9.96E-01 | -5.92E-03 | -1.00E+00 |
| ENSG000002 | TUBG1P | 9.01E-01 | ? | 1.07E+00 | 9.35E-02 | 1.07E+00 |
| ENSG000001 | MIOS | 9.01E-01 | 9.69E-01 | 1.00E+00 | 6.67E-03 | 1.00E+00 |
| ENSG000002 | PPM1N | 9.01E-01 | ? | 1.03E+00 | 4.14E-02 | 1.03E+00 |
| ENSG000002 | --- | 9.01E-01 | ? | 1.03E+00 | 3.57E-02 | 1.03E+00 |
| ENSG000002 | TWF2-DT | 9.01E-01 | ? | 9.54E-01 | -6.86E-02 | -1.05E+00 |
| ENSG000002 | SPAG5-AS1 | 9.01E-01 | 9.70E-01 | 1.02E+00 | 2.66E-02 | 1.02E+00 |
| ENSG000002 | --- | 9.01E-01 | ? | 9.48E-01 | -7.66E-02 | -1.05E+00 |
| ENSG000002 | YWHAEP1 | 9.01E-01 | ? | 1.04E+00 | 5.93E-02 | 1.04E+00 |
| ENSG000001 | NSD3 | 9.01E-01 | 9.70E-01 | 9.95E-01 | -6.82E-03 | -1.00E+00 |
| ENSG000001 | CHRNB2 | 9.01E-01 | 9.70E-01 | 1.02E+00 | 2.93E-02 | 1.02E+00 |
| ENSG000001 | VSIG10L | 9.01E-01 | 9.70E-01 | 9.85E-01 | -2.25E-02 | -1.02E+00 |
| ENSG000002 | --- | 9.01E-01 | ? | 1.05E+00 | 7.56E-02 | 1.05E+00 |
| ENSG000001 | CHST12 | 9.02E-01 | 9.70E-01 | 9.93E-01 | -9.46E-03 | -1.01E+00 |
| ENSG000002 | LINC00513 | 9.02E-01 | ? | 1.04E+00 | 5.18E-02 | 1.04E+00 |
| ENSG000001 | MAN1A2 | 9.02E-01 | 9.70E-01 | 9.96E-01 | -6.21E-03 | -1.00E+00 |
| ENSG000001 | GUSB | 9.02E-01 | 9.70E-01 | 9.94E-01 | -9.15E-03 | -1.01E+00 |
| ENSG000002 | PTENP1 | 9.02E-01 | ? | 9.70E-01 | -4.45E-02 | -1.03E+00 |
| ENSG000001 | CYP4F3 | 9.02E-01 | ? | 1.05E+00 | 6.45E-02 | 1.05E+00 |

| | | | | | | |
|---|---|---|---|---|---|---|
| ENSG000001 | ABCA3 | 9.02E-01 | 9.70E-01 | 1.02E+00 | 2.53E-02 | 1.02E+00 |
| ENSG000002 | MT-TV | 9.02E-01 | ? | 1.07E+00 | 9.32E-02 | 1.07E+00 |
| ENSG000001 | ICAM2 | 9.02E-01 | ? | 9.67E-01 | -4.90E-02 | -1.03E+00 |
| ENSG000001 | AGMO | 9.02E-01 | 9.70E-01 | 1.02E+00 | 2.86E-02 | 1.02E+00 |
| ENSG000002 | LCDR | 9.02E-01 | ? | 1.03E+00 | 3.67E-02 | 1.03E+00 |
| ENSG000002 | CDC37L1-DT | 9.02E-01 | ? | 9.40E-01 | -8.99E-02 | -1.06E+00 |
| ENSG000001 | MDGA1 | 9.02E-01 | ? | 9.77E-01 | -3.34E-02 | -1.02E+00 |
| ENSG000002 | SKIC2 | 9.02E-01 | 9.70E-01 | 9.93E-01 | -9.70E-03 | -1.01E+00 |
| ENSG000001 | DIDO1 | 9.02E-01 | 9.70E-01 | 1.00E+00 | 4.89E-03 | 1.00E+00 |
| ENSG000002 | LINC00632 | 9.02E-01 | 9.70E-01 | 1.02E+00 | 2.87E-02 | 1.02E+00 |
| ENSG000002 | --- | 9.02E-01 | 9.70E-01 | 1.02E+00 | 2.16E-02 | 1.02E+00 |
| ENSG000001 | IGDCC4 | 9.02E-01 | ? | 9.76E-01 | -3.46E-02 | -1.02E+00 |
| ENSG000001 | ARFIP1 | 9.02E-01 | 9.70E-01 | 1.00E+00 | 4.74E-03 | 1.00E+00 |
| ENSG000001 | CCDC174 | 9.02E-01 | 9.70E-01 | 1.01E+00 | 7.47E-03 | 1.01E+00 |
| ENSG000001 | ENGASE | 9.03E-01 | 9.70E-01 | 1.01E+00 | 1.22E-02 | 1.01E+00 |
| ENSG000002 | KRT18P3 | 9.03E-01 | ? | 1.07E+00 | 1.03E-01 | 1.07E+00 |
| ENSG000001 | TMC2 | 9.03E-01 | ? | 9.41E-01 | -8.76E-02 | -1.06E+00 |
| ENSG000002 | --- | 9.03E-01 | ? | 1.05E+00 | 7.67E-02 | 1.05E+00 |
| ENSG000002 | NBPF10 | 9.03E-01 | 9.70E-01 | 1.01E+00 | 1.33E-02 | 1.01E+00 |
| ENSG000001 | PSG6 | 9.03E-01 | ? | 1.06E+00 | 7.91E-02 | 1.06E+00 |
| ENSG000001 | LMOD3 | 9.03E-01 | ? | 1.03E+00 | 4.26E-02 | 1.03E+00 |
| ENSG000001 | SH3BP5L | 9.03E-01 | 9.70E-01 | 1.01E+00 | 7.87E-03 | 1.01E+00 |
| ENSG000002 | --- | 9.03E-01 | 9.70E-01 | 1.01E+00 | 1.18E-02 | 1.01E+00 |
| ENSG000001 | STK32B | 9.03E-01 | 9.70E-01 | 9.81E-01 | -2.75E-02 | -1.02E+00 |
| ENSG000001 | LRRC28 | 9.03E-01 | 9.70E-01 | 9.94E-01 | -8.61E-03 | -1.01E+00 |
| ENSG000002 | SMG1P7 | 9.03E-01 | 9.70E-01 | 1.02E+00 | 2.30E-02 | 1.02E+00 |
| ENSG000001 | CYP2S1 | 9.03E-01 | 9.70E-01 | 9.93E-01 | -9.48E-03 | -1.01E+00 |
| ENSG000001 | FRS3 | 9.03E-01 | 9.70E-01 | 9.87E-01 | -1.89E-02 | -1.01E+00 |
| ENSG00000C | PFKP | 9.03E-01 | 9.70E-01 | 9.96E-01 | -6.39E-03 | -1.00E+00 |
| ENSG000002 | --- | 9.03E-01 | 9.70E-01 | 1.02E+00 | 2.70E-02 | 1.02E+00 |
| ENSG000001 | KCNK7 | 9.03E-01 | ? | 9.39E-01 | -9.08E-02 | -1.06E+00 |
| ENSG000002 | --- | 9.03E-01 | 9.70E-01 | 9.84E-01 | -2.36E-02 | -1.02E+00 |
| ENSG000002 | OSER1-DT | 9.03E-01 | 9.70E-01 | 9.84E-01 | -2.26E-02 | -1.02E+00 |
| ENSG000001 | CLCA2 | 9.04E-01 | 9.70E-01 | 1.01E+00 | 9.30E-03 | 1.01E+00 |
| ENSG00000C | PTPN3 | 9.04E-01 | 9.70E-01 | 1.00E+00 | 4.86E-03 | 1.00E+00 |
| ENSG000001 | MTMR6 | 9.04E-01 | 9.70E-01 | 9.95E-01 | -7.75E-03 | -1.01E+00 |
| ENSG000001 | SPOCK2 | 9.04E-01 | ? | 9.64E-01 | -5.35E-02 | -1.04E+00 |
| ENSG000001 | ZFP3 | 9.04E-01 | 9.70E-01 | 1.01E+00 | 2.00E-02 | 1.01E+00 |
| ENSG000002 | INAFM1 | 9.04E-01 | 9.70E-01 | 9.86E-01 | -2.11E-02 | -1.01E+00 |
| ENSG000002 | NSUN5P2 | 9.04E-01 | 9.70E-01 | 1.01E+00 | 1.87E-02 | 1.01E+00 |
| ENSG000001 | CRYBG1 | 9.04E-01 | 9.70E-01 | 1.00E+00 | 5.56E-03 | 1.00E+00 |
| ENSG000001 | PDE1B | 9.04E-01 | ? | 9.41E-01 | -8.71E-02 | -1.06E+00 |
| ENSG00000C | IZUMO4 | 9.04E-01 | 9.70E-01 | 1.02E+00 | 2.52E-02 | 1.02E+00 |
| ENSG00000C | ALS2 | 9.04E-01 | 9.70E-01 | 9.96E-01 | -5.84E-03 | -1.00E+00 |
| ENSG000001 | DUSP1 | 9.04E-01 | 9.70E-01 | 9.82E-01 | -2.61E-02 | -1.02E+00 |

| | | | | | | |
|---|---|---|---|---|---|---|
| ENSG00000C | NEDD4 | 9.04E-01 | 9.70E-01 | 1.01E+00 | 8.36E-03 | 1.01E+00 |
| ENSG000002 | --- | 9.04E-01 | ? | 1.05E+00 | 7.66E-02 | 1.05E+00 |
| ENSG000001 | NECAB1 | 9.04E-01 | ? | 1.04E+00 | 6.27E-02 | 1.04E+00 |
| ENSG00000C | SLC9A7 | 9.04E-01 | 9.70E-01 | 9.95E-01 | -7.49E-03 | -1.01E+00 |
| ENSG000001 | SLC7A5 | 9.04E-01 | 9.70E-01 | 9.92E-01 | -1.20E-02 | -1.01E+00 |
| ENSG00000C | MARK3 | 9.04E-01 | 9.70E-01 | 1.00E+00 | 5.40E-03 | 1.00E+00 |
| ENSG000001 | CACHD1 | 9.04E-01 | 9.70E-01 | 1.01E+00 | 9.04E-03 | 1.01E+00 |
| ENSG00000C | VPS35 | 9.04E-01 | 9.70E-01 | 1.00E+00 | 3.73E-03 | 1.00E+00 |
| ENSG000001 | VPS13C | 9.04E-01 | 9.70E-01 | 1.01E+00 | 1.00E-02 | 1.01E+00 |
| ENSG000001 | PLOD3 | 9.04E-01 | 9.70E-01 | 9.96E-01 | -6.13E-03 | -1.00E+00 |
| ENSG000001 | BRF1 | 9.05E-01 | 9.70E-01 | 9.95E-01 | -7.71E-03 | -1.01E+00 |
| ENSG000001 | ZBTB20 | 9.05E-01 | 9.70E-01 | 9.80E-01 | -2.89E-02 | -1.02E+00 |
| ENSG00000C | ISOC1 | 9.05E-01 | 9.70E-01 | 1.01E+00 | 8.53E-03 | 1.01E+00 |
| ENSG000002 | --- | 9.05E-01 | ? | 9.59E-01 | -6.05E-02 | -1.04E+00 |
| ENSG000001 | IL12A | 9.05E-01 | ? | 9.76E-01 | -3.52E-02 | -1.02E+00 |
| ENSG00000C | RNF126 | 9.05E-01 | 9.70E-01 | 1.01E+00 | 1.24E-02 | 1.01E+00 |
| ENSG000001 | EZH1 | 9.05E-01 | 9.70E-01 | 9.95E-01 | -7.50E-03 | -1.01E+00 |
| ENSG000001 | JAM3 | 9.05E-01 | 9.70E-01 | 1.01E+00 | 2.01E-02 | 1.01E+00 |
| ENSG000001 | CD8A | 9.05E-01 | ? | 1.02E+00 | 3.28E-02 | 1.02E+00 |
| ENSG000002 | KDM7A-DT | 9.06E-01 | 9.70E-01 | 9.90E-01 | -1.51E-02 | -1.01E+00 |
| ENSG000002 | NCLP1 | 9.06E-01 | ? | 9.65E-01 | -5.12E-02 | -1.04E+00 |
| ENSG000001 | ESYT2 | 9.06E-01 | 9.71E-01 | 9.97E-01 | -4.05E-03 | -1.00E+00 |
| ENSG000001 | LDLRAD4 | 9.06E-01 | ? | 1.04E+00 | 5.64E-02 | 1.04E+00 |
| ENSG000001 | TCTA | 9.06E-01 | 9.71E-01 | 9.95E-01 | -7.94E-03 | -1.01E+00 |
| ENSG000001 | MAGI1 | 9.06E-01 | 9.71E-01 | 1.01E+00 | 8.71E-03 | 1.01E+00 |
| ENSG000002 | --- | 9.06E-01 | ? | 9.37E-01 | -9.44E-02 | -1.07E+00 |
| ENSG000001 | H19 | 9.06E-01 | 9.71E-01 | 1.01E+00 | 1.41E-02 | 1.01E+00 |
| ENSG000001 | ZNF649 | 9.07E-01 | 9.71E-01 | 9.94E-01 | -8.54E-03 | -1.01E+00 |
| ENSG000002 | BMS1P4 | 9.07E-01 | 9.71E-01 | 9.62E-01 | -5.61E-02 | -1.04E+00 |
| ENSG000001 | ADPRH | 9.07E-01 | 9.71E-01 | 9.92E-01 | -1.13E-02 | -1.01E+00 |
| ENSG000002 | PSMD10P2 | 9.07E-01 | ? | 1.03E+00 | 3.74E-02 | 1.03E+00 |
| ENSG000002 | --- | 9.07E-01 | ? | 1.04E+00 | 5.76E-02 | 1.04E+00 |
| ENSG000002 | ARHGEF4-AS | 9.07E-01 | ? | 9.53E-01 | -6.90E-02 | -1.05E+00 |
| ENSG000002 | FAM156A | 9.07E-01 | 9.71E-01 | 9.90E-01 | -1.50E-02 | -1.01E+00 |
| ENSG000002 | --- | 9.07E-01 | ? | 1.06E+00 | 8.72E-02 | 1.06E+00 |
| ENSG000001 | MGAT5B | 9.07E-01 | 9.71E-01 | 1.01E+00 | 1.68E-02 | 1.01E+00 |
| ENSG00000C | MKRN2 | 9.07E-01 | 9.71E-01 | 1.00E+00 | 5.06E-03 | 1.00E+00 |
| ENSG000001 | PTPN9 | 9.07E-01 | 9.71E-01 | 9.97E-01 | -4.95E-03 | -1.00E+00 |
| ENSG000001 | PLAGL1 | 9.07E-01 | 9.71E-01 | 9.94E-01 | -8.76E-03 | -1.01E+00 |
| ENSG000001 | GPR176 | 9.07E-01 | 9.71E-01 | 1.03E+00 | 3.94E-02 | 1.03E+00 |
| ENSG000001 | LAX1 | 9.07E-01 | ? | 9.41E-01 | -8.71E-02 | -1.06E+00 |
| ENSG000001 | MYCBPAP | 9.07E-01 | ? | 9.81E-01 | -2.72E-02 | -1.02E+00 |
| ENSG000001 | RAC1 | 9.07E-01 | 9.71E-01 | 9.98E-01 | -3.00E-03 | -1.00E+00 |
| ENSG000002 | MIF4GD-DT | 9.07E-01 | ? | 9.76E-01 | -3.57E-02 | -1.03E+00 |
| ENSG000001 | PARP11 | 9.07E-01 | 9.71E-01 | 9.94E-01 | -8.52E-03 | -1.01E+00 |

| | | | | | | |
|---|---|---|---|---|---|---|
| ENSG000002 | KCNJ2-AS1 | 9.07E-01 | ? | 9.62E-01 | -5.52E-02 | -1.04E+00 |
| ENSG000001 | PPP6R3 | 9.07E-01 | 9.71E-01 | 1.00E+00 | 5.26E-03 | 1.00E+00 |
| ENSG000001 | GPR35 | 9.07E-01 | ? | 9.69E-01 | -4.56E-02 | -1.03E+00 |
| ENSG00000C | KAT6A | 9.07E-01 | 9.71E-01 | 9.95E-01 | -6.58E-03 | -1.00E+00 |
| ENSG000001 | PRR13P5 | 9.07E-01 | ? | 1.02E+00 | 3.56E-02 | 1.02E+00 |
| ENSG000002 | --- | 9.08E-01 | ? | 9.81E-01 | -2.83E-02 | -1.02E+00 |
| ENSG000001 | FZD4 | 9.08E-01 | 9.71E-01 | 9.92E-01 | -1.16E-02 | -1.01E+00 |
| ENSG000001 | PLXNB3 | 9.08E-01 | 9.71E-01 | 1.01E+00 | 1.50E-02 | 1.01E+00 |
| ENSG000002 | --- | 9.08E-01 | ? | 1.06E+00 | 8.80E-02 | 1.06E+00 |
| ENSG000001 | TMEM41A | 9.08E-01 | 9.71E-01 | 1.00E+00 | 6.25E-03 | 1.00E+00 |
| ENSG000002 | --- | 9.08E-01 | 9.71E-01 | 9.82E-01 | -2.59E-02 | -1.02E+00 |
| ENSG000002 | SIPA1L1-AS1 | 9.08E-01 | ? | 1.03E+00 | 4.11E-02 | 1.03E+00 |
| ENSG000001 | CACNA2D3 | 9.08E-01 | ? | 1.03E+00 | 4.87E-02 | 1.03E+00 |
| ENSG000002 | --- | 9.08E-01 | ? | 9.01E-01 | -1.50E-01 | -1.11E+00 |
| ENSG000002 | --- | 9.08E-01 | ? | 1.05E+00 | 7.64E-02 | 1.05E+00 |
| ENSG000001 | TYRP1 | 9.08E-01 | ? | 1.05E+00 | 7.02E-02 | 1.05E+00 |
| ENSG000001 | CLN5 | 9.08E-01 | 9.71E-01 | 9.97E-01 | -4.88E-03 | -1.00E+00 |
| ENSG000002 | H2BC15 | 9.08E-01 | 9.71E-01 | 1.01E+00 | 1.74E-02 | 1.01E+00 |
| ENSG000001 | UBE2E2 | 9.08E-01 | 9.71E-01 | 9.96E-01 | -6.27E-03 | -1.00E+00 |
| ENSG000001 | FLOT1 | 9.08E-01 | 9.71E-01 | 9.92E-01 | -1.10E-02 | -1.01E+00 |
| ENSG000002 | --- | 9.08E-01 | ? | 1.04E+00 | 5.43E-02 | 1.04E+00 |
| ENSG000001 | UBE2J2 | 9.08E-01 | 9.72E-01 | 1.01E+00 | 8.99E-03 | 1.01E+00 |
| ENSG00000C | SNCAIP | 9.08E-01 | ? | 9.55E-01 | -6.66E-02 | -1.05E+00 |
| ENSG000001 | NEURL2 | 9.09E-01 | ? | 9.65E-01 | -5.09E-02 | -1.04E+00 |
| ENSG000002 | LINC00665 | 9.09E-01 | 9.72E-01 | 1.01E+00 | 8.10E-03 | 1.01E+00 |
| ENSG000001 | HABP2 | 9.09E-01 | ? | 9.60E-01 | -5.91E-02 | -1.04E+00 |
| ENSG000001 | CDC42BPA | 9.09E-01 | 9.72E-01 | 9.95E-01 | -6.58E-03 | -1.00E+00 |
| ENSG000001 | SLC8A2 | 9.09E-01 | ? | 9.77E-01 | -3.40E-02 | -1.02E+00 |
| ENSG000002 | --- | 9.09E-01 | 9.72E-01 | 9.81E-01 | -2.72E-02 | -1.02E+00 |
| ENSG000001 | FBXL14 | 9.09E-01 | 9.72E-01 | 1.01E+00 | 1.68E-02 | 1.01E+00 |
| ENSG000002 | C4A | 9.09E-01 | 9.72E-01 | 9.85E-01 | -2.18E-02 | -1.02E+00 |
| ENSG000002 | --- | 9.09E-01 | ? | 9.32E-01 | -1.02E-01 | -1.07E+00 |
| ENSG00000C | SEPHS1 | 9.09E-01 | 9.72E-01 | 1.00E+00 | 4.89E-03 | 1.00E+00 |
| ENSG000002 | --- | 9.09E-01 | ? | 1.03E+00 | 3.94E-02 | 1.03E+00 |
| ENSG000002 | --- | 9.09E-01 | ? | 1.04E+00 | 5.88E-02 | 1.04E+00 |
| ENSG000002 | RPL21P119 | 9.09E-01 | ? | 1.07E+00 | 9.99E-02 | 1.07E+00 |
| ENSG000002 | --- | 9.09E-01 | ? | 9.77E-01 | -3.31E-02 | -1.02E+00 |
| ENSG000002 | SOX2-OT | 9.09E-01 | ? | 1.03E+00 | 4.21E-02 | 1.03E+00 |
| ENSG000002 | MICOS10-NB | 9.09E-01 | ? | 1.09E+00 | 1.21E-01 | 1.09E+00 |
| ENSG000001 | ZBED10P | 9.09E-01 | 9.72E-01 | 9.87E-01 | -1.87E-02 | -1.01E+00 |
| ENSG000001 | SETDB2 | 9.10E-01 | 9.72E-01 | 1.01E+00 | 7.51E-03 | 1.01E+00 |
| ENSG000001 | AJAP1 | 9.10E-01 | ? | 1.04E+00 | 5.57E-02 | 1.04E+00 |
| ENSG000002 | LIME1 | 9.10E-01 | 9.72E-01 | 9.84E-01 | -2.28E-02 | -1.02E+00 |
| ENSG00000C | TRAPPC6A | 9.10E-01 | 9.72E-01 | 1.02E+00 | 2.59E-02 | 1.02E+00 |
| ENSG000002 | LINC01426 | 9.10E-01 | ? | 9.81E-01 | -2.77E-02 | -1.02E+00 |

| | | | | | | |
|---|---|---|---|---|---|---|
| ENSG000001 | CLK3 | 9.10E-01 | 9.72E-01 | 9.96E-01 | -6.07E-03 | -1.00E+00 |
| ENSG000001 | IRX2 | 9.10E-01 | 9.72E-01 | 9.95E-01 | -6.55E-03 | -1.00E+00 |
| ENSG000001 | RAB19 | 9.10E-01 | ? | 9.71E-01 | -4.30E-02 | -1.03E+00 |
| ENSG000002 | PFN1P6 | 9.10E-01 | ? | 1.02E+00 | 3.53E-02 | 1.02E+00 |
| ENSG000001 | ITGA10 | 9.10E-01 | 9.72E-01 | 9.84E-01 | -2.29E-02 | -1.02E+00 |
| ENSG000001 | LRRK2 | 9.10E-01 | 9.72E-01 | 1.02E+00 | 2.53E-02 | 1.02E+00 |
| ENSG000002 | PLEKHM1P1 | 9.10E-01 | ? | 1.10E+00 | 1.41E-01 | 1.10E+00 |
| ENSG000001 | BASP1 | 9.10E-01 | 9.73E-01 | 9.95E-01 | -7.06E-03 | -1.00E+00 |
| ENSG000002 | --- | 9.11E-01 | 9.73E-01 | 9.78E-01 | -3.20E-02 | -1.02E+00 |
| ENSG000001 | PPIB | 9.11E-01 | 9.73E-01 | 9.97E-01 | -4.05E-03 | -1.00E+00 |
| ENSG000001 | EIF2B4 | 9.11E-01 | 9.73E-01 | 9.96E-01 | -5.76E-03 | -1.00E+00 |
| ENSG000002 | ITSN1 | 9.11E-01 | 9.73E-01 | 9.95E-01 | -6.84E-03 | -1.00E+00 |
| ENSG000001 | GNAL | 9.11E-01 | 9.73E-01 | 1.01E+00 | 7.81E-03 | 1.01E+00 |
| ENSG000002 | ASH1L-AS1 | 9.11E-01 | ? | 9.81E-01 | -2.84E-02 | -1.02E+00 |
| ENSG000001 | --- | 9.11E-01 | 9.73E-01 | 1.02E+00 | 2.58E-02 | 1.02E+00 |
| ENSG000002 | --- | 9.11E-01 | ? | 9.67E-01 | -4.90E-02 | -1.03E+00 |
| ENSG000001 | MTX3 | 9.11E-01 | 9.73E-01 | 1.01E+00 | 7.28E-03 | 1.01E+00 |
| ENSG000002 | CDH13-AS2 | 9.11E-01 | 9.73E-01 | 9.96E-01 | -6.27E-03 | -1.00E+00 |
| ENSG000001 | NFYB | 9.11E-01 | 9.73E-01 | 9.94E-01 | -8.42E-03 | -1.01E+00 |
| ENSG000002 | --- | 9.11E-01 | ? | 9.68E-01 | -4.64E-02 | -1.03E+00 |
| ENSG000002 | SDHAP1 | 9.11E-01 | 9.73E-01 | 1.02E+00 | 2.22E-02 | 1.02E+00 |
| ENSG000001 | ENAH | 9.11E-01 | 9.73E-01 | 1.00E+00 | 6.12E-03 | 1.00E+00 |
| ENSG000001 | MLPH | 9.11E-01 | 9.73E-01 | 9.90E-01 | -1.38E-02 | -1.01E+00 |
| ENSG000001 | ZFP82 | 9.11E-01 | 9.73E-01 | 1.01E+00 | 1.05E-02 | 1.01E+00 |
| ENSG000002 | URB1-AS1 | 9.11E-01 | 9.73E-01 | 9.87E-01 | -1.93E-02 | -1.01E+00 |
| ENSG000001 | KCNS2 | 9.12E-01 | ? | 9.66E-01 | -5.05E-02 | -1.04E+00 |
| ENSG000002 | PNMA2 | 9.12E-01 | ? | 1.04E+00 | 6.25E-02 | 1.04E+00 |
| ENSG000001 | RALGPS2 | 9.12E-01 | 9.73E-01 | 9.94E-01 | -7.98E-03 | -1.01E+00 |
| ENSG000001 | DFFA | 9.12E-01 | 9.73E-01 | 1.00E+00 | 3.99E-03 | 1.00E+00 |
| ENSG000002 | BIVM-ERCC5 | 9.12E-01 | 9.73E-01 | 9.75E-01 | -3.59E-02 | -1.03E+00 |
| ENSG000001 | HUNK | 9.12E-01 | ? | 1.05E+00 | 6.41E-02 | 1.05E+00 |
| ENSG000002 | MIR137HG | 9.12E-01 | ? | 9.77E-01 | -3.34E-02 | -1.02E+00 |
| ENSG000001 | GRM8 | 9.12E-01 | ? | 1.06E+00 | 8.11E-02 | 1.06E+00 |
| ENSG000001 | KNSTRN | 9.12E-01 | 9.73E-01 | 1.01E+00 | 7.73E-03 | 1.01E+00 |
| ENSG000000 | ATP11A | 9.12E-01 | 9.73E-01 | 1.00E+00 | 5.18E-03 | 1.00E+00 |
| ENSG000001 | SPATA1 | 9.13E-01 | ? | 9.59E-01 | -6.07E-02 | -1.04E+00 |
| ENSG000002 | --- | 9.13E-01 | 9.73E-01 | 9.84E-01 | -2.28E-02 | -1.02E+00 |
| ENSG000001 | KAT6B | 9.13E-01 | 9.73E-01 | 9.95E-01 | -7.30E-03 | -1.01E+00 |
| ENSG000000 | GBA2 | 9.13E-01 | 9.73E-01 | 1.00E+00 | 6.94E-03 | 1.00E+00 |
| ENSG000001 | KLHDC1 | 9.13E-01 | ? | 9.79E-01 | -3.04E-02 | -1.02E+00 |
| ENSG000002 | SUGCT-AS1 | 9.13E-01 | ? | 9.55E-01 | -6.59E-02 | -1.05E+00 |
| ENSG000002 | WARS2-AS1 | 9.13E-01 | 9.74E-01 | 1.00E+00 | 6.20E-03 | 1.00E+00 |
| ENSG000002 | --- | 9.13E-01 | ? | 1.03E+00 | 3.85E-02 | 1.03E+00 |
| ENSG000001 | DCLRE1B | 9.13E-01 | 9.74E-01 | 1.01E+00 | 8.15E-03 | 1.01E+00 |
| ENSG000002 | HMGB1P3 | 9.13E-01 | ? | 1.05E+00 | 6.76E-02 | 1.05E+00 |

| | | | | | | |
|---|---|---|---|---|---|---|
| ENSG00000C | SELENOO | 9.13E-01 | 9.74E-01 | 1.01E+00 | 1.04E-02 | 1.01E+00 |
| ENSG000001 | MTRFR | 9.13E-01 | 9.74E-01 | 9.95E-01 | -6.69E-03 | -1.00E+00 |
| ENSG000002 | --- | 9.13E-01 | ? | 1.04E+00 | 5.28E-02 | 1.04E+00 |
| ENSG000002 | --- | 9.13E-01 | ? | 1.05E+00 | 6.96E-02 | 1.05E+00 |
| ENSG000002 | LRRC69 | 9.13E-01 | ? | 9.73E-01 | -4.02E-02 | -1.03E+00 |
| ENSG000002 | ATP13A3-DT | 9.13E-01 | 9.74E-01 | 1.02E+00 | 2.57E-02 | 1.02E+00 |
| ENSG000001 | FAM118B | 9.13E-01 | 9.74E-01 | 1.00E+00 | 6.10E-03 | 1.00E+00 |
| ENSG000002 | EEF1A1P13 | 9.13E-01 | ? | 1.03E+00 | 4.22E-02 | 1.03E+00 |
| ENSG000001 | ACYP1 | 9.13E-01 | 9.74E-01 | 9.91E-01 | -1.34E-02 | -1.01E+00 |
| ENSG000002 | CCDC183-AS1 | 9.14E-01 | ? | 1.02E+00 | 2.84E-02 | 1.02E+00 |
| ENSG000002 | --- | 9.14E-01 | ? | 9.44E-01 | -8.39E-02 | -1.06E+00 |
| ENSG000002 | DNAJC3-DT | 9.14E-01 | 9.74E-01 | 1.02E+00 | 2.40E-02 | 1.02E+00 |
| ENSG000001 | TEX261 | 9.14E-01 | 9.74E-01 | 9.97E-01 | -4.78E-03 | -1.00E+00 |
| ENSG000002 | --- | 9.14E-01 | ? | 1.04E+00 | 5.78E-02 | 1.04E+00 |
| ENSG000001 | MZT2A | 9.14E-01 | 9.74E-01 | 1.01E+00 | 1.15E-02 | 1.01E+00 |
| ENSG000002 | DANT2 | 9.14E-01 | ? | 1.04E+00 | 5.58E-02 | 1.04E+00 |
| ENSG000002 | --- | 9.14E-01 | 9.74E-01 | 1.01E+00 | 1.81E-02 | 1.01E+00 |
| ENSG000002 | --- | 9.14E-01 | ? | 1.04E+00 | 5.47E-02 | 1.04E+00 |
| ENSG000002 | --- | 9.14E-01 | ? | 9.53E-01 | -6.99E-02 | -1.05E+00 |
| ENSG000001 | VPS33B | 9.14E-01 | 9.74E-01 | 1.01E+00 | 8.28E-03 | 1.01E+00 |
| ENSG000001 | MOB1A | 9.14E-01 | 9.74E-01 | 1.00E+00 | 4.35E-03 | 1.00E+00 |
| ENSG000001 | TMEM222 | 9.14E-01 | 9.74E-01 | 1.01E+00 | 8.20E-03 | 1.01E+00 |
| ENSG00000C | NFIX | 9.14E-01 | 9.74E-01 | 1.01E+00 | 7.58E-03 | 1.01E+00 |
| ENSG000001 | LURAP1 | 9.15E-01 | 9.74E-01 | 1.02E+00 | 2.88E-02 | 1.02E+00 |
| ENSG000001 | RUSC1 | 9.15E-01 | 9.74E-01 | 1.00E+00 | 5.27E-03 | 1.00E+00 |
| ENSG000002 | C1QTNF9B | 9.15E-01 | ? | 9.51E-01 | -7.20E-02 | -1.05E+00 |
| ENSG000001 | SLX9 | 9.15E-01 | 9.74E-01 | 9.92E-01 | -1.10E-02 | -1.01E+00 |
| ENSG000002 | TLE1-DT | 9.15E-01 | ? | 1.05E+00 | 7.46E-02 | 1.05E+00 |
| ENSG00000C | ZMYND11 | 9.15E-01 | 9.75E-01 | 1.00E+00 | 5.97E-03 | 1.00E+00 |
| ENSG00000C | CRLF1 | 9.15E-01 | 9.75E-01 | 9.88E-01 | -1.79E-02 | -1.01E+00 |
| ENSG000001 | C1orf159 | 9.15E-01 | 9.75E-01 | 1.01E+00 | 9.02E-03 | 1.01E+00 |
| ENSG000002 | --- | 9.15E-01 | ? | 9.62E-01 | -5.62E-02 | -1.04E+00 |
| ENSG000002 | --- | 9.16E-01 | 9.75E-01 | 9.83E-01 | -2.50E-02 | -1.02E+00 |
| ENSG000001 | GOLGA8F | 9.16E-01 | ? | 1.10E+00 | 1.44E-01 | 1.10E+00 |
| ENSG000001 | MADD | 9.16E-01 | 9.75E-01 | 9.96E-01 | -5.41E-03 | -1.00E+00 |
| ENSG000002 | LINC03049 | 9.16E-01 | 9.75E-01 | 1.02E+00 | 2.77E-02 | 1.02E+00 |
| ENSG000002 | --- | 9.16E-01 | 9.75E-01 | 9.86E-01 | -2.11E-02 | -1.01E+00 |
| ENSG000002 | --- | 9.16E-01 | 9.75E-01 | 9.90E-01 | -1.40E-02 | -1.01E+00 |
| ENSG000002 | PKMP1 | 9.16E-01 | ? | 1.05E+00 | 7.49E-02 | 1.05E+00 |
| ENSG000002 | HNRNPK-AS1 | 9.16E-01 | ? | 1.02E+00 | 3.36E-02 | 1.02E+00 |
| ENSG000001 | KLHL17 | 9.16E-01 | 9.75E-01 | 9.90E-01 | -1.45E-02 | -1.01E+00 |
| ENSG000002 | --- | 9.17E-01 | ? | 9.61E-01 | -5.69E-02 | -1.04E+00 |
| ENSG000001 | SBSPON | 9.17E-01 | ? | 1.02E+00 | 3.10E-02 | 1.02E+00 |
| ENSG000001 | NDUFA5 | 9.17E-01 | 9.75E-01 | 1.01E+00 | 7.25E-03 | 1.01E+00 |
| ENSG000001 | LY6K | 9.17E-01 | 9.75E-01 | 9.88E-01 | -1.74E-02 | -1.01E+00 |

| | | | | | | |
|---|---|---|---|---|---|---|
| ENSG000001 | TPM4 | 9.17E-01 | 9.75E-01 | 9.97E-01 | -5.00E-03 | -1.00E+00 |
| ENSG000002 | ASAH1-AS1 | 9.17E-01 | ? | 1.03E+00 | 4.12E-02 | 1.03E+00 |
| ENSG000001 | RBM4 | 9.17E-01 | 9.75E-01 | 1.00E+00 | 4.80E-03 | 1.00E+00 |
| ENSG000002 | SLC10A5 | 9.17E-01 | ? | 9.67E-01 | -4.90E-02 | -1.03E+00 |
| ENSG000002 | --- | 9.17E-01 | ? | 9.73E-01 | -3.95E-02 | -1.03E+00 |
| ENSG000002 | BMS1P4-AG/ | 9.17E-01 | 9.75E-01 | 9.67E-01 | -4.80E-02 | -1.03E+00 |
| ENSG000001 | TMEM217 | 9.17E-01 | 9.75E-01 | 9.91E-01 | -1.37E-02 | -1.01E+00 |
| ENSG00000C | CHDH | 9.17E-01 | ? | 1.03E+00 | 3.88E-02 | 1.03E+00 |
| ENSG000001 | PWWP3A | 9.17E-01 | 9.76E-01 | 9.96E-01 | -5.31E-03 | -1.00E+00 |
| ENSG000001 | C7orf31 | 9.17E-01 | 9.76E-01 | 1.01E+00 | 1.33E-02 | 1.01E+00 |
| ENSG000001 | VWDE | 9.17E-01 | 9.76E-01 | 9.91E-01 | -1.31E-02 | -1.01E+00 |
| ENSG000002 | --- | 9.17E-01 | ? | 1.07E+00 | 9.56E-02 | 1.07E+00 |
| ENSG00000C | BLTP2 | 9.17E-01 | 9.76E-01 | 9.97E-01 | -4.06E-03 | -1.00E+00 |
| ENSG000001 | AGAP3 | 9.17E-01 | 9.76E-01 | 1.01E+00 | 8.62E-03 | 1.01E+00 |
| ENSG000002 | --- | 9.18E-01 | 9.76E-01 | 1.01E+00 | 1.94E-02 | 1.01E+00 |
| ENSG000001 | KANK4 | 9.18E-01 | 9.76E-01 | 1.01E+00 | 1.15E-02 | 1.01E+00 |
| ENSG000001 | DNAH12 | 9.18E-01 | ? | 1.03E+00 | 4.85E-02 | 1.03E+00 |
| ENSG000002 | RPS3AP21 | 9.18E-01 | ? | 1.03E+00 | 4.27E-02 | 1.03E+00 |
| ENSG000002 | ZNF230-DT | 9.18E-01 | ? | 9.76E-01 | -3.50E-02 | -1.02E+00 |
| ENSG000002 | PRH1 | 9.18E-01 | ? | 9.75E-01 | -3.61E-02 | -1.03E+00 |
| ENSG000002 | AGBL1-AS1 | 9.18E-01 | 9.76E-01 | 1.01E+00 | 1.45E-02 | 1.01E+00 |
| ENSG000002 | --- | 9.18E-01 | ? | 9.67E-01 | -4.87E-02 | -1.03E+00 |
| ENSG000001 | MAP7D3 | 9.18E-01 | 9.76E-01 | 9.95E-01 | -7.29E-03 | -1.01E+00 |
| ENSG000002 | LINC01355 | 9.18E-01 | 9.76E-01 | 9.86E-01 | -2.06E-02 | -1.01E+00 |
| ENSG000001 | HID1 | 9.18E-01 | 9.76E-01 | 9.87E-01 | -1.91E-02 | -1.01E+00 |
| ENSG000001 | LSP1P4 | 9.18E-01 | 9.76E-01 | 1.02E+00 | 2.63E-02 | 1.02E+00 |
| ENSG000001 | SLCO4C1 | 9.18E-01 | ? | 1.04E+00 | 6.34E-02 | 1.04E+00 |
| ENSG000002 | --- | 9.18E-01 | ? | 1.03E+00 | 3.67E-02 | 1.03E+00 |
| ENSG000002 | --- | 9.18E-01 | 9.76E-01 | 1.00E+00 | 6.37E-03 | 1.00E+00 |
| ENSG000002 | --- | 9.18E-01 | 9.76E-01 | 9.84E-01 | -2.33E-02 | -1.02E+00 |
| ENSG000002 | --- | 9.18E-01 | 9.76E-01 | 1.02E+00 | 2.27E-02 | 1.02E+00 |
| ENSG000002 | --- | 9.18E-01 | ? | 1.04E+00 | 5.98E-02 | 1.04E+00 |
| ENSG000002 | LINC00964 | 9.18E-01 | 9.76E-01 | 1.01E+00 | 1.79E-02 | 1.01E+00 |
| ENSG000001 | CAT | 9.18E-01 | 9.76E-01 | 1.00E+00 | 5.73E-03 | 1.00E+00 |
| ENSG000002 | C1orf226 | 9.18E-01 | ? | 1.02E+00 | 2.79E-02 | 1.02E+00 |
| ENSG000001 | USH1G | 9.19E-01 | ? | 9.51E-01 | -7.21E-02 | -1.05E+00 |
| ENSG00000C | ORC1 | 9.19E-01 | 9.76E-01 | 1.02E+00 | 2.48E-02 | 1.02E+00 |
| ENSG000001 | LARP1 | 9.19E-01 | 9.76E-01 | 1.00E+00 | 3.84E-03 | 1.00E+00 |
| ENSG000001 | TPM2 | 9.19E-01 | 9.76E-01 | 9.94E-01 | -8.37E-03 | -1.01E+00 |
| ENSG000002 | --- | 9.19E-01 | ? | 1.05E+00 | 6.82E-02 | 1.05E+00 |
| ENSG000002 | MBL1P | 9.19E-01 | ? | 1.02E+00 | 2.94E-02 | 1.02E+00 |
| ENSG00000C | CAPZB | 9.19E-01 | 9.76E-01 | 9.98E-01 | -3.22E-03 | -1.00E+00 |
| ENSG000001 | UNC50 | 9.19E-01 | 9.76E-01 | 1.00E+00 | 6.94E-03 | 1.00E+00 |
| ENSG000002 | --- | 9.19E-01 | ? | 9.48E-01 | -7.70E-02 | -1.05E+00 |
| ENSG000001 | RPS3AP5 | 9.19E-01 | ? | 9.70E-01 | -4.34E-02 | -1.03E+00 |

| | | | | | | |
|---|---|---|---|---|---|---|
| ENSG000002 | SIRPAP1 | 9.19E-01 | ? | 9.81E-01 | -2.84E-02 | -1.02E+00 |
| ENSG000001 | ROBO4 | 9.19E-01 | ? | 9.82E-01 | -2.63E-02 | -1.02E+00 |
| ENSG000002 | ZNF271P | 9.19E-01 | 9.76E-01 | 9.96E-01 | -6.18E-03 | -1.00E+00 |
| ENSG000001 | MAP7D1 | 9.19E-01 | 9.76E-01 | 9.96E-01 | -5.62E-03 | -1.00E+00 |
| ENSG000001 | MMP7 | 9.19E-01 | ? | 9.67E-01 | -4.86E-02 | -1.03E+00 |
| ENSG000002 | --- | 9.19E-01 | ? | 9.60E-01 | -5.94E-02 | -1.04E+00 |
| ENSG000002 | --- | 9.19E-01 | ? | 1.02E+00 | 3.55E-02 | 1.02E+00 |
| ENSG000001 | DEGS2 | 9.19E-01 | ? | 1.03E+00 | 4.27E-02 | 1.03E+00 |
| ENSG000001 | NHP2P1 | 9.19E-01 | ? | 9.55E-01 | -6.67E-02 | -1.05E+00 |
| ENSG000002 | --- | 9.19E-01 | ? | 9.77E-01 | -3.36E-02 | -1.02E+00 |
| ENSG000001 | TRIR | 9.20E-01 | 9.76E-01 | 1.00E+00 | 6.68E-03 | 1.00E+00 |
| ENSG000001 | RIOK1 | 9.20E-01 | 9.76E-01 | 9.96E-01 | -6.48E-03 | -1.00E+00 |
| ENSG000001 | ZFTA | 9.20E-01 | 9.76E-01 | 1.01E+00 | 7.85E-03 | 1.01E+00 |
| ENSG000002 | ZNF350 | 9.20E-01 | 9.76E-01 | 1.01E+00 | 9.80E-03 | 1.01E+00 |
| ENSG00000C | CHERP | 9.20E-01 | 9.76E-01 | 1.00E+00 | 5.64E-03 | 1.00E+00 |
| ENSG000001 | MAN2C1 | 9.20E-01 | 9.76E-01 | 1.01E+00 | 1.14E-02 | 1.01E+00 |
| ENSG000001 | SDF2L1 | 9.20E-01 | 9.76E-01 | 9.92E-01 | -1.17E-02 | -1.01E+00 |
| ENSG000001 | NSMCE1 | 9.20E-01 | 9.76E-01 | 1.00E+00 | 5.97E-03 | 1.00E+00 |
| ENSG000002 | --- | 9.20E-01 | 9.76E-01 | 9.89E-01 | -1.59E-02 | -1.01E+00 |
| ENSG000001 | SAP30 | 9.20E-01 | 9.76E-01 | 1.01E+00 | 1.09E-02 | 1.01E+00 |
| ENSG000001 | GFAP | 9.20E-01 | ? | 1.04E+00 | 5.13E-02 | 1.04E+00 |
| ENSG000001 | CKM | 9.20E-01 | ? | 9.47E-01 | -7.92E-02 | -1.06E+00 |
| ENSG000002 | APOBEC3G | 9.20E-01 | ? | 9.78E-01 | -3.20E-02 | -1.02E+00 |
| ENSG00000C | ECHDC1 | 9.20E-01 | 9.76E-01 | 9.96E-01 | -5.18E-03 | -1.00E+00 |
| ENSG000001 | FAM90A1 | 9.20E-01 | ? | 1.05E+00 | 6.36E-02 | 1.05E+00 |
| ENSG000002 | --- | 9.20E-01 | ? | 1.03E+00 | 4.16E-02 | 1.03E+00 |
| ENSG000001 | SULT1A2 | 9.20E-01 | ? | 9.80E-01 | -2.84E-02 | -1.02E+00 |
| ENSG000002 | --- | 9.20E-01 | 9.76E-01 | 1.02E+00 | 2.31E-02 | 1.02E+00 |
| ENSG000001 | TRIM17 | 9.20E-01 | 9.76E-01 | 9.82E-01 | -2.55E-02 | -1.02E+00 |
| ENSG000002 | --- | 9.20E-01 | ? | 9.71E-01 | -4.27E-02 | -1.03E+00 |
| ENSG000001 | ARSI | 9.20E-01 | 9.76E-01 | 1.01E+00 | 1.32E-02 | 1.01E+00 |
| ENSG000001 | PSTK | 9.20E-01 | 9.76E-01 | 9.93E-01 | -9.71E-03 | -1.01E+00 |
| ENSG000002 | TRIM13 | 9.20E-01 | 9.76E-01 | 9.96E-01 | -5.21E-03 | -1.00E+00 |
| ENSG000002 | --- | 9.20E-01 | ? | 9.59E-01 | -6.07E-02 | -1.04E+00 |
| ENSG000001 | BUD31 | 9.20E-01 | 9.76E-01 | 9.96E-01 | -5.97E-03 | -1.00E+00 |
| ENSG000002 | AP1G2-AS1 | 9.21E-01 | 9.76E-01 | 9.91E-01 | -1.27E-02 | -1.01E+00 |
| ENSG000001 | RHOJ | 9.21E-01 | ? | 1.02E+00 | 2.93E-02 | 1.02E+00 |
| ENSG000002 | --- | 9.21E-01 | ? | 1.03E+00 | 3.90E-02 | 1.03E+00 |
| ENSG000002 | LINC02975 | 9.21E-01 | ? | 1.03E+00 | 4.13E-02 | 1.03E+00 |
| ENSG000001 | SLC35B1 | 9.21E-01 | 9.76E-01 | 9.96E-01 | -5.29E-03 | -1.00E+00 |
| ENSG000001 | GAK | 9.21E-01 | 9.76E-01 | 9.96E-01 | -5.47E-03 | -1.00E+00 |
| ENSG000001 | FYCO1 | 9.21E-01 | 9.76E-01 | 9.95E-01 | -6.61E-03 | -1.00E+00 |
| ENSG000002 | KCTD13-DT | 9.21E-01 | ? | 9.74E-01 | -3.78E-02 | -1.03E+00 |
| ENSG000001 | MAPKAP1 | 9.21E-01 | 9.76E-01 | 1.00E+00 | 2.97E-03 | 1.00E+00 |
| ENSG000001 | MTERF4 | 9.21E-01 | 9.76E-01 | 1.00E+00 | 6.06E-03 | 1.00E+00 |

| | | | | | | |
|---|---|---|---|---|---|---|
| ENSG000001 | MICAL2 | 9.21E-01 | 9.76E-01 | 1.00E+00 | 4.47E-03 | 1.00E+00 |
| ENSG000002 | SYT15 | 9.21E-01 | ? | 9.81E-01 | -2.70E-02 | -1.02E+00 |
| ENSG000001 | DGKI | 9.21E-01 | 9.76E-01 | 1.01E+00 | 1.12E-02 | 1.01E+00 |
| ENSG000002 | LINC01504 | 9.22E-01 | 9.76E-01 | 9.84E-01 | -2.26E-02 | -1.02E+00 |
| ENSG000001 | SLC39A7 | 9.22E-01 | 9.76E-01 | 1.00E+00 | 4.51E-03 | 1.00E+00 |
| ENSG000001 | NF2 | 9.22E-01 | 9.76E-01 | 1.00E+00 | 4.86E-03 | 1.00E+00 |
| ENSG000001 | SLC25A42 | 9.22E-01 | 9.76E-01 | 9.89E-01 | -1.54E-02 | -1.01E+00 |
| ENSG00000C | FKBP7 | 9.22E-01 | 9.76E-01 | 1.01E+00 | 1.56E-02 | 1.01E+00 |
| ENSG000001 | MAU2 | 9.22E-01 | 9.76E-01 | 9.95E-01 | -7.48E-03 | -1.01E+00 |
| ENSG000001 | MACROD2 | 9.22E-01 | 9.76E-01 | 9.90E-01 | -1.52E-02 | -1.01E+00 |
| ENSG000002 | LINC02004 | 9.22E-01 | ? | 1.04E+00 | 5.42E-02 | 1.04E+00 |
| ENSG000001 | TMOD3 | 9.22E-01 | 9.76E-01 | 9.97E-01 | -4.83E-03 | -1.00E+00 |
| ENSG000001 | MDM4 | 9.22E-01 | 9.76E-01 | 1.01E+00 | 1.04E-02 | 1.01E+00 |
| ENSG000001 | BHLHE41 | 9.22E-01 | 9.76E-01 | 9.86E-01 | -2.04E-02 | -1.01E+00 |
| ENSG000001 | ISG15 | 9.22E-01 | 9.76E-01 | 1.01E+00 | 2.10E-02 | 1.01E+00 |
| ENSG000002 | CALCRL-AS1 | 9.22E-01 | ? | 9.63E-01 | -5.51E-02 | -1.04E+00 |
| ENSG000002 | ZNF593OS | 9.22E-01 | ? | 1.03E+00 | 4.00E-02 | 1.03E+00 |
| ENSG000001 | ZFC3H1 | 9.22E-01 | 9.76E-01 | 9.96E-01 | -5.20E-03 | -1.00E+00 |
| ENSG000001 | RAD1 | 9.22E-01 | 9.76E-01 | 1.00E+00 | 5.39E-03 | 1.00E+00 |
| ENSG000002 | SEC61G-DT | 9.22E-01 | ? | 9.61E-01 | -5.73E-02 | -1.04E+00 |
| ENSG000001 | NUP62CL | 9.22E-01 | ? | 9.82E-01 | -2.58E-02 | -1.02E+00 |
| ENSG000002 | --- | 9.22E-01 | ? | 9.60E-01 | -5.90E-02 | -1.04E+00 |
| ENSG000002 | --- | 9.22E-01 | 9.77E-01 | 9.85E-01 | -2.25E-02 | -1.02E+00 |
| ENSG000001 | PRPF4 | 9.22E-01 | 9.77E-01 | 1.00E+00 | 4.81E-03 | 1.00E+00 |
| ENSG000001 | CALHM5 | 9.22E-01 | 9.77E-01 | 1.01E+00 | 1.81E-02 | 1.01E+00 |
| ENSG000002 | --- | 9.23E-01 | ? | 1.03E+00 | 4.29E-02 | 1.03E+00 |
| ENSG000001 | RHBDD3 | 9.23E-01 | 9.77E-01 | 1.01E+00 | 1.19E-02 | 1.01E+00 |
| ENSG000002 | --- | 9.23E-01 | ? | 1.05E+00 | 6.80E-02 | 1.05E+00 |
| ENSG000002 | PRKDC | 9.23E-01 | 9.77E-01 | 1.01E+00 | 7.95E-03 | 1.01E+00 |
| ENSG000002 | RWDD3-DT | 9.23E-01 | ? | 1.05E+00 | 7.58E-02 | 1.05E+00 |
| ENSG000001 | SLC39A1 | 9.23E-01 | 9.77E-01 | 1.00E+00 | 6.05E-03 | 1.00E+00 |
| ENSG000002 | --- | 9.23E-01 | ? | 1.04E+00 | 5.22E-02 | 1.04E+00 |
| ENSG000002 | SERBP1P5 | 9.23E-01 | 9.77E-01 | 1.01E+00 | 1.76E-02 | 1.01E+00 |
| ENSG000001 | NLGN3 | 9.23E-01 | 9.77E-01 | 1.02E+00 | 2.22E-02 | 1.02E+00 |
| ENSG000002 | --- | 9.23E-01 | ? | 1.03E+00 | 4.26E-02 | 1.03E+00 |
| ENSG000001 | MKRN1 | 9.23E-01 | 9.77E-01 | 1.00E+00 | 3.11E-03 | 1.00E+00 |
| ENSG000002 | --- | 9.23E-01 | ? | 9.78E-01 | -3.16E-02 | -1.02E+00 |
| ENSG000002 | LINC00672 | 9.23E-01 | 9.77E-01 | 9.84E-01 | -2.28E-02 | -1.02E+00 |
| ENSG000002 | LINC02482 | 9.23E-01 | ? | 9.79E-01 | -3.09E-02 | -1.02E+00 |
| ENSG000001 | MYH10 | 9.23E-01 | 9.77E-01 | 9.97E-01 | -4.80E-03 | -1.00E+00 |
| ENSG000002 | --- | 9.23E-01 | ? | 1.03E+00 | 4.77E-02 | 1.03E+00 |
| ENSG000001 | KCNAB1 | 9.23E-01 | 9.77E-01 | 9.89E-01 | -1.54E-02 | -1.01E+00 |
| ENSG000002 | LMNTD2-AS1 | 9.23E-01 | 9.77E-01 | 9.88E-01 | -1.74E-02 | -1.01E+00 |
| ENSG000002 | NIFK-AS1 | 9.24E-01 | 9.77E-01 | 9.89E-01 | -1.66E-02 | -1.01E+00 |
| ENSG000002 | WEE2-AS1 | 9.24E-01 | 9.77E-01 | 9.88E-01 | -1.73E-02 | -1.01E+00 |

| | | | | | | |
|---|---|---|---|---|---|---|
| ENSG000001 | TBL2 | 9.24E-01 | 9.77E-01 | 1.00E+00 | 6.24E-03 | 1.00E+00 |
| ENSG000001 | TRIP4 | 9.24E-01 | 9.77E-01 | 1.00E+00 | 4.44E-03 | 1.00E+00 |
| ENSG000002 | --- | 9.24E-01 | ? | 1.03E+00 | 4.44E-02 | 1.03E+00 |
| ENSG000002 | LINC00683 | 9.24E-01 | 9.77E-01 | 9.86E-01 | -2.08E-02 | -1.01E+00 |
| ENSG000002 | DUX4L26 | 9.24E-01 | ? | 9.57E-01 | -6.30E-02 | -1.04E+00 |
| ENSG00000C | PEX3 | 9.24E-01 | 9.77E-01 | 1.00E+00 | 6.99E-03 | 1.00E+00 |
| ENSG000001 | PEX19 | 9.24E-01 | 9.77E-01 | 1.00E+00 | 4.94E-03 | 1.00E+00 |
| ENSG000002 | --- | 9.24E-01 | 9.77E-01 | 9.84E-01 | -2.39E-02 | -1.02E+00 |
| ENSG000002 | --- | 9.24E-01 | ? | 1.04E+00 | 5.73E-02 | 1.04E+00 |
| ENSG000001 | KAZALD1 | 9.24E-01 | ? | 1.03E+00 | 4.89E-02 | 1.03E+00 |
| ENSG000001 | RSPH3 | 9.24E-01 | 9.77E-01 | 1.00E+00 | 5.44E-03 | 1.00E+00 |
| ENSG000001 | --- | 9.24E-01 | ? | 9.36E-01 | -9.61E-02 | -1.07E+00 |
| ENSG000001 | ZNF17 | 9.24E-01 | 9.77E-01 | 1.01E+00 | 1.01E-02 | 1.01E+00 |
| ENSG000002 | --- | 9.24E-01 | 9.77E-01 | 9.87E-01 | -1.92E-02 | -1.01E+00 |
| ENSG000002 | --- | 9.24E-01 | ? | 1.03E+00 | 4.57E-02 | 1.03E+00 |
| ENSG000001 | DNAH7 | 9.24E-01 | ? | 1.03E+00 | 3.76E-02 | 1.03E+00 |
| ENSG000001 | GPR157 | 9.24E-01 | 9.77E-01 | 9.88E-01 | -1.73E-02 | -1.01E+00 |
| ENSG000002 | --- | 9.24E-01 | ? | 1.03E+00 | 4.09E-02 | 1.03E+00 |
| ENSG000001 | S100A13 | 9.24E-01 | 9.77E-01 | 1.00E+00 | 5.20E-03 | 1.00E+00 |
| ENSG000002 | --- | 9.25E-01 | ? | 9.75E-01 | -3.59E-02 | -1.03E+00 |
| ENSG000001 | KRT18P28 | 9.25E-01 | ? | 9.67E-01 | -4.79E-02 | -1.03E+00 |
| ENSG000002 | DERPC | 9.25E-01 | 9.77E-01 | 1.01E+00 | 1.13E-02 | 1.01E+00 |
| ENSG000002 | SERPINB9P1 | 9.25E-01 | 9.77E-01 | 1.01E+00 | 1.77E-02 | 1.01E+00 |
| ENSG000001 | MAPK3 | 9.25E-01 | 9.77E-01 | 1.01E+00 | 9.24E-03 | 1.01E+00 |
| ENSG000001 | TENM2 | 9.25E-01 | 9.77E-01 | 9.96E-01 | -6.17E-03 | -1.00E+00 |
| ENSG000001 | PDE10A | 9.25E-01 | ? | 9.50E-01 | -7.41E-02 | -1.05E+00 |
| ENSG000001 | PSMG1 | 9.25E-01 | 9.77E-01 | 9.96E-01 | -5.09E-03 | -1.00E+00 |
| ENSG000001 | TMC6 | 9.25E-01 | 9.77E-01 | 1.01E+00 | 8.67E-03 | 1.01E+00 |
| ENSG000001 | CHST9 | 9.25E-01 | 9.77E-01 | 1.02E+00 | 2.25E-02 | 1.02E+00 |
| ENSG000001 | ELP4 | 9.25E-01 | 9.77E-01 | 1.00E+00 | 5.58E-03 | 1.00E+00 |
| ENSG000001 | SOCS4 | 9.25E-01 | 9.77E-01 | 9.96E-01 | -5.75E-03 | -1.00E+00 |
| ENSG000001 | CHD9 | 9.25E-01 | 9.77E-01 | 1.00E+00 | 6.79E-03 | 1.00E+00 |
| ENSG000001 | MAPT | 9.26E-01 | ? | 1.02E+00 | 2.74E-02 | 1.02E+00 |
| ENSG000001 | TKT | 9.26E-01 | 9.78E-01 | 9.97E-01 | -5.04E-03 | -1.00E+00 |
| ENSG000001 | TAF7L | 9.26E-01 | ? | 1.03E+00 | 4.65E-02 | 1.03E+00 |
| ENSG000001 | MSLN | 9.26E-01 | 9.78E-01 | 1.01E+00 | 1.88E-02 | 1.01E+00 |
| ENSG000002 | --- | 9.26E-01 | 9.78E-01 | 9.92E-01 | -1.22E-02 | -1.01E+00 |
| ENSG000001 | BPHL | 9.26E-01 | 9.78E-01 | 1.00E+00 | 6.83E-03 | 1.00E+00 |
| ENSG000001 | CCDC171 | 9.26E-01 | 9.78E-01 | 1.01E+00 | 2.06E-02 | 1.01E+00 |
| ENSG000001 | FGF12 | 9.26E-01 | ? | 9.52E-01 | -7.03E-02 | -1.05E+00 |
| ENSG000001 | PHYKPL | 9.26E-01 | 9.78E-01 | 1.01E+00 | 9.39E-03 | 1.01E+00 |
| ENSG000001 | ANKRD22 | 9.26E-01 | 9.78E-01 | 1.02E+00 | 2.16E-02 | 1.02E+00 |
| ENSG000001 | TMEM104 | 9.26E-01 | 9.78E-01 | 9.95E-01 | -6.55E-03 | -1.00E+00 |
| ENSG000002 | TMPO-AS1 | 9.27E-01 | ? | 1.03E+00 | 3.88E-02 | 1.03E+00 |
| ENSG000001 | ZNF136 | 9.27E-01 | 9.78E-01 | 9.94E-01 | -8.82E-03 | -1.01E+00 |

| | | | | | | |
|---|---|---|---|---|---|---|
| ENSG000001 | TRPM6 | 9.27E-01 | 9.78E-01 | 9.86E-01 | -2.01E-02 | -1.01E+00 |
| ENSG000001 | GRK5 | 9.27E-01 | 9.78E-01 | 9.97E-01 | -4.98E-03 | -1.00E+00 |
| ENSG000001 | TAF1C | 9.27E-01 | 9.78E-01 | 1.01E+00 | 9.78E-03 | 1.01E+00 |
| ENSG000001 | ZNF419 | 9.27E-01 | 9.78E-01 | 9.95E-01 | -7.95E-03 | -1.01E+00 |
| ENSG000001 | UCHL3 | 9.27E-01 | 9.78E-01 | 1.00E+00 | 5.09E-03 | 1.00E+00 |
| ENSG000002 | --- | 9.27E-01 | ? | 9.64E-01 | -5.35E-02 | -1.04E+00 |
| ENSG000001 | KY | 9.27E-01 | ? | 1.02E+00 | 2.73E-02 | 1.02E+00 |
| ENSG000002 | --- | 9.27E-01 | ? | 1.03E+00 | 3.97E-02 | 1.03E+00 |
| ENSG000001 | ZNF384 | 9.27E-01 | 9.78E-01 | 1.00E+00 | 3.97E-03 | 1.00E+00 |
| ENSG000002 | C4orf47 | 9.27E-01 | ? | 9.83E-01 | -2.48E-02 | -1.02E+00 |
| ENSG000002 | --- | 9.27E-01 | ? | 1.03E+00 | 4.65E-02 | 1.03E+00 |
| ENSG000001 | TSNARE1 | 9.27E-01 | 9.78E-01 | 9.90E-01 | -1.45E-02 | -1.01E+00 |
| ENSG000001 | RGS14 | 9.27E-01 | 9.78E-01 | 9.93E-01 | -9.65E-03 | -1.01E+00 |
| ENSG000002 | WASH5P | 9.27E-01 | 9.78E-01 | 1.01E+00 | 1.06E-02 | 1.01E+00 |
| ENSG000001 | HELB | 9.28E-01 | 9.78E-01 | 1.01E+00 | 1.22E-02 | 1.01E+00 |
| ENSG000002 | --- | 9.28E-01 | 9.78E-01 | 1.01E+00 | 1.97E-02 | 1.01E+00 |
| ENSG000001 | ZBTB44 | 9.28E-01 | 9.78E-01 | 9.96E-01 | -5.56E-03 | -1.00E+00 |
| ENSG000001 | GPS1 | 9.28E-01 | 9.78E-01 | 9.96E-01 | -6.43E-03 | -1.00E+00 |
| ENSG000002 | TMEM158 | 9.28E-01 | 9.78E-01 | 9.93E-01 | -1.01E-02 | -1.01E+00 |
| ENSG000001 | PIWIL1 | 9.28E-01 | ? | 9.83E-01 | -2.48E-02 | -1.02E+00 |
| ENSG000001 | MAZ | 9.28E-01 | 9.78E-01 | 9.95E-01 | -6.56E-03 | -1.00E+00 |
| ENSG000001 | CSNK2A1 | 9.28E-01 | 9.78E-01 | 1.00E+00 | 3.62E-03 | 1.00E+00 |
| ENSG000001 | ZNF235 | 9.28E-01 | 9.78E-01 | 9.92E-01 | -1.13E-02 | -1.01E+00 |
| ENSG000002 | TIGD1 | 9.28E-01 | 9.78E-01 | 1.01E+00 | 1.41E-02 | 1.01E+00 |
| ENSG000001 | SLC47A2 | 9.28E-01 | 9.78E-01 | 9.88E-01 | -1.77E-02 | -1.01E+00 |
| ENSG000001 | ASH1L | 9.28E-01 | 9.78E-01 | 9.95E-01 | -7.31E-03 | -1.01E+00 |
| ENSG000001 | ALAD | 9.28E-01 | 9.78E-01 | 1.00E+00 | 6.20E-03 | 1.00E+00 |
| ENSG000002 | --- | 9.28E-01 | ? | 9.72E-01 | -4.16E-02 | -1.03E+00 |
| ENSG000002 | --- | 9.28E-01 | ? | 9.60E-01 | -5.94E-02 | -1.04E+00 |
| ENSG000002 | --- | 9.28E-01 | ? | 9.75E-01 | -3.66E-02 | -1.03E+00 |
| ENSG000002 | ZNF285 | 9.28E-01 | 9.78E-01 | 1.01E+00 | 9.75E-03 | 1.01E+00 |
| ENSG00000C | NCAPD2 | 9.28E-01 | 9.78E-01 | 9.96E-01 | -6.17E-03 | -1.00E+00 |
| ENSG00000C | CMTM6 | 9.28E-01 | 9.78E-01 | 1.00E+00 | 3.48E-03 | 1.00E+00 |
| ENSG000002 | PHLDA1-DT | 9.29E-01 | ? | 9.68E-01 | -4.68E-02 | -1.03E+00 |
| ENSG000002 | --- | 9.29E-01 | ? | 9.69E-01 | -4.50E-02 | -1.03E+00 |
| ENSG000002 | LY6G5C | 9.29E-01 | ? | 9.84E-01 | -2.31E-02 | -1.02E+00 |
| ENSG000002 | ZC3H11B | 9.29E-01 | ? | 1.03E+00 | 4.90E-02 | 1.03E+00 |
| ENSG000001 | CNOT1 | 9.29E-01 | 9.78E-01 | 9.97E-01 | -4.69E-03 | -1.00E+00 |
| ENSG00000C | TXNDC16 | 9.29E-01 | 9.78E-01 | 9.92E-01 | -1.15E-02 | -1.01E+00 |
| ENSG000002 | DNAAF4-CCP | 9.29E-01 | 9.78E-01 | 1.03E+00 | 3.67E-02 | 1.03E+00 |
| ENSG000001 | TAF10 | 9.29E-01 | 9.78E-01 | 1.00E+00 | 6.30E-03 | 1.00E+00 |
| ENSG00000C | PIGQ | 9.29E-01 | 9.79E-01 | 1.01E+00 | 1.14E-02 | 1.01E+00 |
| ENSG000002 | --- | 9.29E-01 | ? | 1.05E+00 | 7.57E-02 | 1.05E+00 |
| ENSG000002 | --- | 9.29E-01 | ? | 9.84E-01 | -2.30E-02 | -1.02E+00 |
| ENSG000002 | --- | 9.29E-01 | ? | 1.04E+00 | 5.41E-02 | 1.04E+00 |

| | | | | | | |
|---|---|---|---|---|---|---|
| ENSG000001 | SZT2 | 9.29E-01 | 9.79E-01 | 9.96E-01 | -6.12E-03 | -1.00E+00 |
| ENSG000001 | SFXN3 | 9.29E-01 | 9.79E-01 | 1.00E+00 | 4.03E-03 | 1.00E+00 |
| ENSG000002 | KRT18P10 | 9.30E-01 | ? | 9.48E-01 | -7.72E-02 | -1.05E+00 |
| ENSG000001 | GPN1 | 9.30E-01 | 9.79E-01 | 9.97E-01 | -4.34E-03 | -1.00E+00 |
| ENSG000002 | --- | 9.30E-01 | ? | 9.81E-01 | -2.71E-02 | -1.02E+00 |
| ENSG000002 | LILRB3 | 9.30E-01 | ? | 1.03E+00 | 4.92E-02 | 1.03E+00 |
| ENSG000001 | NUDT5 | 9.30E-01 | 9.79E-01 | 9.97E-01 | -3.62E-03 | -1.00E+00 |
| ENSG000001 | POP4 | 9.30E-01 | 9.79E-01 | 9.97E-01 | -4.22E-03 | -1.00E+00 |
| ENSG000002 | ZYG11A | 9.30E-01 | ? | 9.72E-01 | -4.14E-02 | -1.03E+00 |
| ENSG00000C | ACHE | 9.30E-01 | ? | 9.77E-01 | -3.38E-02 | -1.02E+00 |
| ENSG000001 | TOB2P1 | 9.30E-01 | ? | 9.74E-01 | -3.81E-02 | -1.03E+00 |
| ENSG000001 | PHF1 | 9.31E-01 | 9.80E-01 | 1.00E+00 | 6.59E-03 | 1.00E+00 |
| ENSG000002 | --- | 9.31E-01 | ? | 9.66E-01 | -4.95E-02 | -1.03E+00 |
| ENSG000002 | CTAGE11P | 9.31E-01 | ? | 1.04E+00 | 6.29E-02 | 1.04E+00 |
| ENSG000001 | GPATCH11 | 9.31E-01 | 9.80E-01 | 1.00E+00 | 5.94E-03 | 1.00E+00 |
| ENSG000001 | ARPC5L | 9.31E-01 | 9.80E-01 | 1.00E+00 | 3.99E-03 | 1.00E+00 |
| ENSG000001 | ARMC1 | 9.31E-01 | 9.80E-01 | 9.97E-01 | -3.73E-03 | -1.00E+00 |
| ENSG000001 | CDK7 | 9.31E-01 | 9.80E-01 | 9.97E-01 | -3.70E-03 | -1.00E+00 |
| ENSG000002 | --- | 9.31E-01 | ? | 1.05E+00 | 7.03E-02 | 1.05E+00 |
| ENSG000001 | HAP1 | 9.31E-01 | 9.80E-01 | 9.91E-01 | -1.36E-02 | -1.01E+00 |
| ENSG000001 | SPRR1B | 9.31E-01 | 9.80E-01 | 1.02E+00 | 2.54E-02 | 1.02E+00 |
| ENSG000001 | GPR161 | 9.31E-01 | 9.80E-01 | 1.00E+00 | 5.76E-03 | 1.00E+00 |
| ENSG000001 | TSPAN5 | 9.31E-01 | 9.80E-01 | 1.00E+00 | 6.34E-03 | 1.00E+00 |
| ENSG000001 | CFAP53 | 9.31E-01 | 9.80E-01 | 1.01E+00 | 1.75E-02 | 1.01E+00 |
| ENSG000001 | CREG2 | 9.32E-01 | ? | 9.82E-01 | -2.65E-02 | -1.02E+00 |
| ENSG000002 | CYP51A1P2 | 9.32E-01 | ? | 1.01E+00 | 2.03E-02 | 1.01E+00 |
| ENSG000001 | CNOT2 | 9.32E-01 | 9.80E-01 | 1.00E+00 | 2.82E-03 | 1.00E+00 |
| ENSG000002 | FOXL2NB | 9.32E-01 | 9.80E-01 | 1.01E+00 | 8.46E-03 | 1.01E+00 |
| ENSG000002 | --- | 9.32E-01 | 9.80E-01 | 1.01E+00 | 1.73E-02 | 1.01E+00 |
| ENSG000001 | KCNC3 | 9.32E-01 | 9.80E-01 | 1.01E+00 | 1.98E-02 | 1.01E+00 |
| ENSG000001 | SERPINB12 | 9.32E-01 | ? | 1.03E+00 | 4.63E-02 | 1.03E+00 |
| ENSG000001 | GATD1 | 9.32E-01 | 9.80E-01 | 1.00E+00 | 5.82E-03 | 1.00E+00 |
| ENSG000001 | DNAAF8 | 9.32E-01 | ? | 9.80E-01 | -2.94E-02 | -1.02E+00 |
| ENSG000001 | CRB3 | 9.32E-01 | 9.80E-01 | 1.01E+00 | 1.16E-02 | 1.01E+00 |
| ENSG000002 | --- | 9.32E-01 | 9.80E-01 | 1.01E+00 | 1.98E-02 | 1.01E+00 |
| ENSG000001 | DLK2 | 9.32E-01 | 9.80E-01 | 9.90E-01 | -1.41E-02 | -1.01E+00 |
| ENSG000001 | ZNF613 | 9.32E-01 | 9.80E-01 | 9.94E-01 | -8.22E-03 | -1.01E+00 |
| ENSG00000C | BCKDHB | 9.32E-01 | 9.80E-01 | 1.00E+00 | 4.71E-03 | 1.00E+00 |
| ENSG00000C | OXCT1 | 9.32E-01 | 9.80E-01 | 9.96E-01 | -5.08E-03 | -1.00E+00 |
| ENSG000001 | RBM44 | 9.32E-01 | ? | 9.63E-01 | -5.46E-02 | -1.04E+00 |
| ENSG000001 | ARMC10 | 9.32E-01 | 9.80E-01 | 9.98E-01 | -3.30E-03 | -1.00E+00 |
| ENSG000002 | ZFP28-DT | 9.32E-01 | ? | 9.61E-01 | -5.66E-02 | -1.04E+00 |
| ENSG000002 | RN7SKP23 | 9.33E-01 | ? | 9.49E-01 | -7.51E-02 | -1.05E+00 |
| ENSG00000C | COL9A2 | 9.33E-01 | ? | 9.83E-01 | -2.45E-02 | -1.02E+00 |
| ENSG000001 | TVP23A | 9.33E-01 | ? | 1.02E+00 | 3.50E-02 | 1.02E+00 |

| | | | | | | |
|---|---|---|---|---|---|---|
| ENSG000002 | LINC02869 | 9.33E-01 | ? | 1.04E+00 | 5.15E-02 | 1.04E+00 |
| ENSG000002 | NEPRO-AS1 | 9.33E-01 | 9.80E-01 | 9.88E-01 | -1.72E-02 | -1.01E+00 |
| ENSG000002 | NECTIN1-DT | 9.33E-01 | ? | 9.61E-01 | -5.71E-02 | -1.04E+00 |
| ENSG000001 | C12orf29 | 9.33E-01 | 9.80E-01 | 9.96E-01 | -5.64E-03 | -1.00E+00 |
| ENSG000001 | ARL8A | 9.33E-01 | 9.80E-01 | 1.00E+00 | 4.26E-03 | 1.00E+00 |
| ENSG000001 | PRELID1 | 9.33E-01 | 9.80E-01 | 9.97E-01 | -4.39E-03 | -1.00E+00 |
| ENSG00000C | GRAMD4 | 9.33E-01 | 9.80E-01 | 9.95E-01 | -7.09E-03 | -1.00E+00 |
| ENSG000002 | --- | 9.33E-01 | ? | 1.02E+00 | 2.64E-02 | 1.02E+00 |
| ENSG000001 | KLHL31 | 9.33E-01 | 9.80E-01 | 1.01E+00 | 1.61E-02 | 1.01E+00 |
| ENSG000001 | ZNF444 | 9.33E-01 | 9.80E-01 | 9.95E-01 | -7.76E-03 | -1.01E+00 |
| ENSG000002 | FOXCUT | 9.33E-01 | ? | 1.04E+00 | 5.68E-02 | 1.04E+00 |
| ENSG000001 | IFIT5 | 9.33E-01 | 9.80E-01 | 9.96E-01 | -6.37E-03 | -1.00E+00 |
| ENSG000001 | PIANP | 9.33E-01 | ? | 1.03E+00 | 3.86E-02 | 1.03E+00 |
| ENSG000002 | UQCRHL | 9.33E-01 | 9.80E-01 | 1.01E+00 | 8.85E-03 | 1.01E+00 |
| ENSG000001 | ZNF534 | 9.33E-01 | ? | 9.56E-01 | -6.48E-02 | -1.05E+00 |
| ENSG000002 | LINC02352 | 9.33E-01 | ? | 9.61E-01 | -5.80E-02 | -1.04E+00 |
| ENSG000002 | GABPAP | 9.33E-01 | ? | 9.72E-01 | -4.11E-02 | -1.03E+00 |
| ENSG000001 | CCNB3 | 9.33E-01 | 9.80E-01 | 1.01E+00 | 1.96E-02 | 1.01E+00 |
| ENSG000001 | ACTA2 | 9.33E-01 | 9.80E-01 | 1.00E+00 | 5.00E-03 | 1.00E+00 |
| ENSG000001 | TUBB2B | 9.34E-01 | 9.80E-01 | 1.01E+00 | 1.12E-02 | 1.01E+00 |
| ENSG000001 | RGPD6 | 9.34E-01 | 9.80E-01 | 1.00E+00 | 6.02E-03 | 1.00E+00 |
| ENSG00000C | LZTS3 | 9.34E-01 | 9.80E-01 | 1.01E+00 | 9.30E-03 | 1.01E+00 |
| ENSG000002 | --- | 9.34E-01 | ? | 1.03E+00 | 3.74E-02 | 1.03E+00 |
| ENSG000001 | TAP1 | 9.34E-01 | 9.80E-01 | 1.00E+00 | 6.38E-03 | 1.00E+00 |
| ENSG000002 | CTSO | 9.34E-01 | 9.80E-01 | 9.96E-01 | -6.38E-03 | -1.00E+00 |
| ENSG000001 | ZNF660 | 9.34E-01 | ? | 9.82E-01 | -2.67E-02 | -1.02E+00 |
| ENSG000002 | SNHG5 | 9.34E-01 | 9.80E-01 | 1.01E+00 | 7.94E-03 | 1.01E+00 |
| ENSG000002 | --- | 9.34E-01 | ? | 1.04E+00 | 6.26E-02 | 1.04E+00 |
| ENSG000001 | SGSM2 | 9.34E-01 | 9.80E-01 | 1.00E+00 | 6.59E-03 | 1.00E+00 |
| ENSG000001 | SNRNP25 | 9.34E-01 | 9.80E-01 | 1.01E+00 | 7.78E-03 | 1.01E+00 |
| ENSG000001 | TREM1 | 9.34E-01 | 9.80E-01 | 9.81E-01 | -2.84E-02 | -1.02E+00 |
| ENSG000002 | --- | 9.34E-01 | ? | 9.72E-01 | -4.16E-02 | -1.03E+00 |
| ENSG000002 | NPEPPSP1 | 9.34E-01 | 9.80E-01 | 1.01E+00 | 8.48E-03 | 1.01E+00 |
| ENSG000001 | ERN1 | 9.34E-01 | 9.80E-01 | 9.97E-01 | -4.25E-03 | -1.00E+00 |
| ENSG000001 | SLC44A5 | 9.34E-01 | 9.80E-01 | 9.93E-01 | -1.01E-02 | -1.01E+00 |
| ENSG000002 | UBE2L5 | 9.34E-01 | ? | 1.03E+00 | 4.78E-02 | 1.03E+00 |
| ENSG000001 | CGAS | 9.34E-01 | 9.80E-01 | 1.01E+00 | 1.22E-02 | 1.01E+00 |
| ENSG000001 | FBXL16 | 9.34E-01 | 9.80E-01 | 9.94E-01 | -8.49E-03 | -1.01E+00 |
| ENSG000002 | ARHGEF35-A | 9.34E-01 | 9.80E-01 | 9.96E-01 | -6.27E-03 | -1.00E+00 |
| ENSG000002 | SMARCA5-AS | 9.34E-01 | 9.80E-01 | 1.01E+00 | 1.27E-02 | 1.01E+00 |
| ENSG000001 | SIVA1 | 9.34E-01 | 9.80E-01 | 1.01E+00 | 1.17E-02 | 1.01E+00 |
| ENSG000002 | --- | 9.34E-01 | ? | 9.77E-01 | -3.41E-02 | -1.02E+00 |
| ENSG000002 | RAD21L1 | 9.34E-01 | ? | 1.03E+00 | 4.96E-02 | 1.03E+00 |
| ENSG00000C | COPZ2 | 9.34E-01 | 9.80E-01 | 1.01E+00 | 8.97E-03 | 1.01E+00 |
| ENSG000002 | PITPNA-AS1 | 9.34E-01 | 9.80E-01 | 9.89E-01 | -1.53E-02 | -1.01E+00 |

| | | | | | | |
|---|---|---|---|---|---|---|
| ENSG000002 | --- | 9.34E-01 | ? | 1.03E+00 | 3.78E-02 | 1.03E+00 |
| ENSG000002 | --- | 9.35E-01 | ? | 1.03E+00 | 4.57E-02 | 1.03E+00 |
| ENSG000001 | HS3ST3B1 | 9.35E-01 | 9.80E-01 | 9.94E-01 | -9.01E-03 | -1.01E+00 |
| ENSG000002 | ELL2P1 | 9.35E-01 | ? | 9.87E-01 | -1.92E-02 | -1.01E+00 |
| ENSG000001 | CCNY | 9.35E-01 | 9.80E-01 | 9.98E-01 | -3.43E-03 | -1.00E+00 |
| ENSG000001 | NSUN7 | 9.35E-01 | ? | 9.71E-01 | -4.32E-02 | -1.03E+00 |
| ENSG000001 | GINS2 | 9.35E-01 | 9.80E-01 | 1.01E+00 | 1.50E-02 | 1.01E+00 |
| ENSG00000C | ZW10 | 9.35E-01 | 9.80E-01 | 9.97E-01 | -3.81E-03 | -1.00E+00 |
| ENSG000002 | --- | 9.35E-01 | ? | 1.03E+00 | 4.84E-02 | 1.03E+00 |
| ENSG000002 | USP27X | 9.35E-01 | 9.81E-01 | 9.93E-01 | -1.01E-02 | -1.01E+00 |
| ENSG000002 | --- | 9.35E-01 | ? | 1.05E+00 | 7.22E-02 | 1.05E+00 |
| ENSG000001 | SETD4 | 9.35E-01 | 9.81E-01 | 1.01E+00 | 7.62E-03 | 1.01E+00 |
| ENSG000001 | TMEM80 | 9.36E-01 | 9.81E-01 | 9.95E-01 | -6.55E-03 | -1.00E+00 |
| ENSG000002 | --- | 9.36E-01 | ? | 9.72E-01 | -4.14E-02 | -1.03E+00 |
| ENSG000002 | --- | 9.36E-01 | ? | 1.03E+00 | 4.45E-02 | 1.03E+00 |
| ENSG000001 | C19orf12 | 9.36E-01 | 9.81E-01 | 1.00E+00 | 6.08E-03 | 1.00E+00 |
| ENSG000001 | HHIPL2 | 9.36E-01 | ? | 9.73E-01 | -3.94E-02 | -1.03E+00 |
| ENSG000001 | C8G | 9.36E-01 | ? | 1.04E+00 | 6.02E-02 | 1.04E+00 |
| ENSG000002 | --- | 9.36E-01 | 9.81E-01 | 1.01E+00 | 9.10E-03 | 1.01E+00 |
| ENSG000001 | TSR1 | 9.36E-01 | 9.81E-01 | 1.00E+00 | 3.73E-03 | 1.00E+00 |
| ENSG000001 | NEK5 | 9.36E-01 | 9.81E-01 | 9.87E-01 | -1.82E-02 | -1.01E+00 |
| ENSG000001 | KLHL30 | 9.36E-01 | ? | 1.02E+00 | 3.26E-02 | 1.02E+00 |
| ENSG000001 | RHOQ | 9.36E-01 | 9.81E-01 | 9.97E-01 | -4.79E-03 | -1.00E+00 |
| ENSG000001 | BBOF1 | 9.36E-01 | 9.81E-01 | 1.01E+00 | 1.02E-02 | 1.01E+00 |
| ENSG000001 | MAP6 | 9.36E-01 | 9.81E-01 | 9.94E-01 | -8.77E-03 | -1.01E+00 |
| ENSG000001 | VASH2 | 9.36E-01 | ? | 9.72E-01 | -4.07E-02 | -1.03E+00 |
| ENSG000002 | --- | 9.36E-01 | ? | 9.84E-01 | -2.27E-02 | -1.02E+00 |
| ENSG000001 | ZBTB17 | 9.36E-01 | 9.81E-01 | 9.95E-01 | -6.69E-03 | -1.00E+00 |
| ENSG000001 | ADGRB1 | 9.36E-01 | ? | 9.80E-01 | -2.91E-02 | -1.02E+00 |
| ENSG000001 | ACOT11 | 9.37E-01 | 9.81E-01 | 9.96E-01 | -5.64E-03 | -1.00E+00 |
| ENSG000002 | --- | 9.37E-01 | ? | 1.03E+00 | 3.98E-02 | 1.03E+00 |
| ENSG000001 | ELMOD2 | 9.37E-01 | 9.81E-01 | 1.00E+00 | 4.46E-03 | 1.00E+00 |
| ENSG000001 | ABCC1 | 9.37E-01 | 9.81E-01 | 1.00E+00 | 3.82E-03 | 1.00E+00 |
| ENSG000001 | RIPPLY3 | 9.37E-01 | ? | 9.62E-01 | -5.65E-02 | -1.04E+00 |
| ENSG000002 | ODAD4 | 9.37E-01 | ? | 9.68E-01 | -4.72E-02 | -1.03E+00 |
| ENSG000001 | UAP1L1 | 9.37E-01 | 9.81E-01 | 9.93E-01 | -1.03E-02 | -1.01E+00 |
| ENSG000002 | --- | 9.37E-01 | ? | 9.76E-01 | -3.53E-02 | -1.02E+00 |
| ENSG000002 | MYL12-AS1 | 9.37E-01 | ? | 9.73E-01 | -4.01E-02 | -1.03E+00 |
| ENSG00000C | MYO15A | 9.37E-01 | ? | 1.01E+00 | 2.04E-02 | 1.01E+00 |
| ENSG000001 | PSMB7 | 9.37E-01 | 9.81E-01 | 1.00E+00 | 3.06E-03 | 1.00E+00 |
| ENSG000002 | C6orf226 | 9.37E-01 | ? | 9.85E-01 | -2.13E-02 | -1.01E+00 |
| ENSG000002 | --- | 9.37E-01 | ? | 1.02E+00 | 3.28E-02 | 1.02E+00 |
| ENSG000001 | TMEM223 | 9.37E-01 | 9.81E-01 | 1.01E+00 | 7.85E-03 | 1.01E+00 |
| ENSG000001 | IQCA1 | 9.37E-01 | 9.81E-01 | 9.97E-01 | -4.76E-03 | -1.00E+00 |
| ENSG000001 | --- | 9.37E-01 | ? | 9.79E-01 | -3.01E-02 | -1.02E+00 |

| | | | | | | |
|---|---|---|---|---|---|---|
| ENSG000002 | HSPA1A | 9.37E-01 | 9.81E-01 | 9.96E-01 | -5.44E-03 | -1.00E+00 |
| ENSG000001 | SH3BGRL2 | 9.37E-01 | 9.81E-01 | 1.00E+00 | 6.55E-03 | 1.00E+00 |
| ENSG000002 | SNHG30 | 9.37E-01 | 9.81E-01 | 9.88E-01 | -1.74E-02 | -1.01E+00 |
| ENSG000002 | SMIM14-DT | 9.37E-01 | ? | 1.01E+00 | 2.13E-02 | 1.01E+00 |
| ENSG000001 | FBXW12 | 9.37E-01 | ? | 9.75E-01 | -3.68E-02 | -1.03E+00 |
| ENSG000002 | --- | 9.38E-01 | ? | 1.03E+00 | 4.39E-02 | 1.03E+00 |
| ENSG000002 | NPIPB7 | 9.38E-01 | ? | 1.02E+00 | 2.26E-02 | 1.02E+00 |
| ENSG000001 | EGFLAM | 9.38E-01 | ? | 1.02E+00 | 3.04E-02 | 1.02E+00 |
| ENSG00000C | PDS5B | 9.38E-01 | 9.81E-01 | 1.00E+00 | 4.89E-03 | 1.00E+00 |
| ENSG000001 | GDF11 | 9.38E-01 | 9.81E-01 | 1.00E+00 | 6.16E-03 | 1.00E+00 |
| ENSG000002 | ZNF717 | 9.38E-01 | 9.81E-01 | 1.00E+00 | 6.04E-03 | 1.00E+00 |
| ENSG000002 | --- | 9.38E-01 | 9.81E-01 | 9.91E-01 | -1.30E-02 | -1.01E+00 |
| ENSG000001 | ZNF45 | 9.38E-01 | 9.81E-01 | 1.00E+00 | 5.07E-03 | 1.00E+00 |
| ENSG000001 | PGRMC1 | 9.38E-01 | 9.81E-01 | 1.00E+00 | 3.27E-03 | 1.00E+00 |
| ENSG000001 | ANKRD40 | 9.38E-01 | 9.81E-01 | 1.00E+00 | 3.71E-03 | 1.00E+00 |
| ENSG000002 | --- | 9.38E-01 | ? | 1.02E+00 | 3.12E-02 | 1.02E+00 |
| ENSG00000C | DNM1L | 9.38E-01 | 9.82E-01 | 9.98E-01 | -3.42E-03 | -1.00E+00 |
| ENSG000001 | TUBD1 | 9.38E-01 | 9.82E-01 | 1.01E+00 | 7.68E-03 | 1.01E+00 |
| ENSG000002 | --- | 9.39E-01 | ? | 9.64E-01 | -5.32E-02 | -1.04E+00 |
| ENSG000001 | ZNF852 | 9.39E-01 | 9.82E-01 | 1.01E+00 | 1.15E-02 | 1.01E+00 |
| ENSG000001 | CCDC152 | 9.39E-01 | ? | 9.77E-01 | -3.31E-02 | -1.02E+00 |
| ENSG000001 | DDIAS | 9.39E-01 | 9.82E-01 | 1.01E+00 | 1.22E-02 | 1.01E+00 |
| ENSG00000C | SPAG9 | 9.39E-01 | 9.82E-01 | 9.98E-01 | -3.36E-03 | -1.00E+00 |
| ENSG000001 | NXPH4 | 9.39E-01 | 9.82E-01 | 9.93E-01 | -1.04E-02 | -1.01E+00 |
| ENSG00000C | TFB1M | 9.39E-01 | 9.82E-01 | 9.96E-01 | -5.68E-03 | -1.00E+00 |
| ENSG000001 | PKN1 | 9.39E-01 | 9.82E-01 | 1.01E+00 | 1.08E-02 | 1.01E+00 |
| ENSG000002 | NBPF8 | 9.39E-01 | 9.82E-01 | 1.00E+00 | 4.16E-03 | 1.00E+00 |
| ENSG000001 | CCDC127 | 9.39E-01 | 9.82E-01 | 9.98E-01 | -3.49E-03 | -1.00E+00 |
| ENSG000001 | HCN3 | 9.39E-01 | 9.82E-01 | 9.91E-01 | -1.31E-02 | -1.01E+00 |
| ENSG000002 | INTS6-AS1 | 9.39E-01 | 9.82E-01 | 9.92E-01 | -1.14E-02 | -1.01E+00 |
| ENSG000001 | CLIC6 | 9.39E-01 | ? | 9.62E-01 | -5.55E-02 | -1.04E+00 |
| ENSG000001 | PHF2 | 9.39E-01 | 9.82E-01 | 9.97E-01 | -4.34E-03 | -1.00E+00 |
| ENSG000002 | --- | 9.39E-01 | ? | 1.02E+00 | 3.14E-02 | 1.02E+00 |
| ENSG000002 | --- | 9.39E-01 | ? | 9.67E-01 | -4.87E-02 | -1.03E+00 |
| ENSG000001 | SNCG | 9.39E-01 | 9.82E-01 | 9.94E-01 | -8.92E-03 | -1.01E+00 |
| ENSG000002 | THBS3-AS1 | 9.39E-01 | 9.82E-01 | 9.86E-01 | -2.07E-02 | -1.01E+00 |
| ENSG000002 | --- | 9.39E-01 | 9.82E-01 | 1.01E+00 | 1.05E-02 | 1.01E+00 |
| ENSG000001 | NGDN | 9.39E-01 | 9.82E-01 | 9.97E-01 | -3.94E-03 | -1.00E+00 |
| ENSG000001 | CFAP46 | 9.39E-01 | ? | 9.81E-01 | -2.74E-02 | -1.02E+00 |
| ENSG000001 | KLK11 | 9.39E-01 | 9.82E-01 | 9.91E-01 | -1.25E-02 | -1.01E+00 |
| ENSG000002 | --- | 9.39E-01 | ? | 1.01E+00 | 2.07E-02 | 1.01E+00 |
| ENSG000001 | KRT34 | 9.40E-01 | ? | 9.87E-01 | -1.95E-02 | -1.01E+00 |
| ENSG000001 | TJP1 | 9.40E-01 | 9.82E-01 | 9.96E-01 | -5.28E-03 | -1.00E+00 |
| ENSG000001 | PLCB2 | 9.40E-01 | 9.82E-01 | 9.89E-01 | -1.58E-02 | -1.01E+00 |
| ENSG000002 | LINC01508 | 9.40E-01 | ? | 9.67E-01 | -4.83E-02 | -1.03E+00 |

| | | | | | | |
|---|---|---|---|---|---|---|
| ENSG000001 | DNM3 | 9.40E-01 | 9.82E-01 | 1.01E+00 | 8.07E-03 | 1.01E+00 |
| ENSG000001 | BCAM | 9.40E-01 | 9.82E-01 | 1.00E+00 | 4.66E-03 | 1.00E+00 |
| ENSG000001 | MAGT1 | 9.40E-01 | 9.82E-01 | 1.00E+00 | 3.42E-03 | 1.00E+00 |
| ENSG000002 | --- | 9.40E-01 | ? | 9.73E-01 | -4.02E-02 | -1.03E+00 |
| ENSG000002 | --- | 9.40E-01 | ? | 1.03E+00 | 3.97E-02 | 1.03E+00 |
| ENSG000001 | CD68 | 9.40E-01 | 9.82E-01 | 9.95E-01 | -7.76E-03 | -1.01E+00 |
| ENSG000001 | USPL1 | 9.40E-01 | 9.82E-01 | 9.96E-01 | -5.52E-03 | -1.00E+00 |
| ENSG000001 | TIMM17A | 9.41E-01 | 9.82E-01 | 9.98E-01 | -3.15E-03 | -1.00E+00 |
| ENSG00000C | CYP51A1 | 9.41E-01 | 9.82E-01 | 9.98E-01 | -2.50E-03 | -1.00E+00 |
| ENSG000001 | METRNL | 9.41E-01 | 9.82E-01 | 9.92E-01 | -1.18E-02 | -1.01E+00 |
| ENSG000001 | TMEM135 | 9.41E-01 | 9.82E-01 | 1.00E+00 | 4.64E-03 | 1.00E+00 |
| ENSG000001 | HS6ST3 | 9.41E-01 | ? | 9.86E-01 | -2.07E-02 | -1.01E+00 |
| ENSG000002 | --- | 9.41E-01 | ? | 9.86E-01 | -2.03E-02 | -1.01E+00 |
| ENSG000002 | TTN-AS1 | 9.41E-01 | 9.83E-01 | 1.00E+00 | 6.87E-03 | 1.00E+00 |
| ENSG000002 | LINC01273 | 9.41E-01 | ? | 1.03E+00 | 3.81E-02 | 1.03E+00 |
| ENSG000001 | LYSMD1 | 9.41E-01 | 9.83E-01 | 9.96E-01 | -6.06E-03 | -1.00E+00 |
| ENSG000002 | RANBP1P1 | 9.41E-01 | ? | 1.03E+00 | 4.22E-02 | 1.03E+00 |
| ENSG000001 | SLC25A51 | 9.41E-01 | 9.83E-01 | 9.97E-01 | -3.82E-03 | -1.00E+00 |
| ENSG000002 | FOXD2-AS1 | 9.41E-01 | 9.83E-01 | 1.01E+00 | 1.28E-02 | 1.01E+00 |
| ENSG000001 | ZC3H18 | 9.42E-01 | 9.83E-01 | 9.98E-01 | -3.10E-03 | -1.00E+00 |
| ENSG000001 | ASXL1 | 9.42E-01 | 9.83E-01 | 9.98E-01 | -3.57E-03 | -1.00E+00 |
| ENSG000001 | PRUNE2 | 9.42E-01 | 9.83E-01 | 1.01E+00 | 1.42E-02 | 1.01E+00 |
| ENSG000001 | CMYA5 | 9.42E-01 | 9.83E-01 | 1.01E+00 | 1.23E-02 | 1.01E+00 |
| ENSG000001 | MEIS3 | 9.42E-01 | 9.83E-01 | 9.95E-01 | -6.89E-03 | -1.00E+00 |
| ENSG000001 | ZNF615 | 9.42E-01 | 9.83E-01 | 1.00E+00 | 6.67E-03 | 1.00E+00 |
| ENSG000001 | CCN2 | 9.42E-01 | 9.83E-01 | 1.00E+00 | 6.17E-03 | 1.00E+00 |
| ENSG000001 | RSPO4 | 9.42E-01 | ? | 1.02E+00 | 3.55E-02 | 1.02E+00 |
| ENSG000001 | KIF26B | 9.42E-01 | 9.83E-01 | 9.96E-01 | -6.18E-03 | -1.00E+00 |
| ENSG000002 | TPTEP2-CSNI | 9.42E-01 | ? | 1.03E+00 | 3.71E-02 | 1.03E+00 |
| ENSG000002 | --- | 9.43E-01 | 9.83E-01 | 1.01E+00 | 1.23E-02 | 1.01E+00 |
| ENSG000001 | WDR59 | 9.43E-01 | 9.83E-01 | 1.00E+00 | 3.89E-03 | 1.00E+00 |
| ENSG000001 | PEX12 | 9.43E-01 | 9.83E-01 | 1.00E+00 | 6.18E-03 | 1.00E+00 |
| ENSG00000C | TNRC6C | 9.43E-01 | 9.83E-01 | 1.01E+00 | 9.71E-03 | 1.01E+00 |
| ENSG000001 | COG3 | 9.43E-01 | 9.83E-01 | 1.00E+00 | 3.74E-03 | 1.00E+00 |
| ENSG000001 | DNAJC17 | 9.43E-01 | 9.83E-01 | 1.00E+00 | 5.51E-03 | 1.00E+00 |
| ENSG000001 | EWSR1 | 9.43E-01 | 9.83E-01 | 9.98E-01 | -2.80E-03 | -1.00E+00 |
| ENSG000002 | --- | 9.43E-01 | ? | 1.03E+00 | 3.87E-02 | 1.03E+00 |
| ENSG000001 | HEPHL1 | 9.43E-01 | 9.83E-01 | 9.91E-01 | -1.33E-02 | -1.01E+00 |
| ENSG000001 | UCKL1 | 9.43E-01 | 9.83E-01 | 1.00E+00 | 6.44E-03 | 1.00E+00 |
| ENSG00000C | TNFRSF1A | 9.43E-01 | 9.83E-01 | 9.98E-01 | -2.95E-03 | -1.00E+00 |
| ENSG000002 | --- | 9.43E-01 | 9.83E-01 | 1.01E+00 | 2.10E-02 | 1.01E+00 |
| ENSG000002 | LINC00667 | 9.43E-01 | 9.83E-01 | 9.97E-01 | -4.46E-03 | -1.00E+00 |
| ENSG000002 | SLC30A6-DT | 9.43E-01 | ? | 9.70E-01 | -4.35E-02 | -1.03E+00 |
| ENSG000001 | CARD9 | 9.43E-01 | 9.83E-01 | 1.01E+00 | 1.50E-02 | 1.01E+00 |
| ENSG000001 | MPP3 | 9.43E-01 | 9.83E-01 | 1.01E+00 | 1.04E-02 | 1.01E+00 |

| | | | | | | |
|---|---|---|---|---|---|---|
| ENSG000001 | MED8 | 9.43E-01 | 9.83E-01 | 9.97E-01 | -3.64E-03 | -1.00E+00 |
| ENSG000002 | PAM16 | 9.43E-01 | 9.83E-01 | 1.01E+00 | 8.56E-03 | 1.01E+00 |
| ENSG000001 | HENMT1 | 9.43E-01 | 9.83E-01 | 9.91E-01 | -1.25E-02 | -1.01E+00 |
| ENSG000002 | ZNF812P | 9.44E-01 | ? | 9.41E-01 | -8.74E-02 | -1.06E+00 |
| ENSG000001 | CHKB | 9.44E-01 | 9.83E-01 | 1.01E+00 | 7.21E-03 | 1.01E+00 |
| ENSG000001 | HSPA2 | 9.44E-01 | 9.83E-01 | 9.95E-01 | -7.05E-03 | -1.00E+00 |
| ENSG000002 | --- | 9.44E-01 | ? | 1.04E+00 | 5.47E-02 | 1.04E+00 |
| ENSG000001 | ZC3H6 | 9.44E-01 | 9.83E-01 | 1.01E+00 | 7.51E-03 | 1.01E+00 |
| ENSG000001 | RBMX2 | 9.44E-01 | 9.83E-01 | 1.00E+00 | 3.98E-03 | 1.00E+00 |
| ENSG00000C | CACNB1 | 9.44E-01 | 9.83E-01 | 1.01E+00 | 7.59E-03 | 1.01E+00 |
| ENSG000001 | DLG4 | 9.44E-01 | 9.83E-01 | 1.00E+00 | 7.03E-03 | 1.00E+00 |
| ENSG000002 | PRELID1P6 | 9.44E-01 | ? | 9.80E-01 | -2.85E-02 | -1.02E+00 |
| ENSG000002 | THBS1-IT1 | 9.44E-01 | 9.83E-01 | 1.01E+00 | 1.04E-02 | 1.01E+00 |
| ENSG000001 | HAUS1 | 9.44E-01 | 9.83E-01 | 9.95E-01 | -6.71E-03 | -1.00E+00 |
| ENSG000002 | RNASEH1-DT | 9.45E-01 | 9.83E-01 | 9.95E-01 | -7.64E-03 | -1.01E+00 |
| ENSG000002 | RPL6P19 | 9.45E-01 | ? | 9.85E-01 | -2.24E-02 | -1.02E+00 |
| ENSG000001 | TRIM11 | 9.45E-01 | 9.83E-01 | 9.97E-01 | -4.32E-03 | -1.00E+00 |
| ENSG000002 | TP53RK-DT | 9.45E-01 | ? | 9.75E-01 | -3.70E-02 | -1.03E+00 |
| ENSG000002 | --- | 9.45E-01 | ? | 1.01E+00 | 1.96E-02 | 1.01E+00 |
| ENSG000001 | PITPNA | 9.45E-01 | 9.83E-01 | 1.00E+00 | 2.68E-03 | 1.00E+00 |
| ENSG000002 | --- | 9.45E-01 | ? | 9.70E-01 | -4.35E-02 | -1.03E+00 |
| ENSG000002 | --- | 9.45E-01 | ? | 1.03E+00 | 3.93E-02 | 1.03E+00 |
| ENSG000001 | HAUS2 | 9.45E-01 | 9.83E-01 | 1.00E+00 | 3.32E-03 | 1.00E+00 |
| ENSG000002 | NR2F2-AS1 | 9.45E-01 | ? | 9.85E-01 | -2.24E-02 | -1.02E+00 |
| ENSG000001 | GPHN | 9.45E-01 | 9.83E-01 | 1.00E+00 | 4.37E-03 | 1.00E+00 |
| ENSG000001 | SERTAD2 | 9.45E-01 | 9.83E-01 | 1.00E+00 | 3.48E-03 | 1.00E+00 |
| ENSG00000C | ESR1 | 9.45E-01 | 9.83E-01 | 1.01E+00 | 1.43E-02 | 1.01E+00 |
| ENSG000002 | PDPR2P | 9.45E-01 | 9.83E-01 | 1.01E+00 | 1.99E-02 | 1.01E+00 |
| ENSG000002 | --- | 9.45E-01 | 9.83E-01 | 1.01E+00 | 7.81E-03 | 1.01E+00 |
| ENSG000001 | MTX2 | 9.45E-01 | 9.84E-01 | 9.97E-01 | -3.64E-03 | -1.00E+00 |
| ENSG000001 | ZNF25 | 9.45E-01 | 9.84E-01 | 9.95E-01 | -7.46E-03 | -1.01E+00 |
| ENSG000001 | B2M | 9.46E-01 | 9.84E-01 | 1.00E+00 | 4.34E-03 | 1.00E+00 |
| ENSG000001 | KIAA0825 | 9.46E-01 | 9.84E-01 | 1.01E+00 | 1.08E-02 | 1.01E+00 |
| ENSG000001 | TMEM107 | 9.46E-01 | 9.84E-01 | 1.00E+00 | 7.19E-03 | 1.00E+00 |
| ENSG000001 | FOXC2 | 9.46E-01 | ? | 1.01E+00 | 1.83E-02 | 1.01E+00 |
| ENSG00000C | GIPR | 9.46E-01 | 9.84E-01 | 1.01E+00 | 1.45E-02 | 1.01E+00 |
| ENSG000002 | --- | 9.46E-01 | ? | 1.02E+00 | 2.96E-02 | 1.02E+00 |
| ENSG000001 | WASHC1 | 9.46E-01 | 9.84E-01 | 9.95E-01 | -7.61E-03 | -1.01E+00 |
| ENSG000001 | RFK | 9.46E-01 | 9.84E-01 | 1.00E+00 | 5.50E-03 | 1.00E+00 |
| ENSG000001 | PLK4 | 9.46E-01 | 9.84E-01 | 9.93E-01 | -1.07E-02 | -1.01E+00 |
| ENSG000001 | NUTM2G | 9.46E-01 | 9.84E-01 | 9.94E-01 | -9.15E-03 | -1.01E+00 |
| ENSG00000C | DGKA | 9.46E-01 | 9.84E-01 | 9.98E-01 | -3.45E-03 | -1.00E+00 |
| ENSG000002 | --- | 9.46E-01 | ? | 1.01E+00 | 1.75E-02 | 1.01E+00 |
| ENSG000001 | SLC6A4 | 9.46E-01 | ? | 9.71E-01 | -4.18E-02 | -1.03E+00 |
| ENSG000001 | DNPEP | 9.46E-01 | 9.84E-01 | 9.97E-01 | -4.07E-03 | -1.00E+00 |

| | | | | | | |
|---|---|---|---|---|---|---|
| ENSG000002 | --- | 9.46E-01 | ? | 1.02E+00 | 2.47E-02 | 1.02E+00 |
| ENSG000001 | RPL22L1 | 9.46E-01 | 9.84E-01 | 1.00E+00 | 4.71E-03 | 1.00E+00 |
| ENSG00000C | HBQ1 | 9.46E-01 | ? | 9.69E-01 | -4.59E-02 | -1.03E+00 |
| ENSG000001 | ALPL | 9.46E-01 | ? | 1.02E+00 | 3.33E-02 | 1.02E+00 |
| ENSG000001 | ZNF131 | 9.46E-01 | 9.84E-01 | 9.97E-01 | -4.48E-03 | -1.00E+00 |
| ENSG000001 | WDR75 | 9.46E-01 | 9.84E-01 | 9.98E-01 | -3.43E-03 | -1.00E+00 |
| ENSG000002 | UCKL1-AS1 | 9.47E-01 | 9.84E-01 | 1.01E+00 | 1.52E-02 | 1.01E+00 |
| ENSG000001 | KCNS3 | 9.47E-01 | 9.84E-01 | 9.93E-01 | -1.01E-02 | -1.01E+00 |
| ENSG000001 | A2M | 9.47E-01 | ? | 1.02E+00 | 3.04E-02 | 1.02E+00 |
| ENSG000002 | SMIM5 | 9.47E-01 | ? | 1.02E+00 | 2.66E-02 | 1.02E+00 |
| ENSG000002 | ASMTL-AS1 | 9.47E-01 | 9.84E-01 | 1.01E+00 | 1.05E-02 | 1.01E+00 |
| ENSG000001 | FBXL6 | 9.47E-01 | 9.84E-01 | 9.94E-01 | -8.80E-03 | -1.01E+00 |
| ENSG000001 | SDHAF4 | 9.47E-01 | 9.84E-01 | 1.00E+00 | 5.02E-03 | 1.00E+00 |
| ENSG00000C | PER3 | 9.47E-01 | 9.84E-01 | 1.02E+00 | 2.19E-02 | 1.02E+00 |
| ENSG000002 | --- | 9.47E-01 | ? | 1.02E+00 | 3.25E-02 | 1.02E+00 |
| ENSG000001 | RNF166 | 9.47E-01 | 9.84E-01 | 1.00E+00 | 6.70E-03 | 1.00E+00 |
| ENSG000002 | PBX2P1 | 9.47E-01 | ? | 9.66E-01 | -5.06E-02 | -1.04E+00 |
| ENSG000002 | --- | 9.47E-01 | ? | 1.04E+00 | 5.34E-02 | 1.04E+00 |
| ENSG000002 | ATP6V1FNB | 9.47E-01 | 9.84E-01 | 9.89E-01 | -1.63E-02 | -1.01E+00 |
| ENSG000001 | PRMT3 | 9.47E-01 | 9.84E-01 | 9.97E-01 | -3.89E-03 | -1.00E+00 |
| ENSG000002 | --- | 9.47E-01 | 9.84E-01 | 9.93E-01 | -9.55E-03 | -1.01E+00 |
| ENSG000001 | RGS2 | 9.47E-01 | 9.84E-01 | 9.96E-01 | -6.49E-03 | -1.00E+00 |
| ENSG00000C | P2RX7 | 9.47E-01 | ? | 9.84E-01 | -2.30E-02 | -1.02E+00 |
| ENSG000001 | TRIM24 | 9.48E-01 | 9.84E-01 | 1.00E+00 | 4.51E-03 | 1.00E+00 |
| ENSG000001 | NDUFV2 | 9.48E-01 | 9.84E-01 | 9.98E-01 | -3.20E-03 | -1.00E+00 |
| ENSG000002 | --- | 9.48E-01 | ? | 1.03E+00 | 3.97E-02 | 1.03E+00 |
| ENSG000002 | PANO1 | 9.48E-01 | ? | 1.01E+00 | 1.94E-02 | 1.01E+00 |
| ENSG000002 | --- | 9.48E-01 | ? | 9.85E-01 | -2.12E-02 | -1.01E+00 |
| ENSG000001 | BICDL2 | 9.48E-01 | 9.84E-01 | 1.01E+00 | 1.95E-02 | 1.01E+00 |
| ENSG000001 | ACAD10 | 9.48E-01 | 9.84E-01 | 1.00E+00 | 4.84E-03 | 1.00E+00 |
| ENSG000001 | ZNF501 | 9.48E-01 | 9.84E-01 | 9.95E-01 | -7.85E-03 | -1.01E+00 |
| ENSG000001 | ABHD12B | 9.48E-01 | ? | 1.02E+00 | 3.13E-02 | 1.02E+00 |
| ENSG000001 | S100A6 | 9.48E-01 | 9.85E-01 | 9.96E-01 | -5.18E-03 | -1.00E+00 |
| ENSG000001 | TNFRSF13C | 9.48E-01 | ? | 9.66E-01 | -4.97E-02 | -1.04E+00 |
| ENSG000001 | FAM25G | 9.48E-01 | ? | 9.79E-01 | -3.02E-02 | -1.02E+00 |
| ENSG000002 | FAM25C | 9.48E-01 | ? | 9.79E-01 | -3.02E-02 | -1.02E+00 |
| ENSG000001 | ACOXL | 9.49E-01 | ? | 1.02E+00 | 2.78E-02 | 1.02E+00 |
| ENSG00000C | PSMC5 | 9.49E-01 | 9.85E-01 | 1.00E+00 | 2.71E-03 | 1.00E+00 |
| ENSG000002 | --- | 9.49E-01 | ? | 9.84E-01 | -2.32E-02 | -1.02E+00 |
| ENSG000002 | --- | 9.49E-01 | ? | 1.02E+00 | 2.79E-02 | 1.02E+00 |
| ENSG000001 | HAS1 | 9.49E-01 | ? | 1.03E+00 | 4.35E-02 | 1.03E+00 |
| ENSG000002 | --- | 9.49E-01 | ? | 1.01E+00 | 1.97E-02 | 1.01E+00 |
| ENSG000002 | ALKBH6 | 9.49E-01 | 9.85E-01 | 9.96E-01 | -5.84E-03 | -1.00E+00 |
| ENSG000001 | SH3GLB2 | 9.49E-01 | 9.85E-01 | 9.96E-01 | -5.33E-03 | -1.00E+00 |
| ENSG000002 | GOT1-DT | 9.49E-01 | 9.85E-01 | 1.01E+00 | 1.43E-02 | 1.01E+00 |

| | | | | | | |
|---|---|---|---|---|---|---|
| ENSG000002 | SPACA6-AS1 | 9.49E-01 | ? | 9.83E-01 | -2.45E-02 | -1.02E+00 |
| ENSG000002 | ERVK-28 | 9.49E-01 | ? | 9.66E-01 | -4.98E-02 | -1.04E+00 |
| ENSG000001 | SEC31A | 9.49E-01 | 9.85E-01 | 9.98E-01 | -2.69E-03 | -1.00E+00 |
| ENSG000002 | --- | 9.49E-01 | 9.85E-01 | 1.01E+00 | 1.98E-02 | 1.01E+00 |
| ENSG000001 | CRYBB2P1 | 9.49E-01 | ? | 1.05E+00 | 7.57E-02 | 1.05E+00 |
| ENSG000002 | LINC01530 | 9.49E-01 | ? | 1.03E+00 | 3.84E-02 | 1.03E+00 |
| ENSG000002 | --- | 9.49E-01 | ? | 1.02E+00 | 2.24E-02 | 1.02E+00 |
| ENSG000002 | FAM133DP | 9.49E-01 | ? | 1.02E+00 | 2.83E-02 | 1.02E+00 |
| ENSG00000C | SDHA | 9.49E-01 | 9.85E-01 | 9.98E-01 | -3.18E-03 | -1.00E+00 |
| ENSG000001 | IFT27 | 9.50E-01 | 9.85E-01 | 9.96E-01 | -6.23E-03 | -1.00E+00 |
| ENSG000002 | --- | 9.50E-01 | ? | 9.85E-01 | -2.12E-02 | -1.01E+00 |
| ENSG00000C | ELP1 | 9.50E-01 | 9.85E-01 | 1.00E+00 | 3.29E-03 | 1.00E+00 |
| ENSG000002 | --- | 9.50E-01 | 9.85E-01 | 1.01E+00 | 2.04E-02 | 1.01E+00 |
| ENSG000001 | HMGN2 | 9.50E-01 | 9.85E-01 | 1.00E+00 | 4.40E-03 | 1.00E+00 |
| ENSG000001 | SERPINB7 | 9.50E-01 | 9.85E-01 | 9.92E-01 | -1.15E-02 | -1.01E+00 |
| ENSG000002 | SNRPGP2 | 9.50E-01 | ? | 9.76E-01 | -3.47E-02 | -1.02E+00 |
| ENSG000002 | HPS1-AS1 | 9.50E-01 | ? | 9.80E-01 | -2.88E-02 | -1.02E+00 |
| ENSG000001 | PIP5K1A | 9.50E-01 | 9.85E-01 | 1.00E+00 | 2.12E-03 | 1.00E+00 |
| ENSG000001 | ZNF337 | 9.50E-01 | 9.85E-01 | 1.00E+00 | 5.11E-03 | 1.00E+00 |
| ENSG000001 | INTS11 | 9.50E-01 | 9.85E-01 | 1.00E+00 | 4.02E-03 | 1.00E+00 |
| ENSG000001 | FBRS | 9.50E-01 | 9.85E-01 | 1.00E+00 | 4.05E-03 | 1.00E+00 |
| ENSG000001 | GPR180 | 9.50E-01 | 9.85E-01 | 1.00E+00 | 4.62E-03 | 1.00E+00 |
| ENSG00000C | GATB | 9.50E-01 | 9.85E-01 | 1.00E+00 | 3.81E-03 | 1.00E+00 |
| ENSG000001 | ARHGAP12 | 9.50E-01 | 9.85E-01 | 1.00E+00 | 4.44E-03 | 1.00E+00 |
| ENSG000001 | STOM | 9.50E-01 | 9.85E-01 | 1.00E+00 | 4.05E-03 | 1.00E+00 |
| ENSG000002 | NOX5 | 9.50E-01 | ? | 1.03E+00 | 4.07E-02 | 1.03E+00 |
| ENSG000002 | UQCRFS1P1 | 9.50E-01 | ? | 1.03E+00 | 3.92E-02 | 1.03E+00 |
| ENSG00000C | FER1L4 | 9.50E-01 | 9.85E-01 | 1.01E+00 | 7.24E-03 | 1.01E+00 |
| ENSG000002 | --- | 9.50E-01 | ? | 9.74E-01 | -3.73E-02 | -1.03E+00 |
| ENSG00000C | POU2F2 | 9.50E-01 | 9.85E-01 | 1.00E+00 | 5.01E-03 | 1.00E+00 |
| ENSG000002 | --- | 9.50E-01 | ? | 9.85E-01 | -2.21E-02 | -1.02E+00 |
| ENSG000002 | RNU1-28P | 9.51E-01 | ? | 1.05E+00 | 6.89E-02 | 1.05E+00 |
| ENSG000002 | RNU1-27P | 9.51E-01 | ? | 1.05E+00 | 6.89E-02 | 1.05E+00 |
| ENSG000001 | LRAT | 9.51E-01 | 9.85E-01 | 1.01E+00 | 1.34E-02 | 1.01E+00 |
| ENSG000001 | IRX3 | 9.51E-01 | 9.85E-01 | 1.01E+00 | 7.57E-03 | 1.01E+00 |
| ENSG000002 | --- | 9.51E-01 | ? | 1.01E+00 | 1.61E-02 | 1.01E+00 |
| ENSG000001 | PRKG1 | 9.51E-01 | 9.86E-01 | 1.01E+00 | 1.51E-02 | 1.01E+00 |
| ENSG000001 | SSBP2 | 9.51E-01 | 9.86E-01 | 1.00E+00 | 4.29E-03 | 1.00E+00 |
| ENSG000001 | OVOL2 | 9.51E-01 | 9.86E-01 | 9.93E-01 | -9.81E-03 | -1.01E+00 |
| ENSG000001 | PIGO | 9.51E-01 | 9.86E-01 | 9.97E-01 | -4.85E-03 | -1.00E+00 |
| ENSG000001 | ZNF331 | 9.51E-01 | 9.86E-01 | 1.00E+00 | 5.15E-03 | 1.00E+00 |
| ENSG00000C | DPM1 | 9.51E-01 | 9.86E-01 | 1.00E+00 | 3.05E-03 | 1.00E+00 |
| ENSG000001 | ARL15 | 9.52E-01 | 9.86E-01 | 1.00E+00 | 3.75E-03 | 1.00E+00 |
| ENSG000001 | DMRTA1 | 9.52E-01 | 9.86E-01 | 9.95E-01 | -6.73E-03 | -1.00E+00 |
| ENSG000002 | --- | 9.52E-01 | 9.86E-01 | 1.01E+00 | 1.07E-02 | 1.01E+00 |

| | | | | | | |
|---|---|---|---|---|---|---|
| ENSG000002 | F8A1 | 9.52E-01 | 9.86E-01 | 1.00E+00 | 4.31E-03 | 1.00E+00 |
| ENSG00000C | ZC3H15 | 9.52E-01 | 9.86E-01 | 9.98E-01 | -2.81E-03 | -1.00E+00 |
| ENSG00000C | SPHK2 | 9.52E-01 | 9.86E-01 | 1.01E+00 | 7.34E-03 | 1.01E+00 |
| ENSG000001 | SEC14L4 | 9.52E-01 ? | | 9.86E-01 | -1.97E-02 | -1.01E+00 |
| ENSG00000C | HAL | 9.52E-01 ? | | 1.01E+00 | 2.10E-02 | 1.01E+00 |
| ENSG000001 | SCNN1G | 9.52E-01 ? | | 9.74E-01 | -3.76E-02 | -1.03E+00 |
| ENSG000001 | GLRB | 9.52E-01 | 9.86E-01 | 9.91E-01 | -1.24E-02 | -1.01E+00 |
| ENSG000001 | BCAS3 | 9.52E-01 | 9.86E-01 | 1.00E+00 | 5.12E-03 | 1.00E+00 |
| ENSG000002 | H2BC7 | 9.52E-01 ? | | 9.86E-01 | -2.03E-02 | -1.01E+00 |
| ENSG000001 | CKB | 9.52E-01 | 9.86E-01 | 9.95E-01 | -6.78E-03 | -1.00E+00 |
| ENSG000002 | --- | 9.53E-01 ? | | 1.03E+00 | 3.89E-02 | 1.03E+00 |
| ENSG000002 | MMP12 | 9.53E-01 ? | | 9.88E-01 | -1.81E-02 | -1.01E+00 |
| ENSG000002 | --- | 9.53E-01 ? | | 9.84E-01 | -2.37E-02 | -1.02E+00 |
| ENSG00000C | CTDP1 | 9.53E-01 | 9.86E-01 | 1.00E+00 | 4.72E-03 | 1.00E+00 |
| ENSG000002 | LINC00887 | 9.53E-01 ? | | 9.72E-01 | -4.09E-02 | -1.03E+00 |
| ENSG000002 | --- | 9.53E-01 | 9.86E-01 | 1.00E+00 | 6.27E-03 | 1.00E+00 |
| ENSG000001 | KLHL38 | 9.53E-01 ? | | 1.02E+00 | 2.67E-02 | 1.02E+00 |
| ENSG000001 | MIER1 | 9.53E-01 | 9.86E-01 | 9.97E-01 | -4.06E-03 | -1.00E+00 |
| ENSG000001 | PERM1 | 9.53E-01 ? | | 1.02E+00 | 2.55E-02 | 1.02E+00 |
| ENSG000001 | NR2C2 | 9.53E-01 | 9.86E-01 | 9.97E-01 | -3.70E-03 | -1.00E+00 |
| ENSG000001 | TUBB6 | 9.53E-01 | 9.86E-01 | 9.98E-01 | -3.29E-03 | -1.00E+00 |
| ENSG000002 | XNDC1N | 9.53E-01 | 9.86E-01 | 9.92E-01 | -1.22E-02 | -1.01E+00 |
| ENSG000001 | ZMAT1 | 9.53E-01 ? | | 9.89E-01 | -1.66E-02 | -1.01E+00 |
| ENSG000001 | MTHFS | 9.53E-01 | 9.86E-01 | 1.00E+00 | 7.04E-03 | 1.00E+00 |
| ENSG000001 | CCDC191 | 9.53E-01 | 9.86E-01 | 1.01E+00 | 7.53E-03 | 1.01E+00 |
| ENSG000002 | --- | 9.53E-01 ? | | 9.79E-01 | -3.13E-02 | -1.02E+00 |
| ENSG000002 | --- | 9.53E-01 ? | | 9.74E-01 | -3.86E-02 | -1.03E+00 |
| ENSG000002 | GPR155-DT | 9.54E-01 ? | | 1.02E+00 | 3.32E-02 | 1.02E+00 |
| ENSG000001 | TMEM120A | 9.54E-01 | 9.87E-01 | 9.97E-01 | -4.80E-03 | -1.00E+00 |
| ENSG000001 | MGST3 | 9.54E-01 | 9.87E-01 | 9.99E-01 | -1.94E-03 | -1.00E+00 |
| ENSG000002 | --- | 9.54E-01 ? | | 1.03E+00 | 3.93E-02 | 1.03E+00 |
| ENSG000002 | SCAT8 | 9.54E-01 | 9.87E-01 | 9.93E-01 | -9.51E-03 | -1.01E+00 |
| ENSG000002 | --- | 9.54E-01 ? | | 9.78E-01 | -3.16E-02 | -1.02E+00 |
| ENSG000002 | --- | 9.54E-01 ? | | 9.83E-01 | -2.47E-02 | -1.02E+00 |
| ENSG000001 | CEP164 | 9.54E-01 | 9.87E-01 | 1.00E+00 | 2.99E-03 | 1.00E+00 |
| ENSG000001 | RPS6KC1 | 9.54E-01 | 9.87E-01 | 9.98E-01 | -3.23E-03 | -1.00E+00 |
| ENSG000001 | TCHH | 9.54E-01 ? | | 9.87E-01 | -1.82E-02 | -1.01E+00 |
| ENSG000002 | --- | 9.54E-01 ? | | 1.02E+00 | 3.41E-02 | 1.02E+00 |
| ENSG000001 | SHISA4 | 9.54E-01 | 9.87E-01 | 1.01E+00 | 8.89E-03 | 1.01E+00 |
| ENSG000002 | UHRF1 | 9.55E-01 | 9.87E-01 | 9.96E-01 | -5.28E-03 | -1.00E+00 |
| ENSG000001 | LTBP2 | 9.55E-01 | 9.87E-01 | 1.00E+00 | 3.04E-03 | 1.00E+00 |
| ENSG000002 | FAM185A | 9.55E-01 | 9.87E-01 | 9.95E-01 | -7.57E-03 | -1.01E+00 |
| ENSG000001 | UBE2S | 9.55E-01 | 9.87E-01 | 1.00E+00 | 5.03E-03 | 1.00E+00 |
| ENSG000002 | LINC02021 | 9.55E-01 ? | | 1.02E+00 | 3.27E-02 | 1.02E+00 |
| ENSG000001 | KIAA1143 | 9.55E-01 | 9.87E-01 | 9.98E-01 | -3.46E-03 | -1.00E+00 |

| | | | | | | | |
|---|---|---|---|---|---|---|---|
| ENSG000002 | --- | 9.55E-01 | ? | | 1.01E+00 | 1.91E-02 | 1.01E+00 |
| ENSG000001 | H2AC7 | 9.55E-01 | ? | | 9.86E-01 | -2.04E-02 | -1.01E+00 |
| ENSG000002 | --- | 9.55E-01 | ? | | 9.75E-01 | -3.65E-02 | -1.03E+00 |
| ENSG00000C | PPP2R3C | 9.55E-01 | | 9.87E-01 | 1.00E+00 | 3.50E-03 | 1.00E+00 |
| ENSG00000C | MADCAM1 | 9.55E-01 | ? | | 9.78E-01 | -3.17E-02 | -1.02E+00 |
| ENSG000001 | TLR6 | 9.55E-01 | | 9.87E-01 | 1.01E+00 | 8.50E-03 | 1.01E+00 |
| ENSG000002 | ZNF790-AS1 | 9.55E-01 | | 9.87E-01 | 9.94E-01 | -8.85E-03 | -1.01E+00 |
| ENSG000001 | PSMF1 | 9.55E-01 | | 9.87E-01 | 1.00E+00 | 1.94E-03 | 1.00E+00 |
| ENSG000002 | --- | 9.55E-01 | | 9.87E-01 | 1.01E+00 | 9.67E-03 | 1.01E+00 |
| ENSG000002 | --- | 9.55E-01 | ? | | 9.71E-01 | -4.30E-02 | -1.03E+00 |
| ENSG000002 | --- | 9.56E-01 | ? | | 9.73E-01 | -3.98E-02 | -1.03E+00 |
| ENSG000002 | LINC01003 | 9.56E-01 | | 9.87E-01 | 1.01E+00 | 1.20E-02 | 1.01E+00 |
| ENSG000002 | LINC02725 | 9.56E-01 | | 9.87E-01 | 9.93E-01 | -1.02E-02 | -1.01E+00 |
| ENSG000001 | EHBP1L1 | 9.56E-01 | | 9.87E-01 | 1.00E+00 | 3.43E-03 | 1.00E+00 |
| ENSG000002 | CEACAM16-A | 9.56E-01 | ? | | 9.74E-01 | -3.80E-02 | -1.03E+00 |
| ENSG000002 | ANKRD10-IT1 | 9.56E-01 | | 9.87E-01 | 9.91E-01 | -1.36E-02 | -1.01E+00 |
| ENSG000001 | KLHL7 | 9.56E-01 | | 9.87E-01 | 1.00E+00 | 2.52E-03 | 1.00E+00 |
| ENSG000001 | SERINC2 | 9.56E-01 | | 9.87E-01 | 1.00E+00 | 3.58E-03 | 1.00E+00 |
| ENSG000001 | CAPRIN1 | 9.56E-01 | | 9.87E-01 | 1.00E+00 | 1.70E-03 | 1.00E+00 |
| ENSG000001 | RNF165 | 9.56E-01 | | 9.87E-01 | 9.89E-01 | -1.55E-02 | -1.01E+00 |
| ENSG000001 | ZNF652 | 9.56E-01 | | 9.87E-01 | 9.98E-01 | -3.53E-03 | -1.00E+00 |
| ENSG000002 | ABCF2-H2BK | 9.56E-01 | | 9.87E-01 | 9.97E-01 | -3.72E-03 | -1.00E+00 |
| ENSG000002 | LINC02560 | 9.56E-01 | ? | | 1.03E+00 | 4.45E-02 | 1.03E+00 |
| ENSG000002 | --- | 9.56E-01 | | 9.87E-01 | 1.01E+00 | 1.17E-02 | 1.01E+00 |
| ENSG000001 | EIF3M | 9.56E-01 | | 9.87E-01 | 9.98E-01 | -2.25E-03 | -1.00E+00 |
| ENSG000002 | C14orf132 | 9.56E-01 | | 9.87E-01 | 1.01E+00 | 7.56E-03 | 1.01E+00 |
| ENSG000001 | PA2G4 | 9.56E-01 | | 9.87E-01 | 1.00E+00 | 2.95E-03 | 1.00E+00 |
| ENSG000001 | KCNJ2 | 9.56E-01 | | 9.87E-01 | 1.01E+00 | 9.80E-03 | 1.01E+00 |
| ENSG000002 | LINC02795 | 9.56E-01 | ? | | 9.85E-01 | -2.17E-02 | -1.02E+00 |
| ENSG000002 | ZNF213-AS1 | 9.56E-01 | | 9.87E-01 | 9.94E-01 | -8.67E-03 | -1.01E+00 |
| ENSG000002 | --- | 9.57E-01 | | 9.87E-01 | 9.92E-01 | -1.19E-02 | -1.01E+00 |
| ENSG000002 | LINC02942 | 9.57E-01 | ? | | 1.03E+00 | 3.84E-02 | 1.03E+00 |
| ENSG000002 | ENSAP2 | 9.57E-01 | ? | | 9.86E-01 | -2.01E-02 | -1.01E+00 |
| ENSG000001 | TSNAX | 9.57E-01 | | 9.87E-01 | 1.00E+00 | 3.69E-03 | 1.00E+00 |
| ENSG000002 | TEN1-CDK3 | 9.57E-01 | | 9.87E-01 | 9.88E-01 | -1.81E-02 | -1.01E+00 |
| ENSG000002 | --- | 9.57E-01 | ? | | 1.02E+00 | 2.64E-02 | 1.02E+00 |
| ENSG000001 | CEP72 | 9.57E-01 | | 9.87E-01 | 9.95E-01 | -6.79E-03 | -1.00E+00 |
| ENSG000002 | RANBP17 | 9.57E-01 | | 9.87E-01 | 1.00E+00 | 7.13E-03 | 1.00E+00 |
| ENSG000001 | PXK | 9.57E-01 | | 9.87E-01 | 1.00E+00 | 3.50E-03 | 1.00E+00 |
| ENSG000002 | --- | 9.57E-01 | | 9.87E-01 | 1.01E+00 | 9.40E-03 | 1.01E+00 |
| ENSG000002 | --- | 9.57E-01 | ? | | 9.83E-01 | -2.46E-02 | -1.02E+00 |
| ENSG000002 | --- | 9.57E-01 | | 9.87E-01 | 1.00E+00 | 5.03E-03 | 1.00E+00 |
| ENSG000002 | --- | 9.57E-01 | | 9.88E-01 | 1.00E+00 | 6.71E-03 | 1.00E+00 |
| ENSG000001 | EFHC2 | 9.57E-01 | ? | | 1.02E+00 | 3.19E-02 | 1.02E+00 |
| ENSG000002 | SLC7A14-AS1 | 9.58E-01 | ? | | 1.03E+00 | 4.03E-02 | 1.03E+00 |

| | | | | | | | |
|---|---|---|---|---|---|---|---|
| ENSG000001 | ARHGAP35 | 9.58E-01 | | 9.88E-01 | 1.00E+00 | 2.25E-03 | 1.00E+00 |
| ENSG000000 | ASTE1 | 9.58E-01 | | 9.88E-01 | 9.96E-01 | -5.60E-03 | -1.00E+00 |
| ENSG000001 | NOC4L | 9.58E-01 | | 9.88E-01 | 9.97E-01 | -4.98E-03 | -1.00E+00 |
| ENSG000001 | TMEM86B | 9.58E-01 | ? | | 1.01E+00 | 1.45E-02 | 1.01E+00 |
| ENSG000001 | SLC38A10 | 9.58E-01 | | 9.88E-01 | 9.97E-01 | -4.35E-03 | -1.00E+00 |
| ENSG000002 | --- | 9.58E-01 | ? | | 9.78E-01 | -3.15E-02 | -1.02E+00 |
| ENSG000001 | NME2P1 | 9.58E-01 | ? | | 1.02E+00 | 3.39E-02 | 1.02E+00 |
| ENSG000001 | STAC | 9.58E-01 | ? | | 9.84E-01 | -2.30E-02 | -1.02E+00 |
| ENSG000002 | MILIP | 9.58E-01 | ? | | 9.86E-01 | -2.11E-02 | -1.01E+00 |
| ENSG000001 | GGT7 | 9.58E-01 | | 9.88E-01 | 1.00E+00 | 7.05E-03 | 1.00E+00 |
| ENSG000002 | --- | 9.58E-01 | ? | | 9.83E-01 | -2.53E-02 | -1.02E+00 |
| ENSG000001 | CHPT1 | 9.58E-01 | | 9.88E-01 | 1.01E+00 | 9.01E-03 | 1.01E+00 |
| ENSG000001 | SPECC1 | 9.58E-01 | | 9.88E-01 | 1.00E+00 | 2.75E-03 | 1.00E+00 |
| ENSG000002 | LINC02005 | 9.58E-01 | ? | | 9.89E-01 | -1.55E-02 | -1.01E+00 |
| ENSG000002 | USP51 | 9.58E-01 | | 9.88E-01 | 1.01E+00 | 9.83E-03 | 1.01E+00 |
| ENSG000002 | MAP3K2-DT | 9.58E-01 | ? | | 1.01E+00 | 1.38E-02 | 1.01E+00 |
| ENSG000002 | DENND3-AS1 | 9.58E-01 | | 9.88E-01 | 9.94E-01 | -9.36E-03 | -1.01E+00 |
| ENSG000002 | SH3TC2-DT | 9.58E-01 | | 9.88E-01 | 1.01E+00 | 9.41E-03 | 1.01E+00 |
| ENSG000001 | RBFOX3 | 9.58E-01 | ? | | 1.02E+00 | 2.68E-02 | 1.02E+00 |
| ENSG000000 | RPL26L1 | 9.58E-01 | | 9.88E-01 | 9.98E-01 | -2.89E-03 | -1.00E+00 |
| ENSG000001 | CTTNBP2NL | 9.58E-01 | | 9.88E-01 | 9.98E-01 | -2.70E-03 | -1.00E+00 |
| ENSG000000 | TBC1D23 | 9.58E-01 | | 9.88E-01 | 1.00E+00 | 2.18E-03 | 1.00E+00 |
| ENSG000001 | CDK5R2 | 9.58E-01 | ? | | 9.87E-01 | -1.86E-02 | -1.01E+00 |
| ENSG000001 | PRKCD | 9.58E-01 | | 9.88E-01 | 1.00E+00 | 1.92E-03 | 1.00E+00 |
| ENSG000001 | WASHC2C | 9.59E-01 | | 9.88E-01 | 1.00E+00 | 2.43E-03 | 1.00E+00 |
| ENSG000002 | --- | 9.59E-01 | ? | | 9.80E-01 | -2.88E-02 | -1.02E+00 |
| ENSG000002 | SPTBN1-AS2 | 9.59E-01 | ? | | 9.89E-01 | -1.55E-02 | -1.01E+00 |
| ENSG000001 | RAB17 | 9.59E-01 | | 9.88E-01 | 1.01E+00 | 7.71E-03 | 1.01E+00 |
| ENSG000002 | KPNB1-DT | 9.59E-01 | ? | | 1.01E+00 | 1.47E-02 | 1.01E+00 |
| ENSG000001 | ARSK | 9.59E-01 | | 9.88E-01 | 1.00E+00 | 4.38E-03 | 1.00E+00 |
| ENSG000002 | --- | 9.59E-01 | ? | | 9.87E-01 | -1.89E-02 | -1.01E+00 |
| ENSG000002 | TERF1P7 | 9.59E-01 | ? | | 9.73E-01 | -3.99E-02 | -1.03E+00 |
| ENSG000002 | FLRT2-AS1 | 9.59E-01 | | 9.88E-01 | 1.00E+00 | 4.02E-03 | 1.00E+00 |
| ENSG000001 | ATG10 | 9.59E-01 | | 9.88E-01 | 9.97E-01 | -4.84E-03 | -1.00E+00 |
| ENSG000002 | HSD17B2-AS | 9.59E-01 | ? | | 9.86E-01 | -1.97E-02 | -1.01E+00 |
| ENSG000002 | IKBKG | 9.59E-01 | | 9.88E-01 | 9.97E-01 | -4.46E-03 | -1.00E+00 |
| ENSG000002 | KRT81 | 9.59E-01 | ? | | 1.03E+00 | 3.62E-02 | 1.03E+00 |
| ENSG000001 | PRORSD1P | 9.59E-01 | | 9.88E-01 | 9.91E-01 | -1.23E-02 | -1.01E+00 |
| ENSG000001 | ASL | 9.59E-01 | | 9.88E-01 | 9.97E-01 | -4.20E-03 | -1.00E+00 |
| ENSG000002 | LCAT | 9.59E-01 | | 9.88E-01 | 9.97E-01 | -4.86E-03 | -1.00E+00 |
| ENSG000001 | UBXN10 | 9.59E-01 | ? | | 1.01E+00 | 1.80E-02 | 1.01E+00 |
| ENSG000002 | --- | 9.59E-01 | | 9.88E-01 | 1.02E+00 | 3.24E-02 | 1.02E+00 |
| ENSG000001 | HLA-DQB1 | 9.59E-01 | | 9.88E-01 | 9.93E-01 | -9.75E-03 | -1.01E+00 |
| ENSG000001 | ARHGEF17 | 9.59E-01 | | 9.88E-01 | 9.98E-01 | -2.98E-03 | -1.00E+00 |
| ENSG000001 | FAM86B1 | 9.60E-01 | | 9.88E-01 | 9.95E-01 | -7.32E-03 | -1.01E+00 |

| | | | | | | |
|---|---|---|---|---|---|---|
| ENSG000002 | --- | 9.60E-01 | ? | 1.02E+00 | 2.30E-02 | 1.02E+00 |
| ENSG000002 | --- | 9.60E-01 | 9.88E-01 | 9.89E-01 | -1.53E-02 | -1.01E+00 |
| ENSG000002 | --- | 9.60E-01 | ? | 1.02E+00 | 2.18E-02 | 1.02E+00 |
| ENSG00000C | SPA17 | 9.60E-01 | 9.88E-01 | 1.00E+00 | 4.12E-03 | 1.00E+00 |
| ENSG000001 | PCDHB14 | 9.60E-01 | 9.88E-01 | 9.90E-01 | -1.42E-02 | -1.01E+00 |
| ENSG000002 | --- | 9.60E-01 | ? | 1.01E+00 | 1.56E-02 | 1.01E+00 |
| ENSG000001 | SMG1P3 | 9.60E-01 | 9.88E-01 | 9.92E-01 | -1.16E-02 | -1.01E+00 |
| ENSG000002 | AFTPH-DT | 9.60E-01 | ? | 9.91E-01 | -1.34E-02 | -1.01E+00 |
| ENSG000001 | PITPNB | 9.60E-01 | 9.88E-01 | 9.98E-01 | -2.37E-03 | -1.00E+00 |
| ENSG000001 | YPEL4 | 9.60E-01 | ? | 1.01E+00 | 1.32E-02 | 1.01E+00 |
| ENSG000002 | DUXAP10 | 9.60E-01 | 9.88E-01 | 1.01E+00 | 8.12E-03 | 1.01E+00 |
| ENSG000001 | RPL7AP66 | 9.60E-01 | ? | 1.01E+00 | 1.64E-02 | 1.01E+00 |
| ENSG000001 | C22orf46 | 9.60E-01 | 9.88E-01 | 9.97E-01 | -4.25E-03 | -1.00E+00 |
| ENSG000001 | RAB3GAP1 | 9.60E-01 | 9.88E-01 | 1.00E+00 | 1.90E-03 | 1.00E+00 |
| ENSG000002 | --- | 9.61E-01 | ? | 9.92E-01 | -1.21E-02 | -1.01E+00 |
| ENSG000002 | --- | 9.61E-01 | ? | 1.02E+00 | 2.49E-02 | 1.02E+00 |
| ENSG000002 | SCAF8 | 9.61E-01 | 9.88E-01 | 9.98E-01 | -2.34E-03 | -1.00E+00 |
| ENSG000001 | RPP38 | 9.61E-01 | 9.88E-01 | 1.00E+00 | 2.93E-03 | 1.00E+00 |
| ENSG000001 | SMN1 | 9.61E-01 | 9.88E-01 | 1.00E+00 | 3.71E-03 | 1.00E+00 |
| ENSG000001 | ATG4B | 9.61E-01 | 9.88E-01 | 9.97E-01 | -3.89E-03 | -1.00E+00 |
| ENSG000001 | ARFGAP1 | 9.61E-01 | 9.88E-01 | 9.98E-01 | -3.17E-03 | -1.00E+00 |
| ENSG000002 | LINC01145 | 9.61E-01 | 9.88E-01 | 9.97E-01 | -3.71E-03 | -1.00E+00 |
| ENSG00000C | STRN4 | 9.61E-01 | 9.88E-01 | 9.99E-01 | -2.16E-03 | -1.00E+00 |
| ENSG000001 | PRDX1 | 9.61E-01 | 9.88E-01 | 9.99E-01 | -1.76E-03 | -1.00E+00 |
| ENSG000002 | DIRC3 | 9.61E-01 | 9.88E-01 | 1.01E+00 | 1.68E-02 | 1.01E+00 |
| ENSG000001 | WDR45 | 9.61E-01 | 9.88E-01 | 9.97E-01 | -3.73E-03 | -1.00E+00 |
| ENSG000001 | ANKZF1 | 9.61E-01 | 9.88E-01 | 9.97E-01 | -3.82E-03 | -1.00E+00 |
| ENSG000001 | RNASEH2C | 9.61E-01 | 9.88E-01 | 9.97E-01 | -4.99E-03 | -1.00E+00 |
| ENSG000002 | SNORA71B | 9.61E-01 | ? | 1.02E+00 | 2.79E-02 | 1.02E+00 |
| ENSG000001 | MIB2 | 9.61E-01 | 9.88E-01 | 9.96E-01 | -5.52E-03 | -1.00E+00 |
| ENSG000002 | LINC00862 | 9.62E-01 | 9.88E-01 | 9.96E-01 | -5.23E-03 | -1.00E+00 |
| ENSG000001 | CHRNA10 | 9.62E-01 | ? | 1.01E+00 | 1.42E-02 | 1.01E+00 |
| ENSG000001 | LYST | 9.62E-01 | 9.88E-01 | 1.00E+00 | 6.12E-03 | 1.00E+00 |
| ENSG000001 | B4GALNT2 | 9.62E-01 | 9.89E-01 | 9.97E-01 | -4.41E-03 | -1.00E+00 |
| ENSG000001 | OR5E1P | 9.62E-01 | ? | 1.02E+00 | 3.09E-02 | 1.02E+00 |
| ENSG000002 | TAGLN2P1 | 9.62E-01 | ? | 1.01E+00 | 1.09E-02 | 1.01E+00 |
| ENSG000001 | TCN2 | 9.62E-01 | ? | 9.90E-01 | -1.49E-02 | -1.01E+00 |
| ENSG000002 | LINC01284 | 9.62E-01 | ? | 9.74E-01 | -3.81E-02 | -1.03E+00 |
| ENSG000001 | ACKR2 | 9.62E-01 | ? | 1.01E+00 | 1.21E-02 | 1.01E+00 |
| ENSG000001 | STK38 | 9.62E-01 | 9.89E-01 | 1.00E+00 | 2.12E-03 | 1.00E+00 |
| ENSG000002 | --- | 9.62E-01 | ? | 9.87E-01 | -1.88E-02 | -1.01E+00 |
| ENSG000001 | EPOR | 9.62E-01 | 9.89E-01 | 1.01E+00 | 1.10E-02 | 1.01E+00 |
| ENSG000002 | --- | 9.62E-01 | ? | 9.84E-01 | -2.28E-02 | -1.02E+00 |
| ENSG000001 | SOCS1 | 9.63E-01 | ? | 9.89E-01 | -1.65E-02 | -1.01E+00 |
| ENSG000002 | LINC00886 | 9.63E-01 | 9.89E-01 | 9.93E-01 | -9.72E-03 | -1.01E+00 |

| | | | | | | |
|---|---|---|---|---|---|---|
| ENSG000001 | MELK | 9.63E-01 | 9.89E-01 | 9.97E-01 | -3.71E-03 | -1.00E+00 |
| ENSG000001 | KYAT3 | 9.63E-01 | 9.89E-01 | 9.98E-01 | -2.18E-03 | -1.00E+00 |
| ENSG000002 | LINC01876 | 9.63E-01 | ? | 1.02E+00 | 2.77E-02 | 1.02E+00 |
| ENSG000001 | CDC42BPB | 9.63E-01 | 9.89E-01 | 1.00E+00 | 1.67E-03 | 1.00E+00 |
| ENSG000001 | SLC17A7 | 9.63E-01 | ? | 1.02E+00 | 2.74E-02 | 1.02E+00 |
| ENSG000001 | HEY1 | 9.63E-01 | 9.89E-01 | 1.00E+00 | 6.22E-03 | 1.00E+00 |
| ENSG000001 | TMPRSS11F | 9.63E-01 | ? | 9.90E-01 | -1.52E-02 | -1.01E+00 |
| ENSG000001 | GIMAP8 | 9.63E-01 | 9.89E-01 | 9.93E-01 | -1.04E-02 | -1.01E+00 |
| ENSG000001 | WDFY3 | 9.63E-01 | 9.89E-01 | 9.97E-01 | -3.65E-03 | -1.00E+00 |
| ENSG000002 | --- | 9.63E-01 | ? | 1.02E+00 | 3.13E-02 | 1.02E+00 |
| ENSG000001 | PDIA6 | 9.63E-01 | 9.89E-01 | 9.98E-01 | -2.46E-03 | -1.00E+00 |
| ENSG000001 | UGT2B7 | 9.63E-01 | 9.89E-01 | 9.92E-01 | -1.16E-02 | -1.01E+00 |
| ENSG000002 | --- | 9.63E-01 | ? | 1.01E+00 | 1.75E-02 | 1.01E+00 |
| ENSG000001 | CLCN5 | 9.63E-01 | 9.89E-01 | 9.97E-01 | -4.52E-03 | -1.00E+00 |
| ENSG000002 | --- | 9.63E-01 | ? | 9.89E-01 | -1.65E-02 | -1.01E+00 |
| ENSG000001 | RCAN2 | 9.64E-01 | ? | 9.87E-01 | -1.83E-02 | -1.01E+00 |
| ENSG000002 | --- | 9.64E-01 | 9.89E-01 | 9.94E-01 | -8.77E-03 | -1.01E+00 |
| ENSG000002 | RPL23AP64 | 9.64E-01 | ? | 9.91E-01 | -1.31E-02 | -1.01E+00 |
| ENSG000002 | MIATNB | 9.64E-01 | 9.90E-01 | 1.00E+00 | 5.06E-03 | 1.00E+00 |
| ENSG00000C | MED15 | 9.64E-01 | 9.90E-01 | 9.98E-01 | -2.26E-03 | -1.00E+00 |
| ENSG000001 | IL16 | 9.64E-01 | ? | 1.01E+00 | 1.22E-02 | 1.01E+00 |
| ENSG000002 | --- | 9.64E-01 | 9.90E-01 | 1.01E+00 | 9.01E-03 | 1.01E+00 |
| ENSG000001 | CTU1 | 9.64E-01 | 9.90E-01 | 9.93E-01 | -9.41E-03 | -1.01E+00 |
| ENSG000002 | H1-12P | 9.64E-01 | ? | 9.86E-01 | -1.98E-02 | -1.01E+00 |
| ENSG000001 | CPED1 | 9.64E-01 | ? | 9.88E-01 | -1.76E-02 | -1.01E+00 |
| ENSG000001 | ALDOA | 9.64E-01 | 9.90E-01 | 1.00E+00 | 3.01E-03 | 1.00E+00 |
| ENSG00000C | RNF4 | 9.64E-01 | 9.90E-01 | 9.99E-01 | -1.62E-03 | -1.00E+00 |
| ENSG000002 | --- | 9.64E-01 | 9.90E-01 | 9.91E-01 | -1.24E-02 | -1.01E+00 |
| ENSG000001 | TALDO1 | 9.64E-01 | 9.90E-01 | 1.00E+00 | 2.07E-03 | 1.00E+00 |
| ENSG000001 | NET1 | 9.65E-01 | 9.90E-01 | 9.98E-01 | -3.47E-03 | -1.00E+00 |
| ENSG000002 | --- | 9.65E-01 | ? | 9.74E-01 | -3.82E-02 | -1.03E+00 |
| ENSG000002 | PSMB10 | 9.65E-01 | 9.90E-01 | 9.96E-01 | -6.41E-03 | -1.00E+00 |
| ENSG000001 | B4GALT5 | 9.65E-01 | 9.90E-01 | 1.00E+00 | 1.59E-03 | 1.00E+00 |
| ENSG000002 | --- | 9.65E-01 | ? | 1.02E+00 | 2.62E-02 | 1.02E+00 |
| ENSG000001 | TBC1D10C | 9.65E-01 | ? | 1.02E+00 | 2.99E-02 | 1.02E+00 |
| ENSG000002 | --- | 9.65E-01 | ? | 1.01E+00 | 1.35E-02 | 1.01E+00 |
| ENSG00000C | MMP11 | 9.65E-01 | ? | 9.87E-01 | -1.94E-02 | -1.01E+00 |
| ENSG000002 | --- | 9.65E-01 | ? | 1.01E+00 | 1.68E-02 | 1.01E+00 |
| ENSG00000C | SRPK1 | 9.65E-01 | 9.90E-01 | 9.99E-01 | -1.87E-03 | -1.00E+00 |
| ENSG000001 | SLC25A14 | 9.65E-01 | 9.90E-01 | 9.97E-01 | -4.23E-03 | -1.00E+00 |
| ENSG000001 | GPR141 | 9.65E-01 | 9.90E-01 | 1.03E+00 | 4.51E-02 | 1.03E+00 |
| ENSG000001 | ZBTB46 | 9.65E-01 | 9.90E-01 | 1.01E+00 | 9.61E-03 | 1.01E+00 |
| ENSG000001 | CYC1 | 9.65E-01 | 9.90E-01 | 9.98E-01 | -2.81E-03 | -1.00E+00 |
| ENSG000002 | --- | 9.65E-01 | 9.90E-01 | 9.93E-01 | -9.79E-03 | -1.01E+00 |
| ENSG000001 | MTG2 | 9.65E-01 | 9.90E-01 | 1.00E+00 | 2.49E-03 | 1.00E+00 |

| | | | | | | |
|---|---|---|---|---|---|---|
| ENSG000002 | MRPL53 | 9.65E-01 | 9.90E-01 | 9.97E-01 | -4.44E-03 | -1.00E+00 |
| ENSG000002 | --- | 9.65E-01 | 9.90E-01 | 9.95E-01 | -7.83E-03 | -1.01E+00 |
| ENSG000001 | UBAP2 | 9.65E-01 | 9.90E-01 | 9.99E-01 | -2.12E-03 | -1.00E+00 |
| ENSG000001 | PYGL | 9.65E-01 | 9.90E-01 | 1.00E+00 | 2.42E-03 | 1.00E+00 |
| ENSG000001 | ADH7 | 9.66E-01 | ? | 1.01E+00 | 1.85E-02 | 1.01E+00 |
| ENSG00000C | RNF215 | 9.66E-01 | 9.90E-01 | 1.00E+00 | 4.70E-03 | 1.00E+00 |
| ENSG000002 | LINC02728 | 9.66E-01 | ? | 1.02E+00 | 2.46E-02 | 1.02E+00 |
| ENSG000002 | --- | 9.66E-01 | ? | 1.02E+00 | 2.43E-02 | 1.02E+00 |
| ENSG00000C | SLAMF7 | 9.66E-01 | 9.90E-01 | 9.95E-01 | -7.59E-03 | -1.01E+00 |
| ENSG000001 | ISM2 | 9.66E-01 | ? | 9.93E-01 | -1.07E-02 | -1.01E+00 |
| ENSG000002 | --- | 9.66E-01 | ? | 9.89E-01 | -1.53E-02 | -1.01E+00 |
| ENSG000002 | --- | 9.66E-01 | ? | 1.02E+00 | 2.64E-02 | 1.02E+00 |
| ENSG000001 | COBL | 9.66E-01 | 9.90E-01 | 9.97E-01 | -4.23E-03 | -1.00E+00 |
| ENSG000002 | --- | 9.66E-01 | ? | 1.01E+00 | 2.04E-02 | 1.01E+00 |
| ENSG000001 | LEMD2 | 9.66E-01 | 9.90E-01 | 9.98E-01 | -2.33E-03 | -1.00E+00 |
| ENSG000001 | MTERF1 | 9.66E-01 | 9.90E-01 | 1.00E+00 | 3.23E-03 | 1.00E+00 |
| ENSG00000C | P2RX5 | 9.66E-01 | 9.90E-01 | 9.96E-01 | -6.45E-03 | -1.00E+00 |
| ENSG000002 | HNRNPA1L3 | 9.66E-01 | 9.90E-01 | 9.97E-01 | -4.20E-03 | -1.00E+00 |
| ENSG000002 | MXRA7P1 | 9.67E-01 | ? | 1.02E+00 | 2.36E-02 | 1.02E+00 |
| ENSG000001 | NTRK2 | 9.67E-01 | 9.90E-01 | 9.92E-01 | -1.21E-02 | -1.01E+00 |
| ENSG000002 | PRSS2 | 9.67E-01 | ? | 1.02E+00 | 2.17E-02 | 1.02E+00 |
| ENSG000002 | FDXACB1 | 9.67E-01 | 9.90E-01 | 1.00E+00 | 6.20E-03 | 1.00E+00 |
| ENSG000001 | AIDA | 9.67E-01 | 9.90E-01 | 9.99E-01 | -1.59E-03 | -1.00E+00 |
| ENSG000002 | PRR34-AS1 | 9.67E-01 | 9.90E-01 | 9.97E-01 | -4.25E-03 | -1.00E+00 |
| ENSG000001 | BHMT2 | 9.67E-01 | ? | 9.84E-01 | -2.38E-02 | -1.02E+00 |
| ENSG000001 | INCENP | 9.67E-01 | 9.90E-01 | 9.98E-01 | -3.25E-03 | -1.00E+00 |
| ENSG000001 | GMPR | 9.67E-01 | ? | 9.90E-01 | -1.43E-02 | -1.01E+00 |
| ENSG000002 | ARL2BPP4 | 9.67E-01 | ? | 9.80E-01 | -2.87E-02 | -1.02E+00 |
| ENSG000002 | MFSD14CP | 9.67E-01 | 9.90E-01 | 1.00E+00 | 2.84E-03 | 1.00E+00 |
| ENSG00000C | C12orf4 | 9.67E-01 | 9.90E-01 | 9.98E-01 | -2.35E-03 | -1.00E+00 |
| ENSG000001 | STAC3 | 9.67E-01 | ? | 9.91E-01 | -1.32E-02 | -1.01E+00 |
| ENSG000002 | --- | 9.67E-01 | ? | 1.01E+00 | 1.76E-02 | 1.01E+00 |
| ENSG000001 | SMIM4 | 9.67E-01 | 9.90E-01 | 1.00E+00 | 4.15E-03 | 1.00E+00 |
| ENSG000002 | SCAMP4 | 9.67E-01 | 9.90E-01 | 1.00E+00 | 2.78E-03 | 1.00E+00 |
| ENSG000001 | TTLL11 | 9.67E-01 | 9.90E-01 | 9.98E-01 | -3.08E-03 | -1.00E+00 |
| ENSG000001 | OAZ3 | 9.67E-01 | 9.90E-01 | 9.95E-01 | -7.52E-03 | -1.01E+00 |
| ENSG000002 | --- | 9.67E-01 | ? | 1.01E+00 | 1.27E-02 | 1.01E+00 |
| ENSG000001 | DCP1B | 9.68E-01 | 9.90E-01 | 1.00E+00 | 3.02E-03 | 1.00E+00 |
| ENSG000001 | PTP4A3 | 9.68E-01 | ? | 1.01E+00 | 1.51E-02 | 1.01E+00 |
| ENSG000001 | C19orf18 | 9.68E-01 | ? | 1.01E+00 | 1.31E-02 | 1.01E+00 |
| ENSG00000C | OCEL1 | 9.68E-01 | 9.90E-01 | 9.96E-01 | -5.06E-03 | -1.00E+00 |
| ENSG000001 | ATP2A2 | 9.68E-01 | 9.90E-01 | 1.00E+00 | 1.62E-03 | 1.00E+00 |
| ENSG000001 | FAHD2B | 9.68E-01 | 9.90E-01 | 1.00E+00 | 4.47E-03 | 1.00E+00 |
| ENSG000002 | LINC00869 | 9.68E-01 | 9.91E-01 | 1.00E+00 | 4.37E-03 | 1.00E+00 |
| ENSG000002 | SERPINB10 | 9.68E-01 | ? | 9.85E-01 | -2.14E-02 | -1.01E+00 |

| | | | | | | |
|---|---|---|---|---|---|---|
| ENSG000001 | COMMD2 | 9.68E-01 | 9.91E-01 | 1.00E+00 | 1.84E-03 | 1.00E+00 |
| ENSG000001 | RFLNA | 9.68E-01 | ? | 9.87E-01 | -1.86E-02 | -1.01E+00 |
| ENSG000001 | FBLL1 | 9.68E-01 | ? | 1.02E+00 | 2.26E-02 | 1.02E+00 |
| ENSG000001 | BICD2 | 9.68E-01 | 9.91E-01 | 1.00E+00 | 3.12E-03 | 1.00E+00 |
| ENSG000002 | PCDHA13 | 9.68E-01 | ? | 1.01E+00 | 1.25E-02 | 1.01E+00 |
| ENSG000001 | GPR15LG | 9.68E-01 | ? | 9.84E-01 | -2.40E-02 | -1.02E+00 |
| ENSG000002 | CCDC188BP | 9.68E-01 | ? | 9.90E-01 | -1.45E-02 | -1.01E+00 |
| ENSG00000C | CLXN | 9.68E-01 | ? | 1.01E+00 | 1.11E-02 | 1.01E+00 |
| ENSG000001 | SH2D1A | 9.68E-01 | ? | 1.02E+00 | 3.04E-02 | 1.02E+00 |
| ENSG000001 | FOXD2 | 9.68E-01 | ? | 1.01E+00 | 1.59E-02 | 1.01E+00 |
| ENSG000001 | PRKAB2 | 9.68E-01 | 9.91E-01 | 1.00E+00 | 1.82E-03 | 1.00E+00 |
| ENSG000001 | ASGR1 | 9.68E-01 | ? | 1.01E+00 | 2.11E-02 | 1.01E+00 |
| ENSG00000C | MRPS34 | 9.68E-01 | 9.91E-01 | 9.98E-01 | -2.59E-03 | -1.00E+00 |
| ENSG000001 | SBNO1 | 9.69E-01 | 9.91E-01 | 9.99E-01 | -1.93E-03 | -1.00E+00 |
| ENSG000001 | FNTA | 9.69E-01 | 9.91E-01 | 1.00E+00 | 1.59E-03 | 1.00E+00 |
| ENSG000001 | PRR34 | 9.69E-01 | ? | 1.01E+00 | 1.90E-02 | 1.01E+00 |
| ENSG000001 | RGPD1 | 9.69E-01 | ? | 1.01E+00 | 1.44E-02 | 1.01E+00 |
| ENSG000002 | ADGRG1 | 9.69E-01 | 9.91E-01 | 1.00E+00 | 2.35E-03 | 1.00E+00 |
| ENSG000002 | GAS6-AS1 | 9.69E-01 | 9.91E-01 | 9.96E-01 | -6.10E-03 | -1.00E+00 |
| ENSG00000C | USP13 | 9.69E-01 | 9.91E-01 | 9.97E-01 | -4.09E-03 | -1.00E+00 |
| ENSG000002 | PPP1R14B-A: | 9.69E-01 | 9.91E-01 | 9.96E-01 | -5.85E-03 | -1.00E+00 |
| ENSG000002 | --- | 9.69E-01 | ? | 9.86E-01 | -2.08E-02 | -1.01E+00 |
| ENSG000002 | SWINGN | 9.69E-01 | 9.91E-01 | 9.94E-01 | -9.23E-03 | -1.01E+00 |
| ENSG000002 | CASC19 | 9.69E-01 | 9.91E-01 | 1.00E+00 | 2.86E-03 | 1.00E+00 |
| ENSG000002 | CLCA4-AS1 | 9.69E-01 | 9.91E-01 | 1.01E+00 | 8.54E-03 | 1.01E+00 |
| ENSG000001 | JPH2 | 9.69E-01 | ? | 9.92E-01 | -1.14E-02 | -1.01E+00 |
| ENSG000001 | HGD | 9.69E-01 | ? | 1.01E+00 | 2.04E-02 | 1.01E+00 |
| ENSG00000C | STX7 | 9.69E-01 | 9.91E-01 | 9.99E-01 | -1.68E-03 | -1.00E+00 |
| ENSG000001 | SDE2 | 9.69E-01 | 9.91E-01 | 9.99E-01 | -1.84E-03 | -1.00E+00 |
| ENSG000001 | LMF2 | 9.69E-01 | 9.91E-01 | 1.00E+00 | 4.55E-03 | 1.00E+00 |
| ENSG00000C | FGF22 | 9.69E-01 | ? | 9.83E-01 | -2.51E-02 | -1.02E+00 |
| ENSG000002 | KCNQ1OT1 | 9.70E-01 | 9.91E-01 | 9.95E-01 | -7.03E-03 | -1.00E+00 |
| ENSG000002 | FTH1P3 | 9.70E-01 | ? | 1.02E+00 | 2.40E-02 | 1.02E+00 |
| ENSG000001 | CTSS | 9.70E-01 | 9.91E-01 | 1.00E+00 | 3.76E-03 | 1.00E+00 |
| ENSG000002 | EIF4HP1 | 9.70E-01 | 9.91E-01 | 1.00E+00 | 5.66E-03 | 1.00E+00 |
| ENSG000002 | LIMD1-AS1 | 9.70E-01 | 9.91E-01 | 9.98E-01 | -3.49E-03 | -1.00E+00 |
| ENSG000002 | --- | 9.70E-01 | ? | 9.86E-01 | -2.08E-02 | -1.01E+00 |
| ENSG000001 | IGLON5 | 9.70E-01 | ? | 9.83E-01 | -2.44E-02 | -1.02E+00 |
| ENSG000001 | CDH24 | 9.70E-01 | 9.91E-01 | 9.97E-01 | -4.49E-03 | -1.00E+00 |
| ENSG000002 | MTCO1P28 | 9.70E-01 | ? | 1.02E+00 | 2.91E-02 | 1.02E+00 |
| ENSG000002 | --- | 9.70E-01 | ? | 1.01E+00 | 2.10E-02 | 1.01E+00 |
| ENSG000002 | --- | 9.70E-01 | 9.91E-01 | 9.97E-01 | -3.66E-03 | -1.00E+00 |
| ENSG000002 | EOGT-DT | 9.70E-01 | ? | 1.01E+00 | 9.96E-03 | 1.01E+00 |
| ENSG000001 | NDRG4 | 9.70E-01 | 9.91E-01 | 1.00E+00 | 5.50E-03 | 1.00E+00 |
| ENSG000001 | TRAPPC5 | 9.70E-01 | 9.91E-01 | 9.98E-01 | -3.54E-03 | -1.00E+00 |

| | | | | | | |
|---|---|---|---|---|---|---|
| ENSG000001 | RUNDC1 | 9.70E-01 | 9.91E-01 | 1.00E+00 | 1.75E-03 | 1.00E+00 |
| ENSG000001 | SMIM15 | 9.70E-01 | 9.91E-01 | 1.00E+00 | 1.84E-03 | 1.00E+00 |
| ENSG00000C | COX10 | 9.70E-01 | 9.91E-01 | 9.99E-01 | -2.10E-03 | -1.00E+00 |
| ENSG000001 | DNAH14 | 9.70E-01 | 9.91E-01 | 9.97E-01 | -4.25E-03 | -1.00E+00 |
| ENSG00000C | EED | 9.70E-01 | 9.91E-01 | 9.98E-01 | -2.27E-03 | -1.00E+00 |
| ENSG00000C | ICAM3 | 9.70E-01 | 9.91E-01 | 9.97E-01 | -4.95E-03 | -1.00E+00 |
| ENSG000002 | C1orf220 | 9.71E-01 | ? | 1.01E+00 | 1.97E-02 | 1.01E+00 |
| ENSG000001 | GTF2IRD2B | 9.71E-01 | 9.91E-01 | 1.00E+00 | 2.68E-03 | 1.00E+00 |
| ENSG000001 | NAIPP2 | 9.71E-01 | ? | 9.92E-01 | -1.13E-02 | -1.01E+00 |
| ENSG000001 | APOE | 9.71E-01 | 9.91E-01 | 1.00E+00 | 3.86E-03 | 1.00E+00 |
| ENSG000002 | CCR5AS | 9.71E-01 | ? | 1.02E+00 | 2.72E-02 | 1.02E+00 |
| ENSG000001 | HS2ST1 | 9.71E-01 | 9.91E-01 | 1.00E+00 | 2.01E-03 | 1.00E+00 |
| ENSG000001 | FBXL12 | 9.71E-01 | 9.91E-01 | 9.98E-01 | -2.47E-03 | -1.00E+00 |
| ENSG000001 | BNIP3 | 9.71E-01 | 9.91E-01 | 1.00E+00 | 1.65E-03 | 1.00E+00 |
| ENSG000001 | CDYL | 9.71E-01 | 9.91E-01 | 1.00E+00 | 1.65E-03 | 1.00E+00 |
| ENSG000001 | NR2C1 | 9.71E-01 | 9.91E-01 | 9.98E-01 | -2.17E-03 | -1.00E+00 |
| ENSG000002 | NCKAP5-AS2 | 9.71E-01 | ? | 1.02E+00 | 2.66E-02 | 1.02E+00 |
| ENSG000001 | ZNF563 | 9.71E-01 | 9.91E-01 | 1.00E+00 | 5.33E-03 | 1.00E+00 |
| ENSG000001 | PGF | 9.71E-01 | 9.91E-01 | 1.00E+00 | 3.09E-03 | 1.00E+00 |
| ENSG000001 | OSMR | 9.71E-01 | 9.91E-01 | 1.00E+00 | 1.98E-03 | 1.00E+00 |
| ENSG000002 | --- | 9.71E-01 | ? | 1.01E+00 | 1.11E-02 | 1.01E+00 |
| ENSG000001 | NDUFB6 | 9.71E-01 | 9.91E-01 | 9.98E-01 | -2.73E-03 | -1.00E+00 |
| ENSG000001 | C3orf70 | 9.71E-01 | ? | 9.86E-01 | -2.02E-02 | -1.01E+00 |
| ENSG000001 | CDC123 | 9.72E-01 | 9.91E-01 | 9.99E-01 | -1.69E-03 | -1.00E+00 |
| ENSG00000C | GPKOW | 9.72E-01 | 9.91E-01 | 9.98E-01 | -2.43E-03 | -1.00E+00 |
| ENSG00000C | SPEN | 9.72E-01 | 9.91E-01 | 1.00E+00 | 2.56E-03 | 1.00E+00 |
| ENSG000001 | GRID1 | 9.72E-01 | ? | 1.01E+00 | 1.60E-02 | 1.01E+00 |
| ENSG000001 | PRMT9 | 9.72E-01 | 9.91E-01 | 9.98E-01 | -2.62E-03 | -1.00E+00 |
| ENSG000001 | FBXO3 | 9.72E-01 | 9.91E-01 | 9.99E-01 | -1.85E-03 | -1.00E+00 |
| ENSG000001 | EIF2B1 | 9.72E-01 | 9.91E-01 | 1.00E+00 | 1.52E-03 | 1.00E+00 |
| ENSG000001 | SELENOF | 9.72E-01 | 9.91E-01 | 1.00E+00 | 1.37E-03 | 1.00E+00 |
| ENSG000002 | ZBED3-AS1 | 9.72E-01 | ? | 1.01E+00 | 1.66E-02 | 1.01E+00 |
| ENSG000001 | LANCL2 | 9.72E-01 | 9.91E-01 | 9.99E-01 | -1.61E-03 | -1.00E+00 |
| ENSG000002 | --- | 9.72E-01 | ? | 1.01E+00 | 1.28E-02 | 1.01E+00 |
| ENSG000001 | RBCK1 | 9.72E-01 | 9.91E-01 | 9.98E-01 | -2.63E-03 | -1.00E+00 |
| ENSG000001 | ZNF585A | 9.72E-01 | 9.91E-01 | 9.98E-01 | -2.21E-03 | -1.00E+00 |
| ENSG000002 | PRKAR2A-AS | 9.72E-01 | ? | 9.91E-01 | -1.28E-02 | -1.01E+00 |
| ENSG00000C | SBNO2 | 9.72E-01 | 9.91E-01 | 9.98E-01 | -3.29E-03 | -1.00E+00 |
| ENSG000002 | TGIF2-RAB5I | 9.72E-01 | ? | 1.02E+00 | 2.35E-02 | 1.02E+00 |
| ENSG000001 | CCR7 | 9.72E-01 | ? | 9.91E-01 | -1.31E-02 | -1.01E+00 |
| ENSG000002 | ARHGEF25 | 9.73E-01 | ? | 9.87E-01 | -1.95E-02 | -1.01E+00 |
| ENSG000002 | LINC02166 | 9.73E-01 | ? | 1.02E+00 | 3.41E-02 | 1.02E+00 |
| ENSG000001 | PPP6R2 | 9.73E-01 | 9.92E-01 | 9.98E-01 | -2.37E-03 | -1.00E+00 |
| ENSG000001 | CHN2 | 9.73E-01 | ? | 9.90E-01 | -1.43E-02 | -1.01E+00 |
| ENSG000002 | --- | 9.73E-01 | ? | 1.01E+00 | 1.01E-02 | 1.01E+00 |

| | | | | | | |
|---|---|---|---|---|---|---|
| ENSG000001 | SH3RF3 | 9.73E-01 | 9.92E-01 | 1.01E+00 | 8.21E-03 | 1.01E+00 |
| ENSG000001 | SMS | 9.73E-01 | 9.92E-01 | 9.99E-01 | -1.26E-03 | -1.00E+00 |
| ENSG000001 | DUSP22 | 9.73E-01 | 9.92E-01 | 1.00E+00 | 1.67E-03 | 1.00E+00 |
| ENSG00000C | NNAT | 9.73E-01 | 9.92E-01 | 1.00E+00 | 5.63E-03 | 1.00E+00 |
| ENSG000002 | DANCR | 9.73E-01 | 9.92E-01 | 1.00E+00 | 2.39E-03 | 1.00E+00 |
| ENSG000001 | FBXO24 | 9.73E-01 | ? | 1.01E+00 | 1.87E-02 | 1.01E+00 |
| ENSG000002 | --- | 9.73E-01 | ? | 9.92E-01 | -1.19E-02 | -1.01E+00 |
| ENSG000002 | HCCS-DT | 9.73E-01 | ? | 9.87E-01 | -1.95E-02 | -1.01E+00 |
| ENSG000002 | MIR200CHG | 9.73E-01 | 9.92E-01 | 1.00E+00 | 5.16E-03 | 1.00E+00 |
| ENSG000001 | SELENOT | 9.74E-01 | 9.92E-01 | 1.00E+00 | 1.39E-03 | 1.00E+00 |
| ENSG000001 | NAE1 | 9.74E-01 | 9.92E-01 | 9.99E-01 | -1.48E-03 | -1.00E+00 |
| ENSG000002 | --- | 9.74E-01 | 9.92E-01 | 9.97E-01 | -4.51E-03 | -1.00E+00 |
| ENSG000001 | KIF13A | 9.74E-01 | 9.92E-01 | 9.99E-01 | -1.60E-03 | -1.00E+00 |
| ENSG000002 | --- | 9.74E-01 | ? | 9.92E-01 | -1.23E-02 | -1.01E+00 |
| ENSG000001 | ADAM21 | 9.74E-01 | 9.92E-01 | 1.00E+00 | 4.90E-03 | 1.00E+00 |
| ENSG000002 | MELTF-AS1 | 9.74E-01 | ? | 1.01E+00 | 8.38E-03 | 1.01E+00 |
| ENSG000001 | ETNK1 | 9.74E-01 | 9.92E-01 | 9.98E-01 | -2.44E-03 | -1.00E+00 |
| ENSG000002 | LGMNP1 | 9.74E-01 | ? | 9.86E-01 | -1.98E-02 | -1.01E+00 |
| ENSG000001 | MMACHC | 9.74E-01 | 9.92E-01 | 9.99E-01 | -2.08E-03 | -1.00E+00 |
| ENSG000001 | MARVELD2 | 9.74E-01 | 9.92E-01 | 9.99E-01 | -1.42E-03 | -1.00E+00 |
| ENSG000002 | --- | 9.74E-01 | ? | 9.91E-01 | -1.30E-02 | -1.01E+00 |
| ENSG000002 | LINC00621 | 9.74E-01 | ? | 1.01E+00 | 1.03E-02 | 1.01E+00 |
| ENSG000001 | MYO6 | 9.74E-01 | 9.92E-01 | 1.00E+00 | 1.95E-03 | 1.00E+00 |
| ENSG000001 | EMC7 | 9.74E-01 | 9.92E-01 | 1.00E+00 | 1.50E-03 | 1.00E+00 |
| ENSG000001 | NXNL2 | 9.74E-01 | ? | 1.01E+00 | 1.30E-02 | 1.01E+00 |
| ENSG000001 | GDPD3 | 9.74E-01 | ? | 9.91E-01 | -1.28E-02 | -1.01E+00 |
| ENSG000002 | LINC02603 | 9.74E-01 | ? | 1.02E+00 | 2.60E-02 | 1.02E+00 |
| ENSG000002 | CFB | 9.74E-01 | 9.92E-01 | 1.01E+00 | 1.33E-02 | 1.01E+00 |
| ENSG000001 | ITPKA | 9.75E-01 | ? | 1.01E+00 | 1.83E-02 | 1.01E+00 |
| ENSG000001 | GOLT1A | 9.75E-01 | ? | 9.91E-01 | -1.27E-02 | -1.01E+00 |
| ENSG000002 | WAKMAR2 | 9.75E-01 | 9.92E-01 | 9.95E-01 | -6.69E-03 | -1.00E+00 |
| ENSG000002 | CNTNAP3P2 | 9.75E-01 | 9.92E-01 | 1.01E+00 | 9.21E-03 | 1.01E+00 |
| ENSG000001 | LMO7 | 9.75E-01 | 9.92E-01 | 1.00E+00 | 3.71E-03 | 1.00E+00 |
| ENSG000001 | THAP2 | 9.75E-01 | 9.92E-01 | 1.00E+00 | 3.16E-03 | 1.00E+00 |
| ENSG000002 | --- | 9.75E-01 | 9.92E-01 | 1.00E+00 | 2.89E-03 | 1.00E+00 |
| ENSG000001 | GPR63 | 9.75E-01 | 9.93E-01 | 1.00E+00 | 4.08E-03 | 1.00E+00 |
| ENSG000002 | PABPC1P4 | 9.75E-01 | ? | 9.94E-01 | -8.00E-03 | -1.01E+00 |
| ENSG00000C | ACTB | 9.75E-01 | 9.93E-01 | 9.99E-01 | -1.19E-03 | -1.00E+00 |
| ENSG000002 | LINC-PINT | 9.75E-01 | 9.93E-01 | 1.00E+00 | 4.82E-03 | 1.00E+00 |
| ENSG000002 | --- | 9.75E-01 | ? | 1.01E+00 | 1.44E-02 | 1.01E+00 |
| ENSG000001 | RASSF8 | 9.75E-01 | 9.93E-01 | 9.98E-01 | -2.30E-03 | -1.00E+00 |
| ENSG000001 | MTTP | 9.76E-01 | ? | 1.01E+00 | 1.16E-02 | 1.01E+00 |
| ENSG000002 | CLMAT3 | 9.76E-01 | ? | 9.87E-01 | -1.88E-02 | -1.01E+00 |
| ENSG000002 | PGK1P1 | 9.76E-01 | ? | 1.01E+00 | 1.93E-02 | 1.01E+00 |
| ENSG000001 | MON1A | 9.76E-01 | 9.93E-01 | 9.98E-01 | -2.50E-03 | -1.00E+00 |

| | | | | | | |
|---|---|---|---|---|---|---|
| ENSG000001 | SPTSSA | 9.76E-01 | 9.93E-01 | 1.00E+00 | 2.66E-03 | 1.00E+00 |
| ENSG000002 | --- | 9.76E-01 | 9.93E-01 | 1.00E+00 | 6.37E-03 | 1.00E+00 |
| ENSG000001 | ANXA2 | 9.76E-01 | 9.93E-01 | 9.99E-01 | -7.88E-04 | -1.00E+00 |
| ENSG000001 | PLAAT3 | 9.76E-01 | ? | 1.01E+00 | 9.87E-03 | 1.01E+00 |
| ENSG000001 | PXDC1 | 9.76E-01 | 9.93E-01 | 1.00E+00 | 1.22E-03 | 1.00E+00 |
| ENSG000002 | EMSY-DT | 9.76E-01 | 9.93E-01 | 1.00E+00 | 6.14E-03 | 1.00E+00 |
| ENSG000002 | --- | 9.76E-01 | ? | 1.01E+00 | 9.36E-03 | 1.01E+00 |
| ENSG000002 | --- | 9.76E-01 | ? | 1.01E+00 | 9.64E-03 | 1.01E+00 |
| ENSG000002 | --- | 9.76E-01 | ? | 1.01E+00 | 1.15E-02 | 1.01E+00 |
| ENSG00000C | ELOVL5 | 9.76E-01 | 9.93E-01 | 1.00E+00 | 8.67E-04 | 1.00E+00 |
| ENSG000002 | --- | 9.76E-01 | ? | 9.90E-01 | -1.48E-02 | -1.01E+00 |
| ENSG000001 | UBA7 | 9.76E-01 | 9.93E-01 | 1.00E+00 | 3.99E-03 | 1.00E+00 |
| ENSG000001 | PARP10 | 9.76E-01 | 9.93E-01 | 1.00E+00 | 3.23E-03 | 1.00E+00 |
| ENSG000002 | --- | 9.77E-01 | ? | 1.01E+00 | 1.21E-02 | 1.01E+00 |
| ENSG000002 | --- | 9.77E-01 | ? | 9.93E-01 | -1.04E-02 | -1.01E+00 |
| ENSG000001 | EFL1 | 9.77E-01 | 9.93E-01 | 1.00E+00 | 1.38E-03 | 1.00E+00 |
| ENSG000002 | CBX3P9 | 9.77E-01 | ? | 9.89E-01 | -1.63E-02 | -1.01E+00 |
| ENSG000001 | NPC1 | 9.77E-01 | 9.93E-01 | 9.99E-01 | -1.15E-03 | -1.00E+00 |
| ENSG000001 | SAMD8 | 9.77E-01 | 9.93E-01 | 1.00E+00 | 1.31E-03 | 1.00E+00 |
| ENSG000002 | LINC03009 | 9.77E-01 | 9.93E-01 | 1.00E+00 | 3.46E-03 | 1.00E+00 |
| ENSG000002 | SUPT4H1 | 9.77E-01 | 9.93E-01 | 1.00E+00 | 1.37E-03 | 1.00E+00 |
| ENSG000002 | STAM-DT | 9.77E-01 | ? | 1.01E+00 | 1.03E-02 | 1.01E+00 |
| ENSG000001 | CST3 | 9.77E-01 | 9.93E-01 | 9.99E-01 | -1.79E-03 | -1.00E+00 |
| ENSG000001 | IRF3 | 9.77E-01 | 9.93E-01 | 1.00E+00 | 2.47E-03 | 1.00E+00 |
| ENSG000001 | PPP1R18 | 9.77E-01 | 9.93E-01 | 9.99E-01 | -1.18E-03 | -1.00E+00 |
| ENSG000002 | --- | 9.77E-01 | ? | 9.94E-01 | -9.15E-03 | -1.01E+00 |
| ENSG000001 | MARCHF8 | 9.77E-01 | 9.93E-01 | 1.00E+00 | 1.62E-03 | 1.00E+00 |
| ENSG000001 | VIPAS39 | 9.77E-01 | 9.93E-01 | 1.00E+00 | 1.75E-03 | 1.00E+00 |
| ENSG00000C | STYK1 | 9.77E-01 | 9.93E-01 | 1.00E+00 | 2.75E-03 | 1.00E+00 |
| ENSG000002 | C1orf210 | 9.77E-01 | 9.93E-01 | 9.96E-01 | -5.34E-03 | -1.00E+00 |
| ENSG000002 | --- | 9.77E-01 | ? | 1.01E+00 | 2.03E-02 | 1.01E+00 |
| ENSG000001 | TCF12 | 9.77E-01 | 9.93E-01 | 1.00E+00 | 1.05E-03 | 1.00E+00 |
| ENSG000002 | MALT1-AS1 | 9.77E-01 | ? | 9.85E-01 | -2.23E-02 | -1.02E+00 |
| ENSG000002 | CD44-DT | 9.78E-01 | ? | 9.95E-01 | -7.80E-03 | -1.01E+00 |
| ENSG000001 | STK3 | 9.78E-01 | 9.93E-01 | 9.99E-01 | -1.69E-03 | -1.00E+00 |
| ENSG000002 | NPBWR1 | 9.78E-01 | ? | 1.01E+00 | 1.29E-02 | 1.01E+00 |
| ENSG000001 | PDCD11 | 9.78E-01 | 9.93E-01 | 9.99E-01 | -1.05E-03 | -1.00E+00 |
| ENSG000001 | RALGPS1 | 9.78E-01 | 9.93E-01 | 9.97E-01 | -4.77E-03 | -1.00E+00 |
| ENSG000001 | ZNF341 | 9.78E-01 | 9.93E-01 | 1.00E+00 | 2.34E-03 | 1.00E+00 |
| ENSG000001 | LRRC34 | 9.78E-01 | 9.93E-01 | 1.01E+00 | 7.76E-03 | 1.01E+00 |
| ENSG000002 | SMIM26 | 9.78E-01 | 9.93E-01 | 1.00E+00 | 1.96E-03 | 1.00E+00 |
| ENSG000002 | HMGB3P6 | 9.78E-01 | ? | 1.01E+00 | 1.06E-02 | 1.01E+00 |
| ENSG000002 | MYO15B | 9.78E-01 | ? | 9.94E-01 | -9.37E-03 | -1.01E+00 |
| ENSG000001 | CPQ | 9.78E-01 | 9.93E-01 | 9.98E-01 | -2.37E-03 | -1.00E+00 |
| ENSG000001 | TRAPPC11 | 9.78E-01 | 9.93E-01 | 9.99E-01 | -1.44E-03 | -1.00E+00 |

| | | | | | | | |
|---|---|---|---|---|---|---|---|
| ENSG000002 | INPP5B | 9.78E-01 | | 9.93E-01 | 9.98E-01 | -2.93E-03 | -1.00E+00 |
| ENSG000002 | LLPHP3 | 9.78E-01 | ? | | 1.01E+00 | 1.04E-02 | 1.01E+00 |
| ENSG00000C | NISCH | 9.78E-01 | | 9.93E-01 | 9.99E-01 | -1.98E-03 | -1.00E+00 |
| ENSG000001 | MLIP | 9.78E-01 | ? | | 9.90E-01 | -1.42E-02 | -1.01E+00 |
| ENSG000001 | ADCY4 | 9.78E-01 | | 9.93E-01 | 1.00E+00 | 3.19E-03 | 1.00E+00 |
| ENSG000002 | ARL17B | 9.78E-01 | ? | | 1.01E+00 | 1.24E-02 | 1.01E+00 |
| ENSG000001 | FASTKD3 | 9.78E-01 | | 9.93E-01 | 1.00E+00 | 2.19E-03 | 1.00E+00 |
| ENSG000001 | GPC3 | 9.78E-01 | ? | | 1.01E+00 | 8.61E-03 | 1.01E+00 |
| ENSG000001 | SECTM1 | 9.78E-01 | ? | | 1.01E+00 | 8.72E-03 | 1.01E+00 |
| ENSG000002 | --- | 9.78E-01 | | 9.93E-01 | 1.00E+00 | 4.92E-03 | 1.00E+00 |
| ENSG000001 | MFSD3 | 9.78E-01 | | 9.93E-01 | 1.00E+00 | 3.83E-03 | 1.00E+00 |
| ENSG000001 | ENOX2 | 9.79E-01 | | 9.93E-01 | 1.00E+00 | 1.51E-03 | 1.00E+00 |
| ENSG000001 | UTP20 | 9.79E-01 | | 9.93E-01 | 9.99E-01 | -2.03E-03 | -1.00E+00 |
| ENSG000002 | --- | 9.79E-01 | ? | | 9.89E-01 | -1.60E-02 | -1.01E+00 |
| ENSG00000C | ELOVL1 | 9.79E-01 | | 9.93E-01 | 1.00E+00 | 1.16E-03 | 1.00E+00 |
| ENSG000001 | METTL17 | 9.79E-01 | | 9.93E-01 | 1.00E+00 | 1.51E-03 | 1.00E+00 |
| ENSG000002 | CCDC183 | 9.79E-01 | ? | | 9.90E-01 | -1.48E-02 | -1.01E+00 |
| ENSG000002 | --- | 9.79E-01 | ? | | 9.88E-01 | -1.72E-02 | -1.01E+00 |
| ENSG000001 | E2F5 | 9.79E-01 | | 9.93E-01 | 1.00E+00 | 2.43E-03 | 1.00E+00 |
| ENSG000001 | FCHSD2 | 9.79E-01 | | 9.93E-01 | 9.99E-01 | -1.60E-03 | -1.00E+00 |
| ENSG000001 | USP38 | 9.79E-01 | | 9.93E-01 | 9.99E-01 | -1.14E-03 | -1.00E+00 |
| ENSG000001 | PPP3CB | 9.79E-01 | | 9.93E-01 | 9.99E-01 | -1.27E-03 | -1.00E+00 |
| ENSG000001 | MBTPS1 | 9.79E-01 | | 9.93E-01 | 1.00E+00 | 1.22E-03 | 1.00E+00 |
| ENSG000002 | CICP16 | 9.79E-01 | ? | | 1.01E+00 | 1.79E-02 | 1.01E+00 |
| ENSG000001 | HOMER2 | 9.80E-01 | | 9.94E-01 | 1.00E+00 | 1.42E-03 | 1.00E+00 |
| ENSG000001 | ZNF610 | 9.80E-01 | | 9.94E-01 | 9.98E-01 | -3.01E-03 | -1.00E+00 |
| ENSG000001 | DNASE1L3 | 9.80E-01 | ? | | 9.92E-01 | -1.17E-02 | -1.01E+00 |
| ENSG000002 | --- | 9.80E-01 | ? | | 1.01E+00 | 1.40E-02 | 1.01E+00 |
| ENSG000002 | CERNA2 | 9.80E-01 | | 9.94E-01 | 9.98E-01 | -3.59E-03 | -1.00E+00 |
| ENSG000001 | EFNB3 | 9.80E-01 | ? | | 9.95E-01 | -7.70E-03 | -1.01E+00 |
| ENSG00000C | KCNAB2 | 9.80E-01 | | 9.94E-01 | 1.00E+00 | 3.79E-03 | 1.00E+00 |
| ENSG000001 | STX11 | 9.80E-01 | | 9.94E-01 | 1.00E+00 | 4.67E-03 | 1.00E+00 |
| ENSG000002 | --- | 9.80E-01 | ? | | 9.94E-01 | -8.69E-03 | -1.01E+00 |
| ENSG000002 | NUDT16-DT | 9.80E-01 | ? | | 1.01E+00 | 9.50E-03 | 1.01E+00 |
| ENSG000001 | FIGNL1 | 9.81E-01 | | 9.94E-01 | 1.00E+00 | 1.71E-03 | 1.00E+00 |
| ENSG000001 | CDK5 | 9.81E-01 | | 9.94E-01 | 9.98E-01 | -2.71E-03 | -1.00E+00 |
| ENSG000002 | --- | 9.81E-01 | ? | | 1.01E+00 | 1.39E-02 | 1.01E+00 |
| ENSG000001 | EDC3 | 9.81E-01 | | 9.94E-01 | 1.00E+00 | 1.21E-03 | 1.00E+00 |
| ENSG000002 | HCG25 | 9.81E-01 | | 9.94E-01 | 1.00E+00 | 5.02E-03 | 1.00E+00 |
| ENSG000001 | LAMA5 | 9.81E-01 | | 9.94E-01 | 1.00E+00 | 3.37E-03 | 1.00E+00 |
| ENSG000002 | RPS14P8 | 9.81E-01 | ? | | 1.01E+00 | 1.53E-02 | 1.01E+00 |
| ENSG000001 | CATSPER1 | 9.81E-01 | ? | | 9.89E-01 | -1.60E-02 | -1.01E+00 |
| ENSG000001 | FMC1 | 9.81E-01 | | 9.94E-01 | 9.96E-01 | -5.47E-03 | -1.00E+00 |
| ENSG000001 | ANKS3 | 9.81E-01 | | 9.94E-01 | 9.98E-01 | -2.46E-03 | -1.00E+00 |
| ENSG000002 | FAM106A | 9.81E-01 | | 9.94E-01 | 1.00E+00 | 4.54E-03 | 1.00E+00 |

| | | | | | | |
|---|---|---|---|---|---|---|
| ENSG000001 | RELL1 | 9.81E-01 | 9.94E-01 | 9.99E-01 | -1.69E-03 | -1.00E+00 |
| ENSG000001 | ASTN1 | 9.81E-01 | ? | 1.01E+00 | 1.27E-02 | 1.01E+00 |
| ENSG000000 | PABPC1 | 9.81E-01 | 9.94E-01 | 9.99E-01 | -1.39E-03 | -1.00E+00 |
| ENSG000002 | --- | 9.81E-01 | ? | 1.01E+00 | 9.78E-03 | 1.01E+00 |
| ENSG000002 | C16orf95-DT | 9.81E-01 | ? | 9.89E-01 | -1.56E-02 | -1.01E+00 |
| ENSG000001 | WWOX | 9.81E-01 | 9.94E-01 | 1.00E+00 | 2.74E-03 | 1.00E+00 |
| ENSG000000 | SNX5 | 9.82E-01 | 9.94E-01 | 1.00E+00 | 1.60E-03 | 1.00E+00 |
| ENSG000001 | AGT | 9.82E-01 | ? | 1.01E+00 | 7.45E-03 | 1.01E+00 |
| ENSG000002 | LINC00239 | 9.82E-01 | ? | 1.01E+00 | 1.26E-02 | 1.01E+00 |
| ENSG000002 | AMY2B | 9.82E-01 | 9.94E-01 | 1.00E+00 | 4.84E-03 | 1.00E+00 |
| ENSG000001 | SERHL2 | 9.82E-01 | 9.94E-01 | 1.00E+00 | 5.66E-03 | 1.00E+00 |
| ENSG000001 | ZNF707 | 9.82E-01 | 9.94E-01 | 9.98E-01 | -2.64E-03 | -1.00E+00 |
| ENSG000001 | ZBTB24 | 9.82E-01 | 9.94E-01 | 9.99E-01 | -1.42E-03 | -1.00E+00 |
| ENSG000002 | --- | 9.82E-01 | ? | 9.89E-01 | -1.61E-02 | -1.01E+00 |
| ENSG000001 | C3 | 9.82E-01 | 9.94E-01 | 1.00E+00 | 3.68E-03 | 1.00E+00 |
| ENSG000001 | DIS3L2 | 9.82E-01 | 9.94E-01 | 1.00E+00 | 1.10E-03 | 1.00E+00 |
| ENSG000001 | KRT18P59 | 9.82E-01 | ? | 1.01E+00 | 1.29E-02 | 1.01E+00 |
| ENSG000001 | LRRC45 | 9.82E-01 | 9.94E-01 | 9.98E-01 | -3.21E-03 | -1.00E+00 |
| ENSG000001 | MAP3K21 | 9.82E-01 | 9.94E-01 | 9.99E-01 | -1.30E-03 | -1.00E+00 |
| ENSG000001 | ZNF205 | 9.82E-01 | 9.94E-01 | 1.00E+00 | 2.32E-03 | 1.00E+00 |
| ENSG000001 | MLF1 | 9.82E-01 | 9.94E-01 | 9.99E-01 | -2.05E-03 | -1.00E+00 |
| ENSG000001 | IFT46 | 9.82E-01 | 9.94E-01 | 9.99E-01 | -1.36E-03 | -1.00E+00 |
| ENSG000001 | DPY19L4 | 9.82E-01 | 9.94E-01 | 1.00E+00 | 1.19E-03 | 1.00E+00 |
| ENSG000001 | SHCBP1 | 9.82E-01 | 9.94E-01 | 1.00E+00 | 3.65E-03 | 1.00E+00 |
| ENSG000001 | CZIB | 9.82E-01 | 9.94E-01 | 1.00E+00 | 1.31E-03 | 1.00E+00 |
| ENSG000001 | NXF1 | 9.83E-01 | 9.94E-01 | 1.00E+00 | 1.35E-03 | 1.00E+00 |
| ENSG000001 | CENPC | 9.83E-01 | 9.94E-01 | 9.99E-01 | -1.75E-03 | -1.00E+00 |
| ENSG000002 | GNL1 | 9.83E-01 | 9.94E-01 | 9.99E-01 | -1.16E-03 | -1.00E+00 |
| ENSG000002 | RPL22P1 | 9.83E-01 | 9.94E-01 | 9.96E-01 | -5.11E-03 | -1.00E+00 |
| ENSG000000 | FBXL3 | 9.83E-01 | 9.94E-01 | 9.99E-01 | -9.63E-04 | -1.00E+00 |
| ENSG000002 | --- | 9.83E-01 | ? | 9.96E-01 | -5.39E-03 | -1.00E+00 |
| ENSG000002 | LINC01266 | 9.83E-01 | ? | 9.91E-01 | -1.37E-02 | -1.01E+00 |
| ENSG000000 | CBARP | 9.83E-01 | 9.95E-01 | 9.98E-01 | -2.74E-03 | -1.00E+00 |
| ENSG000001 | C1orf109 | 9.83E-01 | 9.95E-01 | 9.99E-01 | -1.39E-03 | -1.00E+00 |
| ENSG000001 | DTNB | 9.83E-01 | 9.95E-01 | 9.99E-01 | -2.08E-03 | -1.00E+00 |
| ENSG000002 | --- | 9.83E-01 | ? | 9.96E-01 | -5.69E-03 | -1.00E+00 |
| ENSG000001 | ZNF547 | 9.83E-01 | 9.95E-01 | 9.98E-01 | -2.79E-03 | -1.00E+00 |
| ENSG000002 | --- | 9.84E-01 | 9.95E-01 | 9.97E-01 | -4.40E-03 | -1.00E+00 |
| ENSG000000 | CTNS | 9.84E-01 | 9.95E-01 | 1.00E+00 | 1.85E-03 | 1.00E+00 |
| ENSG000002 | --- | 9.84E-01 | ? | 1.01E+00 | 1.16E-02 | 1.01E+00 |
| ENSG000001 | SRD5A3 | 9.84E-01 | 9.95E-01 | 1.00E+00 | 1.17E-03 | 1.00E+00 |
| ENSG000001 | FYB2 | 9.84E-01 | ? | 9.92E-01 | -1.17E-02 | -1.01E+00 |
| ENSG000002 | CEP83-DT | 9.84E-01 | ? | 9.92E-01 | -1.09E-02 | -1.01E+00 |
| ENSG000002 | LINC01331 | 9.84E-01 | ? | 1.01E+00 | 1.34E-02 | 1.01E+00 |
| ENSG000001 | C19orf53 | 9.84E-01 | 9.95E-01 | 9.99E-01 | -9.35E-04 | -1.00E+00 |

| | | | | | | |
|---|---|---|---|---|---|---|
| ENSG000001 | EID2 | 9.84E-01 | 9.95E-01 | 9.99E-01 | -1.55E-03 | -1.00E+00 |
| ENSG000001 | APP | 9.84E-01 | 9.95E-01 | 1.00E+00 | -6.61E-04 | -1.00E+00 |
| ENSG000001 | PCBP2 | 9.84E-01 | 9.95E-01 | 1.00E+00 | 6.78E-04 | 1.00E+00 |
| ENSG000001 | DDIT4 | 9.84E-01 | 9.95E-01 | 1.00E+00 | 2.17E-03 | 1.00E+00 |
| ENSG000002 | --- | 9.85E-01 | ? | 1.00E+00 | 4.91E-03 | 1.00E+00 |
| ENSG000002 | --- | 9.85E-01 | 9.95E-01 | 1.00E+00 | 3.85E-03 | 1.00E+00 |
| ENSG000001 | RIN3 | 9.85E-01 | 9.95E-01 | 1.00E+00 | 1.36E-03 | 1.00E+00 |
| ENSG000002 | RPS10-NUDT | 9.85E-01 | 9.95E-01 | 1.00E+00 | 1.56E-03 | 1.00E+00 |
| ENSG000001 | TSNAXIP1 | 9.85E-01 | ? | 1.00E+00 | 4.93E-03 | 1.00E+00 |
| ENSG000001 | PIPOX | 9.85E-01 | ? | 1.01E+00 | 1.28E-02 | 1.01E+00 |
| ENSG000002 | --- | 9.85E-01 | ? | 1.01E+00 | 8.35E-03 | 1.01E+00 |
| ENSG000001 | CCRL2 | 9.85E-01 | ? | 1.00E+00 | 4.69E-03 | 1.00E+00 |
| ENSG000002 | --- | 9.85E-01 | 9.96E-01 | 1.00E+00 | 3.33E-03 | 1.00E+00 |
| ENSG000001 | RNF25 | 9.85E-01 | 9.96E-01 | 1.00E+00 | 1.34E-03 | 1.00E+00 |
| ENSG000002 | KLHL23 | 9.85E-01 | 9.96E-01 | 9.99E-01 | -1.77E-03 | -1.00E+00 |
| ENSG000002 | --- | 9.85E-01 | ? | 1.00E+00 | 5.03E-03 | 1.00E+00 |
| ENSG000001 | FOXO1 | 9.85E-01 | 9.96E-01 | 1.00E+00 | 1.19E-03 | 1.00E+00 |
| ENSG000001 | NOL11 | 9.85E-01 | 9.96E-01 | 1.00E+00 | 7.52E-04 | 1.00E+00 |
| ENSG000002 | ST13P15 | 9.85E-01 | ? | 9.92E-01 | -1.14E-02 | -1.01E+00 |
| ENSG000002 | --- | 9.86E-01 | ? | 1.01E+00 | 9.37E-03 | 1.01E+00 |
| ENSG000000 | PHGDH | 9.86E-01 | 9.96E-01 | 1.00E+00 | 2.55E-03 | 1.00E+00 |
| ENSG000001 | CRACR2B | 9.86E-01 | ? | 9.96E-01 | -5.23E-03 | -1.00E+00 |
| ENSG000002 | LST1 | 9.86E-01 | ? | 1.01E+00 | 1.09E-02 | 1.01E+00 |
| ENSG000000 | DNAJC11 | 9.86E-01 | 9.96E-01 | 1.00E+00 | 5.80E-04 | 1.00E+00 |
| ENSG000002 | --- | 9.86E-01 | ? | 1.01E+00 | 1.29E-02 | 1.01E+00 |
| ENSG000002 | CCDC7 | 9.86E-01 | 9.96E-01 | 9.97E-01 | -3.80E-03 | -1.00E+00 |
| ENSG000001 | AIF1L | 9.86E-01 | 9.96E-01 | 1.00E+00 | 4.12E-03 | 1.00E+00 |
| ENSG000001 | EME2 | 9.86E-01 | 9.96E-01 | 1.00E+00 | 1.83E-03 | 1.00E+00 |
| ENSG000001 | PDK1 | 9.86E-01 | 9.96E-01 | 1.00E+00 | 1.01E-03 | 1.00E+00 |
| ENSG000001 | ZMYND19 | 9.86E-01 | 9.96E-01 | 9.99E-01 | -1.24E-03 | -1.00E+00 |
| ENSG000002 | --- | 9.86E-01 | ? | 1.01E+00 | 1.17E-02 | 1.01E+00 |
| ENSG000001 | ADRM1 | 9.87E-01 | 9.96E-01 | 9.99E-01 | -9.39E-04 | -1.00E+00 |
| ENSG000002 | PCMTD2 | 9.87E-01 | 9.96E-01 | 1.00E+00 | 1.06E-03 | 1.00E+00 |
| ENSG000001 | GADD45A | 9.87E-01 | 9.96E-01 | 9.99E-01 | -1.55E-03 | -1.00E+00 |
| ENSG000001 | PMM1 | 9.87E-01 | 9.96E-01 | 9.99E-01 | -1.49E-03 | -1.00E+00 |
| ENSG000001 | ARHGEF7 | 9.87E-01 | 9.96E-01 | 9.99E-01 | -7.29E-04 | -1.00E+00 |
| ENSG000002 | MC1R | 9.87E-01 | 9.96E-01 | 9.98E-01 | -2.28E-03 | -1.00E+00 |
| ENSG000001 | RNASEH1 | 9.87E-01 | 9.96E-01 | 1.00E+00 | 6.71E-04 | 1.00E+00 |
| ENSG000000 | VPS9D1 | 9.87E-01 | 9.96E-01 | 1.00E+00 | 2.14E-03 | 1.00E+00 |
| ENSG000002 | VDAC1P2 | 9.87E-01 | ? | 1.01E+00 | 1.02E-02 | 1.01E+00 |
| ENSG000001 | DMPK | 9.87E-01 | 9.96E-01 | 1.00E+00 | 1.84E-03 | 1.00E+00 |
| ENSG000001 | USP5 | 9.87E-01 | 9.96E-01 | 1.00E+00 | 1.06E-03 | 1.00E+00 |
| ENSG000002 | CFAP45 | 9.87E-01 | ? | 9.96E-01 | -6.22E-03 | -1.00E+00 |
| ENSG000001 | POLE | 9.87E-01 | 9.96E-01 | 1.00E+00 | 1.12E-03 | 1.00E+00 |
| ENSG000001 | LZTS2 | 9.87E-01 | 9.96E-01 | 1.00E+00 | 1.27E-03 | 1.00E+00 |

| | | | | | | |
|---|---|---|---|---|---|---|
| ENSG000002 | ANKHD1-EIF4 | 9.87E-01 | 9.96E-01 | 9.99E-01 | -1.10E-03 | -1.00E+00 |
| ENSG000001 | THUMPD3 | 9.87E-01 | 9.96E-01 | 1.00E+00 | 7.29E-04 | 1.00E+00 |
| ENSG000002 | --- | 9.87E-01 | ? | 1.00E+00 | 5.14E-03 | 1.00E+00 |
| ENSG000001 | RAB1B | 9.87E-01 | 9.96E-01 | 9.99E-01 | -7.74E-04 | -1.00E+00 |
| ENSG000002 | LINC01213 | 9.87E-01 | ? | 1.01E+00 | 1.05E-02 | 1.01E+00 |
| ENSG000001 | KIF5C | 9.88E-01 | ? | 1.01E+00 | 7.33E-03 | 1.01E+00 |
| ENSG000002 | --- | 9.88E-01 | ? | 1.00E+00 | 5.10E-03 | 1.00E+00 |
| ENSG000000 | ZNF582 | 9.88E-01 | 9.96E-01 | 1.00E+00 | 1.65E-03 | 1.00E+00 |
| ENSG000001 | LRRC4 | 9.88E-01 | 9.96E-01 | 9.98E-01 | -3.46E-03 | -1.00E+00 |
| ENSG000001 | VWA1 | 9.88E-01 | 9.96E-01 | 1.00E+00 | 1.86E-03 | 1.00E+00 |
| ENSG000002 | --- | 9.88E-01 | ? | 9.93E-01 | -1.06E-02 | -1.01E+00 |
| ENSG000002 | --- | 9.88E-01 | ? | 1.00E+00 | 4.85E-03 | 1.00E+00 |
| ENSG000001 | RFWD3 | 9.88E-01 | 9.96E-01 | 9.99E-01 | -7.49E-04 | -1.00E+00 |
| ENSG000000 | DSG2 | 9.88E-01 | 9.96E-01 | 1.00E+00 | -5.76E-04 | -1.00E+00 |
| ENSG000002 | CLIC4P1 | 9.88E-01 | ? | 1.01E+00 | 9.44E-03 | 1.01E+00 |
| ENSG000001 | PARP8 | 9.88E-01 | 9.96E-01 | 9.99E-01 | -9.39E-04 | -1.00E+00 |
| ENSG000001 | PSPC1 | 9.88E-01 | 9.96E-01 | 1.00E+00 | 9.77E-04 | 1.00E+00 |
| ENSG000001 | ZNF208 | 9.88E-01 | 9.96E-01 | 9.98E-01 | -3.05E-03 | -1.00E+00 |
| ENSG000002 | --- | 9.88E-01 | ? | 1.00E+00 | 4.69E-03 | 1.00E+00 |
| ENSG000001 | BCL9L | 9.88E-01 | 9.96E-01 | 1.00E+00 | -6.60E-04 | -1.00E+00 |
| ENSG000002 | --- | 9.88E-01 | ? | 1.01E+00 | 1.25E-02 | 1.01E+00 |
| ENSG000000 | TRPM7 | 9.88E-01 | 9.96E-01 | 1.00E+00 | 9.48E-04 | 1.00E+00 |
| ENSG000002 | PYCARD-AS1 | 9.88E-01 | ? | 9.92E-01 | -1.11E-02 | -1.01E+00 |
| ENSG000001 | EXOC5P1 | 9.88E-01 | ? | 9.94E-01 | -8.57E-03 | -1.01E+00 |
| ENSG000001 | TESMIN | 9.88E-01 | 9.96E-01 | 9.99E-01 | -1.91E-03 | -1.00E+00 |
| ENSG000001 | RNPC3 | 9.89E-01 | 9.96E-01 | 9.99E-01 | -1.08E-03 | -1.00E+00 |
| ENSG000001 | EIF4G1 | 9.89E-01 | 9.96E-01 | 1.00E+00 | 3.27E-04 | 1.00E+00 |
| ENSG000002 | --- | 9.89E-01 | ? | 1.00E+00 | 7.11E-03 | 1.00E+00 |
| ENSG000001 | FUT10 | 9.89E-01 | 9.97E-01 | 9.99E-01 | -8.91E-04 | -1.00E+00 |
| ENSG000002 | --- | 9.89E-01 | 9.97E-01 | 1.00E+00 | 1.91E-03 | 1.00E+00 |
| ENSG000001 | PRADC1 | 9.89E-01 | 9.97E-01 | 1.00E+00 | 1.50E-03 | 1.00E+00 |
| ENSG000002 | MT1A | 9.89E-01 | ? | 1.01E+00 | 8.03E-03 | 1.01E+00 |
| ENSG000001 | PPM1D | 9.89E-01 | 9.97E-01 | 9.99E-01 | -9.73E-04 | -1.00E+00 |
| ENSG000001 | CERS6 | 9.89E-01 | 9.97E-01 | 1.00E+00 | -4.96E-04 | -1.00E+00 |
| ENSG000002 | HNRNPKP4 | 9.89E-01 | ? | 1.00E+00 | 3.45E-03 | 1.00E+00 |
| ENSG000001 | FAM209A | 9.89E-01 | ? | 9.92E-01 | -1.15E-02 | -1.01E+00 |
| ENSG000002 | --- | 9.89E-01 | ? | 9.97E-01 | -3.89E-03 | -1.00E+00 |
| ENSG000002 | ST13P5 | 9.89E-01 | ? | 1.00E+00 | 6.02E-03 | 1.00E+00 |
| ENSG000002 | MSH5 | 9.89E-01 | 9.97E-01 | 1.00E+00 | 2.30E-03 | 1.00E+00 |
| ENSG000000 | CDC23 | 9.89E-01 | 9.97E-01 | 1.00E+00 | 6.23E-04 | 1.00E+00 |
| ENSG000000 | PUM3 | 9.89E-01 | 9.97E-01 | 9.99E-01 | -9.93E-04 | -1.00E+00 |
| ENSG000002 | DDX11L2 | 9.89E-01 | 9.97E-01 | 9.99E-01 | -2.01E-03 | -1.00E+00 |
| ENSG000000 | MPHOSPH9 | 9.90E-01 | 9.97E-01 | 1.00E+00 | 1.23E-03 | 1.00E+00 |
| ENSG000001 | MIB1 | 9.90E-01 | 9.97E-01 | 1.00E+00 | 7.52E-04 | 1.00E+00 |
| ENSG000002 | PCAT6 | 9.90E-01 | 9.97E-01 | 9.99E-01 | -1.91E-03 | -1.00E+00 |

| | | | | | | |
|---|---|---|---|---|---|---|
| ENSG000001 | KDELR1 | 9.90E-01 | 9.97E-01 | 1.00E+00 | 4.99E-04 | 1.00E+00 |
| ENSG000002 | IGFL2-AS1 | 9.90E-01 | ? | 1.01E+00 | 8.30E-03 | 1.01E+00 |
| ENSG00000C | BRPF3 | 9.90E-01 | 9.97E-01 | 1.00E+00 | -6.05E-04 | -1.00E+00 |
| ENSG000001 | TSSK3 | 9.90E-01 | ? | 9.95E-01 | -6.66E-03 | -1.00E+00 |
| ENSG000001 | TLR2 | 9.90E-01 | 9.97E-01 | 1.00E+00 | 1.55E-03 | 1.00E+00 |
| ENSG000002 | --- | 9.90E-01 | ? | 9.95E-01 | -7.13E-03 | -1.00E+00 |
| ENSG000001 | TMEM221 | 9.90E-01 | ? | 9.95E-01 | -6.87E-03 | -1.00E+00 |
| ENSG000002 | TFIP11-DT | 9.90E-01 | 9.97E-01 | 1.00E+00 | 2.68E-03 | 1.00E+00 |
| ENSG000002 | RN7SL834P | 9.90E-01 | ? | 9.95E-01 | -7.19E-03 | -1.00E+00 |
| ENSG000001 | ARMCX3 | 9.90E-01 | 9.97E-01 | 1.00E+00 | -7.17E-04 | -1.00E+00 |
| ENSG000002 | MIR3150BHC | 9.90E-01 | ? | 9.94E-01 | -9.38E-03 | -1.01E+00 |
| ENSG000001 | HERC2P3 | 9.90E-01 | 9.97E-01 | 9.99E-01 | -1.22E-03 | -1.00E+00 |
| ENSG000001 | MFSD13A | 9.90E-01 | ? | 9.98E-01 | -3.21E-03 | -1.00E+00 |
| ENSG000002 | MCIDAS | 9.90E-01 | ? | 1.01E+00 | 7.79E-03 | 1.01E+00 |
| ENSG00000C | GMIP | 9.90E-01 | 9.97E-01 | 9.99E-01 | -1.02E-03 | -1.00E+00 |
| ENSG000002 | ARFGEF1-DT | 9.90E-01 | ? | 9.96E-01 | -6.09E-03 | -1.00E+00 |
| ENSG000001 | NBEAL2 | 9.91E-01 | 9.97E-01 | 9.99E-01 | -1.01E-03 | -1.00E+00 |
| ENSG000002 | FAM153B | 9.91E-01 | ? | 1.00E+00 | 3.04E-03 | 1.00E+00 |
| ENSG000001 | FAF1 | 9.91E-01 | 9.97E-01 | 1.00E+00 | 5.35E-04 | 1.00E+00 |
| ENSG000002 | LINC02234 | 9.91E-01 | ? | 1.00E+00 | 5.91E-03 | 1.00E+00 |
| ENSG000001 | B4GALT6 | 9.91E-01 | 9.97E-01 | 1.00E+00 | 9.81E-04 | 1.00E+00 |
| ENSG000002 | LINC02763 | 9.91E-01 | ? | 9.96E-01 | -5.95E-03 | -1.00E+00 |
| ENSG000001 | DLST | 9.91E-01 | 9.97E-01 | 1.00E+00 | -5.93E-04 | -1.00E+00 |
| ENSG000001 | EXOG | 9.91E-01 | 9.97E-01 | 1.00E+00 | 1.12E-03 | 1.00E+00 |
| ENSG00000C | ARFGEF1 | 9.91E-01 | 9.97E-01 | 1.00E+00 | 6.71E-04 | 1.00E+00 |
| ENSG00000C | SIRT4 | 9.91E-01 | ? | 9.97E-01 | -3.63E-03 | -1.00E+00 |
| ENSG000001 | TMEM125 | 9.91E-01 | 9.97E-01 | 1.00E+00 | 2.43E-03 | 1.00E+00 |
| ENSG000002 | RALGAPA1P1 | 9.91E-01 | 9.97E-01 | 1.00E+00 | 2.25E-03 | 1.00E+00 |
| ENSG000002 | --- | 9.91E-01 | 9.97E-01 | 1.00E+00 | 2.69E-03 | 1.00E+00 |
| ENSG000001 | GNG5 | 9.91E-01 | 9.97E-01 | 1.00E+00 | -6.50E-04 | -1.00E+00 |
| ENSG000002 | --- | 9.91E-01 | ? | 1.00E+00 | 4.83E-03 | 1.00E+00 |
| ENSG000002 | SPIN3 | 9.92E-01 | 9.97E-01 | 1.00E+00 | 7.32E-04 | 1.00E+00 |
| ENSG000001 | TSEN15 | 9.92E-01 | 9.97E-01 | 9.99E-01 | -7.24E-04 | -1.00E+00 |
| ENSG000002 | --- | 9.92E-01 | ? | 1.00E+00 | 4.28E-03 | 1.00E+00 |
| ENSG000002 | --- | 9.92E-01 | ? | 9.95E-01 | -7.03E-03 | -1.00E+00 |
| ENSG000001 | TBL1X | 9.92E-01 | 9.97E-01 | 1.00E+00 | -6.13E-04 | -1.00E+00 |
| ENSG000001 | FGF1 | 9.92E-01 | 9.97E-01 | 1.00E+00 | 3.50E-03 | 1.00E+00 |
| ENSG000001 | EOLA2 | 9.92E-01 | 9.97E-01 | 1.00E+00 | 7.39E-04 | 1.00E+00 |
| ENSG000002 | --- | 9.92E-01 | ? | 9.97E-01 | -4.86E-03 | -1.00E+00 |
| ENSG000002 | ATXN7L3-AS | 9.92E-01 | ? | 9.96E-01 | -5.34E-03 | -1.00E+00 |
| ENSG000001 | TIMP4 | 9.92E-01 | ? | 1.00E+00 | 2.61E-03 | 1.00E+00 |
| ENSG000001 | GDAP2 | 9.92E-01 | 9.97E-01 | 1.00E+00 | 6.21E-04 | 1.00E+00 |
| ENSG000002 | --- | 9.92E-01 | ? | 1.00E+00 | 5.58E-03 | 1.00E+00 |
| ENSG000001 | FAM8A1 | 9.92E-01 | 9.97E-01 | 1.00E+00 | 5.82E-04 | 1.00E+00 |
| ENSG000001 | GAPVD1 | 9.92E-01 | 9.97E-01 | 1.00E+00 | -4.84E-04 | -1.00E+00 |

| | | | | | | |
|---|---|---|---|---|---|---|
| ENSG000002 | XKR9 | 9.92E-01 | 9.97E-01 | 9.99E-01 | -1.76E-03 | -1.00E+00 |
| ENSG000002 | DUXAP8 | 9.92E-01 | 9.97E-01 | 1.00E+00 | 2.30E-03 | 1.00E+00 |
| ENSG000002 | --- | 9.92E-01 | 9.97E-01 | 1.00E+00 | 1.43E-03 | 1.00E+00 |
| ENSG000001 | WFIKKN1 | 9.92E-01 | ? | 1.00E+00 | 2.67E-03 | 1.00E+00 |
| ENSG00000C | MLLT10 | 9.93E-01 | 9.97E-01 | 1.00E+00 | -6.76E-04 | -1.00E+00 |
| ENSG00000C | HDAC9 | 9.93E-01 | 9.97E-01 | 9.99E-01 | -1.59E-03 | -1.00E+00 |
| ENSG000002 | --- | 9.93E-01 | ? | 1.00E+00 | 5.24E-03 | 1.00E+00 |
| ENSG000002 | --- | 9.93E-01 | 9.97E-01 | 1.00E+00 | 2.01E-03 | 1.00E+00 |
| ENSG000002 | --- | 9.93E-01 | ? | 1.00E+00 | 3.65E-03 | 1.00E+00 |
| ENSG000001 | CDC42EP3 | 9.93E-01 | 9.97E-01 | 1.00E+00 | -6.91E-04 | -1.00E+00 |
| ENSG000001 | EPHX3 | 9.93E-01 | ? | 9.97E-01 | -3.90E-03 | -1.00E+00 |
| ENSG000001 | --- | 9.93E-01 | ? | 9.98E-01 | -2.40E-03 | -1.00E+00 |
| ENSG000001 | FKTN | 9.93E-01 | 9.97E-01 | 1.00E+00 | -5.22E-04 | -1.00E+00 |
| ENSG000002 | TARID | 9.93E-01 | ? | 1.00E+00 | 4.19E-03 | 1.00E+00 |
| ENSG000001 | NXPH3 | 9.93E-01 | ? | 9.96E-01 | -5.22E-03 | -1.00E+00 |
| ENSG000002 | LINC00863 | 9.93E-01 | 9.97E-01 | 1.00E+00 | 6.61E-04 | 1.00E+00 |
| ENSG000001 | CLPB | 9.93E-01 | 9.97E-01 | 1.00E+00 | -4.51E-04 | -1.00E+00 |
| ENSG000001 | RAF1 | 9.93E-01 | 9.97E-01 | 1.00E+00 | -3.19E-04 | -1.00E+00 |
| ENSG000002 | --- | 9.93E-01 | 9.97E-01 | 9.99E-01 | -1.30E-03 | -1.00E+00 |
| ENSG000002 | --- | 9.93E-01 | 9.97E-01 | 1.00E+00 | 2.13E-03 | 1.00E+00 |
| ENSG000001 | WWC1 | 9.93E-01 | 9.97E-01 | 1.00E+00 | 4.09E-04 | 1.00E+00 |
| ENSG000002 | PGLS-DT | 9.93E-01 | 9.97E-01 | 9.99E-01 | -1.73E-03 | -1.00E+00 |
| ENSG000001 | IFIH1 | 9.93E-01 | 9.97E-01 | 1.00E+00 | 8.09E-04 | 1.00E+00 |
| ENSG000001 | CTR9 | 9.93E-01 | 9.97E-01 | 1.00E+00 | 3.23E-04 | 1.00E+00 |
| ENSG000001 | TPP2 | 9.93E-01 | 9.97E-01 | 1.00E+00 | -3.96E-04 | -1.00E+00 |
| ENSG00000C | ANKRD13A | 9.93E-01 | 9.97E-01 | 1.00E+00 | -6.48E-04 | -1.00E+00 |
| ENSG000001 | SLC17A9 | 9.94E-01 | ? | 1.00E+00 | 2.27E-03 | 1.00E+00 |
| ENSG000001 | C5AR1 | 9.94E-01 | 9.98E-01 | 9.99E-01 | -1.06E-03 | -1.00E+00 |
| ENSG00000C | MTHFD2 | 9.94E-01 | 9.98E-01 | 1.00E+00 | -4.77E-04 | -1.00E+00 |
| ENSG000002 | SNHG6 | 9.94E-01 | 9.98E-01 | 1.00E+00 | -4.45E-04 | -1.00E+00 |
| ENSG000002 | PPP3CB-AS1 | 9.94E-01 | 9.98E-01 | 9.99E-01 | -1.29E-03 | -1.00E+00 |
| ENSG000002 | FUNDC2P1 | 9.94E-01 | ? | 9.97E-01 | -3.88E-03 | -1.00E+00 |
| ENSG000002 | --- | 9.94E-01 | ? | 9.97E-01 | -4.39E-03 | -1.00E+00 |
| ENSG000001 | SLAMF9 | 9.94E-01 | 9.98E-01 | 9.99E-01 | -1.95E-03 | -1.00E+00 |
| ENSG000002 | --- | 9.94E-01 | 9.98E-01 | 9.99E-01 | -1.68E-03 | -1.00E+00 |
| ENSG000002 | --- | 9.94E-01 | 9.98E-01 | 1.00E+00 | 3.19E-03 | 1.00E+00 |
| ENSG000002 | LINC01140 | 9.94E-01 | ? | 9.98E-01 | -2.43E-03 | -1.00E+00 |
| ENSG000001 | PCDH17 | 9.94E-01 | ? | 9.98E-01 | -3.06E-03 | -1.00E+00 |
| ENSG000001 | CKS1B | 9.94E-01 | 9.98E-01 | 9.99E-01 | -7.55E-04 | -1.00E+00 |
| ENSG000001 | TMEM88 | 9.95E-01 | ? | 1.00E+00 | 1.95E-03 | 1.00E+00 |
| ENSG000002 | --- | 9.95E-01 | ? | 1.00E+00 | 3.95E-03 | 1.00E+00 |
| ENSG000001 | KDM8 | 9.95E-01 | 9.98E-01 | 9.99E-01 | -7.28E-04 | -1.00E+00 |
| ENSG000001 | STX6 | 9.95E-01 | 9.98E-01 | 1.00E+00 | 2.28E-04 | 1.00E+00 |
| ENSG000002 | PSORS1C2 | 9.95E-01 | ? | 1.00E+00 | 2.05E-03 | 1.00E+00 |
| ENSG000002 | --- | 9.95E-01 | ? | 1.00E+00 | 2.77E-03 | 1.00E+00 |

| | | | | | | |
|---|---|---|---|---|---|---|
| ENSG000001 | LSM11 | 9.95E-01 | 9.98E-01 | 1.00E+00 | -6.42E-04 | -1.00E+00 |
| ENSG000002 | BACH1-IT1 | 9.95E-01 | ? | 9.97E-01 | -4.40E-03 | -1.00E+00 |
| ENSG000002 | --- | 9.95E-01 | ? | 9.98E-01 | -2.53E-03 | -1.00E+00 |
| ENSG000002 | --- | 9.95E-01 | 9.98E-01 | 1.00E+00 | -4.22E-04 | -1.00E+00 |
| ENSG000002 | --- | 9.95E-01 | 9.98E-01 | 1.00E+00 | 2.88E-03 | 1.00E+00 |
| ENSG000001 | SAA2 | 9.95E-01 | 9.98E-01 | 9.99E-01 | -1.19E-03 | -1.00E+00 |
| ENSG000001 | MBNL1 | 9.96E-01 | 9.98E-01 | 1.00E+00 | -3.34E-04 | -1.00E+00 |
| ENSG000001 | NAGK | 9.96E-01 | 9.98E-01 | 1.00E+00 | 2.89E-04 | 1.00E+00 |
| ENSG000002 | SCART1 | 9.96E-01 | ? | 9.98E-01 | -2.51E-03 | -1.00E+00 |
| ENSG000002 | --- | 9.96E-01 | 9.98E-01 | 9.99E-01 | -8.40E-04 | -1.00E+00 |
| ENSG000002 | --- | 9.96E-01 | ? | 9.99E-01 | -2.08E-03 | -1.00E+00 |
| ENSG000001 | TRAK2 | 9.96E-01 | 9.98E-01 | 1.00E+00 | -3.06E-04 | -1.00E+00 |
| ENSG000002 | GUSBP11 | 9.96E-01 | 9.98E-01 | 1.00E+00 | -5.86E-04 | -1.00E+00 |
| ENSG000001 | ZNF385C | 9.96E-01 | ? | 9.98E-01 | -2.33E-03 | -1.00E+00 |
| ENSG000001 | PKN3 | 9.96E-01 | 9.98E-01 | 1.00E+00 | 1.11E-03 | 1.00E+00 |
| ENSG000002 | LINC01220 | 9.96E-01 | ? | 9.97E-01 | -3.62E-03 | -1.00E+00 |
| ENSG000001 | TRAPPC1 | 9.96E-01 | 9.98E-01 | 1.00E+00 | 2.93E-04 | 1.00E+00 |
| ENSG000001 | RRP7A | 9.96E-01 | 9.98E-01 | 1.00E+00 | -3.51E-04 | -1.00E+00 |
| ENSG000002 | --- | 9.96E-01 | ? | 9.98E-01 | -2.72E-03 | -1.00E+00 |
| ENSG000002 | --- | 9.96E-01 | ? | 1.00E+00 | 1.72E-03 | 1.00E+00 |
| ENSG000002 | LINC02821 | 9.96E-01 | ? | 9.97E-01 | -3.76E-03 | -1.00E+00 |
| ENSG000002 | ARHGAP11B- | 9.96E-01 | ? | 1.00E+00 | 2.51E-03 | 1.00E+00 |
| ENSG00000C | SART3 | 9.97E-01 | 9.99E-01 | 1.00E+00 | 1.58E-04 | 1.00E+00 |
| ENSG000001 | RNASEL | 9.97E-01 | 9.99E-01 | 1.00E+00 | -2.80E-04 | -1.00E+00 |
| ENSG000002 | --- | 9.97E-01 | ? | 9.98E-01 | -3.03E-03 | -1.00E+00 |
| ENSG000002 | --- | 9.97E-01 | ? | 9.99E-01 | -1.67E-03 | -1.00E+00 |
| ENSG000002 | DPH6-DT | 9.97E-01 | ? | 1.00E+00 | 1.83E-03 | 1.00E+00 |
| ENSG000001 | EIF2S3B | 9.97E-01 | 9.99E-01 | 9.99E-01 | -8.16E-04 | -1.00E+00 |
| ENSG000001 | LPAR3 | 9.97E-01 | 9.99E-01 | 1.00E+00 | -2.68E-04 | -1.00E+00 |
| ENSG000001 | ZNF428 | 9.97E-01 | 9.99E-01 | 1.00E+00 | 5.13E-04 | 1.00E+00 |
| ENSG000001 | PTEN | 9.97E-01 | 9.99E-01 | 1.00E+00 | 2.00E-04 | 1.00E+00 |
| ENSG000002 | THAP12P7 | 9.97E-01 | ? | 1.00E+00 | 1.30E-03 | 1.00E+00 |
| ENSG000002 | ZNF625 | 9.97E-01 | 9.99E-01 | 1.00E+00 | -5.52E-04 | -1.00E+00 |
| ENSG000002 | GNRHR2 | 9.97E-01 | ? | 1.00E+00 | 9.34E-04 | 1.00E+00 |
| ENSG00000C | TXK | 9.97E-01 | ? | 1.00E+00 | 1.65E-03 | 1.00E+00 |
| ENSG000002 | ATP6V1G2 | 9.97E-01 | ? | 1.00E+00 | 1.26E-03 | 1.00E+00 |
| ENSG00000C | ANOS1 | 9.97E-01 | 9.99E-01 | 1.00E+00 | 2.45E-04 | 1.00E+00 |
| ENSG000002 | --- | 9.97E-01 | ? | 1.00E+00 | 3.47E-03 | 1.00E+00 |
| ENSG000002 | MAP2K4P1 | 9.98E-01 | ? | 9.98E-01 | -2.23E-03 | -1.00E+00 |
| ENSG000002 | NEFL | 9.98E-01 | 9.99E-01 | 1.00E+00 | -3.54E-04 | -1.00E+00 |
| ENSG000001 | ZBTB8A | 9.98E-01 | 9.99E-01 | 1.00E+00 | -2.53E-04 | -1.00E+00 |
| ENSG000002 | CETN4P | 9.98E-01 | ? | 1.00E+00 | 1.28E-03 | 1.00E+00 |
| ENSG000001 | SLC35F2 | 9.98E-01 | 9.99E-01 | 1.00E+00 | -1.07E-04 | -1.00E+00 |
| ENSG000001 | EML5 | 9.98E-01 | ? | 1.00E+00 | 1.04E-03 | 1.00E+00 |
| ENSG000001 | SMPD4 | 9.98E-01 | 9.99E-01 | 1.00E+00 | 1.26E-04 | 1.00E+00 |

| | | | | | | |
|---|---|---|---|---|---|---|
| ENSG000002 | --- | 9.98E-01 | ? | 1.00E+00 | 1.15E-03 | 1.00E+00 |
| ENSG00000C | SLC44A1 | 9.98E-01 | 9.99E-01 | 1.00E+00 | 1.49E-04 | 1.00E+00 |
| ENSG000002 | --- | 9.98E-01 | ? | 1.00E+00 | -5.99E-04 | -1.00E+00 |
| ENSG000001 | CCDC148 | 9.98E-01 | 9.99E-01 | 1.00E+00 | -5.19E-04 | -1.00E+00 |
| ENSG000002 | PPT2-EGFL8 | 9.98E-01 | 9.99E-01 | 1.00E+00 | -4.15E-04 | -1.00E+00 |
| ENSG000001 | FOXK1 | 9.98E-01 | 9.99E-01 | 1.00E+00 | -1.64E-04 | -1.00E+00 |
| ENSG000001 | L3HYPDH | 9.98E-01 | 9.99E-01 | 1.00E+00 | -1.92E-04 | -1.00E+00 |
| ENSG000001 | MAPK12 | 9.98E-01 | 9.99E-01 | 1.00E+00 | 2.22E-04 | 1.00E+00 |
| ENSG000001 | ZNF573 | 9.98E-01 | 9.99E-01 | 1.00E+00 | -3.03E-04 | -1.00E+00 |
| ENSG000002 | SETSIP | 9.98E-01 | ? | 9.99E-01 | -1.24E-03 | -1.00E+00 |
| ENSG000001 | AGFG2 | 9.98E-01 | 9.99E-01 | 1.00E+00 | -1.97E-04 | -1.00E+00 |
| ENSG000002 | GPAA1P2 | 9.99E-01 | ? | 1.00E+00 | 1.10E-03 | 1.00E+00 |
| ENSG000001 | DOCK8-AS1 | 9.99E-01 | ? | 1.00E+00 | -6.48E-04 | -1.00E+00 |
| ENSG000001 | CCDC102A | 9.99E-01 | 1.00E+00 | 1.00E+00 | 2.79E-04 | 1.00E+00 |
| ENSG000001 | FASN | 9.99E-01 | 1.00E+00 | 1.00E+00 | 1.53E-04 | 1.00E+00 |
| ENSG000001 | NIPAL2 | 9.99E-01 | 1.00E+00 | 1.00E+00 | -6.68E-05 | -1.00E+00 |
| ENSG000001 | RNF121 | 9.99E-01 | 1.00E+00 | 1.00E+00 | -8.92E-05 | -1.00E+00 |
| ENSG000002 | --- | 9.99E-01 | ? | 9.99E-01 | -7.63E-04 | -1.00E+00 |
| ENSG000001 | LINS1 | 9.99E-01 | 1.00E+00 | 1.00E+00 | -8.89E-05 | -1.00E+00 |
| ENSG000002 | CCNL2 | 9.99E-01 | 1.00E+00 | 1.00E+00 | -8.45E-05 | -1.00E+00 |
| ENSG000001 | C15orf40 | 9.99E-01 | 1.00E+00 | 1.00E+00 | 6.57E-05 | 1.00E+00 |
| ENSG000002 | PTOV1-AS2 | 9.99E-01 | 1.00E+00 | 1.00E+00 | 1.26E-04 | 1.00E+00 |
| ENSG000001 | SSC5D | 9.99E-01 | ? | 1.00E+00 | 4.63E-04 | 1.00E+00 |
| ENSG000001 | PLA2G4F | 9.99E-01 | ? | 1.00E+00 | -2.11E-04 | -1.00E+00 |
| ENSG000001 | FAM86B3P | 9.99E-01 | ? | 1.00E+00 | -5.32E-04 | -1.00E+00 |
| ENSG000001 | GGT1 | 9.99E-01 | 1.00E+00 | 1.00E+00 | -1.30E-04 | -1.00E+00 |
| ENSG000001 | TMEM151B | 9.99E-01 | ? | 1.00E+00 | -3.92E-04 | -1.00E+00 |
| ENSG00000C | GNB1 | 1.00E+00 | 1.00E+00 | 1.00E+00 | 1.62E-05 | 1.00E+00 |
| ENSG000002 | PSG2 | 1.00E+00 | ? | 1.00E+00 | 3.20E-04 | 1.00E+00 |
| ENSG000001 | MED4 | 1.00E+00 | 1.00E+00 | 1.00E+00 | -2.15E-05 | -1.00E+00 |
| ENSG000001 | SALL4 | 1.00E+00 | ? | 1.00E+00 | 1.78E-04 | 1.00E+00 |
| ENSG000001 | UTP3 | 1.00E+00 | 1.00E+00 | 1.00E+00 | -1.41E-05 | -1.00E+00 |
| ENSG000002 | --- | 1.00E+00 | ? | 1.00E+00 | 1.39E-04 | 1.00E+00 |
| ENSG000001 | C3orf62 | 1.00E+00 | 1.00E+00 | 1.00E+00 | -1.53E-05 | -1.00E+00 |
| ENSG000001 | ARMC2 | 1.00E+00 | 1.00E+00 | 1.00E+00 | 2.79E-05 | 1.00E+00 |

(PLEC-KD vs SCR)
